# Developmental dynamics of the postsynaptic proteome to understand synaptic maturation and dysmaturation

**DOI:** 10.1101/2022.05.05.490828

**Authors:** Takeshi Kaizuka, Takehiro Suzuki, Noriyuki Kishi, Manfred W. Kilimann, Takehiko Ueyama, Masahiko Watanabe, Hideyuki Okano, Naoshi Dohmae, Toru Takumi

## Abstract

The postsynaptic density (PSD) is a protein condensate composed of ∼1,000 proteins beneath the postsynaptic membrane of excitatory synapses. The number, shape, and plasticity of synapses are altered during development. However, the dynamics of synaptic protein composition across development have not been fully understood. Here we show alterations of PSD protein composition in mouse and primate brains during development. Proteins involved in synapse regulation are enriched in the differentially expressed (288 decreased and 267 increased) proteins on mouse PSD after a 2-week-old. We find that the changes in PSD protein abundance in mouse brains correlate with gene expression levels in postnatal mice and perinatal primates. This alteration of PSD composition is likely to be defective in the brains of mouse models or patients with autism spectrum disorder (ASD). Finally, we demonstrate that the brain of the common marmoset (*Callithrix jacchus*) changes PSD composition after the juvenile period. The alteration of PSD composition after 2-month-old is distinct from that observed in mice. Our results provide a comprehensive architecture of the remodeling of PSD composition across development, which may explain the molecular basics of synapse maturation and the pathology of psychiatric disorders, such as ASD.

## Introduction

In the central nervous system, neurons communicate through synapses. The postsynaptic parts of excitatory synapses are formed mainly on small protrusions on the dendrites called dendritic spines^1,2^. The formation of neuronal circuits occurs throughout the developmental period, and changes in dendritic spines are a crucial part of this process. During development, the number, structure, and dynamics of dendritic spines undergo significant changes^2^. In the mouse brain, the number of dendritic spines increases rapidly in the postnatal period and reaches its peak at the early juvenile stage (3-week-old)^3–5^. In the brain of primates, including humans, the number of dendritic spines increases around perinatal and early postnatal periods and reaches its peak at the early juvenile stage^6,7^. The subsequent decrease of dendritic spines at the late developmental stage is known as ‘pruning’, which continues until adulthood^2^. Spines arise from thin protrusions on dendrites, called filopodia^8,9^. Filopodia grow into spines by enlargement of the heads. Thus, thin spines are immature spines, whereas mushroom-shaped spines are considered mature spines. In the mouse cortex, the ratio of mushroom spines increases from about 40% to 80% between 2-week-old and 4-week-old^10^. Immature spines are highly dynamic and have a shorter lifetime than mature ones^9^. The turnover rate of spines in the mouse cortex shows about a 90% decrease from 2-week-old to 8-week-old^4^.

Dendritic spines have been implicated in neuropsychiatric disorders. The data on synaptic molecules for autism spectrum disorder (ASD) have been accumulating since neuroligin 3, a synaptic molecule, was first reported as a risk gene for ASD, and the concept of “synaptopathy” is currently used to broadly describe certain features of psychiatric or neurological diseases^11–13^. Alteration in spine density or spine morphology has been reported in patients and model animals of ASD, schizophrenia (SCZ), and Alzheimer’s diseases (AD)^2^.

Synaptic plasticity also changes during development^14,15^. These changes are essential for strengthening or weakening synaptic connectivity and the formation and elimination of synapses. Proteins expressed on synapses have crucial roles in these processes, and most of these proteins are localized on the postsynaptic density (PSD).

The PSD is a protein complex beneath the plasma membrane of dendritic spines. The PSD is composed of more than 1,000 proteins, including neurotransmitter receptors, cell-cell adhesion molecules, scaffolding proteins, and signaling enzymes^16,17^. The combination of these proteins makes spines functional synaptic compartments. The PSD can be biochemically purified using differential and density gradient centrifugation^18^. Proteome analysis of the PSD has been performed for various species under both physiological and pathological conditions^16^. During development, the composition of PSD also changes^19,20^. The alteration of PSD composition may be involved in synaptic maturation. So far, several studies have shown the developmental trajectory of gene expression and protein expression in rodents and primates^21–29^. In addition, studies focused on specific proteins in the PSD showed differences in the composition and assembly of protein complexes in the mouse brains at distinct developmental stages^30,31^. By contrast, relatively few papers have addressed alterations of PSD composition during the developmental period using a comprehensive approach. Proteome analysis of synaptic membrane showed alteration of synaptic protein composition in mouse visual cortex at postnatal day (P) 34-78 and rat medial prefrontal cortex at P30-46^26,27^. However, changes in PSD protein composition, including younger periods (i.e., around P14-P21), where differences in number, shape, and turnover rate of spines are observed at a higher rate, have not been analyzed using an unbiased approach. Moreover, the developmental alteration of PSD composition in the primate brain has not been reported.

In this study, we performed a quantitative proteome analysis of the PSD of mouse and marmoset brains at four timepoints during postnatal development when synapse maturation takes place. We found that proteins related to synapse regulation are enriched in proteins with significant changes in amounts during this period. Together with systematic bioinformatics analyses using reported transcriptome datasets, we found a positive correlation between the relative abundance of mRNA and proteins located on PSD, suggesting that transcriptional regulation is involved in PSD remodeling. Referring to transcriptome datasets of primate brains, we found that similar mRNA abundance alteration occurs during a perinatal period in humans and macaques. Comparing with the transcriptome of ASD patient brains, we found that PSD remodeling is thought to be defective in patients with ASD. Although the changes in mRNA abundance in postnatal primates are relatively small, we found alteration of PSD composition in the neocortex and cerebellum of postnatal marmoset. Our results uncovered developmental trajectories of PSD proteome in rodents and primates, which would be involved in the maturation of synapses.

## Results

### Proteome analysis of mouse PSD during postnatal development

We prepared brain samples from 2, 3, 6, and 12-week-old mice. Purification of PSD from brain tissue was performed using differential and sucrose density gradient centrifugation as reported previously^18^ (See also Methods) (Extended Data Fig. 1a). We confirmed that a core scaffolding molecule in PSD, PSD-95, was enriched in the PSD fraction, whereas a presynaptic protein, synaptophysin, was mostly eliminated (Extended Data Fig. 1b). The prepared PSD samples were subjected to LC-MS/MS analysis. The experiment was performed four times independently. The resulting data were subjected to label-free quantification (LFQ). In total, 4,144 proteins were detected from 16 PSD samples. We then extracted proteins according to the following three criteria; (1) at least two unique peptides were identified, (2) quantified in all 16 datasets, and (3) coefficient of variation < 100 in all ages. In the case when multiple proteins are encoded by a single gene, we selected a single protein encoded by a single gene whose signal intensity is highest. The result of this selection was 2,186 proteins (Supplementary Table 1). Principal component analysis (PCA) of relative protein abundance showed that protein composition from 2-week-old to 12-week-old gradually changed (Fig. 1a). Heatmap analysis of individual protein abundance showed the subsets of proteins that gradually increased or decreased in abundance as development progressed (Fig. 1b). These data describe the gradual transition of PSD composition from the juvenile to the adult stage.

**Figure 1.**
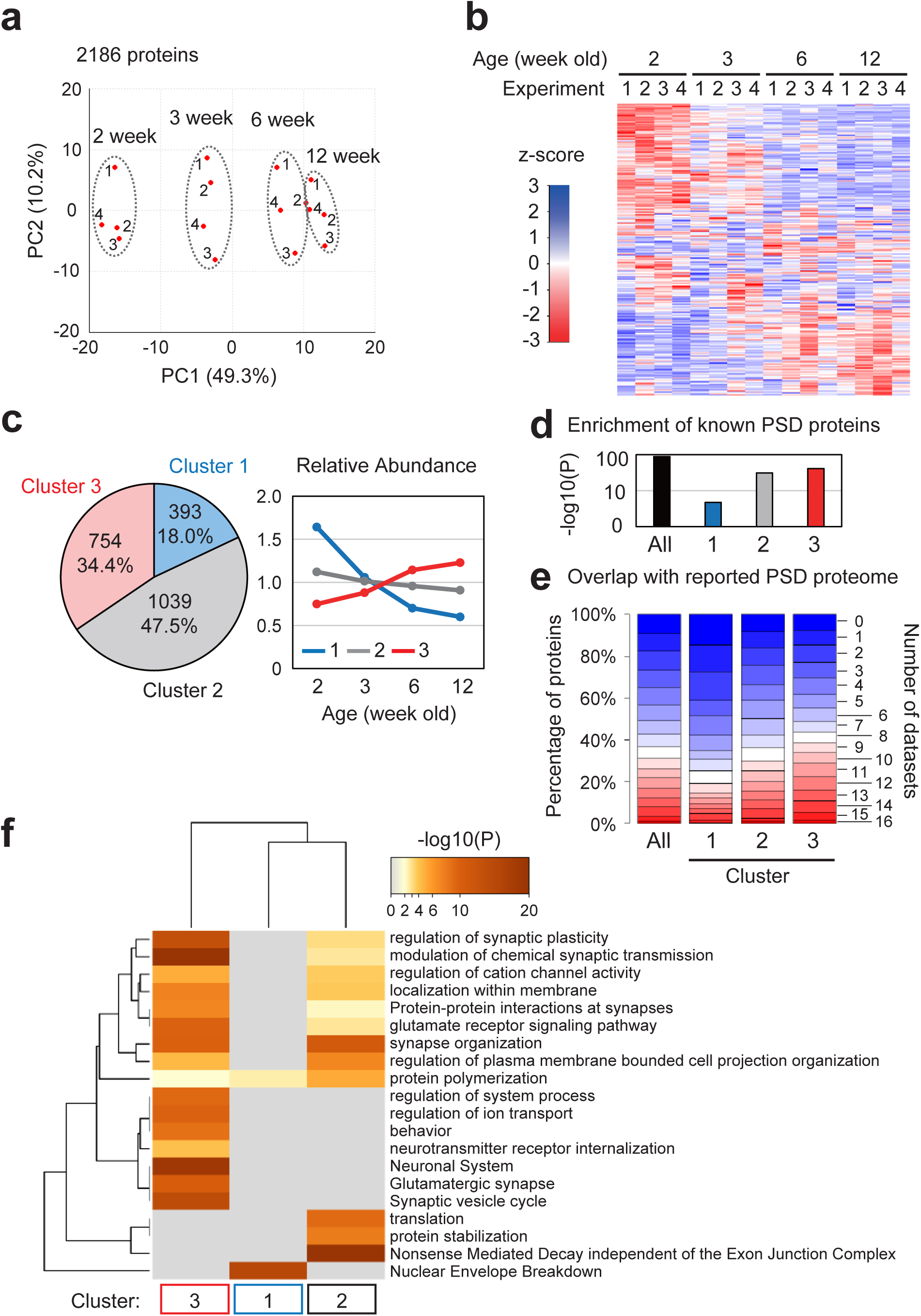
Remodeling of the postsynaptic density (PSD) during postnatal development in mice. (a and b) PSD samples prepared from 2-, 3-, 6-, and 12-week-old mouse brains were subjected to LC-MS/MS to perform label-free quantification. Principal component analysis (PCA) was performed using the relative abundance values of 2,186 proteins (a). The heatmap shows the relative abundance of each protein (b). (c) Identification of PSD protein clusters that are co-regulated during development using k-means clustering based on the mean protein abundance. (left) The pie chart shows the number of proteins in each identified cluster. (right) Expression profile (average of relative abundance) of the proteins in each cluster. (d) Enrichment of proteins reported being expressed on PSD (GOTERM_CC postsynaptic density) of each cluster’s 2,186 proteins or proteins. (e) Overlap between the 2,186 proteins or proteins in each cluster and 16 previously published PSD proteome datasets. (f) Statistically enriched Gene Ontology terms and pathway terms in each cluster identified by Metascape. Top20 terms are displayed as a hierarchically clustered heatmap. The heatmap cells are colored by their P-values; grey cells indicate the lack of enrichment for that term in the corresponding protein list.

### Distinct profile of the co-regulated protein clusters

To identify clusters of co-regulated groups of postsynaptic proteins, we performed k-means clustering. The three clusters were selected based on the best combination of indices provided by the NbClust R package (see Methods for details). The resulting Clusters 1, 2, and 3 were composed of 393, 1039, and 754 proteins, respectively (Fig. 1c and Extended Data Fig. 2, Supplementary Table 1). The abundance of proteins in Cluster 1 gradually decreased but increased in Cluster 3, whereas the abundance of Cluster 2 proteins remained almost unchanged across development (Fig. 1c). We checked whether the proteins in each cluster include proteins reported to localize on PSD using (1) Gene Ontology (GO) analysis and (2) evaluation of the overlap with the proteome of PSD fraction reported in 16 datasets^32,33,34–41,42–46^. We found significant enrichment of the proteins expressed on PSD (GOTERM_CC postsynaptic density) in the 2,186 proteins, and more than 90% of the 2,186 proteins were reported in at least 1 dataset (Fig. 1d, e). Although the enrichment and overlap were also found in each Cluster, the Cluster 1 proteins showed a relatively low enrichment overlap rate (Fig.1 d, e). One possible reason for this result is that the 16 datasets referred to here are proteins detected in adult brains. Indeed, proteins detected in crude PSD fraction of P9 mouse^47^ consist of 16.8%, 19.7%, and 13.6% of Cluster 1, 2, and 3 proteins, respectively. These results suggest that Cluster 1 includes PSD proteins in the juvenile brain, which have not been studied well. To understand what kinds of proteins are included in each cluster, we performed GO analysis and pathway analysis using Metascape and SynGO^48,49^. We found that proteins expressed on synapse (“synapse organization” and “Protein-Protein interaction within synapse”) are enriched in Clusters 2 and 3 (Fig. 1f). In addition, Cluster 3 showed high enrichment of proteins involved in the modulation of synaptic function, such as “regulation of synaptic plasticity”, “modulation of chemical synapse transmission”, and “neurotransmitter internalization” (Fig. 1f). These proteins may contribute to the reorganization of neural circuits based on synaptic plasticity^14,15^. Considering the term “behavior”, they may also be involved in the alteration of behavioral properties during juvenile and adolescence periods^50^. Although the result of Metascape does not provide much insight into the function of Cluster 1 proteins, analysis using SynGO showed that proteins involved in synaptic signaling are enriched in Cluster 1, as well as Cluster 2 and 3 (Extended Data Fig. 1c). These results show the developmental trajectory of the PSD proteome, which is thought to contribute to the synaptic maturation in the developing mouse brain.

### Proteins that regulate synaptic function and dynamics are enriched in differentially expressed proteins

To extract proteins whose expression levels are significantly altered, we used two criteria; (1) Benjamini–Hochberg corrected P-value of one-way analysis of variance (ANOVA) was less than 0.05, (2) fold change (ratio of highest and lowest mean abundance) was more than 1.5. The 690 proteins that meet both criteria were defined as differentially expressed (DE) proteins (Fig. 2a, Supplementary Table 1). We termed DE proteins in Cluster 1 (288) and Cluster 3 (267) as “Decrease” and “Increase”, respectively (Fig. 2b). The classification of 46 major proteins expressed on or associated with PSD is described in Figure 2c. Although 42 proteins belong to Cluster 2 or 3, four proteins are found in a “Decrease” group in Cluster 1. The decrease of DLG3 (also known as SAP-102) and SHANK2 is consistent with previous reports^19,20,51^. We found that key enzymes involved in spine enlargement and synapse stabilization in an “Increase” group, including Ca2+/calmodulin-dependent protein kinase (CaM kinase) (CAMK2A), Protein kinase C (PKC) (PRKCA, PRKCB, PRKCG), and Kalirin-7 (KALRN)^52,53^. To identify signaling pathways affected by the DE proteins on PSD, we performed canonical pathway analysis using Ingenuity Pathway Analysis. We found that the most enriched pathway in “Decrease” and “Increase” proteins is “Synaptogenesis Signaling Pathway” (Fig. 2d). Indeed, we found that at least 126 proteins (55 “Decrease” and 71 “Increase”) have been reported to affect the density, morphology, turnover, transmission, and/or plasticity of synapses (Supplementary Table 2). Among them, 75 proteins (32 in “Decrease” and 43 in “Increase”) have been reported to affect spine density (Supplementary Table 2). This suggests that the increase and decrease of these proteins on PSD are involved in altering synaptic properties across development. Importantly, at least 66 proteins are reported to affect synaptic phenotype by RNAi or overexpression of the protein, suggesting that decreased or increased expression of these proteins can affect synapses (Supplementary Table 2). Other enriched pathways, including “Signaling by Rho Family GTPases”, “RhoGDI Signaling”, and “CREB Signaling in Neurons”, were also involved in synapse regulation^2,54^. To test whether the result of pathway analysis is physiologically relevant, we examined developmental alteration of Rho GTPase signaling. Rho family GTPase signaling plays an essential role in synaptogenesis and synaptic plasticity^2,55,56^. In the dendritic spine, three Rho Family GTPases, RhoA, Rac1, and Cdc42, regulate assembly, disassembly, and severing of actin filament through phosphorylation of cofilin^8,56,57^ (Fig. 2e). As actin filament and cofilin are distributed at cytoplasm within the dendritic spine rather than on PSD, we evaluated phosphorylation of cofilin in synaptosome, an isolated synaptic terminal compartment^58^. We found that phosphorylation of cofilin is decreased upon development in the crude synaptosome fraction obtained from the cortex and cerebellum (Fig. 2f). This result suggests alteration of actin remodeling via cofilin during postnatal development, altering the shape and stability of spines. These results suggest that remodeling of the PSD composition across development plays an essential role in changing synaptic density, dynamics, morphology, and plasticity in the mouse brain.

**Figure 2.**
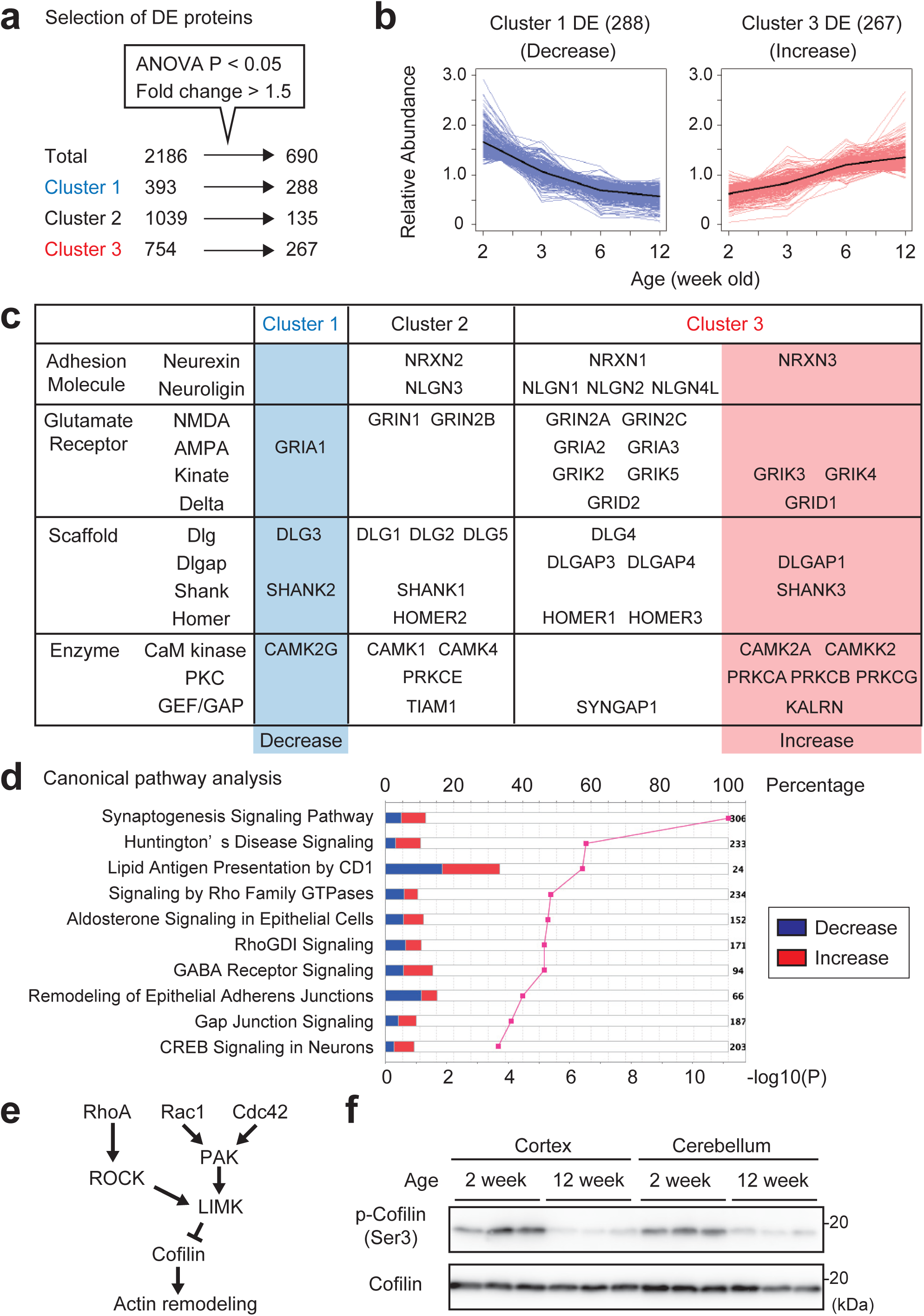
Proteins involved in synapse regulation are enriched in differentially expressed proteins. (a) Selection of differentially expressed (DE) proteins using two criteria; (1) Benjamini–Hochberg corrected P-value of one-way analysis of variance (ANOVA) is less than 0.05, (2) max fold change is more than 1.5. Based on both criteria, 690 proteins were defined as DE proteins (See also Table S1). (b) Expression profile of the DE proteins in Cluster 1 (termed ‘Decrease’) and Cluster 3 (termed ‘Increase’). Black lines indicate the average. (c) Classification of major proteins expressed on or associated with PSD. (d) Canonical pathway analysis using IPA. Pathways with the top 10 lowest P-values were shown. The Magenta line indicates P-value and the bar graph indicates the percentage of proteins included in each term. (e) Overview of Rho family GTPase signaling. Activation of any GTPase results in the inactivation of cofilin by phosphorylation. cofilin affects the morphology of spines through Actin remodeling. (f) Age-dependent reduction of cofilin phosphorylation. Crude synaptosome fractions were obtained from the cortex or cerebellum of mice at indicated ages (n=3 for each sample). 50 μg protein was loaded and phospho-cofilin and total cofilin were detected by immunoblotting.

### Altered abundance of proteins in inhibitory synapses and electrical synapses

In “Decrease” and “Increase” proteins, we also found enrichment of inhibitory synapse proteins (GABA receptor Signaling) and electrical synapse proteins (Gap Junction Signaling) (Fig. 2d). There would be two explanations for this data. One is the alteration of the protein composition of these synapses because the PSD fraction obtained with the method used here also includes proteins in inhibitory synapses and electrical synapses^59,60^. The other is the altered abundance of these proteins in PSD of excitatory synapses because GABA receptor and gap junction protein can be associated with PSD^61,62^. The altered expression of these proteins may be related to the development of excitatory/inhibitory balance^63^.

### Correlation of developmental changes between PSD protein and mRNA levels

What is the upstream mechanism of PSD composition remodeling across development? The simplest explanation would be transcriptional regulation, a decrease or increase in mRNA levels during this period, resulting in a concomitant change in protein abundance. This idea is supported by the global correlation between mRNA abundance and protein abundance^64^. Thus, we referred to two transcriptome datasets of mouse cortex in development^21,28^. Heatmap analysis showed the tendency of correlation between PSD protein and mRNA levels in both datasets. The levels of mRNAs encoding “Decrease” proteins tended to decrease during development, whereas those encoding “Increase” proteins tended to increase during development (Fig. 3a). This correlation supports our idea that transcriptional regulation is involved in altering PSD protein abundance. We found, however, that the timing of the alteration of mRNA abundance is earlier than that of proteins on PSD. The correlation between mRNA changes at P15-adult (in Dataset 1) or P14-adult (in Dataset 2) and PSD protein changes at P14-adult is relatively low (ρ = 0.25 for both datasets) (Extended Data Fig. 3a). On the other hand, mRNA changes from an earlier stage to an adult show a clear positive correlation with PSD protein changes at P14-adult (Extended Data Fig. 3a). The highest correlation was found in both datasets on mRNA changes from P4 to an adult (ρ = 0.53 for both datasets). We focused on the proteins whose changes correlate with mRNA in both datasets. This results in the extraction of 171 proteins in the “Decrease” group and 178 proteins in the “Increase” group (Extended Data Fig. 3b, c). We termed these subgroups as “Decrease A” and “Increase A”, respectively, whereas the other proteins are classified into subgroups “Decrease B” and “Increase B”, respectively (Extended Data Fig. 2, 3c). The plot of mRNA abundance of “Decrease A” and “Increase A” show that they are altered mainly during the early postnatal period before P15 (Fig. 3b). If the abundance of “Decrease A” proteins and “Increase A” proteins are transcriptionally regulated, how can we interpret the time lag between changes in mRNA level and protein level? We thought that the time lag between alteration of mRNA abundance and protein abundance could be due to the slow protein turnover in neurons and referred to the dataset of protein half-life in mouse cortex^65^. The median half-life of the proteins in “Decrease A” and “Increase” A” was 8.32 days and 10.31 days, respectively (Fig. 3c). The long half-life may explain the time lag between the altered abundance of mRNA and protein, as simulated and experimentally described^66^. These data suggest that changes in gene expression levels are involved in the alteration of PSD composition during postnatal development of mice, and the alteration of protein abundance takes several days to become visible after mRNA levels change, as described in Figure 3d.

**Figure 3.**
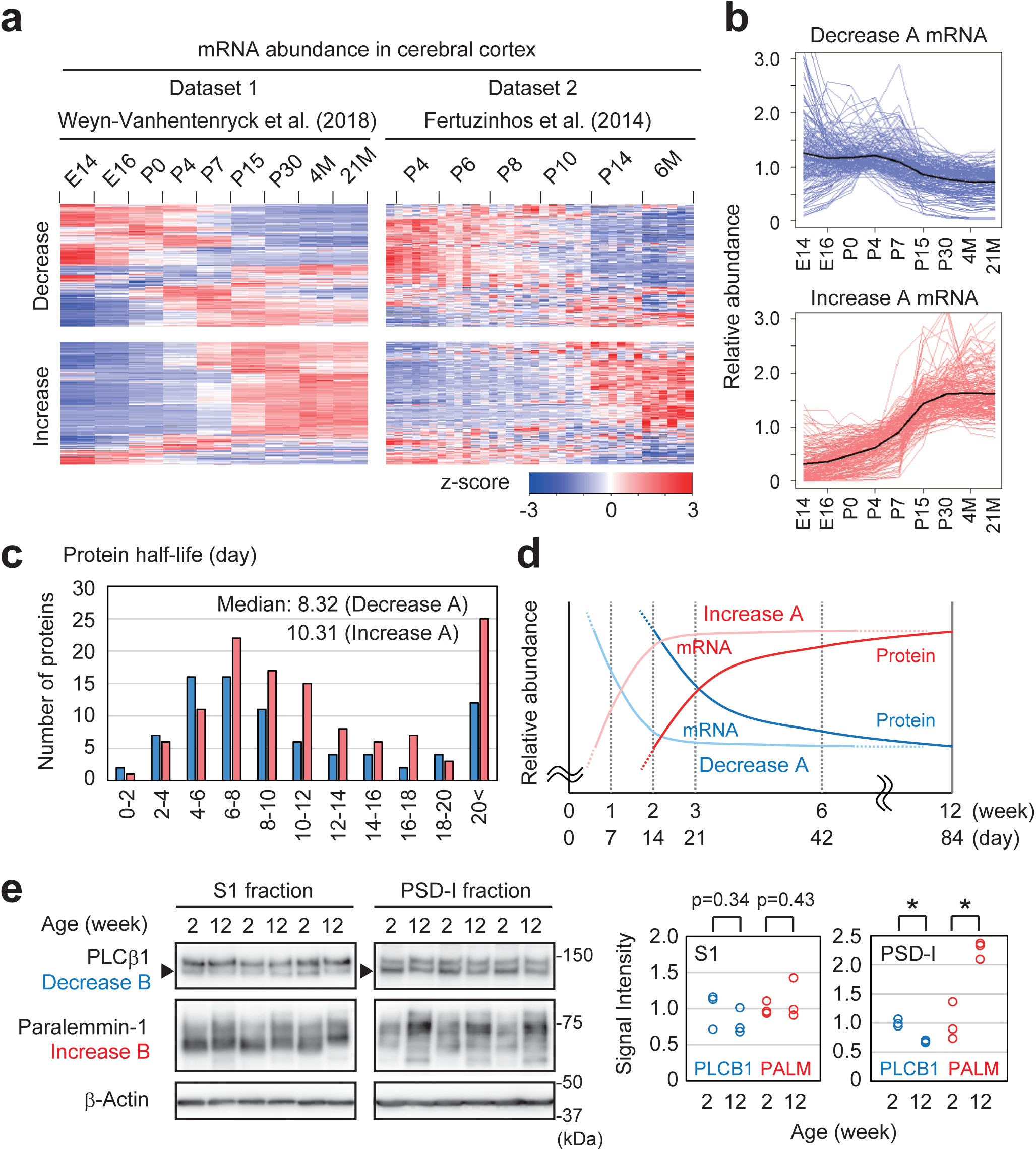
Correlation of protein abundance on PSD and mRNA abundance in mouse cortex during postnatal development. (a) Heatmap of the relative abundance of mRNAs that encode proteins in “Decrease” and “Increase” groups in the mouse cortex^21,28^. Six samples from male and female mice at each timepoint consist of 3 distinct cortical layers (infragranular layers, granular layer, and supragranular layers). (b) Expression profiles of the mRNAs encoding proteins in the groups “Increase A” or “Decrease A”. Relative mRNA abundances (average of 6 samples) were plotted. Black lines indicate the average. (c) The plot of protein half-lives for the proteins in neurons^94^. (d) Model of the time course of mRNA abundance and protein abundance in the PSD. The “Decrease” or “Increase” of protein abundance in the PSD occurred several days or weeks after mRNA. (e) Developmental alteration of PSD localization. Representative of “Decrease” B (PLCB1) and “Increase” B (PALM). 20 μg of proteins obtained from indicated fraction was loaded and analyzed by immunoblotting. Samples were independently prepared from three biological replicates. Quantification of band intensity is shown on the right. Two isoforms of PLCβ1 were detected and the bottom band (shown as an arrowhead) is evaluated because it is thought to be the isoform we detected in the MS dataset. Paralemmin-1 gives multiple bands on Western blots due to differential splicing and phosphorylation (Kutzleb et al., 2007). All immunopositive bands were pooled for quantification. *P < 0.01.

### Alteration of protein localization on PSD

Although relative protein abundance in PSD is primarily correlated with mRNA, other mechanisms should explain the altered abundance of proteins in PSD, particularly proteins in the “Decrease B” and “Increase B” groups. We found that the quantity of some proteins in these groups shows a poor correlation with protein level in synaptosomes reported previously^22^. We hypothesized that the PSD localization of these proteins is altered during development. To test this, we selected phospholipase C beta 1 (PLC®1; PLCB1) and Paralemin-1 (PALM) as representative proteins in “Decrease B” and “Increase B”, respectively. Immunoblotting of S1 and PSD-I fractions showed that the total abundance of these proteins is not significantly altered during development, consistent with the previous report^22^ (Fig.3e). In contrast, the quantity of these proteins in the PSD-I fraction was significantly decreased or increased, respectively (Fig. 3e). These results suggest that alteration of protein localization also contributes to the alteration of PSD composition during development.

### Switching of gene expression in primate brain at the perinatal period

Our next question was whether the PSD remodeling described above is also found in primates, including monkeys and humans. We analyzed transcriptome datasets of developing human and macaque^24,25^. As a result, we found similar changes in mRNA abundance between rodents and primates; mRNAs encoding the homologs of “Decrease A” and “Increase A” tend to be decreased and increased in the neocortex of developing humans and macaques, respectively (Fig. 4a and Extended Data Fig. 4a). Considering the correlation of mRNA and protein levels in the mouse brain, these data suggest that PSD remodeling found in mice also occurs in the primate brain. It should be noted that the alteration of mRNA abundance takes place around the perinatal period rather than the early juvenile period. This suggests that alteration of PSD protein composition occurs around the perinatal or early neonatal period in the primate brain, which is earlier than in mice. We termed the “Decrease A” and “Increase A” proteins whose mRNA abundance is increased and decreased in the developing human neocortex as “Decrease” A1 (117 proteins) and “Increase” A1 (164 proteins), respectively (Extended Data Fig. 2). In the human neocortex, the mRNA abundance of “Decrease A1” and “Increase A1” showed about a 2-3-fold change around the perinatal period (Fig. 4b). We also referred to the mRNA level of other brain regions; hippocampus, striatum, thalamus, amygdala, and cerebellum. The alteration of mRNA abundance was found in all of the regions in human and macaque brains (Fig. 4b and Extended Data Fig. 4). These data suggest that similar PSD remodeling takes place in various brain regions in the primate brain. We showed representative examples of “Decrease A1” and “Increase A1” proteins in Figure 4c. VANGL2 is a Wnt/planar cell polarity pathway component related to synapse formation^67^, and SHANK3 is a scaffolding protein in PSD (Fig. 2c). mRNA of VANGL2 decreased before birth in humans, whereas it occurred between prenatal and juvenile (P15) periods in mice. mRNA of SHANK3 was increased at 35-37 postconceptional weeks in humans, whereas it happened at the neonatal and early juvenile (P0-P15) periods in mice. Thus, the developmental changes in mRNA abundance could be understood as described in Figure 4d. In the mouse brain, the changes in protein levels on PSD were further delayed and occurred between 2-week-old and 12-week-old (Fig. 3d, 4c). The different timing of gene expression changes is thought to be consistent with the timing of synaptogenesis in humans and mice^4,6^.

**Figure 4.**
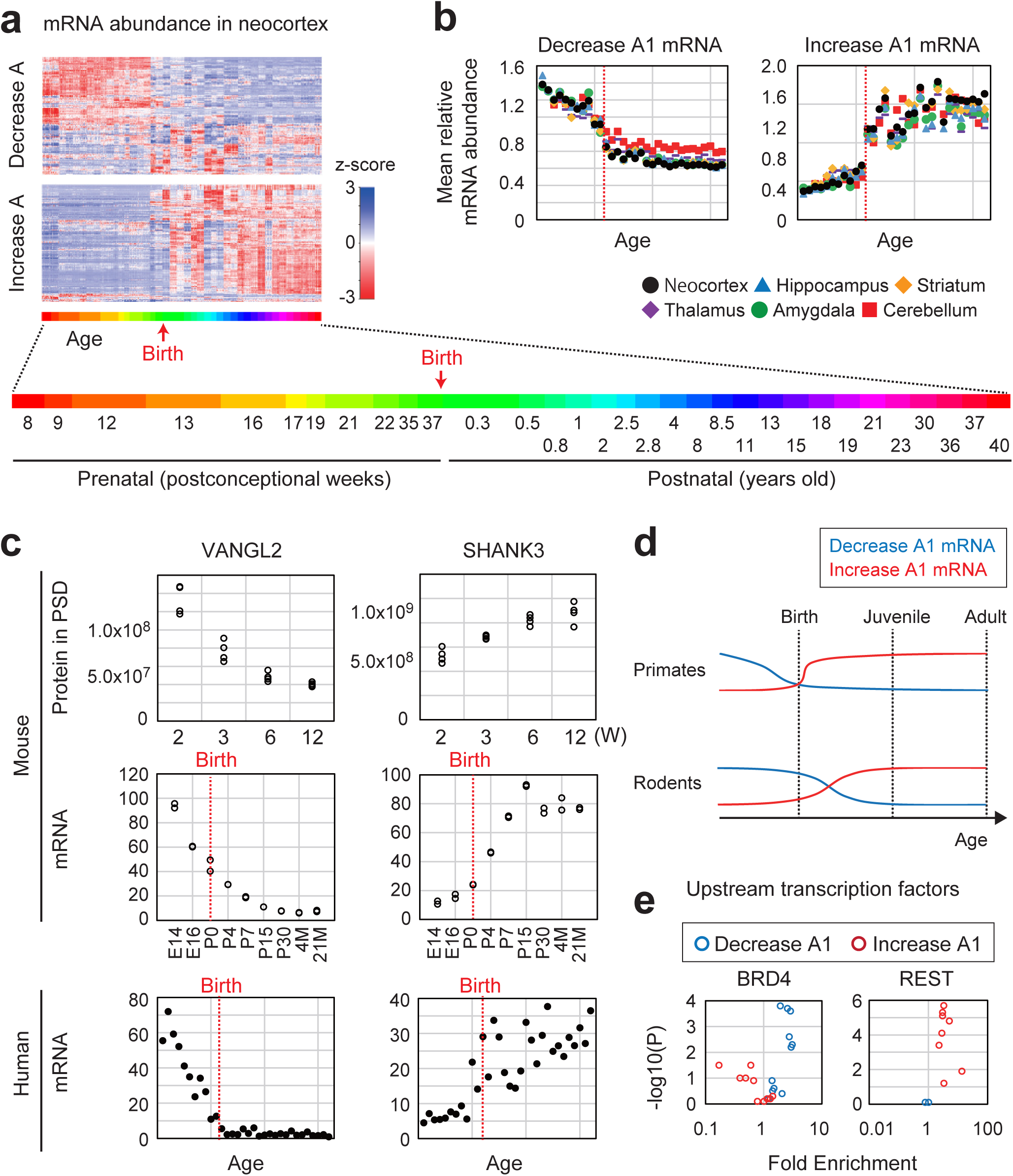
Transcriptional profiles of genes encoding proteins on PSD in developing human brain. (a) Heatmap of the relative abundance of mRNAs that encode “Increase A” and “Decrease A” proteins in cortical regions in the developing human brain^24^. (b) The plot of the mean relative abundance of mRNAs that encode proteins in the groups “Increase A1” (left) or “Decrease A1” (right). The 32-time points of the age of donors were described in (A). Red dotted lines indicate birth. (c) Representative examples of proteins in “Decrease” A1 (left, VANGL2) and “Increase” A1 (right, SHANK3). (top) Protein abundance in the PSD obtained from mouse brains (Table S1). (middle) mRNA abundance in the mouse cortex^28^. (bottom) mRNA abundance in the human brain^24^. The 32-time points of the age of donors were described in (a). Red dotted lines indicate birth. (d) Schematic model of the time course of mRNA abundance of “Increase” A1 and “Decrease” A1 proteins. Similar switching of gene expression occurs mainly in the perinatal period in primates and the early postnatal period in rodents. (e) Representative transcription factors enriched on “Decrease A1” or “Increase A1” genes identified by ChIP-Atlas^68^. See also Extended Data Figure 5a.

### Transcription factors upstream of the genes encoding proteins on PSD

What are the upstream mechanisms that regulate the expression levels of the genes encoding proteins on PSD? We asked whether there are any specific transcription factors involved in regulating the genes encoding proteins on PSD using two methods. We first referred to previously reported ChIP-Seq (chromatin immunoprecipitation sequencing) datasets using ChIP-Atlas^68^. Analysis of datasets of neuronal cells showed enrichment of transcriptional regulators, especially on “Decrease A1” genes (Extended Data Fig. 5a). There are 13 transcription factors enriched on “Decrease A1” and three on “Increase A1” (Extended Data Fig. 5a). BRD4 and REST are representative factors of recurrently detected and significantly enriched on “Decrease A1” genes and “Increase A1” genes, respectively (Fig. 4e). In addition, analysis of transcription factor binding sites showed enrichment of more than 200 transcription factor binding sites around “Decrease A1” and “Increase A1” genes. We described the top 20 items in Extended Data Fig. 5b. These data suggest that the binding of specific transcriptional regulators may contribute to the regulation of gene expression and subsequent remodeling of PSD protein composition.

### Dysregulation of PSD maturation in ASD model mice and patients with ASD

We found that proteins related to various diseases are enriched in our PSD proteome (Extended Data Fig. 6a). Notably, proteins increased during development showed high enrichment of proteins related to neuropsychiatric disorders, including ASD (Extended Data Fig. 6b, c). ASD is a neurodevelopmental disorder that is characterized by impaired social interaction and communication and restricted and repetitive behaviors, activity, or interests^69,70^. Synapse abnormalities are phenotypes of ASD patients and model mice^2,11,13,69,71^. The synaptic defects include alteration of shape, density, and/or turnover of dendritic spines. To test whether the abnormalities in dendritic spines accompany those of PSD composition, we first examined the PSD proteome in a 15q11-13 duplication model mouse (*15q dup*, hereafter), which mimics one of the most frequent copy number variations (CNVs) in ASD^72^. Analysis of PSD proteome in 3-week-old wild type (WT) and *15q dup* mice resulted in quantification of 1,935 proteins (Supplementary Table 4). The difference between WT and *15q dup* mice was relatively small because only 163 proteins were differentially expressed by the t-test (p < 0.05). No proteins were differentially expressed when Benjamini-Hochberg correction was applied (Supplementary Table 3). Nevertheless, PCA of the proteome data showed a cluster of WT and *15q dup* mice and a similar tendency of difference in four biological replicates (Extended Data Fig. 7a). k-means clustering classified these proteins on PSD into 3 clusters, 15q dup-1, 2, and 3 (Extended Data Fig. 7b). About 65% of proteins were classified into 15q dup-2, whose abundance was almost unchanged in *15q dup* mice. The abundance of proteins in 15q dup-1 and 15q dup-3 tended to decrease and increase in *15q dup* mice, respectively. GO- and pathway analysis of each cluster showed enrichment of proteins related to the synapse (“synapse organization”, “regulation of trans-synaptic signaling”, and so on) mainly in 15q dup-2 (Extended Data Fig. 7c). We noticed that GO- and pathway-terms enriched in 15q dup-1 and 15q dup-3 showed some overlap with Cluster 3 and Cluster 1, respectively (Fig. 1f and Extended Data 7c). Indeed, 15q dup-3 showed significant enrichment of Cluster 1 proteins and negative enrichment of Cluster 3 proteins (Extended Data Fig. 7d). Thus, the PSD composition of *15q dup* mice tends to be similar to that of younger mice, suggesting “immaturity” of PSD composition in *15q dup* mice. In *15q dup* mice, enhanced turnover of dendritic spines and increased ratio of thin spines are observed, which could be understood as immature spines^4,73^. Thus, the immaturity of PSD composition we found here might be involved in the immaturity of spine shape and dynamics in *15q dup* mice.

We then wondered whether immaturity of PSD composition is a general feature of ASD because immaturity of the dendritic spine is observed in a wide range of ASD model animals and patients with ASD. Enhanced turnover of spines and increased ratio of thin spines are observed in ASD model mice other than *15q dup*^2,4,73^. In humans, thin dendritic spines are observed in patients with Fragile X syndrome (FXS), a neurodevelopmental disorder with frequent comorbidity of autism^73^. In addition, patients with ASD and several ASD model mice show an increased density of dendritic spines, which suggests disruption of the balance of spine formation and elimination^2,5,74^. Considering that spine number is increased at a younger period, this phenotype can also be understood as immaturity. These phenotypes related to dendritic spines can also be understood as “immaturity”, as dendritic spines observed in patients and model mice of ASD are similar to those in the younger period of normal brains.

To assess the maturation of PSD composition in patients with ASD, we analyzed the transcriptome dataset of ASD patient brains^75^. We focused on genes differentially expressed in a cortical region of patients with ASD using P < 0.05 as a cutoff (P-value before adjusting for multiple testing). In this criterion, approximately 30% of genes were differentially expressed in the neocortex of ASD patients (Fig. 5a). We found that the genes encoding proteins expressed on PSD (2,186 proteins in Supplementary Table 1) are enriched in DE genes in patients with ASD. The enrichment of genes encoding proteins on synapse or PSD in DE genes is consistent with previous reports^76,77^. DE genes were further enriched in the “Increase A1” group, and 86% of them (62/72) were downregulated in ASD patients. Although DE genes were not enriched in the “Decrease A1” group, the ratio of upregulated genes was significantly higher than all PSD proteins (Fig. 5a). The plot of mRNA level alterations during development against that of ASD patients showed a negative correlation between them (ρ = -0.48) (Fig. 5b). These data suggest the possibility that PSD composition in ASD patients is relatively similar to that in the prenatal period compared to healthy subjects. There were 22 “Decrease A1” and 62 “Increase A1” genes, which are upregulated and downregulated in patients with ASD, respectively (Extended Data Fig. 2). We wondered which of these genes are involved in the synaptic pathology of ASD. Simons Foundation Autism Research Initiative (SFARI), a scientific initiative for understanding, diagnosis, and treatment of ASD, provides a list of genes implicated in ASD. We found there were 18 SFARI ASD genes in the 84 (22 “Decrease A1” and the 62 “Increase A1”) genes. (Fig. 5c). They encode proteins involved in various (at least 8) molecular functions in the synapse; fatty acid binding protein (FABP5), scaffold protein (DLGAP1), ion pumps (ATP1A1, ATP2B2), inositol trisphosphate receptor on the endoplasmic reticulum (ITPR1), voltage-gated potassium channel (KCNB1), protein kinase C (PRKCB), small GTPase interacting proteins (ARHGEF9, RAB11FIP5), and syntaxin binding proteins (STXBP1, STXBP5). This suggests that the synaptic pathology in ASD is not simply explained by defects in a single pathway. Instead, it is thought to be caused by the synergistic effect of multiple abnormalities in the synapse. Taken together, these data suggest an “immature” pattern of PSD composition both in the brains of ASD model mice and patients with ASD, which may be involved in synaptic immaturity accompanied by ASD.

**Figure 5.**
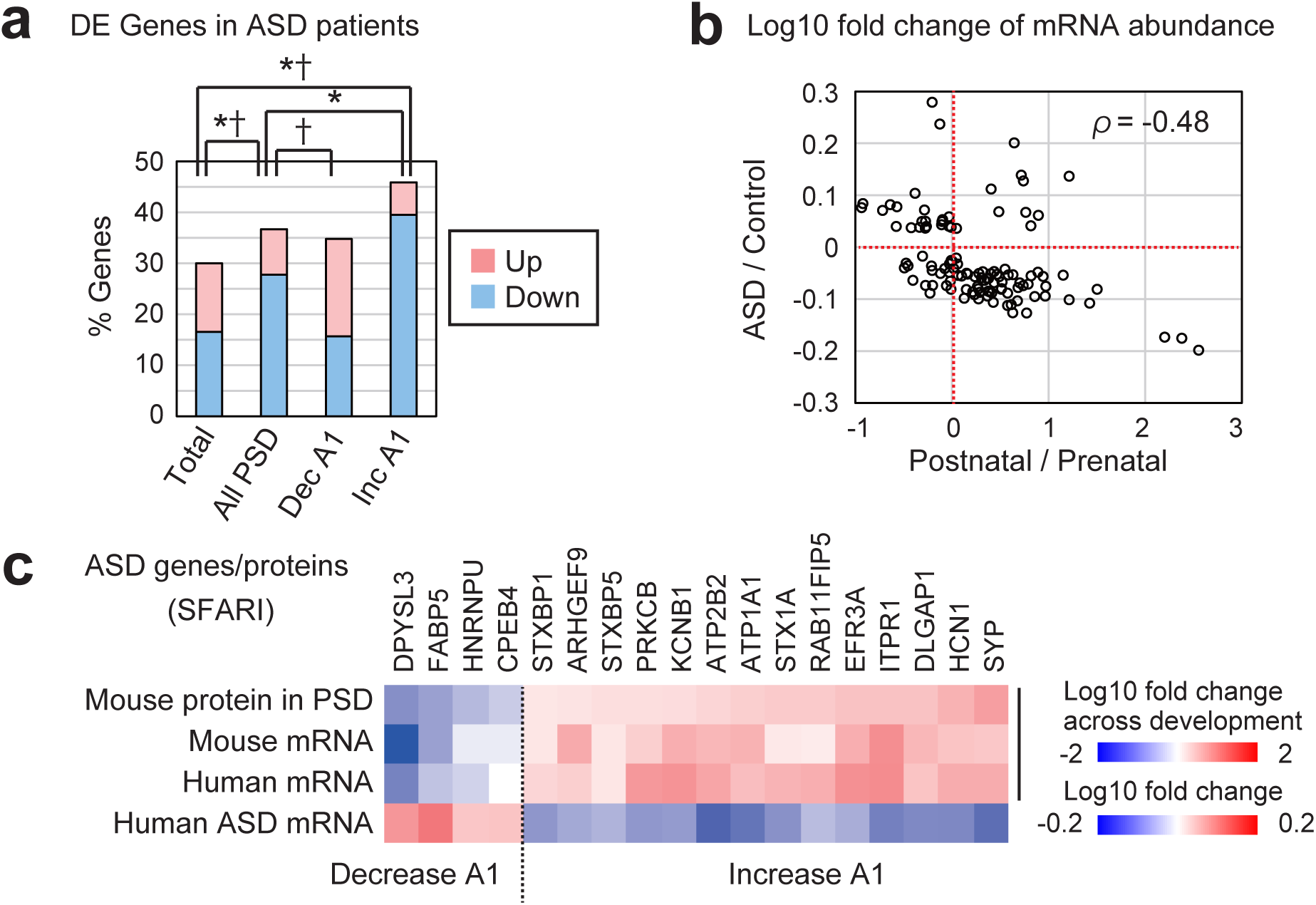
Alteration of PSD composition in ASD model mice and patients with ASD. (a) The percentage of DE (upregulated and downregulated) genes in the cortical region of ASD patients (P < 0.05)^75^. Gene set enrichment analysis was performed using Fisher’s exact test. Asterisk and dagger indicate significant enrichment (P < 0.01) of DE genes and up- or down-regulated genes, respectively. (b) Log10 fold change of mRNA abundance in the developmental human cortex (postnatal vs. prenatal) ^24^ was plotted against that of ASD patients (ASD vs. control)^75^. Red dotted lines indicate zero. (c) SFARI ASD genes showed a negative correlation: “Decrease A1” genes/proteins were increased in ASD, whereas “Increase A1” genes/proteins were decreased in ASD. Heatmap describes the relative abundance of protein or mRNA; Mouse protein: protein abundance (P12w / P2w) (Table S1), Mouse mRNA: mRNA abundance (P180 / P4)^21^, Human mRNA: mRNA abundance (Postnatal / Prenatal), Human ASD mRNA (ASD / Control)^75^.

### Proteome analysis of PSD in the primate brain

The alteration of PSD protein composition in the postnatal mouse brain may explain the changes in synaptic properties during development. This may also be the case in the primate brain. In the brain of the human and common marmoset (*Callithrix jacchus*), more than a 2-fold increase and subsequent decrease in spine density is observed during the postnatal period^6,7^. Our analysis of transcriptome datasets suggested that alteration of PSD composition mediated by gene expression changes occurs during the perinatal period in the primate brain (Fig. 4 and Extended Data Fig. 4). However, alteration of the PSD composition in the later developmental stages is still unknown. We, therefore, decided to perform proteome analysis of the PSD in primate brains. Although the postmortem human brain was a candidate sample for this experiment, there is a problem with protein degradation during the postmortem interval (PMI)^78^. In particular, the PSD proteome of brain regions other than the neocortex is challenging to analyze. We thus decided to use marmoset to obtain PSD from fresh non-human primate brains. The common marmoset is a small New World primate that lives in stable extended families and has well-developed vocal communication^79^. Because of its similarity to humans, the marmoset is regarded as a useful model animal to study cognitive processes and mental illness^80^. To isolate PSD from limited brain samples, we used a 3-step method^78^ instead of a classical purification method. We confirmed enrichment of PSD-95 and elimination of synaptophysin in a PSD fraction prepared with this method (Extended Data Fig. 8a). We found partial degradation of GluN2B (GRIN2B) and significant loss of PSD-95 in PSD fractions 8-24 h after death in the mouse brain, confirming the importance of fresh brain samples for PSD proteome analysis (Extended Data Fig. 8b). Brain samples were prepared from marmosets described in Extended Data Figure 8c. We first analyzed PSD protein composition across seven brain regions in an adult (24-month-old) marmoset (Fig. 6a). Each sample was analyzed with LC-MS/MS 3 times as a technical replicate. Among the detected proteins, 1,960 proteins met the four criteria used for the mouse PSD proteome experiment described above (Supplementary Table 4). PCA showed that the forebrain and hindbrain could be distinguished with the first principal component (Fig. 6b). Heatmap analysis, together with PCA, showed that the PSD composition of the hippocampus and cerebellum is different from that of the other regions (Fig. 6c). Among these proteins, 1,432 proteins were found to be differentially expressed (Benjamini-Hochberg corrected P < 0.05, fold change > 1.5) (Supplementary Table 4). The proteins were classified into 3 clusters, Region-1, 2, and 3 (Fig. 6d). Proteins in Region-1 and Region-3 were enriched in the hippocampus and cerebellum, respectively, whereas Region-2 proteins were ubiquitously expressed (Fig. 6d). GO and pathway analysis showed high enrichment of proteins related to the synapse (“synapse organization”, “synaptic signaling,” and so on) in Region-1 (Fig. 6e). These proteins may be involved in synaptic properties in the processing (Fig. 6e). These proteins might be related to regional variation of splicing^81^.

**Figure 6.**
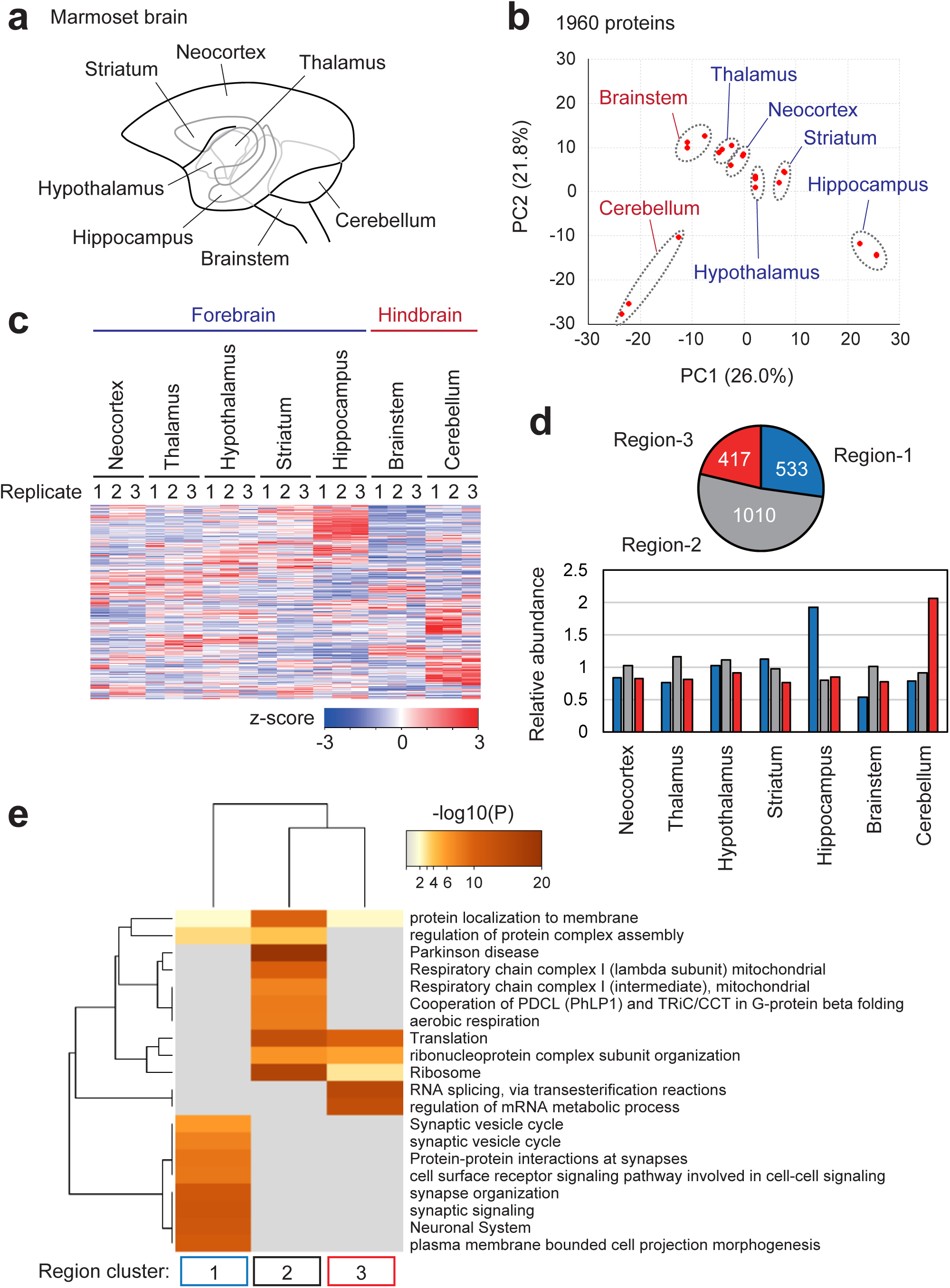
Signatures of PSD proteome composition in marmoset brain regions. (a) Seven brain regions in marmoset. (b and c) PSD samples prepared from individual brain regions of adult (24-month-old) marmoset were subjected to LC-MS/MS to perform label-free quantification. PCA was performed using the relative abundance values of 1,960 proteins (b). The relative abundance of each protein was described using a heatmap (c). (d) Identification of clusters of proteins on PSD using k-means clustering based on the mean protein abundance. (top) The pie chart shows the number of proteins in each identified cluster. (bottom) Expression profile (average of relative abundance) of the proteins in each cluster. (e) Statistically enriched Gene Ontology terms and pathway terms in each cluster identified by Metascape. Top20 terms are displayed as a hierarchically clustered heatmap. The heatmap cells are colored by their P-values; grey cells indicate the lack of enrichment for that term in the corresponding protein list.

### Alteration of PSD protein composition during postnatal development in primates

We then analyzed the alteration of PSD composition during postnatal development in marmoset. The neocortex and cerebellum were selected as representatives of the forebrain and hindbrain, respectively. Brain samples were prepared from 2 months (M), 3 M, 6 M, and 24 M marmosets because the number of dendritic spines in the marmoset neocortex is altered during this period^7^. As a result, 2,361 and 1,740 proteins were found in the neocortex and cerebellum, respectively (Supplementary Table 5, 6). In both regions, 1,335 proteins were found (Extended Data Fig. 8d). PCA and heatmap analysis showed age-dependent alteration of PSD composition in both regions (Fig. 7a, b and Extended Data 8e, f). In the neocortex and cerebellum, 797 and 257 DE proteins were differentially expressed (Benjamini-Hochberg corrected P < 0.05, fold change > 1.5) (Supplementary Table 5, 6). The alteration of protein abundance between the neocortex and cerebellum was relatively low (ρ = 0.28), suggesting that the alteration is highly dependent on brain regions (Extended Data Fig. 8g). k-means clustering classified the proteins into 2 and 3 clusters in the neocortex (NCX1-2) and cerebellum (CB1-3), respectively (Fig. 7c). GO and pathway analysis showed proteins differentially enriched in each cluster (Fig. 7d). All clusters showed enrichment of “cellular protein complex disassembly”, which may be related to turnover of PSD. In NCX1 and CB1, which include “Increase” proteins during development, “regulation of neurotransmitter receptor activity” and “synapse organization” were enriched. They may be involved in altering synaptic numbers and functions during development. Finally, we asked whether the alteration of PSD composition in the postnatal marmoset is similar to that in the postnatal mouse. “Decrease” and “Increase” proteins, which are decreased or increased in mice (Fig. 2b), included both increased and decreased proteins in the marmoset neocortex and cerebellum (Fig. 7e). Although “Increase” proteins showed a weak correlation in the neocortex (ρ = -0.21), the others did not show a significant correlation. The weak correlation between mouse and marmoset proteome data may reflect differences in experimental settings, including brain region and purification method. Nevertheless, this result may suggest different alterations of PSD composition at different developmental stages in the primate brain, as described in Figure 7f. In this model, changes in PSD composition observed in postanal mice are thought to occur in a perinatal period in primates, which is mainly due to altered gene expression (Fig. 4, 7f; Phase 1). At the juvenile-adult stage of primate, there may be changes in PSD composition, which are different from those in Phase 1 (Fig. 7f; Phase 2). Thus, our data revealed the alteration of PSD protein composition in the primate brain.

**Figure 7.**
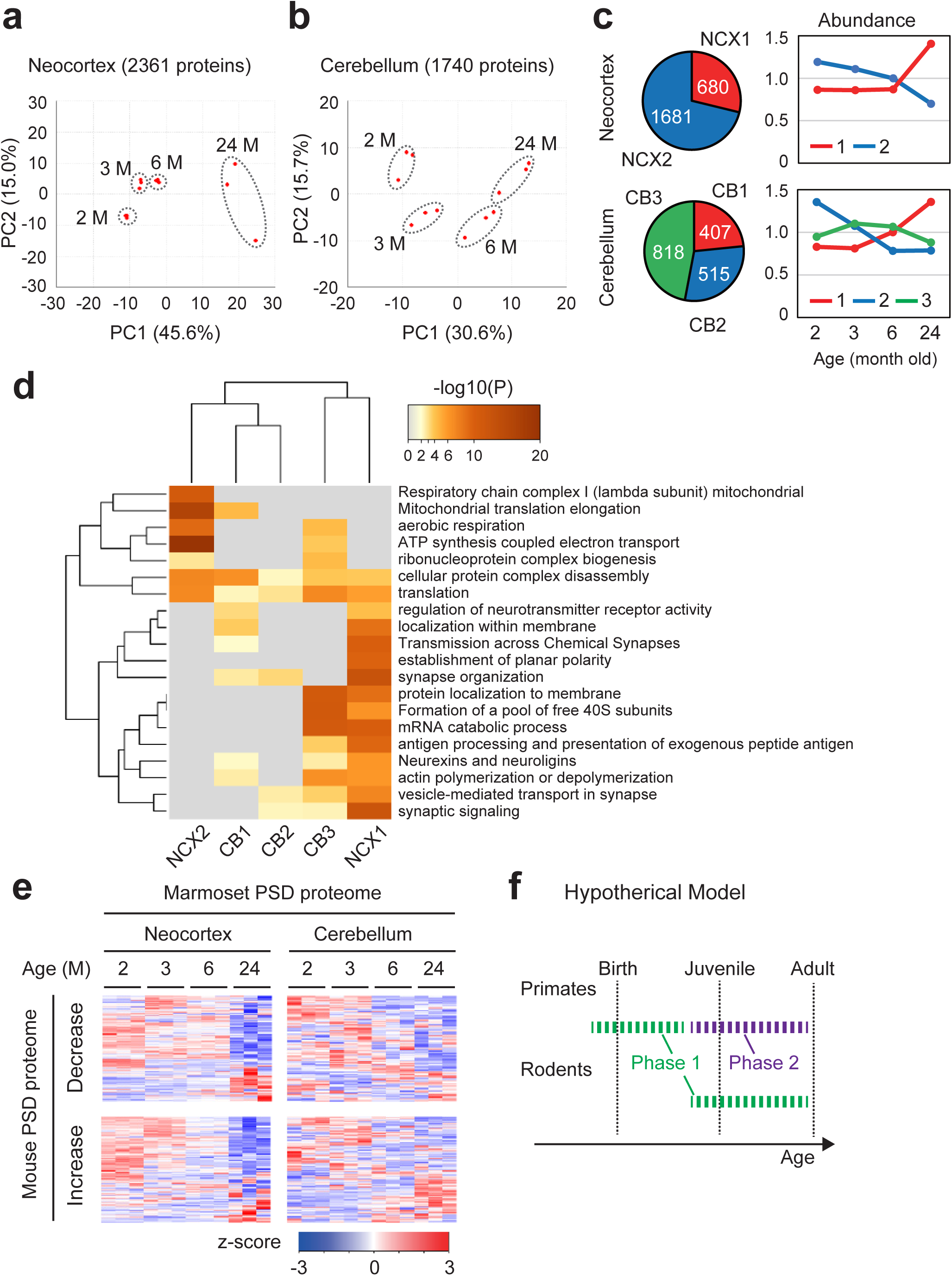
Remodeling of the PSD during postnatal development in marmoset. (a and b) PSD samples prepared from the neocortex (a) or cerebellum (b) of 2-, 3-, 6-, and 24-month-old marmoset brains were subjected to LC-MS/MS to perform label-free quantification. PCA was performed using the relative abundance values of the 2,361 (a) or 1,740 (b) proteins. (c) Identification of clusters of the proteins on PSD using k-means clustering based on the mean protein abundance. (left) The pie charts show the number of proteins in each identified cluster. (right) Expression profile (average of relative abundance) of the proteins in each cluster. (d) Statistically enriched Gene Ontology terms and pathway terms in each cluster identified by Metascape. Top20 terms are displayed as a hierarchically clustered heatmap. The heatmap cells are colored by their P-values; grey cells indicate the lack of enrichment for that term in the corresponding protein list. (e) The relative abundance of proteins whose mouse homologs belong to “Increase” and “Decrease” was described using a heatmap. (f) Hypothetical model of the alteration of PSD composition during development. Phase 1 change occurs during the postnatal period in rodents and the perinatal period in primates. Transcriptional regulation is thought to be involved in Phase 1 change. Phase 2 change occurs during the juvenile-adult period in primates. It is presumably not based on transcriptional regulation. Alteration in Phase 2 is different from that of Phase 1.

## Discussion

In the present study, we revealed the alterations of PSD composition in postnatal mouse and marmoset brains, which occur along with synapse maturation. Comprehensive bioinformatics analyses revealed possible upstream mechanisms and downstream events of this PSD remodeling. In particular, our study is the first to describe the alteration of PSD composition in primate brains during postnatal development. The proteome datasets obtained here will aid in understanding the molecular mechanisms of synapse alteration in primate brains during postnatal development, especially synapse pruning. Considering the observation of the synapse number in previous reports, synapse pruning may occur more significantly in the postnatal primate brain compared to the mouse brain^3,4,6,7^. The proteins in marmoset PSD whose abundance is significantly altered during postnatal development may include critical protein(s) that regulate synapse pruning in primate brains.

We should, however, be aware of the limitations of our proteome datasets of marmoset PSD, which is mainly due to the limited sample availability. First, we used only one marmoset per age. Developmental alteration of the PSD composition we described here may include individual differences. Second, we used a crude PSD fraction obtained by a 3-step method, including contaminant proteins that are not genuine PSD constituents. Third, we did not analyze marmosets younger than 2-months. These issues should be considered in the interpretation of these datasets.

The identification of specific transcription regulators upstream of the genes encoding proteins on PSD may explain their regulatory mechanism. Reference of ChIP-seq data showed that BRD4 is enriched on “Decrease A1” genes, whereas it is less enriched in “Increase A1” genes (Fig. 4d). BRD4 is a transcription factor involved in neuronal development and function^82^. On the other hand, transcriptional repressor REST is enriched only on “Increase A1” genes (Fig. 4d). Downregulation of REST expression during the prenatal period (https://hbatlas.org/) may be involved in the upregulation of “Increase A1” genes and subsequent maturation of PSD composition.

We also found a potential relationship between PSD composition and ASD. Analysis of proteome data of ASD model mouse and transcriptome data of ASD patient brains suggested that PSD composition is “immature” in brains with ASD (Fig. 5). Although we found a relatively small difference in PSD composition between wild-type and *15q dup* mice at 3-seek old, a more significant defect may be detected at other developmental stages, considering the different synaptic phenotypes observed at a different stage^73^. Because developmental alteration of PSD composition can affect multiple signaling pathways related to synapse maturation (Fig. 2d-f), these signals may also be involved in ASD patients and model animals to cause their synaptic phenotypes. For example, enhanced rho family GTPase signaling is observed in Fmr1 KO mice (model of FXS)^83^, suggesting immaturity in synaptic signaling.

What is the mechanism(s) of altered expression of the genes encoding proteins on PSD in patients with ASD? One possible cause is the abnormality in the balance of neuronal excitation and inhibition (E/I balance), which is considered a pathogenic mechanism of ASD^69,84^. The E/I imbalance may cause changes in gene expression patterns. It is reported that hyper-excitation of neurons causes immaturity of gene expression patterns and is implicated in various neuropsychiatric disorders, including ASD, SCZ, and AD^85^. Murano *et al.* termed the genes related to immaturity as hiI (hyperexcitation-induced immaturity-related) genes and hiM (hyperexcitation-induced maturity-related) genes. We found that 17 of 117 genes in “Decrease A1” are encoded by mouse hiI genes and 36 of 164 in “Increase A1” by mouse hiM genes. The overlapping (Decrease A1-hiI and Increase A1-hiM) genes included 17 genes encoding proteins that we summarized in Supplementary Table 2. Those 17 genes, including Kalrn, Rapgef2, Lgi1, Grm1, Camk2a, Camkk2, and Itpr1, belong to “Increase A1” and hiM genes, whereas Arf6 and Mapk8 belong to “Decrease A1” and hiI genes (Supplementary Table 1, 2)^85^. This suggests that hyperexcitation can contribute to the immaturity of PSD protein composition, which may result in synapses with immature properties. This may explain hyperexcitation-induced synaptic immaturity in patients with ASD and other neuropsychiatric disorders.

Another possible mechanism of the altered gene expression in patients with ASD is the disruption of upstream regulators of the genes (Fig. 4e, Extended Data Fig. 5a, b). For example, BRD4, which binds to the promoter sequence of “Decrease A1” genes, may be involved in ASD. In Fmr1 KO mice, the expression of BRD4 protein is upregulated. Inhibition of BRD4 restores the transcriptional profile and rescues excess dendritic spine formation and abnormal social behaviors in Fmr1 KO mice^86^. Upregulation of “Decrease” A1 genes through BRD4 may cause immaturity of PSD composition and, in turn, cause synaptic and behavioral abnormalities. On the other hand, EGR2, whose binding site is enriched around “Increase” A1 genes, is a co-regulator of MECP2, a causal gene of a neurodevelopmental disorder, Rett syndrome (Extended Data Fig. 5b). If this is the case, these proteins can be considered as therapeutic targets for ASD to ameliorate synaptic phenotypes through re-maturation of PSD protein composition.

## Method

### Animals and tissue preparation

The animal experiments were approved by the Animal Experiments Committee in RIKEN. Whole-brain samples from 2, 3, 6, and 12-week-old male ICR mice purchased from Japan SLC Inc. (Shizuoka, Japan) were used. As an ASD model, we used paternal 15q11-13 duplication (*15q dup*) mice^72^, which were maintained by mating male *15q dup* with WT female C57BL/6J mice. Whole-brain samples from 3-week-old male *15q dup* mice were used. After cervical dislocation, the brain was removed from mice, briefly rinsed with ice-cold HBSS (Hanks’ Balanced Salt solution), frozen with liquid nitrogen, and stored at -80°C before use. PSD purification was performed using whole brains pooled from 24 (2 week-old), 8 (3 week-old), or 4 (6 and 12 week-old) mice, respectively. To obtain PSD-I fraction, 6 (2 week-old) or 3 (12 week-old) brains were used. Note that many young mice should be used due to a low yield of PSD from young mice. For the marmoset brain, we used male common marmosets (*Callithrix jacchus*) described in Figure S9C. The animals were sedated with an i.p. injection of ketamine hydrochloride (10 mg/kg) and euthanized via an overdose of sodium pentobarbital (75 mg/ kg i.p.). The whole brain was dissected and individual brain regions were then dissected on ice.

### PSD purification

PSDs were prepared from mouse brains using a previously described method with minor modification^18^. All steps were performed at 4°C or on ice. Homogenization of brains was performed by 12 strokes with Teflon-glass homogenizer in Solution A (0.32 M sucrose, 1 mM NaHCO_3_, 1 mM MgCl_2_ 0.5 mM CaCl_2_, and cOmplete EDTA-free Protease Inhibitor Cocktail). Brain homogenate was centrifuged at 1,400 g for 10 min at 4°C to obtain the pellet and the supernatant fraction. The pellet fraction was resuspended in Solution A with 3 strokes of the homogenizer and centrifuged at 700 g for 10 min at 4°C. The supernatant of the first and second centrifugation was pooled as an S1 fraction and subjected to the subsequent centrifugation at 13,800 g for 10 min at 4°C. The resulting pellet was resuspended with Solution B (0.32 M Sucrose and 1 mM NaHCO_3_) and 6 strokes of the homogenizer (P2 fraction) and centrifuged in a sucrose density gradient (0.85/1.0/1.2 M) for 2 h at 82,500 g. Purified synaptosomes were collected from the 1.0/1.2 M border and diluted twice with Solution B. The purified synaptosome was lysed by adding an equal volume of solution C (1% TX-100, 0.32 M Sucrose, 12 mM Tris-HCl pH 8.1) and rotation at 4°C for 15 min. The sample was then centrifuged at 32,800 g for 20 min at 4°C. The resulting pellet was used as a PSD-I fraction. To obtain the PSD-II fraction, PSD-I was resuspended with Solution B and centrifuged in a sucrose density gradient (1.0/1.5/2.0 M) for 2 h at 201,800 g. The border between 1.0 M and 1.2 M was collected and twice diluted with Solution B. After adding an equal volume of Solution D (1% TX-100, 150 mM KCl), they were centrifuged at 201,800 g for 20 min at 4°C. The resulting pellet, PSD-II, was stored at -80°C before LC-MS/MS analysis. For PSD purification from the marmoset brain, a 3-step method was used to maximize the yield^78^. Homogenization of brain samples was performed by 6 strokes with Teflon-glass homogenizer in Solution A’ (0.32 M sucrose, 10 mM HEPES (pH 7.4), 2 mM EDTA, 5 mM sodium o-vanadate, 30 mM NaF, and cOmplete EDTA-free Protease Inhibitor Cocktail) (tissue weight: volume ratio is 100 mg:1 ml). Brain homogenate was centrifuged at 800 g for 15 min at 4°C the supernatant fraction (S1 fraction). The S1 fraction was then subjected to another centrifugation at 10,000 g for 15 min at 4°C. The resulting pellet was resuspended with Solution B’ (1% Triton X-100, 50 mM HEPES (pH 7.4), 2 mM EDTA, 5 mM EGTA, 5 mM sodium o-vanadate, 30 mM NaF, and cOmplete EDTA-free Protease Inhibitor Cocktail) and 6 strokes of the homogenizer (P2 fraction) and subjected to another centrifugation at 30,000 g for 30 min at 4°C. The resulting pellet was washed once with Solution C’ (50 mM HEPES (pH 7.4), 2 mM EDTA, 5 mM EGTA, 5 mM sodium o-vanadate, 30 mM NaF, and cOmplete EDTA-free Protease Inhibitor Cocktail) and stored at -80°C before LC-MS/MS analysis.

### Immunoblotting

Samples were boiled in sample buffer (62.5 mM Tris-HCl pH6.8, 4% sodium dodecyl sulfate (SDS), 10% glycerol, 0.008% bromophenol blue, and 25 mM DTT). Proteins were separated using SDS-polyacrylamide gel electrophoresis (SDS-PAGE) using 5% (for PLCβ1), 12% (for PALM and β-actin), or 15% (for cofilin and phospho-cofilin) acrylamide gels. Proteins on the gels were transferred onto Immobilon-FL polyvinylidene difluoride membranes (Millipore). The membrane was blocked in blocking buffer (Tris-buffered saline with 0.1% Tween 20 (TBST) and 5% skim milk) at room temperature and then incubated with the indicated primary antibody (1:1000 dilution) overnight in blocking buffer at 4 °C. The membrane was washed three times with TBST, followed by incubation with the respective secondary antibody in blocking buffer for 1 h at room temperature. The membrane was washed five times with TBST. Signal intensities were analyzed using Odyssey near-infrared fluorescence imaging system (LI-COR Biosciences) to detect fluorescence or ImageQuant800 (AMERSHAM) to detect chemiluminescence. Rabbit anti-PSD-95 antibody (ab18258, Abcam), rabbit anti-Synaptophysin antibody (#4329, Cell Signaling Technology), rabbit anti-p-cofilin Ser3 (Cell Signaling #3313), rabbit anti-cofilin (Cell Signaling #5175), rabbit anti-Paralemmin-1 antisera^87^, rabbit anti-PLCβ1 antibody^88^ and mouse anti-β-Actin antibody (A1978, SIGMA) were used as primary antibodies. For fluorescence-based detection, Alexa Fluor 680-conjugated anti-rabbit IgG antibody (A-21076, Thermo Fisher Scientific) and IRDye800CW-conjugated anti-mouse IgG antibody (610-131-121, Rockland Immunochemicals) were used as secondary antibodies. For chemiluminescence detection, Immobilon Crescendo (Millipore WBLUR0500) or Chemi-Lumi One Super (Nacalai 02230-14) was used.

### Protein analysis by LC-MS/MS

The samples were denatured and reduced with 7 M guanidine-HCl / 1 M Tris (pH8.5) / 10 mM EDTA / 50 mM DTT at 37 °C for 2 h, followed by carboxymethylation with 100 mM sodium iodoacetate at 25 °C for 30 min. The protein was desalted using a PAGE Clean Up Kit (Nacalai tesque) according to the instruction manual. The desalted protein was dissolved using 10 mM Tris-HCl (pH8.0) / 0.03% n-Dodecyl-β-D-maltoside and digested with trypsin (TPCK-treated, Worthington Biochemical) for 12 h. The trypsinized protein fragments were applied to liquid chromatography (LC) (EASY-nLC 1200; Thermo Fisher Scientific, Odense, Denmark) coupled to a Q Exactive HF-X hybrid quadrupole-orbitrap mass spectrometer (Thermo Fisher Scientific, Inc., San Jose, CA, USA) with a nanospray ion source in positive mode. The peptides derived from protein fragments were separated on a NANO-HPLC capillary column C18 (0.075 mm ID x 100 mm length, 3 mm particle size, Nikkyo Technos, Tokyo, Japan). The mobile phase “A” was water with 0.1% formic acid and the mobile phase “B” was 80% acetonitrile with 0.1% formic acid. A linear gradient was used for 250 min at a flow rate of 300 nL/min: 0%-% B. The Q Exactive HF-X-MS was operating in the top-10 data-dependent scan mode. The parameters of Q Exactive HF-X were as follows: spray voltage, 2.0 kV; capillary temperature, 250 °C; mass range (m/z), 200-2000; normalized collision energy, 27%. Raw data were acquired with Xcalibur software.

### Protein identification

The MS and MS/MS data were searched against the NCBI-nr using Proteome Discoverer version 2.2 (Thermo Fisher Scientific) with the MASCOT search engine software version 2.6 (Matrix Science). The search parameters were as follows: enzyme, trypsin; static modifications, carboxymethyl (Cys); dynamic modifications, acetyl (Protein N-term), Gln-> pyro-Glu (N-term Gln), oxidation (Met); precursor mass tolerance, ± 15 ppm; fragment mass tolerance, ± 30 mmu; max. missed cleavages, 3. The proteins were identified when their false discovery rates (FDR) were less than 1%.

### Data analysis and bioinformatics

Statistical analyses were performed using R software. Heatmaps were generated in R using the heatmap.2 function in the gplots package. For k-means clustering, the number of clusters was assessed using the NbClust package^89^. One-way analysis of variance (ANOVA) was performed using the oneway.test function. Correction of p-value with Benjamini-Hochberg method was performed using the p.adjust function. Spearman’s rank correlation coefficient was calculated using the cor.test function. Fisher’s exact test was performed using the fisher.test function. Conversion of protein ID (GI number) into gene ID (Entrez Gene ID and official gene symbol) and identification of homologous gene in other species was performed using db2db of bioDBnet^90^. Genes not converted with db2db were individually checked with NCBI Protein (https://www.ncbi.nlm.nih.gov/protein) and NCBI Gene (https://www.ncbi.nlm.nih.gov/gene/). The lists of mouse Entrez Gene ID were used for the following enrichment analyses. Enrichment of proteins reported to be expressed on PSD was evaluated using DAVID version 6.8 (https://david.ncifcrf.gov/home.jsp)^91^ with no background list. GO analysis and pathway analysis were performed using Metascape (https://metascape.org)^48^ and SynGO (https://www.syngoportal.org/)^49^. For analysis with Metascape, proteins included in previously reported PSD proteome datasets (5,412 proteins in 16 datasets) were used as background to avoid overestimating synaptic protein enrichment. M musculus was selected as the Input species and M.musculus (for mouse data) or H.sapiens (for marmoset data) was chosen as the Analysis species. For other settings, default parameters were used. For analysis with SynGO, brain-expressed proteins were used as background. Enrichment of transcription factor binding on the genes was analyzed using ChIP-Atlas^68^. In Enrichment Analysis of ChIP-Atlas, we input the “Decrease A1” or “Increase A1” list, selected “ChIP: TFs and others” and “Neural” and used “Refseq coding genes” as a background. For other settings, default parameters were used. Enrichment of disease-related genes and transcription binding sites were analyzed using ToppCluster^92^. Canonical pathway analysis and network analysis were performed using Ingenuity Pathway Analysis (IPA) (QIAGEN). In k-means clustering, the optimal number of clusters was determined with the NbClust package of R software. For the mouse PSD proteome dataset, NbClust suggested 2 or 3 clusters as the best number. We adopted 3 clusters, and proteins were classified into three groups as described in the Results.

### Comparison with other datasets

Proteome datasets of PSD fraction^32,33,34–41,42–46^ were described in these articles. For transcriptome of developing mouse brain, we used two datasets; Dataset 1^28^ and Dataset 2^21^. Dataset 1 was downloaded from the NCBI website (https://www.ncbi.nlm.nih.gov//sra/?term=SRP055008) and converted to expression level (TPM) with RSEM ^93^. Transcriptome datasets of the developing human brain^24^ and macaque brain^25^ were downloaded from the PsychENCODE website (human transcriptome: http://development.psychencode.org/) (macaque transcriptome: http://evolution.psychencode.org/) (human histone acetylation: http://development.psychencode.org/#). Genes including zero value were eliminated for the analysis. Alignment of gene list was performed with Entrez Gene ID. To evaluate the change of gene expression before and after birth, mean mRNA abundance after the birth period was divided by that before birth. For transcriptome of human ASD patient brain, data described in^75^ was referred. A list of SFARI ASD genes (released on 01-11-2022) was downloaded from the SFARI website (https://gene.sfari.org/database/human-gene/). The ID of proteins or genes in each dataset was converted to Entrez Gene ID using bioDBnet^90^ and then compared with our proteome dataset.

## Acknowledgments

We thank Kaori Otsuki and Shungo Adachi for the helpful discussion and all the Takumi laboratory technical staff for their assistance in preparing the experimental reagent. This work was supported in part by KAKENHI (16H06316, 16H06463, 18K14830, 21H00202, 21H04813, 21K19351), Japan Society of Promotion of Science (JSPS) and Ministry of Education, Culture, Sports, Science, and Technology; Japan Agency for Medical Research and Development, JP21wm0425011; Intramural Research Grant (30-9) for Neurological and Psychiatric Disorders of NCNP; the Takeda Science Foundation; Smoking Research Foundation; Tokyo Biochemical Research Foundation; Kawano Masanori Memorial Public Interest Incorporated Foundation for Promotion of Pediatrics; Taiju Life Social Welfare Foundation; Naito Foundation; The Tokumori Yasumoto Memorial Trust for Researches on Tuberous Sclerosis Complex and Related Rare Neurological Diseases. T.K. was supported by Grant-in-Aid for JSPS Fellows (16J04376). N.K and H.O. were supported by the Brain/MINDS project of AMED (JP20dm0207001).

## Author contributions

T.K. designed the research, prepared the PSD samples, and performed most data analysis. T.S. and N.D. performed LC-MS/MS and label-free quantification. N.K. performed dissection of marmoset brain. M.W.K. and M.K. contributed antibodies. T.U. provided materials and analyzed Rho family signaling data. T.K. wrote and T.T. revised the manuscript. T.T., N.D., and H.O. supervised the work. All authors commented on the manuscript and approved the final version.

## Declaration of interests

The authors declare no competing interests.

## Notes

### Competing Interest Statement

The authors have declared no competing interest.

