## Supplemental text for "Developmental dynamics of the postsynaptic proteome to understand synaptic maturation and dysmaturation"

**Extended Data Fig. 1. Biochemical purification and analysis of the postsynaptic density (PSD) from mouse brain**

(a) Procedure of PSD purification by differential centrifugation and sucrose density gradient centrifugation (Modified from^1^. S and P indicate supernatant and pellet, respectively. (b) Confirmation of PSD purification. PSD fraction was prepared from adult (12-week-old) ICR mouse brains. S1 and PSD fractions are shown in (a) were subjected to SDS-PAGE followed by Western blotting. (c) Statistically enriched Gene Ontology terms and pathway terms in each cluster identified by SynGO.

**Extended Data Fig. 2. Overall workflow of protein classification**

In this study, 2,186 PSD proteins were analyzed (Fig. 1a, b). The proteins were classified into 3 clusters using k-means clustering (Fig. 1c). Differentially expressed (DE) proteins were identified using criteria: max fold change > 1.5, and P-value < 0.05. DE proteins in Cluster 1 were defined as “Decrease” proteins, whereas those in Cluster 3 were defined as “Increase” proteins (Fig. 2a, b). “Decrease” and “Increase” proteins were divided into two subgroups (A and B), respectively, according to the correlation between the change of protein abundance and that of mRNA abundance in mice (Extended Data Fig. 3b, c). The proteins in “Decrease A” and “Increase A” were further divided into two subgroups (1 and 2) based on the correlation with the human transcriptome (Fig. 4a). Genes encoding “Decrease” A1 and “Increase” A1 proteins, 42 and 72, respectively, were reported to be differentially expressed in patients with ASD (Fig. 5b).

**Extended Data Fig. 3. Correlation between the alteration of PSD protein abundance and mRNA abundance in mice during postnatal development**

(a) Spearman's rank correlation coefficient of changes in relative PSD protein abundance and mRNA abundance. For PSD protein, changes from 2-week-old to 12-week-old were used. For mRNA abundance, changes from indicated age to P110 or P180 in Dataset 1^2^ and Dataset 2^3^ are used, respectively. (b) Fold change of the abundance of DE PSD proteins in the developing mouse brain (12-week old vs. 2-week old) was plotted against mRNA abundance in the developmental mouse cortex. The number of proteins in indicated group and Spearman’s rank correlation coefficient are shown. (c) Extraction of proteins whose abundance is correlated with mRNA abundance. The “Decrease A” is a subgroup of “Decrease” whose mRNA is decreased after P4 in both transcriptome datasets. “Increase A” is a subgroup of “Increase” as well.

**Extended Data Fig. 4. Transcriptional profiles of PSD genes in developing macaque brain**

(a) Heatmap of the relative abundance of mRNAs that encode “Increase” and “Decrease” PSD proteins in cortical regions of the developing macaque brain^4^. (b) The plot of mean relative abundance of mRNAs that encode PSD proteins in the groups “Increase” A1 (left) or “Decrease A1” (right) (subgroups of “Increase A” or “Increase B” whose mRNA was increased or decreased in macaque cortex after birth, respectively). The 12-time points of age are described in (A). Red dotted lines indicate birth.

**Extended Data Fig. 5. Upstream transcription factors of genes encoding differentially expressed proteins on PSD**

(a) Proteins reported binding to “Decrease” A1 genes and “Increase” A1 genes in human neuronal cells (antigen class: TFs and others). The enrichment was analyzed with ChIP-Atlas^5^. (b) Enrichment of transcription factor binding sites analyzed with ToppCluster. (left) Top 5 enriched sites around “Decrease A1” genes (top) and “Increase A1” genes (bottom). (right) Representative transcription factor binding sites and downstream genes.

**Extended Data Fig. 6. Enrichment of genes related to neuropsychiatric disorders in genes encoding proteins on PSD**

(a) Heatmap of diseases significantly enriched in PSD protein clusters. Disease enrichment analysis was performed using ToppCluster. (b) Enrichment of proteins encoded by ASD risk genes^6^ (c) Enrichment of proteins encoded by SFARI ASD genes in PSD protein clusters. Asterisk indicates significant enrichment (*P < 0.05, **P < 0.01).

**Extended Data Fig. 7. PSD proteome analysis of ASD model mouse**

(a) PSD samples prepared from 3-week-old WT or *15q dup* mouse brains were subjected to LC-MS/MS to perform label-free quantification. PCA was performed using the relative abundance values of 1,935 proteins. (b) Identification of PSD protein clusters co-regulated in *15q dup* mice using k-means clustering based on the mean protein abundance. (left) The pie chart shows the number of proteins in each identified cluster. (right) Expression profile (average of relative abundance) of the proteins in each cluster. (c) Statistically enriched Gene Ontology terms and pathway terms in each cluster identified by Metascape. Top20 terms are displayed as a hierarchically clustered heatmap. The heatmap cells are colored by their P-values; grey cells indicate the lack of enrichment for that term in the corresponding protein list. (d) Enrichment of Cluster 1 or Cluster 3 proteins in 15q dup-3 and 15q dup-1 clusters.

**Extended Data Fig. 8. Biochemical purification and analysis of the PSD from marmoset brain**

(a) Purification of PSD from adult ICR mouse brain using a 3-step method (see methods). (b) Degradation of synaptic protein upon prolonged postmortem interval (PMI). Wild-type ICR mice were sacrificed. After the indicated duration at room temperature, whole brains were sampled. (c) Characteristics of the marmosets used in the present study. (d) Venn diagram illustrating the overlap of the analyzed proteins in the neocortex and cerebellum. (e and f) PSD samples prepared from 2-, 3-, 6-, and 24-month-old marmoset neocortex (e) and cerebellum (f) were subjected to LC-MS/MS to perform label-free quantification. The relative abundance of each protein was described using a heatmap. (g) Fold change of the abundance of PSD proteins in the developmental marmoset neocortex (24-month-old vs. 2-month-old) was plotted against the developmental marmoset cerebellum.

**Supplementary Table 1. Quantitative PSD proteomics of mouse brain in postnatal development**

**Supplementary Table 2 List of differentially expressed PSD proteins that have been reported to regulate the number, structure, or function of synapses**

**Supplementary Table 3. Quantitative PSD proteomics in ASD model mice (*15q dup*)**

**Supplementary Table 4. LC-MS/MS quantification and comparison of 1,960 PSD proteins in seven brain regions in adult marmoset**

**Supplementary Table 5. LC-MS/MS quantitation and comparison of 2,361 PSD proteins in marmoset neocortex**

**Supplementary Table 6. LC-MS/MS quantitation and comparison of 1,740 PSD proteins in marmoset cerebellum**
