## Supplemental Figures for "Developmental dynamics of the postsynaptic proteome to understand synaptic maturation and dysmaturation"

### Extended Data Fig. 1

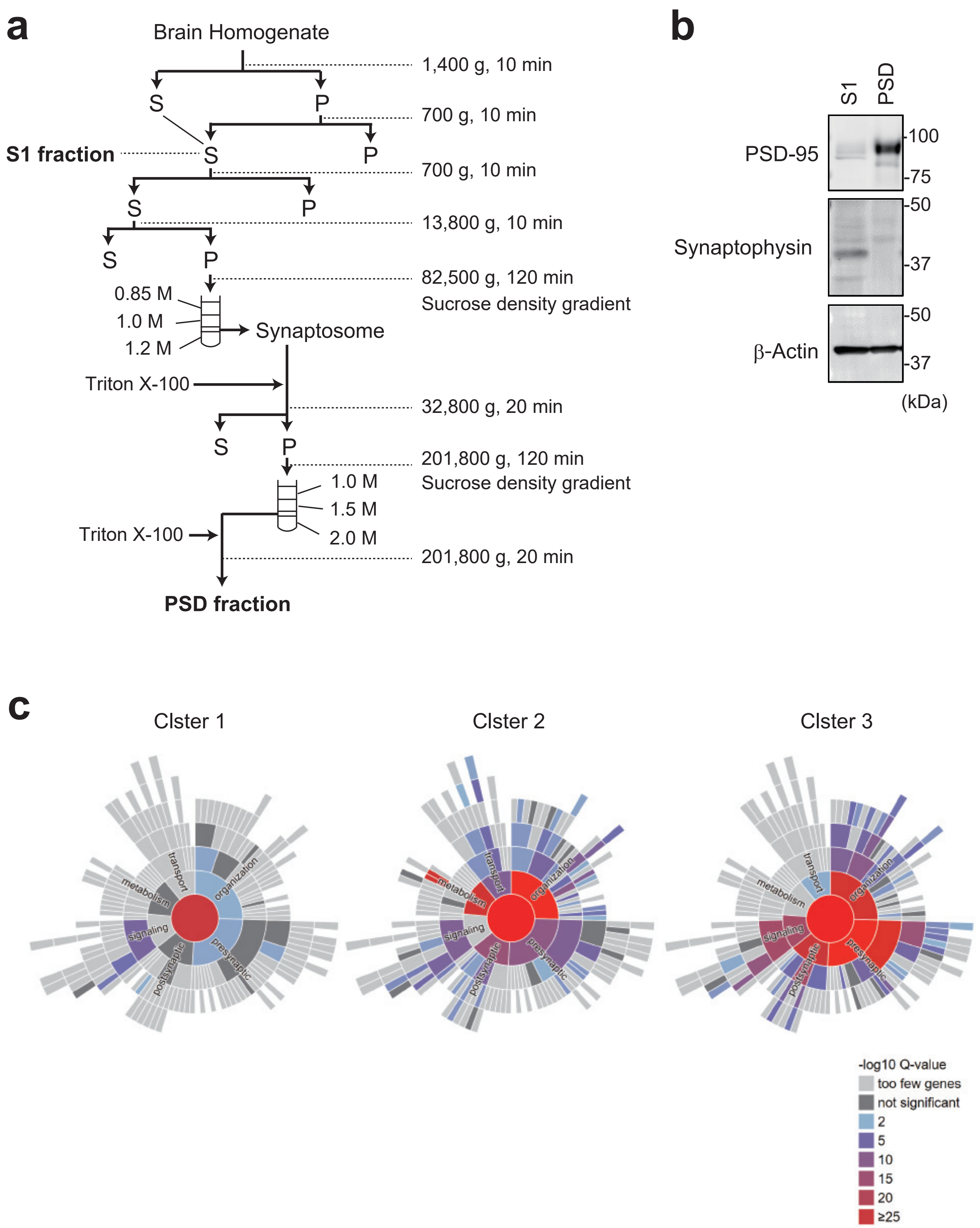

### Extended Data Fig.2

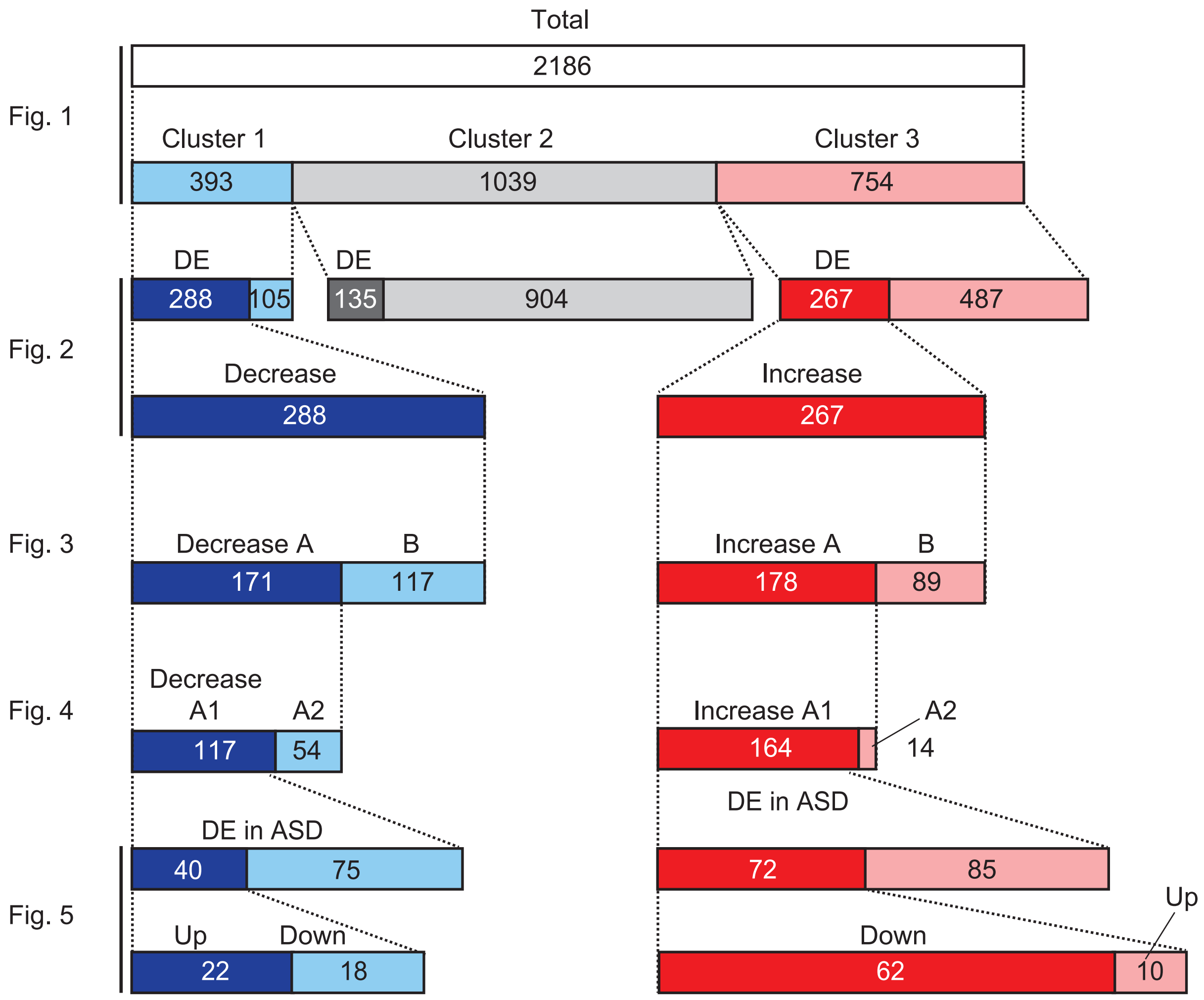

### Extended Data Fig. 3

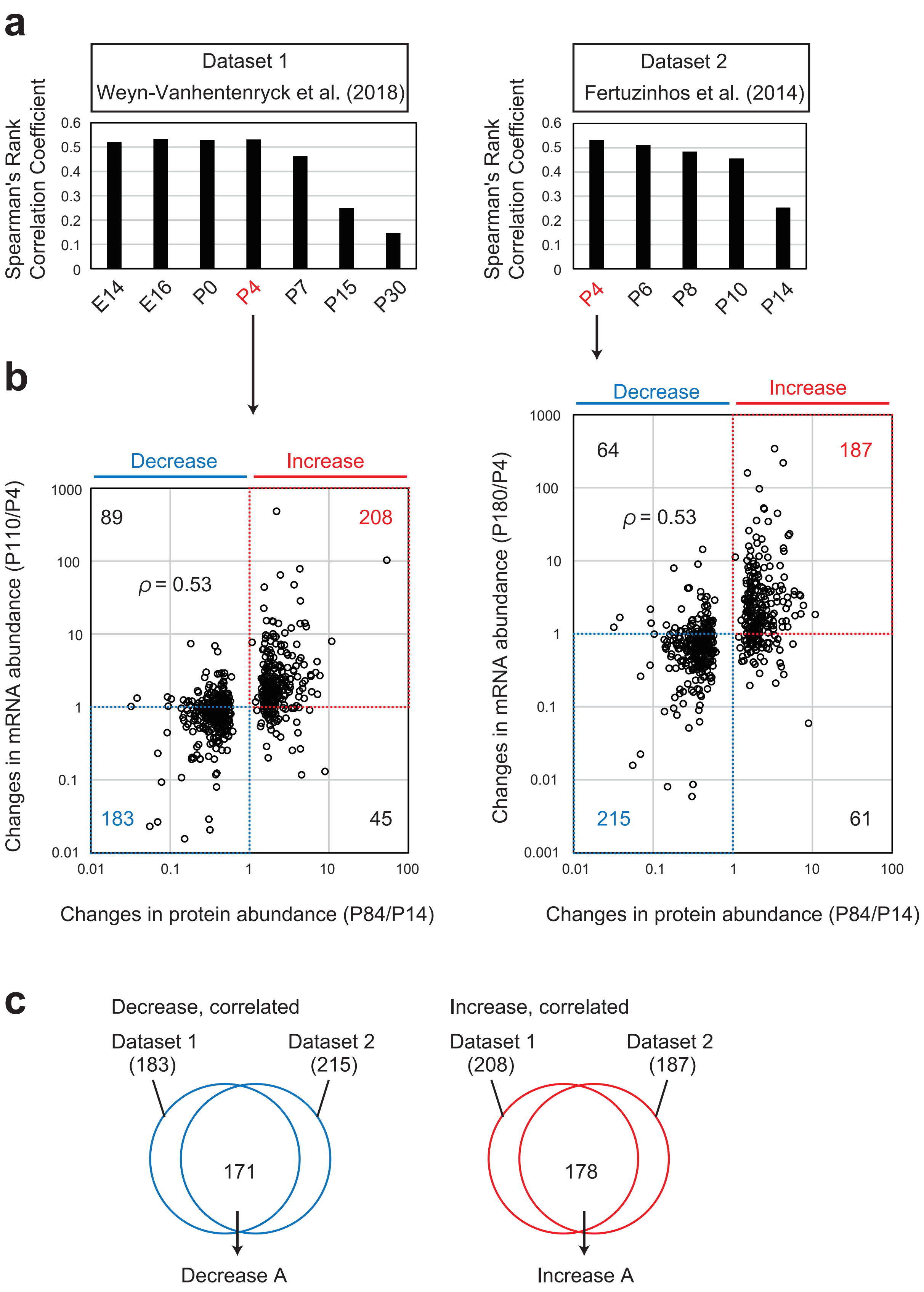

### Extended Data Fig. 4

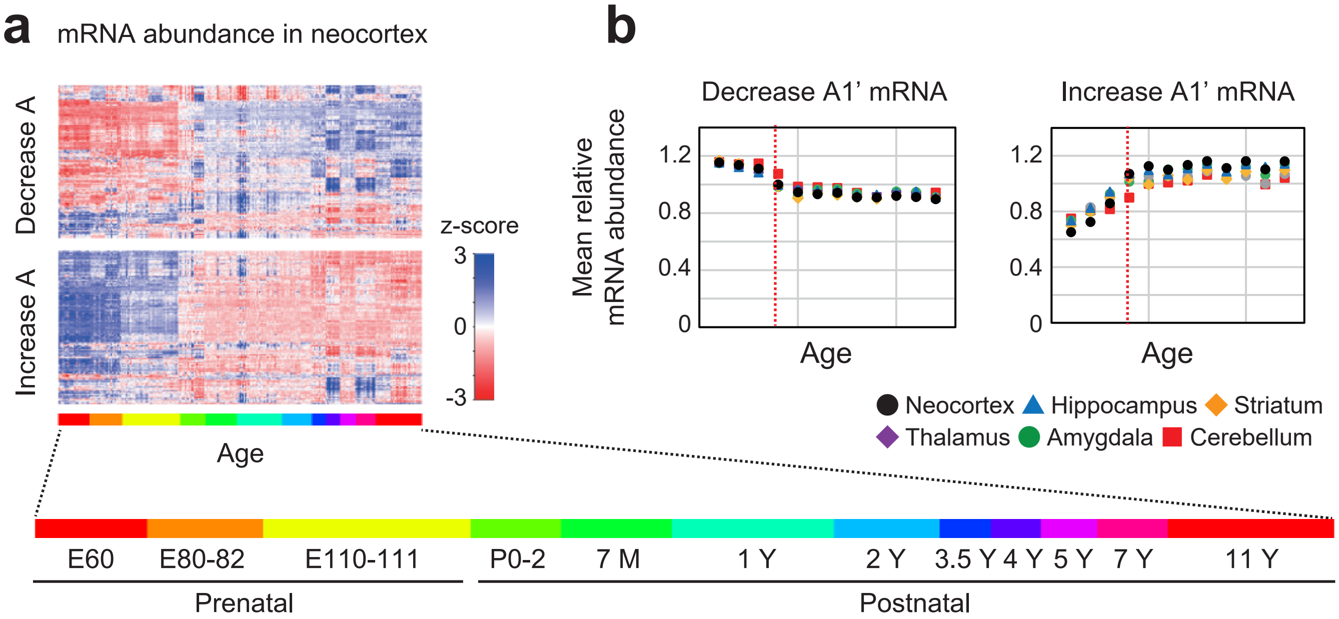

### Extended Data Fig. 5

**a** ChIP-Atlas

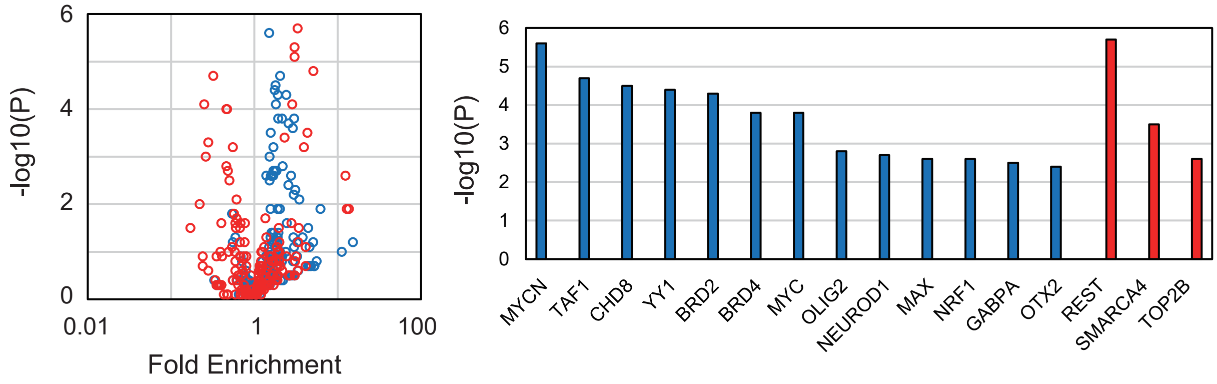


**b** Transcription factor binding site

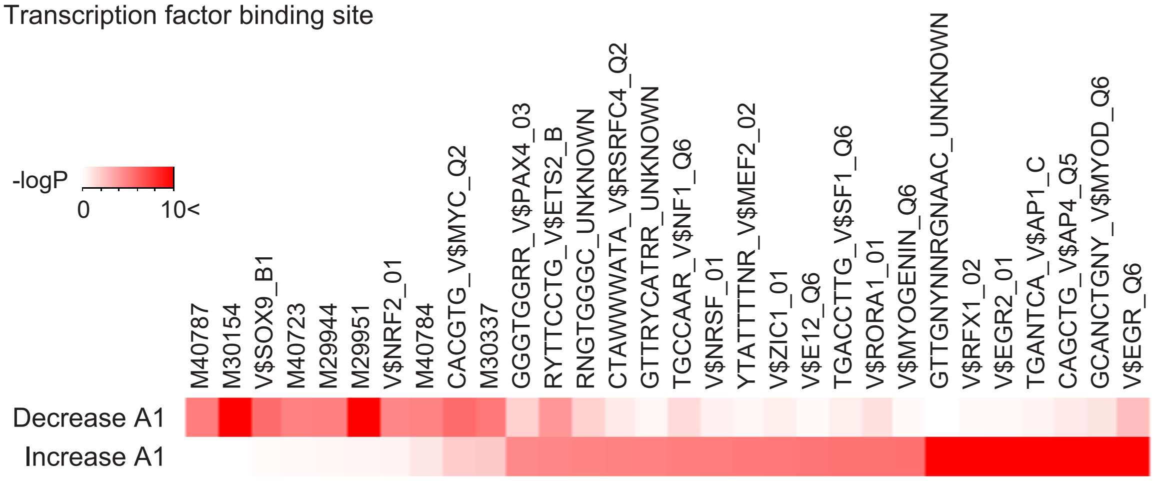

### Extended Data Fig. 6

**a**

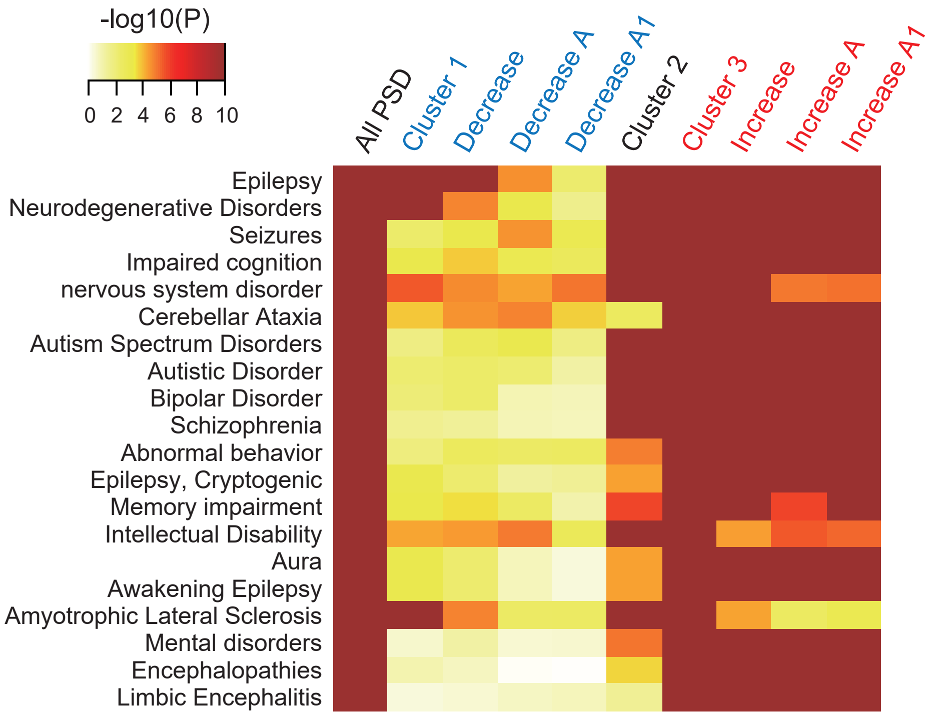


**b**

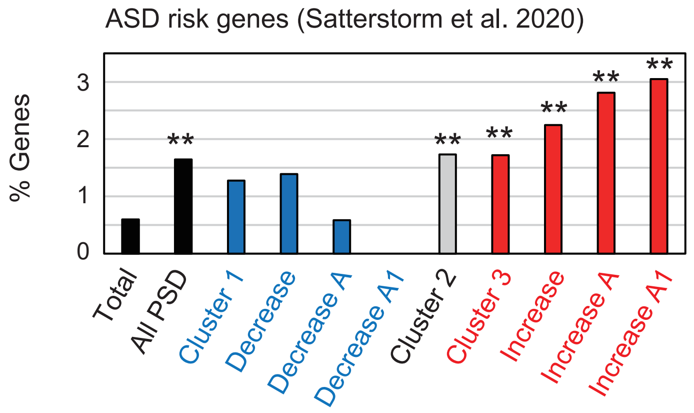


**c**

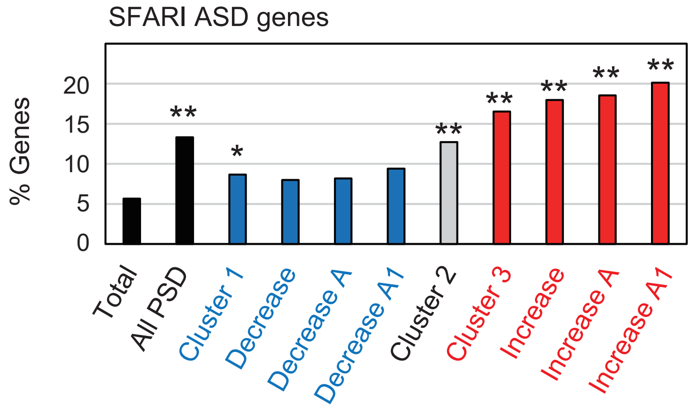

### Extended Data Fig. 7

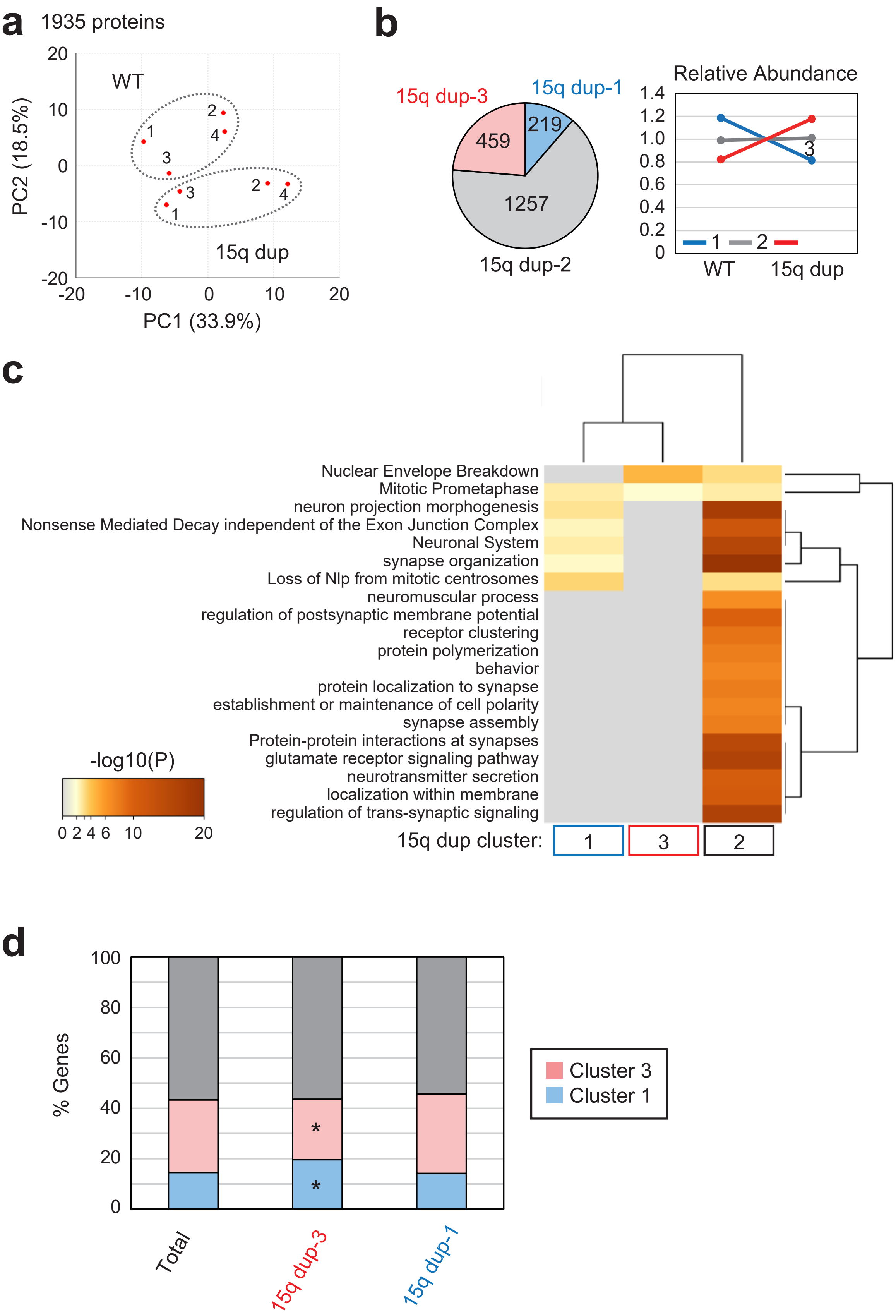

### Extended Data Fig. 8

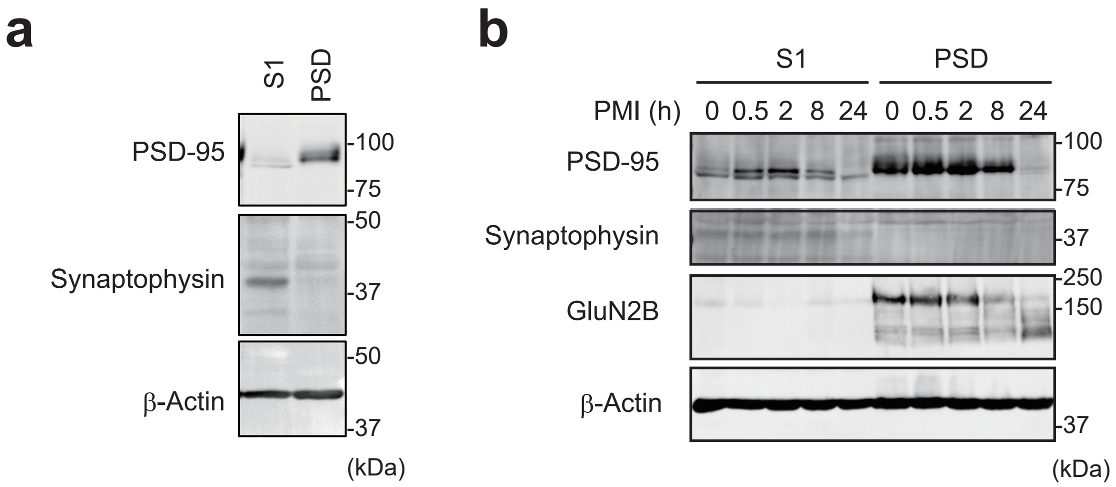


**c**

| Age | Postnatal day | Sex | Body Weight (g) | Animal ID |
|---|---|---|---|---|
| 2 M | 61 | Male | 106.3 | CM-1639M |
| 3 M | 91 | Male | 145.6 | CM-1601M |
| 6 M | 185 | Male | 216 | CM-1553M |
| 24 M | 730 | Male | 489 | CM-1128M |

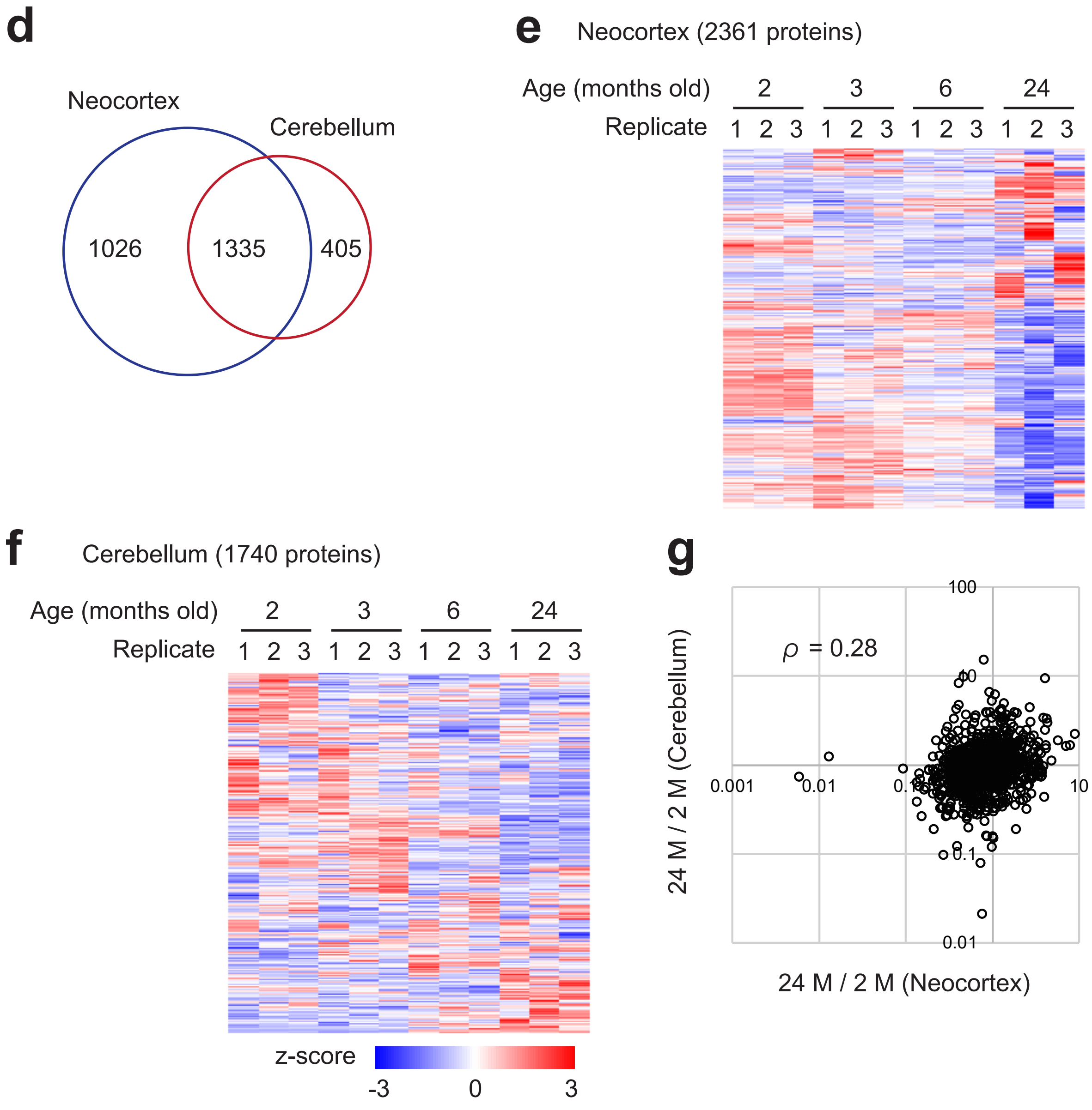
