## Supplemental Tables for "Developmental dynamics of the postsynaptic proteome to understand synaptic maturation and dysmaturation"

**Supplementary Table 1 LC-MS/MS quantitation and comparison of 2,186 PSD proteins in postnatal mouse brains**

| Description of protein | | Detection by LC-MS/MS | | | | | | | | | | |
|---|---|---|---|---|---|---|---|---|---|---|---|---|
| GI Number | Description | value: Cm | PEP Sc | coverage [% | # Peptides | # PSMs | ique Pept | # AAs | MW [kDa] | calc. pI | Mascot: M | Search Er |
| 39104555 | mKIAA0641 protein [Mus musculus] | 0 | 8.417 | 2 | 2 | 7 | 2 | 1415 | 149.5 | 4.55 | 53 | 2 |
| 148682136 | mCG146161, partial [Mus musculus] | 0 | 20.225 | 8 | 3 | 9 | 3 | 756 | 82.9 | 9.35 | 102 | 3 |
| 116089341 | abl interactor 1 isoform 1 [Mus musculus] | 0 | 94.157 | 37 | 14 | 161 | 11 | 481 | 52.3 | 7.64 | 1392 | 14 |
| 116875856 | Abelson tyrosine-protein kinase 2 isoform b [Mus musculus] | 0 | 22.486 | 5 | 5 | 27 | 5 | 1182 | 128.1 | 7.84 | 298 | 5 |
| 77404288 | acid-sensing ion channel 2 isoform MDEG1 [Mus musculus] | 0 | 26.744 | 18 | 7 | 74 | 6 | 512 | 57.7 | 5.19 | 303 | 7 |
| 568991060 | EDICTED: acid-sensing ion channel 1 isoform X2 [Mus muscul | 0 | 84.551 | 24 | 12 | 190 | 11 | 527 | 59.8 | 5.58 | 1590 | 12 |
| 755507681 | TED: microtubule-actin cross-linking factor 1 isoform X1 [Mus m | 0 | 2449.772 | 46 | 275 | 4018 | 18 | 7348 | 830.6 | 5.41 | 78496 | 275 |
| 18079339 | aconitate hydratase, mitochondrial precursor [Mus musculus] | 0 | 125.901 | 30 | 22 | 193 | 22 | 780 | 85.4 | 7.93 | 2806 | 22 |
| 66793429 | roxisomal acyl-coenzyme A oxidase 1 isoform 1 [Mus muscul | 0 | 135.674 | 47 | 20 | 102 | 20 | 661 | 74.7 | 8.12 | 1448 | 20 |
| 157951643 | alpha-actinin-2 [Mus musculus] | 0 | 398.247 | 54 | 44 | 750 | 30 | 894 | 103.8 | 5.45 | 15032 | 44 |
| 568971212 | grin and metalloproteinase domain-containing protein 11 isoforr | 0 | 16.161 | 6 | 3 | 10 | 3 | 854 | 92.8 | 7.74 | 65 | 3 |
| 568932692 | grin and metalloproteinase domain-containing protein 22 isoforr | 0 | 109.325 | 22 | 18 | 153 | 18 | 970 | 107.1 | 7.77 | 2609 | 18 |
| 755550837 | REDICTED: adenylate cyclase type 6 isoform X1 [Mus muscul | 0 | 31.545 | 7 | 7 | 27 | 3 | 1203 | 134 | 7.97 | 139 | 7 |
| 124430532 | adenylate cyclase type 8 isoform 1 [Mus musculus] | 0 | 11.337 | 2 | 2 | 3 | 2 | 1249 | 140 | 6.93 | 113 | 2 |
| 568933631 | PREDICTED: alpha-adducin isoform X1 [Mus musculus] | 0 | 281.949 | 51 | 26 | 520 | 2 | 795 | 87.1 | 5.87 | 13215 | 26 |
| 31560504 | beta-adducin isoform 1 [Mus musculus] | 0 | 246.426 | 48 | 28 | 356 | 2 | 725 | 80.6 | 6.21 | 5609 | 28 |
| 20806109 | poly [ADP-ribose] polymerase 1 [Mus musculus] | 0 | 98.446 | 22 | 18 | 77 | 18 | 1014 | 112.7 | 8.95 | 735 | 18 |
| 66792804 | alpha-1B adrenergic receptor [Mus musculus] | 0.002 | 5.053 | 5 | 2 | 3 | 2 | 515 | 56.5 | 9.17 | 0 | 2 |
| 568947203 | PREDICTED: aggrecan core protein isoform X1 [Mus musculus | 0 | 87.364 | 9 | 14 | 68 | 13 | 2170 | 225.9 | 4.28 | 547 | 14 |
| 10946936 | adenylate kinase isoenzyme 1 isoform 1 [Mus musculus] | 0 | 25.723 | 30 | 5 | 30 | 5 | 210 | 23.1 | 5.83 | 588 | 5 |
| 78711836 | A-kinase anchor protein 2 isoform 1 [Mus musculus] | 0 | 218.396 | 39 | 24 | 265 | 24 | 893 | 98.5 | 5.21 | 5453 | 24 |
| 12698457 | fatty aldehyde dehydrogenase variant form [Mus musculus] | 0 | 56.45 | 33 | 10 | 67 | 10 | 507 | 56.6 | 9.09 | 824 | 10 |
| 60687506 | fructose-bisphosphate aldolase C [Mus musculus] | 0 | 69.355 | 37 | 11 | 89 | 5 | 363 | 39.4 | 7.12 | 990 | 11 |
| 568949368 | PREDICTED: AMP deaminase 3 isoform X3 [Mus musculus] | 0 | 23.914 | 8 | 6 | 41 | 6 | 816 | 94 | 7.55 | 608 | 6 |
| 568953825 | PREDICTED: ankyrin-1 isoform X2 [Mus musculus] | 0 | 285.382 | 27 | 43 | 515 | 38 | 1932 | 212.1 | 6.65 | 7455 | 43 |
| 148747424 | ADP/ATP translocase 1 [Mus musculus] | 0 | 265.022 | 84 | 26 | 2126 | 2 | 298 | 32.9 | 9.72 | 35985 | 26 |
| 22094075 | ADP/ATP translocase 2 [Mus musculus] | 0 | 229.253 | 79 | 25 | 1768 | 12 | 298 | 32.9 | 9.73 | 29685 | 25 |
| 148701560 | annexin A6, isoform CRA_b, partial [Mus musculus] | 0 | 59.634 | 24 | 14 | 78 | 14 | 677 | 76.3 | 5.52 | 561 | 14 |
| 339895913 | AP-1 complex subunit beta-1 isoform 1 [Mus musculus] | 0 | 132.78 | 29 | 25 | 256 | 8 | 953 | 104.9 | 5.06 | 2588 | 25 |
| 6671561 | AP-2 complex subunit alpha-1 isoform a [Mus musculus] | 0 | 244.76 | 45 | 37 | 404 | 25 | 977 | 107.6 | 7.03 | 4684 | 37 |
| 50510693 | mKIAA0899 protein [Mus musculus] | 0 | 236.039 | 46 | 40 | 388 | 18 | 967 | 107.3 | 8.29 | 3782 | 40 |
| 50510363 | mKIAA0109 protein [Mus musculus] | 0 | 122.817 | 46 | 17 | 253 | 17 | 436 | 49.6 | 9.54 | 3361 | 17 |
| 568947213 | EDICTED: AP-3 complex subunit beta-2 isoform X1 [Mus musc | 0 | 65.115 | 13 | 12 | 71 | 8 | 1101 | 121.2 | 5.45 | 705 | 12 |
| 6671565 | AP-3 complex subunit delta-1 [Mus musculus] | 0 | 74.779 | 17 | 14 | 63 | 14 | 1199 | 135 | 7.37 | 469 | 14 |
| 12832639 | unnamed protein product [Mus musculus] | 0 | 9.397 | 15 | 3 | 20 | 3 | 193 | 21.7 | 5.39 | 63 | 3 |
| 39930317 | l beta A4 precursor protein-binding family A member 2 [Mus mu | 0 | 23.82 | 5 | 2 | 18 | 2 | 750 | 82.7 | 4.88 | 378 | 2 |
| 148684830 | ecursor protein-binding, family B, member 1, isoform CRA_c, pa | 0 | 15.541 | 6 | 3 | 24 | 3 | 725 | 78.9 | 5.08 | 148 | 3 |
| 569003029 | CTED: adenomatous polyposis coli protein isoform X1 [Mus mu | 0 | 246.573 | 21 | 42 | 282 | 42 | 2870 | 312.8 | 8.05 | 3617 | 42 |
| 148667988 | SPEG complex locus, isoform CRA_b, partial [Mus musculus] | 0 | 39.729 | 4 | 10 | 41 | 10 | 3293 | 357.1 | 8.21 | 246 | 10 |
| 163644329 | apolipoprotein E precursor [Mus musculus] | 0 | 26.477 | 20 | 5 | 42 | 5 | 311 | 35.8 | 5.68 | 278 | 5 |
| 148665902 | d beta (A4) precursor protein, isoform CRA_a, partial [Mus mu | 0 | 11.986 | 3 | 2 | 24 | 2 | 785 | 88.7 | 5.12 | 141 | 2 |
| 148669655 | aquaporin 4, isoform CRA_c [Mus musculus] | 0 | 13.06 | 10 | 3 | 10 | 3 | 355 | 38 | 6.65 | 55 | 3 |
| 755564427 | TED: serine/threonine-protein kinase A-Raf isoform X1 [Mus m | 0 | 38.229 | 16 | 9 | 93 | 7 | 641 | 71.5 | 9.04 | 959 | 9 |
| 9055166 | ctivity-regulated cytoskeleton-associated protein [Mus musculu | 0 | 93.541 | 40 | 13 | 194 | 4 | 396 | 45.3 | 5.39 | 2142 | 13 |
| 4502201 | ADP-ribosylation factor 1 [Homo sapiens] | 0 | 43.511 | 50 | 7 | 52 | 3 | 181 | 20.7 | 6.8 | 906 | 7 |
| 148704663 | mCG3164, partial [Mus musculus] | 0 | 17.809 | 25 | 4 | 30 | 4 | 191 | 21.5 | 8.38 | 401 | 4 |
| 388327124 | Of Geranylgeranylated Rhoa In Complex With Rhogdi In Its A | 0 | 23.023 | 32 | 5 | 15 | 3 | 195 | 21.9 | 6.37 | 96 | 5 |
| 6753116 | ho-related GTP-binding protein RhoN precursor [Mus musculu | 0.002 | 5.047 | 7 | 2 | 7 | 2 | 227 | 25.4 | 8.35 | 40 | 2 |
| 16307455 | Arpc1b protein [Mus musculus] | 0 | 18.796 | 14 | 3 | 18 | 2 | 376 | 41.5 | 8.35 | 190 | 3 |
| 148665105 | ne deleted in velo-cardio-facial syndrome, isoform CRA_b, par | 0 | 255.692 | 40 | 28 | 331 | 28 | 982 | 106.8 | 6.87 | 7958 | 28 |
| 755495397 | PREDICTED: astrotactin-1 isoform X1 [Mus musculus] | 0 | 108.522 | 19 | 16 | 125 | 16 | 1318 | 146.7 | 5.27 | 2335 | 16 |
| 21450277 | otassium-transporting ATPase subunit alpha-1 precursor [Mus | 0 | 220.754 | 30 | 25 | 338 | 10 | 1023 | 112.9 | 5.45 | 7637 | 25 |
| 6753138 | um/potassium-transporting ATPase subunit beta-1 [Mus musc | 0 | 50.944 | 32 | 9 | 74 | 9 | 304 | 35.2 | 8.65 | 1446 | 9 |
| 7242138 | um/potassium-transporting ATPase subunit beta-2 [Mus musc | 0 | 23.185 | 21 | 5 | 27 | 5 | 290 | 33.3 | 8.31 | 498 | 5 |
| 60360584 | mKIAA4195 protein [Mus musculus] | 0 | 85.412 | 20 | 15 | 101 | 15 | 1061 | 116.5 | 5.5 | 1462 | 15 |
| 148667069 | + transporting, plasma membrane 2, isoform CRA_c, partial [M | 0 | 201.389 | 27 | 25 | 184 | 16 | 1249 | 137.3 | 5.91 | 3528 | 25 |
| [illegible] | [illegible] | [illegible] | [illegible] | [illegible] | [illegible] | [illegible] | [illegible] | [illegible] | [illegible] | [illegible] | [illegible] | [illegible] |

| | | | | | | | | | | | | | | | |
|---|---|---|---|---|---|---|---|---|---|---|---|---|---|---|---|
| 144922638 | kinectin isoform 1 [Mus musculus] | 0 | 180.959 | 29 | 32 | 186 | 32 | 1327 | 152.4 | 5.9 | 1464 | 32 | 88410164.43 | 116643509 | 98668755.85 |
| 112293279 | neural cell adhesion molecule L1 precursor [Mus musculus] | 0 | 121.43 | 19 | 16 | 116 | 16 | 1259 | 140.8 | 5.96 | 2096 | 16 | 195819290.2 | 175077711.2 | 148788694.8 |
| 148237408 | uncharacterized protein LOC100037080 [Xenopus laevis] | 0 | 63.599 | 39 | 9 | 119 | 9 | 295 | 32.8 | 4.87 | 2028 | 9 | 181389725.5 | 215523563.6 | 212154083 |
| 6754508 | LIM and SH3 domain protein 1 [Mus musculus] | 0 | 45.484 | 32 | 8 | 100 | 7 | 263 | 30 | 7.05 | 851 | 8 | 38145896.33 | 26978063.12 | 28793643.92 |
| 257743039 | L-lactate dehydrogenase A chain isoform 2 [Mus musculus] | 0 | 38.471 | 24 | 7 | 55 | 6 | 361 | 39.7 | 8.35 | 669 | 7 | 26740544.12 | 26124404.64 | 20194798.79 |
| 938085832 | -lactate dehydrogenase B chain isoform Ldhbx [Mus musculus | 0 | 39.862 | 24 | 7 | 51 | 6 | 341 | 37.3 | 6.25 | 1053 | 7 | 19181985.74 | 21253542.49 | 16194656.3 |
| 568971910 | PREDICTED: DNA ligase 3 isoform X1 [Mus musculus] | 0 | 21.03 | 6 | 6 | 30 | 6 | 1066 | 118.2 | 9.06 | 278 | 6 | 14919057.72 | 15570015.17 | 13128990.56 |
| 18087805 | 40S ribosomal protein S2 [Mus musculus] | 0 | 75.739 | 48 | 13 | 234 | 3 | 293 | 31.2 | 10.24 | 2617 | 13 | 51534601.03 | 59674007.39 | 56430009.14 |
| 162287370 | prelamin-A/C isoform A precursor [Mus musculus] | 0 | 328.714 | 66 | 43 | 1018 | 41 | 665 | 74.2 | 6.98 | 20711 | 43 | 2618342836 | 2863160703 | 2954252306 |
| 188219589 | lamin-B1 [Mus musculus] | 0 | 383.781 | 68 | 45 | 1289 | 2 | 588 | 66.7 | 5.16 | 26518 | 45 | 53796775.77 | 82523537.55 | 63637286.62 |
| 113195686 | lamin-B2 [Mus musculus] | 0 | 266.775 | 57 | 33 | 818 | 29 | 596 | 67.3 | 5.5 | 18567 | 33 | 955060042.2 | 1136405915 | 1089526619 |
| 568908290 | leucine-rich repeat flightless-interacting protein 1 isoform X27 [l | 0 | 56.554 | 16 | 9 | 74 | 8 | 688 | 77.9 | 6.18 | 1640 | 9 | 26856109.69 | 27302165.74 | 24730914.61 |
| 6678768 | myristoylated alanine-rich C-kinase substrate [Mus musculus] | 0 | 35.42 | 21 | 3 | 42 | 3 | 309 | 29.6 | 4.34 | 1400 | 3 | 6970614.457 | 9899132.092 | 8129693.238 |
| 88014564 | itotic spindle assembly checkpoint protein MAD1 [Mus musculu | 0 | 14.564 | 6 | 4 | 23 | 4 | 717 | 83.5 | 5.66 | 49 | 4 | 4481856.879 | 4625805.464 | 4488635.668 |
| 568944963 | DICTED: myelin-associated glycoprotein isoform X1 [Mus musc | 0 | 8.436 | 5 | 3 | 17 | 3 | 627 | 69.3 | 5.1 | 38 | 3 | 1722685.461 | 1387746.82 | 1426345.935 |
| 568978894 | D: MAP/microtubule affinity-regulating kinase 3 isoform X1 [Mu | 0 | 197.831 | 41 | 24 | 323 | 16 | 806 | 90 | 9.47 | 5354 | 24 | 93921610.86 | 91364964.36 | 97629112.99 |
| 568966823 | megakaryocyte-associated tyrosine-protein kinase isoform X1 [ | 0 | 22.582 | 9 | 4 | 21 | 4 | 506 | 56.2 | 8.9 | 741 | 4 | 24444985 | 24825032.91 | 23596500.95 |
| 755498836 | PREDICTED: matrilin-4 isoform X1 [Mus musculus] | 0 | 44.98 | 17 | 7 | 38 | 7 | 636 | 70.2 | 5.54 | 765 | 7 | 12367009.4 | 17020346.43 | 19060079.1 |
| 37360062 | mKIAA0723 protein [Mus musculus] | 0 | 117.105 | 31 | 20 | 209 | 20 | 852 | 95.2 | 6.27 | 1424 | 20 | 148067464.3 | 145193267.7 | 116281613.1 |
| 568949747 | CTED: myc-associated zinc finger protein isoform X1 [Mus mu | 0 | 12.617 | 5 | 2 | 16 | 2 | 493 | 51.2 | 9.1 | 114 | 2 | 5530566.876 | 4141244.753 | 3204115.298 |
| 568953134 | ED: guanine nucleotide exchange factor DBS isoform X1 [Mus | 0 | 28.194 | 6 | 6 | 19 | 5 | 1211 | 135.6 | 6.04 | 167 | 6 | 2361557.723 | 5318987.884 | 4426835.971 |
| 219519515 | Mdm1 protein [Mus musculus] | 0 | 71.267 | 22 | 13 | 73 | 13 | 719 | 80.8 | 9.47 | 738 | 13 | 85296633.21 | 93012192.28 | 87200469.24 |
| 238637277 | 4F2 cell-surface antigen heavy chain isoform a [Mus musculus] | 0 | 20.805 | 11 | 5 | 25 | 5 | 565 | 62.2 | 5.5 | 66 | 5 | 12865553.63 | 14750438.14 | 11995431.03 |
| 126517469 | methyl-CpG-binding protein 2 isoform 1 [Mus musculus] | 0 | 82.072 | 33 | 12 | 124 | 12 | 501 | 53.5 | 9.88 | 1410 | 12 | 57793919.81 | 68156467.5 | 59547518.28 |
| 568983085 | TED: myocyte-specific enhancer factor 2C isoform X1 [Mus m | 0 | 19.447 | 11 | 4 | 17 | 3 | 474 | 51.2 | 8.03 | 90 | 4 | 11046457.77 | 11428410.44 | 10325028.85 |
| 1363258 | transcription factor MEF2D isoform 1a - mouse | 0 | 16.426 | 6 | 3 | 18 | 2 | 514 | 55 | 8.15 | 64 | 3 | 3623001.067 | 2889335.602 | 2487783.146 |
| 7330691 | MOBP170 [Mus musculus] | 0 | 47.844 | 34 | 6 | 169 | 6 | 170 | 19.2 | 11.14 | 2464 | 6 | 153962292.5 | 150553176 | 159484311.2 |
| 29747959 | myelin oligodendrocyte glycoprotein, partial [Mus musculus] | 0 | 35.722 | 30 | 8 | 129 | 8 | 274 | 31.1 | 8.54 | 2087 | 8 | 54422745.1 | 57169509.1 | 55456751.84 |
| 31982186 | malate dehydrogenase, mitochondrial precursor [Mus musculus | 0 | 34.734 | 29 | 8 | 53 | 8 | 338 | 35.6 | 8.68 | 332 | 8 | 154061898.7 | 193527344.3 | 206890496.8 |
| 148675904 | ehydrogenase 1, NAD (soluble), isoform CRA_c, partial [Mus m | 0 | 27.429 | 16 | 5 | 45 | 5 | 366 | 40 | 7.33 | 360 | 5 | 22357230.83 | 21163061.2 | 29849184.98 |
| 124053457 | multiple PDZ domain protein isoform 2 [Mus musculus] | 0 | 52.922 | 8 | 12 | 50 | 12 | 2055 | 218.6 | 5.02 | 274 | 12 | 6977424.653 | 11182153.26 | 10522110.19 |
| 6678924 | 55 kDa erythrocyte membrane protein [Mus musculus] | 0 | 14.664 | 8 | 3 | 10 | 3 | 466 | 52.2 | 7.2 | 59 | 3 | 2125813.528 | 1617569.095 | 1871169.491 |
| 887222486 | protein Mpv17 isoform 3 [Mus musculus] | 0 | 43.107 | 24 | 3 | 41 | 3 | 178 | 19.9 | 9.57 | 986 | 3 | 53268726.2 | 64795062.99 | 38854757.23 |
| 118200813 | rome c oxidase subunit II (mitochondrion) [Mus musculus dom | 0 | 15.776 | 16 | 3 | 24 | 3 | 227 | 26 | 4.78 | 300 | 3 | 24646682.94 | 22824992.76 | 25956460.68 |
| 291045426 | microtubule-associated protein 1A isoform 2 [Mus musculus] | 0 | 794.142 | 42 | 89 | 1716 | 87 | 2776 | 300 | 5 | 32677 | 89 | 1734850834 | 1730466618 | 1376158823 |
| 187951839 | Microtubule-associated protein 1B [Mus musculus] | 0 | 957.624 | 53 | 90 | 1879 | 88 | 2464 | 270.1 | 4.83 | 40808 | 90 | 3672136282 | 3534140044 | 3026149607 |
| 199023 | microtubule associated protein 2 [Mus musculus domesticus] | 0 | 1010.432 | 71 | 95 | 1990 | 4 | 1828 | 198.9 | 4.88 | 38701 | 95 | 98206410.51 | 92692880.02 | 99609841.13 |
| 148677083 | otubule-associated protein 4, isoform CRA_c, partial [Mus musc | 0 | 186.141 | 32 | 24 | 329 | 10 | 1157 | 120.6 | 5.03 | 4786 | 24 | 24149746.64 | 27441412.7 | 20340795.68 |
| 113204613 | microtubule-associated protein 6 isoform 1 [Mus musculus] | 0 | 460.684 | 74 | 55 | 1615 | 3 | 906 | 96.4 | 9.5 | 30699 | 55 | 59891368.36 | 47140742.29 | 45203576.7 |
| 568965077 | PREDICTED: ermin isoform X1 [Mus musculus] | 0 | 28.168 | 13 | 7 | 33 | 7 | 760 | 85.6 | 9.64 | 351 | 7 | 17867677.04 | 14756097.32 | 10095256.4 |
| 148702269 | tubule-associated protein tau, isoform CRA_g, partial [Mus mus | 0 | 185.701 | 60 | 15 | 489 | 3 | 374 | 39.1 | 9.5 | 7379 | 15 | 102987424.9 | 119716633.5 | 146673714.2 |
| 244790087 | myelin expression factor 2 isoform 1 [Mus musculus] | 0 | 140.467 | 39 | 19 | 188 | 19 | 591 | 63.3 | 8.87 | 3344 | 19 | 236200731 | 226746738.6 | 240607037.5 |
| 74180977 | unnamed protein product [Mus musculus] | 0 | 779.95 | 49 | 92 | 1396 | 75 | 2031 | 234 | 5.97 | 27152 | 92 | 1406621774 | 1392248659 | 1553004894 |
| 127139 | 4; AltName: Full=MLC1EMB; AltName: Full=Myosin light chain 1 | 0 | 15.802 | 19 | 3 | 13 | 3 | 193 | 21.1 | 5.03 | 208 | 3 | 12124401.33 | 2478297.995 | 3468835.54 |
| 954170005 | myosin light polypeptide 6 isoform a [Mus musculus] | 0 | 88.754 | 65 | 10 | 245 | 9 | 158 | 17.7 | 4.65 | 4370 | 10 | 711437206.8 | 680316747.6 | 817997067.7 |
| 568905921 | REDICTED: unconventional myosin-Ib isoform X1 [Mus musculu | 0 | 196.856 | 35 | 32 | 238 | 31 | 1138 | 132.2 | 9.33 | 2379 | 32 | 230162336.5 | 261587868.2 | 184424823.9 |

[illegible]

[illegible]

| | | | | | | | | | | | | | | | | | | | | | | | | | | | | | | | | | | | | | | | | | | | | | | | |
|---|---|---|---|---|---|---|---|---|---|---|---|---|---|---|---|---|---|---|---|---|---|---|---|---|---|---|---|---|---|---|---|---|---|---|---|---|---|---|---|---|---|---|---|---|---|---|---|
| 30519943 | g and assembly machinery component 50 homolog [Mus musc | 0 | 102.489 | 38 | 14 | 141 | 14 | 469 | 51.8 | 6.8 | 1724 | 14 | 81303453.93 | 110566586.5 | 105713181.9 | 132646339 | 51239366.45 | 66091770.93 | 121731464.7 | 91648269.64 | 107989393 | 40806426.67 | 126164018.7 | 100926531.2 | 35929049.72 | 38639615.7 | 47649174.12 | 35848982.77 | 19.58473 | 37.40468 | 39.37723 | 14.10704 | Samm50 | 68653 | SAMM50 | 25813 | 1.33 | 1.02 | 1.16 | 0.49 | 2.72 | 0.00595 | 0.02269 | 2 | Yes | | 12 |
| 148683864 | tripartite motif protein 37 [Mus musculus] | 0 | 36.76 | 8 | 6 | 39 | 6 | 1049 | 117.3 | 5.34 | 615 | 6 | 12725585.09 | 14745021.1 | 8423567.383 | 6320336.5 | 11037459.24 | 14464485.25 | 10773442.02 | 12973803.65 | 13064362.16 | 8481889.875 | 18732026.41 | 14085696.81 | 11358717.25 | 12224251 | 12934460.01 | 13469263.21 | 36.59322 | 14.11664 | 30.94228 | 7.314761 | Trim37 | 68729 | TRIM37 | 4591 | 0.86 | 1.01 | 1.11 | 1.02 | 1.29 | 0.7915 | 0.82157 | 3 | No | | 1 |
| 755544327 | TED: F-actin-uncapping protein LRRC16A isoform X1 [Mus mu | 0 | 43.295 | 10 | 10 | 26 | 10 | 1430 | 158 | 8.02 | 186 | 10 | 8411777.663 | 9525480.617 | 3528216.472 | 14338624.5 | 7539229.502 | 8482808.122 | 6095503.244 | 3245014.962 | 1786503.758 | 3767193.036 | 5091113.204 | 1439842.006 | 3098012.015 | 1574892.803 | 3009836.921 | 2668477.266 | 49.56679 | 36.04291 | 56.90289 | 27.05843 | Carmil1 | 68732 | CARMIL1 | 55604 | 1.71 | 1.21 | 0.58 | 0.50 | 3.46 | 0.05631 | 0.10807 | 1 | No | | 2 |
| 31559970 | transmembrane protein 214 [Mus musculus] | 0 | 17.586 | 7 | 4 | 6 | 4 | 687 | 76.4 | 9.29 | 75 | 4 | 5739912.116 | 5716843.859 | 6565903.735 | 7771729.375 | 4353333.784 | 4260818.763 | 4912215.303 | 6090859.509 | 2686236.263 | 2877315.37 | 3896359.692 | 4034868.374 | 3222027.157 | 3287143.218 | 3520059.628 | 3780096.036 | 14.98714 | 17.16344 | 20.45966 | 7.334426 | Tmem214 | 68796 | TMEM214 | 54867 | 1.42 | 1.08 | 0.74 | 0.76 | 1.91 | 0.00655 | 0.02395 | 1 | Yes | Decrease B | 3 |
| 343531216 | nexilin alpha [Mus musculus] | 0 | 9.008 | 4 | 3 | 23 | 3 | 671 | 80 | 5.25 | 355 | 3 | 3440867.22 | 2321305.914 | 4258503.362 | 4646315.25 | 2969721.076 | 4064251.476 | 4853601.676 | 5385404.322 | 4202035.813 | 3973926.158 | 1836749.671 | 5888066.628 | 3218217.167 | 3042623.661 | 1830276.931 | 4643117.867 | 28.03844 | 24.31898 | 41.80108 | 36.19244 | Nexn | 68810 | NEXN | 91624 | 0.97 | 1.14 | 1.05 | 0.84 | 1.36 | 0.6267 | 0.68191 | 2 | No | | 3 |
| 30794432 | charged multivesicular body protein 2b [Mus musculus] | 0 | 18.285 | 28 | 7 | 64 | 6 | 213 | 23.9 | 8.78 | 650 | 7 | 53190903.03 | 39836262.05 | 41520953.13 | 52244715.5 | 60038270.07 | 43067605.48 | 44097969.93 | 40432245.79 | 26533364.24 | 23500772.53 | 31940413.94 | 33382525.45 | 34480086.71 | 29335192.06 | 38320636.66 | 32071727.48 | 14.98013 | 18.9471 | 16.02505 | 11.35888 | Chmp2b | 68942 | CHMP2B | 25978 | 1.20 | 1.20 | 0.74 | 0.86 | 1.63 | 0.01813 | 0.04887 | 2 | Yes | | 3 |
| 114150032 | l synthase; Short=CDP-DAG synthase; AltName: Full=Mitocho | 0 | 22.329 | 17 | 4 | 23 | 4 | 337 | 37.8 | 8.98 | 141 | 4 | 5571552.035 | 4365829.938 | 5446451.109 | 22344270.63 | 20769561.87 | 5433174.714 | 2758413.08 | 19833529.85 | 3887990.417 | 13951088.41 | 2398470.325 | 4455887.907 | 2273881.099 | 2685852.874 | 3194012.794 | 4649676.493 | 91.44553 | 77.28423 | 85.17465 | 32.38519 | Tamm41 | 68971 | TAMM41 | 132001 | 1.22 | 1.57 | 0.80 | 0.41 | 3.81 | 0.2655 | 0.3467 | 1 | No | | 4 |
| 12840819 | unnamed protein product [Mus musculus] | 0 | 11.973 | 27 | 2 | 19 | 2 | 102 | 11.8 | 9.31 | 70 | 2 | 22146833.4 | 21107760.4 | 19702233.92 | 21132522.13 | 15059151.98 | 12583532.32 | 11191731.39 | 11491366.31 | 7197977.973 | 7598024.413 | 11602836.31 | 7592903.732 | 8510708.876 | 8689287.517 | 9809492.56 | 7057496.928 | 4.777889 | 13.9631 | 24.45769 | 13.26747 | Spcs1 | 69019 | SPCS1 | 28972 | 1.66 | 0.99 | 0.67 | 0.67 | 2.47 | 1.29E-05 | 0.00101 | 1 | Yes | Decrease B | 1 |
| 19526878 | pyrroline-5-carboxylate reductase 2 [Mus musculus] | 0 | 66.886 | 43 | 10 | 87 | 10 | 320 | 33.6 | 7.77 | 785 | 10 | 130158583 | 63591563.42 | 121340101.3 | 128723654 | 110996919 | 105133250.7 | 71859835.61 | 121464416.8 | 60082963.24 | 41748292.29 | 58327252.69 | 59030921.84 | 47664816.99 | 45855312.81 | 52684834.08 | 56132183.68 | 28.66977 | 20.93359 | 15.93004 | 9.27795 | Pycr2 | 69051 | PYCR2 | 29920 | 1.39 | 1.28 | 0.69 | 0.63 | 2.19 | 0.01146 | 0.03527 | 1 | Yes | Decrease B | 5 |
| 12841194 | unnamed protein product [Mus musculus] | 0 | 7.479 | 4 | 2 | 13 | 2 | 433 | 48.2 | 9.61 | 164 | 2 | 13057620.13 | 12254954.22 | 9638461.413 | 9721218 | 10198584.33 | 6462935.586 | 7451570.709 | 7026073.075 | 4582004.968 | 3358792.155 | 4085713.137 | 4155291.438 | 3540733.983 | 3941621.603 | 4802583.327 | 3665628.234 | 15.66738 | 21.31536 | 12.54827 | 14.25721 | Oxa1l | 69089 | OXA1L | 5018 | 1.66 | 1.15 | 0.60 | 0.59 | 2.80 | 0.00107 | 0.0073 | 1 | Yes | Decrease A1 | 2 |
| 194319965 | Chain A, Crystal Structure Of Mouse Vps26b | 0 | 17.384 | 18 | 5 | 25 | 5 | 340 | 39.4 | 7.53 | 67 | 5 | 6722168.899 | 7480835.72 | 4793012.67 | 6187963.625 | 4675286.704 | 5234343.642 | 2496793.381 | 5502289.455 | 5700363.332 | 4762687.953 | 6704169.675 | 4593124.579 | 2419867.912 | 5257271.121 | 4556719.469 | 4867286.255 | 18.00732 | 30.47608 | 17.89102 | 29.69898 | Vps26b | 69091 | VPS26B | 112936 | 1.23 | 0.87 | 1.06 | 0.83 | 1.47 | 0.2077 | 0.28682 | 2 | No | | 5 |
| 148681362 | mCG140375 [Mus musculus] | 0 | 88.097 | 5 | 20 | 67 | 20 | 5189 | 573.6 | 6.2 | 491 | 20 | 23176823.01 | 30247303.11 | 23446277.72 | 22303480.13 | 14906331.86 | 16544017.8 | 18774702.86 | 18071157.32 | 13147591.75 | 7129615.083 | 19812123.26 | 7786201.434 | 17070052.4 | 13517263.97 | 17056766.34 | 9707218.6 | 14.79608 | 10.06876 | 49.1501 | 24.48509 | Ubr4 | 69116 | UBR4 | 23352 | 1.45 | 1.00 | 0.70 | 0.84 | 2.07 | 0.02761 | 0.06676 | 1 | No | | 8 |
| 18017596 | sorting nexin-4 [Mus musculus] | 0 | 14.883 | 8 | 3 | 10 | 3 | 450 | 51.7 | 5.8 | 132 | 3 | 861358.0679 | 3258345.3 | 2612417.358 | 1310948.625 | 2063629.572 | 2786174.923 | 6133374.477 | 2910743.778 | 3693537.111 | 8094567.677 | 8087821.501 | 5899778.196 | 6077032.672 | 9132757.716 | 10449022.13 | 6796175.546 | 55.44884 | 52.17131 | 32.65987 | 25.03281 | Snx4 | 69150 | SNX4 | 8723 | 0.40 | 0.69 | 1.29 | 1.62 | 4.04 | 0.00891 | 0.02936 | 3 | Yes | Increase B | 4 |
| 33468999 | hol O-methyltransferase domain-containing protein 1 [Mus mus | 0 | 14.025 | 24 | 4 | 25 | 4 | 262 | 28.9 | 8.32 | 132 | 4 | 19579785.73 | 15438498.73 | 14173855.06 | 9666743.25 | 10156349.16 | 9298612.37 | 9939170.063 | 7322206.366 | 8703616.923 | 12342498.95 | 13275771.81 | 8674376.116 | 6735733.873 | 13048191.74 | 12120593.47 | 8189111.26 | 27.73514 | 14.05757 | 22.4123 | 30.3269 | Comtd1 | 69156 | COMTD1 | 118881 | 1.32 | 0.82 | 0.96 | 0.90 | 1.60 | 0.2138 | 0.29279 | 2 | No | | 4 |
| 50510421 | mKIAA0227 protein [Mus musculus] | 0 | 16.488 | 20 | 3 | 27 | 3 | 271 | 29.4 | 9.51 | 195 | 3 | 6812610.002 | 5698769.292 | 6407276.263 | 5964605.625 | 7566036.751 | 6917745.644 | 3293247.218 | 7004151.362 | 4729473.658 | 1195154.815 | 1666856.962 | 4323638.417 | 1505751.222 | 3806527.903 | 2571512.255 | 2726206.035 | 7.892426 | 31.57123 | 60.60133 | 35.49174 | Ttc9 | 69480 | TTC9 | 23508 | 1.38 | 1.37 | 0.66 | 0.59 | 2.35 | 0.004 | 0.01748 | 1 | Yes | Decrease A1 | 1 |
| 19223941 | mitotic phosphoprotein 44 [Mus musculus] | 0 | 23.575 | 18 | 4 | 17 | 4 | 325 | 34.8 | 9.25 | 324 | 4 | 6992802.47 | 4155298.269 | 13792410.96 | 18335066.25 | 2807340.956 | 6012739.892 | 6970830.832 | 10272953.07 | 3346469.327 | 1579384.508 | 3474120.437 | 2514335.059 | 1039951.239 | 1979354.465 | 1157864.518 | 3431979.432 | 59.51504 | 47.16025 | 32.11887 | 57.93403 | Nup35 | 69482 | NUP35 | 129401 | 1.97 | 1.19 | 0.50 | 0.35 | 5.69 | 0.06924 | 0.12596 | 1 | No | | 1 |
| 148679826 | RIKEN cDNA 2310022B05 [Mus musculus] | 0 | 30.258 | 25 | 7 | 75 | 7 | 322 | 35.3 | 5.24 | 396 | 7 | 28797538.87 | 25295469.04 | 24020219.39 | 27291302 | 21966753.91 | 25924160.86 | 23547973.57 | 18032777.11 | 22719999.98 | 26882779.82 | 21506115.74 | 22037348.24 | 15009920.29 | 16774104.78 | 14616905.63 | 21535694.07 | 8.025242 | 14.8262 | 10.11591 | 18.70014 | 310022B05Rik | 69551 | C1orf198 | 84886 | 1.19 | 1.01 | 1.05 | 0.76 | 1.55 | 0.01998 | 0.05224 | 2 | No | | 4 |
| 110625761 | AFG3-like protein 2 [Mus musculus] | 0 | 40.759 | 14 | 11 | 50 | 10 | 802 | 89.5 | 8.6 | 414 | 11 | 29627143.14 | 25119776.06 | 14357054.89 | 18015945.34 | 28770577.15 | 16987908.76 | 11446000.73 | 19221201.21 | 26545304.72 | 20635967.7 | 22136331.63 | 12830278.95 | 27628674.94 | 22722723.28 | 24780425.47 | 20292590.3 | 31.58828 | 37.81178 | 27.83883 | 13.04852 | Afg3l2 | 69597 | AFG3L2 | 10939 | 1.02 | 0.90 | 0.96 | 1.12 | 1.25 | 0.6418 | 0.69454 | 2 | No | | 11 |
| 166295212 | isabled homolog 2-interacting protein isoform 2 [Mus musculu | 0 | 243.931 | 41 | 42 | 415 | 35 | 1189 | 131.6 | 8.72 | 4498 | 42 | 221005090.5 | 244068433.5 | 228131629.5 | 203398932.1 | 197969830.3 | 219819748.6 | 205658261.6 | 219856342.3 | 208439953.9 | 194505445.3 | 226008840.6 | 201472435.3 | 201188198.8 | 185290495.4 | 214363774 | 182812519.7 | 7.52349 | 5.15555 | 6.513704 | 7.529005 | Dab2ip | 69601 | DAB2IP | 153090 | 1.07 | 1.01 | 0.99 | 0.93 | 1.14 | 0.2377 | 0.31878 | 2 | No | | 8 |
| 148693600 | ne nucleotide exchange factor (GEF) 12, isoform CRA_c [Mus | 0 | 65.06 | 13 | 17 | 88 | 17 | 1554 | 173.6 | 5.81 | 667 | 17 | 20747300.53 | 25753554.95 | 20036249.06 | 17355524.94 | 24247050.88 | 24938408.75 | 22186171.24 | 19004503.72 | 15958596.95 | 18772717.06 | 26374932.11 | 16474236.83 | 22223950.16 | 27950479.25 | 27700394.38 | 19501691.38 | 16.71483 | 11.78766 | 24.80686 | 17.13726 | Arhgef12 | 69632 | ARHGEF12 | 23365 | 0.96 | 1.04 | 0.89 | 1.12 | 1.26 | 0.5245 | 0.59584 | 2 | No | | 5 |
| 7414392 | death associated protein kinase [Mus musculus] | 0 | 35.618 | 8 | 9 | 49 | 9 | 1430 | 159.7 | 6.93 | 366 | 9 | 31947842.39 | 35881835.2 | 42297307.17 | 40491049.25 | 21497728.31 | 24044312.41 | 39737153.23 | 33762334.33 | 20382124.91 | 18212982.24 | 22230987.61 | 22073086.17 | 13887345.89 | 14183050.64 | 17140906.65 | 17760243.57 | 12.39116 | 28.54475 | 9.03265 | 12.65127 | Dapk1 | 69635 | DAPK1 | 1612 | 1.45 | 1.15 | 0.80 | 0.61 | 2.39 | 0.00101 | 0.0072 | 1 | Yes | Decrease B | 4 |
| 568980339 | ICTED: WD repeat-containing protein 20 isoform X1 [Mus mus | 0 | 27.35 | 14 | 6 | 24 | 6 | 584 | 64.6 | 7.75 | 77 | 6 | 14739711.32 | 12953438.16 | 16391155.28 | 11075322.38 | 14999160.48 | 15638067.8 | 21909298.54 | 23763776.55 | 12338834.42 | 21158706.39 | 29026517.73 | 17702397.44 | 17664291.49 | 14759333.35 | 20977488.68 | 18507438.7 | 16.60897 | 23.13565 | 34.87511 | 14.26732 | Wdr20 | 69641 | WDR20 | 91833 | 0.78 | 1.08 | 1.13 | 1.01 | 1.45 | 0.1471 | 0.22116 | 3 | No | | 2 |
| 568961999 | CTED: muscular LMNA-interacting protein isoform X1 [Mus mu | 0 | 123.567 | 23 | 15 | 166 | 15 | 1036 | 110.8 | 8.35 | 1745 | 15 | 82381080.73 | 61165960.82 | 65403847.6 | 81574823 | 68481587.51 | 80810099.82 | 57073056.26 | 70515111.55 | 89132092.96 | 74135694.05 | 42795952.79 | 84494011.26 | 63006515.86 | 42107309.68 | 37206693.13 | 60721736.47 | 15.05572 | 14.05868 | 28.7171 | 25.62999 | Mlip | 69642 | MLIP | 90523 | 1.10 | 1.04 | 1.10 | 0.77 | 1.43 | 0.1879 | 0.26551 | 2 | No | | 5 |
| 299522842 | dynactin subunit 2 isoform 1 [Mus musculus] | 0 | 66.023 | 28 | 8 | 74 | 8 | 407 | 44.7 | 5.21 | 1590 | 8 | 38788783.61 | 76068129.62 | 49455446.46 | 47931739.5 | 38914168.29 | 40318505.49 | 44525274.31 | 44777321.15 | 47440492.45 | 37743639.54 | 49610192.71 | 38979171.94 | 51166091.09 | 60200467.75 | 48738424.98 | 44684166.43 | 30.23891 | 7.036436 | 13.70986 | 12.83421 | Dctn2 | 69654 | DCTN2 | 10540 | 1.12 | 0.89 | 0.92 | 1.08 | 1.26 | 0.2025 | 0.28124 | 2 | No | | 13 |
| 74152618 | unnamed protein product [Mus musculus] | 0 | 10.76 | 9 | 2 | 8 | 2 | 268 | 28.1 | 6.55 | 208 | 2 | 9957902.977 | 7459788.75 | 8502447.868 | 4562873 | 6511854.581 | 6485252.893 | 6012028.661 | 6811568.614 | 7816173.275 | 7941244.018 | 8637375.715 | 9767001.911 | 9322990.548 | 7283566.807 | 9188514.137 | 12304186.59 | 29.93844 | 5.118528 | 10.46727 | 21.7726 | Emc10 | 69683 | EMC10 | 284361 | 0.95 | 0.80 | 1.06 | 1.19 | 1.48 | 0.02413 | 0.06015 | 3 | No | | 3 |
| 568934506 | PREDICTED: CAD protein isoform X1 [Mus musculus] | 0 | 64.93 | 9 | 17 | 79 | 17 | 2321 | 252.9 | 6.6 | 598 | 17 | 34667979.71 | 39746498.11 | 41631631.12 | 32658423.5 | 28828284.54 | 27037748.76 | 37582548.69 | 24099884.6 | 21463117.91 | 27942181.96 | 37556079.49 | 28728701.53 | 27647936.57 | 29855852.52 | 42154123.53 | 27696927.17 | 11.32303 | 19.73964 | 22.86317 | 21.83999 | Cad | 69719 | CAD | 790 | 1.17 | 0.92 | 0.91 | 1.00 | 1.29 | 0.2128 | 0.29165 | 2 | No | | 4 |
| 29243896 | ubiquitin carboxyl-terminal hydrolase 46 [Mus musculus] | 0 | 31.631 | 19 | 6 | 54 | 4 | 366 | 42.4 | 6.83 | 873 | 6 | 1199590.479 | 9512104.154 | 7488580.37 | 2886469.875 | 12117586.65 | 11945782.25 | 11337711.39 | 11584699.89 | 7137740.773 | 8511912.054 | 11192751.47 | 9138490.454 | 9089056.602 | 9485807.698 | 11325616.52 | 10037513.28 | 73.60382 | 2.992594 | 18.74954 | 9.765209 | Usp46 | 69727 | USP46 | 64854 | 0.59 | 1.31 | 1.00 | 1.11 | 2.23 | 0.01858 | 0.04977 | 3 | Yes | Increase B | 5 |
| 31981560 | stromal membrane-associated protein 2 [Mus musculus] | 0 | 8.334 | 7 | 3 | 9 | 2 | 428 | 46.5 | 8.87 | 22 | 3 | 5076437.855 | 6740448.393 | 4606793.361 | 5786633.938 | 4747368.018 | 5126894.1 | 5268590.433 | 4744795.531 | 5911562.956 | 6460559.49 | 6422967.689 | 6046935.206 | 6931759.018 | 6345020.972 | 6982513.453 | 6033334.497 | 16.72422 | 5.372332 | 4.397763 | 7.024691 | Smap2 | 69780 | SMAP2 | 64744 | 0.95 | 0.85 | 1.07 | 1.13 | 1.32 | 0.00234 | 0.01196 | 3 | No | | 3 |
| 168480117 | mitochondrial folate transporter/carrier [Mus musculus] | 0 | 12.965 | 14 | 3 | 4 | 3 | 316 | 35 | 9.47 | 26 | 3 | 14100238.71 | 10751438.57 | 9924477.498 | 11814346 | 9526309.099 | 10610684.94 | 8533020.766 | 9668146.526 | 7754301.503 | 7177356.146 | 7052019.814 | 7722673.829 | 6524639.389 | 6293844.575 | 5871956.665 | 6235499.961 | 15.52952 | 8.871613 | 4.901264 | 4.336845 | Slc25a32 | 69906 | SLC25A32 | 81034 | 1.34 | 1.10 | 0.85 | 0.71 | 1.87 | 0.00063 | 0.00533 | 2 | Yes | | 0 |
| 37537518 | acylglycerol kinase, mitochondrial precursor [Mus musculus] | 0 | 89.778 | 51 | 15 | 133 | 15 | 421 | 46.9 | 8.4 | 1090 | 15 | 227765939.4 | 232291479.5 | 153277743.9 | 161411816.9 | 230662303.2 | 172587983.4 | 131408137.7 | 163754765.9 | 169593057.2 | 124417656.6 | 140877975.3 | 108971975.6 | 114051853.5 | 123844905.3 | 128420560.3 | 108745271.3 | 21.75461 | 23.68438 | 19.07037 | 7.556776 | Agk | 69923 | AGK | 55750 | 1.24 | 1.12 | 0.87 | 0.76 | 1.63 | 0.04779 | 0.09611 | 2 | No | | 10 |
| 37359832 | mKIAA0193 protein [Mus musculus] | 0 | 46.588 | 24 | 8 | 53 | 8 | 439 | 49.1 | 4.94 | 542 | 8 | 112203668 | 134008654.6 | 132331880.3 | 136806021.6 | 129624856 | 115808558.7 | 148669236.1 | 116350771.4 | 101421590.9 | 121725099.6 | 120350101.9 | 130953024.6 | 118052300.4 | 116394947.4 | 138365223.6 | 129237583.9 | 8.72555 | 12.08563 | 10.44564 | 8.200977 | Scrn1 | 69938 | SCRN1 | 9805 | 1.03 | 1.02 | 0.95 | 1.00 | 1.09 | 0.7207 | 0.75816 | 2 | No | | 6 |
| 89111939 | exocyst complex component 1 isoform 1 [Mus musculus] | 0 | 12.897 | 7 | 4 | 10 | 4 | 901 | 102.6 | 6.47 | 47 | 4 | 3162294.03 | 3329956.922 | 2418400.543 | 2677428.563 | 1702283.119 | 2775009.876 | 2023896.985 | 1287003.745 | 955094.7199 | 3137723.05 | 2728654.839 | 3412674.324 | 1652130.358 | 1741237.488 | 3714561.955 | 3529502.765 | 14.57827 | 32.30629 | 43.19994 | 41.91833 | Exoc1 | 69940 | EXOC1 | 55763 | 1.15 | 0.77 | 1.02 | 1.06 | 1.49 | 0.2567 | 0.33865 | 2 | No | | 8 |
| 68448515 | cell division cycle protein 16 homolog [Mus musculus] | 0 | 8.541 | 6 | 3 | 16 | 3 | 620 | 71.4 | 5.76 | 34 | 3 | 1475372.952 | 1898353.646 | 1282402.263 | 1163735.625 | 1358548.029 | 1495028.792 | 1646136.062 | 1323559.823 | 1408738.212 | 1373781.806 | 1872470.149 | 1297085.945 | 1438275.631 | 1708686.864 | 1953016.353 | 1158973.962 | 22.15067 | 10.08846 | 17.50705 | 21.89486 | Cdc16 | 69957 | CDC16 | 8881 | 0.98 | 0.98 | 1.00 | 1.05 | 1.08 | 0.9567 | 0.9613 | 2 | No | | 0 |
| 17391158 | Transmembrane protein 30A [Mus musculus] | 0 | 7.538 | 7 | 2 | 10 | 2 | 364 | 41.1 | 8.37 | 67 | 2 | 3638775.189 | 4629443.046 | 4221160.211 | 3173813.375 | 3437864.419 | 3407803.003 | 4256068.424 | 3446053.235 | 4338455.686 | 4095278.013 | 4714687.766 | 3798559.113 | 1803289.31 | 4502701.421 | 4778429.1 | 3882490.565 | 16.35082 | 11.35774 | 9.149291 | 35.95958 | Tmem30a | 69981 | TMEM30A | 55754 | 1.01 | 0.94 | 1.09 | 0.96 | 1.16 | 0.3635 | 0.44516 | 2 | No | | 5 |
| 755533203 | D: SAM and SH3 domain-containing protein 1 isoform X1 [Mus | 0 | 34.94 | 8 | 8 | 40 | 8 | 1301 | 143.1 | 7.02 | 468 | 8 | 5945133.112 | 7849624.524 | 5671507.827 | 5691412.75 | 7529234.87 | 8068891.481 | 11831741.98 | 8126465.938 | 10018298.18 | 12441529.53 | 12918559.39 | 9221172.882 | 2387911.994 | 7639595.737 | 13832103.93 | 9830274.771 | 16.65605 | 22.27596 | 16.20745 | 56.6409 | Sash1 | 70097 | SASH1 | 23328 | 0.72 | 1.02 | 1.28 | 0.97 | 1.77 | 0.02551 | 0.06287 | 3 | No | | 2 |
| 219276601 | asparagine--tRNA ligase, cytoplasmic isoform 1 [Mus musculu | 0 | 6.815 | 3 | 2 | 4 | 2 | 559 | 64.2 | 5.86 | 0 | 2 | 6577741.648 | 5147843.931 | 7010261.086 | 5983046.5 | 4821310.985 | 4905913.276 | 5118436.74 | 6845515.055 | 6008509.154 | 5999449.34 | 6849422.747 | 6362291.195 | 7104078.044 | 5775111.547 | 5785802.645 | 6695886.636 | 13.05189 | 17.64185 | 6.350505 | 10.52817 | Nars | 70223 | NARS1 | 4677 | 1.02 | 0.89 | 1.04 | 1.05 | 1.17 | 0.5133 | 0.58411 | 2 | No | | 2 |
| 568934520 | ICTED: actin filament-associated protein 1 isoform X1 [Mus mu | 0 | 38.195 | 13 | 8 | 34 | 8 | 820 | 90 | 8.57 | 410 | 8 | 25105422.09 | 25268468.72 | 20851725.09 | 24481812.75 | 17466835.19 | 20124799.32 | 23802024.04 | 22120008.72 | 19223554.45 | 19306827.81 | 18052114.75 | 22643588.95 | 17096452.47 | 20982199.54 | 15201232.54 | 18499123.63 | 8.684499 | 13.05732 | 9.977666 | 13.56647 | Afap1 | 70292 | AFAP1 | 60312 | 1.16 | 1.01 | 0.96 | 0.87 | 1.33 | 0.05397 | 0.10498 | 2 | No | | 0 |
| 21362321 | E3 ubiquitin-protein ligase RNF126 [Mus musculus] | 0 | 45.316 | 35 | 7 | 28 | 7 | 313 | 34.1 | 5.17 | 475 | 7 | 13336295.17 | 18369957.3 | 16973440.78 | 12748519 | 20014095.05 | 13320385.61 | 24346731.47 | 13774855.36 | 27986341.71 | 22421292.59 | 40352887.32 | 18546476.89 | 20269782.36 | 28359152.99 | 34098339.77 | 23300919.45 | 17.8639 | 29.62352 | 34.79797 | 22.8691 | Rnf126 | 70294 | RNF126 | 55658 | 0.71 | 0.82 | 1.26 | 1.22 | 1.78 | 0.06223 | 0.11587 | 3 | No | | 0 |
| 45598372 | brain acid soluble protein 1 [Mus musculus] | 0 | 61.891 | 58 | 7 | 81 | 7 | 226 | 22.1 | 4.51 | 1593 | 7 | 7847439.126 | 7213763.95 | 8734141.209 | 4650378.25 | 10649892.59 | 8348941.645 | 9634101.961 | 8524829.749 | 15944418.7 | 8058503.51 | 15476855.27 | 7436848.27 | 13970056 | 12637248.87 | 14973784.31 | 9154473.746 | 24.68135 | 11.52531 | 39.28989 | 20.02643 | Basp1 | 70350 | BASP1 | 10409 | 0.70 | 0.91 | 1.15 | 1.24 | 1.78 | 0.07072 | 0.12787 | 3 | No | | 12 |
| 19526912 | hsc70-interacting protein [Mus musculus] | 0 | 15.381 | 12 | 4 | 23 | 4 | 371 | 41.6 | 5.26 | 145 | 4 | 7072054.012 | 7912605.427 | 9228744.902 | 7183269.75 | 2724931.366 | 4634773.947 | 5062654.489 | 4006682.283 | 2922687.194 | 2817709.302 | 3140660.972 | 2447600.484 | 2349528.582 | 2968245.763 | 3063049.62 | 2880152.987 | 12.6434 | 24.79744 | 10.22246 | 11.33372 | St13 | 70356 | ST13 | 6767 | 1.78 | 0.93 | 0.64 | 0.64 | 2.79 | 0.00055 | 0.00488 | 1 | Yes | Decrease A1 | 2 |
| 58037205 | BAG family molecular chaperone regulator 5 [Mus musculus] | 0 | 33.609 | 19 | 5 | 13 | 5 | 447 | 50.9 | 6.05 | 237 | 5 | 11219653.65 | 7181262.515 | 12090803.17 | 10370902.88 | 15258470.19 | 8855345.722 | 12203318.76 | 11822777.69 | 11759665.85 | 9782342.838 | 15205505.01 | 13709658.28 | 209642.0108 | 10538477.43 | 19919139.92 | 14507233.67 | 20.96122 | 21.7599 | 18.68414 | 73.75578 | Bag5 | 70369 | BAG5 | 9529 | 0.89 | 1.04 | 1.09 | 0.98 | 1.23 | 0.5879 | 0.64939 | 2 | No | | 7 |
| 569004475 | PREDICTED: casein kinase I isoform X1 [Mus musculus] | 0 | 33.951 | 20 | 8 | 66 | 4 | 456 | 52.4 | 9.11 | 518 | 8 | 22299562.41 | 24268495.06 | 21223408.76 | 15564228 | 14271466.02 | 15382446.87 | 15857573.21 | 15900414.28 | 13843319.49 | 13939195.9 | 15270687.98 | 13867038.39 | 15689163.71 | 14013193.02 | 14712623.29 | 11843910.36 | 17.9264 | 4.939147 | 4.883667 | 11.56272 | Csnk1g3 | 70425 | CSNK1G3 | 1456 | 1.29 | 0.95 | 0.88 | 0.87 | 1.48 | 0.05574 | 0.10717 | 2 | No | | 5 |
| 21312594 | mitochondrial pyruvate carrier 2 [Mus musculus] | 0 | 42.066 | 33 | 6 | 142 | 6 | 127 | 14.3 | 10.61 | 1186 | 6 | 1294669349 | 1194822225 | 1008356838 | 957075294 | 1163695018 | 811561351.7 | 665168873.1 | 702986265.8 | 712149772.7 | 602027608.8 | 868261599.3 | 690517396.7 | 545029028.7 | 609366321.4 | 632064199.2 | 557571315.4 | 14.19291 | 27.18161 | 15.424 | 7.073559 | Mpc2 | 70456 | MPC2 | 25874 | 1.37 | 1.03 | 0.88 | 0.72 | 1.90 | 0.00616 | 0.0231 | 2 | Yes | | 8 |
| 140971160 | CREB-regulated transcription coactivator 3 [Mus musculus] | 0 | 20.691 | 7 | 3 | 18 | 3 | 619 | 67 | 6.83 | 146 | 3 | 7090817.595 | 6793342.459 | 8277500.249 | 7637540 | 6685119.802 | 6032669.508 | 4758764.007 | 1764957.183 | 5864607.377 | 4919303.089 | 4054324.965 | 4345347.542 | 3882403.918 | 3838040.098 | 5401647.794 | 4784764.14 | 8.768459 | 45.38429 | 16.63872 | 16.8721 | Crtc3 | 70461 | CRTC3 | 64784 | 1.38 | 0.89 | 0.89 | 0.83 | 1.66 | 0.00441 | 0.01865 | 2 | Yes | | 1 |
| [illegible] | | | | | | | | | | | | | | | | | | | | | | | | | | | | | | | | | | | | | | | | | | | | | | | |

| | | | | | | | | | | | | | | | | |
|---|---|---|---|---|---|---|---|---|---|---|---|---|---|---|---|---|
| 16905129 | N-terminal EF-hand calcium-binding protein 2 [Mus musculus] | 0 | 28.104 | 23 | 6 | 32 | 6 | 389 | 43.4 | 5.3 | 274 | 6 | 10625007.54 | 13014821.91 | 9437196.805 | 4892558.469 | 6993276.132 |
| 568966456 | sphatidylinositol 5-phosphate 4-kinase type-2 gamma isoform X | 0 | 28.785 | 11 | 4 | 28 | 4 | 479 | 53.6 | 7.27 | 212 | 4 | 4897867.534 | 6330219.354 | 6800675.955 | 4283621.625 | 3569608.966 |
| 187957332 | Synaptopodin 2 [Mus musculus] | 0 | 19.613 | 6 | 5 | 7 | 5 | 1257 | 135.7 | 8.13 | 77 | 5 | 1859065.731 | 1730408.818 | 1319233.485 | 1496465 | 1339779.641 |
| 568971280 | DICTED: gap junction gamma-2 protein isoform X1 [Mus musc | 0 | 32.964 | 11 | 3 | 29 | 3 | 572 | 61.2 | 7.25 | 344 | 3 | 8177272.371 | 7214762.153 | 6311035.93 | 9526732.625 | 8143617.829 |
| 569003248 | nductance calcium-activated potassium channel protein 2 isofo | 0 | 20.85 | 6 | 3 | 11 | 3 | 852 | 93 | 9.28 | 127 | 3 | 5963017.518 | 5662699.508 | 6626357.756 | 4506411 | 3129715.895 |
| 46402197 | coiled-coil, ANK repeat and PH domain-containing protein 3 [M | 0 | 9.844 | 4 | 3 | 14 | 2 | 833 | 92.7 | 6.02 | 31 | 3 | 19375513.99 | 27129327.46 | 31875983.22 | 20409183.94 | 20590638.92 |
| 1039766579 | REDICTED: ERI1 exoribonuclease 3 isoform X1 [Mus musculu | 0 | 12.425 | 10 | 3 | 12 | 3 | 374 | 41.1 | 6.9 | 89 | 3 | 4716561.655 | 8540823.894 | 7903263.081 | 5741423.875 | 7442325.431 |
| 15593237 | immunoglobulin superfamily receptor PGRL [Mus musculus] | 0 | 105.523 | 29 | 11 | 129 | 11 | 611 | 65.1 | 8.13 | 3008 | 11 | 76402153.07 | 64165999.21 | 54998555.17 | 53374870.88 | 98616201.43 |
| 26341524 | unnamed protein product [Mus musculus] | 0 | 157.923 | 39 | 23 | 300 | 18 | 720 | 82.5 | 5.96 | 2968 | 23 | 132849616.1 | 178961424.8 | 187212849.3 | 177923297.6 | 132029749.3 |
| 568981546 | TED: engulfment and cell motility protein 1 isoform X1 [Mus m | 0 | 85.426 | 21 | 15 | 174 | 10 | 742 | 85.5 | 6 | 1176 | 15 | 50849777.21 | 79569040.48 | 79795522.13 | 69965502 | 56113872.01 |
| 19880961 | age dependent calcium channel gamma 5 subunit [Mus muscu | 0 | 10.329 | 10 | 2 | 3 | 2 | 275 | 30.8 | 7.33 | 34 | 2 | 1762752.104 | 7559601.235 | 7196932.42 | 5090482.625 | 4317479.926 |
| 16554627 | WD repeat-containing protein 5 [Homo sapiens] | 0 | 6.81 | 9 | 3 | 6 | 3 | 334 | 36.6 | 8.27 | 23 | 3 | 6971209.089 | 8522816.876 | 3804200.156 | 8176944.5 | 7751198.614 |
| 188219544 | vesicular glutamate transporter 2 [Mus musculus] | 0 | 23.563 | 12 | 5 | 29 | 4 | 582 | 64.5 | 6.83 | 283 | 5 | 6187154.189 | 11448677.81 | 10290826.15 | 8149960 | 4743626.447 |
| 27923929 | hippocalcin-like protein 4 [Mus musculus] | 0 | 55.704 | 48 | 8 | 43 | 6 | 191 | 22.2 | 4.89 | 1082 | 8 | 18441510.48 | 28128239.84 | 24199615.92 | 20949605.91 | 24909050.12 |
| 18700024 | ate dehydrogenase [NAD] subunit beta, mitochondrial [Mus mu | 0 | 117.499 | 49 | 16 | 166 | 16 | 384 | 42.2 | 8.6 | 2341 | 16 | 173475922.5 | 156965390.5 | 194701637.3 | 212403695.4 | 215587889.3 |
| 194328708 | oxidation resistance protein 1 isoform E [Mus musculus] | 0 | 29.624 | 11 | 6 | 13 | 6 | 866 | 95.9 | 5.33 | 128 | 6 | 9291171.618 | 11068228.25 | 9419771.391 | 9927945.219 | 7995277.915 |
| 568955238 | TED: manganese-transporting ATPase 13A1 isoform X1 [Mus r | 0 | 39.532 | 9 | 8 | 36 | 8 | 1223 | 135 | 7.99 | 163 | 8 | 17023825.08 | 20057029.06 | 17882982.93 | 14056608 | 7936465.494 |
| 37360098 | mKIAA0791 protein [Mus musculus] | 0 | 137.524 | 22 | 22 | 79 | 22 | 1423 | 158.5 | 6.24 | 1026 | 22 | 26049597.05 | 36915630.61 | 62379182.66 | 77735650.25 | 17674149.47 |
| 26972037 | mKIAA0027 protein [Mus musculus] | 0 | 22.604 | 7 | 2 | 27 | 2 | 423 | 46.1 | 8.38 | 870 | 2 | 5098332.82 | 7233340.266 | 4884538.5 | 4902576 | 6321887.725 |
| 118403314 | RNA-binding protein 39 isoform a [Mus musculus] | 0 | 19.196 | 9 | 4 | 21 | 4 | 530 | 59.4 | 10.1 | 234 | 4 | 7528240.213 | 16463551.42 | 7242218.567 | 7068125.875 | 6178205.111 |
| 1039735923 | sphatidylinositol 4,5-bisphosphate 5-phosphatase A isoform X | 0 | 13.812 | 5 | 4 | 17 | 4 | 1039 | 111.2 | 9.38 | 84 | 4 | 1274672.92 | 1973341.448 | 1801791.446 | 1235895.125 | 5712872.034 |
| 226693351 | delphilin isoform 1 [Mus musculus] | 0 | 103.201 | 20 | 17 | 156 | 17 | 1203 | 131.9 | 7.02 | 1953 | 17 | 76458437.55 | 62228843.57 | 89085079.77 | 82588265.7 | 91041695.85 |
| 162417988 | prosaposin receptor GPR37L1 precursor [Mus musculus] | 0 | 23.095 | 6 | 2 | 24 | 2 | 481 | 52.7 | 7.36 | 539 | 2 | 7214960.093 | 7474969.887 | 5026956.515 | 10031915 | 7314595.049 |
| 148707307 | tatic cells 7, protein expressed in, isoform CRA_b, partial [Mus | 0 | 22.018 | 15 | 5 | 27 | 5 | 421 | 47.6 | 7.05 | 132 | 5 | 22475000.38 | 25214371.65 | 17603421.3 | 27009720 | 15066889.93 |
| 568920807 | PREDICTED: neuroligin-1 isoform X1 [Mus musculus] | 0 | 51.565 | 13 | 9 | 119 | 4 | 863 | 96.4 | 6.4 | 2972 | 9 | 22182047.24 | 21034118.09 | 20679766.93 | 23657929 | 16924110.09 |
| 19344056 | BC003940 protein, partial [Mus musculus] | 0 | 22.933 | 39 | 4 | 31 | 4 | 126 | 14.3 | 10.67 | 342 | 4 | 12428168.75 | 15040925.37 | 14208326.17 | 12517686 | 12198985.29 |
| 569009889 | PREDICTED: filamin-A isoform X1 [Mus musculus] | 0 | 147.704 | 16 | 29 | 165 | 26 | 2647 | 281 | 6.04 | 2011 | 29 | 49296329.02 | 71993540.03 | 66023334.45 | 69463502.63 | 35119317.56 |
| 564346383 | PREDICTED: LUC7-like 2 isoform X3 [Rattus norvegicus] | 0 | 31.121 | 20 | 7 | 32 | 4 | 408 | 48.4 | 10.2 | 249 | 7 | 7693266.797 | 9350007.766 | 7991218.289 | 8729375.875 | 8652477.466 |
| 568972299 | breast carcinoma-amplified sequence 3 homolog isoform X1 [M | 0 | 43.007 | 12 | 8 | 36 | 8 | 963 | 104.8 | 6.74 | 155 | 8 | 11098367.45 | 8462474.102 | 12173361.09 | 5690099.688 | 9540405.054 |
| 148705408 | ear receptor binding protein, isoform CRA_b, partial [Mus musc | 0 | 10.506 | 8 | 3 | 8 | 3 | 559 | 62.3 | 5.29 | 23 | 3 | 1905325.894 | 2338048.746 | 1840891.15 | 1966821.313 | 1214657.37 |
| 13435747 | Rho GDP dissociation inhibitor (GDI) alpha [Mus musculus] | 0 | 32.495 | 35 | 4 | 30 | 4 | 204 | 23.4 | 5.2 | 440 | 4 | 16821226.18 | 22960994.2 | 18220412.53 | 14081809.5 | 7325633.146 |
| 755516345 | ): protein-methionine sulfoxide oxidase MICAL3 isoform X1 [Mu | 0 | 137.087 | 18 | 31 | 209 | 31 | 2252 | 253 | 5.8 | 2114 | 31 | 87913684.7 | 101436822.6 | 78580795 | 80853814.44 | 95069917.96 |
| 124487439 | rho guanine nucleotide exchange factor 17 [Mus musculus] | 0 | 38.208 | 6 | 9 | 27 | 9 | 2057 | 221.5 | 6.3 | 120 | 9 | 2662723.701 | 3280947.052 | 1777987.381 | 2266649.375 | 3737561.6 |
| 568994816 | DICTED: syntaxin-binding protein 5-like isoform X1 [Mus musc | 0 | 12.691 | 3 | 3 | 19 | 3 | 1216 | 135.2 | 7.02 | 242 | 3 | 10537920.05 | 14418239.68 | 15139627.31 | 15293640 | 10717149.84 |

[illegible]

| | | | | | | | | | | | | |
|---|---|---|---|---|---|---|---|---|---|---|---|---|
| 12003362 | NADP+-specific isocitrate dehydrogenase [Mus musculus] | 0 | 19.517 | 15 | 6 | 30 | 6 | 452 | 50.9 | 8.69 | 41 | 6 |
| 148684637 | nucleoporin 98, isoform CRA_c [Mus musculus] | 0 | 113.579 | 19 | 24 | 90 | 24 | 1816 | 197.2 | 6.15 | 767 | 24 |
| 157951733 | germ cell-specific gene 1-like protein precursor [Mus musculus] | 0 | 29.023 | 25 | 7 | 92 | 7 | 322 | 35.9 | 7.01 | 1458 | 7 |
| 148698944 | tubule-associated protein 1S, isoform CRA_a, partial [Mus mus | 0 | 56.613 | 19 | 13 | 76 | 13 | 984 | 104 | 7.36 | 209 | 13 |
| 33186863 | 60S ribosomal protein L13 [Mus musculus] | 0 | 31.031 | 32 | 7 | 183 | 7 | 211 | 24.3 | 11.55 | 2974 | 7 |
| 755528856 | PREDICTED: protocadherin Fat 3 isoform X1 [Mus musculus] | 0 | 21.657 | 2 | 6 | 13 | 6 | 4611 | 507.9 | 4.87 | 36 | 6 |
| 568961564 | EDICTED: unconventional myosin-IXa isoform X1 [Mus muscu | 0 | 92.975 | 10 | 22 | 116 | 22 | 2633 | 301.9 | 8.9 | 1022 | 22 |
| 30424655 | ras-related protein Rab-6B [Mus musculus] | 0 | 13.365 | 15 | 3 | 59 | 2 | 208 | 23.4 | 5.53 | 497 | 3 |
| 12832758 | unnamed protein product [Mus musculus] | 0 | 18.044 | 27 | 5 | 27 | 2 | 215 | 23.6 | 8.43 | 205 | 5 |
| 38016190 | ucine-rich repeat-containing protein 4B precursor [Mus muscul | 0 | 48.667 | 20 | 11 | 83 | 10 | 709 | 76.1 | 7.24 | 442 | 11 |
| 227908819 | probable threonine--tRNA ligase 2, cytoplasmic [Mus musculus] | 0 | 11.677 | 5 | 4 | 17 | 4 | 790 | 91.3 | 7.53 | 181 | 4 |
| 148680531 | ryotic translation initiation factor 5A, isoform CRA_c [Mus musc | 0 | 26.537 | 43 | 5 | 52 | 5 | 166 | 18.4 | 5.24 | 572 | 5 |
| 50510921 | mKIAA1415 protein [Mus musculus] | 0 | 21.026 | 4 | 6 | 34 | 6 | 1665 | 186.4 | 6.48 | 722 | 6 |
| 125347500 | C2 domain-containing protein 3 [Mus musculus] | 0 | 27.188 | 3 | 7 | 17 | 7 | 2323 | 255.2 | 6.95 | 141 | 7 |
| 157426849 | protocadherin-19 isoform a precursor [Mus musculus] | 0.002 | 4.968 | 2 | 2 | 10 | 2 | 1145 | 126 | 5.5 | 33 | 2 |
| 160420308 | histone H2B type 1-P isoform 1 [Mus musculus] | 0 | 47.949 | 38 | 5 | 159 | 4 | 138 | 15.6 | 10.24 | 3343 | 5 |
| 28174920 | Rpl17 protein, partial [Mus musculus] | 0 | 54.446 | 35 | 6 | 137 | 6 | 194 | 22.4 | 10.33 | 1952 | 6 |
| 755512491 | 'ED: adhesion G protein-coupled receptor L3 isoform X1 [Mus r | 0 | 147.358 | 18 | 25 | 228 | 24 | 1599 | 177.8 | 6.77 | 3409 | 25 |
| 60359932 | mKIAA1157 protein [Mus musculus] | 0 | 19.597 | 8 | 3 | 25 | 3 | 640 | 70.1 | 9.64 | 236 | 3 |
| 568979802 | ICTED: neuronal cell adhesion molecule isoform X1 [Mus mus | 0 | 62.433 | 9 | 8 | 63 | 8 | 1311 | 144.6 | 6 | 1022 | 8 |
| [illegible] | [illegible] | [illegible] | [illegible] | [illegible] | [illegible] | [illegible] | [illegible] | [illegible] | [illegible] | [illegible] | [illegible] | [illegible] |

**Supplementary Table 2. List of differentially expressed PSD proteins which have been reported to regulate number, shape, or function of synapses**

| Subgruop | Gene Symbol | Entrez ID | fold change (12w/2w) | Representative reference | Method | Sample | Overview of synapse related phenotypes |
|---|---|---|---|---|---|---|---|
| Decrease | Dcx | 13193 | 0.054 | Yoshihara et al. Cell Rep. 2014 8:843-57. | overexpression, RNAi | olfactory bulb granule cells | increased spine density (overexpression), decreased spine density (RNAi) |
| Decrease | Fabp7 | 12140 | 0.070 | Ebrahimi et al. Glia. 2016 64:48-62. | knockout | mPFC | decreased spine density, decreased ratio of mature spines |
| Decrease | Adcy1 | 432530 | 0.147 | Sethna et al. Nat Commun. 2017 8:14359. | knockout | hippocampal CA1 region | decreased spine density (in Fmr1 KO) |
| Decrease | Cdk5r2 | 12570 | 0.148 | Li et al. J Neurosci. 2016 36:11283-11294. | RNAi | primary hippocampal neuron | decreased spine density |
| Decrease | Glra2 | 237213 | 0.198 | Morelli et al. Cereb Cortex. 2017 27:1863-1877. | knockout | cortical layer 5 | increased spine number and density, increased frequency of mEPSC |
| Decrease | Fabp5 | 16592 | 0.217 | Haj-Dahmane et al. Proc Natl Acad Sci U S A. 2018 115:3482-3487. | inhibitor treatment, knockout | dorsal raphe nucleus | abolished eCB-mediated control of excitatory synaptic transmission |
| Decrease | Plcb1 | 18795 | 0.222 | Spires et al. Cereb Cortex. 2005 15:385-93. | knockout | barrel cortex | decreased synapses, decreased percentage of mushroom spines |
| Decrease | Nectin3 | 58998 | 0.235 | Wang et al. Transl Psychiatry. 2017 7:e1228. | RNAi | hippocampal dentate gyrus | decreased spine density |
| Decrease | Myo16 | 244281 | 0.240 | Roesler et al. Front Cell Neurosci. 2019 13:330 | RNAi, knockout | cerebellum, primary Purkinje cells | accelerated F-actin turnover in Purkinje cell dendritic spines (RNAi, knockout),<br>smaller presynaptic terminal area and fewer presynaptic vesicles (knockout) |
| Decrease | Cpeb2 | 231207 | 0.259 | Lu et al. Cell Rep. 2017 21:1783-1794. | knockout | primary cortical and hippocampal neuron, hippocampal CA1 region | increased spine length, decreased amplitude of mEPSC, impaired LTP |
| Decrease | Top1 | 21969 | 0.274 | Mabb et al. Proc Natl Acad Sci U S A. 2014 111:17290-5. | inhibitor treatment | primary cortical neuron | decreased synapse density, decreased frequency of mEPSC |
| Decrease | Ptn | 19242 | 0.280 | Pavlov et al. Mol Cell Neurosci. 2002 20:330-42. | overexpression, knockout | hippocampal CA1 region | impaired LTP (overexpression), enhanced LTP (knockout) |
| Decrease | Hsp90b1 | 22027 | 0.295 | Chen et al. J Neurosci. 2014 34:2464-70. | inhibitor treatment | primary hippocampal neuron, hippocampal DG region | increased spine density, increased synapse protein expression, enhanced LTP |
| Decrease | Clasp2 | 76499 | 0.297 | Beffert et al. J Neurosci. 2012 32:13906-16. | overexpression, RNAi | primary hippocampal neuron | increased number of synapses and the frequency of mEPSC (overexpression),<br>decreased number of synapses and the frequency of mEPSC (RNAi) |
| Decrease | Vangl2 | 93840 | 0.302 | Nagaoka et al. Cell Rep. 2014 6:916-2 | RNAi, mutation | primary hippocampal neuron, hippocampus | decreased spine density |
| Decrease | Tnc | 21923 | 0.319 | Irintchev et al. Cereb Cortex. 2005 15:950-62. | knockout | somatosensory cortex and motor cortex | aberrant distribution of stubby spines |
| Decrease | Ncam1 | 17967 | 0.322 | Irala et al. Development. 2016 143:4224-4235. | overexpression, RNAi | primary hippocampal neuron | increased spine density (overexpression), defect in GDNF dependent spine formation (RNAi) |
| Decrease | Scn2a | 110876 | 0.326 | Spratt et al. Neuron. 2019 103:673-685.e5. | heterozygous knockout | mPFC | increased spine length, decreased spine head volume, decreased mEPSC frequency, impaired LTP |
| Decrease | Srgap2 | 14270 | 0.338 | Charrier et al. Cell. 2012 149:923-35. | overexpression, RNAi, knockout | primary cortical neuron, layer 5 pyramidal neuron | increased width of spine head width and decreased length of spine neck (overexpression)<br>decreased width of spine head width and in creased length of spine neck (RNAi, knockout), increased spine density (knockout) |
| Decrease | Asic2 | 11418 | 0.348 | Zha et al. J Neurosci. 2009 29:8438-46. | knockout | hippocampus | decreased spine density |
| Decrease | Hnrnpm | 76936 | 0.351 | Zhang et al. J Neurosci. 2012 32:599-609. | RNAi | primary hippocampal neuron | increased spine density |
| Decrease | Opa1 | 74143 | 0.359 | Li et al. Cell. 2004 119:873-87. | overexpression | primary hippocampal neuron | decreased spine density |
| Decrease | Gjd2 | 14617 | 0.359 | Miller et al. Elife. 2017 6. pii: e25364. | mutation | hindbrain and spinal cord | impaired function of electrical synapses |
| Decrease | Arl8b | 67166 | 0.364 | Vukoja et al. Neuron. 2018 99:1216-1232.e7. | overexpression, RNAi, mutation | fly motoneurons | defective presynaptic biogenesis (RNAi, mutation) facilitated presynaptic biogenesis and neurotransmission (overexpression) |
| Decrease | Kif21b | 16565 | 0.377 | Muhia et al. Cell Rep. 2016 15:968-977. | knockout | primary hippocampal neuron, hippocampal CA1 region | decreased spine density, impaired LTP, decreased frequency of mEPSC |
| Decrease | Uchl1 | 22223 | 0.382 | Smith et al. Proc Natl Acad Sci U S A. 2009 106:16877-82. | addition of recombinant protein | hippocampus | increased spine density, increased head diameter and length of spines |
| Decrease | Mapk8 | 26419 | 0.387 | Sclip et al. Cell Death Dis. 2014 5:e1019. | inhibitor treatment | hippocampus | increased spine volume and PSD area, restoration of LTD (in TgCRND8) |
| Decrease | Dnm3 | 103967 | 0.389 | Gray et al. J Cell Sci. 2005 118(Pt 6):1279-90. | overexpression | primary hippocampal neuron | increased filopodia density, decreased ratio of mature spines and synapses |
| Decrease | Cfl1 | 12631 | 0.390 | Tsubota et al. PLoS Biol. 2015 13:e1002070. | RNAi | barrel cortex (layer 1) | decreased spine density (upon sensory deprivation) |
| Decrease | Khdrbs1 | 20218 | 0.390 | Klein et al. Proc Natl Acad Sci U S A. 2013 110:3125-30. | RNAi, knockout | primary hippocampal neuron, hippocampal CA1 region | decreased spine density, decreased frequency of mEPSC |
| Decrease | Ptk2b | 19229 | 0.398 | Giralt et al. Nat Commun. 2017 8:15592. | knockout | hippocampal CA1 region (stratum radiatum) | decreased spine density, impaired LTP |
| Decrease | Sept2 | 18000 | 0.398 | Tada et al. Curr Biol. 2007 17:1752-8. | overexpression, RNAi | primary hippocampal neuron | increased spine density (overexpression), decreased spine density (RNAi) |
| Decrease | Gsk3b | 56637 | 0.400 | Ochs et al. Mol Psychiatry. 2015 20:482-9. | knockout | hippocampal CA1 region | decreased spine density and stability, increased ratio of stubby spines and decreased ratio of thin and mushroom spines |
| Decrease | Farp1 | 223254 | 0.411 | Cheadle and Biederer J Cell Biol. 2012 199:985-1001. | overexpression, RNAi | primary hippocampal neuron | increased spine density (overexpression), decreased spine density (RNAi) |
| Decrease | Dapk1 | 69635 | 0.418 | Goodell et al. Cell Rep. 2017 19:2231-2243. | inhibitor treatment | primary hippocampal neuron, hippocampal CA1 region | enhanced LTD-induced CaMKII translocation to spine, impaired LTD |
| Decrease | Vgf | 381677 | 0.419 | Behnke et al. Int J Mol Sci. 2017 18. pii: E612. | overexpression | primary hippocampal neuron | increased spine density, increased number of thinner spines |
| Decrease | Ppp2r2b | 72930 | 0.427 | Dickey and Strack J Neurosci. 2011 31:15716-26. | overexpression, RNAi | primary hippocampal neuron | increased synapse density (overexpression), decreased synapse density (RNAi) |
| Decrease | Pafah1b1 | 18472 | 0.427 | Sudarov et al. EMBO Mol Med. 2013 5:591-607. | heterozygous knockout | barrel cortex, hippocampal CA1 region, primary hippocampal neuror | decreased density of spines and filopodia, increased stability of spines and filopodia |
| Decrease | Rhof | 23912 | 0.434 | Hotulainen et al. J Cell Biol. 2009 185:323-39. | overexpression (active- or inactive-form) | primary hippocampal neuron | decreased spine density (active form), decreased thin spine density (inactive form) |
| Decrease | Rnd2 | 11858 | 0.436 | Wakita et al. J Biol Chem. 2011 286:32672-83. | RNAi | primary hippocampal neuron | decreased spine density |
| Decrease | Kcnn2 | 140492 | 0.444 | Ngo-Anh et al. Nat Neurosci. 2005 8:642-9. | inhibitor treatment | hippocampal CA1 region | increased EPSP |
| Decrease | Hsp90ab1 | 15516 | 0.461 | Chen et al. J Neurosci. 2014 34:2464-70. | inhibitor treatment | primary hippocampal neuron, hippocampal DG region | increased spine density, increased synapse protein expression, enhanced LTP |
| Decrease | Cdh7 | 241201 | 0.467 | Kuwako et al. Cell Rep. 2014 9:311-323. | RNAi | primary pontine nuclei neurons | decreased number of synapses |
| Decrease | Cpeb4 | 67579 | 0.477 | Tsai et al. PLoS One. 2013 8:e84978. | knockout | primary cortical and hippocampal neuron | increased spine length |
| Decrease | Eif4e | 13684 | 0.477 | Santini et al. Sci Signal. 2017 10. pii: eaan0665. | inhibitor treatment | hippocampal CA1 region | decreased spine density (in Fmr1 KO), reduced mGluR-LTD (in Fmr1 KO) |
| Decrease | Pdlim5 | 56376 | 0.484 | Herrick et al. Mol Cell Neurosci. 2010 43:188-200. | overexpression, RNAi | primary hippocampal neuron | increased spine length (overexpression), increased spine width (RNAi) |
| Decrease | Arf6 | 11845 | 0.498 | Choi et al. J Neurosci. 2006 26:4811-9. | RNAi | primary hippocampal neuron | decreased spine density and increased filopodia density |
| Decrease | Cdh6 | 12563 | 0.500 | Basu et al. Neuron. 2017 96:160-176.e8. | knockout | hippocampal CA1 region (stratum oriens) | decrease of mushroom spines, defective LTP |
| Decrease | Shank2 | 210274 | 0.501 | Schmeisser et al. Nature. 2012 486:256-60. | knockout | hippocampal CA1 region | decreased spine density, decreased mEPSC frequency, enhanced LTP |
| Decrease | Dlg3 | 53310 | 0.518 | Murata and Constantine-Paton. J Neurosci. 2013 33:5040-52. | RNAi | primary cortical neuron | increased dendritic filopodia length, decreased density of excitatory synapses |
| Decrease | Gria1 | 14799 | 0.534 | Romberg et al. Eur J Neurosci. 2009 29:1141-52. | knockout | hippocampal CA1 region | attenuated synaptic transmission, impaired LTP induced by single weak tetanus |
| Decrease | Sirt2 | 64383 | 0.536 | Wang et al. Cell Rep. 2017 20:1335-1347. | knockout, inhibitor treatment | hippocampal CA1 region, primary hippocampal neuron | impaired LTP and LTD, increased amplitude of mEPSC |
| Decrease | Psip1 | 101739 | 0.541 | Li et al. PLoS Genet. 2016 12:e1006129. | RNAi | cortex | reduced decay time constant of glutamate-evoked AMPAR-gated current, weaker response upon repetitive stimulations |
| Decrease | Mapk8ip1 | 19099 | 0.567 | Morel et al. J Neurosci. 2018 38:3708-3728. | point mutation | hippocampal CA1 region | decreased threshold for LTP induction |
| Decrease | Sparcl1 | 13602 | 0.592 | Risher et al. Elife. 2014 3. doi: 10.7554/eLife.04047. | knockout | primary visual cortex | decreased number of thalamocortical synapses, smaller PSD, increased spines with multiple excitatory contacts |
| Increase | Rph3a | 19894 | 1.467 | Franchini et al. iScience. 2019 19:927-939. | RNAi | primary hippocampal neuron | decreased spine density |
| Increase | Fam107a | 268709 | 1.512 | Schmidt et al. Proc Natl Acad Sci U S A. 2011 108:17213-8. | overexpression | hippocampal CA1 and CA3 region | decreased spine density, smaller fEPSP amplitude, reduced magnitude of LTP |
| Increase | Agap2 | 216439 | 1.521 | Gross et al. Cell Rep. 2015 11:727-36. | Agap2 heterozygous knockout (with Fmr1 | hippocampal CA1 region | decreased spine density, attenuation of LTD |
| Increase | Gpr158 | 241263 | 1.541 | Condomitti et al. Neuron. 2018 100:201-215.e9. | knockout | hippocampal CA3 region (stratum lucidum) | increased spine density, decreased length of PSD and active zone, impaired synaptic transmission |
| Increase | Nptn | 20320 | 1.577 | Carrott et al. J Neurosci. 2016 36:222-34. | mutation | cochlea | defect in synapse formation |
| Increase | Cabp1 | 29867 | 1.580 | Mikhaylova et al. Neuron. 2018 97:1110-1125.e14. | overexpression, RNAi, knockout | primary hippocampal neuron, hippocampal CA1 region | increased spine density (overexpression), decreased density and volume of spines (RNAi, knockout), impaired LTP (knockout) |
| Increase | Rapgef4 | 56508 | 1.581 | Srivastava et al. J Neurosci. 2012 32:11864-78. | knockout (hetero and homo) | frontal cortex | decreased density of thin spines and filopodia, decreased motility of spines |
| Increase | Syn1 | 20964 | 1.601 | Perlini et al. J Cell Sci. 2011 124:3643-53. | RNAi, mutant expression | primary hippocampal neuron | decreased number of glutamatergic and GABAergic synapses |
| Increase | Setd1a | 233904 | 1.633 | Mukai et al. Neuron. 2019 104:471-487. | heterozygous knockout | mPFC | decreased density and head width of mushroom spines |
| Increase | Vasp | 22323 | 1.637 | Lin et al. J Biol Chem. 2010 285:36010-20. | overexpression, RNAi | primary hippocampal neuron | increased spine density and enlargement of spine head (overexpression), decreased spine density (RNAi) |
| Increase | Nlgn1 | 192167 | 1.656 | Kwon et al. Nat Neurosci. 2012 15:1667-74. | RNAi (sparse) | cortical layer 2/3 | decreased spine density |
| Increase | Kalrn | 545156 | 1.660 | Xie et al. Neuron. 2007 56:640-56. | RNAi | primary cortical neuron | decreased spine density and spine size |
| Increase | Sh3gl2 | 20404 | 1.663 | Yu et al. Nat Commun. 2018 9:2968. | overexpression | primary hippocampal neuron | decreased synapse density (in the presence of Abeta), impaired LTP (in the presence of Abeta) |
| Increase | Herc2 | 15204 | 1.668 | Valnegri et al. Nat Commun. 2017 8:1271. | RNAi | cerebellum | increased density of presynaptic boutons |
| Increase | Shank3 | 58234 | 1.670 | Wang et al. Hum Mol Genet. 2011 20:3093-108. | knockout (major isoforms) | hippocampal CA1 region | decreased spine density and increased spine length (4 weeks old), decreased spine length (10 weeks old) |
| Increase | Slc32a1 | 22348 | 1.672 | Kozorovitskiy et al. Nature. 2012 485:646-50. | knockout | striatum | decreased- or increased-spine density and mEPSC frequency (dependent on Cre expressing neurons) |
| Increase | Map1a | 17754 | 1.682 | Takei et al. J Neurosci. 2015 35:15539-54. | knockout | hippocampal CA1 region, primary hippocampal neuron | impaired LTP and LTD, decreased amplitude and frequency of NMDA mEPSC |
| Increase | Gnao1 | 14681 | 1.696 | Cha et al. Mol Brain. 2019 12:57. | knockout | cerebellum | decreased spine density, increased spine length |
| Increase | Cbln1 | 12404 | 1.706 | Hirai et al. Nat Neurosci. 2005 8:1534-41. | knockout | cerebellum | larger PSD, decreased number of synapses, reduced efficiency of synaptic transmission, impaired LTD |
| Increase | Prkcg | 18752 | 1.834 | Seki et al. Neurobiol Dis. 2009 33:260-73. | mutant expression | primary Purkinje cells | decreased spine density |
| Increase | Lgi1 | 56839 | 1.870 | Thomas et al.eNeuro. 2018 5. pii: ENEURO.0185-18.2018. | knockout | hippocampal CA1 region, primary hippocampal neuron | decreased synapse density, decreased ratio of mushroom spines, decreased length of mushroom spines,<br>decreased mEPSC frequency and amplitude |
| Increase | Cyld | 74256 | 1.873 | Ma et al. Proc Natl Acad Sci U S A. 2017 114:E8760-E8769. | RNAi | primary hippocampal neuron | defect in NMDA-triggered LTD |
| Increase | Usp46 | 69727 | 1.894 | Huo et al. J Neurochem. 2015 134:1067-80. | overexpression, RNAi | primary hippocampal neuron | increase (overexpression) or decrease (RNAi) of surface AMPAR and AMPAR mediated synaptic currents |
| Increase | Tmod1 | 21916 | 1.913 | Omotade et al. J Neurosci. 2018 38:10271-10285. | RNAi | primary hippocampal neuron | decreased spine density, increased filopodia density, decreased frequency and amplitude of mEPSCs |
| Increase | Ppfia3 | 76787 | 1.925 | Wong et al. Proc Natl Acad Sci U S A. 2018 115:2234-2239. | knockout | primary hippocampal neuron | impaired synaptic vesicle exocytosis |
| Increase | Prkca | 18750 | 1.950 | Schaffer et al. Cell Rep. 2018 25:2470-2483.e8. | overexpression | primary hippocampal neuron | decreased spine density |
| Increase | Nptx1 | 18164 | 1.956 | Figueiro-Silva et al. J Neurosci. 2015 35:5504-21. | RNAi | primary cortical neuron | increased spine density and synapse density, increased frequency of mEPSC |
| Increase | Grid1 | 14803 | 1.979 | Gupta et al. Neuropharmacology. 2015 93:274-84. | knockout | mPFC, hippocampal CA1 region | increased spine density and synapse density, higher frequency of mEPSC |
| Increase | Sptbn2 | 20743 | 1.985 | Efimova et al. J Neurosci. 2017 37:6442-6459. | RNAi | primary hippocampal neuron | decreased spine density, increased mEPSC amplitude |
| Increase | Rin1 | 225870 | 1.990 | Szíber et al. Mol Biol Cell 2017 28:285-295. | overexpression, knockout | primary hippocampal neuron | decreased ratio of mushroom spines, increased spine motility (overexpression), increased mEPSC amplitude (knockout) |
| Increase | Stx1b | 56216 | 1.992 | Mishima et al. PLoS One. 2014 9:e90004. | knockout | hippocampus, primary hippocampal neuron | decreased synaptic vesicle docking and neurotransmitter release, decreased frequency of mEPSC |
| Increase | Atp6v1a | 11964 | 2.008 | Fassio et al. Brain. 2018 141:1703-1718. | mutant expression | primary hippocampal neuron | decreased synapse density |
| Increase | Mapre3 | 100732 | 2.013 | Calipari et al. J Neurosci. 2019 39:5634-5646. | overexpression | nucleus accumbens | increased spine density |
| Increase | Camk2a | 12322 | 2.029 | Stephenson et al. J Neurosci. 2017 37:2216-2233. | mutant expression | primary hippocampal neuron | decreased spine density |
| Increase | Dgki | 320127 | 2.043 | Yang et al. EMBO J. 2011 30:165-80. | knockout | hippocampal CA1 region, primary hippocampal neuron | enhanced synaptic transmission, increased amplitude of EPSC, shorter time constant for NMDA-mediated dacay of EPSC |
| Increase | Atp2b4 | 381290 | 2.044 | Rajman et al. EMBO J. 2017 36:1770-1787. | overexpression | primary hippocampal neuron | increased spine size (in the presence of picrotoxin) |
| Increase | Gsg1l | 269994 | 2.051 | Gu et al. Nat Commun. 2016 7:10873. | overexpression, knockout | hippocampal CA1 region | impaired AMPA EPSC (overexpression), enhanced AMPA EPSC, enhanced LTP (knockout) |
| Increase | Atp8a1 | 11980 | 2.095 | Kerr et al. Biochim Biophys Acta. 2016 1862:1755-65. | overexpression | hippocampal CA1 region | fewer excitatory synapses, decreased number of active zones |
| Increase | Snap25 | 20614 | 2.118 | Tomasoni et al. Nat Commun. 2013;4:2136. | overexpression, RNAi | primary hippocampal neuron | increased spine density (overexpression)<br>decreased spine density and increased filopodia density, decreased frequency and amplitude of mEPSCs (RNAi) |
| Increase | Capns1 | 12336 | 2.121 | Amini et al. J Neurosci. 2013 33:5773-84. | knockout | hippocampal CA1 region | decreased spine density and counts in apical dendrite, defective LTP |
| Increase | Snx27 | 76742 | 2.124 | Binda et al. Biochem J. 2019 476:293-306. | RNAi | primary hippocampal neuron | decreased density of inhibitory synapses |
| Increase | Rasgrf1 | 19417 | 2.141 | Megill et al. J Neurosci. 2013 33:9306-18. | RNAi | primary hippocampal neuron | decreased spine density |
| Increase | Stx1a | 20907 | 2.174 | Fujiwara et al. J Neurosci. 2006 26:5767-76. | knockout | hippocampal CA1 region | impaired LTP |
| Increase | Rab11fip5 | 52055 | 2.177 | Bacaj et al. J Neurosci. 2015 35:7460-74. | knockout | acute hippocampal slice, primary hippocampal neuron | impaired LTD |
| Increase | Rasgrf2 | 19418 | 2.236 | Rosa et al. Front Mol Neurosci. 2016 9:66. | RNAi | primary cortical and hippocampal neuron | decreased spine density |
| Increase | Grm1 | 14816 | 2.268 | Ichikawa et al. Proc Natl Acad Sci U S A.. 2016 113:2282-7 | knockout | cerebellum | defective synapse elimination |
| Increase | Rgs14 | 51791 | 2.325 | Evans et al.eNeuro. 2018 5. pii: ENEURO.0353-17.2018. | overexpression, knockout | hippocampal CA2 region | decreased structural plasticity of spines (overexpression), enhanced LTP, enhanced structural plasticity of spines (knockout) |
| Increase | Git1 | 216963 | 2.329 | Zhang et al. J Neurosci. 2005 25:3379-88. | RNAi | primary hippocampal neuron | decreased spine density |
| Increase | Nckipsd | 80987 | 2.366 | Lee et al. EMBO J. 2006 25:4983-95. | overexpression, RNAi | primary hippocampal neuron | increased spine density (overexpression), decreased spine density and spine length (RNAi) |
| Increase | Cdc42ep4 | 56699 | 2.447 | Ageta-Ishihara et al. Nat Commun. 2015 6:10090. | knockout | cerebellum | insufficient glutamate-buffering/clearance capacity |
| Increase | Nrn1 | 68404 | 2.491 | An et al. Cell Death Dis. 2014 5:e1523. | overexpression | primary hippocampal neuron | increased spine density and diameter, increased frequency of mEPSC |
| Increase | Map3k5 | 26408 | 2.505 | Cho et al. Sci Rep. 2016 6:18840. | RNAi | primary cortical and striatal neuron | increased number of PSD-95 positive structure in astrocyte-conditioned media |
| Increase | Itpr1 | 16438 | 2.505 | Sugawara et al. J Neurosci. 2013 33:12186-96. | knockout | cerebellum | increased spine density and length, impaired LTD |
| Increase | Dlgap1 | 224997 | 2.530 | Shin et al. Nat Neurosci. 2012 15:1655-66. | RNAi | primary hippocampal neuron | impaired synaptic scaling |
| Increase | Fga | 14161 | 2.607 | Seeley et al. Neuron. 2019 101:1099-1108.e6. | knockout | plasma injection into somatosensory cortex | defective synapse elimination |
| Increase | Rapgef2 | 76089 | 2.625 | Lee et al. Neuron. 2011 69:957-73. | overexpression, RNAi | primary hippocampal neuron | decreased spine density (overexpression, with Plk2 shRNA), increased spine density (RNAi, with Plk2 overexpression) |
| Increase | Gpm6a | 234267 | 2.699 | Alfonso et al. Proc Natl Acad Sci U S A.. 2005 102:17196-201. | overexpression, RNAi | primary hippocampal neuron | increased filopodia/spine density (overexpression), decreased filopodia/spine density (RNAi) |
| Increase | Idh3a | 67834 | 2.757 | Ugur et al. Cell Rep. 2017 21:3794-3806. | mutation | fly eyes, fly neuromuscular junctions | decreased amplitude of electroretinograms, decreased number of captate projections<br>impaired release of synaptic vesicles upon stimulation |
| Increase | Gad2 | 14417 | 2.878 | Choi et al. J Neurosci. 2002 22:5271-6. | knockout | visual cortex | impaired LTD |
| Increase | Pacsin1 | 23969 | 2.908 | Kehoe et al. J Neurosci. 2014 34:9213-21. | RNAi | hippocampal CA1 region | decreased density and stability of spines |
| Increase | Pde2a | 207728 | 3.223 | Maurin et al. Cereb Cortex. 2018 Aug 23. doi: 10.1093/cercor/bhy192 | inhibitor treatment | primary cortical neuron | increased spine density, enlargement of spine head (in Fmr1 KO), restoration of LTP (in Fmr1 KO) |
| Increase | Hcn1 | 15165 | 3.311 | Nolan et al. Cell. 2004 119:719-32. | knockout | hippocampal CA1 region | enhanced LTP (at synapses in the perforant path) |
| Increase | Arc | 11838 | 3.324 | Peebles et al. Proc Natl Acad Sci U S A.. 2010 107:18173-8. | overexpression, knockout | primary hippocampal neuron, hippocampus | increased spine density, increased ratio of thin spines (overexpression),<br>decreased spine density, decreased ratio of thin spines (knockout) |
| Increase | Ncan | 13004 | 3.468 | Mohan et al. Front Cell Neurosci. 2018 12:346. | addition of recombinant protein | primary cortical neuron | increased spine density (in the presence of Sema3F-Fc) |
| Increase | Dgkz | 104418 | 3.602 | Kim et al. EMBO J. 2009 28:1170-9. | overexpression, RNAi, knockout | hippocampal CA1 region, primary hippocampal neuron | increased spine density (overexpression), decreased spine density (RNAi, knockout)<br>reduced excitatory synaptic transmission (knockout), increased spine elimination (RNAi) |
| Increase | Palm | 18483 | 3.940 | Arstikaitis et al. Mol Biol Cell. 2008 19:2026-38. | overexpression, RNAi | primary hippocampal neuron | increased filopodia/spine density (overexpression), decreased filopodia/spine density (RNAi) |
| Increase | Efhd2 | 27984 | 3.974 | Mielenz et al. Mol Psychiatry. 2018 23:1303-1319. | overexpression, RNAi | primary cortical neuron | increased spine length (overexpression), increased spine number (overexpression, RNAi) |
| Increase | Tnr | 21960 | 4.175 | David et al.J Neurosci. 2013 33:10324-39. | knockout | olfactory bulb | decreased spine density in adult mice |
| Increase | Iqgap2 | 544963 | 4.480 | Dejanovic et al. Neuron. 2018 100:1322-1336.e7. | RNAi | primary hippocampal neuron | decreased spine density |
| Increase | Pde4a | 18577 | 4.605 | Havekes et al. J Neurosci. 2016 36:8936-46. | overexpression | hippocampal CA1 region | impaired LTP |
| Increase | Camkk2 | 207565 | 6.068 | Mairet-Coello et al. Neuron. 2013 78:94-108. | overexpression, knockout | primary cortical and hippocampal neuron, hippocampal CA3 region | decreased spine density (overexpression ;with Abeta), increased spine density (knockout ;with Abeta) |

**Supplementary Table 3 Quantitative PSD proteomics in ASD model mice (*15q dup*)**

| scription of protein | | Detection by LC-MS/MS | | | | | | | | | | | LFQ intensity normalized with total peptide abundance | | | | | | | | efficient of Variation (C | | Mouse Gene ID | | Mean relative abudance | | | | |
|---|---|---|---|---|---|---|---|---|---|---|---|---|---|---|---|---|---|---|---|---|---|---|---|---|---|---|---|---|---|
| Accession | Description | -value: Com | PEP Sc | overage [% | # Peptides | # PSMs | nique Pepti | # AAs | MW [kDa] | calc. pI | Mascot: M | Search Er | WT1 | WT2 | WT3 | WT4 | 15q dup1 | 15q dup2 | 15q dup3 | 15q dup4 | CVWT | CVPat | Gene Symb | Gene ID | WT | 15q dup | Fold | P value | rected P value |
| 148698430 | mCG1040588, isoform CRA_a [Mus musculus] | 0 | 2214.91 | 62 | 266 | 2168 | 2 | 5324 | 607 | 5.48 | 47531 | 266 | 26400085.15 | 14791822.54 | 24235495.94 | 14412535.8 | 22536710.5 | 14109121.97 | 20463926 | 10443587.93 | 31.3195 | 33.1357 | Macf1 | 11426 | 1.08336 | 0.916640755 | 0.84611 | 0.49165 | 0.87078 |
| 71896543 | SH3 and multiple ankyrin repeat domains protein 3 [Mus musculus] | 0 | 827.726 | 69 | 81 | 1184 | 75 | 1730 | 185.3 | 8.88 | 20259 | 81 | 3572270448 | 3634566869 | 3395623596 | 3465263888 | 3195961185 | 3925502012 | 3375395995 | 3203752390 | 3.03971 | 10.0343 | Shank3 | 58234 | 1.01322 | 0.986779431 | 0.9739 | 0.62828 | 0.912476 |
| 226958349 | triosephosphate isomerase [Mus musculus] | 0 | 61.276 | 43 | 8 | 43 | 8 | 299 | 32.2 | 5.74 | 637 | 8 | 55795521.63 | 60892078.39 | 63489221.04 | 39108743.31 | 65823901.38 | 61671918.22 | 61563625 | 31144619.18 | 19.9771 | 29.1739 | Yars | 107271 | 0.99791 | 1.002089923 | 1.00419 | 0.98192 | 0.997464 |
| 28972451 | mKIAA0893 protein [Mus musculus] | 0 | 147.829 | 32 | 23 | 131 | 23 | 1132 | 125.9 | 6.2 | 2190 | 23 | 141009111.8 | 152856229.1 | 175878608.6 | 127140329.7 | 193886436 | 123657490.3 | 172967659.7 | 127499654.4 | 13.8362 | 22.3356 | Arhgap39 | 223666 | 0.98261 | 1.017389941 | 1.0354 | 0.80158 | 0.959776 |
| 148690579 | mCG21690 [Mus musculus] | 0 | 1052.42 | 62 | 69 | 1944 | 2 | 1325 | 146.5 | 8.85 | 51729 | 69 | 375344930.8 | 813472584 | 573699934.9 | 685512246.1 | 562772049.3 | 1021605361 | 598910943.3 | 957877975.3 | 30.3449 | 30.3028 | Syngap1 | 240057 | 0.87599 | 1.124013656 | 1.28314 | 0.29462 | 0.759552 |
| 568955963 | PREDICTED: voltage-dependent P/Q-type calcium channel subunit alpha-1A isoform X2 [Mus musculus] | 0 | 375.189 | 26 | 56 | 520 | 2 | 2464 | 278.3 | 8.94 | 7608 | 56 | 57403562.54 | 26416872.23 | 29621530.44 | 26335763.41 | 31998404.26 | 19153381.27 | 31861147.5 | 16256660.15 | 43.0705 | 33.4345 | Cacna1a | 12286 | 1.16946 | 0.830543446 | 0.7102 | 0.28322 | 0.750719 |
| 568969771 | PREDICTED: calcium/calmodulin-dependent protein kinase type II subunit beta isoform X2 [Mus musculus] | 0 | 523.436 | 82 | 40 | 2137 | 4 | 666 | 72.9 | 7.27 | 54580 | 40 | 1657256459 | 1783629996 | 1428841658 | 1560071369 | 1460668740 | 1620503752 | 1302515608 | 1373747851 | 9.343 | 9.5174 | Camk2b | 12323 | 1.05517 | 0.944830513 | 0.89543 | 0.14925 | 0.693437 |
| 548961515 | peripheral plasma membrane protein CASK isoform 1 [Mus musculus] | 0 | 330.059 | 55 | 45 | 371 | 2 | 926 | 105 | 6.43 | 4294 | 45 | 24939396.89 | 41580313.53 | 29373655.05 | 62969132.12 | 34888840.83 | 50093259.22 | 32903016 | 52636884.38 | 42.8659 | 23.8597 | Cask | 12361 | 0.9646 | 1.035397849 | 1.07339 | 0.77869 | 0.952792 |
| 157951727 | catenin alpha-2 isoform 1 [Mus musculus] | 0 | 740.316 | 74 | 59 | 745 | 2 | 905 | 100.4 | 5.96 | 19403 | 59 | 120216776 | 49520608.45 | 116921017.8 | 48874818.67 | 107497500 | 47603588.77 | 115860212 | 49785945.53 | 47.7746 | 45.5619 | Ctnna2 | 12386 | 1.02253 | 0.977470037 | 0.95593 | 0.89602 | 0.984401 |
| 6671684 | catenin beta-1 [Mus musculus] | 0 | 658.451 | 77 | 40 | 506 | 12 | 781 | 85.4 | 5.86 | 14669 | 40 | 707449973.7 | 859820319.1 | 866728500.6 | 931640330.5 | 861917351.1 | 926509116.9 | 874497115.3 | 844838229.5 | 11.2889 | 4.01512 | Ctnnb1 | 12387 | 0.97932 | 1.0206772 | 1.04223 | 0.50928 | 0.88031 |
| 755497120 | PREDICTED: dynamin-1 isoform X2 [Mus musculus] | 0 | 237.105 | 49 | 36 | 179 | 2 | 868 | 97.7 | 7.34 | 4008 | 36 | 23349270.59 | 1991130.486 | 26362971.81 | 13793298.42 | 32376598.6 | 16508079.07 | 25859607.13 | 13705634.05 | 67.0828 | 38.8568 | Dnm1 | 13429 | 0.8509 | 1.149098772 | 1.35045 | 0.442 | 0.852048 |
| 55153885 | Glyceraldehyde-3-phosphate dehydrogenase [Mus musculus] | 0 | 179.233 | 62 | 15 | 184 | 2 | 333 | 35.8 | 7.71 | 3754 | 15 | 81402075.11 | 39188610.97 | 110650070 | 60565843.29 | 85934948.62 | 63390920.97 | 94509964 | 50991917.06 | 41.7724 | 27.1874 | Gapdh | 14433 | 0.99485 | 1.005149973 | 1.01035 | 0.96831 | 0.99686 |
| 568913023 | PREDICTED: glutamate receptor ionotropic, NMDA 1 isoform X1 [Mus musculus] | 0 | 378.398 | 50 | 51 | 934 | 2 | 959 | 107.9 | 8.97 | 19006 | 51 | 65706601.46 | 136459819.7 | 79070659.12 | 129038716.9 | 95015707.91 | 157066023.4 | 83391388.5 | 127403770.3 | 34.5169 | 28.7469 | Grin1 | 14810 | 0.93976 | 1.06024272 | 1.12821 | 0.60774 | 0.905186 |
| 148678911 | mCG7984 [Mus musculus] | 0 | 18.428 | 13 | 5 | 27 | 5 | 388 | 42.2 | 9.99 | 217 | 5 | 45823244.13 | 62844686.46 | 48509472.41 | 63593668.46 | 33959616.59 | 74751582.02 | 53342769.06 | 79156977.94 | 16.9185 | 34.6085 | Krt2 | 16681 | 0.95576 | 1.044243874 | 1.09258 | 0.67056 | 0.922517 |
| 113195686 | lamin-B2 [Mus musculus] | 0 | 257.053 | 57 | 33 | 597 | 28 | 596 | 67.3 | 5.5 | 11309 | 33 | 460603892.5 | 563320350.8 | 500445618.4 | 540689857.1 | 454622626.4 | 781102415.3 | 581226795.9 | 656963139 | 8.77665 | 22.123 | Lmnb2 | 16907 | 0.90992 | 1.090076567 | 1.19799 | 0.20588 | 0.715071 |
| 1039792234 | PREDICTED: microtubule-associated protein 4 isoform X22 [Mus musculus] | 0 | 145.646 | 34 | 21 | 203 | 16 | 917 | 96.8 | 9.5 | 3270 | 21 | 95781906.76 | 117185658 | 143501286.6 | 107940937.3 | 115954193.6 | 123129373.2 | 136927590.6 | 147584643.8 | 17.4501 | 10.7901 | Map4 | 17758 | 0.9401 | 1.059904532 | 1.12744 | 0.27601 | 0.745606 |
| 6678467 | tubulin alpha-4A chain isoform 1 [Mus musculus] | 0 | 407.375 | 88 | 30 | 708 | 5 | 448 | 49.9 | 5.06 | 14799 | 30 | 536348850.3 | 422783113.3 | 550815220.5 | 438614776 | 515596976.4 | 334669814.7 | 487561193.5 | 368849287.1 | 13.4989 | 20.7095 | Tuba4a | 22145 | 1.06617 | 0.93382521 | 0.87587 | 0.31429 | 0.76536 |
| 14518291 | chapsyn-110 [Mus musculus] | 0 | 525.566 | 69 | 49 | 1215 | 2 | 852 | 94.7 | 6.24 | 21900 | 49 | 132749722.9 | 140209122 | 106053752.9 | 105502042.7 | 99953514.07 | 145419536.1 | 113262465 | 113736536.2 | 14.8492 | 16.3476 | Dlg2 | 23859 | 1.01269 | 0.987310317 | 0.97494 | 0.82566 | 0.968276 |
| 354459836 | Chain A, The Crystal Structure Of The Bet3-Trs31 Heterodimer. | 0 | 10.559 | 18 | 3 | 15 | 3 | 182 | 20.4 | 4.96 | 186 | 3 | 17922753.14 | 15140553.43 | 15783006.5 | 16204691.47 | 24073494.95 | 17231342.95 | 21399856.5 | 16438991.57 | 7.31747 | 18.1563 | Trappc3 | 27096 | 0.90227 | 1.097733706 | 1.21664 | 0.11192 | 0.678142 |
| 568974918 | PREDICTED: SRC kinase signaling inhibitor 1 isoform X21 [Mus musculus] | 0 | 782.321 | 72 | 82 | 2024 | 3 | 1198 | 129.7 | 9.36 | 29895 | 82 | 143414954.3 | 123553026.4 | 102539611.5 | 84346583.14 | 70703692.19 | 71464952.58 | 89005824.5 | 59022746.62 | 22.5619 | 17.078 | Srcin1 | 56013 | 1.21995 | 0.780046162 | 0.63941 | 0.02811 | 0.515682 |
| 359279908 | tropomyosin alpha-3 chain isoform Tpm3.2cy [Mus musculus] | 0 | 85.287 | 38 | 14 | 206 | 2 | 248 | 28.8 | 4.75 | 4488 | 14 | 33529395.58 | 46854899.64 | 24086208.07 | 37015921.62 | 28674720.96 | 19662312.59 | 34032035 | 18681747.79 | 26.5872 | 29.1972 | Tpm3 | 59069 | 1.16672 | 0.833280826 | 0.71421 | 0.14167 | 0.693437 |
| 292630942 | =Myocyte nuclear envelope protein 1; Short=Myne-1; AltName: Full=Nuclear envelope spectrin repeat protein 1; A | 0 | 2686.3 | 54 | 377 | 2154 | 2 | 8799 | 1009.3 | 5.59 | 38448 | 377 | 16458711.32 | 11335335.22 | 14340957.92 | 13566892.73 | 11500293.65 | 11231972.5 | 15034627 | 10757173.33 | 15.1923 | 16.1691 | Syne1 | 64009 | 1.06892 | 0.931076031 | 0.87104 | 0.25955 | 0.745606 |
| 15809016 | myosin regulatory light chain 12B [Homo sapiens] | 0 | 104.805 | 80 | 10 | 88 | 2 | 172 | 19.8 | 4.84 | 2235 | 10 | 12010230.14 | 7726272.129 | 15698912.55 | 4561866.989 | 19073033.05 | 6805733.62 | 15608323 | 5652731.163 | 48.7413 | 55.8859 | Myl12a | 67268 | 0.91803 | 1.081968976 | 1.17858 | 0.67817 | 0.925326 |
| 148710042 | pleckstrin and Sec7 domain containing, isoform CRA_b, partial [Mus musculus] | 0 | 131.712 | 29 | 20 | 114 | 11 | 1029 | 110 | 7.05 | 2633 | 20 | 40333917.89 | 28276551.56 | 39234742.41 | 31137182.74 | 44340443.37 | 37144100.55 | 34945666 | 42628184.65 | 17.128 | 11.1739 | Psd | 73728 | 0.93264 | 1.067359907 | 1.14445 | 0.22524 | 0.718033 |
| 568964398 | PREDICTED: CLIP-associating protein 2 isoform X8 [Mus musculus] | 0 | 307.757 | 41 | 47 | 363 | 2 | 1532 | 167.6 | 8.27 | 5061 | 47 | 19239096.91 | 13740049.4 | 19696714.65 | 13204529.51 | 18613069.11 | 10662751.26 | 20042258 | 11145880.07 | 21.0897 | 32.4297 | Clasp2 | 76499 | 1.04287 | 0.957129605 | 0.91778 | 0.66799 | 0.922517 |
| 159162926 | Chain A, Localisation Of Dynein Light Chains 1 And 2 And Their Pro- Apoptotic Ligands | 0 | 101.514 | 81 | 8 | 145 | 3 | 94 | 10.8 | 7.37 | 2094 | 8 | 211258404.8 | 150690720.4 | 197560157 | 142789316.3 | 180314880.9 | 180035758.6 | 213734158.3 | 174140129.6 | 19.3168 | 9.62906 | Tpgs1 | 110012 | 0.96834 | 1.031661899 | 1.06539 | 0.57172 | 0.895889 |
| 57528312 | regulating synaptic membrane exocytosis protein 1 isoform 1 [Mus musculus] | 0 | 749.588 | 61 | 67 | 1132 | 36 | 1463 | 163.1 | 9.42 | 24711 | 67 | 600357186.1 | 594105699.6 | 629002031.2 | 623922041.6 | 537131271.8 | 636369675.1 | 651471962 | 559425775.4 | 2.81013 | 9.4452 | Rims1 | 116837 | 1.01304 | 0.986963769 | 0.97426 | 0.6119 | 0.905186 |
| 373838741 | regulating synaptic membrane exocytosis protein 2 isoform a [Mus musculus] | 0 | 390.714 | 47 | 54 | 368 | 10 | 1572 | 177.5 | 9.36 | 6917 | 54 | 46793166.79 | 51683094.4 | 54997726.79 | 53955291.99 | 51710214.22 | 67322475.07 | 60365296.09 | 64045812.65 | 7.03609 | 11.0579 | Rims2 | 116838 | 0.92012 | 1.079877287 | 1.17362 | 0.05689 | 0.599194 |
| 13435747 | Rho GDP dissociation inhibitor (GDI) alpha [Mus musculus] | 0 | 23.431 | 30 | 4 | 14 | 4 | 204 | 23.4 | 5.2 | 256 | 4 | 9175500.754 | 9654693.375 | 9892931.711 | 9428740.329 | 9618805.234 | 4363320.834 | 8060750.5 | 5139024.028 | 3.2198 | 36.2768 | Arhgdia | 192662 | 1.16791 | 0.832093478 | 0.71247 | 0.06934 | 0.601634 |
| 568952724 | PREDICTED: SH3 and multiple ankyrin repeat domains protein 2 isoform X6 [Mus musculus] | 0 | 855.75 | 65 | 87 | 1698 | 9 | 1901 | 206.4 | 6.81 | 29274 | 87 | 65361783.19 | 82302202.79 | 76285451.09 | 99759406.68 | 63969832.73 | 108001155.5 | 76150604.75 | 78209105.24 | 17.7693 | 22.9212 | Shank2 | 210274 | 0.99597 | 1.004033377 | 1.0081 | 0.95749 | 0.994906 |
| 189491680 | disks large-associated protein 1 isoform 3 [Mus musculus] | 0 | 324.042 | 63 | 31 | 432 | 2 | 682 | 75.9 | 5.78 | 11269 | 31 | 16081765.12 | 44460724.76 | 19351846.72 | 23388977.62 | 23759931.61 | 29359476.35 | 27377473 | 23141601.33 | 49.4985 | 11.4361 | Dlgap1 | 224997 | 0.99828 | 1.001716436 | 1.00344 | 0.98964 | 0.998251 |
| 1039760383 | PREDICTED: disks large-associated protein 4 isoform X3 [Mus musculus] | 0 | 495.631 | 68 | 47 | 634 | 4 | 935 | 101.5 | 6.64 | 11457 | 47 | 29389013.02 | 66912967.69 | 53786665.43 | 103658079.8 | 93568963.98 | 99670397.66 | 68930590.5 | 133085910 | 48.8609 | 26.7485 | Dlgap4 | 228836 | 0.78196 | 1.218040943 | 1.55768 | 0.13307 | 0.693437 |
| 755525446 | PREDICTED: sorbin and SH3 domain-containing protein 2 isoform X4 [Mus musculus] | 0 | 565.758 | 53 | 66 | 1034 | 2 | 1501 | 167.4 | 7.64 | 13612 | 66 | 24766765.69 | 27172230 | 28260930.92 | 8632991.129 | 23908619.93 | 19275354.45 | 26536485 | 12046843.3 | 41.2781 | 31.0679 | Sorbs2 | 234214 | 1.04142 | 0.958583787 | 0.92046 | 0.76214 | 0.952792 |
| 163644296 | PH and SEC7 domain-containing protein 3 isoform 3 [Mus musculus] | 0 | 143.335 | 57 | 25 | 206 | 2 | 376 | 42.3 | 9.6 | 1509 | 25 | 25676794.65 | 35102411.04 | 37803909.66 | 32528814.83 | 34732471.87 | 30618600.01 | 38976612 | 42314623.48 | 15.8674 | 13.869 | Psd3 | 234353 | 0.94409 | 1.055914096 | 1.11845 | 0.32676 | 0.772948 |
| 124486885 | leucine-rich repeat-containing protein 7 isoform 1 [Mus musculus] | 0 | 1081.01 | 78 | 91 | 1202 | 6 | 1542 | 172.6 | 6.74 | 20497 | 91 | 324302397.2 | 422044084.5 | 277321152.8 | 381658775 | 347334567.7 | 492907828.1 | 339283728.1 | 349894402 | 18.0975 | 19.312 | Lrrc7 | 242274 | 0.95772 | 1.042284435 | 1.0883 | 0.54779 | 0.889301 |
| 38348480 | disks large-associated protein 3 [Mus musculus] | 0 | 488.916 | 63 | 42 | 904 | 2 | 977 | 105.8 | 8.76 | 15455 | 42 | 3494742.593 | 6744763.383 | 15581509.59 | 7818946.57 | 10464623.31 | 11223161.52 | 11223427.5 | 7252500.698 | 60.9069 | 18.8533 | Dlgap3 | 242667 | 0.91161 | 1.088393307 | 1.19393 | 0.57213 | 0.895889 |
| 568947831 | PREDICTED: SH3 and multiple ankyrin repeat domains protein 1 isoform X3 [Mus musculus] | 0 | 909.463 | 56 | 84 | 2239 | 2 | 2158 | 225.3 | 8.47 | 43259 | 84 | 80140032.3 | 53796893.01 | 72966583.57 | 52376512.88 | 65226986.87 | 64709791.11 | 62179796 | 51878850.56 | 21.4031 | 10.2036 | Shank1 | 243961 | 1.03037 | 0.969629758 | 0.94105 | 0.63316 | 0.912586 |
| 755494337 | PREDICTED: neurofascin isoform X1 [Mus musculus] | 0 | 226.295 | 28 | 29 | 171 | 5 | 1466 | 162.9 | 7.09 | 3989 | 29 | 34698424.44 | 40706909.77 | 24233571.68 | 36257738.55 | 29403434.68 | 31886000.13 | 30529580.38 | 31299890.14 | 20.5301 | 3.48464 | Nfasc | 269116 | 1.04933 | 0.950668062 | 0.90597 | 0.40018 | 0.82465 |
| 569000787 | PREDICTED: rho guanine nucleotide exchange factor 33 isoform X6 [Mus musculus] | 0 | 274.433 | 46 | 26 | 242 | 3 | 869 | 97 | 7.15 | 4262 | 26 | 15883622.29 | 13001020 | 16776537.97 | 15455340.64 | 17794010.1 | 15888315.06 | 22263823.25 | 12800629.48 | 10.5725 | 23.0471 | Arhgef33 | 381112 | 0.94124 | 1.058756069 | 1.12485 | 0.40682 | 0.832122 |
| 1039790234 | PREDICTED: protein unc-13 homolog A isoform X2 [Mus musculus] | 0 | 688.771 | 58 | 82 | 695 | 4 | 1735 | 196.2 | 5.27 | 11591 | 82 | 84861829.44 | 46593344.67 | 74669795.46 | 62233440.75 | 68112276.1 | 58480280.28 | 78854778.5 | 36550745.05 | 24.5976 | 29.7598 | Unc13a | 382018 | 1.05165 | 0.948349182 | 0.90177 | 0.60886 | 0.905186 |
| 148708852 | mCG113425, partial [Mus musculus] | 0 | 152.541 | 38 | 23 | 157 | 3 | 857 | 96.2 | 9.77 | 3064 | 23 | 22243494.89 | 18373904.29 | 28602213.58 | 21221464.32 | 19534870.12 | 20430515.35 | 27658957.38 | 19445885.19 | 19.0938 | 18.1586 | Cdkl5 | 382253 | 1.01899 | 0.981010511 | 0.96273 | 0.78308 | 0.952792 |
| 426229900 | PREDICTED: calcium/calmodulin-dependent protein kinase type II subunit alpha isoform X1 [Ovis aries] | 0 | 556.448 | 84 | 40 | 2511 | 4 | 489 | 55.3 | 7.44 | 49369 | 40 | 1737611965 | 1638365398 | 901807382.1 | 1810519733 | 1371713257 | 1627029252 | 1146617002 | 1288685167 | 27.5603 | 14.8473 | CAMK2A | 12322 | 1.05678 | 0.943218193 | 0.89254 | 0.50851 | 0.88031 |
| 116089341 | abl interactor 1 isoform 1 [Mus musculus] | 0 | 131.581 | 40 | 16 | 134 | 12 | 481 | 52.3 | 7.64 | 2386 | 16 | 95671018.35 | 105018428.1 | 102153254.8 | 92664180.55 | 89589006.68 | 105879895.7 | 98011785.13 | 107302098.9 | 5.76057 | 8.1507 | Abi1 | 11308 | 0.99337 | 1.00662561 | 1.01334 | 0.79992 | 0.959609 |
| 148707429 | v-abl Abelson murine leukemia viral oncogene 2 (arg, Abelson-related gene), partial [Mus musculus] | 0.001 | 5.451 | 2 | 2 | 9 | 2 | 1254 | 136.4 | 8.69 | 26 | 2 | 1842824.517 | 1916807.453 | 3966391.577 | 1852251.399 | 3555579.44 | 3046291.312 | 3427689.75 | 2798273.643 | 43.7823 | 10.8482 | Abl2 | 11352 | 0.85497 | 1.145030053 | 1.33926 | 0.19172 | 0.715071 |
| 23956084 | very long-chain specific acyl-CoA dehydrogenase, mitochondrial precursor [Mus musculus] | 0 | 15.721 | 9 | 4 | 6 | 4 | 656 | 70.8 | 8.75 | 25 | 4 | 3186687.354 | 304478.2023 | 2312126.15 | 3720856.376 | 4550582.627 | 6978510.487 | 1161688.75 | 9304239.599 | 63.0492 | 63.33 | Acadvl | 11370 | 0.60434 | 1.395659961 | 2.3094 | 0.15122 | 0.693437 |
| 77404288 | acid-sensing ion channel 2 isoform MDEG1 [Mus musculus] | 0 | 36.425 | 18 | 6 | 57 | 5 | 512 | 57.7 | 5.19 | 433 | 6 | 27305603.16 | 18050644.62 | 21539492.56 | 13868140.73 | 23288464.69 | 15832635.61 | 22364413.81 | 12264344.74 | 28.1617 | 28.6734 | Asic2 | 11418 | 1.04539 | 0.954605835 | 0.91315 | 0.66735 | 0.922517 |
| 568991060 | PREDICTED: acid-sensing ion channel 1 isoform X2 [Mus musculus] | 0 | 94.639 | 38 | 14 | 116 | 13 | 527 | 59.8 | 5.58 | 1376 | 14 | 63302054.21 | 88370681.94 | 89691302.52 | 76754707.71 | 102778899.6 | 102593743.5 | 101088737.7 | 83716924.53 | 15.4419 | 9.48229 | Asic1 | 11419 | 0.89826 | 1.101736353 | 1.22652 | 0.05757 | 0.599194 |
| 18079339 | aconitate hydratase, mitochondrial precursor [Mus musculus] | 0 | 90.492 | 27 | 16 | 83 | 16 | 780 | 85.4 | 7.93 | 981 | 16 | 66781059.92 | 70899659.34 | 101431118.2 | 86732637.24 | 91983324.69 | 81259481.71 | 82386730.38 | 68634831.78 | 19.4568 | 11.8213 | Aco2 | 11429 | 1.00243 | 0.997569474 | 0.99515 | 0.96736 | 0.99686 |
| 66793429 | peroxisomal acyl-coenzyme A oxidase 1 isoform 1 [Mus musculus] | 0 | 93.501 | 35 | 13 | 56 | 13 | 661 | 74.7 | 8.12 | 977 | 13 | 28990274.15 | 27376393.75 | 56272933.91 | 38210356.72 | 52125176.03 | 45879058.44 | 48137623.88 | 55328573.94 | 35.1663 | 8.33082 | Acox1 | 11430 | 0.85632 | 1.143677389 | 1.33557 | 0.11869 | 0.678142 |
| 157951643 | alpha-actinin-2 [Mus musculus] | 0 | 444.664 | 74 | 56 | 662 | 28 | 894 | 103.8 | 5.45 | 10359 | 56 | 483628388.8 | 524075552.8 | 590418374.9 | 516523187.8 | 595359412.7 | 448983176.8 | 589813813.4 | 579905827.2 | 8.46659 | 12.643 | Actn2 | 11472 | 0.97703 | 1.022966837 | 1.04701 | 0.57146 | 0.895889 |
| 568932688 | PREDICTED: disintegrin and metalloproteinase domain-containing protein 22 isoform X1 [Mus musculus] | 0 | 103.697 | 22 | 16 | 86 | 16 | 980 | 108.1 | 7.66 | 1320 | 16 | 90567749.95 | 72809079.01 | 126871054.1 | 82009724.95 | 90416638.85 | 66236402.22 | 96706535.56 | 89998147.54 | 25.4401 | 15.6386 | Adam22 | 11496 | 1.04038 | 0.959615337 | 0.92237 | 0.61454 | 0.905186 |
| 568940354 | PREDICTED: pituitary adenylate cyclase-activating polypeptide type I receptor isoform X1 [Mus musculus] | 0.003 | 3.681 | 3 | 2 | 6 | 2 | 523 | 59.7 | 6.86 | 28 | 2 | 6541881.434 | 5309549.549 | 8029422.642 | 8213263.065 | 9139936.003 | 8542884.125 | 7311227.5 | 6476968.725 | 19.4474 | 15.249 | Adcyap1r1 | 11517 | 0.94331 | 1.056692556 | 1.1202 | 0.38886 | 0.81965 |
| 31560504 | beta-adducin isoform 1 [Mus musculus] | 0 | 258.409 | 56 | 33 | 292 | 30 | 725 | 80.6 | 6.21 | 5672 | 33 | 650926247.3 | 742131936.2 | 790820875.8 | 791944047.4 | 677795483.4 | 464649174.2 | 642006236.9 | 779213267.2 | 8.90166 | 20.4543 | Add2 | 11519 | 1.0744 | 0.925596192 | 0.8615 | 0.21016 | 0.716066 |
| 20806109 | poly [ADP-ribose] polymerase 1 [Mus musculus] | 0 | 86.519 | 22 | 16 | 40 | 16 | 1014 | 112.7 | 8.95 | 403 | 16 | 44114853.84 | 25365434.31 | 47721020.35 | 28991172.42 | 30280838.32 | 29335206.49 | 47750815.75 | 39808773.26 | 30.149 | 23.6518 | Parp1 | 11545 | 0.99665 | 1.00335126 | 1.00673 | 0.97321 | 0.996906 |
| 568947203 | PREDICTED: aggrecan core protein isoform X1 [Mus musculus] | 0 | 19.315 | 3 | 6 | 19 | 6 | 2170 | 225.9 | 4.28 | 81 | 6 | 11917181.84 | 9018735.176 | 10519721.06 | 15260339.31 | 13473682.04 | 12035644.79 | 7416270.031 | 9973612.619 | 22.8171 | 24.543 | Acan | 11595 | 1.04259 | 0.957409363 | 0.9183 | 0.62859 | 0.912476 |
| 10946936 | adenylate kinase isoenzyme 1 isoform 1 [Mus musculus] | 0 | 20.241 | 23 | 3 | 9 | 3 | 210 | 23.1 | 5.83 | 281 | 3 | 10579178.12 | 8622070.026 | 12678426.69 | 11556893.06 | 9959173.89 | 7507107.876 | 12354002.75 | 8767845.622 | 15.843 | 21.3927 | Ak1 | 11636 | 1.05911 | 0.940890514 | 0.88838 | 0.40168 | 0.82599 |
| 568971246 | PREDICTED: fatty aldehyde dehydrogenase isoform X1 [Mus musculus] | 0 | 64.468 | 32 | 11 | 63 | 11 | 507 | 56.6 | 9.09 | 952 | 11 | 56171871.39 | 63120382.25 | 67459144.29 | 56263280.46 | 57792503.12 | 47875966.31 | 51092042.25 | 33322337.49 | 9.10126 | 21.733 | Aldh3a2 | 11671 | 1.12222 | 0.877783118 | 0.78219 | 0.06462 | 0.601634 |
| 6671539 | fructose-bisphosphate aldolase A isoform 2 [Mus musculus] | 0 | 217.05 | 82 | 23 | 323 | 19 | 364 | 39.3 | 8.09 | 4334 | 23 | 356373113.5 | 397553947.5 | 426463219.7 | 421709951.4 | 321067954.6 | 251545599.9 | 397808308.3 | 411934077.4 | 7.99966 | 21.5076 | Aldoa | 11674 | 1.07363 | 0.926370407 | 0.86284 | 0.22347 | 0.718033 |
| 568949368 | PREDICTED: AMP deaminase 3 isoform X3 [Mus musculus] | 0 | 24.582 | 8 | 6 | 23 | 6 | 816 | 94 | 7.55 | 261 | 6 | 14312050.95 | 10592086.62 | 16354602.44 | 11990470.55 | 14388752.41 | 14541379.5 | 14910332.5 | 15783144.68 | 19.1033 | 4.18943 | Ampd3 | 11717 | 0.94353 | 1.056474168 | 1.11971 | 0.26925 | 0.745606 |
| 568953825 | PREDICTED: ankyrin-1 isoform X2 [Mus musculus] | 0 | 226.857 | 27 | 39 | 268 | 33 | 1932 | 212.1 | 6.65 | 4227 | 39 | 138964496.1 | 111937323.5 | 147422495.6 | 144099214.8 | 134265375.4 | 101390231.4 | 120899520.1 | 137259922.5 | 11.9155 | 13.2344 | Ank1 | 11733 | 1.04691 | 0.95309142 | 0.91039 | 0.33091 | 0.776577 |
| 148747424 | ADP/ATP translocase 1 [Mus musculus] | 0 | 397.349 | 86 | 36 | 1282 | 22 | 298 | 32.9 | 9.72 | 22488 | 36 | 8621223030 | 6621238779 | 6180959052 | 6317551160 | 6608081694 | 9023929174 | 6547989944 | 7798923807 | 16.4226 | 15.6238 | Slc25a4 | 11739 | 0.96123 | 1.038772637 | 1.08067 | 0.51891 | 0.880629 |
| 22094075 | ADP/ATP translocase 2 [Mus musculus] | 0 | 319.633 | 79 | 30 | 1070 | 16 | 298 | 32.9 | 9.73 | 18584 | 30 | 3187984830 | 3054898692 | 3151564449 | 3470025840 | 2897314282 | 3291304044 | 2999821199 | 3034655784 | 5.54531 | 5.48116 | Slc25a5 | 11740 | 1.02557 | 0.97443441 | 0.95014 | 0.23788 | 0.731788 |
| 148701560 | annexin A6, isoform CRA_b, partial [Mus musculus] | 0 | 30.585 | 12 | 7 | 23 | 7 | 677 | 76.3 | 5.52 | 197 | 7 | 17390429.05 | 10080659.06 | 14934055.35 | 7948121.051 | 18592981.73 | 12626282.65 | 10156268.94 | 13629281.49 | 34.4385 | 25.761 | Anxa6 | 11749 | 0.95585 | 1.044149915 | 1.09238 | 0.69228 | 0.930425 |
| 339895913 | AP-1 complex subunit beta-1 isoform 1 [Mus musculus] | 0 | 98.001 | 21 | 20 | 94 | 5 | 953 | 104.9 | 5.06 | 859 | 20 | 6421545.791 | 18985512.55 | 10406701.16 | 11515199.5 | 8180034.967 | 11330698.11 | 6172271.438 | 11382797.76 | 44.3408 | 27.5093 | Ap1b1 | 11764 | 1.12161 | 0.878391069 | 0.78315 | 0.41284 | 0.835788 |
| 56744242 | AP-1 complex subunit gamma-1 isoform 1 [Mus musculus] | 0 | 16.344 | 5 | 3 | 9 | 3 | 825 | 91.7 | 6.8 | 39 | 3 | 3950892.218 | 4015476.465 | 7806638.381 | 4928859.595 | 7616263.246 | 4078443.781 | 9246251.125 | 4155099.435 | 34.974 | 41.0983 | Ap1g1 | 11765 | 0.90405 | 1.095947382 | 1.21226 | 0.51166 | 0.88031 |
| 6671561 | AP-2 complex subunit alpha-1 isoform a [Mus musculus] | 0 | 218.048 | 46 | 34 | 209 | 25 | 977 | 107.6 | 7.03 | 2500 | 34 | 208398380.1 | 202510075.1 | 266252211 | 198964047.9 | 250246656.2 | 261220412.5 | 267237001.2 | 232759260.2 | 14.4821 | 5.98652 | Ap2a1 | 11771 | 0.9283 | 1.071699234 | 1.15447 | 0.10251 | 0.674486 |
| 6753074 | AP-2 complex subunit mu isoform 1 [Mus musculus] | 0 | 155.401 | 60 | 20 | 130 | 20 | 435 | 49.6 | 9.54 | 2441 | 20 | 205686210.4 | 213484396.8 | 219877396 | 223922841.4 | 209011281 | 246718153 | 248911604.3 | 253664775.7 | 3.69112 | 8.59101 | Ap2m1 | 11773 | 0.94765 | 1.052345133 | 1.11047 | 0.07408 | 0.62326 |
| 568947213 | PREDICTED: AP-3 complex subunit beta-2 isoform X1 [Mus musculus] | 0 | 63.069 | 18 | 14 | 42 | 12 | 1101 | 121.2 | 5.45 | 681 | 14 | 31464140.55 | 27397421.3 | 23058122.36 | 22810661.52 | 24428307.48 | 25330170.67 | 27581471.06 | 23901180.77 | 15.6706 | 6.42042 | Ap3b2 | 11775 | 1.01694 | 0.983059714 | 0.96668 | 0.70627 | 0.934133 |
| 6671565 | AP-3 complex subunit delta-1 [Mus musculus] | 0 | 62.198 | 12 | 11 | 28 | 11 | 1199 | 135 | 7.37 | 301 | 11 | 34803745.46 | 40234580.03 | 43706942.43 | 29577724.2 | 37289734.61 | 44307421.39 | 34850295.13 | 34594556.55 | 16.7212 | 11.998 | Ap3d1 | 11776 | 0.99092 | 1.00908261 | 1.01833 | 0.8653 | 0.978314 |
| 148684830 | amyloid beta (A4) precursor protein-binding, family B, member 1, isoform CRA_c, partial [Mus musculus] | 0 | 8.254 | 4 | 2 | 8 | 2 | 725 | 78.9 | 5.08 | 19 | 2 | 3772773.754 | 2500344.31 | 3852323.939 | 2534822.09 | 3996691.428 | 2372297.601 | 3875361 | 1887128.352 | 23.6484 | 35.0385 | Apbb1 | 11785 | 1.02133 | 0.978670894 | 0.95823 | 0.84554 | 0.973079 |
| 569003029 | PREDICTED: adenomatous polyposis coli protein isoform X1 [Mus musculus] | 0 | 156.106 | 15 | 30 | 122 | 30 | 2870 | 312.8 | 8.05 | 1248 | 30 | 65318803.74 | 40427755.14 | 63434969.01 | 61805250.14 | 72946476.01 | 47763058.34 | 57919157.44 | 49611790.32 | 20.1481 | 20.1104 | Apc | 11789 | 1.00598 | 0.994019745 | 0.98811 | 0.93577 | 0.994906 |
| 148667988 | SPEG complex locus, isoform CRA_b, partial [Mus musculus] | 0 | 24.828 | 3 | 7 | 14 | 7 | 3293 | 357.1 | 8.21 | 43 | 7 | 10084121.16 | 10591890.96 | 13342415.8 | 8937967.558 | 5230204.294 | 5059043.441 | 4655888.563 | 4952078.905 | 17.3775 | 4.84914 | Speg | 11790 | 1.36687 | 0.633128789 | 0.4632 | 0.00087 | 0.336276 |
| 163644329 | apolipoprotein E precursor [Mus musculus] | 0 | 9.027 | 8 | 2 | 8 | 2 | 311 | 35.8 | 5.68 | 83 | 2 | 1117663.178 | 1887903.352 | 3342452.444 | 2865037.364 | 4165638.251 | 1820596.033 | 3874589.125 | 3703016.618 | 43.2257 | 31.3829 | Apoe | 11816 | 0.80898 | 1.191017413 | 1.47224 | 0.18611 | 0.715071 |
| 755564427 | PREDICTED: serine/threonine-protein kinase A-Raf isoform X1 [Mus musculus] | 0 | 36.024 | 14 | 6 | 39 | 5 | 641 | 71.5 | 9.04 | 699 | 6 | 21701761.1 | 23640525.06 | 23001127.75 | 22389799.15 | 25667436.58 | 25523795.39 | 25419505.73 | 28498398.67 | 3.6922 | 5.64833 | Araf | 11836 | 0.92659 | 1.073405591 | 1.15844 | 0.00552 | 0.38143 |
| 6671569 | 60S acidic ribosomal protein P0 [Mus musculus] | 0 | 126.277 | 67 | 14 | 172 | 8 | 317 | 34.2 | 6.25 | 2295 | 14 | 159171307.6 | 209747709.9 | 139371307.3 | 218984885.1 | 127782050 | 166592217.7 | 151199320.3 | 175854292.4 | 21.2446 | 13.5225 | Rplp0 | 11837 | 1.07848 | 0.921519176 | 0.85446 | 0.27406 | 0.745606 |
| 9055166 | activity-regulated cytoskeleton-associated protein [Mus musculus] | 0 | 87.903 | 44 | 13 | 99 | 13 | 396 | 45.3 | 5.39 | 1448 | 13 | 255703471 | 270910676 | 257078546.6 | 233494057.1 | 223067568.6 | 301938245.8 | 242388059.1 | 247164601 | 6.08569 | 13.3427 | Arc | 11838 | 1.00129 | 0.998706395 | 0.99742 | 0.97297 | 0.996906 |
| 4502201 | ADP-ribosylation factor 1 [Homo sapiens] | 0 | 27.248 | 41 | 7 | 36 | 6 | 181 | 20.7 | 6.8 | 651 | 7 | 14117052.27 | 7045424.998 | 12170745.83 | 9146689.799 | 25925544.16 | 8461053.162 | 23803503.5 | 6868737.496 | 29.5687 | 61.4158 | Arf1 | 11840 | 0.79004 | 1.20996083 | 1.53152 | 0.32239 | 0.771047 |
| 148704663 | mCG3164, partial [Mus musculus] | 0 | 14.295 | 20 | 4 | 22 | 3 | 191 | 21.5 | 8.38 | 200 | 4 | 11712151.55 | 8959953.905 | 13475304.67 | 8075649.363 | 13580029.8 | 8011274.393 | 12420316.75 | 5341281.834 | 23.5625 | 39.0278 | Arf6 | 11845 | 1.03518 | 0.964816146 | 0.93202 | 0.76438 | 0.952792 |
| 4506711 | 40S ribosomal protein S27 [Homo sapiens] | 0 | 24.439 | 39 | 4 | 88 | 2 | 84 | 9.5 | 9.45 | 839 | 4 | 12657562.6 | 20669401.37 | 41043986.71 | 73179735.71 | 45701866.52 | 41543432.18 | 38610306.5 | 77013933.81 | 71.5628 | 35.8142 | Rhoa | 11848 | 0.84696 | 1.153040011 | 1.36139 | 0.44243 | 0.852048 |
| 4757764 | rho-related GTP-binding protein RhoB precursor [Homo sapiens] | 0 | 11.297 | 18 | 3 | 4 | 2 | 196 | 22.1 | 5.24 | 43 | 3 | 9604025.311 | 6285040.356 | 4197455.643 | 2882431.173 | 5324429.016 | 5400781.634 | 3485013.25 | 3736065.074 | 51.0434 | 22.6724 | Rhob | 11852 | 1.12276 | 0.877242237 | 0.78133 | 0.44916 | 0.854594 |
| 40254129 | armadillo repeat protein deleted in velo-cardio-facial syndrome homolog isoform 1 [Mus musculus] | 0 | 264.291 | 45 | 32 | 151 | 32 | 962 | 105 | 6.64 | 4077 | 32 | 285583998.1 | 214995369.7 | 269026563.2 | 202092616.5 | 279956914.8 | 201938222.4 | 283238682.1 | 207954618.5 | 16.7187 | 18.2278 | Arvcf | 11877 | 0.99929 | 1.000714675 | 1.00143 | 0.99115 | 0.998251 |
| 755495397 | PREDICTED: astrotactin-1 isoform X1 [Mus musculus] | 0 | 141.515 | 27 | 22 | 82 | 22 | 1318 | 146.7 | 5.27 | 2086 | 22 | 79009786.75 | 44474859.84 | 66932063.62 | 68939693.29 | 67804581.65 | 53239482.87 | 66249083.56 | 50597205.76 | 22.4684 | 14.8173 | Astn1 | 11899 | 1.04317 | 0.956830187 | 0.91723 | 0.55168 | 0.892044 |
| 21450277 | sodium/potassium-transporting ATPase subunit alpha-1 precursor [Mus musculus] | 0 | 168.162 | 31 | 25 | 201 | 10 | 1023 | 112.9 | 5.45 | 5661 | 25 | 46753633.64 | 59923724.57 | 49160491.34 | 58554593.5 | 53241978.38 | 59273507.09 | 50678668.75 | 60784695.54 | 12.3346 | 8.59777 | Atp1a1 | 11928 | 0.97813 | 1.021868236 | 1.04471 | 0.57918 | 0.897763 |
| 6753138 | sodium/potassium-transporting ATPase subunit beta-1 [Mus musculus] | 0 | 49.213 | 20 | 8 | 77 | 8 | 304 | 35.2 | 8.65 | 1405 | 8 | 84928812.64 | 70213050.78 | 61110574.38 | 76670377.06 | 71823706.07 | 48591906.9 | 65849680.77 | 47756731.4 | 13.7616 | 22.776 | Atp1b1 | 11931 | 1.11834 | 0.881664488 | 0.78837 | 0.11054 | 0.678142 |
| 7242138 | sodium/potassium-transporting ATPase subunit beta-2 [Mus musculus] | 0 | 19.82 | 21 | 4 | 15 | 4 | 290 | 33.3 | 8.31 | 118 | 4 | 3357887.023 | 4614076.568 | 2591543.63 | 4437816.068 | 1905336.639 | 5051324.023 | 2203545.25 | 8444633.748 | 25.3685 | 69.2041 | Atp1b2 | 11932 | 0.92015 | 1.07984737 | 1.17355 | 0.69747 | 0.931521 |
| 60360584 | mKIAA4195 protein [Mus musculus] | 0 | 46.038 | 14 | 10 | 40 | 10 | 1061 | 116.5 | 5.5 | 437 | 10 | 38922653.15 | 42272881.39 | 27680329.85 | 40930662.35 | 32125836.34 | 33158013.38 | 33901915.63 | 32904122.16 | 17.7778 | 2.21825 | Atp2a2 | 11938 | 1.06285 | 0.937151952 | 0.88174 | 0.23418 | 0.730166 |
| 148667089 | ATPase, Ca++ transporting, plasma membrane 2, isoform CRA_c, partial [Mus musculus] | 0 | 98.452 | 19 | 17 | 91 | 10 | 1249 | 137.3 | 5.91 | 1728 | 17 | 34832452.23 | 32978871.41 | 28414422.91 | 14738777.69 | 27385884.94 | 23359427.01 | 24620420.88 | 16301478.74 | 32.7243 | 20.596 | Atp2b2 | 11941 | 1.09523 | 0.904766583 | 0.82609 | 0.38208 | 0.816034 |
| 6680748 | ATP synthase subunit alpha, mitochondrial precursor [Mus musculus] | 0 | 116.2 | 38 | 16 | 105 | 16 | 553 | 59.7 | 9.19 | 1900 | 16 | 139126719.3 | 108910455.6 | 156195541 | 158077082.6 | 162914871 | 145035671.1 | 151040585.3 | 128680998.1 | 16.1956 | 9.69801 | Atp5a1 | 11946 | 0.97795 | 1.022054544 | 1.0451 | 0.6535 | 0.915864 |
| 23272966 | Atp5b protein, partial [Mus musculus] | 0 | 89.46 | 40 | 15 | 59 | 15 | 533 | 56.6 | 5.4 | 783 | 15 | 91154408.92 | 64538521.92 | 77164381.39 | 76324059.68 | 70441509.13 | 60472844.4 | 95129002.5 | 70155260.99 | 14.0887 | 17.2084 | Atp5b | 11947 | 0.9586 | 1.041397532 | 1.08637 | 0.61546 | 0.905186 |
| 12840425 | unnamed protein product [Mus musculus] | 0 | 21.132 | 22 | 4 | 13 | 4 | 298 | 32.8 | 9.01 | 100 | 4 | 1869246.852 | 4252352.395 | 1735964.02 | 4391192.515 | 2865454.053 | 4942396.774 | 2394982.688 | 4550350.033 | 47.5661 | 33.6188 | Atp5c1 | 11949 | 0.90893 | 1.091066833 | 1.20038 | 0.54432 | 0.88733 |
| 74195489 | unnamed protein product [Mus musculus] | 0 | 14.482 | 18 | 4 | 13 | 4 | 256 | 28.9 | 9.26 | 91 | 4 | 8193876.614 | 6675214.913 | 4948398.29 | 13067557.83 | 10075673.1 | 9551134.735 | 34496970.25 | 11311131.33 | 42.4793 | 74.0562 | Atp5f1 | 11950 | 0.66894 | 1.331060578 | 1.98981 | 0.24426 | 0.733806 |
| 31560731 | V-type proton ATPase catalytic subunit A [Mus musculus] | 0 | 139.532 | 48 | 21 | 74 | 21 | 617 | 68.3 | 5.58 | 1409 | 21 | 11862374.2 | 96354161.48 | 90402893.63 | 100278192.3 | 99913493.24 | 118811034.7 | 92945837.5 | 110248833 | 9.50528 | 10.7893 | Atp6v1a | 11964 | 0.9732 | 1.026796211 | 1.05507 | 0.48574 | 0.869 |
| 74191447 | unnamed protein product [Mus musculus] | 0 | 90.67 | 41 | 16 | 89 | 16 | 521 | 57.5 | 5.81 | 1438 | 16 | 103574905 | 99111765.56 | 71932025.77 | 94701175.58 | 106655905.9 | 110622695 | 92135745.56 | 82888890.32 | 15.242 | 13.1233 | Atp6v1b2 | 11966 | 0.96982 | 1.030176822 | 1.06223 | 0.56885 | 0.895889 |
| 31981304 | V-type proton ATPase subunit d 1 [Mus musculus] | 0 | 77.807 | 46 | 11 | 58 | 11 | 351 | 40.3 | 5 | 1196 | 11 | 68091332.96 | 66213424.74 | 34452211.34 | 58085460.36 | 45159183.86 | 62782795.67 | 41513816.94 | 62237087.98 | 27.2634 | 21.1091 | Atp6v0d1 | 11972 | 1.03455 | 0.965454219 | 0.93322 | 0.70503 | 0.933129 |
| 45504359 | V-type proton ATPase subunit E 1 [Mus musculus] | 0 | 53.791 | 54 | 10 | 52 | 10 | 226 | 26.1 | 8.43 | 860 | 10 | 73113187.2 | 53770774.87 | 46573385.74 | 59846392.01 | 70832315.85 | 60728077.5 | 60192727.5 | 66244244.29 | 19.2922 | 7.79945 | Atp6v1e1 | 11973 | 0.94974 | 1.050261693 | 1.10584 | 0.35515 | 0.798662 |
| 12025532 | V-type proton ATPase 116 kDa subunit a isoform 1 [Mus musculus] | 0 | 127.437 | 29 | 17 | 84 | 17 | 838 | 96.2 | 6.57 | 1282 | 17 | 225706574.9 | 269389158.9 | 182890552 | 183466612.9 | 209854390.2 | 180975578.8 | 166285495.6 | 180218392.7 | 19.145 | 9.93056 | Atp6v0a1 | 11975 | 1.07763 | 0.922366732 | 0.85592 | 0.21799 | 0.716066 |
| 116256512 | heterogeneous nuclear ribonucleoprotein D0 isoform a [Mus musculus] | 0 | 30.711 | 17 | 7 | 22 | 5 | 355 | 38.3 | 7.81 | 159 | 7 | 9482655.285 | 15500493.67 | 13337933.62 | 10084235.94 | 12965776.52 | 13174078.23 | 21919449 | 20040915.81 | 23.3776 | 27.2053 | Hnrnpd | 11991 | 0.83095 | 1.169046909 | 1.40687 | 0.11954 | 0.678336 |
| 148709172 | AU RNA binding protein/enoyl-coenzyme A hydratase, isoform CRA_b, partial [Mus musculus] | 0 | 33.713 | 24 | 7 | 18 | 7 | 317 | 33.8 | 9.5 | 203 | 7 | 58254199.17 | 66054990.21 | 61575735.36 | 89395772.23 | 72318619.49 | 55002373.5 | 110861758.5 | 116498000.5 | 20.3386 | 33.6306 | Auh | 11992 | 0.8736 | 1.12639837 | 1.28937 | 0.27237 | 0.745606 |
| 1199931 | pre-acrosome localization protein [Mus musculus] | 0 | 29.194 | 6 | 5 | 13 | 5 | 1060 | 120.3 | 8.65 | 191 | 5 | 2780036.924 | 4789405.533 | 5374045.087 | 2433700.597 | 2852789.161 | 1594988.8 | 3375617.25 | 4331441.115 | 37.8623 | 37.5388 | Cep131 | 12009 | 1.11704 | 0.882959867 | 0.79045 | 0.41711 | 0.837049 |
| 157951688 | brevican core protein isoform 1 precursor [Mus musculus] | 0 | 181.68 | 36 | 23 | 222 | 23 | 883 | 95.8 | 4.93 | 4849 | 23 | 314599326.8 | 324388554 | 304919520.9 | 352661966.5 | 307531747.8 | 284576843.2 | 296329556.1 | 295931761 | 6.35757 | 3.16549 | Bcan | 12032 | 1.04522 | 0.954776253 | 0.91347 | 0.04794 | 0.592616 |
| 126723336 | prohibitin-2 [Mus musculus] | 0 | 71.73 | 49 | 12 | 56 | 12 | 299 | 33.3 | 9.83 | 959 | 12 | 86054330.61 | 83466365.28 | 42816182.82 | 63759084.44 | 101336077.8 | 158464402.4 | 68899582 | 107602488.2 | 29.1342 | 39.9562 | Phb2 | 12034 | 0.77512 | 1.224883382 | 1.58026 | 0.10607 | 0.674486 |
| 26344399 | unnamed protein product [Mus musculus] | 0 | 16.329 | 10 | 3 | 13 | 3 | 412 | 46.5 | 9.04 | 271 | 3 | 4806249.591 | 5785962.969 | 5611567.338 | 7863160.485 | 6065286.631 | 10495953.25 | 5965283.25 | 8827280.569 | 21.6532 | 28.2234 | Bckdk | 12041 | 0.86852 | 1.131482596 | 1.30277 | 0.20577 | 0.715071 |
| 10946572 | fatty acid-binding protein, brain [Mus musculus] | 0 | 16.171 | 30 | 3 | 19 | 3 | 132 | 14.9 | 5.63 | 287 | 3 | 15383193.04 | 14488140.55 | 16072533.35 | 17193571.73 | 29768178.7 | 23383528.41 | 17599348.25 | 24632701.51 | 7.23274 | 20.9575 | Fabp7 | 12140 | 0.79658 | 1.2034196 | 1.51073 | 0.01994 | 0.505785 |
| 124487407 | protein bassoon [Mus musculus] | 0 | 2119.62 | 66 | 173 | 4725 | 6 | 3942 | 418.6 | 7.71 | 103564 | 173 | 119078942.9 | 107071621.9 | 90025236.38 | 101469706.1 | 80835079.83 | 133331370.6 | 98303218.03 | 106669479.6 | 11.5703 | 20.8653 | Bsn | 12217 | 0.99822 | 1.001784976 | 1.00358 | 0.97712 | 0.997464 |
| 31560618 | mitotic checkpoint protein BUB3 [Mus musculus] | 0 | 17.108 | 25 | 5 | 11 | 5 | 326 | 36.9 | 6.84 | 50 | 5 | 6035429.204 | 7317602.449 | 3122411.879 | 3639924.303 | 3970383.298 | 5409094.351 | 4515400.5 | 5912354.68 | 29.4556 | 17.6261 | Bub3 | 12237 | 0.94591 | 1.054087021 | 1.11436 | 0.544 | 0.88733 |
| 124286805 | complement C1q subcomponent subunit A precursor [Mus musculus] | 0 | 20.765 | 24 | 4 | 20 | 4 | 245 | 26 | 9.11 | 354 | 4 | 16584484.66 | 12392192.21 | 19909598.74 | 11713248.19 | 20002553.88 | 9558383.878 | 22283062.63 | 8764638.895 | 25.3156 | 46.1142 | C1qa | 12259 | 0.99992 | 1.000075205 | 1.00015 | 0.99956 | 0.999562 |
| 50223 | precursor polypeptide (AA -25 to 228) [Mus musculus] | 0 | 13.085 | 16 | 4 | 53 | 3 | 253 | 26.8 | 8.41 | 802 | 4 | 10836331.94 | 19354169.72 | 11794138.21 | 13676868.86 | 15949518.37 | 10891473.94 | 17742865.81 | 9647496.88 | 27.4015 | 28.7672 | C1qb | 12260 | 1.01301 | 0.986985927 | 0.97431 | 0.89998 | 0.985548 |
| 112181167 | complement component 1 Q subcomponent-binding protein, mitochondrial [Mus musculus] | 0 | 73.046 | 47 | 7 | 37 | 7 | 279 | 31 | 4.87 | 1302 | 7 | 41409961.89 | 50624425.04 | 66238724.41 | 39397365.7 | 56439135.22 | 61248348.58 | 50761198 | 39547757.56 | 25.0151 | 17.9661 | C1qbp | 12261 | 0.98802 | 1.011983559 | 1.02426 | 0.88098 | 0.980491 |
| 113680120 | complement C1q subcomponent subunit C precursor [Mus musculus] | 0 | 14.629 | 17 | 3 | 19 | 3 | 246 | 26 | 8.54 | 186 | 3 | 3270402.754 | 12905686.3 | 14309443.12 | 12182995.76 | 9070091.329 | 10691574.1 | 12968733.38 | 18846790.55 | 46.9626 | 33.1806 | C1qc | 12262 | 0.90547 | 1.094525902 | 1.20879 | 0.52415 | 0.88292 |
| 148708308 | calcium binding protein 39, isoform CRA_b, partial [Mus musculus] | 0.003 | 3.958 | 5 | 2 | 4 | 2 | 351 | 40.7 | 6.67 | 0 | 2 | 3552879.59 | 3172602.449 | 3274440.323 | 3049951.489 | 3616954.293 | 3427069.922 | 2776977.75 | 3248434.314 | 6.6975 | 11.0146 | Cab39 | 12283 | 0.99829 | 1.001707745 | 1.00342 | 0.95949 | 0.994906 |
| 1039727362 | PREDICTED: voltage-dependent R-type calcium channel subunit alpha-1E isoform X2 [Mus musculus] | 0 | 221.107 | 21 | 39 | 287 | 3 | 2466 | 277.9 | 8.05 | 3780 | 39 | 15126251.09 | 15314490.49 | 14749352.38 | 9557810.16 | 13838851.46 | 17635865.28 | 12858260.5 | 15226605.58 | 20.1855 | 13.9189 | Cacna1e | 12290 | 0.95791 | 1.042094169 | 1.08789 | 0.51213 | 0.88031 |
| 755510852 | PREDICTED: voltage-dependent calcium channel subunit alpha-2/delta-1 isoform X8 [Mus musculus] | 0 | 82.299 | 19 | 16 | 53 | 16 | 1110 | 125.3 | 5.4 | 701 | 16 | 42188390.61 | 38401067.21 | 15304134.12 | 62502197.26 | 52650224.73 | 34097964.3 | 21465294.69 | 62537019.39 | 50.4007 | 43.1408 | Cacna2d1 | 12293 | 0.95099 | 1.049009048 | 1.10307 | 0.7762 | 0.952792 |
| 226693335 | voltage-dependent L-type calcium channel subunit beta-1 isoform B [Mus musculus] | 0 | 122.007 | 49 | 20 | 206 | 7 | 597 | 65.4 | 6.62 | 2869 | 20 | 34497197.91 | 38919959.66 | 39255420.5 | 42614322.23 | 25454547.73 | 13292272.79 | 43946101.5 | 17086132.61 | 8.57978 | 54.7151 | Cacnb1 | 12295 | 1.21762 | 0.782378461 | 0.64255 | 0.09562 | 0.670764 |
| 568912086 | PREDICTED: voltage-dependent L-type calcium channel subunit beta-2 isoform X7 [Mus musculus] | 0 | 40.033 | 22 | 10 | 73 | 6 | 604 | 68.1 | 8.59 | 566 | 10 | 17872661.52 | 4141836.723 | 14574875.75 | 7263758.878 | 11328822.61 | 6264639.63 | 10184005 | 4862231.986 | 57.9288 | 37.8425 | Cacnb2 | 12296 | 1.14659 | 0.853405432 | 0.7443 | 0.45747 | 0.860877 |
| 113680271 | voltage-dependent L-type calcium channel subunit beta-3 isoform 1 [Mus musculus] | 0 | 138.107 | 56 | 17 | 195 | 7 | 484 | 54.5 | 6.27 | 3803 | 17 | 52863317.73 | 64490665.86 | 58903992.01 | 60997130.3 | 31480763.79 | 13659806.7 | 40073993.13 | 97171331.81 | 8.22553 | 63.4706 | Cacnb3 | 12297 | 0.88258 | 1.117421829 | 1.26609 | 0.53447 | 0.886555 |
| 80861468 | voltage-dependent L-type calcium channel subunit beta-4 isoform a [Mus musculus] | 0 | 195.097 | 58 | 19 | 265 | 14 | 519 | 57.9 | 9.28 | 4539 | 19 | 232549386.1 | 251505502.8 | 231309582.2 | 265798291.6 | 224309123.7 | 251615313.9 | 244008225.5 | 219007869.3 | 6.72773 | 6.63166 | Cacnb4 | 12298 | 1.02199 | 0.978010438 | 0.95697 | 0.37898 | 0.81965 |
| 148697713 | calcium channel, voltage-dependent, gamma subunit 2, isoform CRA_a, partial [Mus musculus] | 0 | 100.036 | 30 | 8 | 125 | 6 | 325 | 36.1 | 9.09 | 1928 | 8 | 227143115.5 | 264690296.1 | 206431085.2 | 290155201.1 | 241269772 | 216859637.7 | 206868409.5 | 210938261.7 | 15.1697 | 7.03845 | Cacng2 | 12300 | 1.06033 | 0.939666227 | 0.8862 | 0.21458 | 0.716066 |
| 33468885 | calcyclin-binding protein [Mus musculus] | 0 | 10.547 | 12 | 3 | 17 | 3 | 229 | 26.5 | 7.87 | 167 | 3 | 3712568.944 | 5737166.045 | 6295852.551 | 4324852.154 | 6323698.197 | 6186799.6 | 6391643.688 | 7094780.93 | 23.9565 | 6.24798 | Cacybp | 12301 | 0.87135 | 1.128648189 | 1.29528 | 0.0582 | 0.599194 |
| 12849385 | unnamed protein product [Mus musculus] | 0 | 23.109 | 21 | 5 | 11 | 5 | 339 | 38.6 | 7.65 | 147 | 5 | 7313475.471 | 5122679.26 | 4502891.802 | 4171827.493 | 6209972.258 | 7544057.663 | 5788193.25 | 9220329.633 | 26.7774 | 21.5043 | Anxa2 | 12306 | 0.84658 | 1.153421958 | 1.36245 | 0.11758 | 0.678142 |
| 34098931 | calretinin [Mus musculus] | 0 | 16.435 | 19 | 5 | 21 | 4 | 271 | 31.4 | 5.02 | 119 | 5 | 6220149.342 | 2660068.654 | 8369791.888 | 3590518.544 | 11043121.83 | 6536584.342 | 11765381.25 | 5652819.142 | 49.7173 | 35.4385 | Calb2 | 12308 | 0.74646 | 1.253541815 | 1.67932 | 0.13033 | 0.693437 |
| 74220435 | unnamed protein product [Mus musculus] | 0 | 153.71 | 84 | 10 | 136 | 3 | 149 | 16.8 | 4.22 | 5166 | 10 | 196933714 | 73534624.54 | 231905908.4 | 89807415.67 | 177332390.5 | 59347306.93 | 181290736 | 48315615.35 | 52.8511 | 62.2826 | Calm1 | 12313 | 1.11894 | 0.881058617 | 0.7874 | 0.57689 | 0.895889 |
| 6680836 | calreticulin precursor [Mus musculus] | 0.001 | 6.221 | 5 | 2 | 6 | 2 | 416 | 48 | 4.49 | 43 | 2 | 3353973.192 | 5219077.182 | 3577992.698 | 3177096.955 | 5057995.559 | 4822879.292 | 4366616.344 | 4442182.677 | 24.5074 | 6.9662 | Calr | 12317 | 0.90118 | 1.098816869 | 1.2193 | 0.14179 | 0.693437 |
| 532049867 | REDICTED: calcium/calmodulin-dependent protein kinase type II subunit gamma isoform X4 [Microtus ochrogaste | 0 | 397.611 | 81 | 34 | 1489 | 2 | 565 | 62.8 | 7.94 | 33077 | 34 | 182324215.6 | 245586801.9 | 247474469.4 | 239902944.8 | 248088238.8 | 219257946.6 | 213665723.8 | 195051413 | 13.6198 | 10.0317 | Camk2g | 12325 | 1.0219 | 0.978103067 | 0.95714 | 0.6254 | 0.910246 |
| 148698452 | CAP, adenylate cyclase-associated protein 1 (yeast), isoform CRA_b, partial [Mus musculus] | 0 | 25.912 | 14 | 5 | 19 | 4 | 494 | 53.8 | 7.99 | 202 | 5 | 9724299.761 | 8456949.463 | 8755966.236 | 9050179.203 | 8747408.553 | 7795856.891 | 6760073 | 7335076.817 | 6.0252 | 11.1082 | Cap1 | 12331 | 1.07985 | 0.920153713 | 0.85212 | 0.03875 | 0.568012 |
| 157951598 | calpain-2 catalytic subunit [Mus musculus] | 0 | 8.783 | 3 | 2 | 4 | 2 | 700 | 79.8 | 4.96 | 52 | 2 | 1218022.727 | 2186389.389 | 1275909.064 | 1549181.076 | 1438715.751 | 2079741.641 | 1187321.625 | 2174054.953 | 28.4782 | 28.0536 | Capn2 | 12334 | 0.95039 | 1.049608285 | 1.1044 | 0.63744 | 0.914499 |
| 18202239 | e small subunit; Short=CANP small subunit; AltName: Full=Calcium-dependent protease small subunit; Short=CDF | 0 | 11.222 | 12 | 3 | 5 | 3 | 269 | 28.4 | 5.63 | 71 | 3 | 5394292.702 | 5170165.064 | 3692831.871 | 4844795.153 | 3907534.241 | 2988008.039 | 3885013.625 | 3433359.269 | 15.8355 | 12.2607 | Capns1 | 12336 | 1.14672 | 0.853278616 | 0.7441 | 0.03115 | 0.515682 |
| 568949484 | PREDICTED: calpain-5 isoform X1 [Mus musculus] | 0 | 57.329 | 23 | 12 | 38 | 12 | 653 | 74.2 | 7.36 | 359 | 12 | 34913494.95 | 30142611.54 | 24792336.69 | 21546688.93 | 32827384.43 | 16900258.73 | 26761751 | 20293316.18 | 21.1656 | 29.1799 | Capn5 | 12337 | 1.07019 | 0.929807987 | 0.86882 | 0.45723 | 0.860877 |
| 161086971 | F-actin-capping protein subunit alpha-1 [Mus musculus] | 0 | 52.885 | 43 | 8 | 47 | 6 | 286 | 32.9 | 5.55 | 494 | 8 | 59408138.78 | 47757941.21 | 69613108.8 | 56992131.98 | 75313221.98 | 42714932.23 | 66269568.63 | 44440066.26 | 15.3673 | 28.2512 | Capza1 | 12340 | 1.01088 | 0.989116903 | 0.97847 | 0.89615 | 0.984401 |
| 6671672 | F-actin-capping protein subunit alpha-2 [Mus musculus] | 0 | 163.878 | 75 | 15 | 137 | 13 | 286 | 32.9 | 5.85 | 2936 | 15 | 545834330.3 | 430646386.3 | 511004466.7 | 465301757.1 | 550778584.9 | 483271207.4 | 507776741.1 | 436295187.4 | 10.3629 | 9.66779 | Capza2 | 12343 | 0.99355 | 1.006445019 | 1.01297 | 0.86159 | 0.976238 |
| 4826659 | F-actin-capping protein subunit beta isoform 1 [Homo sapiens] | 0 | 241.918 | 74 | 24 | 293 | 24 | 272 | 30.6 | 6 | 5158 | 24 | 1092244901 | 873115623.7 | 1120012691 | 812684815.1 | 1106037806 | 975926453.2 | 1052242453 | 982922417.2 | 15.8419 | 5.99289 | Capzb | 12345 | 0.97267 | 1.027332 | 1.0562 | 0.53443 | 0.886555 |
| 6753294 | catenin alpha-1 [Mus musculus] | 0 | 207.632 | 40 | 28 | 234 | 17 | 906 | 100 | 6.23 | 4774 | 28 | 47831097.82 | 39169528.01 | 51225735.43 | 43503177.67 | 58492273.4 | 38608086.24 | 53545224.5 | 31914638.42 | 11.5259 | 27.2933 | Ctnna1 | 12385 | 0.99772 | 1.002280283 | 1.00457 | 0.97648 | 0.997464 |
| 568915171 | PREDICTED: catenin delta-1 isoform X1 [Mus musculus] | 0 | 439.027 | 60 | 45 | 447 | 5 | 967 | 108.1 | 6.23 | 9589 | 45 | 102074005.1 | 16176109.94 | 34940687.79 | 46836384.49 | 31465193.99 | 24895584.26 | 26756351.88 | 24000742.74 | 73.8597 | 12.4275 | Ctnnd1 | 12388 | 1.30249 | 0.697506727 | 0.53552 | 0.25694 | 0.744267 |
| 56744247 | cerebellin-1 precursor [Mus musculus] | 0 | 69.99 | 47 | 8 | 59 | 7 | 193 | 21.1 | 7.28 | 1135 | 8 | 67979299.38 | 87480998.9 | 45193956.88 | 128228470.6 | 63564318.52 | 104927957.4 | 81700025.75 | 65211944.15 | 42.8177 | 24.3707 | Cbln1 | 12404 | 1.02092 | 0.979080005 | 0.95902 | 0.87208 | 0.979509 |
| 161353502 | serpin H1 precursor [Mus musculus] | 0 | 41.371 | 23 | 6 | 20 | 6 | 417 | 46.5 | 8.82 | 156 | 6 | 16299881.94 | 10818524.05 | 14337265.97 | 10182042.85 | 22589979.31 | 21086424.46 | 21798326.25 | 6502942.685 | 22.5151 | 42.7108 | Serpinh1 | 12406 | 0.83546 | 1.164542282 | 1.3939 | 0.26206 | 0.745606 |
| 1049108 | carbonyl reductase [Mus musculus] | 0 | 9.861 | 14 | 3 | 11 | 3 | 277 | 30.7 | 7.74 | 21 | 3 | 5932925.349 | 7778562.744 | 5706536.953 | 6129022.723 | 5738350.651 | 6255109.679 | 5271522 | 4947864.589 | 14.7773 | 10.254 | Cbr1 | 12408 | 1.06981 | 0.93018827 | 0.86949 | 0.18118 | 0.715071 |
| 126521835 | T-complex protein 1 subunit beta [Mus musculus] | 0 | 99.556 | 40 | 16 | 53 | 16 | 535 | 57.4 | 6.4 | 872 | 16 | 47190025.09 | 52331704.56 | 51612121.44 | 50654455.88 | 50636804.69 | 49486666.59 | 51009022.13 | 45956561.93 | 4.51456 | 4.67502 | Cct2 | 12461 | 1.01178 | 0.988218806 | 0.97671 | 0.49554 | 0.87488 |
| 6753320 | T-complex protein 1 subunit gamma [Mus musculus] | 0 | 74.657 | 34 | 14 | 67 | 14 | 545 | 60.6 | 6.7 | 1054 | 14 | 52098977.62 | 42339820.99 | 56898071.13 | 45777012.91 | 55052700.59 | 26678817.39 | 52979966.63 | 30553647.91 | 13.1731 | 35.7585 | Cct3 | 12462 | 1.08789 | 0.912112046 | 0.83842 | 0.36185 | 0.804292 |
| 6753322 | T-complex protein 1 subunit delta [Mus musculus] | 0 | 62.85 | 29 | 13 | 66 | 12 | 539 | 58 | 8.02 | 410 | 13 | 34839291.55 | 31274812.56 | 37726399.13 | 27151592.55 | 32031142.49 | 24601853.17 | 34766521.13 | 24763769.12 | 13.9543 | 17.751 | Cct4 | 12464 | 1.06 | 0.940002076 | 0.8868 | 0.32316 | 0.771047 |
| 6671702 | T-complex protein 1 subunit epsilon [Mus musculus] | 0 | 59.672 | 27 | 10 | 44 | 9 | 541 | 59.6 | 6.02 | 1042 | 10 | 16001906.44 | 16721977.37 | 20052369.42 | 15004778.69 | 18817984.66 | 14088808.23 | 14582616.13 | 12684814.21 | 12.911 | 17.5596 | Cct5 | 12465 | 1.05945 | 0.940551026 | 0.88777 | 0.30994 | 0.763986 |
| 62948125 | Chaperonin containing Tcp1, subunit 6a (zeta) [Mus musculus] | 0 | 75.34 | 28 | 12 | 34 | 12 | 531 | 58 | 6.9 | 770 | 12 | 51784872.07 | 56868304.63 | 41567641.02 | 32193343.17 | 38032606 | 29829221.29 | 38042526.03 | 30842391 | 24.0619 | 13.063 | Cct6a | 12466 | 1.14309 | 0.856914128 | 0.74965 | 0.10221 | 0.674486 |
| 148666717 | chaperonin subunit 7 (eta), isoform CRA_b [Mus musculus] | 0 | 68.526 | 27 | 10 | 42 | 10 | 556 | 61.3 | 8.28 | 720 | 10 | 31825072.62 | 26845307.66 | 31292843.93 | 28027737.61 | 27292039.3 | 13504658.29 | 26738797.56 | 17057845.58 | 8.26589 | 32.7778 | Cct7 | 12468 | 1.16486 | 0.835142108 | 0.71695 | 0.06345 | 0.601634 |
| 50510319 | mKIAA0002 protein [Mus musculus] | 0 | 49.143 | 25 | 12 | 41 | 12 | 555 | 60.2 | 5.62 | 538 | 12 | 111127330 | 118975287.2 | 89927342.56 | 94544080.84 | 104976199.1 | 141725360.8 | 99503789.88 | 136534073.6 | 13.2059 | 17.8314 | Cct8 | 12469 | 0.92403 | 1.075966076 | 1.16442 | 0.22987 | 0.723653 |
| 148705649 | CD38 antigen [Mus musculus] | 0 | 11.49 | 13 | 2 | 3 | 2 | 304 | 34.4 | 8.29 | 38 | 2 | 5942011.839 | 7553129.076 | 4832381.545 | 5408664.893 | 7206645.372 | 6895377.215 | 4789347.125 | 4724059.794 | 19.7275 | 22.544 | Cd38 | 12494 | 1.00255 | 0.997449763 | 0.99491 | 0.97393 | 0.99712 |
| 114687888 | cadherin-11 preproprotein [Mus musculus] | 0 | 103.978 | 26 | 14 | 92 | 14 | 796 | 88.1 | 4.89 | 1580 | 14 | 104710064.3 | 101810211.8 | 94279949.62 | 85227311.25 | 92456767.97 | 98389247.12 | 91415333.98 | 94868477.7 | 9.02552 | 3.28415 | Cdh11 | 12552 | 1.01166 | 0.988340923 | 0.97695 | 0.64738 | 0.915705 |
| 1039781739 | PREDICTED: cadherin-13 isoform X2 [Mus musculus] | 0 | 18.315 | 7 | 3 | 18 | 3 | 731 | 80.1 | 5.14 | 191 | 3 | 9800813.174 | 6592453.672 | 5990789.052 | 5326329.788 | 7702279.511 | 6435237.65 | 4378746 | 6066008.426 | 28.6398 | 22.3015 | Cdh13 | 12554 | 1.05982 | 0.940180625 | 0.88711 | 0.54061 | 0.886555 |
| 119964694 | cadherin-15 preproprotein [Mus musculus] | 0 | 29.353 | 14 | 8 | 23 | 8 | 784 | 85.6 | 4.86 | 157 | 8 | 21422069.51 | 19827284.11 | 20313804.38 | 17841123.98 | 21987702.78 | 17926985.76 | 21833979.75 | 12484432.26 | 7.54092 | 24.0538 | Cdh15 | 12555 | 1.03366 | 0.966341646 | 0.93488 | 0.60272 | 0.905186 |
| 116760627 | cadherin-2 preproprotein [Mus musculus] | 0 | 261.97 | 34 | 22 | 208 | 20 | 906 | 99.7 | 4.78 | 4186 | 22 | 759041551.2 | 630417404 | 758325439 | 630403203.4 | 734156930.6 | 617276536.4 | 680383858.3 | 579052637.6 | 10.6629 | 10.4959 | Cdh2 | 12558 | 1.03105 | 0.968952339 | 0.93977 | 0.43871 | 0.852048 |
| 6753376 | cadherin-4 isoform 1 preproprotein [Mus musculus] | 0 | 57.762 | 11 | 7 | 43 | 5 | 913 | 100 | 4.81 | 609 | 7 | 20518214.07 | 36277030.65 | 33434373.04 | 35529142.83 | 39435097.64 | 37685370.82 | 32207758.25 | 44222245.03 | 23.4726 | 12.9225 | Cdh4 | 12561 | 0.9005 | 1.099501584 | 1.22099 | 0.16914 | 0.711915 |
| 110347562 | cadherin-6 preproprotein [Mus musculus] | 0 | 54.799 | 19 | 14 | 101 | 13 | 790 | 88.3 | 5 | 1303 | 14 | 77400698.34 | 87785519.13 | 67379021.75 | 85345853.26 | 75537096.84 | 87304355.61 | 72238517.25 | 84039992.93 | 11.5807 | 8.85256 | Cdh6 | 12563 | 0.9981 | 1.001897663 | 1.0038 | 0.96014 | 0.994906 |
| 1845579 | cadherin-8 [Mus musculus] | 0 | 54.488 | 14 | 8 | 45 | 8 | 799 | 88.1 | 4.72 | 542 | 8 | 43691658.73 | 37898052.01 | 36707500.55 | 45182243.3 | 46510164.66 | 46802826.99 | 38010874.38 | 44928679.19 | 10.2567 | 9.3456 | Cdh8 | 12564 | 0.9624 | 1.037597549 | 1.07813 | 0.31882 | 0.767321 |
| 755550275 | PREDICTED: cadherin-9 isoform X1 [Mus musculus] | 0 | 58.362 | 16 | 10 | 74 | 8 | 867 | 97.3 | 5.21 | 1242 | 10 | 45768506.56 | 32785030 | 47532817.59 | 33360473.61 | 39995667.34 | 36352763.59 | 45607336 | 29775538.99 | 19.7576 | 17.4995 | Cdh9 | 12565 | 1.02479 | 0.97520545 | 0.95161 | 0.72088 | 0.939018 |
| 6680908 | cyclin-dependent kinase 5 [Mus musculus] | 0 | 30.548 | 33 | 8 | 45 | 7 | 292 | 33.3 | 7.66 | 597 | 8 | 31545870.61 | 34110136.19 | 36109047.12 | 24976835.98 | 33419584.21 | 33616465.46 | 33080789 | 26669687.13 | 15.2964 | 10.596 | Cdk5 | 12568 | 0.99982 | 1.000176059 | 1.00035 | 0.9971 | 0.998251 |
| 568982990 | PREDICTED: centrin-3 isoform X1 [Mus musculus] | 0 | 12.556 | 20 | 2 | 6 | 2 | 183 | 21.3 | 5.11 | 70 | 2 | 3510425.045 | 3601803.556 | 3063912.428 | 2660297.852 | 2188055.334 | 116607.4598 | 2497948 | 3824825.311 | 13.5504 | 71.135 | Cetn3 | 12626 | 1.1961 | 0.80390294 | 0.67211 | 0.23506 | 0.730886 |
| 6680924 | cofilin-1 [Mus musculus] | 0 | 62.315 | 72 | 10 | 86 | 8 | 166 | 18.5 | 8.09 | 1447 | 10 | 128917234.8 | 136383405.9 | 138880107.4 | 138651032.8 | 143798381 | 162532183.2 | 130016164.6 | 180593625.2 | 3.43773 | 14.2987 | Cfl1 | 12631 | 0.9361 | 1.063899253 | 1.13652 | 0.15132 | 0.693437 |
| 6680946 | cold-inducible RNA-binding protein [Mus musculus] | 0 | 10.897 | 21 | 3 | 7 | 3 | 172 | 18.6 | 9.61 | 159 | 3 | 10043591.17 | 18942650.57 | 17722087.45 | 13803448.07 | 13611713.89 | 22162567.33 | 19548595 | 26215033.83 | 26.6842 | 25.9161 | Cirbp | 12696 | 0.85198 | 1.148019564 | 1.34747 | 0.16491 | 0.710681 |
| 568936834 | PREDICTED: citron Rho-interacting kinase isoform X18 [Mus musculus] | 0 | 703.437 | 59 | 94 | 1129 | 3 | 1657 | 189.8 | 6.64 | 17492 | 94 | 11081804.23 | 17042052.32 | 13539686.83 | 16467000.57 | 12946595.76 | 18475858.24 | 14135597 | 15294193.18 | 19.0251 | 15.6247 | Cit | 12704 | 0.97713 | 1.022874739 | 1.04682 | 0.72181 | 0.939018 |
| 10946574 | creatine kinase B-type [Mus musculus] | 0 | 151.471 | 63 | 16 | 130 | 16 | 381 | 42.7 | 5.67 | 3089 | 16 | 161994873.5 | 187572392.6 | 178234647.4 | 257432636.4 | 196327825.6 | 213632898.3 | 182335032 | 185012443.9 | 21.4445 | 7.32156 | Ckb | 12709 | 1.00507 | 0.994927275 | 0.98991 | 0.93184 | 0.994387 |
| 148696104 | creatine kinase, mitochondrial 1, ubiquitous, isoform CRA_c, partial [Mus musculus] | 0 | 350.13 | 65 | 27 | 1179 | 25 | 423 | 47.5 | 8.18 | 22670 | 27 | 10020038388 | 12082347284 | 10896005620 | 12533968680 | 11259589775 | 12673616028 | 9863998468 | 14500647842 | 10.0275 | 16.4222 | Ckmt1 | 12716 | 0.97053 | 1.029473375 | 1.06074 | 0.56774 | 0.895889 |
| 31418362 | Coronin, actin binding protein 1A [Mus musculus] | 0 | 11.573 | 6 | 2 | 6 | 2 | 461 | 51 | 6.48 | 319 | 2 | 5136820.337 | 1222098.231 | 5117367.241 | 4755623.954 | 4892551.625 | 4091774.359 | 4696102.938 | 4616922.98 | 46.7892 | 7.47517 | Coro1a | 12721 | 0.94018 | 1.059817156 | 1.12725 | 0.61169 | 0.905186 |
| 122939194 | clathrin light chain A isoform b [Mus musculus] | 0 | 8.563 | 13 | 4 | 18 | 4 | 248 | 27 | 4.5 | 35 | 4 | 7340375.664 | 12678355.3 | 27590874.35 | 38698381.7 | 44659256.25 | 27793321.21 | 31604640 | 24704741.26 | 66.1502 | 27.2704 | Clta | 12757 | 0.8026 | 1.197396115 | 1.49189 | 0.25219 | 0.733806 |
| 568986618 | PREDICTED: clusterin isoform X1 [Mus musculus] | 0 | 51.154 | 26 | 11 | 43 | 11 | 458 | 52.7 | 5.76 | 404 | 11 | 54799045.58 | 55666605.03 | 65203414.33 | 48627842.68 | 75826684.61 | 55187465.85 | 64438348.63 | 72424388.31 | 12.2079 | 13.7244 | Clu | 12759 | 0.91145 | 1.088545917 | 1.1943 | 0.10597 | 0.674486 |
| 247629309 | N-acylneuraminate cytidylyltransferase [Mus musculus] | 0 | 62.091 | 37 | 10 | 34 | 10 | 432 | 48 | 8.1 | 436 | 10 | 47564608.04 | 40018999.7 | 46943922.2 | 43929601.58 | 74671858.36 | 28661249.21 | 48798948.75 | 40256992.62 | 7.73398 | 40.6348 | Cmas | 12764 | 0.96243 | 1.037567914 | 1.07807 | 0.73759 | 0.943908 |
| 226423907 | 2',3'-cyclic-nucleotide 3'-phosphodiesterase isoform 2 [Mus musculus] | 0 | 273.919 | 66 | 30 | 725 | 30 | 420 | 47.1 | 8.97 | 14168 | 30 | 2032707485 | 1986391999 | 1887673102 | 1990020637 | 2045083788 | 1960583962 | 2081444254 | 2049617291 | 3.11 | 2.53979 | Cnp | 12799 | 0.98504 | 1.014964651 | 1.03038 | 0.18552 | 0.715071 |
| 6680954 | contactin-1 precursor [Mus musculus] | 0 | 343.19 | 49 | 39 | 333 | 39 | 1020 | 113.3 | 6.16 | 7365 | 39 | 582440108.8 | 634362042 | 509395602 | 476244648.9 | 735000135.8 | 567264497.6 | 558385106.3 | 519539524.3 | 12.9515 | 16.0619 | Cntn1 | 12805 | 0.96121 | 1.038787072 | 1.0807 | 0.48426 | 0.869 |
| 651165917 | cytochrome c oxidase subunit 4 isoform 1, mitochondrial isoform 1 [Mus musculus] | 0.001 | 6.651 | 10 | 2 | 15 | 2 | 206 | 23.5 | 9.92 | 113 | 2 | 7760390.042 | 8210321.874 | 8987140.953 | 7834895.741 | 11014118.37 | 9382197.977 | 7099753.5 | 11319710.82 | 6.85051 | 19.9225 | Cox4i1 | 12857 | 0.91589 | 1.08411054 | 1.18367 | 0.18532 | 0.715071 |
| 112181182 | cytochrome c oxidase subunit 5A, mitochondrial precursor [Mus musculus] | 0 | 14.985 | 27 | 3 | 8 | 3 | 146 | 16.1 | 6.54 | 99 | 3 | 6744152.277 | 8692207.132 | 6542674.243 | 10209924.44 | 13274354.21 | 7527167.097 | 9649669.5 | 9133242.994 | 21.5891 | 24.5221 | Cox5a | 12858 | 0.89696 | 1.103039239 | 1.22975 | 0.26162 | 0.745606 |
| 148704346 | copine VI, isoform CRA_a [Mus musculus] | 0 | 72.009 | 31 | 16 | 73 | 15 | 595 | 66.1 | 6.2 | 1547 | 16 | 55345923.26 | 53081934.55 | 71301936.21 | 57183917.88 | 89316393.25 | 80583060.3 | 66296925 | 103321178.7 | 13.8818 | 18.2967 | Cpne6 | 12891 | 0.822 | 1.177998403 | 1.43309 | 0.02665 | 0.515682 |
| 755562317 | PREDICTED: carnitine O-palmitoyltransferase 1, liver isoform isoform X2 [Mus musculus] | 0 | 25.441 | 10 | 7 | 21 | 7 | 798 | 90.6 | 8.54 | 181 | 7 | 7269013.492 | 6321404.148 | 9452481.764 | 9410796.993 | 10829021.88 | 10068042.81 | 10096398.06 | 12848777.79 | 19.3584 | 11.9261 | Cpt1a | 12894 | 0.85073 | 1.149268029 | 1.35092 | 0.03172 | 0.515682 |
| 31542421 | crk-like protein isoform 1 [Mus musculus] | 0 | 9.084 | 9 | 2 | 8 | 2 | 303 | 33.8 | 6.74 | 31 | 2 | 4482426.16 | 3303426.836 | 3886569.344 | 3246473.703 | 3899491.694 | 3164666.043 | 3501994.75 | 2543658.641 | 15.5293 | 17.5145 | Crkl | 12929 | 1.06454 | 0.935455999 | 0.87874 | 0.3098 | 0.763986 |
| 148705576 | collapsin response mediator protein 1, isoform CRA_b, partial [Mus musculus] | 0 | 162.176 | 38 | 18 | 133 | 14 | 691 | 74.7 | 6.7 | 1923 | 18 | 161914454 | 149704755.6 | 147620291.9 | 165470227.4 | 160260462.4 | 131575214.9 | 140996842.2 | 97719699.79 | 5.97946 | 19.7241 | Crmp1 | 12933 | 1.08007 | 0.919930164 | 0.85173 | 0.14738 | 0.693437 |
| 568986628 | PREDICTED: dihydropyrimidinase-related protein 2 isoform X1 [Mus musculus] | 0 | 274.029 | 66 | 27 | 197 | 23 | 673 | 73.1 | 6.4 | 4171 | 27 | 641543848.2 | 608176174.7 | 598013250.1 | 613793289.2 | 638648610.9 | 502693741.3 | 495268984.8 | 420781790.9 | 3.02642 | 17.6431 | Dpysl2 | 12934 | 1.08943 | 0.910568573 | 0.83582 | 0.07194 | 0.613219 |
| 162138936 | hyaluronan and proteoglycan link protein 1 precursor [Mus musculus] | 0 | 74.605 | 45 | 14 | 187 | 13 | 356 | 40.5 | 7.8 | 2826 | 14 | 155501008.5 | 78518456.67 | 162564674.4 | 114001572.2 | 88381438.1 | 146384016.2 | 161601714.4 | 82411341.79 | 30.6598 | 33.5542 | Hapln1 | 12950 | 1.03215 | 0.967850868 | 0.9377 | 0.78625 | 0.952792 |
| 6753530 | alpha-crystallin B chain [Mus musculus] | 0 | 10.247 | 13 | 2 | 11 | 2 | 175 | 20.1 | 7.33 | 47 | 2 | 4414693.539 | 5144344.165 | 5670616.726 | 5711384.439 | 2585996.01 | 5669035.354 | 4638416.875 | 5938887.215 | 11.5549 | 32.3203 | Cryab | 12955 | 1.05302 | 0.946982034 | 0.8993 | 0.54347 | 0.88733 |
| 13385942 | citrate synthase, mitochondrial precursor [Mus musculus] | 0 | 18.889 | 12 | 3 | 18 | 3 | 464 | 51.7 | 8.57 | 290 | 3 | 3776193.274 | 5364399.385 | 5249015.168 | 7318281.292 | 4922812.078 | 5480254.456 | 4796840.672 | 6431177.143 | 26.7823 | 13.7597 | Cs | 12974 | 1.00177 | 0.998227813 | 0.99646 | 0.982 | 0.997464 |
| 1262300 | casein kinase II alpha subunit [Mus musculus] | 0 | 147.103 | 51 | 16 | 84 | 15 | 391 | 45.2 | 7.69 | 2079 | 16 | 277009570 | 232973032.7 | 279797619.3 | 235576252 | 206569503.7 | 261727648.5 | 264359656.2 | 196620897.6 | 9.95767 | 15.3776 | Csnk2a1 | 12995 | 1.04915 | 0.950845652 | 0.9063 | 0.31587 | 0.766307 |
| 26344986 | unnamed protein product [Mus musculus] | 0 | 82.209 | 45 | 12 | 49 | 11 | 350 | 41.1 | 8.56 | 705 | 12 | 47016983.66 | 44820420.63 | 53723396.59 | 48800529.58 | 57357433.97 | 55648354.29 | 68940044 | 55010299.67 | 7.79881 | 11.0448 | Csnk2a2 | 13000 | 0.90124 | 1.098755105 | 1.21915 | 0.03048 | 0.515682 |
| 745399120 | casein kinase II subunit beta isoform 1 [Mus musculus] | 0 | 52.694 | 45 | 9 | 64 | 9 | 236 | 27.1 | 5.96 | 1555 | 9 | 106984997.4 | 125666178.8 | 91675030.63 | 118735355.1 | 98645149.55 | 167673894.4 | 103060119.5 | 137016901.5 | 13.8515 | 25.5189 | Csnk2b | 13001 | 0.9343 | 1.065695798 | 1.14063 | 0.41547 | 0.836939 |
| 148668863 | mCG116562, isoform CRA_b, partial [Mus musculus] | 0 | 70.63 | 5 | 14 | 66 | 14 | 3413 | 373.2 | 4.65 | 442 | 14 | 70870614.77 | 90089220.95 | 85968562.26 | 97493530.76 | 70832546.53 | 60430486.48 | 65544661 | 74524052.86 | 13.0307 | 9.32414 | Vcan | 13003 | 1.12052 | 0.879479888 | 0.78489 | 0.02811 | 0.515682 |
| 1709255 | RecName: Full=Neurocan core protein; AltName: Full=Chondroitin sulfate proteoglycan 3; Flags: Precursor | 0 | 79.262 | 15 | 15 | 108 | 15 | 1268 | 137.1 | 5.72 | 1684 | 15 | 97619719.63 | 74566403.6 | 93351772.31 | 100095907.7 | 120096827.4 | 82070106.25 | 93204525.09 | 76465454.6 | 12.6557 | 20.8513 | Ncan | 13004 | 0.99159 | 1.008411331 | 1.01697 | 0.89521 | 0.984401 |
| 148707626 | cysteine and glycine-rich protein 1, isoform CRA_a, partial [Mus musculus] | 0 | 84.17 | 52 | 9 | 38 | 9 | 208 | 22 | 8.48 | 899 | 9 | 27231481.21 | 29138146.94 | 53402950.04 | 35342718.28 | 34032931.27 | 29306504.04 | 37991848.75 | 31445355.79 | 32.8835 | 84.1373 | Csrp1 | 13007 | 0.76803 | 1.231974634 | 1.60408 | 0.41788 | 0.837049 |
| 7304989 | C-terminal-binding protein 1 isoform 3 [Mus musculus] | 0 | 40.244 | 23 | 6 | 7 | 6 | 441 | 47.7 | 6.77 | 243 | 6 | 1573730.396 | 3676804.534 | 6044580.818 | 3158831.938 | 4184210.922 | 3055948.83 | 2503547 | 1530250.72 | 51.2321 | 39.2976 | Ctbp1 | 13016 | 1.1236 | 0.876399178 | 0.77999 | 0.48891 | 0.869525 |
| 509495 | cortactin [Mus musculus] | 0 | 115.704 | 38 | 19 | 227 | 19 | 546 | 61.2 | 5.4 | 2779 | 19 | 169665938.7 | 212424272.6 | 220610690.1 | 370072111.8 | 250890308.4 | 271985769.1 | 227904331.9 | 432357659.1 | 35.9737 | 31.378 | Cttn | 13043 | 0.90242 | 1.097575942 | 1.21625 | 0.4411 | 0.852048 |
| 6753590 | cholesterol 24-hydroxylase precursor [Mus musculus] | 0 | 76.359 | 44 | 16 | 47 | 16 | 500 | 56.8 | 8.82 | 577 | 16 | 35712920.22 | 35194711.82 | 50519824.07 | 40199081.76 | 58577743.28 | 55758905.78 | 70737771.63 | 63028254.08 | 17.5879 | 10.5338 | Cyp46a1 | 13116 | 0.78894 | 1.211056801 | 1.53504 | 0.0042 | 0.38143 |
| 4503253 | dolichyl-diphosphooligosaccharide--protein glycosyltransferase subunit DAD1 [Homo sapiens] | 0.001 | 6.672 | 19 | 2 | 2 | 2 | 113 | 12.5 | 7.08 | 51 | 2 | 728674.3455 | 3172257.734 | 2332550.107 | 3963973.756 | 3286467.362 | 4676553.76 | 2680022 | 5206933.409 | 54.3101 | 36.4906 | Dad1 | 13135 | 0.75687 | 1.243131288 | 1.64247 | 0.16324 | 0.706428 |
| 410812207 | death-associated protein kinase 3 isoform a [Mus musculus] | 0 | 19.285 | 10 | 4 | 17 | 4 | 465 | 53.2 | 8.56 | 279 | 4 | 12213712.21 | 12353327.3 | 13378599.07 | 10376131.49 | 8406594.729 | 7972102.11 | 9078399.5 | 8371388.544 | 10.3414 | 5.42347 | Dapk3 | 13144 | 1.17642 | 0.823575895 | 0.70007 | 0.00159 | 0.38143 |
| 148708608 | drebrin-like, isoform CRA_b, partial [Mus musculus] | 0 | 61.048 | 25 | 7 | 41 | 7 | 443 | 49.3 | 4.94 | 1425 | 7 | 26772062.36 | 16535129.55 | 27482264.58 | 22271753.53 | 21277277.74 | 21645525.08 | 25139606.63 | 29500306.21 | 21.6856 | 15.6834 | Dbnl | 13169 | 0.97639 | 1.023614589 | 1.04837 | 0.73439 | 0.943908 |
| 118601017 | dynactin subunit 1 isoform 1 [Mus musculus] | 0 | 157.213 | 31 | 28 | 107 | 28 | 1281 | 141.6 | 5.9 | 2170 | 28 | 96581052.03 | 87133001.99 | 91409516.27 | 103262505.9 | 93257963.85 | 114133506.9 | 95483357.23 | 90046891.27 | 7.34677 | 11.0295 | Dctn1 | 13191 | 0.98115 | 1.018845451 | 1.03841 | 0.59279 | 0.902243 |
| 26339314 | unnamed protein product [Mus musculus] | 0 | 47.926 | 37 | 11 | 66 | 8 | 379 | 42.5 | 9.72 | 967 | 11 | 29888587.93 | 25535723.06 | 27293879.88 | 31196764.53 | 43211633.08 | 26042456.53 | 34572510.5 | 34913965.15 | 8.93867 | 20.2132 | Dcx | 13193 | 0.90174 | 1.098258727 | 1.21793 | 0.14713 | 0.693437 |
| 569006320 | PREDICTED: DNA damage-binding protein 1 isoform X1 [Mus musculus] | 0 | 27.158 | 8 | 10 | 38 | 10 | 1152 | 128 | 5.27 | 180 | 10 | 10791539.46 | 12340237.68 | 16429415.05 | 22148649.79 | 22306100.46 | 11495501.89 | 15009689.69 | 21035154.82 | 32.8824 | 29.1724 | Ddb1 | 13194 | 0.93815 | 1.061850462 | 1.13186 | 0.59197 | 0.902243 |
| 28981429 | Asap1 protein, partial [Mus musculus] | 0 | 56.008 | 16 | 13 | 49 | 12 | 1167 | 129.1 | 7.83 | 585 | 13 | 39432106.34 | 23137609.85 | 42333544.29 | 23270950.53 | 31155494.95 | 28464800.41 | 31631998.81 | 37161908.68 | 32.0669 | 11.3668 | Asap1 | 13196 | 0.99906 | 1.000935318 | 1.00187 | 0.99158 | 0.998251 |
| 169234955 | dendrin [Mus musculus] | 0 | 86.864 | 36 | 14 | 59 | 14 | 710 | 76.4 | 9.98 | 582 | 14 | 47241105.2 | 47439128.6 | 45186129.71 | 38990317.71 | 37589972.82 | 43472279.51 | 36997460.5 | 28763800.48 | 8.83281 | 16.4764 | Ddn | 13199 | 1.09836 | 0.90164226 | 0.8209 | 0.06844 | 0.601634 |
| 2662377 | oligosaccharyltransferase [Mus musculus] | 0 | 23.446 | 18 | 6 | 35 | 6 | 441 | 49 | 5.83 | 107 | 6 | 42695130.86 | 30595481.47 | 22064190.23 | 44087558.92 | 44673472.82 | 51822596.85 | 57186526.5 | 39930836.72 | 30.0153 | 15.758 | Ddost | 13200 | 0.83735 | 1.162648638 | 1.38848 | 0.0814 | 0.642075 |
| 110835723 | pre-mRNA-splicing factor ATP-dependent RNA helicase DHX15 isoform 2 [Mus musculus] | 0 | 18.75 | 7 | 5 | 17 | 5 | 795 | 90.9 | 7.46 | 357 | 5 | 10678966.32 | 14920877.8 | 13007782.53 | 9140340.992 | 12274580.01 | 9097711.916 | 12345659.31 | 2811170.933 | 16.0566 | 49.0349 | Dhx15 | 13204 | 1.07586 | 0.92414276 | 0.85898 | 0.55457 | 0.894982 |
| 6753620 | ATP-dependent RNA helicase DDX3X [Mus musculus] | 0 | 205.273 | 51 | 27 | 188 | 6 | 662 | 73.1 | 7.18 | 3604 | 27 | 57581736.28 | 34043841.2 | 63066174.95 | 43468650.95 | 46834916.81 | 32252349.83 | 64538184.25 | 27804191.28 | 26.6934 | 38.6924 | Ddx3x | 13205 | 1.07233 | 0.927674564 | 0.86511 | 0.55182 | 0.892044 |
| 74210167 | unnamed protein product [Mus musculus] | 0 | 92.987 | 28 | 19 | 142 | 13 | 690 | 76.7 | 8.6 | 1705 | 19 | 171310040.7 | 184151001.8 | 168722715.4 | 234727535.6 | 162263939.1 | 218633780.4 | 177044027.8 | 209502290 | 16.2069 | 13.867 | Ddx5 | 13207 | 0.99441 | 1.005590274 | 1.01124 | 0.91986 | 0.987994 |
| 148693657 | DEAD (Asp-Glu-Ala-Asp) box polypeptide 6, isoform CRA_a, partial [Mus musculus] | 0 | 56.545 | 30 | 10 | 25 | 10 | 485 | 54.4 | 8.66 | 519 | 10 | 28797199.54 | 24534431.96 | 27309770.96 | 29335494.54 | 18834668.22 | 25044480.15 | 25698284.5 | 34683721.24 | 7.6506 | 25.0342 | Ddx6 | 13206 | 1.02713 | 0.972866495 | 0.94717 | 0.68627 | 0.92798 |
| 148707490 | DEAH (Asp-Glu-Ala-His) box polypeptide 9, isoform CRA_b, partial [Mus musculus] | 0 | 102.613 | 20 | 19 | 61 | 19 | 1384 | 149.6 | 6.83 | 1156 | 19 | 86604071.8 | 91003408.92 | 81292593.68 | 96780782.12 | 37980964.51 | 101017886.3 | 100176368.4 | 93552531.76 | 7.39374 | 36.4482 | Dhx9 | 13211 | 1.03334 | 0.96665773 | 0.93547 | 0.72414 | 0.939018 |
| 23943838 | tricarboxylate transport protein, mitochondrial [Mus musculus] | 0 | 156.868 | 52 | 13 | 147 | 13 | 311 | 33.9 | 9.89 | 2016 | 13 | 460037170.3 | 448423531.9 | 601068175.8 | 541600290.8 | 467639528.2 | 516587350.5 | 581832477.6 | 455388861.3 | 14.0397 | 11.3638 | Slc25a1 | 13358 | 1.00729 | 0.992711998 | 0.98553 | 0.87727 | 0.980491 |
| 755524052 | PREDICTED: 7-dehydrocholesterol reductase isoform X1 [Mus musculus] | 0 | 7.118 | 4 | 2 | 10 | 2 | 474 | 54.2 | 8.35 | 123 | 2 | 2751288.81 | 5382422.763 | 3643340.286 | 4312356.59 | 3211984.701 | 3733680.23 | 2584039 | 5223058.966 | 27.584 | 30.5306 | Dhcr7 | 13360 | 1.04334 | 0.956661756 | 0.91692 | 0.6872 | 0.928578 |

| | | | | | | | | | | | | | | | | | | | | | | | | | | | | | |
|---|---|---|---|---|---|---|---|---|---|---|---|---|---|---|---|---|---|---|---|---|---|---|---|---|---|---|---|---|---|
| 568994562 | PREDICTED: disks large homolog 1 isoform X6 [Mus musculus] | 0 | 300.134 | 54 | 38 | 474 | 3 | 917 | 101.3 | 5.69 | 8530 | 38 | 7995869.904 | 11849491.67 | 10331713.15 | 11223272.88 | 8323863.638 | 9873980.431 | 5770550.75 | 10260153.41 | 16.3141 | 23.8102 | Dlg1 | 13383 | 1.09483 | 0.905171174 | 0.82677 | 0.22417 | 0.718033 |
| 118026923 | MAGUK p55 subfamily member 3 [Mus musculus] | 0 | 17.094 | 9 | 4 | 9 | 4 | 585 | 66.4 | 6.47 | 39 | 4 | 1882668.323 | 1087545.402 | 2628173.878 | 2922844.002 | 2224489.298 | 486726.5747 | 1953051.219 | 485406.1739 | 38.5632 | 72.3878 | Mpp3 | 13384 | 1.24662 | 0.75337709 | 0.60433 | 0.22362 | 0.718033 |
| 568971490 | PREDICTED: disks large homolog 4 isoform X1 [Mus musculus] | 0 | 624.997 | 75 | 50 | 1803 | 46 | 754 | 83.6 | 6.09 | 44171 | 50 | 10252542928 | 10917921033 | 10066448238 | 10232595933 | 10882760101 | 12606548194 | 10430861735 | 10178146225 | 3.63053 | 9.92491 | Dlg4 | 13385 | 0.96928 | 1.030721923 | 1.06339 | 0.29931 | 0.761857 |
| 225637538 | dystrophia myotonica WD repeat-containing protein [Mus musculus] | 0 | 8.317 | 4 | 3 | 11 | 3 | 665 | 69.8 | 7.24 | 160 | 3 | 4104022.734 | 2663331.139 | 4690756.587 | 2662930.965 | 4039665.403 | 2803889.286 | 4594718 | 2667359.674 | 29.1629 | 26.7259 | Dmwd | 13401 | 1.00055 | 0.999454096 | 0.99891 | 0.99577 | 0.998251 |
| 569009599 | PREDICTED: dystrophin isoform X1 [Mus musculus] | 0 | 141.636 | 11 | 33 | 85 | 32 | 3696 | 427.4 | 5.87 | 1102 | 33 | 48010752.73 | 32024475.24 | 50832034.8 | 32388177.4 | 44981756.72 | 33643786.48 | 53108364.5 | 42919866.19 | 24.518 | 18.3168 | Dmd | 13405 | 0.96627 | 1.033731941 | 1.06982 | 0.67199 | 0.922517 |
| 134288917 | cytoplasmic dynein 1 heavy chain 1 [Mus musculus] | 0 | 1337.27 | 51 | 193 | 1236 | 193 | 4644 | 531.7 | 6.42 | 20160 | 193 | 1914059737 | 1802781861 | 1846873236 | 1804463495 | 1669277022 | 2342098035 | 1906867882 | 1861858490 | 2.83188 | 14.6046 | Dync1h1 | 13424 | 0.97281 | 1.027192731 | 1.05591 | 0.50255 | 0.88031 |
| 568939413 | PREDICTED: cytoplasmic dynein 1 intermediate chain 1 isoform X1 [Mus musculus] | 0 | 27.563 | 9 | 4 | 15 | 4 | 645 | 72.8 | 5.15 | 637 | 4 | 4574669.961 | 3094121.707 | 5142341.599 | 1976570.85 | 4328523.418 | 3442782.73 | 5033611.25 | 2024542.336 | 38.8317 | 34.9845 | Dync1i1 | 13426 | 0.99859 | 1.001409845 | 1.00282 | 0.99174 | 0.998251 |
| 311893374 | cytoplasmic dynein 1 intermediate chain 2 isoform 1 [Mus musculus] | 0 | 22.382 | 11 | 4 | 23 | 4 | 655 | 73.3 | 5.15 | 144 | 4 | 4944665.47 | 12934740.44 | 9925521.702 | 7028609.063 | 10007030.51 | 4494889.592 | 8645138.125 | 6391117.092 | 39.9612 | 32.989 | Dync1i2 | 13427 | 1.08226 | 0.917737758 | 0.84798 | 0.55603 | 0.895107 |
| 568958775 | PREDICTED: dynamin-2 isoform X2 [Mus musculus] | 0 | 44.513 | 15 | 12 | 65 | 3 | 880 | 99.2 | 7.12 | 676 | 12 | 2989298.431 | 3670336.06 | 3227630.888 | 4872029.852 | 5776098.802 | 2525858.742 | 6674862.375 | 4582199.945 | 22.6875 | 36.688 | Dnm2 | 13430 | 0.86014 | 1.139858983 | 1.3252 | 0.27098 | 0.745606 |
| 2626980 | Doc2 [Mus musculus] | 0 | 40.007 | 27 | 7 | 32 | 6 | 405 | 44.5 | 6.98 | 553 | 7 | 40563894.78 | 26879148.87 | 38057890.83 | 26652040.06 | 34909309.63 | 26610686.85 | 37443162 | 27292581.36 | 22.1426 | 17.2111 | Doc2a | 13446 | 1.02282 | 0.977178654 | 0.95538 | 0.75723 | 0.952737 |
| 6753670 | dolichol-phosphate mannosyltransferase subunit 1 isoform 1 [Mus musculus] | 0 | 56.197 | 45 | 9 | 35 | 9 | 260 | 29.2 | 9.51 | 602 | 9 | 66783846.08 | 41866658.83 | 74331202.44 | 51861216.84 | 59595877.84 | 47439921.05 | 72291978.28 | 55626210.65 | 24.8742 | 17.6317 | Dpm1 | 13480 | 0.99976 | 1.000236408 | 1.00047 | 0.99763 | 0.998251 |
| 274319588 | dipeptidyl aminopeptidase-like protein 6 isoform 3 [Mus musculus] | 0 | 28.504 | 10 | 7 | 19 | 6 | 859 | 97.2 | 6.48 | 261 | 7 | 15204240.5 | 10628487.78 | 10728426.58 | 12958415.02 | 28680171.66 | 12465515.65 | 13105826.75 | 18125292.36 | 17.5158 | 41.435 | Dpp6 | 13483 | 0.81249 | 1.187513667 | 1.46158 | 0.19344 | 0.715071 |
| 24586704 | Developmentally regulated GTP binding protein 1 [Mus musculus] | 0 | 15.017 | 12 | 3 | 6 | 3 | 367 | 40.5 | 8.9 | 62 | 3 | 2059689.035 | 2129596.159 | 1988899.014 | 1975550.281 | 2437591.385 | 2004164.77 | 2981783.25 | 2135348.767 | 3.48852 | 18.1791 | Drg1 | 13494 | 0.92067 | 1.079330639 | 1.17233 | 0.1616 | 0.702694 |
| 569011258 | PREDICTED: dystrophin-related protein 2 isoform X2 [Mus musculus] | 0 | 33.187 | 9 | 6 | 21 | 6 | 981 | 110.2 | 6.18 | 231 | 6 | 10513254.8 | 10436536.36 | 10117264.58 | 6904347.346 | 7039187.726 | 3706853.846 | 9069976.5 | 2986490.528 | 18.2681 | 50.1158 | Drp2 | 13497 | 1.2496 | 0.750404506 | 0.60052 | 0.06372 | 0.601634 |
| 1039727474 | PREDICTED: dystonin isoform X1 [Mus musculus] | 0 | 1632.05 | 38 | 225 | 1267 | 220 | 7730 | 871.6 | 5.45 | 25735 | 225 | 1574444143 | 1574261655 | 1827675329 | 1624051551 | 1714027227 | 1417591714 | 1837827002 | 1733798230 | 7.31311 | 10.7714 | Dst | 13518 | 0.99227 | 1.007728051 | 1.01558 | 0.82073 | 0.964114 |
| 755560477 | PREDICTED: dystrobrevin alpha isoform X1 [Mus musculus] | 0 | 83.195 | 24 | 15 | 68 | 12 | 783 | 88.5 | 7.12 | 925 | 15 | 47175253.26 | 47719268.3 | 58004909.16 | 25309747.86 | 49705374.09 | 46534693.59 | 54144701.7 | 37677890.29 | 30.889 | 14.8116 | Dtna | 13527 | 0.9731 | 1.026902099 | 1.05529 | 0.76023 | 0.952792 |
| 148669448 | dystrobrevin, beta, isoform CRA_e, partial [Mus musculus] | 0 | 27.533 | 11 | 7 | 21 | 4 | 792 | 87.3 | 8.35 | 160 | 7 | 10223050.25 | 5331181.308 | 11204262.91 | 6921078.172 | 10682311.78 | 13631279.74 | 8694650.25 | 12034130.47 | 32.7349 | 18.5834 | Dtnb | 13528 | 0.85566 | 1.144340941 | 1.33738 | 0.15175 | 0.693437 |
| 24418935 | dual specificity tyrosine-phosphorylation-regulated kinase 1A [Mus musculus] | 0 | 21.614 | 7 | 5 | 17 | 3 | 763 | 85.4 | 8.75 | 284 | 5 | 4781235.261 | 3713367.962 | 3988126.162 | 2555747.111 | 5810289.719 | 7599488.806 | 4756882.875 | 7295505.548 | 24.5099 | 20.8481 | Dyrk1a | 13548 | 0.74263 | 1.257370984 | 1.69313 | 0.018 | 0.483854 |
| 7106301 | microtubule-associated protein RP/EB family member 1 [Mus musculus] | 0 | 45.536 | 40 | 8 | 27 | 7 | 268 | 30 | 5.22 | 320 | 8 | 11825335.39 | 15759010.19 | 18418403.23 | 27754940.17 | 26954963.29 | 16625461.77 | 23588661.97 | 22286134.72 | 36.743 | 19.2351 | Mapre1 | 13589 | 0.90382 | 1.096178254 | 1.21283 | 0.36585 | 0.80688 |
| 13277723 | Sparcl1 protein [Mus musculus] | 0 | 9.206 | 4 | 2 | 8 | 2 | 650 | 72.3 | 4.6 | 120 | 2 | 9329676.395 | 7506773.722 | 8894408.602 | 10035764.21 | 9968911.631 | 2715206.969 | 8670294.25 | 2315530.471 | 11.9212 | 67.0453 | Sparcl1 | 13602 | 1.20352 | 0.796477482 | 0.66179 | 0.19136 | 0.715071 |
| 1912314 | orphan G-protein-coupled receptor [Mus musculus] | 0 | 7.595 | 3 | 2 | 7 | 2 | 382 | 42.5 | 9.42 | 59 | 2 | 8232031.059 | 7580420.967 | 7621776.89 | 7606098.781 | 8393698.687 | 7394469.959 | 9436133.25 | 8028389.545 | 4.06045 | 10.2839 | S1pr1 | 13609 | 0.96559 | 1.034410637 | 1.07127 | 0.27033 | 0.745606 |
| 28460696 | elongation factor 1-alpha 1 [Rattus norvegicus] | 0 | 249.339 | 68 | 20 | 503 | 11 | 462 | 50.1 | 9.01 | 9768 | 20 | 571785743.9 | 459224830.9 | 614124991 | 425875948 | 556806327.5 | 493686955.9 | 677378109.3 | 521620250.9 | 17.3018 | 14.3859 | Eef1a1 | 13627 | 0.95869 | 1.041310033 | 1.08618 | 0.48762 | 0.869525 |
| 6681273 | elongation factor 1-alpha 2 [Mus musculus] | 0 | 237.564 | 65 | 20 | 458 | 11 | 463 | 50.4 | 9.03 | 7710 | 20 | 321013278.7 | 177672879.3 | 335354260.3 | 191359077.1 | 291886782.9 | 191167173.7 | 406980120.7 | 231801881.6 | 32.5105 | 33.498 | Eef1a2 | 13628 | 0.95509 | 1.044911918 | 1.09405 | 0.71426 | 0.936732 |
| 26324898 | unnamed protein product [Mus musculus] | 0 | 197.355 | 43 | 29 | 166 | 28 | 858 | 95.2 | 6.93 | 3152 | 29 | 244278375.9 | 203161596.9 | 251358460.4 | 203489474.8 | 263606940.2 | 207096484.7 | 240125026.7 | 216013774.5 | 11.4599 | 10.9759 | Eef2 | 13629 | 0.98658 | 1.013424041 | 1.02721 | 0.74648 | 0.947173 |
| 4503529 | eukaryotic initiation factor 4A-I isoform 1 [Homo sapiens] | 0 | 34.83 | 21 | 7 | 43 | 6 | 406 | 46.1 | 5.48 | 622 | 7 | 44929773.34 | 40196685.79 | 37640334.09 | 36794562.28 | 32143891.09 | 40804775.88 | 35212866.63 | 37713549.32 | 9.16936 | 10.0915 | Eif4a1 | 13681 | 1.04481 | 0.955191094 | 0.91423 | 0.23532 | 0.730886 |
| 568921780 | PREDICTED: eukaryotic translation initiation factor 4E isoform X1 [Mus musculus] | 0 | 7.176 | 8 | 2 | 14 | 2 | 230 | 26.6 | 7.42 | 67 | 2 | 18812234.93 | 16622677.16 | 20036610.49 | 30916058.49 | 23212682.82 | 29148695.49 | 20321860.25 | 16951969.81 | 29.5008 | 23.0739 | Eif4e | 13684 | 0.98155 | 1.018450039 | 1.03759 | 0.84966 | 0.97368 |
| 148685063 | eukaryotic translation initiation factor 4, gamma 2, isoform CRA_f, partial [Mus musculus] | 0 | 29.474 | 11 | 8 | 19 | 8 | 931 | 104.5 | 8.31 | 111 | 8 | 8988089.577 | 6888646.481 | 10426484.13 | 6984035.164 | 8108188.68 | 10099768.97 | 16437944.88 | 10796437.33 | 20.4837 | 31.4371 | Eif4g2 | 13690 | 0.84561 | 1.154390283 | 1.36516 | 0.17551 | 0.713031 |
| 6679641 | emerin [Mus musculus] | 0 | 48.249 | 35 | 7 | 54 | 7 | 259 | 29.4 | 5.01 | 848 | 7 | 11297852.01 | 13614625.5 | 19635618.96 | 21264565.75 | 21837705.53 | 25502405.14 | 18441190.81 | 26985695.93 | 28.9177 | 16.5384 | Emd | 13726 | 0.83003 | 1.169973471 | 1.40956 | 0.0696 | 0.601634 |
| 569006328 | PREDICTED: serine/threonine-protein kinase MARK2 isoform X3 [Mus musculus] | 0 | 278.225 | 57 | 37 | 439 | 26 | 841 | 94 | 9.72 | 6890 | 37 | 140249026.5 | 210437793.1 | 177477859.9 | 288201638.2 | 226259954.1 | 217505637.5 | 233552204.1 | 230643539.2 | 30.858 | 3.08275 | Mark2 | 13728 | 0.94688 | 1.053119265 | 1.1122 | 0.49703 | 0.875118 |
| 568910444 | PREDICTED: protein enabled homolog isoform X2 [Mus musculus] | 0 | 66.763 | 20 | 12 | 98 | 11 | 829 | 89.1 | 8.09 | 898 | 12 | 51938339.42 | 59902922.11 | 39620571 | 52913471.04 | 69971080.83 | 57319027.99 | 50808871.81 | 61676123.03 | 16.5016 | 13.4111 | Enah | 13800 | 0.9203 | 1.07970228 | 1.17321 | 0.17948 | 0.714602 |
| 2282582 | actin-binding protein [Mus musculus] | 0 | 8.479 | 7 | 3 | 4 | 3 | 589 | 66 | 6.81 | 20 | 3 | 3345205.591 | 1739027.781 | 2584253.375 | 2682404.56 | 2704511.341 | 2073547.293 | 2909798 | 1764826.412 | 25.4683 | 22.6198 | Enc1 | 13803 | 1.04536 | 0.954644134 | 0.91322 | 0.61564 | 0.905186 |
| 1039766553 | PREDICTED: alpha-enolase isoform X2 [Mus musculus] | 0 | 75.096 | 37 | 11 | 48 | 8 | 434 | 47.2 | 6.71 | 1951 | 11 | 39570875.78 | 43253641.44 | 38083299.2 | 34018662.97 | 44189461.46 | 47878730.55 | 40830606.5 | 44025978.41 | 9.86366 | 6.51506 | Eno1 | 13806 | 0.93371 | 1.066289632 | 1.14199 | 0.06122 | 0.601634 |
| 568915341 | PREDICTED: band 4.1-like protein 1 isoform X1 [Mus musculus] | 0 | 342.036 | 35 | 42 | 263 | 38 | 1602 | 174.6 | 5.39 | 5454 | 42 | 691087821.1 | 501867204.6 | 595627333.6 | 501166157.3 | 540273963.9 | 482114272.3 | 547462205.1 | 464464648.2 | 15.8432 | 8.15698 | Epb41l1 | 13821 | 1.05907 | 0.940927458 | 0.88844 | 0.2476 | 0.733806 |
| 568964817 | PREDICTED: band 4.1-like protein 2 isoform X1 [Mus musculus] | 0 | 103.001 | 21 | 17 | 66 | 13 | 1062 | 117.6 | 5.39 | 779 | 17 | 28308469.7 | 13323611.03 | 45688521.23 | 23891746.3 | 36954045.55 | 14257750.96 | 44124990.73 | 24243592.29 | 48.4823 | 44.4098 | Epb41l2 | 13822 | 0.96374 | 1.036257777 | 1.07524 | 0.83231 | 0.968992 |
| 358030313 | dematin isoform 1 [Mus musculus] | 0 | 155.463 | 67 | 20 | 295 | 20 | 405 | 45.4 | 8.41 | 3577 | 20 | 385528495.6 | 460144697.1 | 535376795.6 | 505686205.7 | 548357895.1 | 357042878.9 | 485801860.3 | 482217311.7 | 13.8313 | 17.1199 | Dmtn | 13829 | 1.00354 | 0.996458592 | 0.99294 | 0.95074 | 0.994906 |
| 34328113 | ephrin type-A receptor 4 precursor [Mus musculus] | 0 | 28.537 | 8 | 6 | 17 | 4 | 986 | 109.7 | 6.51 | 337 | 6 | 9212626.8 | 2340050.297 | 8847757.949 | 8024142.009 | 9753588.814 | 8357802.937 | 8131997.5 | 5233828.475 | 45.2573 | 24.1153 | Epha4 | 13838 | 0.94904 | 1.050960755 | 1.10739 | 0.69692 | 0.931521 |
| 145312274 | ephrin type-A receptor 6 [Mus musculus] | 0 | 18.087 | 5 | 5 | 14 | 3 | 1130 | 126.4 | 6.87 | 39 | 5 | 2494779.57 | 6668345.812 | 4875570.274 | 6201569.742 | 4073755.25 | 5896189.665 | 3201057.563 | 6315364.294 | 36.98 | 30.3578 | Epha6 | 13840 | 1.01898 | 0.981022841 | 0.96275 | 0.87961 | 0.980491 |
| 148681559 | Eph receptor B6, isoform CRA_a, partial [Mus musculus] | 0 | 36.218 | 11 | 7 | 14 | 7 | 1033 | 112.6 | 7.02 | 135 | 7 | 12335043.96 | 5519173.036 | 12867151.23 | 8021363.41 | 10531017.97 | 4927823.578 | 10032911.5 | 9436939.827 | 36.3909 | 29.4928 | Ephb6 | 13848 | 1.05177 | 0.948229062 | 0.90155 | 0.6775 | 0.925166 |
| 568955192 | PREDICTED: epidermal growth factor receptor substrate 15-like 1 isoform X1 [Mus musculus] | 0 | 12.241 | 5 | 4 | 5 | 4 | 923 | 100.8 | 5.01 | 24 | 4 | 2386285.777 | 1733052.946 | 2031057.034 | 1541789.738 | 2326082.099 | 1824478.207 | 1975883.875 | 1796022.455 | 19.1702 | 12.2923 | Eps15l1 | 13859 | 0.98525 | 1.01474776 | 1.02994 | 0.8031 | 0.959848 |
| 409264628 | epidermal growth factor receptor kinase substrate 8 isoform 2 [Mus musculus] | 0 | 123.218 | 29 | 17 | 78 | 17 | 847 | 94.4 | 7.46 | 2510 | 17 | 84944271.05 | 51656842.31 | 91094053.32 | 61400856.98 | 78911805.41 | 62625681.17 | 97984992.44 | 57111464.58 | 25.9839 | 24.79 | Eps8 | 13860 | 0.98713 | 1.012869275 | 1.02607 | 0.89066 | 0.984401 |
| 4758302 | enhancer of rudimentary homolog [Homo sapiens] | 0 | 19.643 | 41 | 3 | 11 | 3 | 104 | 12.3 | 5.92 | 567 | 3 | 23361346.2 | 18205282.91 | 18515151.93 | 23996328.65 | 13835898.24 | 22372995.07 | 17258827.75 | 24917549.34 | 14.6731 | 25.4591 | Erh | 13877 | 1.03504 | 0.964959246 | 0.93229 | 0.64468 | 0.915705 |
| 117647236 | KH domain-containing, RNA-binding, signal transduction-associated protein 3 [Mus musculus] | 0 | 8.939 | 9 | 3 | 5 | 2 | 346 | 38.8 | 8.1 | 42 | 3 | 1717098.228 | 3847525.899 | 3264315.583 | 2672539.52 | 2422032.764 | 4697535.058 | 3177997.25 | 3761731.168 | 31.615 | 27.3258 | Khdrbs3 | 13992 | 0.89993 | 1.100068052 | 1.22239 | 0.37085 | 0.80688 |
| 254281243 | ena/VASP-like protein isoform 1 [Mus musculus] | 0 | 39.189 | 27 | 9 | 53 | 8 | 414 | 44.3 | 8.85 | 683 | 9 | 15684713.17 | 5013908.039 | 17355445.13 | 26525315.35 | 21354847.52 | 14536633.35 | 16496826.19 | 27275818.1 | 54.6248 | 28.5313 | Evl | 14026 | 0.89542 | 1.104578319 | 1.23358 | 0.49922 | 0.877125 |
| 568969858 | PREDICTED: RNA-binding protein EWS isoform X1 [Mus musculus] | 0 | 31.363 | 11 | 5 | 43 | 5 | 668 | 69.8 | 9.29 | 398 | 5 | 1577635.115 | 17968486.38 | 27478611.18 | 21561469.39 | 14644792.2 | 26535959.51 | 24661200.5 | 33989732.44 | 64.708 | 31.9295 | Ewsr1 | 14030 | 0.81448 | 1.18552354 | 1.45557 | 0.29634 | 0.760494 |
| 568991307 | PREDICTED: focal adhesion kinase 1 isoform X1 [Mus musculus] | 0 | 7.382 | 2 | 2 | 4 | 2 | 1140 | 129.1 | 6.84 | 69 | 2 | 3463871.047 | 584927.3132 | 3719318.622 | 1258240.194 | 3294092.26 | 2404092.412 | 3641467.375 | 879332.5722 | 69.5437 | 48.2443 | Ptk2 | 14083 | 0.93803 | 1.061969668 | 1.13213 | 0.77515 | 0.952792 |
| 113680348 | fascin [Mus musculus] | 0 | 28.76 | 15 | 6 | 27 | 6 | 493 | 54.5 | 6.89 | 201 | 6 | 18562571.12 | 18311633.01 | 22120146.84 | 22841776.79 | 20613222.32 | 20381097.13 | 17661286.56 | 12804489.19 | 11.5131 | 20.3198 | Fscn1 | 14086 | 1.06769 | 0.932313841 | 0.87321 | 0.27577 | 0.745606 |
| 148702862 | fatty acid synthase, isoform CRA_b [Mus musculus] | 0 | 67.847 | 7 | 13 | 37 | 13 | 2529 | 274.8 | 6.62 | 349 | 13 | 17782046.57 | 3715602.554 | 13724149.48 | 17634420.71 | 25350931.52 | 34945214.3 | 23076797.56 | 29746283.7 | 49.9864 | 18.5128 | Fasn | 14104 | 0.63692 | 1.363083872 | 2.14013 | 0.01171 | 0.461382 |
| 15822914 | fau [Mus musculus domesticus] | 0.001 | 4.244 | 8 | 2 | 4 | 2 | 137 | 14.8 | 10.05 | 21 | 2 | 12018099.54 | 25392236.02 | 79247731.87 | 78412865.19 | 60064931.79 | 90073892.82 | 90798618 | 123110919.8 | 72.0593 | 28.2915 | Fau | 14109 | 0.69778 | 1.302220709 | 1.86624 | 0.10053 | 0.674486 |
| 20452466 | ferrochelatase, mitochondrial [Mus musculus] | 0 | 23.129 | 13 | 4 | 10 | 4 | 422 | 47.4 | 8.91 | 39 | 4 | 21840283.97 | 23658535.39 | 25888753.26 | 21158093.59 | 18156945.83 | 18209995.89 | 18817539.25 | 24347645.13 | 9.14893 | 15.0453 | Fech | 14151 | 1.07563 | 0.924374087 | 0.85938 | 0.12615 | 0.693437 |
| 148683476 | fibrinogen, alpha polypeptide, isoform CRA_a [Mus musculus] | 0 | 81.171 | 19 | 9 | 49 | 9 | 789 | 87.3 | 6.11 | 895 | 9 | 36630157.21 | 36623287.83 | 45346064.78 | 36122876.14 | 54708156.92 | 28541209.68 | 69306675.5 | 37921373.24 | 11.5044 | 37.9276 | Fga | 14161 | 0.89642 | 1.103575184 | 1.23109 | 0.37362 | 0.80688 |
| 11136630 | fibroblast growth factor 12 isoform 1 [Homo sapiens] | 0.001 | 6.01 | 11 | 2 | 6 | 2 | 243 | 27.4 | 9.96 | 33 | 2 | 9257860.504 | 8153590.999 | 10740560.9 | 8686666.215 | 11017890.04 | 8489826.851 | 9894444.5 | 8761654.064 | 12.1151 | 12.1305 | Fgf12 | 14167 | 0.98233 | 1.017667904 | 1.03597 | 0.69455 | 0.930721 |
| 569009667 | PREDICTED: four and a half LIM domains protein 1 isoform X2 [Mus musculus] | 0 | 18.007 | 17 | 5 | 15 | 5 | 339 | 37.9 | 9.09 | 54 | 5 | 4290234.169 | 8995018.74 | 9698768.043 | 7464936.028 | 6018375.742 | 3601610.227 | 7150496.75 | 9355003.058 | 31.5675 | 36.6636 | Fhl1 | 14199 | 1.07642 | 0.923579076 | 0.85801 | 0.54751 | 0.889301 |
| 18204139 | Four and a half LIM domains 2 [Mus musculus] | 0 | 18.305 | 22 | 6 | 13 | 6 | 279 | 32 | 7.3 | 31 | 6 | 7964285.789 | 11557241.36 | 9578651.612 | 13291472.29 | 9347807.052 | 7875580.458 | 8878888.375 | 9528528.129 | 21.8933 | 8.31354 | Fhl2 | 14200 | 1.08665 | 0.91334742 | 0.84051 | 0.2145 | 0.716066 |
| 1039783122 | PREDICTED: peptidyl-prolyl cis-trans isomerase FKBP8 isoform X1 [Mus musculus] | 0 | 44.283 | 23 | 6 | 17 | 6 | 448 | 47.8 | 5.2 | 795 | 6 | 7800865.098 | 5028294.71 | 8855956.214 | 7104058.321 | 9932737.685 | 7817223.602 | 10219885.25 | 7083729.417 | 22.4448 | 17.6844 | Fkbp8 | 14232 | 0.90188 | 1.098122364 | 1.2176 | 0.21127 | 0.716066 |
| 11528490 | protein flightless-1 homolog isoform 1 [Mus musculus] | 0 | 77.01 | 14 | 15 | 53 | 15 | 1271 | 144.7 | 6.06 | 829 | 15 | 39415697.38 | 36197711.3 | 48755086.5 | 29775948.32 | 38320496.83 | 19627437.04 | 32390101.75 | 23570109.26 | 20.5101 | 29.7001 | Flii | 14248 | 1.15011 | 0.849894012 | 0.73897 | 0.13289 | 0.693437 |
| 6679809 | flotillin-1 [Mus musculus] | 0 | 61.96 | 29 | 10 | 59 | 10 | 428 | 47.5 | 7.15 | 1084 | 10 | 39929454.83 | 38740732.32 | 29544552.46 | 32307924.1 | 43297245.3 | 40927497.94 | 35071528.38 | 37798392.15 | 14.2547 | 9.15214 | Flot1 | 14251 | 0.94432 | 1.055682242 | 1.11793 | 0.22747 | 0.722761 |
| 1039735679 | PREDICTED: flotillin-2 isoform X1 [Mus musculus] | 0 | 71.191 | 30 | 11 | 48 | 11 | 494 | 54 | 7.3 | 671 | 11 | 54899518.88 | 29845530.08 | 45147298.8 | 19592884.78 | 68461737.44 | 23509492.08 | 46111641.25 | 27567317.1 | 42.0376 | 49.6039 | Flot2 | 14252 | 0.9487 | 1.051295266 | 1.10814 | 0.76521 | 0.952792 |
| 569009687 | PREDICTED: fragile X mental retardation protein 1 homolog isoform X2 [Mus musculus] | 0 | 58.803 | 26 | 12 | 72 | 10 | 593 | 66.7 | 7.23 | 899 | 12 | 52232986.11 | 41861897.44 | 71947741.95 | 51988494.32 | 76255491.83 | 73448833.66 | 95055439.63 | 86528400.73 | 23.0993 | 11.9602 | Fmr1 | 14265 | 0.79382 | 1.20617708 | 1.51945 | 0.0123 | 0.461382 |
| 46849812 | fibronectin isoform a precursor [Mus musculus] | 0 | 8.3 | 1 | 3 | 5 | 3 | 2477 | 272.4 | 5.59 | 31 | 3 | 2321811.259 | 1873397.989 | 1596285.108 | 1481032.201 | 2213720.002 | 1931266.77 | 2613213.5 | 1855979.25 | 20.5699 | 15.9255 | Fn1 | 14268 | 0.91555 | 1.084451299 | 1.18448 | 0.23433 | 0.730166 |
| 157951723 | SLIT-ROBO Rho GTPase-activating protein 2 [Mus musculus] | 0 | 122.671 | 28 | 21 | 116 | 20 | 1071 | 120.7 | 6.64 | 780 | 21 | 61177477.15 | 53313208.38 | 53323191.24 | 57841311.06 | 69184228.81 | 67443588.25 | 50419637.56 | 58378884.28 | 6.78057 | 14.1712 | Srgap2 | 14270 | 0.95803 | 1.041969702 | 1.08762 | 0.33813 | 0.782483 |
| 163954941 | fragile X mental retardation syndrome-related protein 1 isoform 1 [Mus musculus] | 0 | 63.684 | 24 | 12 | 61 | 9 | 677 | 76.2 | 6.98 | 974 | 12 | 12143347.64 | 16177058.78 | 21988477.03 | 18248480.7 | 19744417.98 | 27961504.82 | 24416135.88 | 27671065.21 | 23.969 | 15.3255 | Fxr1 | 14359 | 0.81446 | 1.185540061 | 1.45561 | 0.03187 | 0.515682 |
| 148672975 | Fyn proto-oncogene, isoform CRA_b [Mus musculus] | 0 | 49.568 | 20 | 9 | 50 | 5 | 572 | 64.6 | 7.14 | 922 | 9 | 19535927.81 | 16084021.84 | 8239248.851 | 9249015.075 | 9447350.513 | 12456238.97 | 14717619.25 | 13281709.97 | 40.9444 | 17.8318 | Fyn | 14360 | 1.03112 | 0.968883995 | 0.93965 | 0.79412 | 0.956891 |
| 148704084 | frizzled homolog 3 (Drosophila), isoform CRA_a, partial [Mus musculus] | 0 | 8.824 | 3 | 2 | 6 | 2 | 676 | 77.2 | 7.53 | 49 | 2 | 1444746.139 | 914456.3545 | 1325896.335 | 1159301.086 | 1233932.921 | 1570891.033 | 1254573.375 | 1180855.391 | 18.976 | 13.483 | Fzd3 | 14365 | 0.96075 | 1.039252994 | 1.08171 | 0.52019 | 0.880629 |
| 568971594 | PREDICTED: gamma-aminobutyric acid receptor subunit alpha-1 isoform X1 [Mus musculus] | 0 | 58.428 | 23 | 10 | 111 | 6 | 553 | 62.3 | 9.67 | 1085 | 10 | 55385001.43 | 68058720.44 | 51115412.37 | 72244662.77 | 61056037.5 | 62310064.91 | 54550120.63 | 56183640.62 | 16.3024 | 6.39437 | Gabra1 | 14394 | 1.02642 | 0.973583205 | 0.94853 | 0.57556 | 0.895889 |
| 1039796303 | PREDICTED: gamma-aminobutyric acid receptor subunit alpha-3 isoform X1 [Mus musculus] | 0 | 27.038 | 13 | 5 | 48 | 3 | 523 | 58.9 | 9.07 | 794 | 5 | 13031827.4 | 11472472.22 | 11542043.79 | 10060205.69 | 16259168.98 | 16045350.95 | 14208145 | 8911937.821 | 10.5295 | 24.6987 | Gabra3 | 14396 | 0.90822 | 1.091775317 | 1.2021 | 0.24679 | 0.733806 |
| 6679907 | gamma-aminobutyric acid receptor subunit beta-1 precursor [Mus musculus] | 0 | 63.225 | 26 | 10 | 60 | 6 | 474 | 54.1 | 8.78 | 665 | 10 | 8800217.201 | 9638387.509 | 18947979.15 | 16101366.8 | 22064806.68 | 9152604.688 | 19422371.25 | 8002391.848 | 36.9861 | 48.579 | Gabrb1 | 14400 | 0.95403 | 1.04596645 | 1.09636 | 0.77632 | 0.952792 |
| 6679909 | gamma-aminobutyric acid receptor subunit beta-2 precursor [Mus musculus] | 0 | 65.344 | 31 | 11 | 59 | 4 | 474 | 54.6 | 9.32 | 280 | 11 | 15625231.95 | 21934112.79 | 22912937.4 | 18446632.82 | 23676169.77 | 20000571.88 | 21920286 | 17136207.59 | 16.9335 | 13.5419 | Gabrb2 | 14401 | 0.9764 | 1.023595852 | 1.04833 | 0.67709 | 0.925166 |
| 6679911 | gamma-aminobutyric acid receptor subunit beta-3 isoform a precursor [Mus musculus] | 0 | 62.349 | 29 | 9 | 63 | 5 | 473 | 54.1 | 9.1 | 1047 | 9 | 117379361.9 | 58392729.62 | 67593018.86 | 59371315.08 | 94587587.69 | 72213614.25 | 86746285 | 62674751.51 | 37.1298 | 18.119 | Gabrb3 | 14402 | 0.97821 | 1.021787906 | 1.04455 | 0.8378 | 0.969831 |
| 14626498 | GABA(A) receptor gamma 2 subunit [Mus musculus] | 0 | 49.056 | 20 | 7 | 58 | 7 | 474 | 55.1 | 8.66 | 507 | 7 | 41249996.82 | 31069692.76 | 51140229.87 | 30829375.92 | 50119066.25 | 42025599.7 | 45017195.06 | 35008845.15 | 25.1072 | 14.6662 | Gabrg2 | 14406 | 0.94523 | 1.054773665 | 1.1159 | 0.46865 | 0.863085 |
| 1039759084 | PREDICTED: glutamate decarboxylase 1 isoform X1 [Mus musculus] | 0 | 13.812 | 8 | 4 | 14 | 4 | 621 | 69.7 | 7.15 | 196 | 4 | 30408036.33 | 41108005 | 36812934.32 | 26990245.36 | 52808941.3 | 27453716.81 | 30516799.38 | 45550555.76 | 18.7232 | 30.9468 | Gad1 | 14415 | 0.92796 | 1.072041308 | 1.15527 | 0.47082 | 0.863085 |
| 124517709 | glutamate decarboxylase 2 [Mus musculus] | 0 | 20.711 | 9 | 4 | 19 | 4 | 585 | 65.2 | 6.9 | 174 | 4 | 12557987.98 | 507597.3282 | 5101525.729 | 10522959.31 | 15402333.28 | 16209164.43 | 13525189.28 | 6391130.023 | 75.9126 | 34.7068 | Gad2 | 14417 | 0.7153 | 1.284696436 | 1.79602 | 0.15619 | 0.69857 |
| 93102411 | galactocerebrosidase precursor [Mus musculus] | 0 | 25.112 | 12 | 5 | 24 | 5 | 684 | 77.2 | 6.74 | 188 | 5 | 16288946.99 | 39155894.7 | 18978568.03 | 24402016.72 | 29035309.09 | 12455491.25 | 22563776.5 | 9856560.868 | 41.3128 | 48.2793 | Galc | 14420 | 1.14423 | 0.85576714 | 0.7479 | 0.39357 | 0.824613 |
| 6679935 | neuromodulin [Mus musculus] | 0 | 19.164 | 22 | 2 | 8 | 2 | 227 | 23.6 | 4.73 | 173 | 2 | 2266072.929 | 7972290.098 | 14548096.06 | 24368439.06 | 11162174.33 | 28635569.74 | 19632909.75 | 17736480.07 | 77.2151 | 37.371 | Gap43 | 14432 | 0.77825 | 1.221752574 | 1.56988 | 0.28439 | 0.751585 |
| 568971608 | PREDICTED: growth arrest-specific protein 7 isoform X1 [Mus musculus] | 0 | 57.754 | 28 | 9 | 42 | 9 | 476 | 54.3 | 7.37 | 573 | 9 | 39453854.74 | 38325761.95 | 31526670.71 | 32566068.29 | 24366488.94 | 37597731.07 | 25490088.63 | 39421651.23 | 11.2788 | 24.8742 | Gas7 | 14457 | 1.0558 | 0.944199108 | 0.8943 | 0.42915 | 0.845779 |
| 160298168 | protein NipSnap homolog 2 [Mus musculus] | 0 | 63.899 | 40 | 8 | 37 | 8 | 281 | 32.9 | 9.33 | 579 | 8 | 67173030.53 | 92405861.37 | 96933323.98 | 85448369.14 | 85575109.36 | 105471173.2 | 85865968.5 | 75565543.98 | 15.3154 | 14.2077 | Nipsnap2 | 14467 | 0.98486 | 1.015144915 | 1.03076 | 0.78137 | 0.952792 |
| 50510861 | mKIAA1258 protein [Mus musculus] | 0 | 58.87 | 35 | 12 | 50 | 12 | 487 | 54.5 | 5.63 | 537 | 12 | 37848981.31 | 31713866.71 | 31558678.59 | 31692777.94 | 44617437.03 | 39750261.92 | 35648755.75 | 29035576.92 | 9.32941 | 17.7045 | Gda | 14544 | 0.94239 | 1.057607898 | 1.12226 | 0.30792 | 0.762471 |
| 6753964 | ganglioside-induced differentiation-associated protein 1 [Mus musculus] | 0 | 84.963 | 41 | 11 | 43 | 11 | 358 | 41.3 | 8.37 | 913 | 11 | 62744121.12 | 37639118.84 | 43175106.84 | 34865111.24 | 66365285.44 | 38080460.37 | 40139588.25 | 44275392.35 | 28.1934 | 27.5844 | Gdap1 | 14545 | 0.97158 | 1.028417418 | 1.0585 | 0.78286 | 0.952792 |
| 148697875 | guanosine diphosphate (GDP) dissociation inhibitor 1, isoform CRA_a, partial [Mus musculus] | 0 | 60.05 | 24 | 9 | 35 | 7 | 498 | 55.6 | 5.03 | 681 | 9 | 29560561.08 | 22077755.74 | 26702320.96 | 17069907.42 | 30571878.83 | 23249427.46 | 30272601.38 | 14340554.76 | 22.9445 | 31.0294 | Gdi1 | 14567 | 0.9844 | 1.015599665 | 1.03169 | 0.87742 | 0.980491 |
| 74150721 | unnamed protein product [Mus musculus] | 0 | 24.452 | 12 | 4 | 10 | 2 | 512 | 57.7 | 7.94 | 337 | 4 | 9898360.86 | 13658459.06 | 8815452.566 | 12605902.11 | 12189256.66 | 17099325.68 | 11229278.78 | 15406272.62 | 20.1446 | 19.6044 | Gdi2 | 14569 | 0.89152 | 1.10848076 | 1.24336 | 0.17468 | 0.713031 |
| 148694984 | glycerol phosphate dehydrogenase 2, mitochondrial [Mus musculus] | 0 | 91.164 | 31 | 18 | 70 | 18 | 727 | 80.9 | 6.61 | 708 | 18 | 67006763.91 | 78245107.73 | 75712097.49 | 60801351.43 | 155447762.1 | 113898190.5 | 90248848.75 | 96366168.07 | 11.3985 | 25.7913 | Gpd2 | 14571 | 0.76387 | 1.236125039 | 1.61823 | 0.02888 | 0.515682 |
| 84000448 | glial fibrillary acidic protein isoform 2 [Mus musculus] | 0 | 233.642 | 65 | 33 | 854 | 31 | 430 | 49.9 | 5.34 | 15738 | 33 | 1932592177 | 1813881243 | 2241975027 | 1998799982 | 1968296764 | 1344534607 | 2594193830 | 2126520453 | 9.03737 | 25.7029 | Gfap | 14580 | 0.99711 | 1.002889821 | 1.0058 | 0.96761 | 0.99686 |
| 33286888 | Gja1 protein [Mus musculus] | 0 | 153.876 | 36 | 12 | 303 | 12 | 434 | 49.5 | 9.51 | 6355 | 12 | 531145341.2 | 713394214.8 | 332519637.6 | 653212677.7 | 469955483.4 | 560270033.2 | 398316446.4 | 271662063.6 | 30.1491 | 28.6675 | Gja1 | 14609 | 1.13486 | 0.865139017 | 0.76233 | 0.24895 | 0.733806 |
| 58331126 | gap junction beta-6 protein [Mus musculus] | 0 | 30.558 | 13 | 2 | 16 | 2 | 261 | 30.3 | 8.38 | 257 | 2 | 9486941.924 | 9504454.887 | 14044068.2 | 7052691.062 | 10032602 | 10739778.9 | 13937833 | 5260758.92 | 29.118 | 35.8567 | Gjb6 | 14623 | 1.00146 | 0.998536291 | 0.99708 | 0.99029 | 0.998251 |
| 31982332 | glutamine synthetase [Mus musculus] | 0 | 166.45 | 54 | 19 | 210 | 9 | 373 | 42.1 | 7.08 | 2442 | 19 | 238653261.4 | 214615506.4 | 244316233.3 | 245304098.5 | 171423191.6 | 152605135.8 | 179677128.2 | 159313112.1 | 6.09745 | 7.30942 | Glul | 14645 | 1.17428 | 0.825724394 | 0.70318 | 0.0003 | 0.195279 |
| 5725305 | unnamed protein product [Mus musculus] | 0 | 55.128 | 25 | 7 | 54 | 7 | 457 | 52.6 | 8.79 | 1080 | 7 | 54252920.21 | 53545140.43 | 51768295.48 | 57763411.12 | 57065018.17 | 52547721.41 | 48304616.55 | 48422306.39 | 4.62853 | 8.04915 | Glra1 | 14654 | 1.02594 | 0.974059717 | 0.94943 | 0.30084 | 0.761857 |
| 568921800 | PREDICTED: glycine receptor subunit beta isoform X4 [Mus musculus] | 0 | 36.597 | 17 | 7 | 32 | 7 | 520 | 58.7 | 8.53 | 420 | 7 | 63608439.73 | 36131054.52 | 51818922.11 | 47290481.44 | 62498510.06 | 50864060.26 | 43380337.63 | 41360202.23 | 22.8724 | 19.3152 | Glrb | 14658 | 1.00188 | 0.998121215 | 0.99625 | 0.98079 | 0.997464 |
| 6680027 | glutamate dehydrogenase 1, mitochondrial precursor [Mus musculus] | 0 | 292.309 | 62 | 29 | 425 | 29 | 558 | 61.3 | 8 | 9339 | 29 | 3068178196 | 2879665162 | 3082008823 | 3216439861 | 2776350114 | 2875197676 | 3037768981 | 2740977469 | 4.52337 | 4.65017 | Glud1 | 14661 | 1.03446 | 0.965535665 | 0.93337 | 0.07766 | 0.627416 |
| 148671281 | guanine nucleotide binding protein, alpha inhibiting 1 [Mus musculus] | 0 | 18.808 | 18 | 6 | 25 | 2 | 354 | 40.4 | 5.97 | 116 | 6 | 7982214.672 | 9661350.93 | 7953023.255 | 11015422.98 | 6969564.175 | 8334386.014 | 8366621.5 | 7196206.051 | 16.1283 | 9.56113 | Gnai1 | 14677 | 1.08514 | 0.914858667 | 0.84308 | 0.13239 | 0.693437 |
| 309255 | inhibitory G protein of adenylate cyclase, alpha chain [Mus musculus] | 0 | 27.944 | 19 | 6 | 29 | 2 | 355 | 40.5 | 5.55 | 154 | 6 | 9820997.939 | 6245876.935 | 8271918.819 | 2950892.31 | 11416955.33 | 7692270.024 | 9072312 | 5701364.952 | 43.4926 | 28.3636 | Gnai2 | 14678 | 0.89222 | 1.107780568 | 1.2416 | 0.42106 | 0.83996 |
| 148679159 | guanine nucleotide binding protein, alpha o, isoform CRA_a, partial [Mus musculus] | 0 | 79.636 | 33 | 13 | 115 | 11 | 461 | 51.4 | 6.89 | 1072 | 13 | 75367682.26 | 107625760.8 | 84171969.31 | 92451514.79 | 86251207.99 | 101232559.7 | 103761220.2 | 68639918.93 | 15.2606 | 17.9889 | Gnao1 | 14681 | 0.99963 | 1.000372452 | 1.00075 | 0.99517 | 0.998251 |
| 699027699 | Chain A, Structure Of A Fragment Of Human Phospholipase C-beta3 Delta472-559, In Complex With Galphaq | 0 | 18.512 | 16 | 4 | 13 | 4 | 379 | 44.6 | 6.06 | 56 | 4 | 2356150.871 | 3303741.54 | 4050824.846 | 7617370.843 | 5696716.387 | 4857396.349 | 5199717.5 | 4062450.667 | 53.0328 | 13.8688 | Gnaq | 14682 | 0.93301 | 1.06698708 | 1.14359 | 0.62246 | 0.908209 |
| 47271396 | protein GNAS isoform XLas [Mus musculus] | 0 | 13.11 | 4 | 4 | 11 | 3 | 1133 | 121.4 | 4.81 | 72 | 4 | 1044867.571 | 678352.5642 | 1173735.943 | 1705865.423 | 2711073.267 | 2176701.988 | 2820907.25 | 1593653.555 | 36.974 | 24.2208 | Gnas | 14683 | 0.66203 | 1.33796917 | 2.02101 | 0.01583 | 0.461382 |
| 27532946 | guanine nucleotide-binding protein G(z) subunit alpha [Mus musculus] | 0 | 60.826 | 41 | 12 | 44 | 12 | 355 | 40.8 | 7.61 | 716 | 12 | 79994997.96 | 95294117.18 | 73951348.21 | 69375663.53 | 94465796.41 | 81168644.81 | 76349427.88 | 91697228.18 | 14.1831 | 9.97987 | Gnaz | 14687 | 0.96215 | 1.037845501 | 1.07867 | 0.4109 | 0.835711 |
| 6680045 | guanine nucleotide-binding protein G(I)/G(S)/G(T) subunit beta-1 [Mus musculus] | 0 | 91.72 | 46 | 11 | 78 | 7 | 340 | 37.4 | 6 | 1206 | 11 | 34340128.74 | 19482484.84 | 36527584.82 | 41070937.42 | 41802492.61 | 28889167.33 | 36024768 | 27333948.72 | 28.4447 | 19.9862 | Gnb1 | 14688 | 0.9901 | 1.009904041 | 1.02001 | 0.9127 | 0.986636 |
| 13937391 | guanine nucleotide-binding protein G(I)/G(S)/G(T) subunit beta-2 [Mus musculus] | 0 | 79.237 | 35 | 9 | 56 | 5 | 340 | 37.3 | 6 | 955 | 9 | 24449314.46 | 11077711.74 | 22998355.4 | 27066258.27 | 18953234.79 | 12080170.73 | 15177581.56 | 10376948.31 | 33.1016 | 26.6514 | Gnb2 | 14693 | 1.20399 | 0.796006533 | 0.66114 | 0.12073 | 0.681064 |
| 5174447 | receptor of activated protein C kinase 1 [Homo sapiens] | 0 | 127.359 | 74 | 16 | 100 | 16 | 317 | 35.1 | 7.69 | 2105 | 16 | 352085136.5 | 386922836 | 195952319.8 | 386177864.2 | 334412851.3 | 306862974.4 | 289980060.5 | 389285229.6 | 27.5572 | 13.1698 | Rack1 | 14694 | 1.00023 | 0.999773992 | 0.99955 | 0.99773 | 0.998251 |
| 41281679 | guanine nucleotide-binding protein subunit beta-5 isoform 2 [Mus musculus] | 0 | 64.418 | 36 | 9 | 54 | 9 | 353 | 38.7 | 6.06 | 1065 | 9 | 43593362.71 | 42668931.85 | 54545441.53 | 38262270.32 | 54554433.29 | 51444690.86 | 58241173.5 | 46691524.31 | 15.4603 | 9.27834 | Gnb5 | 14697 | 0.9183 | 1.08169658 | 1.17793 | 0.10922 | 0.678142 |
| 148237408 | uncharacterized protein LOC100037080 [Xenopus laevis] | 0 | 140.726 | 59 | 15 | 108 | 15 | 295 | 32.8 | 4.87 | 3031 | 15 | 285766082.4 | 264938184.6 | 103938751.5 | 381913345.3 | 281180216.8 | 216102345.8 | 179697954.3 | 295035298.1 | 44.5052 | 22.4091 | Got2 | 14719 | 1.03213 | 0.967867449 | 0.93774 | 0.8087 | 0.961747 |
| 7710028 | glypican-1 precursor [Mus musculus] | 0 | 39.465 | 20 | 7 | 23 | 7 | 557 | 61.3 | 7.05 | 318 | 7 | 13924899.1 | 28570758.85 | 27487414.09 | 14813112.89 | 30092975.77 | 17925101.61 | 17755880 | 19288334.29 | 37.3006 | 27.8609 | Gpc1 | 14733 | 0.99843 | 1.001566638 | 1.00314 | 0.98969 | 0.998251 |
| 194319965 | Chain A, Crystal Structure Of Mouse Vps26b | 0 | 14.439 | 17 | 5 | 17 | 5 | 340 | 39.4 | 7.53 | 151 | 5 | 2985525.346 | 2244550.988 | 5761687.216 | 3632095.798 | 3230628.361 | 4856610.922 | 3425214.75 | 7878053.591 | 41.4107 | 44.278 | Gpi1 | 14751 | 0.85986 | 1.140136322 | 1.32595 | 0.39919 | 0.824613 |
| 148667829 | LanC (bacterial lantibiotic synthetase component C)-like 1, isoform CRA_b, partial [Mus musculus] | 0 | 43.507 | 25 | 8 | 34 | 8 | 425 | 48 | 8 | 554 | 8 | 76861778.35 | 80713232.65 | 63168976.16 | 85951567.6 | 65162626.98 | 277445780.7 | 55966910 | 92239794.29 | 12.7078 | 85.0047 | Lancl1 | 14768 | 0.76913 | 1.23086783 | 1.60033 | 0.41332 | 0.835788 |
| 2673845 | glutathione peroxidase [Mus musculus] | 0 | 25.759 | 37 | 7 | 29 | 7 | 201 | 22.3 | 7.21 | 400 | 7 | 14917053.09 | 24208980.71 | 25069594.35 | 22613701.49 | 25844252.68 | 17438335.74 | 24785519 | 21618698.17 | 21.3642 | 16.842 | Gpx1 | 14775 | 0.9837 | 1.016303337 | 1.03315 | 0.81787 | 0.964114 |
| 15011841 | glutathione peroxidase 3 precursor [Mus musculus] | 0 | 8.2 | 13 | 2 | 11 | 2 | 226 | 25.4 | 8.22 | 82 | 2 | 4725092.915 | 1122573.94 | 4957290.587 | 3626631.127 | 6755674.519 | 3980142.932 | 5935761.25 | 1973844.182 | 48.6593 | 45.8396 | Gpx3 | 14778 | 0.87261 | 1.127394651 | 1.29199 | 0.47501 | 0.863085 |
| 6680083 | growth factor receptor-bound protein 2 [Mus musculus] | 0 | 36.705 | 43 | 9 | 41 | 9 | 217 | 25.2 | 6.32 | 431 | 9 | 23411366.7 | 21322309.69 | 21758543.64 | 20003774.44 | 22850061.13 | 30018830.93 | 28026829.05 | 26244329.84 | 6.50117 | 11.3601 | Grb2 | 14784 | 0.89339 | 1.106612673 | 1.23867 | 0.02168 | 0.505785 |
| 7305103 | probable G-protein coupled receptor 162 [Mus musculus] | 0 | 19.749 | 8 | 3 | 11 | 3 | 588 | 64.1 | 8.79 | 91 | 3 | 1432113.391 | 4367171.833 | 7828952.536 | 5279997.091 | 4822465.973 | 14665109.29 | 3988406.75 | 1182976.44 | 55.8552 | 95.3302 | Gpr162 | 14788 | 0.868 | 1.131996651 | 1.30414 | 0.67104 | 0.922517 |
| 7305107 | protein C10 [Mus musculus] | 0 | 46.986 | 63 | 5 | 29 | 5 | 126 | 13.2 | 5.14 | 673 | 5 | 94024685.07 | 52814660.59 | 66580536.56 | 65178308.15 | 73979334.37 | 57199131.11 | 67748638 | 49645158.01 | 24.9642 | 17.4348 | Grcc10 | 14790 | 1.05696 | 0.943043225 | 0.89222 | 0.4913 | 0.87078 |
| 164664512 | glutamate receptor 1 isoform 1 precursor [Mus musculus] | 0 | 250.16 | 43 | 35 | 374 | 2 | 907 | 101.5 | 7.69 | 6654 | 35 | 19038289.56 | 31400907.45 | 16528217.17 | 28106736.49 | 18465009.54 | 34655866.98 | 13728687.13 | 35553399.16 | 29.9346 | 43.5504 | Gria1 | 14799 | 0.96289 | 1.03711221 | 1.07709 | 0.79103 | 0.954864 |
| 139394522 | glutamate receptor 2 isoform 1 precursor [Mus musculus] | 0 | 457.514 | 54 | 52 | 715 | 39 | 883 | 98.8 | 7.9 | 15444 | 52 | 1576844838 | 1508502892 | 1565246654 | 1609311868 | 1934421521 | 1897296799 | 1660628029 | 1887863196 | 2.68472 | 6.75218 | Gria2 | 14800 | 0.91787 | 1.082132975 | 1.17896 | 0.00532 | 0.38143 |
| 153792501 | glutamate receptor ionotropic, delta-1 precursor [Mus musculus] | 0 | 30.915 | 7 | 7 | 40 | 6 | 1009 | 112.1 | 6.7 | 320 | 7 | 7571176.498 | 3225744.022 | 9584543.838 | 3690226.493 | 9428273.295 | 5737037.413 | 9649828.5 | 5294382.933 | 51.0802 | 30.9756 | Grid1 | 14803 | 0.88856 | 1.111437728 | 1.25083 | 0.46369 | 0.863085 |
| 6680091 | glutamate receptor ionotropic, delta-2 precursor [Mus musculus] | 0 | 305.338 | 46 | 35 | 228 | 34 | 1007 | 113 | 6.1 | 5268 | 35 | 630186508.2 | 483726049.8 | 535994393.9 | 572106749.1 | 519793702.4 | 537462035 | 470842255.1 | 510728120.8 | 11.0901 | 5.53013 | Grid2 | 14804 | 1.04299 | 0.957006695 | 0.91756 | 0.22512 | 0.718033 |
| 755532817 | PREDICTED: glutamate receptor ionotropic, kainate 2 isoform X1 [Mus musculus] | 0 | 121.176 | 28 | 20 | 115 | 19 | 943 | 106.7 | 7.84 | 2398 | 20 | 107858535 | 91769157.64 | 108775175.4 | 93534690.88 | 108880378.3 | 125620765.6 | 126508094.6 | 92237955.96 | 9.03677 | 14.3157 | Grik2 | 14806 | 0.94 | 1.059998305 | 1.12766 | 0.21679 | 0.716066 |
| 124487364 | glutamate receptor ionotropic, kainate 3 precursor [Mus musculus] | 0 | 73.318 | 23 | 13 | 53 | 12 | 919 | 104 | 7.52 | 602 | 13 | 54357158.03 | 21271860.16 | 57359570.39 | 42762451.43 | 50000977.25 | 34646069.37 | 52092102.13 | 30253729.37 | 37.255 | 26.1542 | Grik3 | 14807 | 1.02555 | 0.974446923 | 0.95017 | 0.83127 | 0.968976 |
| 564326486 | PREDICTED: glutamate receptor ionotropic, kainate 5 isoform X1 [Rattus norvegicus] | 0 | 133.642 | 34 | 24 | 105 | 23 | 980 | 109.3 | 8.21 | 1066 | 24 | 101524548.2 | 103084226.5 | 127844832.1 | 98904656.9 | 156602492.1 | 169881025.5 | 164760713.5 | 99566452.36 | 12.4703 | 22.0398 | Grik5 | 14809 | 0.84401 | 1.155994193 | 1.36965 | 0.06423 | 0.601634 |
| 41680705 | glutamate receptor ionotropic, NMDA 2A precursor [Mus musculus] | 0 | 545.589 | 49 | 57 | 811 | 53 | 1464 | 165.3 | 7.01 | 20988 | 57 | 1617140407 | 1636509973 | 1568098232 | 1640765980 | 1826926238 | 2018782371 | 1635438400 | 1697408999 | 2.06194 | 9.43879 | Grin2a | 14811 | 0.94751 | 1.052491585 | 1.1108 | 0.08344 | 0.644421 |
| 117168299 | glutamate receptor ionotropic, NMDA 2B precursor [Mus musculus] | 0 | 894.583 | 55 | 76 | 1143 | 70 | 1482 | 165.9 | 6.87 | 20102 | 76 | 4779715087 | 5592098323 | 4918271398 | 5336168827 | 4945938508 | 6572268765 | 4992027315 | 5160444365 | 7.26255 | 14.3094 | Grin2b | 14812 | 0.97531 | 1.024692696 | 1.05064 | 0.56637 | 0.895889 |
| 755520724 | PREDICTED: glutamate receptor ionotropic, NMDA 2D isoform X1 [Mus musculus] | 0 | 191.995 | 31 | 27 | 202 | 23 | 1340 | 144.7 | 8.43 | 2913 | 27 | 194958595.5 | 142173018.5 | 184546554.2 | 189404087.1 | 195785953.9 | 154765747.4 | 184981772.6 | 155391370.2 | 13.5624 | 12.0747 | Grin2d | 14814 | 1.01438 | 0.985622461 | 0.97165 | 0.7626 | 0.952792 |
| 34328063 | metabotropic glutamate receptor 1 isoform alpha precursor [Mus musculus] | 0 | 81.947 | 19 | 21 | 90 | 19 | 1199 | 133.1 | 6.86 | 629 | 21 | 99473515.82 | 121739591.5 | 91658091.59 | 106663409.8 | 106100879.9 | 111701034.6 | 81717166.28 | 101685752 | 12.2034 | 13.0104 | Grm1 | 14816 | 1.02233 | 0.977666754 | 0.95631 | 0.63396 | 0.912586 |
| 116812904 | metabotropic glutamate receptor 8 isoform 1 precursor [Mus musculus] | 0 | 21.63 | 7 | 6 | 17 | 5 | 908 | 101.6 | 8.15 | 93 | 6 | 8620918.033 | 8786057.303 | 10085640.49 | 12312943.19 | 13705788.21 | 12690812.94 | 9728273.5 | 11973174.91 | 17.1353 | 14.0363 | Grm8 | 14823 | 0.90566 | 1.094336179 | 1.20832 | 0.1347 | 0.693437 |
| 12835845 | unnamed protein product [Mus musculus] | 0 | 111.391 | 42 | 22 | 176 | 20 | 655 | 72.4 | 5.11 | 4226 | 22 | 63640766.98 | 49427503.38 | 46566427.73 | 53393282.93 | 70161989.21 | 52437651.05 | 53796246.86 | 71446804.69 | 14.0208 | 16.5272 | Hspa5 | 14828 | 0.92446 | 1.075541172 | 1.16343 | 0.21871 | 0.716066 |
| 6754086 | glutathione S-transferase Mu 5 [Mus musculus] | 0.001 | 4.202 | 9 | 2 | 7 | 2 | 224 | 26.6 | 7.21 | 23 | 2 | 458792.4576 | 867355.5796 | 975885.9865 | 1779058.602 | 1722624.117 | 1652148.205 | 920573.1875 | 1703472.177 | 54.1699 | 25.8206 | Gstm5 | 14866 | 0.80975 | 1.190252195 | 1.4699 | 0.20517 | 0.715071 |
| 74211870 | unnamed protein product [Mus musculus] | 0 | 14.505 | 6 | 4 | 9 | 4 | 724 | 78.2 | 7.8 | 55 | 4 | 4593523.009 | 2723950.881 | 4353075.353 | 2842726.42 | 5645751.871 | 1813836.368 | 5949985.5 | 3091092.204 | 27.06 | 48.5913 | Gtpbp1 | 14904 | 0.93592 | 1.064080545 | 1.13694 | 0.67179 | 0.922517 |
| 568940748 | ICTED: membrane-associated guanylate kinase, WW and PDZ domain-containing protein 1 isoform X1 [Mus mu | 0 | 78.109 | 13 | 13 | 55 | 12 | 1484 | 163.2 | 7.36 | 1121 | 13 | 39087140.05 | 37586870.3 | 45792768.21 | 41026180.88 | 39959672.68 | 44462584.53 | 32476127.88 | 36589842.97 | 8.73213 | 13.2503 | Magi1 | 14924 | 1.03156 | 0.96843746 | 0.93881 | 0.45139 | 0.857112 |
| 569009723 | PREDICTED: glycerol kinase isoform X1 [Mus musculus] | 0 | 37.561 | 17 | 9 | 27 | 9 | 587 | 63.6 | 5.87 | 235 | 9 | 13878281.09 | 13069061.46 | 12471410.16 | 13168721.06 | 28458024.08 | 18363746.2 | 22325421.75 | 19058635.03 | 4.38658 | 20.8924 | Gk | 14933 | 0.74702 | 1.252983296 | 1.67732 | 0.0086 | 0.426794 |
| 61888838 | 3-hydroxyacyl-CoA dehydrogenase type-2 [Mus musculus] | 0 | 38.109 | 40 | 7 | 35 | 7 | 261 | 27.3 | 8.76 | 712 | 7 | 28888487.63 | 27503483.42 | 23102406.67 | 25479022.9 | 23746395.73 | 19500497.5 | 26789308.75 | 24283324.25 | 9.5982 | 12.8316 | Hsd17b10 | 15108 | 1.05346 | 0.946541633 | 0.89851 | 0.22479 | 0.718033 |
| 755537403 | PREDICTED: huntingtin-associated protein 1 isoform X1 [Mus musculus] | 0 | 55.586 | 29 | 12 | 52 | 12 | 629 | 70.2 | 4.74 | 432 | 12 | 54485240.01 | 45261986.62 | 58408587.73 | 46008934.8 | 84131712.54 | 58202785.42 | 75039900.63 | 43384819.32 | 12.6397 | 27.7264 | Hap1 | 15114 | 0.87827 | 1.121728439 | 1.2772 | 0.1908 | 0.715071 |
| 122441 | RecName: Full=Hemoglobin subunit alpha; AltName: Full=Alpha-globin; AltName: Full=Hemoglobin alpha chain | 0 | 26.611 | 43 | 6 | 28 | 6 | 142 | 15.1 | 8.22 | 86 | 6 | 34005031.27 | 45055668.02 | 44312715.13 | 36804749.11 | 46489847.46 | 35834218.76 | 38803388.19 | 31038598.75 | 13.7007 | 17.027 | Hba-a1 | 15122 | 1.02565 | 0.974348459 | 0.94998 | 0.65365 | 0.915864 |
| 1209637 | holocytochrome c-type synthetase [Mus musculus] | 0 | 9.701 | 10 | 2 | 9 | 2 | 272 | 31 | 6.87 | 79 | 2 | 3605747.379 | 3885312.348 | 4601462.584 | 3078142.579 | 7567450.075 | 6684279.852 | 6059814.5 | 4788785.907 | 16.7327 | 18.6142 | Hccs | 15159 | 0.75343 | 1.246571146 | 1.65453 | 0.00968 | 0.448067 |
| 3023934 | RecName: Full=Histone deacetylase 2; Short=HD2; AltName: Full=YY1 transcription factor-binding protein | 0 | 13.025 | 8 | 3 | 8 | 3 | 488 | 55.3 | 5.91 | 34 | 3 | 8736460.217 | 10038679.28 | 15128015.56 | 8106165.159 | 22218405.57 | 10243921.63 | 16034893 | 13266313.98 | 30.3461 | 33.0281 | Hdac2 | 15182 | 0.80964 | 1.190360129 | 1.47023 | 0.15161 | 0.693437 |
| 7108465 | histone deacetylase-3 [Mus musculus] | 0 | 9.04 | 9 | 3 | 3 | 3 | 428 | 48.7 | 5.11 | 14 | 3 | 3525182.041 | 2319397.561 | 2893723.669 | 1873880.372 | 2820189.026 | 2425198.99 | 3076461.25 | 1720004.571 | 26.9791 | 23.5475 | Hdac3 | 15183 | 1.02761 | 0.972386522 | 0.94626 | 0.76909 | 0.952792 |
| 148705521 | Huntington disease gene homolog, isoform CRA_b, partial [Mus musculus] | 0 | 10.547 | 1 | 2 | 6 | 2 | 3142 | 346.9 | 6.3 | 102 | 2 | 1111730.83 | 468731.1244 | 1239049.957 | 752183.133 | 1080873.53 | 754398.6092 | 1044193.813 | 418240.1338 | 39.1932 | 37.3174 | Htt | 15194 | 1.03989 | 0.960114579 | 0.92329 | 0.77866 | 0.952792 |
| 148689921 | ous to the E6-AP (UBE3A) carboxyl terminus) domain and RCC1 (CHC1)-like domain (RLD) 2, isoform CRA_a [M | 0.001 | 5.445 | 0 | 2 | 6 | 2 | 4902 | 534.5 | 6.3 | 47 | 2 | 1350391.385 | 686598.5097 | 1924392.908 | 1441773.363 | 1791762.906 | 1877416.113 | 1262178.625 | 1912181.313 | 37.7109 | 17.7328 | Herc2 | 15204 | 0.88239 | 1.117613999 | 1.26658 | 0.27011 | 0.745606 |
| 26348757 | unnamed protein product [Mus musculus] | 0.001 | 4.797 | 4 | 2 | 8 | 2 | 528 | 60.5 | 6.54 | 83 | 2 | 1737556.054 | 2031629.853 | 1892021.76 | 1587361.65 | 2182159.845 | 1661279.605 | 2176374.5 | 2131255.14 | 10.5977 | 12.3675 | Hexa | 15211 | 0.94139 | 1.058605299 | 1.12451 | 0.20427 | 0.715071 |
| 226165 | beta hexosaminidase beta | 0 | 8.497 | 4 | 2 | 16 | 2 | 539 | 61.1 | 8.12 | 17 | 2 | 4551887.443 | 11067731.87 | 6498862.243 | 7707448.448 | 5713666.324 | 9849948.749 | 6530407.75 | 9924831.215 | 36.6935 | 27.4783 | Hexb | 15212 | 0.96454 | 1.035458512 | 1.07352 | 0.76536 | 0.952792 |
| 755537437 | PREDICTED: hepatocyte growth factor-regulated tyrosine kinase substrate isoform X1 [Mus musculus] | 0 | 7.388 | 3 | 2 | 7 | 2 | 805 | 89.2 | 6.25 | 49 | 2 | 1832279.299 | 2693614.291 | 4873102.198 | 1544692.014 | 1952309.16 | 6757774.87 | 4492518.625 | 2362045.692 | 55.0487 | 56.8422 | Hgs | 15239 | 0.82568 | 1.174321033 | 1.42225 | 0.42107 | 0.83996 |
| 817939 | histone H2A [Mus musculus domesticus] | 0 | 67.325 | 48 | 4 | 33 | 2 | 137 | 14.7 | 10.59 | 868 | 4 | 259997529.6 | 77978635.78 | 239337602 | 119312026.3 | 128400052.9 | 127206165.5 | 241004176 | 133957561.1 | 51.2241 | 35.3032 | Hist2h2aa1 | 15267 | 1.04977 | 0.950227434 | 0.90517 | 0.76406 | 0.952792 |
| 148671544 | human immunodeficiency virus type I enhancer binding protein 2 [Mus musculus] | 0 | 34.632 | 5 | 11 | 33 | 11 | 2430 | 266.6 | 6.96 | 79 | 11 | 15487748.39 | 12936607.17 | 15896825.61 | 11148182.73 | 13309893.29 | 11571553.49 | 10265267.38 | 10156647.87 | 16.1272 | 12.9851 | Hivep2 | 15273 | 1.10088 | 0.899119511 | 0.81673 | 0.10631 | 0.674486 |
| 568966719 | PREDICTED: hexokinase-1 isoform X1 [Mus musculus] | 0 | 48.397 | 13 | 12 | 31 | 12 | 974 | 108.2 | 6.8 | 251 | 12 | 20008150.17 | 17600206.93 | 24514062.01 | 29743372.85 | 33885695.1 | 25557452.49 | 22317560.53 | 35461425.88 | 23.2952 | 21.7385 | Hk1 | 15275 | 0.87873 | 1.121271192 | 1.27602 | 0.17835 | 0.713031 |
| 74218228 | unnamed protein product [Mus musculus] | 0 | 25.73 | 18 | 5 | 21 | 5 | 334 | 36.9 | 7.36 | 340 | 5 | 54137425.14 | 48998327.29 | 51232828.74 | 43279400.85 | 51329477.88 | 57276697.9 | 66643920.5 | 54139331.54 | 9.30544 | 11.6078 | Hnrnpc | 15381 | 0.92567 | 1.074329427 | 1.1606 | 0.09747 | 0.670764 |
| 672424490 | heterogeneous nuclear ribonucleoprotein K isoform 3 [Mus musculus] | 0 | 84.292 | 47 | 17 | 207 | 17 | 440 | 48.5 | 5.54 | 2607 | 17 | 142794030.3 | 147325723.4 | 144546851 | 194735096.7 | 127480597.6 | 131734387.2 | 124915003.3 | 138319699.5 | 15.8836 | 4.48462 | Hnrnpk | 15387 | 1.09285 | 0.907147732 | 0.83007 | 0.08237 | 0.644421 |
| 197927211 | heterogeneous nuclear ribonucleoprotein L isoform a [Rattus norvegicus] | 0 | 133.615 | 50 | 16 | 69 | 8 | 623 | 67.9 | 8.47 | 1466 | 16 | 30774149.56 | 31690167.26 | 28551434.18 | 35823884.52 | 23986557.15 | 48120230.06 | 36291237.13 | 45400469.69 | 9.59589 | 28.3187 | Hnrnpl | 15388 | 0.90394 | 1.096062707 | 1.21254 | 0.27816 | 0.745606 |
| 12836501 | unnamed protein product [Mus musculus] | 0 | 67.576 | 30 | 15 | 82 | 15 | 554 | 60.8 | 9.67 | 1002 | 15 | 62083189.45 | 57157309.08 | 59402191.75 | 75506903.7 | 58899718.85 | 114404326.6 | 63638438.59 | 80716356.15 | 12.9527 | 31.6538 | Hp1bp3 | 15441 | 0.88893 | 1.111067348 | 1.24989 | 0.27518 | 0.745606 |
| 6754240 | neuron-specific calcium-binding protein hippocalcin [Mus musculus] | 0 | 40.609 | 44 | 9 | 72 | 4 | 193 | 22.4 | 4.97 | 649 | 9 | 17017819.74 | 6400969.072 | 9943490.543 | 6470981.926 | 11374284.03 | 12442778.71 | 11923267.25 | 10790627.8 | 50.093 | 6.11198 | Hpca | 15444 | 0.92245 | 1.077551752 | 1.16814 | 0.531 | 0.886555 |
| 13435621 | Hypoxanthine guanine phosphoribosyl transferase 1 [Mus musculus] | 0.001 | 6.593 | 10 | 2 | 9 | 2 | 218 | 24.5 | 6.68 | 53 | 2 | 2862637.914 | 2486323.48 | 2639965.559 | 2308088.594 | 3945209.464 | 2678458.24 | 3577805.25 | 1403091.897 | 9.13943 | 39.006 | Hprt | 15452 | 0.9403 | 1.059701139 | 1.12698 | 0.59215 | 0.902243 |
| 568907554 | PREDICTED: arf-GAP domain and FG repeat-containing protein 1 isoform X1 [Mus musculus] | 0 | 18.536 | 10 | 4 | 7 | 4 | 577 | 59.7 | 8.65 | 34 | 4 | 12728749.07 | 14246020.18 | 13687087.57 | 15780709.58 | 13199811.38 | 10334707.69 | 13080741.31 | 12597960.39 | 9.05388 | 10.8747 | Agfg1 | 15463 | 1.06842 | 0.931576435 | 0.87192 | 0.09852 | 0.670764 |
| 124339838 | heat shock 70 kDa protein 1-like [Mus musculus] | 0 | 26.195 | 14 | 6 | 147 | 2 | 641 | 70.6 | 6.24 | 3129 | 6 | 680719.8935 | 994478.2732 | 817074.3062 | 963028.0686 | 989966.4018 | 1388114.886 | 785368.25 | 1554903.704 | 16.7258 | 30.001 | Hspa1l | 15482 | 0.84547 | 1.154527306 | 1.36554 | 0.14959 | 0.693437 |
| 31982273 | peroxisomal multifunctional enzyme type 2 [Mus musculus] | 0 | 47.094 | 21 | 11 | 44 | 11 | 735 | 79.4 | 8.57 | 412 | 11 | 28676306.08 | 29677487.82 | 23491456.14 | 26922154.34 | 29628713.29 | 24865023.36 | 26849976.75 | 29747142.85 | 9.99228 | 8.48238 | Hsd17b4 | 15488 | 0.98943 | 1.010567953 | 1.02136 | 0.75762 | 0.952737 |
| 6680297 | dnaJ homolog subfamily A member 1 [Mus musculus] | 0 | 186.39 | 59 | 17 | 220 | 17 | 397 | 44.8 | 7.08 | 2831 | 17 | 266261175.7 | 161559213.8 | 288484548.2 | 251501120.4 | 281656258.2 | 166003280.9 | 213289804.4 | 236894540.1 | 23.0248 | 21.471 | Dnaja1 | 15502 | 1.0375 | 0.962499838 | 0.92771 | 0.65166 | 0.915864 |
| 148673922 | heat shock protein 110, isoform CRA_a, partial [Mus musculus] | 0 | 28.668 | 8 | 5 | 24 | 4 | 884 | 99.1 | 5.62 | 497 | 5 | 13211955.14 | 5417474.05 | 13845129.64 | 7030386.773 | 12120917.74 | 7350462.76 | 10749600.75 | 6378244.956 | 43.2982 | 29.8015 | Hsph1 | 15505 | 1.03818 | 0.961819184 | 0.92645 | 0.78415 | 0.952792 |
| 31560686 | heat shock-related 70 kDa protein 2 [Mus musculus] | 0 | 125.273 | 37 | 20 | 342 | 10 | 633 | 69.6 | 5.67 | 7183 | 20 | 31751412.95 | 18953785.31 | 19750150.04 | 20035992.03 | 22433279.36 | 31019821.12 | 15181893.69 | 19168839.46 | 26.9768 | 30.6771 | Hspa2 | 15512 | 1.01507 | 0.984926644 | 0.9703 | 0.8873 | 0.984249 |
| 40556608 | heat shock protein HSP 90-beta [Mus musculus] | 0 | 123.572 | 38 | 20 | 147 | 13 | 724 | 83.2 | 5.03 | 2135 | 20 | 56565856.16 | 47219990.02 | 53850918.97 | 55798545.93 | 66651778.47 | 65300611.34 | 51803409.75 | 54633842.28 | 7.96327 | 12.5443 | Hsp90ab1 | 15516 | 0.94477 | 1.055230085 | 1.11692 | 0.19698 | 0.715071 |
| 6754254 | heat shock protein HSP 90-alpha [Mus musculus] | 0 | 118.217 | 34 | 20 | 144 | 14 | 733 | 84.7 | 5.01 | 1747 | 20 | 30470343.77 | 37039398.76 | 36532246.21 | 42798862.23 | 40593449.9 | 41784757.9 | 34810274.19 | 39334315.94 | 13.724 | 7.79212 | Hsp90aa1 | 15519 | 0.96808 | 1.031915317 | 1.06593 | 0.44249 | 0.852048 |
| 60360216 | mKIAA4025 protein [Mus musculus] | 0 | 10.947 | 4 | 3 | 15 | 2 | 930 | 103.2 | 5.88 | 215 | 3 | 81367192.92 | 177223568 | 79432670.34 | 126110943.7 | 77218125.33 | 224163897.3 | 76725272 | 197504156.3 | 39.7658 | 54.2368 | Hspa4 | 15525 | 0.89278 | 1.107215699 | 1.24018 | 0.56129 | 0.895889 |
| 162461907 | stress-70 protein, mitochondrial [Mus musculus] | 0 | 10.994 | 6 | 4 | 12 | 4 | 679 | 73.4 | 6.07 | 89 | 4 | 5116290.506 | 5851411.53 | 6040226.515 | 6449574.375 | 6344134.712 | 7346984.532 | 5979695.75 | 6552963.227 | 9.51044 | 8.8183 | Hspa9 | 15526 | 0.94432 | 1.055680434 | 1.11793 | 0.13588 | 0.693437 |
| 1655901 | elav G homolog [Mus musculus] | 0 | 8.063 | 11 | 4 | 10 | 4 | 326 | 36 | 9.17 | 31 | 4 | 7328622.612 | 13740068.05 | 8529060.895 | 15946007.69 | 8781715.802 | 16925597.35 | 16218189.5 | 21745716.76 | 36.198 | 33.6349 | Elavl1 | 15568 | 0.83402 | 1.165979616 | 1.39802 | 0.22831 | 0.723653 |
| 755507811 | PREDICTED: ELAV-like protein 2 isoform X1 [Mus musculus] | 0 | 51.745 | 19 | 10 | 42 | 2 | 530 | 57.6 | 9.79 | 690 | 10 | 8823195.607 | 9395266.529 | 11771854.67 | 12718411.52 | 7635630.572 | 11997171.45 | 12981342 | 11308795.35 | 17.4762 | 21.2491 | Elavl2 | 15569 | 0.98598 | 1.014015788 | 1.02843 | 0.84569 | 0.973079 |
| 27229298 | ELAV-like protein 3 [Rattus norvegicus] | 0 | 32.78 | 19 | 7 | 29 | 3 | 367 | 39.5 | 9.28 | 497 | 7 | 8498179.166 | 3916773.592 | 4237836.46 | 4565514.706 | 8906274.206 | 7360623.595 | 11880532 | 3029509.165 | 40.4458 | 47.3292 | Elavl3 | 15571 | 0.80993 | 1.190067542 | 1.46934 | 0.28756 | 0.752733 |
| 145386533 | islet cell autoantigen 1 isoform 1 [Mus musculus] | 0 | 7.888 | 4 | 2 | 4 | 2 | 478 | 54.3 | 5.8 | 35 | 2 | 1720456.418 | 1971893.475 | 1524779.02 | 1815075.282 | 1869648.122 | 2106789.324 | 1594134.125 | 1799173.266 | 10.6318 | 11.4773 | Ica1 | 15893 | 0.97656 | 1.02343715 | 1.048 | 0.57171 | 0.895889 |
| 159110562 | intercellular adhesion molecule 5 precursor [Mus musculus] | 0 | 20.431 | 6 | 5 | 10 | 5 | 917 | 96.9 | 6.32 | 52 | 5 | 14073530.43 | 13232833.57 | 17322226.01 | 14490475.21 | 19773610.55 | 20832350.49 | 20103313.38 | 17254237.52 | 12.0018 | 7.97977 | Icam5 | 15898 | 0.86253 | 1.137467847 | 1.31875 | 0.00717 | 0.396316 |
| 6680345 | isocitrate dehydrogenase [NAD] subunit gamma 1, mitochondrial precursor [Mus musculus] | 0 | 81.685 | 39 | 11 | 61 | 11 | 393 | 42.8 | 9.01 | 838 | 11 | 84807709.21 | 56587215.52 | 86189933.74 | 56442099.92 | 82920403.37 | 58134415.75 | 90787327.75 | 61571644.52 | 23.5803 | 21.7823 | Idh3g | 15929 | 0.98374 | 1.016255924 | 1.03305 | 0.846 | 0.973079 |
| 226693333 | interleukin-1 receptor accessory protein isoform d precursor [Mus musculus] | 0 | 24.22 | 12 | 7 | 11 | 7 | 685 | 78.6 | 6.79 | 26 | 7 | 6580700.717 | 1473249.64 | 9751836.372 | 3146463.619 | 8819250.116 | 10401195.16 | 8579891.875 | 6824288.825 | 70.3399 | 16.9173 | Il1rap | 16180 | 0.75399 | 1.24600835 | 1.65255 | 0.13543 | 0.693437 |
| 568949719 | PREDICTED: integrin-linked protein kinase isoform X1 [Mus musculus] | 0 | 8.057 | 8 | 3 | 4 | 3 | 484 | 54.9 | 7.91 | 22 | 3 | 3770725.388 | 1439595.868 | 3634830.516 | 1938149.481 | 3979730.796 | 1533421.225 | 3465102 | 3080225.387 | 43.8349 | 34.9623 | Ilk | 16202 | 0.94417 | 1.055826565 | 1.11825 | 0.70115 | 0.931521 |
| 10863157 | imprinted and ancient [Mus musculus] | 0 | 19.508 | 19 | 4 | 14 | 4 | 318 | 36.2 | 5.05 | 57 | 4 | 9275824.2 | 3728105.288 | 5183391.244 | 2791972.179 | 6583211.104 | 4972599.043 | 4445456.313 | 3781238.179 | 54.5642 | 24.1749 | Impact | 16210 | 1.02936 | 0.970639463 | 0.94295 | 0.85337 | 0.975398 |
| 1669535 | scg [Mus musculus] | 0 | 98.092 | 26 | 14 | 60 | 14 | 876 | 97.1 | 4.78 | 1732 | 14 | 43372582 | 35367537.49 | 47813135.33 | 35593960.39 | 49587140.85 | 44356874.55 | 52130230.38 | 38318515.99 | 15.0824 | 13.2613 | Kpnb1 | 16211 | 0.93581 | 1.064193305 | 1.13719 | 0.24568 | 0.733806 |
| 148674210 | mCG5221, partial [Mus musculus] | 0 | 37.015 | 5 | 11 | 20 | 11 | 2498 | 285.1 | 5.06 | 128 | 11 | 11293121.38 | 5201773.136 | 14224657.64 | 11182449.7 | 11031258.41 | 2484787.162 | 9682396 | 9850064.396 | 36.1563 | 47.1798 | Cep250 | 16328 | 1.11812 | 0.881875438 | 0.78871 | 0.44649 | 0.852048 |
| 4503521 | eukaryotic translation initiation factor 3 subunit E [Homo sapiens] | 0 | 18.229 | 14 | 5 | 14 | 5 | 445 | 52.2 | 6.04 | 72 | 5 | 2694271.798 | 3116878.813 | 4464468.526 | 5455073.891 | 5153810.461 | 4718003.162 | 6271850.563 | 7031639.285 | 32.163 | 18.178 | Eif3e | 16341 | 0.80865 | 1.191348664 | 1.47325 | 0.06441 | 0.601634 |
| 6680502 | integral membrane protein 2B [Mus musculus] | 0 | 39.726 | 41 | 6 | 18 | 6 | 266 | 30.2 | 5.3 | 257 | 6 | 12504323.77 | 13856855.43 | 16142894.56 | 13222061.94 | 23946753.24 | 25938947.33 | 16709685.88 | 20753332.15 | 11.3008 | 18.4582 | Itm2b | 16432 | 0.77898 | 1.221021248 | 1.56746 | 0.01066 | 0.458267 |
| 568940836 | PREDICTED: inositol 1,4,5-trisphosphate receptor type 1 isoform X1 [Mus musculus] | 0 | 250.098 | 22 | 49 | 187 | 41 | 2758 | 314 | 6.02 | 2997 | 49 | 132296082 | 116072216.4 | 146551093.4 | 129753872.4 | 145644537.4 | 195508460.5 | 163698089.1 | 163130481.6 | 9.52002 | 12.4398 | Itpr1 | 16438 | 0.87984 | 1.120159077 | 1.27314 | 0.02541 | 0.515682 |
| 568997326 | PREDICTED: intersectin-1 isoform X1 [Mus musculus] | 0 | 173.028 | 24 | 29 | 112 | 26 | 1743 | 197.5 | 7.99 | 1890 | 29 | 66589351.13 | 55182376.12 | 86594919.33 | 56245314.16 | 62687171.42 | 85989797.27 | 83648010.33 | 86292669.13 | 22.0199 | 14.278 | Itsn1 | 16443 | 0.9074 | 1.092597644 | 1.20409 | 0.19429 | 0.715071 |
| 30520029 | caytaxin [Mus musculus] | 0 | 28.356 | 22 | 5 | 21 | 5 | 372 | 42.2 | 4.68 | 328 | 5 | 19772955.95 | 16331749.82 | 14684016.03 | 13433424.52 | 27456047.73 | 13539135.9 | 18060833.5 | 2964063.252 | 17.1147 | 65.6142 | Atcay | 16467 | 1.01744 | 0.98255682 | 0.96571 | 0.92019 | 0.987994 |
| 28395018 | junction plakoglobin [Mus musculus] | 0 | 247.876 | 60 | 28 | 194 | 24 | 745 | 81.7 | 6.14 | 4804 | 28 | 184651506.7 | 136785076.2 | 155414343.8 | 153458062.8 | 154047344.1 | 148818245.2 | 157478799.8 | 141224793.9 | 12.6231 | 4.70329 | Jup | 16480 | 1.02333 | 0.976669928 | 0.9544 | 0.52147 | 0.881256 |
| 119395752 | potassium voltage-gated channel subfamily A member 1 [Mus musculus] | 0 | 30.453 | 17 | 6 | 24 | 3 | 495 | 56.4 | 5.14 | 170 | 6 | 4489913.719 | 6806611.945 | 8208501.516 | 4828572.311 | 8886275.594 | 4919942.18 | 9617284.25 | 4474994.461 | 28.7126 | 38.0313 | Kcna1 | 16485 | 0.93175 | 1.068251081 | 1.1465 | 0.59497 | 0.902243 |
| 25742772 | potassium voltage-gated channel subfamily A member 2 [Rattus norvegicus] | 0 | 30.748 | 17 | 5 | 27 | 2 | 499 | 56.7 | 4.86 | 390 | 5 | 14038204.24 | 9471183.132 | 12967725.4 | 11714596.22 | 12136343.34 | 13200027.5 | 10145666.94 | 13107007.23 | 16.2917 | 11.677 | Kcna2 | 16490 | 0.99589 | 1.004105527 | 1.00824 | 0.93729 | 0.994906 |
| 31543026 | potassium voltage-gated channel subfamily A member 4 [Mus musculus] | 0 | 20.329 | 9 | 4 | 9 | 3 | 654 | 73.4 | 5.29 | 96 | 4 | 4148112.178 | 1807644.302 | 3144646.969 | 2254322.898 | 6726471.724 | 3212391.424 | 5915180.25 | 3563968.797 | 36.4553 | 35.662 | Kcna4 | 16492 | 0.73797 | 1.262026918 | 1.71013 | 0.09257 | 0.662454 |
| 1695270 | K+ channel beta2 subunit [Mus musculus] | 0 | 71.346 | 46 | 12 | 63 | 10 | 367 | 41 | 9 | 438 | 12 | 58623743.68 | 57192147.57 | 75104533.78 | 54466348.17 | 80056590.71 | 60180722.78 | 75765846 | 55957180.45 | 15.2129 | 17.2329 | Kcnab2 | 16498 | 0.94863 | 1.051365062 | 1.10829 | 0.40922 | 0.834385 |
| 200976 | potassium channel protein [Mus musculus] | 0 | 12.694 | 4 | 3 | 14 | 3 | 857 | 95.6 | 8.25 | 105 | 3 | 8845608.105 | 10409716.46 | 8286703.44 | 11644735.35 | 7719068.969 | 7905337.494 | 5609373.75 | 9117874.514 | 15.5652 | 19.2088 | Kcnb1 | 16500 | 1.12705 | 0.872946367 | 0.77454 | 0.08113 | 0.642075 |
| 26006243 | mKIAA1044 protein [Mus musculus] | 0 | 19.175 | 12 | 5 | 14 | 5 | 636 | 71.2 | 8.07 | 138 | 5 | 17935747.07 | 7460445.213 | 8182832.572 | 5102803.772 | 7834393.561 | 5934496.628 | 7784502.875 | 8934569.662 | 58.5802 | 16.3202 | Kcnd2 | 16508 | 1.11846 | 0.88153982 | 0.78817 | 0.50644 | 0.88031 |
| 390013285 | ATP-sensitive inward rectifier potassium channel 12 isoform 1 [Mus musculus] | 0 | 30.841 | 15 | 7 | 28 | 4 | 529 | 59.3 | 6.92 | 492 | 7 | 12005075.15 | 1681120.768 | 8814726.784 | 1983242.833 | 8672970.432 | 2222101.112 | 8331440.75 | 2167409.213 | 83.6825 | 68.1379 | Kcnj12 | 16515 | 1.06736 | 0.932642265 | 0.87379 | 0.81404 | 0.961921 |
| 158711668 | inward rectifier potassium channel 4 [Mus musculus] | 0 | 87.667 | 35 | 12 | 93 | 10 | 445 | 49.6 | 6.21 | 1850 | 12 | 294612452.6 | 249939580 | 288996829.7 | 222823352.8 | 316347318.5 | 317923512.6 | 303748882.8 | 240993630.5 | 12.8504 | 12.3482 | Kcnj4 | 16520 | 0.94514 | 1.054863524 | 1.1161 | 0.26396 | 0.745606 |
| 1039745588 | PREDICTED: calcium-activated potassium channel subunit alpha-1 isoform X12 [Mus musculus] | 0 | 185.487 | 28 | 24 | 140 | 24 | 1262 | 140.3 | 7.03 | 2201 | 24 | 121614040.3 | 99188733.01 | 139725574 | 131156908.1 | 120820918.7 | 125374533.8 | 135185443.8 | 111020594.1 | 14.2088 | 8.15504 | Kcnma1 | 16531 | 0.99927 | 1.000727817 | 1.00146 | 0.98639 | 0.998251 |
| 1039759174 | PREDICTED: potassium voltage-gated channel subfamily KQT member 2 isoform X1 [Mus musculus] | 0 | 38.829 | 13 | 10 | 33 | 10 | 937 | 103 | 9.29 | 157 | 10 | 19894403.22 | 25962649.59 | 26483503.76 | 11879105.12 | 18025861.73 | 5546410.349 | 23027892.75 | 7317834.572 | 32.3404 | 62.4729 | Kcnq2 | 16536 | 1.21936 | 0.780641557 | 0.64021 | 0.21132 | 0.716066 |
| 568908175 | PREDICTED: kinesin-like protein KIF1A isoform X1 [Mus musculus] | 0 | 18.846 | 3 | 4 | 14 | 4 | 1800 | 203.1 | 5.47 | 201 | 4 | 3809189.746 | 3815706.911 | 2231887.556 | 3846851.38 | 6423066.189 | 2486380.847 | 6234898.75 | 1905936.968 | 23.2401 | 56.2819 | Kif1a | 16560 | 0.89118 | 1.108820187 | 1.24422 | 0.53256 | 0.886555 |
| 568984232 | PREDICTED: kinesin-like protein KIF2A isoform X1 [Mus musculus] | 0 | 195.623 | 43 | 24 | 192 | 24 | 743 | 83.8 | 6.42 | 3736 | 24 | 323146654.1 | 305067186.7 | 293449005.2 | 302139896.7 | 364002228.3 | 286632896.2 | 324371271.5 | 263374815.8 | 4.07928 | 14.2537 | Kif2a | 16563 | 0.99408 | 1.00592095 | 1.01191 | 0.87893 | 0.980491 |
| 341940868 | RecName: Full=Kinesin-like protein KIF21B; AltName: Full=Kinesin-like protein KIF6 | 0 | 14.637 | 3 | 5 | 23 | 3 | 1668 | 186.1 | 7.08 | 98 | 5 | 3308926.742 | 4346058.573 | 3969860.6 | 3876560.05 | 4030586.489 | 3477603.417 | 3106652.938 | 4353907.274 | 11.0621 | 14.8886 | Kif21b | 16565 | 1.01748 | 0.982518769 | 0.96564 | 0.71785 | 0.938535 |
| 568971862 | PREDICTED: kinesin-like protein KIF3A isoform X1 [Mus musculus] | 0 | 19.798 | 7 | 4 | 12 | 4 | 725 | 82.6 | 6.39 | 698 | 4 | 3596399.795 | 2063604.554 | 5570942.422 | 2707495.351 | 5308189.109 | 3188490.266 | 4919549.625 | 2680045.786 | 43.8001 | 31.9385 | Kif3a | 16568 | 0.92816 | 1.071844615 | 1.15481 | 0.60817 | 0.905186 |
| 26006171 | mKIAA0531 protein [Mus musculus] | 0 | 96.3 | 28 | 22 | 84 | 10 | 987 | 112.6 | 6.42 | 942 | 22 | 12633049.1 | 8873386.412 | 12407294.81 | 9128293.164 | 9593074.98 | 10511457.2 | 12599327.63 | 26678776.73 | 18.927 | 53.8097 | Kif5c | 16574 | 0.84046 | 1.159537879 | 1.37964 | 0.35992 | 0.80334 |
| 568910514 | PREDICTED: kinesin-associated protein 3 isoform X1 [Mus musculus] | 0 | 13.736 | 6 | 4 | 7 | 4 | 809 | 92.8 | 5.08 | 29 | 4 | 3584106.265 | 2024255.582 | 5015698.273 | 4455667.625 | 3872612.572 | 4799955.444 | 4238786.563 | 4672018.286 | 34.5991 | 9.63374 | Kifap3 | 16579 | 0.92335 | 1.076650565 | 1.16603 | 0.39653 | 0.824613 |
| 12850652 | unnamed protein product [Mus musculus] | 0 | 17.616 | 12 | 4 | 18 | 4 | 419 | 46.4 | 6.04 | 97 | 4 | 5692260.59 | 4134455.551 | 4268452.954 | 2510898.179 | 5846484.923 | 3580285.46 | 5886329.813 | 2249984.209 | 31.3431 | 40.732 | Uhmk1 | 16589 | 0.97199 | 1.02800822 | 1.05763 | 0.83588 | 0.969252 |
| 238537882 | Chain A, Crystal Structure Of The Nuclear Export Complex Crm1- Snurportin1-Rangtp | 0 | 22.272 | 7 | 6 | 19 | 6 | 1073 | 123.2 | 6.07 | 319 | 6 | 5156551.587 | 4194289.794 | 8286476.405 | 10191716.62 | 6472816.261 | 4198343.095 | 7370322.625 | 3539468.506 | 39.8892 | 33.7358 | Fabp5 | 16592 | 1.12645 | 0.873546128 | 0.77548 | 0.38288 | 0.816843 |
| 131412178 | kinesin light chain 1 isoform 1D [Mus musculus] | 0 | 44.576 | 21 | 12 | 28 | 8 | 616 | 69.5 | 5.96 | 241 | 12 | 80897689.88 | 78984478.17 | 99175427.35 | 39301236.26 | 98695840 | 108095972 | 101049185.5 | 126673675.5 | 33.8178 | 11.6688 | Klc1 | 16593 | 0.81422 | 1.18578355 | 1.45635 | 0.05247 | 0.595816 |
| 1039757216 | PREDICTED: kinesin light chain 2 isoform X1 [Mus musculus] | 0 | 38.897 | 20 | 11 | 29 | 7 | 636 | 70.2 | 7.06 | 356 | 11 | 12248046.93 | 12338645.11 | 11325044.16 | 9992728.299 | 13189984.74 | 13168528.49 | 9790686.875 | 11653989.92 | 9.49638 | 13.4687 | Klc2 | 16594 | 0.97974 | 1.020262224 | 1.04136 | 0.64261 | 0.915705 |
| 1197836 | SRP1 [Mus musculus] | 0 | 46.878 | 19 | 7 | 29 | 3 | 538 | 60.1 | 5.01 | 715 | 7 | 4583264.859 | 835074.9972 | 3676327.099 | 3496109.544 | 1317033.131 | 1385639.268 | 1497499.375 | 348424.879 | 51.259 | 46.7002 | Kpna1 | 16646 | 1.46922 | 0.530777481 | 0.36126 | 0.05573 | 0.599194 |
| 568928522 | PREDICTED: importin subunit alpha-7 isoform X1 [Mus musculus] | 0 | 39.127 | 17 | 7 | 28 | 3 | 541 | 60.5 | 4.98 | 696 | 7 | 2496092.997 | 1165715.521 | 3342542.19 | 757625.3237 | 1153139.003 | 1417618.616 | 984646.625 | 719100.5166 | 61.509 | 27.4509 | Kpna6 | 16650 | 1.28974 | 0.710258228 | 0.5507 | 0.20578 | 0.715071 |
| 755537500 | PREDICTED: keratin, type I cytoskeletal 10 isoform X1 [Mus musculus] | 0 | 37.699 | 11 | 8 | 120 | 5 | 601 | 61.6 | 5.12 | 2369 | 8 | 34160796.53 | 40781186.17 | 61136297.19 | 31106945.67 | 30820677.25 | 83394014.22 | 14862278.78 | 82139192.72 | 32.3255 | 66.6797 | Krt10 | 16661 | 0.88364 | 1.116360401 | 1.26337 | 0.58064 | 0.898781 |
| 755537506 | PREDICTED: kinase suppressor of Ras 1 isoform X1 [Mus musculus] | 0 | 12.503 | 4 | 3 | 5 | 3 | 914 | 101.8 | 8.72 | 126 | 3 | 7461139.257 | 6013120.638 | 6736916.065 | 2435854.671 | 5788834.416 | 3507752.328 | 3352115 | 3539039.832 | 39.3936 | 28.766 | Ksr1 | 16706 | 1.16633 | 0.833672539 | 0.71478 | 0.24659 | 0.733806 |
| 568986880 | PREDICTED: kinectin isoform X3 [Mus musculus] | 0 | 173.789 | 29 | 28 | 98 | 28 | 1304 | 149.9 | 5.94 | 1706 | 28 | 65931320.83 | 43649648.54 | 63315806.44 | 61707086.74 | 70248139.16 | 69241971.15 | 77026318.38 | 65329450.08 | 17.3079 | 6.90268 | Ktn1 | 16709 | 0.90853 | 1.091474567 | 1.20137 | 0.08065 | 0.642075 |
| 112293279 | neural cell adhesion molecule L1 precursor [Mus musculus] | 0 | 111.344 | 20 | 15 | 74 | 15 | 1259 | 140.8 | 5.96 | 1120 | 15 | 151975003 | 138265638.3 | 180865043 | 147803458.6 | 141191059 | 75613832.25 | 139603674.6 | 89481897.98 | 11.8567 | 30.395 | L1cam | 16728 | 1.16249 | 0.837510572 | 0.72045 | 0.06587 | 0.601634 |
| 6754508 | LIM and SH3 domain protein 1 [Mus musculus] | 0 | 15.26 | 10 | 2 | 14 | 2 | 263 | 30 | 7.05 | 476 | 2 | 12998026.03 | 12573163.47 | 16905189.11 | 12954760.63 | 16805210.82 | 14067273.99 | 15570050.5 | 21363568.99 | 14.725 | 18.566 | Lasp1 | 16796 | 0.89958 | 1.100415789 | 1.22325 | 0.15011 | 0.693437 |
| 568921915 | PREDICTED: rho guanine nucleotide exchange factor 2 isoform X1 [Mus musculus] | 0 | 272.794 | 43 | 37 | 228 | 37 | 1185 | 133.8 | 6.57 | 4307 | 37 | 297530550.1 | 246084019.5 | 285047692.6 | 248352286.1 | 280345584.3 | 234719346.9 | 259520911.3 | 239140920.7 | 9.64378 | 8.26372 | Arhgef2 | 16800 | 1.03027 | 0.9697295 | 0.94124 | 0.37946 | 0.81223 |
| 257743039 | L-lactate dehydrogenase A chain isoform 2 [Mus musculus] | 0 | 36.216 | 25 | 7 | 34 | 6 | 361 | 39.7 | 8.35 | 633 | 7 | 14067797.78 | 12147918.98 | 16809870.94 | 12229137.23 | 20809514.04 | 12816266.54 | 17240302 | 11487981.69 | 15.8202 | 27.3389 | Ldha | 16828 | 0.93964 | 1.06036402 | 1.12848 | 0.48658 | 0.869379 |
| 18087805 | 40S ribosomal protein S2 [Mus musculus] | 0 | 96.759 | 48 | 16 | 182 | 16 | 293 | 31.2 | 10.24 | 2176 | 16 | 321122729 | 354496640.7 | 344436039.6 | 366952247.3 | 393349582.8 | 369952014.5 | 388221623.4 | 417968284.7 | 5.59768 | 5.04598 | Rps2 | 16898 | 0.93828 | 1.06172295 | 1.13157 | 0.0166 | 0.461382 |
| 12835914 | unnamed protein product [Mus musculus] | 0 | 280.941 | 41 | [illegible] | [illegible] | [illegible] | [illegible] | [illegible] | [illegible] | 18057 | 41 | [illegible] | [illegible] | [illegible] | [illegible] | [illegible] | [illegible] | [illegible] | [illegible] | [illegible] | [illegible] | [illegible] | [illegible] | [illegible] | [illegible] | [illegible] | [illegible] | [illegible] |
| 188219589 | lamin-B1 [Mus musculus] | 0 | 361.477 | [illegible] | [illegible] | [illegible] | [illegible] | [illegible] | [illegible] | [illegible] | [illegible] | [illegible] | [illegible] | [illegible] | [illegible] | [illegible] | [illegible] | [illegible] | [illegible] | [illegible] | [illegible] | [illegible] | [illegible] | [illegible] | [illegible] | [illegible] | [illegible] | [illegible] | [illegible] |
| 124494256 | prolow-density lipoprotein receptor-related protein 1 precursor [Mus musculus] | 0 | 16.529 | [illegible] | [illegible] | [illegible] | [illegible] | 4545 | 504.4 | 5.36 | 59 | [illegible] | [illegible] | [illegible] | [illegible] | [illegible] | [illegible] | [illegible] | [illegible] | [illegible] | [illegible] | [illegible] | Lrp1 | [illegible] | [illegible] | [illegible] | [illegible] | [illegible] | [illegible] |

| | | | | | | | | | | | | | | | | | | | | | | | | | | | | | |
|---|---|---|---|---|---|---|---|---|---|---|---|---|---|---|---|---|---|---|---|---|---|---|---|---|---|---|---|---|---|
| 568908290 | PREDICTED: leucine-rich repeat flightless-interacting protein 1 isoform X27 [Mus musculus] | 0 | 47.24 | 17 | 8 | 44 | 7 | 688 | 77.9 | 6.18 | 1143 | 8 | 29142276.73 | 21621690.31 | 27658075.29 | 21384036.69 | 35021777.08 | 16335292.9 | 31188257.5 | 23136837.74 | 16.148 | 31.6072 | Lrrfip1 | 16978 | 0.9714 | 1.028595729 | 1.05888 | 0.76208 | 0.952792 |
| 568978894 | PREDICTED: MAP/microtubule affinity-regulating kinase 3 isoform X1 [Mus musculus] | 0 | 203.623 | 38 | 27 | 245 | 14 | 806 | 90 | 9.47 | 3937 | 27 | 76442331.86 | 79226520.19 | 71473691.91 | 75964542.93 | 75530037.59 | 69026168.57 | 75520433.88 | 60911615.88 | 4.23497 | 9.87492 | Mark3 | 17169 | 1.03787 | 0.962131472 | 0.92703 | 0.19806 | 0.715071 |
| 568966823 | PREDICTED: megakaryocyte-associated tyrosine-protein kinase isoform X1 [Mus musculus] | 0 | 15.735 | 9 | 4 | 20 | 4 | 506 | 56.2 | 8.9 | 239 | 4 | 8046100.64 | 10956112.22 | 6360540.086 | 11735249.27 | 13667927.76 | 4635402.349 | 7647608.406 | 21137891 | 27.0512 | 61.8913 | Matk | 17179 | 0.88133 | 1.118674466 | 1.26931 | 0.54081 | 0.886555 |
| 755498836 | PREDICTED: matrilin-4 isoform X1 [Mus musculus] | 0 | 25.415 | 10 | 5 | 21 | 5 | 636 | 70.2 | 5.54 | 343 | 5 | 5915270.207 | 5871187.079 | 4598508.134 | 2979229.633 | 6368229.063 | 8710284.351 | 5703994 | 7507510.718 | 28.5738 | 18.6873 | Matn4 | 17183 | 0.8127 | 1.187303965 | 1.46094 | 0.05842 | 0.599194 |
| 37360062 | mKIAA0723 protein [Mus musculus] | 0 | 118.49 | 34 | 17 | 85 | 17 | 852 | 95.2 | 6.27 | 1214 | 17 | 76999804.6 | 81191718.67 | 83667257.45 | 83071146.95 | 56611827.04 | 92511269.67 | 83862167.75 | 77548947.5 | 3.70853 | 19.7051 | Matr3 | 17184 | 1.02265 | 0.97734614 | 0.9557 | 0.66059 | 0.919605 |
| 568953134 | PREDICTED: guanine nucleotide exchange factor DBS isoform X1 [Mus musculus] | 0 | 17.039 | 4 | 4 | 5 | 3 | 1211 | 135.6 | 6.04 | 34 | 4 | 705996.4056 | 621271.0658 | 677558.1811 | 1252071.245 | 1859084.174 | 737859.6208 | 2308845.5 | 596642.2527 | 36.1096 | 61.084 | Mcf2l | 17207 | 0.74364 | 1.256359225 | 1.68947 | 0.25405 | 0.737014 |
| 251823814 | nuclear protein MDM1 isoform 3 [Mus musculus] | 0 | 46.244 | 16 | 9 | 27 | 9 | 718 | 80.8 | 9.44 | 145 | 9 | 26838865.41 | 59960082.6 | 52548563.73 | 55634808.36 | 39221913.07 | 68679982.43 | 46783499.81 | 88173943.13 | 30.6027 | 36.5019 | Mdm1 | 17245 | 0.89065 | 1.109347791 | 1.24555 | 0.40473 | 0.828732 |
| 126517469 | methyl-CpG-binding protein 2 isoform 1 [Mus musculus] | 0 | 29.476 | 21 | 7 | 42 | 7 | 501 | 53.5 | 9.88 | 603 | 7 | 7748005.095 | 6901460.61 | 19402790.32 | 20854582.73 | 19741972.58 | 26555911.86 | 21490945.5 | 33490730.38 | 54.0853 | 24.353 | Mecp2 | 17257 | 0.70309 | 1.296906273 | 1.84457 | 0.05311 | 0.595816 |
| 7330691 | MOBP170 [Mus musculus] | 0 | 43.173 | 28 | 5 | 82 | 5 | 170 | 19.2 | 11.14 | 757 | 5 | 285544927.5 | 277443705.7 | 191749256.6 | 224072853.8 | 234511564.7 | 249778600.4 | 255383375.3 | 222892210.3 | 18.2296 | 6.1329 | Mobp | 17433 | 1.00837 | 0.991632229 | 0.9834 | 0.86844 | 0.97927 |
| 29747959 | myelin oligodendrocyte glycoprotein, partial [Mus musculus] | 0 | 24.9 | 23 | 6 | 40 | 6 | 274 | 31.1 | 8.54 | 670 | 6 | 113287445.1 | 107448337.3 | 75980089.83 | 87564913.94 | 96373101.77 | 108211996.8 | 93568678.75 | 120901873.8 | 18.0475 | 11.9228 | Mog | 17441 | 0.95671 | 1.043288046 | 1.09049 | 0.44691 | 0.852048 |
| 31982186 | malate dehydrogenase, mitochondrial precursor [Mus musculus] | 0 | 27.996 | 25 | 7 | 21 | 7 | 338 | 35.6 | 8.68 | 262 | 7 | 13448535.81 | 14677236.58 | 19637736.65 | 14827369.24 | 16683426.89 | 16760750.53 | 15540644.25 | 14984541.92 | 17.4516 | 5.46048 | Mdh2 | 17448 | 0.98911 | 1.01089193 | 1.02202 | 0.81802 | 0.964114 |
| 148675904 | malate dehydrogenase 1, NAD (soluble), isoform CRA_c, partial [Mus musculus] | 0 | 39.216 | 28 | 7 | 28 | 7 | 366 | 40 | 7.33 | 658 | 7 | 41762315.18 | 29145612.93 | 19708603.57 | 21532690.75 | 18497409.54 | 26451726.02 | 28550469.25 | 23921365.95 | 35.7429 | 17.8181 | Mdh1 | 17449 | 1.07028 | 0.92972163 | 0.86867 | 0.52521 | 0.883731 |
| 765826460 | multiple PDZ domain protein isoform 1 [Mus musculus] | 0 | 10.523 | 2 | 3 | 6 | 3 | 2069 | 220.1 | 5.02 | 66 | 3 | 15809874.71 | 14939288.44 | 15814847.13 | 12154745.63 | 13744344.12 | 15393548.18 | 16003200.38 | 6033610.857 | 11.8046 | 36.0069 | Mpdz | 17475 | 1.06865 | 0.931351216 | 0.87152 | 0.47251 | 0.863085 |
| 6678924 | 55 kDa erythrocyte membrane protein [Mus musculus] | 0 | 7.203 | 8 | 3 | 6 | 3 | 466 | 52.2 | 7.2 | 0 | 3 | 796983.1459 | 2743644.004 | 1780123.444 | 2566609.799 | 2247960.652 | 3239995.706 | 1942337.625 | 3974816.2 | 45.0421 | 32.6729 | Mpp1 | 17524 | 0.81766 | 1.182337963 | 1.446 | 0.22077 | 0.718033 |
| 887222486 | protein Mpv17 isoform 3 [Mus musculus] | 0 | 17.154 | 19 | 2 | 9 | 2 | 178 | 19.9 | 9.57 | 435 | 2 | 12964036.17 | 18077955.42 | 12005409.6 | 11062728.83 | 12271509.41 | 32583674.12 | 9063257 | 22244163.79 | 23.148 | 55.8324 | Mpv17 | 17527 | 0.83072 | 1.169279277 | 1.40755 | 0.35819 | 0.801262 |
| 124244033 | microtubule-associated protein 1A isoform 1 [Mus musculus] | 0 | 663.17 | 40 | 89 | 918 | 87 | 3014 | 325.7 | 5.01 | 17698 | 89 | 1489121362 | 1522967068 | 1399199144 | 1534652326 | 1188614069 | 1299589910 | 1390657122 | 1540282849 | 4.12451 | 10.9776 | Map1a | 17754 | 1.04635 | 0.953647861 | 0.9114 | 0.15266 | 0.693437 |
| 171543853 | microtubule-associated protein 1B [Mus musculus] | 0 | 1032.54 | 58 | 101 | 1170 | 99 | 2464 | 270.1 | 4.83 | 27324 | 101 | 2613730542 | 2364730919 | 2569312245 | 2331431241 | 2411844869 | 2005934725 | 2627024288 | 2374474105 | 5.76426 | 10.9497 | Map1b | 17755 | 1.02383 | 0.976167714 | 0.95344 | 0.46462 | 0.863085 |
| 148667815 | mCG121680 [Mus musculus] | 0 | 947.721 | 66 | 92 | 1154 | 2 | 1828 | 198.9 | 4.88 | 19728 | 92 | 19391069.02 | 48160740.15 | 36284102.61 | 58276339.76 | 43082418.24 | 49516561.28 | 31365151.5 | 56862480.49 | 41.2398 | 23.9112 | Map2 | 17756 | 0.94543 | 1.054570543 | 1.11544 | 0.65488 | 0.916931 |
| 113204613 | microtubule-associated protein 6 isoform 1 [Mus musculus] | 0 | 294.707 | 66 | 48 | 683 | 48 | 906 | 96.4 | 9.5 | 10995 | 48 | 488479384.3 | 904330224.1 | 1045510641 | 1236323740 | 870242483.1 | 973340710.6 | 1016708092 | 1544824214 | 34.5522 | 27.4236 | Map6 | 17760 | 0.90959 | 1.090407581 | 1.19879 | 0.43642 | 0.852048 |
| 568965077 | PREDICTED: ensconsin isoform X1 [Mus musculus] | 0 | 17.205 | 5 | 3 | 17 | 3 | 760 | 85.6 | 9.64 | 310 | 3 | 3745986.431 | 5790500.274 | 7777260.614 | 5427103.265 | 8089198.762 | 4689298.118 | 7337523.125 | 4857774.253 | 29.1069 | 27.6481 | Map7 | 17761 | 0.9532 | 1.046797868 | 1.09819 | 0.65705 | 0.918974 |
| 148702269 | microtubule-associated protein tau, isoform CRA_g, partial [Mus musculus] | 0 | 135.644 | 56 | 15 | 157 | 3 | 374 | 39.1 | 9.5 | 2689 | 15 | 10781772.75 | 36420484.35 | 133455535.4 | 155465700.9 | 137419126.7 | 138001631.4 | 149244366.8 | 175838926.8 | 84.646 | 11.98 | Mapt | 17762 | 0.71773 | 1.282268613 | 1.78656 | 0.12166 | 0.684314 |
| 244790087 | myelin expression factor 2 isoform 1 [Mus musculus] | 0 | 52.61 | 26 | 13 | 51 | 13 | 591 | 63.3 | 8.87 | 388 | 13 | 66899837.17 | 94514270.75 | 94304498.07 | 99647498.55 | 98429713.77 | 74207619.35 | 108963040.8 | 91692816.79 | 16.6983 | 15.6355 | Myef2 | 17876 | 0.9754 | 1.024602837 | 1.05045 | 0.68168 | 0.926957 |
| 114326446 | myosin-9 [Mus musculus] | 0 | 808.633 | 53 | 105 | 758 | 83 | 1960 | 226.2 | 5.66 | 15641 | 105 | 756083422 | 684748044.8 | 824426150.1 | 694254584.6 | 1116753459 | 538298266.7 | 750488642.3 | 627344451.3 | 8.73547 | 33.5472 | Myh9 | 17886 | 0.98776 | 1.012244285 | 1.02479 | 0.8934 | 0.984401 |
| 148702277 | myosin, light polypeptide 4, isoform CRA_a [Mus musculus] | 0 | 21.121 | 26 | 4 | 20 | 4 | 205 | 22.3 | 5.05 | 300 | 4 | 6716114.557 | 8645141.332 | 6725027.409 | 5041773.984 | 11623484.13 | 6035474.55 | 9262464.25 | 4172285.468 | 21.716 | 42.6848 | Myl4 | 17896 | 0.93189 | 1.068112862 | 1.14618 | 0.60463 | 0.905186 |
| 33620739 | myosin light polypeptide 6 isoform MLC3sm [Mus musculus] | 0 | 98.458 | 84 | 11 | 175 | 10 | 151 | 17 | 4.55 | 4060 | 11 | 568688026.5 | 583288553.6 | 572897550.9 | 503887819.1 | 802080376.1 | 362434389.3 | 514559460.3 | 394123988.1 | 6.47194 | 38.6294 | Myl6 | 17904 | 1.03616 | 0.963838871 | 0.9302 | 0.71539 | 0.936732 |
| 568905921 | PREDICTED: unconventional myosin-Ib isoform X1 [Mus musculus] | 0 | 222.814 | 40 | 36 | 148 | 36 | 1138 | 132.2 | 9.33 | 2083 | 36 | 253962391 | 182960852.7 | 222964737.2 | 204323843.4 | 270296201.5 | 223134247.6 | 224440952.6 | 185405587.9 | 13.9306 | 15.3839 | Myo1b | 17912 | 0.9779 | 1.022102072 | 1.0452 | 0.68569 | 0.92798 |
| 568972076 | PREDICTED: unconventional myosin-Ic isoform X1 [Mus musculus] | 0 | 49.034 | 14 | 13 | 40 | 13 | 1063 | 121.9 | 9.35 | 299 | 13 | 19901421.49 | 13638191.61 | 16554050.59 | 13884961.24 | 18037164.45 | 12295054.55 | 16024466.81 | 10836153.35 | 18.2562 | 23.1814 | Myo1c | 17913 | 1.056 | 0.943998483 | 0.89394 | 0.47157 | 0.863085 |
| 568960781 | PREDICTED: unconventional myosin-Va isoform X3 [Mus musculus] | 0 | 971.774 | 61 | 109 | 1350 | 103 | 1855 | 215.4 | 8.5 | 24144 | 109 | 3571855354 | 3294201053 | 3522130258 | 3402804604 | 3469413446 | 3207307982 | 3348376141 | 3065854555 | 3.61226 | 5.33582 | Myo5a | 17918 | 1.02604 | 0.97395876 | 0.94924 | 0.15383 | 0.697078 |
| 755529764 | PREDICTED: unconventional myosin-VI isoform X1 [Mus musculus] | 0 | 199.836 | 36 | 36 | 159 | 36 | 1297 | 149.5 | 8.34 | 3435 | 36 | 153258767.2 | 123917284.3 | 551072374 | 280723608.3 | 181689661.3 | 158605410.8 | 163020293.4 | 176817467.9 | 70.2739 | 6.4575 | Myo6 | 17920 | 1.23969 | 0.760305163 | 0.6133 | 0.31398 | 0.76536 |
| 568966888 | PREDICTED: protein phosphatase 1 regulatory subunit 12A isoform X1 [Mus musculus] | 0 | 71.345 | 16 | 12 | 38 | 11 | 1029 | 114.9 | 5.49 | 918 | 12 | 39433694.66 | 29833326.32 | 35203936.43 | 30582203.69 | 30801997.83 | 25688208.63 | 30919909.5 | 29900587.52 | 13.2227 | 8.4175 | Ppp1r12a | 17931 | 1.07031 | 0.929694936 | 0.86863 | 0.1327 | 0.693437 |
| 29789104 | beta-soluble NSF attachment protein [Mus musculus] | 0 | 28.55 | 24 | 5 | 18 | 4 | 298 | 33.5 | 5.47 | 306 | 5 | 11711573.54 | 12548254.14 | 14600224.46 | 13807443.49 | 15132307.23 | 14297110.16 | 14463144 | 12965152.17 | 9.7709 | 6.38593 | Napb | 17957 | 0.96174 | 1.038258022 | 1.07956 | 0.23164 | 0.727645 |
| 190610036 | cytoplasmic protein NCK2 [Mus musculus] | 0 | 10.945 | 7 | 3 | 18 | 3 | 380 | 42.9 | 6.95 | 111 | 3 | 9481005.972 | 7079949.405 | 8049104.729 | 6379148.21 | 6387752.406 | 7797874.671 | 7764319.25 | 7526851.81 | 17.34 | 9.02847 | Nck2 | 17974 | 1.02501 | 0.974987431 | 0.9512 | 0.63194 | 0.912586 |
| 13529464 | Nucleolin [Mus musculus] | 0 | 55.168 | 19 | 14 | 54 | 14 | 707 | 76.7 | 4.75 | 600 | 14 | 47253693.01 | 26421664 | 36764380.5 | 35565650.65 | 20087009.98 | 72962416.88 | 38385612 | 41862092.94 | 23.3623 | 50.6559 | Ncl | 17975 | 0.91453 | 1.085472998 | 1.18692 | 0.5833 | 0.898781 |
| 33563266 | cytochrome c oxidase subunit NDUFA4 [Mus musculus] | 0 | 39.464 | 78 | 8 | 112 | 8 | 82 | 9.3 | 9.52 | 1198 | 8 | 329740322.2 | 307473469.9 | 305652757.4 | 247271360.1 | 479998432.5 | 541079231.3 | 330274653.9 | 427141230.1 | 11.8483 | 20.0898 | Ndufa4 | 17992 | 0.80181 | 1.198190866 | 1.49436 | 0.02213 | 0.505785 |
| 1374782 | Nedd4 [Mus musculus] | 0 | 55.581 | 16 | 12 | 54 | 7 | 957 | 109.9 | 5.55 | 753 | 12 | 16289560.56 | 14850638.76 | 16443555.82 | 11928383.86 | 15171970.89 | 14851159.02 | 15991504.91 | 12963881.62 | 14.0691 | 8.68513 | Nedd4 | 17999 | 1.0045 | 0.995496501 | 0.99103 | 0.91696 | 0.987992 |
| 755493895 | PREDICTED: septin-2 isoform X1 [Mus musculus] | 0 | 56.991 | 35 | 8 | 20 | 8 | 395 | 45.2 | 7.58 | 462 | 8 | 29031905.37 | 39058996.87 | 18551905.01 | 20629280.12 | 22651071 | 31379657.57 | 23862521.25 | 27301386.89 | 34.805 | 14.8996 | 43710 | 18000 | 1.00978 | 0.990222231 | 0.98063 | 0.92162 | 0.987994 |
| 568972092 | PREDICTED: neurofibromin isoform X1 [Mus musculus] | 0 | 196.991 | 18 | 38 | 117 | 38 | 2852 | 320.5 | 7.42 | 1746 | 38 | 78914033.48 | 73085720.2 | 98445347.7 | 71858963.38 | 92959880.94 | 69035489.51 | 78001612.06 | 65178254.13 | 15.2701 | 16.1758 | Nf1 | 18015 | 1.0273 | 0.972702163 | 0.94686 | 0.64057 | 0.915433 |
| 568956312 | PREDICTED: nuclear factor 1 X-type isoform X5 [Mus musculus] | 0 | 22.626 | 11 | 4 | 16 | 4 | 583 | 64.1 | 9.13 | 86 | 4 | 8075446.143 | 7756634.619 | 10138196.82 | 10829332.94 | 11609157.6 | 10907658.92 | 10664967.69 | 10505972.75 | 16.4642 | 4.459 | Nfix | 18032 | 0.91442 | 1.085580466 | 1.18718 | 0.0736 | 0.621937 |
| 39204499 | neurofilament light polypeptide [Mus musculus] | 0 | 630.505 | 68 | 45 | 1348 | 24 | 543 | 61.5 | 4.64 | 25269 | 45 | 5191431850 | 4905813419 | 4910733608 | 4178703088 | 4382294930 | 3299012835 | 6505751967 | 4387382189 | 9.02851 | 28.9175 | Nefl | 18039 | 1.01621 | 0.983786498 | 0.96809 | 0.83543 | 0.969252 |
| 112363107 | neurofilament medium polypeptide [Mus musculus] | 0 | 796.249 | 73 | 69 | 2329 | 63 | 848 | 95.9 | 4.78 | 41312 | 69 | 12447116311 | 14279945480 | 14571588798 | 13340930089 | 12912924098 | 8916212425 | 16484704529 | 13064724688 | 7.05757 | 24.0951 | Nefm | 18040 | 1.03076 | 0.969240974 | 0.94032 | 0.63289 | 0.912586 |
| 568969995 | PREDICTED: protein NipSnap homolog 1 isoform X1 [Mus musculus] | 0 | 73.154 | 48 | 9 | 50 | 9 | 287 | 33.7 | 8.66 | 1211 | 9 | 197620288.7 | 204767246 | 202165567.2 | 214661751.2 | 209641502.4 | 237446152 | 204817704.5 | 203767803.1 | 3.51809 | 7.42919 | Nipsnap1 | 18082 | 0.97823 | 1.02176761 | 1.0445 | 0.33642 | 0.782483 |
| 149408132 | serine/threonine-protein kinase NLK [Mus musculus] | 0 | 88.623 | 30 | 16 | 78 | 16 | 527 | 58.3 | 8.13 | 974 | 16 | 241145645.4 | 212219529.5 | 223622837.4 | 226856221.7 | 216414658.8 | 243388434.2 | 232027553.5 | 214178474.3 | 5.2716 | 6.08191 | Nlk | 18099 | 0.9988 | 1.001196167 | 1.0024 | 0.95453 | 0.994906 |
| 26335275 | unnamed protein product [Mus musculus] | 0 | 18.53 | 25 | 4 | 10 | 4 | 207 | 23.3 | 6.3 | 233 | 4 | 8670544.309 | 13883309.33 | 7450597.931 | 2369058.231 | 5533908.895 | 5770407.529 | 3026448 | 369367.0907 | 58.4075 | 68.8275 | Emc8 | 18117 | 1.37544 | 0.62455893 | 0.45408 | 0.1504 | 0.693437 |
| 568936908 | PREDICTED: nitric oxide synthase, brain isoform X1 [Mus musculus] | 0 | 11 | 3 | 4 | 12 | 4 | 1464 | 164.4 | 7.31 | 116 | 4 | 3880593.734 | 1860756.714 | 1334069.525 | 4614247.221 | 1740935.423 | 3871784.505 | 1963089.563 | 3185597.435 | 53.852 | 37.6093 | Nos1 | 18125 | 1.04135 | 0.958654082 | 0.92059 | 0.81235 | 0.961921 |
| 148680456 | mCG125894, isoform CRA_a [Mus musculus] | 0 | 141.278 | 45 | 9 | 128 | 5 | 292 | 32.6 | 4.77 | 2563 | 9 | 83491457.61 | 36584436.34 | 85151378.6 | 45196849.44 | 114241158.7 | 56705090.44 | 95147540.13 | 28144575.72 | 40.458 | 52.4612 | Npm1 | 18148 | 0.91956 | 1.080442924 | 1.17496 | 0.65186 | 0.915864 |
| 112363090 | catenin delta-2 [Mus musculus] | 0 | 645.061 | 64 | 61 | 755 | 57 | 1246 | 134.9 | 7.65 | 13278 | 61 | 832079453.7 | 842170397.6 | 824106252.2 | 895337653.3 | 875870377.1 | 836955705 | 875269560.7 | 754074065.2 | 3.78793 | 6.85567 | Ctnnd2 | 18163 | 1.00765 | 0.992350788 | 0.98482 | 0.70844 | 0.935089 |
| 6679120 | neuronal pentraxin-1 precursor [Mus musculus] | 0.001 | 6.194 | 5 | 2 | 9 | 2 | 432 | 47.1 | 6.44 | 79 | 2 | 2933110.801 | 2649483.163 | 2265321.47 | 3412124.741 | 2448539.558 | 3152200.533 | 2300675 | 3884261.439 | 17.1603 | 24.6832 | Nptx1 | 18164 | 0.97719 | 1.022808419 | 1.04668 | 0.77356 | 0.952792 |
| 28394197 | mKIAA0578 protein [Mus musculus] | 0 | 317.234 | 33 | 40 | 250 | 33 | 1553 | 170.2 | 6.43 | 5012 | 40 | 336571132.9 | 392269826.2 | 384717932.9 | 426207488.5 | 363482590.1 | 497134468.6 | 398603467.2 | 459162678.3 | 9.59935 | 13.9454 | Nrxn1 | 18189 | 0.94518 | 1.054821554 | 1.116 | 0.25151 | 0.733806 |
| 327365334 | neurexin-2-beta isoform 1 precursor [Mus musculus] | 0 | 69.727 | 9 | 15 | 60 | 10 | 1703 | 183.9 | 5.82 | 546 | 15 | 17679281.21 | 22997954.33 | 19055452.09 | 26495352.11 | 21041474.56 | 19897746.7 | 19870555.06 | 15929405.16 | 18.5086 | 11.6645 | Nrxn2 | 18190 | 1.05823 | 0.941774437 | 0.88996 | 0.33966 | 0.78329 |
| 568978942 | PREDICTED: neurexin 3 isoform X1 [Mus musculus] | 0 | 182.803 | 25 | 30 | 158 | 26 | 1696 | 186.4 | 5.74 | 2342 | 30 | 144231303.1 | 135644933.9 | 159884096.9 | 158281581.2 | 140330268.7 | 149447052.9 | 161640579.5 | 153859031.8 | 7.76813 | 5.87682 | Nrxn3 | 18191 | 0.99399 | 1.006012553 | 1.0121 | 0.81292 | 0.961921 |
| 31543349 | vesicle-fusing ATPase [Mus musculus] | 0 | 231.361 | 62 | 35 | 218 | 35 | 744 | 82.6 | 6.95 | 2740 | 35 | 387103512.4 | 404528338.2 | 433638796.3 | 372880784 | 419917193.7 | 393910429.9 | 403793963.7 | 338010195.7 | 6.54749 | 9.14986 | Nsf | 18195 | 1.01348 | 0.986517891 | 0.97339 | 0.64731 | 0.915705 |
| 68215970 | BDNF/NT-3 growth factors receptor isoform a precursor [Mus musculus] | 0 | 31.227 | 13 | 8 | 24 | 4 | 821 | 92.1 | 6.55 | 104 | 8 | 12023631.34 | 7120871.613 | 6433955.797 | 6840723.9 | 6696933.5 | 6303813.985 | 4651516.25 | 3288636.282 | 32.422 | 30.0169 | Ntrk2 | 18212 | 1.21511 | 0.784890089 | 0.64594 | 0.11 | 0.678142 |
| 9506939 | NT-3 growth factor receptor isoform 3 precursor [Rattus norvegicus] | 0 | 55.812 | 22 | 13 | 47 | 3 | 825 | 92.7 | 6.67 | 523 | 13 | 187181509.1 | 252530610.2 | 79823776.67 | 282340360.1 | 143503345.5 | 111809090.3 | 140631579.1 | 96413770.36 | 44.7514 | 18.5455 | Ntrk3 | 18213 | 1.23915 | 0.760848146 | 0.61401 | 0.1456 | 0.693437 |
| 568945015 | PREDICTED: numb-like protein isoform X2 [Mus musculus] | 0 | 34.132 | 19 | 9 | 30 | 9 | 608 | 64.7 | 8.57 | 181 | 9 | 27880702.45 | 109380475.4 | 80966833.34 | 26128156.11 | 35093851.84 | 27674910.63 | 31036907.63 | 33113743.94 | 67.1766 | 9.99186 | Numbl | 18223 | 1.31631 | 0.683693841 | 0.5194 | 0.2036 | 0.715071 |
| 16716349 | nuclear pore glycoprotein p62 [Mus musculus] | 0 | 55.088 | 23 | 8 | 45 | 8 | 526 | 53.2 | 5.31 | 614 | 8 | 43092048.12 | 36196928.57 | 38009185.27 | 32420653.09 | 48421312.28 | 41254044.99 | 44096974.63 | 41223607.42 | 11.8489 | 7.75748 | Nup62 | 18226 | 0.92216 | 1.077844089 | 1.16883 | 0.06427 | 0.601634 |
| 148681975 | neurexophilin 1, isoform CRA_a, partial [Mus musculus] | 0.001 | 4.463 | 7 | 2 | 2 | 2 | 305 | 35 | 9.04 | 0 | 2 | 2382448.03 | 2399872.63 | 3086096.198 | 2453181.634 | 1928745.793 | 1640754.438 | 1383280.375 | 1451908.905 | 13.117 | 15.2397 | Nxph1 | 18231 | 1.23418 | 0.765823177 | 0.62051 | 0.00335 | 0.38143 |
| 1039759427 | PREDICTED: outer dense fiber protein 2 isoform X1 [Mus musculus] | 0 | 60.955 | 19 | 12 | 30 | 12 | 875 | 100.9 | 8.44 | 509 | 12 | 24819131.15 | 13483641.92 | 29982595.29 | 21479788.8 | 26522150.55 | 9484242.156 | 19618949.63 | 16371774.54 | 30.8393 | 39.3399 | Odf2 | 18286 | 1.10984 | 0.890159554 | 0.80206 | 0.40414 | 0.828732 |
| 60360580 | mKIAA4192 protein [Mus musculus] | 0 | 22.745 | 7 | 7 | 20 | 7 | 1066 | 120.9 | 6.84 | 109 | 7 | 4138365.066 | 4014787.002 | 3853076.684 | 3874036.861 | 5008646.023 | 4573302.743 | 4682791.313 | 3853493.302 | 3.35529 | 10.7552 | Ogdh | 18293 | 0.93417 | 1.065825487 | 1.14093 | 0.06861 | 0.601634 |
| 226874869 | oligodendrocyte-myelin glycoprotein precursor [Mus musculus] | 0 | 34.552 | 14 | 5 | 27 | 5 | 443 | 49.6 | 8.41 | 691 | 5 | 162728295.8 | 127864814.9 | 176075478.9 | 137563749.1 | 145928796.2 | 116890009.1 | 146090442 | 101643112.2 | 14.7141 | 17.3209 | Omg | 18377 | 1.08403 | 0.915965855 | 0.84496 | 0.18576 | 0.715071 |
| 19070182 | myb-binding protein 1a [Mus musculus] | 0 | 37.684 | 8 | 7 | 13 | 7 | 1344 | 151.9 | 8.92 | 448 | 7 | 2467558.313 | 4898203.504 | 3437509.591 | 1408845.857 | 5193174.127 | 4476144.263 | 5621320.188 | 3862279.93 | 48.5774 | 16.2345 | Mybbp1a | 18432 | 0.77871 | 1.221291035 | 1.56835 | 0.08359 | 0.644421 |
| 26354649 | unnamed protein product [Mus musculus] | 0 | 85.721 | 33 | 19 | 100 | 16 | 636 | 70.6 | 9.55 | 781 | 19 | 130539622.1 | 137616082.3 | 148709553.1 | 161058228.6 | 105069904.4 | 140624814.4 | 175796043.9 | 152514655.8 | 9.23571 | 20.5521 | Pabpc1 | 18458 | 1.0034 | 0.99659869 | 0.99322 | 0.95371 | 0.994906 |
| 7305363 | platelet-activating factor acetylhydrolase IB subunit alpha [Mus musculus] | 0 | 64.164 | 40 | 12 | 71 | 12 | 410 | 46.6 | 7.37 | 477 | 12 | 41617137.74 | 47154926.62 | 48303708.92 | 41204182.29 | 54380742.63 | 46634890.95 | 44373486.31 | 66661228.58 | 8.26108 | 18.9721 | Pafah1b1 | 18472 | 0.91348 | 1.086517477 | 1.18942 | 0.16598 | 0.711253 |
| 12846314 | unnamed protein product [Mus musculus] | 0 | 43.563 | 61 | 9 | 75 | 9 | 199 | 22.2 | 8.12 | 784 | 9 | 208986496.6 | 353013359.1 | 291724803.9 | 306512682.1 | 211930735.2 | 309375907.9 | 294285203.8 | 308244930 | 20.6945 | 16.5604 | Prdx1 | 18477 | 1.01594 | 0.984063317 | 0.96863 | 0.81858 | 0.964114 |
| 239985639 | paralemmin-1 isoform 1 [Mus musculus] | 0 | 48.54 | 24 | 6 | 32 | 6 | 383 | 41.6 | 4.84 | 1634 | 6 | 29135634.05 | 37427333.75 | 40863625.39 | 26440178.78 | 35366218.75 | 30825491.84 | 31045546 | 40745560.09 | 20.3049 | 13.5137 | Palm | 18483 | 0.98486 | 1.01514089 | 1.03075 | 0.81112 | 0.961921 |
| 589938267 | PREDICTED: poly(rC)-binding protein 2 isoform X1 [Peromyscus maniculatus bairdii] | 0 | 65.027 | 33 | 8 | 138 | 6 | 471 | 49.1 | 7.64 | 3197 | 8 | 23199971.55 | 27149922.7 | 32397078.22 | 38225564.52 | 28365415.84 | 58762177.33 | 42418784.75 | 29429164.69 | 21.5587 | 35.7227 | Pcbp2 | 18521 | 0.86425 | 1.135750192 | 1.31415 | 0.26956 | 0.745606 |
| 28972738 | mKIAA1400 protein [Mus musculus] | 0 | 46.546 | 10 | 9 | 37 | 9 | 1098 | 119.6 | 4.94 | 592 | 9 | 29972564.63 | 24093017.9 | 26200955.4 | 24611375.99 | 19643607.87 | 27044126.99 | 20437020.75 | 14212397.53 | 10.1372 | 25.8652 | Pcdh10 | 18526 | 1.12642 | 0.873582943 | 0.77554 | 0.09276 | 0.662454 |
| 148703569 | pericentriolar material 1, isoform CRA_a [Mus musculus] | 0 | 52.479 | 8 | 12 | 26 | 12 | 2064 | 232.7 | 4.97 | 252 | 12 | 13125826.38 | 10971962.41 | 14005825.26 | 13461173.69 | 15842138.51 | 18398476.54 | 12744385.38 | 16836912.76 | 10.3162 | 14.9498 | Pcm1 | 18536 | 0.89377 | 1.1062265 | 1.2377 | 0.06598 | 0.601634 |
| 755534236 | PREDICTED: pericentrin isoform X4 [Mus musculus] | 0 | 141.107 | 14 | 32 | 99 | 32 | 2994 | 340.4 | 5.45 | 963 | 32 | 71167893.78 | 100434142.2 | 147224222.7 | 114375869.1 | 136760171.9 | 119575826.7 | 121343040.7 | 163977831.1 | 29.1634 | 15.1735 | Pcnt | 18541 | 0.88875 | 1.111251722 | 1.25036 | 0.20016 | 0.715071 |
| 6679233 | cyclin-dependent kinase 18 [Mus musculus] | 0 | 45.843 | 30 | 11 | 36 | 9 | 451 | 51.8 | 8.51 | 357 | 11 | 30465963.91 | 22593722.4 | 34975295.89 | 14353333.9 | 31240040.57 | 24704435.56 | 34102224.5 | 27459963.53 | 35.4556 | 14.079 | Cdk18 | 18557 | 0.93123 | 1.068769523 | 1.1477 | 0.47709 | 0.863587 |
| 26346917 | unnamed protein product [Mus musculus] | 0 | 11.054 | 13 | 2 | 5 | 2 | 191 | 21.8 | 5.4 | 124 | 2 | 14920151.14 | 8452958.888 | 8798985.178 | 8555373.873 | 12995106.26 | 7839443.44 | 8672529 | 9108118.706 | 31.0571 | 23.7096 | Pdcd6 | 18570 | 1.02662 | 0.973377859 | 0.94814 | 0.7958 | 0.958226 |
| 6679243 | calcium/calmodulin-dependent 3',5'-cyclic nucleotide phosphodiesterase 1B isoform 1 [Mus musculus] | 0.001 | 6.112 | 3 | 2 | 8 | 2 | 535 | 61.2 | 5.72 | 25 | 2 | 3085744.49 | 1409826.047 | 3172172.246 | 1946656.409 | 3897684.735 | 3942468.529 | 3696116.875 | 3249811.384 | 36.0496 | 8.56196 | Pde1b | 18574 | 0.78805 | 1.21195018 | 1.53791 | 0.03104 | 0.515682 |
| 41282015 | cAMP-specific 3',5'-cyclic phosphodiesterase 4A isoform 5 [Mus musculus] | 0 | 69.17 | 21 | 13 | 53 | 5 | 844 | 93.5 | 5.26 | 574 | 13 | 9763765.66 | 7234844.93 | 11785580.05 | 6115293.147 | 13210035.76 | 7717106.916 | 6192168.25 | 8624355.692 | 29.2042 | 33.8071 | Pde4a | 18577 | 0.98805 | 1.01194996 | 1.02419 | 0.91843 | 0.987994 |
| 112181198 | cAMP-specific 3',5'-cyclic phosphodiesterase 4B isoform 1 [Mus musculus] | 0 | 149.794 | 45 | 23 | 147 | 2 | 721 | 82 | 5.63 | 1825 | 23 | 3841487.247 | 5020255.458 | 7418527.398 | 3982242.724 | 9434924.124 | 8796028.702 | 7505058 | 9391145.794 | 32.658 | 10.2446 | Pde4b | 18578 | 0.73164 | 1.268364912 | 1.7336 | 0.00757 | 0.4058 |
| 6679245 | retinal rod rhodopsin-sensitive cGMP 3',5'-cyclic phosphodiesterase subunit delta [Mus musculus] | 0 | 16.018 | 25 | 3 | 16 | 3 | 150 | 17.3 | 5.67 | 260 | 3 | 12960361.13 | 11625178.69 | 12004145.08 | 11314613.58 | 14577757.47 | 10999998.33 | 12544998.5 | 11165915.37 | 5.9636 | 13.4354 | Pde6d | 18582 | 0.98576 | 1.014243532 | 1.0289 | 0.71429 | 0.936732 |
| 32816620 | platelet-derived growth factor alpha [Mus musculus] | 0.001 | 5.86 | 9 | 2 | 8 | 2 | 211 | 24.1 | 9.23 | 15 | 2 | 1989149.65 | 3132405.671 | 3570809.024 | 4365284.317 | 3739821.915 | 4963182.576 | 4479005.5 | 5158415.457 | 30.3746 | 13.7787 | Pdgfa | 18590 | 0.83175 | 1.168251618 | 1.40457 | 0.06575 | 0.601634 |
| 568930759 | PREDICTED: protein-arginine deiminase type-2 isoform X1 [Mus musculus] | 0 | 19.155 | 9 | 5 | 11 | 5 | 695 | 78.7 | 5.68 | 89 | 5 | 10248833.86 | 4169338.458 | 8143865.676 | 3069298.491 | 12939212.83 | 1043617.289 | 6552512.875 | 3214630.625 | 52.4783 | 87.3879 | Padi2 | 18600 | 1.0381 | 0.961903137 | 0.9266 | 0.88406 | 0.983141 |
| 241666481 | peroxisomal membrane protein 11B isoform 1 [Mus musculus] | 0 | 25.135 | 28 | 5 | 21 | 5 | 259 | 28.7 | 9.95 | 212 | 5 | 14418280.03 | 6769262.01 | 12883585.99 | 1811996.696 | 12096053.33 | 2738484.894 | 13793038.5 | 2220882.172 | 64.7056 | 78.9038 | Pex11b | 18632 | 1.07545 | 0.924553479 | 0.85969 | 0.77478 | 0.952792 |
| 568991691 | PREDICTED: ATP-dependent 6-phosphofructokinase, muscle type isoform X1 [Mus musculus] | 0 | 247.368 | 46 | 30 | 190 | 25 | 850 | 93.2 | 7.25 | 3393 | 30 | 261933685 | 241048764.1 | 298388674.7 | 249517883.8 | 262056555.3 | 289955412.3 | 304090381.3 | 287804501 | 9.62126 | 6.12171 | Pfkm | 18642 | 0.95762 | 1.042381091 | 1.08851 | 0.18114 | 0.715071 |
| 26389590 | unnamed protein product [Mus musculus] | 0 | 15.996 | 36 | 4 | 12 | 4 | 140 | 14.9 | 8.28 | 69 | 4 | 44479878.22 | 75368196.43 | 84669829.94 | 44123266.56 | 89116871.97 | 123733501.9 | 100329515 | 91691385.99 | 33.7333 | 15.5683 | Pfn1 | 18643 | 0.76094 | 1.23906217 | 1.62834 | 0.0247 | 0.515682 |
| 163310986 | Chain A, Mouse Profilin Iia In Complex With A Double Repeat From The Fh1 Domain Of Mdia1 | 0 | 23.278 | 44 | 4 | 50 | 4 | 140 | 15 | 7.02 | 1809 | 4 | 41346902.04 | 41051614.92 | 39210267.32 | 43009852.06 | 41268321.41 | 49794259.39 | 34669660.5 | 36277637.38 | 3.78257 | 16.7936 | Pfn2 | 18645 | 1.00799 | 0.992013074 | 0.98415 | 0.85787 | 0.975398 |
| 114326546 | phosphoglycerate mutase 1 [Mus musculus] | 0 | 66.939 | 53 | 10 | 37 | 10 | 254 | 28.8 | 7.18 | 727 | 10 | 27495573.74 | 18874295.31 | 26605971.95 | 22092569.59 | 29928001.43 | 22517972.08 | 32658946.63 | 15945361.39 | 16.953 | 29.8694 | Pgam1 | 18648 | 0.9695 | 1.030501279 | 1.06292 | 0.73855 | 0.943908 |
| 202423 | phosphoglycerate kinase [Mus musculus] | 0 | 47.419 | 25 | 7 | 33 | 7 | 417 | 44.5 | 7.65 | 726 | 7 | 24966558.86 | 18669615.59 | 30280616.28 | 17772760.25 | 23464482.66 | 19292420.77 | 29143333.75 | 20838830.69 | 25.5531 | 18.6744 | Pgk1 | 18655 | 0.99431 | 1.00569064 | 1.01145 | 0.94491 | 0.994906 |
| 148684042 | mCG8461, isoform CRA_c [Mus musculus] | 0 | 103.699 | 66 | 14 | 99 | 14 | 274 | 30 | 5.76 | 1219 | 14 | 169194245.9 | 104009503.8 | 88816165.62 | 89829443.7 | 180507712.3 | 217536377.1 | 111848140.4 | 161318425.4 | 33.7494 | 26.2209 | Phb | 18673 | 0.80468 | 1.195324643 | 1.48547 | 0.10856 | 0.678142 |
| 148689581 | mCG10343, isoform CRA_c, partial [Mus musculus] | 0 | 123.92 | 43 | 18 | 349 | 18 | 375 | 41.5 | 9.13 | 5916 | 18 | 1900330184 | 1001607344 | 666188058.2 | 832300652.2 | 1125882044 | 2077977068 | 669086693.6 | 1248951190 | 50.0658 | 45.8644 | Slc25a3 | 18674 | 0.92423 | 1.075766254 | 1.16395 | 0.66985 | 0.922517 |
| 26354981 | unnamed protein product [Mus musculus] | 0 | 12.156 | 5 | 2 | 10 | 2 | 530 | 58.9 | 6.87 | 309 | 2 | 3579394.464 | 1874339.422 | 4154338.814 | 1854723.259 | 4477986.203 | 3557404.164 | 3927118.125 | 3815420.353 | 41.165 | 9.83416 | Pick1 | 18693 | 0.84159 | 1.158407418 | 1.37645 | 0.13298 | 0.693437 |
| 39104624 | mKIAA0589 protein [Mus musculus] | 0 | 35.382 | 13 | 7 | 16 | 5 | 705 | 77.2 | 6.24 | 234 | 7 | 16654374.43 | 20963127.53 | 14088712.66 | 11151434.32 | 5550577.547 | 3626771.69 | 5501076.375 | 5241735.436 | 26.4673 | 18.3189 | Pip5k1c | 18717 | 1.51871 | 0.481292294 | 0.31691 | 0.00235 | 0.38143 |
| 125490368 | phosphatidylinositol 5-phosphate 4-kinase type-2 alpha [Mus musculus] | 0 | 35.942 | 24 | 8 | 26 | 4 | 405 | 46.1 | 6.99 | 133 | 8 | 7333904.876 | 3735772.396 | 10313889.16 | 5389251.679 | 9253404.656 | 4425652.041 | 10581620.75 | 1301906.856 | 42.2287 | 67.3138 | Pip4k2a | 18718 | 1.02312 | 0.976875428 | 0.9548 | 0.91025 | 0.98561 |
| 6679339 | membrane-associated phosphatidylinositol transfer protein 1 [Mus musculus] | 0.001 | 6.366 | 2 | 2 | 4 | 2 | 1243 | 134.9 | 6.06 | 25 | 2 | 3054645.847 | 1051723.173 | 1292583.688 | 1641439.147 | 3041689.476 | 3431764.622 | 3138690.5 | 2348892.851 | 50.9258 | 15.3366 | Pitpnm1 | 18739 | 0.74104 | 1.258961866 | 1.69892 | 0.05023 | 0.592616 |
| 594542497 | E3 ubiquitin-protein ligase Praja-1 isoform 3 [Mus musculus] | 0 | 37.944 | 16 | 6 | 35 | 6 | 615 | 67.8 | 5.19 | 511 | 6 | 15196414.2 | 17170339.3 | 20437488.31 | 13573973.53 | 19247481.63 | 19554775.39 | 17771127.38 | 18578558.67 | 17.801 | 4.21005 | Pja1 | 18744 | 0.93801 | 1.06199193 | 1.13218 | 0.20142 | 0.715071 |
| 359807367 | pyruvate kinase PKM isoform M1 [Mus musculus] | 0 | 75.014 | 34 | 15 | 61 | 15 | 531 | 57.9 | 7.14 | 1340 | 15 | 70109495.95 | 49873706.5 | 85677024.66 | 70512286.61 | 92680489.24 | 80760392.44 | 68247813.13 | 121045723.9 | 21.2776 | 24.8844 | Pkm | 18746 | 0.86452 | 1.135484373 | 1.31343 | 0.1591 | 0.700858 |
| 255958156 | cAMP-dependent protein kinase catalytic subunit beta isoform 4 [Mus musculus] | 0 | 84.957 | 39 | 13 | 85 | 5 | 398 | 46 | 8.56 | 579 | 13 | 11096634.33 | 3824677.766 | 12518612.44 | 5413535.114 | 15484625.75 | 9146578.57 | 17670541.25 | 3408230.802 | 51.6305 | 56.4744 | Prkacb | 18749 | 0.83635 | 1.163645041 | 1.39133 | 0.43703 | 0.852048 |
| 164663791 | protein kinase C alpha type [Mus musculus] | 0 | 97.023 | 31 | 16 | 61 | 6 | 672 | 76.8 | 7.05 | 1208 | 16 | 15162616.57 | 10583129.35 | 16535365.55 | 9873299.822 | 12484953.21 | 4362111.312 | 11886967 | 8569331.478 | 25.3548 | 40.0019 | Prkca | 18750 | 1.16601 | 0.833988176 | 0.71525 | 0.18688 | 0.715071 |
| 6679345 | protein kinase C beta type isoform 1 [Mus musculus] | 0 | 136.644 | 52 | 24 | 86 | 15 | 673 | 76.8 | 7.02 | 1506 | 24 | 75027313.73 | 69993247.09 | 62212189.95 | 52566676.45 | 66087711.03 | 58877350.59 | 61802825.59 | 48284332.8 | 15.0809 | 12.9119 | Prkcb | 18751 | 1.05001 | 0.949990654 | 0.90474 | 0.35651 | 0.798662 |
| 6755080 | protein kinase C gamma type isoform 1 [Mus musculus] | 0 | 385.496 | 76 | 39 | 467 | 37 | 697 | 78.3 | 7.46 | 10295 | 39 | 1273295041 | 1380923598 | 1110901015 | 1092254181 | 1150144503 | 1206927954 | 1003682274 | 1173148928 | 11.3341 | 7.90635 | Prkcg | 18752 | 1.03444 | 0.965556318 | 0.93341 | 0.36277 | 0.804292 |
| 893548269 | protein kinase C delta type isoform 1 [Mus musculus] | 0 | 7.256 | 3 | 3 | 14 | 2 | 700 | 80.2 | 7.39 | 48 | 3 | 3333863.492 | 2627506.066 | 2353510.78 | 2360158.558 | 2679044.624 | 1748193.607 | 1814710.625 | 2573115.84 | 17.2891 | 22.2488 | Prkcd | 18753 | 1.09543 | 0.904568322 | 0.82576 | 0.21643 | 0.716066 |
| 6755084 | protein kinase C epsilon type [Mus musculus] | 0 | 21.853 | 8 | 6 | 19 | 5 | 737 | 83.5 | 7.03 | 122 | 6 | 2830677.252 | 1989400.434 | 3101767.727 | 2011758.768 | 3842331.965 | 3005976.445 | 2948170.5 | 2574163.758 | 22.8906 | 17.3047 | Prkce | 18754 | 0.89074 | 1.109263429 | 1.24533 | 0.16961 | 0.711915 |
| 15030028 | Plasminogen activator, tissue [Mus musculus] | 0 | 17.173 | 7 | 3 | 8 | 3 | 559 | 63.1 | 8.15 | 195 | 3 | 1225202.005 | 7842339.733 | 4919466.42 | 3304618.669 | 3585972.347 | 1297396.042 | 7283488.5 | 443223.6571 | 64.5712 | 96.968 | Plat | 18791 | 1.15656 | 0.843435486 | 0.72926 | 0.59226 | 0.902243 |
| 568915825 | PREDICTED: 1-phosphatidylinositol 4,5-bisphosphate phosphodiesterase beta-4 isoform X1 [Mus musculus] | 0 | 21.712 | 7 | 8 | 23 | 8 | 1207 | 137.5 | 6.73 | 89 | 8 | 7929103.604 | 6584130.847 | 11334309.67 | 6311612.505 | 8973964.594 | 10695090.37 | 10154879.38 | 9758531.184 | 28.6992 | 7.32121 | Plcb4 | 18798 | 0.89653 | 1.103472833 | 1.23083 | 0.17575 | 0.713031 |
| 7242181 | phospholipase D3 [Mus musculus] | 0 | 19.448 | 11 | 5 | 15 | 5 | 488 | 54.4 | 6.52 | 153 | 5 | 17381986.18 | 30129421.39 | 15901401.61 | 23119616.15 | 26730647.4 | 29299978.11 | 25089299 | 26409892.32 | 29.8768 | 6.55174 | Pld3 | 18807 | 0.8918 | 1.108199263 | 1.24265 | 0.16811 | 0.711915 |
| 254675117 | plectin isoform 1c2alpha3alpha [Mus musculus] | 0 | 2382.92 | 65 | 290 | 4912 | 289 | 4589 | 521.6 | 5.8 | 83020 | 290 | 10676517623 | 10352589941 | 11392099542 | 10388718786 | 10951161055 | 8336725786 | 11500648567 | 10792504961 | 4.5041 | 13.5206 | Plec | 18810 | 1.01456 | 0.985438186 | 0.97129 | 0.6936 | 0.930425 |
| 13591880 | myelin proteolipid protein [Rattus norvegicus] | 0 | 44.603 | 36 | 7 | 202 | 7 | 277 | 30.1 | 8.35 | 3595 | 7 | 2045929948 | 2131116069 | 1317318340 | 1566763253 | 1595837864 | 1755690870 | 1661607587 | 1790131299 | 22.005 | 5.20885 | Plp1 | 18823 | 1.0186 | 0.981401281 | 0.96348 | 0.75722 | 0.952737 |
| 158508501 | septin-5 [Mus musculus] | 0 | 141.351 | 56 | 14 | 117 | 12 | 369 | 42.7 | 6.67 | 1713 | 14 | 362677170.9 | 351401955.1 | 344959338.9 | 317756370.6 | 309186762 | 334197730.6 | 322396157.6 | 377969892.2 | 5.54579 | 8.87835 | 43713 | 18951 | 1.01214 | 0.987855033 | 0.976 | 0.65729 | 0.918974 |
| 568972207 | PREDICTED: septin-4 isoform X1 [Mus musculus] | 0 | 51.533 | 12 | 8 | 34 | 6 | 971 | 110 | 8.35 | 700 | 8 | 14230479.11 | 19248713.6 | 17329694.16 | 14289253.79 | 16995700.38 | 19710141.73 | 16449268.38 | 26853319.99 | 15.0839 | 23.9219 | 43712 | 18952 | 0.89725 | 1.102754063 | 1.22904 | 0.21498 | 0.716066 |
| 21729749 | DNA polymerase beta [Mus musculus] | 0.003 | 3.867 | 6 | 2 | 2 | 2 | 335 | 38.3 | 8.85 | 16 | 2 | 1270336.642 | 1786341.552 | 2065033.917 | 2696862.775 | 1852138.424 | 1390030.977 | 2397417.563 | 2509202.612 | 30.4059 | 25.4373 | Polb | 18970 | 0.97932 | 1.020680601 | 1.04223 | 0.84107 | 0.970469 |
| 71774133 | peptidyl-prolyl cis-trans isomerase B precursor [Mus musculus] | 0 | 57.862 | 56 | 13 | 64 | 13 | 216 | 23.7 | 9.55 | 1111 | 13 | 167283005.2 | 212911160.5 | 212903090.5 | 309878612.5 | 208089399.2 | 147213891.8 | 209854768.8 | 423949818.3 | 26.6106 | 49.0659 | Ppib | 19035 | 0.95448 | 1.045522303 | 1.09539 | 0.76119 | 0.952792 |
| 13994195 | serine/threonine-protein phosphatase PP1-alpha catalytic subunit [Mus musculus] | 0 | 226.701 | 77 | 24 | 313 | 7 | 330 | 37.5 | 6.33 | 4590 | 24 | 283618753 | 274906560.6 | 238654724.4 | 383847103.4 | 305489890 | 324063539.5 | 295658787.8 | 295524877.5 | 21.062 | 4.39844 | Ppp1ca | 19045 | 0.98347 | 1.01653366 | 1.03362 | 0.76553 | 0.952792 |
| 4506005 | serine/threonine-protein phosphatase PP1-beta catalytic subunit isoform 1 [Homo sapiens] | 0 | 237.547 | 73 | 22 | 180 | 9 | 327 | 37.2 | 6.19 | 3698 | 22 | 118700395.1 | 102657561.5 | 101851995.6 | 121635220.8 | 106742684.2 | 132643968.2 | 104585398.4 | 133381037.4 | 9.36637 | 13.254 | Ppp1cb | 19046 | 0.96475 | 1.035250462 | 1.07308 | 0.42372 | 0.840684 |
| 401709959 | serine/threonine-protein phosphatase PP1-gamma catalytic subunit isoform 2 [Rattus norvegicus] | 0 | 187.058 | 57 | 21 | 251 | 3 | 337 | 38.5 | 6.13 | 5165 | 21 | 102924646.2 | 104694972.5 | 124017248.5 | 109140240 | 122431862.7 | 115479695.5 | 131936910.6 | 105911381.7 | 8.69289 | 9.24622 | Ppp1cc | 19047 | 0.96183 | 1.038168394 | 1.07937 | 0.27561 | 0.745606 |
| 8394030 | serine/threonine-protein phosphatase 2B catalytic subunit alpha isoform [Rattus norvegicus] | 0 | 180.562 | 66 | 23 | 239 | 13 | 521 | 58.6 | 5.86 | 3415 | 23 | 105058259.8 | 141296674.3 | 126804018.6 | 157014835 | 128387567 | 152004030.7 | 114898409.3 | 132217853 | 16.6659 | 11.6297 | Ppp3ca | 19055 | 1.00252 | 0.997479461 | 0.99497 | 0.96208 | 0.99605 |
| 568987032 | PREDICTED: serine/threonine-protein phosphatase 2B catalytic subunit beta isoform isoform X1 [Mus musculus] | 0 | 175.711 | 60 | 21 | 157 | 11 | 534 | 60.2 | 6.1 | 3135 | 21 | 83442700.46 | 105043938.1 | 112275881.5 | 111589068.9 | 126184974 | 120984587.7 | 92790611.13 | 110546019.6 | 13.092 | 13.0843 | Ppp3cb | 19056 | 0.95578 | 1.044218879 | 1.09253 | 0.3766 | 0.807893 |
| 28436800 | Ppp3cc protein, partial [Mus musculus] | 0 | 38.773 | 16 | 6 | 41 | 2 | 538 | 61.4 | 7.39 | 911 | 6 | 2865835.775 | 3550125.003 | 3652190.152 | 954653.6026 | 3686469.555 | 5611105.854 | 4415968.625 | 4079351.981 | 45.376 | 18.6725 | Ppp3cc | 19057 | 0.76506 | 1.234944539 | 1.61419 | 0.065 | 0.601634 |
| 255079 | calmodulin-dependent protein phosphatase regulatory subunit beta 1 isoform [Mus sp.] | 0 | 76.597 | 61 | 7 | 56 | 7 | 170 | 19.3 | 4.81 | 1917 | 7 | 118205202.1 | 160055362 | 178057362.7 | 131941848.7 | 192428328 | 152005837.3 | 152740344.5 | 135993682.8 | 18.3755 | 15.1831 | Ppp3r1 | 19058 | 0.96323 | 1.036767143 | 1.07634 | 0.55751 | 0.895256 |
| 148680682 | nucleoporin 88, isoform CRA_d, partial [Mus musculus] | 0 | 35.324 | 12 | 7 | 26 | 7 | 755 | 85.2 | 6.33 | 368 | 7 | 20175364.62 | 9593706.626 | 17803002.78 | 18281785.38 | 14778314.31 | 14890439.93 | 20885796.13 | 13133776.3 | 28.5055 | 21.3865 | Nup88 | 19069 | 1.01672 | 0.983283185 | 0.96712 | 0.85802 | 0.975398 |
| 12861034 | unnamed protein product [Mus musculus] | 0 | 51.403 | 45 | 7 | 52 | 7 | 225 | 26 | 5.78 | 951 | 7 | 36319083.2 | 58536552.41 | 49183844.42 | 51898460.01 | 52623780.9 | 67947719.17 | 47357001.13 | 65101514.14 | 19.0118 | 16.9145 | Mob4 | 19070 | 0.91353 | 1.086468173 | 1.18931 | 0.22037 | 0.718033 |
| 30794476 | cAMP-dependent protein kinase type I-alpha regulatory subunit [Mus musculus] | 0 | 83.456 | 42 | 17 | 102 | 14 | 381 | 43.2 | 5.35 | 1343 | 17 | 86872386.48 | 82164483.53 | 63511093.66 | 77461255 | 63194666.69 | 92586882.08 | 67322212.63 | 53872120.58 | 13.0661 | 23.8976 | Prkar1a | 19084 | 1.05628 | 0.943723722 | 0.89344 | 0.42678 | 0.84306 |
| 254675178 | cAMP-dependent protein kinase type I-beta regulatory subunit [Mus musculus] | 0 | 89.864 | 40 | 13 | 84 | 10 | 381 | 43.2 | 5.96 | 1338 | 13 | 98823283.1 | 81575686.02 | 96384722.42 | 75598368.72 | 80246381.53 | 90942744.24 | 73096684.06 | 60262912.78 | 12.8172 | 16.9108 | Prkar1b | 19085 | 1.07281 | 0.92718664 | 0.86426 | 0.21202 | 0.716066 |
| 22550094 | cAMP-dependent protein kinase type II-alpha regulatory subunit [Mus musculus] | 0 | 76.369 | 49 | 13 | 33 | 10 | 402 | 45.6 | 4.93 | 498 | 13 | 16996633.03 | 20110707.84 | 19456905.9 | 20258353.58 | 14352339.56 | 9773846.919 | 12941324.38 | 15379769.6 | 7.87943 | 18.6054 | Prkar2a | 19087 | 1.18856 | 0.811438521 | 0.68271 | 0.00541 | 0.38143 |
| 45598396 | cAMP-dependent protein kinase type II-beta regulatory subunit [Mus musculus] | 0 | 73.273 | 39 | 11 | 56 | 8 | 416 | 46.1 | 4.98 | 1463 | 11 | 53535299.8 | 30423936.52 | 45804106.28 | 36798613.73 | 36130732.61 | 29922875.87 | 33707320.38 | 31877498.37 | 24.3362 | 8.03903 | Prkar2b | 19088 | 1.11711 | 0.8828857 | 0.79033 | 0.14651 | 0.693437 |
| 568915859 | PREDICTED: C-Jun-amino-terminal kinase-interacting protein 1 isoform X1 [Mus musculus] | 0 | 11.631 | 6 | 4 | 9 | 4 | 711 | 77.2 | 5.01 | 14 | 4 | 5338754.681 | 7565980.007 | 5638536.554 | 6409197.53 | 7172775.445 | 5575627.001 | 4818766.875 | 6597228.561 | 15.9255 | 17.3631 | Mapk8ip1 | 19099 | 1.01604 | 0.983955188 | 0.96842 | 0.79419 | 0.956891 |
| 13173473 | major prion protein precursor [Mus musculus] | 0 | 12.94 | 12 | 3 | 35 | 3 | 254 | 28 | 9.33 | 726 | 3 | 23815520.55 | 35621839.44 | 16830442.41 | 22958070.72 | 22718398.96 | 26330350.1 | 19431615.5 | 23794512.57 | 31.6553 | 12.392 | Prnp | 19122 | 1.0363 | 0.963702513 | 0.92995 | 0.69195 | 0.930425 |
| 254675339 | peripherin isoform 3 [Mus musculus] | 0 | 199.558 | 62 | 30 | 439 | 27 | 475 | 54.3 | 5.52 | 6703 | 30 | 626797053.9 | 681988956.6 | 576729610.3 | 554333510.3 | 602882751.5 | 612548537 | 866862013.3 | 668299725 | 9.30796 | 17.8733 | Prph | 19132 | 0.94013 | 1.059868483 | 1.12736 | 0.29481 | 0.759552 |
| 12843957 | unnamed protein product [Mus musculus] | 0 | 13.789 | 14 | 4 | 10 | 3 | 318 | 34.8 | 6.98 | 50 | 4 | 852448.9585 | 615901.503 | 833534.8561 | 615681.493 | 867608.6755 | 665787.9239 | 754047.1875 | 599440.9422 | 18.0152 | 16.0809 | Prps1 | 19139 | 1.00529 | 0.994714043 | 0.98948 | 0.93312 | 0.994812 |
| 225735645 | prosaposin isoform B precursor [Mus musculus] | 0 | 11.864 | 8 | 4 | 7 | 4 | 557 | 61.4 | 5.19 | 16 | 4 | 17819456.21 | 14059993.14 | 31323296.69 | 11383829.67 | 29272261.94 | 35258027.79 | 24711552.75 | 22190807.98 | 47.4818 | 20.5976 | Psap | 19156 | 0.80192 | 1.198076702 | 1.494 | 0.13139 | 0.693437 |
| 10121123 | cytohesin-1 [Mus musculus] | 0 | 66.171 | 38 | 12 | 48 | 8 | 418 | 48.7 | 5.64 | 595 | 12 | 31160905.28 | 31636460.29 | 28852109.16 | 24583334.13 | 31288812.95 | 26232799.63 | 30173121.75 | 23365512.61 | 11.0866 | 13.1381 | Cyth1 | 19157 | 1.02276 | 0.977242763 | 0.9555 | 0.61422 | 0.905186 |
| 341940420 | binding site opener; Short=Protein ARNO; AltName: Full=PH, SEC7 and coiled-coil domain-containing protein 2; SI | 0 | 81.981 | 39 | 13 | 68 | 9 | 400 | 46.6 | 5.63 | 866 | 13 | 82049007.37 | 82372854.74 | 97604470.12 | 58840317.18 | 88910763.23 | 88845585 | 79330207 | 91409069.46 | 19.9368 | 6.11897 | Cyth2 | 19158 | 0.95872 | 1.041276565 | 1.08611 | 0.44384 | 0.852048 |
| 60360638 | mKIAA4241 protein [Mus musculus] | 0 | 62.607 | 34 | 11 | 32 | 8 | 453 | 51.9 | 7.4 | 424 | 11 | 36864575.42 | 37582072.74 | 44956526.52 | 32894684.22 | 39437282.94 | 38145420.71 | 42821455.25 | 32578028.58 | 13.2106 | 11.1434 | Cyth3 | 19159 | 0.99776 | 1.002241642 | 1.00449 | 0.96029 | 0.994906 |
| 6679501 | 26S protease regulatory subunit 4 [Mus musculus] | 0 | 9.844 | 8 | 3 | 10 | 2 | 440 | 49.2 | 6.21 | 38 | 3 | 1079616.503 | 1121209.509 | 1674203.792 | 3542679.469 | 2553969.942 | 2272371.609 | 764888 | 2294832.503 | 62.4275 | 41.3136 | Psmc1 | 19179 | 0.9694 | 1.030603749 | 1.06314 | 0.87403 | 0.979509 |
| 7110703 | 26S protease regulatory subunit 8 [Mus musculus] | 0 | 16.823 | 15 | 4 | 15 | 3 | 406 | 45.6 | 7.55 | 72 | 4 | 5231764.9 | 6345400.975 | 5606041.129 | 3048769.879 | 4729169.33 | 2411442.938 | 3965156.75 | 2307768.763 | 28.018 | 35.4837 | Psmc5 | 19184 | 1.20266 | 0.797344782 | 0.66299 | 0.115 | 0.678142 |
| 50510331 | mKIAA0024 protein [Mus musculus] | 0 | 12.284 | 7 | 2 | 9 | 2 | 487 | 57.1 | 8.82 | 133 | 2 | 4051252.176 | 7239372.783 | 6138968.974 | 6898422.952 | 7344354.499 | 9012585.738 | 6854231.75 | 6919687.717 | 23.5092 | 13.4111 | Ptdss1 | 19210 | 0.89345 | 1.106554582 | 1.23853 | 0.14854 | 0.693437 |
| 148704056 | PTK2 protein tyrosine kinase 2 beta, isoform CRA_c [Mus musculus] | 0 | 43.965 | 14 | 11 | 29 | 11 | 1024 | 117.4 | 6.49 | 230 | 11 | 12249965.36 | 11296789.04 | 13972240.12 | 16312716.65 | 17896659.32 | 16572956.09 | 12529640.63 | 14749645.43 | 16.3608 | 15.0836 | Ptk2b | 19229 | 0.9315 | 1.068499293 | 1.14707 | 0.26291 | 0.745606 |
| 15929276 | Twinfilin, actin-binding protein, homolog 1 (Drosophila) [Mus musculus] | 0 | 11.342 | 13 | 4 | 11 | 4 | 350 | 40.1 | 6.71 | 36 | 4 | 21064437.63 | 32297191.96 | 24342046.81 | 25215176.51 | 24462242.28 | 40559708.15 | 26006064.75 | 37241543.24 | 18.3791 | 25.0427 | Twf1 | 19230 | 0.89035 | 1.109653877 | 1.24632 | 0.22268 | 0.718033 |
| 1039766996 | PREDICTED: receptor-type tyrosine-protein phosphatase delta isoform X18 [Mus musculus] | 0 | 274.055 | 28 | 36 | 164 | 23 | 1919 | 215.1 | 6.6 | 3493 | 36 | 83582219.33 | 66040115.45 | 91790251.61 | 95162860.83 | 81406989.12 | 157640194.2 | 87766567.09 | 101080130.7 | 15.4641 | 32.4924 | Ptprd | 19266 | 0.88055 | 1.119453365 | 1.27132 | 0.26465 | 0.745606 |
| 568928624 | PREDICTED: receptor-type tyrosine-protein phosphatase F isoform X1 [Mus musculus] | 0 | 56.778 | 7 | 11 | 42 | 3 | 1924 | 214.1 | 6.58 | 790 | 11 | 5548191.61 | 5626599.141 | 2938436.906 | 5246443.756 | 4370198.658 | 4763214.244 | 4337815.25 | 5648288.664 | 26.4096 | 12.7697 | Ptprf | 19268 | 1.00624 | 0.993758845 | 0.9876 | 0.9352 | 0.994906 |
| 126632011 | Protein tyrosine phosphatase, receptor type, N polypeptide 2 [Mus musculus] | 0 | 8.986 | 2 | 2 | 3 | 2 | 1001 | 111.5 | 5.95 | 36 | 2 | 3469651.498 | 2305612.066 | 2810599.27 | 2671741.103 | 3455234.706 | 2747740.904 | 2719418.063 | 2028414.687 | 17.2683 | 21.2818 | Ptprn2 | 19276 | 1.01381 | 0.986185614 | 0.97275 | 0.84646 | 0.973079 |
| 568999464 | PREDICTED: receptor-type tyrosine-protein phosphatase S isoform X1 [Mus musculus] | 0 | 295.469 | 33 | 39 | 168 | 28 | 1920 | 213.4 | 6.9 | 3604 | 39 | 164517114.2 | 155714636.7 | 145825458.6 | 160034124.9 | 132776257.5 | 155751581.8 | 153655208.9 | 135528873 | 5.10208 | 8.2732 | Ptprs | 19280 | 1.04019 | 0.959811196 | 0.92273 | 0.14334 | 0.693437 |
| 568939473 | PREDICTED: receptor-type tyrosine-protein phosphatase zeta isoform X1 [Mus musculus] | 0 | 81.223 | 8 | 14 | 83 | 14 | 2318 | 255 | 4.88 | 1150 | 14 | 117063039.3 | 161354369.4 | 143830284.2 | 165860035.4 | 203604078.3 | 223824165.6 | 182686818.1 | 202362114.1 | 15.0455 | 8.2723 | Ptprz1 | 19283 | 0.8398 | 1.160196963 | 1.38151 | 0.00681 | 0.396316 |
| 6679573 | transcriptional activator protein Pur-alpha [Mus musculus] | 0 | 138.069 | 67 | 12 | 63 | 12 | 321 | 34.9 | 6.44 | 1842 | 12 | 147689652.3 | 112869587.3 | 140080623.5 | 156372092.9 | 139027588.5 | 153959553.7 | 147249050.6 | 145853309.9 | 13.505 | 4.17862 | Pura | 19290 | 0.97456 | 1.025437416 | 1.0522 | 0.49 | 0.870399 |
| 6755252 | transcriptional activator protein Pur-beta [Mus musculus] | 0 | 82.039 | 52 | 8 | 110 | 7 | 324 | 33.9 | 5.43 | 3862 | 8 | 41976240.94 | 49037077.08 | 50649312.27 | 55509401.23 | 37762289.91 | 62774007.47 | 47826143.94 | 55847142.3 | 11.3609 | 21.0791 | Purb | 19291 | 0.98247 | 1.017533319 | 1.03569 | 0.78154 | 0.952792 |
| 148680385 | ATP-binding cassette, sub-family D (ALD), member 3, isoform CRA_b, partial [Mus musculus] | 0 | 83.809 | 23 | 10 | 39 | 10 | 685 | 77.9 | 9.26 | 638 | 10 | 73636115.2 | 59827443.86 | 60337910.99 | 52595880.12 | 77978807.36 | 56153905.4 | 75341582.5 | 54962457.98 | 14.2353 | 18.5154 | Abcd3 | 19299 | 0.96469 | 1.035313623 | 1.07321 | 0.57106 | 0.895889 |
| 113930737 | peroxisomal biogenesis factor 5 isoform 1 [Mus musculus] | 0 | 87.561 | 41 | 16 | 61 | 16 | 639 | 70.7 | 4.55 | 1001 | 16 | 65833415.3 | 53418953.59 | 73513829.98 | 35399782.98 | 72162216.28 | 46905076.09 | 57784657.5 | 54279382.9 | 29.1621 | 18.3523 | Pex5 | 19305 | 0.99354 | 1.006456278 | 1.013 | 0.94253 | 0.994906 |
| 14249144 | ras-related protein Rab-11B [Rattus norvegicus] | 0 | 7.305 | 15 | 3 | 9 | 3 | 218 | 24.5 | 5.94 | 74 | 3 | 2528798.933 | 613124.3868 | 2141543.695 | 1803993.761 | 3828547.884 | 5185614.206 | 3851835.5 | 5946311.771 | 46.6915 | 22.1917 | Rab11b | 19326 | 0.5473 | 1.452700874 | 2.65431 | 0.00456 | 0.38143 |
| 6755260 | ras-related protein Rab-33A [Mus musculus] | 0.001 | 6.23 | 9 | 2 | 8 | 2 | 237 | 26.5 | 7.88 | 0 | 2 | 2811521.361 | 4176457.041 | 2231277.701 | 3199912.065 | 2482264.564 | 4789515.76 | 2636370.5 | 4023975.498 | 26.341 | 31.9617 | Rab33a | 19337 | 0.94259 | 1.057414946 | 1.12182 | 0.6037 | 0.905186 |
| 6679593 | ras-related protein Rab-3A [Mus musculus] | 0 | 36.467 | 35 | 6 | 42 | 5 | 220 | 25 | 5.03 | 604 | 6 | 20434424.15 | 26805311.13 | 22213655.51 | 25468587.89 | 31858005.68 | 31582884.14 | 27610786.88 | 26729992.71 | 12.3195 | 9.01277 | Rab3a | 19339 | 0.89253 | 1.10747202 | 1.24083 | 0.02752 | 0.515682 |
| 148687100 | RAS-related C3 botulinum substrate 1, isoform CRA_b, partial [Mus musculus] | 0 | 36.536 | 26 | 7 | 79 | 3 | 256 | 28.1 | 9.45 | 1036 | 7 | 42127392.46 | 43423047.62 | 41518894.17 | 48914959.51 | 46093718.28 | 49757362.4 | 41674789 | 44155328.25 | 7.66893 | 7.50855 | Rac1 | 19353 | 0.98407 | 1.015928023 | 1.03237 | 0.57437 | 0.895889 |
| 5453555 | GTP-binding nuclear protein Ran isoform 1 [Homo sapiens] | 0 | 35.284 | 38 | 6 | 33 | 6 | 216 | 24.4 | 7.49 | 333 | 6 | 51950289.04 | 50358067.92 | 57513420.45 | 49298659.08 | 63016485.73 | 87300408.98 | 67507488 | 56455196.16 | 6.99141 | 19.3761 | Ran | 19384 | 0.86521 | 1.134793422 | 1.31159 | 0.05595 | 0.599194 |
| 10442646 | Ran-binding protein 2 [Mus musculus] | 0 | 189.507 | 17 | 41 | 158 | 41 | 3053 | 340.9 | 6.18 | 2032 | 41 | 95908589.24 | 127619619.5 | 140370337.7 | 123065388.6 | 119250886 | 174211750.4 | 143686736.9 | 167126780 | 15.3722 | 16.4839 | Ranbp2 | 19386 | 0.8925 | 1.107503582 | 1.24091 | 0.1087 | 0.678142 |
| 148672614 | RAN GTPase activating protein 1, isoform CRA_b, partial [Mus musculus] | 0 | 76.804 | 31 | 16 | 45 | 16 | 680 | 72.8 | 4.74 | 543 | 16 | 68301697.31 | 90672189.84 | 34559605.54 | 39129833.07 | 98703222.39 | 61343385.52 | 46706874.28 | 125559841.5 | 45.2586 | 43.0873 | Rangap1 | 19387 | 0.82362 | 1.176378967 | 1.4283 | 0.30501 | 0.761857 |
| 31980729 | rasGAP-activating-like protein 1 [Mus musculus] | 0 | 77.922 | 20 | 11 | 47 | 11 | 799 | 89.3 | 6.37 | 786 | 11 | 37659663.32 | 22964211.58 | 22516337.81 | 18670254.51 | 29692939.49 | 16917823.43 | 26753032.63 | 28548087.9 | 32.8577 | 22.8967 | Rasal1 | 19415 | 0.9995 | 1.000497816 | 1.001 | 0.99619 | 0.998251 |
| 6755288 | ras-specific guanine nucleotide-releasing factor 1 isoform 1 [Mus musculus] | 0 | 100.131 | 23 | 21 | 70 | 19 | 1262 | 144 | 7.17 | 642 | 21 | 48314642.97 | 29085119.72 | 48418309.44 | 35962099.58 | 49597722.32 | 33465265.51 | 41239600.25 | 33631743.2 | 23.6572 | 19.3904 | Rasgrf1 | 19417 | 1.01203 | 0.987971017 | 0.97623 | 0.88046 | 0.980491 |
| 7242199 | ras-specific guanine nucleotide-releasing factor 2 [Mus musculus] | 0 | 93.494 | 23 | 18 | 51 | 16 | 1189 | 135.6 | 7.85 | 418 | 18 | 48353806.3 | 24003757.91 | 42398989.99 | 26271042.77 | 35842038.4 | 43079525.05 | 35624772.44 | 29607213.39 | 33.9537 | 15.2949 | Rasgrf2 | 19418 | 0.98904 | 1.010961287 | 1.02217 | 0.90947 | 0.98561 |
| 7242201 | RAS guanyl-releasing protein 1 [Mus musculus] | 0 | 53.948 | 17 | 8 | 28 | 8 | 795 | 90.2 | 8.1 | 548 | 8 | 14705145.8 | 10079327.59 | 13762993.63 | 10104724.81 | 14149694.73 | 10643820.74 | 12096879.93 | 10144410.89 | 19.9141 | 15.2764 | Rasgrp1 | 19419 | 1.0169 | 0.983097123 | 0.96676 | 0.79755 | 0.958226 |
| 1016275 | retinoblastoma-binding protein mRbAp48 [Mus musculus] | 0 | 14.604 | 7 | 4 | 11 | 4 | 461 | 51.7 | 5.11 | 180 | 4 | 8823690.871 | 8602034.472 | 8105397.526 | 10341384.65 | 5846194.48 | 7659005.864 | 5112030.5 | 3453655.577 | 10.7437 | 31.5957 | Rbbp4 | 19646 | 1.23819 | 0.761808549 | 0.61526 | 0.01339 | 0.461382 |
| 568936964 | PREDICTED: membrane-associated phosphatidylinositol transfer protein 2 isoform X1 [Mus musculus] | 0 | 25.179 | 5 | 6 | 22 | 6 | 1385 | 153.3 | 7.55 | 214 | 6 | 6175379.149 | 6396576.452 | 6412312.587 | 4217133.982 | 7233767.677 | 6816044.182 | 4131245.5 | 6029846.087 | 18.2921 | 22.7135 | Pitpnm2 | 19679 | 0.97871 | 1.021291968 | 1.04351 | 0.7811 | 0.952792 |
| 148696866 | UPF1 regulator of nonsense transcripts homolog (yeast), isoform CRA_a, partial [Mus musculus] | 0 | 79.887 | 21 | 19 | 73 | 19 | 1131 | 124.6 | 6.81 | 889 | 19 | 69224028.45 | 53425852.81 | 75466460.99 | 74573011.26 | 60925575.6 | 65262814.98 | 73928389.06 | 52167914.8 | 14.9769 | 14.3623 | Upf1 | 19704 | 1.03887 | 0.961132061 | 0.92517 | 0.48304 | 0.868779 |
| 146134422 | regulator of G-protein signaling 9 isoform 1 [Mus musculus] | 0 | 19.209 | 8 | 5 | 7 | 5 | 675 | 76.9 | 9.33 | 55 | 5 | 3936319.689 | 5246677.538 | 9958413.625 | 10829539.79 | 4451641.442 | 6089972.642 | 9196940.188 | 10933741.13 | 45.5253 | 38.272 | Rgs9 | 19739 | 0.98843 | 1.011565093 | 1.0234 | 0.94042 | 0.994906 |
| 755513554 | PREDICTED: rabphilin-3A isoform X1 [Mus musculus] | 0 | 306.246 | 54 | 29 | 281 | 28 | 684 | 75.8 | 8.18 | 4936 | 29 | 823453206.5 | 943914975.3 | 962405188.1 | 970676134.1 | 776733889.9 | 929757776.2 | 1021619382 | 1003565697 | 7.42504 | 11.9462 | Rph3a | 19894 | 0.9958 | 1.004201656 | 1.00844 | 0.90896 | 0.98561 |
| 13592009 | 60S ribosomal protein L10a [Rattus norvegicus] | 0 | 106.859 | 60 | 14 | 115 | 14 | 217 | 24.8 | 9.94 | 1253 | 14 | 411400226.6 | 481135911.3 | 387657638.5 | 488163331.3 | 388724929.2 | 477490442.3 | 420718178.4 | 425724229.2 | 11.3491 | 8.58165 | Rpl10a | 19896 | 1.016 | 0.984000163 | 0.9685 | 0.67002 | 0.922517 |
| 12840700 | unnamed protein product [Mus musculus] | 0 | 35.213 | 34 | 7 | 105 | 7 | 188 | 21.7 | 11.82 | 1319 | 7 | 105039257.7 | 420042276.5 | 208043711.2 | 306899459.7 | 197343266.2 | 224121933.1 | 248255811.3 | 337505816.6 | 51.8508 | 24.1452 | Rpl18 | 19899 | 1.01602 | 0.983979556 | 0.96846 | 0.91532 | 0.987992 |
| 198643 | ribosomal protein L19 [Mus musculus] | 0 | 34.878 | 26 | 6 | 134 | 6 | 196 | 23.5 | 11.47 | 1396 | 6 | 97570321.24 | 121931864.8 | 108501684.6 | 168991330.1 | 102252314.7 | 132606037 | 121503687.3 | 102835382.5 | 25.3107 | 12.9456 | Rpl19 | 19921 | 1.03953 | 0.960470539 | 0.92395 | 0.60648 | 0.905186 |
| 459683845 | 60S ribosomal protein L22 isoform b [Mus musculus] | 0 | 42.1 | 44 | 5 | 34 | 5 | 151 | 17.6 | 9.19 | 1046 | 5 | 114083027 | 169370180.7 | 163764194.9 | 178506977.5 | 120989601.4 | 116253684.8 | 134236947 | 115814105.3 | 18.4608 | 7.0601 | Rpl22 | 19934 | 1.12437 | 0.875626493 | 0.77877 | 0.06135 | 0.601634 |
| 4506621 | 60S ribosomal protein L26 [Homo sapiens] | 0 | 11.679 | 19 | 5 | 63 | 5 | 145 | 17.2 | 10.55 | 567 | 5 | 69180765.74 | 142668222.8 | 93936406.54 | 130876321.5 | 88496054.02 | 152454408 | 101498110 | 148853375.4 | 30.9494 | 26.5434 | Rpl26 | 19941 | 0.94112 | 1.058881865 | 1.12513 | 0.58183 | 0.898781 |
| 4506623 | 60S ribosomal protein L27 [Homo sapiens] | 0 | 30.928 | 43 | 5 | 49 | 5 | 136 | 15.8 | 10.56 | 337 | 5 | 276074817.1 | 425440247.6 | 410551869 | 448700355.8 | 191630957.1 | 553058428.5 | 440734644.1 | 605151726.9 | 19.9084 | 41.0926 | Rpl27 | 19942 | 0.93143 | 1.068571051 | 1.14724 | 0.58591 | 0.899211 |
| 148667873 | mCG112980, isoform CRA_b, partial [Mus musculus] | 0 | 23.558 | 31 | 3 | 30 | 3 | 123 | 13.7 | 10.45 | 445 | 3 | 82614882.07 | 132086239.9 | 87294016.83 | 128082132.5 | 78422799.84 | 111935546.8 | 109529051 | 134198243.7 | 24.346 | 21.1331 | Rpl37a | 19981 | 0.99536 | 1.004638443 | 1.00932 | 0.95595 | 0.994906 |
| 148704677 | mCG7611, isoform CRA_a, partial [Mus musculus] | 0 | 9.759 | 19 | 3 | 100 | 3 | 134 | 15.3 | 10.49 | 942 | 3 | 2875257.701 | 8639596.536 | 25628002.27 | 41410054.04 | 25466984.44 | 52081473.85 | 31556811.15 | 49464707.09 | 88.777 | 33.1318 | Rpl36a | 19982 | 0.66255 | 1.33744978 | 2.01864 | 0.11652 | 0.678142 |
| 38571759 | Ribosomal protein L6 [Mus musculus] | 0 | 95.952 | 39 | 16 | 189 | 16 | 296 | 33.5 | 10.7 | 3571 | 16 | 333753111.4 | 352359395.7 | 226872739 | 331255839.3 | 298942724.6 | 334964437.5 | 371009853.8 | 395219293.9 | 18.2952 | 12.0295 | Rpl6 | 19988 | 0.94105 | 1.05895347 | 1.12529 | 0.31307 | 0.76536 |
| 148682392 | mCG13479, isoform CRA_a [Mus musculus] | 0 | 77.761 | 51 | 15 | 229 | 15 | 270 | 31.4 | 10.78 | 1741 | 15 | 325838422 | 280433971.1 | 305433093.6 | 383178157.6 | 304083403.6 | 311576016.6 | 346611200.8 | 343529302.3 | 13.5214 | 6.66373 | Rpl7 | 19989 | 0.9958 | 1.004197465 | 1.00843 | 0.91473 | 0.987992 |
| 14149647 | 60S ribosomal protein L9 [Mus musculus] | 0 | 71.201 | 47 | 8 | 53 | 8 | 192 | 21.9 | 9.95 | 847 | 8 | 333359417.3 | 322643948 | 311412734.1 | 287734959.5 | 333515205 | 336845112 | 358991155.1 | 320127320.7 | 6.22832 | 4.77923 | Rpl9 | 20005 | 0.96378 | 1.036215408 | 1.07515 | 0.11199 | 0.678142 |
| 568916002 | PREDICTED: dolichyl-diphosphooligosaccharide--protein glycosyltransferase subunit 2 isoform X5 [Mus musculus] | 0 | 73.1 | 26 | 9 | 28 | 9 | 669 | 73.6 | 6.49 | 643 | 9 | 19236160.62 | 15371614.71 | 25318365.51 | 17781612.97 | 26412252.5 | 22027309.84 | 29214716.13 | 27048164.72 | 21.8183 | 11.5165 | Rpn2 | 20014 | 0.85201 | 1.147988927 | 1.34739 | 0.04094 | 0.585646 |
| 6755364 | activated RNA polymerase II transcriptional coactivator p15 [Mus musculus] | 0 | 18.836 | 27 | 3 | 26 | 3 | 127 | 14.4 | 9.6 | 348 | 3 | 28204219.57 | 44500986.68 | 53217391.82 | 54305376.54 | 28808960.56 | 46611660.24 | 57560188.25 | 79170543.63 | 26.7698 | 39.7244 | Sub1 | 20024 | 0.91864 | 1.081358461 | 1.17713 | 0.53527 | 0.886555 |
| 14277700 | 40S ribosomal protein S12 [Homo sapiens] | 0 | 94.148 | 64 | 7 | 123 | 7 | 132 | 14.5 | 7.21 | 1612 | 7 | 206610113.1 | 182522947.7 | 136615843 | 196184106.3 | 206729077.2 | 163082249.1 | 181411279.8 | 225188477.8 | 17.1004 | 14.1062 | Rps12 | 20042 | 0.96364 | 1.036358854 | 1.07546 | 0.53363 | 0.886555 |
| 3097244 | ribosomal protein S14 [Mus musculus] | 0 | 39.407 | 40 | 7 | 109 | 7 | 151 | 16.3 | 10.13 | 925 | 7 | 179142790 | 200894113.7 | 158283399.3 | 227289590.8 | 169458756.5 | 178949710.9 | 175248623.7 | 207751265.7 | 15.455 | 9.326 | Rps14 | 20044 | 1.02285 | 0.97715356 | 0.95533 | 0.63433 | 0.912586 |
| 4506687 | 40S ribosomal protein S15 isoform 2 [Homo sapiens] | 0 | 43.059 | 55 | 4 | 31 | 4 | 145 | 17 | 10.39 | 612 | 4 | 27799781.75 | 22244451.96 | 35326593.8 | 31439890.48 | 30940882.23 | 31771466.26 | 41811995 | 4279771.582 | 19.0549 | 59.0407 | Rps15 | 20054 | 1.03549 | 0.964512072 | 0.93146 | 0.82162 | 0.964126 |
| 60334810 | Rps16 protein [Mus musculus] | 0 | 20.435 | 35 | 6 | 39 | 6 | 172 | 19.3 | 10.2 | 238 | 6 | 167373963.1 | 242003489.3 | 167344316.9 | 225233815.6 | 157861942.4 | 220362664.3 | 188258229 | 221148381.8 | 19.384 | 15.34 | Rps16 | 20055 | 1.00901 | 0.990988622 | 0.98214 | 0.88907 | 0.984401 |
| 26377673 | unnamed protein product [Mus musculus] | 0 | 96.036 | 48 | 8 | 51 | 5 | 141 | 16.1 | 9.94 | 1817 | 8 | 93835994.39 | 118287639.2 | 86551732.87 | 124782333.2 | 107299328.6 | 115025444.8 | 98865805.5 | 97407272.59 | 17.5019 | 7.81356 | Rps17 | 20068 | 1.00577 | 0.99422859 | 0.98852 | 0.90841 | 0.98561 |
| 148692364 | mCG126277, isoform CRA_b [Mus musculus] | 0 | 31.019 | 31 | 7 | 63 | 7 | 162 | 17.8 | 10.37 | 983 | 7 | 243204156.3 | 366019207.6 | 339812488.4 | 393692980.1 | 295764162.8 | 382883102.1 | 301768448.3 | 502485844.5 | 19.5004 | 26.0049 | Rps19 | 20085 | 0.95039 | 1.049607594 | 1.10439 | 0.56955 | 0.895889 |
| 568987089 | PREDICTED: 40S ribosomal protein S24 isoform X1 [Mus musculus] | 0 | 35.51 | 27 | 5 | 42 | 5 | 175 | 20.2 | 10.55 | 573 | 5 | 71245351.52 | 132566292.6 | 81923804.9 | 106807762.1 | 90375231.46 | 141489855.6 | 112255451.5 | 159061462.5 | 27.8852 | 24.2462 | Rps24 | 20088 | 0.87648 | 1.123518673 | 1.28185 | 0.22572 | 0.718361 |
| 4506717 | 40S ribosomal protein S29 isoform 1 [Homo sapiens] | 0.001 | 4.197 | 32 | 2 | 7 | 2 | 56 | 6.7 | 10.13 | 24 | 2 | 3381206.178 | 14025932.47 | 13779671.43 | 22383187.32 | 9645537.297 | 24417823.73 | 16983274 | 35445315.78 | 58.0961 | 50.9316 | Rps29 | 20090 | 0.76495 | 1.235052802 | 1.61456 | 0.26797 | 0.745606 |
| 254553321 | 40S ribosomal protein S3a [Mus musculus] | 0 | 80.185 | 45 | 14 | 170 | 14 | 264 | 29.9 | 9.73 | 2237 | 14 | 449745527.7 | 532238781.7 | 449176699.8 | 518907290.8 | 418994312 | 451437592.7 | 532226566.8 | 550783860 | 9.08268 | 12.9484 | Rps3a1 | 20091 | 0.99914 | 1.000864358 | 1.00173 | 0.98327 | 0.997711 |
| 12851918 | unnamed protein product [Mus musculus] | 0 | 127.994 | 67 | 20 | 229 | 20 | 263 | 29.6 | 10.15 | 2174 | 20 | 316134943.9 | 499168742.8 | 327463685.4 | 344462414.2 | 370833770.9 | 345353781.9 | 401194661.6 | 304855180.9 | 23.0501 | 11.4715 | Rps4x | 20102 | 1.02234 | 0.977661755 | 0.9563 | 0.74374 | 0.947173 |
| 254675270 | 40S ribosomal protein S5 [Mus musculus] | 0 | 89.334 | 56 | 9 | 102 | 9 | 204 | 22.9 | 9.72 | 1933 | 9 | 97567712.65 | 105864045.5 | 100948146.4 | 103589650.3 | 107494995.1 | 110337956.6 | 109208915 | 108761195.3 | 3.49909 | 1.07902 | Rps5 | 20103 | 0.96701 | 1.032986976 | 1.06822 | 0.01004 | 0.449633 |
| 26389925 | unnamed protein product [Mus musculus] | 0 | 43.609 | 31 | 7 | 57 | 7 | 249 | 28.7 | 10.77 | 961 | 7 | 160879161.6 | 124649206.1 | 130889960 | 193076379 | 155126260.1 | 134172423.4 | 143655184.4 | 175252916.1 | 20.612 | 11.6287 | Rps6 | 20104 | 1.00106 | 0.998942333 | 0.99789 | 0.98632 | 0.998251 |
| 29294760 | Rps6ka1 protein, partial [Mus musculus] | 0 | 15.05 | 5 | 3 | 7 | 3 | 773 | 86.4 | 8.5 | 34 | 3 | 9402686.249 | 9581013.74 | 10947527.49 | 11734053.55 | 12520662.71 | 9212472.653 | 9598554 | 9217716.584 | 10.7246 | 15.7746 | Rps6ka1 | 20111 | 1.01357 | 0.986427303 | 0.97322 | 0.78448 | 0.952792 |
| 148705008 | mCG19129, partial [Mus musculus] | 0 | 51.822 | 51 | 9 | 96 | 9 | 213 | 24 | 10.18 | 896 | 9 | 115466317.7 | 153025965.8 | 147263750.4 | 201386063.2 | 174163599.8 | 169750576.1 | 175588847.3 | 180397472.4 | 22.9957 | 2.50712 | Rps7 | 20115 | 0.93716 | 1.062836539 | 1.1341 | 0.29107 | 0.754456 |
| 54607016 | reticulon-3 isoform 1 [Mus musculus] | 0 | 189.816 | 43 | 28 | 112 | 25 | 964 | 103.8 | 4.92 | 1472 | 28 | 117938797.3 | 91856811.16 | 137023705.3 | 92090670.03 | 113601150.5 | 102634977.3 | 149471582.6 | 74328363.65 | 19.9869 | 28.2512 | Rtn3 | 20168 | 0.99872 | 1.001281182 | 1.00257 | 0.98867 | 0.998251 |
| 6755382 | ruvB-like 2 [Mus musculus] | 0 | 20.451 | 14 | 5 | 15 | 5 | 463 | 51.1 | 5.64 | 137 | 5 | 5041293.887 | 5096105.504 | 7214332.514 | 7491832.341 | 10508924.56 | 7426444.492 | 7783571.375 | 8182572.343 | 21.3163 | 16.4055 | Ruvbl2 | 20174 | 0.84581 | 1.154190768 | 1.3646 | 0.05637 | 0.599194 |
| 1039743392 | PREDICTED: ryanodine receptor 2 isoform X7 [Mus musculus] | 0 | 114.783 | 7 | 26 | 79 | 26 | 4984 | 566.3 | 6.1 | 682 | 26 | 56181328.41 | 67255653.03 | 67177095.1 | 56073641.85 | 60604509.25 | 49099432.94 | 55972018.19 | 50804590.14 | 10.3814 | 9.64118 | Ryr2 | 20191 | 1.06522 | 0.934781439 | 0.87755 | 0.1172 | 0.678142 |
| 110626031 | KH domain-containing, RNA-binding, signal transduction-associated protein 1 [Mus musculus] | 0 | 48.643 | 21 | 6 | 48 | 5 | 443 | 48.3 | 8.72 | 593 | 6 | 72087237.71 | 69711157.35 | 86377897.35 | 62166908.12 | 88503753.45 | 47642510.98 | 101256248.8 | 52742934.22 | 13.9431 | 36.399 | Khdrbs1 | 20218 | 1.00034 | 0.999659333 | 0.99932 | 0.99732 | 0.998251 |
| 124244104 | ataxin-2 [Mus musculus] | 0 | 46.567 | 11 | 9 | 26 | 9 | 1286 | 136.4 | 9.54 | 534 | 9 | 9619533.905 | 4285470.087 | 11169218.32 | 10154669.47 | 15323953.34 | 16182647.29 | 20084542.5 | 17215827.98 | 34.9968 | 12.0433 | Atxn2 | 20239 | 0.67725 | 1.32275485 | 1.95314 | 0.00401 | 0.38143 |
| 117414174 | sodium channel protein type 8 subunit alpha isoform 1 [Mus musculus] | 0 | 55.934 | 8 | 12 | 46 | 8 | 1978 | 225 | 6.3 | 549 | 12 | 38005730.9 | 24419952.69 | 37814651.48 | 9862859.859 | 49966351.71 | 20970332.34 | 35567017.75 | 21344906.93 | 48.6199 | 43.1523 | Scn8a | 20273 | 0.92542 | 1.074575664 | 1.16117 | 0.66057 | 0.919605 |
| 148672612 | mCG8782 [Mus musculus] | 0 | 12.679 | 4 | 2 | 4 | 2 | 989 | 111 | 7.42 | 49 | 2 | 3100368.626 | 3755365.197 | 2069390.369 | 1561733.798 | 4097437.835 | 1086022.238 | 2880717.25 | 1239117.331 | 37.7782 | 61.6331 | Zc3h7b | 20286 | 1.05981 | 0.94019433 | 0.88714 | 0.7457 | 0.947173 |
| 568960895 | PREDICTED: neuroplastin isoform X1 [Mus musculus] | 0 | 25.835 | 16 | 5 | 28 | 5 | 397 | 44.3 | 7.74 | 249 | 5 | 15689259.88 | 19103557.35 | 14000334.09 | 16675881.47 | 15315741.27 | 18592569.27 | 12849988.63 | 16242988.77 | 13.03 | 15.0791 | Nptn | 20320 | 1.01921 | 0.980791323 | 0.96231 | 0.71242 | 0.936509 |
| 14290512 | SEC22 vesicle trafficking protein homolog B (S. cerevisiae) [Mus musculus] | 0 | 16.252 | 20 | 4 | 9 | 4 | 215 | 24.7 | 8.51 | 121 | 4 | 2864083.724 | 3565990.763 | 4360922.189 | 3287661.128 | 5051397.671 | 4974196.295 | 5291220.25 | 4198429.244 | 17.9203 | 9.70003 | Sec22b | 20333 | 0.83817 | 1.161832519 | 1.38616 | 0.01368 | 0.461382 |
| 67906177 | protein transport protein Sec23A [Mus musculus] | 0 | 86.456 | 27 | 15 | 50 | 12 | 765 | 86.1 | 7.08 | 993 | 15 | 43491556.81 | 38099987.43 | 53492096.66 | 46065465.05 | 51606061.51 | 49732004.88 | 50119822.13 | 55175187.73 | 14.1278 | 4.80046 | Sec23a | 20334 | 0.93428 | 1.065717229 | 1.14068 | 0.1127 | 0.678142 |
| 50511055 | mKIAA1699 protein [Mus musculus] | 0 | 23.937 | 10 | 8 | 10 | 8 | 977 | 110.7 | 6.55 | 27 | 8 | 5533181.679 | 3699194.097 | 5935399.184 | 3681156.495 | 8814988.4 | 3982047.563 | 5584147.875 | 10451317.75 | 25.2866 | 40.9556 | Exoc4 | 20336 | 0.79062 | 1.209380665 | 1.52966 | 0.16793 | 0.711915 |
| 568922184 | PREDICTED: semaphorin-4A isoform X1 [Mus musculus] | 0 | 9.761 | 7 | 3 | 6 | 3 | 798 | 87.6 | 7.99 | 36 | 3 | 5536929.607 | 4708686.338 | 4643118.339 | 4283374.103 | 2175895.247 | 4656143.571 | 5178766.75 | 7389919.051 | 11.0579 | 44.1363 | Sema4a | 20351 | 0.99407 | 1.005926872 | 1.01192 | 0.96034 | 0.994906 |
| 755537830 | PREDICTED: septin-8 isoform X1 [Mus musculus] | 0 | 136.742 | 52 | 18 | 103 | 3 | 484 | 55.8 | 6.1 | 1350 | 18 | 13643644.24 | 14890199.62 | 14268001.83 | 11178390.94 | 15142919.97 | 18795629.65 | 17292083.13 | 20952215.21 | 12.0498 | 13.5754 | 43716 | 20362 | 0.85572 | 1.144278427 | 1.33721 | 0.02124 | 0.505785 |
| 26345390 | unnamed protein product [Mus musculus] | 0 | 20.885 | 16 | 3 | 11 | 3 | 254 | 29 | 11.22 | 340 | 3 | 2615990.967 | 772117.7885 | 8924197.324 | 12934226.57 | 5891123.075 | 8415227.047 | 9382202 | 13445293.06 | 89.1711 | 33.8314 | Srsf2 | 20382 | 0.80944 | 1.190561727 | 1.47085 | 0.392 | 0.824486 |
| 31560792 | endophilin-A1 [Mus musculus] | 0 | 46.613 | 30 | 9 | 46 | 7 | 352 | 39.9 | 5.39 | 466 | 9 | 35935480.33 | 36659886.86 | 39514020.65 | 34419368.34 | 31171572.15 | 15704958.99 | 30398034.25 | 11291789.65 | 5.83098 | 45.8258 | Sh3gl2 | 20404 | 1.24655 | 0.75345127 | 0.60443 | 0.03137 | 0.515682 |
| 7305485 | endophilin-A2 isoform 1 [Mus musculus] | 0 | 29.111 | 26 | 7 | 31 | 5 | 368 | 41.5 | 5.72 | 471 | 7 | 3855870.687 | 802960.148 | 3640056.877 | 4021942.412 | 3319398.448 | 3150286.697 | 3065905.5 | 2217487.974 | 49.5484 | 16.7427 | Sh3gl1 | 20405 | 1.02358 | 0.976416314 | 0.95392 | 0.86531 | 0.978314 |
| 6755504 | vinexin isoform a [Mus musculus] | 0 | 13.641 | 6 | 3 | 10 | 3 | 733 | 82.3 | 9.2 | 39 | 3 | 3326923.415 | 1199705.99 | 1388013.755 | 2850816.628 | 4296143.806 | 1090383.832 | 3425121.656 | 2227125.219 | 48.2447 | 50.7031 | Sorbs3 | 20410 | 0.88521 | 1.114789327 | 1.25935 | 0.54089 | 0.886555 |
| 569006631 | PREDICTED: sorbin and SH3 domain-containing protein 1 isoform X1 [Mus musculus] | 0 | 416.037 | 38 | 62 | 552 | 61 | 2079 | 228.1 | 7.96 | 9311 | 62 | 1075178880 | 865756229 | 1051178493 | 915102060.6 | 1016057907 | 770769440.6 | 932620713.4 | 860071457.3 | 10.4649 | 11.6709 | Sorbs1 | 20411 | 1.04377 | 0.956229767 | 0.91613 | 0.30507 | 0.761857 |
| 258547119 | cytoplasmic FMR1-interacting protein 1 isoform b [Mus musculus] | 0 | 274.441 | 44 | 45 | 282 | 25 | 1251 | 144.8 | 6.95 | 4296 | 45 | 93356965.97 | 56805118.35 | 114955412.6 | 76574519.37 | 107271788.8 | 79884103.87 | 131535633.8 | 85389850.64 | 28.9317 | 23.2947 | Cyfip1 | 20430 | 0.91634 | 1.083657263 | 1.18259 | 0.39591 | 0.824613 |
| 26328241 | unnamed protein product [Mus musculus] | 0 | 16.241 | 10 | 4 | 6 | 4 | 678 | 76.1 | 8.76 | 32 | 4 | 9283858.833 | 6839807.494 | 8793628.496 | 8610980.689 | 10576953.99 | 8652602.242 | 10792310.25 | 6362031.528 | 12.726 | 22.6572 | Clpb | 20480 | 0.95915 | 1.040845855 | 1.08517 | 0.56097 | 0.895889 |

| | | | | | | | | | | | | | | | | | | | | | | | | | | | | | |
|---|---|---|---|---|---|---|---|---|---|---|---|---|---|---|---|---|---|---|---|---|---|---|---|---|---|---|---|---|---|
| 124517716 | solute carrier family 12 member 2 [Mus musculus] | 0 | 8.742 | 2 | 2 | 10 | 2 | 1206 | 130.6 | 6.52 | 0 | 2 | 5493332.706 | 5437801.244 | 5406763.901 | 5661051.866 | 6224102.798 | 4793485.966 | 6709602.25 | 4037647.352 | 2.06098 | 22.7853 | Slc12a2 | 20496 | 1.00535 | 0.994650569 | 0.98936 | 0.92815 | 0.991703 |
| 882343 | high affinity glutamate trasporter [Mus musculus] | 0 | 88.402 | 19 | 11 | 145 | 11 | 574 | 62.4 | 6.4 | 2373 | 11 | 106705577.6 | 132248598.6 | 230490818.1 | 108189705 | 141854176.2 | 125104529.9 | 110779278.8 | 147369401.1 | 40.5585 | 12.6635 | Slc1a2 | 20511 | 1.04763 | 0.952366638 | 0.90906 | 0.68126 | 0.926957 |
| 24233554 | excitatory amino acid transporter 1 [Mus musculus] | 0 | 69.834 | 16 | 6 | 52 | 5 | 543 | 59.6 | 8.4 | 1322 | 6 | 109254586.4 | 74122794.74 | 108905737.7 | 46930708.08 | 127750723.8 | 76044914.32 | 130544193.3 | 57240680.74 | 35.5537 | 37.7059 | Slc1a3 | 20512 | 0.92834 | 1.071657212 | 1.15438 | 0.60259 | 0.905186 |
| 568967097 | PREDICTED: excitatory amino acid transporter 4 isoform X1 [Mus musculus] | 0 | 25.483 | 11 | 4 | 21 | 3 | 574 | 62.1 | 8.62 | 267 | 4 | 6743931.979 | 3984300.433 | 5204020.58 | 6025993.283 | 5529647.781 | 5396618.782 | 4816330.875 | 5043990.543 | 21.5758 | 6.27322 | Slc1a6 | 20513 | 1.02741 | 0.972589476 | 0.94664 | 0.65032 | 0.915864 |
| 568999605 | PREDICTED: sodium/calcium exchanger 1 isoform X11 [Mus musculus] | 0 | 54.302 | 15 | 9 | 37 | 9 | 935 | 104 | 4.92 | 467 | 9 | 20131379.88 | 15868431.23 | 29385637.75 | 15018872.82 | 23210150.66 | 19499881.58 | 24967307.06 | 14505795.42 | 32.742 | 22.5199 | Slc8a1 | 20541 | 0.98906 | 1.010940654 | 1.02212 | 0.91558 | 0.987992 |
| 755514082 | PREDICTED: nuclear receptor corepressor 2 isoform X1 [Mus musculus] | 0 | 71.179 | 9 | 16 | 43 | 16 | 2546 | 277.3 | 7.4 | 257 | 16 | 25474917.15 | 21661784.23 | 27716183.84 | 21659462.19 | 23421831.48 | 18299174.69 | 30658155.63 | 30622978.55 | 12.4026 | 23.3835 | Ncor2 | 20602 | 0.96747 | 1.032527928 | 1.06724 | 0.64649 | 0.915705 |
| 6755588 | synaptosomal-associated protein 25 isoform a [Mus musculus] | 0 | 151.675 | 75 | 19 | 239 | 5 | 206 | 23.3 | 4.77 | 5099 | 19 | 82075787.41 | 87960423.74 | 99500416.96 | 71901412.84 | 83593670.2 | 69490332.51 | 94228601.5 | 84334310.31 | 13.5044 | 12.2741 | Snap25 | 20614 | 1.01455 | 0.985453357 | 0.97132 | 0.76099 | 0.952792 |
| 7305507 | clathrin coat assembly protein AP180 isoform 1 [Mus musculus] | 0 | 31.016 | 10 | 6 | 17 | 5 | 901 | 91.8 | 4.88 | 363 | 6 | 16362468.15 | 15008269.76 | 7627346.995 | 16447724.69 | 17146202.77 | 15121227.17 | 13795410.94 | 5490508.314 | 30.358 | 39.7316 | Snap91 | 20616 | 1.03638 | 0.963621579 | 0.9298 | 0.77893 | 0.952792 |
| 223462828 | SNF related kinase [Mus musculus] | 0 | 69.385 | 25 | 14 | 51 | 14 | 748 | 81.9 | 7.62 | 648 | 14 | 79160110.3 | 63417933.37 | 76710499.56 | 74693481.36 | 76213359.77 | 76473766.31 | 62777135.75 | 64570847.88 | 9.47296 | 10.5017 | Snrk | 20623 | 1.0243 | 0.975702964 | 0.95256 | 0.51674 | 0.88031 |
| 148674121 | syntrophin, acidic 1, isoform CRA_a, partial [Mus musculus] | 0 | 25.92 | 18 | 6 | 17 | 6 | 519 | 55.5 | 7.21 | 265 | 6 | 16034708.45 | 15199358.05 | 14294094.81 | 11179448.01 | 16591980.59 | 16997541.45 | 16076086.38 | 15771156.58 | 14.9606 | 3.32499 | Snta1 | 20648 | 0.92853 | 1.071465884 | 1.15393 | 0.09328 | 0.662454 |
| 219519541 | Sntb2 protein [Mus musculus] | 0.001 | 6.408 | 4 | 2 | 10 | 2 | 531 | 57.4 | 9.33 | 59 | 2 | 1480013.86 | 1355138.415 | 1672213.492 | 2035414.609 | 817550.5135 | 1948760.451 | 1693523.625 | 1529475.919 | 18.1379 | 32.3851 | Sntb2 | 20650 | 1.04416 | 0.955835792 | 0.91541 | 0.64367 | 0.915705 |
| 6678077 | SPARC precursor [Mus musculus] | 0 | 32.026 | 26 | 8 | 22 | 8 | 302 | 34.4 | 4.86 | 111 | 8 | 27322325.84 | 32730744.29 | 22863356.26 | 30847767.76 | 36286285.73 | 49557487.98 | 27264131.13 | 54248249.6 | 15.2663 | 29.4939 | Sparc | 20692 | 0.80936 | 1.190637073 | 1.47108 | 0.08644 | 0.655499 |
| 14715029 | Serine (or cysteine) peptidase inhibitor, clade E, member 2 [Mus musculus] | 0 | 13.202 | 9 | 4 | 18 | 4 | 397 | 44.2 | 9.85 | 107 | 4 | 11184309.58 | 9570299.317 | 9112830.485 | 9318773.444 | 10068659.1 | 7066365.404 | 10819925.5 | 7390056.746 | 9.63496 | 21.3507 | Serpine2 | 20720 | 1.05154 | 0.948461784 | 0.90198 | 0.39769 | 0.824613 |
| 568913208 | PREDICTED: spectrin alpha chain, non-erythrocytic 1 isoform X3 [Mus musculus] | 0 | 3160.87 | 78 | 226 | 7078 | 2 | 2490 | 286.7 | 5.39 | 159385 | 226 | 39140399.68 | 32075098.04 | 47431359.96 | 42154633.77 | 23671518.95 | 32892564.75 | 42149894.25 | 33251796.92 | 15.9442 | 22.8719 | Sptan1 | 20740 | 1.09849 | 0.90150635 | 0.82068 | 0.19557 | 0.715071 |
| 187956419 | Spectrin beta 1 [Mus musculus] | 0 | 1212.86 | 67 | 141 | 1334 | 132 | 2329 | 267.9 | 5.49 | 24359 | 141 | 1774062838 | 1680990001 | 2037682325 | 1912482075 | 2458490118 | 1723368622 | 2063625308 | 1882797398 | 8.45212 | 15.5724 | Sptb | 20741 | 0.95345 | 1.046548702 | 1.09764 | 0.34528 | 0.78974 |
| 117938332 | spectrin beta chain, non-erythrocytic 1 isoform 1 [Mus musculus] | 0 | 3001.78 | 78 | 220 | 4920 | 21 | 2363 | 274.1 | 5.58 | 100933 | 220 | 2592785910 | 2045691458 | 2839678963 | 2477774244 | 2531854248 | 1828081295 | 2639377358 | 2279053419 | 13.3332 | 15.5547 | Sptbn1 | 20742 | 1.03523 | 0.964773121 | 0.93194 | 0.51535 | 0.88031 |
| 55926127 | spectrin beta chain, non-erythrocytic 2 [Mus musculus] | 0 | 2142.62 | 76 | 169 | 3038 | 154 | 2388 | 270.8 | 5.86 | 64434 | 169 | 10272554110 | 8943642186 | 10623264574 | 9823143417 | 8844228420 | 7702138674 | 9774862745 | 9061628526 | 7.32229 | 9.71982 | Sptbn2 | 20743 | 1.05703 | 0.94297129 | 0.8921 | 0.10594 | 0.674486 |
| 568916084 | PREDICTED: neuronal proto-oncogene tyrosine-protein kinase Src isoform X1 [Mus musculus] | 0 | 28.738 | 15 | 7 | 31 | 4 | 552 | 62.1 | 8.02 | 303 | 7 | 6519698.482 | 7728895.249 | 7749727.392 | 11189600.24 | 9258013.364 | 10280115.71 | 6772777.25 | 13545432.64 | 24.2535 | 28.154 | Src | 20779 | 0.90871 | 1.091292835 | 1.20093 | 0.37146 | 0.80688 |
| 2980671 | protein kinase [Mus musculus] | 0 | 8.745 | 6 | 3 | 3 | 2 | 648 | 73.1 | 6.19 | 0 | 3 | 1452075.138 | 1357701.543 | 2702831.882 | 1505930.647 | 1308717.188 | 2573658.316 | 2762408.563 | 2454557.472 | 36.1952 | 28.8562 | Srpk1 | 20815 | 0.8709 | 1.129099002 | 1.29647 | 0.2981 | 0.761852 |
| 755510887 | PREDICTED: SRSF protein kinase 2 isoform X1 [Mus musculus] | 0 | 28.743 | 12 | 7 | 24 | 6 | 750 | 84.9 | 5.21 | 153 | 7 | 18318874.87 | 15456481.4 | 21982009.46 | 13017722.04 | 17255069.9 | 9885948.111 | 16610774.5 | 16452349.53 | 22.4379 | 22.9941 | Srpk2 | 20817 | 1.06645 | 0.933547861 | 0.87538 | 0.43995 | 0.852048 |
| 12842535 | unnamed protein product [Mus musculus] | 0 | 26.424 | 23 | 4 | 12 | 4 | 269 | 29.6 | 9.28 | 124 | 4 | 524643.7719 | 2992883.118 | 1890290.719 | 4805532.555 | 3157465.959 | 2925179.977 | 2482183.125 | 4374791.709 | 70.8606 | 25.0366 | Srprb | 20818 | 0.88225 | 1.117750358 | 1.26693 | 0.51736 | 0.88031 |
| 148695104 | Sjogren syndrome antigen B, isoform CRA_b, partial [Mus musculus] | 0 | 24.307 | 14 | 5 | 16 | 5 | 422 | 48.5 | 9.74 | 681 | 5 | 10408384.51 | 1640489.854 | 3895669.626 | 9201203.466 | 5778411.646 | 5098971.645 | 7526997.5 | 6628314.719 | 66.7264 | 16.8119 | Ssb | 20823 | 1.00225 | 0.997747002 | 0.9955 | 0.99 | 0.998251 |
| 74201485 | unnamed protein product [Mus musculus] | 0 | 15.243 | 7 | 4 | 18 | 4 | 553 | 60.2 | 4.84 | 184 | 4 | 10258355.37 | 16355164.18 | 11152132.54 | 8511414.064 | 12173029.59 | 6629459.261 | 9437680.25 | 6167844.507 | 29.1618 | 32.3748 | Stam | 20844 | 1.1471 | 0.852896563 | 0.74352 | 0.22374 | 0.718033 |
| 568950016 | PREDICTED: stromal interaction molecule 1 isoform X1 [Mus musculus] | 0.001 | 6.014 | 3 | 2 | 4 | 2 | 793 | 88.9 | 6.55 | 23 | 2 | 1923578.128 | 2826610.867 | 2865532.137 | 1167051.795 | 2851403.795 | 2919085.244 | 2890025 | 2318264.535 | 36.9895 | 10.4068 | Stim1 | 20866 | 0.88887 | 1.111125164 | 1.25004 | 0.24934 | 0.733806 |
| 4063383 | serine-threonine kinase receptor-associated protein [Mus musculus] | 0 | 23.684 | 17 | 4 | 10 | 4 | 351 | 38.5 | 5.12 | 285 | 4 | 4984444.955 | 7160769.3 | 6501590.048 | 7288521.371 | 6730458.372 | 5754613.03 | 5602614.875 | 5493028.159 | 16.3082 | 9.61954 | Strap | 20901 | 1.04755 | 0.952447505 | 0.90921 | 0.36439 | 0.805816 |
| 15011853 | syntaxin-1A [Mus musculus] | 0 | 35.908 | 28 | 6 | 22 | 6 | 288 | 33 | 5.24 | 527 | 6 | 19594207.73 | 12843402.67 | 20142970.92 | 12882820.72 | 23190594.3 | 22351561.18 | 22505590 | 11955224.77 | 24.7518 | 26.8793 | Stx1a | 20907 | 0.90005 | 1.099951404 | 1.2221 | 0.32164 | 0.771047 |
| 6981602 | syntaxin-binding protein 1 [Rattus norvegicus] | 0 | 147.844 | 50 | 22 | 139 | 22 | 594 | 67.5 | 6.96 | 2037 | 22 | 117603118.6 | 102466980.4 | 132628601.2 | 126545318.1 | 158302192.2 | 108652644.1 | 130693526.7 | 93097857.09 | 10.9392 | 23.0782 | Stxbp1 | 20910 | 0.98814 | 1.011858059 | 1.024 | 0.85983 | 0.976238 |
| 6755690 | syntaxin-binding protein 3 [Mus musculus] | 0 | 21.908 | 10 | 5 | 12 | 5 | 592 | 67.9 | 8.02 | 47 | 5 | 9196656.192 | 7601383.996 | 10933232.1 | 5651549.88 | 11578106 | 6538983.145 | 9811183.25 | 6313088.811 | 26.9994 | 30.0167 | Stxbp3 | 20912 | 0.9873 | 1.01269574 | 1.02572 | 0.90415 | 0.98561 |
| 568987140 | PREDICTED: succinyl-CoA ligase [ADP-forming] subunit beta, mitochondrial isoform X1 [Mus musculus] | 0 | 25.027 | 12 | 5 | 13 | 5 | 500 | 53.8 | 7.53 | 74 | 5 | 274601868.4 | 206395254.6 | 226276312 | 152676059.1 | 305777112.7 | 203104877.2 | 207310501.9 | 222919796.5 | 23.4699 | 20.4849 | Sucla2 | 20916 | 0.956 | 1.044002266 | 1.09206 | 0.59075 | 0.902243 |
| 160707901 | synapsin-1 isoform a [Mus musculus] | 0 | 476.382 | 76 | 34 | 1075 | 32 | 706 | 74.1 | 9.8 | 21575 | 34 | 2768632784 | 2406315304 | 2521483068 | 2778212356 | 2570320299 | 2047708029 | 2379593296 | 2936937754 | 7.05805 | 14.9546 | Syn1 | 20964 | 1.02646 | 0.973537225 | 0.94844 | 0.53921 | 0.886555 |
| 161168987 | synapsin-2 isoform IIa [Mus musculus] | 0 | 429.414 | 69 | 27 | 633 | 2 | 586 | 63.3 | 8.43 | 12317 | 27 | 28967507.71 | 12238886.27 | 29079691.54 | 57441224.08 | 43759620.57 | 9311850.519 | 34329516.69 | 50631128.72 | 58.7404 | 52.3874 | Syn2 | 20965 | 0.96123 | 1.038774944 | 1.08068 | 0.84974 | 0.97368 |
| 31560541 | synaptogyrin-3 [Mus musculus] | 0 | 7.405 | 9 | 2 | 21 | 2 | 229 | 24.5 | 8.18 | 74 | 2 | 7003477.46 | 10429578.21 | 6126674.637 | 9241709.053 | 6542587.406 | 11050500.37 | 5282102.313 | 10306566.72 | 24.1713 | 33.945 | Syngr3 | 20974 | 0.99424 | 1.005763853 | 1.01159 | 0.95775 | 0.994906 |
| 6678197 | synaptotagmin-1 isoform 1 [Mus musculus] | 0 | 245.649 | 65 | 29 | 241 | 3 | 421 | 47.4 | 8.53 | 3783 | 29 | 33740565.59 | 78925153.92 | 33246387.57 | 12260543.74 | 52779486.56 | 44005687.12 | 35314170.06 | 52826902.28 | 71.0582 | 18.1195 | Syt1 | 20979 | 0.92202 | 1.077976337 | 1.16914 | 0.66428 | 0.922517 |
| 31543797 | synaptotagmin-2 [Mus musculus] | 0 | 132.873 | 52 | 20 | 112 | 13 | 422 | 47.2 | 7.99 | 1578 | 20 | 75995229.08 | 46475114.72 | 55482702.41 | 45690461.96 | 66810824.72 | 52970534.28 | 59342464.94 | 56673428.16 | 25.2319 | 9.93421 | Syt2 | 20980 | 0.97355 | 1.026453343 | 1.05434 | 0.70452 | 0.933129 |
| 1321655 | synaptotagumin III [Mus musculus] | 0 | 80.92 | 28 | 12 | 41 | 10 | 587 | 63.2 | 6.62 | 910 | 12 | 38513469.9 | 40007929.84 | 49624154.01 | 41822234.4 | 54172072.08 | 51238365.8 | 51056359.75 | 43521312.42 | 11.634 | 9.09479 | Syt3 | 20981 | 0.91885 | 1.081145677 | 1.17662 | 0.06683 | 0.601634 |
| 54607084 | contactin-2 precursor [Mus musculus] | 0 | 13.067 | 4 | 3 | 10 | 3 | 1040 | 113.1 | 7.62 | 94 | 3 | 2892390.969 | 1662470.062 | 2188271.068 | 1994240.611 | 2706461.75 | 1302784.357 | 2050877 | 1774076.728 | 23.7858 | 29.9457 | Cntn2 | 21367 | 1.0545 | 0.945498662 | 0.89663 | 0.58534 | 0.899211 |
| 56090475 | S-phase kinase-associated protein 1 [Rattus norvegicus] | 0 | 77.327 | 83 | 11 | 75 | 11 | 163 | 18.7 | 4.54 | 1191 | 11 | 94009665.23 | 105005834.7 | 86838369.03 | 98373271.28 | 103658189.7 | 94149678.8 | 78480204.14 | 76299192.71 | 7.94255 | 14.7982 | Skp1a | 21402 | 1.04294 | 0.957058556 | 0.91765 | 0.33548 | 0.782118 |
| 1039748586 | PREDICTED: alpha-globin transcription factor CP2 isoform X1 [Mus musculus] | 0 | 8.794 | 5 | 2 | 5 | 2 | 566 | 64 | 5.8 | 43 | 2 | 4203720.505 | 2617301.453 | 1889260.791 | 1590294.615 | 1916358.209 | 908092.7122 | 2419135 | 1585179.634 | 45.3663 | 37.1092 | Tfcp2 | 21422 | 1.20268 | 0.797317864 | 0.66295 | 0.23932 | 0.732938 |
| 1039736493 | PREDICTED: nucleolar transcription factor 1 isoform X1 [Mus musculus] | 0 | 30.212 | 9 | 6 | 14 | 6 | 795 | 93 | 6.06 | 103 | 6 | 12015895.95 | 19161705.09 | 15217053.35 | 13716943.99 | 23420304.54 | 11024367.39 | 18933182.25 | 16177673.55 | 20.2983 | 29.8335 | Ubtf | 21429 | 0.92717 | 1.072832102 | 1.15711 | 0.46252 | 0.863085 |
| 110625624 | T-complex protein 1 subunit alpha isoform 1 [Mus musculus] | 0 | 80.502 | 30 | 15 | 68 | 15 | 556 | 60.4 | 6.16 | 864 | 15 | 53193358.06 | 71241196.28 | 64719090.21 | 75339496.24 | 66483197.29 | 72351011.52 | 55446601.13 | 60454803.8 | 14.6179 | 11.5117 | Tcp1 | 21454 | 1.01879 | 0.981207653 | 0.96311 | 0.7015 | 0.931521 |
| 12846252 | unnamed protein product [Mus musculus] | 0 | 81.183 | 62 | 8 | 47 | 8 | 198 | 21.8 | 5.41 | 871 | 8 | 103649603.9 | 97119325.66 | 97477383.34 | 102944712.1 | 104960394.4 | 109064244.7 | 90031375 | 88158706.12 | 3.46809 | 10.7158 | Prdx2 | 21672 | 1.01131 | 0.988686363 | 0.97763 | 0.69917 | 0.931521 |
| 6755763 | THO complex subunit 4 [Mus musculus] | 0 | 46.141 | 37 | 5 | 28 | 5 | 255 | 26.9 | 11.15 | 916 | 5 | 21117537.41 | 26909494.67 | 30088789.91 | 41675518.9 | 20967019.1 | 32865349.88 | 32052474.75 | 32406299.74 | 28.9026 | 19.4328 | Alyref | 21681 | 1.0063 | 0.993698833 | 0.98748 | 0.94479 | 0.994906 |
| 755556338 | DICTED: serine/threonine-protein phosphatase 2A 56 kDa regulatory subunit delta isoform isoform X1 [Mus musc | 0 | 119.186 | 33 | 15 | 80 | 8 | 595 | 69.1 | 7.96 | 1315 | 15 | 34711289.52 | 33855482.46 | 31607066.46 | 34163412.67 | 34533299.52 | 42217777.71 | 35502308.25 | 36364660.31 | 4.06396 | 9.30549 | Ppp2r5d | 21770 | 0.94953 | 1.050470144 | 1.10631 | 0.10313 | 0.674486 |
| 6678303 | transcription factor A, mitochondrial precursor [Mus musculus] | 0 | 25.242 | 24 | 6 | 35 | 6 | 243 | 28 | 9.69 | 278 | 6 | 77870208.18 | 64902107.53 | 50825725.29 | 75063197.29 | 61235165.55 | 95742658.11 | 66428158.88 | 77534325.74 | 18.216 | 20.2948 | Tfam | 21780 | 0.94333 | 1.056669562 | 1.12015 | 0.44099 | 0.852048 |
| 38325830 | CD90.1 [Mus musculus] | 0 | 67.247 | 37 | 7 | 96 | 7 | 162 | 18.1 | 9.19 | 1956 | 7 | 801840262.8 | 974867090.7 | 499367547.6 | 836937132.2 | 827913133.2 | 864926736.9 | 632866490.3 | 762726867.3 | 25.7451 | 13.2093 | Thy1 | 21838 | 1.00396 | 0.9960366 | 0.9921 | 0.95818 | 0.994906 |
| 148706135 | tripartite motif protein 28 [Mus musculus] | 0 | 17.841 | 7 | 5 | 12 | 5 | 834 | 88.8 | 5.77 | 39 | 5 | 54350613.26 | 40972038.52 | 50273337.07 | 36689915.38 | 54902841.15 | 39781317.24 | 40633422.63 | 48605005.52 | 17.8818 | 15.5576 | Trim28 | 21849 | 0.99553 | 1.004469264 | 1.00898 | 0.94229 | 0.994906 |
| 568947657 | PREDICTED: tight junction protein ZO-1 isoform X7 [Mus musculus] | 0 | 357.84 | 41 | 53 | 313 | 7 | 1795 | 201.7 | 7.47 | 4453 | 53 | 33246431.71 | 26781104.96 | 24985865.17 | 36309284.12 | 22988001.14 | 34171831.2 | 28604041.1 | 26934277.3 | 17.592 | 16.469 | Tjp1 | 21872 | 1.03685 | 0.963146294 | 0.92891 | 0.56433 | 0.895889 |
| 148709667 | tight junction protein 2, isoform CRA_b [Mus musculus] | 0 | 64.948 | 17 | 15 | 64 | 15 | 1229 | 138 | 7.17 | 483 | 15 | 11098981.01 | 12837589.03 | 31840435.32 | 19781353.46 | 29331058.78 | 21819086.61 | 16664936.97 | 22283269.31 | 49.8341 | 23.1 | Tjp2 | 21873 | 0.91223 | 1.087771831 | 1.19243 | 0.52428 | 0.88292 |
| 6678359 | transketolase [Mus musculus] | 0 | 22.578 | 9 | 3 | 11 | 3 | 623 | 67.6 | 7.5 | 317 | 3 | 4655573.565 | 2675424.675 | 3692671.429 | 3439800.137 | 4046511.688 | 883113.2261 | 3148096.25 | 475232.6449 | 22.594 | 81.0137 | Tkt | 21881 | 1.2568 | 0.743204414 | 0.59135 | 0.17377 | 0.713031 |
| 11230778 | tropomodulin-1 [Mus musculus] | 0 | 119.755 | 60 | 17 | 72 | 16 | 359 | 40.4 | 5.1 | 1673 | 17 | 100925646.3 | 79890850.97 | 118175546.6 | 65536422.53 | 94798617.17 | 62851467.82 | 97609871.28 | 83373455.94 | 25.4106 | 18.6488 | Tmod1 | 21916 | 1.03683 | 0.96317341 | 0.92896 | 0.66038 | 0.919605 |
| 121949765 | lamina-associated polypeptide 2 isoform epsilon [Mus musculus] | 0 | 112.735 | 52 | 15 | 107 | 15 | 452 | 50.3 | 9.45 | 1976 | 15 | 153089298.5 | 110161497.6 | 140314856.6 | 173997863.2 | 126531613.8 | 209164925.2 | 171894288.9 | 180614731.9 | 18.4994 | 19.9171 | Tmpo | 21917 | 0.91259 | 1.087410924 | 1.19157 | 0.25003 | 0.733806 |
| 568926394 | PREDICTED: tenascin isoform X1 [Mus musculus] | 0 | 176.021 | 22 | 33 | 154 | 33 | 2110 | 231.7 | 4.89 | 2223 | 33 | 71386151.02 | 66458333.17 | 66997222.74 | 66689502.55 | 89132041.36 | 130459646 | 84488795.06 | 96611993.81 | 3.45592 | 20.7636 | Tnc | 21923 | 0.80786 | 1.192140309 | 1.47568 | 0.02152 | 0.505785 |
| 226958549 | tenascin-R precursor [Mus musculus] | 0 | 285.54 | 35 | 33 | 221 | 33 | 1358 | 149.5 | 4.94 | 6621 | 33 | 620862831.1 | 548086919.7 | 541798633.4 | 494276218.7 | 501825558 | 478222983.9 | 538223821.6 | 442408210.9 | 9.47899 | 8.21883 | Tnr | 21960 | 1.05866 | 0.941343898 | 0.88919 | 0.11353 | 0.678142 |
| 1039728359 | PREDICTED: tensin-1 isoform X5 [Mus musculus] | 0 | 20.096 | 3 | 4 | 7 | 3 | 1901 | 203.1 | 8.44 | 88 | 4 | 972265.813 | 1379932.336 | 1835135.311 | 2801490.891 | 2107661.675 | 1336320.416 | 1413549.625 | 1247304.879 | 45.0017 | 25.7855 | Tns1 | 21961 | 1.06751 | 0.932487349 | 0.87351 | 0.6331 | 0.912586 |
| 112734855 | DNA topoisomerase 1 [Mus musculus] | 0 | 72.732 | 23 | 14 | 66 | 14 | 767 | 90.8 | 9.33 | 615 | 14 | 27676743.54 | 16995227.49 | 27392893.08 | 26646103.85 | 26749300.62 | 14328387.26 | 22692444.63 | 25539305.99 | 20.8288 | 25.0689 | Top1 | 21969 | 1.05 | 0.949997292 | 0.90476 | 0.55893 | 0.895889 |
| 34328148 | DNA topoisomerase 2-beta [Mus musculus] | 0 | 50.663 | 10 | 15 | 51 | 15 | 1612 | 181.8 | 8.29 | 219 | 15 | 22654484.38 | 23875918.4 | 25403308.98 | 27944302.06 | 31746182.56 | 23221734.17 | 25429466.75 | 20968962.24 | 9.13045 | 18.3171 | Top2b | 21974 | 0.9926 | 1.007395645 | 1.0149 | 0.89029 | 0.984401 |
| 148665014 | topoisomerase (DNA) III beta, isoform CRA_a, partial [Mus musculus] | 0 | 13.622 | 7 | 4 | 6 | 4 | 867 | 97.4 | 7.99 | 15 | 4 | 4761117.129 | 1737803.877 | 1866226.975 | 4093514.206 | 2548606.35 | 2654764.175 | 3284941.375 | 1802513.141 | 49.473 | 23.6199 | Top3b | 21976 | 1.09529 | 0.904708307 | 0.826 | 0.53713 | 0.886555 |
| 568979504 | PREDICTED: apoptosis-stimulating of p53 protein 1 isoform X1 [Mus musculus] | 0 | 38.158 | 10 | 8 | 39 | 6 | 1122 | 122.8 | 6.67 | 359 | 8 | 11147464.96 | 7528731.465 | 11371084.21 | 12830641.12 | 8905969.058 | 9470973.935 | 9097510.75 | 10335763.41 | 21.03 | 6.70535 | Ppp1r13b | 21981 | 1.06281 | 0.937193934 | 0.88181 | 0.32079 | 0.771047 |
| 1039764113 | PREDICTED: tumor protein D52 isoform X4 [Mus musculus] | 0 | 18.612 | 13 | 4 | 7 | 4 | 316 | 34.6 | 5.55 | 19 | 4 | 5089865.655 | 9072677.104 | 10064840.88 | 1485738.542 | 10914134.87 | 1456586.915 | 10309439.75 | 1339545.403 | 61.2039 | 88.6852 | Tpd52 | 21985 | 1.03405 | 0.96594975 | 0.93414 | 0.90242 | 0.98561 |
| 568965335 | PREDICTED: tumor protein D53 isoform X1 [Mus musculus] | 0 | 7.747 | 7 | 2 | 9 | 2 | 342 | 37.2 | 8.21 | 114 | 2 | 1148100.994 | 3686646.464 | 2170724.516 | 4457044.994 | 3430607.064 | 2333447.511 | 2986475.625 | 4126996.75 | 51.9142 | 23.4341 | Tpd52l1 | 21987 | 0.94186 | 1.058135063 | 1.12345 | 0.68624 | 0.92798 |
| 124246 | RecName: Full=IgE-binding protein | 0 | 43.717 | 20 | 8 | 24 | 6 | 557 | 62.7 | 9.31 | 367 | 8 | 8930562.343 | 4609417.263 | 8154300.553 | 4840477.107 | 4386290.904 | 11524054.53 | 10354879.25 | 3565656.212 | 33.5965 | 54.4731 | Tpi1 | 21991 | 0.94152 | 1.058477536 | 1.12422 | 0.73426 | 0.943908 |
| 256000796 | tropomyosin alpha-1 chain isoform Tpm1.12br [Mus musculus] | 0 | 71.545 | 40 | 12 | 158 | 9 | 245 | 28.3 | 4.77 | 3887 | 12 | 70894537.12 | 99187855.01 | 127795955.6 | 96942237.03 | 107289285.8 | 95912230.99 | 124747314.9 | 114924206.1 | 23.5649 | 11.0061 | Tpm1 | 22003 | 0.94264 | 1.057362802 | 1.12171 | 0.39548 | 0.824613 |
| 6678419 | tripeptidyl-peptidase 2 isoform 1 [Mus musculus] | 0 | 31.721 | 8 | 9 | 29 | 9 | 1262 | 139.8 | 6.58 | 170 | 9 | 11429710.76 | 12835022.25 | 17296269.66 | 9476370.795 | 18366735.2 | 11933436.98 | 17100929.88 | 8506153.726 | 26.0469 | 32.8221 | Tpp2 | 22019 | 0.95446 | 1.045536483 | 1.09542 | 0.68232 | 0.927174 |
| 114842405 | TNF receptor-associated factor 3 isoform a [Mus musculus] | 0 | 356.139 | 68 | 38 | 627 | 38 | 567 | 64.3 | 8.05 | 9462 | 38 | 1013289793 | 1220789698 | 1207530103 | 1570486091 | 1506152797 | 1918827636 | 1389815689 | 1759030022 | 18.509 | 14.5853 | Traf3 | 22031 | 0.8652 | 1.134795529 | 1.31159 | 0.05775 | 0.599194 |
| 148690398 | tuberous sclerosis 2, isoform CRA_e [Mus musculus] | 0 | 17.878 | 3 | 3 | 8 | 3 | 1814 | 202.1 | 7.05 | 135 | 3 | 6020203.598 | 2957291.088 | 4891978.006 | 2735864.668 | 6386988.499 | 4073115.881 | 4979552.25 | 3240069.746 | 38.0126 | 28.8484 | Tsc2 | 22084 | 0.94121 | 1.058789437 | 1.12492 | 0.63496 | 0.912586 |
| 11230780 | tumor susceptibility gene 101 protein [Mus musculus] | 0 | 19.109 | 17 | 5 | 10 | 5 | 391 | 44.1 | 6.71 | 71 | 5 | 17369202.63 | 6593633.213 | 9074780.846 | 6531978.992 | 13776192.92 | 4607836.839 | 14497982.5 | 7067474.066 | 51.7905 | 49.1082 | Tsg101 | 22088 | 0.99522 | 1.004777352 | 1.0096 | 0.9795 | 0.997464 |
| 31981945 | 60S ribosomal protein L13a [Mus musculus] | 0 | 37.4 | 33 | 8 | 171 | 8 | 203 | 23.4 | 11.02 | 1383 | 8 | 159614001 | 222124742 | 183850661.8 | 284075540.7 | 202716238.7 | 250414716.9 | 213083210.4 | 249479777.7 | 25.5455 | 10.7657 | Rpl13a | 22121 | 0.9626 | 1.037402592 | 1.07771 | 0.59963 | 0.905186 |
| 19705424 | 26S proteasome non-ATPase regulatory subunit 3 [Mus musculus] | 0 | 9.23 | 4 | 2 | 7 | 2 | 530 | 60.7 | 8.44 | 152 | 2 | 2391531.052 | 983410.6432 | 1662156.485 | 1022173.042 | 1984495.924 | 950322.9998 | 1980602 | 876308.0614 | 43.7132 | 42.6859 | Psmd3 | 22123 | 1.02258 | 0.977424502 | 0.95585 | 0.88743 | 0.984249 |
| 568997436 | PREDICTED: E3 ubiquitin-protein ligase TTC3 isoform X1 [Mus musculus] | 0 | 25.939 | 4 | 5 | 15 | 5 | 1998 | 225.8 | 6.32 | 98 | 5 | 11116713.42 | 3806418.99 | 7338597.118 | 7418929.404 | 9153925.901 | 4887268.271 | 10560144.88 | 6176359.524 | 40.2279 | 33.996 | Ttc3 | 22129 | 0.98185 | 1.018145376 | 1.03696 | 0.8946 | 0.984401 |
| 6755901 | tubulin alpha-1A chain [Mus musculus] | 0 | 496.846 | 89 | 33 | 798 | 7 | 451 | 50.1 | 5.06 | 17032 | 33 | 2457604729 | 1983426017 | 2417164814 | 1806974722 | 3194860522 | 1738130673 | 2579263883 | 1619743158 | 14.8473 | 32.544 | Tuba1a | 22142 | 0.97377 | 1.026230462 | 1.05387 | 0.78281 | 0.952792 |
| 4507729 | tubulin beta-2A chain isoform 1 [Homo sapiens] | 0 | 543.319 | 81 | 32 | 1309 | 3 | 445 | 49.9 | 4.89 | 29666 | 32 | 215287991.2 | 273647193.8 | 215713526.5 | 377855032.6 | 364731046.5 | 221179570.6 | 222279437.6 | 328429342.6 | 28.2904 | 25.8984 | Tubb2a | 22151 | 0.97561 | 1.024386052 | 1.04999 | 0.80739 | 0.961747 |
| 12963615 | tubulin beta-3 chain [Mus musculus] | 0 | 377.659 | 71 | 26 | 881 | 10 | 450 | 50.4 | 4.93 | 16643 | 26 | 943741556.8 | 596422138.3 | 801278783.8 | 652202630.3 | 755168568.9 | 412508906.5 | 845594883.1 | 369984145.4 | 20.8866 | 40.2325 | Tubb3 | 22152 | 1.11352 | 0.886479492 | 0.79611 | 0.32724 | 0.773144 |
| 12851187 | unnamed protein product [Mus musculus] | 0 | 516.508 | 83 | 32 | 1153 | 3 | 444 | 49.5 | 4.91 | 25948 | 32 | 1164557239 | 689475498.9 | 1243725598 | 465384384 | 1252839132 | 769273418.8 | 1242303500 | 707505249.6 | 42.0545 | 29.7173 | Tubb4a | 22153 | 0.94575 | 1.05425018 | 1.11472 | 0.68318 | 0.927189 |
| 7106439 | tubulin beta-5 chain [Mus musculus] | 0 | 517.148 | 80 | 30 | 1276 | 5 | 444 | 49.6 | 4.89 | 29623 | 30 | 589779209.2 | 604216993.5 | 597365546.4 | 673239829.4 | 853880277.7 | 482288839.2 | 366633064 | 572117540.3 | 6.2507 | 36.5509 | Tubb5 | 22154 | 1.04002 | 0.959978689 | 0.92304 | 0.66946 | 0.922517 |
| 529280788 | Chain A, Murine Epidermal Fatty Acid-binding Protein (fabp5), Apo Form, Poly-his Tag-mediated Crystal Packing | 0.001 | 6.456 | 22 | 2 | 4 | 2 | 155 | 17.3 | 7.5 | 14 | 2 | 1759962.977 | 2381490.185 | 3869666.903 | 2773576.311 | 3353376.597 | 2779887.233 | 4120177.25 | 1910916.431 | 32.8865 | 30.6563 | U2af2 | 22185 | 0.93988 | 1.060118432 | 1.12793 | 0.61113 | 0.905186 |
| 444189294 | ubiquitin-like modifier-activating enzyme 1 isoform 1 [Mus musculus] | 0 | 53.156 | 14 | 12 | 28 | 12 | 1118 | 124.2 | 5.99 | 295 | 12 | 20992780.03 | 23324717.71 | 22948111.52 | 30766447.01 | 28234158.98 | 27065008.91 | 23851271.75 | 21857895.09 | 17.5276 | 11.5822 | Uba1 | 22201 | 0.98505 | 1.014953138 | 1.03036 | 0.78424 | 0.952792 |
| 148705826 | ubiquitin carboxy-terminal hydrolase L1, isoform CRA_b, partial [Mus musculus] | 0 | 38.169 | 28 | 5 | 20 | 5 | 233 | 25.8 | 5.35 | 222 | 5 | 22736550.26 | 32477251.69 | 30206483.98 | 30775208.46 | 34327387.7 | 36056631.73 | 34946876.38 | 27597911.76 | 14.8625 | 11.506 | Uchl1 | 22223 | 0.93283 | 1.06716853 | 1.14401 | 0.19705 | 0.715071 |
| 7305619 | ubiquitin carboxyl-terminal hydrolase 5 isoform 1 [Mus musculus] | 0 | 45.332 | 16 | 10 | 32 | 10 | 858 | 95.8 | 5.01 | 494 | 10 | 9967657.259 | 3955539.82 | 11576176.87 | 9326292.909 | 12860516.09 | 7117286.141 | 9734788.938 | 7034277.901 | 37.9676 | 29.9512 | Usp5 | 22225 | 0.97316 | 1.026842729 | 1.05517 | 0.83068 | 0.968871 |
| 1246787 | UDP-galactose ceramide galactosyltransferase [Mus musculus] | 0 | 78.833 | 28 | 12 | 44 | 12 | 541 | 61.1 | 9.36 | 726 | 12 | 75897627.06 | 66657688.33 | 77247328.87 | 61887901.8 | 81782420.87 | 62108792.97 | 88572805.8 | 67822871.94 | 10.4852 | 16.2761 | Ugt8a | 22239 | 0.96805 | 1.031953722 | 1.06602 | 0.53899 | 0.886555 |
| 612407885 | dihydropyrimidinase-related protein 3 CRMP4b [Mus musculus] | 0 | 171.728 | 47 | 20 | 136 | 17 | 683 | 73.8 | 6.46 | 2562 | 20 | 211523183.7 | 205151286.1 | 213584365.6 | 240563226.4 | 232908916.2 | 161922176.2 | 163847746.4 | 142750775.5 | 7.19116 | 22.5437 | Dpysl3 | 22240 | 1.10774 | 0.892261239 | 0.80548 | 0.09346 | 0.662454 |
| 569008676 | PREDICTED: probable ubiquitin carboxyl-terminal hydrolase FAF-X isoform X1 [Mus musculus] | 0 | 132.759 | 14 | 28 | 107 | 27 | 2575 | 292.4 | 5.83 | 1245 | 28 | 46372130.54 | 108602301.3 | 73934115.25 | 90829979.13 | 66535039.49 | 106614691.4 | 62394460.59 | 108046377.1 | 33.1225 | 28.8868 | Usp9x | 22284 | 0.96404 | 1.035958083 | 1.0746 | 0.75358 | 0.95243 |
| 568965341 | PREDICTED: utrophin isoform X1 [Mus musculus] | 0 | 24.404 | 3 | 8 | 18 | 7 | 3435 | 393.1 | 5.38 | 172 | 8 | 27931627.6 | 18457140.17 | 26276116.9 | 13429161.53 | 17596742.57 | 32113236.25 | 28263468.72 | 30606168.07 | 31.5765 | 24.165 | Utrn | 22288 | 0.8845 | 1.115503912 | 1.26117 | 0.27889 | 0.745606 |
| 255069795 | valine--tRNA ligase [Mus musculus] | 0.001 | 6.709 | 2 | 3 | 5 | 3 | 1263 | 140.1 | 7.77 | 0 | 3 | 1700911.741 | 4124735.842 | 2769934.562 | 2856727.903 | 3140485.641 | 3723332.115 | 3696477.563 | 3953468.845 | 34.6418 | 9.51223 | Vars | 22321 | 0.8821 | 1.117902078 | 1.26732 | 0.19522 | 0.715071 |
| 568987488 | PREDICTED: vinculin isoform X1 [Mus musculus] | 0 | 81.761 | 21 | 18 | 66 | 18 | 1134 | 123.8 | 5.71 | 613 | 18 | 53046310.12 | 88988720.64 | 76420903.54 | 99318325.84 | 76648104.22 | 85587746.54 | 77652020.97 | 104701890 | 25.0918 | 15.0903 | Vcl | 22330 | 0.95952 | 1.040484538 | 1.08439 | 0.59358 | 0.902243 |
| 6755963 | voltage-dependent anion-selective channel protein 1 [Mus musculus] | 0 | 147.93 | 70 | 16 | 184 | 15 | 283 | 30.7 | 8.54 | 5944 | 16 | 701225246 | 681715032 | 524212521.3 | 707836919.5 | 1021339985 | 889915781.8 | 622930837.3 | 946798296.1 | 13.3179 | 19.9297 | Vdac1 | 22333 | 0.85794 | 1.142060169 | 1.33117 | 0.06715 | 0.601634 |
| 6755965 | voltage-dependent anion-selective channel protein 2 [Mus musculus] | 0 | 79.711 | 41 | 11 | 157 | 11 | 295 | 31.7 | 7.49 | 6709 | 11 | 836074509.6 | 825292563.1 | 712580703.9 | 811473669.9 | 1052758935 | 1015509362 | 877943437.4 | 836001588.9 | 7.12623 | 11.0852 | Vdac2 | 22334 | 0.91435 | 1.085652003 | 1.18735 | 0.04631 | 0.591577 |
| 312222784 | voltage-dependent anion-selective channel protein 3 isoform 1 [Mus musculus] | 0 | 153.748 | 58 | 14 | 143 | 13 | 284 | 30.9 | 8.79 | 3728 | 14 | 853422823.2 | 739680883.4 | 661348910.2 | 710649715.7 | 983656567.6 | 902872709.2 | 795096868 | 731404716.7 | 10.9888 | 13.1374 | Vdac3 | 22335 | 0.92977 | 1.070228783 | 1.15107 | 0.15716 | 0.700461 |
| 6755971 | protein lin-7 homolog B [Mus musculus] | 0 | 44.523 | 51 | 8 | 74 | 5 | 207 | 22.9 | 8.68 | 982 | 8 | 23331267.05 | 18575404.21 | 37108758.68 | 22749992.45 | 21478964.14 | 21425785.31 | 22612612 | 21418530.86 | 31.6868 | 2.69798 | Lin7b | 22342 | 1.07859 | 0.921412683 | 0.85428 | 0.39434 | 0.824613 |
| 6755973 | protein lin-7 homolog C [Mus musculus] | 0 | 47.343 | 54 | 8 | 88 | 5 | 197 | 21.8 | 8.43 | 1091 | 8 | 57113537.7 | 58898528.84 | 53324758.47 | 55055871.69 | 54060565.73 | 66880580.7 | 51143913.75 | 61774887.9 | 4.32392 | 12.2855 | Lin7c | 22343 | 0.97934 | 1.020659458 | 1.04219 | 0.55536 | 0.895107 |
| 12832989 | unnamed protein product [Mus musculus] | 0.001 | 6.942 | 4 | 3 | 13 | 2 | 586 | 69.4 | 6.2 | 70 | 3 | 2581197.024 | 1681730.389 | 2216975.676 | 1992840.263 | 2799908.929 | 2556702.823 | 2289071.25 | 2144635.189 | 17.8806 | 11.8636 | Ezr | 22350 | 0.92786 | 1.072144248 | 1.15551 | 0.21669 | 0.716066 |
| 568928766 | PREDICTED: nuclease-sensitive element-binding protein 1 isoform X1 [Mus musculus] | 0 | 33.645 | 39 | 5 | 25 | 5 | 326 | 36.2 | 9.88 | 446 | 5 | 225293.8866 | 5078070.72 | 4446046.592 | 13690016.22 | 5811734.11 | 5711023.524 | 14150420.44 | 18972326.11 | 96.3695 | 58.5848 | Ybx1 | 22608 | 0.68853 | 1.311465052 | 1.90472 | 0.2657 | 0.745606 |
| 5803225 | 14-3-3 protein epsilon [Homo sapiens] | 0 | 146.903 | 68 | 17 | 276 | 15 | 255 | 29.2 | 4.74 | 5646 | 17 | 481014414.9 | 510700565 | 567478124 | 551624209.3 | 609384290.1 | 628004574.3 | 589005337.3 | 575162295.4 | 7.43909 | 3.85792 | Ywhae | 22627 | 0.93557 | 1.064431538 | 1.13774 | 0.01885 | 0.492977 |
| 74215924 | unnamed protein product [Mus musculus] | 0 | 137.016 | 51 | 14 | 225 | 10 | 310 | 35.1 | 5.49 | 5104 | 14 | 364432890.5 | 264199687.3 | 382680472.2 | 318406171.2 | 381868229.4 | 265158227.7 | 333365864 | 292182872.1 | 15.9189 | 16.0006 | Ywhag | 22628 | 1.02196 | 0.978040906 | 0.95703 | 0.71063 | 0.935294 |
| 1526541 | 14-3-3 eta [Mus musculus] | 0 | 139.648 | 64 | 16 | 262 | 13 | 246 | 28.2 | 4.86 | 4619 | 16 | 165277901 | 164516497.7 | 166062519.8 | 171922484.4 | 159415638.9 | 170305121.9 | 148055101.5 | 150712599 | 2.02337 | 6.3893 | Ywhah | 22629 | 1.03031 | 0.96968918 | 0.94116 | 0.11305 | 0.678142 |
| 55153870 | Ywhaq protein, partial [Mus musculus] | 0 | 149.297 | 43 | 16 | 267 | 13 | 284 | 32.2 | 5.94 | 8476 | 16 | 230996458.8 | 214661702.8 | 285187895.4 | 236093319.4 | 297834069.5 | 243371382.9 | 261802672.4 | 228157052.7 | 12.5662 | 11.6495 | Ywhaq | 22630 | 0.96786 | 1.032143364 | 1.06642 | 0.48062 | 0.86754 |
| 6756041 | 14-3-3 protein zeta/delta isoform 1 [Mus musculus] | 0 | 132.146 | 60 | 15 | 348 | 13 | 245 | 27.8 | 4.79 | 9764 | 15 | 550778350.7 | 580702075.2 | 593882580.7 | 647444558 | 625665912.1 | 618536005.5 | 617948298.4 | 551539559 | 6.81206 | 5.76147 | Ywhaz | 22631 | 0.99146 | 1.008541154 | 1.01723 | 0.7146 | 0.936732 |
| 194328715 | BUB3-interacting and GLEBS motif-containing protein ZNF207 isoform 1 [Mus musculus] | 0.001 | 4.444 | 5 | 2 | 7 | 2 | 495 | 52.8 | 9.1 | 23 | 2 | 5543287.661 | 9104778.657 | 9074024.584 | 12685730.6 | 9172972.504 | 8908244.261 | 8693708.75 | 11523460.22 | 32.0366 | 13.7232 | Zfp207 | 22680 | 0.97469 | 1.025306655 | 1.05193 | 0.77753 | 0.952792 |
| 31542413 | coronin-1C [Mus musculus] | 0 | 44.646 | 21 | 7 | 68 | 6 | 474 | 53.1 | 7.08 | 1260 | 7 | 91783775.51 | 41176132.06 | 81068133.49 | 60220080.77 | 80935213.53 | 141307325 | 82998018 | 67009696.3 | 32.7789 | 35.3931 | Coro1c | 23790 | 0.84841 | 1.151589138 | 1.35735 | 0.26512 | 0.745606 |
| 568906339 | PREDICTED: disintegrin and metalloproteinase domain-containing protein 23 isoform X1 [Mus musculus] | 0 | 27.152 | 10 | 6 | 21 | 6 | 829 | 91.6 | 7.62 | 94 | 6 | 15409229.99 | 18675840.98 | 20124762.02 | 19239695.48 | 16281612.12 | 7370402.187 | 17987520.5 | 9746975.286 | 11.2028 | 39.6469 | Adam23 | 23792 | 1.17674 | 0.82326403 | 0.69962 | 0.09137 | 0.662454 |
| 568967554 | PREDICTED: adenomatous polyposis coli protein 2 isoform X1 [Mus musculus] | 0 | 31.696 | 4 | 9 | 28 | 9 | 2317 | 247.5 | 8.94 | 323 | 9 | 27658146.79 | 32405562.62 | 33315976.56 | 20264651.35 | 28041087.92 | 39933582.91 | 26097644.75 | 48013743.98 | 21.0151 | 29.0883 | Apc2 | 23805 | 0.88878 | 1.111217605 | 1.25027 | 0.27839 | 0.745606 |
| 6753178 | barrier-to-autointegration factor [Mus musculus] | 0 | 52.757 | 51 | 5 | 36 | 5 | 89 | 10.1 | 6.09 | 791 | 5 | 127980147.2 | 95089342.36 | 143104481.3 | 155848983.8 | 114667451.2 | 179594946.4 | 180489447.2 | 158437277.9 | 20.0874 | 19.4699 | Banf1 | 23825 | 0.90377 | 1.096230095 | 1.21295 | 0.21864 | 0.716066 |
| 31981400 | phenylalanine--tRNA ligase beta subunit [Mus musculus] | 0 | 49.139 | 20 | 12 | 46 | 12 | 589 | 65.7 | 7.12 | 430 | 12 | 60572410.9 | 50688009.28 | 57740462.95 | 58305372.96 | 67242057.86 | 55882048.07 | 57353771.81 | 75533137.91 | 7.51661 | 14.3653 | Farsb | 23874 | 0.94061 | 1.059391131 | 1.12628 | 0.20661 | 0.715071 |
| 28201958 | rho-related GTP-binding protein RhoF precursor [Mus musculus] | 0 | 13.904 | 18 | 3 | 13 | 3 | 211 | 23.6 | 8.43 | 68 | 3 | 48671825 | 47634638.31 | 56791915.59 | 46541427.38 | 58755888.93 | 59834876.17 | 61098157.5 | 49832124.19 | 9.35626 | 8.92698 | Rhof | 23912 | 0.93037 | 1.069627135 | 1.14968 | 0.07454 | 0.624381 |
| 12861758 | unnamed protein product [Mus musculus] | 0 | 9.732 | 12 | 2 | 11 | 2 | 199 | 21.4 | 9.88 | 129 | 2 | 2795090.915 | 3221320.272 | 3048732.613 | 6132667.985 | 4432443.635 | 263265.9986 | 3250762.5 | 1939124.69 | 41.1979 | 72.4285 | Ly6h | 23934 | 1.21178 | 0.788217979 | 0.65046 | 0.30671 | 0.761857 |
| 7106349 | ly-6/neurotoxin-like protein 1 precursor [Mus musculus] | 0 | 16.069 | 25 | 3 | 13 | 3 | 116 | 12.8 | 8.19 | 432 | 3 | 34205236 | 17611061 | 15357678.6 | 15560607.02 | 27173209.4 | 13811815.23 | 17189407.75 | 13722879.19 | 43.8591 | 35.2798 | Lynx1 | 23936 | 1.07008 | 0.929915681 | 0.86901 | 0.64183 | 0.915705 |
| 83816907 | dnaJ homolog subfamily B member 6 isoform a [Mus musculus] | 0 | 82.501 | 27 | 10 | 119 | 9 | 365 | 39.8 | 9.36 | 1732 | 10 | 115028017.9 | 31082373.26 | 82497251.74 | 66975678.56 | 107088373 | 26046216.66 | 94802554.31 | 22001331.29 | 47.1814 | 71.5754 | Dnajb6 | 23950 | 1.08367 | 0.916328098 | 0.84558 | 0.70129 | 0.931521 |
| 7657413 | teneurin-1 [Mus musculus] | 0 | 65.797 | 8 | 15 | 45 | 15 | 2731 | 305.6 | 6.37 | 354 | 15 | 31137872.14 | 32405369.35 | 31602302.63 | 21453993.58 | 31900669.36 | 32087002.37 | 34527761.63 | 18542578.72 | 17.6921 | 24.7654 | Tenm1 | 23963 | 0.99804 | 1.001962164 | 1.00393 | 0.98028 | 0.997464 |
| 568973887 | PREDICTED: teneurin-2 isoform X1 [Mus musculus] | 0 | 215.16 | 21 | 42 | 179 | 41 | 2774 | 307.4 | 6.68 | 2573 | 42 | 109313556 | 125955510.7 | 115607506.1 | 112422335.2 | 141536917 | 150000022.8 | 134155578.8 | 128909010.2 | 6.23886 | 6.61346 | Tenm2 | 23964 | 0.9103 | 1.089697005 | 1.19707 | 0.00789 | 0.4058 |
| 568954791 | PREDICTED: teneurin-3 isoform X1 [Mus musculus] | 0 | 29.112 | 3 | 7 | 14 | 7 | 2722 | 303.6 | 6.54 | 137 | 7 | 9593581.267 | 12732899.92 | 12518115.71 | 9215449.848 | 13134485.04 | 13190519.83 | 13402295.25 | 13326452.46 | 16.9595 | 0.92543 | Tenm3 | 23965 | 0.90739 | 1.092609968 | 1.20412 | 0.05315 | 0.595816 |
| 124248484 | teneurin-4 isoform 2 [Mus musculus] | 0 | 138.229 | 13 | 28 | 102 | 27 | 2796 | 311 | 6.49 | 1040 | 28 | 72058093.27 | 59932272.24 | 69216144.37 | 55811927.95 | 58028662.26 | 86281438.32 | 57756787.13 | 80361225.75 | 11.9025 | 21.0733 | Tenm4 | 23966 | 0.9529 | 1.047103231 | 1.09886 | 0.47637 | 0.863085 |
| 148690599 | protein kinase C and casein kinase substrate in neurons 1 [Mus musculus] | 0 | 35.432 | 21 | 7 | 14 | 7 | 441 | 50.5 | 5.24 | 299 | 7 | 11141145.66 | 26334207.85 | 8524930.892 | 11341547.41 | 6385610.719 | 10552569.59 | 7367435.813 | 7982468.091 | 56.5132 | 22.0473 | Pacsin1 | 23969 | 1.27952 | 0.72047561 | 0.56308 | 0.18172 | 0.715071 |
| 148672882 | mCG7191 [Mus musculus] | 0 | 22.717 | 28 | 3 | 8 | 3 | 187 | 20.8 | 6.54 | 154 | 3 | 9675146.311 | 7799617.147 | 9578646.142 | 7876838.35 | 13460044.55 | 8072189.046 | 14124027.5 | 8072498.997 | 11.8398 | 30.3084 | Pebp1 | 23980 | 0.88814 | 1.111856383 | 1.25189 | 0.25196 | 0.733806 |
| 6754994 | poly(rC)-binding protein 1 [Mus musculus] | 0 | 40.063 | 29 | 6 | 114 | 4 | 356 | 37.5 | 7.09 | 2243 | 6 | 43241260.94 | 29116527.46 | 39606781.14 | 32189527.43 | 45234625.95 | 27985129.75 | 38597200 | 25312895.77 | 18.0782 | 27.0861 | Pcbp1 | 23983 | 1.02497 | 0.975027919 | 0.95127 | 0.76732 | 0.952792 |
| 160333193 | enoyl-CoA delta isomerase 2, mitochondrial isoform a [Mus musculus] | 0 | 41.912 | 29 | 8 | 20 | 8 | 391 | 43.2 | 8.92 | 484 | 8 | 28388948.96 | 15798290.44 | 19156420.62 | 19338782.36 | 16311191.92 | 24464396.98 | 14836389 | 28002628.63 | 26.1089 | 30.3786 | Eci2 | 23986 | 0.99439 | 1.005605416 | 1.01127 | 0.95722 | 0.994906 |
| 12963653 | peptidyl-prolyl cis-trans isomerase NIMA-interacting 1 [Mus musculus] | 0 | 89.743 | 70 | 7 | 50 | 7 | 165 | 18.4 | 8.79 | 1976 | 7 | 52296211.25 | 26196068.63 | 39613338.44 | 28723710.11 | 43004548.39 | 23748749.06 | 47415407.25 | 14312949.95 | 32.4511 | 48.8826 | Pin1 | 23988 | 1.06664 | 0.93335655 | 0.87504 | 0.65801 | 0.919309 |
| 755499566 | DICTED: interferon-inducible double-stranded RNA-dependent protein kinase activator A isoform X1 [Mus muscu | 0 | 46.554 | 24 | 7 | 46 | 7 | 365 | 39.9 | 9.51 | 715 | 7 | 64663040.45 | 69901410.8 | 48712764.3 | 65440140.65 | 47271065.94 | 67101860 | 49714548.5 | 65131976.21 | 14.908 | 17.8966 | Prkra | 23992 | 1.0408 | 0.959204009 | 0.92161 | 0.50712 | 0.88031 |
| 6755224 | twinfilin-2 [Mus musculus] | 0 | 43.781 | 34 | 7 | 25 | 7 | 349 | 39.4 | 6.8 | 557 | 7 | 56263694.31 | 31587850.27 | 35969654.87 | 32331817.86 | 60881214.19 | 28933192.7 | 41965670 | 54146648.28 | 29.8223 | 30.2839 | Twf2 | 23999 | 0.91296 | 1.087037331 | 1.19067 | 0.44623 | 0.852048 |
| 755495684 | PREDICTED: regulator of G-protein signaling 7 isoform X2 [Mus musculus] | 0 | 42.388 | 18 | 8 | 31 | 8 | 488 | 57 | 8.09 | 168 | 8 | 22832692.25 | 23020018.79 | 21006531.63 | 21036297.64 | 20575674.77 | 19819769.14 | 18164251.13 | 21069549.32 | 5.0175 | 6.38268 | Rgs7 | 24012 | 1.04934 | 0.950656281 | 0.90595 | 0.04933 | 0.592616 |
| 148672573 | septin 3, isoform CRA_c, partial [Mus musculus] | 0 | 81.972 | 41 | 12 | 52 | 12 | 397 | 45 | 9.26 | 233 | 12 | 49588574.42 | 33128745.35 | 38805039.17 | 42676150.82 | 33690405.12 | 57406916.8 | 42618001.38 | 38035602.92 | 16.8388 | 24.0159 | 43711 | 24050 | 0.97752 | 1.022480813 | 1.046 | 0.77126 | 0.952792 |
| 3004994 | G protein-coupled receptor kinase 6 [Mus musculus] | 0 | 18.324 | 10 | 4 | 9 | 4 | 589 | 67 | 8.59 | 222 | 4 | 7123994.714 | 5695243.515 | 8756544.374 | 6561447.519 | 9071364.062 | 5707733.827 | 11635061.75 | 5788883.984 | 18.336 | 35.4927 | Grk6 | 26385 | 0.93262 | 1.067381423 | 1.1445 | 0.5407 | 0.886555 |
| 6678794 | dual specificity mitogen-activated protein kinase kinase 1 [Mus musculus] | 0 | 28.655 | 12 | 6 | 34 | 6 | 393 | 43.4 | 6.7 | 332 | 6 | 23194099.8 | 26999306.24 | 23774263.12 | 31735370.7 | 35456333.69 | 24028907.18 | 25065912 | 24041819.93 | 14.8172 | 20.4801 | Map2k1 | 26395 | 0.98651 | 1.013485706 | 1.02734 | 0.83877 | 0.969831 |
| 939699106 | dual specificity mitogen-activated protein kinase kinase 4 isoform a [Mus musculus] | 0 | 13.029 | 9 | 3 | 10 | 3 | 408 | 45.4 | 8.03 | 122 | 3 | 8049905.273 | 7422310.472 | 8681592.514 | 7445738.866 | 5887482.686 | 4762934.573 | 6826106.75 | 3343138.589 | 7.55238 | 28.8311 | Map2k4 | 26398 | 1.20565 | 0.794352403 | 0.65886 | 0.01566 | 0.461382 |
| 255918231 | dual specificity mitogen-activated protein kinase kinase 7 isoform 3 [Mus musculus] | 0 | 31.273 | 22 | 8 | 37 | 8 | 435 | 49.1 | 9.16 | 190 | 8 | 35420021.3 | 37665528.11 | 30018046.47 | 35818781.14 | 32235754.2 | 34696786.73 | 26299616 | 34413064.17 | 9.47424 | 12.2192 | Map2k7 | 26400 | 1.04231 | 0.957694949 | 0.91882 | 0.31143 | 0.76536 |
| 171846249 | mitogen-activated protein kinase kinase kinase 5 [Mus musculus] | 0 | 28.8 | 7 | 8 | 12 | 8 | 1380 | 154.4 | 5.78 | 61 | 8 | 8128624.661 | 10808427.47 | 10177210.71 | 9022462.722 | 10332405.25 | 11114670.36 | 10138336.25 | 9962726.671 | 12.52 | 4.89114 | Map3k5 | 26408 | 0.95719 | 1.042811304 | 1.08945 | 0.23659 | 0.73104 |
| 6754632 | mitogen-activated protein kinase 1 [Mus musculus] | 0 | 50.814 | 32 | 9 | 28 | 9 | 358 | 41.2 | 6.98 | 293 | 9 | 50444756.65 | 31494998.35 | 53028188.27 | 63807345.99 | 55184732.16 | 44378083.67 | 51983526.25 | 59753283.27 | 27.0499 | 12.2502 | Mapk1 | 26413 | 0.96946 | 1.030541581 | 1.06301 | 0.68928 | 0.930096 |
| 970112998 | mitogen-activated protein kinase 10 isoform 1x [Mus musculus] | 0 | 24.62 | 17 | 7 | 26 | 3 | 478 | 54.1 | 7.27 | 112 | 7 | 12198540.12 | 13355680.79 | 9456288.93 | 13512151.04 | 51589887.58 | 12257740.93 | 14057996.25 | 10426952.32 | 15.4707 | 89.3303 | Mapk10 | 26414 | 0.70911 | 1.290890702 | 1.82044 | 0.35392 | 0.798662 |
| 158854037 | neurobeachin [Mus musculus] | 0 | 77.994 | 7 | 17 | 64 | 13 | 2936 | 326.5 | 6.2 | 767 | 17 | 29465868.51 | 30410754.4 | 30181343.55 | 33629172.96 | 27653715.26 | 23292474.23 | 24209397.5 | 25710437.52 | 5.98037 | 7.5591 | Nbea | 26422 | 1.10163 | 0.898370994 | 0.81549 | 0.00511 | 0.38143 |
| 6755196 | proteasome subunit alpha type-4 [Mus musculus] | 0.003 | 4.068 | 7 | 2 | 3 | 2 | 261 | 29.5 | 7.72 | 14 | 2 | 1052252.596 | 1814529.976 | 1302872.68 | 1544987.705 | 997939.63 | 1763491.007 | 968994.0625 | 1803129.817 | 22.8581 | 33.4125 | Psma4 | 26441 | 1.0161 | 0.98390067 | 0.96831 | 0.87814 | 0.980491 |
| 7106387 | proteasome subunit alpha type-5 [Mus musculus] | 0.001 | 6.121 | 8 | 2 | 7 | 2 | 241 | 26.4 | 4.79 | 23 | 2 | 930950.0197 | 1892536.442 | 1416627.542 | 1441122.522 | 1050112.44 | 1405026.363 | 843774.625 | 1604113.782 | 27.6572 | 27.9472 | Psma5 | 26442 | 1.07353 | 0.926474875 | 0.86302 | 0.48355 | 0.868779 |
| 148696995 | solute carrier family 27 (fatty acid transporter), member 1, isoform CRA_b, partial [Mus musculus] | 0 | 19.869 | 9 | 5 | 21 | 4 | 668 | 73.6 | 8.43 | 76 | 5 | 5121027.871 | 5673043.159 | 6088732.524 | 5662379.941 | 6577758.541 | 7396767.319 | 6081999.75 | 9080639.192 | 7.03919 | 18.0465 | Slc27a1 | 26457 | 0.87245 | 1.127548022 | 1.29239 | 0.05327 | 0.595816 |
| 171846263 | homer protein homolog 1 isoform L [Mus musculus] | 0 | 288.691 | 78 | 29 | 650 | 28 | 366 | 41.4 | 5.53 | 12171 | 29 | 4425924048 | 4210233494 | 4077058440 | 4182851119 | 4335931095 | 4841754508 | 4590360185 | 4032112933 | 3.46452 | 7.79322 | Homer1 | 26556 | 0.97394 | 1.026057348 | 1.05351 | 0.27503 | 0.745606 |
| 58372172 | homer protein homolog 2 isoform 1 [Mus musculus] | 0 | 169.139 | 54 | 26 | 328 | 25 | 354 | 40.5 | 6.01 | 5750 | 26 | 559902313.5 | 547281667.6 | 566600862.3 | 635680683 | 667282940.3 | 711732819.2 | 578886905.6 | 694368844.4 | 6.87479 | 8.90196 | Homer2 | 26557 | 0.93091 | 1.069089914 | 1.14844 | 0.0526 | 0.595816 |
| 3834613 | homer-3 [Mus musculus] | 0 | 79.161 | 42 | 16 | 252 | 15 | 358 | 39.9 | 5.29 | 4968 | 16 | 187246008.1 | 223316536.4 | 166467796.9 | 203827763.7 | 196982391 | 229921883.2 | 162426395.2 | 214404934.9 | 12.3855 | 14.4246 | Homer3 | 26558 | 0.98556 | 1.014437454 | 1.0293 | 0.77208 | 0.952792 |
| 568929156 | PREDICTED: neurochondrin isoform X1 [Mus musculus] | 0 | 135.451 | 38 | 19 | 95 | 19 | 712 | 77.3 | 5.54 | 2186 | 19 | 106270687.5 | 77492055.8 | 110829878.9 | 75792987.27 | 128584216.6 | 90393990.79 | 107887885.8 | 74992985.78 | 20.0102 | 22.9587 | Ncdn | 26562 | 0.95924 | 1.040755825 | 1.08497 | 0.6139 | 0.905186 |
| 45597453 | long-chain fatty acid transport protein 4 [Mus musculus] | 0 | 40.766 | 18 | 9 | 30 | 8 | 643 | 72.3 | 8.59 | 278 | 9 | 21429660.28 | 10812436.69 | 16540004.81 | 22032352.03 | 18905242.38 | 14468351.71 | 21104107.13 | 19850036.78 | 29.4347 | 15.5339 | Slc27a4 | 26569 | 0.97579 | 1.024205809 | 1.04961 | 0.77802 | 0.952792 |
| 74216868 | unnamed protein product [Mus musculus] | 0 | 87.782 | 39 | 15 | 56 | 13 | 593 | 64.3 | 7.09 | 1083 | 15 | 53208626.92 | 44455862.16 | 47013037.19 | 41798478.9 | 51891663.83 | 36318240.28 | 43747889.25 | 30912318.41 | 10.4716 | 22.399 | Dpysl4 | 26757 | 1.06757 | 0.93242835 | 0.87341 | 0.29737 | 0.761852 |
| 160707976 | protein piccolo isoform 1 [Mus musculus] | 0 | 2175.92 | 56 | 202 | 3228 | 197 | 5068 | 550.5 | 6.51 | 62985 | 202 | 5304212809 | 5148888459 | 4951545983 | 5483307263 | 4908870485 | 5822825558 | 5107670005 | 5372285523 | 4.33213 | 7.4522 | Pclo | 26875 | 0.99231 | 1.007688838 | 1.0155 | 0.73443 | 0.943908 |
| 25141235 | ATP-dependent RNA helicase DDX3Y [Mus musculus] | 0 | 135.468 | 44 | 25 | 166 | 4 | 658 | 73.4 | 7.53 | 2686 | 25 | 12178713.64 | 11795485.25 | 15843702.19 | 10855393.36 | 21423734.57 | 8180323.933 | 12939794.5 | 10567273.29 | 17.277 | 43.4393 | Ddx3y | 26900 | 0.97651 | 1.023489382 | 1.04811 | 0.84989 | 0.97368 |
| 74188195 | unnamed protein product [Mus musculus] | 0 | 49.83 | 18 | 13 | 53 | 13 | 932 | 95.5 | 7.93 | 221 | 13 | 29589420.44 | 22919477.07 | 21843541.21 | 28331499.66 | 37206441.01 | 42916919.72 | 25905677.13 | 44232319.24 | 15.0287 | 22.2278 | Gprin1 | 26913 | 0.81191 | 1.188093709 | 1.46334 | 0.04142 | 0.585646 |
| 41152517 | core histone macro-H2A.1 isoform 1 [Mus musculus] | 0 | 62.718 | 37 | 8 | 45 | 7 | 372 | 39.7 | 9.8 | 934 | 8 | 32292659.33 | 18482778.38 | 33485303.75 | 25722952.17 | 19447888.41 | 23826635.67 | 36833251.13 | 23667647.15 | 25.1325 | 29.053 | H2afy | 26914 | 1.02904 | 0.970956695 | 0.94355 | 0.77172 | 0.952792 |
| 1039729111 | PREDICTED: mitogen-activated protein kinase kinase kinase kinase 4 isoform X11 [Mus musculus] | 0 | 109.203 | 21 | 21 | 116 | 11 | 1298 | 147.5 | 7.42 | 1472 | 21 | 23457928.52 | 15183319.63 | 48683248.2 | 24601065.23 | 40841818.18 | 18664588.17 | 32405474.88 | 32242022.89 | 51.5523 | 29.5597 | Map4k4 | 26921 | 0.9482 | 1.05179757 | 1.10925 | 0.73285 | 0.943908 |
| 6755004 | apoptosis-inducing factor 1, mitochondrial isoform 1 precursor [Mus musculus] | 0 | 48.678 | 22 | 9 | 23 | 9 | 612 | 66.7 | 9.17 | 335 | 9 | 14367819.88 | 12742383.88 | 17499375.09 | 14734206.07 | 14431020.1 | 17431107.39 | 12567819.75 | 11823438.1 | 13.3148 | 17.7676 | Aifm1 | 26926 | 1.02673 | 0.973265785 | 0.94792 | 0.6448 | 0.915705 |
| 568979761 | ICTED: serine/threonine-protein phosphatase 2A 56 kDa regulatory subunit gamma isoform isoform X1 [Mus mus | 0 | 152.304 | 36 | 18 | 88 | 10 | 606 | 70.1 | 7.97 | 1643 | 18 | 24644000.58 | 14265355.34 | 17802506.25 | 12863587.88 | 22347513.86 | 24774678.44 | 24831277 | 13886863.73 | 30.2478 | 24.1372 | Ppp2r5c | 26931 | 0.89535 | 1.104653998 | 1.23377 | 0.3129 | 0.76536 |
| 33859660 | serine/threonine-protein phosphatase 2A 56 kDa regulatory subunit epsilon isoform [Mus musculus] | 0 | 62.657 | 28 | 10 | 60 | 8 | 467 | 54.7 | 6.95 | 1197 | 10 | 32014826.71 | 23876667.1 | 29835896.76 | 16338528.1 | 25313818.27 | 22067849.42 | 27558843 | 16153046.43 | 27.5087 | 21.7615 | Ppp2r5e | 26932 | 1.0568 | 0.943195323 | 0.8925 | 0.54674 | 0.889301 |
| 568974184 | PREDICTED: myosin phosphatase Rho-interacting protein isoform X1 [Mus musculus] | 0 | 724.334 | 57 | 100 | 726 | 2 | 2423 | 273.9 | 5.8 | 13417 | 100 | 5238021.1 | 4937677.784 | 8463170.501 | 3370882.767 | 8041356.317 | 4749325.504 | 7006955.5 | 5836432.095 | 38.8345 | 22.259 | Mprip | 26936 | 0.92393 | 1.076071086 | 1.16467 | 0.50705 | 0.88031 |
| 6755983 | visinin-like protein 1 [Mus musculus] | 0 | 56.58 | 65 | 12 | 92 | 10 | 191 | 22.1 | 5.15 | 1399 | 12 | 71231601.09 | 85066846.54 | 68773617.63 | 44768821.96 | 114574612.7 | 82204183.48 | 66349748.14 | 76103369.57 | 24.817 | 24.6334 | Vsnl1 | 26950 | 0.88607 | 1.113928953 | 1.25716 | 0.2426 | 0.733806 |
| 4506663 | 60S ribosomal protein L8 [Homo sapiens] | 0 | 93.032 | 42 | 10 | 217 | 10 | 257 | 28 | 11.03 | 2883 | 10 | 78155695.48 | 121275602.4 | 108124783.5 | 188785258.2 | 126424795.1 | 150541365.4 | 143894326.3 | 160929941.9 | 37.6802 | 9.96274 | Rpl8 | 26961 | 0.92074 | 1.079256629 | 1.17216 | 0.41632 | 0.836939 |
| 148684443 | ribosomal protein S3, isoform CRA_e [Mus musculus] | 0 | 95.182 | 78 | 18 | 189 | 18 | 243 | 26.6 | 9.58 | 2062 | 18 | 545003502.2 | 738885353 | 598207729.6 | 781960002 | 598678332.4 | 663683170 | 635286742.9 | 800667812 | 16.8974 | 13.0704 | Rps3 | 27050 | 0.99361 | 1.006388864 | 1.01286 | 0.90854 | 0.98561 |
| 31980969 | protein transport protein Sec23B isoform 1 [Mus musculus] | 0 | 38.31 | 13 | 8 | 16 | 5 | 767 | 86.4 | 6.96 | 237 | 8 | 4008447.499 | 11348499.06 | 4922168.174 | 4099134.854 | 5753710.875 | 2131249.703 | 5360619 | 4935238.173 | 57.8657 | 36.1621 | Sec23b | 27054 | 1.14562 | 0.854380459 | 0.74578 | 0.45614 | 0.860877 |
| 28972638 | mKIAA1121 protein [Mus musculus] | 0 | 110.858 | 21 | 20 | 63 | 14 | 1392 | 156.7 | 5.82 | 572 | 20 | 50978340.49 | 47271157.43 | 50449654.66 | 45377855.5 | 45370785.61 | 53444203.91 | 48168263.25 | 41020015.53 | 5.47886 | 11.0764 | Cadps | 27062 | 1.0159 | 0.984103498 | 0.9687 | 0.62199 | 0.908209 |
| 30410942 | Rpl7a protein, partial [Mus musculus] | 0 | 132.389 | 50 | 18 | 214 | 18 | 270 | 30.5 | 10.55 | 1900 | 18 | 276529821.9 | 397054931.3 | 312502889 | 514975141.9 | 294893567.1 | 379369091.7 | 330706577.3 | 432310184 | 28.2355 | 16.62 | Rpl7a | 27176 | 1.02171 | 0.978292738 | 0.95751 | 0.80193 | 0.959776 |
| 20514282 | SYN3 [Mus musculus] | 0 | 151.238 | 51 | 17 | 143 | 16 | 579 | 63.3 | 9.38 | 1885 | 17 | 197566542.9 | 163571934.1 | 213391638.7 | 185311653.6 | 170616579.2 | 124900293 | 229287209.4 | 179616186 | 11.0626 | 24.2975 | Syn3 | 27204 | 1.03785 | 0.962150558 | 0.92706 | 0.58221 | 0.898781 |
| 12836870 | unnamed protein product [Mus musculus] | 0 | 47.35 | 45 | 8 | 110 | 8 | 166 | 19.4 | 10.14 | 1375 | 8 | 162794100.1 | 205931407.7 | 165333612.2 | 243451511.2 | 209398912.7 | 181014120.1 | 184782553.6 | 204821266.3 | 19.6639 | 7.27507 | Rps11 | 27207 | 0.99839 | 1.00160909 | 1.00322 | 0.97648 | 0.997464 |
| 148684523 | pleckstrin homology domain containing, family B (evectins) member 1, isoform CRA_c [Mus musculus] | 0.001 | 4.745 | 10 | 2 | 6 | 2 | 209 | 23.5 | 5.26 | 34 | 2 | 2259614.925 | 1108286.74 | 420497.4496 | 796686.2171 | 2206465.694 | 174172.2667 | 529103.125 | 1250696.833 | 69.2429 | 86.2745 | Plekhb1 | 27276 | 1.04856 | 0.951444025 | 0.90739 | 0.86516 | 0.978314 |
| 148680841 | mCG1492, isoform CRA_d, partial [Mus musculus] | 0 | 12.501 | 10 | 3 | 10 | 3 | 348 | 38 | 7.62 | 65 | 3 | 5449144.535 | 7204665.881 | 18940013 | 7557685.971 | 14631830.69 | 13764272.74 | 12446312 | 17033287.19 | 63.0444 | 13.3481 | Srr | 27364 | 0.80702 | 1.192978782 | 1.47825 | 0.1978 | 0.715071 |
| 255308899 | 60S ribosomal protein L3 [Mus musculus] | 0 | 166.89 | 46 | 18 | 290 | 18 | 403 | 46.1 | 10.21 | 4788 | 18 | 242917717.9 | 388859643.2 | 261785736.3 | 411226754 | 246873672.8 | 384174764.5 | 321380862.1 | 458759938.6 | 26.3958 | 25.573 | Rpl3 | 27367 | 0.96082 | 1.039175333 | 1.08155 | 0.68457 | 0.927621 |
| 148687432 | transducin (beta)-like 2, isoform CRA_a, partial [Mus musculus] | 0 | 21.221 | 15 | 6 | 17 | 6 | 446 | 49.9 | 8.98 | 77 | 6 | 13189667.1 | 16066193.24 | 13613423.15 | 9015159.923 | 18900085.54 | 17836545.85 | 16364055.5 | 14530258.89 | 22.56 | 11.2116 | Tbl2 | 27368 | 0.86825 | 1.131752927 | 1.30349 | 0.06471 | 0.601634 |
| 148672382 | mCG6805 [Mus musculus] | 0 | 14.246 | 31 | 3 | 46 | 3 | 115 | 13 | 11 | 237 | 3 | 37661481.94 | 75418850.29 | 34806173.35 | 85611712.74 | 44121640.69 | 51154068.85 | 49348581.14 | 33197590.85 | 44.4176 | 18.1665 | Rps26 | 27370 | 1.13536 | 0.864639883 | 0.76156 | 0.34487 | 0.789724 |
| 148672704 | casein kinase 1, epsilon, isoform CRA_b, partial [Mus musculus] | 0 | 95.317 | 41 | 15 | 128 | 4 | 417 | 47.4 | 9.66 | 1764 | 15 | 21663227.97 | 8116933.614 | 25101022.46 | 12650873.25 | 24418843.17 | 7368301.253 | 27078815 | 7080336.518 | 46.5309 | 65.2092 | Csnk1e | 27373 | 1.01188 | 0.988119711 | 0.97652 | 0.95445 | 0.994906 |
| 254826790 | mitochondrial dicarboxylate carrier [Mus musculus] | 0 | 83.344 | 49 | 10 | 48 | 10 | 287 | 31.7 | 9.32 | 1295 | 10 | 176496768.7 | 146990565.4 | 140441934.5 | 118572629.8 | 178618395 | 212338774.8 | 152848074.5 | 151989303.6 | 16.4119 | 16.338 | Slc25a10 | 27376 | 0.91137 | 1.088627837 | 1.19449 | 0.17799 | 0.713031 |
| 23956078 | ATP-binding cassette sub-family F member 2 isoform 1 [Mus musculus] | 0 | 60.631 | 23 | 12 | 35 | 12 | 628 | 71.7 | 7.05 | 491 | 12 | 33322395.53 | 36980209.21 | 38218278.92 | 37194295.13 | 33758574.25 | 37233050.16 | 35746925.06 | 28898781.89 | 5.87516 | 10.7079 | Abcf2 | 27407 | 1.03582 | 0.964180709 | 0.93084 | 0.27698 | 0.745606 |
| 590121980 | angiomotin isoform 1 [Mus musculus] | 0 | 23.697 | 8 | 7 | 20 | 6 | 1126 | 120.8 | 7.5 | 104 | 7 | 9504239.965 | 8253883.082 | 6942916.287 | 10648300.18 | 9434550.326 | 11131738.95 | 10860573.25 | 18694948.81 | 18.0739 | 33.3311 | Amot | 27494 | 0.82716 | 1.172835766 | 1.4179 | 0.14966 | 0.693437 |
| 568995527 | PREDICTED: probable palmitoyltransferase ZDHHC8 isoform X1 [Mus musculus] | 0 | 39.503 | 17 | 10 | 36 | 10 | 775 | 83.5 | 9.35 | 303 | 10 | 71639822.71 | 72753460 | 51962915.73 | 56152156.44 | 49339640.62 | 46464046.6 | 65923460.5 | 33036264.2 | 16.8249 | 27.7426 | Zdhhc8 | 27801 | 1.1291 | 0.870895176 | 0.77131 | 0.14391 | 0.693437 |
| 23271707 | Eif3b protein [Mus musculus] | 0 | 13.929 | 6 | 5 | 10 | 5 | 970 | 108.9 | 5.54 | 46 | 5 | 649957.5454 | 5114733.398 | 2171108.313 | 3332950.277 | 2791028.745 | 5513311.218 | 6231633 | 2843035.907 | 66.9087 | 41.1622 | Eif3b | 27979 | 0.78671 | 1.213289265 | 1.54223 | 0.28421 | 0.751585 |
| 148681465 | EF hand domain containing 2 [Mus musculus] | 0 | 9.618 | 19 | 4 | 14 | 4 | 242 | 26.9 | 5.14 | 119 | 4 | 9935436.115 | 5696402.827 | 5010660.978 | 4222168.812 | 4878369.681 | 1801758.801 | 3566868 | 835430.4897 | 41.0482 | 65.1022 | Efhd2 | 27984 | 1.3834 | 0.616596515 | 0.44571 | 0.06959 | 0.601634 |
| 148709446 | PRP19/PSO4 pre-mRNA processing factor 19 homolog (S. cerevisiae), isoform CRA_b, partial [Mus musculus] | 0 | 7.39 | 5 | 2 | 4 | 2 | 533 | 58.4 | 6.52 | 32 | 2 | 3454146.78 | 15791075.32 | 8323956.047 | 1181928.533 | 6912562.335 | 12489902.26 | 9964981.5 | 10209111.26 | 89.919 | 23.1448 | Prpf19 | 28000 | 0.84157 | 1.158434374 | 1.37652 | 0.45993 | 0.862375 |
| 148671850 | mCG13557, isoform CRA_c [Mus musculus] | 0 | 25.879 | 27 | 4 | 9 | 4 | 215 | 23.6 | 9.73 | 228 | 4 | 30706641.18 | 4948272.035 | 18790223.12 | 18665018.32 | 18254883.67 | 4880365.707 | 21306326 | 2845686.48 | 57.605 | 78.7628 | Atp5o | 28080 | 1.21448 | 0.785519544 | 0.64679 | 0.39373 | 0.824613 |
| 21703842 | tRNA-splicing ligase RtcB homolog [Mus musculus] | 0 | 119.894 | 43 | 20 | 81 | 20 | 505 | 55.2 | 7.23 | 828 | 20 | 112467605.4 | 100007663.7 | 111091049.7 | 136789071.2 | 105899995.5 | 101621407.1 | 112857537.5 | 116519214 | 13.472 | 6.14783 | Rtcb | 28088 | 1.02614 | 0.973856625 | 0.94905 | 0.51355 | 0.88031 |
| 148699794 | 1-acylglycerol-3-phosphate O-acyltransferase 3, isoform CRA_b, partial [Mus musculus] | 0 | 6.963 | 8 | 3 | 4 | 3 | 379 | 43.5 | 8.51 | 20 | 3 | 8341627.363 | 2893604.559 | 8457673.209 | 8997550.005 | 10474592.07 | 15520521 | 10672325.06 | 2647458.177 | 39.9708 | 54.1872 | Agpat3 | 28169 | 0.84377 | 1.156229487 | 1.37031 | 0.41358 | 0.835788 |
| 28972369 | mKIAA0719 protein [Mus musculus] | 0 | 21.806 | 14 | 7 | 20 | 7 | 626 | 69.5 | 8.37 | 143 | 7 | 21666030.45 | 16663810.86 | 20800524.24 | 16206940.38 | 29393269.4 | 24782086.51 | 24245223 | 20836243.29 | 14.8598 | 14.175 | Tomm70a | 28185 | 0.863 | 1.137000691 | 1.3175 | 0.03749 | 0.56531 |
| 256220343 | dehydrogenase/reductase SDR family member 4 isoform 1 [Mus musculus] | 0 | 13.061 | 12 | 4 | 15 | 4 | 279 | 29.9 | 9.38 | 48 | 4 | 22946356.12 | 33563617.55 | 27317538.08 | 25939713.59 | 23465379.59 | 31453730.77 | 27763684 | 30615851.46 | 16.291 | 12.7241 | Dhrs4 | 28200 | 0.98417 | 1.01583129 | 1.03217 | 0.76887 | 0.952792 |
| 115270960 | BAG family molecular chaperone regulator 3 [Mus musculus] | 0 | 21.621 | 7 | 3 | 15 | 3 | 577 | 61.8 | 7.27 | 298 | 3 | 13819293.26 | 14740527.3 | 24053802.91 | 17938503.62 | 15321079.8 | 14752956.16 | 22447573 | 17430701.4 | 26.2338 | 20.0209 | Bag3 | 29810 | 1.00427 | 0.995730977 | 0.9915 | 0.96046 | 0.994906 |
| 1039746315 | PREDICTED: protein NDRG2 isoform X3 [Mus musculus] | 0 | 25.577 | 18 | 5 | 13 | 5 | 374 | 41.1 | 5.4 | 143 | 5 | 15536325.26 | 12177397.54 | 19977384.21 | 13227899.37 | 19089366.4 | 8087663.362 | 15709125.13 | 9078628.323 | 22.7324 | 40.7139 | Ndrg2 | 29811 | 1.07932 | 0.920677511 | 0.85301 | 0.50537 | 0.88031 |
| 7305307 | protein NDRG3 isoform 2 [Mus musculus] | 0 | 16.35 | 19 | 4 | 12 | 4 | 375 | 41.5 | 5.25 | 43 | 4 | 11412445.21 | 7538804.067 | 10973473.42 | 5575114.157 | 10520971.94 | 4097029.903 | 8625245 | 7603912.446 | 31.5474 | 34.9585 | Ndrg3 | 29812 | 1.07013 | 0.929873576 | 0.86894 | 0.57138 | 0.895889 |
| 37360028 | mKIAA0655 protein [Mus musculus] | 0 | 8.588 | 3 | 3 | 11 | 3 | 1083 | 120.9 | 6.58 | 21 | 3 | 2074652.866 | 1032868.042 | 2448244.633 | 2189205.809 | 2128087.696 | 2666245.608 | 2448060.438 | 2509472.215 | 32.1341 | 9.27375 | Hip1r | 29816 | 0.8853 | 1.114700421 | 1.25912 | 0.18036 | 0.715071 |
| 49022835 | mKIAA0623 protein [Mus musculus] | 0 | 12.9 | 4 | 3 | 7 | 3 | 1056 | 114.6 | 8.54 | 60 | 3 | 5777470.89 | 4833181.311 | 4304968.447 | 4045080.028 | 3647167.119 | 3698521.62 | 3424899.75 | 4028472.91 | 16.1456 | 6.73675 | Ulk2 | 29869 | 1.12327 | 0.876727826 | 0.78051 | 0.04148 | 0.585646 |
| 223461282 | Cttnbp2 protein [Mus musculus] | 0 | 123.419 | 17 | 21 | 171 | 21 | 1650 | 178.8 | 8.19 | 2835 | 21 | 120351584.1 | 121470686 | 94228002.1 | 141439140.3 | 110973078.9 | 153143143.8 | 130943023.4 | 132906205.6 | 16.219 | 13.0574 | Cttnbp2 | 30785 | 0.9498 | 1.050202193 | 1.10571 | 0.36783 | 0.80688 |
| 251823786 | PDZ and LIM domain protein 4 [Mus musculus] | 0 | 21.174 | 26 | 6 | 19 | 6 | 330 | 35.5 | 8.05 | 173 | 6 | 12423231.62 | 11853555.72 | 9286182.883 | 7226327.274 | 8015204.337 | 6473450.485 | 10537061.63 | 19231866.29 | 23.5864 | 51.4875 | Pdlim4 | 30794 | 0.95922 | 1.040780864 | 1.08503 | 0.78856 | 0.953636 |
| 21703770 | myeloid leukemia factor 2 [Mus musculus] | 0 | 59.565 | 41 | 7 | 29 | 7 | 247 | 28 | 6.98 | 427 | 7 | 34356038.05 | 18528693.82 | 21520916.4 | 25029834.37 | 22026993.42 | 9889546.682 | 25966973.41 | 8687314.915 | 27.6212 | 52.0162 | Mlf2 | 30853 | 1.19797 | 0.802027682 | 0.66949 | 0.18753 | 0.715071 |
| 569003953 | PREDICTED: myc box-dependent-interacting protein 1 isoform X1 [Mus musculus] | 0 | 59.983 | 27 | 11 | 40 | 10 | 603 | 66.4 | 5.4 | 314 | 11 | 32526641.71 | 24254833.19 | 31309447.7 | 25270261.76 | 35481168.69 | 28300778.55 | 36846768 | 26824091.35 | 14.755 | 15.7973 | Bin1 | 30948 | 0.94148 | 1.058516626 | 1.12431 | 0.32299 | 0.771047 |
| 254540192 | C-Jun-amino-terminal kinase-interacting protein 3 isoform a [Mus musculus] | 0 | 88.913 | 18 | 15 | 42 | 15 | 1337 | 147.5 | 5.45 | 851 | 15 | 36372752.12 | 26608082.71 | 22493328.35 | 17846594.16 | 42800102.96 | 25086680.87 | 29224940.88 | 24966947.84 | 30.5348 | 27.5983 | Mapk8ip3 | 30957 | 0.91678 | 1.083220775 | 1.18155 | 0.44738 | 0.852048 |
| 94721328 | vesicle-associated membrane protein-associated protein A [Mus musculus] | 0 | 73.926 | 39 | 8 | 60 | 7 | 249 | 27.8 | 8.4 | 693 | 8 | 114235816.4 | 102937995 | 81550467.21 | 105605747 | 94651873.66 | 90062467.48 | 80133243.44 | 94162487.05 | 13.7367 | 7.50389 | Vapa | 30960 | 1.05937 | 0.940629407 | 0.88791 | 0.1924 | 0.715071 |
| 13430890 | histone H1.4 [Mus musculus] | 0 | 28.344 | 19 | 5 | 74 | 3 | 219 | 22 | 11.11 | 1170 | 5 | 4731234.648 | 20656041.02 | 64803602.42 | 95022071.33 | 45242312.18 | 106261355.5 | 60072996.5 | 150434564.4 | 89.0614 | 52.6621 | Hist1h1e | 50709 | 0.67692 | 1.323081995 | 1.95457 | 0.2103 | 0.716066 |
| 568988117 | PREDICTED: sacsin isoform X1 [Mus musculus] | 0 | 290.194 | 17 | 64 | 217 | 64 | 4591 | 521.4 | 7.05 | 2672 | 64 | 140243950.6 | 106147958.2 | 172950349.1 | 121329539.1 | 136524853.5 | 123161142.8 | 158568180 | 114087271.4 | 21.3009 | 14.5224 | Sacs | 50720 | 1.00776 | 0.99223649 | 0.98459 | 0.90832 | 0.98561 |
| 282721034 | membrane-associated guanylate kinase, WW and PDZ domain-containing protein 2 isoform 1 [Mus musculus] | 0 | 210.736 | 37 | 37 | 191 | 36 | 1275 | 140.8 | 6.19 | 2215 | 37 | 192906251.8 | 156730346.2 | 219271904.6 | 184792071.2 | 245605613.2 | 203061775.7 | 202904557.2 | 192072878.2 | 13.6665 | 11.2344 | Magi2 | 50791 | 0.94369 | 1.05630858 | 1.11934 | 0.24612 | 0.733806 |
| 244790106 | spastin isoform 1 [Mus musculus] | 0 | 10.09 | 7 | 3 | 5 | 3 | 614 | 66.4 | 9.69 | 0 | 3 | 5063535.314 | 9234930.674 | 5416700.942 | 8184264.543 | 5450622.587 | 7090468.901 | 1396299 | 8523435.295 | 29.4426 | 54.8523 | Spast | 50850 | 1.10799 | 0.892005999 | 0.80506 | 0.4903 | 0.870399 |
| 37359786 | mKIAA0132 protein [Mus musculus] | 0 | 26.086 | 12 | 7 | 23 | 7 | 637 | 71 | 6.44 | 390 | 7 | 8194254.113 | 8436901.306 | 8077889.444 | 2093274.472 | 6337273.855 | 6920511.001 | 6690244 | 4775248.088 | 45.8942 | 15.6494 | Keap1 | 50868 | 1.04035 | 0.959650299 | 0.92243 | 0.75804 | 0.952737 |
| 74141836 | unnamed protein product [Mus musculus] | 0 | 14.386 | 14 | 4 | 8 | 3 | 352 | 39.4 | 5.19 | 42 | 4 | 2997177.359 | 2542369.935 | 4376575.743 | 3628087.176 | 5143870.514 | 2492419.676 | 3353616 | 2231246.862 | 23.5199 | 39.8206 | Tmod3 | 50875 | 1.01207 | 0.987930029 | 0.97615 | 0.9198 | 0.987994 |
| 31543876 | tropomodulin-2 [Mus musculus] | 0 | 232.511 | 66 | 21 | 198 | 20 | 351 | 39.5 | 5.35 | 5056 | 21 | 558506086.5 | 622594247.1 | 708424366.9 | 636240111.1 | 619116255.9 | 516300551.6 | 623874091.9 | 596413724.8 | 9.73983 | 8.46926 | Tmod2 | 50876 | 1.03484 | 0.965162091 | 0.93267 | 0.32418 | 0.772052 |
| 595582321 | nck-associated protein 1 isoform a [Mus musculus] | 0 | 434.558 | 53 | 51 | 385 | 51 | 1134 | 129.4 | 6.68 | 7267 | 51 | 653268244.4 | 698361417.6 | 859701809.7 | 739907590.3 | 832989378 | 938461874.5 | 968091643.6 | 1031917779 | 12.0125 | 8.80122 | Nckap1 | 50884 | 0.87799 | 1.122007811 | 1.27792 | 0.0149 | 0.461382 |
| 158749640 | prolactin regulatory element-binding protein isoform 1 [Mus musculus] | 0.001 | 6.798 | 7 | 2 | 6 | 2 | 417 | 45.4 | 8.76 | 19 | 2 | 4234498.083 | 2020998.36 | 6368690.759 | 5706649.864 | 6818130.125 | 6580259.532 | 6086314.25 | 2183005.611 | 42.0429 | 40.196 | Preb | 50907 | 0.91658 | 1.083424843 | 1.18204 | 0.58692 | 0.899913 |
| 148699227 | myeloid-associated differentiation marker, isoform CRA_b, partial [Mus musculus] | 0 | 20.897 | 12 | 4 | 19 | 4 | 347 | 37.9 | 8.57 | 364 | 4 | 23232099.9 | 13314397.28 | 8711788.935 | 10976842.14 | 16828091.53 | 15538765.76 | 11101947 | 17494775.71 | 45.5069 | 18.8721 | Myadm | 50918 | 0.95965 | 1.040345596 | 1.08408 | 0.74756 | 0.947921 |
| 568974701 | PREDICTED: misshapen-like kinase 1 isoform X1 [Mus musculus] | 0 | 245.671 | 34 | 33 | 268 | 22 | 1374 | 154.9 | 7.65 | 4137 | 33 | 174140286.1 | 157346571.1 | 170860605.3 | 184795563.6 | 166694023.6 | 226137483.5 | 176218820.9 | 210236342.9 | 6.58718 | 14.3829 | Mink1 | 50932 | 0.93716 | 1.062835365 | 1.1341 | 0.17821 | 0.713031 |
| 568974714 | PREDICTED: MAGUK p55 subfamily member 2 isoform X1 [Mus musculus] | 0 | 66.739 | 34 | 13 | 48 | 12 | 569 | 63.3 | 6.11 | 505 | 13 | 66770252.65 | 93692050.46 | 67128779.49 | 92968075.72 | 74170379.45 | 81105967.21 | 69974856.81 | 62635680.78 | 19.0098 | 10.7451 | Mpp2 | 50997 | 1.0537 | 0.946302101 | 0.89808 | 0.37591 | 0.807546 |
| 26353870 | unnamed protein product [Mus musculus] | 0 | 273.334 | 75 | 33 | 259 | 33 | 589 | 65.3 | 5.11 | 3389 | 33 | 497925879.4 | 441095120.1 | 485782984.2 | 483759167.8 | 443920395.2 | 428635114 | 448198929.1 | 402793798.2 | 5.20422 | 4.76351 | Ppp2r1a | 51792 | 1.05094 | 0.949061332 | 0.90306 | 0.02837 | 0.515682 |
| 160333923 | heterogeneous nuclear ribonucleoprotein U [Mus musculus] | 0 | 83.244 | 29 | 17 | 88 | 17 | 800 | 87.9 | 6.24 | 1038 | 17 | 119715739.2 | 70400050.35 | 50959956.06 | 65296764.9 | 44265371.07 | 64196544.34 | 67933056.63 | 58238188.53 | 39.0415 | 17.7174 | Hnrnpu | 51810 | 1.1326 | 0.867396307 | 0.76584 | 0.30043 | 0.761857 |
| 569005874 | PREDICTED: serine/threonine-protein phosphatase 6 regulatory subunit 3 isoform X1 [Mus musculus] | 0 | 50.432 | 11 | 8 | 24 | 8 | 879 | 98.6 | 4.6 | 679 | 8 | 6896786.102 | 9127689.27 | 9072848.998 | 6890260.011 | 12303720.37 | 3357611.073 | 7465607.125 | 5173359.515 | 15.9345 | 54.697 | Ppp6r3 | 52036 | 1.06116 | 0.938838683 | 0.88473 | 0.6668 | 0.922517 |
| 52421788 | rab11 family-interacting protein 5 isoform 1 [Mus musculus] | 0 | 61.303 | 11 | 9 | 30 | 9 | 1318 | 139.5 | 5.12 | 868 | 9 | 35710915.98 | 13904124.65 | 26032303.26 | 20350653.43 | 29760284.69 | 20751510.95 | 34793952.69 | 21593077.07 | 38.5299 | 25.2254 | Rab11fip5 | 52055 | 0.94627 | 1.053725967 | 1.11355 | 0.65072 | 0.915864 |
| 26000252 | 1-acylglycerolphosphate acyltransferase-epsilon [Mus musculus] | 0 | 19.66 | 13 | 4 | 40 | 4 | 365 | 42.3 | 9.38 | 287 | 4 | 59171082.84 | 94004239.85 | 59172577.14 | 85959690.81 | 97018835.82 | 57091167.41 | 51556576.93 | 53089629.23 | 24.2553 | 33.5128 | Agpat5 | 52123 | 1.071 | 0.929000267 | 0.86741 | 0.50991 | 0.88031 |
| 19527140 | calcium/calmodulin-dependent protein kinase type 1 [Mus musculus] | 0.001 | 6.103 | 5 | 2 | 6 | 2 | 374 | 41.6 | 5.35 | 34 | 2 | 4914811.785 | 3501527.778 | 4028925.294 | 2922908.526 | 4610737.021 | 1607486.133 | 4666024.25 | 1391970.533 | 22.0163 | 59.1184 | Camk1 | 52163 | 1.11185 | 0.888152522 | 0.79881 | 0.46927 | 0.863085 |
| 30519931 | receptor expression-enhancing protein 1 [Mus musculus] | 0 | 11.562 | 16 | 3 | 9 | 3 | 201 | 22.3 | 9.5 | 51 | 3 | 2326870.777 | 2913858.307 | 2047197.091 | 2213815.777 | 2352606.273 | 2864526.066 | 2230551.75 | 8843255.806 | 15.8658 | 78.3794 | Reep1 | 52250 | 0.73678 | 1.26322187 | 1.71452 | 0.33159 | 0.776788 |
| 46092406 | mKIAA1828 protein [Mus musculus] | 0 | 56.555 | 24 | 8 | 45 | 8 | 587 | 64.6 | 7.28 | 1146 | 8 | 110335225.9 | 76686665.36 | 111221887.1 | 120176463.1 | 93579753.43 | 139951222.7 | 108223712.5 | 109265687.2 | 18.2933 | 17.2887 | Adgra1 | 52389 | 0.9625 | 1.037495527 | 1.07791 | 0.57251 | 0.895889 |
| 157821523 | septin-11 [Rattus norvegicus] | 0 | 127.704 | 53 | 20 | 153 | 10 | 432 | 49.8 | 6.68 | 1907 | 20 | 48490140.32 | 53519407.69 | 52200269.69 | 61117559.99 | 57256893.15 | 161234992.9 | 54358272.7 | 199105669.8 | 9.85161 | 62.2567 | 43719 | 52398 | 0.6266 | 1.373395492 | 2.19181 | 0.13209 | 0.693437 |

| | | | | | | | | | | | | | | | | | | | | | | | | | | | | | |
|---|---|---|---|---|---|---|---|---|---|---|---|---|---|---|---|---|---|---|---|---|---|---|---|---|---|---|---|---|---|
| 20810027 | Acetyl-Coenzyme A acyltransferase 2 (mitochondrial 3-oxoacyl-Coenzyme A thiolase) [Mus musculus] | 0 | 36.393 | 19 | 6 | 35 | 6 | 397 | 41.8 | 8.09 | 476 | 6 | 23425552.55 | 28132955.81 | 29556306.59 | 16354940.83 | 27798288.19 | 11931164.74 | 29343132.5 | 8423286.865 | 24.4159 | 55.405 | Acaa2 | 52538 | 1.11416 | 0.885841101 | 0.79508 | 0.44687 | 0.852048 |
| 12834933 | unnamed protein product [Mus musculus] | 0 | 77.53 | 50 | 13 | 111 | 13 | 313 | 34 | 8.48 | 2134 | 13 | 85289333.59 | 94236688.82 | 65533555.15 | 79199013.28 | 97812159 | 112210977.1 | 65699209.19 | 114062904.7 | 14.8722 | 22.9629 | Dhrs1 | 52585 | 0.90823 | 1.091768398 | 1.20208 | 0.24486 | 0.733806 |
| 15029877 | Ncald protein [Mus musculus] | 0 | 37.293 | 47 | 7 | 73 | 4 | 193 | 22.2 | 5.22 | 697 | 7 | 42848728.02 | 24850029.47 | 37166917.91 | 22824059.47 | 40762874.8 | 35876924.38 | 34358762.5 | 29648843.62 | 30.2469 | 13.0246 | Ncald | 52589 | 0.95171 | 1.04828877 | 1.10148 | 0.56656 | 0.895889 |
| 19527228 | CDGSH iron-sulfur domain-containing protein 1 [Mus musculus] | 0 | 26.818 | 35 | 3 | 20 | 3 | 108 | 12.1 | 9.06 | 309 | 3 | 26265867.72 | 29095906.36 | 19127787.91 | 19438023.45 | 37192480.33 | 42181521.26 | 22237833.88 | 36536779.5 | 21.2331 | 24.8377 | Cisd1 | 52637 | 0.80945 | 1.190545301 | 1.4708 | 0.06741 | 0.601634 |
| 21326440 | ZW10 interactor [Mus musculus] | 0 | 14.032 | 13 | 3 | 13 | 3 | 252 | 28.7 | 8.43 | 51 | 3 | 4510546.016 | 10711421.66 | 8640414.144 | 5592065.637 | 10706123.51 | 8529056.074 | 7559773 | 11789207.86 | 38.5039 | 20.1353 | Zwint | 52696 | 0.86582 | 1.1341843 | 1.30996 | 0.23238 | 0.728784 |
| 568935984 | PREDICTED: protein RUFY3 isoform X1 [Mus musculus] | 0 | 32.459 | 13 | 8 | 36 | 8 | 687 | 76.8 | 5.68 | 483 | 8 | 32432817.84 | 26833124.92 | 41227415.57 | 19171226.14 | 47770008.52 | 36037431.8 | 37942408.5 | 20559613.84 | 31.0729 | 31.6342 | Rufy3 | 52822 | 0.91356 | 1.086439395 | 1.18924 | 0.46741 | 0.863085 |
| 37360266 | mKIAA1162 protein [Mus musculus] | 0 | 12.409 | 12 | 3 | 4 | 3 | 354 | 39.1 | 4.37 | 66 | 3 | 4160753.902 | 5138568.462 | 4301511.733 | 6539289.464 | 6050156.087 | 6289105.983 | 6081257.25 | 5448840.767 | 21.6836 | 6.05924 | Tmx4 | 52837 | 0.91526 | 1.084737112 | 1.18516 | 0.15609 | 0.69857 |
| 568936011 | PREDICTED: small G protein signaling modulator 1 isoform X1 [Mus musculus] | 0 | 16.615 | 4 | 4 | 13 | 4 | 1148 | 129 | 5.77 | 148 | 4 | 8366472.73 | 4535283.028 | 4769744.992 | 3884229.048 | 5681408.369 | 4228229.465 | 2769211.375 | 5618916.137 | 37.4854 | 30.1196 | Sgsm1 | 52850 | 1.08175 | 0.918251475 | 0.84886 | 0.53005 | 0.886555 |
| 568951027 | PREDICTED: CDP-diacylglycerol--inositol 3-phosphatidyltransferase isoform X1 [Mus musculus] | 0.001 | 6.002 | 10 | 2 | 6 | 2 | 220 | 24.3 | 8.19 | 50 | 2 | 5366226.561 | 4527613.808 | 5355592.479 | 3893085.871 | 6488087.314 | 5189005.255 | 5359015 | 4523789.327 | 14.899 | 15.1389 | Cdipt | 52858 | 0.94061 | 1.059391516 | 1.12628 | 0.30734 | 0.762436 |
| 111600355 | Rgs7bp protein [Mus musculus] | 0 | 13.68 | 11 | 2 | 6 | 2 | 257 | 29 | 8.46 | 152 | 2 | 1505719.41 | 3388221.552 | 4382746.585 | 2247450.054 | 4547956.012 | 1206296.357 | 4644331 | 1555351.391 | 43.929 | 62.314 | Rgs7bp | 52882 | 0.98169 | 1.018306322 | 1.0373 | 0.92707 | 0.991095 |
| 171906571 | jouberin isoform 1 [Mus musculus] | 0 | 52.758 | 14 | 13 | 42 | 13 | 1047 | 119.6 | 7.2 | 306 | 13 | 17583769.04 | 21662239.08 | 36577194.87 | 33041215.16 | 72527137.24 | 37473438.92 | 35742784.25 | 53702148.82 | 33.2173 | 34.3742 | Ahi1 | 52906 | 0.7062 | 1.293798814 | 1.83205 | 0.05806 | 0.599194 |
| 28972680 | mKIAA1232 protein [Mus musculus] | 0 | 628.182 | 71 | 58 | 941 | 5 | 950 | 103.8 | 6.7 | 14958 | 58 | 18363136.41 | 36794151.83 | 27000490.57 | 20554713.13 | 23956498.2 | 43816174.81 | 21782620 | 45491529.65 | 32.1981 | 37.4003 | Dlg3 | 53310 | 0.864 | 1.135996063 | 1.3148 | 0.32528 | 0.772052 |
| 26346549 | unnamed protein product [Mus musculus] | 0 | 53.209 | 21 | 9 | 33 | 9 | 618 | 70.3 | 8.72 | 584 | 9 | 12976958.98 | 18382446.57 | 18185531.68 | 12948847.32 | 23305444.17 | 22031021.92 | 22154578.75 | 35592615.62 | 19.6705 | 25.5051 | Nxf1 | 53319 | 0.75486 | 1.245141335 | 1.6495 | 0.03129 | 0.515682 |
| 116063560 | contactin-associated protein 1 precursor [Mus musculus] | 0 | 251.113 | 37 | 37 | 222 | 37 | 1385 | 156.2 | 7.06 | 3393 | 37 | 478568774.3 | 433859879.9 | 407844948.4 | 377356388.9 | 586120302.5 | 447282548 | 493182102.4 | 410535325.5 | 10.0988 | 15.6616 | Cntnap1 | 53321 | 0.93411 | 1.065889062 | 1.14107 | 0.21841 | 0.716066 |
| 922959942 | myotubularin-related protein 1 isoform D [Mus musculus] | 0 | 33.111 | 13 | 6 | 19 | 5 | 703 | 79.3 | 6.7 | 105 | 6 | 12689777.42 | 5836977.88 | 10151336.74 | 6976954.516 | 5013585.081 | 6135255.325 | 5915119.75 | 5060360.561 | 34.8847 | 10.4482 | Mtmr1 | 53332 | 1.23418 | 0.765820803 | 0.62051 | 0.07625 | 0.627416 |
| 157909797 | mitochondrial import receptor subunit TOM40 homolog [Mus musculus] | 0 | 29.515 | 19 | 5 | 18 | 5 | 361 | 37.9 | 7.74 | 509 | 5 | 20439222.03 | 15043901.62 | 14327289.46 | 13811863.43 | 25866177.78 | 20610711.18 | 21506465.25 | 17327610.24 | 19.266 | 16.4984 | Tomm40 | 53333 | 0.85437 | 1.145626912 | 1.3409 | 0.05912 | 0.600904 |
| 228480241 | metaxin-2 [Mus musculus] | 0 | 61.202 | 50 | 7 | 33 | 7 | 263 | 29.7 | 5.63 | 900 | 7 | 33928070.06 | 18686480.66 | 28672991.73 | 21067718.61 | 32610562.81 | 28108571.61 | 35846015.09 | 23926119.71 | 27.3685 | 17.2924 | Mtx2 | 53375 | 0.91862 | 1.081383399 | 1.17719 | 0.33887 | 0.782483 |
| 148666221 | mCG119114, isoform CRA_a, partial [Mus musculus] | 0 | 75.98 | 40 | 14 | 87 | 12 | 357 | 38.1 | 9.31 | 964 | 14 | 35927000.2 | 80266372.55 | 64300096.89 | 108594092.4 | 47705300.76 | 84204611.17 | 85950885.13 | 112230920.1 | 42.0272 | 32.1326 | Hnrnpa2b1 | 53379 | 0.93378 | 1.066223397 | 1.14184 | 0.62926 | 0.912586 |
| 568974812 | PREDICTED: exocyst complex component 7 isoform X1 [Mus musculus] | 0 | 9.828 | 5 | 3 | 4 | 3 | 720 | 82.5 | 7.06 | 18 | 3 | 2771933.55 | 864460.9221 | 2389423.321 | 3677720.648 | 1342223.477 | 2160490.497 | 2706401.625 | 1091407.092 | 48.3443 | 40.7648 | Exoc7 | 53413 | 1.14132 | 0.858679892 | 0.75236 | 0.42022 | 0.83996 |
| 8394392 | synaptotagmin-5 [Mus musculus] | 0 | 25.238 | 22 | 8 | 27 | 5 | 386 | 43.1 | 9.6 | 163 | 8 | 729453.2128 | 3636309.861 | 5624728.001 | 3582425.114 | 4320774.322 | 4708908.889 | 4230660.25 | 5236464.05 | 59.3574 | 9.90198 | Syt5 | 53420 | 0.84646 | 1.153537067 | 1.36277 | 0.27829 | 0.745606 |
| 12848217 | unnamed protein product [Mus musculus] | 0 | 23.756 | 35 | 5 | 9 | 5 | 186 | 21 | 6.06 | 109 | 5 | 10222140.37 | 17563204.59 | 12342517.58 | 10748291.08 | 12921097.13 | 16354218.75 | 13397905 | 15362717.78 | 26.3614 | 11.1771 | Dctn3 | 53598 | 0.93426 | 1.065739119 | 1.14073 | 0.37358 | 0.80688 |
| 7949055 | hippocalcin-like protein 1 [Mus musculus] | 0 | 48.512 | 49 | 9 | 70 | 5 | 193 | 22.3 | 5.5 | 528 | 9 | 48361455.68 | 36129349.17 | 33383610.88 | 27950169.45 | 42010201.63 | 30816562.23 | 37497837.25 | 26863216.03 | 23.6831 | 19.7117 | Hpcal1 | 53602 | 1.03052 | 0.969482722 | 0.94077 | 0.70734 | 0.934268 |
| 226443026 | nucleosome assembly protein 1-like 1 isoform 1 precursor [Mus musculus] | 0 | 30.465 | 16 | 5 | 34 | 4 | 418 | 48.5 | 4.48 | 486 | 5 | 35140192.39 | 20487056.79 | 29595197.32 | 20544439.31 | 27409703.16 | 16290435.14 | 25817750 | 18597111.67 | 27.258 | 24.5883 | Nap1l1 | 53605 | 1.09104 | 0.908955466 | 0.83311 | 0.36542 | 0.80688 |
| 74141990 | unnamed protein product [Mus musculus] | 0 | 93.648 | 38 | 13 | 131 | 12 | 528 | 60.4 | 9.61 | 1560 | 13 | 77307317.22 | 99293247.13 | 102605090.1 | 118740027.2 | 78151956.74 | 118669072.2 | 111528768 | 91620506.88 | 17.1407 | 18.5216 | Nono | 53610 | 0.99746 | 1.002537388 | 1.00509 | 0.96923 | 0.99686 |
| 125858934 | Gria3 protein [Mus musculus] | 0 | 322.831 | 45 | 33 | 435 | 3 | 888 | 100.4 | 8.29 | 9570 | 33 | 265679040.4 | 285878243.4 | 189695471.5 | 222663706.4 | 222957503.6 | 303618654.7 | 189921258.5 | 225927762.9 | 17.9153 | 20.4531 | Gria3 | 53623 | 1.01127 | 0.988726427 | 0.9777 | 0.87354 | 0.979509 |
| 148702018 | porcupine homolog (Drosophila), isoform CRA_b, partial [Mus musculus] | 0.001 | 5.647 | 4 | 2 | 4 | 2 | 516 | 58.3 | 9.35 | 0 | 2 | 5039561.112 | 6592791.44 | 5261421.524 | 5115329.718 | 6238706.668 | 9215939.603 | 6075209.063 | 6855029.329 | 13.3185 | 20.4685 | Porcn | 53627 | 0.87348 | 1.126518679 | 1.28969 | 0.09776 | 0.670764 |
| 92110037 | protein PRRC2A isoform 1 [Mus musculus] | 0 | 23.789 | 5 | 6 | 9 | 6 | 2158 | 229.1 | 9.39 | 57 | 6 | 11114045.57 | 14384259.56 | 10264683.68 | 9283775.953 | 10292942.52 | 12837137.98 | 11976988.25 | 11759909.85 | 19.6415 | 9.02173 | Prrc2a | 53761 | 0.9802 | 1.0198035 | 1.04041 | 0.72322 | 0.939018 |
| 9790069 | spliceosome RNA helicase Ddx39b [Mus musculus] | 0 | 8.777 | 7 | 2 | 6 | 2 | 428 | 49 | 5.67 | 17 | 2 | 1315008.653 | 1306667.727 | 1051127.274 | 1244439.726 | 1509096.494 | 1566349.109 | 1090767.375 | 1717389.072 | 9.99673 | 18.2351 | Ddx39b | 53817 | 0.91053 | 1.089470651 | 1.19652 | 0.1526 | 0.693437 |
| 49022852 | mKIAA0991 protein [Mus musculus] | 0 | 21.585 | 11 | 5 | 15 | 5 | 605 | 67.8 | 8.12 | 64 | 5 | 7474704.738 | 6045799.421 | 7901491.418 | 5917699.113 | 14192149.85 | 13815264.8 | 11493854.13 | 17969741.35 | 14.6573 | 18.6626 | 43717 | 53860 | 0.64472 | 1.355277267 | 2.10211 | 0.0019 | 0.38143 |
| 74198630 | unnamed protein product [Mus musculus] | 0 | 17.207 | 12 | 4 | 10 | 4 | 345 | 38.8 | 8.24 | 130 | 4 | 4229949.603 | 6679847.31 | 14706291.43 | 13680431.49 | 16758568.11 | 12760923 | 13368570 | 19485165.86 | 52.5274 | 20.1019 | Zranb2 | 53861 | 0.77302 | 1.226977128 | 1.58725 | 0.10456 | 0.674486 |
| 114050895 | cadherin EGF LAG seven-pass G-type receptor 2 isoform 1 precursor [Mus musculus] | 0 | 32.61 | 4 | 8 | 20 | 8 | 2919 | 316.8 | 5.47 | 74 | 8 | 6821161.429 | 9621400.939 | 8887590.066 | 15179516.03 | 13346335.92 | 14408732.78 | 8935557.188 | 16454956.89 | 35.2571 | 23.893 | Celsr2 | 53883 | 0.86508 | 1.134919443 | 1.31192 | 0.23422 | 0.730166 |
| 162417957 | ephexin-1 isoform 1 [Mus musculus] | 0 | 39.904 | 15 | 7 | 35 | 7 | 710 | 82.2 | 5.99 | 531 | 7 | 19869075.23 | 29270279.18 | 26942838.76 | 21041245.89 | 22357643.18 | 28439096 | 24563072.13 | 34540996.67 | 18.7137 | 19.4332 | Ngef | 53972 | 0.93828 | 1.06171919 | 1.13156 | 0.3973 | 0.824613 |
| 755524532 | PREDICTED: rho guanine nucleotide exchange factor 7 isoform X1 [Mus musculus] | 0 | 191.047 | 48 | 29 | 134 | 5 | 862 | 97 | 6.8 | 2663 | 29 | 13988719.79 | 9090616.103 | 12755706.61 | 13623704.55 | 15998621.11 | 11241649.4 | 13453390 | 9713688.559 | 18.1415 | 21.71 | Arhgef7 | 54126 | 0.9905 | 1.009498739 | 1.01918 | 0.89774 | 0.985273 |
| 4506715 | 40S ribosomal protein S28 [Homo sapiens] | 0 | 8.122 | 30 | 2 | 21 | 2 | 69 | 7.8 | 10.7 | 386 | 2 | 69105643.2 | 131604773.6 | 77006021.5 | 126812272.7 | 54819568.34 | 113354517.5 | 82369272 | 143596124.1 | 32.2731 | 38.9657 | Rps28 | 54127 | 1.01301 | 0.986991808 | 0.97432 | 0.92126 | 0.987994 |
| 5031569 | alpha-centractin [Homo sapiens] | 0 | 28.267 | 20 | 6 | 21 | 2 | 376 | 42.6 | 6.64 | 331 | 6 | 3972816.864 | 5580663.665 | 4673166.139 | 4681595.645 | 4944631.185 | 5967181.572 | 5093867 | 6214647.872 | 13.9391 | 11.3387 | Actr1a | 54130 | 0.91947 | 1.080530038 | 1.17517 | 0.11916 | 0.678142 |
| 13431714 | se anchor protein 8-like; Short=AKAP8-like protein; AltName: Full=Neighbor of A-kinase-anchoring protein 95; Sho | 0 | 16.464 | 8 | 4 | 4 | 4 | 642 | 71.4 | 5.05 | 73 | 4 | 4263890.094 | 3705950.65 | 4442047.366 | 4818579.757 | 5291865.779 | 4091592.543 | 4436726.125 | 4956198.541 | 10.7481 | 11.3747 | Akap8l | 54194 | 0.95707 | 1.042933916 | 1.08972 | 0.31603 | 0.766307 |
| 8567358 | guanylate cyclase soluble subunit beta-1 isoform 1 [Mus musculus] | 0 | 48.721 | 20 | 11 | 44 | 11 | 620 | 70.6 | 5.31 | 507 | 11 | 39732819.25 | 32169330.02 | 40828252.24 | 29977175.07 | 35961834.12 | 32850553.6 | 33007815.56 | 27765188.59 | 15.1615 | 10.5042 | Gucy1b1 | 54195 | 1.04819 | 0.95180858 | 0.90805 | 0.34417 | 0.789054 |
| 148706231 | mCG5564, partial [Mus musculus] | 0 | 17.551 | 27 | 4 | 29 | 4 | 118 | 13.6 | 11.56 | 374 | 4 | 125964718 | 209342809.8 | 144765026.7 | 272803449.9 | 148998141.5 | 188841920 | 157500405 | 211740392.3 | 35.4612 | 16.3672 | Rpl36 | 54217 | 1.03137 | 0.968632535 | 0.93917 | 0.7636 | 0.952792 |
| 755506621 | PREDICTED: band 4.1-like protein 4B isoform X3 [Mus musculus] | 0 | 96.406 | 27 | 16 | 86 | 16 | 863 | 95.8 | 9.25 | 1422 | 16 | 70245950.05 | 48353779.74 | 75682132.01 | 58095274.69 | 59451802.51 | 55072134.78 | 60595396.63 | 59897546.88 | 19.4517 | 4.25405 | Epb41l4b | 54357 | 1.03562 | 0.964381476 | 0.93121 | 0.51421 | 0.88031 |
| 269784644 | DBIRD complex subunit ZNF326 [Mus musculus] | 0 | 22.624 | 12 | 6 | 15 | 6 | 580 | 65.1 | 5.21 | 135 | 6 | 8508474.985 | 4264593.453 | 8439161.986 | 8697593.945 | 6128555.142 | 8889304.976 | 9271403.875 | 10363220.72 | 28.6821 | 20.7943 | Zfp326 | 54367 | 0.92654 | 1.073458654 | 1.15857 | 0.42966 | 0.845779 |
| 21687008 | voltage-dependent calcium channel gamma-3 subunit [Rattus norvegicus] | 0 | 82.002 | 27 | 8 | 110 | 6 | 315 | 35.5 | 9.48 | 1414 | 8 | 442453695 | 397051960.8 | 322634638.9 | 350343162.3 | 361178915.9 | 411338222.1 | 339117260 | 375037925.8 | 13.9503 | 8.153 | Cacng3 | 54376 | 1.00861 | 0.991393867 | 0.98293 | 0.83902 | 0.969831 |
| 9506455 | voltage-dependent calcium channel gamma-4 subunit [Mus musculus] | 0 | 29.034 | 25 | 6 | 24 | 6 | 327 | 36.5 | 9.11 | 444 | 6 | 58230819.16 | 57939810.03 | 48956725.26 | 74542310.83 | 59618571.45 | 74617656.68 | 60577173.25 | 57605527.09 | 17.7878 | 12.3201 | Cacng4 | 54377 | 0.97409 | 1.025908471 | 1.0532 | 0.64607 | 0.915705 |
| 11093516 | GABA-B receptor [Mus musculus] | 0 | 151.801 | 29 | 21 | 146 | 21 | 960 | 108.2 | 8.21 | 2811 | 21 | 203801818.7 | 226279136.1 | 245333878.8 | 192846235.4 | 238016069.4 | 256250160.3 | 223887124.8 | 217494698.3 | 10.7927 | 7.34632 | Gabbr1 | 54393 | 0.96264 | 1.037356086 | 1.07761 | 0.29023 | 0.753822 |
| 3065925 | 14-3-3 protein beta [Mus musculus] | 0 | 122.038 | 53 | 15 | 198 | 11 | 246 | 28.1 | 4.84 | 5632 | 15 | 90515726.4 | 101677625.2 | 108341059.9 | 113099611.9 | 105560094.4 | 117363277.7 | 92692933.69 | 99155851.56 | 9.46624 | 10.1444 | Ywhab | 54401 | 0.99863 | 1.001373884 | 1.00275 | 0.96969 | 0.99686 |
| 1039772072 | PREDICTED: electrogenic sodium bicarbonate cotransporter 1 isoform X2 [Mus musculus] | 0 | 19.562 | 4 | 3 | 12 | 3 | 1131 | 127.2 | 6.89 | 61 | 3 | 7085422.246 | 3787880.012 | 5885546.466 | 2732845.619 | 7178858.425 | 4359775.302 | 6349688 | 4341345.956 | 40.4884 | 25.8052 | Slc4a4 | 54403 | 0.93437 | 1.065625215 | 1.14047 | 0.59494 | 0.902243 |
| 568923259 | PREDICTED: synaptotagmin-6 isoform X1 [Mus musculus] | 0 | 37.773 | 23 | 12 | 41 | 11 | 529 | 59.1 | 8.03 | 277 | 12 | 31751823.39 | 22591976.84 | 29352363.85 | 35487313.69 | 35907318.28 | 29568996.38 | 24772287.31 | 34257323.98 | 18.2092 | 16.1127 | Syt6 | 54524 | 0.97816 | 1.021841115 | 1.04466 | 0.73105 | 0.943908 |
| 569007526 | PREDICTED: synaptotagmin-7 isoform X1 [Mus musculus] | 0 | 80.221 | 29 | 13 | 62 | 13 | 687 | 75.7 | 9.44 | 1003 | 13 | 118801207.5 | 98096582.94 | 90716299.06 | 101607157 | 112072278.6 | 88508334.72 | 117977446.8 | 95216933.59 | 11.629 | 13.4001 | Syt7 | 54525 | 0.99447 | 1.005533132 | 1.01113 | 0.90488 | 0.98561 |
| 172073152 | nuclear pore membrane glycoprotein 210 precursor [Mus musculus] | 0 | 134.223 | 19 | 22 | 71 | 22 | 1886 | 204 | 6.65 | 1526 | 22 | 97937412.44 | 85469067.05 | 93519036.45 | 92953081.1 | 95904950.43 | 94639259.6 | 101800731.5 | 86319156.64 | 5.59286 | 6.73939 | Nup210 | 54563 | 0.98826 | 1.011736807 | 1.02375 | 0.61197 | 0.905186 |
| 1039797914 | PREDICTED: WD repeat domain phosphoinositide-interacting protein 4 isoform X1 [Mus musculus] | 0 | 12.042 | 8 | 3 | 8 | 3 | 491 | 53.6 | 7.94 | 61 | 3 | 4410311.914 | 3564981.871 | 4683059.415 | 4239933.402 | 5013792.836 | 1906886.497 | 4842032.125 | 4856286.885 | 11.2697 | 36.1176 | Wdr45 | 54636 | 1.00833 | 0.991667336 | 0.98347 | 0.93221 | 0.994387 |
| 6755114 | peroxiredoxin-5, mitochondrial precursor [Mus musculus] | 0 | 63.116 | 57 | 11 | 51 | 11 | 210 | 21.9 | 8.85 | 945 | 11 | 178216681.2 | 161835108.5 | 189384543.9 | 152396166.6 | 168591747.4 | 159102616.8 | 160093238.9 | 149221408.9 | 9.69309 | 4.97845 | Prdx5 | 54683 | 1.03399 | 0.966012983 | 0.93426 | 0.26721 | 0.745606 |
| 568968076 | PREDICTED: plexin-C1 isoform X1 [Mus musculus] | 0 | 52.794 | 10 | 12 | 35 | 12 | 1752 | 196.1 | 8.94 | 240 | 12 | 19688788.06 | 13100517.98 | 14973319.88 | 17836823.16 | 22399933.87 | 15942603.31 | 21038782.77 | 16148226.39 | 17.8826 | 17.6022 | Plxnc1 | 54712 | 0.92964 | 1.07036185 | 1.15137 | 0.30549 | 0.761857 |
| 148697712 | eukaryotic translation initiation factor 3, subunit 7 (zeta), isoform CRA_c, partial [Mus musculus] | 0 | 19.698 | 12 | 6 | 20 | 6 | 549 | 64.1 | 6.19 | 75 | 6 | 15621779.72 | 21022703.5 | 18750746.15 | 20987605.02 | 17999131.67 | 20099874.09 | 19995756.5 | 22574507.71 | 13.3443 | 9.28959 | Eif3d | 55944 | 0.97271 | 1.02729308 | 1.05612 | 0.52326 | 0.882739 |
| 12852262 | unnamed protein product [Mus musculus] | 0 | 32.711 | 59 | 5 | 58 | 5 | 109 | 12.5 | 9.61 | 1027 | 5 | 55592207.27 | 69068538.17 | 47630312.59 | 75401654.2 | 91552986.74 | 122639467.3 | 53891883.25 | 77450265.21 | 20.3651 | 33.2619 | Mpc1 | 55951 | 0.83507 | 1.164931533 | 1.39501 | 0.17 | 0.711915 |
| 568920613 | PREDICTED: inositol monophosphatase 1 isoform X1 [Mus musculus] | 0 | 7.997 | 7 | 2 | 12 | 2 | 330 | 36.1 | 6.87 | 76 | 2 | 3282952.891 | 3444495.244 | 3157658.1 | 3791320.673 | 3431750.317 | 3893309.036 | 2533800.313 | 3174827.099 | 8.02895 | 17.4076 | Impa1 | 55980 | 1.02406 | 0.975936451 | 0.953 | 0.62827 | 0.912476 |
| 256985136 | E3 ubiquitin-protein ligase PDZRN3 [Mus musculus] | 0 | 19.399 | 4 | 4 | 11 | 4 | 1063 | 119.3 | 6.05 | 147 | 4 | 5058508.841 | 5347812.087 | 4332275.285 | 4159564.4 | 3680561.776 | 1575221.169 | 3591775.875 | 1875086.521 | 12.0546 | 41.4333 | Pdzrn3 | 55983 | 1.27601 | 0.723994165 | 0.56739 | 0.01693 | 0.461382 |
| 120407050 | nucleolar protein 58 [Mus musculus] | 0 | 11.041 | 7 | 3 | 4 | 3 | 536 | 60.3 | 8.34 | 0 | 3 | 3981784.638 | 1874352.467 | 2178329.728 | 1984616.404 | 1271123.717 | 2319412.489 | 501194.0313 | 2406432.896 | 39.6308 | 55.9788 | Nop58 | 55989 | 1.21317 | 0.786833712 | 0.64858 | 0.23885 | 0.732938 |
| 568951059 | PREDICTED: tripartite motif-containing protein 3 isoform X1 [Mus musculus] | 0 | 29.337 | 13 | 8 | 21 | 8 | 750 | 81.5 | 7.71 | 87 | 8 | 1704841.482 | 2916432.463 | 13769119.97 | 8424883.247 | 23234491.74 | 6316919.859 | 5097202.625 | 25958803.81 | 82.7022 | 72.4253 | Trim3 | 55992 | 0.61346 | 1.386537395 | 2.26018 | 0.21849 | 0.716066 |
| 9256519 | 60S acidic ribosomal protein P1 [Mus musculus] | 0 | 47.408 | 66 | 3 | 22 | 3 | 114 | 11.5 | 4.32 | 975 | 3 | 33504035.17 | 7530213.295 | 33175670.5 | 18036209.41 | 25619547.34 | 12843680.98 | 27789833.5 | 6537815.52 | 54.7245 | 56.0143 | Rplp1 | 56040 | 1.11788 | 0.882115826 | 0.78909 | 0.57069 | 0.895889 |
| 568936047 | PREDICTED: general vesicular transport factor p115 isoform X1 [Mus musculus] | 0.001 | 5.789 | 3 | 3 | 4 | 3 | 966 | 107.8 | 4.96 | 23 | 3 | 789253.1907 | 1183033.242 | 1131229.881 | 571264.377 | 1156778.189 | 1832892.429 | 1175369 | 1176152.854 | 31.5796 | 24.8522 | Uso1 | 56041 | 0.81517 | 1.184828831 | 1.45347 | 0.10761 | 0.678142 |
| 13591862 | protein SET isoform 1 [Mus musculus] | 0 | 51.996 | 38 | 9 | 68 | 7 | 289 | 33.4 | 4.32 | 841 | 9 | 23050042.06 | 30871913.16 | 31198296.02 | 42468743.46 | 32231450.41 | 20364422.19 | 25760143.13 | 42264249.2 | 25.0524 | 31.233 | Set | 56086 | 1.02808 | 0.971923973 | 0.94538 | 0.78735 | 0.952792 |
| 103472019 | general receptor for phosphoinositides 1-associated scaffold protein [Mus musculus] | 0 | 107.764 | 48 | 12 | 76 | 12 | 392 | 42.3 | 9.1 | 1733 | 12 | 57883542.63 | 65767910.89 | 53092042.58 | 59919965.7 | 72792911.88 | 67707109.6 | 64963714.81 | 75942133.81 | 8.87357 | 7.02313 | Grasp | 56149 | 0.91364 | 1.086363746 | 1.18905 | 0.02103 | 0.505785 |
| 13124378 | ltName: Full=Neuronal olfactomedin-related ER localized protein; AltName: Full=Olfactomedin-1; AltName: Full=Pa | 0 | 97.258 | 29 | 12 | 89 | 12 | 485 | 55.4 | 6.95 | 946 | 12 | 227890756.2 | 300152599.1 | 262671721.1 | 321573217.4 | 227011796.7 | 360445041.7 | 267098583.8 | 314192658.8 | 14.8786 | 19.78 | Olfm1 | 56177 | 0.97525 | 1.024751814 | 1.05076 | 0.70498 | 0.933129 |
| 9625037 | rho-related GTP-binding protein RhoG precursor [Mus musculus] | 0 | 67.521 | 70 | 11 | 99 | 10 | 191 | 21.3 | 8.12 | 2486 | 11 | 72905554.98 | 75769579.08 | 73267411.33 | 65726511.68 | 71137679.78 | 57879972.64 | 55142091.88 | 64196382.94 | 6.00573 | 11.4754 | Rhog | 56212 | 1.07334 | 0.926658428 | 0.86334 | 0.05635 | 0.599194 |
| 15488756 | HtrA serine peptidase 1 [Mus musculus] | 0 | 14.338 | 9 | 5 | 27 | 5 | 480 | 51.2 | 7.81 | 98 | 5 | 17533306.51 | 24798116.07 | 17215065.02 | 17471725.96 | 20533540.44 | 22687618.7 | 18188615.13 | 21237986.76 | 19.2073 | 9.08418 | Htra1 | 56213 | 0.96474 | 1.035258279 | 1.07309 | 0.52263 | 0.882454 |
| 568951073 | PREDICTED: syntaxin-1B isoform X1 [Mus musculus] | 0 | 65.354 | 27 | 10 | 48 | 10 | 406 | 46.4 | 6.18 | 1102 | 10 | 59276913.33 | 82784268.66 | 53097588.84 | 55101112.7 | 86414499.67 | 82216895.96 | 71147731.25 | 55600892.1 | 21.9342 | 18.6368 | Stx1b | 56216 | 0.91731 | 1.082692148 | 1.18029 | 0.28982 | 0.753767 |
| 9625023 | MAGUK p55 subfamily member 5 [Mus musculus] | 0 | 11.865 | 6 | 2 | 2 | 2 | 675 | 77.2 | 6.09 | 29 | 2 | 491467.8994 | 2126544.497 | 1912700.091 | 2202282.626 | 2571270.548 | 2524653.427 | 1737930.125 | 2511057.363 | 47.7605 | 17.1087 | Mpp5 | 56217 | 0.83755 | 1.162453758 | 1.38793 | 0.19601 | 0.715071 |
| 148688440 | heterogeneous nuclear ribonucleoprotein H2, isoform CRA_b, partial [Mus musculus] | 0 | 54.14 | 26 | 8 | 85 | 3 | 460 | 50.3 | 6.54 | 1037 | 8 | 13451575.93 | 10950129.94 | 15582924.16 | 13052264.48 | 11752597.77 | 10494096.05 | 15275686.5 | 12012948.56 | 14.3173 | 16.4633 | Hnrnph2 | 56258 | 1.03414 | 0.965862438 | 0.93398 | 0.55285 | 0.892955 |
| 12836486 | unnamed protein product [Mus musculus] | 0 | 56.579 | 17 | 12 | 71 | 9 | 669 | 69.4 | 9.72 | 872 | 12 | 50871147 | 70423193.37 | 55554509.84 | 66922947.45 | 54916993.88 | 72071544.48 | 56748545 | 82009918.69 | 15.1611 | 19.4483 | Rbm14 | 56275 | 0.95687 | 1.04312933 | 1.09015 | 0.51497 | 0.88031 |
| 148702808 | mitochondrial ribosomal protein L12, partial [Mus musculus] | 0 | 10.849 | 18 | 3 | 4 | 3 | 214 | 22.9 | 9.04 | 17 | 3 | 1421781.527 | 180578.9541 | 933173.7308 | 3728099.255 | 3464415.158 | 502618.8592 | 1077957.75 | 1578711.682 | 97.6556 | 77.4989 | Mrpl12 | 56282 | 0.97206 | 1.027939828 | 1.05749 | 0.93108 | 0.99428 |
| 293651556 | drebrin isoform 1 [Mus musculus] | 0 | 241.761 | 50 | 27 | 412 | 27 | 706 | 77.2 | 4.49 | 6688 | 27 | 815750213.1 | 804183310.1 | 889135647.8 | 1027158013 | 851341146.9 | 1030802451 | 864891122.8 | 946737647.7 | 11.5999 | 8.99434 | Dbn1 | 56320 | 0.97821 | 1.021790483 | 1.04455 | 0.57238 | 0.895889 |
| 568927081 | PREDICTED: dnaJ homolog subfamily B member 5 isoform X3 [Mus musculus] | 0 | 67.259 | 34 | 12 | 56 | 12 | 420 | 47.2 | 9.38 | 939 | 12 | 64744595.15 | 61771305.88 | 65398129.23 | 69204377.73 | 67063813.74 | 75270623.61 | 68024638.5 | 82615448.16 | 4.68097 | 9.88843 | Dnajb5 | 56323 | 0.94251 | 1.05749237 | 1.122 | 0.08913 | 0.662454 |
| 19263839 | Eukaryotic translation initiation factor 3, subunit C [Mus musculus] | 0 | 25.488 | 8 | 7 | 24 | 7 | 911 | 105.5 | 5.83 | 325 | 7 | 11051463.31 | 4863682.185 | 10830058.17 | 9067646.799 | 4689337.213 | 5755327.282 | 8904450.25 | 8300096.343 | 32.0243 | 29.1439 | Eif3c | 56347 | 1.12864 | 0.871361894 | 0.77205 | 0.28829 | 0.752733 |
| 74204320 | unnamed protein product [Mus musculus] | 0 | 63.292 | 42 | 9 | 30 | 9 | 312 | 34.7 | 9.58 | 638 | 9 | 34776988.26 | 20394651.15 | 44483415.07 | 21386705.88 | 57795116.9 | 20038239.58 | 52991732.5 | 36833725.96 | 38.1 | 40.8449 | Hsd17b12 | 56348 | 0.83853 | 1.161471984 | 1.38513 | 0.3019 | 0.761857 |
| 31980998 | acyl-coenzyme A thioesterase 9, mitochondrial isoform 1 [Mus musculus] | 0 | 101.542 | 44 | 16 | 131 | 16 | 439 | 50.5 | 8.59 | 1009 | 16 | 68794315.61 | 67750451.73 | 81877840.67 | 71395621.03 | 84011222.33 | 59286537.03 | 93223776.42 | 114425484.5 | 8.92486 | 26.0382 | Acot9 | 56360 | 0.9046 | 1.095399667 | 1.21092 | 0.24541 | 0.733806 |
| 9790125 | transgelin-3 [Mus musculus] | 0 | 26.248 | 30 | 5 | 15 | 5 | 199 | 22.5 | 7.33 | 274 | 5 | 16624438.75 | 7014523.278 | 9530988.127 | 10804629.07 | 8762017.593 | 8478527.944 | 7851332.188 | 13777652.34 | 37.0287 | 28.1296 | Tagln3 | 56370 | 1.06162 | 0.938377644 | 0.88391 | 0.62131 | 0.908209 |
| 568923335 | PREDICTED: PDZ and LIM domain protein 5 isoform X1 [Mus musculus] | 0 | 27.259 | 12 | 7 | 28 | 7 | 734 | 78.1 | 8.48 | 143 | 7 | 8719321.687 | 13821520.88 | 10109115.42 | 4670964.296 | 15811200.28 | 19222966.62 | 17684706.75 | 24982870.46 | 40.5112 | 20.38 | Pdlim5 | 56376 | 0.64893 | 1.351068386 | 2.08199 | 0.01022 | 0.449633 |
| 9790141 | actin-related protein 2/3 complex subunit 3 [Mus musculus] | 0 | 23.293 | 27 | 6 | 51 | 6 | 178 | 20.5 | 8.59 | 283 | 6 | 67558585.28 | 90953686.28 | 79946543.27 | 107915294.9 | 91440895.77 | 100990196.6 | 67521201.94 | 89829853.94 | 19.7802 | 16.2006 | Arpc3 | 56378 | 0.9951 | 1.004895508 | 1.00984 | 0.94139 | 0.994906 |
| 9789997 | LETM1 and EF-hand domain-containing protein 1, mitochondrial precursor [Mus musculus] | 0 | 32.694 | 14 | 7 | 14 | 7 | 738 | 82.9 | 6.52 | 43 | 7 | 17249692.42 | 7157358.98 | 9426942.294 | 10203918.26 | 10119888.99 | 14175162.34 | 8504486 | 10606862.98 | 39.5668 | 22.0334 | Letm1 | 56384 | 1.00722 | 0.992778128 | 0.98566 | 0.9514 | 0.994906 |
| 31543701 | leucine-rich repeat protein SHOC-2 [Mus musculus] | 0 | 25.51 | 12 | 4 | 18 | 4 | 582 | 64.9 | 8.46 | 482 | 4 | 33591901.56 | 34246464.07 | 38688927.44 | 28619240.81 | 32725146.84 | 48409345.71 | 30789734 | 56438963.77 | 12.2014 | 29.4583 | Shoc2 | 56392 | 0.89056 | 1.109441819 | 1.24578 | 0.25076 | 0.733806 |
| 9790225 | calcineurin B homologous protein 1 [Mus musculus] | 0 | 14.895 | 26 | 4 | 15 | 4 | 195 | 22.4 | 5.1 | 135 | 4 | 6399695.598 | 3534822.483 | 5545374.361 | 1802441.617 | 6765177.099 | 6926097.113 | 6612980.625 | 6130974.443 | 47.7732 | 5.19386 | Chp1 | 56398 | 0.79064 | 1.209364258 | 1.52961 | 0.07135 | 0.610935 |
| 33416526 | Syncrip protein, partial [Mus musculus] | 0 | 41.545 | 17 | 10 | 50 | 7 | 667 | 74.7 | 9.03 | 340 | 10 | 35774565.33 | 38311133.83 | 44831279.14 | 18862264.49 | 29368953.92 | 39579926.54 | 40599039.75 | 42093914.08 | 32.1286 | 15.2654 | Syncrip | 56403 | 0.9521 | 1.047897657 | 1.10061 | 0.59895 | 0.905186 |
| 31127103 | Dusp14 protein [Mus musculus] | 0 | 7.594 | 14 | 2 | 8 | 2 | 198 | 22.3 | 9.26 | 96 | 2 | 8640805.192 | 3704850.006 | 7737650.517 | 4617982.556 | 9824338.572 | 5586493.431 | 5843460 | 5147294.979 | 38.6026 | 32.853 | Dusp14 | 56405 | 0.96673 | 1.033272075 | 1.06883 | 0.80075 | 0.959776 |
| 148704285 | cerebellin 3 precursor protein, isoform CRA_b, partial [Mus musculus] | 0 | 8.285 | 12 | 2 | 5 | 2 | 211 | 22.8 | 6.79 | 100 | 2 | 11314668.66 | 11404888.67 | 13587886.74 | 18258383.49 | 11887402.47 | 15834373.32 | 13037423 | 20365247.43 | 23.8423 | 24.6904 | Cbln3 | 56410 | 0.94331 | 1.056691185 | 1.1202 | 0.53476 | 0.886555 |
| 597517915 | ATP-dependent 6-phosphofructokinase, platelet type isoform 2 [Mus musculus] | 0 | 172.005 | 38 | 23 | 169 | 14 | 784 | 85.5 | 6.89 | 2887 | 23 | 58810957.85 | 34210575.89 | 54767656.48 | 31894742.27 | 52113797.67 | 45325018.48 | 47027691.17 | 42602837.96 | 30.8002 | 8.55973 | Pfkp | 56421 | 0.97986 | 1.020137278 | 1.0411 | 0.80622 | 0.961747 |
| 12832963 | unnamed protein product [Mus musculus] | 0 | 9.403 | 11 | 3 | 7 | 3 | 304 | 34.9 | 6.01 | 30 | 3 | 1522126.195 | 2031263.854 | 3039765.452 | 1690146.343 | 4565178.038 | 2756947.9 | 4244991.188 | 2798255.693 | 32.8276 | 26.3653 | Stub1 | 56424 | 0.73169 | 1.268310348 | 1.7334 | 0.0403 | 0.582811 |
| 954794740 | mitochondrial carrier homolog 2 isoform 1x [Mus musculus] | 0 | 88.301 | 48 | 12 | 102 | 12 | 314 | 34.6 | 8.32 | 1870 | 12 | 308770725.8 | 277544665 | 295597629.5 | 264866583.4 | 348116199.4 | 384313988.3 | 307661953.5 | 322481630.5 | 6.75995 | 9.85452 | Mtch2 | 56428 | 0.914 | 1.085995926 | 1.18817 | 0.03185 | 0.515682 |
| 148696488 | destrin, isoform CRA_b [Mus musculus] | 0 | 11.566 | 18 | 4 | 7 | 3 | 222 | 25 | 8.76 | 43 | 4 | 163380034 | 315464098.6 | 335964156.9 | 217177147.4 | 295504429.4 | 462813903.5 | 300057907.8 | 513864996.2 | 31.6476 | 28.4921 | Dstn | 56431 | 0.79255 | 1.207453447 | 1.52351 | 0.09919 | 0.671097 |
| 13124572 | RecName: Full=Sorting nexin-1 | 0 | 7.719 | 5 | 2 | 6 | 2 | 522 | 58.9 | 5.22 | 22 | 2 | 14998537.09 | 18271189.35 | 16695586.03 | 17526871.06 | 15204448.03 | 19261547.37 | 16102105 | 16161876.63 | 8.33075 | 10.6355 | Snx1 | 56440 | 1.00568 | 0.994321307 | 0.98871 | 0.87183 | 0.979509 |
| 9790221 | actin-related protein 2/3 complex subunit 1A [Mus musculus] | 0 | 165.672 | 65 | 19 | 157 | 18 | 370 | 41.6 | 8.18 | 2230 | 19 | 420835107.4 | 420921387.3 | 383563848.3 | 415418855.8 | 401247789.4 | 420341701.3 | 399417123.6 | 448166430.1 | 4.37192 | 5.42853 | Arpc1a | 56443 | 0.99141 | 1.008590513 | 1.01733 | 0.64015 | 0.915433 |
| 226246593 | actin-related protein 10 [Mus musculus] | 0 | 53.896 | 32 | 9 | 35 | 9 | 417 | 46.2 | 7.61 | 525 | 9 | 12151461.75 | 13389049.73 | 17476663 | 10812747.58 | 19692213.46 | 14440533.05 | 15414376.69 | 12631369.41 | 21.3902 | 19.2726 | Actr10 | 56444 | 0.92803 | 1.071965215 | 1.15509 | 0.35382 | 0.798662 |
| 9789937 | dnaJ homolog subfamily A member 2 [Mus musculus] | 0 | 192.055 | 61 | 20 | 227 | 2 | 412 | 45.7 | 6.48 | 4803 | 20 | 10888002.23 | 5139923.692 | 12562671.21 | 3048819.058 | 6231689.727 | 2133735.806 | 5407710.5 | 2508049.326 | 57.39 | 50.4521 | Dnaja2 | 56445 | 1.32049 | 0.679506719 | 0.51459 | 0.17419 | 0.713031 |
| 255958286 | succinyl-CoA ligase [ADP/GDP-forming] subunit alpha, mitochondrial precursor [Mus musculus] | 0 | 26.865 | 20 | 5 | 23 | 5 | 346 | 36.1 | 9.39 | 461 | 5 | 28972268.17 | 17254511.57 | 51859026.5 | 40633011.28 | 34497185.73 | 35937594.57 | 38718688 | 31729852.55 | 42.9889 | 8.2711 | Suclg1 | 56451 | 0.99226 | 1.007741369 | 1.0156 | 0.94552 | 0.994906 |
| 4505813 | dynein light chain 1, cytoplasmic [Homo sapiens] | 0 | 102.031 | 84 | 9 | 132 | 4 | 89 | 10.4 | 7.4 | 1806 | 9 | 288315026 | 196301205.2 | 265631598.4 | 181279495.2 | 229762002.9 | 226755814.3 | 225687049.5 | 206225328.5 | 22.3761 | 4.82999 | Dynll1 | 56455 | 1.02368 | 0.976319706 | 0.95373 | 0.6995 | 0.931521 |
| 9845273 | mitochondrial carrier homolog 1 [Mus musculus] | 0 | 95.186 | 42 | 13 | 76 | 13 | 389 | 41.5 | 9.32 | 1253 | 13 | 70314771.45 | 71384026.35 | 56813813.77 | 64422612.64 | 79098142.48 | 129911581.5 | 75578957.5 | 81170326.39 | 10.1747 | 28.1623 | Mtch1 | 56462 | 0.83645 | 1.163551339 | 1.39106 | 0.10151 | 0.674486 |
| 74204693 | unnamed protein product [Mus musculus] | 0 | 204.669 | 46 | 29 | 167 | 29 | 910 | 102 | 7.56 | 2148 | 29 | 142046430.6 | 143365770.9 | 101296721.6 | 156092695.8 | 126099373.6 | 132432442 | 130924127 | 142162769 | 17.5343 | 5.0692 | Snd1 | 56463 | 1.01041 | 0.989591683 | 0.9794 | 0.82863 | 0.968871 |
| 9790071 | fatty acid desaturase 2 [Mus musculus] | 0 | 9.876 | 7 | 3 | 9 | 3 | 444 | 52.4 | 8.82 | 36 | 3 | 2662909.817 | 1982062.932 | 3342437.595 | 645079.1662 | 4404670.525 | 2210636.262 | 2753688 | 958427.4593 | 53.3556 | 55.345 | Fads2 | 56473 | 0.9106 | 1.089395603 | 1.19634 | 0.66049 | 0.919605 |
| 148692474 | TANK-binding kinase 1, isoform CRA_b, partial [Mus musculus] | 0 | 99.887 | 28 | 15 | 79 | 15 | 732 | 83.7 | 6.8 | 1775 | 15 | 58290658.85 | 48256283 | 68477905.22 | 79994450.37 | 75061226.56 | 87549710.4 | 60866539.5 | 65139380.42 | 21.3551 | 16.4392 | Tbk1 | 56480 | 0.9382 | 1.061801555 | 1.13175 | 0.38811 | 0.81965 |
| 24638341 | embrane protein-associated protein B; Short=VAMP-B; Short=VAMP-associated protein B; Short=VAP-B; AltNam | 0 | 27.664 | 23 | 5 | 14 | 4 | 243 | 26.9 | 7.78 | 134 | 5 | 11258880.15 | 8006213.328 | 8363036.526 | 10032051.74 | 10077193.63 | 8485468.915 | 8008625.25 | 8074261.4 | 16.0752 | 11.1665 | Vapb | 56491 | 1.04169 | 0.958307142 | 0.91995 | 0.43351 | 0.849034 |
| 74226963 | unnamed protein product [Mus musculus] | 0 | 34.653 | 23 | 8 | 29 | 8 | 456 | 50.2 | 6.42 | 134 | 8 | 15274010.14 | 10244655.63 | 19060139.98 | 17115784.61 | 17405432.82 | 13540112.78 | 17410530.75 | 11495897.1 | 24.5268 | 19.6752 | Ruvbl1 | 56505 | 1.01516 | 0.984840239 | 0.97013 | 0.85393 | 0.975398 |
| 323637461 | rap guanine nucleotide exchange factor 4 isoform 1 [Mus musculus] | 0 | 359.2 | 56 | 48 | 608 | 48 | 1011 | 115.4 | 6.92 | 10312 | 48 | 1062249611 | 1062005211 | 1015578336 | 1055194972 | 1018519001 | 1209357604 | 1095402624 | 1186510412 | 2.132 | 7.78135 | Rapgef4 | 56508 | 0.96384 | 1.03615946 | 1.07503 | 0.13281 | 0.693437 |
| 257900522 | MAGUK p55 subfamily member 6 isoform a [Mus musculus] | 0 | 15.803 | 10 | 5 | 18 | 4 | 553 | 62.6 | 6.4 | 63 | 5 | 8529211.312 | 18406826.31 | 26637342.36 | 16571018.05 | 22175035.88 | 30285267.56 | 20203660.13 | 38740922.14 | 42.3738 | 30.411 | Mpp6 | 56524 | 0.77273 | 1.227268799 | 1.58822 | 0.11683 | 0.678142 |
| 50510373 | mKIAA0128 protein [Mus musculus] | 0 | 100.295 | 41 | 15 | 74 | 7 | 449 | 50.7 | 6.2 | 1164 | 15 | 33607880.12 | 29381717.28 | 35482199.07 | 25811673.85 | 32471113.03 | 33723180.59 | 37735486.63 | 50167512.46 | 13.9559 | 20.9748 | 43714 | 56526 | 0.8929 | 1.10709728 | 1.23989 | 0.15517 | 0.698266 |
| 125628634 | microtubule-associated serine/threonine-protein kinase 1 [Mus musculus] | 0 | 86.995 | 17 | 18 | 53 | 16 | 1570 | 170.9 | 8.44 | 640 | 18 | 40018875.8 | 20486271.17 | 35614439.33 | 19665138.64 | 45981887.67 | 27445593.81 | 33407989.94 | 29597258.24 | 35.9452 | 24.3075 | Mast1 | 56527 | 0.91813 | 1.081865883 | 1.17833 | 0.46722 | 0.863085 |
| 148675018 | Sec11-like 1 (S. cerevisiae), isoform CRA_d [Mus musculus] | 0 | 11.443 | 12 | 3 | 6 | 3 | 247 | 27.9 | 7.68 | 100 | 3 | 9513423.141 | 9786602.667 | 4004076.241 | 8987005.814 | 8537062.516 | 5493422.879 | 11024222 | 12280380.91 | 33.8507 | 32.0956 | Sec11a | 56529 | 0.92756 | 1.072443717 | 1.1562 | 0.55684 | 0.895143 |
| 160333783 | intracellular hyaluronan-binding protein 4 [Mus musculus] | 0 | 30.094 | 23 | 6 | 24 | 6 | 412 | 45.9 | 6.84 | 176 | 6 | 27672570.65 | 14727164.96 | 34647544.47 | 32347580.82 | 39504540.64 | 12060186.55 | 19529051.25 | 25382908.2 | 32.5454 | 48.1609 | Habp4 | 56541 | 1.06275 | 0.937251289 | 0.88191 | 0.6744 | 0.923206 |
| 568995942 | PREDICTED: glycogen synthase kinase-3 beta isoform X1 [Mus musculus] | 0 | 126.452 | 61 | 19 | 134 | 14 | 433 | 48 | 8.78 | 1903 | 19 | 203839594.9 | 190638048.7 | 190851183.5 | 209756802.7 | 208942859 | 156116197 | 190390912.1 | 200819932.1 | 4.81889 | 12.2934 | Gsk3b | 56637 | 1.02502 | 0.974979475 | 0.95118 | 0.46941 | 0.863085 |
| 37747507 | Pnkd protein [Mus musculus] | 0 | 22.423 | 13 | 4 | 12 | 4 | 424 | 47.4 | 9.11 | 147 | 4 | 5120776.16 | 2787101.928 | 4882966.818 | 2292033.698 | 5428117.681 | 2121426.943 | 4979111.563 | 2300175.453 | 38.1674 | 46.9121 | Pnkd | 56695 | 1.00849 | 0.991506772 | 0.98316 | 0.95695 | 0.994906 |
| 21426893 | histone H1.5 [Mus musculus] | 0 | 17.847 | 22 | 5 | 39 | 5 | 223 | 22.6 | 10.92 | 167 | 5 | 1447564.19 | 7580071.037 | 30757953.73 | 33691188.37 | 15393877.64 | 44406923.2 | 30340717.16 | 66870843.13 | 88.3965 | 55.7756 | Hist1h1b | 56702 | 0.63757 | 1.36242742 | 2.1369 | 0.17633 | 0.713031 |
| 26349771 | unnamed protein product [Mus musculus] | 0.001 | 5.286 | 6 | 2 | 2 | 2 | 378 | 42.4 | 8.5 | 27 | 2 | 1187009.62 | 1515174.06 | 1367364.132 | 2026960.044 | 1229214.051 | 1645555.703 | 1518815.375 | 1786146.193 | 23.6912 | 15.3479 | Dnajb12 | 56709 | 0.99322 | 1.006779231 | 1.01365 | 0.9264 | 0.990922 |
| 6573173 | DBCCR1 [Mus musculus] | 0 | 43.426 | 17 | 12 | 30 | 12 | 761 | 88.7 | 9.01 | 153 | 12 | 39761668.27 | 26476622.79 | 42815527.16 | 25924948.14 | 31770373.77 | 39914015.42 | 44148183.09 | 34480644.98 | 26.0859 | 14.7323 | Brinp1 | 56710 | 0.94625 | 1.053750024 | 1.11361 | 0.48875 | 0.869525 |
| 1039754011 | PREDICTED: target of rapamycin complex subunit LST8 isoform X1 [Mus musculus] | 0 | 14.179 | 13 | 3 | 9 | 3 | 328 | 36 | 5.86 | 139 | 3 | 2943000.598 | 2768624.454 | 3123735.663 | 3392454.198 | 2210760.565 | 1620185.378 | 3151160.75 | 1303810.911 | 8.71932 | 39.1996 | Mlst8 | 56716 | 1.19216 | 0.80784105 | 0.67763 | 0.06059 | 0.601634 |
| 148682872 | FK506 binding protein 12-rapamycin associated protein 1, isoform CRA_a, partial [Mus musculus] | 0 | 50.313 | 6 | 13 | 43 | 13 | 2580 | 292.3 | 7.52 | 247 | 13 | 21814379.74 | 19411910.24 | 28390809.23 | 14319588.64 | 23437540.17 | 24639901.31 | 23197527 | 20765212.46 | 27.8467 | 7.05845 | Mtor | 56717 | 0.95395 | 1.046048626 | 1.09654 | 0.52894 | 0.886555 |
| 148678139 | ring finger protein 14, isoform CRA_c, partial [Mus musculus] | 0 | 12.267 | 6 | 2 | 14 | 2 | 505 | 57.1 | 4.68 | 429 | 2 | 6200815.377 | 3506785.676 | 7086573.878 | 1065493.288 | 5268018.488 | 4387515.563 | 3727694 | 3230100.339 | 61.1458 | 21.2229 | Rnf14 | 56736 | 1.03615 | 0.96384591 | 0.93021 | 0.83524 | 0.969252 |
| 12836608 | unnamed protein product [Mus musculus] | 0 | 87.557 | 41 | 12 | 75 | 12 | 415 | 47.3 | 7.97 | 1497 | 12 | 111839052.3 | 74010954.65 | 111200698.3 | 68137348.27 | 99374270.89 | 75475640.32 | 102634698.7 | 38342514.13 | 25.7134 | 37.5649 | Alg2 | 56737 | 1.07248 | 0.927518606 | 0.86483 | 0.53828 | 0.886555 |
| 568923410 | PREDICTED: muscleblind-like protein 1 isoform X1 [Mus musculus] | 0.001 | 6.867 | 4 | 2 | 10 | 2 | 436 | 47.1 | 8.62 | 40 | 2 | 11664237.49 | 8631972.798 | 13147186.9 | 9080331.485 | 11683816.87 | 13195064.56 | 13120930.5 | 9730245.752 | 20.1743 | 13.6151 | Mbnl1 | 56758 | 0.94231 | 1.057685436 | 1.12243 | 0.37065 | 0.80688 |
| 229577332 | dnaJ homolog subfamily B member 2 isoform 3 [Mus musculus] | 0 | 26.246 | 19 | 4 | 81 | 3 | 324 | 35.6 | 5.91 | 830 | 4 | 7941844.7 | 5284405.319 | 10904431.21 | 7545883.445 | 8971944.398 | 7302329.653 | 8203645.938 | 5799761.725 | 29.1565 | 18.0051 | Dnajb2 | 56812 | 1.02258 | 0.977420709 | 0.95584 | 0.80291 | 0.959848 |
| 9938002 | leucine-rich glioma-inactivated protein 1 precursor [Mus musculus] | 0 | 73.237 | 33 | 15 | 89 | 2 | 557 | 63.6 | 8.02 | 1090 | 15 | 26955582.18 | 13602219.09 | 15783924.41 | 12964504.14 | 17115118.78 | 6576260.064 | 16722326 | 7111662.522 | 37.6986 | 49.0094 | Lgi1 | 56839 | 1.18643 | 0.813570447 | 0.68573 | 0.25971 | 0.745606 |
| 1039761441 | PREDICTED: NMDA receptor synaptonuclear signaling and neuronal migration factor isoform X1 [Mus musculus] | 0 | 10.467 | 7 | 3 | 5 | 3 | 590 | 66.7 | 8.95 | 65 | 3 | 9792054.616 | 2881862.37 | 7640505.249 | 3136893.042 | 3352411.486 | 2786818.937 | 2768209.75 | 2456036.219 | 58.1893 | 13.139 | Nsmf | 56876 | 1.3472 | 0.652795886 | 0.48456 | 0.1287 | 0.693437 |
| 28972652 | mKIAA1176 protein [Mus musculus] | 0 | 135.858 | 25 | 22 | 163 | 22 | 1164 | 128.6 | 6.81 | 1989 | 22 | 189217023.6 | 167133435 | 140981148.2 | 129242933.3 | 177013129 | 159158518.4 | 155907722.2 | 96776643.93 | 17.1583 | 23.6945 | Slc12a5 | 57138 | 1.03103 | 0.968966942 | 0.9398 | 0.6834 | 0.927189 |
| 568938514 | PREDICTED: PERQ amino acid-rich with GYF domain-containing protein 1 isoform X1 [Mus musculus] | 0 | 20.284 | 6 | 5 | 11 | 5 | 1075 | 119 | 5.48 | 39 | 5 | 23567645.78 | 20123884.06 | 20646227.34 | 19221733.1 | 24380195.95 | 17595801.8 | 21533388.69 | 19377634.09 | 8.9979 | 14.103 | Gigyf1 | 57330 | 1.00404 | 0.995959845 | 0.99195 | 0.92606 | 0.990922 |
| 31981106 | mannosyl-oligosaccharide glucosidase [Mus musculus] | 0 | 58.536 | 17 | 9 | 37 | 9 | 834 | 91.8 | 9 | 705 | 9 | 20307665.32 | 7475663.26 | 22997209.54 | 21201475.26 | 17988103.02 | 34914670.95 | 20971883.13 | 27300683.41 | 39.4645 | 29.6419 | Mogs | 57377 | 0.83141 | 1.168594209 | 1.40556 | 0.20725 | 0.715071 |
| 74195524 | unnamed protein product [Mus musculus] | 0 | 14.061 | 19 | 4 | 37 | 2 | 121 | 14.5 | 9 | 54 | 4 | 6312655.325 | 3314686.099 | 2829862.815 | 8726112.411 | 2228558.032 | 8063115.927 | 2304787.766 | 7176241.198 | 52.0639 | 62.9525 | Gabarapl1 | 57436 | 1.03444 | 0.965557841 | 0.93341 | 0.87087 | 0.979509 |
| 148686123 | cell cycle exit and neuronal differentiation 1, isoform CRA_b, partial [Mus musculus] | 0 | 66.886 | 59 | 7 | 34 | 7 | 174 | 17.6 | 8.88 | 268 | 7 | 39245605.79 | 51233178.74 | 43858762.7 | 53197588.96 | 56100881.68 | 68129125.4 | 44067714.25 | 55368044.74 | 13.838 | 17.5795 | Cend1 | 57754 | 0.91213 | 1.087866125 | 1.19266 | 0.17596 | 0.713031 |
| 148665412 | mCG130125, isoform CRA_b, partial [Mus musculus] | 0.001 | 4.546 | 12 | 2 | 10 | 2 | 130 | 14.5 | 11.06 | 70 | 2 | 66701817.46 | 135796658 | 62094685.14 | 98524145.88 | 64825902.86 | 795152.171 | 69786646.81 | 112683291.3 | 37.5671 | 74.3731 | Rpl35a | 57808 | 1.18819 | 0.811805053 | 0.68323 | 0.35477 | 0.798662 |
| 171184435 | very-long-chain (3R)-3-hydroxyacyl-CoA dehydratase 3 [Mus musculus] | 0 | 62.49 | 30 | 8 | 33 | 8 | 362 | 43.1 | 9.13 | 496 | 8 | 60739799.13 | 45413027.3 | 55827323.19 | 43946701.37 | 55924724.8 | 53252257.95 | 61628570 | 43461949.28 | 15.7885 | 14.1665 | Hacd3 | 57874 | 0.98015 | 1.019849508 | 1.0405 | 0.72054 | 0.939018 |
| 569012144 | PREDICTED: SH3 domain-containing kinase-binding protein 1 isoform X1 [Mus musculus] | 0 | 140.113 | 46 | 22 | 103 | 22 | 709 | 78.1 | 7.55 | 2511 | 22 | 159718492 | 144058044.7 | 170310340.4 | 144858201.2 | 178554857.8 | 177049769.9 | 173644958.3 | 188044974.5 | 8.16589 | 3.43866 | Sh3kbp1 | 58194 | 0.9264 | 1.07360168 | 1.1589 | 0.01287 | 0.461382 |
| 568961808 | PREDICTED: dnaJ homolog subfamily A member 4 isoform X1 [Mus musculus] | 0 | 24.981 | 12 | 6 | 37 | 6 | 555 | 61.8 | 8.46 | 193 | 6 | 12937107.83 | 18917079.78 | 15378574.67 | 14224571.88 | 10844671.14 | 12940285.36 | 10924207 | 11736565.8 | 16.7262 | 8.38192 | Dnaja4 | 58233 | 1.13912 | 0.860878789 | 0.75574 | 0.03412 | 0.528235 |
| 359465599 | acidic fibroblast growth factor intracellular-binding protein isoform 1 [Mus musculus] | 0 | 11.869 | 10 | 3 | 3 | 3 | 364 | 41.8 | 6.48 | 55 | 3 | 2826167.324 | 520612.6686 | 4183288.041 | 1566909.453 | 1070083.044 | 1202451.615 | 1397533.625 | 1344657.72 | 69.6377 | 11.7684 | Fibp | 58249 | 1.28928 | 0.710718732 | 0.55125 | 0.24674 | 0.733806 |
| 755502742 | PREDICTED: PEX5-related protein isoform X1 [Mus musculus] | 0 | 21.317 | 13 | 7 | 17 | 7 | 673 | 74.8 | 5.5 | 83 | 7 | 3978172.969 | 10645861.49 | 8186714.569 | 7250031.45 | 4147617.054 | 7374972.581 | 6238947.75 | 7147480.889 | 36.7096 | 23.6176 | Pex5l | 58869 | 1.09372 | 0.906280133 | 0.82862 | 0.44149 | 0.852048 |
| 568975090 | PREDICTED: rho GTPase-activating protein 23 isoform X4 [Mus musculus] | 0 | 89.196 | 16 | 18 | 79 | 17 | 1489 | 162.2 | 9.03 | 879 | 18 | 29316878.78 | 28290252.36 | 29408536.73 | 26093133.69 | 28013179.32 | 21307487.76 | 23862498.09 | 24417361.16 | 5.45235 | 11.325 | Arhgap23 | 58996 | 1.0736 | 0.926399674 | 0.86289 | 0.04976 | 0.592616 |
| 133777819 | Trappc2l protein, partial [Mus musculus] | 0.001 | 5.966 | 15 | 2 | 12 | 2 | 140 | 16.1 | 7.33 | 82 | 2 | 5420816.483 | 6433393.268 | 5296600.793 | 5700295.207 | 5908766.731 | 5681467.334 | 5666260.5 | 1754989.18 | 8.91354 | 42.1148 | Trappc2l | 59005 | 1.09172 | 0.908280351 | 0.83197 | 0.3885 | 0.81965 |
| 148701752 | heterogeneous nuclear ribonucleoprotein H1, isoform CRA_b, partial [Mus musculus] | 0 | 65.587 | 25 | 8 | 84 | 2 | 485 | 52.5 | 6.67 | 1065 | 8 | 13505365.09 | 20947927.1 | 14148982.02 | 16904250.32 | 1474908.919 | 14629592.28 | 11897322.94 | 10655247.3 | 20.6718 | 59.0463 | Hnrnph1 | 59013 | 1.25776 | 0.742237652 | 0.59013 | 0.08949 | 0.662454 |
| 148695543 | nucleoporin 160, isoform CRA_b, partial [Mus musculus] | 0 | 120.71 | 20 | 18 | 82 | 18 | 1420 | 160 | 5.62 | 1696 | 18 | 20730607.35 | 23395038.38 | 41877661.74 | 30439906.63 | 33956870.46 | 27182607.12 | 42540887.44 | 29057360.95 | 32.447 | 20.6735 | Nup160 | 59015 | 0.93461 | 1.065392288 | 1.13994 | 0.51137 | 0.88031 |
| 568975132 | PREDICTED: mitochondrial Rho GTPase 1 isoform X1 [Mus musculus] | 0 | 39.659 | 19 | 11 | 30 | 11 | 752 | 86.1 | 6.67 | 121 | 11 | 33623186.37 | 30956992.21 | 36271467.33 | 26860368.41 | 46008997.81 | 43169927.43 | 42463387 | 53772608.47 | 12.7555 | 11.7703 | Rhot1 | 59040 | 0.81572 | 1.1842796 | 1.45182 | 0.00455 | 0.38143 |
| 34576555 | serine/threonine-protein kinase 25 [Rattus norvegicus] | 0 | 27.14 | 15 | 6 | 17 | 2 | 426 | 48.1 | 6.81 | 166 | 6 | 2403773.757 | 2771882.495 | 2535262.296 | 2912081.265 | 2353790.503 | 2305252.917 | 2806550.25 | 3560944.699 | 8.61948 | 21.105 | Stk25 | 59041 | 0.98136 | 1.018639592 | 1.03799 | 0.75785 | 0.952737 |
| 54607114 | protein LAP2 isoform 1 [Mus musculus] | 0 | 61.837 | 12 | 13 | 83 | 10 | 1450 | 162.1 | 5.86 | 1247 | 13 | 20048041.3 | 17257160.75 | 18259043.24 | 17248427.04 | 22297063.52 | 18908546.18 | 22435874 | 20186839.14 | 7.24173 | 8.15946 | Erbin | 59079 | 0.92968 | 1.070324186 | 1.15129 | 0.04343 | 0.588297 |
| 11140825 | trafficking protein particle complex subunit 4 [Mus musculus] | 0 | 20.904 | 27 | 5 | 17 | 5 | 219 | 24.4 | 6.21 | 232 | 5 | 2338009.169 | 4947303.41 | 4602134.505 | 2309590.599 | 6478077.58 | 4175191.944 | 5717054.125 | 4831086.073 | 40.0671 | 19.0135 | Trappc4 | 60409 | 0.80213 | 1.197872295 | 1.49337 | 0.09124 | 0.662454 |
| 31543799 | synaptotagmin-9 [Mus musculus] | 0 | 16.618 | 11 | 5 | 14 | 3 | 491 | 56.2 | 7.24 | 30 | 5 | 2842011.26 | 924487.6801 | 2957665.77 | 3024059.941 | 3094039.025 | 3788406.571 | 3340187.125 | 946594.4143 | 41.4919 | 45.253 | Syt9 | 60510 | 0.93207 | 1.067933824 | 1.14577 | 0.67602 | 0.924447 |
| 568946340 | PREDICTED: alpha-actinin-4 isoform X3 [Mus musculus] | 0 | 415.877 | 67 | 53 | 635 | 32 | 907 | 104.4 | 5.48 | 10752 | 53 | 585696397.6 | 465772278 | 534753887.5 | 439766826.9 | 584414502.7 | 441527741.4 | 576980358.8 | 475224208.1 | 13.0876 | 13.8611 | Actn4 | 60595 | 0.98729 | 1.012708501 | 1.02574 | 0.79882 | 0.958881 |
| 20988692 | Mitogen-activated protein kinase 8 interacting protein 2 [Mus musculus] | 0 | 24.406 | 9 | 6 | 21 | 6 | 830 | 89.8 | 4.44 | 554 | 6 | 11933630.91 | 4054048.602 | 9118803.99 | 6435943.542 | 8527504.958 | 3953878.634 | 14085159 | 10769605.2 | 43.1225 | 45.277 | Mapk8ip2 | 60597 | 0.91392 | 1.086081859 | 1.18838 | 0.6045 | 0.905186 |
| 148681139 | dual specificity phosphatase 10, partial [Mus musculus] | 0 | 61.214 | 21 | 8 | 31 | 8 | 493 | 53.6 | 7.74 | 799 | 8 | 15228756.38 | 14054121.78 | 16324931.14 | 13905235.06 | 19947270.76 | 19611510.98 | 17396653.81 | 17281167.67 | 7.60586 | 7.63208 | Dusp10 | 63953 | 0.88992 | 1.110082973 | 1.2474 | 0.00665 | 0.396316 |
| 1039729723 | PREDICTED: integral membrane protein 2C isoform X1 [Mus musculus] | 0 | 28.85 | 13 | 5 | 14 | 5 | 564 | 60.9 | 7.52 | 179 | 5 | 13154222.43 | 33091677.42 | 19136827.1 | 8937269.494 | 13314920.95 | 38535154.84 | 10952672.75 | 17501066.4 | 56.7315 | 62.7642 | Itm2c | 64294 | 0.9613 | 1.038699205 | 1.08051 | 0.86148 | 0.976238 |
| 14916467 | 72 kDa inositol polyphosphate 5-phosphatase isoform a [Mus musculus] | 0 | 27.909 | 11 | 5 | 22 | 5 | 647 | 71.9 | 9.19 | 230 | 5 | 14720350.9 | 5450864.035 | 9238046.929 | 4923846.822 | 9975817.542 | 4940862.026 | 9102196.25 | 5087078.972 | 52.6625 | 36.246 | Inpp5e | 64436 | 1.0824 | 0.917603546 | 0.84775 | 0.63527 | 0.912586 |
| 568968173 | PREDICTED: disco-interacting protein 2 homolog A isoform X1 [Mus musculus] | 0 | 68.322 | 13 | 16 | 52 | 11 | 1572 | 170.5 | 7.83 | 643 | 16 | 7002566.962 | 13089048.09 | 9136835.841 | 20910749.85 | 8647598.803 | 14946233.81 | 9226268.625 | 16638736.95 | 48.8798 | 32.5511 | Dip2a | 64451 | 1.00683 | 0.993168916 | 0.98643 | 0.96449 | 0.996581 |
| 114158672 | nischarin [Mus musculus] | 0.001 | 6.676 | 2 | 2 | 5 | 2 | 1593 | 174.9 | 5.16 | 26 | 2 | 11415425.91 | 2894804.737 | 2949620.686 | 3258085.956 | 8463522.906 | 2053852.575 | 3793952.938 | 3342452.298 | 81.7563 | 63.4177 | Nisch | 64652 | 1.07503 | 0.924966527 | 0.86041 | 0.78593 | 0.952792 |
| 568914069 | PREDICTED: hamartin isoform X1 [Mus musculus] | 0 | 12.595 | 3 | 2 | 2 | 2 | 1174 | 130.2 | 6.44 | 75 | 2 | 1900446.805 | 1416771.587 | 2829133.223 | 1762646.254 | 3117492.999 | 1718060.547 | 2599075.875 | 1870239.261 | 30.5104 | 28.0616 | Tsc1 | 64930 | 0.91891 | 1.081089906 | 1.17649 | 0.4622 | 0.863085 |
| 14994229 | adaptor-related protein complex AP-3 mu2 subunit [Mus musculus] | 0 | 49 | 32 | 10 | 50 | 7 | 418 | 46.9 | 7.56 | 695 | 10 | 28473196.74 | 35601946.05 | 33877522.59 | 27019287.77 | 33993815.6 | 36450634.85 | 29935624 | 42855739.25 | 13.2553 | 15.1127 | Ap3m2 | 64933 | 0.9319 | 1.068095942 | 1.14614 | 0.22874 | 0.723653 |
| 148682691 | claudin 12, isoform CRA_a, partial [Mus musculus] | 0 | 19.749 | 27 | 4 | 19 | 4 | 252 | 27.8 | 8.47 | 652 | 4 | 34219035.65 | 41442645.42 | 29483263.79 | 37129988.54 | 43642489.94 | 49327493.55 | 30912687 | 38121257.31 | 14.1316 | 19.4088 | Cldn12 | 64945 | 0.93516 | 1.064838531 | 1.13867 | 0.3311 | 0.776577 |
| 12849613 | unnamed protein product [Mus musculus] | 0 | 58.168 | 46 | 6 | 98 | 6 | 140 | 14.8 | 10.51 | 1382 | 6 | 225331363.7 | 286302412.2 | 210900296 | 226516264.1 | 206085693.1 | 221958237.2 | 220064968.3 | 219817624.2 | 14.1004 | 3.37661 | Rpl23 | 65019 | 1.04465 | 0.955352314 | 0.91452 | 0.28105 | 0.748062 |
| 13928670 | vacuolar protein sorting-associated protein 35 [Mus musculus] | 0 | 35.796 | 13 | 9 | 34 | 9 | 796 | 91.7 | 5.44 | 126 | 9 | 13849606.85 | 16587785.12 | 18238287.36 | 12468540.91 | 20073147.75 | 21406278.91 | 19654566.25 | 15145964.34 | 17.0642 | 14.2662 | Vps35 | 65114 | 0.88986 | 1.110138822 | 1.24756 | 0.09142 | 0.662454 |
| 148705344 | dihydropyrimidinase-like 5, isoform CRA_c, partial [Mus musculus] | 0 | 109.183 | 48 | 16 | 77 | 16 | 583 | 63.5 | 6.92 | 915 | 16 | 100376407.2 | 52596643.07 | 90822195.01 | 97645443.86 | 122179447.2 | 74165092.84 | 100098371.4 | 55960704.2 | 26.0178 | 32.9786 | Dpysl5 | 65254 | 0.9842 | 1.015800269 | 1.03211 | 0.88576 | 0.9839 |
| 13905234 | Rogdi protein [Mus musculus] | 0 | 17.614 | 18 | 4 | 11 | 4 | 309 | 34.4 | 8.54 | 95 | 4 | 6828943.127 | 5782508.789 | 11221731.96 | 2084058.355 | 7799841.012 | 13679472.96 | 9057070.625 | 399533.0181 | 58.0335 | 71.1727 | Rogdi | 66049 | 0.91173 | 1.088274344 | 1.19364 | 0.71955 | 0.939018 |
| 21313623 | GTP-binding protein 8 isoform 1 [Mus musculus] | 0 | 15.179 | 15 | 4 | 8 | 4 | 285 | 31.9 | 9.28 | 126 | 4 | 1562104.866 | 1623126.793 | 2620313.365 | 2033148.519 | 3170665.077 | 4196210.003 | 3777923.75 | 4639350.109 | 24.8801 | 15.8445 | Gtpbp8 | 66067 | 0.66365 | 1.336346289 | 2.01362 | 0.00243 | 0.38143 |
| 568942582 | PREDICTED: MICOS complex subunit Mic19 isoform X1 [Mus musculus] | 0 | 60.314 | 35 | 9 | 84 | 9 | 232 | 26.9 | 7.81 | 951 | 9 | 80592755.5 | 127128589.1 | 134885894.6 | 86126200.67 | 145960187.7 | 178217496.4 | 133271272.2 | 202758097.5 | 25.9211 | 19.062 | Chchd3 | 66075 | 0.78743 | 1.212567735 | 1.5399 | 0.03297 | 0.520743 |
| 225637531 | eukaryotic translation initiation factor 3 subunit F [Mus musculus] | 0 | 27.636 | 22 | 5 | 10 | 5 | 361 | 38 | 5.58 | 159 | 5 | 3207135.075 | 3984792.528 | 3337406.226 | 2237186.208 | 3127999.058 | 3158110.607 | 3070675.688 | 1998153.903 | 22.6051 | 19.782 | Eif3f | 66085 | 1.05852 | 0.941480291 | 0.88943 | 0.46943 | 0.863085 |
| 148667067 | mCG132973, isoform CRA_a, partial [Mus musculus] | 0 | 9.267 | 7 | 2 | 4 | 2 | 311 | 35.3 | 9.57 | 64 | 2 | 797310.9416 | 1626732.842 | 1190673.292 | 1591448.989 | 1138088.139 | 1810215.85 | 1399301 | 2276241.767 | 29.9656 | 30.0448 | Emc3 | 66087 | 0.88016 | 1.119837629 | 1.27231 | 0.30503 | 0.761857 |
| 15030153 | Coiled-coil-helix-coiled-coil-helix domain containing 6 [Mus musculus] | 0 | 70.966 | 33 | 10 | 71 | 10 | 273 | 29.8 | 8.16 | 1475 | 10 | 66668387.12 | 38163803.23 | 25585369.14 | 33904349 | 60854113 | 52037217.92 | 61308248.63 | 41937708.31 | 43.428 | 16.8854 | Chchd6 | 66098 | 0.86381 | 1.136191686 | 1.31533 | 0.24359 | 0.733806 |
| 12833615 | unnamed protein product [Mus musculus] | 0 | 34.312 | 25 | 7 | 21 | 7 | 377 | 42.5 | 9.74 | 375 | 7 | 14385237.05 | 11162959.83 | 16982943.61 | 18261487.09 | 21137927.43 | 19801641.67 | 17756170.56 | 19526208.7 | 20.6374 | 7.11215 | Ndufa9 | 66108 | 0.87462 | 1.12537765 | 1.2867 | 0.04408 | 0.588297 |
| 16758754 | V-type proton ATPase subunit F [Rattus norvegicus] | 0 | 13.713 | 34 | 4 | 15 | 4 | 119 | 13.4 | 5.82 | 197 | 4 | 5143738.18 | 7617130.53 | 5548553.501 | 8018982.082 | 9676981.277 | 9352447.16 | 6624261.25 | 14219810.86 | 21.9691 | 31.5758 | Atp6v1f | 66144 | 0.7954 | 1.204602818 | 1.51447 | 0.09835 | 0.670764 |
| 12963559 | V-type proton ATPase subunit G 2 [Mus musculus] | 0 | 33.372 | 41 | 4 | 21 | 4 | 118 | 13.6 | 10.26 | 590 | 4 | 18998281.82 | 16506452.9 | 19923140.2 | 14772298.41 | 22199297.21 | 16085344.9 | 17403911 | 25925249.76 | 13.3767 | 22.1653 | Atp6v1g2 | 66237 | 0.92482 | 1.075181678 | 1.16259 | 0.30555 | 0.761857 |
| 755551991 | PREDICTED: ADP-ribosylation factor GTPase-activating protein 3 isoform X1 [Mus musculus] | 0 | 78.742 | 33 | 15 | 64 | 14 | 525 | 57.6 | 8.47 | 1124 | 15 | 49545746.93 | 33950073.07 | 52610170.62 | 36058395.39 | 44969330.33 | 25103905.4 | 57749854.13 | 38328439.31 | 21.8479 | 32.7441 | Arfgap3 | 66251 | 1.01777 | 0.982227094 | 0.96507 | 0.86171 | 0.976238 |
| 13384852 | nicolin-1 [Mus musculus] | 0.001 | 5.769 | 9 | 2 | 5 | 2 | 213 | 24.3 | 7.08 | 78 | 2 | 906872.2296 | 1879565.528 | 1876404.924 | 2673002.697 | 3004588.186 | 2637673.709 | 2503183.25 | 4028478.815 | 39.4125 | 22.6717 | Nicn1 | 66257 | 0.75202 | 1.247982357 | 1.65951 | 0.05181 | 0.595816 |
| 13384870 | transmembrane protein 126A [Mus musculus] | 0 | 10.496 | 24 | 4 | 7 | 4 | 196 | 21.5 | 9.41 | 37 | 4 | 16446812.4 | 37059897.14 | 30260718.19 | 27573702.86 | 29507102.09 | 66407107.81 | 32340787.47 | 72262187.65 | 30.8167 | 44.5545 | Tmem126a | 66271 | 0.71405 | 1.285950542 | 1.80093 | 0.11166 | 0.678142 |
| 568920383 | PREDICTED: tumor protein D54 isoform X1 [Mus musculus] | 0 | 29.206 | 30 | 5 | 26 | 5 | 230 | 25.1 | 6.55 | 182 | 5 | 41943000.32 | 42650819.43 | 32328041.31 | 20442679.03 | 44878309.5 | 43832263.89 | 36467077.25 | 44645352.03 | 30.2647 | 9.46291 | Tpd52l2 | 66314 | 0.89434 | 1.105663343 | 1.23629 | 0.19551 | 0.715071 |
| 61743961 | neuroblast differentiation-associated protein AHNAK isoform 1 [Mus musculus] | 0 | 103.892 | 22 | 25 | 86 | 25 | 5656 | 603.9 | 6.3 | 680 | 25 | 41699559.53 | 28725896.61 | 37761132.21 | 32854285.81 | 36432088.91 | 77485801.31 | 49959792.13 | 69665318.58 | 16.0603 | 31.9662 | Ahnak | 66395 | 0.75305 | 1.246946366 | 1.65585 | 0.05542 | 0.599194 |
| 12833936 | unnamed protein product [Mus musculus] | 0 | 25.001 | 42 | 3 | 16 | 3 | 146 | 16.3 | 4.97 | 448 | 3 | 22912870.37 | 13586158.61 | 21811982.09 | 14093614.66 | 29382849.3 | 23161184.93 | 25193538 | 11069537.97 | 27.3204 | 35.4051 | Cyb5b | 66427 | 0.89826 | 1.101744978 | 1.22654 | 0.41116 | 0.835711 |
| 12833077 | unnamed protein product [Mus musculus] | 0 | 19.798 | 11 | 2 | 10 | 2 | 325 | 35.4 | 9.28 | 203 | 2 | 9714765.195 | 7674507.197 | 10003560.61 | 4964027.027 | 14896323.49 | 10765585.97 | 13103051.5 | 6873735.795 | 28.7677 | 30.3669 | Cyc1 | 66445 | 0.82971 | 1.170289659 | 1.41048 | 0.16269 | 0.705822 |
| 13385010 | microsomal glutathione S-transferase 3 [Mus musculus] | 0 | 97.721 | 75 | 8 | 36 | 8 | 153 | 16.9 | 9.5 | 786 | 8 | 77580511.95 | 31118185.31 | 13994177.58 | 33287892.83 | 57959503.55 | 33592210.9 | 57749252.56 | 1434652.903 | 69.5788 | 70.9485 | Mgst3 | 66447 | 1.0171 | 0.98289903 | 0.96637 | 0.94735 | 0.994906 |
| 12849578 | unnamed protein product [Mus musculus] | 0 | 11.76 | 14 | 3 | 9 | 3 | 279 | 30.9 | 10.73 | 119 | 3 | 11854391.65 | 13088782.53 | 11455470.23 | 10289052.83 | 13648365.51 | 10686718.95 | 14128270.56 | 10114992.96 | 9.89293 | 16.7681 | Ptpmt1 | 66461 | 0.98015 | 1.019846008 | 1.0405 | 0.70035 | 0.931521 |
| 27754130 | prostamide/prostaglandin F synthase [Mus musculus] | 0 | 9.382 | 13 | 3 | 8 | 3 | 201 | 21.7 | 6.74 | 56 | 3 | 5816062.668 | 4938584.968 | 6287167.848 | 3778809.764 | 3073137.869 | 1741145.218 | 3215181.75 | 2415717.517 | 21.1895 | 25.9108 | Prxl2b | 66469 | 1.33185 | 0.66815369 | 0.50167 | 0.00704 | 0.396316 |
| 12849405 | unnamed protein product [Mus musculus] | 0 | 23.155 | 34 | 4 | 31 | 4 | 145 | 16 | 10.43 | 729 | 4 | 50679008.38 | 30284013.81 | 33752231.89 | 34195193.55 | 32709279.26 | 25549767.43 | 41108471.69 | 27106896.49 | 24.5423 | 22.2465 | Rps23 | 66475 | 1.08147 | 0.918528446 | 0.84933 | 0.36818 | 0.80688 |
| 77404294 | up-regulated during skeletal muscle growth protein 5 [Mus musculus] | 0 | 29.109 | 45 | 4 | 26 | 4 | 58 | 6.4 | 9.83 | 938 | 4 | 84360488.94 | 99606046.84 | 71838780.31 | 92745258.94 | 57945895.2 | 142246984.4 | 70856514 | 100470612 | 13.7182 | 40.2855 | Atp5md | 66477 | 0.9681 | 1.03189886 | 1.0659 | 0.77984 | 0.952792 |
| 12846287 | unnamed protein product [Mus musculus] | 0 | 31.917 | 31 | 7 | 88 | 7 | 204 | 24 | 11.58 | 1030 | 7 | 190859850.5 | 172634263.4 | 173032824.7 | 230483943.1 | 207327251.3 | 235169893.9 | 224833627.3 | 157401654.8 | 14.1764 | 16.7286 | Rpl15 | 66480 | 0.96374 | 1.036263099 | 1.07526 | 0.53545 | 0.886555 |
| 21536222 | 40S ribosomal protein S21 [Mus musculus] | 0 | 43.964 | 58 | 4 | 45 | 4 | 83 | 9.1 | 8.51 | 982 | 4 | 161657850.5 | 275752904.3 | 163180668.1 | 260648569.6 | 87717051.19 | 50617111.68 | 54955045.88 | 174420042.6 | 28.5108 | 62.4785 | Rps21 | 66481 | 1.40159 | 0.598412429 | 0.42695 | 0.0261 | 0.515682 |
| 13385044 | 60S ribosomal protein L35 [Mus musculus] | 0 | 25.182 | 27 | 5 | 106 | 5 | 123 | 14.5 | 11.05 | 522 | 5 | 58034325.03 | 93756187.27 | 55992337.75 | 77786381.91 | 103981487.9 | 69933658.02 | 87519853.75 | 133245635.1 | 25.0126 | 27.2811 | Rpl35 | 66489 | 0.8396 | 1.160399007 | 1.38208 | 0.1422 | 0.693437 |
| 22094989 | mitochondrial import inner membrane translocase subunit TIM50 precursor [Mus musculus] | 0 | 15.808 | 10 | 3 | 10 | 3 | 353 | 39.8 | 8.13 | 127 | 3 | 3448766.606 | 2180241.133 | 5068144.047 | 2856019.52 | 5446801.448 | 3761351.085 | 5590470 | 3461289.894 | 36.4195 | 24.3053 | Timm50 | 66525 | 0.85205 | 1.147949855 | 1.34728 | 0.20593 | 0.715071 |
| 12847642 | unnamed protein product [Mus musculus] | 0 | 12.693 | 13 | 4 | 11 | 4 | 341 | 38.7 | 7.88 | 71 | 4 | 3082360.735 | 5317018.754 | 4472228.937 | 3041341.33 | 4882119.863 | 4425832.128 | 4130692.125 | 9460413.781 | 27.9788 | 43.8366 | Gdpd1 | 66569 | 0.82 | 1.179998628 | 1.43902 | 0.25032 | 0.733806 |
| 13905140 | Farsa protein [Mus musculus] | 0 | 76.868 | 34 | 13 | 57 | 13 | 508 | 57.6 | 8.02 | 972 | 13 | 39098888.68 | 37208071.23 | 40601554.13 | 30449143.5 | 45479855.11 | 116315378.3 | 51655365.63 | 30722801.87 | 12.1628 | 62.0543 | Farsa | 66590 | 0.75273 | 1.247274747 | 1.65701 | 0.25142 | 0.733806 |
| 13385134 | signal peptidase complex subunit 2 [Mus musculus] | 0 | 12.886 | 20 | 3 | 8 | 3 | 226 | 25 | 8.57 | 41 | 3 | 3555936.841 | 6329001.466 | 6040752.877 | 7392556.342 | 6607794.16 | 5421242.362 | 5415924.625 | 5921716.415 | 27.8479 | 9.64001 | Spcs2 | 66624 | 0.99896 | 1.001037381 | 1.00208 | 0.98921 | 0.998251 |
| 225543409 | paraspeckle component 1 [Mus musculus] | 0 | 52.522 | 27 | 9 | 42 | 9 | 523 | 58.7 | 6.67 | 922 | 9 | 32665144.89 | 31778270.53 | 34401887.98 | 37329666.52 | 33450734.27 | 31604185.5 | 35367354.25 | 35040764.56 | 7.18682 | 5.09236 | Pspc1 | 66645 | 1.00262 | 0.997379117 | 0.99477 | 0.90922 | 0.98561 |
| 114152892 | RecName: Full=Tubulin polyglutamylase complex subunit 2; Short=PGs2 | 0 | 16.446 | 11 | 2 | 14 | 2 | 296 | 33.2 | 7.24 | 256 | 2 | 8902423.585 | 8159083.002 | 9236984.307 | 5796142.916 | 9512472.841 | 6896298.012 | 8299709.75 | 6590555.896 | 19.3407 | 17.2399 | Tpgs2 | 66648 | 1.01255 | 0.987449894 | 0.97521 | 0.85298 | 0.975398 |
| 56699438 | elongation factor 1-delta isoform a [Mus musculus] | 0 | 21.305 | 10 | 5 | 11 | 5 | 660 | 72.9 | 6.51 | 78 | 5 | 7956516.465 | 9321064.488 | 10449289.44 | 11171300.28 | 9699033.508 | 11954172.96 | 12535841.5 | 13193437.75 | 14.4297 | 12.8145 | Eef1d | 66656 | 0.90167 | 1.098333919 | 1.21812 | 0.08599 | 0.655052 |
| 148705973 | mCG15863, isoform CRA_a [Mus musculus] | 0.001 | 6.192 | 3 | 2 | 7 | 2 | 681 | 76.4 | 8.97 | 45 | 2 | 1497900.451 | 1255516.196 | 2581891.587 | 4909141.253 | 2247548.485 | 796535.8661 | 2716024.813 | 1019683.088 | 65.1358 | 55.042 | Srp72 | 66661 | 1.20351 | 0.79648684 | 0.6618 | 0.39969 | 0.824613 |
| 28201956 | mRNA export factor [Mus musculus] | 0 | 53.543 | 38 | 10 | 43 | 10 | 368 | 40.9 | 7.83 | 714 | 10 | 40145072.19 | 40891632.71 | 44262693.27 | 48783476 | 44611988.41 | 35439199.98 | 49818867.31 | 32746055.58 | 9.05155 | 19.544 | Rae1 | 66679 | 1.03406 | 0.965943578 | 0.93413 | 0.54188 | 0.886555 |
| 148690000 | trafficking protein particle complex 5, partial [Mus musculus] | 0 | 13.826 | 20 | 3 | 10 | 3 | 223 | 24.8 | 10.04 | 84 | 3 | 1858786.182 | 1374530.32 | 1909916.797 | 1499448.323 | 2762730.165 | 2175374.562 | 2409533.75 | 2351634.237 | 15.9031 | 10.1576 | Trappc5 | 66682 | 0.81296 | 1.187039515 | 1.46014 | 0.00549 | 0.38143 |
| 5031571 | actin-related protein 2 isoform b [Homo sapiens] | 0 | 156.825 | 55 | 16 | 127 | 16 | 394 | 44.7 | 6.74 | 2922 | 16 | 333083646.9 | 245774739.8 | 307173690.7 | 300843736.5 | 305652724 | 263557758.3 | 328250726.4 | 285164471.9 | 12.3737 | 9.37071 | Actr2 | 66713 | 1.00179 | 0.998206317 | 0.99642 | 0.96465 | 0.996581 |
| 146231954 | leucine-rich repeat serine/threonine-protein kinase 2 [Mus musculus] | 0.001 | 6.22 | 1 | 2 | 3 | 2 | 2527 | 284.5 | 6.83 | 80 | 2 | 592279.3666 | 486771.0129 | 926960.9467 | 270651.51 | 837400.5289 | 467433.5471 | 707951.5 | 284862.2068 | 48.0576 | 42.9126 | Lrrk2 | 66725 | 0.99541 | 1.004587565 | 1.00922 | 0.97819 | 0.997464 |
| 12833586 | unnamed protein product [Mus musculus] | 0 | 11.934 | 22 | 3 | 7 | 2 | 121 | 14.3 | 8.68 | 89 | 3 | 100538661 | 68328417.07 | 106130061.2 | 59528232.28 | 106909515.1 | 51127277.28 | 111472280.2 | 60483359.25 | 27.676 | 37.7145 | Map1lc3a | 66734 | 1.00682 | 0.993178603 | 0.98645 | 0.95529 | 0.994906 |
| 13385196 | ER membrane protein complex subunit 2 [Mus musculus] | 0 | 10.492 | 11 | 3 | 21 | 3 | 297 | 34.9 | 6.81 | 126 | 3 | 3608164.949 | 3625798.729 | 3932728.674 | 4646464.118 | 4381298.771 | 3932439.742 | 5205562.75 | 4587033.919 | 12.2822 | 11.6814 | Emc2 | 66736 | 0.93239 | 1.067606518 | 1.14502 | 0.16134 | 0.702694 |
| 37360128 | mKIAA0868 protein [Mus musculus] | 0 | 10.317 | 2 | 2 | 7 | 2 | 1399 | 155.2 | 6.9 | 180 | 2 | 4359604.273 | 3950692.789 | 4364654.783 | 2867302.63 | 3964417.899 | 3439882.133 | 3963786.125 | 3736728.75 | 18.17 | 6.58114 | Cntnap2 | 66797 | 1.01427 | 0.985726545 | 0.97185 | 0.77996 | 0.952792 |
| 268837281 | nucleus accumbens-associated protein 1 [Mus musculus] | 0.001 | 5.401 | 4 | 2 | 2 | 2 | 514 | 56.5 | 5.73 | 27 | 2 | 1469685.541 | 1462353.226 | 1629280.453 | 985634.1858 | 2053274.685 | 1687793.733 | 1830718.625 | 1554758.215 | 20.0668 | 11.9715 | Nacc1 | 66830 | 0.87536 | 1.124637394 | 1.28477 | 0.06521 | 0.601634 |
| 27229002 | WD repeat domain phosphoinositide-interacting protein 3 [Mus musculus] | 0 | 8.744 | 7 | 2 | 4 | 2 | 344 | 38 | 7.56 | 25 | 2 | 3651032.281 | 3588172.908 | 5865008.081 | 2588719.058 | 9264176.893 | 4209415.613 | 9907191.75 | 6735797.129 | 35.2506 | 34.5626 | Wdr45b | 66840 | 0.68514 | 1.314861431 | 1.91912 | 0.04998 | 0.592616 |
| 148679790 | transcription factor 25 (basic helix-loop-helix), isoform CRA_c, partial [Mus musculus] | 0 | 14.516 | 9 | 5 | 14 | 5 | 692 | 78.3 | 6.7 | 79 | 5 | 1204349.273 | 7389299.341 | 6991021.11 | 5814729.824 | 8300832.66 | 6395319.049 | 4076991.75 | 9507680.302 | 53.1484 | 33.5407 | Tcf25 | 66855 | 0.86149 | 1.138514358 | 1.32157 | 0.38859 | 0.81965 |
| 568918098 | PREDICTED: protein TANC1 isoform X1 [Mus musculus] | 0 | 17.579 | 3 | 5 | 8 | 5 | 1874 | 202.6 | 8.24 | 45 | 5 | 4350694.158 | 2768972.241 | 3760879.291 | 4775544 | 3479398.91 | 1974417.306 | 4527588.625 | 4645066.997 | 22.2121 | 33.8524 | Tanc1 | 66860 | 1.034 | 0.965999645 | 0.93424 | 0.7452 | 0.947173 |
| 21630253 | copine-8 isoform 1 [Mus musculus] | 0 | 76.143 | 32 | 14 | 42 | 8 | 577 | 64.6 | 5.97 | 650 | 14 | 16733578.68 | 10119608.29 | 13888631.63 | 10073673.57 | 16567412.5 | 17121877.65 | 14273088.88 | 18797847.98 | 25.4009 | 11.2018 | Cpne8 | 66871 | 0.86439 | 1.13561248 | 1.31378 | 0.07639 | 0.627416 |
| 13385320 | vacuolar protein sorting-associated protein 28 homolog isoform 1 [Mus musculus] | 0 | 19.188 | 27 | 3 | 5 | 3 | 221 | 25.4 | 5.54 | 111 | 3 | 6153737.287 | 3890478.746 | 5722521.355 | 6291845.905 | 5987490.052 | 6323206.689 | 5516528 | 4465741.37 | 20.1209 | 14.5172 | Vps28 | 66914 | 0.99472 | 1.005284659 | 1.01063 | 0.93477 | 0.994906 |
| 54607098 | succinate dehydrogenase [ubiquinone] flavoprotein subunit, mitochondrial precursor [Mus musculus] | 0.003 | 3.835 | 3 | 2 | 3 | 2 | 664 | 72.5 | 7.37 | 14 | 2 | 1694870.536 | 1579122.076 | 1232261.836 | 2054645.529 | 1376207.173 | 1827323.265 | 1092158.375 | 1339763.046 | 20.6719 | 21.7329 | Sdha | 66945 | 1.07588 | 0.924120907 | 0.85894 | 0.35022 | 0.797109 |
| 148695348 | thioredoxin domain containing 14, isoform CRA_b [Mus musculus] | 0 | 19.056 | 17 | 5 | 18 | 5 | 317 | 36.5 | 8.12 | 161 | 5 | 12255845.9 | 10732007.44 | 6756522.865 | 9544560.409 | 11603183.91 | 17629260.68 | 8145541.25 | 16947087.89 | 23.6773 | 33.2496 | Tmx2 | 66958 | 0.83938 | 1.160618449 | 1.38271 | 0.18934 | 0.715071 |
| 148706281 | lute carrier family 25 (mitochondrial carrier; phosphate carrier), member 23, isoform CRA_b, partial [Mus muscul | 0 | 20.835 | 14 | 6 | 15 | 6 | 508 | 56.5 | 7.97 | 47 | 6 | 10493272.11 | 8430616.493 | 8995050.116 | 6136258.275 | 14536284.17 | 13071367.23 | 12509812.75 | 13580034.51 | 21.2393 | 6.41031 | Slc25a23 | 66972 | 0.77616 | 1.223837017 | 1.57678 | 0.0027 | 0.38143 |
| 13385408 | 60S ribosomal protein L11 [Mus musculus] | 0 | 26.503 | 36 | 6 | 78 | 6 | 178 | 20.2 | 9.6 | 1063 | 6 | 230448755.5 | 276201816.7 | 233422398.1 | 279301074.4 | 229677395 | 280631735.4 | 243809052.8 | 286763723.5 | 10.4024 | 10.6944 | Rpl11 | 67025 | 0.98956 | 1.010439413 | 1.0211 | 0.78903 | 0.953636 |
| 11037478 | Makorin RING zinc-finger protein 2 [Mus musculus] | 0.001 | 4.818 | 5 | 2 | 2 | 2 | 416 | 46.5 | 7.34 | 0 | 2 | 1633693.014 | 1968851.926 | 1925567.224 | 2023517.791 | 1487361.244 | 1734302.154 | 1266692.875 | 1467605.445 | 9.22454 | 12.8632 | Mkrn2 | 67027 | 1.11813 | 0.881868783 | 0.7887 | 0.02161 | 0.505785 |
| 21313156 | dnaJ homolog subfamily B member 4 [Mus musculus] | 0 | 71.995 | 45 | 13 | 44 | 12 | 337 | 37.8 | 8.59 | 520 | 13 | 49973858.08 | 50961498.45 | 59598421.46 | 59863825.7 | 49658836.53 | 45699674.75 | 59960070.75 | 55855307.76 | 9.73608 | 12.0238 | Dnajb4 | 67035 | 1.02137 | 0.978627611 | 0.95815 | 0.59902 | 0.905186 |
| 40068493 | probable ATP-dependent RNA helicase DDX17 isoform 1 [Mus musculus] | 0 | 103.603 | 37 | 22 | 166 | 16 | 652 | 72.5 | 8.68 | 2406 | 22 | 46031359.84 | 51346497.63 | 46032467.66 | 74992132.29 | 39905311.02 | 55278547.82 | 42831999.13 | 60885054.53 | 25.3171 | 20.088 | Ddx17 | 67040 | 1.04673 | 0.953267703 | 0.91071 | 0.58826 | 0.901256 |

| | | | | | | | | | | | | | | | | | | | | | | | | | | | | | |
|---|---|---|---|---|---|---|---|---|---|---|---|---|---|---|---|---|---|---|---|---|---|---|---|---|---|---|---|---|---|
| 18266680 | succinyl-CoA:3-ketoacid coenzyme A transferase 1, mitochondrial precursor [Mus musculus] | 0 | 25.854 | 18 | 6 | 13 | 6 | 520 | 56 | 8.53 | 73 | 6 | 10282586.1 | 14387102.88 | 11795688.25 | 7890137.241 | 20968726.2 | 10256168.77 | 11298785.38 | 12368628.52 | 24.5648 | 35.756 | Oxct1 | 67041 | 0.89383 | 1.106166504 | 1.23755 | 0.3841 | 0.816875 |
| 755558856 | PREDICTED: 40S ribosomal protein S10 isoform X1 [Mus musculus] | 0 | 41.958 | 28 | 6 | 88 | 6 | 222 | 24.9 | 10.81 | 1537 | 6 | 121758546.9 | 119952777.5 | 108140661.8 | 70585428.46 | 95270250.41 | 108913111.6 | 133587984.1 | 131840242 | 22.6384 | 15.8004 | Rps10 | 67097 | 0.94475 | 1.055248838 | 1.11696 | 0.44625 | 0.852048 |
| 12846159 | unnamed protein product [Mus musculus] | 0 | 27.698 | 25 | 5 | 66 | 5 | 227 | 26.3 | 11.9 | 627 | 5 | 246640333.6 | 228184546.4 | 220810303 | 245355447.5 | 234768907.9 | 210894733.5 | 250891988.6 | 215849013.1 | 5.43424 | 8.04424 | Rpl14 | 67115 | 1.01542 | 0.984576421 | 0.96962 | 0.54635 | 0.889301 |
| 148676911 | Metadherin, isoform CRA_b, partial [Mus musculus] | 0 | 74.419 | 27 | 13 | 72 | 13 | 609 | 66.6 | 9.39 | 1218 | 13 | 23295023.9 | 25384725.86 | 30142447.64 | 40588553.27 | 33361430.49 | 46693151.39 | 37129888.38 | 42376129.52 | 25.8251 | 14.6679 | Mtdh | 67154 | 0.85608 | 1.143921048 | 1.33623 | 0.0834 | 0.644421 |
| 110625979 | elongation factor 1-gamma [Mus musculus] | 0 | 71.005 | 30 | 13 | 68 | 13 | 437 | 50 | 6.74 | 420 | 13 | 62035646.32 | 58158624.77 | 65934166.56 | 63643716.87 | 55719297.14 | 70268870.1 | 74911636 | 69611275.98 | 5.24273 | 12.2465 | Eef1g | 67160 | 0.96014 | 1.039860836 | 1.08303 | 0.28847 | 0.752733 |
| 568955576 | PREDICTED: NADH dehydrogenase [ubiquinone] 1 alpha subcomplex subunit 13 isoform X1 [Mus musculus] | 0.001 | 6.293 | 6 | 2 | 4 | 2 | 311 | 34 | 8.6 | 0 | 2 | 11779162.9 | 4036310.751 | 2696365.231 | 1794085.257 | 7013787.353 | 4426184.011 | 6615284.75 | 3845108.658 | 89.8736 | 28.7334 | Ndufa13 | 67184 | 0.96222 | 1.037777324 | 1.07852 | 0.87422 | 0.979509 |
| 83745120 | 60S acidic ribosomal protein P2 [Mus musculus] | 0 | 88.661 | 70 | 7 | 59 | 7 | 115 | 11.6 | 4.54 | 2347 | 7 | 200305673.5 | 246982240.9 | 355575416.1 | 429594957.3 | 382775234.7 | 380751560 | 407915305.6 | 459155836 | 33.7131 | 8.95204 | Rplp2 | 67186 | 0.86094 | 1.139061065 | 1.32305 | 0.12059 | 0.681064 |
| 148667641 | RIKEN cDNA 2810022L02, isoform CRA_c, partial [Mus musculus] | 0 | 7.152 | 4 | 2 | 5 | 2 | 611 | 67.3 | 9.74 | 0 | 2 | 2270891.887 | 2327292.253 | 3095760.094 | 1882780.851 | 2486650.946 | 2296465.231 | 2707081.5 | 2391733.76 | 21.2075 | 7.11628 | Spats2l | 67198 | 0.98432 | 1.015684863 | 1.03187 | 0.78595 | 0.952792 |
| 166197652 | nuclear distribution protein nudE homolog 1 isoform a [Mus musculus] | 0 | 33.125 | 24 | 7 | 49 | 5 | 344 | 38.5 | 5.34 | 781 | 7 | 1291957.834 | 2161265.766 | 2814050.448 | 1054129.848 | 5217052.635 | 3189266.761 | 2811543 | 4297840.072 | 44.2679 | 28.1664 | Nde1 | 67203 | 0.64118 | 1.358815099 | 2.11923 | 0.02364 | 0.515682 |
| 19526848 | mitochondrial amidoxime reducing component 2 precursor [Mus musculus] | 0 | 32.518 | 24 | 8 | 31 | 8 | 338 | 38.2 | 8.68 | 352 | 8 | 79281645.04 | 75240666.4 | 61723835.06 | 72257701.88 | 78563128.5 | 98862747.92 | 64137068.78 | 83344816.71 | 10.4101 | 17.6222 | 43526 | 67247 | 0.94065 | 1.059346633 | 1.12618 | 0.30314 | 0.761857 |
| 148705735 | TBC1 domain family, member 19, isoform CRA_a [Mus musculus] | 0 | 16.077 | 6 | 3 | 11 | 3 | 592 | 67.8 | 6.95 | 516 | 3 | 6410442.212 | 5566319.498 | 6403045.448 | 6260647.275 | 9683624.888 | 5126826.884 | 6889043.75 | 5357583.831 | 6.52294 | 31.0081 | Tbc1d19 | 67249 | 0.95325 | 1.046745458 | 1.09808 | 0.59205 | 0.902243 |
| 21313024 | mitochondrial thiamine pyrophosphate carrier isoform 1 [Mus musculus] | 0 | 30.342 | 25 | 7 | 39 | 7 | 318 | 35.6 | 9.2 | 340 | 7 | 22394326.7 | 28364056.61 | 22533629.39 | 20303959.13 | 21938168.58 | 36230479.12 | 19577261.5 | 19184667.43 | 14.8023 | 33.3867 | Slc25a19 | 67283 | 0.9825 | 1.017502048 | 1.03563 | 0.85599 | 0.975398 |
| 26006165 | mKIAA0443 protein [Mus musculus] | 0 | 83.326 | 17 | 16 | 50 | 16 | 1350 | 152 | 5.08 | 478 | 16 | 64565035.79 | 19160752.79 | 37499147.5 | 34532955.82 | 54322951.58 | 45241680.71 | 45314198.66 | 32636072.64 | 48.4854 | 20.087 | Gprasp1 | 67298 | 0.93472 | 1.065282891 | 1.13968 | 0.62101 | 0.908209 |
| 568929555 | PREDICTED: dedicator of cytokinesis protein 7 isoform X1 [Mus musculus] | 0 | 80.101 | 10 | 17 | 51 | 17 | 2142 | 242.6 | 6.71 | 480 | 17 | 35893854.21 | 26271958.06 | 35064216.91 | 25443437.25 | 37601161.23 | 21529177.76 | 32892490.19 | 23979434.4 | 18.1799 | 25.9687 | Dock7 | 67299 | 1.02795 | 0.972049094 | 0.94562 | 0.73404 | 0.943908 |
| 568975333 | PREDICTED: clathrin heavy chain 1 isoform X1 [Mus musculus] | 0 | 563.649 | 57 | 71 | 414 | 71 | 1683 | 192.3 | 5.67 | 8694 | 71 | 725952061.9 | 543998171.9 | 650044730.7 | 593255441.3 | 500561648.3 | 1123973363 | 713968757.8 | 666665211 | 12.4453 | 35.2421 | Cltc | 67300 | 0.91086 | 1.089141209 | 1.19573 | 0.40728 | 0.832197 |
| 15079335 | Small nuclear ribonucleoprotein D3 [Mus musculus] | 0 | 7.622 | 15 | 2 | 12 | 2 | 126 | 14 | 10.46 | 98 | 2 | 8419501.934 | 2195178.662 | 3506311.059 | 2503860.841 | 7885569.041 | 2856425.442 | 10260709.5 | 2947045.824 | 69.6975 | 61.6763 | Snrpd3 | 67332 | 0.81947 | 1.180529122 | 1.4406 | 0.46486 | 0.863085 |
| 262050556 | FUN14 domain-containing protein 2 [Mus musculus] | 0 | 10.899 | 15 | 3 | 11 | 3 | 151 | 16.6 | 9.7 | 49 | 3 | 1120577.235 | 1974866.462 | 1662588.671 | 1942390.458 | 3068821.54 | 7808968.861 | 2404206.75 | 8413291.448 | 23.6028 | 57.6094 | Fundc2 | 67391 | 0.47193 | 1.528067973 | 3.2379 | 0.05473 | 0.599194 |
| 120953219 | protein SOGA3 precursor [Mus musculus] | 0 | 33.044 | 10 | 9 | 29 | 8 | 945 | 103.4 | 6.14 | 386 | 9 | 39623473.36 | 13777440.17 | 23336783.12 | 16261151.6 | 95232497.84 | 26437878.28 | 19910272.44 | 24785417.31 | 50.0776 | 86.2384 | Soga3 | 67412 | 0.71713 | 1.282872561 | 1.7889 | 0.36823 | 0.80688 |
| 12852375 | unnamed protein product [Mus musculus] | 0 | 15.063 | 7 | 3 | 15 | 3 | 520 | 60.1 | 9.35 | 220 | 3 | 6181790.646 | 6043918.13 | 6053985.637 | 6106453.586 | 8617892.129 | 8886463.85 | 7669802.75 | 5032891.085 | 1.03506 | 23.2856 | Far1 | 67420 | 0.89338 | 1.106623207 | 1.2387 | 0.1492 | 0.693437 |
| 4506697 | 40S ribosomal protein S20 isoform 2 [Homo sapiens] | 0 | 20.502 | 18 | 3 | 50 | 3 | 119 | 13.4 | 9.94 | 541 | 3 | 55268279.73 | 105678014.3 | 73839643.92 | 135032980 | 75323977.91 | 114142324.6 | 68745943.63 | 132478998.5 | 38.0727 | 31.3841 | Rps20 | 67427 | 0.97255 | 1.027445165 | 1.05644 | 0.8305 | 0.968871 |
| 13195630 | coiled-coil domain-containing protein 127 isoform 1 [Mus musculus] | 0 | 39.11 | 28 | 5 | 20 | 5 | 260 | 30.5 | 9.13 | 852 | 5 | 17825202.7 | 9223032.551 | 17136781.82 | 17634326.95 | 14704789.61 | 14476898.35 | 14561513.94 | 7604279.948 | 26.9473 | 27.1847 | Ccdc127 | 67433 | 1.09253 | 0.907465266 | 0.83061 | 0.37244 | 0.80688 |
| 21312960 | plakophilin-2 [Mus musculus] | 0 | 40.975 | 14 | 7 | 34 | 7 | 795 | 88 | 9.33 | 536 | 7 | 11224052.49 | 9324957.136 | 16846823.44 | 10142461.89 | 10066462.04 | 12647242.36 | 12161117.75 | 5057256.987 | 28.5948 | 34.754 | Pkp2 | 67451 | 1.08696 | 0.913042377 | 0.84 | 0.46336 | 0.863085 |
| 12835918 | unnamed protein product [Mus musculus] | 0 | 16.662 | 6 | 4 | 18 | 4 | 803 | 90.6 | 9.14 | 212 | 4 | 7276920.365 | 13287486.17 | 8466092.517 | 10068840.71 | 9580602.736 | 9733803.951 | 9608087.375 | 6198893.821 | 26.6628 | 19.6149 | Pnpla8 | 67452 | 1.0536 | 0.946403761 | 0.89826 | 0.54783 | 0.889301 |
| 13385672 | solute carrier family 25 member 46 [Mus musculus] | 0 | 9.976 | 11 | 3 | 7 | 3 | 418 | 46.2 | 7.64 | 45 | 3 | 1609964.437 | 1804874.228 | 1488209.081 | 1915544.323 | 2183715.199 | 2909433.944 | 1680386.75 | 2777577.565 | 11.2501 | 23.7662 | Slc25a46 | 67453 | 0.83307 | 1.166925507 | 1.40075 | 0.06273 | 0.601634 |
| 13385676 | FERM domain-containing protein 8 [Mus musculus] | 0 | 7.407 | 6 | 3 | 4 | 3 | 466 | 51.8 | 6.44 | 24 | 3 | 631289.167 | 1357671.548 | 934570.7833 | 1040086.091 | 730349.9516 | 650758.7647 | 702986 | 977204.1902 | 30.2436 | 18.9541 | Frmd8 | 67457 | 1.12845 | 0.871554533 | 0.77235 | 0.22419 | 0.718033 |
| 12832971 | unnamed protein product [Mus musculus] | 0 | 30.409 | 23 | 6 | 19 | 6 | 335 | 36.2 | 8.95 | 156 | 6 | 106254009.8 | 70786772.43 | 53697366.09 | 50582273.48 | 69452862.41 | 48417722.47 | 51990734.75 | 46902178.9 | 36.319 | 19.1838 | Decr1 | 67460 | 1.12961 | 0.870389465 | 0.77052 | 0.2862 | 0.752733 |
| 12849983 | unnamed protein product [Mus musculus] | 0 | 21.868 | 9 | 5 | 19 | 5 | 590 | 64.5 | 8.31 | 145 | 5 | 13562053.9 | 10443983.63 | 75957772.94 | 98514845.27 | 17849949.64 | 6231625.421 | 9155095.5 | 16035876.94 | 89.5203 | 44.8271 | Poc5 | 67463 | 1.60224 | 0.39775829 | 0.24825 | 0.14663 | 0.693437 |
| 12836691 | unnamed protein product [Mus musculus] | 0 | 21.775 | 23 | 4 | 14 | 4 | 260 | 29.7 | 5.38 | 141 | 4 | 12333869.85 | 5952986.374 | 6976529.856 | 6456196.028 | 12186772.16 | 2402798.029 | 11517409.06 | 6373692.395 | 37.3974 | 56.7992 | Snap29 | 67474 | 0.98815 | 1.01185493 | 1.02399 | 0.94693 | 0.994906 |
| 21312944 | calcium-binding and coiled-coil domain-containing protein 1 [Mus musculus] | 0 | 31.846 | 13 | 7 | 25 | 7 | 691 | 77.2 | 4.82 | 217 | 7 | 69430927.12 | 94448906.19 | 104512242.8 | 83361371.59 | 102921302.4 | 139502665.8 | 108340963.7 | 110434424.9 | 17.1274 | 14.261 | Calcoco1 | 67488 | 0.86537 | 1.13462763 | 1.31114 | 0.04951 | 0.592616 |
| 148677255 | WD repeat domain 48, isoform CRA_d [Mus musculus] | 0 | 82.349 | 23 | 12 | 34 | 12 | 755 | 84.4 | 7.39 | 1022 | 12 | 20209731.63 | 16880551.19 | 29964221.29 | 18719435.31 | 26266915.83 | 24786964.53 | 26947344.44 | 17643417.1 | 27.2409 | 17.8775 | Wdr48 | 67561 | 0.94559 | 1.054408443 | 1.11508 | 0.52079 | 0.880875 |
| 12837563 | unnamed protein product [Mus musculus] | 0 | 12.074 | 15 | 3 | 20 | 3 | 167 | 18.5 | 10.08 | 274 | 3 | 12097326.84 | 11250114.89 | 13873311.66 | 9589209.804 | 8421648 | 11214604.16 | 11393739 | 12042042.63 | 15.2374 | 14.897 | Tmem35a | 67564 | 1.04159 | 0.958412924 | 0.92015 | 0.46546 | 0.863085 |
| 148691828 | cytoplasmic polyadenylation element binding protein 4, isoform CRA_a [Mus musculus] | 0 | 83 | 25 | 12 | 40 | 6 | 729 | 80 | 7.18 | 838 | 12 | 24969559.13 | 19053634.17 | 27368392.33 | 15352412.17 | 25718169.54 | 15461706.49 | 28599270.5 | 18006556.14 | 25.2731 | 28.3412 | Cpeb4 | 67579 | 0.99403 | 1.00596864 | 1.01201 | 0.95194 | 0.994906 |
| 569004387 | PREDICTED: dynactin subunit 4 isoform X1 [Mus musculus] | 0 | 20.659 | 14 | 5 | 17 | 5 | 467 | 53 | 7.72 | 90 | 5 | 10085027.65 | 7508560.615 | 10437978.89 | 9267616.253 | 9847928.108 | 4941504.28 | 6219468.5 | 8826677.828 | 14.0087 | 30.4266 | Dctn4 | 67665 | 1.11117 | 0.888826527 | 0.7999 | 0.204 | 0.715071 |
| 12963655 | 60S ribosomal protein L38 [Mus musculus] | 0 | 29.135 | 50 | 4 | 43 | 4 | 70 | 8.2 | 10.1 | 649 | 4 | 181741431.4 | 116669898.3 | 82338971.06 | 164533324.6 | 102910352.9 | 72527134.93 | 111309544.3 | 50736627.87 | 33.2383 | 33.1087 | Rpl38 | 67671 | 1.2354 | 0.764603912 | 0.61891 | 0.09879 | 0.670764 |
| 13385800 | transcription elongation factor B polypeptide 2 [Mus musculus] | 0 | 22.489 | 53 | 5 | 20 | 5 | 118 | 13.2 | 5.01 | 389 | 5 | 9091717.361 | 12935066.8 | 17068118.03 | 19583479.93 | 9077730.264 | 10955619.86 | 9481638.125 | 11362782.63 | 31.4906 | 10.8627 | Elob | 67673 | 1.1788 | 0.82120029 | 0.69664 | 0.11017 | 0.678142 |
| 148681386 | succinate dehydrogenase complex, subunit B, iron sulfur (Ip), isoform CRA_c [Mus musculus] | 0 | 13.213 | 8 | 3 | 7 | 3 | 386 | 42.8 | 8.07 | 119 | 3 | 4613764.209 | 6017226.282 | 1784693.97 | 5561214.102 | 5168956.446 | 9110627.217 | 6072174.75 | 3548169.045 | 42.2449 | 39.1084 | Sdhb | 67680 | 0.85856 | 1.141439299 | 1.32948 | 0.36328 | 0.804292 |
| 568959967 | PREDICTED: kin of IRRE-like protein 3 isoform X1 [Mus musculus] | 0.001 | 4.441 | 3 | 2 | 12 | 2 | 803 | 88 | 7.81 | 14 | 2 | 3402234.706 | 5153869.595 | 4063584.193 | 6599656.787 | 5139836.807 | 5476516.218 | 3493461.25 | 8459746.914 | 29.0876 | 36.6591 | Kirrel3 | 67703 | 0.91983 | 1.080169983 | 1.17431 | 0.52718 | 0.886268 |
| 12842068 | unnamed protein product [Mus musculus] | 0 | 29.021 | 44 | 3 | 24 | 3 | 151 | 16.3 | 5.67 | 757 | 3 | 64998612.44 | 67375728.86 | 66707422.36 | 60246910.18 | 68920909.67 | 77239638.3 | 65352365.5 | 82215356.57 | 4.96135 | 10.4684 | Arpc5 | 67771 | 0.9378 | 1.062199013 | 1.13265 | 0.08458 | 0.648205 |
| 13385872 | interleukin enhancer-binding factor 2 [Mus musculus] | 0 | 37.333 | 26 | 7 | 28 | 7 | 390 | 43 | 5.26 | 345 | 7 | 35818367.4 | 34586507.14 | 39804628.03 | 37847792.95 | 27798050.22 | 51921614.12 | 42952295.25 | 46344284.75 | 6.20109 | 24.4293 | Ilf2 | 67781 | 0.9339 | 1.066101223 | 1.14156 | 0.35995 | 0.80334 |
| 40789289 | regulator of G-protein signaling 8 [Mus musculus] | 0 | 15.467 | 24 | 4 | 9 | 4 | 180 | 21 | 9.26 | 87 | 4 | 9315889.8 | 2092966.974 | 6248418.152 | 17168086.05 | 4581306.685 | 20110044.56 | 5422524.5 | 20018520.86 | 73.1711 | 69.4411 | Rgs8 | 67792 | 0.81983 | 1.180172312 | 1.43954 | 0.50458 | 0.88031 |
| 21362303 | synaptosomal-associated protein 47 [Mus musculus] | 0 | 176.878 | 72 | 24 | 152 | 24 | 413 | 46.5 | 5.76 | 3524 | 24 | 267168269.4 | 174594424.6 | 239583309.1 | 165218996.2 | 237161724.6 | 165903310.9 | 194948825.3 | 163148957.7 | 23.4534 | 18.0774 | Snap47 | 67826 | 1.05312 | 0.9468802 | 0.89912 | 0.50604 | 0.88031 |
| 148693873 | isocitrate dehydrogenase 3 (NAD+) alpha, isoform CRA_c, partial [Mus musculus] | 0 | 23.497 | 17 | 5 | 17 | 5 | 375 | 40.5 | 6.54 | 138 | 5 | 12294969.79 | 9120608.624 | 14766808.41 | 12419898.08 | 16322025.12 | 9515216.211 | 16111153.75 | 7878423.809 | 19.0769 | 35.2702 | Idh3a | 67834 | 0.98756 | 1.012440772 | 1.02519 | 0.90593 | 0.98561 |
| 312433993 | G-protein-signaling modulator 1 isoform 3 [Mus musculus] | 0 | 16.892 | 7 | 4 | 8 | 4 | 705 | 78.1 | 6.62 | 73 | 4 | 6550009.864 | 4455350.182 | 3463074.542 | 2766938.546 | 5207873.72 | 4365330.17 | 4198012.875 | 5437909.236 | 38.222 | 12.7494 | Gpsm1 | 67839 | 0.94584 | 1.054157772 | 1.11452 | 0.59469 | 0.902243 |
| 170172564 | E3 ubiquitin-protein ligase RNF115 [Mus musculus] | 0.001 | 5.921 | 5 | 2 | 6 | 2 | 305 | 33.8 | 5.83 | 0 | 2 | 2709188.034 | 4207330.53 | 3099138.785 | 3953505.827 | 2977133.059 | 3365279.22 | 2445143.125 | 3326530.262 | 20.1924 | 14.0761 | Rnf115 | 67845 | 1.07112 | 0.928878587 | 0.8672 | 0.30332 | 0.761857 |
| 21312758 | serine/threonine-protein phosphatase 6 catalytic subunit [Mus musculus] | 0 | 26.855 | 20 | 5 | 18 | 5 | 305 | 35.1 | 5.69 | 300 | 5 | 37706923.7 | 16662780.54 | 16544953.92 | 28812459.41 | 39869258.13 | 14065669.18 | 20252002.5 | 11974816.1 | 41.2289 | 59.0264 | Ppp6c | 67857 | 1.07298 | 0.927024291 | 0.86398 | 0.69268 | 0.930425 |
| 21312994 | mitochondrial 2-oxoglutarate/malate carrier protein [Mus musculus] | 0 | 71.983 | 44 | 11 | 56 | 11 | 314 | 34.1 | 9.94 | 1868 | 11 | 368007230.6 | 270328541.5 | 223877534.9 | 218332761.9 | 279426904.4 | 481960637 | 215925165.5 | 264956392.1 | 25.649 | 37.8176 | Slc25a11 | 67863 | 0.93038 | 1.069623719 | 1.14967 | 0.57485 | 0.895889 |
| 148705834 | transmembrane protein 33, isoform CRA_e, partial [Mus musculus] | 0 | 27.641 | 21 | 5 | 35 | 5 | 264 | 29.8 | 9.58 | 359 | 5 | 43163303.14 | 37863344.47 | 26641138.58 | 35746141.83 | 33149524.13 | 39634141.02 | 29542551.38 | 32455522.81 | 19.2128 | 12.6319 | Tmem33 | 67878 | 1.03103 | 0.968970801 | 0.93981 | 0.61317 | 0.905186 |
| 568909575 | PREDICTED: calmodulin-regulated spectrin-associated protein 2 isoform X1 [Mus musculus] | 0 | 35.78 | 7 | 8 | 18 | 8 | 1481 | 166.6 | 7.06 | 325 | 8 | 19886667.48 | 13089746.78 | 15645620.05 | 11203150.13 | 15615717.65 | 2841334.343 | 8694039.531 | 1482279.781 | 25.1224 | 90.0802 | Camsap2 | 67886 | 1.35261 | 0.647385006 | 0.47862 | 0.08163 | 0.642075 |
| 30794450 | 60S ribosomal protein L4 [Mus musculus] | 0 | 166.183 | 45 | 20 | 309 | 20 | 419 | 47.1 | 11 | 4929 | 20 | 626631034.6 | 598263970 | 483532179.7 | 638440144.8 | 492992044.7 | 460165625.6 | 532070832.7 | 474722416.8 | 12.0715 | 6.34794 | Rpl4 | 67891 | 1.08984 | 0.910161891 | 0.83513 | 0.04647 | 0.591577 |
| 18017590 | phospholipid phosphatase 3 [Mus musculus] | 0 | 42.993 | 31 | 7 | 29 | 7 | 312 | 35.2 | 9.07 | 257 | 7 | 61008003.46 | 58302953.46 | 70125033.77 | 59732750.94 | 77072440.5 | 51024654.25 | 60473980 | 54786932.67 | 8.56852 | 18.8988 | Plpp3 | 67916 | 1.0118 | 0.988202196 | 0.97668 | 0.82633 | 0.968473 |
| 882939105 | transcription elongation factor B polypeptide 1 isoform 1 [Mus musculus] | 0 | 16.218 | 25 | 3 | 11 | 3 | 134 | 14.9 | 4.73 | 38 | 3 | 18094733.1 | 23163135.9 | 15843022.51 | 21032232.01 | 16457614.99 | 20956274.02 | 15529390.88 | 17684039.59 | 16.486 | 13.4228 | Eloc | 67923 | 1.05046 | 0.949544355 | 0.90394 | 0.38416 | 0.816875 |
| 13324686 | mitochondrial import receptor subunit TOM20 homolog [Mus musculus] | 0 | 29.451 | 44 | 3 | 14 | 3 | 145 | 16.3 | 8.6 | 106 | 3 | 6275873.121 | 8831305.491 | 21213202.8 | 2460572.55 | 19599614.44 | 21903307.91 | 20003573.69 | 12504806.3 | 83.676 | 22.2825 | Tomm20 | 67952 | 0.68765 | 1.312347226 | 1.90844 | 0.10104 | 0.674486 |
| 568968333 | PREDICTED: plasma membrane calcium-transporting ATPase 1 isoform X1 [Mus musculus] | 0 | 145.656 | 23 | 22 | 107 | 10 | 1249 | 137.8 | 6 | 1986 | 22 | 28679511.29 | 29489628.05 | 33132772.76 | 38305595.71 | 34931057.59 | 44672221.76 | 33883915.13 | 40364230.7 | 13.5379 | 13.0525 | Atp2b1 | 67972 | 0.91447 | 1.085528853 | 1.18706 | 0.11891 | 0.678142 |
| 31377644 | ATPase family AAA domain-containing protein 1 isoform 1 [Homo sapiens] | 0 | 29.472 | 23 | 6 | 31 | 6 | 361 | 40.7 | 6.9 | 499 | 6 | 22262126.35 | 25562857.92 | 25431209.2 | 25834498.78 | 43117195.25 | 44707261.8 | 27580714.75 | 38997387.23 | 6.79014 | 20.0271 | Atad1 | 67979 | 0.7818 | 1.218198577 | 1.55819 | 0.01289 | 0.461382 |
| 13386010 | 60S ribosomal protein L22-like 1 [Mus musculus] | 0 | 42.232 | 39 | 3 | 27 | 3 | 122 | 14.5 | 9.45 | 336 | 3 | 16988907.76 | 19549887.76 | 23622971.95 | 27113119.97 | 12452520.85 | 22318859.84 | 18469042.75 | 7072593.587 | 20.4559 | 44.4742 | Rpl22l1 | 68028 | 1.18268 | 0.817316531 | 0.69107 | 0.14525 | 0.693437 |
| 12834476 | unnamed protein product [Mus musculus] | 0 | 12.883 | 21 | 3 | 15 | 3 | 183 | 20.2 | 8.92 | 93 | 3 | 5171784.195 | 5177221.324 | 4962309.321 | 6208208.197 | 5648846.136 | 5991567.819 | 3027445.75 | 9049714.701 | 10.4316 | 41.5858 | Emc4 | 68032 | 0.95141 | 1.048589576 | 1.10214 | 0.67905 | 0.925326 |
| 13386026 | UPF0568 protein C14orf166 homolog [Mus musculus] | 0 | 85.2 | 50 | 10 | 58 | 10 | 244 | 28.1 | 6.89 | 1619 | 10 | 90876077.64 | 94448024.96 | 82792645.63 | 69715612.44 | 68630201.78 | 58682277.39 | 61178789.06 | 75316965.72 | 12.9901 | 11.431 | Rtraf | 68045 | 1.12304 | 0.876962879 | 0.78089 | 0.03198 | 0.515682 |
| 4506685 | 40S ribosomal protein S13 [Homo sapiens] | 0 | 33.354 | 32 | 7 | 30 | 7 | 151 | 17.2 | 10.54 | 384 | 7 | 167312789.7 | 212548937 | 200111846.4 | 189674263.4 | 182584157.1 | 246823225.9 | 226710226 | 235611965.6 | 9.96202 | 12.6179 | Rps13 | 68052 | 0.92652 | 1.073482243 | 1.15862 | 0.1231 | 0.688884 |
| 27754054 | dephospho-CoA kinase domain-containing protein [Mus musculus] | 0 | 24.37 | 42 | 8 | 15 | 8 | 231 | 26.5 | 9.58 | 90 | 8 | 14411438.88 | 12270902.09 | 20101233.11 | 13774218.69 | 18579880.2 | 15546113.17 | 20152171.63 | 18355615.38 | 22.639 | 10.5528 | Dcakd | 68087 | 0.90933 | 1.090666304 | 1.19941 | 0.17505 | 0.713031 |
| 12857620 | unnamed protein product [Mus musculus] | 0 | 105.195 | 66 | 8 | 65 | 8 | 168 | 19.6 | 8.43 | 1260 | 8 | 418525480.1 | 310719684.3 | 321096256.5 | 343195853.2 | 349018297.6 | 351239845.2 | 357976219.8 | 346882257.1 | 13.9738 | 1.36818 | Arpc4 | 68089 | 0.99586 | 1.00413747 | 1.00831 | 0.90965 | 0.98561 |
| 56554097 | Chain A, Crystal Structure Of Mouse Phosphoglucose Isomerase | 0 | 32.334 | 16 | 8 | 47 | 8 | 564 | 63.5 | 7.97 | 410 | 8 | 11876576.37 | 18810376.34 | 10714222.8 | 10369850.22 | 28754632.36 | 24634653.36 | 16326107.94 | 26255792.37 | 30.6311 | 22.4432 | Dynll2 | 68097 | 0.70083 | 1.299170831 | 1.85376 | 0.0163 | 0.461382 |
| 569004429 | PREDICTED: protein spire homolog 1 isoform X6 [Mus musculus] | 0 | 24.73 | 9 | 7 | 33 | 7 | 802 | 90.6 | 8.81 | 440 | 7 | 26652629.02 | 21714038.67 | 30005520.08 | 18177636.39 | 25144242.16 | 22550487.97 | 23455211.13 | 23140057.51 | 21.68 | 4.72118 | Spire1 | 68166 | 1.01184 | 0.988158523 | 0.97659 | 0.83972 | 0.969881 |
| 74203647 | unnamed protein product [Mus musculus] | 0 | 32.492 | 31 | 6 | 52 | 6 | 160 | 18.2 | 11.05 | 647 | 6 | 171069987.1 | 216519074.9 | 281544674.7 | 304266024.1 | 264553375.4 | 278467187.1 | 280646860 | 334521053.5 | 25.0116 | 10.6437 | Rpl24 | 68193 | 0.91331 | 1.086690624 | 1.18984 | 0.22442 | 0.718033 |
| 148679646 | heat shock factor binding protein 1, isoform CRA_a [Mus musculus] | 0 | 190.792 | 64 | 6 | 74 | 6 | 88 | 10.1 | 4.41 | 2899 | 6 | 490405153.4 | 158220790.7 | 605840532.7 | 353523912.6 | 508908474.7 | 187454604.1 | 513643618 | 179990060.8 | 47.8807 | 54.4311 | Hsbp1 | 68196 | 1.07271 | 0.927286669 | 0.86443 | 0.70029 | 0.931521 |
| 71725385 | GTP-binding protein Di-Ras2 [Mus musculus] | 0 | 35.845 | 43 | 7 | 53 | 6 | 199 | 22.5 | 8.76 | 617 | 7 | 24870483.23 | 20324419.58 | 19284594.11 | 12612103.56 | 18632759.7 | 27006379.88 | 25987218 | 22382035.38 | 26.2535 | 16.1876 | Diras2 | 68203 | 0.90113 | 1.098870796 | 1.21944 | 0.22995 | 0.723653 |
| 61098124 | protein FAM98B [Mus musculus] | 0 | 36.459 | 25 | 7 | 15 | 2 | 429 | 45.3 | 8.5 | 157 | 7 | 1016965.633 | 1672531.131 | 1181305.466 | 3200103.873 | 1632432.433 | 1436020.597 | 1097700.125 | 4188727.403 | 56.2702 | 67.8558 | Fam98b | 68215 | 0.91676 | 1.08323559 | 1.18159 | 0.72355 | 0.939018 |
| 13386106 | cleavage and polyadenylation specificity factor subunit 5 [Mus musculus] | 0 | 12.966 | 15 | 3 | 8 | 3 | 227 | 26.2 | 8.82 | 59 | 3 | 4156201.206 | 6021247.168 | 8146276.83 | 7317605.669 | 4606621.763 | 11048083.52 | 7775031.25 | 11841280.49 | 27.1233 | 37.5771 | Nudt21 | 68219 | 0.84191 | 1.158090871 | 1.37555 | 0.2456 | 0.733806 |
| 148670128 | lycerol-3-phosphate O-acyltransferase 1 (lysophosphatidic acid acyltransferase, delta), isoform CRA_a [Mus mu | 0 | 17.487 | 15 | 5 | 15 | 5 | 378 | 43.8 | 8.46 | 55 | 5 | 10986298.25 | 7535205.453 | 10881018.64 | 7492052.075 | 11877785.72 | 12495807.11 | 12852429.5 | 9540267.291 | 21.4135 | 12.7412 | Agpat4 | 68262 | 0.882 | 1.117996811 | 1.26757 | 0.09304 | 0.662454 |
| 18152793 | pyruvate dehydrogenase E1 component subunit beta, mitochondrial precursor [Mus musculus] | 0 | 15.506 | 13 | 4 | 6 | 4 | 359 | 38.9 | 6.87 | 21 | 4 | 2646397.234 | 1802075.765 | 2794261.811 | 4463911.086 | 4946361.941 | 383432.7302 | 3618639.688 | 3362636.98 | 38.0685 | 62.5676 | Pdhb | 68263 | 0.97483 | 1.025165816 | 1.05163 | 0.89639 | 0.984401 |
| 21311845 | mitochondrial glutamate carrier 1 [Mus musculus] | 0 | 214.117 | 82 | 19 | 385 | 16 | 323 | 34.6 | 9.09 | 6354 | 19 | 1684656221 | 1542140632 | 1445184295 | 1428336929 | 1809166787 | 2241974148 | 1525060582 | 1755896126 | 7.7119 | 16.3239 | Slc25a22 | 68267 | 0.9083 | 1.091702013 | 1.20192 | 0.10389 | 0.674486 |
| 110287975 | rase subunit STT3B; Short=Oligosaccharyl transferase subunit STT3B; Short=STT3-B; AltName: Full=B6dom1 a | 0 | 21.531 | 7 | 5 | 18 | 5 | 823 | 93.2 | 8.95 | 313 | 5 | 12742610.8 | 13773801.22 | 16174972.26 | 49299743.33 | 5977977.045 | 16210590.38 | 17413933.25 | 14231839.47 | 76.5009 | 38.3156 | Stt3b | 68292 | 1.26166 | 0.738339345 | 0.58521 | 0.33816 | 0.782483 |
| 12859600 | unnamed protein product [Mus musculus] | 0 | 13.257 | 5 | 3 | 7 | 3 | 832 | 94.4 | 6.61 | 34 | 3 | 2681434.458 | 3026863.819 | 2414880.991 | 2771645.961 | 2176114.9 | 2007791.567 | 2098598 | 2972138.228 | 9.27273 | 19.2051 | Vps53 | 68299 | 1.0814 | 0.918599214 | 0.84945 | 0.15972 | 0.700858 |
| 755565726 | PREDICTED: MICOS complex subunit Mic26 isoform X1 [Mus musculus] | 0.001 | 6.486 | 11 | 2 | 5 | 2 | 230 | 26.4 | 8.78 | 13 | 2 | 8259042.624 | 6154291.196 | 4281021.021 | 4840902.613 | 8657890.423 | 12741639.24 | 5147266.313 | 7407796.04 | 30.0389 | 37.5312 | Apoo | 68316 | 0.81876 | 1.181237813 | 1.44271 | 0.20268 | 0.715071 |
| 148686612 | cysteine rich protein 2, isoform CRA_a, partial [Mus musculus] | 0 | 33.422 | 37 | 4 | 21 | 4 | 209 | 22.8 | 8.65 | 509 | 4 | 5500639.131 | 9503421.418 | 19465763.78 | 25302335.96 | 19463706.86 | 13004809.01 | 21209277.5 | 43600616.57 | 60.6632 | 54.8102 | Crip2 | 68337 | 0.76118 | 1.238816386 | 1.62749 | 0.28886 | 0.752733 |
| 148676697 | RAB14, member RAS oncogene family, isoform CRA_a, partial [Mus musculus] | 0 | 15.172 | 23 | 4 | 21 | 3 | 218 | 24.5 | 9.8 | 204 | 4 | 2619374.994 | 1612500.466 | 5528072.427 | 1423726.585 | 7267083.323 | 1592300.906 | 3335478.375 | 1614948.123 | 67.7967 | 77.3634 | Rab14 | 68365 | 0.89493 | 1.105072831 | 1.23482 | 0.70237 | 0.931521 |
| 568981958 | PREDICTED: neuritin isoform X1 [Mus musculus] | 0 | 29.045 | 34 | 4 | 16 | 4 | 151 | 16.3 | 5.6 | 545 | 4 | 19483101.91 | 38168474.83 | 23145197.6 | 40587078.85 | 16575789.01 | 40268210.77 | 24799437.5 | 27190557.52 | 34.8706 | 36.099 | Nrn1 | 68404 | 1.05451 | 0.945487031 | 0.89661 | 0.67903 | 0.925326 |
| 148680264 | mCG5258, isoform CRA_b, partial [Mus musculus] | 0 | 8.979 | 21 | 4 | 82 | 4 | 119 | 13.4 | 11.47 | 715 | 4 | 94395794.68 | 193837659.1 | 81839245.28 | 157805682.6 | 93316329.48 | 134496339 | 90099336.2 | 145713912.1 | 40.1441 | 24.4554 | Rpl34 | 68436 | 1.0648 | 0.935196988 | 0.87828 | 0.61209 | 0.905186 |
| 148685561 | TBC1 domain family, member 10b, partial [Mus musculus] | 0 | 37.338 | 13 | 6 | 17 | 6 | 817 | 88.7 | 8.78 | 132 | 6 | 9925083.862 | 23660199.34 | 55669097.49 | 10734154.79 | 46464840.14 | 55333674.33 | 38162676 | 52819682.15 | 85.5873 | 15.892 | Tbc1d10b | 68449 | 0.68305 | 1.316946834 | 1.92803 | 0.08724 | 0.655499 |
| 58037123 | complex I assembly factor TMEM126B, mitochondrial [Mus musculus] | 0 | 8.447 | 10 | 2 | 5 | 2 | 230 | 25.4 | 9.51 | 26 | 2 | 8414077.156 | 5474382.757 | 5269649.161 | 4795509.619 | 12633929.89 | 15782211.73 | 11167087.5 | 6941788.853 | 27.4183 | 31.5668 | Tmem126b | 68472 | 0.67974 | 1.320258739 | 1.9423 | 0.03092 | 0.515682 |
| 148666524 | required for meiotic nuclear division 5 homolog A (S. cerevisiae), isoform CRA_a, partial [Mus musculus] | 0 | 10.508 | 6 | 2 | 6 | 2 | 407 | 45.3 | 6.04 | 99 | 2 | 805038.313 | 731665.2418 | 1418266.316 | 1098072.241 | 1048520.856 | 1188365.566 | 1056181.5 | 757065.4696 | 30.8887 | 17.9755 | Rmnd5a | 68477 | 1.00036 | 0.999641039 | 0.99928 | 0.99693 | 0.998251 |
| 124486662 | vacuolar protein sorting-associated protein 51 homolog [Mus musculus] | 0 | 15.82 | 8 | 5 | 9 | 5 | 782 | 86.1 | 6.24 | 106 | 5 | 11986701.73 | 14329893.06 | 10076954.29 | 9962107.239 | 11762112.91 | 12217498.49 | 9282345 | 29338792.27 | 17.6867 | 58.8902 | Vps51 | 68505 | 0.8509 | 1.149097176 | 1.35044 | 0.42267 | 0.840085 |
| 221554491 | liprin-alpha-4 [Mus musculus] | 0 | 277.369 | 40 | 35 | 257 | 25 | 1187 | 133.6 | 6.27 | 4246 | 35 | 151500694 | 126372094.7 | 148996855.6 | 139917478.9 | 152349792.6 | 156995417.7 | 151717011 | 135382861.7 | 8.01992 | 6.33754 | Ppfia4 | 68507 | 0.9745 | 1.025496165 | 1.05233 | 0.35443 | 0.798662 |
| 148684225 | RIKEN cDNA 5730509C05, partial [Mus musculus] | 0 | 7.393 | 6 | 2 | 3 | 2 | 451 | 47.6 | 10.76 | 27 | 2 | 1468282.567 | 2393547.676 | 7564924.908 | 6480692.293 | 6050711.17 | 5502946.703 | 7515913 | 9068620.27 | 66.9407 | 22.7492 | Wipf2 | 68524 | 0.77781 | 1.22218703 | 1.57131 | 0.18286 | 0.715071 |
| 28972720 | mKIAA1321 protein [Mus musculus] | 0 | 9.152 | 3 | 2 | 4 | 2 | 716 | 78.2 | 8.7 | 47 | 2 | 847774.2826 | 2006120.57 | 2000212.984 | 2789774.986 | 1719353.916 | 2806727.029 | 1788628.5 | 3998985.362 | 41.8603 | 41.4813 | Nufip2 | 68564 | 0.85133 | 1.148673281 | 1.34927 | 0.35615 | 0.798662 |
| 34610235 | reticulon-4 isoform A [Mus musculus] | 0 | 249.714 | 42 | 27 | 196 | 25 | 1162 | 126.5 | 4.54 | 4042 | 27 | 234307883.2 | 218048054.5 | 251564774.5 | 249905097.2 | 202367692.8 | 238084889.7 | 227315760 | 248358511.2 | 6.57128 | 8.62036 | Rtn4 | 68585 | 1.02016 | 0.97983962 | 0.96048 | 0.48284 | 0.868779 |
| 19527322 | LETM1 domain-containing protein 1 [Mus musculus] | 0 | 17.287 | 13 | 4 | 13 | 4 | 360 | 41.7 | 10.54 | 74 | 4 | 16319346.69 | 19934083.01 | 17171497.32 | 14003529.61 | 17334067.02 | 24551895.47 | 16171300.69 | 21058241.9 | 14.5304 | 19.2324 | Letmd1 | 68614 | 0.92025 | 1.079751143 | 1.17333 | 0.24402 | 0.733806 |
| 30519943 | sorting and assembly machinery component 50 homolog [Mus musculus] | 0 | 94.516 | 37 | 13 | 92 | 13 | 469 | 51.8 | 6.8 | 1487 | 13 | 204335066.4 | 184371052.6 | 200903502.4 | 209187060.9 | 255538622.3 | 212726871.5 | 187384708.9 | 189966639.4 | 5.3926 | 14.9238 | Samm50 | 68653 | 0.97153 | 1.028472254 | 1.05861 | 0.50885 | 0.88031 |
| 148683864 | tripartite motif protein 37 [Mus musculus] | 0 | 40.292 | 10 | 9 | 24 | 9 | 1049 | 117.3 | 5.34 | 452 | 9 | 27911961.9 | 10842617.21 | 25191368.8 | 14177626.47 | 20042195.82 | 16007936.44 | 18894759.81 | 55309011.78 | 42.4718 | 67.3889 | Trim37 | 68729 | 0.82944 | 1.17056354 | 1.41128 | 0.45975 | 0.862375 |
| 755544327 | PREDICTED: F-actin-uncapping protein LRRC16A isoform X1 [Mus musculus] | 0 | 29.343 | 7 | 7 | 20 | 7 | 1430 | 158 | 8.02 | 105 | 7 | 3519646.093 | 2450211.628 | 3785118.619 | 3450309.612 | 4174976.392 | 3087375.907 | 3762465 | 3733918.871 | 17.7343 | 12.1765 | Carmil1 | 68732 | 0.94445 | 1.055551745 | 1.11764 | 0.3331 | 0.778439 |
| 31559970 | transmembrane protein 214 [Mus musculus] | 0 | 23.618 | 10 | 5 | 15 | 5 | 687 | 76.4 | 9.29 | 198 | 5 | 5099657.031 | 8522523.427 | 7678031.783 | 7933931.29 | 9632445.531 | 8710050.128 | 8193734 | 5341383.747 | 20.7215 | 23.2178 | Tmem214 | 68796 | 0.95674 | 1.043256323 | 1.09042 | 0.60039 | 0.905186 |
| 568968377 | PREDICTED: mitochondrial inner membrane protease ATP23 homolog isoform X1 [Mus musculus] | 0.001 | 6.893 | 8 | 2 | 5 | 2 | 272 | 29.6 | 7.37 | 15 | 2 | 1722750.852 | 2644707.907 | 3005960.831 | 2354156.366 | 2187414.011 | 2526485.511 | 2646199 | 2405326.788 | 22.3185 | 8.01941 | Atp23 | 68876 | 0.99806 | 1.001941689 | 1.00389 | 0.9749 | 0.997464 |
| 30794432 | charged multivesicular body protein 2b [Mus musculus] | 0 | 9.988 | 12 | 3 | 31 | 3 | 213 | 23.9 | 8.78 | 126 | 3 | 1750396.047 | 2871572.299 | 11149380.32 | 15279928.76 | 8357291.45 | 3172073.631 | 12723269.59 | 14136897.31 | 84.1605 | 51.4688 | Chmp2b | 68942 | 0.89432 | 1.105676397 | 1.23633 | 0.66989 | 0.922517 |
| 114150032 | e: Full=CDP-diacylglycerol synthase; Short=CDP-DAG synthase; AltName: Full=Mitochondrial translocator assem | 0 | 26.202 | 26 | 7 | 28 | 7 | 337 | 37.8 | 8.98 | 251 | 7 | 36437303.03 | 55801343.71 | 48553058.42 | 46994447.72 | 53167770.06 | 47486353.45 | 54241023 | 50856768.61 | 17.0156 | 5.81088 | Tamm41 | 68971 | 0.95435 | 1.045651904 | 1.09567 | 0.3328 | 0.778439 |
| 12840819 | unnamed protein product [Mus musculus] | 0 | 11.112 | 27 | 3 | 5 | 3 | 102 | 11.8 | 9.31 | 35 | 3 | 16394346.95 | 20129781.39 | 10392284.58 | 20379086.88 | 12976159.65 | 16803245.47 | 10167229 | 24070941.16 | 27.6927 | 37.6542 | Spcs1 | 69019 | 1.02496 | 0.975037333 | 0.95129 | 0.83677 | 0.969547 |
| 19526878 | pyrroline-5-carboxylate reductase 2 [Mus musculus] | 0 | 64.42 | 43 | 12 | 76 | 12 | 320 | 33.6 | 7.77 | 1105 | 12 | 75425664.58 | 122401818.8 | 110828836.5 | 139115543 | 92910845.51 | 115464678.5 | 94239598.81 | 134677740 | 24.0945 | 18.1217 | Pycr2 | 69051 | 1.01184 | 0.988160188 | 0.9766 | 0.88073 | 0.980491 |
| 118200813 | cytochrome c oxidase subunit II (mitochondrion) [Mus musculus domesticus] | 0 | 14.355 | 13 | 2 | 9 | 2 | 227 | 26 | 4.78 | 167 | 2 | 16830332 | 10432640.62 | 19137708.67 | 11770914.71 | 25093147.83 | 13399864.49 | 16788449 | 15555218.21 | 28.3299 | 28.9 | Vps26b | 69091 | 0.90183 | 1.098172644 | 1.21772 | 0.37235 | 0.80688 |
| 148681362 | mCG140375 [Mus musculus] | 0 | 35.158 | 3 | 10 | 20 | 10 | 5189 | 573.6 | 6.2 | 173 | 10 | 14396368.28 | 19712237.9 | 16087288.27 | 14227050.53 | 19526443.25 | 16800747.78 | 14750134.31 | 14958874.29 | 15.813 | 13.4022 | Ubr4 | 69116 | 0.98763 | 1.012365976 | 1.02504 | 0.81899 | 0.964114 |
| 50510421 | mKIAA0227 protein [Mus musculus] | 0 | 14.173 | 17 | 2 | 9 | 2 | 271 | 29.4 | 9.51 | 130 | 2 | 3574134.026 | 1986650.399 | 9171383.862 | 11798371.72 | 9194230.881 | 6048077.594 | 7087069.375 | 7938681.03 | 69.6754 | 17.6041 | Ttc9 | 69480 | 0.9342 | 1.065803012 | 1.14088 | 0.71102 | 0.935294 |
| 148679826 | RIKEN cDNA 2310022B05 [Mus musculus] | 0 | 25.067 | 21 | 5 | 15 | 5 | 322 | 35.3 | 5.24 | 106 | 5 | 7283332.406 | 13799938.59 | 18033603.66 | 23229584.42 | 12744489.18 | 10281305.36 | 14691778 | 21684748.15 | 43.2829 | 32.9988 | 10022B05F | 69551 | 1.02418 | 0.975817923 | 0.95278 | 0.86568 | 0.978314 |
| 110625761 | AFG3-like protein 2 [Mus musculus] | 0 | 10.048 | 3 | 3 | 13 | 2 | 802 | 89.5 | 8.6 | 54 | 3 | 3342688.615 | 3573990.945 | 1628103.532 | 3709087.455 | 3728913.046 | 4360270.327 | 2391846.75 | 3948383.689 | 31.6241 | 23.606 | Afg3l2 | 69597 | 0.91847 | 1.081532065 | 1.17754 | 0.43137 | 0.8471 |
| 166295212 | disabled homolog 2-interacting protein isoform 2 [Mus musculus] | 0 | 249.771 | 47 | 40 | 223 | 14 | 1189 | 131.6 | 8.72 | 4027 | 40 | 73586885.06 | 71670824.37 | 75018748.33 | 61596536.11 | 82437782.52 | 84385470.65 | 76025512.13 | 76136628.58 | 8.61582 | 5.40035 | Dab2ip | 69601 | 0.93823 | 1.061765635 | 1.13166 | 0.04698 | 0.591743 |
| 148693600 | Rho guanine nucleotide exchange factor (GEF) 12, isoform CRA_c [Mus musculus] | 0 | 24.385 | 5 | 7 | 24 | 7 | 1554 | 173.6 | 5.81 | 127 | 7 | 4657137.139 | 2201320.144 | 5685550.88 | 7701461.716 | 4340165.545 | 7838106.992 | 7592358.688 | 5471347.549 | 45.2025 | 26.7727 | Arhgef12 | 69632 | 0.89016 | 1.109843683 | 1.2468 | 0.41351 | 0.835788 |
| 32027990 | death-associated protein kinase-beta [Mus musculus] | 0 | 44.507 | 11 | 12 | 38 | 12 | 1442 | 161.4 | 6.89 | 270 | 12 | 31448031.57 | 23760648.15 | 41335253.13 | 37373579.07 | 38040822.01 | 19795727.97 | 33011849.38 | 28720511.38 | 22.8429 | 25.8729 | Dapk1 | 69635 | 1.05661 | 0.943394991 | 0.89285 | 0.53399 | 0.886555 |
| 568961999 | PREDICTED: muscular LMNA-interacting protein isoform X1 [Mus musculus] | 0 | 116.003 | 31 | 18 | 83 | 18 | 1036 | 110.8 | 8.35 | 1259 | 18 | 75017373.81 | 73863378.61 | 78751140.28 | 63232333.97 | 78158026.82 | 34409189.93 | 73161489.5 | 53720864.85 | 9.15577 | 33.3707 | Mlip | 69642 | 1.09695 | 0.903048618 | 0.82323 | 0.26793 | 0.745606 |
| 299522842 | dynactin subunit 2 isoform 1 [Mus musculus] | 0 | 52.299 | 26 | 8 | 34 | 8 | 407 | 44.7 | 5.21 | 824 | 8 | 43584202.38 | 36012403.37 | 42654360.34 | 40293996.09 | 49895172 | 67037566.61 | 45167718.25 | 42435924.3 | 8.31613 | 21.5923 | Dctn2 | 69654 | 0.88561 | 1.114392681 | 1.25834 | 0.11889 | 0.678142 |
| 568953480 | PREDICTED: calmodulin-regulated spectrin-associated protein 3 isoform X1 [Mus musculus] | 0 | 71.082 | 18 | 17 | 72 | 17 | 1280 | 138.3 | 8.54 | 766 | 17 | 29154334.39 | 22443678.44 | 27080249.44 | 31007885.5 | 28989289.65 | 16400995.01 | 27254094.61 | 18412053.51 | 13.442 | 27.5905 | Camsap3 | 69697 | 1.0928 | 0.907195997 | 0.83015 | 0.24811 | 0.733806 |
| 568934506 | PREDICTED: CAD protein isoform X1 [Mus musculus] | 0 | 25.285 | 3 | 6 | 19 | 6 | 2321 | 252.9 | 6.6 | 236 | 6 | 5118349.071 | 4615431.672 | 6071535.967 | 3946362.688 | 7063977.267 | 4434577.018 | 5269192 | 6823727.416 | 18.1316 | 21.3427 | Cad | 69719 | 0.91141 | 1.088590562 | 1.1944 | 0.26027 | 0.745606 |
| 31541888 | NF-kappa-B inhibitor-interacting Ras-like protein 1 isoform 1 [Mus musculus] | 0 | 12.033 | 18 | 3 | 3 | 3 | 192 | 21.6 | 7.33 | 35 | 3 | 9020378.647 | 4232204.419 | 5345724.472 | 3465378.541 | 7092250.981 | 3888456.075 | 5001655.25 | 4082153.298 | 44.608 | 29.2415 | Nkiras1 | 69721 | 1.04745 | 0.952545554 | 0.90939 | 0.73905 | 0.943908 |
| 29243896 | ubiquitin carboxyl-terminal hydrolase 46 [Mus musculus] | 0 | 31.378 | 15 | 5 | 24 | 5 | 366 | 42.4 | 6.83 | 614 | 5 | 14738028.07 | 11269959.5 | 17478237.66 | 9502462.82 | 16415975.11 | 19086328.6 | 17541395.34 | 11584455.87 | 26.8864 | 20.0468 | Usp46 | 69727 | 0.90104 | 1.098960885 | 1.21966 | 0.27219 | 0.745606 |
| 12845764 | unnamed protein product [Mus musculus] | 0 | 23.908 | 23 | 6 | 11 | 6 | 326 | 36.7 | 6.09 | 84 | 6 | 6602174.976 | 6945190.891 | 8160407.179 | 8340305.568 | 9114983.761 | 11459663.72 | 8452637.25 | 8906880.889 | 11.5428 | 14.1944 | Nup37 | 69736 | 0.884 | 1.116002158 | 1.26245 | 0.04895 | 0.592616 |
| 31981560 | stromal membrane-associated protein 2 [Mus musculus] | 0 | 13.371 | 7 | 3 | 18 | 3 | 428 | 46.5 | 8.87 | 133 | 3 | 6872493.045 | 6989162.713 | 6427918.076 | 3453672.993 | 8470033.762 | 6758267.702 | 7052341.25 | 6423297.372 | 28.1736 | 12.5441 | Smap2 | 69780 | 0.90542 | 1.094584544 | 1.20893 | 0.23939 | 0.732938 |
| 148699134 | tripartite motif protein 32, isoform CRA_a, partial [Mus musculus] | 0.001 | 6.512 | 3 | 2 | 7 | 2 | 666 | 73.1 | 7.01 | 22 | 2 | 356076.8038 | 609667.9856 | 382296.0631 | 410139.3808 | 441995.7487 | 629471.7944 | 443551.9063 | 362656.4592 | 26.2871 | 24.1129 | Trim32 | 69807 | 0.96713 | 1.032865897 | 1.06797 | 0.72452 | 0.939018 |
| 168480117 | mitochondrial folate transporter/carrier [Mus musculus] | 0 | 10.789 | 8 | 2 | 8 | 2 | 316 | 35 | 9.47 | 138 | 2 | 7253186.349 | 7852813.56 | 10600492.49 | 8645579.835 | 12361870.48 | 9601956.344 | 9638677.5 | 10427007.35 | 16.975 | 12.3111 | Slc25a32 | 69906 | 0.89949 | 1.100514274 | 1.22349 | 0.0964 | 0.670764 |
| 148671597 | nucleoporin 43, isoform CRA_a, partial [Mus musculus] | 0 | 20.692 | 15 | 4 | 8 | 4 | 396 | 43.7 | 5.33 | 20 | 4 | 14491845.79 | 5129352.757 | 12134118.45 | 9325188.405 | 12629273.19 | 7530805.355 | 13075566 | 6140288.47 | 39.1978 | 35.8049 | Nup43 | 69912 | 1.02119 | 0.978813723 | 0.95851 | 0.87867 | 0.980491 |
| 37537518 | acylglycerol kinase, mitochondrial precursor [Mus musculus] | 0 | 146.693 | 62 | 20 | 79 | 20 | 421 | 46.9 | 8.4 | 1382 | 20 | 300968817.4 | 246453753.1 | 271418658.5 | 262235066.1 | 288212841.7 | 327853046 | 248198747.9 | 265281486 | 8.47918 | 12.2032 | Agk | 69923 | 0.97807 | 1.021925873 | 1.04483 | 0.57949 | 0.897763 |
| 37359832 | mKIAA0193 protein [Mus musculus] | 0 | 23.713 | 13 | 4 | 19 | 4 | 439 | 49.1 | 4.94 | 251 | 4 | 12823781.35 | 12049386.69 | 7547402.973 | 9129205.247 | 9108950.581 | 14103792.39 | 9709796.5 | 15798248.51 | 23.8073 | 26.9501 | Scrn1 | 69938 | 0.92056 | 1.079439093 | 1.17259 | 0.41652 | 0.836939 |
| 89111939 | exocyst complex component 1 isoform 1 [Mus musculus] | 0 | 34.471 | 12 | 7 | 13 | 7 | 901 | 102.6 | 6.47 | 130 | 7 | 2493427.115 | 2465848.198 | 2248482.594 | 5211766.736 | 3894423.157 | 1827329.971 | 3432496.688 | 1845006.741 | 45.3755 | 38.9748 | Exoc1 | 69940 | 1.06065 | 0.939353458 | 0.88564 | 0.70219 | 0.931521 |
| 219276601 | asparagine--tRNA ligase, cytoplasmic isoform 1 [Mus musculus] | 0.001 | 6.219 | 3 | 2 | 8 | 2 | 559 | 64.2 | 5.86 | 23 | 2 | 2504721.007 | 1630456.48 | 1463239.151 | 2490716.891 | 948126.0546 | 3238192.031 | 1831519.375 | 1159567.318 | 27.3574 | 57.6041 | Nars | 70223 | 1.05972 | 0.940279274 | 0.88729 | 0.71082 | 0.935294 |
| 568934520 | PREDICTED: actin filament-associated protein 1 isoform X1 [Mus musculus] | 0 | 19.974 | 8 | 7 | 21 | 6 | 820 | 90 | 8.57 | 193 | 7 | 9147276.936 | 7184026 | 9732209.382 | 10815511.91 | 12459488.73 | 12028857.5 | 12721931.75 | 11720668.15 | 16.5188 | 3.63911 | Afap1 | 70292 | 0.85955 | 1.140448969 | 1.3268 | 0.00899 | 0.434958 |
| 21362321 | E3 ubiquitin-protein ligase RNF126 [Mus musculus] | 0 | 22.302 | 18 | 4 | 15 | 4 | 313 | 34.1 | 5.17 | 95 | 4 | 19721054.96 | 14089028.07 | 17405263.72 | 9809512.304 | 16539229.9 | 7356706.915 | 11991924.63 | 10179244.87 | 28.2137 | 33.459 | Rnf126 | 70294 | 1.13967 | 0.860327965 | 0.75489 | 0.24306 | 0.733806 |
| 19526912 | hsc70-interacting protein [Mus musculus] | 0 | 7.452 | 8 | 3 | 6 | 3 | 371 | 41.6 | 5.26 | 31 | 3 | 3335412.515 | 2987113.401 | 2907437.983 | 831983.3116 | 3774042.11 | 3583174.412 | 1351433 | 3264756.623 | 45.2245 | 37.2355 | St13 | 70356 | 0.91325 | 1.0867451 | 1.18997 | 0.5704 | 0.895889 |
| 58037205 | BAG family molecular chaperone regulator 5 [Mus musculus] | 0 | 49.641 | 29 | 8 | 28 | 8 | 447 | 50.9 | 6.05 | 359 | 8 | 14035934.01 | 10175793.79 | 15125943.59 | 8531586.03 | 19142420.37 | 11760388.61 | 10567264.44 | 13810921.84 | 26.1008 | 27.4421 | Bag5 | 70369 | 0.92815 | 1.071853802 | 1.15483 | 0.47924 | 0.865853 |
| 1039741861 | PREDICTED: inverted formin-2 isoform X2 [Mus musculus] | 0 | 50.31 | 10 | 10 | 28 | 10 | 1296 | 141.3 | 5.35 | 275 | 10 | 14670765.56 | 19399052.26 | 21876890.36 | 15098762.95 | 16690584.81 | 24597078.97 | 19358174.25 | 26858768.83 | 19.5741 | 21.3529 | Inf2 | 70435 | 0.89619 | 1.103810329 | 1.23167 | 0.20729 | 0.715071 |
| 21312594 | mitochondrial pyruvate carrier 2 [Mus musculus] | 0 | 44.958 | 46 | 6 | 103 | 6 | 127 | 14.3 | 10.61 | 693 | 6 | 415170178.1 | 335099144.1 | 261381976.9 | 232597044.7 | 330461402.7 | 585732009.8 | 244936793.6 | 405263943.6 | 26.2757 | 37.0415 | Mpc2 | 70456 | 0.88538 | 1.114616429 | 1.25891 | 0.37069 | 0.80688 |
| 28972684 | mKIAA1246 protein [Mus musculus] | 0.001 | 4.819 | 2 | 2 | 10 | 2 | 833 | 89.8 | 7.01 | 39 | 2 | 2551302.95 | 2366015.252 | 2695202.507 | 1836615.448 | 2940061.916 | 2617198.654 | 2795110.813 | 2218507.049 | 15.8938 | 11.8103 | Lrfn2 | 70530 | 0.94397 | 1.056031042 | 1.11871 | 0.29438 | 0.759552 |
| 148694207 | mCG141703, isoform CRA_c [Mus musculus] | 0 | 184.699 | 21 | 34 | 105 | 32 | 2585 | 276 | 5.62 | 1483 | 34 | 78224491.28 | 47892454.27 | 78389542.89 | 58415887.93 | 82045849.15 | 68636562.28 | 83541044.81 | 53147761.38 | 23.04 | 19.6969 | Tln2 | 70549 | 0.95557 | 1.044428722 | 1.09299 | 0.57685 | 0.895889 |
| 941355852 | redox-regulatory protein FAM213A isoform a [Mus musculus] | 0 | 24.029 | 21 | 4 | 21 | 4 | 232 | 25.9 | 9.03 | 202 | 4 | 15002102.52 | 11634012.2 | 11443027.66 | 12230711.66 | 13099581.69 | 16156608.04 | 12545046.25 | 13210987.15 | 13.1257 | 11.8419 | Prxl2a | 70564 | 0.95535 | 1.044647516 | 1.09347 | 0.34977 | 0.797109 |
| 568988757 | PREDICTED: importin-5 isoform X1 [Mus musculus] | 0 | 29.61 | 8 | 6 | 22 | 6 | 1123 | 126.4 | 4.97 | 166 | 6 | 6612570.676 | 6885725.861 | 8959334.195 | 7622560.769 | 10133380.43 | 9480546.012 | 9971029.188 | 6378279.646 | 13.9636 | 19.6162 | Ipo5 | 70572 | 0.91092 | 1.089078415 | 1.19558 | 0.20182 | 0.715071 |
| 226371690 | dnaJ homolog subfamily B member 14 [Mus musculus] | 0 | 7.19 | 5 | 2 | 15 | 2 | 379 | 42.3 | 8.59 | 156 | 2 | 3489997.068 | 4394771.117 | 3891619.86 | 1579459.183 | 4735003.579 | 5192374.358 | 4539824.75 | 5019023.907 | 36.8383 | 5.96545 | Dnajb14 | 70604 | 0.81334 | 1.186662371 | 1.459 | 0.0515 | 0.595816 |
| 160333312 | serine/threonine-protein kinase SIK3 [Mus musculus] | 0 | 110.126 | 20 | 19 | 56 | 19 | 1369 | 150.6 | 7.02 | 766 | 19 | 54835232.43 | 49768680.32 | 60665446.74 | 46208069.43 | 44942700.48 | 50706450.59 | 46131010.84 | 48558803.45 | 11.8942 | 5.39719 | Sik3 | 70661 | 1.05261 | 0.94739273 | 0.90004 | 0.17071 | 0.711915 |
| 226437676 | nuclear pore complex protein Nup205 [Mus musculus] | 0 | 177.06 | 22 | 33 | 123 | 31 | 2008 | 227.3 | 6.34 | 1885 | 33 | 48798927.17 | 39904245.89 | 62537783.5 | 54621175.28 | 68423477.21 | 63935849.09 | 75219323.69 | 70750409.97 | 18.5466 | 6.75947 | Nup205 | 70699 | 0.85033 | 1.149665936 | 1.35202 | 0.01441 | 0.461382 |
| 158508485 | carboxyl-terminal PDZ ligand of neuronal nitric oxide synthase protein isoform 1 [Mus musculus] | 0 | 26.691 | 16 | 7 | 21 | 7 | 503 | 55.8 | 6.09 | 78 | 7 | 24000705.63 | 26941267.27 | 25926808.95 | 23380979.64 | 26753785.6 | 36382524.77 | 26075588.63 | 28305859.15 | 6.60939 | 16.2057 | Nos1ap | 70729 | 0.9207 | 1.079295556 | 1.17225 | 0.1376 | 0.693437 |
| 568924022 | PREDICTED: serine/threonine-protein kinase DCLK2 isoform X1 [Mus musculus] | 0 | 122.703 | 39 | 21 | 102 | 20 | 772 | 84.6 | 8.62 | 1976 | 21 | 175285549.6 | 100644110.8 | 166112026.3 | 113789174 | 190270454 | 125421265.3 | 166556550.6 | 118724689.2 | 26.7931 | 22.6621 | Dclk2 | 70762 | 0.96098 | 1.039023123 | 1.08122 | 0.6703 | 0.922517 |
| 242397483 | phytanoyl-CoA hydroxylase-interacting protein-like isoform 1 [Mus musculus] | 0 | 18.862 | 17 | 6 | 13 | 5 | 375 | 42.3 | 6.35 | 22 | 6 | 11091264.68 | 23233620.22 | 10487894.92 | 15939966.22 | 15187187.93 | 18140769.19 | 16523668.13 | 14120853.51 | 38.7979 | 10.8593 | Phyhipl | 70911 | 0.97419 | 1.025814607 | 1.053 | 0.80205 | 0.959776 |
| 166851842 | storkhead-box protein 2 isoform 1 [Mus musculus] | 0 | 33.017 | 13 | 5 | 12 | 5 | 926 | 102.7 | 8.46 | 73 | 5 | 7485845.509 | 7853015.716 | 4593122.992 | 4405290.284 | 10575436.6 | 12447899.43 | 6988359.375 | 6485137.378 | 30.2098 | 31.4349 | Stox2 | 71069 | 0.80012 | 1.199880607 | 1.49963 | 0.12456 | 0.692614 |
| 58037239 | golgin subfamily A member 7B [Mus musculus] | 0 | 14.146 | 20 | 3 | 17 | 3 | 167 | 18.3 | 6.01 | 209 | 3 | 52970147.04 | 42595830.58 | 48994290.22 | 41900466.21 | 50922257.01 | 47148732.57 | 41092086.5 | 40189661.81 | 11.3804 | 11.3718 | Golga7b | 71146 | 1.01943 | 0.980569343 | 0.96188 | 0.64621 | 0.915705 |
| 1039746878 | PREDICTED: disks large homolog 5 isoform X1 [Mus musculus] | 0 | 56.145 | 9 | 13 | 26 | 13 | 1931 | 215.3 | 7.46 | 181 | 13 | 14764552.4 | 8046158.246 | 15123758.69 | 9698325.562 | 14474157.04 | 9625265.484 | 15191485.5 | 9281355.219 | 30.0039 | 25.7167 | Dlg5 | 71228 | 0.99023 | 1.00976527 | 1.01972 | 0.92437 | 0.989849 |
| 87159826 | rho GTPase-activating protein 26 [Mus musculus] | 0 | 49.839 | 14 | 9 | 43 | 9 | 814 | 92 | 6.68 | 1105 | 9 | 50336290.02 | 39352036.4 | 46544123.15 | 38479268.37 | 36299446.41 | 33809495.72 | 47120969.75 | 44516248.17 | 13.1047 | 15.7937 | Arhgap26 | 71302 | 1.03854 | 0.961464544 | 0.92579 | 0.47832 | 0.864994 |
| 568914296 | PREDICTED: formin-like protein 2 isoform X1 [Mus musculus] | 0 | 24.988 | 7 | 6 | 18 | 6 | 1098 | 124.7 | 7.65 | 116 | 6 | 6615971.022 | 5726534.583 | 6855136.501 | 5383624.816 | 9582962.104 | 23641148.95 | 11112370.38 | 5550808.845 | 11.433 | 62.5971 | Fmnl2 | 71409 | 0.66018 | 1.339821588 | 2.02948 | 0.15761 | 0.700461 |
| 187956405 | Arhgap21 protein [Mus musculus] | 0 | 153.341 | 21 | 29 | 131 | 28 | 1954 | 216.7 | 7.64 | 2587 | 29 | 69779015.7 | 71513894 | 84465679.96 | 80431459.67 | 85720734.87 | 68180548.11 | 88573949.72 | 82360029.89 | 9.20455 | 11.1431 | Arhgap21 | 71435 | 0.97045 | 1.029547489 | 1.06089 | 0.44734 | 0.852048 |
| 568993338 | PREDICTED: serine/threonine-protein phosphatase 6 regulatory subunit 2 isoform X1 [Mus musculus] | 0 | 44.996 | 15 | 8 | 18 | 8 | 978 | 106.4 | 4.86 | 346 | 8 | 15105230.69 | 6783197.704 | 17633974.6 | 17444633.77 | 15000746.12 | 9010856.744 | 7934592.5 | 7278443.334 | 35.8357 | 36.0582 | Ppp6r2 | 71474 | 1.18445 | 0.815551627 | 0.68855 | 0.20299 | 0.715071 |
| 23956214 | splicing factor, proline- and glutamine-rich [Mus musculus] | 0 | 120.785 | 33 | 18 | 291 | 17 | 699 | 75.4 | 9.44 | 3599 | 18 | 261462854.1 | 321724481.4 | 417952107.1 | 381697590.1 | 364485285 | 356554916.3 | 451301199.5 | 383918351 | 19.8917 | 11.066 | Sfpq | 71514 | 0.94099 | 1.059005447 | 1.12541 | 0.32628 | 0.772772 |
| 50510765 | mKIAA1026 protein [Mus musculus] | 0 | 35.443 | 9 | 5 | 16 | 5 | 808 | 89.9 | 7.46 | 670 | 5 | 3427495.92 | 5254092.739 | 3687744.101 | 6626763.628 | 3278540.19 | 3212647.106 | 6377517.5 | 4471870.428 | 31.3594 | 34.1271 | Kazn | 71529 | 1.04556 | 0.954439383 | 0.91285 | 0.70698 | 0.934268 |
| 755530462 | PREDICTED: unconventional myosin-Ie isoform X1 [Mus musculus] | 0 | 56.552 | 15 | 13 | 30 | 13 | 1119 | 128 | 9.07 | 361 | 13 | 18972855.52 | 17356402.13 | 21380184.61 | 16412048.12 | 22192939.02 | 13682509.81 | 22108417.75 | 15724640.35 | 11.7337 | 23.7682 | Myo1e | 71602 | 1.00279 | 0.997206362 | 0.99443 | 0.96769 | 0.99686 |
| 37360572 | mKIAA1841 protein [Mus musculus] | 0 | 17.655 | 9 | 5 | 13 | 5 | 719 | 82.1 | 6.71 | 120 | 5 | 9657159.429 | 6372920.512 | 7567243.06 | 8790722.361 | 6872137.034 | 5632403.066 | 5036989.75 | 6419009.396 | 17.7093 | 13.625 | 10010F05F | 71675 | 1.14956 | 0.850439791 | 0.7398 | 0.04326 | 0.588297 |
| 148693635 | vacuolar protein sorting 11 (yeast), isoform CRA_a [Mus musculus] | 0 | 62.97 | 20 | 14 | 34 | 14 | 946 | 108.2 | 6.95 | 234 | 14 | 23033723.37 | 14572967.69 | 21549721.99 | 25079750.79 | 41983574.11 | 14079333.48 | 18369909.63 | 13734178.69 | 21.6525 | 61.0693 | Vps11 | 71732 | 0.9772 | 1.022800232 | 1.04666 | 0.89453 | 0.984401 |
| 28972564 | mKIAA1002 protein [Mus musculus] | 0 | 39.563 | 9 | 7 | 17 | 7 | 1070 | 117.5 | 8.82 | 191 | 7 | 22947203.32 | 13107574.4 | 17417748.04 | 3283059.514 | 18376988.45 | 7864989.219 | 13228458.19 | 7496973.85 | 58.5771 | 43.782 | R3hdm2 | 71750 | 1.09437 | 0.905631576 | 0.82754 | 0.63436 | 0.912586 |
| 568960085 | PREDICTED: C2 domain-containing protein 2-like isoform X1 [Mus musculus] | 0 | 38.49 | 20 | 7 | 21 | 7 | 707 | 76.3 | 7.01 | 204 | 7 | 16479181.11 | 28971180.56 | 13661361.38 | 13544122.88 | 12438438.44 | 9372452.55 | 11394644.5 | 12843785.34 | 40.3627 | 13.4786 | C2cd2l | 71764 | 1.22414 | 0.775860429 | 0.6338 | 0.12621 | 0.693437 |
| 78711838 | AP-2 complex subunit beta isoform a [Mus musculus] | 0 | 216.691 | 43 | 32 | 194 | 17 | 951 | 105.7 | 5.34 | 2520 | 32 | 131471191.1 | 195973674.3 | 155651635.5 | 107556422.8 | 145764808.2 | 168453114.6 | 162454030.3 | 139159286.4 | 25.5448 | 8.94073 | Ap2b1 | 71770 | 0.97913 | 1.020869164 | 1.04263 | 0.76449 | 0.952792 |
| 124486670 | mitochondrial glutamate carrier 2 [Mus musculus] | 0 | 129.32 | 73 | 18 | 255 | 15 | 320 | 34.1 | 9.19 | 4275 | 18 | 355718684.6 | 335950701.4 | 311627111.7 | 393267575.4 | 329327080.9 | 423042590.5 | 304589033.6 | 393232182.6 | 9.8827 | 15.1612 | Slc25a18 | 71803 | 0.98116 | 1.018837875 | 1.0384 | 0.69385 | 0.930425 |
| 148679174 | nucleoporin 93, isoform CRA_b [Mus musculus] | 0 | 164.111 | 44 | 27 | 136 | 27 | 822 | 93.6 | 5.71 | 2229 | 27 | 79594968.23 | 68280088.67 | 90581740.97 | 98615537.65 | 117574811.3 | 111115899.1 | 117457039.9 | 108896105.4 | 15.671 | 3.89396 | Nup93 | 71805 | 0.85107 | 1.14893209 | 1.34999 | 0.00547 | 0.38143 |
| 12847728 | unnamed protein product [Mus musculus] | 0 | 18.448 | 7 | 4 | 15 | 4 | 723 | 81.6 | 7.78 | 98 | 4 | 6026088.827 | 3848335.029 | 3719937.53 | 13053894.08 | 8045938.422 | 11502978.73 | 9962298.75 | 10826102.49 | 65.9051 | 14.8555 | Tars2 | 71807 | 0.79564 | 1.204358374 | 1.51369 | 0.19056 | 0.715071 |
| 58037257 | DDB1- and CUL4-associated factor 7 [Mus musculus] | 0 | 28.928 | 21 | 6 | 20 | 6 | 342 | 38.9 | 5.52 | 436 | 6 | 31932678.48 | 16725060.76 | 30079254.7 | 28046719.18 | 33325770.72 | 30430846.9 | 33801984.75 | 29270256.09 | 25.5996 | 6.95177 | Dcaf7 | 71833 | 0.91419 | 1.08580508 | 1.18772 | 0.21225 | 0.716066 |
| 19526936 | lanC-like protein 2 [Mus musculus] | 0 | 81.935 | 33 | 12 | 63 | 12 | 450 | 50.7 | 7.28 | 1682 | 12 | 57208213.69 | 66607724.53 | 82940351.68 | 45821722.04 | 73197522.73 | 44251495.29 | 68531800.52 | 51698611.67 | 24.8589 | 23.0456 | Lancl2 | 71835 | 1.03039 | 0.969610697 | 0.94101 | 0.73288 | 0.943908 |
| 148702793 | RIKEN cDNA 2310003H01, isoform CRA_a [Mus musculus] | 0 | 33.83 | 12 | 8 | 19 | 8 | 916 | 98 | 5.33 | 158 | 8 | 28964356.46 | 28903813.02 | 33898309.27 | 23928501.02 | 33533435.15 | 43119793.98 | 30156148.98 | 38989267.59 | 14.0723 | 15.7645 | Faap100 | 71885 | 0.88488 | 1.115121988 | 1.2602 | 0.07641 | 0.627416 |
| 37360114 | mKIAA0829 protein [Mus musculus] | 0 | 16.631 | 3 | 3 | 12 | 3 | 1332 | 146.6 | 5.97 | 309 | 3 | 726228.3582 | 3296722.428 | 1833702.243 | 2133818.574 | 3570258.511 | 3184021.254 | 1490764.25 | 2727211.545 | 52.8955 | 32.9259 | Cand1 | 71902 | 0.84276 | 1.157244469 | 1.37317 | 0.32471 | 0.772052 |
| 170014720 | D-beta-hydroxybutyrate dehydrogenase, mitochondrial precursor [Mus musculus] | 0 | 164.784 | 52 | 14 | 264 | 14 | 343 | 38.3 | 9.01 | 4468 | 14 | 1254113635 | 1054026630 | 1069100506 | 1082946682 | 1252495925 | 1340098814 | 1162713090 | 1234490626 | 8.38174 | 5.84655 | Bdh1 | 71911 | 0.94396 | 1.056043576 | 1.11874 | 0.06692 | 0.601634 |
| 148690793 | myosin, heavy polypeptide 14, isoform CRA_a [Mus musculus] | 0 | 359.218 | 36 | 59 | 331 | 49 | 2008 | 229.3 | 5.57 | 6910 | 59 | 175250988.2 | 182671174.6 | 261345834.3 | 151015245.4 | 236636864.7 | 133844828.8 | 176037406 | 157950695.6 | 24.8226 | 24.9222 | Myh14 | 71960 | 1.04463 | 0.955373242 | 0.91456 | 0.6302 | 0.912586 |
| 60360266 | mKIAA4213 protein [Mus musculus] | 0 | 30.618 | 15 | 6 | 33 | 4 | 509 | 58.4 | 7.36 | 382 | 6 | 43345600.54 | 92272487.06 | 83296292.1 | 83173015.34 | 84408687.75 | 169026280 | 68057681.25 | 148808699.8 | 28.9584 | 41.5936 | Ppp2r2a | 71978 | 0.78222 | 1.217784056 | 1.55684 | 0.16747 | 0.711915 |
| 1039738320 | PREDICTED: BTB/POZ domain-containing protein 17 isoform X1 [Mus musculus] | 0 | 92.923 | 39 | 12 | 78 | 12 | 485 | 52.8 | 9.33 | 1630 | 12 | 71938229.7 | 46241599.37 | 47070301.53 | 40823956.71 | 51243795.77 | 47304660.88 | 50649854 | 41483698.26 | 26.9652 | 9.38567 | Btbd17 | 72014 | 1.03879 | 0.961205187 | 0.92531 | 0.6169 | 0.905686 |
| 755492792 | PREDICTED: uncharacterized protein KIAA1211-like homolog isoform X1 [Mus musculus] | 0 | 219.803 | 52 | 33 | 267 | 32 | 1204 | 129.1 | 7.2 | 3207 | 33 | 135687225.4 | 135209704.9 | 170696941 | 192192514.3 | 180056204.3 | 172385629.3 | 163232107.5 | 189018006.2 | 17.652 | 6.23426 | 10300C02F | 72097 | 0.94703 | 1.052974755 | 1.11188 | 0.28271 | 0.750719 |
| 569004573 | PREDICTED: nucleoporin SEH1 isoform X1 [Mus musculus] | 0 | 28.456 | 17 | 5 | 22 | 5 | 417 | 46.1 | 7.84 | 372 | 5 | 18199839.74 | 14170239.55 | 22872105.05 | 17492073.05 | 19921961.7 | 18665964.67 | 22021758.5 | 17137599.42 | 19.7184 | 10.6263 | Seh1l | 72124 | 0.96669 | 1.033313235 | 1.06892 | 0.56687 | 0.895889 |
| 568936214 | PREDICTED: WD repeat and FYVE domain-containing protein 3 isoform X1 [Mus musculus] | 0 | 27.743 | 3 | 8 | 14 | 8 | 3527 | 394.1 | 6.77 | 34 | 8 | 7645374.127 | 4766607.845 | 10041636.46 | 9481446.173 | 7797029.904 | 8205504.965 | 6966000.5 | 8368182.585 | 29.766 | 7.99922 | Wdfy3 | 72145 | 1.00946 | 0.990543231 | 0.98126 | 0.90709 | 0.98561 |
| 17863988 | DEAD/H box polypeptide 36 protein [Mus musculus] | 0 | 18.843 | 5 | 4 | 6 | 4 | 1001 | 113.8 | 8.41 | 113 | 4 | 48207011.46 | 22859234.36 | 26360174.35 | 23156963.95 | 22069057.76 | 19764271.6 | 20984591.38 | 15422816.49 | 40.2863 | 14.9001 | Dhx36 | 72162 | 1.21297 | 0.787034659 | 0.64885 | 0.14098 | 0.693437 |
| 148665046 | RIKEN cDNA 2810401C16, isoform CRA_c, partial [Mus musculus] | 0 | 19.844 | 10 | 4 | 12 | 4 | 634 | 69.7 | 9.11 | 80 | 4 | 7867134.095 | 3853310.298 | 12325296.67 | 6224676.707 | 9733891.941 | 8668350.896 | 13139927.75 | 6740052.925 | 47.2303 | 28.0322 | Aifm3 | 72168 | 0.88313 | 1.116870854 | 1.26467 | 0.40459 | 0.828732 |
| 29789219 | U7 snRNA-associated Sm-like protein LSm11 [Mus musculus] | 0 | 78.105 | 40 | 13 | 81 | 13 | 361 | 39.9 | 10.73 | 1105 | 13 | 50702758.04 | 58834625.37 | 48792947.43 | 66151559.48 | 46279495.59 | 47885989.02 | 40187124.75 | 36000671.95 | 14.2191 | 12.9217 | Lsm11 | 72290 | 1.13709 | 0.86290834 | 0.75887 | 0.03149 | 0.515682 |
| 134031976 | leucine-rich PPR motif-containing protein, mitochondrial precursor [Mus musculus] | 0 | 14.388 | 4 | 6 | 13 | 6 | 1392 | 156.5 | 6.83 | 0 | 6 | 4441425.72 | 4749743.653 | 6009095.383 | 3347416.278 | 6113007.284 | 6967531.762 | 6131345.625 | 3880594.331 | 23.6134 | 22.919 | Lrpprc | 72416 | 0.89086 | 1.109144585 | 1.24503 | 0.23398 | 0.730166 |
| 26329665 | unnamed protein product [Mus musculus] | 0 | 11.343 | 6 | 2 | 4 | 2 | 406 | 44.8 | 6.99 | 37 | 2 | 4899402.87 | 3207251.991 | 1361613.233 | 1768613.23 | 4155573.637 | 1179457.781 | 3529933.75 | 992667.9117 | 57.0506 | 65.482 | Tspyl4 | 72480 | 1.06538 | 0.934615786 | 0.87726 | 0.77196 | 0.952792 |
| 254587960 | serine/threonine-protein phosphatase PGAM5, mitochondrial isoform 2 [Mus musculus] | 0 | 50.169 | 45 | 11 | 139 | 11 | 287 | 31.8 | 9.04 | 1541 | 11 | 46568188.78 | 59912801.63 | 71544401.91 | 77873772.14 | 83904418.75 | 112531268.6 | 81797770.25 | 90731529.92 | 21.5455 | 15.236 | Pgam5 | 72542 | 0.81906 | 1.180944646 | 1.44184 | 0.02836 | 0.515682 |
| 568957623 | PREDICTED: inactive hydroxysteroid dehydrogenase-like protein 1 isoform X1 [Mus musculus] | 0 | 12.235 | 15 | 4 | 19 | 4 | 354 | 39.1 | 8.9 | 23 | 4 | 7556353.107 | 11518450.68 | 7448606.98 | 9982798.965 | 9330212.193 | 15174302.03 | 8532103.75 | 13919506.93 | 21.6711 | 28.0995 | Hsdl1 | 72552 | 0.87479 | 1.125205163 | 1.28625 | 0.22316 | 0.718033 |
| 291463307 | protein shisa-9 isoform 1 precursor [Mus musculus] | 0 | 70.931 | 35 | 12 | 139 | 2 | 424 | 46.8 | 8.62 | 1159 | 12 | 16262228.4 | 9781712.218 | 13833309.85 | 15314750.4 | 14699148.58 | 12037981.59 | 9857795 | 10879473.65 | 20.7133 | 17.5813 | Shisa9 | 72555 | 1.07517 | 0.924828356 | 0.86017 | 0.31736 | 0.766675 |
| 148671484 | mCG2822, isoform CRA_b, partial [Mus musculus] | 0 | 8.527 | 3 | 3 | 9 | 2 | 919 | 106 | 9.99 | 0 | 3 | 1975276.976 | 3958466.49 | 3500942.459 | 3755025.336 | 2253750.583 | 5893928.701 | 3280689.125 | 7047830.659 | 27.3269 | 48.2645 | Bclaf1 | 72567 | 0.83305 | 1.166945708 | 1.4008 | 0.3138 | 0.76536 |
| 148682229 | PDZ domain containing 11, isoform CRA_c, partial [Mus musculus] | 0 | 48.348 | 64 | 6 | 52 | 6 | 149 | 17.2 | 7.15 | 1373 | 6 | 74750157.66 | 91847664.36 | 67706948.04 | 64057552.46 | 64082351.51 | 74144367.89 | 52207017.75 | 63957302.81 | 16.5316 | 14.1031 | Pdzd11 | 72621 | 1.07955 | 0.920450448 | 0.85262 | 0.19943 | 0.715071 |
| 148706782 | mCG5013, isoform CRA_b, partial [Mus musculus] | 0 | 44.408 | 25 | 10 | 31 | 10 | 566 | 62.8 | 5.01 | 273 | 10 | 25317523.99 | 19704270.36 | 25132935.06 | 24103010.8 | 31372070.51 | 24353803.21 | 25109192.47 | 20701291.29 | 11.1538 | 17.4588 | Tdrkh | 72634 | 0.96283 | 1.037174855 | 1.07722 | 0.50648 | 0.88031 |
| 26354779 | unnamed protein product [Mus musculus] | 0 | 41.196 | 16 | 6 | 12 | 6 | 622 | 69.9 | 8.84 | 233 | 6 | 7680662.042 | 14572028.21 | 10609414.46 | 10474364.13 | 10741631.54 | 14283316.98 | 11733452.75 | 13248597.06 | 26.1589 | 12.5798 | Tmem209 | 72649 | 0.92854 | 1.0714622 | 1.15392 | 0.34315 | 0.787666 |
| 257153388 | putative tyrosine-protein phosphatase auxilin isoform a [Mus musculus] | 0 | 27.304 | 9 | 7 | 18 | 7 | 968 | 105.3 | 7.47 | 142 | 7 | 22570526.11 | 23858069.96 | 20854080.64 | 22412492.88 | 19113648.46 | 18496283.49 | 20081065.25 | 18918626.21 | 5.48773 | 3.50132 | Dnajc6 | 72685 | 1.07868 | 0.921315881 | 0.85411 | 0.00344 | 0.38143 |
| 148693689 | sodium channel, voltage-gated, type II, beta [Mus musculus] | 0 | 7.51 | 9 | 2 | 10 | 2 | 232 | 25.8 | 7.31 | 37 | 2 | 5141404.786 | 9802826.235 | 6716209.39 | 7204070.246 | 8195898.134 | 9336551.706 | 7149319.5 | 9030103.887 | 26.8303 | 11.6193 | Scn2b | 72821 | 0.92254 | 1.07746313 | 1.16794 | 0.30669 | 0.761857 |
| 28849905 | ubiquitin-associated and SH3 domain-containing protein B [Mus musculus] | 0 | 99.057 | 26 | 14 | 84 | 14 | 638 | 71.4 | 6.81 | 1742 | 14 | 129708701.8 | 141251065.9 | 164887151.8 | 116014414 | 128586178.4 | 141787796.5 | 160047482.8 | 141094462.9 | 15.0045 | 9.0664 | Ubash3b | 72828 | 0.9825 | 1.017495981 | 1.03562 | 0.70131 | 0.931521 |
| 19111156 | putative ATP-dependent RNA helicase DHX30 isoform 2 [Mus musculus] | 0 | 132.101 | 29 | 29 | 94 | 29 | 1217 | 136.6 | 8.75 | 1464 | 29 | 73718055.56 | 60053117.93 | 68611279.99 | 62880560.15 | 64918366.78 | 70829170.21 | 75572662.88 | 61885496.02 | 9.17592 | 8.94058 | Dhx30 | 72831 | 0.98525 | 1.014750499 | 1.02994 | 0.66126 | 0.919869 |
| 148678076 | protein phosphatase 2 (formerly 2A), regulatory subunit B (PR 52), beta isoform, isoform CRA_a [Mus musculus] | 0 | 13.489 | 9 | 3 | 10 | 2 | 446 | 51.5 | 5.81 | 127 | 3 | 2244069.003 | 2876897.687 | 2455356.699 | 3010788.554 | 2564333.354 | 2788416.433 | 2618316.875 | 3524959.262 | 13.5221 | 15.461 | Ppp2r2b | 72930 | 0.95884 | 1.041158733 | 1.08585 | 0.45609 | 0.860877 |
| 41152116 | leucine-rich repeat-containing protein 47 [Mus musculus] | 0 | 8.15 | 6 | 3 | 6 | 3 | 581 | 63.6 | 8.1 | 36 | 3 | 2814608.189 | 2963335.367 | 2459790.262 | 2410629.462 | 3179842.99 | 2844875.974 | 3651791.5 | 3424972.854 | 10.1301 | 10.5545 | Lrrc47 | 72946 | 0.89671 | 1.103289932 | 1.23038 | 0.03126 | 0.515682 |
| 33469051 | tubulin polymerization-promoting protein [Mus musculus] | 0 | 62.663 | 36 | 6 | 47 | 6 | 218 | 23.6 | 9.42 | 629 | 6 | 113009218.6 | 75991023.83 | 130236944.4 | 69812597.79 | 132052687.7 | 68755608.79 | 87916943.5 | 149046859.3 | 29.9239 | 34.1821 | Tppp | 72948 | 0.94107 | 1.05892722 | 1.12523 | 0.62565 | 0.910246 |
| 148690869 | solute carrier family 17 (sodium-dependent inorganic phosphate cotransporter), member 7 [Mus musculus] | 0 | 18.88 | 11 | 5 | 18 | 2 | 577 | 63.4 | 7.08 | 124 | 5 | 2551059.806 | 1464000.978 | 7695023.188 | 8283544.377 | 13195496.93 | 9442405.816 | 8312625.75 | 14174673.78 | 69.8271 | 25.1918 | Slc17a7 | 72961 | 0.61407 | 1.385934048 | 2.25698 | 0.0315 | 0.515682 |
| 19526956 | ER membrane protein complex subunit 7 precursor [Mus musculus] | 0 | 10.31 | 16 | 2 | 8 | 2 | 241 | 26.3 | 9.23 | 112 | 2 | 5327631.927 | 6836225.737 | 5584435.432 | 6754340.737 | 4385094.642 | 6424762.608 | 3815116 | 8756501.478 | 12.7498 | 38.3361 | Emc7 | 73024 | 1.02341 | 0.976585988 | 0.95424 | 0.82112 | 0.964114 |
| 56090652 | mitochondrial coenzyme A transporter SLC25A42 [Mus musculus] | 0 | 82.011 | 43 | 9 | 34 | 9 | 318 | 35.2 | 10.05 | 952 | 9 | 23626198.73 | 9375662.637 | 29620027.23 | 19110313.16 | 22021306.63 | 19130559.47 | 30472350 | 20661941.53 | 41.7776 | 21.9891 | Slc25a42 | 73095 | 0.93935 | 1.06064852 | 1.12913 | 0.61422 | 0.905186 |
| 12851679 | unnamed protein product [Mus musculus] | 0 | 15.554 | 21 | 5 | 17 | 5 | 252 | 27.5 | 9.7 | 280 | 5 | 13087096.17 | 8399866.83 | 12583834.04 | 8026874.114 | 9961560.398 | 9000856.549 | 11945103.5 | 7144094.515 | 25.4721 | 21.0156 | Cenpv | 73139 | 1.05048 | 0.9495185 | 0.90389 | 0.56738 | 0.895889 |
| 568976065 | PREDICTED: la-related protein 1 isoform X1 [Mus musculus] | 0 | 17.648 | 5 | 5 | 23 | 5 | 1091 | 123.1 | 9.19 | 263 | 5 | 6293605.096 | 4267151.445 | 4410011.351 | 2209344.055 | 7681803.089 | 4832188.097 | 3958530.063 | 9543960.261 | 38.8628 | 39.6013 | Larp1 | 73158 | 0.79544 | 1.204561722 | 1.51434 | 0.20007 | 0.715071 |
| 268370040 | neural Wiskott-Aldrich syndrome protein isoform 1 [Mus musculus] | 0 | 27.785 | 24 | 9 | 18 | 9 | 501 | 54.2 | 7.93 | 35 | 9 | 19330932.21 | 19581342.64 | 22485112.11 | 19113922.75 | 17530247.71 | 20134156.53 | 24494162.5 | 21881344.13 | 7.86516 | 13.9511 | Wasl | 73178 | 0.97856 | 1.021443786 | 1.04383 | 0.6154 | 0.905186 |
| 218505739 | alpha-tubulin N-acetyltransferase 1 isoform 1 [Mus musculus] | 0 | 31.269 | 20 | 7 | 25 | 7 | 421 | 47.1 | 9.98 | 174 | 7 | 29434828.27 | 48655147.77 | 45271016.66 | 46114156.37 | 34312481.76 | 49135800.6 | 41030880.75 | 52163077.46 | 20.6326 | 18.2835 | Atat1 | 73242 | 0.97929 | 1.020707112 | 1.04229 | 0.77347 | 0.952792 |
| 110625952 | centriole, cilia and spindle-associated protein [Mus musculus] | 0 | 32.231 | 28 | 5 | 39 | 5 | 252 | 28.4 | 9.22 | 331 | 5 | 41139680.88 | 27381609.49 | 37187068.65 | 36992361.95 | 36842830.98 | 23277194.27 | 33098649 | 30238297.47 | 16.3977 | 18.5833 | Ccsap | 73420 | 1.0723 | 0.927697941 | 0.86515 | 0.28471 | 0.751585 |
| 569007894 | PREDICTED: heat shock 70 kDa protein 12A isoform X1 [Mus musculus] | 0 | 146.589 | 45 | 25 | 157 | 25 | 695 | 77.4 | 6.87 | 2826 | 25 | 164203802.8 | 188640349.5 | 253041448.7 | 178644146.4 | 255829088.4 | 176169014.8 | 242117567.8 | 204161844.3 | 20.0084 | 16.5137 | Hspa12a | 73442 | 0.94362 | 1.056379215 | 1.1195 | 0.41407 | 0.835788 |
| 22087520 | memory-related protein [Mus musculus] | 0 | 20.415 | 13 | 5 | 9 | 5 | 485 | 53.6 | 9.14 | 69 | 5 | 3950210.041 | 6112845.952 | 6745201.145 | 6638799.621 | 7195184.896 | 9136547.557 | 7375933.125 | 8059796.781 | 22.246 | 11.0705 | Wdr13 | 73447 | 0.84931 | 1.150692349 | 1.35486 | 0.03827 | 0.56531 |
| 21746161 | tubulin beta-2B chain [Mus musculus] | 0 | 543.963 | 81 | 32 | 1288 | 2 | 445 | 49.9 | 4.89 | 28961 | 32 | 131402691.1 | 177745210.1 | 110338107.7 | 208899859 | 169863994.5 | 146328799.5 | 103208652 | 145400523 | 28.364 | 19.6475 | Tubb2b | 73710 | 1.05329 | 0.946710906 | 0.89881 | 0.56691 | 0.895889 |
| 60359854 | mKIAA1649 protein [Mus musculus] | 0.003 | 3.935 | 5 | 2 | 3 | 2 | 453 | 49.7 | 10.42 | 0 | 2 | 2041452.414 | 1053349.752 | 1495284.806 | 994877.5395 | 2017000.872 | 806724.1346 | 1635932.125 | 730880.184 | 34.7139 | 48.6199 | Poldip3 | 73826 | 1.0366 | 0.963395933 | 0.92938 | 0.81249 | 0.961921 |
| 12963799 | V-type proton ATPase subunit D [Mus musculus] | 0 | 21.798 | 20 | 5 | 20 | 5 | 247 | 28.4 | 9.45 | 171 | 5 | 11150075.09 | 7318823.332 | 9658303.798 | 13348025.79 | 14045066.62 | 14415143.25 | 9140021.125 | 6770696.947 | 24.4576 | 33.8301 | Atp6v1d | 73834 | 0.96627 | 1.033731271 | 1.06982 | 0.76006 | 0.952792 |
| 15042971 | hyaluronan and proteoglycan link protein 2 precursor [Mus musculus] | 0.001 | 5.368 | 6 | 2 | 3 | 2 | 341 | 37.9 | 9.09 | 17 | 2 | 5506076.827 | 5230674.717 | 5437432.757 | 4224753.916 | 4364193.486 | 1593546.64 | 3733447.75 | 3940046.047 | 11.6663 | 36.3189 | Hapln2 | 73940 | 1.19887 | 0.801126357 | 0.66823 | 0.04885 | 0.592616 |
| 568996375 | PREDICTED: dynamin-1-like protein isoform X1 [Mus musculus] | 0 | 130.431 | 38 | 21 | 76 | 21 | 755 | 84 | 7.37 | 2240 | 21 | 44892643.97 | 44485016 | 49051680.03 | 44973701.47 | 66239640.4 | 31170720.94 | 42974040.44 | 23008448.57 | 4.67742 | 46.0422 | Dnm1l | 74006 | 1.0577 | 0.942299803 | 0.89089 | 0.61609 | 0.905186 |
| 205360836 | ankyrin repeat and BTB/POZ domain-containing protein BTBD11 isoform 1 [Mus musculus] | 0 | 197.217 | 37 | 28 | 166 | 28 | 1109 | 121.5 | 6.81 | 2743 | 28 | 193003085.8 | 164437742.2 | 183302706 | 178231826.4 | 239009845.8 | 205952520.5 | 198376189.9 | 175181736.7 | 6.62214 | 12.8987 | Btbd11 | 74007 | 0.93526 | 1.064744855 | 1.13845 | 0.13641 | 0.693437 |
| 13386338 | ras-related protein Rap-2b precursor [Mus musculus] | 0 | 15.081 | 21 | 3 | 10 | 2 | 183 | 20.5 | 4.81 | 129 | 3 | 4940764.15 | 3294606.811 | 4272217.786 | 3039394.408 | 4579436.49 | 5053821.71 | 4768388.625 | 3752079.34 | 22.6653 | 12.3242 | Rap2b | 74012 | 0.92265 | 1.077349797 | 1.16767 | 0.25817 | 0.745606 |
| 568963185 | PREDICTED: copine-4 isoform X1 [Mus musculus] | 0 | 34.429 | 15 | 7 | 18 | 6 | 575 | 64.4 | 6.64 | 248 | 7 | 20815846.48 | 13114192.38 | 21614473.47 | 17789244.56 | 22246805.56 | 14755277.73 | 10601590.5 | 24121109.91 | 20.9991 | 35.3812 | Cpne4 | 74020 | 1.01109 | 0.988908111 | 0.97806 | 0.91721 | 0.987992 |
| 119392074 | putative oxidoreductase GLYR1 isoform 1 [Mus musculus] | 0 | 28.45 | 17 | 7 | 13 | 7 | 552 | 60.4 | 9.13 | 98 | 7 | 19874559.1 | 16852250 | 22324084.21 | 19360497.49 | 21769298.1 | 28631822.41 | 24508021.63 | 24338514.21 | 11.4461 | 11.4387 | Glyr1 | 74022 | 0.88272 | 1.117282322 | 1.26573 | 0.02807 | 0.515682 |
| 568968797 | PREDICTED: glutamate receptor-interacting protein 1 isoform X8 [Mus musculus] | 0 | 117.266 | 26 | 22 | 72 | 21 | 1129 | 122.2 | 6.29 | 367 | 22 | 60854763.17 | 66002023.56 | 64146709.88 | 56157019.06 | 63506374.93 | 84850745.59 | 60539789.94 | 76545685.8 | 6.98598 | 15.9294 | Grip1 | 74053 | 0.92812 | 1.071877313 | 1.15489 | 0.16651 | 0.711253 |
| 1039794414 | PREDICTED: cAMP-regulated phosphoprotein 21 isoform X1 [Mus musculus] | 0 | 20.624 | 6 | 4 | 12 | 4 | 844 | 92.4 | 8.56 | 88 | 4 | 15513249.93 | 42141317.24 | 33237802.51 | 48319353.85 | 46381180.64 | 44394167.5 | 37590114.63 | 74064780.75 | 41.0086 | 31.7834 | Arpp21 | 74100 | 0.81496 | 1.185043192 | 1.45412 | 0.19198 | 0.715071 |

| | | | | | | | | | | | | | | | | | | | | | | | | | | | | |
|---|---|---|---|---|---|---|---|---|---|---|---|---|---|---|---|---|---|---|---|---|---|---|---|---|---|---|---|---|
| 21311871 | nebulette [Mus musculus] | 0 | 44.275 | 37 | 9 | 51 | 9 | 270 | 31.1 | 8.31 | 733 | 9 | 25120624.42 | 24704982.71 | 35474648.41 | 35090423.94 | 31406791.44 | 33832633.4 | 37004421.19 | 84929905.65 | 19.9066 | 54.5534 | Nebl | 74103 | 0.78286 | 1.217135226 | 1.55472 | 0.24996 | 0.733806 |
| 23956222 | actin-related protein 3 [Mus musculus] | 0 | 162.765 | 71 | 22 | 181 | 19 | 418 | 47.3 | 5.88 | 2559 | 22 | 395529600.3 | 356056270.7 | 396678979 | 362575361.5 | 384923468.6 | 325077234.2 | 314353099.4 | 350742042.3 | 5.66865 | 9.13254 | Actr3 | 74117 | 1.04704 | 0.952963488 | 0.91015 | 0.12433 | 0.692614 |
| 260763997 | armadillo repeat-containing protein 8 isoform 1 [Mus musculus] | 0 | 9.467 | 5 | 3 | 9 | 3 | 673 | 75.4 | 6.73 | 88 | 3 | 4533701.273 | 3408889.917 | 3839142.435 | 3512537.621 | 4546254.267 | 4257988.695 | 4004771 | 3894993.513 | 13.2778 | 6.94161 | Armc8 | 74125 | 0.95594 | 1.044056627 | 1.09217 | 0.27332 | 0.745606 |
| 568996433 | PREDICTED: dynamin-like 120 kDa protein, mitochondrial isoform X2 [Mus musculus] | 0 | 168.848 | 38 | 27 | 117 | 27 | 997 | 115.5 | 7.55 | 2291 | 27 | 169150093 | 139966135.4 | 139812824.3 | 117237693.6 | 210711670.5 | 170236828.7 | 140479229.8 | 122834081.6 | 15.0337 | 23.8698 | Opa1 | 74143 | 0.93548 | 1.064518524 | 1.13794 | 0.40842 | 0.833638 |
| 21311939 | cap-specific mRNA (nucleoside-2'-O-)-methyltransferase 1 [Mus musculus] | 0 | 7.7 | 2 | 2 | 7 | 2 | 837 | 95.6 | 7.27 | 51 | 2 | 1715104.8 | 1630388.044 | 1401226.717 | 942670.2413 | 1499203.405 | 1212856.671 | 1508657.5 | 1310396.896 | 24.3387 | 10.5225 | Cmtr1 | 74157 | 1.01411 | 0.985894098 | 0.97218 | 0.84006 | 0.969881 |
| 21312654 | actin-related protein 2/3 complex subunit 5-like protein [Mus musculus] | 0 | 70.004 | 58 | 9 | 50 | 9 | 153 | 17 | 6.8 | 641 | 9 | 144408991.1 | 112661202.4 | 142021595.8 | 116895048.4 | 142830679.7 | 138302635.3 | 147014086 | 138315199.1 | 12.8202 | 2.95366 | Arpc5l | 74192 | 0.95337 | 1.046631058 | 1.09782 | 0.1895 | 0.715071 |
| 148704074 | elongation protein 3 homolog (S. cerevisiae), isoform CRA_c, partial [Mus musculus] | 0 | 14.427 | 7 | 3 | 6 | 3 | 591 | 66.9 | 9.07 | 43 | 3 | 2572891.369 | 3452695.691 | 5329316.611 | 2367964.123 | 6138629.048 | 4938348.977 | 4509379.688 | 3234877.718 | 39.3611 | 25.4714 | Elp3 | 74195 | 0.84334 | 1.156660259 | 1.37152 | 0.20768 | 0.715071 |
| 21312662 | centrosomal protein of 97 kDa isoform 1 [Mus musculus] | 0 | 8.849 | 4 | 3 | 7 | 3 | 856 | 94.6 | 5 | 26 | 3 | 5049302.819 | 2886828.187 | 5200248.389 | 5175556.945 | 2390577.42 | 2556586.991 | 2217900.875 | 497994.269 | 24.6697 | 49.8622 | Cep97 | 74201 | 1.40997 | 0.590033553 | 0.41847 | 0.01137 | 0.461382 |
| 75992920 | long-chain-fatty-acid--CoA ligase 3 isoform a [Mus musculus] | 0 | 65.438 | 25 | 13 | 48 | 13 | 720 | 80.4 | 8.54 | 708 | 13 | 22773265.22 | 18484767.24 | 30166727.61 | 12717684.11 | 27374793.12 | 17607024.87 | 30379921.13 | 18898347.94 | 34.9433 | 26.6404 | Acsl3 | 74205 | 0.94329 | 1.05671244 | 1.12024 | 0.6195 | 0.908128 |
| 124486630 | signal-induced proliferation-associated 1-like protein 3 [Mus musculus] | 0 | 183.738 | 25 | 31 | 163 | 31 | 1776 | 194.9 | 8.32 | 3259 | 31 | 123377964.5 | 115554996.1 | 191218041 | 160168140 | 175641530.2 | 115570785.6 | 164313544.3 | 140113247.6 | 23.713 | 17.939 | Sipa1l3 | 74206 | 0.99551 | 1.004485795 | 1.00901 | 0.95378 | 0.994906 |
| 15214774 | Tubulin, gamma complex associated protein 2 [Mus musculus] | 0 | 34.492 | 12 | 8 | 15 | 8 | 905 | 103.2 | 6.77 | 384 | 8 | 4350998.938 | 3031516.445 | 6853369.979 | 7244151.248 | 6209377.135 | 4897840.306 | 6578403.125 | 3499394.608 | 37.5833 | 26.4003 | Tubgcp2 | 74237 | 1.00691 | 0.993085173 | 0.98627 | 0.95404 | 0.994906 |
| 12834195 | unnamed protein product [Mus musculus] | 0.001 | 5.16 | 4 | 2 | 4 | 2 | 513 | 57.5 | 7.03 | 17 | 2 | 920492.445 | 690118.9828 | 855530.4069 | 769335.9348 | 740069.8449 | 1026235.537 | 1396712.375 | 1034308.298 | 12.4249 | 25.6349 | Smu1 | 74255 | 0.87059 | 1.129405849 | 1.29728 | 0.145 | 0.693437 |
| 189491655 | ubiquitin carboxyl-terminal hydrolase CYLD isoform a [Mus musculus] | 0 | 296.369 | 51 | 36 | 262 | 36 | 955 | 106.8 | 5.63 | 7147 | 36 | 632417458.4 | 558785956.6 | 689869059.9 | 601338771.4 | 608236332.9 | 754859179.1 | 672263212.7 | 621887547.2 | 8.88908 | 9.98754 | Cyld | 74256 | 0.96598 | 1.034016863 | 1.07043 | 0.35011 | 0.797109 |
| 33859724 | heterogeneous nuclear ribonucleoprotein R isoform a [Mus musculus] | 0 | 42.777 | 19 | 10 | 44 | 7 | 632 | 70.8 | 8.13 | 575 | 10 | 23535342.3 | 25583379.33 | 21941618.62 | 19858845.58 | 24984960.35 | 32145632.22 | 25851267 | 15682649.29 | 10.6729 | 27.5117 | Hnrnpr | 74326 | 0.95915 | 1.040854373 | 1.08519 | 0.61034 | 0.905186 |
| 27753993 | leucine-rich repeat transmembrane neuronal protein 1 precursor [Mus musculus] | 0 | 62.607 | 30 | 14 | 195 | 14 | 522 | 58.7 | 7.11 | 2039 | 14 | 383427735 | 373784035 | 425464060.3 | 407987541.9 | 504133484.7 | 461740442.7 | 411623423.5 | 429958496.8 | 5.90162 | 8.9702 | Lrrtm1 | 74342 | 0.9362 | 1.063797784 | 1.13629 | 0.05991 | 0.601634 |
| 568924220 | PREDICTED: ubiquitin-associated protein 2-like isoform X1 [Mus musculus] | 0 | 69.178 | 15 | 11 | 53 | 11 | 1140 | 120.4 | 7.14 | 1075 | 11 | 57573222.98 | 46857354.69 | 62285054.47 | 54772727.24 | 48050599.05 | 52339846.08 | 64420560.06 | 66565069.41 | 11.6805 | 15.6394 | Ubap2l | 74383 | 0.97817 | 1.02183372 | 1.04464 | 0.67222 | 0.922517 |
| 148679073 | mCG14322, isoform CRA_b, partial [Mus musculus] | 0 | 72.157 | 33 | 16 | 82 | 15 | 605 | 67.6 | 8.22 | 419 | 16 | 36518383.21 | 48897701.35 | 38670148.28 | 33313446.57 | 37541112.25 | 48583238.95 | 40309240.44 | 39332723.54 | 17.1155 | 11.817 | Neto2 | 74513 | 0.97411 | 1.025889592 | 1.05316 | 0.63331 | 0.912586 |
| 37360058 | mKIAA0711 protein [Mus musculus] | 0 | 13.176 | 5 | 3 | 10 | 3 | 634 | 68 | 5.39 | 91 | 3 | 5203521.975 | 5142129.14 | 8819901.043 | 4998342.758 | 7622894.958 | 4911703.289 | 6616467.25 | 5268876.263 | 30.7006 | 20.4776 | Kbtbd11 | 74901 | 0.99473 | 1.005270206 | 1.0106 | 0.95621 | 0.994906 |
| 256773180 | rab11 family-interacting protein 2 isoform 1 [Mus musculus] | 0 | 43.007 | 21 | 9 | 32 | 9 | 529 | 60 | 9.28 | 275 | 9 | 24530904.39 | 28266202.94 | 26508764.09 | 33069235.79 | 21081770.52 | 21967554.06 | 24755334 | 21729153.19 | 12.9961 | 7.25955 | Rab11fip2 | 74998 | 1.11313 | 0.886873272 | 0.79674 | 0.02889 | 0.515682 |
| 109730895 | Purine-rich element binding protein G [Mus musculus] | 0 | 43.987 | 35 | 8 | 19 | 7 | 350 | 39.9 | 9.51 | 273 | 8 | 37852992.23 | 44762936.33 | 36193905.81 | 44358145.85 | 28508343.27 | 52163827.52 | 36306197.38 | 47062727.2 | 10.8037 | 25.9383 | Purg | 75029 | 0.99733 | 1.002668371 | 1.00535 | 0.97099 | 0.99686 |
| 568988974 | PREDICTED: SLIT and NTRK-like protein 5 isoform X1 [Mus musculus] | 0 | 60.6 | 13 | 10 | 35 | 10 | 982 | 109.7 | 7.2 | 528 | 10 | 35222181.17 | 24253940.06 | 33120988.31 | 22732289.35 | 35879279.96 | 29264364.28 | 30043508.75 | 29832282.75 | 21.6962 | 9.91991 | Slitrk5 | 75409 | 0.95968 | 1.040316554 | 1.08402 | 0.51367 | 0.88031 |
| 755544831 | PREDICTED: serine/threonine-protein kinase WNK2 isoform X1 [Mus musculus] | 0 | 36.559 | 6 | 10 | 40 | 8 | 2249 | 237.8 | 6.09 | 223 | 10 | 8485375.619 | 9906058.032 | 9879543.948 | 13102113.24 | 7014498.987 | 11486168.85 | 8915409.75 | 9577172.221 | 18.9037 | 19.9543 | Wnk2 | 75607 | 1.05589 | 0.944110686 | 0.89414 | 0.44647 | 0.852048 |
| 28077049 | charged multivesicular body protein 4b [Mus musculus] | 0 | 17.644 | 17 | 3 | 14 | 3 | 224 | 24.9 | 4.82 | 366 | 3 | 5226105.919 | 5950908.85 | 8139515.672 | 6993965.61 | 8587843.876 | 11323039.69 | 5577728.5 | 15926701.09 | 19.2949 | 42.44 | Chmp4b | 75608 | 0.77697 | 1.223028965 | 1.5741 | 0.14978 | 0.693437 |
| 148693644 | mCG10725, isoform CRA_b, partial [Mus musculus] | 0 | 20.292 | 28 | 5 | 33 | 5 | 138 | 15.1 | 10.01 | 336 | 5 | 181749790.5 | 218136818.9 | 212702452.5 | 247853586.2 | 234151780.5 | 228053038.4 | 242148533.5 | 251026577.9 | 12.5889 | 4.17141 | Rps25 | 75617 | 0.94772 | 1.052283347 | 1.11034 | 0.15105 | 0.693437 |
| 124486789 | phosphoinositide 3-kinase regulatory subunit 4 [Mus musculus] | 0 | 12.346 | 3 | 4 | 5 | 4 | 1358 | 152.5 | 7.12 | 45 | 4 | 3494065.243 | 928728.0839 | 1983188.169 | 1084936.193 | 1901990.435 | 1546747.025 | 2549497.875 | 981725.9882 | 62.8245 | 37.6333 | Pik3r4 | 75669 | 1.03531 | 0.964690717 | 0.93179 | 0.85587 | 0.975398 |
| 167234372 | eukaryotic translation initiation factor 4B [Mus musculus] | 0.003 | 3.896 | 3 | 2 | 6 | 2 | 611 | 68.8 | 5.67 | 19 | 2 | 1491243.125 | 5907285.13 | 4774261.918 | 7299036.101 | 3097508.424 | 2712252.419 | 2664230.688 | 4954612.803 | 50.8765 | 32.2442 | Eif4b | 75705 | 1.18368 | 0.816317847 | 0.68964 | 0.30635 | 0.761857 |
| 568960194 | PREDICTED: angiomotin-like protein 1 isoform X1 [Mus musculus] | 0 | 20.755 | 6 | 5 | 8 | 4 | 1005 | 111.7 | 6.6 | 38 | 5 | 1993237 | 1003906.091 | 1552031.803 | 2329296.665 | 2781175.696 | 385725.8831 | 1707478.375 | 1094037.412 | 33.3538 | 68.0241 | Amotl1 | 75723 | 1.07084 | 0.929161512 | 0.8677 | 0.70979 | 0.935294 |
| 568907820 | PREDICTED: mitochondrial fission factor isoform X2 [Mus musculus] | 0 | 52.456 | 39 | 8 | 40 | 8 | 328 | 36.9 | 7.44 | 355 | 8 | 52143407.71 | 27611463.75 | 49926852.99 | 24300774.85 | 38879982.77 | 26430404.18 | 50094975.56 | 29284777.62 | 37.8496 | 29.5823 | Mff | 75734 | 1.03111 | 0.968887961 | 0.93965 | 0.80576 | 0.961747 |
| 124007123 | K2; AltName: Full=Brain-specific serine/threonine-protein kinase 2; Short=BR serine/threonine-protein kinase 2; A | 0 | 36.431 | 15 | 7 | 25 | 6 | 735 | 81.7 | 8.79 | 280 | 7 | 9484076.414 | 7689395.825 | 10284022.93 | 8833712.315 | 8034959.098 | 7841501.09 | 6645337.75 | 10335601.62 | 12.086 | 18.7713 | Brsk2 | 75770 | 1.04966 | 0.950341618 | 0.90538 | 0.39916 | 0.824613 |
| 568918744 | PREDICTED: cytoskeleton-associated protein 5 isoform X1 [Mus musculus] | 0 | 303.046 | 32 | 54 | 267 | 54 | 2040 | 226.6 | 8.02 | 3809 | 54 | 259325484.3 | 202543307.9 | 242623584 | 232375421.3 | 200899630.4 | 266877049.9 | 263241282.3 | 247037761.6 | 10.1866 | 12.4035 | Ckap5 | 75786 | 0.97849 | 1.021508915 | 1.04396 | 0.61274 | 0.905186 |
| 569001947 | PREDICTED: serine/arginine repetitive matrix protein 2 isoform X1 [Mus musculus] | 0 | 41.573 | 4 | 8 | 16 | 8 | 2703 | 294.7 | 12.03 | 236 | 8 | 3030679.276 | 5833680.705 | 11598511.78 | 13206825.98 | 11011410.47 | 9061299.83 | 13494243.44 | 17327731.56 | 56.8719 | 28.0217 | Srrm2 | 75956 | 0.79631 | 1.203690805 | 1.51159 | 0.19916 | 0.715071 |
| 569009115 | PREDICTED: dedicator of cytokinesis protein 11 isoform X1 [Mus musculus] | 0 | 34.542 | 5 | 7 | 20 | 6 | 2086 | 239.1 | 7.75 | 551 | 7 | 6813622.671 | 6253510.992 | 5638686.869 | 2860336.261 | 8390402.283 | 7649869.617 | 9142997 | 8897160.863 | 32.5392 | 7.73896 | Dock11 | 75974 | 0.77511 | 1.224888421 | 1.58027 | 0.01564 | 0.461382 |
| 254588016 | janus kinase and microtubule-interacting protein 1 [Mus musculus] | 0 | 11.473 | 8 | 5 | 7 | 5 | 626 | 73.1 | 6.09 | 24 | 5 | 4726794.243 | 8297510.835 | 5863618.778 | 5672162.58 | 12421042.78 | 4370979.105 | 6126631.5 | 4663685.248 | 24.7843 | 54.5689 | Jakmip1 | 76071 | 0.94204 | 1.057961481 | 1.12306 | 0.72247 | 0.939018 |
| 755504644 | PREDICTED: rap guanine nucleotide exchange factor 2 isoform X3 [Mus musculus] | 0 | 210.482 | 29 | 30 | 137 | 5 | 1649 | 184.5 | 6.49 | 3139 | 30 | 2855089.841 | 12033910.87 | 12887001.54 | 2925998.774 | 8811820.74 | 8612178.251 | 9718767.25 | 9184358.13 | 72.1286 | 5.35549 | Rapgef2 | 76089 | 0.91608 | 1.083920584 | 1.18322 | 0.63085 | 0.912586 |
| 40254160 | ras-related protein Rap-2a precursor [Mus musculus] | 0 | 18.208 | 26 | 4 | 15 | 3 | 183 | 20.6 | 4.82 | 161 | 4 | 4936040.519 | 7810303.465 | 6016322.465 | 4380247.445 | 5316571.096 | 6795458.479 | 6385214 | 6996303.146 | 26.1161 | 11.7526 | Rap2a | 76108 | 0.95167 | 1.048330672 | 1.10157 | 0.51194 | 0.88031 |
| 164519055 | protein FAM131B isoform a [Mus musculus] | 0 | 25.128 | 19 | 6 | 19 | 6 | 360 | 38.9 | 4.48 | 200 | 6 | 15048060.49 | 9297987.953 | 11215225.61 | 19320014.04 | 14589491.21 | 13019329.37 | 13096987.25 | 12910396.85 | 32.3091 | 5.92351 | Fam131b | 76156 | 1.01166 | 0.988339976 | 0.97695 | 0.8929 | 0.984401 |
| 226443061 | ubiquitin carboxyl-terminal hydrolase 31 [Mus musculus] | 0 | 34.482 | 9 | 9 | 15 | 9 | 1344 | 146.2 | 9.17 | 268 | 9 | 15753029.18 | 19502121.34 | 23243438.18 | 14745531.26 | 21289545.58 | 31326327.24 | 20563806.5 | 27538733.02 | 21.1519 | 20.4811 | Usp31 | 76179 | 0.84207 | 1.15793224 | 1.37511 | 0.07722 | 0.627416 |
| 12805339 | Abhd12 protein, partial [Mus musculus] | 0 | 49.634 | 27 | 8 | 31 | 8 | 449 | 50.7 | 9.33 | 574 | 8 | 32572464.03 | 16961885.46 | 28989808.75 | 39923448.26 | 25419240.73 | 20096367.09 | 35568171.25 | 17471828.32 | 32.3621 | 32.474 | Abhd12 | 76192 | 1.09167 | 0.908333159 | 0.83206 | 0.45599 | 0.860877 |
| 12837642 | unnamed protein product [Mus musculus] | 0 | 9.5 | 11 | 2 | 12 | 2 | 216 | 24.2 | 6.39 | 91 | 2 | 6289556.638 | 4753464.77 | 5756547.931 | 5168954.703 | 7044430.261 | 5887272.918 | 6517069 | 5773062.563 | 12.2408 | 9.37882 | Rab2b | 76338 | 0.93106 | 1.068940156 | 1.14809 | 0.11916 | 0.678142 |
| 112821631 | KIAA1768 [Mus musculus] | 0.001 | 5.995 | 1 | 2 | 8 | 2 | 1742 | 191.2 | 6.23 | 28 | 2 | 1293210.543 | 2087758.481 | 2697728.808 | 1859934.36 | 3085316.128 | 2853666.342 | 3092053.375 | 2944466.225 | 29.2756 | 3.8617 | Kndc1 | 76484 | 0.79729 | 1.202713802 | 1.50851 | 0.01437 | 0.461382 |
| 568993605 | PREDICTED: trafficking protein particle complex subunit 9 isoform X1 [Mus musculus] | 0 | 78.687 | 20 | 18 | 51 | 18 | 1148 | 128.2 | 6.47 | 628 | 18 | 30090688.04 | 16668811.96 | 39617805.81 | 16427610.22 | 21571909.45 | 19119215.67 | 23883579.13 | 11254531.53 | 43.8204 | 31.566 | Trappc9 | 76510 | 1.13827 | 0.861733343 | 0.75706 | 0.36792 | 0.80688 |
| 568980664 | PREDICTED: autophagy-related protein 2 homolog B isoform X1 [Mus musculus] | 0 | 11.288 | 2 | 3 | 6 | 3 | 2076 | 231.3 | 5.88 | 128 | 3 | 3432886.191 | 2002381.556 | 4834382.265 | 3050209.432 | 4918782.014 | 3035642.858 | 3744782.813 | 3760163.734 | 35.1693 | 20.1738 | Atg2b | 76559 | 0.92566 | 1.074342222 | 1.16063 | 0.47585 | 0.863085 |
| 568932043 | PREDICTED: E3 ubiquitin-protein ligase MIB2 isoform X1 [Mus musculus] | 0 | 37.572 | 13 | 11 | 34 | 11 | 982 | 106.8 | 8.13 | 183 | 11 | 66673228.21 | 36516627.31 | 58364445.56 | 36071267.43 | 79818139.92 | 49327340.78 | 59317947.25 | 32965893.26 | 31.4075 | 35.3952 | Mib2 | 76580 | 0.9432 | 1.056803424 | 1.12045 | 0.65079 | 0.915864 |
| 358439483 | MICOS complex subunit Mic60 isoform 2 [Mus musculus] | 0 | 243.759 | 50 | 33 | 353 | 33 | 746 | 82.9 | 6.61 | 6277 | 33 | 510083185.5 | 548437758.1 | 482547665 | 546872485.4 | 719622617.9 | 711369833.2 | 616707375.3 | 588487209.5 | 6.0744 | 10.0567 | Immt | 76614 | 0.88395 | 1.116052302 | 1.26258 | 0.00973 | 0.448067 |
| 568909760 | PREDICTED: CLIP-associating protein 1 isoform X1 [Mus musculus] | 0 | 236.394 | 35 | 41 | 200 | 36 | 1615 | 177.7 | 8.97 | 4048 | 41 | 151302462 | 138621356.2 | 177750038.6 | 137827735.1 | 179105733.9 | 134483465.4 | 159101831.1 | 136039335.3 | 12.3107 | 13.9217 | Clasp1 | 76707 | 0.99734 | 1.002659108 | 1.00533 | 0.95623 | 0.994906 |
| 112363072 | actin-related protein 2/3 complex subunit 2 [Mus musculus] | 0 | 108.237 | 52 | 17 | 183 | 5 | 300 | 34.3 | 7.36 | 2094 | 17 | 79575548.59 | 76965631.44 | 76188964.59 | 66176021.45 | 74232925.93 | 95639686.85 | 70495037 | 79784351.9 | 7.87071 | 13.8424 | Arpc2 | 76709 | 0.96568 | 1.034319611 | 1.07108 | 0.42954 | 0.845779 |
| 126723792 | sorting nexin-27 isoform 1 [Mus musculus] | 0 | 19.392 | 11 | 5 | 11 | 5 | 539 | 61 | 6.4 | 51 | 5 | 12639063.11 | 13069813.09 | 11542381.02 | 12291155.4 | 11118814.93 | 12114170.48 | 9600782.625 | 11765748.96 | 5.21653 | 9.97345 | Snx27 | 76742 | 1.0525 | 0.94749528 | 0.90023 | 0.10303 | 0.674486 |
| 161016786 | liprin-alpha-3 [Mus musculus] | 0 | 379.902 | 55 | 51 | 343 | 45 | 1194 | 133.3 | 5.64 | 6964 | 51 | 532524472.9 | 503681906.8 | 571924790.4 | 522131090.8 | 420946081 | 656033001.5 | 580251837.6 | 549198382.2 | 5.4121 | 17.7636 | Ppfia3 | 76787 | 0.98244 | 1.017563398 | 1.03575 | 0.72206 | 0.939018 |
| 148696938 | mCG14783, isoform CRA_a, partial [Mus musculus] | 0 | 38.209 | 39 | 10 | 136 | 10 | 190 | 22.1 | 10.48 | 1251 | 10 | 163751166.3 | 208832724.9 | 164602250.4 | 203643891.8 | 206758392 | 209335573.2 | 198498120.4 | 197757714.2 | 13.1631 | 2.87091 | Rpl18a | 76808 | 0.95395 | 1.046047318 | 1.09654 | 0.20359 | 0.715071 |
| 148687609 | Bri3 binding protein, isoform CRA_a, partial [Mus musculus] | 0 | 15.187 | 12 | 2 | 27 | 2 | 283 | 31.3 | 9.19 | 655 | 2 | 3688056.813 | 6457289.332 | 4426792.003 | 5749993.127 | 7372676.274 | 7833593.386 | 6981645.109 | 7062088.19 | 24.6627 | 5.28065 | Bri3bp | 76809 | 0.8199 | 1.180098594 | 1.43932 | 0.01441 | 0.461382 |
| 148698057 | RIKEN cDNA 2410166I05, isoform CRA_a [Mus musculus] | 0 | 40.085 | 34 | 7 | 20 | 7 | 295 | 32.5 | 6.55 | 382 | 7 | 67451897.73 | 44732531.93 | 57241965.28 | 39840521.27 | 60610773.09 | 37139743.44 | 15258205.63 | 37624356.34 | 23.8357 | 49.1762 | Mtfr1l | 76824 | 1.16292 | 0.837082971 | 0.71981 | 0.23712 | 0.73104 |
| 1024864169 | 30S ribosomal protein S4 [Lactobacillus harbinensis] | 0 | 27.622 | 28 | 7 | 37 | 7 | 194 | 22.6 | 10.65 | 409 | 7 | 19673109.3 | 391518562.2 | 346750585.8 | 404083495.2 | 327723789.6 | 426430209.6 | 365305910.3 | 423986186.4 | 28.4551 | 12.4305 | Rps9 | 76846 | 0.9291 | 1.070896876 | 1.15261 | 0.37503 | 0.807546 |
| 39104558 | mKIAA1168 protein [Mus musculus] | 0 | 457.722 | 55 | 63 | 581 | 12 | 1259 | 146.1 | 6.99 | 8414 | 63 | 170055695.4 | 193207875.3 | 213812562.1 | 204174658 | 201069018.1 | 167399378.6 | 199512014.9 | 198832322.8 | 9.63832 | 8.46599 | Cyfip2 | 76884 | 1.00933 | 0.990673473 | 0.98152 | 0.78125 | 0.952792 |
| 58037475 | protein FAM81A [Mus musculus] | 0 | 167.237 | 61 | 22 | 222 | 22 | 364 | 41.7 | 8.75 | 3484 | 22 | 684997004.9 | 812684626.2 | 709641593.5 | 727353499.6 | 675725964.9 | 973304247.5 | 738406887.1 | 786063771.8 | 7.5687 | 16.1564 | Fam81a | 76886 | 0.9609 | 1.039099084 | 1.08138 | 0.42538 | 0.841626 |
| 21313308 | heterogeneous nuclear ribonucleoprotein M isoform a [Mus musculus] | 0 | 271.678 | 63 | 36 | 427 | 36 | 729 | 77.6 | 8.63 | 8442 | 36 | 528157940.2 | 718943367.4 | 800193021.6 | 803711111.8 | 735946839.1 | 518532639.3 | 751758500.8 | 579816492.5 | 18.1188 | 17.8386 | Hnrnpm | 76936 | 1.04873 | 0.951269441 | 0.90707 | 0.47321 | 0.863085 |
| 568918827 | PREDICTED: breast carcinoma-amplified sequence 1 homolog isoform X1 [Mus musculus] | 0 | 56.724 | 23 | 10 | 95 | 10 | 653 | 69.4 | 6.68 | 925 | 10 | 6709281.705 | 23254284.05 | 63919692.63 | 103871608.8 | 73238871.54 | 93064809.14 | 69974246.88 | 131744860.8 | 88.0444 | 30.8563 | Bcas1 | 76960 | 0.69906 | 1.300944938 | 1.861 | 0.15249 | 0.693437 |
| 50511177 | mKIAA1910 protein [Mus musculus] | 0 | 7.311 | 3 | 2 | 5 | 2 | 760 | 84.9 | 6.21 | 15 | 2 | 2110449.303 | 1890745.39 | 2302180.316 | 1998116.579 | 1711293.455 | 1700490.661 | 1560038.25 | 1838097.774 | 8.47126 | 6.6768 | Slitrk1 | 76965 | 1.0987 | 0.90129503 | 0.82032 | 0.01189 | 0.461382 |
| 12851222 | unnamed protein product [Mus musculus] | 0 | 8.055 | 13 | 2 | 8 | 2 | 180 | 20.1 | 9.6 | 49 | 2 | 30303359 | 38860441.32 | 27997962.78 | 28140137.68 | 33394398.28 | 47407780.39 | 32656608.5 | 43086682.49 | 16.3856 | 18.6007 | Arxes2 | 76976 | 0.88915 | 1.110852795 | 1.24935 | 0.13 | 0.693437 |
| 148695611 | zinc finger protein 289, isoform CRA_b [Mus musculus] | 0 | 47.025 | 20 | 11 | 47 | 10 | 534 | 58.1 | 8.12 | 952 | 11 | 40143470.27 | 35979231.8 | 56106291.61 | 49415235.54 | 33033345.71 | 38567449.67 | 54600863.94 | 47978025.46 | 19.9866 | 22.0731 | Arfgap2 | 77038 | 1.01298 | 0.979021802 | 0.95891 | 0.78718 | 0.952792 |
| 568938849 | PREDICTED: SUN domain-containing protein 1 isoform X18 [Mus musculus] | 0 | 98.881 | 29 | 18 | 76 | 18 | 819 | 91.4 | 6.87 | 774 | 18 | 63438870.04 | 65964829.77 | 64479554.31 | 64971653.46 | 64960397.73 | 99103760.36 | 74905616.13 | 84267436.84 | 1.6236 | 17.971 | Sun1 | 77053 | 0.88939 | 1.110605008 | 1.24872 | 0.06906 | 0.601634 |
| 568976364 | PREDICTED: protein TANC2 isoform X5 [Mus musculus] | 0 | 405.703 | 40 | 61 | 397 | 61 | 2042 | 225.4 | 8.25 | 5988 | 61 | 620374534.8 | 481315514.2 | 492916914.1 | 570784133 | 529982095 | 548642153.1 | 682678674.2 | 588851015.3 | 12.1886 | 11.5766 | Tanc2 | 77097 | 0.95908 | 1.040917063 | 1.08533 | 0.36728 | 0.80688 |
| 18031941 | Myotubularin-related protein 2 [Mus musculus] | 0 | 17.75 | 7 | 4 | 17 | 3 | 643 | 73.2 | 7.25 | 140 | 4 | 5615076.967 | 3202908.146 | 8287192.678 | 6257320.184 | 8162022.125 | 3130362.694 | 8141631.625 | 6935948.972 | 35.8704 | 36.0744 | Mtmr2 | 77116 | 0.93953 | 1.060472923 | 1.12873 | 0.65195 | 0.915864 |
| 226443091 | heterogeneous nuclear ribonucleoprotein A0 [Mus musculus] | 0 | 38.469 | 29 | 6 | 56 | 5 | 305 | 30.5 | 9.31 | 610 | 6 | 51620764.37 | 33772238.7 | 49579942.13 | 84218278.7 | 59770406.51 | 34155914.04 | 65540625.75 | 138075871 | 38.6394 | 59.9539 | Hnrnpa0 | 77134 | 0.84837 | 1.151628473 | 1.35746 | 0.45769 | 0.860877 |
| 568907061 | PREDICTED: ras-associated and pleckstrin homology domains-containing protein 1 isoform X1 [Mus musculus] | 0 | 107.431 | 23 | 22 | 83 | 22 | 1318 | 142.5 | 8.94 | 1384 | 22 | 98959112.65 | 97930798.06 | 109757147.5 | 101018634.4 | 99844657.93 | 87386999.36 | 117746459.7 | 105650428.9 | 5.28129 | 12.2926 | Raph1 | 77300 | 0.99638 | 1.003620768 | 1.00727 | 0.91753 | 0.987992 |
| 5803135 | ras-related protein Rab-35 isoform 1 [Homo sapiens] | 0 | 13.521 | 21 | 4 | 33 | 2 | 201 | 23 | 8.29 | 179 | 4 | 8010622.276 | 7633581.586 | 6615647.982 | 7069633.796 | 8784801.109 | 6699613.708 | 5782857.25 | 8569596.712 | 8.38303 | 19.5463 | Rab35 | 77407 | 0.99142 | 1.008575535 | 1.0173 | 0.87788 | 0.980491 |
| 75832043 | microtubule-associated tumor suppressor candidate 2 homolog [Mus musculus] | 0 | 17.26 | 4 | 4 | 15 | 4 | 1353 | 147.3 | 8.24 | 139 | 4 | 2125917.712 | 5077790.675 | 5150597.306 | 8528876.98 | 8734202.504 | 3797940.576 | 4533033.75 | 5644027.081 | 50.1277 | 38.304 | Mtus2 | 77521 | 0.95811 | 1.041888536 | 1.08744 | 0.79744 | 0.958226 |
| 134053896 | LIM and calponin homology domains-containing protein 1 isoform 1 [Mus musculus] | 0 | 116.467 | 29 | 21 | 112 | 21 | 1057 | 118.1 | 5.48 | 1955 | 21 | 47564109.82 | 52060866.01 | 80020384.97 | 64520500.49 | 75166125.25 | 32614432.79 | 56262268.5 | 62556769.58 | 23.8266 | 31.5048 | Limch1 | 77569 | 1.03731 | 0.962685739 | 0.92806 | 0.71598 | 0.936732 |
| 148687681 | vacuolar protein sorting 33A (yeast) [Mus musculus] | 0 | 77.988 | 32 | 14 | 49 | 14 | 598 | 67.5 | 7.08 | 698 | 14 | 34777747.4 | 24380293.48 | 32612911.6 | 24653553.58 | 31474045.53 | 19937799.97 | 22402771.13 | 23320362.27 | 18.4618 | 20.5973 | Vps33a | 77573 | 1.09032 | 0.909676092 | 0.83432 | 0.23691 | 0.73104 |
| 1039738673 | PREDICTED: myosin-10 isoform X2 [Mus musculus] | 0 | 1317.61 | 64 | 134 | 1573 | 4 | 1997 | 231.2 | 5.53 | 35102 | 134 | 43559569.93 | 36945957.58 | 41506444.5 | 23282732.59 | 68353139.07 | 31109085.12 | 42374759.39 | 29740356.17 | 25.1151 | 41.7111 | Myh10 | 77579 | 0.91706 | 1.082944001 | 1.18089 | 0.53715 | 0.886555 |
| 1039738680 | PREDICTED: histone H2A.V isoform X1 [Mus musculus] | 0 | 29.852 | 46 | 4 | 35 | 3 | 131 | 14.2 | 9.35 | 784 | 4 | 7161538.247 | 3385591.28 | 16429769.06 | 30571863.53 | 12925979.5 | 16014973.18 | 15860303.94 | 27578910.3 | 84.1151 | 35.8127 | H2afv | 77605 | 0.88585 | 1.114150135 | 1.25772 | 0.60851 | 0.905186 |
| 37360372 | mKIAA1418 protein [Mus musculus] | 0 | 28.741 | 11 | 8 | 25 | 8 | 857 | 97.2 | 8.43 | 402 | 8 | 9455066.661 | 7571683.998 | 12296551.42 | 12575966.73 | 13386377.13 | 13948622.11 | 13912739.88 | 15572780.84 | 22.8601 | 6.66869 | Smpd4 | 77626 | 0.84885 | 1.151147519 | 1.35612 | 0.02743 | 0.515682 |
| 568907856 | PREDICTED: A-kinase anchor protein SPHKAP isoform X1 [Mus musculus] | 0 | 103.554 | 16 | 18 | 64 | 17 | 1712 | 187.6 | 5.08 | 1095 | 18 | 47739643.96 | 27772821.01 | 35901201.84 | 29271290.95 | 35127136.81 | 45194118.38 | 25543210.13 | 32231724.54 | 25.8533 | 23.6574 | Sphkap | 77629 | 1.00929 | 0.990713978 | 0.9816 | 0.91911 | 0.987994 |
| 1039749702 | PREDICTED: myotubularin-related protein 5 isoform X1 [Mus musculus] | 0 | 148.675 | 17 | 25 | 94 | 24 | 1927 | 214.8 | 7.14 | 1747 | 25 | 148223703 | 259960789.8 | 175158014.4 | 158472969.4 | 180271654.8 | 228447646.1 | 208999842 | 236048874.1 | 27.4443 | 11.6541 | Sbf1 | 77980 | 0.92984 | 1.070164013 | 1.15092 | 0.36125 | 0.804292 |
| 58037519 | trafficking protein particle complex subunit 6B isoform 1 [Mus musculus] | 0 | 20.159 | 27 | 3 | 13 | 3 | 158 | 17.9 | 8.68 | 469 | 3 | 7193740.617 | 7726381.21 | 11333813.33 | 8188849.431 | 11329000.48 | 8454855.522 | 10850258 | 4744811.789 | 21.6055 | 41.0433 | Trappc6b | 78232 | 0.95034 | 1.049663762 | 1.10452 | 0.6917 | 0.930425 |
| 124487457 | MAP7 domain-containing protein 2 isoform 1 [Mus musculus] | 0 | 16.946 | 7 | 5 | 21 | 5 | 781 | 86 | 8.81 | 304 | 5 | 5273890.425 | 5034928.476 | 8312774.937 | 8765831.894 | 8417211.886 | 8276192.845 | 6909014.813 | 5546980.099 | 28.7055 | 18.4555 | Map7d2 | 78283 | 0.96883 | 1.031165066 | 1.06434 | 0.72415 | 0.939018 |
| 13195690 | ubiquitin-40S ribosomal protein S27a precursor [Mus musculus] | 0 | 59.173 | 49 | 7 | 98 | 7 | 156 | 17.9 | 9.64 | 1432 | 7 | 422257954.7 | 398211027.2 | 594413764 | 573076663.8 | 573260921.3 | 664156054.8 | 583803704 | 635705928.3 | 20.3289 | 7.00759 | Rps27a | 78294 | 0.89449 | 1.105507137 | 1.2359 | 0.07666 | 0.627416 |
| 1039772910 | PREDICTED: protein tweety homolog 3 isoform X2 [Mus musculus] | 0 | 12.618 | 6 | 3 | 15 | 3 | 526 | 58.1 | 5.74 | 267 | 3 | 4074716.564 | 2317517.404 | 3521252.963 | 1668193.292 | 3918793.478 | 3124043.371 | 4783066.406 | 2461448.843 | 37.9516 | 28.0943 | Ttyh3 | 78339 | 0.89541 | 1.104591151 | 1.23362 | 0.39834 | 0.824613 |
| 283945600 | partner of Y14 and mago isoform 2 [Mus musculus] | 0 | 24.886 | 36 | 5 | 10 | 5 | 202 | 22.7 | 9.48 | 55 | 5 | 11644700.77 | 21932987.95 | 17769315.52 | 22658034.33 | 18442345.27 | 34653959.55 | 17566521.5 | 36489962.67 | 27.3126 | 37.989 | Pym1 | 78428 | 0.81702 | 1.182977191 | 1.44791 | 0.19493 | 0.715071 |
| 148703544 | mCG116006, partial [Mus musculus] | 0 | 32.263 | 17 | 7 | 16 | 7 | 528 | 60.3 | 6.96 | 399 | 7 | 22434003.3 | 13014611.5 | 18646470.32 | 17926572.89 | 21407257.96 | 11262298.57 | 18976141.13 | 7841874.247 | 21.4952 | 42.8809 | Micu3 | 78506 | 1.09531 | 0.904690446 | 0.82597 | 0.43304 | 0.848968 |
| 568990302 | PREDICTED: rapamycin-insensitive companion of mTOR isoform X1 [Mus musculus] | 0 | 21.13 | 4 | 6 | 17 | 6 | 1732 | 194.2 | 7.44 | 143 | 6 | 28128587.76 | 20677694.08 | 26555011.73 | 5758626.84 | 46999783.9 | 12230619.54 | 26214111.19 | 21105586.25 | 50.2866 | 55.3847 | Rictor | 78757 | 0.8645 | 1.135504756 | 1.31349 | 0.50495 | 0.88031 |
| 27753958 | spermatogenesis-associated protein 2-like protein [Mus musculus] | 0 | 118.902 | 56 | 15 | 149 | 15 | 426 | 46.7 | 5.2 | 2247 | 15 | 161748358.5 | 183157700.6 | 224197861.7 | 218005468.6 | 278868667.4 | 236287869.6 | 240732487.4 | 211319507.1 | 15.0039 | 11.5062 | Spata2l | 78779 | 0.89743 | 1.102568413 | 1.22858 | 0.06841 | 0.601634 |
| 568989101 | PREDICTED: inactive ubiquitin carboxyl-terminal hydrolase 54 isoform X1 [Mus musculus] | 0 | 15.378 | 3 | 4 | 8 | 4 | 1645 | 182.9 | 7.39 | 70 | 4 | 1858001.447 | 3397365.859 | 10414292.37 | 2240725.439 | 3790498.595 | 6366573.403 | 8091641.75 | 5471116.207 | 89.5912 | 30.2469 | Usp54 | 78787 | 0.86045 | 1.139548748 | 1.32436 | 0.53314 | 0.886555 |
| 568965809 | PREDICTED: syntaxin-binding protein 5 isoform X1 [Mus musculus] | 0 | 11.096 | 4 | 3 | 15 | 3 | 1152 | 127.6 | 7.21 | 97 | 3 | 5614409.476 | 5985627.694 | 5630545.786 | 3942184.431 | 6101016.726 | 6000578.633 | 4810036.75 | 6910416.026 | 17.3207 | 14.5324 | Stxbp5 | 78808 | 0.94112 | 1.058879689 | 1.12513 | 0.33389 | 0.779342 |
| 11907574 | TSC22-related inducible leucine zipper 2 [Mus musculus] | 0 | 7.287 | 5 | 2 | 5 | 2 | 387 | 40 | 8.15 | 21 | 2 | 3584253.062 | 7111316.439 | 5336582.118 | 6276777.667 | 8231331.971 | 7645061.616 | 5924993 | 7890760.752 | 27.1383 | 13.8383 | Tsc22d4 | 78829 | 0.85802 | 1.141982023 | 1.33095 | 0.09013 | 0.662454 |
| 27369581 | calcium-binding mitochondrial carrier protein Aralar1 [Mus musculus] | 0 | 209.178 | 57 | 27 | 165 | 27 | 677 | 74.5 | 8.25 | 4013 | 27 | 376215142.8 | 303214098.3 | 255865275 | 226380356.1 | 321229394.8 | 342550524 | 256115038.2 | 260141764.3 | 22.508 | 14.7446 | Slc25a12 | 78830 | 0.99216 | 1.007841209 | 1.01581 | 0.91073 | 0.98561 |
| 148677354 | CCR4-NOT transcription complex, subunit 10, isoform CRA_b, partial [Mus musculus] | 0.001 | 4.274 | 3 | 2 | 4 | 2 | 799 | 87.4 | 7.94 | 23 | 2 | 2205789.927 | 1833374.206 | 1906185.635 | 620120.2622 | 1239689.763 | 1920175.78 | 2713295.125 | 1205446.584 | 42.6258 | 40.121 | Cnot10 | 78893 | 0.96239 | 1.037608789 | 1.07816 | 0.80548 | 0.961747 |
| 21313536 | yllysine-residue succinyltransferase component of 2-oxoglutarate dehydrogenase complex, mitochondrial [Mus | 0 | 8.453 | 6 | 3 | 10 | 3 | 454 | 49 | 8.95 | 117 | 3 | 1655137.559 | 1882008.799 | 1764201.174 | 1724625.341 | 2343375.486 | 2654925.486 | 1747318.125 | 1905172.559 | 5.41121 | 19.1354 | Dlst | 78920 | 0.89635 | 1.103645033 | 1.23126 | 0.10423 | 0.674486 |
| 41281720 | GAS2-like protein 1 isoform beta [Mus musculus] | 0 | 17.413 | 5 | 2 | 7 | 2 | 678 | 72.4 | 10.01 | 105 | 2 | 4683726.194 | 3911597.048 | 3665271.295 | 2958940.857 | 5201924.23 | 3358449.569 | 4175757 | 2374340.535 | 18.7007 | 31.8103 | Gas2l1 | 78926 | 1.0036 | 0.996404087 | 0.99283 | 0.97012 | 0.99686 |
| 1039791016 | PREDICTED: tumor suppressor candidate 3 isoform X1 [Mus musculus] | 0 | 12.192 | 10 | 3 | 5 | 3 | 348 | 39.6 | 9.91 | 53 | 3 | 651636.7082 | 4047441.587 | 2348838.264 | 3606761.718 | 1412872.031 | 5210545.835 | 4718442.375 | 14639833.06 | 71.5482 | 87.5298 | Tusc3 | 80286 | 0.58164 | 1.418355158 | 2.43852 | 0.24064 | 0.733806 |
| 116174793 | spectrin beta chain, non-erythrocytic 4 isoform sigma1 [Mus musculus] | 0 | 1176.83 | 72 | 138 | 1380 | 133 | 2561 | 288.6 | 6 | 23784 | 138 | 2130503782 | 1809192737 | 2204464042 | 1876009243 | 2536020117 | 1549810652 | 2225367396 | 1859389948 | 9.57257 | 21.0241 | Sptbn4 | 80297 | 0.99071 | 1.009290381 | 1.01876 | 0.87822 | 0.980491 |
| 1039762360 | PREDICTED: vacuolar protein sorting-associated protein 16 homolog isoform X1 [Mus musculus] | 0 | 108.872 | 29 | 17 | 60 | 17 | 911 | 102.6 | 7.69 | 909 | 17 | 57277960.07 | 36019927.12 | 55076996.29 | 34497791.26 | 57665674.79 | 46793610.11 | 56203888.38 | 33162156.66 | 26.5249 | 23.2725 | Vps16 | 80743 | 0.97092 | 1.029075427 | 1.05989 | 0.75211 | 0.951816 |
| 213972562 | leucine-rich repeat and fibronectin type III domain-containing protein 1 isoform 1 precursor [Mus musculus] | 0 | 8.635 | 4 | 2 | 5 | 2 | 766 | 81.9 | 7.59 | 20 | 2 | 2022953.475 | 2964185.332 | 1859201.529 | 2805101.167 | 3693207.283 | 1219111.068 | 2251166.25 | 3015341.561 | 22.9057 | 41.7361 | Lrfn1 | 80749 | 0.97341 | 1.026594934 | 1.05464 | 0.83298 | 0.968992 |
| 13507666 | serine beta-lactamase-like protein LACTB, mitochondrial precursor [Mus musculus] | 0 | 18.505 | 8 | 4 | 12 | 4 | 551 | 60.7 | 8.9 | 157 | 4 | 2657348.304 | 2341371.373 | 3281620.838 | 3525106.839 | 3663075.073 | 9187941.983 | 9190882.688 | 4640794.786 | 18.5251 | 44.0083 | Lactb | 80907 | 0.61346 | 1.38654106 | 2.2602 | 0.04709 | 0.591743 |
| 28972067 | mKIAA0099 protein [Mus musculus] | 0 | 17.22 | 7 | 4 | 7 | 4 | 1218 | 129.5 | 7.15 | 55 | 4 | 1624130.843 | 2696151.236 | 2273699.032 | 1670216.093 | 4084908.906 | 1985311.352 | 2817339.375 | 3571446.066 | 24.8711 | 29.3891 | Pum1 | 80912 | 0.79758 | 1.202402857 | 1.50759 | 0.09268 | 0.662454 |
| 49258190 | NCK-interacting protein with SH3 domain [Mus musculus] | 0 | 81.507 | 22 | 14 | 70 | 14 | 714 | 78.5 | 6.05 | 1263 | 14 | 73243079 | 54085770.19 | 79775918.03 | 52761347.03 | 74701252.96 | 66079396.87 | 77339843.72 | 53809302.55 | 20.9395 | 15.5951 | Nckipsd | 80987 | 0.97732 | 1.022684796 | 1.04642 | 0.73849 | 0.943908 |
| 9055242 | dnaJ homolog subfamily B member 1 isoform 1 [Mus musculus] | 0 | 20.493 | 18 | 5 | 22 | 4 | 340 | 38.1 | 8.63 | 202 | 5 | 7286219.538 | 9081695.252 | 5331982.545 | 6117192.347 | 8503597.805 | 3811668.768 | 8579444.875 | 5128965.934 | 23.4358 | 37.0654 | Dnajb1 | 81489 | 1.03331 | 0.966690439 | 0.93553 | 0.76843 | 0.952792 |
| 148673379 | ankyrin repeat domain 17, isoform CRA_c, partial [Mus musculus] | 0 | 9.839 | 1 | 3 | 9 | 3 | 2652 | 280.2 | 6.93 | 41 | 3 | 3125560.147 | 1595006.363 | 2539511.091 | 1966922.782 | 2088515.205 | 2162188.582 | 2887553 | 1460811.5 | 29.0451 | 27.1621 | Ankrd17 | 81702 | 1.03523 | 0.964774505 | 0.93195 | 0.73597 | 0.943908 |
| 15214281 | it 1; AltName: Full=Pre-mRNA-splicing factor SF3b 155 kDa subunit; Short=SF3b155; AltName: Full=Spliceosome | 0 | 11.161 | 2 | 3 | 3 | 3 | 1304 | 145.7 | 7.09 | 34 | 3 | 1118981.908 | 855428.091 | 1674682.349 | 1238916.324 | 391581.001 | 2603087 | 3184386.125 | 2170457.814 | 27.9591 | 57.701 | Sf3b1 | 81898 | 0.73851 | 1.261491813 | 1.70816 | 0.2161 | 0.716066 |
| 21361975 | voltage-dependent calcium channel gamma-7 subunit [Homo sapiens] | 0 | 36.106 | 21 | 4 | 17 | 4 | 275 | 31 | 7.12 | 661 | 4 | 31197879.77 | 27241929.71 | 30787814.8 | 22224259.21 | 33130896.65 | 31048832.69 | 29627687.25 | 24843503.89 | 14.9216 | 11.8675 | Cacng7 | 81904 | 0.96871 | 1.031286177 | 1.06459 | 0.53331 | 0.886555 |
| 20532015 | subunit; AltName: Full=Neuronal voltage-gated calcium channel gamma-8 subunit; AltName: Full=Transmembran | 0 | 104.79 | 41 | 10 | 101 | 10 | 423 | 43.4 | 9.2 | 3415 | 10 | 229859211.3 | 294115437.7 | 245640989.1 | 349503114.7 | 299577213.8 | 340638425.8 | 234442954.4 | 322138156.5 | 19.2748 | 15.4817 | Cacng8 | 81905 | 0.96646 | 1.033540946 | 1.06941 | 0.60454 | 0.905186 |
| 23822106 | RecName: Full=Ribosome-binding protein 1; AltName: Full=Ribosome receptor protein; Short=RRp; Short=mRRp | 0 | 94.396 | 15 | 17 | 73 | 17 | 1605 | 172.8 | 9.33 | 686 | 17 | 33769223.72 | 33071166.21 | 35875848.52 | 52243858.4 | 36793831.85 | 42309239.67 | 48972116.38 | 50876225.02 | 23.4413 | 14.4062 | Rrbp1 | 81910 | 0.92815 | 1.071849323 | 1.15482 | 0.32278 | 0.771047 |
| 18959268 | nuclear distribution protein nudE-like 1 [Rattus norvegicus] | 0 | 38.937 | 30 | 9 | 73 | 7 | 345 | 38.3 | 5.24 | 1305 | 9 | 35418982.09 | 29193136.61 | 30846326.5 | 21342672.19 | 27439415.85 | 12152557.99 | 36574701.5 | 14948763.11 | 20.0785 | 49.8145 | Ndel1 | 83431 | 1.12354 | 0.876461599 | 0.78009 | 0.35346 | 0.798662 |
| 148706996 | phosphodiesterase 4D interacting protein (myomegalin), isoform CRA_b [Mus musculus] | 0 | 42.403 | 5 | 11 | 30 | 11 | 2506 | 280.4 | 5.55 | 277 | 11 | 20198725.28 | 15824197.1 | 25386013.11 | 23713890.52 | 18077860.07 | 11363949.35 | 16612089.75 | 10707939.96 | 19.8845 | 26.0812 | Pde4dip | 83679 | 1.19989 | 0.800112388 | 0.66682 | 0.04514 | 0.591743 |
| 13994209 | wiskott-Aldrich syndrome protein family member 1 [Mus musculus] | 0 | 91.378 | 22 | 12 | 121 | 11 | 559 | 61.5 | 6.37 | 2153 | 12 | 227201617.2 | 246718217.2 | 239657046.4 | 241387649.6 | 218461818.8 | 218283829.7 | 198540682.6 | 210431093.7 | 3.45943 | 4.43311 | Wasf1 | 83767 | 1.06067 | 0.939330149 | 0.8856 | 0.00471 | 0.38143 |
| 167466245 | E3 ubiquitin-protein ligase NEDD4-like isoform 1 [Mus musculus] | 0 | 232.182 | 45 | 35 | 276 | 29 | 976 | 112.2 | 5.87 | 4877 | 35 | 193490631.2 | 221874111.1 | 185193054.7 | 189106135.6 | 221922302.5 | 235147377.2 | 192792905.1 | 214262888.2 | 8.43855 | 8.2081 | Nedd4l | 83814 | 0.95494 | 1.04505703 | 1.09437 | 0.17657 | 0.713031 |
| 148664825 | DnaJ (Hsp40) homolog, subfamily A, member 3, isoform CRA_a, partial [Mus musculus] | 0 | 117.515 | 34 | 13 | 76 | 13 | 486 | 53.1 | 9.28 | 2578 | 13 | 102787761.9 | 84441291.9 | 132863377.4 | 108989680.3 | 147919561.2 | 97919469.4 | 129581228 | 104237805.1 | 18.6384 | 19.3024 | Dnaja3 | 83945 | 0.94434 | 1.055655019 | 1.11787 | 0.43999 | 0.852048 |
| 568993681 | PREDICTED: RNA binding protein fox-1 homolog 2 isoform X1 [Mus musculus] | 0 | 16.293 | 7 | 4 | 14 | 4 | 488 | 52.4 | 6.64 | 211 | 4 | 13382370.98 | 15338519.39 | 14063391.1 | 18762462.63 | 13797599.96 | 12520025.2 | 12156313.25 | 13042225.39 | 15.5466 | 5.52977 | Rbfox2 | 93686 | 1.08872 | 0.91128319 | 0.83703 | 0.09123 | 0.662454 |
| 148677820 | casein kinase 1, alpha 1, isoform CRA_e [Mus musculus] | 0 | 105.284 | 52 | 14 | 98 | 14 | 334 | 38.5 | 9.57 | 1733 | 14 | 135168990.4 | 79951144.92 | 145249201.6 | 109185668.2 | 140486416.1 | 101027525 | 151575225.4 | 65735464.57 | 24.8906 | 34.1715 | Csnk1a1 | 93687 | 1.01156 | 0.988441825 | 0.97715 | 0.91619 | 0.987992 |
| 18087751 | protocadherin gamma-C5 precursor [Mus musculus] | 0 | 49.076 | 13 | 8 | 26 | 8 | 944 | 101.8 | 5.27 | 532 | 8 | 35013763.65 | 41908943.71 | 31208226.87 | 35120757.79 | 30087173.48 | 37541901.93 | 27961897.22 | 27973816.95 | 12.4335 | 14.7128 | Pcdhgc5 | 93708 | 1.07378 | 0.926215568 | 0.86257 | 0.17281 | 0.713031 |
| 6005768 | gamma-aminobutyric acid receptor-associated protein-like 2 [Homo sapiens] | 0 | 36.215 | 57 | 9 | 86 | 7 | 117 | 13.7 | 8.1 | 631 | 9 | 85548788.94 | 113118744.2 | 100656143.6 | 111445092.9 | 93318516.75 | 92644329.57 | 82000884.53 | 127005503.6 | 12.3606 | 19.7911 | Gabarapl2 | 93739 | 1.01961 | 0.980391225 | 0.96154 | 0.74616 | 0.947173 |
| 29789289 | enoyl-CoA hydratase, mitochondrial precursor [Mus musculus] | 0 | 87.305 | 58 | 13 | 117 | 13 | 290 | 31.5 | 8.48 | 1370 | 13 | 148746520.4 | 126272133.5 | 85909689.91 | 136495985 | 126220025.8 | 124589570.5 | 115827791.2 | 168792312.2 | 21.8951 | 17.7299 | Echs1 | 93747 | 0.9632 | 1.036796459 | 1.0764 | 0.61768 | 0.906154 |
| 37360286 | mKIAA1215 protein [Mus musculus] | 0 | 92.908 | 37 | 15 | 64 | 15 | 575 | 65.7 | 9.5 | 1133 | 15 | 60222079.87 | 75238402.47 | 66297781.62 | 88719619.82 | 62074209.34 | 87640607.05 | 57620534.09 | 68129023.62 | 17.0466 | 19.2207 | Vangl2 | 93840 | 1.02653 | 0.973471658 | 0.94831 | 0.69312 | 0.930425 |
| 100816764 | calmin isoform a [Mus musculus] | 0 | 186.148 | 39 | 25 | 114 | 25 | 1052 | 117.2 | 5.03 | 2076 | 25 | 163226656.5 | 140125328.4 | 190296111.8 | 141826269.8 | 145174264 | 145030385.2 | 177244004.4 | 142316691.6 | 14.7549 | 10.8811 | Clmn | 94040 | 1.02065 | 0.979354159 | 0.95954 | 0.67012 | 0.922517 |
| 15808998 | cat eye syndrome critical region protein 6 homolog [Mus musculus] | 0 | 58.054 | 26 | 9 | 42 | 9 | 572 | 58.1 | 9.01 | 774 | 9 | 111820620.1 | 98586273.16 | 106685387.2 | 65245084.76 | 111045046.8 | 121630359.7 | 103150388.9 | 78341491.94 | 21.9147 | 17.797 | Tmem121b | 94047 | 0.96004 | 1.039962004 | 1.08325 | 0.58908 | 0.901795 |
| 16418469 | striatin-3 isoform 1 [Mus musculus] | 0 | 65.955 | 19 | 12 | 80 | 10 | 796 | 87.1 | 5.29 | 999 | 12 | 28417497.31 | 40851982.68 | 31944175.87 | 31698174.64 | 29382003.98 | 63675537.03 | 38154462.25 | 61785014.04 | 16.0433 | 35.478 | Strn3 | 94186 | 0.81564 | 1.184361938 | 1.45207 | 0.14482 | 0.693437 |
| 60360534 | mKIAA4136 protein [Mus musculus] | 0 | 32.623 | 7 | 6 | 47 | 5 | 1138 | 128 | 6.28 | 914 | 6 | 17398775.91 | 10956984.24 | 14741914.81 | 10017349.78 | 19516223.65 | 14924001.17 | 16694229.13 | 12394741.93 | 25.7751 | 18.8701 | Slc4a10 | 94229 | 0.91072 | 1.089281502 | 1.19607 | 0.29598 | 0.760494 |
| 163644255 | E3 ubiquitin-protein ligase HECW1 [Mus musculus] | 0 | 175.697 | 25 | 29 | 177 | 27 | 1604 | 179.4 | 5.39 | 3175 | 29 | 65402140 | 85094567.85 | 91771893.51 | 76250407.74 | 105656184.4 | 74698215.48 | 75868879.09 | 69938519.26 | 14.3394 | 19.9661 | Hecw1 | 94253 | 0.98814 | 1.011855152 | 1.02399 | 0.85395 | 0.975398 |
| 21619320 | Melanoma antigen, family D, 1 [Mus musculus] | 0 | 43.859 | 14 | 8 | 28 | 7 | 775 | 85.6 | 7.5 | 419 | 8 | 22187671.81 | 12781420.26 | 17580599 | 10441323.54 | 16602822.18 | 6733150.925 | 14059081.44 | 7116444.033 | 33.1565 | 44.6233 | Maged1 | 94275 | 1.1719 | 0.828101544 | 0.70663 | 0.24705 | 0.733806 |
| 16716499 | sideroflexin-3 isoform 1 [Mus musculus] | 0 | 66.638 | 45 | 9 | 34 | 9 | 321 | 35.4 | 9.51 | 1049 | 9 | 231257709.6 | 225661425 | 179319449.2 | 124314610.5 | 229504380.4 | 317183994.1 | 187058229.6 | 213705768.5 | 26.1252 | 23.7856 | Sfxn3 | 94280 | 0.89057 | 1.109425392 | 1.24574 | 0.25984 | 0.745606 |
| 30519995 | sideroflexin-5 [Mus musculus] | 0 | 70.915 | 52 | 11 | 56 | 11 | 342 | 37.3 | 9.44 | 1014 | 11 | 106494646.2 | 58729242.57 | 79543159.87 | 83184211.3 | 101981448 | 152351169 | 64152690.31 | 111149860.8 | 23.8694 | 33.716 | Sfxn5 | 94282 | 0.86578 | 1.134220868 | 1.31006 | 0.26295 | 0.745606 |
| 33859811 | trifunctional enzyme subunit alpha, mitochondrial precursor [Mus musculus] | 0 | 216.571 | 52 | 24 | 133 | 24 | 763 | 82.6 | 9.14 | 3639 | 24 | 208114724.2 | 160538307.7 | 173744708.7 | 191925285.9 | 185062836.7 | 229220421.6 | 192895467.4 | 166989481.9 | 11.336 | 13.5074 | Hadha | 97212 | 0.97359 | 1.026413929 | 1.05426 | 0.57281 | 0.895889 |
| 15778440 | zinedin [Mus musculus] | 0 | 75.74 | 26 | 13 | 89 | 11 | 760 | 81.6 | 5.4 | 856 | 13 | 71648483.17 | 26351465.77 | 73530982.12 | 49455996.11 | 65543185.71 | 56197797.52 | 68741675.88 | 62702543.73 | 40.092 | 8.11251 | Strn4 | 97387 | 0.93349 | 1.066506429 | 1.14249 | 0.51449 | 0.88031 |
| 74196946 | unnamed protein product [Mus musculus] | 0 | 77.29 | 35 | 10 | 52 | 10 | 386 | 42.7 | 9.98 | 865 | 10 | 90689468.51 | 84461749.35 | 100298299.7 | 105850554.8 | 103509432.3 | 90099392.27 | 116402342.3 | 82848534.89 | 10.0442 | 15.111 | Lrrc59 | 98238 | 0.98507 | 1.014931839 | 1.03032 | 0.7546 | 0.952474 |
| 148681171 | lamin B receptor, partial [Mus musculus] | 0 | 16.469 | 7 | 3 | 11 | 3 | 627 | 71.5 | 9.41 | 176 | 3 | 8888900.804 | 10513664.5 | 3184278.852 | 7956400.303 | 5576862.381 | 6946634.838 | 6592610.188 | 9773048.399 | 41.2555 | 24.8796 | Lbr | 98386 | 1.02783 | 0.972168571 | 0.94584 | 0.82718 | 0.968871 |
| 37360088 | mKIAA0778 protein [Mus musculus] | 0 | 153.12 | 26 | 23 | 203 | 6 | 1022 | 112.2 | 5.6 | 5352 | 23 | 12547795.97 | 13527909.92 | 16377783.44 | 16550196.81 | 16465951.52 | 26966028.32 | 15007729.78 | 20771098.37 | 13.6198 | 27.0965 | Atp1a2 | 98660 | 0.8538 | 1.146201178 | 1.34247 | 0.12849 | 0.693437 |
| 26345420 | unnamed protein product [Mus musculus] | 0 | 49.503 | 23 | 7 | 53 | 5 | 421 | 46.5 | 5.59 | 412 | 7 | 15505083.42 | 4827487.466 | 10205698.51 | 13312454.11 | 15809738.24 | 12644216.61 | 13890767.88 | 8976186.961 | 42.2532 | 22.4503 | Hnrnpf | 98758 | 0.92151 | 1.078491731 | 1.17035 | 0.51906 | 0.880629 |
| 198278553 | myosin regulatory light polypeptide 9 [Mus musculus] | 0 | 84.006 | 73 | 8 | 74 | 3 | 172 | 19.8 | 4.92 | 1652 | 8 | 1786756.692 | 5182950.143 | 3782308.362 | 2779520.193 | 5582879.819 | 2203437.597 | 4180486.625 | 585146.921 | 42.8763 | 69.9508 | Myl9 | 98932 | 1.03756 | 0.962444268 | 0.92761 | 0.85845 | 0.975398 |
| 60360532 | mKIAA4128 protein [Mus musculus] | 0 | 17.402 | 5 | 5 | 17 | 5 | 1002 | 113.3 | 7.2 | 131 | 5 | 36511309.94 | 49052010.89 | 33888689.24 | 33207050.29 | 32382489.54 | 50690747.98 | 38807569.63 | 56780457.81 | 19.3807 | 24.8055 | Osbpl6 | 99031 | 0.92152 | 1.07848056 | 1.17033 | 0.36678 | 0.80688 |
| 148229140 | brefeldin A-inhibited guanine nucleotide-exchange protein 2 [Mus musculus] | 0 | 13.615 | 3 | 4 | 7 | 4 | 1792 | 202.1 | 6.55 | 0 | 4 | 1850291.27 | 1530307.948 | 1747282.581 | 1586985.86 | 1885398.588 | 2438685.948 | 2134009 | 2086867.228 | 8.73991 | 10.7032 | Arfgef2 | 99371 | 0.88007 | 1.119928813 | 1.27254 | 0.01508 | 0.461382 |
| 148675629 | embrane associated guanylate kinase, WW and PDZ domain containing 3, isoform CRA_c, partial [Mus musculu | 0 | 145.968 | 31 | 23 | 98 | 23 | 1146 | 125 | 6.46 | 1290 | 23 | 150839965.6 | 168053805 | 168409510.2 | 140423216.8 | 172834799.8 | 171924792.1 | 150178366.6 | 179430233.8 | 8.74549 | 7.54673 | Magi3 | 99470 | 0.96418 | 1.035820509 | 1.0743 | 0.25917 | 0.745606 |
| 157836767 | Chain A, The Armadillo Repeat Region From Murine Beta-Catenin | 0 | 512.202 | 81 | 38 | 304 | 10 | 470 | 50.8 | 8.28 | 8092 | 38 | 96100258.57 | 82936133.47 | 74356768.75 | 49830586.21 | 72106801.58 | 121736448.5 | 79219128 | 94938362.11 | 25.7097 | 23.9128 | Wdr47 | 99512 | 0.90349 | 1.096505706 | 1.21363 | 0.3127 | 0.76536 |
| 952977828 | fibrinogen gamma chain isoform 2 precursor [Mus musculus] | 0 | 42.198 | 28 | 8 | 34 | 8 | 443 | 50.3 | 5.62 | 307 | 8 | 37764862.71 | 38394974.35 | 46342560.33 | 40108827.04 | 48768084.44 | 27088005.03 | 70307478.5 | 38253403.33 | 9.64347 | 39.9187 | Fgg | 99571 | 0.93716 | 1.062835663 | 1.1341 | 0.58339 | 0.898781 |
| 568924767 | PREDICTED: adhesion G protein-coupled receptor L2 isoform X1 [Mus musculus] | 0 | 100.303 | 16 | 18 | 72 | 17 | 1493 | 167.3 | 6.71 | 1099 | 18 | 59021335.74 | 54443006.49 | 67539731.25 | 46212967.43 | 66030516.56 | 70336686.49 | 62295940 | 62625125.9 | 15.6716 | 5.73385 | Adgrl2 | 99633 | 0.93024 | 1.069756915 | 1.14998 | 0.12825 | 0.693437 |
| 568924855 | PREDICTED: protein transport protein Sec24B isoform X1 [Mus musculus] | 0 | 56.621 | 10 | 11 | 37 | 11 | 1293 | 140.2 | 7.37 | 633 | 11 | 16237944.59 | 9970193.899 | 13396731.12 | 13243270.28 | 17574999.07 | 6066402.882 | 17555951.38 | 12105543.3 | 19.3959 | 41.1341 | Sec24b | 99683 | 0.99572 | 1.004284054 | 1.0086 | 0.97124 | 0.99686 |
| 568921145 | PREDICTED: ankyrin repeat domain-containing protein 50 isoform X1 [Mus musculus] | 0 | 8.466 | 2 | 3 | 7 | 3 | 1427 | 155.3 | 6.55 | 44 | 3 | 4236786.33 | 4066670.09 | 3442893.951 | 1073876.347 | 2782119.951 | 1108175.138 | 1736238.75 | 2601655.346 | 45.5905 | 37.9226 | Ankrd50 | 99696 | 1.21817 | 0.781834531 | 0.64181 | 0.21502 | 0.716066 |
| 39930509 | microtubule-associated protein RP/EB family member 3 isoform 1 [Mus musculus] | 0 | 65.032 | 45 | 12 | 65 | 2 | 281 | 31.9 | 5.54 | 927 | 12 | 3981837.944 | 6851859.219 | 5844410.82 | 7622991.261 | 2741461.628 | 7017414.285 | 4266223 | 7373512.071 | 25.9115 | 41.5943 | Mapre3 | 100732 | 1.06351 | 0.936487819 | 0.88056 | 0.61356 | 0.905186 |
| 24047300 | Usp30 protein, partial [Mus musculus] | 0 | 16.595 | 11 | 3 | 13 | 3 | 557 | 61.9 | 8.73 | 172 | 3 | 2002670.528 | 2175680.606 | 5782619.949 | 6739057.865 | 7396526.907 | 3311568.11 | 5273876.25 | 5364207.369 | 58.4664 | 31.2601 | Usp30 | 100756 | 0.87788 | 1.122118602 | 1.27821 | 0.46192 | 0.863085 |
| 568932670 | PREDICTED: A-kinase anchor protein 9 isoform X2 [Mus musculus] | 0 | 468.057 | 33 | 90 | 312 | 90 | 3813 | 437.1 | 5.05 | 4352 | 90 | 630269388.6 | 859891437.3 | 753856236.6 | 874799267.1 | 330437096.5 | 165435982.7 | 284151587 | 253019551.4 | 14.5238 | 26.9413 | Akap9 | 100986 | 1.50237 | 0.497629535 | 0.33123 | 0.00023 | 0.195279 |
| 19527154 | protein BRICK1 [Mus musculus] | 0 | 47.248 | 77 | 9 | 132 | 9 | 75 | 8.8 | 5.45 | 845 | 9 | 121794340.4 | 105257869.1 | 129129190.2 | 119792819.3 | 129819687.5 | 109749452.9 | 141585476.3 | 109794530.7 | 8.40218 | 12.8101 | Brk1 | 101314 | 0.98451 | 1.015487193 | 1.03146 | 0.70172 | 0.931521 |
| 148685191 | RIKEN cDNA 6330503K22, isoform CRA_a [Mus musculus] | 0 | 15.546 | 5 | 5 | 14 | 5 | 1004 | 111.1 | 8.85 | 79 | 5 | 3951731.596 | 6376907.653 | 4659063.404 | 5510310.404 | 3205466.291 | 5253641.382 | 3931624.75 | 4741339.651 | 20.4956 | 21.0424 | Ccp110 | 101565 | 1.08945 | 0.910551864 | 0.83579 | 0.26964 | 0.745606 |
| 19527168 | PC4 and SFRS1-interacting protein isoform 1 [Mus musculus] | 0 | 29.075 | 17 | 7 | 25 | 6 | 528 | 59.7 | 9.13 | 431 | 7 | 4748605.136 | 3263751.029 | 19991625.93 | 23754800.74 | 15234328.64 | 9403256.791 | 15948434.75 | 18542773.1 | 80.7354 | 26.0952 | Psip1 | 101739 | 0.93354 | 1.066463807 | 1.14239 | 0.75197 | 0.951816 |
| 25470894 | copine-7 [Mus musculus] | 0 | 55.023 | 28 | 13 | 42 | 12 | 557 | 61.9 | 5.4 | 556 | 13 | 36015180.49 | 26725842.94 | 24371765.42 | 30056252.13 | 32899977.51 | 33191170.73 | 28262414.66 | 37479405.77 | 17.2481 | 11.4267 | Cpne7 | 102278 | 0.94111 | 1.05889081 | 1.12515 | 0.28879 | 0.752733 |
| 118582292 | RecName: Full=Acyl-CoA dehydrogenase family member 11; Short=ACAD-11 | 0 | 16.659 | 6 | 5 | 15 | 5 | 779 | 87.3 | 8.43 | 202 | 5 | 9307513.092 | 12268176.54 | 8710702.606 | 10796358.94 | 11586901.62 | 20189782.52 | 10891807.63 | 13515796.29 | 15.525 | 30.2117 | Acad11 | 102632 | 0.84474 | 1.155258523 | 1.36759 | 0.14684 | 0.693437 |
| 568958665 | PREDICTED: pleckstrin homology-like domain family B member 1 isoform X1 [Mus musculus] | 0 | 128.58 | 23 | 26 | 88 | 26 | 1555 | 170 | 8.75 | 1172 | 26 | 89912831.82 | 84572242.71 | 105186787.4 | 73628137.92 | 108875229 | 75061342.36 | 93450922.25 | 117494544.6 | 14.8617 | 18.8885 | Phldb1 | 102693 | 0.94442 | 1.055577435 | 1.1177 | 0.39706 | 0.824613 |
| 148694050 | leucine rich repeat containing 49, isoform CRA_a, partial [Mus musculus] | 0 | 32.985 | 13 | 7 | 22 | 7 | 770 | 87.5 | 8.5 | 228 | 7 | 16453585.3 | 9986050.49 | 17413776.55 | 11394666.25 | 15814952.17 | 11015104.55 | 16070049.38 | 10891970.58 | 26.5794 | 21.4359 | Lrrc49 | 102747 | 1.01335 | 0.986647103 | 0.97365 | 0.88118 | 0.980491 |
| 26338349 | unnamed protein product [Mus musculus] | 0 | 38.979 | 27 | 7 | 49 | 7 | 333 | 34.8 | 6.39 | 825 | 7 | 24532923.38 | 37135553.91 | 20364266.5 | 36779026.97 | 29483030.33 | 36782889.21 | 28367579 | 35462205.28 | 28.7816 | 12.9592 | 30019K06f | 102941 | 0.95467 | 1.045333846 | 1.09497 | 0.57551 | 0.895889 |
| 111494223 | 5'-nucleotidase domain-containing protein 3 [Mus musculus] | 0 | 15.311 | 7 | 2 | 2 | 2 | 546 | 63.1 | 8.56 | 100 | 2 | 2530524.331 | 2427149.019 | 1715820.861 | 2045740.754 | 1895899.568 | 2504043.033 | 2199284 | 2403303.224 | 17.1126 | 11.9223 | Nt5dc3 | 103466 | 0.98401 | 1.015985702 | 1.03249 | 0.76831 | 0.952792 |
| 29789351 | nuclear pore complex protein Nup107 [Mus musculus] | 0 | 60.033 | 18 | 14 | 43 | 14 | 926 | 106.7 | 5.43 | 640 | 14 | 37718240.37 | 20083233.39 | 34950470.74 | 26061907.05 | 36814943.4 | 26231244.81 | 43185808.63 | 15970558.37 | 27.3213 | 39.1863 | Nup107 | 103468 | 0.98592 | 1.014078462 | 1.02856 | 0.91045 | 0.98561 |
| 297342902 | n A, Crystal Structure Of Mouse Mitochondrial Aspartate AminotransferaseKYNURENINE AMINOTRANSFERAS | 0 | 53.137 | 34 | 10 | 27 | 10 | 401 | 44.5 | 8.69 | 245 | 10 | 44413488.46 | 24475031.21 | 35749987.36 | 37730700 | 28777702.88 | 15627868.36 | 19667500.63 | 19891326.88 | 23.2821 | 26.4325 | Xpo1 | 103573 | 1.25805 | 0.741952557 | 0.58977 | 0.02635 | 0.515682 |
| 148671945 | mCG20203, isoform CRA_c [Mus musculus] | 0 | 67.171 | 30 | 9 | 41 | 9 | 524 | 59 | 6.83 | 765 | 9 | 27235683.53 | 20724580.97 | 33377998.05 | 23722608.86 | 35224604.39 | 13999669.6 | 28862508 | 26210581.36 | 20.702 | 34.1107 | Tubg1 | 103733 | 1.00365 | 0.996353102 | 0.99273 | 0.97198 | 0.99686 |
| 26325850 | unnamed protein product [Mus musculus] | 0 | 91.75 | 36 | 16 | 100 | 16 | 608 | 68.5 | 6.46 | 1729 | 16 | 76816506.64 | 75602851.54 | 54316065.93 | 56086331.81 | 60465701.97 | 74911459.07 | 68326194.95 | 65845799.26 | 18.5082 | 8.89465 | Rpn1 | 103963 | 0.98736 | 1.012636677 | 1.0256 | 0.81233 | 0.961921 |
| 568910356 | PREDICTED: dynamin-3 isoform X1 [Mus musculus] | 0 | 105.03 | 24 | 18 | 109 | 10 | 873 | 98.2 | 8.47 | 1917 | 18 | 49438265.42 | 59820483.32 | 58094491.94 | 39108612.95 | 46121605.63 | 61316598.76 | 50915189.75 | 66704650.97 | 18.3955 | 16.7362 | Dnm3 | 103967 | 0.95691 | 1.043094633 | 1.09007 | 0.51287 | 0.88031 |
| 148704589 | reticulon 1, isoform CRA_b, partial [Mus musculus] | 0 | 61.04 | 15 | 10 | 45 | 8 | 804 | 85.8 | 4.6 | 297 | 10 | 26095175.76 | 25276249.58 | 36343949.94 | 24302613.17 | 25151477.22 | 33956860.26 | 27651864.63 | 38343359.37 | 20.0243 | 19.1644 | Rtn1 | 104001 | 0.94481 | 1.055185084 | 1.11682 | 0.45576 | 0.860877 |
| 1039758701 | PREDICTED: cadherin-22 isoform X1 [Mus musculus] | 0.001 | 6.641 | 3 | 3 | 7 | 3 | 851 | 91 | 4.97 | 47 | 3 | 4533313.711 | 5721756.964 | 3883093.743 | 2662287.797 | 6500246.589 | 3649196.367 | 7256713.875 | 2252109.45 | 30.4028 | 47.9933 | Cdh22 | 104010 | 0.92162 | 1.078384929 | 1.1701 | 0.61335 | 0.905186 |
| 1039750357 | PREDICTED: synaptojanin-1 isoform X13 [Mus musculus] | 0 | 164.213 | 30 | 33 | 159 | 33 | 1456 | 160.5 | 8.48 | 1739 | 33 | 213330946.6 | 129151453.7 | 151929071.8 | 150498712.5 | 164280656.8 | 150012373.2 | 162567086.8 | 153237684.8 | 22.4924 | 4.42683 | Synj1 | 104015 | 1.01162 | 0.988382517 | 0.97703 | 0.84767 | 0.973079 |
| 158303337 | synaptopodin isoform B [Mus musculus] | 0 | 363.445 | 68 | 29 | 456 | 29 | 690 | 74 | 9.2 | 8506 | 29 | 1168786945 | 1204775881 | 1101190266 | 1204412387 | 1005443057 | 793913685.6 | 1144859110 | 1058666681 | 4.1674 | 14.9252 | Synpo | 104027 | 1.07789 | 0.922105597 | 0.85547 | 0.07488 | 0.624526 |
| 568971038 | PREDICTED: neurexophilin-3 isoform X1 [Mus musculus] | 0 | 7.504 | 10 | 2 | 2 | 2 | 277 | 31 | 8.85 | 18 | 2 | 3391210.543 | 5682872.58 | 6345010.877 | 3315644.067 | 6057923.499 | 9449517.12 | 6131929 | 7036616.416 | 33.3062 | 22.0968 | Nxph3 | 104079 | 0.79032 | 1.20968353 | 1.53063 | 0.06674 | 0.601634 |
| 226494598 | WD repeat-containing protein 7 [Mus musculus] | 0 | 88.713 | 18 | 18 | 48 | 18 | 1489 | 163.3 | 6.9 | 983 | 18 | 29907517.61 | 17767436.84 | 23005992.85 | 23621839.2 | 30190223.27 | 17684895.87 | 33776307.56 | 16162005.71 | 21.0884 | 36.1467 | Wdr7 | 104082 | 0.98173 | 1.018273553 | 1.03723 | 0.86828 | 0.97927 |
| 148670608 | ATP citrate lyase, isoform CRA_a [Mus musculus] | 0 | 64.175 | 16 | 14 | 55 | 14 | 1102 | 120.9 | 7.34 | 975 | 14 | 28223302.3 | 34281284.78 | 31868287.74 | 27556062.79 | 30265611.51 | 32347883.99 | 30714450.75 | 33379852.38 | 10.3775 | 4.56322 | Acly | 104112 | 0.98078 | 1.019220253 | 1.03919 | 0.51772 | 0.88031 |
| 20544149 | casein kinase I isoform delta isoform 1 [Mus musculus] | 0 | 113.263 | 48 | 17 | 123 | 6 | 415 | 47.3 | 9.74 | 1563 | 17 | 50144149.8 | 31660518.94 | 34472837.16 | 50078562.53 | 50365737.25 | 39907610.93 | 55177394.63 | 46831211.45 | 23.8225 | 13.3713 | Csnk1d | 104318 | 0.92771 | 1.072289842 | 1.15585 | 0.31445 | 0.76536 |
| 568914984 | PREDICTED: diacylglycerol kinase zeta isoform X5 [Mus musculus] | 0 | 74.541 | 19 | 12 | 55 | 11 | 960 | 107.3 | 8 | 1793 | 12 | 34621008.48 | 72848799.87 | 41920378.39 | 19834803.37 | 36932837.14 | 73865595.48 | 34472404.56 | 80551843.85 | 52.8019 | 42.7582 | Dgkz | 104418 | 0.85673 | 1.143268003 | 1.33445 | 0.4226 | 0.840085 |
| 21312428 | cdc42 effector protein 1 [Mus musculus] | 0 | 13.878 | 12 | 3 | 13 | 3 | 409 | 43 | 7.21 | 246 | 3 | 4840414.052 | 5096097.942 | 8495375.327 | 7624864.76 | 7932510.545 | 9081688.194 | 6663638.25 | 9484909.585 | 27.9868 | 15.3005 | Cdc42ep1 | 104445 | 0.88001 | 1.119994168 | 1.27271 | 0.16078 | 0.702266 |
| 12847471 | unnamed protein product [Mus musculus] | 0.001 | 6.697 | 3 | 2 | 3 | 2 | 660 | 75.7 | 7.56 | 29 | 2 | 1333822.265 | 512045.7176 | 1354836.574 | 954065.1373 | 1051492.569 | 413250.0282 | 965073.5625 | 623321.5462 | 38.1702 | 39.004 | Rars | 104458 | 1.15284 | 0.847163414 | 0.73485 | 0.30911 | 0.7639 |
| 568978508 | PREDICTED: tetratricopeptide repeat protein 7B isoform X1 [Mus musculus] | 0 | 77.455 | 26 | 17 | 55 | 17 | 937 | 104.3 | 6.65 | 787 | 17 | 40029387.29 | 39004701.7 | 44115694.15 | 37947327.27 | 38971590.67 | 36931229.6 | 35482106.63 | 29015667.76 | 6.69989 | 11.2548 | Ttc7b | 104718 | 1.06865 | 0.931354318 | 0.87153 | 0.08769 | 0.655499 |
| 19527256 | ATP-dependent RNA helicase DDX1 [Mus musculus] | 0 | 191.954 | 48 | 26 | 159 | 26 | 740 | 82.4 | 7.21 | 2640 | 26 | 142910816 | 170603039.9 | 168852127.5 | 208019597.4 | 188839684.8 | 163362487.6 | 182891283.9 | 221339860.2 | 15.5251 | 12.7368 | Ddx1 | 104721 | 0.95435 | 1.045650313 | 1.09567 | 0.39472 | 0.824613 |
| 12963791 | retinol dehydrogenase 14 [Mus musculus] | 0 | 11.182 | 9 | 2 | 7 | 2 | 334 | 36.3 | 8.18 | 32 | 2 | 2789972.514 | 2268959.249 | 2492895.998 | 2740521.983 | 3638965.882 | 3161858.051 | 3228485.125 | 2769957.685 | 9.3596 | 11.121 | Rdh14 | 105014 | 0.89144 | 1.108563946 | 1.24357 | 0.02673 | 0.515682 |
| 568982882 | PREDICTED: sodium/hydrogen exchanger 3 isoform X1 [Mus musculus] | 0 | 9.226 | 3 | 2 | 9 | 2 | 837 | 93.9 | 7.34 | 65 | 2 | 3675961.347 | 1966977.063 | 3338372.98 | 2224698.164 | 3359801.089 | 1921328.331 | 3103103 | 1433777.074 | 29.7367 | 37.6824 | Slc9a3 | 105243 | 1.06602 | 0.93398027 | 0.87614 | 0.59734 | 0.904419 |
| 568986286 | PREDICTED: dedicator of cytokinesis protein 9 isoform X1 [Mus musculus] | 0 | 98.682 | 14 | 24 | 57 | 23 | 2142 | 244.6 | 7.3 | 529 | 24 | 50876198.49 | 30024107.79 | 46050446.09 | 34303216.91 | 38950379.05 | 28129829.19 | 48456608.16 | 26041127.25 | 24.2401 | 29.3398 | Dock9 | 105445 | 1.06497 | 0.935026578 | 0.8798 | 0.51602 | 0.88031 |
| 755547553 | PREDICTED: exocyst complex component 5 isoform X1 [Mus musculus] | 0 | 11.044 | 6 | 4 | 12 | 4 | 723 | 83.4 | 6.8 | 17 | 4 | 12504904.23 | 4126790.05 | 5824543.9 | 4316033.923 | 5990220.679 | 5056600.256 | 4857451.469 | 3502109.837 | 58.9913 | 21.1531 | Exoc5 | 105504 | 1.15951 | 0.840491457 | 0.72487 | 0.40143 | 0.82599 |
| 568986342 | EDICTED: serine/threonine-protein phosphatase 6 regulatory ankyrin repeat subunit A isoform X1 [Mus musculu | 0 | 13.793 | 4 | 4 | 3 | 3 | 1083 | 116 | 6.1 | 83 | 3 | 15372795.64 | 15254959.42 | 1518776.989 | 8409036.156 | 1761984.574 | 2249226.829 | 12284895.13 | 12886685.65 | 83.8391 | 15.4223 | Ankrd28 | 105522 | 1.16308 | 0.836922234 | 0.71958 | 0.55089 | 0.892023 |
| 77046155 | Phyhip protein, partial [Mus musculus] | 0 | 21.238 | 16 | 6 | 17 | 5 | 431 | 48.4 | 8.48 | 85 | 6 | 10231277.04 | 9747817.723 | 7925150.197 | 9787430.971 | 8434925.996 | 8296503.511 | 10067090 | 7129637.172 | 10.8489 | 14.2422 | Phyhip | 105653 | 1.05255 | 0.947451438 | 0.90015 | 0.27933 | 0.745606 |
| 755547588 | PREDICTED: E3 ubiquitin-protein ligase MYCBP2 isoform X1 [Mus musculus] | 0 | 50.593 | 3 | 11 | 17 | 11 | 4797 | 526 | 7.15 | 187 | 11 | 5637172.99 | 5063192.237 | 6447137.888 | 9715682.178 | 7643847.451 | 2086903.164 | 13511892.97 | 5417917.25 | 30.956 | 67.1032 | Mycbp2 | 105689 | 0.96763 | 1.032371295 | 1.06691 | 0.86942 | 0.979509 |
| 28972079 | mKIAA0147 protein [Mus musculus] | 0 | 90.833 | 15 | 15 | 45 | 15 | 1694 | 182.8 | 5.29 | 709 | 15 | 32129661.15 | 24489723.54 | 32294446.01 | 28508381.33 | 29937465.74 | 28208187.54 | 29055735.5 | 21291022.89 | 12.5513 | 25.981 | Scrib | 105782 | 0.99557 | 1.00443324 | 1.0089 | 0.95312 | 0.994906 |
| 84794611 | prickle-like protein 1 [Mus musculus] | 0 | 217.235 | 44 | 22 | 118 | 22 | 832 | 94.1 | 6.28 | 2356 | 22 | 150944695 | 145727045 | 134624124.3 | 135810833.9 | 137785789.6 | 159568287.2 | 150313271.1 | 126089750.3 | 5.5599 | 10.1856 | Prickle1 | 106042 | 0.99417 | 1.005829267 | 1.01173 | 0.84786 | 0.973079 |
| 148678962 | mCG11048, isoform CRA_b, partial [Mus musculus] | 0 | 26.382 | 17 | 5 | 37 | 5 | 349 | 40.1 | 9.29 | 375 | 5 | 23850689.85 | 19990107.52 | 27345736.25 | 27217008.94 | 31026490.89 | 25358603.51 | 37033738.13 | 26104849.84 | 14.1203 | 18.0413 | Tecr | 106259 | 0.90309 | 1.096913728 | 1.21463 | 0.15073 | 0.693437 |
| 19527352 | nurim [Mus musculus] | 0 | 7.821 | 11 | 2 | 10 | 2 | 262 | 29.4 | 8.21 | 61 | 2 | 6772566.136 | 6792824.339 | 5727276.192 | 6003729.35 | 7130676.073 | 13897314.26 | 10268086 | 10914878.55 | 8.56245 | 26.3005 | Nrm | 106582 | 0.74944 | 1.250558771 | 1.66865 | 0.02429 | 0.515682 |
| 254939654 | putative ATP-dependent RNA helicase DHX57 isoform 1 [Mus musculus] | 0 | 31.394 | 7 | 7 | 9 | 6 | 1388 | 155.7 | 7.87 | 177 | 7 | 6241857.429 | 6287107.608 | 16692600.84 | 2766405.705 | 16114501 | 7259737.18 | 14509948.5 | 13879374.7 | 75.3667 | 30.1568 | Dhx57 | 106794 | 0.76388 | 1.236122123 | 1.61822 | 0.21766 | 0.716066 |
| 28972213 | mKIAA0416 protein [Mus musculus] | 0 | 67.448 | 30 | 15 | 85 | 14 | 528 | 60.3 | 8.28 | 1210 | 15 | 120943578.9 | 117651282.6 | 125882877.6 | 96096753.5 | 125065241.2 | 131405076.1 | 129158304.8 | 108715521.8 | 11.4125 | 8.29803 | Lrrtm2 | 107065 | 0.96464 | 1.035363904 | 1.07332 | 0.35017 | 0.797109 |
| 19527388 | ubiquitin thioesterase OTUB1 [Mus musculus] | 0 | 23.707 | 22 | 5 | 19 | 5 | 271 | 31.3 | 4.94 | 601 | 5 | 13498860.11 | 7285547.406 | 16916679.71 | 6720684.363 | 12895689.27 | 4967908.855 | 8064815 | 4116468.857 | 44.5148 | 52.8593 | Otub1 | 107260 | 1.19306 | 0.8069352 | 0.67635 | 0.30017 | 0.761857 |
| 148709219 | unc-5 homolog A (C. elegans), isoform CRA_b [Mus musculus] | 0 | 42.994 | 13 | 10 | 32 | 10 | 920 | 100.9 | 7.36 | 187 | 10 | 23957607.94 | 19134965.79 | 19329043.71 | 19739976.99 | 16278425.48 | 19235762.18 | 16363307.88 | 9740367.453 | 11.1588 | 26.0863 | Unc5a | 107448 | 1.14288 | 0.857116366 | 0.74996 | 0.06816 | 0.601634 |
| 568911608 | PREDICTED: bifunctional glutamate/proline--tRNA ligase isoform X1 [Mus musculus] | 0 | 12.077 | 3 | 4 | 5 | 4 | 1519 | 170.8 | 7.84 | 41 | 4 | 1669741.433 | 2348883.682 | 1669232.731 | 3094669.321 | 1044614.757 | 7221775.15 | 2841049.25 | 6710178.98 | 30.9506 | 67.3197 | Eprs | 107508 | 0.66034 | 1.339663219 | 2.02876 | 0.19215 | 0.715071 |
| 62548848 | translocon-associated protein alpha, muscle specific isoform [Mus musculus] | 0 | 9.749 | 8 | 2 | 5 | 2 | 298 | 33.7 | 4.65 | 126 | 2 | 2449977.284 | 440787.676 | 2599049.198 | 1888260.406 | 2490444.812 | 1477662.645 | 2534673.5 | 2017032.163 | 53.3794 | 23.1911 | Ssr1 | 107513 | 0.92818 | 1.071816997 | 1.15475 | 0.62284 | 0.908209 |
| 404501474 | coronin-2A isoform 1 [Mus musculus] | 0 | 40.028 | 18 | 9 | 32 | 9 | 543 | 61.7 | 8.15 | 431 | 9 | 36390429.53 | 42076227.12 | 50273770.75 | 22676997.18 | 29397287.39 | 34601220.99 | 46402477.13 | 31585899.25 | 30.6772 | 21.3458 | Coro2a | 107684 | 1.03214 | 0.96785821 | 0.93772 | 0.7454 | 0.947173 |
| 568912732 | PREDICTED: rap guanine nucleotide exchange factor 1 isoform X1 [Mus musculus] | 0 | 22.292 | 7 | 6 | 18 | 6 | 1257 | 140 | 6.07 | 82 | 6 | 12617073.67 | 3466129.712 | 15993169.81 | 8851965.324 | 16337806.74 | 9104890.89 | 9897699.125 | 8497712.58 | 52.4973 | 33.1314 | Rapgef1 | 107746 | 0.96567 | 1.034326919 | 1.07109 | 0.8299 | 0.968871 |
| 23271467 | Aldh1l1 protein [Mus musculus] | 0 | 19.15 | 6 | 5 | 13 | 5 | 902 | 98.7 | 5.99 | 162 | 5 | 7039991.022 | 7121309.829 | 3916915.297 | 7890516.828 | 10656557.29 | 7283001.328 | 8220630.25 | 4380663.711 | 27.0956 | 33.9749 | Aldh1l1 | 107747 | 0.91909 | 1.080908744 | 1.17606 | 0.49325 | 0.871733 |
| 568991001 | PREDICTED: brain-specific angiogenesis inhibitor 1 isoform X1 [Mus musculus] | 0 | 278.594 | 30 | 39 | 296 | 39 | 1620 | 177 | 7.43 | 4677 | 39 | 357356021.9 | 367217027.8 | 372180125.8 | 352896371.8 | 367832739.8 | 465424754.3 | 379894283.9 | 431705289.5 | 2.44014 | 11.0744 | Adgrb1 | 107831 | 0.93692 | 1.063081953 | 1.13466 | 0.08004 | 0.642075 |
| 148691633 | fibroblast growth factor receptor substrate 3, partial [Mus musculus] | 0 | 35.265 | 19 | 8 | 30 | 8 | 500 | 54.7 | 6.73 | 302 | 8 | 10782703.03 | 11834720.73 | 12650317.35 | 11025608.84 | 10507671.22 | 16673042.3 | 13759491.31 | 12937286.67 | 7.3204 | 18.8808 | Frs3 | 107971 | 0.92429 | 1.075712068 | 1.16383 | 0.2069 | 0.715071 |
| 54291704 | phosphofurin acidic cluster sorting protein 1 [Mus musculus] | 0 | 89.118 | 25 | 18 | 82 | 17 | 961 | 104.8 | 7.74 | 1430 | 18 | 105724795.1 | 70286358.59 | 112749128.3 | 73506890.11 | 96783138.96 | 99438521.52 | 122465490 | 99601106.05 | 24.0574 | 11.4741 | Pacs1 | 107975 | 0.92823 | 1.071770796 | 1.15464 | 0.30313 | 0.761857 |
| 755560365 | PREDICTED: CUGBP Elav-like family member 4 isoform X1 [Mus musculus] | 0 | 10.317 | 5 | 2 | 6 | 2 | 564 | 60.1 | 8.54 | 107 | 2 | 5549950.461 | 1913670.12 | 2388448.882 | 3163548.445 | 4985495.927 | 2447744.68 | 3068598.25 | 2172074.552 | 49.6344 | 40.0206 | Celf4 | 108013 | 1.0133 | 0.986698687 | 0.97375 | 0.93639 | 0.994906 |
| 86991442 | protein lin-7 homolog A isoform 1 [Mus musculus] | 0 | 38.041 | 38 | 8 | 78 | 4 | 233 | 26 | 8.72 | 836 | 8 | 23948010.95 | 22277393.77 | 22608666.24 | 23080333.68 | 22571267.73 | 25251199.22 | 24341370.75 | 23097939.92 | 3.15688 | 5.08564 | Lin7a | 108030 | 0.98212 | 1.017883541 | 1.03642 | 0.28063 | 0.747972 |
| 1010228567 | calcium/calmodulin-dependent protein kinase type II subunit delta isoform 9 [Homo sapiens] | 0 | 328.528 | 71 | 30 | 1761 | 3 | 533 | 60 | 7.27 | 38038 | 30 | 64813492.94 | 63534476.36 | 77748671.61 | 60200612.47 | 76421094.3 | 66120783.75 | 52050017.38 | 48828405.91 | 11.5648 | 21.0481 | Camk2d | 108058 | 1.04488 | 0.955118375 | 0.91409 | 0.47306 | 0.863085 |
| 148689213 | mCG19511 [Mus musculus] | 0 | 38.522 | 13 | 10 | 48 | 7 | 872 | 95.8 | 8.06 | 309 | 10 | 18614903.93 | 11392280.22 | 23807961.58 | 21339186.67 | 28001132.61 | 14792544.6 | 25209460.44 | 25060017.78 | 28.5687 | 24.9659 | Grm2 | 108068 | 0.89354 | 1.106462314 | 1.23829 | 0.30076 | 0.761857 |
| 32469489 | metabotropic glutamate receptor 3 precursor [Mus musculus] | 0 | 31.374 | 10 | 7 | 32 | 5 | 879 | 99.1 | 7.75 | 260 | 7 | 1751151.044 | 7231884.673 | 5718656.617 | 8077083.469 | 3874062.845 | 8182693.718 | 5401700.188 | 8968190.191 | 49.2426 | 36.01 | Grm3 | 108069 | 0.92586 | 1.074135551 | 1.16014 | 0.63755 | 0.914499 |
| 219801747 | metabotropic glutamate receptor 5 isoform b precursor [Mus musculus] | 0 | 87.542 | 18 | 19 | 94 | 19 | 1203 | 131.8 | 7.84 | 975 | 19 | 57931076.48 | 94374379.23 | 74814492.56 | 72569344.04 | 69654852.87 | 108190186.3 | 59479022.53 | 85363988.22 | 19.9856 | 26.2935 | Grm5 | 108071 | 0.96305 | 1.036953077 | 1.07674 | 0.67336 | 0.923206 |
| 60460877 | metabotropic glutamate receptor 7 precursor [Mus musculus] | 0 | 64.388 | 20 | 13 | 38 | 12 | 915 | 102.2 | 7.88 | 345 | 13 | 88201011.43 | 65534571.32 | 52909520.47 | 51436170.62 | 68482550.9 | 65753298.84 | 65012007.44 | 49109564.76 | 26.3606 | 14.1424 | Grm7 | 108073 | 1.0192 | 0.980799548 | 0.96232 | 0.80797 | 0.961747 |
| 33563294 | phosphatidylinositol 5-phosphate 4-kinase type-2 beta [Mus musculus] | 0 | 55.587 | 28 | 10 | 41 | 6 | 416 | 47.3 | 7.33 | 789 | 10 | 26546301.09 | 24963273.69 | 26975764.22 | 26446059.52 | 23821369.17 | 23837098.39 | 25816988 | 20704417.54 | 3.34319 | 8.97372 | Pip4k2b | 108083 | 1.054 | 0.946002427 | 0.89754 | 0.05707 | 0.599194 |
| 262231776 | brain-specific angiogenesis inhibitor 1-associated protein 2 isoform a [Mus musculus] | 0 | 395.766 | 75 | 39 | 703 | 3 | 535 | 59.2 | 9.04 | 11814 | 39 | 261840460.4 | 93199075.33 | 166187065.2 | 111883178.3 | 160182524.2 | 69706409.64 | 171717124.5 | 48347631.95 | 47.8052 | 55.5811 | Baiap2 | 108100 | 1.16911 | 0.830890715 | 0.7107 | 0.38682 | 0.81965 |
| 755560383 | PREDICTED: gamma-soluble NSF attachment protein isoform X1 [Mus musculus] | 0 | 24.49 | 14 | 4 | 22 | 4 | 362 | 40.3 | 5.55 | 398 | 4 | 9018487.549 | 8649798.699 | 11322371.71 | 9885623.948 | 13526044.42 | 8205181.431 | 12070522.88 | 8352222.7 | 12.2211 | 25.4008 | Napg | 108123 | 0.95955 | 1.040450195 | 1.08431 | 0.596 | 0.903104 |
| 46909607 | UDP-N-acetylglucosamine--peptide N-acetylglucosaminyltransferase 110 kDa subunit isoform 1 [Mus musculus] | 0 | 208.273 | 41 | 33 | 186 | 2 | 1046 | 116.9 | 6.7 | 3269 | 33 | 4814844.013 | 3230153.908 | 5228769.01 | 3271233.914 | 3976742.749 | 5991689.868 | 6149090.688 | 5343872.18 | 25.0603 | 18.4359 | Ogt | 108155 | 0.87064 | 1.129357032 | 1.29715 | 0.13704 | 0.693437 |
| 148704520 | rogenase (NADP+ dependent), methenyltetrahydrofolate cyclohydrolase, formyltetrahydrofolate synthase, isofor | 0.001 | 6.596 | 2 | 2 | 4 | 2 | 949 | 102.6 | 7.4 | 0 | 2 | 1585999.108 | 5208486.901 | 2184496.662 | 4075239.867 | 2489853.401 | 6690844.189 | 3603252.25 | 6773160.699 | 51.3344 | 41.3808 | Mthfd1 | 108156 | 0.79255 | 1.20745003 | 1.5235 | 0.24525 | 0.733806 |
| 14318722 | ATPase, H+ transporting, lysosomal V1 subunit H [Mus musculus] | 0 | 55.656 | 29 | 10 | 29 | 10 | 483 | 55.8 | 6.61 | 250 | 10 | 21696099.53 | 21644049.34 | 26386219.18 | 21593914.97 | 27211804.47 | 23519755.96 | 23534551 | 22830709.81 | 10.386 | 8.18044 | Atp6v1h | 108664 | 0.96934 | 1.030658247 | 1.06326 | 0.38641 | 0.81965 |
| 568969741 | PREDICTED: girdin isoform X1 [Mus musculus] | 0 | 99.734 | 13 | 22 | 89 | 22 | 2016 | 231.5 | 5.94 | 1371 | 22 | 45560104.89 | 46615472.96 | 49581690.22 | 48878491.7 | 49398435.85 | 43301962.53 | 44231218.5 | 51868481.85 | 3.95862 | 8.70418 | Ccdc88a | 108686 | 1.00484 | 0.995162131 | 0.99037 | 0.84583 | 0.973079 |
| 30425078 | retinol dehydrogenase 13 isoform 1 precursor [Mus musculus] | 0 | 20.899 | 17 | 4 | 14 | 4 | 334 | 36.4 | 8.85 | 155 | 4 | 6369777.25 | 4245040.478 | 6198354.551 | 9741654.746 | 10363240.67 | 5591828.708 | 5402075.438 | 3423526.258 | 34.3759 | 47.5633 | Rdh13 | 108841 | 1.03456 | 0.965439978 | 0.93319 | 0.81979 | 0.964114 |

| | | | | | | | | | | | | | | | | | | | | | | | | | | | | | |
|---|---|---|---|---|---|---|---|---|---|---|---|---|---|---|---|---|---|---|---|---|---|---|---|---|---|---|---|---|---|
| 239985513 | ATPase family AAA domain-containing protein 3 [Mus musculus] | 0 | 104.074 | 42 | 19 | 109 | 19 | 591 | 66.7 | 9.29 | 1759 | 19 | 152964185.7 | 148573303.6 | 154027105.7 | 146802951.9 | 174788745.8 | 201746840.7 | 163670638 | 170693545.1 | 2.29582 | 9.37387 | Atad3a | 108888 | 0.91736 | 1.082642903 | 1.18018 | 0.01885 | 0.492977 |
| 26350029 | unnamed protein product [Mus musculus] | 0 | 11.57 | 5 | 2 | 8 | 2 | 592 | 65.1 | 8.63 | 32 | 2 | 1604874.05 | 2032501.874 | 1170380.177 | 2549979.48 | 1342855.164 | 3483065.651 | 2073057.75 | 2462546.862 | 32.0828 | 38.1149 | Slc30a9 | 109108 | 0.88015 | 1.119849187 | 1.27234 | 0.38506 | 0.817874 |
| 18314535 | Methylmalonic aciduria (cobalamin deficiency) type A [Mus musculus] | 0 | 55.419 | 30 | 10 | 52 | 10 | 415 | 45.9 | 9.32 | 633 | 10 | 59718461.33 | 55254633.72 | 60278234.29 | 64737857.83 | 56068382.08 | 60792210.19 | 55618342.88 | 74492702.24 | 6.46404 | 14.2787 | Mmaa | 109136 | 0.98566 | 1.014338834 | 1.02909 | 0.72942 | 0.943709 |
| 1039727293 | PREDICTED: saccharopine dehydrogenase-like oxidoreductase isoform X1 [Mus musculus] | 0 | 16.462 | 10 | 5 | 19 | 5 | 517 | 56.3 | 9.11 | 81 | 5 | 50004857.19 | 56955241.77 | 44858219.05 | 55398485.34 | 58563017.01 | 63440794.22 | 50741820.5 | 52660115.42 | 10.6278 | 10.258 | Sccpdh | 109232 | 0.95796 | 1.042043448 | 1.08778 | 0.29803 | 0.761852 |
| 568905212 | EDICTED: phosphatidylinositol 3,4,5-trisphosphate-dependent Rac exchanger 2 protein isoform X1 [Mus muscul | 0 | 10.464 | 3 | 4 | 11 | 4 | 1617 | 183.7 | 7.44 | 19 | 4 | 3934796.231 | 9577998.848 | 9599963.323 | 5014879.513 | 3783224.768 | 7538242.815 | 10050065.78 | 6631836.427 | 42.4552 | 36.9509 | Prex2 | 109294 | 1.00221 | 0.997786106 | 0.99558 | 0.98796 | 0.998251 |
| 30425144 | protein FAM163B [Mus musculus] | 0 | 49.112 | 57 | 4 | 22 | 4 | 167 | 18.3 | 5.15 | 1002 | 4 | 65378425.23 | 54143312.27 | 63048644.68 | 53953869.54 | 66459680.08 | 58325729.88 | 63131515.75 | 43375698.02 | 10.0553 | 17.6295 | Fam163b | 109349 | 1.01118 | 0.988816932 | 0.97788 | 0.83193 | 0.968992 |
| 568978634 | PREDICTED: alpha-actinin-1 isoform X4 [Mus musculus] | 0 | 755.957 | 74 | 65 | 874 | 41 | 914 | 105.5 | 5.39 | 16384 | 65 | 1888679394 | 1664025502 | 2085852087 | 1732609295 | 1540755136 | 1068411048 | 1938874688 | 1691722244 | 10.1657 | 23.4933 | Actn1 | 109711 | 1.08312 | 0.916875393 | 0.84651 | 0.21843 | 0.716066 |
| 37574113 | active breakpoint cluster region-related protein isoform 1 [Mus musculus] | 0 | 56.926 | 16 | 12 | 37 | 10 | 871 | 98.9 | 6.68 | 330 | 12 | 18618637.99 | 28097734.57 | 24986680.7 | 15995853.23 | 26691329.39 | 32386331.29 | 24109451.25 | 28712868.56 | 25.4717 | 12.4847 | Abr | 109934 | 0.87875 | 1.121248542 | 1.27596 | 0.11584 | 0.678142 |
| 568949334 | PREDICTED: synaptotagmin-17 isoform X1 [Mus musculus] | 0 | 12.631 | 6 | 3 | 5 | 3 | 582 | 65.3 | 8.51 | 29 | 3 | 6098325.869 | 5556267.251 | 4852553.197 | 5856471.264 | 4447905.865 | 5782147.204 | 4535047.688 | 5813721.605 | 9.65614 | 14.6798 | Syt17 | 110058 | 1.04156 | 0.958437499 | 0.92019 | 0.37354 | 0.80688 |
| 24418919 | glycogen phosphorylase, brain form [Mus musculus] | 0 | 23.513 | 8 | 6 | 19 | 6 | 843 | 96.7 | 6.73 | 154 | 6 | 8720128.904 | 10414652.53 | 8794662.527 | 8819024.085 | 11440756.9 | 10930656.16 | 11271188.25 | 9582311.433 | 8.91942 | 7.80165 | Pygb | 110078 | 0.91902 | 1.080982505 | 1.17624 | 0.0331 | 0.520743 |
| 33859809 | fibrinogen beta chain preproprotein [Mus musculus] | 0 | 80.118 | 41 | 14 | 61 | 14 | 481 | 54.7 | 7.08 | 1200 | 14 | 32897905.9 | 31333386.03 | 36157408.35 | 36038697.61 | 37682537.7 | 34381144.03 | 66562550.28 | 47296169.25 | 6.99801 | 31.1216 | Fgb | 110135 | 0.84646 | 1.15354439 | 1.36279 | 0.14238 | 0.693437 |
| 568940298 | PREDICTED: RAF proto-oncogene serine/threonine-protein kinase isoform X1 [Mus musculus] | 0 | 26.343 | 12 | 6 | 20 | 4 | 668 | 75.1 | 9.32 | 151 | 6 | 6464402.992 | 6777081.586 | 7168744.743 | 5742043.706 | 6967857.083 | 5053691.735 | 5424493.25 | 5266035.444 | 9.23615 | 15.3787 | Raf1 | 110157 | 1.0704 | 0.929597028 | 0.86846 | 0.15632 | 0.69857 |
| 1039750424 | PREDICTED: diacylglycerol kinase gamma isoform X1 [Mus musculus] | 0 | 11.826 | 6 | 3 | 6 | 3 | 834 | 93.9 | 6.62 | 28 | 3 | 3456969.319 | 1983853.102 | 4162808.791 | 3254849.244 | 3132104.423 | 3148809.849 | 3250554.375 | 2372808.706 | 28.2506 | 13.6276 | Dgkg | 110197 | 1.03853 | 0.961466202 | 0.92579 | 0.64843 | 0.915864 |
| 124487229 | breakpoint cluster region protein [Mus musculus] | 0 | 83.07 | 19 | 16 | 54 | 14 | 1270 | 143 | 6.92 | 1007 | 16 | 35489859.47 | 40734490.02 | 40720767.18 | 43224702.68 | 37681813.3 | 47327787.12 | 41628912.69 | 34080574.41 | 8.12974 | 14.1253 | Bcr | 110279 | 0.99829 | 1.001711708 | 1.00343 | 0.96788 | 0.99686 |
| 13385090 | cytochrome c oxidase subunit 6B1 [Mus musculus] | 0 | 8.742 | 26 | 2 | 14 | 2 | 86 | 10.1 | 8.72 | 200 | 2 | 12190641.4 | 7327573.283 | 11340225.28 | 8250456.521 | 14124989.01 | 9124122.161 | 11436845.5 | 8518261.453 | 24.0586 | 23.5897 | Cox6b1 | 110323 | 0.95025 | 1.049752967 | 1.10472 | 0.57641 | 0.895889 |
| 568930433 | PREDICTED: rap1 GTPase-activating protein 1 isoform X1 [Mus musculus] | 0 | 37.104 | 15 | 9 | 34 | 8 | 755 | 83.8 | 7.28 | 452 | 9 | 26329288.55 | 22673145.69 | 24123556.15 | 16759489.62 | 19744637.7 | 19985052.94 | 15790472.06 | 22545837.36 | 18.2182 | 14.2893 | Rap1gap | 110351 | 1.07037 | 0.929625628 | 0.86851 | 0.27788 | 0.745606 |
| 595763175 | beta-adrenergic receptor kinase 1 isoform 1 [Mus musculus] | 0 | 80.2 | 29 | 15 | 60 | 10 | 689 | 79.6 | 7.28 | 960 | 15 | 46510990.07 | 29497913.44 | 51965175.2 | 31115288.26 | 48493479.03 | 32024118.22 | 51704696.13 | 32253383.72 | 28.0952 | 25.4198 | Grk2 | 110355 | 0.98335 | 1.016646761 | 1.03386 | 0.86607 | 0.978314 |
| 12805321 | Sec13 protein [Mus musculus] | 0 | 30.066 | 19 | 3 | 14 | 3 | 322 | 35.5 | 5.38 | 573 | 3 | 6282087.71 | 10660445.41 | 10628242.72 | 9836337.264 | 10520192.79 | 11774804.69 | 12693912.75 | 11760995.23 | 22.2594 | 7.63451 | Sec13 | 110379 | 0.88898 | 1.1110162 | 1.24976 | 0.08475 | 0.648205 |
| 125347788 | protein Shroom2 isoform 1 [Mus musculus] | 0 | 44.407 | 9 | 9 | 29 | 9 | 1487 | 165.2 | 6.7 | 623 | 9 | 6517827.952 | 11854011.33 | 6972600.805 | 4996726.972 | 4919413.166 | 2987632.563 | 6124055.438 | 10520973.2 | 39.1361 | 52.0495 | Shroom2 | 110380 | 1.10546 | 0.894539063 | 0.8092 | 0.53153 | 0.886555 |
| 21450129 | acetyl-CoA acetyltransferase, mitochondrial precursor [Mus musculus] | 0 | 86.873 | 43 | 13 | 54 | 13 | 424 | 44.8 | 8.51 | 901 | 13 | 147783280.7 | 119834682.9 | 62317906.8 | 132202529.9 | 131167008 | 46741617.97 | 142021113 | 95138661.8 | 32.2632 | 41.4146 | Acat1 | 110446 | 1.05366 | 0.946341046 | 0.89815 | 0.69347 | 0.930425 |
| 148708002 | high density lipoprotein (HDL) binding protein, isoform CRA_a [Mus musculus] | 0 | 159.82 | 30 | 26 | 79 | 26 | 1285 | 143.4 | 6.8 | 2089 | 26 | 50983074.04 | 61591015.45 | 56200541.57 | 53807434.54 | 58738005.72 | 44382849.7 | 69554218.09 | 48070896.92 | 8.0885 | 20.5647 | Hdlbp | 110611 | 1.00414 | 0.995858382 | 0.99175 | 0.9425 | 0.994906 |
| 26383576 | unnamed protein product [Mus musculus] | 0 | 10.454 | 13 | 3 | 7 | 3 | 248 | 27.8 | 10.17 | 45 | 3 | 1652759.969 | 2182897.526 | 2596087.975 | 2874950.99 | 2012160.878 | 3610658.229 | 3277886.125 | 5563066.578 | 22.8511 | 40.6377 | Srsf1 | 110809 | 0.78305 | 1.21695304 | 1.55413 | 0.15002 | 0.693437 |
| 150010574 | sodium channel protein type 2 subunit alpha [Mus musculus] | 0 | 49.827 | 8 | 14 | 43 | 10 | 2006 | 227.8 | 5.67 | 309 | 14 | 12764036.14 | 11901781.4 | 18021459.47 | 10907076.94 | 15752263.67 | 14720597.62 | 16647532.5 | 18971176.28 | 23.6886 | 10.9676 | Scn2a | 110876 | 0.89558 | 1.104416758 | 1.23318 | 0.13818 | 0.693437 |
| 148710162 | solute carrier family 8 (sodium/calcium exchanger), member 2, isoform CRA_a, partial [Mus musculus] | 0 | 119.48 | 22 | 13 | 59 | 13 | 1014 | 109.9 | 5.27 | 1258 | 13 | 133468355.8 | 130260280.6 | 133762421.2 | 124617322.2 | 150995068.2 | 121925684 | 142816187 | 121624833.5 | 3.25385 | 11.0829 | Slc8a2 | 110891 | 0.9856 | 1.01439719 | 1.02921 | 0.6398 | 0.915433 |
| 13592053 | 60S ribosomal protein L10 [Rattus norvegicus] | 0 | 89.038 | 43 | 10 | 149 | 10 | 214 | 24.6 | 10.08 | 1692 | 10 | 195304189.3 | 151533736.6 | 164480019.3 | 191582268.3 | 186337388.3 | 147265454.9 | 225556872.4 | 123108168.9 | 12.056 | 26.3672 | Rpl10 | 110954 | 1.0149 | 0.985104819 | 0.97065 | 0.84247 | 0.971507 |
| 120300971 | ELKS/Rab6-interacting/CAST family member 1 isoform 1 [Mus musculus] | 0 | 291.965 | 41 | 46 | 388 | 4 | 1120 | 128.3 | 5.91 | 6220 | 46 | 2412237.326 | 8518655.166 | 14361547.5 | 14114850.05 | 12294508.17 | 21806175.76 | 12422923 | 22644182.64 | 57.3103 | 33.0033 | Erc1 | 111173 | 0.7259 | 1.274100647 | 1.7552 | 0.11323 | 0.678142 |
| 18700004 | 3-ketoacyl-CoA thiolase A, peroxisomal precursor [Mus musculus] | 0 | 20.943 | 19 | 4 | 12 | 4 | 424 | 43.9 | 8.44 | 120 | 4 | 5654981.627 | 5663380.227 | 12667522.05 | 6446205.431 | 12744995.86 | 6118758.524 | 13176951.88 | 10253943.9 | 44.6022 | 30.6166 | Acaa1a | 113868 | 0.83689 | 1.163111407 | 1.3898 | 0.25288 | 0.73472 |
| 12852446 | unnamed protein product [Mus musculus] | 0 | 21.343 | 13 | 5 | 13 | 5 | 551 | 62.6 | 5.72 | 87 | 5 | 7902047.163 | 7544714.625 | 14837247.83 | 12699389.29 | 12976707.92 | 9740489.85 | 11849921 | 14749367.56 | 33.5058 | 17.0184 | Palmd | 114301 | 0.93139 | 1.06861425 | 1.14734 | 0.47612 | 0.863085 |
| 148672373 | mCG49427 [Mus musculus] | 0 | 13.359 | 20 | 3 | 74 | 2 | 156 | 17.8 | 10.7 | 651 | 3 | 61619295.56 | 87692006.98 | 54670821.32 | 94315014.81 | 56157041.27 | 89323710.96 | 72655800 | 78061036.89 | 25.9759 | 18.6388 | Rpl31 | 114641 | 1.00353 | 0.996468346 | 0.99296 | 0.96623 | 0.99686 |
| 111955212 | phospholipase DDHD1 isoform 3 [Mus musculus] | 0 | 13.524 | 7 | 5 | 6 | 5 | 918 | 102.1 | 5.9 | 45 | 5 | 2903766.386 | 4658836.725 | 4372760.176 | 840162.978 | 6693534.556 | 3750002.188 | 4581036.25 | 5931589.607 | 54.7102 | 25.2392 | Ddhd1 | 114874 | 0.75748 | 1.242520805 | 1.64034 | 0.1112 | 0.678142 |
| 18699726 | vacuolar protein sorting-associated protein 4A [Mus musculus] | 0 | 7.666 | 4 | 2 | 6 | 2 | 437 | 48.9 | 7.8 | 23 | 2 | 371723.6957 | 5705329.224 | 4219749.673 | 974041.1911 | 2865389.147 | 5506234.358 | 3419585.75 | 4960252.99 | 90.9122 | 29.8047 | Vps4a | 116733 | 0.80442 | 1.195580566 | 1.48627 | 0.37336 | 0.80688 |
| 68533242 | ubiquitin-protein ligase E3B [Mus musculus] | 0 | 80.341 | 17 | 16 | 74 | 16 | 1070 | 122.7 | 8.32 | 1116 | 16 | 34045478.52 | 28630701.26 | 43335257.87 | 26648980.89 | 45306479.66 | 29860566.86 | 45768212.81 | 31734770.48 | 22.5119 | 22.3917 | Ube3b | 117146 | 0.92987 | 1.070127851 | 1.15083 | 0.41191 | 0.835788 |
| 16905129 | N-terminal EF-hand calcium-binding protein 2 [Mus musculus] | 0 | 17.009 | 16 | 4 | 15 | 4 | 389 | 43.4 | 5.3 | 96 | 4 | 4668886.315 | 3792540.199 | 9229071.757 | 7810096.268 | 4452633.959 | 11473936.23 | 7383863.938 | 12149007.15 | 40.2827 | 40.8046 | Necab2 | 117148 | 0.83663 | 1.163366812 | 1.39053 | 0.30457 | 0.761857 |
| 187957332 | Synaptopodin 2 [Mus musculus] | 0.001 | 6.573 | 2 | 2 | 3 | 2 | 1257 | 135.7 | 8.13 | 40 | 2 | 10852585.87 | 13804692.11 | 9393316.905 | 10027102.19 | 9783146.423 | 12794753.4 | 9045797 | 12042062 | 17.7014 | 16.3686 | Synpo2 | 118449 | 1.00469 | 0.995305197 | 0.99065 | 0.94047 | 0.994906 |
| 568971280 | PREDICTED: gap junction gamma-2 protein isoform X1 [Mus musculus] | 0 | 39.689 | 15 | 3 | 16 | 3 | 572 | 61.2 | 7.25 | 278 | 3 | 12794397.49 | 12265026.79 | 13225616.84 | 11000343.85 | 18995359.46 | 10035566.36 | 11381815.25 | 8120253.71 | 7.82637 | 39.2843 | Gjc2 | 118454 | 1.00769 | 0.992308294 | 0.98473 | 0.94085 | 0.994906 |
| 921274049 | small conductance calcium-activated potassium channel protein 2 isoform SK2-L [Mus musculus] | 0 | 25.233 | 7 | 5 | 17 | 3 | 849 | 92.7 | 9.19 | 83 | 5 | 3672035.686 | 552752.0942 | 2522515.649 | 880776.8979 | 2490839.466 | 1270973.452 | 2531836 | 909734.5063 | 76.4786 | 46.2962 | Kcnn2 | 140492 | 1.02863 | 0.971365139 | 0.94432 | 0.90354 | 0.98561 |
| 32189434 | immunoglobulin superfamily member 8 precursor [Mus musculus] | 0 | 91.407 | 43 | 11 | 34 | 11 | 611 | 65 | 7.84 | 787 | 11 | 66925999.69 | 42047377.71 | 24188368.42 | 26725481.13 | 27597917.77 | 21994709.39 | 18672117.5 | 30295429.98 | 49.097 | 21.3915 | Igsf8 | 140559 | 1.23729 | 0.762709735 | 0.61644 | 0.18203 | 0.715071 |
| 26341524 | unnamed protein product [Mus musculus] | 0 | 197.764 | 47 | 26 | 148 | 21 | 720 | 82.5 | 5.96 | 3070 | 26 | 181356705 | 133453091.4 | 218602660.7 | 214538849.3 | 249509593.1 | 174942949.4 | 230306598.6 | 159203764.5 | 21.069 | 21.2568 | Elmo2 | 140579 | 0.95774 | 1.042263268 | 1.08826 | 0.59311 | 0.902243 |
| 148700731 | engulfment and cell motility 1, ced-12 homolog (C. elegans), isoform CRA_b [Mus musculus] | 0 | 77.702 | 20 | 15 | 71 | 10 | 817 | 93.4 | 6.95 | 680 | 15 | 44207643.62 | 36521333.06 | 57733957.68 | 41558700.58 | 51638099.14 | 56433300.94 | 50470354 | 50972450.67 | 20.1416 | 5.24099 | Elmo1 | 140580 | 0.92429 | 1.075712083 | 1.16383 | 0.17049 | 0.711915 |
| 16554627 | WD repeat-containing protein 5 [Homo sapiens] | 0.001 | 6.145 | 7 | 2 | 6 | 2 | 334 | 36.6 | 8.27 | 15 | 2 | 2873661.261 | 3606057.713 | 2843962.13 | 2834614.347 | 5110571.713 | 3018548.886 | 5842964 | 7276681.905 | 12.4367 | 33.3969 | Wdr5 | 140858 | 0.72789 | 1.272112258 | 1.74768 | 0.04617 | 0.591577 |
| 27923929 | hippocalcin-like protein 4 [Mus musculus] | 0 | 32.833 | 41 | 7 | 21 | 5 | 191 | 22.2 | 4.89 | 271 | 7 | 15444561.9 | 13367066.65 | 14389160.31 | 13521801.75 | 7319467.898 | 15105524.57 | 10194866 | 8622695.615 | 6.73561 | 33.0329 | Hpcal4 | 170638 | 1.15802 | 0.841984243 | 0.72709 | 0.07126 | 0.610935 |
| 18700024 | isocitrate dehydrogenase [NAD] subunit beta, mitochondrial [Mus musculus] | 0 | 108.201 | 51 | 18 | 109 | 18 | 384 | 42.2 | 8.6 | 1543 | 18 | 140101614.7 | 202798229.6 | 255148210 | 177908264.6 | 219477427.7 | 187980257 | 262396382.7 | 196169078.9 | 24.866 | 15.4162 | Idh3b | 170718 | 0.94515 | 1.054852591 | 1.11607 | 0.47178 | 0.863085 |
| 37360098 | mKIAA0791 protein [Mus musculus] | 0 | 160.984 | 31 | 27 | 81 | 27 | 1423 | 158.5 | 6.24 | 1221 | 27 | 60515731.81 | 52979673.58 | 58016643.2 | 52582631.1 | 80364128.74 | 50638379.2 | 65700417.25 | 42677749.24 | 6.93285 | 27.8741 | Nup155 | 170762 | 0.96702 | 1.032981246 | 1.06821 | 0.67108 | 0.922517 |
| 28972037 | mKIAA0027 protein [Mus musculus] | 0 | 14.087 | 9 | 2 | 10 | 2 | 423 | 46.1 | 8.38 | 229 | 2 | 7848852.599 | 9581266.184 | 7008048.074 | 5773527.746 | 7769025.777 | 15477267.44 | 6153111.5 | 6510319.296 | 21.163 | 48.8815 | Mlc1 | 170790 | 0.91382 | 1.086175244 | 1.1886 | 0.56422 | 0.895889 |
| 118403314 | RNA-binding protein 39 isoform a [Mus musculus] | 0 | 25.24 | 12 | 5 | 15 | 5 | 530 | 59.4 | 10.1 | 673 | 5 | 12069256.21 | 9764619.403 | 11572253.88 | 14195375.41 | 10943662.26 | 11448773.42 | 12284487.75 | 10339833.65 | 15.3148 | 7.31259 | Rbm39 | 170791 | 1.02791 | 0.97209246 | 0.9457 | 0.54201 | 0.886555 |
| 1039735923 | PREDICTED: phosphatidylinositol 4,5-bisphosphate 5-phosphatase A isoform X1 [Mus musculus] | 0 | 19.513 | 6 | 4 | 14 | 4 | 1039 | 111.2 | 9.38 | 85 | 4 | 10843827.79 | 4706372.376 | 11724600.93 | 16746654.88 | 20628780.27 | 6227821.037 | 19320950.5 | 6701855.154 | 44.8787 | 59.1585 | Inpp5j | 170835 | 0.90859 | 1.091412509 | 1.20122 | 0.64901 | 0.915864 |
| 226693351 | delphilin isoform 1 [Mus musculus] | 0 | 160.653 | 27 | 23 | 156 | 23 | 1203 | 131.9 | 7.02 | 2935 | 23 | 144302001.3 | 96343155.79 | 139761430.4 | 109096515.1 | 103513869.6 | 123811802.5 | 111837629.8 | 115208046.6 | 19.0887 | 7.39523 | Grid2ip | 170935 | 1.03722 | 0.962779209 | 0.92823 | 0.50573 | 0.88031 |
| 148707307 | non-metastatic cells 7, protein expressed in, isoform CRA_b, partial [Mus musculus] | 0 | 7.029 | 8 | 3 | 11 | 3 | 421 | 47.6 | 7.05 | 23 | 3 | 5884263.219 | 4493112.843 | 7392518.684 | 7836594.234 | 6196225.409 | 5445484.795 | 6449962.5 | 6594418.682 | 23.778 | 8.28392 | Nme7 | 171567 | 1.0183 | 0.981699138 | 0.96406 | 0.78406 | 0.952792 |
| 568920807 | PREDICTED: neuroligin-1 isoform X1 [Mus musculus] | 0 | 78.174 | 19 | 12 | 112 | 8 | 863 | 96.4 | 6.4 | 2993 | 12 | 55535126.58 | 58924737.22 | 53034119.7 | 37700932.29 | 43512670.95 | 45141497.36 | 46557320.25 | 39958186.33 | 18.2872 | 6.4915 | Nlgn1 | 192167 | 1.07894 | 0.921061945 | 0.85367 | 0.17652 | 0.713031 |
| 19344056 | BC003940 protein, partial [Mus musculus] | 0 | 12.196 | 30 | 3 | 5 | 3 | 126 | 14.3 | 10.67 | 40 | 3 | 8112577.425 | 11351976.08 | 16772827.91 | 9640030.795 | 14844851.36 | 18185198.25 | 16220814.5 | 16288081.92 | 32.915 | 8.37511 | Mcrip1 | 192173 | 0.82353 | 1.176469004 | 1.42857 | 0.04997 | 0.592616 |
| 569009889 | PREDICTED: filamin-A isoform X1 [Mus musculus] | 0 | 65.619 | 8 | 14 | 41 | 12 | 2647 | 281 | 6.04 | 511 | 14 | 9783555.18 | 13167477.52 | 14461107.31 | 12537911.21 | 9538506.773 | 6978981.737 | 20098408.5 | 13285634.7 | 15.7952 | 45.7211 | Flna | 192176 | 1.00049 | 0.999514084 | 0.99903 | 0.99692 | 0.998251 |
| 12659086 | CGI-74-like SR-rich protein [Mus musculus] | 0 | 16.557 | 11 | 3 | 12 | 3 | 392 | 46.6 | 10.02 | 222 | 3 | 5128106.179 | 3842556.931 | 4251663.053 | 3645376.621 | 12670910.66 | 4835342.485 | 9642471 | 4179349.017 | 15.6001 | 51.6094 | Luc7l2 | 192196 | 0.69997 | 1.300033978 | 1.85728 | 0.12792 | 0.693437 |
| 568972299 | PREDICTED: breast carcinoma-amplified sequence 3 homolog isoform X1 [Mus musculus] | 0 | 23.453 | 8 | 5 | 19 | 5 | 963 | 104.8 | 6.74 | 89 | 5 | 7347359.12 | 11555304.67 | 8151628.627 | 9478309.239 | 5640071.567 | 3646314.243 | 7541192.75 | 2030300.733 | 20.1281 | 50.7785 | Bcas3 | 192197 | 1.31909 | 0.680906862 | 0.51619 | 0.02636 | 0.515682 |
| 126116574 | NAC-alpha domain-containing protein 1 [Mus musculus] | 0 | 7.63 | 2 | 2 | 5 | 2 | 1504 | 156.7 | 4.56 | 29 | 2 | 2401711.826 | 1621399.892 | 1567325.07 | 2390212.634 | 2197950.017 | 2194301.137 | 2020826.375 | 1512732.731 | 23.2239 | 16.3126 | Nacad | 192950 | 1.00345 | 0.996552397 | 0.99313 | 0.96287 | 0.996334 |
| 755516345 | PREDICTED: protein-methionine sulfoxide oxidase MICAL3 isoform X1 [Mus musculus] | 0 | 103.677 | 14 | 27 | 76 | 27 | 2252 | 253 | 5.8 | 651 | 27 | 55434044.53 | 58033508.77 | 49624803.14 | 53121499.01 | 43379864.99 | 57799119.91 | 47567081.13 | 63592343.85 | 6.60402 | 17.4446 | Mical3 | 194401 | 1.00904 | 0.990956889 | 0.98208 | 0.85165 | 0.975109 |
| 568994816 | PREDICTED: syntaxin-binding protein 5-like isoform X1 [Mus musculus] | 0 | 9.063 | 2 | 2 | 5 | 2 | 1216 | 135.2 | 7.02 | 15 | 2 | 2535407.067 | 1976199.006 | 4442950.946 | 1162776.56 | 4938752.551 | 2385177.299 | 2016628.125 | 1785184.099 | 55.1403 | 52.465 | Stxbp5l | 207227 | 0.95253 | 1.047469986 | 1.09967 | 0.81109 | 0.961921 |
| 126723307 | constitutive coactivator of PPAR-gamma-like protein 2 [Mus musculus] | 0 | 54.698 | 15 | 12 | 41 | 11 | 1091 | 119.6 | 8.46 | 538 | 12 | 30603507.5 | 33561571.79 | 37550332.8 | 33953464.55 | 31913486.26 | 33263073.01 | 37951858.75 | 38982134.24 | 8.39255 | 9.75054 | Fam120c | 207375 | 0.97681 | 1.023189894 | 1.04748 | 0.49954 | 0.877125 |
| 62990174 | protein phosphatase 1 regulatory subunit 29 precursor [Mus musculus] | 0 | 167.186 | 41 | 21 | 170 | 18 | 823 | 90 | 7.52 | 1858 | 21 | 209552762.1 | 229938914.5 | 180497344 | 203229632.7 | 206919018.1 | 250506308.2 | 183710472.2 | 194531955.1 | 9.89293 | 14.026 | Elfn2 | 207393 | 0.9925 | 1.007504492 | 1.01512 | 0.86725 | 0.979074 |
| 74226654 | unnamed protein product [Mus musculus] | 0 | 76.867 | 37 | 11 | 45 | 11 | 496 | 55 | 7.37 | 896 | 11 | 55844209.1 | 40402723.39 | 45221091.04 | 45878338.87 | 38781797.61 | 35496248.98 | 45485647.94 | 42521724.65 | 13.8402 | 10.7356 | Wdr37 | 207615 | 1.07168 | 0.928321896 | 0.86623 | 0.15973 | 0.700858 |
| 74177284 | unnamed protein product [Mus musculus] | 0 | 56.641 | 15 | 7 | 15 | 7 | 945 | 106.3 | 5.29 | 324 | 7 | 30259501.69 | 9446675.041 | 11671636.55 | 12044642.64 | 7221030.859 | 6055368.376 | 7864940 | 3149195.015 | 60.9932 | 34.3821 | Pde2a | 207728 | 1.44614 | 0.55386401 | 0.383 | 0.09534 | 0.670764 |
| 298493223 | transmembrane protein 132B precursor [Mus musculus] | 0 | 203.966 | 39 | 26 | 171 | 7 | 1078 | 119.3 | 4.87 | 3464 | 26 | 33842195.19 | 18315632.91 | 22624118.71 | 14084428.54 | 17657474.96 | 11700036.38 | 17437470.5 | 14722729.86 | 38.2527 | 18.1574 | Tmem132b | 208151 | 1.18186 | 0.81814124 | 0.69225 | 0.17722 | 0.713031 |
| 287327708 | nuclease EXOG, mitochondrial isoform 1 precursor [Mus musculus] | 0 | 31.772 | 19 | 6 | 24 | 6 | 392 | 43.6 | 7.94 | 535 | 6 | 30195134.33 | 20165177.55 | 23294120.23 | 25161242.71 | 26199312.85 | 20648292.64 | 25810939.5 | 13553956.75 | 17.0062 | 27.3862 | Exog | 208194 | 1.06811 | 0.931885114 | 0.87246 | 0.41785 | 0.837049 |
| 568981372 | PREDICTED: disco-interacting protein 2 homolog C isoform X1 [Mus musculus] | 0 | 57.151 | 10 | 14 | 56 | 10 | 1616 | 177.3 | 7.66 | 511 | 14 | 23337527.33 | 16825187.62 | 23261502.11 | 13984545.28 | 21635488.59 | 24806702.93 | 20029737.25 | 19166693.31 | 24.3037 | 11.607 | Dip2c | 208440 | 0.94952 | 1.050475264 | 1.10632 | 0.46856 | 0.863085 |
| 124486588 | sickle tail protein isoform c [Mus musculus] | 0 | 320.29 | 30 | 48 | 410 | 3 | 1997 | 218.3 | 8.28 | 5441 | 48 | 1524600.335 | 2887682.752 | 1625314.377 | 2129155.735 | 601177.2238 | 898934.1221 | 1576948.938 | 2733673.593 | 30.5103 | 65.1612 | Etl4 | 208618 | 1.16856 | 0.831441853 | 0.71151 | 0.3386 | 0.782483 |
| 148665170 | eukaryotic translation initiation factor 4, gamma 1 [Mus musculus] | 0 | 21.484 | 4 | 6 | 14 | 3 | 1628 | 178.5 | 5.26 | 165 | 6 | 548907.1497 | 1269655.78 | 2156633.113 | 652772.4089 | 1957434.916 | 2009853.728 | 2196791 | 1525605.811 | 63.8261 | 14.7622 | Eif4g1 | 208643 | 0.75144 | 1.248563324 | 1.66157 | 0.10113 | 0.674486 |
| 21644583 | GTP-binding protein Di-Ras1 [Mus musculus] | 0 | 29.187 | 39 | 6 | 24 | 5 | 198 | 22.3 | 8.9 | 160 | 6 | 6419940.061 | 5327061.045 | 17802569.81 | 6193723.667 | 13829242.07 | 6439845.294 | 19585352.5 | 5819072.704 | 66.361 | 57.3508 | Diras1 | 208666 | 0.87803 | 1.121967666 | 1.27782 | 0.59446 | 0.902243 |
| 37360034 | mKIAA0666 protein [Mus musculus] | 0 | 25.299 | 6 | 5 | 15 | 5 | 1087 | 124.4 | 7.34 | 197 | 5 | 3588321.034 | 5242448.383 | 6981097.354 | 2653920.389 | 5312350.891 | 3850035.841 | 7076365.75 | 4217504.452 | 41.2748 | 28.3173 | Daam1 | 208846 | 0.94886 | 1.051139909 | 1.10779 | 0.69199 | 0.930425 |
| 148277096 | dedicator of cytokinesis protein 3 [Mus musculus] | 0 | 576.625 | 45 | 72 | 421 | 62 | 2030 | 233.1 | 6.99 | 7623 | 72 | 427304179.8 | 331699371.8 | 433653733 | 339458517.2 | 475831117.7 | 456259662.7 | 477782361.7 | 271622112.7 | 14.3444 | 23.7034 | Dock3 | 208869 | 0.95352 | 1.046483364 | 1.0975 | 0.53591 | 0.886555 |
| 124487217 | protein unc-13 homolog C [Mus musculus] | 0 | 152.686 | 13 | 26 | 176 | 17 | 2210 | 249.7 | 6.07 | 2643 | 26 | 24658987.46 | 21032849.74 | 26930234.15 | 23840927.73 | 29435086.42 | 32570754.68 | 29608674 | 32088705.27 | 10.0996 | 5.2854 | Unc13c | 208898 | 0.87627 | 1.123725708 | 1.28239 | 0.00353 | 0.38143 |
| 26006231 | mKIAA0940 protein [Mus musculus] | 0 | 78.632 | 26 | 14 | 47 | 5 | 722 | 78.9 | 7.42 | 714 | 14 | 17372365.4 | 14956035.92 | 21745337.71 | 14921592.96 | 18490650.32 | 8798633.623 | 24084221.94 | 15945160.63 | 18.6083 | 37.6841 | Cpeb3 | 208922 | 1.0123 | 0.987699976 | 0.9757 | 0.90996 | 0.98561 |
| 148665188 | mCG141781, isoform CRA_b, partial [Mus musculus] | 0 | 15.295 | 4 | 5 | 10 | 5 | 1434 | 161.8 | 5.6 | 0 | 5 | 4600684.608 | 5673170.578 | 2879666.654 | 3916257.897 | 3114336.836 | 3713378.069 | 3248316.125 | 2937479.317 | 27.5157 | 10.206 | Vps8 | 209018 | 1.13483 | 0.865165367 | 0.76237 | 0.14757 | 0.693437 |
| 568912322 | PREDICTED: FERM domain-containing protein 4A isoform X1 [Mus musculus] | 0 | 73.388 | 20 | 13 | 54 | 13 | 1061 | 118.6 | 9.19 | 517 | 13 | 35153486.68 | 35946281.8 | 49458716.87 | 31558641.71 | 55730983.95 | 38022084.86 | 41666173.13 | 43330276.01 | 20.6555 | 17.2059 | Frmd4a | 209630 | 0.91951 | 1.080492825 | 1.17508 | 0.27126 | 0.745606 |
| 23943816 | UPF0258 protein KIAA1024 [Mus musculus] | 0 | 29.754 | 10 | 7 | 18 | 7 | 917 | 102.7 | 7.24 | 308 | 7 | 7025364.71 | 6381668.856 | 7867963.068 | 3936469.184 | 7674356.936 | 4904990.932 | 4060644.125 | 4972804.743 | 26.8277 | 29.0556 | Minar1 | 209743 | 1.07685 | 0.923145205 | 0.85726 | 0.46513 | 0.863085 |
| 568956402 | PREDICTED: piggyBac transposable element-derived protein 5 isoform X1 [Mus musculus] | 0.003 | 4.056 | 3 | 2 | 3 | 2 | 523 | 58.3 | 8.85 | 16 | 2 | 834608.6705 | 2359703.657 | 2812511.873 | 1650814.963 | 1994403.519 | 2524385.521 | 1223878.25 | 1524811.555 | 45.1399 | 31.2813 | Pgbd5 | 209966 | 1.02614 | 0.973858812 | 0.94905 | 0.85662 | 0.975398 |
| 28972596 | mKIAA1060 protein [Mus musculus] | 0 | 26.374 | 7 | 6 | 20 | 5 | 1115 | 125.9 | 8.43 | 211 | 6 | 8040038.764 | 3292654.185 | 9019506.514 | 7282065.44 | 10433914.63 | 5538247.663 | 8232137.625 | 5107670.775 | 36.38 | 33.9765 | Adcy2 | 210044 | 0.97054 | 1.029461224 | 1.06071 | 0.82046 | 0.964114 |
| 26333111 | unnamed protein product [Mus musculus] | 0 | 7.993 | 10 | 2 | 13 | 2 | 325 | 35.6 | 6.68 | 66 | 2 | 377180.1926 | 3003446.176 | 1520496.358 | 2252178.798 | 2054363.727 | 2115009.193 | 1492408 | 7203924.302 | 62.558 | 83.1068 | Iglon5 | 210094 | 0.71465 | 1.285349007 | 1.79857 | 0.36236 | 0.804292 |
| 169790905 | adhesion G protein-coupled receptor B3 precursor [Mus musculus] | 0 | 287.73 | 30 | 35 | 352 | 35 | 1522 | 171.2 | 7.02 | 4683 | 35 | 603655526.1 | 761858159.5 | 720153567.1 | 804231379.9 | 778425692.5 | 902166201.9 | 685319738.3 | 764304430 | 11.9492 | 11.4562 | Adgrb3 | 210933 | 0.96008 | 1.039919078 | 1.08316 | 0.37161 | 0.80688 |

[illegible]

| | | | | | | | | | | | | | | | | | | | | | | | | | | | | | |
|---|---|---|---|---|---|---|---|---|---|---|---|---|---|---|---|---|---|---|---|---|---|---|---|---|---|---|---|---|---|
| 755502295 | PREDICTED: sterile alpha motif domain-containing protein 10 isoform X1 [Mus musculus] | 0 | 8.911 | 10 | 3 | 48 | 2 | 336 | 36.8 | 10.49 | 67 | 3 | 1516569.731 | 3782056.668 | 2741138.906 | 3733568.298 | 4005771.011 | 3247587.161 | 2299123.75 | 3194296.586 | 36.193 | 21.9097 | Samd10 | 229011 | 0.9603 | 1.039699855 | 1.08268 | 0.71553 | 0.936732 |
| 568920862 | PREDICTED: YTH domain-containing family protein 3 isoform X1 [Mus musculus] | 0 | 42.034 | 16 | 8 | 29 | 5 | 600 | 66.1 | 8.78 | 388 | 8 | 10840560.71 | 15023502.14 | 13263923.73 | 15649319.03 | 11315622.69 | 15818916.32 | 14338708.5 | 17649498.3 | 15.7285 | 18.1161 | Ythdf3 | 229096 | 0.96185 | 1.038151345 | 1.07933 | 0.55054 | 0.892023 |
| 54020676 | eukaryotic translation initiation factor 2A [Mus musculus] | 0 | 31.277 | 17 | 8 | 33 | 8 | 581 | 64.4 | 8.91 | 192 | 8 | 13754385.62 | 18289247.58 | 14828817.06 | 24462451.33 | 15445954.18 | 18273605.69 | 17753623.38 | 18560024.67 | 27.0509 | 8.08094 | Eif2a | 229317 | 1.00921 | 0.990792169 | 0.98175 | 0.90122 | 0.98561 |
| 37360236 | mKIAA1089 protein [Mus musculus] | 0 | 18.712 | 5 | 7 | 19 | 7 | 1064 | 119.2 | 6.27 | 67 | 7 | 12631598.67 | 5571728.446 | 15737416.13 | 7977177.882 | 13647046.39 | 8992608.697 | 11797035.38 | 11775878.71 | 43.5987 | 16.6118 | Smg5 | 229512 | 0.95127 | 1.048730559 | 1.10245 | 0.67993 | 0.925878 |
| 27532986 | PRUNEM1 [Mus musculus] | 0.001 | 5.037 | 5 | 2 | 4 | 2 | 454 | 50.3 | 5.15 | 0 | 2 | 646127.7544 | 1380959.251 | 945559.5711 | 2086722.637 | 1611616.037 | 1918072.133 | 1245922.375 | 2013260.78 | 49.4521 | 20.4007 | Prune1 | 229589 | 0.85403 | 1.145971219 | 1.34184 | 0.27194 | 0.745606 |
| 568922614 | PREDICTED: vang-like protein 1 isoform X1 [Mus musculus] | 0 | 8.931 | 4 | 2 | 2 | 2 | 539 | 61.4 | 8.82 | 14 | 2 | 2004903.7 | 1329547.085 | 1391960.619 | 1456166.132 | 2004239.723 | 1809069.853 | 2151783.75 | 1053716.112 | 20.0891 | 27.808 | Vangl1 | 229658 | 0.93666 | 1.063344246 | 1.13526 | 0.49693 | 0.875118 |
| 148675659 | cold shock domain containing E1, RNA binding, isoform CRA_b, partial [Mus musculus] | 0 | 25.737 | 9 | 8 | 24 | 8 | 854 | 95.3 | 6.32 | 102 | 8 | 5551352.755 | 4467952.419 | 6172059.144 | 6022742.531 | 6255108.24 | 5020569.011 | 5115757.406 | 4619672.16 | 13.8749 | 13.3633 | Csde1 | 229663 | 1.02783 | 0.972169021 | 0.94585 | 0.58488 | 0.899211 |
| 169646306 | striatin-interacting protein 1 [Mus musculus] | 0 | 27.204 | 9 | 6 | 18 | 5 | 837 | 95.5 | 6.25 | 243 | 6 | 3766668.023 | 5112387.046 | 7492492.64 | 7215220.856 | 6623461.07 | 12478210.14 | 6628668 | 10605395.1 | 30.0783 | 32.357 | Strip1 | 229707 | 0.78724 | 1.212757564 | 1.54051 | 0.11271 | 0.678142 |
| 21361647 | adenosylhomocysteinase 2 isoform a [Homo sapiens] | 0 | 79.32 | 37 | 16 | 83 | 16 | 530 | 58.9 | 6.89 | 487 | 16 | 56466734.57 | 81237581.67 | 67405891.71 | 81215173.76 | 91822855.67 | 79258238.43 | 63313635.27 | 78651602.4 | 16.763 | 14.9102 | Ahcyl1 | 229709 | 0.95542 | 1.0445816 | 1.09332 | 0.45516 | 0.860877 |
| 21955259 | noelin-3 precursor [Rattus norvegicus] | 0 | 39.457 | 23 | 10 | 45 | 10 | 478 | 54.9 | 8.15 | 485 | 10 | 74744129.51 | 87066495.23 | 75429208.9 | 85067259.83 | 82582465.54 | 102067448.4 | 81748927.13 | 104018115.8 | 7.94009 | 13.0496 | Olfm3 | 229759 | 0.93055 | 1.069450257 | 1.14927 | 0.12903 | 0.693437 |
| 37359962 | mKIAA0455 protein [Mus musculus] | 0 | 68.683 | 25 | 15 | 91 | 15 | 795 | 86.3 | 9.07 | 1283 | 15 | 17960702.65 | 26606141.68 | 29112310.64 | 44678650.81 | 42144699.01 | 51984201.13 | 28613365.81 | 58685445.32 | 37.6354 | 28.8092 | Plppr4 | 229791 | 0.78962 | 1.21038343 | 1.53287 | 0.11589 | 0.678142 |
| 100818161 | rap1 GTPase-GDP dissociation stimulator 1 isoform a [Mus musculus] | 0 | 61.452 | 24 | 11 | 42 | 11 | 607 | 66 | 5.35 | 675 | 11 | 27354705 | 37247723.58 | 32582617.88 | 34678773.47 | 35725773.71 | 38147565.23 | 32670121.63 | 34949035.14 | 12.7381 | 6.38594 | Rap1gds1 | 229877 | 0.96478 | 1.035223901 | 1.07302 | 0.35162 | 0.797646 |
| 755503815 | PREDICTED: adenylate kinase isoenzyme 5 isoform X1 [Mus musculus] | 0 | 23.561 | 12 | 5 | 21 | 5 | 566 | 63.6 | 5.72 | 233 | 5 | 13076733.72 | 9034763.55 | 12839812.67 | 8995135.303 | 13339863.43 | 16300264.94 | 31156460.88 | 7876987.752 | 20.7415 | 57.9943 | Ak5 | 229949 | 0.78044 | 1.219562483 | 1.56267 | 0.27163 | 0.745606 |
| 568926465 | PREDICTED: PHD finger protein 24 isoform X1 [Mus musculus] | 0 | 8.122 | 6 | 2 | 6 | 2 | 431 | 49.1 | 4.96 | 39 | 2 | 5579627.364 | 4882632.093 | 3899082.509 | 1117252.6 | 5696128.551 | 8276152.304 | 4375958.875 | 7570142.413 | 50.6548 | 27.3997 | Phf24 | 230085 | 0.74781 | 1.252187198 | 1.67447 | 0.09585 | 0.670764 |
| 294610683 | solute carrier family 25 member 51 [Mus musculus] | 0 | 63.305 | 54 | 13 | 57 | 13 | 298 | 33.7 | 9.66 | 428 | 13 | 52297935.02 | 27483179.29 | 53425498.59 | 37690895.97 | 55597109.32 | 37910255.72 | 62712213.69 | 37852953.8 | 29.1023 | 26.0126 | Slc25a51 | 230125 | 0.9365 | 1.063498427 | 1.13561 | 0.53733 | 0.886555 |
| 148670309 | of kappa light polypeptide enhancer in B-cells, kinase complex-associated protein, isoform CRA_a, partial [Mus m | 0 | 14.434 | 3 | 3 | 6 | 3 | 1371 | 153.6 | 6.15 | 59 | 3 | 2625567.009 | 1640194.402 | 1834306.757 | 1779933.398 | 1641656.669 | 2137639.927 | 2414965.5 | 1749400.362 | 22.5699 | 17.958 | Elp1 | 230233 | 0.99598 | 1.004023145 | 1.00808 | 0.95728 | 0.994906 |
| 68533246 | thyroid hormone receptor-associated protein 3 [Mus musculus] | 0 | 15.458 | 7 | 6 | 16 | 5 | 951 | 108.1 | 10.17 | 30 | 6 | 3553062.424 | 5271520.612 | 6581237.74 | 9796831.837 | 2186842.618 | 8954093.421 | 4929057.625 | 8917727.395 | 41.9019 | 52.8364 | Thrap3 | 230753 | 1.00428 | 0.995717674 | 0.99147 | 0.98054 | 0.997464 |
| 755508137 | PREDICTED: adhesion G protein-coupled receptor B2 isoform X1 [Mus musculus] | 0 | 283.27 | 30 | 33 | 318 | 33 | 1581 | 172.1 | 7.46 | 6951 | 33 | 249196573.7 | 310637440.2 | 321033025.6 | 315332868.1 | 345423685.1 | 340232198.9 | 302400557.3 | 337212365.7 | 11.2043 | 5.90786 | Adgrb2 | 230775 | 0.94881 | 1.051187984 | 1.1079 | 0.14736 | 0.693437 |
| 1039767501 | PREDICTED: eukaryotic translation initiation factor 4 gamma 3 isoform X1 [Mus musculus] | 0 | 46.79 | 8 | 12 | 31 | 9 | 1795 | 196.8 | 6.25 | 271 | 12 | 5360148.401 | 4821373.995 | 7217770.926 | 8878412.117 | 9047469.845 | 7713975.694 | 6034349.344 | 16244967.04 | 28.1639 | 46.0598 | Eif4g3 | 230861 | 0.8046 | 1.195397366 | 1.4857 | 0.23726 | 0.73104 |
| 22477590 | Sh2d5 protein, partial [Mus musculus] | 0 | 21.93 | 15 | 5 | 9 | 5 | 486 | 53.3 | 8.79 | 111 | 5 | 5392412.007 | 9153403.609 | 7796500.225 | 2595280.253 | 7197073.14 | 11977135.83 | 6691437.75 | 1658482.586 | 46.2228 | 61.3027 | Sh2d5 | 230863 | 0.9507 | 1.049303243 | 1.10372 | 0.8086 | 0.961747 |
| 120537280 | C230096C10Rik protein [Mus musculus] | 0 | 42.446 | 13 | 9 | 24 | 9 | 997 | 111.6 | 7.43 | 292 | 9 | 12406470.38 | 8443215.547 | 11460723.68 | 16233198.51 | 21141995.63 | 11038772.52 | 17594754 | 12659850.35 | 26.4678 | 29.6237 | Emc1 | 230866 | 0.87483 | 1.125174734 | 1.28617 | 0.26345 | 0.745606 |
| 568931100 | PREDICTED: rootletin isoform X1 [Mus musculus] | 0 | 408.931 | 44 | 75 | 432 | 74 | 1935 | 219 | 5.6 | 8038 | 75 | 377286197 | 330752873 | 601183319.9 | 595404265.3 | 491604092.3 | 273376850.4 | 448735990.5 | 564680671.9 | 29.8904 | 27.8404 | Crocc | 230872 | 1.03427 | 0.965726794 | 0.93373 | 0.74931 | 0.949513 |
| 33859801 | F-box only protein 2 [Mus musculus] | 0 | 33.659 | 29 | 6 | 31 | 6 | 297 | 33.7 | 4.28 | 450 | 6 | 10227980.31 | 25166119.26 | 26031616.03 | 14435193.86 | 29242267.3 | 16114448.16 | 25381248.5 | 11627963.76 | 41.4338 | 39.47 | Fbxo2 | 230904 | 0.95889 | 1.041111978 | 1.08575 | 0.78325 | 0.952792 |
| 148682887 | mCG16669, isoform CRA_e, partial [Mus musculus] | 0 | 23.771 | 15 | 5 | 20 | 5 | 416 | 44.7 | 6.99 | 140 | 5 | 3753000.825 | 1785974.843 | 3309768.409 | 5014881.93 | 4136809.689 | 3663034.172 | 2834367.5 | 2400691.74 | 38.4518 | 24.103 | Tardbp | 230908 | 1.03081 | 0.969190772 | 0.94022 | 0.79778 | 0.958226 |
| 546231934 | transmembrane protein 201 isoform c [Mus musculus] | 0 | 36.559 | 9 | 6 | 51 | 6 | 664 | 72.5 | 9.2 | 611 | 6 | 62089414.95 | 70579279.53 | 46977609.85 | 78993039.95 | 58334069.54 | 100790027.3 | 58566275.5 | 81235871.28 | 21.1252 | 27.3312 | Tmem201 | 230917 | 0.92775 | 1.072254996 | 1.15576 | 0.44368 | 0.852048 |
| 1039767586 | PREDICTED: dnaJ homolog subfamily C member 11 isoform X1 [Mus musculus] | 0 | 65.996 | 28 | 12 | 48 | 12 | 597 | 67.1 | 8.53 | 489 | 12 | 39026241.64 | 25768895.65 | 35270141.58 | 21553304.73 | 44466353.52 | 35097740.52 | 45966298.56 | 24811467.15 | 26.709 | 26.0209 | Dnajc11 | 230935 | 0.89438 | 1.10561564 | 1.23618 | 0.30171 | 0.761857 |
| 150010629 | adherens junction-associated protein 1 [Mus musculus] | 0 | 50.954 | 20 | 6 | 30 | 6 | 412 | 44.7 | 9.42 | 1175 | 6 | 38381934.34 | 38169933.78 | 37404131.45 | 35633375.28 | 36274676.79 | 39585221.61 | 35332846.25 | 33311315.42 | 3.33907 | 7.24299 | Ajap1 | 230959 | 1.01729 | 0.982708506 | 0.966 | 0.41422 | 0.835788 |
| 148705321 | mCG11629, isoform CRA_d, partial [Mus musculus] | 0 | 87.004 | 30 | 13 | 77 | 13 | 509 | 55.1 | 9.42 | 1408 | 13 | 129705420.2 | 94595406.04 | 121335761.7 | 137985359.4 | 98248297.94 | 94397844.22 | 108222343.2 | 87667697.49 | 15.5585 | 8.76209 | Hadhb | 231086 | 1.10877 | 0.891230721 | 0.8038 | 0.0613 | 0.601634 |
| 568934124 | PREDICTED: actin-binding LIM protein 2 isoform X1 [Mus musculus] | 0 | 331.529 | 62 | 31 | 335 | 2 | 715 | 78.8 | 8.18 | 5648 | 31 | 19028657.62 | 14097318.12 | 22142136.79 | 24262556.45 | 17786307.93 | 12533704.5 | 20326866.13 | 16107597.82 | 22.2075 | 19.5832 | Ablim2 | 231148 | 1.08734 | 0.912662406 | 0.83936 | 0.28903 | 0.752733 |
| 112807186 | eIF-2-alpha kinase activator GCN1 [Mus musculus] | 0 | 119.749 | 13 | 29 | 86 | 29 | 2671 | 292.8 | 7.36 | 762 | 29 | 31368013.44 | 23542352.25 | 31637053.72 | 29791398.75 | 34441219.06 | 44177232.73 | 38213152.5 | 36585707.28 | 13.0086 | 10.8931 | Gcn1 | 231659 | 0.86255 | 1.137451903 | 1.31871 | 0.01663 | 0.461382 |
| 568937790 | PREDICTED: RIMS-binding protein 2 isoform X2 [Mus musculus] | 0 | 316.901 | 40 | 34 | 328 | 34 | 1363 | 150.2 | 5.78 | 5071 | 34 | 404425502.5 | 396301349 | 421331583.3 | 426431734 | 420615053.6 | 443760100.7 | 385415089.2 | 479538523.9 | 3.429 | 9.15287 | Rimbp2 | 231760 | 0.97607 | 1.023932188 | 1.04904 | 0.37323 | 0.80688 |
| 123785178 | RecName: Full=Uncharacterized protein C7orf43 homolog | 0 | 19.212 | 8 | 3 | 10 | 3 | 580 | 62.7 | 8.68 | 258 | 3 | 6703042.979 | 3697546.407 | 6241316.506 | 4873259.453 | 4061698.115 | 4241643.45 | 2881720.75 | 3959109.741 | 25.3549 | 16.2198 | Map11 | 231807 | 1.17379 | 0.826210899 | 0.70388 | 0.07723 | 0.627416 |
| 148368976 | arf-GAP with dual PH domain-containing protein 1 [Mus musculus] | 0 | 37.705 | 33 | 8 | 26 | 8 | 374 | 43.3 | 8.65 | 236 | 8 | 24846558.97 | 17934174.4 | 23164505.07 | 25782464.72 | 27229024.6 | 12024545.9 | 25163333 | 18400076.75 | 15.2777 | 33.3583 | Adap1 | 231821 | 1.05105 | 0.948948758 | 0.90286 | 0.586 | 0.899211 |
| 568938676 | PREDICTED: ras-associating and dilute domain-containing protein isoform X1 [Mus musculus] | 0 | 50.918 | 12 | 9 | 35 | 9 | 1122 | 122.7 | 6.57 | 348 | 9 | 43429758.23 | 37825409.61 | 37903500.73 | 33542159.75 | 36431127.97 | 29150108.39 | 28051628.38 | 28157805.51 | 10.6159 | 13.2014 | Radil | 231858 | 1.11261 | 0.887391201 | 0.79758 | 0.03522 | 0.540817 |
| 124487401 | serine/threonine-protein kinase LMTK2 precursor [Mus musculus] | 0 | 27.401 | 5 | 6 | 18 | 6 | 1471 | 160.4 | 4.51 | 293 | 6 | 12919711.88 | 3425074.131 | 12194051.49 | 7165403.095 | 11041209.92 | 7026953.899 | 13345243.5 | 2577814.953 | 50.0977 | 55.6853 | Lmtk2 | 231876 | 1.02458 | 0.97542138 | 0.95202 | 0.89964 | 0.985548 |
| 568941572 | PREDICTED: IQ motif and SEC7 domain-containing protein 1 isoform X1 [Mus musculus] | 0 | 543.565 | 54 | 53 | 868 | 4 | 1203 | 133.9 | 8.38 | 19472 | 53 | 134257330.8 | 142462399.6 | 119645482.5 | 142884448.3 | 163194272.6 | 180965099.4 | 126010894.4 | 173174112.1 | 8.05797 | 15.1268 | Iqsec1 | 232227 | 0.91198 | 1.088022359 | 1.19304 | 0.09859 | 0.670764 |
| 21450317 | histone deacetylase 11 [Mus musculus] | 0 | 33.727 | 37 | 8 | 18 | 8 | 347 | 39.1 | 7.14 | 60 | 8 | 22788679.1 | 27131541.96 | 24489008.2 | 24250495.6 | 23769944.36 | 21398854.68 | 25208133.13 | 2250253.872 | 7.33047 | 59.0415 | Hdac11 | 232232 | 1.15198 | 0.848017933 | 0.73614 | 0.27635 | 0.745606 |
| 568941650 | PREDICTED: ankyrin repeat domain-containing protein 26 isoform X1 [Mus musculus] | 0 | 12.94 | 3 | 4 | 7 | 4 | 1686 | 192.7 | 5.95 | 30 | 4 | 6113567.385 | 3119958.801 | 6575280.321 | 4857700.111 | 6916440.56 | 1407190.775 | 4929525.438 | 3611760.236 | 29.9123 | 54.8699 | Ankrd26 | 232339 | 1.10129 | 0.898709155 | 0.81605 | 0.51997 | 0.880629 |
| 158563816 | RecName: Full=Coiled-coil domain-containing protein 136 | 0 | 49.425 | 11 | 9 | 28 | 9 | 1136 | 131.8 | 4.84 | 510 | 9 | 40326114.78 | 76762290.06 | 64179635.02 | 98365955.75 | 95026964.1 | 108634487.7 | 82069548.91 | 92455701.36 | 34.6936 | 11.5661 | Ccdc136 | 232664 | 0.85018 | 1.149816974 | 1.35243 | 0.11345 | 0.678142 |
| 568945295 | PREDICTED: CCR4-NOT transcription complex subunit 3 isoform X1 [Mus musculus] | 0.001 | 6.558 | 3 | 2 | 2 | 2 | 752 | 82 | 6.2 | 18 | 2 | 947292.6871 | 1660815.378 | 2384672.652 | 5851696.794 | 982134.618 | 1458271.039 | 5162860.875 | 5572374.819 | 80.203 | 73.1106 | Cnot3 | 232791 | 0.90295 | 1.097050471 | 1.21496 | 0.73172 | 0.943908 |
| 594009839 | protein shisa-7 isoform 2 precursor [Mus musculus] | 0 | 169.24 | 38 | 17 | 478 | 17 | 541 | 56.4 | 9.86 | 8106 | 17 | 612382773.3 | 832475593 | 606660073.5 | 878685281.1 | 664439839.5 | 925585600.3 | 651579836 | 771343024.4 | 19.5657 | 16.8368 | Shisa7 | 232813 | 0.98608 | 1.013922676 | 1.02824 | 0.83601 | 0.969252 |
| 75677442 | rho GTPase-activating protein 35 [Mus musculus] | 0 | 11.778 | 3 | 4 | 11 | 4 | 1499 | 170.3 | 6.61 | 46 | 4 | 3203880.414 | 3621713.44 | 4086745.333 | 3756785.821 | 4878361.459 | 3149718.332 | 3562589.75 | 3552019.668 | 9.9666 | 19.901 | Arhgap35 | 232906 | 0.98411 | 1.015885118 | 1.03228 | 0.78685 | 0.952792 |
| 56961624 | AP-2 complex subunit sigma [Rattus norvegicus] | 0 | 12.416 | 21 | 3 | 26 | 3 | 142 | 17 | 6.18 | 142 | 3 | 16978195.31 | 34193426.96 | 26225119.7 | 20747476.42 | 21841110.03 | 31545550.04 | 25871445.25 | 44416098.09 | 30.4622 | 31.8241 | Ap2s1 | 232910 | 0.88491 | 1.11509407 | 1.26013 | 0.3414 | 0.78458 |
| 568945451 | PREDICTED: MAP/microtubule affinity-regulating kinase 4 isoform X1 [Mus musculus] | 0 | 85.793 | 31 | 17 | 88 | 16 | 688 | 75.4 | 9.79 | 734 | 17 | 54865171.04 | 62249598.26 | 61339106.79 | 64296692.71 | 69925785.42 | 65890860.71 | 59663798.69 | 63937856.23 | 6.71294 | 6.57655 | Mark4 | 232944 | 0.96681 | 1.033191488 | 1.06866 | 0.20738 | 0.715071 |
| 755519436 | PREDICTED: sodium/potassium-transporting ATPase subunit alpha-3 isoform X1 [Mus musculus] | 0 | 331.121 | 46 | 42 | 408 | 25 | 1066 | 117.2 | 5.64 | 7810 | 42 | 176743886.7 | 214847229.7 | 235095182.9 | 260205369.6 | 252024621.8 | 312427717.6 | 208409743.4 | 268168601.9 | 15.9034 | 16.5121 | Atp1a3 | 232975 | 0.92005 | 1.079950842 | 1.1738 | 0.21494 | 0.716066 |
| 74209147 | unnamed protein product [Mus musculus] | 0 | 121.627 | 23 | 18 | 81 | 16 | 1324 | 142 | 9.47 | 1185 | 18 | 53377346.31 | 54064486.72 | 64101786.61 | 64353854.13 | 64253688.91 | 61046207.41 | 59656064.66 | 45739403.96 | 10.2986 | 14.1934 | Arhgap33 | 233071 | 1.01115 | 0.98885086 | 0.97795 | 0.80711 | 0.961747 |
| 148675233 | desmuslin, isoform CRA_b [Mus musculus] | 0 | 52.534 | 9 | 11 | 44 | 11 | 1561 | 173.1 | 5.12 | 721 | 11 | 35630962.37 | 11194732.64 | 26748006.69 | 12874177.65 | 36287809.8 | 119284651.1 | 52008832 | 132707037 | 53.9458 | 56.4255 | Synm | 233335 | 0.40516 | 1.594841602 | 3.93634 | 0.04237 | 0.585646 |
| 158937308 | vacuolar protein sorting-associated protein 33B [Mus musculus] | 0 | 24.517 | 11 | 5 | 6 | 5 | 617 | 70.5 | 6.86 | 92 | 5 | 1911098.025 | 8184163.774 | 2621987.145 | 2515738.017 | 4145922.528 | 7846423.916 | 2864092.5 | 8915524.095 | 77.0441 | 48.7371 | Vps33b | 233405 | 0.78108 | 1.218920319 | 1.56056 | 0.34033 | 0.78329 |
| 21704174 | spondin-1 precursor [Mus musculus] | 0 | 15.544 | 6 | 4 | 11 | 4 | 807 | 90.8 | 6.02 | 94 | 4 | 1930633.93 | 3923058.089 | 3698304.414 | 1435904.515 | 3561949.386 | 2436186.189 | 4513581.25 | 1716552.508 | 45.4367 | 40.3237 | Spon1 | 233744 | 0.94657 | 1.053426916 | 1.11288 | 0.73578 | 0.943908 |
| 1039778399 | PREDICTED: pleckstrin homology domain-containing family A member 7 isoform X2 [Mus musculus] | 0 | 93.06 | 19 | 20 | 68 | 20 | 1359 | 154.3 | 9.26 | 886 | 20 | 110564562.3 | 44960406.7 | 108088687.6 | 76641818.7 | 108615010.9 | 76949104.45 | 102537020 | 72759370.99 | 36.2953 | 19.9428 | Plekha7 | 233765 | 0.97061 | 1.029388905 | 1.06056 | 0.78281 | 0.952792 |
| 568950403 | PREDICTED: ataxin-2-like protein isoform X2 [Mus musculus] | 0 | 39.444 | 13 | 10 | 28 | 10 | 1096 | 115.2 | 8.66 | 134 | 10 | 20550994.19 | 22659017.7 | 19902747.83 | 26084975.35 | 20987090.71 | 14743162.21 | 25609681.63 | 16678049.97 | 12.487 | 24.7889 | Atxn2l | 233871 | 1.06686 | 0.933141749 | 0.87466 | 0.35509 | 0.798662 |
| 148685673 | fusion, derived from t(12;16) malignant liposarcoma (human), isoform CRA_f, partial [Mus musculus] | 0 | 28.679 | 13 | 5 | 12 | 5 | 531 | 54.1 | 9.38 | 117 | 5 | 9549846.884 | 19659930.75 | 21889396.43 | 23410080.13 | 17074029.58 | 24953304.29 | 24583582 | 39169875.4 | 33.5236 | 34.892 | Fus | 233908 | 0.82655 | 1.173451269 | 1.4197 | 0.21009 | 0.716066 |
| 755524176 | PREDICTED: liprin-alpha-1 isoform X1 [Mus musculus] | 0 | 158.54 | 24 | 27 | 154 | 10 | 1287 | 145 | 6.3 | 1565 | 27 | 33063662.5 | 12471822.42 | 31323325.02 | 25512603.24 | 29774433.54 | 36830046.17 | 25019903.44 | 26197048.55 | 36.4326 | 18.0474 | Ppfia1 | 233977 | 0.92983 | 1.070165852 | 1.15092 | 0.49875 | 0.877125 |
| 148679188 | copine II, isoform CRA_a, partial [Mus musculus] | 0 | 15.462 | 10 | 6 | 8 | 5 | 565 | 62.8 | 5.96 | 25 | 6 | 3864990.086 | 4535324.366 | 7925743.38 | 7018931.266 | 7562919.041 | 7177735.884 | 3505507.5 | 5456714.486 | 33.3173 | 31.3077 | Cpne2 | 234577 | 0.99239 | 1.007606887 | 1.01533 | 0.94908 | 0.994906 |
| 568956537 | PREDICTED: CCR4-NOT transcription complex subunit 1 isoform X1 [Mus musculus] | 0 | 71.547 | 9 | 19 | 49 | 19 | 2383 | 267.6 | 7.11 | 400 | 19 | 20738297.6 | 14824577.32 | 24607918.85 | 15876324.32 | 22646363 | 21339127.39 | 22645834.38 | 19782740.34 | 23.8464 | 6.3011 | Cnot1 | 234594 | 0.93619 | 1.06381184 | 1.13632 | 0.3155 | 0.766307 |
| 148679269 | dynein, cytoplasmic 1 light intermediate chain 2, isoform CRA_a, partial [Mus musculus] | 0 | 22.416 | 9 | 4 | 18 | 3 | 527 | 57.6 | 7.15 | 160 | 4 | 7724510.45 | 5723715.891 | 7773262.951 | 5756063.142 | 6919275.865 | 7017033.871 | 7547698.75 | 7330710.841 | 17.2016 | 4.009 | Dync1li2 | 234663 | 0.96707 | 1.032928698 | 1.0681 | 0.47144 | 0.863085 |
| 568956599 | PREDICTED: enhancer of mRNA-decapping protein 4 isoform X1 [Mus musculus] | 0 | 95.811 | 20 | 17 | 59 | 17 | 1408 | 152.5 | 5.82 | 492 | 17 | 30721402.66 | 25397390.13 | 38300982.09 | 37243954.22 | 36857299.76 | 24301931.1 | 37510480.38 | 42497945.28 | 18.3187 | 21.9531 | Edc4 | 234699 | 0.96517 | 1.034834436 | 1.07218 | 0.64553 | 0.915705 |
| 299890868 | ATP-dependent RNA helicase DDX19B isoform 1 [Mus musculus] | 0.001 | 6.594 | 4 | 2 | 9 | 2 | 494 | 55.6 | 8.54 | 74 | 2 | 5256135.333 | 12266827.99 | 9367786.617 | 9646220.497 | 9005116.728 | 9239125.961 | 11605660 | 13161141.51 | 31.7106 | 18.5013 | Ddx19b | 234733 | 0.91861 | 1.081385737 | 1.17719 | 0.39248 | 0.824582 |
| 28972255 | mKIAA0513 protein [Mus musculus] | 0 | 10.469 | 7 | 3 | 15 | 3 | 445 | 50.2 | 5.11 | 70 | 3 | 4348009.647 | 4175285.314 | 4368138.134 | 3991133.612 | 4284363.927 | 3314030.071 | 3296304.625 | 2009862.803 | 4.16506 | 28.9219 | 30548M08f | 234797 | 1.13355 | 0.866452205 | 0.76437 | 0.08105 | 0.642075 |
| 148679813 | nucleoporin 133 [Mus musculus] | 0 | 102.978 | 22 | 17 | 65 | 17 | 1155 | 128.5 | 5.2 | 1298 | 17 | 32101312.53 | 31251560.9 | 40801481.2 | 34227795.1 | 38837861.48 | 35207286.35 | 37856474.31 | 35866309.52 | 12.4944 | 4.58237 | Nup133 | 234865 | 0.9672 | 1.032800207 | 1.06783 | 0.35107 | 0.79732 |
| 109659843 | guanylate cyclase soluble subunit alpha-2 [Mus musculus] | 0 | 70.933 | 25 | 15 | 66 | 15 | 730 | 81.8 | 6.87 | 687 | 15 | 72146285.82 | 74240561.64 | 61163534.34 | 52420913.81 | 57962684.31 | 61298868.42 | 54350224.5 | 37427967.96 | 15.6257 | 20.1059 | Gucy1a2 | 234889 | 1.10389 | 0.896113793 | 0.81178 | 0.14677 | 0.693437 |
| 592743503 | phospholipid phosphatase-related protein type 2 isoform 2 [Mus musculus] | 0 | 36.099 | 18 | 6 | 28 | 6 | 452 | 48.4 | 9.45 | 1121 | 6 | 27654624.23 | 20996364.89 | 26678782.28 | 18707296.05 | 29383408.26 | 19313399.88 | 30773008.25 | 16020056.46 | 18.4766 | 30.633 | Plppr2 | 235044 | 0.99233 | 1.00766543 | 1.01545 | 0.93472 | 0.994906 |
| 28173550 | septin-7 isoform 1 [Mus musculus] | 0 | 176.345 | 57 | 19 | 157 | 18 | 437 | 50.6 | 8.57 | 2470 | 19 | 162690944.4 | 181521870.1 | 147690685.6 | 183694516.1 | 151579023.2 | 193644247.2 | 163965133.4 | 233038125.7 | 10.0627 | 19.5318 | 43715 | 235072 | 0.95301 | 1.046993483 | 1.09862 | 0.43714 | 0.852048 |
| 755528617 | PREDICTED: protein turtle homolog B isoform X6 [Mus musculus] | 0 | 200.408 | 30 | 28 | 184 | 28 | 1366 | 148.9 | 6.76 | 2850 | 28 | 161693646 | 181336075.4 | 206553102.7 | 224881735.3 | 231323650.9 | 283705574 | 211632243.1 | 213502937.2 | 14.3466 | 14.3102 | Igsf9b | 235086 | 0.90336 | 1.096638893 | 1.21395 | 0.10628 | 0.674486 |
| 568959241 | PREDICTED: neurotrimin isoform X2 [Mus musculus] | 0 | 27.986 | 17 | 4 | 19 | 4 | 367 | 40.5 | 7.59 | 639 | 4 | 7905777.276 | 6165447.616 | 3873917.913 | 7374252.408 | 6218938.07 | 4261306.345 | 4874344.875 | 5158155.537 | 28.3091 | 15.9478 | Ntm | 235106 | 1.10488 | 0.895124897 | 0.81016 | 0.26819 | 0.745606 |
| 148693715 | RIKEN cDNA D130054N24, isoform CRA_f, partial [Mus musculus] | 0 | 21.667 | 11 | 5 | 15 | 5 | 723 | 79.2 | 6.8 | 49 | 5 | 6547828.945 | 17118918.56 | 13140226.84 | 9069176.003 | 11399222.5 | 11307585.77 | 11873815.38 | 13454209.95 | 40.489 | 8.28655 | Rnf214 | 235315 | 0.97701 | 1.022986483 | 1.04705 | 0.82776 | 0.968871 |
| 189491684 | dmX-like protein 2 [Mus musculus] | 0 | 106.993 | 11 | 26 | 81 | 24 | 3054 | 340.5 | 6.4 | 377 | 26 | 84540071.3 | 57560426.68 | 90452438.81 | 50144396.91 | 119164129.6 | 76136289.49 | 87203197.5 | 70643229.32 | 28.0249 | 24.5864 | Dmxl2 | 235380 | 0.8892 | 1.110796819 | 1.2492 | 0.27582 | 0.745606 |
| 148528987 | coronin-2B [Mus musculus] | 0 | 114.118 | 43 | 17 | 98 | 17 | 480 | 54.9 | 8.27 | 1529 | 17 | 162242938.2 | 125334348.2 | 157356302.3 | 156718321.7 | 143989794.5 | 110685782.3 | 161695490.8 | 108174413.9 | 11.2359 | 19.9074 | Coro2b | 235431 | 1.06847 | 0.931533824 | 0.87184 | 0.26137 | 0.745606 |
| 568962910 | PREDICTED: dnaJ homolog subfamily C member 13 isoform X1 [Mus musculus] | 0.001 | 5.372 | 1 | 2 | 2 | 2 | 2248 | 254.9 | 6.77 | 0 | 2 | 2776111.331 | 2279645.567 | 6331024.461 | 2551143.483 | 2996123.913 | 3425325.078 | 3950528 | 2216522.455 | 54.772 | 23.2897 | Dnajc13 | 235567 | 1.05087 | 0.949129013 | 0.90318 | 0.7526 | 0.951816 |
| 148689290 | CaM kinase-like vesicle-associated, partial [Mus musculus] | 0 | 119.103 | 38 | 15 | 83 | 15 | 559 | 59.8 | 6 | 1765 | 15 | 94939486.63 | 60980880.2 | 98505940.96 | 61795808.3 | 90495582.33 | 109013465.3 | 89180425.44 | 75021365.68 | 25.8741 | 15.3358 | Camkv | 235604 | 0.93016 | 1.069843243 | 1.15018 | 0.37453 | 0.807546 |
| 22122795 | cytoplasmic dynein 1 light intermediate chain 1 [Mus musculus] | 0 | 117.801 | 49 | 18 | 91 | 17 | 523 | 56.6 | 6.42 | 1652 | 18 | 67577653.12 | 79622845.14 | 95223244.99 | 83537563.75 | 80678394.87 | 91140598.35 | 82954666.88 | 93228992.57 | 13.9886 | 7.03158 | Dync1li1 | 235661 | 0.9673 | 1.032704042 | 1.06762 | 0.42698 | 0.84306 |
| 568941707 | PREDICTED: Alstrom syndrome protein 1 homolog isoform X1 [Mus musculus] | 0 | 11.125 | 1 | 2 | 3 | 2 | 3733 | 413.8 | 6.01 | 36 | 2 | 2502570.855 | 656049.6805 | 2780664.467 | 1207546.717 | 2929397.768 | 389282.7544 | 2478493.75 | 1581950.529 | 57.0244 | 60.7307 | Alms1 | 236266 | 0.98401 | 1.015991586 | 1.0325 | 0.94136 | 0.994906 |
| 148697796 | pyruvate dehydrogenase kinase, isoenzyme 3, isoform CRA_a, partial [Mus musculus] | 0 | 26.357 | 16 | 5 | 23 | 5 | 437 | 50 | 8.69 | 764 | 5 | 12299082.02 | 9717957.159 | 12397081.89 | 7684907.392 | 10613070.99 | 6814758.029 | 12112994.25 | 6351084.763 | 21.5059 | 31.5753 | Pdk3 | 236900 | 1.07959 | 0.920412285 | 0.85256 | 0.42494 | 0.84162 |
| 148682288 | mCG115438, isoform CRA_a [Mus musculus] | 0 | 63.239 | 24 | 12 | 53 | 10 | 532 | 62.5 | 5.72 | 671 | 12 | 60254557.01 | 56945028.83 | 56472318.73 | 47283147.57 | 59481447.69 | 55461583.49 | 54886658.5 | 53422813.52 | 10.0731 | 4.64369 | Arhgef9 | 236915 | 0.99483 | 1.005172022 | 1.0104 | 0.85772 | 0.975398 |
| 569011973 | PREDICTED: oral-facial-digital syndrome 1 protein homolog isoform X1 [Mus musculus] | 0 | 15.097 | 4 | 3 | 6 | 3 | 1040 | 119.9 | 6.09 | 113 | 3 | 3428212.229 | 3397986.949 | 4991795.2 | 2965904.809 | 3430518.226 | 2739466.564 | 3221727.75 | 2563175.572 | 24.0617 | 13.5595 | Ofd1 | 237222 | 1.1058 | 0.894198228 | 0.80864 | 0.19795 | 0.715071 |
| 26325880 | unnamed protein product [Mus musculus] | 0 | 21.378 | 18 | 5 | 28 | 5 | 419 | 44.6 | 9.73 | 167 | 5 | 27471367.71 | 26709390.18 | 31768586.01 | 28107486.23 | 24868065.48 | 22713105.23 | 29224457.19 | 26502908.36 | 7.86837 | 10.6308 | C2cd4c | 237397 | 1.04945 | 0.950551948 | 0.90576 | 0.18039 | 0.715071 |
| 568967519 | PREDICTED: cyclin-dependent kinase 17 isoform X2 [Mus musculus] | 0 | 44.577 | 23 | 11 | 44 | 9 | 539 | 61.4 | 9.01 | 758 | 11 | 26430619.59 | 17927923.49 | 25989773.62 | 29381665.08 | 23964072.57 | 17025114.33 | 26094490.75 | 19931332.67 | 19.6792 | 18.6554 | Cdk17 | 237459 | 1.06809 | 0.931912994 | 0.87251 | 0.35661 | 0.798662 |
| 51243034 | oxysterol-binding protein-related protein 8 isoform a [Mus musculus] | 0.001 | 6.06 | 2 | 2 | 9 | 2 | 889 | 101.2 | 6.96 | 25 | 2 | 3336194.166 | 2033581.408 | 2953423.618 | 2447166.323 | 2751077.978 | 1209120.475 | 2264834.813 | 700778.7348 | 21.1918 | 54.3761 | Osbpl8 | 237542 | 1.21725 | 0.782746669 | 0.64304 | 0.1314 | 0.693437 |
| 148689814 | TRH-degrading enzyme, isoform CRA_b [Mus musculus] | 0.001 | 5.506 | 2 | 2 | 8 | 2 | 1066 | 122.1 | 7.49 | 79 | 2 | 2137095.608 | 2355457.299 | 2810643.231 | 1483973.261 | 2691242.436 | 2007985.615 | 2530520.5 | 2077888.995 | 25.1214 | 14.4234 | Trhde | 237553 | 0.97124 | 1.028763399 | 1.05923 | 0.70099 | 0.931521 |
| 281183137 | sterile alpha and TIR motif-containing protein 1 isoform 1 [Mus musculus] | 0 | 19.473 | 6 | 4 | 20 | 4 | 764 | 84 | 6.64 | 45 | 4 | 8854236.048 | 6682750.538 | 7283686.85 | 5136244.017 | 9102458.174 | 8778417.116 | 8204169.75 | 6973785.677 | 21.9993 | 11.3416 | Sarm1 | 237868 | 0.91638 | 1.083616335 | 1.18249 | 0.20637 | 0.715071 |
| 40254254 | protein sidekick-2 precursor [Mus musculus] | 0 | 19.471 | 3 | 6 | 15 | 6 | 2176 | 239.8 | 7.36 | 102 | 6 | 9792338.08 | 17483728.84 | 14255012.39 | 12021611.02 | 18300895.09 | 24210107.29 | 15046112.5 | 24732128.76 | 24.5172 | 22.8404 | Sdk2 | 237979 | 0.78846 | 1.211544053 | 1.5366 | 0.0461 | 0.591577 |
| 62543571 | dedicator of cytokinesis protein 4 [Mus musculus] | 0 | 416.605 | 43 | 60 | 332 | 59 | 1978 | 226.4 | 7.65 | 6182 | 60 | 238782118.6 | 184803222.7 | 251040650 | 199330736 | 253214888.3 | 234084626.3 | 252868136.8 | 198331471.7 | 14.4086 | 10.9935 | Dock4 | 238130 | 0.96439 | 1.035610465 | 1.07385 | 0.45802 | 0.860877 |
| 148704518 | mCG52906 [Mus musculus] | 0 | 72.239 | 30 | 9 | 40 | 9 | 721 | 76.9 | 4.79 | 943 | 9 | 27447558.09 | 29417803.8 | 37172226.59 | 29705992.81 | 28886798.98 | 40761777.88 | 35364345.88 | 29787609.05 | 13.8253 | 16.353 | Akap5 | 238276 | 0.95723 | 1.042766205 | 1.08935 | 0.45824 | 0.860877 |
| 568984455 | PREDICTED: cAMP-specific 3',5'-cyclic phosphodiesterase 4D isoform X1 [Mus musculus] | 0 | 116.536 | 35 | 21 | 118 | 14 | 803 | 90.7 | 5.49 | 1325 | 21 | 55043531.77 | 29154250.83 | 48728332.14 | 27240122.06 | 52432209.59 | 38561795.4 | 46716029.75 | 25066320.37 | 34.8126 | 29.1734 | Pde4d | 238871 | 0.99192 | 1.008082298 | 1.0163 | 0.94549 | 0.994906 |
| 30425250 | beta-actin-like protein 2 [Mus musculus] | 0 | 123.331 | 44 | 13 | 592 | 4 | 376 | 42 | 5.49 | 11106 | 13 | 1831172.232 | 3882536.036 | 5464345.834 | 7470142.756 | 10373678.28 | 12579652.66 | 6615505.5 | 9291151.71 | 51.2864 | 25.5105 | Actbl2 | 238880 | 0.64854 | 1.351459384 | 2.08385 | 0.02613 | 0.515682 |
| 568987548 | PREDICTED: ERC protein 2 isoform X21 [Mus musculus] | 0 | 462.144 | 56 | 57 | 800 | 2 | 727 | 84.4 | 7.5 | 13628 | 57 | 2778008.548 | 3543994.896 | 4287511.64 | 6602061.285 | 2818734.408 | 4677363.847 | 3022801 | 5461211.13 | 38.3935 | 32.1341 | Erc2 | 238988 | 1.0371 | 0.962898361 | 0.92845 | 0.77844 | 0.952792 |
| 123701966 | BTB/POZ domain-containing protein KCTD12 [Mus musculus] | 0 | 52.703 | 36 | 10 | 51 | 7 | 327 | 35.9 | 5.81 | 886 | 10 | 88721019.69 | 77731634.75 | 115561571 | 86035244.13 | 114187618.1 | 112462411.5 | 116620119.8 | 89471345.46 | 17.8036 | 11.6393 | Kctd12 | 239217 | 0.91921 | 1.080785159 | 1.17577 | 0.16851 | 0.711915 |
| 66792511 | Eif2c2 protein, partial [Mus musculus] | 0 | 45.726 | 12 | 8 | 30 | 5 | 883 | 99.3 | 9.29 | 486 | 8 | 2980457.465 | 1038516.2 | 5549718.21 | 6642871.885 | 2837945.183 | 391086.8559 | 3879003.313 | 392148.0818 | 62.3959 | 94.1233 | Ago2 | 239528 | 1.36739 | 0.63261331 | 0.46264 | 0.20753 | 0.715071 |
| 755551662 | PREDICTED: disco-interacting protein 2 homolog B isoform X2 [Mus musculus] | 0 | 150.218 | 25 | 27 | 96 | 26 | 1618 | 176 | 8.27 | 1492 | 27 | 82672150.46 | 69338293.33 | 93844973.61 | 72696458.2 | 79723778.1 | 117831508.9 | 97892368.94 | 95386995.26 | 13.8564 | 16.0066 | Dip2b | 239667 | 0.89811 | 1.101894768 | 1.22691 | 0.10792 | 0.678142 |
| 568995287 | PREDICTED: MKL/myocardin-like protein 2 isoform X1 [Mus musculus] | 0 | 20.329 | 7 | 5 | 16 | 5 | 1112 | 120.5 | 6.04 | 162 | 5 | 3969289.78 | 2301513.008 | 6901965.213 | 1302110.506 | 4929233.795 | 2980336.051 | 5510951.5 | 3361772.547 | 67.6975 | 28.9977 | Mrtfb | 239719 | 0.92618 | 1.073820349 | 1.15941 | 0.68787 | 0.928842 |
| 86262157 | protein MB21D2 [Mus musculus] | 0 | 30.433 | 21 | 6 | 15 | 6 | 491 | 55.6 | 7.5 | 148 | 6 | 11843782.52 | 7522302.03 | 12299234.81 | 10361218.11 | 11947786.41 | 6102966.068 | 11114641.34 | 6390386.929 | 20.5084 | 34.5617 | Mb21d2 | 239796 | 1.08341 | 0.916594955 | 0.84603 | 0.4217 | 0.840085 |
| 569000237 | PREDICTED: copine-5 isoform X1 [Mus musculus] | 0 | 29.608 | 15 | 8 | 18 | 2 | 610 | 67.3 | 5.69 | 112 | 8 | 4501497.467 | 914958.9101 | 752680.9359 | 3233388.907 | 3410881.447 | 2998570.808 | 3329209.063 | 4314541.455 | 77.7482 | 16.0282 | Cpne5 | 240058 | 0.80173 | 1.198274654 | 1.49462 | 0.26967 | 0.745606 |
| 34147224 | fibronectin type III and SPRY domain-containing protein 1 [Mus musculus] | 0 | 23.328 | 11 | 3 | 6 | 3 | 496 | 55.5 | 6.67 | 140 | 3 | 1650354.094 | 3133775.899 | 2108075.654 | 2768662.972 | 2553687.588 | 3095257.915 | 2382599 | 3508572.593 | 27.4707 | 17.8457 | Fsd1 | 240121 | 0.91136 | 1.088639675 | 1.19452 | 0.30598 | 0.761857 |
| 126506316 | junctional protein associated with coronary artery disease [Mus musculus] | 0 | 317.902 | 48 | 42 | 292 | 42 | 1320 | 144.7 | 6.52 | 4682 | 42 | 360309863.1 | 336175441.6 | 408187114.4 | 347861127.1 | 395247871.4 | 433432746.2 | 344195737.2 | 399242680.4 | 8.70509 | 9.36081 | Jcad | 240185 | 0.96046 | 1.039536934 | 1.08233 | 0.2638 | 0.745606 |
| 244793002 | probable ATP-dependent RNA helicase YTHDC2 [Mus musculus] | 0 | 24.208 | 4 | 4 | 8 | 4 | 1445 | 161 | 8.51 | 63 | 4 | 4820037.511 | 6736918.474 | 4320737.536 | 4859385.599 | 5354144.981 | 5121041.939 | 5384157.938 | 4209063.745 | 20.5186 | 10.99 | Ythdc2 | 240255 | 1.01639 | 0.983613221 | 0.96775 | 0.78958 | 0.953707 |
| 38348478 | coiled-coil domain-containing protein 85B isoform 1 [Mus musculus] | 0 | 7.388 | 14 | 3 | 13 | 3 | 202 | 22.1 | 5.08 | 66 | 3 | 5986573.293 | 5937395.618 | 6357291.341 | 5469078.557 | 7086267.065 | 6637123.629 | 6912836.75 | 6848605.129 | 6.13478 | 2.70044 | Ccdc85b | 240514 | 0.92711 | 1.072889261 | 1.15724 | 0.00382 | 0.38143 |
| 1039728814 | PREDICTED: pleckstrin homology domain-containing family A member 6 isoform X4 [Mus musculus] | 0 | 228.919 | 38 | 38 | 292 | 2 | 1276 | 143.1 | 8.51 | 4737 | 38 | 18672941.93 | 16050467.88 | 25865231.25 | 14505493.05 | 19526014.6 | 22231495.68 | 25771343.25 | 22435646.98 | 26.7985 | 11.3711 | Plekha6 | 240753 | 0.90991 | 1.090091418 | 1.19802 | 0.23577 | 0.73104 |
| 28972830 | mKIAA1747 protein [Mus musculus] | 0 | 9.419 | 4 | 3 | 8 | 3 | 788 | 89.7 | 7.96 | 56 | 3 | 1382498.81 | 2966095.484 | 3312140.747 | 2056456.805 | 3759046.326 | 3598767.364 | 2948282.375 | 2841278.286 | 36.0616 | 13.98 | Brinp2 | 240843 | 0.84998 | 1.150021762 | 1.353 | 0.13366 | 0.693437 |
| 254028203 | GPI inositol-deacylase [Mus musculus] | 0 | 46.03 | 12 | 9 | 32 | 9 | 922 | 104.5 | 8.91 | 347 | 9 | 29664493.07 | 15817237.57 | 37406622.84 | 19280133.48 | 37439646.46 | 22966472.32 | 44681022.81 | 21809408.21 | 38.5954 | 35.268 | Pgap1 | 241062 | 0.89205 | 1.107952148 | 1.24203 | 0.43876 | 0.852048 |
| 33942102 | acid-sensing ion channel 4 [Mus musculus] | 0 | 8.463 | 6 | 2 | 2 | 2 | 539 | 59.2 | 6.79 | 22 | 2 | 5507394.402 | 5530574.212 | 1763443.218 | 1010171.344 | 1912103.378 | 1186184.769 | 2421106.75 | 978440.2581 | 69.6653 | 40.9388 | Asic4 | 241118 | 1.36012 | 0.639883925 | 0.47046 | 0.1932 | 0.715071 |
| 568907669 | PREDICTED: neuronal tyrosine-phosphorylated phosphoinositide-3-kinase adapter 2 isoform X1 [Mus musculus] | 0 | 30.083 | 16 | 7 | 18 | 7 | 748 | 81.6 | 8.88 | 272 | 7 | 13706245.95 | 8000086.623 | 16419897.82 | 22308951.53 | 14719502.44 | 11528875.42 | 9990123.875 | 10570930.09 | 39.3541 | 18.023 | Nyap2 | 241134 | 1.12705 | 0.872946998 | 0.77454 | 0.32168 | 0.771047 |
| 27370290 | cadherin-7 isoform 1 preproprotein [Mus musculus] | 0 | 35.46 | 16 | 8 | 19 | 8 | 785 | 87.1 | 4.82 | 120 | 8 | 22954138.09 | 20862717.1 | 20471897.88 | 13740867.36 | 17516164.83 | 7438304.631 | 19615298 | 12877493.96 | 20.4837 | 37.6442 | Cdh7 | 241201 | 1.15193 | 0.848074751 | 0.73622 | 0.1767 | 0.713031 |
| 52350563 | probable G-protein coupled receptor 158 precursor [Mus musculus] | 0 | 95.686 | 22 | 19 | 72 | 19 | 1200 | 134.3 | 8.09 | 1090 | 19 | 51814693.39 | 65288591.66 | 90878728.76 | 72478555.87 | 84675701.86 | 87887100.23 | 80299160.53 | 82783823.48 | 23.216 | 3.81293 | Gpr158 | 241263 | 0.91043 | 1.089570925 | 1.19677 | 0.14732 | 0.693437 |
| 29244192 | volume-regulated anion channel subunit LRRC8A [Mus musculus] | 0 | 43.486 | 16 | 11 | 33 | 10 | 810 | 94.1 | 7.94 | 352 | 11 | 18514508.69 | 15583494.94 | 24885972.3 | 16420265.38 | 25034893.45 | 37581993.67 | 20748129.13 | 26128701.57 | 22.3218 | 26.2697 | Lrrc8a | 241296 | 0.81563 | 1.184369133 | 1.45209 | 0.08675 | 0.655499 |
| 157820475 | leucine-rich repeat-containing protein 4C precursor [Rattus norvegicus] | 0 | 16.19 | 8 | 5 | 19 | 3 | 640 | 71.9 | 7.15 | 127 | 5 | 9409645.772 | 8377613.982 | 9317426.295 | 7784323.793 | 9485835.03 | 11267083.5 | 9393065.5 | 10172747.93 | 8.94271 | 8.5778 | Lrrc4c | 241568 | 0.9278 | 1.072196319 | 1.15563 | 0.05853 | 0.599194 |
| 755499611 | PREDICTED: UPF0606 protein KIAA1549L homolog isoform X1 [Mus musculus] | 0 | 178.741 | 14 | 18 | 144 | 18 | 2147 | 230.8 | 9.29 | 3577 | 18 | 235291489.4 | 195162960.7 | 243312079.5 | 139799825.7 | 248871410.2 | 233132472.6 | 211773541.4 | 166175168.5 | 23.2747 | 16.707 | 30041D05I | 241589 | 0.97228 | 1.027717784 | 1.05702 | 0.70979 | 0.935294 |
| 161086990 | leucine zipper putative tumor suppressor 3 isoform a [Mus musculus] | 0 | 109.005 | 38 | 19 | 165 | 19 | 700 | 74.9 | 6.99 | 2063 | 19 | 140488383.1 | 152420908.8 | 150841715.6 | 139503501 | 132598189.8 | 136967333.4 | 145769985 | 130123782.9 | 4.63641 | 5.04449 | Lzts3 | 241638 | 1.03349 | 0.966514791 | 0.9352 | 0.09778 | 0.670764 |
| 568917008 | PREDICTED: serine/threonine-protein kinase PAK 7 isoform X1 [Mus musculus] | 0 | 142 | 38 | 24 | 154 | 22 | 736 | 82.9 | 8.21 | 3094 | 24 | 169618339.6 | 164801764.3 | 211736868.3 | 158817580.4 | 174851448.7 | 173641569.7 | 189234332.9 | 174483623 | 13.6578 | 4.19627 | Pak7 | 241656 | 0.99489 | 1.005106192 | 1.01026 | 0.89055 | 0.984401 |
| 568917014 | PREDICTED: double zinc ribbon and ankyrin repeat-containing protein 1 isoform X1 [Mus musculus] | 0 | 127.169 | 35 | 21 | 123 | 21 | 792 | 86.4 | 8.18 | 1487 | 21 | 79439781.58 | 83907122.89 | 87975893.36 | 85904239.73 | 81735186.1 | 78254109.86 | 78079295.72 | 89316415.73 | 4.32369 | 6.42306 | Dzank1 | 241688 | 1.01481 | 0.985191315 | 0.97081 | 0.47093 | 0.863085 |
| 755499655 | PREDICTED: syntaphilin isoform X1 [Mus musculus] | 0 | 121.725 | 48 | 18 | 118 | 18 | 572 | 61.6 | 6.42 | 1566 | 18 | 96984537.71 | 116904186.1 | 73404115.62 | 113330627.7 | 105086102.5 | 120471652.6 | 99051484.13 | 87731252.42 | 19.8 | 13.2337 | Snph | 241727 | 0.98559 | 1.014412723 | 1.02925 | 0.81581 | 0.963141 |
| 33942097 | regulating synaptic membrane exocytosis protein 4 [Mus musculus] | 0 | 41.279 | 32 | 6 | 29 | 3 | 269 | 29.3 | 5.8 | 895 | 6 | 9637662.96 | 6358857.997 | 7001680.156 | 4804070.829 | 6391925.418 | 5771522.434 | 2973315.25 | 4696343.565 | 28.9904 | 30.1973 | Rims4 | 241770 | 1.1673 | 0.832704903 | 0.71336 | 0.16356 | 0.706428 |
| 148673748 | RIKEN cDNA 1700012H17, isoform CRA_c, partial [Mus musculus] | 0 | 22.584 | 11 | 3 | 21 | 3 | 405 | 44.3 | 8.95 | 348 | 3 | 4638908.134 | 1648831.049 | 3412159.967 | 4425544.842 | 1220872.974 | 1330723.208 | 4937518.688 | 2483747.193 | 38.6352 | 69.2578 | Fam110b | 242297 | 1.17232 | 0.827681584 | 0.70602 | 0.38186 | 0.816034 |
| 124487263 | gamma-aminobutyric acid type B receptor subunit 2 precursor [Mus musculus] | 0 | 143.069 | 31 | 20 | 276 | 20 | 940 | 105.6 | 8.72 | 4237 | 20 | 159580167.2 | 203996793.8 | 174765423.7 | 195987203.2 | 185331230 | 236999810 | 181834617.5 | 205509355.3 | 11.0048 | 12.5015 | Gabbr2 | 242425 | 0.9512 | 1.048798705 | 1.1026 | 0.28883 | 0.752733 |
| 27777697 | paralemmin-2 [Mus musculus] | 0 | 43.336 | 22 | 6 | 25 | 3 | 376 | 42.1 | 5.15 | 446 | 6 | 6353864.709 | 12669154.56 | 12424161.7 | 8437123.362 | 10932562.68 | 13789814.86 | 11014056.5 | 17229389.19 | 31.0384 | 22.4439 | Palm2 | 242481 | 0.85911 | 1.14088854 | 1.32799 | 0.17825 | 0.713031 |
| 52345394 | actin-related protein 3B isoform 1 [Mus musculus] | 0 | 129.68 | 53 | 16 | 120 | 13 | 418 | 47.5 | 6.02 | 2042 | 16 | 145254011.8 | 121545617.7 | 144540987.7 | 119878287.6 | 148436316.9 | 142131772.2 | 143293798.1 | 144301388.2 | 10.5291 | 1.89852 | Actr3b | 242894 | 0.95768 | 1.042315779 | 1.08837 | 0.15062 | 0.693437 |
| 26335191 | unnamed protein product [Mus musculus] | 0 | 35.677 | 26 | 10 | 34 | 8 | 477 | 52.9 | 8.5 | 167 | 10 | 34268273.14 | 23862597.69 | 25432853.54 | 26936484.27 | 22481263.48 | 28267893.83 | 23797276.13 | 20783496.36 | 16.6631 | 13.4437 | Kctd8 | 243043 | 1.0737 | 0.926297096 | 0.86271 | 0.22498 | 0.718033 |
| 568935627 | PREDICTED: uncharacterized protein KIAA1671 isoform X1 [Mus musculus] | 0 | 9.963 | 2 | 3 | 7 | 3 | 1767 | 192.6 | 9.04 | 58 | 3 | 1773901.197 | 4498648.958 | 7279023.387 | 7515968.928 | 5589633.19 | 3957000.59 | 6875749.5 | 1858575.619 | 51.2967 | 47.4088 | 00026A02F | 243219 | 1.07082 | 0.929181964 | 0.86773 | 0.7014 | 0.931521 |
| 148687294 | RIKEN cDNA 6430598A04, isoform CRA_a, partial [Mus musculus] | 0 | 45.967 | 17 | 9 | 34 | 9 | 845 | 88.8 | 9.44 | 656 | 9 | 24558882.95 | 28734594.38 | 27167765.39 | 29243263.36 | 30666648.74 | 33435283.64 | 25528892.25 | 32852276.89 | 7.67771 | 11.7492 | Nyap1 | 243300 | 0.94496 | 1.055035648 | 1.11648 | 0.17624 | 0.713031 |
| 30425288 | protein ELFN1 precursor [Mus musculus] | 0 | 114.691 | 35 | 19 | 66 | 16 | 828 | 90.8 | 8.18 | 836 | 19 | 41410152.2 | 44294686.93 | 41103641.74 | 48519919.83 | 46386589.28 | 57042118.26 | 47428608.69 | 35879149.77 | 7.84804 | 18.5375 | Elfn1 | 243312 | 0.96849 | 1.031508347 | 1.06507 | 0.56268 | 0.895889 |
| 197927164 | leucine-rich repeat transmembrane neuronal protein 4 isoform 3 precursor [Mus musculus] | 0 | 95.68 | 35 | 19 | 167 | 15 | 591 | 67.2 | 9 | 2133 | 19 | 149099666.7 | 161285908 | 150954450 | 184673044.3 | 150673008 | 203793886.5 | 156253975.8 | 175827254.2 | 10.1239 | 13.982 | Lrrtm4 | 243499 | 0.96958 | 1.030418907 | 1.06275 | 0.51134 | 0.88031 |
| 38348566 | H1 histone family, member X [Mus musculus] | 0 | 9.89 | 12 | 2 | 10 | 2 | 188 | 20.1 | 11.22 | 219 | 2 | 14205319 | 13082365.58 | 17135804.97 | 12514934.83 | 17768431.92 | 16488665.97 | 17243024 | 9679263.115 | 14.4558 | 24.7167 | H1fx | 243529 | 0.9641 | 1.035904498 | 1.07448 | 0.63976 | 0.915433 |
| 568941827 | PREDICTED: glutamate receptor-interacting protein 2 isoform X2 [Mus musculus] | 0 | 75.168 | 20 | 15 | 57 | 14 | 1049 | 112.8 | 6.64 | 384 | 15 | 37073803.8 | 28024836.07 | 37969388.52 | 44457385.09 | 50884454.04 | 46404976.05 | 39078505.25 | 39179724.97 | 18.3265 | 13.1938 | Grip2 | 243547 | 0.91326 | 1.086736559 | 1.18995 | 0.1665 | 0.711253 |
| 568941833 | PREDICTED: prickle-like protein 2 isoform X1 [Mus musculus] | 0 | 421.396 | 57 | 37 | 686 | 37 | 937 | 105.8 | 7.9 | 12446 | 37 | 649519813.2 | 771734037.9 | 702162437.3 | 746379997.1 | 752982582.3 | 896033191.1 | 772920121.8 | 764155687.8 | 7.47656 | 8.39148 | Prickle2 | 243548 | 0.94777 | 1.052229385 | 1.11022 | 0.11453 | 0.678142 |
| 84781690 | IQ motif and SEC7 domain-containing protein 3 isoform 1 [Mus musculus] | 0 | 206.165 | 35 | 33 | 229 | 32 | 1195 | 129 | 6.19 | 2644 | 33 | 251100592.9 | 206725124.7 | 264467307 | 232549839.6 | 287282537.1 | 239105404.7 | 244271188.1 | 198952387 | 10.4812 | 14.9056 | Iqsec3 | 243621 | 0.99233 | 1.007674203 | 1.01547 | 0.87209 | 0.979509 |
| 568939590 | PREDICTED: neurabin-1 isoform X1 [Mus musculus] | 0 | 323.459 | 44 | 38 | 415 | 37 | 1322 | 147.6 | 5.49 | 8300 | 38 | 563354826.4 | 500868923.9 | 510543830.2 | 531691383.8 | 486505687.4 | 459339615.9 | 458499368.5 | 462547803.9 | 5.25413 | 2.85035 | Ppp1r9a | 243725 | 1.06029 | 0.939706695 | 0.88627 | 0.00797 | 0.4058 |
| 171543899 | plexin-A4 precursor [Mus musculus] | 0 | 9.745 | 2 | 3 | 4 | 2 | 1893 | 212.4 | 6.83 | 44 | 3 | 1157805.271 | 919176.0294 | 1302324.044 | 800161.6911 | 1450609.353 | 1133417.561 | 1223013.75 | 859208.0584 | 21.7345 | 20.9645 | Plxna4 | 243743 | 0.94497 | 1.05503022 | 1.11647 | 0.49331 | 0.871733 |
| 225543212 | disks large-associated protein 2 isoform 2 [Mus musculus] | 0 | 406.356 | 49 | 37 | 599 | 2 | 1060 | 119.2 | 6.9 | 9659 | 37 | 7981158.328 | 10106742.65 | 6501603.204 | 11778917.23 | 13691892.57 | 16084244.38 | 8659618.5 | 16825804.44 | 25.5531 | 26.696 | Dlgap2 | 244310 | 0.79381 | 1.206189483 | 1.51949 | 0.07335 | 0.621937 |
| 148703516 | DNA segment, Chr 8, ERATO Doi 82, expressed, isoform CRA_a [Mus musculus] | 0 | 72.882 | 18 | 18 | 43 | 18 | 1374 | 148.1 | 7.36 | 101 | 18 | 36267388.57 | 25782198.28 | 36606723.21 | 26881153.12 | 31665605.25 | 19628631.74 | 26726198.13 | 21011647.33 | 18.6501 | 22.3595 | Prag1 | 244418 | 1.11803 | 0.881972506 | 0.78886 | 0.15107 | 0.693437 |
| 124487493 | signal-induced proliferation-associated 1-like protein 2 [Mus musculus] | 0 | 248.508 | 33 | 37 | 161 | 36 | 1722 | 189.3 | 6.79 | 2888 | 37 | 119588260.7 | 89274296.92 | 135786782.2 | 104320259 | 132415804.3 | 94001438.39 | 115590670.6 | 95066795.76 | 17.8081 | 16.7966 | Sipa1l2 | 244668 | 1.01342 | 0.986575288 | 0.97351 | 0.83378 | 0.968992 |
| 568959424 | PREDICTED: noelin-2 isoform X1 [Mus musculus] | 0 | 39.566 | 21 | 9 | 41 | 9 | 478 | 53.8 | 8.37 | 655 | 9 | 108737210.7 | 118752810.6 | 101513606.8 | 116740411.6 | 121833956.4 | 137491535.1 | 105557650.3 | 124569262.5 | 7.09267 | 10.7226 | Olfm2 | 244723 | 0.95326 | 1.046737094 | 1.09806 | 0.20355 | 0.715071 |
| 148693542 | expressed sequence AW551984, isoform CRA_a, partial [Mus musculus] | 0.001 | 5.67 | 3 | 2 | 6 | 2 | 840 | 93.3 | 7.08 | 64 | 2 | 1185477.595 | 581045.189 | 947265.003 | 667428.2615 | 1117204.162 | 395516.0886 | 710196.9375 | 653076.6909 | 32.5837 | 41.5413 | AW551984 | 244810 | 1.08074 | 0.919257596 | 0.85058 | 0.55698 | 0.895143 |
| 124486797 | S phase cyclin A-associated protein in the endoplasmic reticulum [Mus musculus] | 0 | 9.841 | 3 | 3 | 6 | 3 | 1398 | 157.7 | 7.31 | 20 | 3 | 3659905.068 | 2405070.978 | 4319148.488 | 3545452.927 | 4099300.61 | 4355165.951 | 4678834.375 | 3727618.432 | 22.8303 | 9.54573 | Scaper | 244891 | 0.9048 | 1.095202813 | 1.21044 | 0.1511 | 0.693437 |
| 157909795 | pseudopodium-enriched atypical kinase 1 [Mus musculus] | 0 | 143.992 | 24 | 27 | 85 | 27 | 1735 | 191 | 6.87 | 1381 | 27 | 81404617.93 | 63014409.43 | 78268282.55 | 72419229.06 | 74430147.07 | 59474260.36 | 68030083.47 | 54446724.24 | 10.9567 | 13.8596 | Peak1 | 244895 | 1.07022 | 0.929780265 | 0.86878 | 0.15806 | 0.700461 |
| 148677228 | myosin VIIA and Rab interacting protein, partial [Mus musculus] | 0 | 16.276 | 6 | 4 | 7 | 4 | 865 | 95.9 | 6.07 | 71 | 4 | 10018581.05 | 14355334.66 | 12273435.81 | 14062152.87 | 12609427.19 | 12895207.21 | 11590355.63 | 15410156 | 15.7534 | 12.3593 | Myrip | 245049 | 0.9826 | 1.017397159 | 1.03541 | 0.73906 | 0.943908 |
| 148710252 | SLIT and NTRK-like family, member 4, isoform CRA_b [Mus musculus] | 0 | 19.479 | 7 | 5 | 10 | 5 | 929 | 104.7 | 7.56 | 58 | 5 | 7657213.122 | 1663071.806 | 6155368.303 | 1655410.899 | 1797578.104 | 4741801.536 | 947411.7188 | 4099840.188 | 72.1686 | 62.5857 | Slitrk4 | 245446 | 1.19307 | 0.806933236 | 0.67635 | 0.46851 | 0.863085 |
| 50510949 | mKIAA1480 protein [Mus musculus] | 0 | 160.795 | 33 | 21 | 143 | 18 | 876 | 96.5 | 6.2 | 2376 | 21 | 305948442.7 | 293949703 | 280790901.8 | 298665345.4 | 283250375.7 | 354546033.7 | 255448537.2 | 265312282.6 | 3.59049 | 15.459 | Nlgn3 | 245537 | 1.0089 | 0.991104384 | 0.98237 | 0.8287 | 0.968871 |
| 244793066 | G-protein coupled receptor-associated sorting protein 2 [Mus musculus] | 0 | 18.766 | 7 | 4 | 9 | 4 | 826 | 92.7 | 5.11 | 66 | 4 | 1978915.842 | 4823840.582 | 4820031.915 | 6715244.551 | 5957258.644 | 5132354.532 | 5319523.25 | 6756527.1 | 42.5986 | 12.6728 | Gprasp2 | 245607 | 0.88368 | 1.116318089 | 1.26326 | 0.29126 | 0.754456 |
| 1004126094 | FERM and PDZ domain-containing protein 3 isoform 1 [Mus musculus] | 0 | 203.278 | 27 | 33 | 187 | 33 | 1774 | 195.2 | 8.02 | 3533 | 33 | 166811925.2 | 172043937.5 | 155857503.1 | 175948535.5 | 190501240.9 | 186103280.3 | 172047317.3 | 206301203.9 | 5.1988 | 7.47483 | Frmpd3 | 245643 | 0.94087 | 1.059126163 | 1.12568 | 0.044 | 0.588297 |
| 29244248 | connector enhancer of kinase suppressor of ras 2 isoform 1 [Mus musculus] | 0 | 426.147 | 62 | 48 | 475 | 4 | 1032 | 117.3 | 6.79 | 9988 | 48 | 44237473.37 | 39195017.85 | 37086877.89 | 34810944.13 | 32842992.25 | 33648444.63 | 30776833.75 | 26368627.94 | 10.3611 | 10.5461 | Cnksr2 | 245684 | 1.11361 | 0.88639018 | 0.79596 | 0.02222 | 0.505785 |
| 71725377 | protocadherin Fat 2 precursor [Mus musculus] | 0 | 118.003 | 8 | 26 | 93 | 26 | 4351 | 479.8 | 5.12 | 762 | 26 | 58108383.41 | 51363424.96 | 41283150.55 | 40950298.87 | 47612546.9 | 68099638.06 | 48004362.69 | 41859181.03 | 17.3858 | 22.3488 | Fat2 | 245827 | 0.96509 | 1.034913503 | 1.07235 | 0.64237 | 0.915705 |
| 66730565 | trafficking protein particle complex subunit 1 [Mus musculus] | 0.004 | 3.166 | 12 | 2 | 6 | 2 | 145 | 16.9 | 9.16 | 0 | 2 | 4686875.916 | 2847495.756 | 6164843.135 | 2986985.866 | 4393004.969 | 2281582.263 | 3756987.625 | 2302569.413 | 37.6369 | 33.3481 | Trappc1 | 245828 | 1.13433 | 0.86566927 | 0.76315 | 0.33731 | 0.782483 |
| 148698349 | arginine/proline rich coiled-coil 1, isoform CRA_b, partial [Mus musculus] | 0 | 38.391 | 14 | 8 | 35 | 8 | 900 | 98.4 | 10.23 | 260 | 8 | 15026694 | 21231121.83 | 16457397.14 | 16257373.15 | 16973612.05 | 10364960.48 | 11729878.75 | 9350043.965 | 15.849 | 27.9999 | Map7d1 | 245877 | 1.17509 | 0.824909264 | 0.702 | 0.05625 | 0.599194 |
| 21553113 | wiskott-Aldrich syndrome protein family member 3 [Mus musculus] | 0 | 71.061 | 24 | 8 | 28 | 8 | 501 | 55.2 | 6.37 | 686 | 8 | 48139925.73 | 38148367.37 | 38847124.77 | 35617273.12 | 45399739.47 | 29227025 | 48661858.5 | 28770493.52 | 13.6352 | 27.6137 | Wasf3 | 245880 | 1.02779 | 0.972208292 | 0.94592 | 0.72616 | 0.940505 |
| 28972872 | mKIAA1916 protein [Mus musculus] | 0 | 23.697 | 9 | 5 | 16 | 5 | 588 | 66.1 | 7.27 | 328 | 5 | 6666825.12 | 3530624.458 | 13641906.72 | 9506177.133 | 11388468.98 | 4040443.801 | 9328341.25 | 5858608.928 | 51.5484 | 43.3524 | Lgi2 | 246316 | 1.04268 | 0.957323157 | 0.91814 | 0.8099 | 0.961921 |
| 251823964 | neuropilin and tolloid-like protein 1 precursor [Mus musculus] | 0 | 108.624 | 41 | 16 | 104 | 15 | 533 | 60.2 | 6.99 | 1166 | 16 | 66290852.41 | 61413949.18 | 81627752.97 | 66128366.32 | 86975592.31 | 93563598.13 | 79072038.03 | 73296098.82 | 12.7841 | 10.6746 | Neto1 | 246317 | 0.90557 | 1.094427029 | 1.20855 | 0.06139 | 0.601634 |
| 568953324 | PREDICTED: gamma-tubulin complex component 3 isoform X1 [Mus musculus] | 0 | 33.773 | 11 | 8 | 15 | 8 | 930 | 106.1 | 8.29 | 77 | 8 | 10141261.72 | 45674899.56 | 113710241.8 | 123501247.6 | 130456705 | 76523061.45 | 132969183.5 | 133751407.9 | 74.373 | 23.6183 | Tubgcp3 | 259279 | 0.76436 | 1.235641199 | 1.61657 | 0.19065 | 0.715071 |
| 568941911 | PREDICTED: SLIT-ROBO Rho GTPase-activating protein 3 isoform X1 [Mus musculus] | 0 | 115.05 | 24 | 20 | 97 | 18 | 1104 | 125 | 6.71 | 1393 | 20 | 39611513.14 | 31936829.76 | 44525877.4 | 52674755.41 | 42763661.7 | 49330580.94 | 41785566.56 | 36519241.82 | 20.6265 | 12.3448 | Srgap3 | 259302 | 0.99513 | 1.004865354 | 1.00978 | 0.93797 | 0.994906 |
| 755556707 | PREDICTED: proline-rich transmembrane protein 1 isoform X1 [Mus musculus] | 0 | 26.846 | 20 | 4 | 48 | 4 | 343 | 35.3 | 7.25 | 536 | 4 | 24816961.34 | 36849606.25 | 26243640.69 | 46051039.45 | 19720812.2 | 82480468.02 | 43629890.75 | 70820512.84 | 29.6996 | 51.9651 | Prrt1 | 260297 | 0.76415 | 1.235845369 | 1.61727 | 0.21534 | 0.716066 |
| 124486648 | neuron navigator 3 [Mus musculus] | 0 | 92.59 | 11 | 19 | 48 | 16 | 2366 | 252.9 | 8.76 | 656 | 19 | 55274172.88 | 68634293.29 | 55069820.14 | 58799574.1 | 63139002.95 | 76346789.53 | 60164522.31 | 73753914.91 | 10.7012 | 11.5604 | Nav3 | 260315 | 0.93031 | 1.069694087 | 1.14983 | 0.12961 | 0.693437 |
| 148674567 | spermatogenesis associated 2, partial [Mus musculus] | 0 | 60.962 | 29 | 13 | 48 | 13 | 540 | 60.3 | 8.47 | 663 | 13 | 56615902.7 | 66842325.51 | 65521584.86 | 63163554.81 | 60303535.45 | 62028380.88 | 65809548.19 | 49297386.09 | 7.20606 | 11.9465 | Spata2 | 263876 | 1.03003 | 0.969965173 | 0.94168 | 0.41621 | 0.836939 |
| 148685167 | ribosomal protein S15a, isoform CRA_a [Mus musculus] | 0 | 47.198 | 53 | 8 | 101 | 8 | 157 | 17.8 | 9.94 | 781 | 8 | 239814354.4 | 306545404.9 | 269303291.3 | 306478075.4 | 272632228 | 339534062.3 | 307964543.6 | 307696748 | 11.5211 | 8.90399 | Rps15a | 267019 | 0.95503 | 1.044973559 | 1.09418 | 0.25838 | 0.745606 |
| 6679439 | peptidyl-prolyl cis-trans isomerase A [Mus musculus] | 0 | 59.899 | 48 | 9 | 50 | 9 | 164 | 18 | 7.9 | 841 | 9 | 117613768.9 | 100772295.2 | 94847986.69 | 146034054.5 | 107363581.7 | 88092722.92 | 99880192.69 | 75671142.03 | 19.9772 | 14.9622 | Ppia | 268373 | 1.1063 | 0.893697405 | 0.80782 | 0.15084 | 0.693437 |
| 27370460 | keratin-like protein KRT222 [Mus musculus] | 0 | 55.501 | 42 | 9 | 27 | 8 | 294 | 34.2 | 5.78 | 400 | 9 | 28394722 | 21119083.87 | 40849913.16 | 11236505.33 | 32173756.05 | 26097244.2 | 46926710.5 | 28836330.7 | 49.0994 | 27.7068 | Krt222 | 268481 | 0.86236 | 1.137644739 | 1.31923 | 0.33712 | 0.782483 |
| 22748747 | protein LSM12 homolog [Homo sapiens] | 0 | 16.183 | 19 | 2 | 9 | 2 | 195 | 21.7 | 7.74 | 269 | 2 | 10243725.93 | 9999953.07 | 5258939.575 | 2370589.506 | 6671062.003 | 7457339.965 | 13507210 | 6454467.494 | 54.9466 | 39.3187 | Lsm12 | 268490 | 0.89967 | 1.100331529 | 1.22304 | 0.56366 | 0.895889 |
| 568979715 | PREDICTED: gephyrin isoform X1 [Mus musculus] | 0 | 429.563 | 63 | 38 | 328 | 38 | 839 | 91.2 | 6.21 | 8978 | 38 | 664786552 | 497837482.5 | 697498145.1 | 564811583.3 | 766727458.1 | 641272588.2 | 576652419.3 | 538587452.9 | 15.1251 | 15.8582 | Gphn | 268566 | 0.98013 | 1.019867159 | 1.04054 | 0.72961 | 0.943709 |
| 568985430 | PREDICTED: protein FAM107A isoform X1 [Mus musculus] | 0 | 13.206 | 17 | 4 | 14 | 4 | 248 | 28.4 | 10.26 | 53 | 4 | 8097709.736 | 20894094.98 | 19987703.29 | 17108608.1 | 2685772.226 | 248762.0729 | 4758607.75 | 6486743.074 | 35.3679 | 75.9203 | Fam107a | 268709 | 1.64669 | 0.353313522 | 0.21456 | 0.00685 | 0.396316 |
| 148669547 | mCG121327 [Mus musculus] | 0 | 17.348 | 2 | 3 | 4 | 3 | 1852 | 199.1 | 6.81 | 168 | 3 | 1710111.1 | 883504.5355 | 4818295.125 | 1264279.635 | 3244476.68 | 3246164.723 | 3562513.188 | 2781352.963 | 82.9019 | 10.0264 | Zswim8 | 268721 | 0.80669 | 1.193313911 | 1.47928 | 0.29844 | 0.761852 |
| 568987748 | PREDICTED: leucine-rich repeat-containing protein 16B isoform X1 [Mus musculus] | 0 | 13.188 | 2 | 2 | 6 | 2 | 1386 | 151.4 | 7.72 | 188 | 2 | 3463009.708 | 2258931.39 | 3278210.821 | 2048888.544 | 3169852.908 | 1010259.001 | 3753102.188 | 3218331.33 | 25.7647 | 43.5524 | Carmil3 | 268747 | 0.99538 | 1.004617219 | 1.00928 | 0.97213 | 0.99686 |
| 37202121 | 4-aminobutyrate aminotransferase, mitochondrial isoform 1 precursor [Mus musculus] | 0 | 14.281 | 7 | 3 | 10 | 3 | 500 | 56.4 | 8.09 | 180 | 3 | 1086597.715 | 763663.8889 | 2019884.912 | 679155.5982 | 1867345.305 | 641122.0965 | 1565120.75 | 760888.9311 | 53.9872 | 49.7245 | Abat | 268860 | 0.96961 | 1.030390204 | 1.06269 | 0.87364 | 0.979509 |
| 22267973 | F-box protein 45 [Mus musculus] | 0.001 | 5.273 | 7 | 2 | 2 | 2 | 286 | 30.6 | 7.83 | 0 | 2 | 1228174.4 | 663937.3779 | 989741.5618 | 801230.8257 | 1351362.284 | 664469.9477 | 1257976.75 | 1286139.402 | 26.5661 | 28.0192 | Fbxo45 | 268882 | 0.89362 | 1.106376411 | 1.23808 | 0.31764 | 0.766675 |
| 568995480 | PREDICTED: limbic system-associated membrane protein isoform X1 [Mus musculus] | 0 | 9.839 | 10 | 2 | 6 | 2 | 361 | 39.8 | 7.37 | 25 | 2 | 3868401.718 | 3903126.764 | 4463698.392 | 3618741.55 | 5556737.465 | 3184573.891 | 3618124.5 | 2878149.59 | 9.00012 | 31.6008 | Lsamp | 268890 | 1.01982 | 0.980175228 | 0.96112 | 0.81428 | 0.961921 |
| 568997544 | PREDICTED: roundabout homolog 2 isoform X1 [Mus musculus] | 0 | 16.443 | 4 | 5 | 9 | 5 | 1598 | 174.7 | 6.71 | 35 | 5 | 2899921.228 | 5943821.272 | 8159671.497 | 5594623.428 | 7867612.817 | 7589917.43 | 6878078 | 9763635.057 | 38.1705 | 15.3505 | Robo2 | 268902 | 0.82629 | 1.173705269 | 1.42044 | 0.10429 | 0.674486 |
| 569000372 | PREDICTED: caskin-1 isoform X1 [Mus musculus] | 0 | 344.184 | 44 | 37 | 371 | 37 | 1438 | 151.1 | 9.17 | 7344 | 37 | 246189587.5 | 244523799.8 | 263974262.6 | 305057030.2 | 289187500.2 | 298671385.9 | 273890207.6 | 300860871.4 | 10.6284 | 4.22142 | Caskin1 | 268932 | 0.95371 | 1.04628662 | 1.09707 | 0.14506 | 0.693437 |
| 158705916 | RecName: Full=Metabotropic glutamate receptor 4; Short=mGluR4; Flags: Precursor | 0 | 101.69 | 26 | 18 | 64 | 17 | 912 | 101.8 | 8.82 | 1226 | 18 | 110554686.1 | 86015978.25 | 107305750.5 | 68196253.66 | 96370850.72 | 114074572.1 | 117699221.6 | 97869837.78 | 21.2925 | 10.2837 | Grm4 | 268934 | 0.93241 | 1.067588876 | 1.14498 | 0.27838 | 0.745606 |
| 569000463 | PREDICTED: striatin isoform X1 [Mus musculus] | 0 | 123.059 | 40 | 21 | 78 | 20 | 809 | 89.2 | 5.41 | 1492 | 21 | 115336829.8 | 74677039.82 | 112313471.4 | 121714012.4 | 86590577.67 | 127982547.1 | 105108721.4 | 124223204.8 | 20.0482 | 17.2058 | Strn | 268980 | 0.97711 | 1.022885857 | 1.04684 | 0.73999 | 0.943908 |
| 145279233 | CBP80/20-dependent translation initiation factor [Mus musculus] | 0.001 | 6.016 | 3 | 2 | 10 | 2 | 623 | 70.2 | 6.61 | 46 | 2 | 2911112.111 | 2567040.785 | 2693193.776 | 2251458.446 | 2734969.756 | 2367624.117 | 2247697.688 | 1841404.259 | 10.5779 | 16.0249 | Ctif | 269037 | 1.06277 | 0.937234738 | 0.88188 | 0.2293 | 0.723653 |
| 154091036 | sn1-specific diacylglycerol lipase alpha [Mus musculus] | 0 | 43.008 | 12 | 10 | 32 | 10 | 1044 | 115.3 | 6.42 | 471 | 10 | 23140565.78 | 15785867.39 | 22325626.04 | 17901818.68 | 29565563.36 | 17033782.35 | 27394816.5 | 16557841.74 | 17.8079 | 30.0676 | Dagla | 269060 | 0.93284 | 1.067164001 | 1.144 | 0.48524 | 0.869 |
| 238776842 | inactive dipeptidyl peptidase 10 [Mus musculus] | 0 | 14.117 | 5 | 4 | 8 | 3 | 800 | 91.1 | 6.48 | 39 | 4 | 2933786.791 | 2220524.449 | 2735069.012 | 1635306.756 | 3095204.337 | 3220928.854 | 3435222.375 | 3010126.209 | 24.4005 | 5.7918 | Dpp10 | 269109 | 0.85476 | 1.145237829 | 1.33983 | 0.0378 | 0.56531 |
| 568935667 | PREDICTED: nuclear pore complex protein Nup54 isoform X1 [Mus musculus] | 0 | 13.918 | 5 | 2 | 8 | 2 | 564 | 60.9 | 6.81 | 176 | 2 | 9554370.887 | 4890704.366 | 9714861.078 | 3757743.937 | 10378968.14 | 9125643.818 | 9393695.5 | 10683869.06 | 44.4355 | 7.60664 | Nup54 | 269113 | 0.82719 | 1.172807718 | 1.41782 | 0.11739 | 0.678142 |
| 160333553 | 60S ribosomal protein L12 [Mus musculus] | 0 | 64.737 | 55 | 6 | 89 | 6 | 165 | 17.8 | 9.42 | 2459 | 6 | 340333292.6 | 412703394.6 | 370152365.8 | 335560099 | 333264553.5 | 429540956.4 | 394376046.8 | 422307608.5 | 9.72944 | 11.0874 | Rpl12 | 269261 | 0.96026 | 1.039740139 | 1.08277 | 0.32524 | 0.772052 |
| 148696043 | vacuolar protein sorting 39 (yeast), isoform CRA_b, partial [Mus musculus] | 0 | 29.831 | 9 | 7 | 32 | 7 | 922 | 105.4 | 7.11 | 382 | 7 | 15066747.23 | 9544233.788 | 17419572.39 | 16130453.63 | 18611760.41 | 10067625.93 | 12490788.13 | 10905821.67 | 23.8429 | 29.6607 | Vps39 | 269338 | 1.0552 | 0.94480065 | 0.89538 | 0.57905 | 0.897763 |
| 40889614 | Chain A, The Crystal Structure Of Murine P97VCP AT 3.6A | 0 | 50.123 | 17 | 11 | 38 | 11 | 816 | 90.5 | 5.43 | 517 | 11 | 18283015.14 | 17941870.66 | 22464105.59 | 20313287.11 | 39224180.86 | 21625142.27 | 20577743 | 24216257.14 | 10.5823 | 32.8579 | Vcp | 269523 | 0.85572 | 1.144282044 | 1.33722 | 0.18622 | 0.715071 |
| 568931328 | PREDICTED: protein 4.1 isoform X12 [Mus musculus] | 0 | 149.757 | 26 | 23 | 192 | 19 | 1025 | 113.1 | 5.31 | 1614 | 23 | 96429518.72 | 157674243.6 | 72389008.2 | 118303527.3 | 143295228.8 | 72417883.75 | 78836893.84 | 115767887.3 | 32.5685 | 32.3574 | Epb41 | 269587 | 1.04032 | 0.959679766 | 0.92248 | 0.73764 | 0.943908 |
| 119433663 | leucine zipper protein 1 [Mus musculus] | 0 | 112.752 | 31 | 25 | 107 | 25 | 1068 | 119.2 | 7.99 | 1228 | 25 | 56423609.74 | 75504838.88 | 91538873.49 | 99097010 | 59926284.96 | 52648361.9 | 64108119.63 | 83287597.46 | 23.444 | 20.1308 | Luzp1 | 269593 | 1.10745 | 0.892548944 | 0.80595 | 0.22233 | 0.718033 |
| 50511119 | mKIAA1805 protein [Mus musculus] | 0 | 23.368 | 13 | 5 | 10 | 5 | 591 | 66.9 | 9.48 | 310 | 5 | 6716961.753 | 5035205.213 | 6265228.502 | 6040859.126 | 15689162.88 | 8481528.46 | 16072665.75 | 10461403.27 | 11.8192 | 29.9063 | Zfp512 | 269639 | 0.64359 | 1.356412937 | 2.10758 | 0.01356 | 0.461382 |
| 568934255 | CTED: serine/threonine-protein phosphatase 2A 55 kDa regulatory subunit B gamma isoform isoform X1 [Mus mu | 0 | 25.553 | 12 | 4 | 22 | 3 | 499 | 56.8 | 6.87 | 451 | 4 | 18060845.78 | 2937225.285 | 12256801.69 | 8365727.418 | 18747260.04 | 1663013.659 | 16848288 | 1081831.196 | 61.2792 | 99.2976 | Ppp2r2c | 269643 | 1.04102 | 0.958977407 | 0.92119 | 0.89085 | 0.984401 |
| 148688086 | golgi autoantigen, golgin subfamily a, 3, isoform CRA_a [Mus musculus] | 0 | 41.677 | 9 | 11 | 17 | 11 | 1487 | 167.2 | 5.38 | 116 | 11 | 2604870.322 | 2684745.806 | 5456005.656 | 12422253.86 | 5406433.5 | 2971401.625 | 6244011.563 | 8409651.088 | 79.6739 | 39.0366 | Golga3 | 269682 | 1.00295 | 0.997048058 | 0.99411 | 0.98983 | 0.998251 |
| 359718915 | probable E3 ubiquitin-protein ligase HECTD4 [Mus musculus] | 0 | 50.204 | 4 | 14 | 30 | 14 | 4418 | 483.8 | 6.06 | 124 | 14 | 8525988.293 | 7032310.848 | 7988027.218 | 5205650.534 | 6290524.196 | 9516560.95 | 7553748.688 | 6844701.306 | 20.2948 | 18.6514 | Hectd4 | 269700 | 0.97535 | 1.024654335 | 1.05056 | 0.7323 | 0.943908 |
| 148687481 | cytoplasmic linker 2, isoform CRA_c, partial [Mus musculus] | 0 | 220.107 | 45 | 35 | 214 | 5 | 1060 | 117 | 6.48 | 3349 | 35 | 18201134.16 | 6233242.056 | 17242554.17 | 9984743.739 | 15699233.1 | 12861642.62 | 16440759.78 | 7685942.626 | 44.6808 | 30.1342 | Clip2 | 269713 | 0.99017 | 1.009831446 | 1.01986 | 0.944 | 0.994906 |
| 187954745 | AP2 associated kinase 1 [Mus musculus] | 0 | 13.38 | 5 | 4 | 14 | 4 | 959 | 103.3 | 6.7 | 149 | 4 | 2550382.525 | 2635473.109 | 1290950.808 | 2369059.458 | 2188874.712 | 5830355.722 | 5050927.188 | 4253236.582 | 28.2009 | 36.1691 | Aak1 | 269774 | 0.67605 | 1.323949866 | 1.95836 | 0.04565 | 0.591577 |
| 148699324 | cDNA sequence BC047219, isoform CRA_b, partial [Mus musculus] | 0 | 9.229 | 18 | 3 | 6 | 3 | 239 | 25.2 | 11.21 | 33 | 3 | 2221516.983 | 3883971.423 | 4617428.554 | 2299901.869 | 4549792.517 | 3064266.79 | 3195743 | 3250486.692 | 36.4815 | 19.75 | Nat14 | 269854 | 0.96169 | 1.03830691 | 1.07967 | 0.71911 | 0.939018 |
| 388327124 | hain A, Crystal Structure Of Geranylgeranylated Rhoa In Complex With Rhogdi In Its Active Gppnhp-Bound Forr | 0 | 10.322 | 23 | 3 | 4 | 2 | 195 | 21.9 | 6.37 | 26 | 3 | 3887471.121 | 3228174.438 | 3079487.535 | 3346830.729 | 6740981.544 | 830367.4163 | 3134850.5 | 6616680.512 | 10.3995 | 66.2799 | Idh2 | 269951 | 0.8775 | 1.122499118 | 1.2792 | 0.53753 | 0.886555 |
| 148684637 | nucleoporin 98, isoform CRA_c [Mus musculus] | 0 | 137.841 | 19 | 24 | 115 | 24 | 1816 | 197.2 | 6.15 | 1179 | 24 | 67996917.98 | 56128958.32 | 71581164.89 | 58329685.16 | 83436008.12 | 93802128.99 | 88720311.88 | 70340905.8 | 11.7335 | 11.9974 | Nup98 | 269966 | 0.86065 | 1.139348807 | 1.32382 | 0.01683 | 0.461382 |
| 157951733 | germ cell-specific gene 1-like protein precursor [Mus musculus] | 0 | 27.866 | 22 | 7 | 44 | 7 | 322 | 35.9 | 7.01 | 145 | 7 | 25186524.76 | 27885831.01 | 26459837.03 | 23257023.2 | 27794208.49 | 28209712.85 | 26394798.13 | 21401777.98 | 7.64781 | 12.0618 | Gsg1l | 269994 | 0.9951 | 1.00489512 | 1.00984 | 0.89565 | 0.984401 |
| 148696944 | microtubule-associated protein 1S, isoform CRA_a, partial [Mus musculus] | 0 | 26.715 | 9 | 7 | 26 | 7 | 984 | 104 | 7.36 | 115 | 7 | 10588140.59 | 10069309.91 | 10501736.49 | 10661110.76 | 10636882.58 | 9962226.736 | 9908515.141 | 10100007.8 | 2.53751 | 3.28237 | Map1s | 270058 | 1.01471 | 0.985288172 | 0.971 | 0.20444 | 0.715071 |
| 148679020 | glutaryl-Coenzyme A dehydrogenase, isoform CRA_a, partial [Mus musculus] | 0 | 31.205 | 22 | 5 | 13 | 5 | 454 | 50.3 | 8.48 | 98 | 5 | 21461592.79 | 14369159.76 | 19381581.38 | 17942247.47 | 18625880.93 | 12794792.66 | 21452308.5 | 12193783.26 | 16.3258 | 27.7433 | Gcdh | 270076 | 1.05851 | 0.941486491 | 0.88944 | 0.48312 | 0.868779 |
| 899445 | 60S ribosomal protein [Mus musculus] | 0 | 21.084 | 20 | 4 | 139 | 4 | 213 | 24.7 | 11.55 | 3199 | 4 | 283839182.7 | 353357537.4 | 315943319 | 369264012.3 | 297916269.1 | 381281669.8 | 326790169.1 | 385995842 | 11.6028 | 12.3147 | Rpl13 | 270106 | 0.97437 | 1.025633733 | 1.05262 | 0.56741 | 0.895889 |
| 74205490 | unnamed protein product [Mus musculus] | 0 | 22.041 | 12 | 4 | 15 | 4 | 447 | 50.2 | 8.73 | 225 | 4 | 132737223 | 80023892.17 | 106186521.4 | 62769234.4 | 155751225.5 | 74771122.77 | 98098089.75 | 102822315.3 | 32.0801 | 31.7085 | Elmod1 | 270162 | 0.93885 | 1.06115144 | 1.13027 | 0.60756 | 0.905186 |
| 568961564 | PREDICTED: unconventional myosin-IXa isoform X1 [Mus musculus] | 0 | 25.634 | 3 | 9 | 29 | 9 | 2633 | 301.9 | 8.9 | 43 | 9 | 8627549.132 | 7068350.61 | 9990258.295 | 6114334.474 | 11715600.49 | 12486999.29 | 11072779.56 | 13887056.1 | 21.5047 | 9.85461 | Myo9a | 270163 | 0.78556 | 1.214443121 | 1.54596 | 0.00606 | 0.396316 |
| 30424655 | ras-related protein Rab-6B [Mus musculus] | 0 | 10.269 | 15 | 3 | 23 | 2 | 208 | 23.4 | 5.53 | 178 | 3 | 4297159.356 | 5061075.958 | 4590974.562 | 4421149.076 | 6335673.848 | 6870200.633 | 6280622.5 | 5466323.565 | 7.2887 | 9.28519 | Rab6b | 270192 | 0.84806 | 1.151938562 | 1.35832 | 0.00266 | 0.38143 |
| 38016190 | leucine-rich repeat-containing protein 4B precursor [Mus musculus] | 0 | 75.685 | 24 | 13 | 67 | 11 | 709 | 76.1 | 7.24 | 916 | 13 | 104578719.4 | 71614584.98 | 93971943.42 | 94949955.57 | 112475367.2 | 102236197.6 | 95475707 | 81146861.49 | 15.2893 | 13.4283 | Lrrc4b | 272381 | 0.96534 | 1.034660523 | 1.07181 | 0.51952 | 0.880629 |
| 227908819 | probable threonine--tRNA ligase 2, cytoplasmic [Mus musculus] | 0 | 12.611 | 5 | 3 | 12 | 3 | 790 | 91.3 | 7.53 | 113 | 3 | 1562081.918 | 1613453.652 | 1471515.699 | 1474981.521 | 1089197.161 | 1653394.183 | 1551027.75 | 1465149.37 | 4.53304 | 17.0875 | Tarsl2 | 272396 | 1.03058 | 0.969424257 | 0.94066 | 0.504 | 0.88031 |
| 148680533 | eukaryotic translation initiation factor 5A, isoform CRA_d, partial [Mus musculus] | 0 | 21.278 | 31 | 3 | 14 | 3 | 174 | 19.2 | 6.21 | 288 | 3 | 18319480.36 | 16270400.09 | 15828628.53 | 7494034.29 | 18735942.88 | 13403645.58 | 16360184 | 6347273.11 | 33.0212 | 39.1808 | Eif5a | 276770 | 1.02719 | 0.972813862 | 0.94707 | 0.83828 | 0.969831 |
| 160420308 | histone H2B type 1-P isoform 1 [Mus musculus] | 0 | 40.625 | 33 | 5 | 106 | 2 | 138 | 15.6 | 10.24 | 2540 | 5 | 83106816.37 | 57825894.75 | 42139749.9 | 38129180.67 | 5226334.492 | 21470727.94 | 27402828.75 | 585031.4271 | 36.8768 | 93.6686 | Hist1h2bp | 319188 | 1.60357 | 0.396430488 | 0.24722 | 0.01351 | 0.461382 |
| 28174920 | Rpl17 protein, partial [Mus musculus] | 0 | 80.222 | 45 | 10 | 130 | 10 | 194 | 22.4 | 10.33 | 1446 | 10 | 283560971.9 | 302843119.1 | 256358622.2 | 311634674 | 247164977.6 | 298943373.8 | 316663871.1 | 297229128.6 | 8.48364 | 10.3027 | Rpl17 | 319195 | 0.99758 | 1.002421348 | 1.00485 | 0.94453 | 0.994906 |
| 28316794 | repressor of yield of DENV protein homolog [Mus musculus] | 0 | 11.631 | 12 | 3 | 11 | 3 | 290 | 33 | 7.28 | 77 | 3 | 4970767.693 | 4643036.628 | 5643652.068 | 4128301.476 | 3777592.258 | 4025309.906 | 6062714.25 | 4212620.23 | 13.0939 | 23.1015 | Shfl | 319278 | 1.0349 | 0.965099258 | 0.93255 | 0.61184 | 0.905186 |
| 755512491 | PREDICTED: adhesion G protein-coupled receptor L3 isoform X1 [Mus musculus] | 0 | 194.889 | 22 | 26 | 174 | 26 | 1599 | 177.8 | 6.77 | 2763 | 26 | 130911647.5 | 133632605.2 | 114123721 | 141041584.4 | 143162883 | 109005267.8 | 126069460.5 | 97447688.63 | 8.75271 | 16.8052 | Adgrl3 | 319387 | 1.04423 | 0.955772066 | 0.91529 | 0.37538 | 0.807546 |
| 60359932 | mKIAA1157 protein [Mus musculus] | 0 | 12.176 | 5 | 2 | 4 | 2 | 640 | 70.1 | 9.64 | 60 | 2 | 2422155.238 | 2624027.191 | 4690219.675 | 554025.4537 | 5007283.584 | 2024304.341 | 4714140.125 | 2001736.191 | 65.7532 | 47.9649 | Ppm1h | 319468 | 0.85618 | 1.143816135 | 1.33595 | 0.49187 | 0.87078 |
| 568979802 | PREDICTED: neuronal cell adhesion molecule isoform X1 [Mus musculus] | 0 | 49.647 | 10 | 9 | 42 | 9 | 1311 | 144.6 | 6 | 659 | 9 | 17891887.58 | 20889878.8 | 27532999.75 | 25187436.78 | 29660904.43 | 113539030.3 | 19998881 | 96387533.23 | 18.856 | 72.358 | Nrcam | 319504 | 0.52125 | 1.478751429 | 2.83694 | 0.12499 | 0.692997 |
| 568926814 | PREDICTED: FSD1-like protein isoform X1 [Mus musculus] | 0 | 7.333 | 7 | 2 | 5 | 2 | 508 | 57 | 6.86 | 22 | 2 | 381337.9834 | 1127787.663 | 1140448.045 | 815633.4158 | 1072713.336 | 1464333.278 | 1129565.5 | 1398003.257 | 41.1526 | 15.3102 | Fsd1l | 319636 | 0.81249 | 1.187507802 | 1.46156 | 0.09626 | 0.670764 |
| 124487111 | echinoderm microtubule-associated protein-like 5 [Mus musculus] | 0 | 24.447 | 3 | 6 | 13 | 6 | 1977 | 220 | 7.8 | 307 | 6 | 12516975.96 | 9798386.604 | 12982905.61 | 9005434.058 | 24871223.52 | 31225464.17 | 28551274 | 25150947.83 | 17.7783 | 11.0103 | Eml5 | 319670 | 0.57499 | 1.425010365 | 2.47832 | 0.0001 | 0.193964 |
| 222418589 | NACHT and WD repeat domain-containing protein 2 [Mus musculus] | 0 | 419.316 | 44 | 57 | 345 | 57 | 1742 | 197.3 | 6.15 | 7001 | 57 | 507496125.5 | 569691310 | 646868550.8 | 577557943.2 | 541814359.4 | 635581665.4 | 606726110 | 505505651.4 | 9.91063 | 10.375 | Nwd2 | 319807 | 1.00261 | 0.997389346 | 0.99479 | 0.94434 | 0.994906 |
| 28892913 | brorin precursor [Mus musculus] | 0 | 17.197 | 14 | 3 | 12 | 3 | 324 | 35.4 | 5.38 | 122 | 3 | 4154814.279 | 8151339.045 | 7202338.937 | 3710953.972 | 6675392.368 | 4922854.603 | 7815678.188 | 9455734.389 | 37.9593 | 26.4426 | Vwc2 | 319922 | 0.89153 | 1.108472074 | 1.24334 | 0.3699 | 0.80688 |

| | | | | | | | | | | | | | | | | | | | | | | | | | | | | | |
|---|---|---|---|---|---|---|---|---|---|---|---|---|---|---|---|---|---|---|---|---|---|---|---|---|---|---|---|---|---|
| 74188559 | unnamed protein product [Mus musculus] | 0 | 13.831 | 6 | 3 | 5 | 3 | 843 | 93 | 4.88 | 45 | 3 | 2463650.194 | 9698384.145 | 1664027.08 | 6243529.306 | 4049854.75 | 3960740.109 | 1780834.5 | 3273384.299 | 73.8437 | 32.1249 | Apba1 | 319924 | 1.2114 | 0.788595051 | 0.65098 | 0.39813 | 0.824613 |
| 568970326 | PREDICTED: tensin-3 isoform X1 [Mus musculus] | 0 | 9.857 | 2 | 3 | 7 | 2 | 1589 | 171.7 | 7.2 | 87 | 3 | 2532250.274 | 5651717.848 | 3762757.702 | 3137330.248 | 4766171.622 | 6081640.111 | 4671811.656 | 6724250.339 | 35.8181 | 18.1217 | Tns3 | 319939 | 0.80819 | 1.191808592 | 1.47466 | 0.07782 | 0.627416 |
| 31982212 | junctophilin-4 [Mus musculus] | 0 | 10.996 | 6 | 3 | 7 | 3 | 628 | 66 | 6.71 | 45 | 3 | 1908487.218 | 2418901.057 | 2854267.664 | 3019971.297 | 3100060.707 | 3858120.371 | 2384972.625 | 3742578.197 | 19.5022 | 20.7422 | Jph4 | 319984 | 0.87615 | 1.123848507 | 1.28271 | 0.13736 | 0.693437 |
| 257796249 | exophilin-5 [Mus musculus] | 0 | 9.789 | 2 | 4 | 8 | 4 | 1960 | 217.5 | 7.5 | 27 | 4 | 3152130.3 | 2420721.726 | 2859497.868 | 2337716.244 | 1669416.945 | 1625073.089 | 1913213.875 | 1450633.867 | 14.2055 | 11.4603 | Exph5 | 320051 | 1.23592 | 0.764078892 | 0.61823 | 0.00297 | 0.38143 |
| 1039775244 | PREDICTED: diacylglycerol kinase iota isoform X4 [Mus musculus] | 0 | 129.872 | 30 | 21 | 117 | 20 | 1091 | 120.5 | 7.37 | 1805 | 21 | 68602088.28 | 67628481.09 | 71473300.28 | 71089421.56 | 73915373.55 | 46355803.25 | 59666732.06 | 70971751.09 | 2.69333 | 19.9654 | Dgki | 320127 | 1.05264 | 0.947359872 | 0.89998 | 0.31312 | 0.76536 |
| 74145171 | unnamed protein product [Mus musculus] | 0 | 37.251 | 12 | 7 | 26 | 2 | 688 | 79.6 | 7.59 | 502 | 7 | 1620642.65 | 1237665.357 | 1589300.627 | 1155140.476 | 587831.3129 | 962160.7047 | 1498079.5 | 918414.1727 | 17.0362 | 37.9912 | Grk3 | 320129 | 1.17099 | 0.829007914 | 0.70795 | 0.11624 | 0.678142 |
| 755497651 | PREDICTED: protein SCAI isoform X1 [Mus musculus] | 0 | 196.129 | 56 | 27 | 194 | 27 | 565 | 65.5 | 7.12 | 3810 | 27 | 613773698 | 534174330.9 | 565671113 | 651822827.3 | 623801415.6 | 733797835.3 | 650995649.4 | 758548261.4 | 8.7804 | 9.33385 | Scai | 320271 | 0.92174 | 1.078264886 | 1.16982 | 0.05158 | 0.595816 |
| 148665529 | IQ calmodulin-binding motif containing 1, isoform CRA_f, partial [Mus musculus] | 0 | 48.783 | 19 | 10 | 33 | 10 | 600 | 68.9 | 9.38 | 239 | 10 | 38138929.38 | 32831382.9 | 308011987.5 | 277813434.9 | 45178862.48 | 24175101.33 | 44987034.38 | 46137558.44 | 90.8362 | 26.5247 | Iqcb1 | 320299 | 1.60728 | 0.392716517 | 0.24434 | 0.14807 | 0.693437 |
| 568939281 | PREDICTED: protein furry homolog isoform X1 [Mus musculus] | 0 | 115.909 | 9 | 22 | 69 | 22 | 3074 | 345.2 | 6.19 | 722 | 22 | 56387663.24 | 41807894.64 | 63829999.35 | 43967988.39 | 54965984.33 | 50821926.48 | 56813067.88 | 42844717.49 | 20.2602 | 12.0828 | Fry | 320365 | 1.00133 | 0.998668456 | 0.99734 | 0.98273 | 0.997682 |
| 70887782 | von Willebrand factor C domain-containing protein 2-like isoform 1 precursor [Mus musculus] | 0 | 10.291 | 11 | 2 | 8 | 2 | 222 | 24.5 | 5.2 | 32 | 2 | 4570916.322 | 5725969.357 | 7901696.96 | 3353592.478 | 9334646.703 | 8566197.653 | 7682891 | 11231099.85 | 35.9235 | 16.4142 | Vwc2l | 320460 | 0.73851 | 1.261494636 | 1.70817 | 0.0209 | 0.505785 |
| 54873617 | protein phosphatase 1E [Mus musculus] | 0 | 119.518 | 28 | 14 | 70 | 14 | 749 | 83.4 | 4.97 | 1298 | 14 | 106366379.6 | 62156447.7 | 122059008.7 | 79562023.58 | 122229671.5 | 85689546.34 | 94973130.34 | 65040894.29 | 28.9577 | 25.7978 | Ppm1e | 320472 | 1.003 | 0.997004897 | 0.99403 | 0.97637 | 0.997464 |
| 83627727 | striatin-interacting proteins 2 isoform a [Mus musculus] | 0 | 39.787 | 16 | 7 | 23 | 6 | 844 | 96.2 | 5.77 | 501 | 7 | 5803732.625 | 7065720.18 | 7946051.083 | 2690441.599 | 4640682.199 | 9165935.762 | 7387145 | 8006144.809 | 39.1187 | 26.3023 | Strip2 | 320609 | 0.89197 | 1.108032826 | 1.24223 | 0.37855 | 0.811177 |
| 148667391 | RIKEN cDNA A930037G23, isoform CRA_a [Mus musculus] | 0.004 | 3.408 | 3 | 2 | 6 | 2 | 587 | 65 | 5.11 | 31 | 2 | 8245629.584 | 8753933.442 | 4738339.347 | 5100821.941 | 5745629.959 | 6298000.844 | 6147701.25 | 6172344.517 | 31.04 | 3.93072 | Iffo1 | 320678 | 1.04834 | 0.95166149 | 0.90778 | 0.57653 | 0.895889 |
| 257467641 | protein SOGA1 [Mus musculus] | 0 | 23.436 | 4 | 6 | 21 | 5 | 1654 | 183 | 6.52 | 147 | 6 | 5277745.714 | 3961803.081 | 4285470.866 | 3923187.209 | 3526412.006 | 1944772.657 | 3774498 | 1898302.238 | 14.4819 | 36.0193 | Soga1 | 320706 | 1.22049 | 0.779512462 | 0.63869 | 0.03761 | 0.56531 |
| 569010236 | PREDICTED: plasma membrane calcium-transporting ATPase 3 isoform X1 [Mus musculus] | 0 | 47.684 | 10 | 8 | 51 | 3 | 1249 | 137.5 | 5.9 | 1259 | 8 | 2534585.164 | 3158357.833 | 2875712.04 | 1547810.803 | 1645908.425 | 2272735.685 | 2036682.625 | 1622072.018 | 27.7628 | 16.6735 | Atp2b3 | 320707 | 1.1435 | 0.85650014 | 0.74902 | 0.15026 | 0.693437 |
| 74200521 | unnamed protein product [Mus musculus] | 0 | 16.555 | 4 | 3 | 9 | 3 | 1025 | 114.9 | 7.93 | 89 | 3 | 6378907.188 | 4223917.891 | 3636115.483 | 3971867.936 | 6474303.032 | 7512277.24 | 5607572.625 | 6641906.684 | 27.2597 | 11.9011 | Mdga2 | 320772 | 0.81944 | 1.180558303 | 1.44069 | 0.03388 | 0.528235 |
| 568922988 | PREDICTED: neuronal growth regulator 1 isoform X1 [Mus musculus] | 0 | 78.897 | 36 | 9 | 72 | 9 | 393 | 43.1 | 7.69 | 1598 | 9 | 121351064.5 | 75666248.41 | 79666372.01 | 70765053.97 | 80572487.36 | 58050166.75 | 72341498.25 | 66297531.44 | 26.7999 | 13.7343 | Negr1 | 320840 | 1.11235 | 0.887648657 | 0.79799 | 0.21237 | 0.716066 |
| 145587090 | cadherin-18 precursor [Mus musculus] | 0 | 41.431 | 14 | 8 | 36 | 8 | 790 | 87.9 | 5.16 | 385 | 8 | 21518990.91 | 14655546.59 | 20739386.86 | 21139144.9 | 20849252.84 | 28374360.5 | 18919945.88 | 27470064.4 | 16.6763 | 19.7518 | Cdh18 | 320865 | 0.89888 | 1.101116421 | 1.22498 | 0.17659 | 0.713031 |
| 148676983 | mCG123390, isoform CRA_d, partial [Mus musculus] | 0 | 63.179 | 16 | 11 | 102 | 9 | 789 | 88.4 | 4.94 | 1722 | 11 | 99129217.76 | 91491604.7 | 116018934 | 81338402.1 | 111968416.1 | 112202331.7 | 114815259 | 96423465.23 | 15.0806 | 7.70384 | Cdh10 | 320873 | 0.94239 | 1.057605084 | 1.12225 | 0.20918 | 0.716066 |
| 28893409 | ectonucleotide pyrophosphatase/phosphodiesterase family member 6 precursor [Mus musculus] | 0 | 62.771 | 25 | 9 | 49 | 9 | 440 | 50.6 | 7.31 | 1320 | 9 | 51862014.15 | 36909919.54 | 37304808.97 | 35853819.8 | 44549170.57 | 30700086.86 | 35256455.25 | 28872345.3 | 18.8005 | 20.1024 | Enpp6 | 320981 | 1.07485 | 0.925151479 | 0.86073 | 0.31748 | 0.766675 |
| 47894398 | tropomyosin alpha-4 chain [Mus musculus] | 0 | 17.233 | 15 | 5 | 20 | 5 | 248 | 28.5 | 4.68 | 103 | 5 | 19099889.38 | 17910771.44 | 19609747.27 | 8628265.761 | 19209035.08 | 10295477.73 | 25628896.25 | 2871212.163 | 31.7053 | 68.8374 | Tpm4 | 326618 | 1.05877 | 0.941226296 | 0.88898 | 0.75825 | 0.952737 |
| 148700589 | mCG17007, partial [Mus musculus] | 0 | 64.075 | 53 | 9 | 91 | 9 | 104 | 11.5 | 11.19 | 1801 | 9 | 1212613880 | 967188219 | 1178737117 | 1309234450 | 734394572.4 | 1489667125 | 1113824965 | 1507046741 | 12.3564 | 30.2181 | Hist1h4a | 326619 | 0.98138 | 1.018623483 | 1.03795 | 0.82932 | 0.968871 |
| 568967830 | PREDICTED: liprin-alpha-2 isoform X6 [Mus musculus] | 0 | 427.216 | 55 | 55 | 559 | 6 | 1258 | 143.3 | 6.07 | 11724 | 55 | 111050013.8 | 74414680.06 | 83403823.79 | 73350608.82 | 66548169.35 | 63541422.64 | 62030668.5 | 57621441.63 | 20.5539 | 5.95535 | Ppfia2 | 327814 | 1.15622 | 0.843777798 | 0.72977 | 0.04219 | 0.585646 |
| 145699114 | colorectal mutant cancer protein isoform 1 [Mus musculus] | 0 | 20.357 | 7 | 5 | 8 | 5 | 1003 | 111.9 | 5.39 | 103 | 5 | 3535050.929 | 3019254.78 | 3545228.849 | 3454450.658 | 4363054.043 | 1710574.925 | 1759431.75 | 1998601.589 | 7.36279 | 51.9266 | Mcc | 328949 | 1.15917 | 0.840828743 | 0.72537 | 0.20234 | 0.715071 |
| 49457861 | E3 ubiquitin-protein ligase HECW2 isoform 1 [Mus musculus] | 0 | 81.721 | 15 | 17 | 65 | 15 | 1578 | 176.1 | 5.41 | 365 | 17 | 23874984.37 | 30237048.27 | 35985564.57 | 25046929.24 | 42938293.84 | 39303249.33 | 33272752.63 | 20523090.14 | 19.2415 | 28.9181 | Hecw2 | 329152 | 0.91682 | 1.083178201 | 1.18145 | 0.39041 | 0.822031 |
| 568906557 | PREDICTED: abl interactor 2 isoform X1 [Mus musculus] | 0 | 105.466 | 28 | 12 | 67 | 8 | 542 | 58.8 | 6.02 | 1503 | 12 | 109114976.9 | 98086310.89 | 106969879.4 | 105258551.4 | 132895741.5 | 119617999.6 | 116579673.1 | 102036045.3 | 4.56033 | 10.7525 | Abi2 | 329165 | 0.94195 | 1.058053122 | 1.12326 | 0.10478 | 0.674486 |
| 755492562 | PREDICTED: protein unc-80 homolog isoform X1 [Mus musculus] | 0 | 43.22 | 4 | 9 | 25 | 9 | 3341 | 372 | 7.03 | 392 | 9 | 9703405.825 | 7649944.987 | 12564754.64 | 7291681.936 | 13237092.29 | 6285334.459 | 13077114.88 | 6161298.555 | 26.0207 | 41.3208 | Unc80 | 329178 | 0.97958 | 1.02041648 | 1.04168 | 0.8738 | 0.979509 |
| 755508335 | PREDICTED: CUB and sushi domain-containing protein 2 isoform X1 [Mus musculus] | 0 | 31.538 | 3 | 9 | 23 | 9 | 3612 | 392.3 | 6.14 | 121 | 9 | 9106205.375 | 4742613.475 | 7252493.326 | 3788418.147 | 6086635.116 | 7526628.116 | 5240030.75 | 5035577.203 | 38.8054 | 18.9505 | Csmd2 | 329942 | 1.02052 | 0.979481594 | 0.95979 | 0.85733 | 0.975398 |
| 257467657 | UPF0606 protein KIAA1549 precursor [Mus musculus] | 0 | 117.457 | 12 | 19 | 77 | 19 | 1940 | 209.1 | 6.1 | 1104 | 19 | 101374207.5 | 90411176.12 | 86032196.83 | 94703408.64 | 97244675.7 | 84638708.53 | 74343141.25 | 95956990.4 | 7.01965 | 12.2076 | i30045J12f | 330286 | 1.02806 | 0.971936875 | 0.94541 | 0.44977 | 0.854909 |
| 47564092 | F-box only protein 41 [Mus musculus] | 0 | 233.284 | 47 | 31 | 226 | 31 | 873 | 94.3 | 8.24 | 3340 | 31 | 555517606.4 | 460118291.1 | 518409039.3 | 463057438.7 | 529832367.8 | 480553686 | 507394832.9 | 409408010 | 9.23242 | 10.8542 | Fbxo41 | 330369 | 1.01782 | 0.982184432 | 0.96499 | 0.63391 | 0.912586 |
| 148696832 | hyaluronan and proteoglycan link protein 4, partial [Mus musculus] | 0 | 88.566 | 44 | 12 | 65 | 11 | 413 | 44.2 | 9.35 | 1735 | 12 | 77584181.31 | 68043174.36 | 83851017.24 | 65480113.78 | 72984472.23 | 64710003.87 | 63201690.38 | 57725507.29 | 11.552 | 9.76208 | Hapln4 | 330790 | 1.06564 | 0.934360341 | 0.87681 | 0.13742 | 0.693437 |
| 568957049 | PREDICTED: adhesion G protein-coupled receptor L1 isoform X2 [Mus musculus] | 0 | 279.752 | 38 | 36 | 191 | 35 | 1472 | 162.3 | 6.43 | 3509 | 36 | 287803530.9 | 236636756.8 | 243911728.3 | 240361463.3 | 268149684.1 | 265897915.1 | 249459924.6 | 248273052.2 | 9.49135 | 4.08401 | Adgrl1 | 330814 | 0.9887 | 1.011304662 | 1.02287 | 0.67463 | 0.923206 |
| 568959727 | PREDICTED: rho GTPase-activating protein 32 isoform X1 [Mus musculus] | 0 | 600.657 | 57 | 77 | 610 | 74 | 2103 | 231.1 | 6.87 | 9226 | 77 | 579917254.5 | 597575793.7 | 624453575.4 | 580009471.7 | 573613185.5 | 586931731.7 | 581704235.7 | 568596253.5 | 3.52971 | 1.41629 | Arhgap32 | 330914 | 1.01515 | 0.984846858 | 0.97015 | 0.16602 | 0.711253 |
| 237874261 | interleukin-1 receptor accessory protein-like 1 precursor [Mus musculus] | 0 | 116.739 | 33 | 22 | 83 | 22 | 696 | 79.7 | 6.32 | 1661 | 22 | 144575552.6 | 125680941.1 | 137175164.8 | 120389024.5 | 134695866.2 | 148020147.8 | 138972796.4 | 128235667.9 | 8.29777 | 6.03518 | Il1rapl1 | 331461 | 0.97949 | 1.020509297 | 1.04188 | 0.45181 | 0.857112 |
| 145580629 | keratin Kb40 [Mus musculus] | 0 | 21.937 | 2 | 3 | 39 | 2 | 1068 | 112.2 | 7.97 | 314 | 3 | 43189476.21 | 25773153.91 | 58176098.67 | 11806882.29 | 18940083.03 | 50573148.62 | 16150372.25 | 59314207.83 | 58.2201 | 60.4631 | Krt78 | 332131 | 0.97875 | 1.021245872 | 1.04341 | 0.92273 | 0.988641 |
| 569012064 | PREDICTED: FERM and PDZ domain-containing protein 4 isoform X1 [Mus musculus] | 0 | 18.348 | 3 | 4 | 7 | 4 | 1778 | 195.6 | 5.34 | 95 | 4 | 6756041.859 | 3502819.992 | 4823695.865 | 4623650.465 | 4982138.961 | 3530034.349 | 5274964.5 | 1007798.555 | 27.4248 | 52.7062 | Frmpd4 | 333605 | 1.14235 | 0.857649018 | 0.75078 | 0.34044 | 0.78329 |
| 118026911 | unconventional myosin-Id [Mus musculus] | 0 | 39.272 | 13 | 11 | 33 | 11 | 1006 | 116 | 9.41 | 198 | 11 | 21801626.75 | 15699874.98 | 21523163.89 | 13964546.75 | 22640557.86 | 15961136.49 | 21631314.81 | 12893899.53 | 21.9657 | 25.3892 | Myo1d | 338367 | 0.99906 | 1.000942376 | 1.00189 | 0.99141 | 0.998251 |
| 30017461 | arf-GAP with GTPase, ANK repeat and PH domain-containing protein 1 isoform 1 [Mus musculus] | 0 | 83.63 | 20 | 12 | 82 | 10 | 857 | 94.4 | 7.94 | 2073 | 12 | 17706450.72 | 33908213.94 | 40960790.78 | 23179759.52 | 21148029.45 | 30193722.75 | 22998486.06 | 14203476.51 | 36.1629 | 29.6891 | Agap1 | 347722 | 1.13319 | 0.866805469 | 0.76492 | 0.31309 | 0.76536 |
| 568974533 | PREDICTED: unconventional myosin-XVIIIa isoform X1 [Mus musculus] | 0 | 429.728 | 44 | 68 | 452 | 68 | 2105 | 238.4 | 6.07 | 7491 | 68 | 351626347.5 | 347017138.8 | 376216235 | 332511265.5 | 358542801.5 | 333154608.2 | 364708178.5 | 349878889.4 | 5.16596 | 3.89731 | Myo18a | 360013 | 1.00039 | 0.999613844 | 0.99923 | 0.98173 | 0.997464 |
| 51492004 | tripartite motif-containing protein 46 isoform 2 [Mus musculus] | 0 | 116.428 | 38 | 20 | 96 | 20 | 759 | 83.4 | 7.65 | 2338 | 20 | 101446025 | 111111211.3 | 114974102 | 96931290.38 | 115877117.8 | 75744283.95 | 101007848.3 | 85410769.68 | 7.87663 | 18.6668 | Trim46 | 360213 | 1.05785 | 0.942152703 | 0.89063 | 0.27936 | 0.745606 |
| 84370019 | headcase protein homolog [Mus musculus] | 0 | 10.082 | 5 | 2 | 4 | 2 | 544 | 59.1 | 8.6 | 47 | 2 | 1162108.567 | 1660428.835 | 1142147.485 | 1838390.473 | 1546068.573 | 1565948.211 | 1357209.625 | 1821606.712 | 24.2978 | 12.1281 | Heca | 380629 | 0.95967 | 1.040330863 | 1.08405 | 0.56518 | 0.895889 |
| 152031637 | RecName: Full=Inner nuclear membrane protein Man1; AltName: Full=LEM domain-containing protein 3 | 0 | 75.45 | 21 | 17 | 90 | 17 | 921 | 100.2 | 7.55 | 1412 | 17 | 48393372.62 | 31035474.92 | 52508954.14 | 55816502.92 | 55120296.13 | 67177452.11 | 60591714.8 | 43192170.21 | 23.4952 | 17.9779 | Lemd3 | 380664 | 0.90739 | 1.09261479 | 1.20414 | 0.24848 | 0.733806 |
| 124286811 | neurofilament heavy polypeptide [Mus musculus] | 0 | 439.572 | 40 | 44 | 997 | 4 | 1090 | 116.9 | 5.81 | 17643 | 44 | 45307842.39 | 20255278.1 | 29891189.49 | 45878793.01 | 44032256.91 | 23375806.61 | 34731034.9 | 42419966.18 | 35.3363 | 26.0863 | Nefh | 380684 | 0.98872 | 1.01128384 | 1.02283 | 0.92124 | 0.987994 |
| 568974579 | PREDICTED: protein shisa-6 homolog isoform X1 [Mus musculus] | 0 | 138.885 | 43 | 17 | 165 | 16 | 557 | 61.8 | 9.52 | 1942 | 17 | 192579035.2 | 168812520.3 | 203204495.1 | 218655427.6 | 239005887.5 | 196401747.3 | 188125768.3 | 204903315.6 | 10.6957 | 10.7868 | Shisa6 | 380702 | 0.97196 | 1.028035969 | 1.05769 | 0.48849 | 0.869525 |
| 568974591 | PREDICTED: rap1 GTPase-activating protein 2 isoform X3 [Mus musculus] | 0 | 29.682 | 11 | 8 | 17 | 7 | 790 | 87 | 6.9 | 125 | 8 | 6048213.577 | 6326702.754 | 6658074.393 | 5096230.992 | 5346619.482 | 19578108.92 | 12851586.5 | 7405566.229 | 11.14 | 56.351 | Rap1gap2 | 380711 | 0.69626 | 1.303741515 | 1.8725 | 0.15115 | 0.693437 |
| 158937310 | coiled-coil domain-containing protein 177 [Mus musculus] | 0 | 46.009 | 17 | 10 | 41 | 10 | 706 | 79.8 | 10.8 | 582 | 10 | 24806961.36 | 47071805.49 | 47552091.75 | 18342526.68 | 51746745.25 | 14125755.3 | 58222125.57 | 13199040.8 | 43.8203 | 69.9414 | Ccdc177 | 380768 | 1.00174 | 0.998255995 | 0.99652 | 0.99353 | 0.998251 |
| 1039746424 | PREDICTED: LIM domain only protein 7 isoform X10 [Mus musculus] | 0 | 286.415 | 43 | 42 | 299 | 2 | 1466 | 166.4 | 6.64 | 5455 | 42 | 5106093.345 | 10198527.01 | 6461745.622 | 5523196.641 | 5693364.903 | 5024372.846 | 5210120.875 | 6589137.966 | 34.0212 | 12.4225 | Lmo7 | 380928 | 1.09582 | 0.904177962 | 0.82511 | 0.36294 | 0.804292 |
| 54261813 | presenilins-associated rhomboid-like protein, mitochondrial precursor [Mus musculus] | 0 | 21.672 | 13 | 4 | 13 | 4 | 377 | 41.9 | 9.92 | 137 | 4 | 8465031.477 | 20200842.95 | 14337622.35 | 15352520.2 | 16400124.91 | 30890065.91 | 13403342.5 | 25030514.88 | 33.0255 | 37.3469 | Parl | 381038 | 0.81005 | 1.189950159 | 1.46898 | 0.19333 | 0.715071 |
| 134288888 | GRB2-associated and regulator of MAPK protein [Mus musculus] | 0 | 7.132 | 3 | 2 | 3 | 2 | 876 | 97.2 | 6.6 | 20 | 2 | 1953676.318 | 1175954.612 | 1899497.941 | 862758.4308 | 2129961.768 | 1053905.173 | 1935629.75 | 760122.9871 | 36.635 | 45.2878 | Garem1 | 381126 | 1.00104 | 0.998957854 | 0.99792 | 0.99452 | 0.998251 |
| 755494464 | PREDICTED: plasma membrane calcium-transporting ATPase 4 isoform X5 [Mus musculus] | 0 | 86.092 | 18 | 16 | 72 | 7 | 1239 | 136.9 | 6.24 | 1356 | 16 | 16153691.3 | 14789645.71 | 11328908.57 | 9079028.192 | 20985909.16 | 27168753.77 | 20887978.25 | 27245200.53 | 25.1243 | 15.0397 | Atp2b4 | 381290 | 0.69563 | 1.304367632 | 1.87508 | 0.00356 | 0.38143 |
| 148762971 | uncharacterized protein C9orf172 homolog [Mus musculus] | 0 | 377.071 | 60 | 47 | 630 | 47 | 974 | 107.1 | 9.11 | 11440 | 47 | 1589719736 | 1550385418 | 1689182510 | 1702638489 | 1718494781 | 1678175386 | 1698347954 | 1607069942 | 4.56954 | 2.89537 | Ajm1 | 381353 | 0.98714 | 1.012857921 | 1.02605 | 0.37602 | 0.807546 |
| 594190788 | [Pyruvate dehydrogenase [acetyl-transferring]]-phosphatase 1, mitochondrial isoform d [Mus musculus] | 0 | 24.172 | 9 | 4 | 18 | 4 | 578 | 65.4 | 6.9 | 303 | 4 | 6893335.849 | 4042507.278 | 9916673.203 | 4033964.902 | 12378734.45 | 4092138.169 | 10456644.75 | 4389729.036 | 45.118 | 53.8859 | Pdp1 | 381511 | 0.88558 | 1.114418837 | 1.2584 | 0.54915 | 0.890708 |
| 168480088 | transmembrane protein 240 [Mus musculus] | 0 | 27.898 | 39 | 4 | 26 | 4 | 173 | 20 | 7.8 | 178 | 4 | 18211546.42 | 22312796.38 | 21240227.37 | 14106533.53 | 27243624.13 | 20683637.01 | 18361721.5 | 9980522.412 | 19.3845 | 37.3963 | Tmem240 | 381582 | 0.99738 | 1.002618639 | 1.00525 | 0.981 | 0.997464 |
| 148705959 | centrosomal protein 135, isoform CRA_a, partial [Mus musculus] | 0 | 33.82 | 11 | 11 | 19 | 11 | 1152 | 134.4 | 6.57 | 75 | 11 | 18449635.05 | 7975486.793 | 11379896.2 | 23768348.33 | 18031408.34 | 7121144.539 | 12666171.25 | 12748370.62 | 46.0298 | 35.2392 | Cep135 | 381644 | 1.09815 | 0.901852806 | 0.82125 | 0.53525 | 0.886555 |
| 86476054 | neurosecretory protein VGF precursor [Mus musculus] | 0 | 20.717 | 11 | 5 | 13 | 5 | 617 | 68.2 | 4.7 | 151 | 5 | 8145967.782 | 11266966.48 | 13566665.3 | 13767605.24 | 14370505.54 | 13807992.06 | 9809097.5 | 13393283.74 | 22.4098 | 16.0629 | Vgf | 381677 | 0.95278 | 1.047220672 | 1.09912 | 0.51312 | 0.88031 |
| 357527369 | protein arginine N-methyltransferase 8 [Mus musculus] | 0 | 15.671 | 10 | 5 | 12 | 5 | 394 | 45.2 | 6.93 | 57 | 5 | 18123438.99 | 5265540.672 | 14905702.17 | 6090087.195 | 23609515.08 | 7750417.043 | 16762276.75 | 7225447.732 | 57.6944 | 56.7248 | Prmt8 | 381813 | 0.89008 | 1.109923002 | 1.247 | 0.60788 | 0.905186 |
| 37360134 | mKIAA0881 protein [Mus musculus] | 0 | 85.452 | 22 | 19 | 76 | 17 | 1070 | 121.3 | 7.37 | 1755 | 19 | 50888708.44 | 41801520.75 | 45909818.83 | 49410792.55 | 49216880.22 | 44278988.61 | 69944201.78 | 45491336.92 | 8.61119 | 22.9607 | Taok2 | 381921 | 0.9473 | 1.052704309 | 1.11127 | 0.44019 | 0.852048 |
| 1039779092 | PREDICTED: serine/threonine-protein kinase BRSK1 isoform X1 [Mus musculus] | 0 | 88.761 | 24 | 14 | 68 | 13 | 806 | 88.1 | 9.33 | 1726 | 14 | 52737029.83 | 38053657.72 | 48751918.08 | 42212566.1 | 46309770.65 | 44584948.74 | 56169175.94 | 41689739.93 | 14.442 | 13.3153 | Brsk1 | 381979 | 0.98111 | 1.018888791 | 1.0385 | 0.71341 | 0.936732 |
| 568948028 | PREDICTED: serine/threonine-protein kinase LMTK3 isoform X1 [Mus musculus] | 0 | 75.514 | 11 | 12 | 54 | 12 | 1450 | 153.7 | 4.92 | 819 | 12 | 45222318.7 | 33787640.29 | 40758799.95 | 23848677.59 | 32539125.08 | 36632181.12 | 44331985.75 | 31440516.66 | 25.9391 | 16.1204 | Lmtk3 | 381983 | 0.9954 | 1.004596501 | 1.00924 | 0.95386 | 0.994906 |
| 755526940 | PREDICTED: genetic suppressor element 1 isoform X1 [Mus musculus] | 0.001 | 5.838 | 1 | 2 | 4 | 2 | 1579 | 175.5 | 7.53 | 0 | 2 | 1370458.963 | 1250398.51 | 1189031.425 | 1197545.724 | 1456716.331 | 1403994.612 | 1120607.625 | 1138885.411 | 6.67782 | 13.6749 | Gse1 | 382034 | 0.98887 | 1.011134812 | 1.02252 | 0.78109 | 0.952792 |
| 755525967 | PREDICTED: CREB-regulated transcription coactivator 1 isoform X1 [Mus musculus] | 0 | 9.815 | 4 | 2 | 7 | 2 | 657 | 69.6 | 6.02 | 28 | 2 | 2837878.287 | 5781408.144 | 7289720.077 | 5785350.334 | 5562771.982 | 6835191.324 | 4800697.375 | 9589648.837 | 34.3746 | 31.4031 | Crtc1 | 382056 | 0.89493 | 1.105067503 | 1.23481 | 0.39973 | 0.824613 |
| 247494171 | BTB/POZ domain-containing protein KCTD16 [Mus musculus] | 0 | 136.122 | 55 | 27 | 172 | 25 | 427 | 48.9 | 8.19 | 1744 | 27 | 490782060.2 | 651126089 | 575254683.6 | 622200260.8 | 564719479.2 | 768576457.1 | 549586932.1 | 664427796.2 | 11.9806 | 15.9444 | Kctd16 | 383348 | 0.95745 | 1.042554011 | 1.08889 | 0.43165 | 0.8471 |
| 126517480 | ankyrin repeat domain-containing protein 63 [Mus musculus] | 0 | 27.23 | 20 | 5 | 13 | 5 | 390 | 41 | 10.83 | 154 | 5 | 26107017.95 | 19238327.46 | 22711135.82 | 19419160.34 | 22568088.71 | 17563165.93 | 23147184.75 | 15848604.79 | 14.8389 | 18.3388 | Ankrd63 | 383787 | 1.05011 | 0.949889179 | 0.90456 | 0.42403 | 0.840684 |
| 29467630 | neuronal transmembrane protein Slitrk3 [Mus musculus] | 0.001 | 6.935 | 2 | 2 | 5 | 2 | 980 | 109.6 | 7.56 | 54 | 2 | 3060381.218 | 2664504.882 | 3461924.632 | 2555602.124 | 3164834.706 | 3011942.118 | 2596476.625 | 2286066.305 | 14.0518 | 14.4467 | Slitrk3 | 386750 | 1.02996 | 0.970042053 | 0.94183 | 0.57369 | 0.895889 |
| 41462393 | leucine-rich repeat transmembrane protein FLRT2 precursor [Mus musculus] | 0 | 43.009 | 19 | 10 | 29 | 10 | 660 | 73.9 | 7.91 | 140 | 10 | 45259253.82 | 38262685.23 | 45804116.57 | 37841037.88 | 50467041.22 | 48470203.91 | 40253634.38 | 55268707.25 | 10.3552 | 12.8807 | Flrt2 | 399558 | 0.92453 | 1.075471459 | 1.16326 | 0.12318 | 0.688884 |
| 110625722 | optic atrophy 3 protein homolog [Mus musculus] | 0 | 32.484 | 28 | 3 | 19 | 3 | 179 | 20.1 | 8.34 | 144 | 3 | 35816961.87 | 5673885.562 | 22546584.12 | 25825906.57 | 25862903.83 | 7343520.57 | 25891963.5 | 20210119.07 | 55.8041 | 44.0846 | Opa3 | 403187 | 1.06239 | 0.937608819 | 0.88255 | 0.74164 | 0.945377 |
| 46250738 | core histone macro-H2A.2 [Mus musculus] | 0 | 40.521 | 27 | 6 | 22 | 5 | 372 | 40.1 | 9.69 | 497 | 6 | 58882644.35 | 22304503.93 | 53185195.72 | 20144723.98 | 43317635.12 | 22672852.79 | 46719436 | 17773113.13 | 52.4227 | 44.5157 | H2afy2 | 404634 | 1.08433 | 0.915670098 | 0.84446 | 0.64672 | 0.915705 |
| 61888832 | adenylate cyclase type 1 [Mus musculus] | 0 | 13.913 | 3 | 3 | 6 | 3 | 1118 | 123.3 | 8.47 | 86 | 3 | 4866604.122 | 11172768.96 | 2843662.022 | 2630830.457 | 4169128.974 | 4874893.658 | 4018768.5 | 2121333.413 | 74.2238 | 31.0112 | Adcy1 | 432530 | 1.17248 | 0.827518047 | 0.70578 | 0.47582 | 0.863085 |
| 529250185 | cytospin-B isoform 1 [Mus musculus] | 0 | 58.787 | 16 | 12 | 47 | 12 | 1058 | 117 | 6.57 | 570 | 12 | 38873917.73 | 20615711.54 | 28660245.9 | 19585387.57 | 45509411.52 | 13022431.94 | 31714299.84 | 17330498.94 | 33.1713 | 54.885 | Specc1 | 432572 | 1.00074 | 0.999263299 | 0.99853 | 0.99648 | 0.998251 |
| 71892426 | proline-rich protein 7 [Mus musculus] | 0 | 58.833 | 41 | 9 | 61 | 9 | 269 | 30.3 | 8.6 | 860 | 9 | 184863598.6 | 153082226.3 | 172188879.5 | 143665905.6 | 188243789 | 182430131.6 | 177901934.4 | 136451618.5 | 11.3573 | 13.7725 | Prr7 | 432763 | 0.97668 | 1.023324028 | 1.04776 | 0.62157 | 0.908209 |
| 58219050 | OCIA domain-containing protein 2 isoform 1 [Mus musculus] | 0 | 21.826 | 38 | 4 | 21 | 4 | 154 | 16.9 | 9.41 | 369 | 4 | 11663645.74 | 11223547.44 | 9389232.1 | 7158945.169 | 14747865.86 | 24970188.7 | 11413463.69 | 17424169.38 | 20.8118 | 33.671 | Ociad2 | 433904 | 0.73035 | 1.269654893 | 1.73843 | 0.05497 | 0.599194 |
| 305632805 | transmembrane protein 178B precursor [Rattus norvegicus] | 0 | 13.016 | 12 | 3 | 9 | 3 | 294 | 33.3 | 9.17 | 141 | 3 | 4045368.987 | 2918619.573 | 4684413.939 | 5099246.511 | 2558835.203 | 2802839.49 | 4829825.75 | 2339984.707 | 22.6938 | 36.6115 | Tmem178b | 434008 | 1.144 | 0.856001074 | 0.74825 | 0.20671 | 0.715071 |
| 187957004 | EG434080 protein [Mus musculus] | 0 | 34.423 | 26 | 9 | 28 | 8 | 411 | 46.9 | 7.75 | 186 | 9 | 9597719.612 | 8481002.753 | 12041529.38 | 19401305.14 | 9072916.608 | 13635799.29 | 14471521.75 | 14004523.77 | 39.6683 | 19.581 | Gm5580 | 434080 | 0.98348 | 1.016515394 | 1.03359 | 0.88505 | 0.983671 |
| 153791731 | PNMA-like protein 2 [Mus musculus] | 0.001 | 5.563 | 4 | 2 | 5 | 2 | 661 | 71.7 | 5.17 | 13 | 2 | 3599459.085 | 4724182.318 | 3123316.372 | 4371595.549 | 5115798.74 | 5920816.903 | 4254011 | 5994841.048 | 18.3702 | 15.3223 | Pnmal2 | 434128 | 0.85266 | 1.147340213 | 1.3456 | 0.04634 | 0.591577 |
| 333108269 | putative protein shisa-8 precursor [Mus musculus] | 0 | 11.291 | 9 | 3 | 9 | 3 | 399 | 42.4 | 9.96 | 88 | 3 | 15239725.94 | 41005560.28 | 26535673.19 | 39110158.31 | 25757474.92 | 55291382.59 | 29081590.5 | 77217865.42 | 39.4291 | 51.6266 | Shisa8 | 435145 | 0.78833 | 1.211671569 | 1.53701 | 0.27103 | 0.745606 |
| 108773813 | nuclear pore complex protein Nup85 [Mus musculus] | 0 | 44.47 | 19 | 8 | 22 | 8 | 656 | 74.7 | 5.57 | 615 | 8 | 7405358.771 | 8470889.191 | 12193720.12 | 4789747.954 | 12517201.87 | 12106936.63 | 14903656.25 | 1224755.939 | 37.3768 | 59.8882 | Nup85 | 445007 | 0.89278 | 1.107221731 | 1.2402 | 0.58444 | 0.899211 |
| 59858539 | XK-related protein 4 [Mus musculus] | 0 | 16.364 | 8 | 4 | 19 | 4 | 647 | 71.5 | 8.03 | 225 | 4 | 8613038.935 | 14373110.45 | 6548213.478 | 7804294.323 | 8167989.061 | 10570867.75 | 7446352.75 | 12698250.53 | 37.1168 | 24.612 | Xkr4 | 497097 | 0.97973 | 1.020267121 | 1.04137 | 0.86049 | 0.976238 |
| 148690464 | mCG17651 [Mus musculus] | 0 | 9.433 | 3 | 4 | 10 | 4 | 1234 | 138.2 | 6.24 | 13 | 4 | 5727462.629 | 2590972.705 | 8264192.208 | 3285598.43 | 8741187.292 | 2895723.31 | 7040909.625 | 4340196.449 | 51.8813 | 45.7004 | Baiap3 | 545192 | 0.92655 | 1.073445247 | 1.15853 | 0.68377 | 0.927189 |
| 1039729570 | PREDICTED: centrosomal protein of 170 kDa isoform X1 [Mus musculus] | 0 | 143.829 | 22 | 24 | 100 | 22 | 1650 | 181.9 | 7.36 | 1737 | 24 | 62690685.21 | 61926709.76 | 61410760.18 | 47304085.15 | 59507311.3 | 42671450.45 | 65905214.88 | 39762188.43 | 12.6368 | 24.5074 | Cep170 | 545389 | 1.05777 | 0.942231816 | 0.89077 | 0.41974 | 0.839919 |
| 225480 | tubulin Mbeta 1 | 0 | 51.252 | 18 | 7 | 234 | 2 | 451 | 50.4 | 5.07 | 4523 | 7 | 174068683.6 | 83843953.76 | 186588309.1 | 27671890.62 | 164031771.7 | 73565539.94 | 155957252 | 65616470.68 | 64.0962 | 45.6467 | Tubb1 | 545486 | 1.01396 | 0.98603974 | 0.97246 | 0.94598 | 0.994906 |
| 120953245 | ankyrin repeat domain-containing protein 34A [Mus musculus] | 0 | 9.013 | 5 | 2 | 8 | 2 | 495 | 52.4 | 9.45 | 65 | 2 | 2488844.842 | 2031174.749 | 1836054.913 | 2619305.132 | 2406989.921 | 1629950.679 | 1947196.25 | 1791087.513 | 16.526 | 17.2267 | Ankrd34a | 545554 | 1.07165 | 0.928351523 | 0.86628 | 0.27499 | 0.745606 |
| 1039790376 | PREDICTED: microtubule-associated serine/threonine-protein kinase 3 isoform X10 [Mus musculus] | 0 | 131.841 | 29 | 20 | 61 | 18 | 1312 | 143.2 | 8.59 | 1063 | 20 | 77007487.06 | 66384688.39 | 62423302.04 | 40243528.72 | 61503201.52 | 49006748.18 | 45446783.56 | 40382014.75 | 25.1321 | 18.3431 | Mast3 | 546071 | 1.11239 | 0.887611866 | 0.79793 | 0.21403 | 0.716066 |
| 110347404 | probable ribonuclease ZC3H12B [Mus musculus] | 0 | 15.537 | 5 | 3 | 13 | 2 | 835 | 93.8 | 7.18 | 128 | 3 | 4155512.434 | 7248433.531 | 8412943.43 | 6002719.523 | 7044816.493 | 10488312.78 | 4933988.5 | 11033911.88 | 28.2213 | 34.568 | Zc3h12b | 547176 | 0.87051 | 1.129489853 | 1.2975 | 0.30442 | 0.761857 |
| 85701578 | protein FAM196B [Mus musculus] | 0.003 | 3.856 | 4 | 2 | 4 | 2 | 535 | 59.1 | 7.84 | 0 | 2 | 3189123.826 | 2943961.359 | 3010021.277 | 3069898.235 | 3900977.93 | 4510300.141 | 3084133.125 | 4533339.656 | 3.41164 | 17.0066 | Insyn2b | 574403 | 0.86489 | 1.135110091 | 1.31243 | 0.03253 | 0.520251 |
| 4506635 | 60S ribosomal protein L32 [Homo sapiens] | 0 | 46.081 | 40 | 8 | 46 | 8 | 135 | 15.9 | 11.33 | 698 | 8 | 96689162.88 | 110673629.5 | 107542724.7 | 158297314.2 | 130770475.3 | 104701588.6 | 119309858.2 | 96223546.26 | 23.1017 | 13.6025 | Rpl32l | 621697 | 1.02402 | 0.975982294 | 0.95309 | 0.73534 | 0.943908 |
| 85677491 | docking protein 6 [Mus musculus] | 0 | 10.522 | 12 | 3 | 8 | 2 | 331 | 38.3 | 8.48 | 37 | 3 | 4451747.036 | 2941934.878 | 5149307.649 | 3671997.038 | 5108878.925 | 3785559.806 | 4752910.625 | 3302689.632 | 23.5776 | 19.7586 | Dok6 | 623279 | 0.97784 | 1.02216348 | 1.04533 | 0.78213 | 0.952792 |
| 90903233 | phospholipid hydroperoxide glutathione peroxidase, nuclear isoform 1 [Mus musculus] | 0 | 36.645 | 31 | 8 | 49 | 8 | 253 | 29.2 | 10.23 | 352 | 8 | 113842350.6 | 113866826.1 | 119564911.3 | 115502364.3 | 117328735.4 | 131623622.9 | 118293330.5 | 85909136.47 | 2.32935 | 17.11 | Gpx4 | 625249 | 1.0105 | 0.989495252 | 0.97921 | 0.81401 | 0.961921 |
| 1039779567 | PREDICTED: protein FAM196A isoform X1 [Mus musculus] | 0 | 45.028 | 25 | 9 | 27 | 9 | 488 | 53.7 | 7.58 | 324 | 9 | 13750471.93 | 12885203.96 | 14982509.65 | 13549907.11 | 22466928.66 | 21712234.28 | 17881628.88 | 20415477.71 | 6.34846 | 9.75804 | Insyn2a | 627214 | 0.8016 | 1.198396625 | 1.495 | 0.0008 | 0.336276 |
| 569001273 | PREDICTED: patched domain-containing protein 4 isoform X1 [Mus musculus] | 0 | 23.997 | 5 | 3 | 6 | 3 | 904 | 103 | 8.68 | 165 | 3 | 1769738.783 | 3114927.762 | 5090802.467 | 2882821.54 | 391940.5279 | 1033496.691 | 1725042.906 | 215812.0989 | 42.9842 | 81.4941 | Ptchd4 | 627626 | 1.58504 | 0.414961946 | 0.2618 | 0.02176 | 0.505785 |
| 126090620 | dapper homolog 3 [Mus musculus] | 0 | 118.588 | 39 | 14 | 89 | 14 | 610 | 63.2 | 10.42 | 2321 | 14 | 132729333.1 | 104262921.3 | 124178829.4 | 122933486.2 | 109723312.6 | 138861931.7 | 141714906.4 | 137768694.5 | 9.91002 | 11.3284 | Dact3 | 629378 | 0.95656 | 1.043435517 | 1.09082 | 0.29518 | 0.759552 |
| 90787451 | putative gag-pol protein [Mus musculus] | 0 | 22.333 | 5 | 5 | 18 | 3 | 1067 | 119.7 | 9.26 | 103 | 5 | 8840709.731 | 7353492.803 | 5839458.516 | 9817637.573 | 4518131.891 | 10558348.84 | 6236831.5 | 5746108.322 | 21.8614 | 38.8814 | BC094435 | 639390 | 1.08134 | 0.918658639 | 0.84955 | 0.47626 | 0.863085 |
| 256017197 | RNA-binding protein Nova-1 [Mus musculus] | 0 | 24.533 | 16 | 5 | 14 | 5 | 507 | 51.7 | 8.72 | 181 | 5 | 7050560.854 | 6530766.181 | 9435143.946 | 6667653.29 | 3200785.189 | 8067487.762 | 9098154 | 8521102.744 | 18.335 | 37.5762 | Nova1 | 664883 | 1.0136 | 0.986399657 | 0.97316 | 0.8999 | 0.985548 |
| 244792359 | traf2 and NCK-interacting protein kinase isoform 1 [Mus musculus] | 0 | 292.592 | 43 | 43 | 261 | 31 | 1360 | 154.9 | 7.09 | 3309 | 43 | 462179843.4 | 540349945.1 | 448322710.9 | 567098846.2 | 427347523.7 | 524911575.7 | 474013566.3 | 654660438 | 11.5302 | 18.8523 | Tnik | 665113 | 0.98463 | 1.015365585 | 1.03121 | 0.79169 | 0.955057 |
| 126723102 | protein EFR3 homolog B [Mus musculus] | 0 | 55.251 | 20 | 11 | 34 | 10 | 817 | 92.3 | 6.54 | 643 | 11 | 15577106.08 | 23520035.28 | 22312137.46 | 17384811.87 | 26476426.49 | 16741354.17 | 16966976.38 | 18831859.96 | 19.3917 | 23.1768 | Efr3b | 668212 | 0.99859 | 1.001410084 | 1.00282 | 0.98572 | 0.998251 |
| 262072980 | uncharacterized protein C19orf48 homolog [Mus musculus] | 0 | 62.477 | 46 | 8 | 32 | 8 | 284 | 32.4 | 6.32 | 1168 | 8 | 19718127.3 | 22801574.68 | 22339918.81 | 22363247.9 | 27317253.04 | 23431215.44 | 31122279.63 | 20780255.45 | 6.45625 | 17.6233 | 10002F23F | 668661 | 0.91875 | 1.081254649 | 1.17688 | 0.15452 | 0.698266 |
| 186915027 | Rap1A-retro1 [Mus musculus] | 0 | 10.402 | 20 | 3 | 10 | 3 | 184 | 21 | 6.67 | 95 | 3 | 2162319.765 | 2683810.706 | 6407731.901 | 6213845.494 | 7836251.992 | 10296447.54 | 6947143.5 | 10978002.92 | 51.662 | 21.3969 | Gm9392 | 668838 | 0.65269 | 1.347313325 | 2.06426 | 0.02028 | 0.505785 |
| 148696314 | mCG1040076 [Mus musculus] | 0 | 34.19 | 32 | 7 | 67 | 7 | 148 | 16.6 | 10.84 | 328 | 7 | 189651758.3 | 185855892 | 176094811.4 | 232558231.9 | 134094216.6 | 187650358.9 | 181204797.5 | 183603570.3 | 12.7556 | 14.6647 | Gm14044 | 668899 | 1.06637 | 0.933632389 | 0.87553 | 0.21811 | 0.716066 |
| 148698365 | mCG120681 [Mus musculus] | 0 | 15.033 | 29 | 4 | 67 | 4 | 137 | 15.7 | 11.88 | 435 | 4 | 95512074.37 | 109305391.6 | 74264956.13 | 93769444.34 | 81075781.02 | 88700230.36 | 87473321.62 | 80055876.52 | 15.4672 | 5.20692 | Rpl28-ps3 | 100039645 | 1.05005 | 0.94994536 | 0.90466 | 0.28292 | 0.750719 |
| 148668546 | mCG114988 [Mus musculus] | 0 | 40.103 | 57 | 10 | 150 | 10 | 152 | 17.7 | 10.86 | 2951 | 10 | 221024263.4 | 254358113.9 | 248957438 | 317194982 | 269082264.1 | 311915094.9 | 277863988.2 | 298739542.9 | 15.59 | 6.736 | Gm10260 | 100039740 | 0.94722 | 1.05277805 | 1.11144 | 0.24495 | 0.733806 |
| 568910344 | PREDICTED: immunoglobulin-like domain-containing receptor 2 isoform X3 [Mus musculus] | 0 | 184.102 | 48 | 22 | 222 | 22 | 602 | 67.5 | 7.66 | 3710 | 22 | 297811282.7 | 351050116.2 | 307295066.4 | 403869752.6 | 330535327 | 388523058.7 | 326460178.1 | 370941578.5 | 14.2586 | 8.60853 | Ildr2 | 100039795 | 0.97967 | 1.020325662 | 1.04149 | 0.63972 | 0.915433 |
| 148692922 | mCG1034428 [Mus musculus] | 0 | 54.89 | 44 | 10 | 99 | 10 | 156 | 17.7 | 10.42 | 965 | 10 | 217886744.1 | 258613025.5 | 252699068.3 | 275751979.8 | 232580168.9 | 297669857.1 | 246021951.2 | 331684836.7 | 9.66747 | 16.6103 | Rpl23a-ps3 | 100041163 | 0.95125 | 1.048750828 | 1.1025 | 0.36043 | 0.803491 |
| 1039742895 | PREDICTED: LOW QUALITY PROTEIN: protein AHNAK2 [Mus musculus] | 0 | 115.697 | 42 | 15 | 55 | 15 | 3501 | 368.2 | 5.86 | 1362 | 15 | 100653896.8 | 125069446.1 | 107935971.1 | 86205316.56 | 91163581.99 | 167252018.8 | 131621934 | 132379308.1 | 15.3963 | 23.8173 | Ahnak2 | 100041194 | 0.89117 | 1.108833947 | 1.24425 | 0.19385 | 0.715071 |
| 148696763 | mCG1048875, isoform CRA_c [Mus musculus] | 0 | 175.221 | 55 | 22 | 336 | 22 | 452 | 49.5 | 7.56 | 5327 | 22 | 521603723.7 | 451474426 | 533266073.3 | 562682452.8 | 501953407.6 | 372189237.3 | 525839261.6 | 428462151.9 | 9.1131 | 15.3484 | Gm9755 | 100041265 | 1.06173 | 0.938272117 | 0.88372 | 0.20454 | 0.715071 |
| 569002888 | PREDICTED: pro-neuregulin-2, membrane-bound isoform isoform X1 [Mus musculus] | 0 | 26.082 | 8 | 6 | 28 | 6 | 859 | 92.6 | 9.28 | 330 | 6 | 3821387.745 | 4214008.896 | 7535595.726 | 13136497.98 | 30541737.53 | 9727720.68 | 8883619.875 | 8246463.251 | 60.03 | 75.3428 | Nrg2 | 100042150 | 0.66679 | 1.333213797 | 1.99946 | 0.26381 | 0.745606 |
| 755565071 | PREDICTED: NHS-like protein 2 isoform X1 [Mus musculus] | 0 | 124.231 | 24 | 18 | 63 | 18 | 1242 | 134.7 | 8.51 | 1297 | 18 | 56248999.19 | 50410541.91 | 44259982.44 | 53033241.35 | 63561960.19 | 41960439.74 | 56390638.13 | 43120131.2 | 9.96577 | 20.4737 | Nhsl2 | 100042480 | 0.99736 | 1.002641666 | 1.0053 | 0.96455 | 0.996581 |
| 148670962 | mCG19976 [Mus musculus] | 0 | 28.228 | 27 | 4 | 20 | 4 | 160 | 18.7 | 10.55 | 370 | 4 | 142644066.6 | 145510949.8 | 117038018.3 | 175467745.6 | 123089622.9 | 123367230.8 | 148211802 | 108260274.8 | 16.4747 | 13.1747 | Gm10095 | 100042650 | 1.07174 | 0.9282645 | 0.86613 | 0.23 | 0.723653 |
| 148666048 | mCG1038069, partial [Mus musculus] | 0 | 17.795 | 26 | 4 | 22 | 4 | 206 | 23.2 | 7.88 | 144 | 4 | 10053311.49 | 11303755.19 | 13254605.27 | 9708510.805 | 15893091.8 | 7972096.257 | 10639892.5 | 6788451.711 | 14.4723 | 39.2069 | - | 100046119 | 1.03535 | 0.964647598 | 0.93171 | 0.74 | 0.943908 |
| 152002336 | neuroligin 4* [Mus musculus] | 0 | 37.166 | 12 | 8 | 37 | 5 | 945 | 97.2 | 6.34 | 540 | 8 | 40014315.49 | 31036378.36 | 30943306.21 | 10791637.41 | 24881408.99 | 13143778.41 | 25922428 | 32509749.91 | 43.8302 | 33.4072 | Nlgn4l | 100113365 | 1.07803 | 0.921965027 | 0.85523 | 0.59998 | 0.905186 |
| 12846616 | unnamed protein product [Mus musculus] | 0 | 69.328 | 69 | 8 | 48 | 8 | 147 | 15.8 | 7.69 | 858 | 8 | 127119302.5 | 128907919.8 | 133798447.7 | 121030264.6 | 152823647.1 | 112181660.5 | 145891033 | 106220521.6 | 4.1302 | 18.1644 | Hbb-bs | 100503605 | 0.99391 | 1.006090558 | 1.01226 | 0.90076 | 0.98561 |
| 568966130 | EDICTED: voltage-dependent calcium channel beta subunit-associated regulatory protein isoform X2 [Mus muscu | 0 | 12.906 | 6 | 3 | 6 | 3 | 716 | 76 | 6.33 | 48 | 3 | 3044955.101 | 2285293.788 | 2472864.566 | 2487061.425 | 2920276.607 | 3770621.909 | 3074051.75 | 3282149.73 | 12.7535 | 11.3504 | Cbarp | 100503659 | 0.88187 | 1.118133978 | 1.26792 | 0.03172 | 0.515682 |
| 23956082 | 60S ribosomal protein L5 [Mus musculus] | 0 | 69.427 | 39 | 12 | 128 | 12 | 297 | 34.4 | 9.77 | 978 | 12 | 239341396.5 | 250871584.7 | 212149406.1 | 336381190.9 | 196700799.6 | 222799491.6 | 214354914.7 | 201531787.7 | 20.6578 | 5.70614 | Rpl5 | 100503670 | 1.10851 | 0.891492841 | 0.80423 | 0.11375 | 0.678142 |
| 148694624 | mCG10153 [Mus musculus] | 0 | 53.393 | 58 | 6 | 98 | 6 | 115 | 12.8 | 9.63 | 1164 | 6 | 170947856.7 | 173530049 | 176049600.7 | 205042418.9 | 172897124.8 | 178745409.8 | 207153593.8 | 175777704 | 8.76751 | 8.63317 | Rpl30-ps1( | 100504405 | 0.99383 | 1.006166452 | 1.01241 | 0.84773 | 0.973079 |

**Supplementary Table 4 LC-MS/MS quantification and comparison of 1,960 PSD proteins in seven brain regions in adult marmoset**

| Description of protein | | Detection by LC-MS/MS | | | | | | | | | | | |
|---|---|---|---|---|---|---|---|---|---|---|---|---|---|
| GI Number | Description | -value: Com | PEP Sc | coverage [% | # Peptides | # PSMs | nique Pepti | # AAs | MW [kDa] | calc. pI | Mascot: M | Search En | NCX1 … |
| 296206333 | PREDICTED: abl interactor 1 isoform X1 [Callithrix jacchus] | 0 | 126.05 | 37 | 16 | 140 | 11 | 508 | 55.1 | 7.06 | 1805 | 16 | 18949838.57 … [illegible] |
| 532503413 | very long-chain specific acyl-CoA dehydrogenase, mitochondrial isoform 1 precursor [Callithrix jacchus] | 0 | 116.328 | 34 | 19 | 108 | 19 | 655 | 70.2 | 8.76 | 1734 | 19 | 32886811.83 … [illegible] |
| 1061016719 | PREDICTED: LOW QUALITY PROTEIN: acid-sensing ion channel 1 [Callithrix jacchus] | 0 | 40.843 | 18 | 10 | 70 | 9 | 528 | 59.8 | 5.73 | 238 | 10 | 17006792.17 … [illegible] |
| 1060937427 | PREDICTED: aconitate hydratase, mitochondrial [Callithrix jacchus] | 0 | 264.801 | 44 | 34 | 409 | 34 | 857 | 94.2 | 7.91 | 6774 | 34 | 185066745.3 … [illegible] |
| 390474818 | PREDICTED: low molecular weight phosphotyrosine protein phosphatase isoform X1 [Callithrix jacchus] | 0 | 12.832 | 21 | 3 | 11 | 3 | 158 | 18 | 6.32 | 78 | 3 | 1041971.353 … [illegible] |
| 675750393 | PREDICTED: alpha-actinin-2 [Callithrix jacchus] | 0 | 500.593 | 70 | 56 | 879 | 37 | 894 | 103.8 | 5.45 | 14765 | 56 | 233464841.8 … [illegible] |
| 675709781 | PREDICTED: actin, aortic smooth muscle [Callithrix jacchus] | 0 | 428.793 | 85 | 27 | 3176 | 3 | 377 | 42 | 5.39 | 44097 | 27 | 5136083.964 … [illegible] |
| 532273828 | aspartoacylase [Callithrix jacchus] | 0 | 42.589 | 36 | 9 | 31 | 9 | 313 | 35.6 | 6.92 | 173 | 9 | 17847031.68 … [illegible] |
| 532281686 | disintegrin and metalloproteinase domain-containing protein 11 preproprotein [Callithrix jacchus] | 0 | 53.441 | 16 | 8 | 31 | 8 | 769 | 83.6 | 7.34 | 435 | 8 | 3835518.276 … [illegible] |
| 675673469 | PREDICTED: disintegrin and metalloproteinase domain-containing protein 22 isoform X1 [Callithrix jacchus] | 0 | 258.002 | 31 | 26 | 208 | 26 | 1024 | 113.5 | 7.14 | 3480 | 26 | 42233125.08 … [illegible] |
| 296227203 | PREDICTED: LOW QUALITY PROTEIN: adenylate cyclase type 8 [Callithrix jacchus] | 0.009 | 3.767 | 1 | 2 | 10 | 2 | 1252 | 140.1 | 6.99 | 16 | 2 | 1276829.549 … [illegible] |
| 296223606 | PREDICTED: beta-adducin [Callithrix jacchus] | 0 | 393.519 | 57 | 36 | 712 | 34 | 726 | 80.8 | 5.92 | 13673 | 36 | 379931280.1 … [illegible] |
| 296230268 | PREDICTED: poly [ADP-ribose] polymerase 1 [Callithrix jacchus] | 0 | 120.307 | 28 | 23 | 121 | 23 | 1013 | 113 | 8.9 | 515 | 23 | 14671649.81 … [illegible] |
| 675680730 | PREDICTED: advillin isoform X1 [Callithrix jacchus] | 0 | 49.995 | 16 | 11 | 23 | 11 | 862 | 96.6 | 5.8 | 196 | 11 | 37607299.34 … [illegible] |
| 1061002891 | PREDICTED: LOW QUALITY PROTEIN: aggrecan core protein [Callithrix jacchus] | 0 | 261.202 | 19 | 32 | 311 | 32 | 2264 | 234.8 | 4.31 | 3876 | 32 | 115969918.5 … [illegible] |
| 296218936 | PREDICTED: AH receptor-interacting protein isoform X1 [Callithrix jacchus] | 0 | 13.6 | 14 | 4 | 30 | 4 | 330 | 37.5 | 6.54 | 193 | 4 | 5717051.798 … [illegible] |
| 532524482 | adenylate kinase isoenzyme 1 [Callithrix jacchus] | 0 | 119.425 | 67 | 14 | 185 | 13 | 194 | 21.6 | 8.62 | 2694 | 14 | 29587223.29 … [illegible] |
| 296196349 | PREDICTED: serum albumin [Callithrix jacchus] | 0 | 161.62 | 44 | 26 | 236 | 26 | 608 | 69.4 | 5.99 | 2195 | 26 | 16015128.11 … [illegible] |
| 1061018166 | PREDICTED: aldehyde dehydrogenase, mitochondrial [Callithrix jacchus] | 0 | 84.885 | 35 | 14 | 94 | 14 | 517 | 56.2 | 6.65 | 1095 | 14 | 22167645.54 … [illegible] |
| 1060946290 | PREDICTED: fructose-bisphosphate aldolase A isoform X2 [Callithrix jacchus] | 0 | 257.175 | 84 | 28 | 558 | 22 | 364 | 39.4 | 8.24 | 8542 | 28 | 158990083.2 … [illegible] |
| 296202196 | PREDICTED: fructose-bisphosphate aldolase C [Callithrix jacchus] | 0 | 227.901 | 76 | 26 | 428 | 20 | 364 | 39.4 | 6.44 | 8828 | 26 | 63783736.68 … [illegible] |
| 1060950263 | PREDICTED: ankyrin-1 isoform X3 [Callithrix jacchus] | 0 | 394.697 | 42 | 62 | 497 | 56 | 1908 | 209.1 | 6.73 | 5780 | 62 | 41955118.79 … [illegible] |
| 532519378 | ankyrin-3 isoform 1 [Callithrix jacchus] | 0 | 895.288 | 33 | 114 | 1515 | 69 | 4382 | 480.6 | 6.52 | 21663 | 114 | 155419469.7 … [illegible] |
| 675657852 | PREDICTED: rabankyrin-5 [Callithrix jacchus] | 0 | 42.989 | 12 | 10 | 36 | 10 | 1169 | 128.7 | 6.06 | 60 | 10 | 2271862.483 … [illegible] |
| 296213563 | PREDICTED: acidic leucine-rich nuclear phosphoprotein 32 family member A [Callithrix jacchus] | 0 | 56.006 | 38 | 12 | 108 | 6 | 249 | 28.6 | 4.09 | 1775 | 12 | 35188708.69 … [illegible] |
| 532520188 | ADP/ATP translocase 1 [Callithrix jacchus] | 0 | 162.852 | 61 | 21 | 608 | 6 | 298 | 33 | 9.76 | 7010 | 21 | 83537989.92 … [illegible] |
| 296236267 | PREDICTED: ADP/ATP translocase 2 [Callithrix jacchus] | 0 | 196.106 | 58 | 22 | 641 | 9 | 298 | 32.9 | 9.74 | 8163 | 22 | 199808615.8 … [illegible] |
| 296195660 | PREDICTED: annexin A5 [Callithrix jacchus] | 0 | 28.263 | 22 | 7 | 25 | 7 | 321 | 36.1 | 5.11 | 195 | 7 | 8180776.045 … [illegible] |
| 532503130 | annexin A6 isoform 1 [Callithrix jacchus] | 0 | 114.185 | 37 | 23 | 123 | 23 | 673 | 75.8 | 5.55 | 1022 | 23 | 25555586.33 … [illegible] |
| 675708723 | PREDICTED: annexin A7 isoform X1 [Callithrix jacchus] | 0 | 22.784 | 14 | 6 | 24 | 6 | 491 | 53 | 6.29 | 119 | 6 | 2868345.901 … [illegible] |
| 296221338 | PREDICTED: thioredoxin-dependent peroxide reductase, mitochondrial [Callithrix jacchus] | 0 | 31.756 | 34 | 6 | 44 | 6 | 256 | 28 | 7.83 | 497 | 6 | 43368264.58 … [illegible] |
| 296229779 | PREDICTED: peroxiredoxin-6 [Callithrix jacchus] | 0 | 67.92 | 61 | 10 | 49 | 10 | 224 | 24.9 | 6.01 | 321 | 10 | 8256052.087 … [illegible] |
| 675752514 | PREDICTED: AP-1 complex subunit gamma-1 isoform X1 [Callithrix jacchus] | 0 | 87.629 | 22 | 15 | 127 | 15 | 825 | 91.7 | 6.8 | 1043 | 15 | 35581932.39 … [illegible] |
| 675642271 | PREDICTED: AP-1 complex subunit sigma-1A isoform X1 [Callithrix jacchus] | 0 | 25.631 | 31 | 5 | 41 | 3 | 161 | 19.1 | 5.73 | 1129 | 5 | 13916389.81 … [illegible] |
| 1060971792 | PREDICTED: AP-2 complex subunit alpha-1 isoform X2 [Callithrix jacchus] | 0 | 592.06 | 71 | 53 | 1722 | 9 | 955 | 105.3 | 7.66 | 31977 | 53 | 519385163.2 … [illegible] |
| 532291188 | AP-2 complex subunit alpha-2 isoform 2 [Callithrix jacchus] | 0 | 605.542 | 62 | 50 | 1507 | 6 | 939 | 103.9 | 6.96 | 29251 | 50 | 33501410.39 … [illegible] |
| 296224700 | PREDICTED: AP-2 complex subunit mu isoform X1 [Callithrix jacchus] | 0 | 363.271 | 69 | 28 | 774 | 28 | 435 | 49.6 | 9.54 | 15727 | 28 | 1239827568 … [illegible] |
| 296204176 | PREDICTED: AP-3 complex subunit beta-2 isoform X2 [Callithrix jacchus] | 0 | 200.055 | 35 | 31 | 256 | 27 | 1084 | 119.3 | 5.59 | 4643 | 31 | 58896026.45 … [illegible] |
| 532282834 | AP-3 complex subunit delta-1 isoform 3 [Callithrix jacchus] | 0 | 201.212 | 27 | 25 | 245 | 25 | 1217 | 136.3 | 7.44 | 3421 | 25 | 54234656.2 … [illegible] |
| 296193924 | PREDICTED: AP-3 complex subunit sigma-1 isoform X2 [Callithrix jacchus] | 0 | 28.252 | 33 | 6 | 45 | 6 | 193 | 21.7 | 5.39 | 521 | 6 | 5462987.02 … [illegible] |
| 532347965 | amyloid beta A4 precursor protein-binding family A member 2 isoform a [Callithrix jacchus] | 0 | 27.625 | 10 | 5 | 23 | 4 | 751 | 82.5 | 4.92 | 434 | 5 | 3066768.026 … [illegible] |
| 1060942657 | PREDICTED: amyloid beta A4 precursor protein-binding family B member 1 [Callithrix jacchus] | 0 | 62.315 | 21 | 10 | 47 | 10 | 711 | 77.3 | 5.06 | 648 | 10 | 1325703.853 … [illegible] |

[illegible]

| | | | | | | | | | | | | | | |
|---|---|---|---|---|---|---|---|---|---|---|---|---|---|---|
| 1060933597 | PREDICTED: cytochrome c oxidase subunit NDUFA4-like [Callithrix jacchus] | 0 | 33.425 | 56 | 6 | 178 | 6 | 82 | 9.4 | 9.5 | 1715 | 6 | 112388363.3 | 325892175.6 |
| 296194616 | PREDICTED: NADH dehydrogenase [ubiquinone] iron-sulfur protein 4, mitochondrial isoform X1 [Callithrix jacchus] | 0 | 94.487 | 35 | 6 | 56 | 6 | 175 | 20.1 | 10.04 | 1061 | 6 | 81374697.07 | 75195377.58 |
| 296218952 | PREDICTED: NADH dehydrogenase [ubiquinone] flavoprotein 1, mitochondrial isoform X1 [Callithrix jacchus] | 0 | 293.064 | 74 | 27 | 315 | 27 | 464 | 50.6 | 8.07 | 5210 | 27 | 660853698.4 | 392580026.6 |
| 675666542 | PREDICTED: septin-2 isoform X1 [Callithrix jacchus] | 0 | 351.988 | 80 | 24 | 522 | 23 | 379 | 43.3 | 6.48 | 11118 | 24 | 577564064.5 | 558348059.3 |
| 1060996977 | PREDICTED: neurofibromin isoform X3 [Callithrix jacchus] | 0 | 346.63 | 25 | 63 | 370 | 63 | 2857 | 321.3 | 7.42 | 3668 | 63 | 76103308.78 | 63935642.28 |
| 296221730 | PREDICTED: neurofilament light polypeptide [Callithrix jacchus] | 0 | 522.558 | 73 | 44 | 976 | 40 | 543 | 61.5 | 4.68 | 17375 | 44 | 751007740.6 | 821963373.5 |
| 1060949795 | PREDICTED: neurofilament medium polypeptide isoform X1 [Callithrix jacchus] | 0 | 492.085 | 59 | 52 | 1175 | 47 | 921 | 103.2 | 4.98 | 14589 | 52 | 1300422600 | 951126896.4 |
| 296191340 | PREDICTED: 39S ribosomal protein L40, mitochondrial [Callithrix jacchus] | 0 | 38.87 | 39 | 6 | 33 | 6 | 207 | 24.5 | 9.69 | 360 | 6 | 17680508.24 | 13104735.35 |
| 325910861 | nucleoside diphosphate kinase A [Callithrix jacchus] | 0 | 113.957 | 66 | 11 | 294 | 6 | 152 | 17.2 | 6.54 | 4270 | 11 | 54495067.5 | 56260840.44 |
| 325910863 | nucleoside diphosphate kinase B [Callithrix jacchus] | 0 | 115.727 | 69 | 10 | 295 | 5 | 152 | 17.3 | 8.41 | 4585 | 10 | 408143172.5 | 337897632.4 |
| 296201692 | PREDICTED: glycylpeptide N-tetradecanoyltransferase 1 isoform X1 [Callithrix jacchus] | 0 | 17.834 | 9 | 3 | 11 | 3 | 496 | 56.9 | 7.99 | 60 | 3 | 2574842.687 | 3215444.904 |
| 296194659 | PREDICTED: NAD(P) transhydrogenase, mitochondrial [Callithrix jacchus] | 0 | 190.906 | 34 | 29 | 187 | 29 | 1086 | 113.8 | 8.28 | 2301 | 29 | 42599454.34 | 65679116.3 |
| 532504078 | 39S ribosomal protein L49, mitochondrial [Callithrix jacchus] | 0 | 28.065 | 27 | 4 | 42 | 4 | 166 | 19.1 | 9 | 954 | 4 | 5519338.862 | 10984253.69 |
| 532515720 | nitric oxide synthase, brain isoform 1 [Callithrix jacchus] | 0 | 91.167 | 19 | 21 | 80 | 21 | 1434 | 160.9 | 7.34 | 717 | 21 | 12261793.38 | 4237166.496 |
| 532291672 | Niemann-Pick C1 protein precursor [Callithrix jacchus] | 0 | 19.203 | 5 | 5 | 22 | 5 | 1277 | 142 | 5.3 | 155 | 5 | 8469804.833 | 1647709.143 |
| 296193667 | PREDICTED: nucleophosmin [Callithrix jacchus] | 0 | 270.642 | 60 | 16 | 392 | 16 | 296 | 32.8 | 4.74 | 10716 | 16 | 387250447.3 | 297727544.3 |
| 532510748 | catenin delta-2 [Callithrix jacchus] | 0 | 498.24 | 54 | 50 | 801 | 45 | 1225 | 132.6 | 7.75 | 12943 | 50 | 169539244.5 | 164548099 |
| 675661613 | PREDICTED: neuronal pentraxin-1 [Callithrix jacchus] | 0 | 102.466 | 40 | 15 | 108 | 14 | 432 | 47.1 | 6.55 | 2187 | 15 | 41537535.9 | 65300231.23 |
| 1060950908 | PREDICTED: LOW QUALITY PROTEIN: neurexin-1 [Callithrix jacchus] | 0 | 324.756 | 33 | 37 | 362 | 31 | 1467 | 161.1 | 6.13 | 5032 | 37 | 137123238.5 | 89990901.22 |
| 1060940948 | PREDICTED: neurexin 3 [Callithrix jacchus] | 0 | 398.583 | 31 | 39 | 456 | 35 | 1615 | 177.9 | 5.85 | 8228 | 39 | 126590588.3 | 151428823.8 |
| 532524258 | vesicle-fusing ATPase [Callithrix jacchus] | 0 | 521.039 | 72 | 55 | 1378 | 55 | 744 | 82.6 | 6.95 | 21467 | 55 | 1291249310 | 1136489526 |
| 675663148 | PREDICTED: NT-3 growth factor receptor isoform X7 [Callithrix jacchus] | 0 | 32.32 | 10 | 7 | 30 | 5 | 864 | 97.1 | 6.74 | 216 | 7 | 1914686.066 | 11203090.45 |
| 532213481 | protein numb homolog isoform 1 [Callithrix jacchus] | 0 | 120.681 | 31 | 17 | 184 | 13 | 652 | 70.8 | 8.5 | 1378 | 17 | 51623467.78 | 48273528.14 |
| 532276178 | numb-like protein [Callithrix jacchus] | 0 | 104.319 | 34 | 18 | 180 | 14 | 598 | 63.6 | 8.85 | 1828 | 18 | 45452937.08 | 45450086.8 |
| 296202091 | PREDICTED: oligodendrocyte-myelin glycoprotein [Callithrix jacchus] | 0 | 85.655 | 21 | 11 | 289 | 11 | 440 | 49.7 | 8.4 | 4017 | 11 | 684984293.4 | 590437612.4 |
| 296195612 | PREDICTED: heat shock 70 kDa protein 4L [Callithrix jacchus] | 0 | 274.9 | 55 | 37 | 328 | 33 | 840 | 94.5 | 5.81 | 5128 | 37 | 54486670.86 | 251491420.7 |
| 296227579 | PREDICTED: claudin-11 [Callithrix jacchus] | 0 | 101.654 | 36 | 5 | 521 | 5 | 207 | 21.9 | 7.72 | 11524 | 5 | 1122885317 | 2005001824 |
| 296213129 | PREDICTED: P2X purinoceptor 7 isoform X1 [Callithrix jacchus] | 0 | 68.479 | 20 | 9 | 53 | 9 | 595 | 68.6 | 8.31 | 389 | 9 | 11665460.08 | 15782944.97 |
| 675777894 | PREDICTED: protein disulfide-isomerase isoform X1 [Callithrix jacchus] | 0 | 29.098 | 14 | 8 | 58 | 8 | 531 | 59.5 | 4.91 | 410 | 8 | 5924392.082 | 6733016.826 |
| 532218439 | polyadenylate-binding protein 1 [Callithrix jacchus] | 0 | 217.856 | 42 | 28 | 399 | 19 | 636 | 70.7 | 9.5 | 4651 | 28 | 161308791.4 | 195490817 |
| 532507082 | platelet-activating factor acetylhydrolase IB subunit beta isoform a [Callithrix jacchus] | 0 | 13.766 | 12 | 2 | 25 | 2 | 229 | 25.5 | 6.21 | 305 | 2 | 6438089.647 | 3243591.655 |
| 1061011331 | PREDICTED: peroxiredoxin-1 [Callithrix jacchus] | 0 | 233.39 | 92 | 19 | 540 | 16 | 199 | 22.1 | 8.05 | 7057 | 19 | 370253060.4 | 305980530.2 |
| 532519804 | serine/threonine-protein kinase PAK 1 isoform 2 [Callithrix jacchus] | 0 | 83.904 | 27 | 12 | 80 | 6 | 547 | 60.8 | 5.58 | 1042 | 12 | 2476598.508 | 8466209.671 |
| 675769253 | PREDICTED: serine/threonine-protein kinase PAK 3 isoform X2 [Callithrix jacchus] | 0 | 87.036 | 30 | 14 | 75 | 8 | 559 | 62.3 | 5.45 | 845 | 14 | 3984578.877 | 5251875.468 |
| 532199877 | paralemmin-1 isoform 1 [Callithrix jacchus] | 0 | 300.139 | 64 | 24 | 483 | 23 | 379 | 41.4 | 4.92 | 17058 | 24 | 168668022.8 | 248414603.8 |
| 1061016920 | PREDICTED: LOW QUALITY PROTEIN: poly(rC)-binding protein 2 [Callithrix jacchus] | 0 | 148.245 | 38 | 12 | 196 | 8 | 466 | 48.6 | 7.64 | 4286 | 12 | 27627914.26 | 32224021.73 |
| 532284048 | protocadherin-10 isoform 1 precursor [Callithrix jacchus] | 0 | 132.845 | 24 | 20 | 143 | 20 | 1041 | 112.9 | 4.87 | 2624 | 20 | 48172356.8 | 23306378.33 |
| 296199435 | PREDICTED: protein-L-isoaspartate(D-aspartate) O-methyltransferase isoform X1 [Callithrix jacchus] | 0 | 154.747 | 53 | 12 | 202 | 12 | 286 | 30.3 | 6.73 | 3206 | 12 | 36689836.55 | 32589153.75 |
| 532503730 | cyclin-dependent kinase 18 isoform b [Callithrix jacchus] | 0 | 51.536 | 34 | 14 | 81 | 12 | 474 | 54.5 | 8.73 | 556 | 14 | 16900801.56 | 19052717.8 |
| 532290248 | pyruvate carboxylase, mitochondrial precursor [Callithrix jacchus] | 0 | 130.383 | 27 | 24 | 143 | 24 | 1178 | 129.5 | 6.83 | 1813 | 24 | 153261036.8 | 64170753.16 |
| 532501925 | programmed cell death protein 6 isoform 1 [Callithrix jacchus] | 0 | 17.122 | 23 | 4 | 22 | 4 | 191 | 21.8 | 5.4 | 194 | 4 | 10262017.68 | 15322656.43 |
| 296211904 | PREDICTED: calcium/calmodulin-dependent 3',5'-cyclic nucleotide phosphodiesterase 1B isoform X1 [Callithrix jacchus] | 0 | 95.948 | 35 | 16 | 81 | 14 | 536 | 61.3 | 5.58 | 885 | 16 | 18674046.78 | 11277695.61 |
| 1061020067 | PREDICTED: pyruvate dehydrogenase E1 component subunit alpha, somatic form, mitochondrial [Callithrix jacchus] | 0 | 236.583 | 62 | 20 | 563 | 16 | 390 | 43.3 | 8.06 | 8156 | 20 | 645887524.5 | 595123629.5 |
| 532504907 | protein-arginine deiminase type-2 [Callithrix jacchus] | 0 | 165.906 | 40 | 19 | 236 | 19 | 665 | 75.4 | 5.6 | 5769 | 19 | 102371425.4 | 176843292.3 |
| 675660315 | PREDICTED: LOW QUALITY PROTEIN: pyruvate dehydrogenase kinase, isozyme 2 [Callithrix jacchus] | 0 | 80.325 | 32 | 10 | 73 | 9 | 414 | 46.8 | 6.74 | 758 | 10 | 31131339.33 | 52170711.14 |
| 1060945431 | PREDICTED: 3-phosphoinositide-dependent protein kinase 1 isoform X1 [Callithrix jacchus] | 0 | 17.223 | 9 | 5 | 19 | 5 | 650 | 73.6 | 8.13 | 185 | 5 | 726143.4387 | 1843642.559 |

| | | | | | | | | | | | | | | | | | |
|---|---|---|---|---|---|---|---|---|---|---|---|---|---|---|---|---|---|
| 1060997397 | PREDICTED: misshapen-like kinase 1 isoform X1 [Callithrix jacchus] | 0 | 222.594 | 36 | 43 | 344 | 2 | 1367 | 154.1 | 7.55 | 3646 | 43 | 3234084.199 | 2774114.29 | 3493489.008 | 1603109.179 | [illegible] |
| 1060995551 | PREDICTED: MAGUK p55 subfamily member 2 isoform X1 [Callithrix jacchus] | 0 | 173.899 | 47 | 22 | 260 | 21 | 569 | 63.2 | 6.1 | 3253 | 22 | 110401933.1 | 94327689.34 | 104989340.9 | 76905002.87 | [illegible] |
| 296234514 | PREDICTED: serine/threonine-protein phosphatase 2A 65 kDa regulatory subunit A alpha isoform [Callithrix jacchus] | 0 | 357.203 | 63 | 31 | 553 | 31 | 589 | 65.3 | 5.11 | 11104 | 31 | 226135426.4 | 315109462.1 | 322492843.6 | 251626119.9 | [illegible] |
| 1061011077 | PREDICTED: CTP synthase 1 [Callithrix jacchus] | 0 | 155.472 | 46 | 23 | 246 | 19 | 591 | 66.7 | 6.54 | 3207 | 23 | 46650922.54 | 61378818.73 | 52108048.21 | 68120013.81 | [illegible] |
| 532513516 | RUN domain-containing protein 3A isoform 2 [Callithrix jacchus] | 0 | 109.81 | 43 | 10 | 67 | 10 | 405 | 45.7 | 5.26 | 1284 | 10 | 22959616.95 | 25471759.12 | 23135068.07 | 15098611.1 | [illegible] |
| 532287088 | heterogeneous nuclear ribonucleoprotein U isoform a [Callithrix jacchus] | 0 | 167.439 | 33 | 24 | 246 | 24 | 825 | 90.6 | 6.01 | 3492 | 24 | 93245125.81 | 85940216.85 | 89472579.15 | 62759450.67 | [illegible] |
| 1060944358 | PREDICTED: serine/threonine-protein phosphatase 6 regulatory subunit 3 isoform X1 [Callithrix jacchus] | 0 | 30 | 11 | 8 | 25 | 8 | 901 | 101 | 4.68 | 89 | 8 | 16477282.75 | 4294183.683 | 5421241.32 | 1996221.676 | [illegible] |
| 675720865 | PREDICTED: rab11 family-interacting protein 5 isoform X1 [Callithrix jacchus] | 0 | 184.322 | 27 | 24 | 216 | 24 | 1309 | 137.5 | 5.14 | 5043 | 24 | 94440813.76 | 99352681.89 | 91673046.85 | 103198651.5 | [illegible] |
| 1060955003 | PREDICTED: LOW QUALITY PROTEIN: calcium/calmodulin-dependent protein kinase type 1 [Callithrix jacchus] | 0 | 36.935 | 15 | 7 | 43 | 3 | 537 | 58.7 | 6.98 | 990 | 7 | 2348218.173 | 4500721.531 | 5250804.333 | 1544106.465 | [illegible] |
| 675648939 | PREDICTED: septin-11 isoform X1 [Callithrix jacchus] | 0 | 358.556 | 64 | 32 | 1003 | 17 | 437 | 50.3 | 6.39 | 17156 | 32 | 697879144.7 | 655102974.7 | 667725107.3 | 790507161 | [illegible] |
| 675712749 | [illegible]ICTED: serine/threonine-protein phosphatase 2A 55 kDa regulatory subunit B delta isoform isoform X1 [Callithrix jac[illegible] | 0 | 52.51 | 23 | 8 | 51 | 3 | 475 | 54.6 | 7.01 | 273 | 8 | 2344598.738 | 3422046.973 | 3359798.732 | 3928748.604 | [illegible] |
| 296217097 | PREDICTED: 39S ribosomal protein L48, mitochondrial isoform X1 [Callithrix jacchus] | 0 | 27.302 | 18 | 4 | 31 | 4 | 261 | 29.2 | 9.35 | 455 | 4 | 5786380.438 | 18560750.23 | 12413919.72 | 13546120.2 | [illegible] |
| 296222651 | PREDICTED: LOW QUALITY PROTEIN: 3-ketoacyl-CoA thiolase, mitochondrial [Callithrix jacchus] | 0 | 74.766 | 47 | 12 | 75 | 12 | 397 | 41.9 | 8.4 | 377 | 12 | 10648414.77 | 10368423.73 | 10091312.7 | 17845977.33 | [illegible] |
| 675674052 | PREDICTED: LOW QUALITY PROTEIN: extended synaptotagmin-2 [Callithrix jacchus] | 0 | 67.643 | 23 | 15 | 56 | 15 | 894 | 99.1 | 8.29 | 593 | 15 | 21245962.7 | 12511528.56 | 23437261.08 | 14354519.27 | [illegible] |
| 296220643 | PREDICTED: CDGSH iron-sulfur domain-containing protein 1 [Callithrix jacchus] | 0 | 101.243 | 50 | 7 | 201 | 7 | 108 | 12.2 | 9.09 | 3904 | 7 | 386354005.6 | 304474367.8 | 294815440.8 | 283040587 | [illegible] |
| 1060966945 | PREDICTED: probable D-lactate dehydrogenase, mitochondrial isoform X1 [Callithrix jacchus] | 0 | 30.469 | 15 | 6 | 47 | 6 | 484 | 51.7 | 6.32 | 315 | 6 | 24985585.14 | 12126497.53 | 10571057.67 | 25297722.67 | [illegible] |
| 532285020 | protein RUFY3 isoform 1 [Callithrix jacchus] | 0 | 117.53 | 39 | 24 | 216 | 22 | 620 | 70.6 | 5.66 | 2141 | 24 | 22273725.91 | 63503719.78 | 58309400.83 | 32002469.96 | [illegible] |
| 296191527 | PREDICTED: small G protein signaling modulator 1 isoform X1 [Callithrix jacchus] | 0 | 56.68 | 12 | 10 | 28 | 10 | 1147 | 129.4 | 5.83 | 430 | 10 | 19828868.09 | 24129831.56 | 21317530.8 | 5710026.548 | [illegible] |
| 296194482 | PREDICTED: regulator of G-protein signaling 7-binding protein [Callithrix jacchus] | 0 | 66.758 | 54 | 8 | 58 | 8 | 257 | 28.9 | 8.46 | 1605 | 8 | 41291734.19 | 22674796.62 | 19570565.39 | 36355794.38 | [illegible] |
| 532333879 | disks large homolog 3 isoform a [Callithrix jacchus] | 0 | 333.965 | 58 | 41 | 467 | 37 | 831 | 91.6 | 6.95 | 5495 | 41 | 138800875.8 | 164911215.5 | 171065201.7 | 122094833.9 | [illegible] |
| 532524094 | contactin-associated protein 1 precursor [Callithrix jacchus] | 0 | 573.435 | 47 | 54 | 1228 | 54 | 1377 | 155.5 | 7.08 | 27578 | 54 | 1086308505 | 1102371880 | 1125692153 | 1285970995 | [illegible] |
| 532511504 | membrane-associated progesterone receptor component 1 [Callithrix jacchus] | 0 | 46.33 | 52 | 6 | 27 | 6 | 195 | 21.7 | 4.7 | 434 | 6 | 10189322.18 | 21267531.05 | 24035162.66 | 12244270.99 | [illegible] |
| 1060988498 | PREDICTED: syntaxin-7 isoform X1 [Callithrix jacchus] | 0 | 64.079 | 49 | 9 | 134 | 9 | 261 | 29.8 | 5.55 | 3328 | 9 | 20823557.41 | 17606632.42 | 17267286.78 | 36267335.62 | [illegible] |
| 675770674 | PREDICTED: myotubularin-related protein 1 isoform X1 [Callithrix jacchus] | 0 | 67.635 | 22 | 13 | 44 | 12 | 689 | 77.3 | 6.54 | 191 | 13 | 2558425.587 | 6624391.393 | 5739966.199 | 9908859.684 | [illegible] |
| 296234064 | PREDICTED: LOW QUALITY PROTEIN: mitochondrial import receptor subunit TOM40 homolog [Callithrix jacchus] | 0 | 120.248 | 47 | 12 | 110 | 12 | 361 | 37.9 | 7.25 | 1671 | 12 | 53102844.12 | 89628074.35 | 88730788.67 | 70813330.16 | [illegible] |
| 532510219 | eukaryotic translation initiation factor 3 subunit G [Callithrix jacchus] | 0 | 63.798 | 47 | 11 | 70 | 11 | 320 | 35.6 | 6.13 | 801 | 11 | 15399723.37 | 22620441.53 | 21940626.51 | 26860290.08 | [illegible] |
| 1060956168 | PREDICTED: syntenin-1 isoform X1 [Callithrix jacchus] | 0 | 55.83 | 37 | 7 | 53 | 7 | 298 | 32.5 | 7.49 | 789 | 7 | 15679859.81 | 29940131.67 | 31570588.36 | 34866422.91 | [illegible] |
| 675672750 | PREDICTED: heterogeneous nuclear ribonucleoproteins A2/B1 isoform X1 [Callithrix jacchus] | 0 | 218.632 | 53 | 23 | 474 | 18 | 396 | 41.1 | 8.9 | 8080 | 23 | 302731192.6 | 336450873.4 | 339726564 | 137045757.6 | [illegible] |
| 1060949073 | PREDICTED: thioredoxin-like protein 1 isoform X1 [Callithrix jacchus] | 0 | 47.741 | 35 | 8 | 61 | 8 | 315 | 35.1 | 4.97 | 932 | 8 | 7509130.906 | 5286150.744 | 5631247.36 | 9228016.441 | [illegible] |
| 1060999365 | PREDICTED: exocyst complex component 7 isoform X4 [Callithrix jacchus] | 0 | 208.181 | 50 | 33 | 255 | 33 | 698 | 79.6 | 6.65 | 2893 | 33 | 78590582.86 | 116404323.6 | 101841702.9 | 55560109.87 | [illegible] |
| 296190175 | PREDICTED: dynactin subunit 3 isoform X1 [Callithrix jacchus] | 0 | 47.507 | 41 | 8 | 113 | 8 | 186 | 21.2 | 5.64 | 1312 | 8 | 43391792.22 | 35621137.08 | 31610180.21 | 34081788.65 | [illegible] |
| 675724679 | PREDICTED: hippocalcin-like protein 1 isoform X2 [Callithrix jacchus] | 0 | 77.476 | 55 | 11 | 81 | 8 | 193 | 22.3 | 5.35 | 959 | 11 | 41054791.78 | 45020374.77 | 35524439.18 | 17624523.95 | [illegible] |
| 532348827 | protein PRRC2A [Callithrix jacchus] | 0 | 42.206 | 7 | 10 | 46 | 10 | 2163 | 229 | 9.45 | 272 | 10 | 5842013.004 | 3065766.829 | 2752926.132 | 6040869.626 | [illegible] |
| 675640399 | PREDICTED: tubulin alpha-8 chain [Callithrix jacchus] | 0 | 390.17 | 61 | 24 | 1845 | 9 | 449 | 50.1 | 5.06 | 52951 | 24 | 11156744.38 | 64069126.38 | 62006259.02 | 162126136.6 | [illegible] |

[illegible]

| | | | | | | | | | | | | | | |
|---|---|---|---|---|---|---|---|---|---|---|---|---|---|---|
| 532517230 | serine/threonine-protein phosphatase 6 regulatory subunit 2 isoform 1 [Callithrix jacchus] | 0 | 48.369 | 15 | 10 | 41 | 10 | 958 | 104.2 | 4.79 | 246 | 10 | 6183571.935 | 12070704.13 |
| 532444837 | splicing factor, proline- and glutamine-rich [Callithrix jacchus] | 0 | 203.404 | 42 | 22 | 460 | 21 | 708 | 76.3 | 9.44 | 6785 | 22 | 174473678.1 | 114108620.5 |
| 296213360 | PREDICTED: unconventional myosin-Ie [Callithrix jacchus] | 0 | 69.177 | 16 | 15 | 60 | 11 | 1108 | 127 | 9.04 | 534 | 15 | 3533341.672 | 14135217.27 |
| 675661236 | PREDICTED: ATP synthase subunit d, mitochondrial isoform X1 [Callithrix jacchus] | 0 | 124.571 | 69 | 13 | 138 | 13 | 177 | 20.3 | 5.48 | 2254 | 13 | 86408051.21 | 90840519.9 |
| 675650388 | PREDICTED: regulator of G-protein signaling 12 [Callithrix jacchus] | 0 | 16.494 | 3 | 4 | 25 | 4 | 1440 | 155.7 | 7.53 | 377 | 4 | 1700674.581 | 2553547.233 |
| 296216373 | PREDICTED: vacuolar protein sorting-associated protein 11 homolog [Callithrix jacchus] | 0 | 163.33 | 39 | 28 | 168 | 28 | 941 | 107.7 | 7.01 | 1620 | 28 | 70796946.51 | 35978831.18 |
| 296216381 | PREDICTED: C2 domain-containing protein 2-like isoform X1 [Callithrix jacchus] | 0 | 105.102 | 35 | 16 | 114 | 16 | 706 | 76.5 | 7.91 | 1434 | 16 | 35825078.81 | 30428206.21 |
| 675659488 | PREDICTED: AP-2 complex subunit beta isoform X1 [Callithrix jacchus] | 0 | 824.41 | 70 | 67 | 1780 | 39 | 951 | 105.6 | 5.34 | 29781 | 67 | 1170852337 | 1050277603 |
| 532288490 | inositol-3-phosphate synthase 1 isoform 1 [Callithrix jacchus] | 0 | 18.582 | 8 | 4 | 11 | 4 | 559 | 61.1 | 5.66 | 143 | 4 | 1326833.399 | 1844599.582 |
| 296203036 | PREDICTED: cytoplasmic phosphatidylinositol transfer protein 1 isoform X1 [Callithrix jacchus] | 0 | 22.539 | 19 | 6 | 27 | 6 | 332 | 38.3 | 6.4 | 396 | 6 | 858897.325 | 263607.5864 |
| 1060936515 | PREDICTED: mitochondrial glutamate carrier 2 isoform X1 [Callithrix jacchus] | 0 | 43.69 | 37 | 11 | 142 | 7 | 315 | 33.7 | 9.11 | 1765 | 11 | 9623788.514 | 4701869.006 |
| 532442163 | threonine--tRNA ligase, mitochondrial isoform a [Callithrix jacchus] | 0 | 46.851 | 18 | 11 | 43 | 11 | 718 | 81 | 7.56 | 176 | 11 | 10067783.57 | 16067509.82 |
| 1061010434 | PREDICTED: transmembrane protein 50A isoform X1 [Callithrix jacchus] | 0.003 | 5.065 | 12 | 2 | 17 | 2 | 157 | 17.4 | 5.39 | 129 | 2 | 4875634.791 | 4460377.455 |
| 296201773 | PREDICTED: DDB1- and CUL4-associated factor 7 [Callithrix jacchus] | 0 | 25.909 | 17 | 5 | 25 | 5 | 342 | 38.9 | 5.52 | 558 | 5 | 6837847.557 | 29206549.06 |
| 296224489 | PREDICTED: protein disulfide-isomerase A6 [Callithrix jacchus] | 0 | 79.272 | 31 | 10 | 89 | 10 | 440 | 47.9 | 5.01 | 1182 | 10 | 44724379.47 | 41777078.68 |
| 296212284 | PREDICTED: cullin-associated NEDD8-dissociated protein 1 [Callithrix jacchus] | 0 | 196.541 | 32 | 32 | 251 | 32 | 1230 | 136.3 | 5.78 | 1922 | 32 | 60494373.25 | 107422282.4 |
| 532509699 | uncharacterized protein KIAA0195 [Callithrix jacchus] | 0 | 19.444 | 5 | 6 | 32 | 6 | 1360 | 151.6 | 6.47 | 439 | 6 | 8294268.537 | 6285761.029 |
| 675752497 | PREDICTED: IST1 homolog isoform X1 [Callithrix jacchus] | 0 | 133.895 | 52 | 16 | 201 | 16 | 366 | 39.9 | 5.44 | 2597 | 16 | 48890058.53 | 69828038.27 |
| 296221714 | ICTED: serine/threonine-protein phosphatase 2A 55 kDa regulatory subunit B alpha isoform isoform X1 [Callithrix jac | 0 | 128.98 | 45 | 13 | 100 | 7 | 447 | 51.7 | 6.2 | 970 | 13 | 22865507.58 | 5369735.001 |
| 296208595 | PREDICTED: calponin-3 isoform X2 [Callithrix jacchus] | 0 | 51.063 | 34 | 8 | 45 | 8 | 329 | 36.4 | 6.05 | 254 | 8 | 2368473.891 | 2020207.81 |
| 532511604 | BTB/POZ domain-containing protein 17 precursor [Callithrix jacchus] | 0 | 159.09 | 54 | 17 | 218 | 17 | 478 | 52.4 | 9.13 | 3623 | 17 | 36459241.59 | 88796479.3 |
| 296230448 | PREDICTED: NADH-cytochrome b5 reductase 1 [Callithrix jacchus] | 0 | 113.322 | 52 | 14 | 115 | 14 | 305 | 34.1 | 9.38 | 1627 | 14 | 63671919.21 | 69179084.65 |
| 675772684 | PREDICTED: mitotic-spindle organizing protein 2B isoform X1 [Callithrix jacchus] | 0.008 | 3.996 | 11 | 2 | 9 | 2 | 193 | 20.7 | 11.94 | 20 | 2 | 2446932.791 | 2253074.585 |
| 1060950393 | PREDICTED: nucleoporin SEH1 isoform X1 [Callithrix jacchus] | 0 | 57.302 | 23 | 7 | 40 | 7 | 423 | 46.7 | 7.9 | 507 | 7 | 4676918.829 | 7885208.839 |
| 675662040 | PREDICTED: APC membrane recruitment protein 2 isoform X1 [Callithrix jacchus] | 0 | 103.723 | 35 | 17 | 141 | 4 | 676 | 70 | 7.25 | 2267 | 17 | 614319.5237 | 2085996.74 |
| 1060980692 | PREDICTED: WD repeat and FYVE domain-containing protein 3 isoform X1 [Callithrix jacchus] | 0 | 47.232 | 5 | 15 | 43 | 15 | 3544 | 397.1 | 6.77 | 108 | 15 | 4479216.944 | 6563136.134 |
| 532508846 | transmembrane protein 163 [Callithrix jacchus] | 0 | 23.525 | 19 | 4 | 38 | 4 | 289 | 31.5 | 7.53 | 885 | 4 | 3019913.676 | 6353537.109 |
| 296227769 | PREDICTED: ATP-dependent RNA helicase DHX36 [Callithrix jacchus] | 0 | 34.613 | 10 | 9 | 27 | 9 | 1010 | 114.9 | 7.77 | 120 | 9 | 31464307.39 | 23888205.82 |
| 1060976447 | PREDICTED: LOW QUALITY PROTEIN: echinoderm microtubule-associated protein-like 2 [Callithrix jacchus] | 0 | 128.024 | 31 | 23 | 143 | 23 | 849 | 91.6 | 6.86 | 1279 | 23 | 30752384.03 | 18490122.32 |
| 532291718 | dual specificity protein phosphatase 3 [Callithrix jacchus] | 0 | 36.305 | 43 | 6 | 36 | 6 | 185 | 20.5 | 7.8 | 591 | 6 | 8966655.542 | 10908930.22 |
| 532438335 | leucine-rich PPR motif-containing protein, mitochondrial precursor [Callithrix jacchus] | 0 | 254.17 | 39 | 47 | 344 | 47 | 1398 | 158.4 | 6.25 | 3859 | 47 | 168002853.1 | 89009160.33 |
| 296201694 | ICTED: LOW QUALITY PROTEIN: 1-phosphatidylinositol 4,5-bisphosphate phosphodiesterase delta-3 [Callithrix jac | 0 | 105.112 | 29 | 15 | 102 | 15 | 791 | 89.6 | 6.76 | 1464 | 15 | 56160923.27 | 48196615.06 |
| 1061018644 | PREDICTED: serine/threonine-protein phosphatase PGAM5, mitochondrial isoform X3 [Callithrix jacchus] | 0 | 58.665 | 42 | 10 | 207 | 10 | 288 | 31.9 | 8.68 | 2991 | 10 | 54729948.45 | 78560789.36 |
| 1060967110 | PREDICTED: inactive hydroxysteroid dehydrogenase-like protein 1 [Callithrix jacchus] | 0 | 35.299 | 25 | 8 | 62 | 8 | 330 | 36.9 | 8.66 | 496 | 8 | 22318948.12 | 18415644.54 |
| 296192766 | PREDICTED: pterin-4-alpha-carbinolamine dehydratase 2 [Callithrix jacchus] | 0 | 32.654 | 48 | 5 | 34 | 5 | 130 | 14.6 | 9.11 | 419 | 5 | 12308579.51 | 26774409.93 |
| 1060988630 | PREDICTED: bcl-2-associated transcription factor 1 isoform X1 [Callithrix jacchus] | 0 | 91.674 | 20 | 15 | 96 | 14 | 921 | 106.2 | 9.98 | 762 | 15 | 19758781.89 | 22559172.15 |
| 296202435 | PREDICTED: carbonic anhydrase-related protein 10 [Callithrix jacchus] | 0 | 17.902 | 16 | 4 | 18 | 4 | 328 | 37.5 | 8.09 | 227 | 4 | 2483269.587 | 3392027.786 |
| 675742900 | PREDICTED: tudor and KH domain-containing protein [Callithrix jacchus] | 0 | 78.024 | 27 | 11 | 91 | 11 | 560 | 61.7 | 5.07 | 2106 | 11 | 15212336.32 | 18647763.67 |
| 296224117 | PREDICTED: protein FAM98A [Callithrix jacchus] | 0 | 171.324 | 38 | 11 | 111 | 8 | 516 | 55.1 | 8.95 | 2783 | 11 | 10018122.76 | 14056334.82 |
| 1061010204 | PREDICTED: rho guanine nucleotide exchange factor 10-like protein isoform X1 [Callithrix jacchus] | 0 | 53.665 | 14 | 11 | 18 | 11 | 1279 | 140.5 | 6.02 | 83 | 11 | 8899777.599 | 5615370.045 |
| 296216300 | PREDICTED: sodium channel subunit beta-2 [Callithrix jacchus] | 0 | 44.848 | 38 | 8 | 65 | 8 | 215 | 24.3 | 6.99 | 1011 | 8 | 16380083.21 | 33335882.67 |
| 1060954174 | PREDICTED: putative ATP-dependent RNA helicase DHX30 isoform X2 [Callithrix jacchus] | 0 | 251.614 | 46 | 45 | 315 | 45 | 1194 | 133.9 | 8.78 | 3085 | 45 | 108546590.6 | 273903593.8 |

| | | | | | | | | | | | | | | | |
|---|---|---|---|---|---|---|---|---|---|---|---|---|---|---|---|
| 1060958455 | PREDICTED: inactive phospholipase C-like protein 2 isoform X1 [Callithrix jacchus] | 0 | 87.933 | 20 | 18 | 63 | 17 | 1157 | 129 | 6.98 | 657 | 18 | 15507366.32 | 28634549.78 | 28662556.82 |
| 1060964546 | PREDICTED: E3 ubiquitin-protein ligase Praja-2 [Callithrix jacchus] | 0 | 41.044 | 17 | 8 | 19 | 7 | 709 | 78.2 | 4.42 | 283 | 8 | 1813634.51 | 4080612.872 | 2706190.985 |
| 532287860 | disks large-associated protein 1 isoform 3 [Callithrix jacchus] | 0 | 239.667 | 43 | 31 | 273 | 11 | 929 | 103.6 | 6.77 | 5952 | 31 | 32164820.98 | 25823332.37 | 28357635.36 |
| 296224047 | PREDICTED: serine/arginine-rich splicing factor 7 isoform X1 [Callithrix jacchus] | 0 | 19.47 | 19 | 5 | 49 | 4 | 238 | 27.4 | 11.82 | 760 | 5 | 18363929.56 | 13931424.03 | 13580199.56 |
| 296224030 | PREDICTED: mitogen-activated protein kinase kinase kinase 3 isoform X1 [Callithrix jacchus] | 0 | 45.42 | 13 | 10 | 46 | 10 | 893 | 100.8 | 7.56 | 799 | 10 | 21052409.96 | 6769835.902 | 4822976.867 |
| 296222395 | PREDICTED: ankyrin repeat domain-containing protein 29 isoform X1 [Callithrix jacchus] | 0 | 16.652 | 20 | 6 | 13 | 6 | 301 | 32.4 | 9.6 | 50 | 6 | 2534319.127 | 5340019.865 | 3707821.173 |
| 296218693 | [illegible]: LOW QUALITY PROTEIN: serine/threonine-protein phosphatase 2A 56 kDa regulatory subunit beta isoform [Callit[illegible] | 0 | 49.002 | 17 | 7 | 88 | 6 | 497 | 57.4 | 6.71 | 1021 | 7 | 8184458.778 | 8499526.095 | 11478698.3 |
| 532508388 | abhydrolase domain-containing protein FAM108B1 isoform 2 precursor [Callithrix jacchus] | 0 | 18.728 | 19 | 4 | 25 | 4 | 288 | 32.1 | 6.21 | 65 | 4 | 3841464.871 | 4555704.173 | 4352717.333 |
| 675637567 | PREDICTED: transient receptor potential cation channel subfamily M member 3 isoform X9 [Callithrix jacchus] | 0 | 104.614 | 15 | 19 | 62 | 19 | 1744 | 199 | 7.21 | 909 | 19 | 10866250.68 | 16912532.82 | 14739940.49 |
| 675710443 | PREDICTED: opalin isoform X1 [Callithrix jacchus] | 0 | 65.501 | 47 | 5 | 61 | 5 | 141 | 15.6 | 5.55 | 718 | 5 | 32317779.34 | 17562059.67 | 18412767.43 |
| 296221128 | PREDICTED: alpha-internexin [Callithrix jacchus] | 0 | 363.106 | 74 | 36 | 1110 | 30 | 499 | 55.3 | 5.4 | 19509 | 36 | 990239905.1 | 1223962703 | 1227103584 |
| 675711715 | PREDICTED: actin-binding LIM protein 1 isoform X3 [Callithrix jacchus] | 0 | 284.741 | 50 | 31 | 508 | 31 | 788 | 89 | 8.46 | 7781 | 31 | 120441153.1 | 111144858.7 | 124452528.7 |
| 1061004761 | PREDICTED: LOW QUALITY PROTEIN: rab3 GTPase-activating protein catalytic subunit [Callithrix jacchus] | 0 | 47.819 | 12 | 10 | 27 | 10 | 989 | 111.3 | 5.45 | 102 | 10 | 2470749.781 | 1523588.497 | 1541892.995 |
| 296204889 | PREDICTED: aspartate--tRNA ligase, cytoplasmic [Callithrix jacchus] | 0 | 130.642 | 52 | 21 | 171 | 21 | 501 | 57.1 | 6.44 | 2190 | 21 | 29952163.29 | 33800071.67 | 27308803.79 |
| 675746008 | PREDICTED: ras GTPase-activating protein nGAP isoform X2 [Callithrix jacchus] | 0 | 222.434 | 35 | 36 | 255 | 32 | 1287 | 144.6 | 7.36 | 3563 | 36 | 50975443.86 | 39383150.73 | 40746523.34 |
| 675746367 | PREDICTED: protein PRRC2C isoform X1 [Callithrix jacchus] | 0 | 96.563 | 12 | 24 | 87 | 24 | 2902 | 317.6 | 9.06 | 349 | 24 | 2892720.416 | 7717184.241 | 14270809.3 |
| 296229450 | PREDICTED: NADH dehydrogenase [ubiquinone] iron-sulfur protein 2, mitochondrial [Callithrix jacchus] | 0 | 337.34 | 65 | 23 | 325 | 23 | 463 | 52.6 | 7.18 | 4928 | 23 | 452809114.4 | 462407348.2 | 475832908.3 |
| 675747956 | PREDICTED: serine/threonine-protein kinase MRCK alpha isoform X1 [Callithrix jacchus] | 0 | 259.299 | 29 | 43 | 290 | 39 | 1845 | 209.7 | 6.47 | 2928 | 43 | 52030936.41 | 46507573.33 | 44787178.28 |
| 390477223 | PREDICTED: WD repeat-containing protein 26 [Callithrix jacchus] | 0 | 35.299 | 16 | 8 | 35 | 8 | 658 | 71.9 | 6.16 | 283 | 8 | 2933432.133 | 3561883.123 | 3607534.492 |
| 532509686 | serine/threonine-protein kinase MARK1 [Callithrix jacchus] | 0 | 196.674 | 38 | 28 | 224 | 19 | 795 | 88.8 | 9.35 | 1492 | 28 | 28190518.64 | 16266343.22 | 26252239.81 |
| 675645703 | PREDICTED: beta-centractin [Callithrix jacchus] | 0 | 144.387 | 55 | 15 | 166 | 6 | 376 | 42.3 | 6.4 | 3041 | 15 | 14655260.77 | 8697471.162 | 7590357.215 |
| 532511542 | zinc transporter ZIP10 precursor [Callithrix jacchus] | 0 | 24.672 | 10 | 7 | 35 | 7 | 831 | 94.1 | 6.8 | 248 | 7 | 24925482.79 | 14212380.36 | 9286268.379 |
| 675665262 | PREDICTED: NADH-ubiquinone oxidoreductase 75 kDa subunit, mitochondrial isoform X1 [Callithrix jacchus] | 0 | 452.123 | 65 | 40 | 749 | 40 | 729 | 79.5 | 6.35 | 13771 | 40 | 868932412.7 | 755960872.6 | 785896452.9 |
| 675717582 | PREDICTED: IlsH domain and HEAT repeat-containing protein KIAA1468 homolog isoform X1 [Callithrix jacchus] | 0 | 50.455 | 11 | 11 | 50 | 11 | 1227 | 135.6 | 5.44 | 575 | 11 | 12195088.61 | 11776316.9 | 11008363.36 |
| 532516950 | calcium/calmodulin-dependent protein kinase type 1D isoform 2 [Callithrix jacchus] | 0 | 47.672 | 27 | 9 | 53 | 5 | 385 | 42.9 | 7.21 | 1161 | 9 | 5167697.302 | 10691072.64 | 9875913.615 |
| 532218343 | tubulin beta-4B chain [Callithrix jacchus] | 0 | 629.545 | 80 | 33 | 3900 | 3 | 445 | 49.8 | 4.89 | 92623 | 33 | 698959914.3 | 1722238440 | 1663131658 |
| 532280676 | rab-like protein 6 isoform 1 [Callithrix jacchus] | 0 | 36.34 | 18 | 10 | 43 | 10 | 732 | 79.8 | 5.35 | 223 | 10 | 30803606.67 | 11456813.16 | 10922869.35 |
| 1060936357 | PREDICTED: protein transport protein Sec16A isoform X2 [Callithrix jacchus] | 0 | 103.732 | 10 | 20 | 109 | 20 | 2376 | 253.3 | 5.99 | 1196 | 20 | 4923876.464 | 19250304.74 | 16842383.14 |
| 675639520 | PREDICTED: calcium-binding mitochondrial carrier protein SCaMC-2 isoform X3 [Callithrix jacchus] | 0 | 88.326 | 34 | 13 | 74 | 13 | 501 | 56 | 8.95 | 1325 | 13 | 11234805.37 | 21193363.93 | 17743591.66 |
| 532276392 | gelsolin isoform c [Callithrix jacchus] | 0 | 235.049 | 41 | 24 | 310 | 24 | 742 | 82 | 5.77 | 3673 | 24 | 110220173.2 | 110763499.7 | 106990641.1 |
| 1060935650 | PREDICTED: rab GTPase-activating protein 1 isoform X1 [Callithrix jacchus] | 0 | 55.836 | 14 | 13 | 43 | 11 | 1069 | 121.6 | 5.25 | 203 | 13 | 21332052.31 | 6489902.838 | 6873224.312 |
| 1060935666 | PREDICTED: DENN domain-containing protein 1A isoform X1 [Callithrix jacchus] | 0 | 112.334 | 23 | 17 | 142 | 17 | 1074 | 118.3 | 6.89 | 1658 | 17 | 10156775.08 | 27164587.2 | 17558249.79 |
| 675663679 | PREDICTED: alkyldihydroxyacetonephosphate synthase, peroxisomal isoform X1 [Callithrix jacchus] | 0 | 61.152 | 20 | 9 | 47 | 9 | 678 | 74.7 | 8.12 | 580 | 9 | 8824059.894 | 4857549.389 | 6422583.09 |
| 1060943644 | PREDICTED: palmitoyltransferase ZDHHC5 isoform X1 [Callithrix jacchus] | 0 | 51.6 | 23 | 12 | 39 | 12 | 715 | 77.4 | 9.01 | 260 | 12 | 7466473.232 | 27832086.69 | 23911561.25 |
| 532288512 | 182 kDa tankyrase-1-binding protein [Callithrix jacchus] | 0 | 113.589 | 20 | 25 | 74 | 25 | 1731 | 182 | 4.86 | 1228 | 25 | 60345729.3 | 23228928.85 | 14755002.71 |
| 1060943466 | PREDICTED: MAP kinase-activating death domain protein isoform X1 [Callithrix jacchus] | 0 | 109.725 | 18 | 20 | 92 | 20 | 1660 | 184.3 | 6.3 | 875 | 20 | 25877768.13 | 30078581.35 | 18736080.26 |
| 675700186 | PREDICTED: rho GTPase-activating protein 1 [Callithrix jacchus] | 0 | 11.958 | 8 | 3 | 12 | 3 | 439 | 50.4 | 6.6 | 40 | 3 | 599073.6191 | 6688136.335 | 5109144.711 |
| 532332949 | vacuolar protein sorting-associated protein 18 homolog [Callithrix jacchus] | 0 | 123.142 | 24 | 18 | 109 | 18 | 973 | 110.2 | 6.14 | 1562 | 18 | 43125884.25 | 36386453.39 | 32661246.54 |

[illegible]

**Supplementary Table 5 LC-MS/MS quantitation and comparison of 2,361 PSD proteins in marmoset neocortex**

| Description of protein | | Detection by LC-MS/MS | | | | | | | | | | | | LFQ intensity normalized with total peptide abundance | | | | | | | | | | | | Coefficient of Variation (CV) | | | | Marmoset Gene ID | | Mouse Gene ID | | Mean relative abundance | | | | | | | | |
|---|---|---|---|---|---|---|---|---|---|---|---|---|---|---|---|---|---|---|---|---|---|---|---|---|---|---|---|---|---|---|---|---|---|---|---|---|---|---|---|---|---|---|
| GI Number | Description | -value: Co | m PEP Sc | overage [% | # Peptides | # PSMs | nique Pepti | rotein Grou | # AAs | MW [kDa] | calc. pI | Mascot: M | Search En | rmalized): F1: S | rmalized): F2: S | rmalized): F3: S | rmalized): F4: S | rmalized): F5: S | rmalized): F6: S | rmalized): F7: S | rmalized): F8: S | rmalized): F9: S | malized): F10: S | malized): F11: S | malized): F12: S | CV 2M | CV 3M | CV 6M | CV 24M | Gene ID | Gene Symbol | Gene ID | Gene Symbol | 2M | 3M | 6M | 24M | P value (ANOVA) | Corrected P value | Fold | Cluster | DE |
| 296206333 | PREDICTED: abl interactor 1 isoform X1 [Callithrix jacchus] | 0 | 132.723 | 39 | 17 | 133 | 12 | 1 | 508 | 55.1 | 7.06 | 1704 | 17 | 155645675 | 147052533.4 | 147138903.1 | 164456057.8 | 174057253.8 | 149557765.3 | 156564363.7 | 157098850.6 | 163392607.5 | 113754008 | 84636544.58 | 96141725.75 | 3.292198 | 7.587922 | 2.388029 | 14.93733 | 100408906 | ABI1 | 11308 | Abi1 | 1.05 | 1.14 | 1.12 | 0.69 | 0.01454 | 0.041729623 | 1.657106 | 2 | Yes |
| 1060959086 | PREDICTED: Abelson tyrosine-protein kinase 2 [Callithrix jacchus] | 0 | 26.186 | 8 | 7 | 27 | 7 | 1 | 1080 | 118 | 7.75 | 286 | 7 | 17926595.26 | 13744444.25 | 12120743.53 | 16296018 | 14205377.3 | 13103346.17 | 13621976.75 | 17654041.31 | 13719414.85 | 5929755.245 | 4971827.964 | 1466641.836 | 20.52032 | 11.1569 | 15.3369 | 56.99087 | 100388523 | ABL2 | 11352 | Abl2 | 1.21 | 1.20 | 1.24 | 0.34 | 0.01425 | 0.041197674 | 3.637986 | 2 | Yes |
| 296202026 | PREDICTED: acid-sensing ion channel 2 [Callithrix jacchus] | 0 | 15.995 | 9 | 4 | 27 | 3 | 1 | 512 | 57.6 | 5.19 | 151 | 4 | 2492077.957 | 2582529 | 2198941.032 | 2998526.414 | 2968740.113 | 2238832.458 | 1664558.28 | 1865030.339 | 1971387.9 | 1660878.274 | 1705754.89 | 1846594.703 | 8.27053 | 15.72984 | 8.496777 | 5.576333 | 100413424 | ASIC2 | 11418 | Asic2 | 1.11 | 1.25 | 0.84 | 0.80 | 0.02246 | 0.054974632 | 1.574092 | 2 | No |
| 1061016719 | PREDICTED: LOW QUALITY PROTEIN: acid-sensing ion channel 1 [Callithrix jacchus] | 0 | 25.21 | 13 | 7 | 48 | 6 | 1 | 528 | 59.8 | 5.73 | 165 | 7 | 27079982.6 | 27747352 | 25539829.57 | 29449597 | 22437757.92 | 27107731.74 | 23128429.06 | 22600653.99 | 23000362.2 | 13907395.57 | 8567434.194 | 8915338.374 | 4.226174 | 13.55687 | 1.201634 | 28.55351 | 100399222 | ASIC1 | 11419 | Asic1 | 1.24 | 1.22 | 1.06 | 0.48 | 0.009917 | 0.033305202 | 2.560265 | 2 | Yes |
| 1060937427 | PREDICTED: aconitate hydratase, mitochondrial [Callithrix jacchus] | 0 | 224.813 | 37 | 29 | 219 | 29 | 1 | 857 | 94.2 | 7.91 | 3449 | 29 | 126157568.5 | 120407042.5 | 122565718.1 | 165278172.7 | 167313342.1 | 161091880.7 | 152502187 | 157416287.7 | 151608621.8 | 195218716.8 | 198820443.3 | 172091500.7 | 2.360854 | 1.927616 | 2.032722 | 7.686104 | 100389367 | ACO2 | 11429 | Aco2 | 0.78 | 1.04 | 0.98 | 1.20 | 0.0002836 | 0.006955849 | 1.533688 | 1 | Yes |
| 675750393 | PREDICTED: alpha-actinin-2 [Callithrix jacchus] | 0 | 522.039 | 68 | 56 | 976 | 36 | 1 | 894 | 103.8 | 5.45 | 18596 | 56 | 708690061.6 | 708937782.7 | 747656803.5 | 614220490.4 | 619845699.2 | 588469058.6 | 489876208 | 481501585.7 | 530158368.4 | 338666111.7 | 433316388 | 294359311.1 | 3.107161 | 2.753787 | 5.197413 | 19.96983 | 100407884 | ACTN2 | 11472 | Actn2 | 1.32 | 1.11 | 0.92 | 0.65 | 0.001329 | 0.011524484 | 2.030573 | 2 | Yes |
| 532273828 | aspartoacylase [Callithrix jacchus] | 0 | 9.902 | 12 | 3 | 9 | 3 | 1 | 313 | 35.6 | 6.92 | 39 | 3 | 1206894.594 | 952033.625 | 1324856.996 | 2294884.628 | 2641196.283 | 2591104.548 | 2925205.53 | 2027141.976 | 1848676.562 | 3171539.386 | 999162.5236 | 3157055.149 | 16.40928 | 7.45961 | 25.45019 | 51.17782 | 100896561 | ASPA | 11484 | Aspa | 0.55 | 1.20 | 1.08 | 1.17 | 0.006807 | 0.026845569 | 2.160634 | 1 | Yes |
| 532281686 | disintegrin and metalloproteinase domain-containing protein 11 preproprotein [Callithrix jacchus] | 0 | 24.147 | 6 | 3 | 7 | 3 | 1 | 769 | 83.6 | 7.34 | 112 | 3 | 3582930.18 | 3666917.938 | 3290649.905 | 2542624.762 | 4898493.457 | 2232260.363 | 2270732.199 | 3411687.366 | 2967200.441 | 3661368.092 | 3359125.51 | 9596889.669 | 5.621439 | 45.21805 | 19.94642 | 63.50048 | 100398230 | ADAM11 | 11488 | Adam11 | 0.93 | 0.85 | 0.76 | 1.46 | 0.4673 | 0.506778053 | 1.921169 | 1 | No |
| 675650426 | PREDICTED: alpha-adducin isoform X6 [Callithrix jacchus] | 0 | 450.034 | 71 | 37 | 595 | 36 | 1 | 662 | 73.3 | 6.37 | 14198 | 37 | 1007491969 | 936027165.6 | 951446773.9 | 999862692.2 | 915076454.5 | 964847768.8 | 1013726529 | 1041955811 | 1104246173 | 820238528.3 | 803915318.4 | 766192140.7 | 3.89722 | 4.438516 | 4.397146 | 3.479035 | 100401343 | ADD1 | 11518 | Add1 | 1.02 | 1.02 | 1.12 | 0.84 | 0.004142 | 0.020727551 | 1.321954 | 2 | No |
| 296223606 | PREDICTED: beta-adducin [Callithrix jacchus] | 0 | 309.311 | 57 | 37 | 428 | 34 | 1 | 726 | 80.8 | 5.92 | 7383 | 37 | 755223156.6 | 795885955 | 848642101.1 | 765802548.7 | 760026356.1 | 733033988.8 | 807381589.5 | 794127133.9 | 761675685.9 | 703585161.6 | 322909161.1 | 410343681 | 5.855576 | 2.323059 | 2.985201 | 41.63199 | 100402289 | ADD2 | 11519 | Add2 | 1.13 | 1.07 | 1.12 | 0.68 | 0.1552 | 0.208759909 | 1.670161 | 2 | No |
| 296230268 | PREDICTED: poly [ADP-ribose] polymerase 1 [Callithrix jacchus] | 0 | 65.177 | 17 | 14 | 63 | 14 | 1 | 1013 | 113 | 8.9 | 259 | 14 | 44846663.79 | 53500732.72 | 47702691.69 | 46856630.79 | 52207616.2 | 49968695.63 | 38409952.9 | 34963972.44 | 32321985.99 | 19924706.14 | 10761708.77 | 26676953.06 | 9.057701 | 5.409554 | 8.664911 | 41.77576 | 100403519 | PARP1 | 11545 | Parp1 | 1.28 | 1.30 | 0.92 | 0.50 | 0.007299 | 0.027522762 | 2.598051 | 2 | Yes |
| 1061002891 | PREDICTED: LOW QUALITY PROTEIN: aggrecan core protein [Callithrix jacchus] | 0 | 72.943 | 8 | 14 | 83 | 14 | 1 | 2264 | 234.8 | 4.31 | 755 | 14 | 15719338.61 | 11286915.5 | 13687785.05 | 19993581.24 | 17731342.54 | 21190700.69 | 42174294.13 | 42258065.43 | 50295284.61 | 59254323.24 | 54777815.07 | 49048688.22 | 16.357 | 8.94565 | 10.38687 | 9.410573 | 100385452 | ACAN | 11595 | Acan | 0.41 | 0.59 | 1.36 | 1.64 | 0.0006863 | 0.008906816 | 4.007487 | 1 | Yes |
| 296218936 | PREDICTED: AH receptor-interacting protein isoform X1 [Callithrix jacchus] | 0 | 22.965 | 25 | 7 | 13 | 7 | 1 | 330 | 37.5 | 6.54 | 40 | 7 | 10379772.37 | 10011887.38 | 10088192.97 | 5548901.058 | 5903705.406 | 7747762.37 | 4194621.881 | 3364560.743 | 4877467.886 | 5338599.361 | 1719706.629 | 1680705.834 | 1.910998 | 18.44489 | 18.27614 | 72.11517 | 100407721 | AIP | 11632 | Aip | 1.72 | 1.08 | 0.70 | 0.49 | 0.001849 | 0.013777091 | 3.487792 | 2 | Yes |
| 532524482 | adenylate kinase isoenzyme 1 [Callithrix jacchus] | 0 | 57.428 | 50 | 10 | 90 | 9 | 1 | 194 | 21.6 | 8.62 | 1320 | 10 | 66386803.52 | 69555330.72 | 76932245.48 | 100952818.7 | 91622885.47 | 67224037.8 | 62442630.04 | 48329067.29 | 59607620.65 | 56689416.91 | 40297431.86 | 45388462.61 | 7.625434 | 20.11132 | 13.14574 | 17.6781 | 100400464 | AK1 | 11636 | Ak1 | 1.08 | 1.32 | 0.87 | 0.73 | 0.05042 | 0.092606563 | 1.824753 | 2 | No |
| 675660140 | PREDICTED: A-kinase anchor protein 1, mitochondrial [Callithrix jacchus] | 0 | 12.323 | 5 | 3 | 13 | 3 | 1 | 876 | 93.7 | 5.01 | 179 | 3 | 1996690.809 | 1773319.875 | 2547043.654 | 1434314.808 | 1136644.29 | 1246674.668 | 1273558.332 | 824388.4793 | 1332966.522 | 1038884.509 | 700275.1586 | 987448.8711 | 18.91123 | 11.82764 | 24.31444 | 20.07653 | 100391330 | AKAP1 | 11640 | Akap1 | 1.55 | 0.94 | 0.84 | 0.67 | 0.05175 | 0.094098152 | 2.316817 | 2 | No |
| 296196349 | PREDICTED: serum albumin [Callithrix jacchus] | 0 | 79.698 | 32 | 16 | 92 | 16 | 1 | 608 | 69.4 | 5.99 | 1024 | 16 | 9758853.785 | 11485855.7 | 14671523.33 | 43276376.34 | 41086103.99 | 44658846.3 | 21782752.42 | 17348558.4 | 23646425.33 | 58765720.26 | 26191105.38 | 45006726.74 | 20.81648 | 4.188906 | 15.46018 | 37.74733 | 100391910 | ALB | 11657 | Alb | 0.40 | 1.44 | 0.70 | 1.45 | 0.000345 | 0.007128395 | 3.618518 | 1 | Yes |
| 1061018166 | PREDICTED: aldehyde dehydrogenase, mitochondrial [Callithrix jacchus] | 0 | 51.288 | 25 | 12 | 54 | 12 | 1 | 517 | 56.2 | 6.65 | 386 | 12 | 21303414.74 | 18667369.31 | 15948157.98 | 18777965.98 | 24008907.37 | 25785962.94 | 22773854.81 | 18735777.66 | 19317828.07 | 15898694.96 | 31931023.06 | 11389569.56 | 14.36581 | 15.9382 | 10.76577 | 54.69154 | 100388478 | ALDH2 | 11669 | Aldh2 | 0.91 | 1.12 | 0.99 | 0.97 | 0.6142 | 0.639938421 | 1.22629 | 2 | No |
| 1060946290 | PREDICTED: fructose-bisphosphate aldolase A isoform X2 [Callithrix jacchus] | 0 | 222.31 | 78 | 26 | 336 | 22 | 1 | 364 | 39.4 | 8.24 | 4929 | 26 | 254780068.7 | 265070263.7 | 256924078.5 | 297982639.1 | 316109260.8 | 312129624.8 | 258288738.3 | 272414891.8 | 247401676.5 | 245739307.5 | 261075615 | 229774892 | 2.096754 | 3.085669 | 4.835404 | 6.374543 | 103796602 | ALDOA | 11674 | Aldoa | 0.97 | 1.15 | 0.97 | 0.92 | 0.008137 | 0.029523186 | 1.257445 | 2 | No |
| 296202196 | PREDICTED: fructose-bisphosphate aldolase C [Callithrix jacchus] | 0 | 145.17 | 54 | 18 | 160 | 14 | 1 | 364 | 39.4 | 6.44 | 2777 | 18 | 39965621.19 | 46112123.44 | 74863371.16 | 68475667.34 | 119526844 | 55716174.91 | 60038097.83 | 52583157.26 | 53941099.13 | 51684154.52 | 75172594.92 | 117911228.2 | 34.72536 | 41.56341 | 7.151539 | 41.15316 | 100408520 | ALDOC | 11676 | Aldoc | 0.79 | 1.19 | 0.82 | 1.20 | 0.5441 | 0.575744111 | 1.520854 | 1 | No |
| 1060950259 | PREDICTED: ankyrin-1 isoform X1 [Callithrix jacchus] | 0 | 164.317 | 20 | 34 | 208 | 28 | 1 | 1933 | 211.9 | 6.61 | 2074 | 34 | 27360837.76 | 29309366.59 | 34709179.82 | 17574159.23 | 27369972.18 | 29094513.62 | 18886333.21 | 23065958.42 | 19076652.5 | 41307840.34 | 21980409.29 | 36629963 | 12.49797 | 25.17699 | 11.60148 | 30.27476 | 100388861 | ANK1 | 11733 | Ank1 | 1.12 | 0.91 | 0.75 | 1.22 | 0.08604 | 0.13352594 | 1.637227 | 1 | No |
| 532519378 | ankyrin-3 isoform 1 [Callithrix jacchus] | 0 | 866.777 | 36 | 132 | 1024 | 85 | 1 | 4382 | 480.6 | 6.52 | 12388 | 132 | 306003503.2 | 294918552.5 | 296971690 | 272023094 | 257461509.4 | 261097682.9 | 291024799.4 | 280398058.2 | 273284969 | 252523174.8 | 272980691.7 | 273697110.2 | 1.97036 | 2.875892 | 3.170702 | 4.513246 | 100408683 | ANK3 | 11735 | Ank3 | 1.08 | 0.95 | 1.01 | 0.96 | 0.01401 | 0.041005725 | 1.135737 | 2 | No |
| 675657852 | PREDICTED: rabankyrin-5 [Callithrix jacchus] | 0 | 9.611 | 2 | 3 | 9 | 3 | 1 | 1169 | 128.7 | 6.06 | 55 | 3 | 2014395.439 | 3250869.125 | 3404069.216 | 3550827.886 | 3977864.989 | 3169332.326 | 2441240.333 | 2430744.249 | 3149030.843 | 2537482.18 | 1352840.214 | 1378160.257 | 26.36755 | 11.34266 | 15.39854 | 38.53648 | 100415707 | ANKFY1 | 11736 | Ankfy1 | 1.06 | 1.31 | 0.98 | 0.65 | 0.08298 | 0.130365439 | 2.03057 | 2 | No |
| 296213563 | PREDICTED: acidic leucine-rich nuclear phosphoprotein 32 family member A [Callithrix jacchus] | 0 | 61.465 | 38 | 12 | 77 | 6 | 1 | 249 | 28.6 | 4.09 | 697 | 12 | 26851423.03 | 26987788.25 | 15189809.14 | 9861579.511 | 22032779.3 | 21253374.81 | 25751267.48 | 31946088.13 | 30766691.31 | 28782870.06 | 57289643.51 | 26435689.93 | 29.43347 | 38.45811 | 11.15507 | 45.79961 | 100398495 | ANP32A | 11737 | Anp32a | 0.85 | 0.66 | 1.10 | 1.39 | 0.2119 | 0.264818942 | 2.116896 | 1 | No |
| 532520188 | ADP/ATP translocase 1 [Callithrix jacchus] | 0 | 160.528 | 61 | 19 | 437 | 7 | 1 | 298 | 33 | 9.76 | 6015 | 19 | 262897945.5 | 257734386.1 | 257865410.6 | 374678616.9 | 343852916.6 | 273874047.2 | 196801105.6 | 206978911.3 | 196365258.9 | 183584595.3 | 148345588.9 | 233209498 | 1.134527 | 15.61479 | 3.002239 | 22.63229 | 100411371 | SLC25A4 | 11739 | Slc25a4 | 1.06 | 1.35 | 0.82 | 0.77 | 0.001205 | 0.011268514 | 1.756036 | 2 | Yes |
| 296236267 | PREDICTED: ADP/ATP translocase 2 [Callithrix jacchus] | 0 | 192.944 | 60 | 21 | 526 | 7 | 1 | 298 | 32.9 | 9.74 | 7232 | 21 | 601380803.4 | 566355000 | 578429532.2 | 649752934.1 | 617243924.4 | 588251511.1 | 412333011.7 | 385060014.2 | 452850813.6 | 377097825 | 388778115.3 | 319145355.8 | 3.05678 | 4.975204 | 8.18489 | 10.31063 | 100387000 | SLC25A5 | 11740 | Slc25a5 | 1.18 | 1.25 | 0.84 | 0.73 | 0.001809 | 0.01361535 | 1.709873 | 2 | Yes |
| 532503130 | annexin A6 isoform 1 [Callithrix jacchus] | 0 | 46.202 | 20 | 12 | 49 | 12 | 1 | 673 | 75.8 | 5.55 | 381 | 12 | 11820668.56 | 15586755.69 | 12630833.25 | 15412035.87 | 12484286.95 | 9747426.997 | 7788235.639 | 8552206.303 | 6319867.44 | 19638990.77 | 23565736.54 | 14137584.42 | 14.8531 | 22.57618 | 15.02019 | 24.77721 | 100411129 | ANXA6 | 11749 | Anxa6 | 1.02 | 0.95 | 0.57 | 1.45 | 0.02945 | 0.065192971 | 2.530518 | 1 | No |
| 296221338 | PREDICTED: thioredoxin-dependent peroxide reductase, mitochondrial [Callithrix jacchus] | 0 | 17.443 | 21 | 5 | 23 | 5 | 1 | 256 | 28 | 7.83 | 165 | 5 | 18983761.96 | 27431941.06 | 32319689.74 | 38042407.96 | 34137297.38 | 33010810.84 | 30203023.47 | 32242032.16 | 33577556.52 | 28316381.73 | 14452435.6 | 57071911.18 | 25.70653 | 7.530837 | 5.309511 | 65.32109 | 100399960 | PRDX3 | 11757 | Prdx3 | 0.83 | 1.11 | 1.01 | 1.05 | 0.3653 | 0.409704938 | 1.336 | 1 | No |
| 296229779 | PREDICTED: peroxiredoxin-6 [Callithrix jacchus] | 0 | 16.64 | 25 | 5 | 13 | 5 | 1 | 224 | 24.9 | 6.01 | 31 | 5 | 8247119.039 | 3878644.188 | 4179214.13 | 12115720.29 | 11220949.63 | 9307836.065 | 2944736.105 | 4631909.103 | 6476262.951 | 8274310.566 | 3952170.546 | 5927238.407 | 44.89437 | 13.1819 | 37.70777 | 35.75691 | 100403759 | PRDX6 | 11758 | Prdx6 | 0.80 | 1.61 | 0.69 | 0.89 | 0.03822 | 0.077890975 | 2.322972 | 2 | No |
| 675752514 | PREDICTED: AP-1 complex subunit gamma-1 isoform X1 [Callithrix jacchus] | 0 | 111.893 | 28 | 19 | 107 | 19 | 1 | 825 | 91.7 | 6.8 | 817 | 19 | 83499001.5 | 83687104.47 | 81420964.43 | 67603725.58 | 73692112.19 | 69530713.82 | 62432080.15 | 60233704.53 | 59478441.22 | 51825922.57 | 41588102.51 | 53841471.28 | 1.517549 | 4.427969 | 2.527314 | 13.3857 | 100394696 | AP1G1 | 11765 | Ap1g1 | 1.26 | 1.07 | 0.92 | 0.75 | 0.0002364 | 0.006753763 | 1.68827 | 2 | Yes |
| 532516586 | AP-1 complex subunit mu-1 isoform 1 [Callithrix jacchus] | 0 | 54.857 | 29 | 11 | 62 | 11 | 1 | 435 | 49.7 | 7.58 | 466 | 11 | 85320649.73 | 81103590.94 | 91791505.88 | 94009021.72 | 83167897.83 | 77467630.96 | 65903497.81 | 64564423.14 | 62880267.87 | 65828823.32 | 26647881.12 | 126146845.1 | 6.254557 | 9.899427 | 2.350517 | 68.77888 | 100414742 | AP1M1 | 11767 | Ap1m1 | 1.12 | 1.10 | 0.84 | 0.95 | 0.01794 | 0.047880542 | 1.335496 | 2 | No |
| 675642271 | PREDICTED: AP-1 complex subunit sigma-1A isoform X1 [Callithrix jacchus] | 0 | 24.018 | 31 | 6 | 24 | 5 | 1 | 161 | 19.1 | 5.73 | 518 | 6 | 14380174.86 | 14207175 | 12493433.37 | 13651959.91 | 12581764.66 | 11864820.83 | 11108473.35 | 10226436.83 | 11193553.27 | 8724167.789 | 5956886.727 | 7542475.914 | 7.616431 | 7.08192 | 4.938722 | 18.74425 | 100384920 | AP1S1 | 11769 | Ap1s1 | 1.23 | 1.14 | 0.97 | 0.66 | 0.01539 | 0.042917568 | 1.848526 | 2 | Yes |
| 532452551 | AP-2 complex subunit alpha-1 isoform 2 [Callithrix jacchus] | 0 | 559.528 | 79 | 58 | 999 | 11 | 1 | 955 | 105.3 | 7.66 | 18789 | 58 | 378639234.3 | 383035386.6 | 394649220.1 | 544230273.8 | 514227930.8 | 416308690.2 | 471444525.6 | 457848769.9 | 431459655.3 | 430259253.3 | 526308543.1 | 543384271.7 | 2.146036 | 13.60857 | 4.482228 | 12.1972 | 100414497 | AP2A1 | 11771 | Ap2a1 | 0.84 | 1.07 | 0.99 | 1.09 | 0.02682 | 0.061563499 | 1.297173 | 1 | No |
| 532291194 | AP-2 complex subunit alpha-2 isoform 2 [Callithrix jacchus] | 0 | 571.193 | 67 | 54 | 877 | 7 | 1 | 939 | 103.8 | 6.96 | 16466 | 54 | 96873638.71 | 108425087.1 | 110078649 | 75132741.12 | 87854222.57 | 112320623.1 | 133707028 | 118425438.6 | 123160024.2 | 118051293.6 | 127977355.8 | 85577570.09 | 6.843458 | 20.59572 | 6.253408 | 20.0629 | 100407362 | AP2A2 | 11772 | Ap2a2 | 0.97 | 0.85 | 1.16 | 1.02 | 0.1157 | 0.166230779 | 1.363175 | 1 | No |
| 296224700 | PREDICTED: AP-2 complex subunit mu isoform X1 [Callithrix jacchus] | 0 | 324.575 | 69 | 28 | 638 | 28 | 1 | 435 | 49.6 | 9.54 | 12663 | 28 | 2395068147 | 2451264994 | 2394035036 | 2017716185 | 2262243613 | 2598645226 | 2691758012 | 2598249486 | 3018908714 | 2281933436 | 2892311389 | 2274805460 | 1.356875 | 12.72087 | 7.975016 | 14.27608 | 100385882 | AP2M1 | 11773 | Ap2m1 | 0.97 | 0.92 | 1.11 | 1.00 | 0.2759 | 0.324701445 | 1.207936 | 2 | No |
| 296204176 | PREDICTED: AP-3 complex subunit beta-2 isoform X2 [Callithrix jacchus] | 0 | 172.836 | 32 | 30 | 204 | 25 | 1 | 1084 | 119.3 | 5.59 | 3426 | 30 | 107639814.6 | 105929372.8 | 104933203 | 116119812.4 | 129632164.6 | 121114101.4 | 103142929.4 | 99211644.33 | 99225740.19 | 110417070 | 73565950.29 | 96872588.75 | 1.2894 | 5.587046 | 2.253798 | 19.91039 | 100394963 | AP3B2 | 11775 | Ap3b2 | 1.00 | 1.16 | 0.95 | 0.89 | 0.03482 | 0.072847511 | 1.306244 | 2 | No |
| 532282834 | AP-3 complex subunit delta-1 isoform 3 [Callithrix jacchus] | 0 | 175.028 | 29 | 27 | 152 | 27 | 1 | 1217 | 136.3 | 7.44 | 2680 | 27 | 71299642.25 | 70711354.25 | 66305998.46 | 67615558.94 | 76274732.5 | 75896614.77 | 63986003.58 | 61598629.88 | 73627580.28 | 68415858.84 | 87697382.89 | 102272250.6 | 3.930291 | 6.679938 | 9.590693 | 19.71779 | 100407169 | AP3D1 | 11776 | Ap3d1 | 0.94 | 0.99 | 0.90 | 1.17 | 0.3948 | 0.439659406 | 1.297036 | 1 | No |
| 296193924 | PREDICTED: AP-3 complex subunit sigma-1 isoform X2 [Callithrix jacchus] | 0 | 21.454 | 33 | 6 | 23 | 6 | 1 | 193 | 21.7 | 5.39 | 329 | 6 | 14966649.59 | 15377345.72 | 12834822.33 | 11018446.99 | 11087053.6 | 12433577.93 | 11458373.42 | 10895142.36 | 11433640.68 | 11641924.32 | 5535089.025 | 5577574.288 | 9.483153 | 6.930922 | 2.826064 | 46.32362 | 100406901 | AP3S1 | 11777 | Ap3s1 | 1.29 | 1.03 | 1.01 | 0.68 | 0.0962 | 0.144866322 | 1.897587 | 2 | No |
| 532347965 | amyloid beta A4 precursor protein-binding family A member 2 isoform a [Callithrix jacchus] | 0 | 37.206 | 10 | 7 | 25 | 6 | 1 | 751 | 82.5 | 4.92 | 301 | 7 | 12442918.54 | 12315911.19 | 6307507.115 | 2429247.585 | 2465935.007 | 5409979.215 | 8138211.41 | 9129272.23 | 7859760.022 | 9596280.853 | 3353709.661 | 7490967.194 | 33.85844 | 49.79351 | 7.966546 | 46.61117 | 100403686 | APBA2 | 11784 | Apba2 | 1.43 | 0.47 | 1.16 | 0.94 | 0.06739 | 0.113372635 | 3.014638 | 2 | No |
| 1060942657 | PREDICTED: amyloid beta A4 precursor protein-binding family B member 1 [Callithrix jacchus] | 0 | 49.918 | 17 | 9 | 37 | 9 | 1 | 711 | 77.3 | 5.06 | 423 | 9 | 9586689.833 | 10236107.13 | 3266367.639 | 6873836.039 | 7376621.62 | 4851854.345 | 6214408.175 | 5556424.833 | 4527925.427 | 4208231.348 | 2624911.811 | 6200753.268 | 50.02641 | 20.9879 | 15.64532 | 41.24213 | 100401543 | APBB1 | 11785 | Apbb1 | 1.29 | 1.07 | 0.91 | 0.73 | 0.5244 | 0.558949819 | 1.771471 | 2 | No |
| 1060964515 | PREDICTED: adenomatous polyposis coli protein isoform X3 [Callithrix jacchus] | 0 | 234.36 | 20 | 44 | 208 | 43 | 1 | 2860 | 313.2 | 7.68 | 2392 | 44 | 108828352.9 | 98613126.25 | 103762804.5 | 67164429.3 | 74795867.44 | 86838328.35 | 68264157.3 | 57377954.88 | 74542334.15 | 49220771.39 | 36325935.19 | 29177073.97 | 4.92378 | 13.00579 | 13.015 | 26.56343 | 100385653 | APC | 11789 | Apc | 1.46 | 1.07 | 0.94 | 0.54 | 0.002597 | 0.016420321 | 2.71264 | 2 | Yes |
| 1061006705 | EDICTED: LOW QUALITY PROTEIN: striated muscle preferentially expressed protein kinase [Callithrix jacch | 0 | 59.396 | 6 | 15 | 39 | 15 | 1 | 3271 | 355.1 | 8.38 | 234 | 15 | 10297816.55 | 9291937.344 | 9816387.5 | 9751178.758 | 8319497.419 | 7756934.653 | 10215990.02 | 10387955.18 | 14243979.58 | 3523059.457 | 8072043.288 | 2810708.316 | 5.132529 | 11.94306 | 19.60698 | 59.4407 | 100412488 | SPEG | 11790 | Speg | 1.13 | 0.99 | 1.33 | 0.55 | 0.1353 | 0.18721652 | 2.419019 | 2 | No |
| 532200015 | DNA-(apurinic or apyrimidinic site) lyase [Callithrix jacchus] | 0 | 13.722 | 11 | 3 | 14 | 3 | 1 | 318 | 35.5 | 7.91 | 40 | 3 | 5418474.312 | 5414605.531 | 5454644.055 | 4831004.507 | 3918963.187 | 6466147.842 | 3105921.487 | 2529335.803 | 4863165.095 | 6475217.156 | 3405541.157 | 4314170.549 | 0.406766 | 25.4451 | 34.73864 | 33.32553 | 100400557 | APEX1 | 11792 | Apex1 | 1.16 | 1.08 | 0.75 | 1.01 | 0.279 | 0.327977214 | 1.551445 | 2 | No |
| 675699772 | PREDICTED: apoptosis inhibitor 5 isoform X1 [Callithrix jacchus] | 0 | 13.415 | 7 | 3 | 5 | 3 | 1 | 524 | 59 | 7.34 | 59 | 3 | 1228438.124 | 1264486 | 1058341.613 | 5335720.275 | 5149000.722 | 4018269.76 | 3188498.506 | 3145137.306 | 3277327.966 | 3665441.149 | 2887698.365 | 1426008.208 | 9.300636 | 14.74597 | 2.10341 | 42.74856 | 100398155 | API5 | 11800 | Api5 | 0.40 | 1.63 | 1.08 | 0.90 | 0.0001207 | 0.005998496 | 4.083893 | 2 | Yes |
| 296216647 | PREDICTED: amyloid-like protein 2 isoform X1 [Callithrix jacchus] | 0 | 64.475 | 20 | 13 | 51 | 12 | 1 | 768 | 87.4 | 4.73 | 280 | 13 | 16634546.76 | 17943882.04 | 15304830.79 | 13678153.57 | 11251886.48 | 13483678.85 | 11371905.25 | 15123577.04 | 15561787.22 | 15959757.74 | 11100888.57 | 18380363.61 | 7.935761 | 10.52887 | 16.42743 | 24.47462 | 100414190 | APLP2 | 11804 | Aplp2 | 1.14 | 0.87 | 0.96 | 1.03 | 0.1379 | 0.190034889 | 1.298579 | 2 | No |
| 296234066 | PREDICTED: apolipoprotein E [Callithrix jacchus] | 0 | 38.449 | 28 | 10 | 40 | 10 | 1 | 320 | 36.4 | 5.36 | 551 | 10 | 10911896.1 | 18540948.75 | 21118125.56 | 13076902.84 | 17869552.43 | 29716911.35 | 23570411.93 | 26829227.32 | 30482190.42 | 19732318.58 | 11519373.5 | 8719594.442 | 31.4849 | 42.35978 | 12.82524 | 42.95934 | 100401608 | APOE | 11816 | Apoe | 0.87 | 1.05 | 1.39 | 0.69 | 0.09861 | 0.147977649 | 2.023498 | 2 | No |
| 532451145 | amyloid beta A4 protein isoform a precursor [Callithrix jacchus] | 0 | 88.572 | 26 | 17 | 77 | 16 | 1 | 770 | 87 | 4.81 | 861 | 17 | 38511963.76 | 39171859.16 | 45369782.73 | 26517401.99 | 26628279.07 | 27100751.8 | 17606191.3 | 23584607.68 | 22681558.57 | 13546695.72 | 37313431.81 | 11198872.64 | 9.223501 | 1.158147 | 15.13678 | 69.83964 | 100407052 | APP | 11820 | App | 1.50 | 0.97 | 0.78 | 0.75 | 0.02611 | 0.060604558 | 1.982849 | 2 | No |
| 675716223 | PREDICTED: aquaporin-4 isoform X1 [Callithrix jacchus] | 0 | 25.338 | 21 | 7 | 95 | 7 | 1 | 358 | 38.8 | 8.72 | 1321 | 7 | 216463532.2 | 207564137.9 | 318272436.9 | 115403103 | 125536274.1 | 177747179.6 | 237153208.9 | 233523548.1 | 254423031.2 | 113136608.3 | 86759104.61 | 171997032 | 24.85905 | 23.97143 | 4.620186 | 35.20236 | 100409781 | AQP4 | 11829 | Aqp4 | 1.31 | 0.74 | 1.28 | 0.66 | 0.0272 | 0.062133849 | 1.996006 | 2 | No |
| 296213086 | PREDICTED: 60S acidic ribosomal protein P0 [Callithrix jacchus] | 0 | 96.029 | 54 | 14 | 195 | 14 | 1 | 318 | 34.3 | 5.97 | 2908 | 14 | 392525101.3 | 394585933.5 | 319320063.5 | 413197164.7 | 269511669.2 | 292398300.4 | 266436307.9 | 380665928.3 | 291124691.2 | 294199526.3 | 253379688.7 | 245355685.6 | 11.62447 | 23.75211 | 19.21865 | 9.909803 | 100410963 | RPLP0 | 11837 | Rplp0 | 1.16 | 1.02 | 0.98 | 0.83 | 0.1278 | 0.178617515 | 1.395362 | 2 | No |
| 675735821 | PREDICTED: activity-regulated cytoskeleton-associated protein [Callithrix jacchus] | 0 | 62.607 | 32 | 12 | 96 | 12 | 1 | 398 | 45.4 | 5.54 | 819 | 12 | 54021809.97 | 60870480.31 | 58418297.98 | 55672239.9 | 106357699.1 | 139237007.7 | 55287777.73 | 125443573.1 | 126831220 | 39726480.51 | 34766811.43 | 32896964.37 | 6.006607 | 41.92034 | 39.90498 | 9.859353 | 100387688 | ARC | 11838 | Arc | 0.78 | 1.35 | 1.38 | 0.48 | 0.009222 | 0.031892187 | 2.863971 | 2 | Yes |
| 1061016646 | PREDICTED: ADP-ribosylation factor 3 [Callithrix jacchus] | 0 | 68.706 | 59 | 9 | 82 | 4 | 1 | 181 | 20.6 | 7.43 | 809 | 9 | 59413283.51 | 60624894.25 | 64265563.5 | 51606387.94 | 49598158.97 | 55048010.16 | 45905608.74 | 43791667.81 | 62888277.93 | 55761214.1 | 61726934.18 | 102529607.5 | 4.11078 | 5.291747 | 20.58259 | 34.70851 | 100408060 | ARF3 | 11842 | Arf3 | 1.03 | 0.88 | 0.86 | 1.23 | 0.06971 | 0.115663411 | 1.44193 | 1 | No |
| 296225497 | PREDICTED: ADP-ribosylation factor 4 [Callithrix jacchus] | 0 | 47.572 | 48 | 8 | 35 | 4 | 1 | 180 | 20.5 | 7.14 | 258 | 8 | 8744448.234 | 8174957.5 | 7150461.057 | 6054725.699 | 6633177.629 | 6602591.109 | 8402173.879 | 5767033.575 | 6194516.019 | 8957507.155 | 8326521.157 | 6031577.946 | 10.0675 | 5.062068 | 20.83471 | 19.81253 | 100407980 | ARF4 | 11843 | Arf4 | 1.11 | 0.89 | 0.94 | 1.07 | 0.1757 | 0.22915704 | 1.247758 | 1 | No |
| 675689900 | PREDICTED: ADP-ribosylation factor 6 [Callithrix jacchus] | 0 | 40.951 | 38 | 5 | 45 | 5 | 1 | 175 | 20.1 | 8.95 | 279 | 5 | 32949200.67 | 34325493.38 | 32082541.48 | 41875789.86 | 39034664.38 | 35794018.97 | 41179046.05 | 39916827.43 | 44262357.77 | 44945812.49 | 47551848.94 | 37676662.55 | 3.415205 | 7.822507 | 5.349768 | 11.79449 | 100409406 | ARF6 | 11845 | Arf6 | 0.84 | 0.99 | 1.06 | 1.10 | 0.01866 | 0.049190759 | 1.310165 | 1 | No |
| 296225196 | PREDICTED: transforming protein RhoA [Callithrix jacchus] | 0 | 144.432 | 56 | 11 | 223 | 9 | 1 | 193 | 21.8 | 6.1 | 3826 | 11 | 285057803.1 | 248715600.3 | 270915243.6 | 296625216.5 | 294832319.2 | 315069566.5 | 362503987.7 | 344146194.2 | 359409913.1 | 332582349.4 | 327218701.7 | 228036959.2 | 6.829729 | 3.707228 | 2.765759 | 19.89284 | 100393236 | RHOA | 11848 | Rhoa | 0.88 | 0.99 | 1.16 | 0.97 | 0.007672 | 0.028447824 | 1.324811 | 2 | No |
| 390474730 | PREDICTED: rho-related GTP-binding protein RhoB [Callithrix jacchus] | 0 | 76.413 | 49 | 9 | 105 | 7 | 1 | 196 | 22.1 | 5.24 | 2078 | 9 | 45597526.12 | 45026065.06 | 44711815.78 | 64266495.95 | 58893156.74 | 58273565.87 | 62984867.12 | 69205865.75 | 60481644.28 | 76534854.41 | 112933466.2 | 72444137.58 | 0.995386 | 5.449529 | 6.994563 | 25.53106 | 100390079 | RHOB | 11852 | Rhob | 0.70 | 0.94 | 1.00 | 1.36 | 0.006183 | 0.025266862 | 1.935284 | 1 | Yes |
| 1060939397 | PREDICTED: rho GTPase-activating protein 5 [Callithrix jacchus] | 0 | 125.29 | 22 | 27 | 108 | 27 | 1 | 1501 | 172 | 6.67 | 923 | 27 | 41755508.99 | 34053013.06 | 33178647.92 | 33100373.64 | 31560079.65 | 33508078.1 | 35779874.45 | 44200832.28 | 38680647.73 | 58347714.03 | 43148454.33 | 37062646.57 | 12.99163 | 3.139733 | 10.81525 | 23.73623 | 100388004 | ARHGAP5 | 11855 | Arhgap5 | 0.94 | 0.85 | 1.02 | 1.19 | 0.1666 | 0.220329899 | 1.411438 | 1 | No |
| 532337601 | armadillo repeat protein deleted in velo-cardio-facial syndrome [Callithrix jacchus] | 0 | 151.567 | 34 | 24 | 95 | 24 | 1 | 962 | 104.6 | 6.95 | 1684 | 24 | 59278340.72 | 74319443.05 | 65528836.12 | 72466196.37 | 40120769.93 | 65302126.32 | 69566003.18 | 58775089.29 | 60365284.8 | 45437719.07 | 7667517.672 | 59330060.71 | 11.38403 | 28.65013 | 9.261374 | 71.33471 | 100412657 | ARVCF | 11877 | Arvcf | 1.17 | 1.05 | 1.11 | 0.66 | 0.5241 | 0.558882257 | 1.771033 | 2 | No |
| 1060959163 | PREDICTED: astrotactin-1 [Callithrix jacchus] | 0 | 217.865 | 32 | 27 | 183 | 27 | 1 | 1294 | 144 | 5.19 | 3085 | 27 | 95964204.24 | 105852061 | 104463917.9 | 98563675.6 | 86359202.75 | 87744568.04 | 71670822.67 | 71685428.31 | 70299537.46 | 36644193.05 | 82258739.55 | 49300470.78 | 5.243454 | 7.35218 | 1.117632 | 41.99957 | 100389868 | ASTN1 | 11899 | Astn1 | 1.28 | 1.14 | 0.89 | 0.70 | 0.006359 | 0.025807488 | 1.820892 | 2 | Yes |
| 296208963 | PREDICTED: sodium/potassium-transporting ATPase subunit alpha-1 [Callithrix jacchus] | 0 | 643.149 | 56 | 60 | 1114 | 39 | 1 | 1023 | 112.9 | 5.54 | 22814 | 60 | 1298710807 | 1285813604 | 1356922099 | 1023076698 | 1043891279 | 1100952955 | 1500903965 | 1499153282 | 1600286781 | 1855172068 | 1775419604 | 1688335593 | 2.883531 | 3.818234 | 3.775198 | 4.706499 | 100393318 | ATP1A1 | 11928 | Atp1a1 | 0.93 | 0.74 | 1.08 | 1.25 | 0.0004882 | 0.007832903 | 1.678996 | 1 | Yes |
| 296229852 | PREDICTED: sodium/potassium-transporting ATPase subunit beta-1 [Callithrix jacchus] | 0 | 148.041 | 46 | 16 | 284 | 16 | 1 | 303 | 35.1 | 8.53 | 2967 | 16 | 829129056 | 843826556.5 | 848899600 | 783584594.3 | 737078301.8 | 674148405.4 | 906541490 | 875021999.9 | 960139620.9 | 1286385380 | 1182182089 | 1188637675 | 1.221516 | 7.50722 | 4.708761 | 4.78952 | 100414358 | ATP1B1 | 11931 | Atp1b1 | 0.91 | 0.79 | 0.99 | 1.32 | 0.004056 | 0.020427019 | 1.666296 | 1 | Yes |
| 1060995112 | PREDICTED: sodium/potassium-transporting ATPase subunit beta-2 [Callithrix jacchus] | 0 | 88.964 | 36 | 10 | 113 | 10 | 1 | 290 | 33.4 | 8.31 | 2035 | 10 | 112419303.6 | 117211961.3 | 116371480.3 | 111090779.4 | 108027019.2 | 106308996.6 | 134265967.8 | 112691847.3 | 115116596.2 | 195606735.6 | 238161754.1 | 174568146.8 | 2.218904 | 2.232988 | 9.792084 | 15.9769 | 100392174 | ATP1B2 | 11932 | Atp1b2 | 0.84 | 0.79 | 0.88 | 1.48 | 0.02838 | 0.063897137 | 1.86935 | 1 | No |
| 532447777 | sarcoplasmic/endoplasmic reticulum calcium ATPase 2 isoform b [Callithrix jacchus] | 0 | 224.793 | 38 | 35 | 212 | 35 | 1 | 1042 | 114.7 | 5.38 | 2418 | 35 | 183415975.1 | 179179496 | 176392584.7 | 165595182.5 | 165121711.8 | 174976859.2 | 167061071 | 160833599.2 | 165858790.4 | 91005940.15 | 64675370.42 | 76015666.43 | 1.968436 | 3.297393 | 2.007187 | 17.10085 | 100398261 | ATP2A2 | 11938 | Atp2a2 | 1.22 | 1.14 | 1.12 | 0.52 | 0.001164 | 0.011131045 | 2.326262 | 2 | Yes |
| 532524566 | plasma membrane calcium-transporting ATPase 2 isoform 1 [Callithrix jacchus] | 0 | 468.122 | 47 | 54 | 497 | 27 | 1 | 1243 | 136.7 | 5.91 | 9523 | 54 | 209430738.2 | 227923713.8 | 187540508.2 | 192786955 | 209933535.4 | 215263286.3 | 210351998.9 | 229093711.3 | 226073323.6 | 150381454.1 | 223026741.7 | 125320012.7 | 9.705025 | 5.701354 | 4.535975 | 30.52631 | 100404341 | ATP2B2 | 11941 | Atp2b2 | 1.04 | 1.03 | 1.11 | 0.83 | 0.3306 | 0.375783061 | 1.334432 | 2 | No |
| 675716768 | PREDICTED: ATP synthase subunit alpha, mitochondrial [Callithrix jacchus] | 0 | 208.845 | 50 | 26 | 394 | 26 | 1 | 553 | 59.6 | 9.13 | 7740 | 26 | 796265167.7 | 774142817.3 | 786514184.4 | 1009164819 | 976234714.5 | 891247919.3 | 854239860.4 | 907826629.4 | 912947477.2 | 928783485.9 | 917126569 | 916678316.1 | 1.411206 | 6.345244 | 3.64681 | 0.745299 | 100395383 | LOC100395383 | 11946 | Atp5a1 | 0.88 | 1.08 | 1.00 | 1.04 | 0.000549 | 0.008104613 | 1.22051 | 1 | No |
| 1061018738 | PREDICTED: LOW QUALITY PROTEIN: ATP synthase subunit beta, mitochondrial [Callithrix jacchus] | 0 | 257.781 | 46 | 17 | 210 | 3 | 1 | 502 | 53.6 | 5.62 | 5391 | 17 | 51463456.64 | 50086243.5 | 52991608.32 | 73640394.15 | 70661499.56 | 67182751.93 | 60492455.56 | 61458526.39 | 63819910.04 | 60924966.2 | 18860117.89 | 128492155.6 | 2.821257 | 4.584792 | 2.764359 | 79.66521 | 100394984 | ATP5F1B | 11947 | Atp5b | 0.81 | 1.11 | 0.98 | 1.10 | 0.002914 | 0.017337199 | 1.368467 | 1 | No |
| 296206131 | PREDICTED: ATP synthase subunit gamma, mitochondrial isoform X1 [Callithrix jacchus] | 0 | 72.227 | 45 | 12 | 117 | 12 | 1 | 298 | 33 | 9.25 | 1488 | 12 | 141718960.6 | 133567929.2 | 152849407.6 | 131512332.1 | 128573065 | 150507935.6 | 139618778.4 | 135934847.7 | 134711566.3 | 158264453.5 | 106181693 | 105492207.8 | 6.782206 | 8.699598 | 1.867835 | 24.54814 | 100388706 | ATP5F1C | 11949 | Atp5c1 | 1.06 | 1.01 | 1.01 | 0.91 | 0.7568 | 0.774506759 | 1.157318 | 2 | No |
| 296208831 | PREDICTED: ATP synthase F(0) complex subunit B1, mitochondrial [Callithrix jacchus] | 0 | 52.6 | 34 | 10 | 103 | 10 | 1 | 256 | 28.7 | 9.38 | 1502 | 10 | 118699840.6 | 116594464.6 | 143105183.1 | 84726119.3 | 125428371.2 | 121410757.9 | 123005072.1 | 133836282.4 | 143917585.9 | 150479956.1 | 105811354.3 | 69637527.15 | 11.68277 | 20.29447 | 7.829019 | 37.27396 | 100392086 | ATP5PB | 11950 | Atp5pb | 1.05 | 0.92 | 1.12 | 0.91 | 0.5286 | 0.562664804 | 1.22959 | 2 | No |
| 675754462 | PREDICTED: ATP synthase-coupling factor 6, mitochondrial isoform X1 [Callithrix jacchus] | 0 | 34.168 | 17 | 3 | 23 | 3 | 1 | 170 | 19.4 | 9.98 | 1203 | 3 | 1913274.237 | 18833551.5 | 16754068.84 | 2660414.755 | 3003035.072 | 1838082.383 | 1638345.19 | 7265782.041 | 6083933.91 | 4081693.663 | 13824553.93 | 3280925.401 | 73.81742 | 23.94358 | 59.39263 | 83.11457 | 100411036 | ATP5PF | 11957 | Atp5j | 1.85 | 0.37 | 0.74 | 1.04 | 0.3163 | 0.363199125 | 4.999098 | 2 | No |
| [illegible] | [illegible] | [illegible] | [illegible] | [illegible] | [illegible] | [illegible] | [illegible] | [illegible] | [illegible] | [illegible] | [illegible] | [illegible] | [illegible] | [illegible] | [illegible] | [illegible] | [illegible] | [illegible] | [illegible] | [illegible] | [illegible] | [illegible] | [illegible] | [illegible] | [illegible] | [illegible] | [illegible] | [illegible] | [illegible] | [illegible] | [illegible] | [illegible] | [illegible] | [illegible] | [illegible] | [illegible] | [illegible] | [illegible] | [illegible] | [illegible] | [illegible] | [illegible] |

| | | | |
|---|---|---|---|
| 675775484 | PREDICTED: LOW QUALITY PROTEIN: X-ray repair cross-complementing protein 6 [Callithrix jacchus] | 0 | 102.853 |
| 675771045 | PREDICTED: LOW QUALITY PROTEIN: glucose-6-phosphate 1-dehydrogenase [Callithrix jacchus] | 0 | 23.098 |
| 675642911 | PREDICTED: gamma-aminobutyric acid receptor subunit alpha-1 [Callithrix jacchus] | 0 | 56.249 |
| 1060982081 | PREDICTED: gamma-aminobutyric acid receptor subunit alpha-2 isoform X1 [Callithrix jacchus] | 0 | 39.243 |
| 1061020844 | PREDICTED: gamma-aminobutyric acid receptor subunit alpha-3 isoform X1 [Callithrix jacchus] | 0 | 32.243 |
| 296196621 | PREDICTED: gamma-aminobutyric acid receptor subunit alpha-4 isoform X1 [Callithrix jacchus] | 0 | 72.311 |
| 390460895 | PREDICTED: gamma-aminobutyric acid receptor subunit beta-1 [Callithrix jacchus] | 0 | 56.097 |
| 390459183 | PREDICTED: gamma-aminobutyric acid receptor subunit beta-2 isoform X2 [Callithrix jacchus] | 0 | 93.405 |
| 532508175 | gamma-aminobutyric acid receptor subunit beta-3 isoform 1 precursor [Callithrix jacchus] | 0 | 65.256 |
| 675667718 | PREDICTED: gamma-aminobutyric acid receptor subunit delta [Callithrix jacchus] | 0 | 11.55 |
| 675663880 | PREDICTED: glutamate decarboxylase 1 isoform X1 [Callithrix jacchus] | 0 | 37.843 |
| 296206328 | PREDICTED: glutamate decarboxylase 2 [Callithrix jacchus] | 0 | 73.817 |
| 390475484 | PREDICTED: neuromodulin [Callithrix jacchus] | 0 | 463.498 |
| 675658116 | PREDICTED: growth arrest-specific protein 7 isoform X1 [Callithrix jacchus] | 0 | 234.341 |
| 1060958918 | PREDICTED: protein NipSnap homolog 2 isoform X3 [Callithrix jacchus] | 0 | 38.541 |
| 675675169 | PREDICTED: myotrophin isoform X2 [Callithrix jacchus] | 0 | 13.211 |
| 1060934392 | PREDICTED: guanine deaminase [Callithrix jacchus] | 0 | 123.859 |
| 296226731 | PREDICTED: ganglioside-induced differentiation-associated protein 1 [Callithrix jacchus] | 0 | 156.392 |
| 167045824 | ATPase, H+ transporting, lysosomal accessory protein 1 precursor (predicted) [Callithrix jacchus] | 0 | 159.892 |
| 296206102 | PREDICTED: rab GDP dissociation inhibitor beta isoform X1 [Callithrix jacchus] | 0 | 106.134 |
| 532443013 | glycerol-3-phosphate dehydrogenase, mitochondrial precursor [Callithrix jacchus] | 0 | 248.566 |
| 532504008 | glial fibrillary acidic protein isoform 1 [Callithrix jacchus] | 0 | 305.428 |
| 1060949939 | PREDICTED: GDNF family receptor alpha-2 [Callithrix jacchus] | 0 | 71.339 |
| 1060988084 | PREDICTED: gap junction alpha-1 protein [Callithrix jacchus] | 0 | 251.005 |
| 675767717 | PREDICTED: gap junction beta-1 protein [Callithrix jacchus] | 0 | 8.334 |
| 675661907 | PREDICTED: gap junction beta-6 protein [Callithrix jacchus] | 0 | 22.365 |
| 532208183 | glutamine synthetase [Callithrix jacchus] | 0 | 124.352 |
| 532519176 | hydroxyacylglutathione hydrolase, mitochondrial isoform 1 precursor [Callithrix jacchus] | 0 | 22.925 |
| 675664811 | PREDICTED: glutaminase kidney isoform, mitochondrial isoform X1 [Callithrix jacchus] | 0 | 87.216 |
| 1060946636 | PREDICTED: glutamate dehydrogenase 1, mitochondrial [Callithrix jacchus] | 0 | 139.826 |
| 296197592 | PREDICTED: guanine nucleotide-binding protein-like 1 isoform X1 [Callithrix jacchus] | 0 | 63.425 |
| 296232500 | PREDICTED: guanine nucleotide-binding protein subunit alpha-11 isoform X2 [Callithrix jacchus] | 0 | 139.158 |
| 296192593 | PREDICTED: guanine nucleotide-binding protein subunit alpha-12 [Callithrix jacchus] | 0 | 29.189 |
| 296202997 | PREDICTED: guanine nucleotide-binding protein subunit alpha-13 isoform X1 [Callithrix jacchus] | 0 | 74.913 |
| 296209836 | PREDICTED: guanine nucleotide-binding protein G(i) subunit alpha-1 isoform X1 [Callithrix jacchus] | 0 | 254.728 |
| 296225271 | PREDICTED: guanine nucleotide-binding protein G(i) subunit alpha-2 [Callithrix jacchus] | 0 | 256.88 |
| 1061012643 | PREDICTED: guanine nucleotide-binding protein G(k) subunit alpha [Callithrix jacchus] | 0 | 166.58 |
| 675715429 | PREDICTED: guanine nucleotide-binding protein G(olf) subunit alpha isoform X2 [Callithrix jacchus] | 0 | 96.853 |
| 296231109 | PREDICTED: guanine nucleotide-binding protein G(o) subunit alpha [Callithrix jacchus] | 0 | 380.591 |
| 296189680 | PREDICTED: guanine nucleotide-binding protein G(q) subunit alpha [Callithrix jacchus] | 0 | 168.384 |
| 675657292 | PREDICTED: guanine nucleotide-binding protein G(s) subunit alpha isoforms XLas isoform X1 [Callithrix jacchus] | 0 | 117.083 |
| 296191481 | PREDICTED: guanine nucleotide-binding protein G(z) subunit alpha [Callithrix jacchus] | 0 | 184.834 |
| 1061009740 | PREDICTED: guanine nucleotide-binding protein G(I)/G(S)/G(T) subunit beta-1 [Callithrix jacchus] | 0 | 322.453 |
| 675642346 | PREDICTED: guanine nucleotide-binding protein G(I)/G(S)/G(T) subunit beta-2 [Callithrix jacchus] | 0 | 247.229 |
| 390459533 | PREDICTED: receptor of activated protein C kinase 1 [Callithrix jacchus] | 0 | 89.237 |
| 675725314 | PREDICTED: guanine nucleotide-binding protein subunit beta-4 isoform X1 [Callithrix jacchus] | 0 | 176.466 |
| 296214036 | PREDICTED: guanine nucleotide-binding protein subunit beta-5 isoform X2 [Callithrix jacchus] | 0 | 151.04 |
| 326368250 | guanine nucleotide-binding protein G(I)/G(S)/G(O) subunit gamma-10 [Callithrix jacchus] | 0 | 36.999 |
| 675670701 | PREDICTED: guanine nucleotide-binding protein G(I)/G(S)/G(O) subunit gamma-12 [Callithrix jacchus] | 0 | 48.425 |
| 1060939669 | PREDICTED: guanine nucleotide-binding protein G(I)/G(S)/G(O) subunit gamma-2 [Callithrix jacchus] | 0 | 48.369 |
| 1060943898 | PREDICTED: guanine nucleotide-binding protein G(I)/G(S)/G(O) subunit gamma-3 [Callithrix jacchus] | 0 | 83.533 |
| 1060943893 | PREDICTED: seipin isoform X1 [Callithrix jacchus] | 0 | 25.577 |
| 296230916 | PREDICTED: guanine nucleotide-binding protein G(I)/G(S)/G(O) subunit gamma-4 [Callithrix jacchus] | 0 | 25.373 |
| 1060975883 | PREDICTED: guanine nucleotide-binding protein G(I)/G(S)/G(O) subunit gamma-7 [Callithrix jacchus] | 0 | 48.237 |
| 296220967 | PREDICTED: aspartate aminotransferase, cytoplasmic [Callithrix jacchus] | 0 | 25.807 |
| 296231243 | PREDICTED: aspartate aminotransferase, mitochondrial [Callithrix jacchus] | 0 | 79.991 |
| 532217405 | glypican-1 precursor [Callithrix jacchus] | 0 | 139.961 |
| 296236446 | PREDICTED: glypican-4 [Callithrix jacchus] | 0 | 75.493 |
| 532446899 | glucose-6-phosphate isomerase isoform 2 [Callithrix jacchus] | 0 | 60.985 |
| 296210736 | PREDICTED: prosaposin receptor GPR37 [Callithrix jacchus] | 0 | 16.611 |
| 390464779 | PREDICTED: lanC-like protein 1 isoform X1 [Callithrix jacchus] | 0 | 130.803 |
| 63115346 | cytosolic glutathione peroxidase [Callithrix jacchus] | 0 | 39.027 |
| 296203176 | PREDICTED: growth factor receptor-bound protein 2 [Callithrix jacchus] | 0 | 62.4 |
| 532511194 | putative G-protein coupled receptor 162 isoform 2 [Callithrix jacchus] | 0 | 15.105 |
| 296211239 | PREDICTED: protein C10 isoform X1 [Callithrix jacchus] | 0 | 49.494 |
| 296193304 | PREDICTED: glutamate receptor 1 isoform X2 [Callithrix jacchus] | 0 | 174.058 |
| 532338867 | glutamate receptor, ionotropic kainate 3 precursor [Callithrix jacchus] | 0 | 48.63 |
| 532287888 | glutamate [NMDA] receptor subunit zeta-1 isoform NR1-3 precursor [Callithrix jacchus] | 0 | 206.156 |
| 390467285 | PREDICTED: LOW QUALITY PROTEIN: glutamate receptor ionotropic, NMDA 2B [Callithrix jacchus] | 0 | 432.211 |
| 532342533 | metabotropic glutamate receptor 1 isoform alpha precursor [Callithrix jacchus] | 0 | 56.842 |
| 296213817 | PREDICTED: protein disulfide-isomerase A3 [Callithrix jacchus] | 0 | 60.808 |
| 675639373 | PREDICTED: 78 kDa glucose-regulated protein [Callithrix jacchus] | 0 | 279.562 |
| 532507207 | general transcription factor II-I isoform 1 [Callithrix jacchus] | 0 | 40.8 |
| 532524034 | GTP-binding protein 1 [Callithrix jacchus] | 0 | 44.225 |
| 1060954796 | PREDICTED: membrane-associated guanylate kinase, WW and PDZ domain-containing protein 1 [Callithrix jacchus] | 0 | 127.798 |
| 675766132 | PREDICTED: glycerol kinase isoform X4 [Callithrix jacchus] | 0 | 135.242 |
| 296235560 | PREDICTED: 3-hydroxyacyl-CoA dehydrogenase type-2 [Callithrix jacchus] | 0 | 68.365 |
| 675765307 | PREDICTED: cytochrome c-type heme lyase [Callithrix jacchus] | 0 | 38.193 |
| 1060973540 | PREDICTED: potassium/sodium hyperpolarization-activated cyclic nucleotide-gated channel 2 [Callithrix jacchus] | 0 | 14.659 |
| 391852050 | huntingtin [Callithrix jacchus] | 0 | 22.033 |
| 532444263 | hepatocyte growth factor-regulated tyrosine kinase substrate [Callithrix jacchus] | 0 | 17.183 |
| 532447223 | hexokinase-1 isoform HKI [Callithrix jacchus] | 0 | 701.198 |
| 390477571 | PREDICTED: dimethyladenosine transferase 2, mitochondrial isoform X1 [Callithrix jacchus] | 0.001 | 6.022 |
| 675705699 | PREDICTED: heme oxygenase 2 [Callithrix jacchus] | 0 | 11.281 |
| 532343747 | heterogeneous nuclear ribonucleoproteins C1/C2 isoform b [Callithrix jacchus] | 0 | 70.005 |
| 532345905 | heterogeneous nuclear ribonucleoprotein A1 isoform a [Callithrix jacchus] | 0 | 167.754 |
| 296233743 | PREDICTED: LOW QUALITY PROTEIN: heterogeneous nuclear ribonucleoprotein L [Callithrix jacchus] | 0 | 159.076 |
| 532517785 | heterochromatin protein 1-binding protein 3 [Callithrix jacchus] | 0 | 45.724 |
| 296207386 | PREDICTED: neuron-specific calcium-binding protein hippocalcin [Callithrix jacchus] | 0 | 61.219 |
| 532501705 | hypoxanthine-guanine phosphoribosyltransferase [Callithrix jacchus] | 0 | 19.728 |
| 675763277 | PREDICTED: LOW QUALITY PROTEIN: protein arginine N-methyltransferase 1 [Callithrix jacchus] | 0 | 71.471 |
| 675737386 | PREDICTED: ribonuclease UK114 [Callithrix jacchus] | 0 | 14.585 |
| 675694793 | PREDICTED: heat shock cognate 71 kDa protein isoform X1 [Callithrix jacchus] | 0 | 479.631 |
| 296193889 | PREDICTED: peroxisomal multifunctional enzyme type 2 isoform X1 [Callithrix jacchus] | 0 | 301.078 |
| 675638135 | PREDICTED: dnaJ homolog subfamily A member 1 [Callithrix jacchus] | 0 | 241.064 |
| 296203664 | PREDICTED: heat shock protein 105 kDa isoform X1 [Callithrix jacchus] | 0 | 333.077 |
| 296192276 | PREDICTED: heat shock protein beta-1 [Callithrix jacchus] | 0 | 55.022 |
| 675664919 | PREDICTED: 60 kDa heat shock protein, mitochondrial [Callithrix jacchus] | 0 | 268.371 |
| 1060939881 | PREDICTED: heat shock-related 70 kDa protein 2 [Callithrix jacchus] | 0 | 202.119 |
| 675692814 | PREDICTED: heat shock protein HSP 90-alpha [Callithrix jacchus] | 0 | 108.258 |
| 296193719 | PREDICTED: heat shock 70 kDa protein 4 isoform X1 [Callithrix jacchus] | 0 | 265.933 |
| 296192860 | PREDICTED: stress-70 protein, mitochondrial [Callithrix jacchus] | 0 | 273.55 |
| 532213181 | ELAV-like protein 1 [Callithrix jacchus] | 0 | 46.925 |
| 675638037 | PREDICTED: ELAV-like protein 2 isoform X3 [Callithrix jacchus] | 0 | 95.954 |
| 1060973103 | PREDICTED: ELAV-like protein 3 isoform X1 [Callithrix jacchus] | 0 | 44.85 |
| 675670045 | PREDICTED: ELAV-like protein 4 isoform X1 [Callithrix jacchus] | 0 | 100.317 |
| 1061013984 | PREDICTED: islet cell autoantigen 1 isoform X6 [Callithrix jacchus] | 0 | 14.316 |
| 1060975487 | PREDICTED: intercellular adhesion molecule 5 [Callithrix jacchus] | 0 | 147.747 |
| 296205433 | PREDICTED: isocitrate dehydrogenase [NADP] cytoplasmic [Callithrix jacchus] | 0 | 19.856 |
| 296236714 | PREDICTED: isocitrate dehydrogenase [NAD] subunit gamma, mitochondrial [Callithrix jacchus] | 0 | 56.882 |
| 675771036 | PREDICTED: NF-kappa-B essential modulator isoform X1 [Callithrix jacchus] | 0 | 20.378 |
| 390478548 | PREDICTED: LOW QUALITY PROTEIN: interleukin enhancer-binding factor 3 [Callithrix jacchus] | 0 | 79.398 |
| 296202645 | PREDICTED: importin subunit beta-1 [Callithrix jacchus] | 0 | 131.158 |
| 532524052 | eukaryotic translation initiation factor 3 subunit E [Callithrix jacchus] | 0 | 61.202 |
| 532524710 | E3 ubiquitin-protein ligase Itchy homolog isoform 2 [Callithrix jacchus] | 0 | 37.913 |
| 1060953090 | PREDICTED: integrin beta-1-binding protein 1 isoform X1 [Callithrix jacchus] | 0 | 11.85 |
| 1060955751 | PREDICTED: leukocyte surface antigen CD47 isoform X1 [Callithrix jacchus] | 0 | 24.66 |
| 675636795 | PREDICTED: integral membrane protein 2B [Callithrix jacchus] | 0 | 30.166 |
| 675729887 | PREDICTED: inositol 1,4,5-trisphosphate receptor type 1 isoform X1 [Callithrix jacchus] | 0 | 184.265 |
| 532346167 | intersectin-1 isoform ITSN-l [Callithrix jacchus] | 0 | 537.703 |
| 675670552 | PREDICTED: tyrosine-protein kinase JAK1 [Callithrix jacchus] | 0 | 18.773 |
| 675756670 | PREDICTED: caytaxin [Callithrix jacchus] | 0 | 37.089 |
| 532348611 | potassium voltage-gated channel subfamily A member 1 [Callithrix jacchus] | 0 | 53.724 |
| 1061012684 | PREDICTED: potassium voltage-gated channel subfamily A member 2 [Callithrix jacchus] | 0 | 58.729 |
| 532441105 | voltage-gated potassium channel subunit beta-2 isoform 1 [Callithrix jacchus] | 0 | 145.855 |
| 532347043 | potassium voltage-gated channel subfamily B member 1 [Callithrix jacchus] | 0 | 88.371 |
| 675763240 | PREDICTED: potassium voltage-gated channel subfamily C member 3 [Callithrix jacchus] | 0 | 7.523 |
| 1061015467 | PREDICTED: potassium voltage-gated channel subfamily D member 2 [Callithrix jacchus] | 0 | 60.356 |
| 532333915 | potassium voltage-gated channel subfamily H member 1 isoform 1 [Callithrix jacchus] | 0 | 15.429 |
| 296191890 | PREDICTED: inward rectifier potassium channel 4 [Callithrix jacchus] | 0 | 17.137 |
| 532285268 | calcium-activated potassium channel subunit alpha-1 isoform f [Callithrix jacchus] | 0 | 153.132 |
| 1060972295 | PREDICTED: LOW QUALITY PROTEIN: far upstream element-binding protein 2, partial [Callithrix jacchus] | 0 | 60.658 |
| 296206929 | PREDICTED: dolichyl-diphosphooligosaccharide--protein glycosyltransferase 48 kDa subunit [Callithrix jacchus] | 0 | 22.785 |
| 532292208 | kinesin-like protein KIF21A isoform 1 [Callithrix jacchus] | 0 | 132.573 |
| 1060961243 | PREDICTED: kinesin-like protein KIF21B [Callithrix jacchus] | 0 | 58.19 |
| 675644673 | PREDICTED: kinesin-like protein KIF3A isoform X1 [Callithrix jacchus] | 0 | 102.017 |
| 296199864 | PREDICTED: kinesin-like protein KIF3B isoform X1 [Callithrix jacchus] | 0 | 47.727 |
| 296206410 | PREDICTED: kinesin-1 heavy chain [Callithrix jacchus] | 0 | 157.954 |
| 390477127 | PREDICTED: LOW QUALITY PROTEIN: kinesin-associated protein 3 [Callithrix jacchus] | 0 | 23.407 |
| 675693059 | PREDICTED: kinesin light chain 1 isoform X21 [Callithrix jacchus] | 0 | 119.658 |
| 1060955425 | PREDICTED: importin subunit alpha-5 [Callithrix jacchus] | 0 | 42.243 |
| 1060933739 | PREDICTED: importin subunit alpha-4 [Callithrix jacchus] | 0 | 31.609 |
| 296227675 | PREDICTED: importin subunit alpha-3 [Callithrix jacchus] | 0 | 35.017 |
| 296211766 | PREDICTED: LOW QUALITY PROTEIN: keratin, type II cytoskeletal 1 [Callithrix jacchus] | 0 | 88.288 |
| 1060939751 | PREDICTED: kinectin [Callithrix jacchus] | 0 | 311.568 |
| 167045807 | L1 cell adhesion molecule isoform 1 precursor (predicted) [Callithrix jacchus] | 0 | 200.871 |
| 675668758 | PREDICTED: stathmin [Callithrix jacchus] | 0 | 17.9 |
| 390463601 | PREDICTED: LOW QUALITY PROTEIN: LIM and SH3 domain protein 1 [Callithrix jacchus] | 0 | 95.058 |
| 532337435 | rho guanine nucleotide exchange factor 2 isoform 1 [Callithrix jacchus] | 0 | 408.149 |
| 402239590 | lactate dehydrogenase A [Callithrix jacchus] | 0 | 82.931 |
| 532292138 | L-lactate dehydrogenase B chain [Callithrix jacchus] | 0 | 139.628 |
| 390469123 | PREDICTED: galectin-3 [Callithrix jacchus] | 0 | 25.759 |
| 532510754 | lethal(2) giant larvae protein homolog 1 [Callithrix jacchus] | 0 | 155.91 |
| 296219361 | PREDICTED: 40S ribosomal protein S2 [Callithrix jacchus] | 0 | 103.233 |
| 296229150 | PREDICTED: lamin [Callithrix jacchus] | 0 | 98.338 |
| 296193836 | PREDICTED: lamin-B1 [Callithrix jacchus] | 0 | 97.845 |
| 532338430 | lamin-B2 [Callithrix jacchus] | 0 | 156.162 |
| 532341497 | prolow-density lipoprotein receptor-related protein 1 precursor [Callithrix jacchus] | 0 | 136.106 |
| 532445993 | leucine-rich repeat flightless-interacting protein 1 isoform 1 [Callithrix jacchus] | 0 | 58.291 |
| 675733207 | PREDICTED: tyrosine-protein kinase Lyn isoform X2 [Callithrix jacchus] | 0 | 48.334 |
| 296199048 | PREDICTED: myristoylated alanine-rich C-kinase substrate [Callithrix jacchus] | 0 | 106.46 |
| 675766444 | PREDICTED: amine oxidase [flavin-containing] A [Callithrix jacchus] | 0 | 106.059 |
| 1060940704 | PREDICTED: MAP/microtubule affinity-regulating kinase 3 [Callithrix jacchus] | 0 | 244.719 |
| 1060962144 | PREDICTED: matrin-3 isoform X1 [Callithrix jacchus] | 0 | 172.199 |
| 1060949316 | PREDICTED: myelin basic protein [Callithrix jacchus] | 0 | 235.923 |
| 1060933370 | PREDICTED: LOW QUALITY PROTEIN: guanine nucleotide exchange factor DBS [Callithrix jacchus] | 0 | 42.676 |
| 390470681 | PREDICTED: 4F2 cell-surface antigen heavy chain [Callithrix jacchus] | 0 | 32.297 |
| 390480372 | PREDICTED: methyl-CpG-binding protein 2 [Callithrix jacchus] | 0 | 72.643 |
| 532208605 | lactadherin isoform a preproprotein [Callithrix jacchus] | 0.001 | 6.712 |
| 296236904 | PREDICTED: macrophage migration inhibitory factor [Callithrix jacchus] | 0 | 10.041 |
| 1061010775 | PREDICTED: MARCKS-related protein [Callithrix jacchus] | 0 | 50.779 |
| 1060958752 | PREDICTED: myelin-associated oligodendrocyte basic protein isoform X1 [Callithrix jacchus] | 0 | 50.752 |
| 380848769 | myelin-oligodendrocyte glycoprotein isoform 1 precursor [Callithrix jacchus] | 0 | 192.888 |
| 296192270 | PREDICTED: malate dehydrogenase, mitochondrial [Callithrix jacchus] | 0 | 123.8 |
| 532209689 | malate dehydrogenase, cytoplasmic isoform 2 [Callithrix jacchus] | 0 | 75.807 |
| 1060958138 | PREDICTED: ras-related protein M-Ras isoform X1 [Callithrix jacchus] | 0 | 51.467 |
| 532522948 | moesin [Callithrix jacchus] | 0 | 113.974 |
| 301170770 | cytochrome c oxidase subunit I (mitochondrion) [Callithrix jacchus] | 0 | 56.251 |
| 722490420 | cytochrome c oxidase subunit II (mitochondrion) [Callithrix jacchus] | 0 | 62.051 |
| 722490417 | NADH dehydrogenase subunit 1 (mitochondrion) [Callithrix jacchus] | 0 | 13.734 |
| 722490418 | NADH dehydrogenase subunit 2 (mitochondrion) [Callithrix jacchus] | 0 | 37.391 |
| 301170777 | NADH dehydrogenase subunit 4 (mitochondrion) [Callithrix jacchus] | 0 | 87.311 |
| 301170778 | NADH dehydrogenase subunit 5 (mitochondrion) [Callithrix jacchus] | 0 | 56.377 |
| 296194378 | PREDICTED: LOW QUALITY PROTEIN: microtubule-associated protein 1B [Callithrix jacchus] | 0 | 1281.964 |
| 532284634 | microtubule-associated protein 2 isoform 1 [Callithrix jacchus] | 0 | 1385.833 |
| 1060954170 | PREDICTED: microtubule-associated protein 4 isoform X2 [Callithrix jacchus] | 0 | 213.842 |
| 675696856 | PREDICTED: microtubule-associated protein 6 [Callithrix jacchus] | 0 | 474.823 |
| 1060995948 | PREDICTED: microtubule-associated protein tau [Callithrix jacchus] | 0 | 178.732 |
| 296236635 | PREDICTED: myotubularin isoform X1 [Callithrix jacchus] | 0 | 20.321 |
| 296229087 | PREDICTED: metaxin-1 isoform X1 [Callithrix jacchus] | 0 | 83.264 |
| 675687000 | PREDICTED: myelin expression factor 2 isoform X1 [Callithrix jacchus] | 0 | 51.208 |
| 1060937228 | PREDICTED: myosin-9 [Callithrix jacchus] | 0 | 924.941 |
| 675680090 | PREDICTED: myosin light polypeptide 6 [Callithrix jacchus] | 0 | 106.47 |
| 675664833 | PREDICTED: unconventional myosin-Ib isoform X1 [Callithrix jacchus] | 0 | 261.57 |
| 1060976396 | PREDICTED: unconventional myosin-If [Callithrix jacchus] | 0 | 10.47 |
| 1060938715 | PREDICTED: LOW QUALITY PROTEIN: unconventional myosin-Va [Callithrix jacchus] | 0 | 1086.934 |
| 675653306 | PREDICTED: LOW QUALITY PROTEIN: unconventional myosin-VI [Callithrix jacchus] | 0 | 369.882 |
| 390467981 | PREDICTED: protein phosphatase 1 regulatory subunit 12A isoform X3 [Callithrix jacchus] | 0 | 169.858 |
| 1060944489 | PREDICTED: nucleosome assembly protein 1-like 4 isoform X1 [Callithrix jacchus] | 0 | 53.424 |
| 675656437 | PREDICTED: beta-soluble NSF attachment protein isoform X1 [Callithrix jacchus] | 0 | 191.939 |
| 1060941385 | PREDICTED: neural cell adhesion molecule 1 [Callithrix jacchus] | 0 | 418.169 |
| 1060969353 | PREDICTED: neural cell adhesion molecule 2 [Callithrix jacchus] | 0 | 201.195 |
| 1060951385 | PREDICTED: cytoplasmic protein NCK2 isoform X1 [Callithrix jacchus] | 0 | 61.491 |
| 1061007157 | PREDICTED: nucleolin [Callithrix jacchus] | 0 | 170.829 |
| 390459299 | PREDICTED: NADH dehydrogenase [ubiquinone] 1 alpha subcomplex subunit 2 isoform X1 [Callithrix jacchus] | 0 | 30.81 |
| 1060933597 | PREDICTED: cytochrome c oxidase subunit NDUFA4-like [Callithrix jacchus] | 0 | 33.275 |
| 296194616 | PREDICTED: NADH dehydrogenase [ubiquinone] iron-sulfur protein 4, mitochondrial isoform X1 [Callithrix jacchus] | 0 | 50.35 |
| 296218952 | PREDICTED: NADH dehydrogenase [ubiquinone] flavoprotein 1, mitochondrial isoform X1 [Callithrix jacchus] | 0 | 238.268 |
| 675666542 | PREDICTED: septin-2 isoform X1 [Callithrix jacchus] | 0 | 322.681 |
| 675686079 | PREDICTED: neogenin isoform X1 [Callithrix jacchus] | 0 | 48.077 |
| 675659669 | PREDICTED: neurofibromin isoform X1 [Callithrix jacchus] | 0 | 293.715 |
| 1060936958 | PREDICTED: LOW QUALITY PROTEIN: merlin [Callithrix jacchus] | 0 | 29.378 |
| 296221730 | PREDICTED: neurofilament light polypeptide [Callithrix jacchus] | 0 | 381.902 |
| 1060949795 | PREDICTED: neurofilament medium polypeptide isoform X1 [Callithrix jacchus] | 0 | 487.035 |
| 532508568 | cysteine desulfurase, mitochondrial isoform a precursor [Callithrix jacchus] | 0 | 13.499 |
| 296191617 | PREDICTED: protein NipSnap homolog 1 [Callithrix jacchus] | 0 | 44.53 |
| 296191340 | PREDICTED: 39S ribosomal protein L40, mitochondrial [Callithrix jacchus] | 0 | 29.839 |
| 325910861 | nucleoside diphosphate kinase A [Callithrix jacchus] | 0 | 87.303 |
| 325910863 | nucleoside diphosphate kinase B [Callithrix jacchus] | 0 | 104.072 |
| 296201692 | PREDICTED: glycylpeptide N-tetradecanoyltransferase 1 isoform X1 [Callithrix jacchus] | 0 | 16.697 |
| 296194659 | PREDICTED: NAD(P) transhydrogenase, mitochondrial [Callithrix jacchus] | 0 | 128.468 |
| 532504078 | 39S ribosomal protein L49, mitochondrial [Callithrix jacchus] | 0 | 13.969 |
| 296193667 | PREDICTED: nucleophosmin [Callithrix jacchus] | 0 | 195.042 |
| 675661613 | PREDICTED: neuronal pentraxin-1 [Callithrix jacchus] | 0 | 42.36 |
| 1060950908 | PREDICTED: LOW QUALITY PROTEIN: neurexin-1 [Callithrix jacchus] | 0 | 237.858 |
| 1060944004 | PREDICTED: LOW QUALITY PROTEIN: neurexin-2-beta [Callithrix jacchus] | 0 | 113.008 |
| 1060940948 | PREDICTED: neurexin 3 [Callithrix jacchus] | 0 | 314.65 |
| 532524258 | vesicle-fusing ATPase [Callithrix jacchus] | 0 | 438.286 |
| 1060934192 | PREDICTED: BDNF/NT-3 growth factors receptor [Callithrix jacchus] | 0 | 53.912 |
| 296207169 | PREDICTED: nuclear migration protein nudC [Callithrix jacchus] | 0 | 19.351 |
| 532213481 | protein numb homolog isoform 1 [Callithrix jacchus] | 0 | 112.025 |
| 532276178 | numb-like protein [Callithrix jacchus] | 0 | 114.229 |
| 296209608 | PREDICTED: neurexophilin-1 [Callithrix jacchus] | 0 | 8.973 |
| 1061013363 | PREDICTED: 2-oxoglutarate dehydrogenase, mitochondrial isoform X1 [Callithrix jacchus] | 0 | 612.144 |
| 296202091 | PREDICTED: oligodendrocyte-myelin glycoprotein [Callithrix jacchus] | 0 | 84.835 |
| 296195612 | PREDICTED: heat shock 70 kDa protein 4L [Callithrix jacchus] | 0 | 176.441 |
| 296227579 | PREDICTED: claudin-11 [Callithrix jacchus] | 0 | 118.53 |
| 675777894 | PREDICTED: protein disulfide-isomerase isoform X1 [Callithrix jacchus] | 0 | 19.95 |
| 532218439 | polyadenylate-binding protein 1 [Callithrix jacchus] | 0 | 167.023 |
| 675657675 | PREDICTED: platelet-activating factor acetylhydrolase IB subunit alpha [Callithrix jacchus] | 0 | 79.915 |
| 532507082 | platelet-activating factor acetylhydrolase IB subunit beta isoform a [Callithrix jacchus] | 0 | 10.924 |
| 1061011331 | PREDICTED: peroxiredoxin-1 [Callithrix jacchus] | 0 | 143.769 |
| 532519804 | serine/threonine-protein kinase PAK 1 isoform 2 [Callithrix jacchus] | 0 | 92.832 |
| 675769246 | PREDICTED: serine/threonine-protein kinase PAK 3 isoform X1 [Callithrix jacchus] | 0 | 65.046 |
| 532199877 | paralemmin-1 isoform 1 [Callithrix jacchus] | 0 | 295.548 |
| 1061020512 | PREDICTED: immunoglobulin-binding protein 1 [Callithrix jacchus] | 0 | 35.869 |
| 1061016920 | PREDICTED: LOW QUALITY PROTEIN: poly(rC)-binding protein 2 [Callithrix jacchus] | 0 | 120.766 |
| 532284048 | protocadherin-10 isoform 1 precursor [Callithrix jacchus] | 0 | 94.993 |
| 1060949979 | PREDICTED: pericentriolar material 1 protein [Callithrix jacchus] | 0 | 72.919 |
| 296199435 | PREDICTED: protein-L-isoaspartate(D-aspartate) O-methyltransferase isoform X1 [Callithrix jacchus] | 0 | 97.87 |
| 675766528 | PREDICTED: cyclin-dependent kinase 16 [Callithrix jacchus] | 0 | 45.538 |
| 532290248 | pyruvate carboxylase, mitochondrial precursor [Callithrix jacchus] | 0 | 68.123 |
| 532501925 | programmed cell death protein 6 isoform 1 [Callithrix jacchus] | 0 | 9.718 |
| 532521936 | programmed cell death 6-interacting protein isoform 2 [Callithrix jacchus] | 0 | 212.604 |
| 675663542 | PREDICTED: calcium/calmodulin-dependent 3',5'-cyclic nucleotide phosphodiesterase 1A isoform X1 [Callithrix jacchus] | 0 | 15.301 |
| 296211904 | PREDICTED: calcium/calmodulin-dependent 3',5'-cyclic nucleotide phosphodiesterase 1B isoform X1 [Callithrix jacchus] | 0 | 31.4 |
| 532337035 | cAMP-specific 3',5'-cyclic phosphodiesterase 4A isoform 1 [Callithrix jacchus] | 0 | 26.1 |
| 1061020067 | PREDICTED: pyruvate dehydrogenase E1 component subunit alpha, somatic form, mitochondrial [Callithrix jacchus] | 0 | 255.445 |
| 532504907 | protein-arginine deiminase type-2 [Callithrix jacchus] | 0 | 95.797 |
| 675660315 | PREDICTED: LOW QUALITY PROTEIN: pyruvate dehydrogenase kinase, isozyme 2 [Callithrix jacchus] | 0 | 64.41 |
| 1060960585 | PREDICTED: astrocytic phosphoprotein PEA-15 isoform X3 [Callithrix jacchus] | 0 | 26.043 |
| 675755692 | PREDICTED: ATP-dependent 6-phosphofructokinase, liver type isoform X1 [Callithrix jacchus] | 0 | 120.773 |
| 1061015727 | PREDICTED: ATP-dependent 6-phosphofructokinase, muscle type [Callithrix jacchus] | 0 | 245.928 |
| 390463383 | PREDICTED: profilin-1 [Callithrix jacchus] | 0 | 19.419 |
| 359338981 | phosphoglycerate mutase 1 [Callithrix jacchus] | 0 | 113.45 |
| 675768055 | PREDICTED: phosphoglycerate kinase 1 [Callithrix jacchus] | 0 | 124.777 |
| 1060998022 | PREDICTED: prohibitin [Callithrix jacchus] | 0 | 130.672 |
| 675682353 | PREDICTED: phosphate carrier protein, mitochondrial isoform X2 [Callithrix jacchus] | 0 | 90.306 |
| 1060937323 | PREDICTED: PRKCA-binding protein isoform X1 [Callithrix jacchus] | 0 | 21.529 |
| 1060965272 | PREDICTED: phosphatidylinositol 3-kinase regulatory subunit alpha isoform X1 [Callithrix jacchus] | 0 | 18.579 |
| 675756643 | PREDICTED: phosphatidylinositol 4-phosphate 5-kinase type-1 gamma [Callithrix jacchus] | 0 | 146.493 |

| | | | | |
|---|---|---|---|---|
| 296206295 | PREDICTED: phosphatidylinositol 5-phosphate 4-kinase type-2 alpha isoform X1 [Callithrix jacchus] | 0 | 99.796 | [illegible] |
| 675742731 | PREDICTED: phosphatidylinositol 4-phosphate 5-kinase type-1 alpha isoform X1 [Callithrix jacchus] | 0 | 87.484 | [illegible] |
| 675703149 | PREDICTED: membrane-associated phosphatidylinositol transfer protein 1 isoform X1 [Callithrix jacchus] | 0 | 57.71 | [illegible] |
| 1060938161 | PREDICTED: pyruvate kinase PKM isoform X1 [Callithrix jacchus] | 0 | 247.594 | [illegible] |
| 532521172 | cAMP-dependent protein kinase catalytic subunit alpha isoform 1 [Callithrix jacchus] | 0 | 116.425 | [illegible] |
| 675670970 | PREDICTED: cAMP-dependent protein kinase catalytic subunit beta isoform X2 [Callithrix jacchus] | 0 | 119.18 | [illegible] |
| 1060998649 | PREDICTED: protein kinase C alpha type [Callithrix jacchus] | 0 | 141.649 | [illegible] |
| 532344533 | protein kinase C beta type isoform 2 [Callithrix jacchus] | 0 | 237.181 | [illegible] |
| 675763884 | PREDICTED: protein kinase C gamma type [Callithrix jacchus] | 0 | 417.144 | [illegible] |
| 1060954545 | PREDICTED: protein kinase C delta type [Callithrix jacchus] | 0 | 19.292 | [illegible] |
| 532521246 | protein kinase C epsilon type [Callithrix jacchus] | 0 | 76.092 | [illegible] |
| 1060992206 | PREDICTED: 1-phosphatidylinositol 4,5-bisphosphate phosphodiesterase beta-1 [Callithrix jacchus] | 0 | 767.97 | [illegible] |
| 1060943976 | PREDICTED: 1-phosphatidylinositol 4,5-bisphosphate phosphodiesterase beta-3 [Callithrix jacchus] | 0 | 102.274 | [illegible] |
| 1060955933 | PREDICTED: LOW QUALITY PROTEIN: plectin [Callithrix jacchus] | 0 | 1953.23 | [illegible] |
| 532513324 | proliferation-associated protein 2G4 [Callithrix jacchus] | 0 | 39.061 | [illegible] |
| 296236083 | PREDICTED: myelin proteolipid protein isoform X1 [Callithrix jacchus] | 0 | 126.126 | [illegible] |
| 296214868 | PREDICTED: pinin isoform X1 [Callithrix jacchus] | 0 | 16.159 | [illegible] |
| 532518720 | septin-5 isoform 1 [Callithrix jacchus] | 0 | 304.293 | [illegible] |
| 1060936574 | PREDICTED: septin-5 [Callithrix jacchus] | 0 | 242.884 | [illegible] |
| 1061014415 | PREDICTED: synaptophysin-like protein 1 isoform X1 [Callithrix jacchus] | 0 | 10.218 | [illegible] |
| 296213430 | PREDICTED: peptidyl-prolyl cis-trans isomerase B [Callithrix jacchus] | 0 | 67.662 | [illegible] |
| 532449211 | serine/threonine-protein phosphatase PP1-alpha catalytic subunit isoform 1 [Callithrix jacchus] | 0 | 194.802 | [illegible] |
| 675723479 | PREDICTED: serine/threonine-protein phosphatase PP1-beta catalytic subunit isoform X1 [Callithrix jacchus] | 0 | 182.077 | [illegible] |
| 390468177 | PREDICTED: serine/threonine-protein phosphatase PP1-gamma catalytic subunit [Callithrix jacchus] | 0 | 197.358 | [illegible] |
| 296202728 | PREDICTED: protein phosphatase 1 regulatory subunit 1B isoform X1 [Callithrix jacchus] | 0 | 16.785 | [illegible] |
| 390459640 | PREDICTED: serine/threonine-protein phosphatase 2A catalytic subunit alpha isoform [Callithrix jacchus] | 0 | 146.753 | [illegible] |
| 296221947 | PREDICTED: serine/threonine-protein phosphatase 2A catalytic subunit beta isoform [Callithrix jacchus] | 0 | 147.305 | [illegible] |
| 296195900 | PREDICTED: serine/threonine-protein phosphatase 2B catalytic subunit alpha isoform [Callithrix jacchus] | 0 | 145.909 | [illegible] |
| 675708690 | PREDICTED: serine/threonine-protein phosphatase 2B catalytic subunit beta isoform isoform X1 [Callithrix jacchus] | 0 | 107.4 | [illegible] |
| 296221785 | PREDICTED: serine/threonine-protein phosphatase 2B catalytic subunit gamma isoform isoform X1 [Callithrix jacchus] | 0 | 37.352 | [illegible] |
| 296207603 | PREDICTED: palmitoyl-protein thioesterase 1 [Callithrix jacchus] | 0 | 25.213 | [illegible] |
| 1060998742 | PREDICTED: cAMP-dependent protein kinase type I-alpha regulatory subunit [Callithrix jacchus] | 0 | 138.932 | [illegible] |
| 1060963768 | PREDICTED: LOW QUALITY PROTEIN: cAMP-dependent protein kinase type I-beta regulatory subunit [Callithrix jacchus] | 0 | 86.014 | [illegible] |
| 390475024 | PREDICTED: cAMP-dependent protein kinase type II-alpha regulatory subunit isoform X1 [Callithrix jacchus] | 0 | 131.131 | [illegible] |
| 296209931 | PREDICTED: cAMP-dependent protein kinase type II-beta regulatory subunit [Callithrix jacchus] | 0 | 216.257 | [illegible] |
| 532285030 | C-Jun-amino-terminal kinase-interacting protein 1 [Callithrix jacchus] | 0 | 35.087 | [illegible] |
| 675656015 | PREDICTED: major prion protein [Callithrix jacchus] | 0 | 74.727 | [illegible] |
| 1061020560 | PREDICTED: ribose-phosphate pyrophosphokinase 1 [Callithrix jacchus] | 0 | 75.052 | [illegible] |
| 532519674 | puromycin-sensitive aminopeptidase [Callithrix jacchus] | 0 | 58.911 | [illegible] |
| 675708947 | PREDICTED: prosaposin isoform X1 [Callithrix jacchus] | 0 | 25.294 | [illegible] |
| 1060999653 | PREDICTED: cytohesin-1 isoform X1 [Callithrix jacchus] | 0 | 19.653 | [illegible] |
| 675762721 | PREDICTED: cytohesin-2 isoform X1 [Callithrix jacchus] | 0 | 25.476 | [illegible] |
| 532513268 | cytohesin-3 [Callithrix jacchus] | 0 | 51.687 | [illegible] |
| 296215138 | PREDICTED: proteasome subunit alpha type-3 isoform X1 [Callithrix jacchus] | 0 | 35.091 | [illegible] |
| 296199648 | PREDICTED: proteasome subunit beta type-1 [Callithrix jacchus] | 0 | 39.433 | [illegible] |
| 296228789 | PREDICTED: proteasome subunit beta type-4 [Callithrix jacchus] | 0 | 59.1 | [illegible] |
| 675688767 | PREDICTED: proteasome subunit beta type-5 [Callithrix jacchus] | 0 | 51.842 | [illegible] |
| 296202288 | PREDICTED: proteasome subunit beta type-6 isoform X1 [Callithrix jacchus] | 0 | 25.684 | [illegible] |
| 296190795 | PREDICTED: proteasome subunit beta type-7 [Callithrix jacchus] | 0 | 16.549 | [illegible] |
| 296215716 | PREDICTED: 26S protease regulatory subunit 4 [Callithrix jacchus] | 0 | 58.874 | [illegible] |
| 532522624 | 26S protease regulatory subunit 7 isoform 1 [Callithrix jacchus] | 0 | 82.641 | [illegible] |
| 532327639 | 26S protease regulatory subunit 6A [Callithrix jacchus] | 0 | 63.281 | [illegible] |
| 1060996156 | PREDICTED: LOW QUALITY PROTEIN: 26S protease regulatory subunit 8 [Callithrix jacchus] | 0 | 92.691 | [illegible] |
| 532508558 | phosphatidylserine synthase 1 [Callithrix jacchus] | 0 | 45.117 | [illegible] |
| 390473546 | PREDICTED: LOW QUALITY PROTEIN: protein-tyrosine kinase 2-beta [Callithrix jacchus] | 0 | 85.447 | [illegible] |
| 296210532 | PREDICTED: pleiotrophin isoform X1 [Callithrix jacchus] | 0 | 34.834 | [illegible] |
| 296198505 | PREDICTED: protein tyrosine phosphatase type IVA 1 [Callithrix jacchus] | 0 | 24.871 | [illegible] |
| 532504542 | protein tyrosine phosphatase type IVA 3 isoform 1 [Callithrix jacchus] | 0 | 14.01 | [illegible] |
| 675683661 | PREDICTED: tyrosine-protein phosphatase non-receptor type 11 isoform X1 [Callithrix jacchus] | 0 | 76.031 | [illegible] |
| 296204996 | PREDICTED: tyrosine-protein phosphatase non-receptor type 4 isoform X1 [Callithrix jacchus] | 0.001 | 6.712 | [illegible] |
| 532348905 | receptor-type tyrosine-protein phosphatase delta isoform 1 precursor [Callithrix jacchus] | 0 | 383.909 | [illegible] |
| 675669716 | PREDICTED: receptor-type tyrosine-protein phosphatase F [Callithrix jacchus] | 0 | 108.796 | [illegible] |
| 1060988305 | PREDICTED: receptor-type tyrosine-protein phosphatase kappa isoform X1 [Callithrix jacchus] | 0 | 46.493 | [illegible] |
| 296212367 | PREDICTED: receptor-type tyrosine-protein phosphatase R isoform X1 [Callithrix jacchus] | 0 | 9.738 | [illegible] |
| 1060975013 | PREDICTED: LOW QUALITY PROTEIN: receptor-type tyrosine-protein phosphatase S [Callithrix jacchus] | 0 | 386.225 | [illegible] |
| 532283714 | receptor-type tyrosine-protein phosphatase zeta isoform 1 precursor [Callithrix jacchus] | 0 | 265.938 | [illegible] |
| 1060963262 | PREDICTED: LOW QUALITY PROTEIN: transcriptional activator protein Pur-alpha [Callithrix jacchus] | 0 | 104.806 | [illegible] |
| 675672416 | PREDICTED: transcriptional activator protein Pur-beta [Callithrix jacchus] | 0 | 128.895 | [illegible] |
| 296208572 | PREDICTED: ATP-binding cassette sub-family D member 3 isoform X1 [Callithrix jacchus] | 0 | 101.922 | [illegible] |
| 1060944007 | PREDICTED: glycogen phosphorylase, muscle form [Callithrix jacchus] | 0 | 71.857 | [illegible] |
| 296223703 | PREDICTED: ras-related protein Rab-1A [Callithrix jacchus] | 0 | 48.778 | [illegible] |
| 532208303 | ras-related protein Rab-11B [Callithrix jacchus] | 0 | 32.303 | [illegible] |
| 532509257 | ras-related protein Rab-18 isoform 1 [Callithrix jacchus] | 0 | 24.525 | [illegible] |
| 296233287 | PREDICTED: ras-related protein Rab-3A [Callithrix jacchus] | 0 | 67.751 | [illegible] |
| 390467774 | PREDICTED: ras-related protein Rab-5B [Callithrix jacchus] | 0 | 36.263 | [illegible] |
| 1060998562 | PREDICTED: ras-related protein Rab-5C isoform X2 [Callithrix jacchus] | 0 | 33.486 | [illegible] |
| 296225985 | PREDICTED: ras-related protein Rab-7a [Callithrix jacchus] | 0 | 38.983 | [illegible] |
| 532215943 | ras-related C3 botulinum toxin substrate 1 isoform Rac1b [Callithrix jacchus] | 0 | 105.78 | [illegible] |
| 532515580 | ras-specific guanine nucleotide-releasing factor 1 isoform 1 [Callithrix jacchus] | 0 | 105.426 | [illegible] |
| 675645551 | PREDICTED: ras-specific guanine nucleotide-releasing factor 2 [Callithrix jacchus] | 0 | 193.961 | [illegible] |
| 390465661 | PREDICTED: histone-binding protein RBBP4 isoform X1 [Callithrix jacchus] | 0 | 26.977 | [illegible] |
| 1060944242 | PREDICTED: RNA-binding protein 4 isoform X1 [Callithrix jacchus] | 0 | 21.203 | [illegible] |
| 532339727 | membrane-associated phosphatidylinositol transfer protein 2 [Callithrix jacchus] | 0 | 68.433 | [illegible] |
| 1060941272 | PREDICTED: radixin isoform X1 [Callithrix jacchus] | 0 | 148.602 | [illegible] |
| 1060972513 | PREDICTED: regulator of nonsense transcripts 1 isoform X1 [Callithrix jacchus] | 0 | 171.204 | [illegible] |
| 532511170 | ralBP1-associated Eps domain-containing protein 1 isoform a [Callithrix jacchus] | 0 | 82.14 | [illegible] |
| 1060950446 | PREDICTED: ralA-binding protein 1 [Callithrix jacchus] | 0 | 48.903 | [illegible] |
| 1060953002 | PREDICTED: LOW QUALITY PROTEIN: rho-associated protein kinase 2 [Callithrix jacchus] | 0 | 131.749 | [illegible] |
| 532512390 | rabphilin-3A isoform 1 [Callithrix jacchus] | 0 | 305.465 | [illegible] |
| 675652340 | PREDICTED: 60S ribosomal protein L10a [Callithrix jacchus] | 0 | 93.322 | [illegible] |
| 390479205 | PREDICTED: 60S ribosomal protein L18 [Callithrix jacchus] | 0 | 57.742 | [illegible] |
| 296202709 | PREDICTED: 60S ribosomal protein L19 isoform X1 [Callithrix jacchus] | 0 | 43.624 | [illegible] |
| 1061009858 | PREDICTED: 60S ribosomal protein L22 isoform X1 [Callithrix jacchus] | 0 | 32.845 | [illegible] |
| 1060944570 | PREDICTED: 39S ribosomal protein L23, mitochondrial [Callithrix jacchus] | 0 | 22.314 | [illegible] |
| 675658525 | PREDICTED: 60S ribosomal protein L27 isoform X1 [Callithrix jacchus] | 0 | 32.745 | [illegible] |
| 1060971006 | PREDICTED: 60S ribosomal protein L28 isoform X1 [Callithrix jacchus] | 0 | 47.746 | [illegible] |
| 532279836 | 60S ribosomal protein L29 [Callithrix jacchus] | 0 | 12.106 | [illegible] |
| 675665684 | PREDICTED: 60S ribosomal protein L37a [Callithrix jacchus] | 0 | 33.58 | [illegible] |
| 675683659 | PREDICTED: 60S ribosomal protein L6 [Callithrix jacchus] | 0 | 129.867 | [illegible] |
| 675649807 | PREDICTED: 60S ribosomal protein L9 isoform X1 [Callithrix jacchus] | 0 | 101.27 | [illegible] |
| 296194833 | PREDICTED: activated RNA polymerase II transcriptional coactivator p15 [Callithrix jacchus] | 0 | 10.049 | [illegible] |
| 296199284 | PREDICTED: 40S ribosomal protein S12 [Callithrix jacchus] | 0 | 101.198 | [illegible] |
| 532512576 | 40S ribosomal protein S14 [Callithrix jacchus] | 0 | 56.385 | [illegible] |
| 296233780 | PREDICTED: 40S ribosomal protein S16 [Callithrix jacchus] | 0 | 24.428 | [illegible] |
| 675663222 | PREDICTED: 40S ribosomal protein S17 [Callithrix jacchus] | 0 | 90.13 | [illegible] |
| 296197873 | PREDICTED: 40S ribosomal protein S18 [Callithrix jacchus] | 0 | 55.331 | [illegible] |
| 1060975100 | PREDICTED: 40S ribosomal protein S19 isoform X1 [Callithrix jacchus] | 0 | 40.67 | [illegible] |
| 675760722 | PREDICTED: 40S ribosomal protein S24 [Callithrix jacchus] | 0 | 42.332 | [illegible] |
| 675673819 | PREDICTED: 40S ribosomal protein S3a [Callithrix jacchus] | 0 | 104.047 | [illegible] |
| 296234784 | PREDICTED: 40S ribosomal protein S5 [Callithrix jacchus] | 0 | 93.006 | [illegible] |
| 296189985 | PREDICTED: 40S ribosomal protein S6 [Callithrix jacchus] | 0 | 44.532 | [illegible] |
| 390462649 | PREDICTED: 40S ribosomal protein S7 [Callithrix jacchus] | 0 | 37.522 | [illegible] |
| 296207770 | PREDICTED: 40S ribosomal protein S8 [Callithrix jacchus] | 0 | 89.542 | [illegible] |
| 532283964 | reticulon-3 isoform e [Callithrix jacchus] | 0 | 182.563 | [illegible] |
| 532504387 | ruvB-like 2 [Callithrix jacchus] | 0 | 74.051 | [illegible] |
| 1060961953 | PREDICTED: ryanodine receptor 2 [Callithrix jacchus] | 0 | 385.879 | [illegible] |
| 296232338 | PREDICTED: protein S100-B [Callithrix jacchus] | 0 | 16.659 | [illegible] |
| 296207324 | PREDICTED: KH domain-containing, RNA-binding, signal transduction-associated protein 1 [Callithrix jacchus] | 0 | 31.913 | [illegible] |
| 675709111 | PREDICTED: GTP-binding protein SAR1a [Callithrix jacchus] | 0 | 16.278 | [illegible] |
| 296208728 | PREDICTED: serine--tRNA ligase, cytoplasmic isoform X1 [Callithrix jacchus] | 0 | 16.411 | [illegible] |
| 675683417 | PREDICTED: ataxin-2 isoform X1 [Callithrix jacchus] | 0 | 86.346 | [illegible] |
| 675641614 | PREDICTED: zinc finger CCCH domain-containing protein 7B [Callithrix jacchus] | 0 | 24.244 | [illegible] |
| 675659787 | PREDICTED: stromal cell-derived factor 2 isoform X1 [Callithrix jacchus] | 0 | 11.495 | [illegible] |
| 675686116 | PREDICTED: neuroplastin isoform X2 [Callithrix jacchus] | 0 | 109.059 | [illegible] |
| 1060959717 | PREDICTED: vesicle-trafficking protein SEC22b [Callithrix jacchus] | 0 | 34.49 | [illegible] |
| 296210590 | PREDICTED: exocyst complex component 4 isoform X1 [Callithrix jacchus] | 0 | 221.442 | [illegible] |
| 675752696 | PREDICTED: Golgi apparatus protein 1 [Callithrix jacchus] | 0 | 220.88 | [illegible] |
| 545561432 | semaphorin 7A [Callithrix jacchus] | 0 | 304.539 | [illegible] |
| 532340023 | serine/arginine-rich splicing factor 2 [Callithrix jacchus] | 0 | 26.012 | [illegible] |
| 532520286 | serine/arginine-rich splicing factor 3 [Callithrix jacchus] | 0 | 40.82 | [illegible] |
| 675691073 | PREDICTED: serine/arginine-rich splicing factor 5 isoform X1 [Callithrix jacchus] | 0 | 33.395 | [illegible] |
| 675688012 | PREDICTED: neuroendocrine protein 7B2 isoform X1 [Callithrix jacchus] | 0 | 10.713 | [illegible] |
| 1060946954 | PREDICTED: sphingosine-1-phosphate lyase 1 isoform X1 [Callithrix jacchus] | 0 | 20.344 | [illegible] |
| 675724067 | PREDICTED: LOW QUALITY PROTEIN: intersectin-2 [Callithrix jacchus] | 0 | 193.889 | [illegible] |
| 675637929 | PREDICTED: endophilin-A1 isoform X1 [Callithrix jacchus] | 0 | 122.801 | [illegible] |
| 296204201 | PREDICTED: endophilin-A3 isoform X1 [Callithrix jacchus] | 0 | 58.876 | [illegible] |
| 1060933941 | PREDICTED: SHC-transforming protein 3 isoform X1 [Callithrix jacchus] | 0 | 34.511 | [illegible] |
| 675725871 | PREDICTED: transformer-2 protein homolog beta isoform X1 [Callithrix jacchus] | 0 | 21.322 | [illegible] |
| 675697291 | PREDICTED: caseinolytic peptidase B protein homolog isoform X2 [Callithrix jacchus] | 0 | 94.099 | [illegible] |
| 296193820 | PREDICTED: solute carrier family 12 member 2 isoform X1 [Callithrix jacchus] | 0 | 141.82 | [illegible] |
| 1061017270 | PREDICTED: monocarboxylate transporter 2 isoform X1 [Callithrix jacchus] | 0 | 16.026 | [illegible] |
| 532275886 | excitatory amino acid transporter 2 isoform 1 [Callithrix jacchus] | 0 | 213.197 | [illegible] |
| 1060965913 | PREDICTED: excitatory amino acid transporter 1 [Callithrix jacchus] | 0 | 132.242 | [illegible] |
| 390463103 | PREDICTED: LOW QUALITY PROTEIN: band 3 anion transport protein [Callithrix jacchus] | 0 | 26.348 | [illegible] |
| 1060952386 | PREDICTED: sodium/calcium exchanger 1 isoform X1 [Callithrix jacchus] | 0 | 59.859 | [illegible] |
| 296207180 | PREDICTED: sodium/hydrogen exchanger 1 [Callithrix jacchus] | 0 | 42.787 | [illegible] |
| 296200179 | PREDICTED: synaptosomal-associated protein 25 isoform X1 [Callithrix jacchus] | 0 | 310.486 | [illegible] |
| 532348863 | clathrin coat assembly protein AP180 isoform a [Callithrix jacchus] | 0 | 254.346 | [illegible] |
| 296220175 | PREDICTED: gamma-synuclein isoform X4 [Callithrix jacchus] | 0 | 41.331 | [illegible] |
| 675775854 | PREDICTED: synaptosomal-associated protein 23 isoform X1 [Callithrix jacchus] | 0 | 40.19 | [illegible] |
| 1060950647 | PREDICTED: small nuclear ribonucleoprotein Sm D1 isoform X1 [Callithrix jacchus] | 0 | 18.119 | [illegible] |
| 532440329 | small nuclear ribonucleoprotein-associated protein N [Callithrix jacchus] | 0 | 8.642 | [illegible] |
| 23503524 | Cu,Zn-superoxide dismutase [Callithrix jacchus] | 0 | 36.618 | [illegible] |
| 532334195 | superoxide dismutase [Mn], mitochondrial isoform A precursor [Callithrix jacchus] | 0 | 33.291 | [illegible] |
| 675665961 | PREDICTED: glia-derived nexin [Callithrix jacchus] | 0 | 70.418 | [illegible] |
| 532292070 | spectrin alpha chain, non-erythrocytic 1 isoform 1 [Callithrix jacchus] | 0 | 3181.75 | [illegible] |
| 1060939885 | PREDICTED: spectrin beta chain, erythrocytic [Callithrix jacchus] | 0 | 565.949 | [illegible] |
| 1060952217 | PREDICTED: spectrin beta chain, non-erythrocytic 1 isoform X1 [Callithrix jacchus] | 0 | 2787.821 | [illegible] |
| 532290632 | spectrin beta chain, non-erythrocytic 2 [Callithrix jacchus] | 0 | 1275.397 | [illegible] |
| 390459232 | PREDICTED: testican-1 isoform X1 [Callithrix jacchus] | 0 | 25.994 | [illegible] |
| 1060992804 | PREDICTED: proto-oncogene tyrosine-protein kinase Src isoform X1 [Callithrix jacchus] | 0 | 181.842 | [illegible] |
| 675687851 | PREDICTED: signal recognition particle 14 kDa protein [Callithrix jacchus] | 0 | 22.57 | [illegible] |
| 532520200 | SRSF protein kinase 1 [Callithrix jacchus] | 0 | 46.623 | [illegible] |
| 1061014366 | PREDICTED: SRSF protein kinase 2 isoform X1 [Callithrix jacchus] | 0 | 101.355 | [illegible] |
| 390476349 | PREDICTED: signal recognition particle receptor subunit beta isoform X1 [Callithrix jacchus] | 0 | 32.509 | [illegible] |
| 1060959474 | PREDICTED: 60 kDa SS-A/Ro ribonucleoprotein isoform X1 [Callithrix jacchus] | 0 | 26.846 | [illegible] |
| 675775482 | PREDICTED: NHP2-like protein 1 isoform X4 [Callithrix jacchus] | 0 | 14.552 | [illegible] |
| 296206236 | PREDICTED: signal transducing adapter molecule 1 isoform X1 [Callithrix jacchus] | 0 | 17.395 | [illegible] |
| 675657056 | PREDICTED: double-stranded RNA-binding protein Staufen homolog 1 isoform X1 [Callithrix jacchus] | 0 | 63.114 | [illegible] |
| 675697490 | PREDICTED: stromal interaction molecule 1 [Callithrix jacchus] | 0 | 16.7 | [illegible] |
| 296218619 | PREDICTED: stress-induced-phosphoprotein 1 [Callithrix jacchus] | 0 | 126.652 | [illegible] |
| 296221154 | PREDICTED: STE20-like serine/threonine-protein kinase isoform X1 [Callithrix jacchus] | 0 | 59.851 | [illegible] |
| 296210875 | PREDICTED: serine-threonine kinase receptor-associated protein [Callithrix jacchus] | 0 | 75.718 | [illegible] |
| 296192233 | PREDICTED: syntaxin-1A isoform X1 [Callithrix jacchus] | 0 | 203.713 | [illegible] |
| 390471525 | PREDICTED: syntaxin-4 isoform X1 [Callithrix jacchus] | 0 | 11.699 | [illegible] |
| 296190870 | PREDICTED: syntaxin-binding protein 1 isoform X2 [Callithrix jacchus] | 0 | 458.302 | [illegible] |
| 296208700 | PREDICTED: syntaxin-binding protein 3 isoform X1 [Callithrix jacchus] | 0 | 23.793 | [illegible] |
| 532523116 | succinyl-CoA ligase [ADP-forming] subunit beta, mitochondrial precursor [Callithrix jacchus] | 0 | 44.448 | [illegible] |
| 296238204 | PREDICTED: eukaryotic translation initiation factor 1 [Callithrix jacchus] | 0 | 30.66 | [illegible] |
| 390479710 | PREDICTED: LOW QUALITY PROTEIN: synapsin-1 [Callithrix jacchus] | 0 | 665.938 | [illegible] |
| 1060955115 | PREDICTED: LOW QUALITY PROTEIN: synapsin-2 [Callithrix jacchus] | 0 | 302.924 | [illegible] |
| 296236925 | PREDICTED: synaptogyrin-1 [Callithrix jacchus] | 0 | 14.505 | [illegible] |
| 675705070 | PREDICTED: synaptogyrin-3 [Callithrix jacchus] | 0 | 24.94 | [illegible] |
| 1061020300 | PREDICTED: synaptophysin [Callithrix jacchus] | 0 | 21.244 | [illegible] |
| 390467977 | PREDICTED: synaptotagmin-1 isoform X1 [Callithrix jacchus] | 0 | 372.743 | [illegible] |
| 532521304 | synaptotagmin-2 [Callithrix jacchus] | 0 | 136.349 | [illegible] |
| 296234439 | PREDICTED: synaptotagmin-3 [Callithrix jacchus] | 0 | 44.695 | [illegible] |
| 532288306 | transaldolase [Callithrix jacchus] | 0 | 12.473 | [illegible] |
| 532521118 | contactin-2 precursor [Callithrix jacchus] | 0 | 162.308 | [illegible] |
| 675672433 | PREDICTED: LOW QUALITY PROTEIN: protein TBRG4 [Callithrix jacchus] | 0 | 7.728 | [illegible] |
| 675644606 | PREDICTED: S-phase kinase-associated protein 1 [Callithrix jacchus] | 0 | 102.389 | [illegible] |
| 1061016774 | PREDICTED: alpha-globin transcription factor CP2 isoform X1 [Callithrix jacchus] | 0 | 19.16 | [illegible] |
| 532344285 | nucleolar transcription factor 1 isoform b [Callithrix jacchus] | 0 | 51.664 | [illegible] |
| 296199555 | PREDICTED: T-complex protein 1 subunit alpha [Callithrix jacchus] | 0 | 269.259 | [illegible] |
| 296233039 | PREDICTED: peroxiredoxin-2 [Callithrix jacchus] | 0 | 105.793 | [illegible] |
| 532205695 | THO complex subunit 4 [Callithrix jacchus] | 0 | 51.76 | [illegible] |
| 532518045 | 26S proteasome non-ATPase regulatory subunit 2 [Callithrix jacchus] | 0 | 126.908 | [illegible] |
| 675652664 | PREDICTED: serine/threonine-protein phosphatase 2A 56 kDa regulatory subunit delta isoform [Callithrix jacchus] | 0 | 83.793 | [illegible] |
| 675709637 | PREDICTED: transcription factor A, mitochondrial [Callithrix jacchus] | 0 | 74.917 | [illegible] |
| 675732583 | PREDICTED: protein TFG isoform X1 [Callithrix jacchus] | 0.001 | 6.155 | [illegible] |
| 675694619 | PREDICTED: thy-1 membrane glycoprotein [Callithrix jacchus] | 0 | 97.688 | [illegible] |
| 1060969214 | PREDICTED: T-lymphoma invasion and metastasis-inducing protein 1 [Callithrix jacchus] | 0 | 57.905 | [illegible] |
| 532447689 | transcription intermediary factor 1-beta [Callithrix jacchus] | 0 | 63.99 | [illegible] |
| 675637604 | PREDICTED: tight junction protein ZO-2 isoform X1 [Callithrix jacchus] | 0 | 139.595 | [illegible] |
| 532515268 | transketolase isoform 1 [Callithrix jacchus] | 0 | 95.194 | [illegible] |
| 1060935144 | PREDICTED: tropomodulin-1 isoform X1 [Callithrix jacchus] | 0 | 69.843 | [illegible] |
| 390468059 | PREDICTED: thymopoietin isoform X2 [Callithrix jacchus] | 0 | 31.981 | [illegible] |
| 296190669 | PREDICTED: tenascin isoform X1 [Callithrix jacchus] | 0 | 475.041 | [illegible] |
| 1060959175 | PREDICTED: tenascin-R isoform X1 [Callithrix jacchus] | 0 | 475.835 | [illegible] |
| 296191752 | PREDICTED: target of Myb protein 1 [Callithrix jacchus] | 0 | 32.64 | [illegible] |
| 296200465 | PREDICTED: DNA topoisomerase 1 [Callithrix jacchus] | 0 | 18.061 | [illegible] |
| 675740399 | PREDICTED: DNA topoisomerase 2-beta isoform X1 [Callithrix jacchus] | 0 | 79.106 | [illegible] |
| 1060940711 | PREDICTED: apoptosis-stimulating of p53 protein 1 isoform X1 [Callithrix jacchus] | 0 | 40.443 | [illegible] |
| 532514560 | tumor protein D52 isoform 2 [Callithrix jacchus] | 0 | 59.88 | [illegible] |
| 675654138 | PREDICTED: tumor protein D53 isoform X3 [Callithrix jacchus] | 0 | 13.206 | [illegible] |
| 675677637 | PREDICTED: triosephosphate isomerase [Callithrix jacchus] | 0 | 141.106 | [illegible] |
| 532520228 | tripeptidyl-peptidase 2 [Callithrix jacchus] | 0 | 160.999 | [illegible] |
| 296212726 | PREDICTED: endoplasmin [Callithrix jacchus] | 0 | 95.549 | [illegible] |
| 1060940686 | PREDICTED: TNF receptor-associated factor 3 isoform X1 [Callithrix jacchus] | 0 | 47.932 | [illegible] |
| 532444707 | transferrin receptor protein 1 [Callithrix jacchus] | 0 | 22.79 | [illegible] |
| 1061001251 | PREDICTED: translationally-controlled tumor protein isoform X1 [Callithrix jacchus] | 0 | 25.432 | [illegible] |
| 1060945490 | PREDICTED: tuberin isoform X1 [Callithrix jacchus] | 0 | 157.825 | [illegible] |
| 675698965 | PREDICTED: tumor susceptibility gene 101 protein [Callithrix jacchus] | 0 | 43.525 | [illegible] |
| 296234343 | PREDICTED: 60S ribosomal protein L13a isoform X1 [Callithrix jacchus] | 0 | 40.186 | [illegible] |
| 532452637 | 26S proteasome non-ATPase regulatory subunit 3 [Callithrix jacchus] | 0 | 70.19 | [illegible] |
| 296205655 | PREDICTED: tubulin alpha-4A chain [Callithrix jacchus] | 0 | 590.38 | [illegible] |
| 1060972304 | PREDICTED: tubulin beta-4A chain [Callithrix jacchus] | 0 | 649.471 | [illegible] |
| 296190552 | PREDICTED: thioredoxin [Callithrix jacchus] | 0 | 23.186 | [illegible] |
| 675704634 | PREDICTED: SUMO-conjugating enzyme UBC9 [Callithrix jacchus] | 0 | 9.017 | [illegible] |
| 1061018940 | PREDICTED: ubiquitin-like modifier-activating enzyme 1 [Callithrix jacchus] | 0 | 107.093 | [illegible] |
| 296203592 | PREDICTED: ubiquitin carboxyl-terminal hydrolase 12 isoform X1 [Callithrix jacchus] | 0 | 44.108 | [illegible] |
| 296196670 | PREDICTED: ubiquitin carboxyl-terminal hydrolase isozyme L1 [Callithrix jacchus] | 0 | 75.809 | [illegible] |
| 532507048 | ubiquitin carboxyl-terminal hydrolase 10 isoform 2 [Callithrix jacchus] | 0 | 47.505 | [illegible] |
| 532519798 | ubiquitin carboxyl-terminal hydrolase 5 isoform 1 [Callithrix jacchus] | 0 | 148.846 | [illegible] |
| 675648041 | PREDICTED: 2-hydroxyacylsphingosine 1-beta-galactosyltransferase isoform X1 [Callithrix jacchus] | 0 | 28.814 | [illegible] |
| 296193157 | PREDICTED: dihydropyrimidinase-related protein 3 isoform X1 [Callithrix jacchus] | 0 | 199.814 | [illegible] |
| 675638345 | PREDICTED: protein unc-13 homolog B isoform X9 [Callithrix jacchus] | 0 | 115.279 | [illegible] |
| 675644655 | PREDICTED: cytochrome b-c1 complex subunit 8 [Callithrix jacchus] | 0 | 60.342 | [illegible] |
| 1060954212 | PREDICTED: cytochrome b-c1 complex subunit 1, mitochondrial isoform X1 [Callithrix jacchus] | 0 | 255.362 | [illegible] |
| 1061019526 | PREDICTED: probable ubiquitin carboxyl-terminal hydrolase FAF-X isoform X1 [Callithrix jacchus] | 0 | 179.86 | [illegible] |
| 296201313 | PREDICTED: vesicle-associated membrane protein 2 isoform X2 [Callithrix jacchus] | 0 | 99.913 | [illegible] |
| 532286538 | valine--tRNA ligase [Callithrix jacchus] | 0 | 66.569 | [illegible] |
| 532443671 | vasodilator-stimulated phosphoprotein [Callithrix jacchus] | 0 | 42.824 | [illegible] |
| 532522126 | guanine nucleotide exchange factor VAV2 isoform 1 [Callithrix jacchus] | 0 | 63.953 | [illegible] |
| 675708582 | PREDICTED: vinculin isoform X1 [Callithrix jacchus] | 0 | 101.535 | [illegible] |
| 296220295 | PREDICTED: voltage-dependent anion-selective channel protein 2 isoform X1 [Callithrix jacchus] | 0 | 257.464 | [illegible] |
| 1060950304 | PREDICTED: voltage-dependent anion-selective channel protein 3 isoform X1 [Callithrix jacchus] | 0 | 248.655 | [illegible] |
| 532503521 | protein lin-7 homolog B [Callithrix jacchus] | 0 | 31.345 | [illegible] |
| 296198490 | PREDICTED: protein lin-7 homolog C [Callithrix jacchus] | 0 | 63.741 | [illegible] |
| 296200450 | PREDICTED: vesicular inhibitory amino acid transporter [Callithrix jacchus] | 0 | 41.362 | [illegible] |
| 532448065 | ezrin [Callithrix jacchus] | 0 | 134.502 | [illegible] |
| 675742333 | PREDICTED: vacuolar protein sorting-associated protein 45 isoform X1 [Callithrix jacchus] | 0 | 56.212 | [illegible] |
| 1060940597 | PREDICTED: tryptophan--tRNA ligase, cytoplasmic [Callithrix jacchus] | 0 | 21.623 | [illegible] |
| 296236993 | PREDICTED: eukaryotic translation initiation factor 4H [Callithrix jacchus] | 0 | 16.489 | [illegible] |
| 675650188 | PREDICTED: WD repeat-containing protein 1 [Callithrix jacchus] | 0 | 171.532 | [illegible] |
| 296196929 | PREDICTED: wolframin [Callithrix jacchus] | 0 | 38.175 | [illegible] |
| 296221926 | PREDICTED: dynactin subunit 6 isoform X2 [Callithrix jacchus] | 0 | 25.476 | [illegible] |
| 1061006340 | PREDICTED: X-ray repair cross-complementing protein 5 [Callithrix jacchus] | 0 | 112.9 | [illegible] |
| 532507749 | nuclease-sensitive element-binding protein 1 [Callithrix jacchus] | 0 | 100.735 | [illegible] |
| 296201052 | PREDICTED: 14-3-3 protein epsilon isoform X1 [Callithrix jacchus] | 0 | 239.464 | [illegible] |
| 296192280 | PREDICTED: 14-3-3 protein gamma [Callithrix jacchus] | 0 | 172.252 | [illegible] |
| 296191727 | PREDICTED: 14-3-3 protein eta [Callithrix jacchus] | 0 | 138.98 | [illegible] |
| 675724732 | PREDICTED: 14-3-3 protein theta [Callithrix jacchus] | 0 | 183.27 | [illegible] |
| 1060957442 | PREDICTED: 14-3-3 protein zeta/delta [Callithrix jacchus] | 0 | 170.675 | [illegible] |
| 675659591 | PREDICTED: BUB3-interacting and GLEBS motif-containing protein ZNF207 isoform X1 [Callithrix jacchus] | 0 | 8.482 | [illegible] |
| 390474658 | PREDICTED: zinc transporter 3 isoform X3 [Callithrix jacchus] | 0 | 19.285 | [illegible] |
| 532501852 | coronin-1B [Callithrix jacchus] | 0 | 96.161 | [illegible] |
| 296212827 | PREDICTED: coronin-1C [Callithrix jacchus] | 0 | 136.833 | [illegible] |
| 675665338 | PREDICTED: disintegrin and metalloproteinase domain-containing protein 23 isoform X1 [Callithrix jacchus] | 0 | 124.237 | [illegible] |
| 1060941136 | PREDICTED: barrier-to-autointegration factor isoform X1 [Callithrix jacchus] | 0 | 46.217 | [illegible] |
| 296205707 | PREDICTED: phenylalanine--tRNA ligase beta subunit isoform X1 [Callithrix jacchus] | 0 | 94.748 | [illegible] |

| | | | | | | | | | | | | | | | | | | | |
|---|---|---|---|---|---|---|---|---|---|---|---|---|---|---|---|---|---|---|---|
| 296201373 | PREDICTED: fragile X mental retardation syndrome-related protein 2 isoform X1 [Callithrix jacchus] | 0 | 72.205 | 25 | 12 | 63 | 9 | 1 | 673 | 74.1 | 6.23 | 892 | 12 | 17969820.55 | 18155555.69 | 16397073.53 | 20721703.89 | 18867619.85 | 20770662.57 |
| 675649043 | PREDICTED: ras GTPase-activating protein-binding protein 2 [Callithrix jacchus] | 0 | 134.929 | 50 | 19 | 230 | 18 | 1 | 482 | 54.1 | 5.55 | 2252 | 19 | 224444069.3 | 223999846.9 | 200183609 | 184346537.3 | 175603742.7 | 187157544 |
| 296188827 | PREDICTED: glypican-6 isoform X1 [Callithrix jacchus] | 0 | 32.927 | 15 | 6 | 21 | 5 | 1 | 555 | 62.6 | 5.41 | 237 | 6 | 9017325.56 | 9346894.625 | 7493156.018 | 10989669.1 | 11309337.15 | 8988124.337 |
| 296228983 | PREDICTED: HCLS1-associated protein X-1 isoform X1 [Callithrix jacchus] | 0 | 36.293 | 33 | 7 | 39 | 7 | 1 | 283 | 32 | 5.05 | 378 | 7 | 5553001.026 | 4409238.469 | 4474343.044 | 11465868.83 | 11726609.87 | 10632596.29 |
| 675735786 | PREDICTED: ly-6/neurotoxin-like protein 1 [Callithrix jacchus] | 0 | 15.421 | 30 | 3 | 39 | 3 | 1 | 116 | 12.5 | 7.5 | 1265 | 3 | 135486570.4 | 158564897.8 | 121596217.5 | 161112267.2 | 138394630.6 | 142307343.8 |
| 296212001 | PREDICTED: extended synaptotagmin-1 [Callithrix jacchus] | 0 | 53.469 | 15 | 12 | 24 | 12 | 1 | 1104 | 122.7 | 5.77 | 202 | 12 | 8328039.948 | 8088672.813 | 7811246.504 | 8788356.349 | 8623251.702 | 6732197.377 |
| 532522796 | monoglyceride lipase isoform 1 [Callithrix jacchus] | 0 | 12.643 | 10 | 3 | 11 | 3 | 1 | 313 | 34.4 | 6.65 | 47 | 3 | 8928994.599 | 8717077.5 | 4502388.164 | 6919505.47 | 6684385.809 | 6009855.676 |
| 1061014603 | PREDICTED: dnaJ homolog subfamily B member 6 isoform X1 [Callithrix jacchus] | 0 | 50.584 | 19 | 8 | 118 | 7 | 1 | 336 | 36.8 | 6.67 | 571 | 8 | 23519620.11 | 25006804.28 | 32575800.89 | 13461014.09 | 21576166.03 | 27813785.06 |
| 390461864 | PREDICTED: LOW QUALITY PROTEIN: 5'-nucleotidase [Callithrix jacchus] | 0 | 95.17 | 35 | 16 | 101 | 16 | 1 | 574 | 63.5 | 7.18 | 1215 | 16 | 73580039.48 | 65958462.81 | 68758481.68 | 81192014.7 | 105346193.8 | 118869617.1 |
| 1060978080 | PREDICTED: teneurin-3 isoform X1 [Callithrix jacchus] | 0 | 48.484 | 5 | 15 | 43 | 14 | 1 | 2721 | 303.5 | 6.57 | 151 | 15 | 23055421.16 | 25480249.63 | 22329412.03 | 21862169.63 | 22164360.85 | 19976758.3 |
| 296197946 | PREDICTED: protein kinase C and casein kinase substrate in neurons protein 1 [Callithrix jacchus] | 0 | 290.524 | 67 | 32 | 397 | 30 | 1 | 444 | 50.9 | 5.24 | 7047 | 32 | 769998958.5 | 739611716.5 | 714536097 | 772284254.8 | 781181844.9 | 869806418.3 |
| 1060937451 | REDICTED: protein kinase C and casein kinase substrate in neurons protein 2 isoform X1 [Callithrix jacchus | 0 | 38.781 | 18 | 11 | 47 | 9 | 1 | 553 | 63.6 | 5.58 | 126 | 11 | 46969916.83 | 41002932.25 | 37168362.61 | 19490809.62 | 16553723.17 | 25701205.87 |
| 1061018287 | PREDICTED: phosphatidylethanolamine-binding protein 1 [Callithrix jacchus] | 0 | 92.028 | 79 | 10 | 59 | 10 | 1 | 187 | 21 | 7.99 | 1058 | 10 | 35026281.82 | 34059758.31 | 31839558.74 | 50823448.51 | 39218831.66 | 48569142.32 |
| 296223628 | PREDICTED: poly(rC)-binding protein 1 [Callithrix jacchus] | 0 | 89.776 | 48 | 12 | 137 | 7 | 1 | 356 | 37.5 | 7.09 | 1746 | 12 | 46542049.89 | 44994191.5 | 54402213.08 | 55451706.2 | 48464588.37 | 43042800.23 |
| 1060974389 | PREDICTED: peptidyl-prolyl cis-trans isomerase NIMA-interacting 1 [Callithrix jacchus] | 0 | 37.652 | 50 | 5 | 25 | 5 | 1 | 163 | 18.2 | 8.82 | 497 | 5 | 14901545.55 | 17475309.13 | 10205947.53 | 8310305.23 | 11473137.42 | 5546946.308 |
| 675663625 | REDICTED: interferon-inducible double-stranded RNA-dependent protein kinase activator A [Callithrix jacchu | 0 | 125.221 | 58 | 17 | 127 | 17 | 1 | 313 | 34.4 | 8.41 | 1673 | 17 | 185616239.1 | 185303668.9 | 199827106.2 | 176047758.3 | 182183435.1 | 178148612 |
| 296233807 | PREDICTED: 26S protease regulatory subunit 6B [Callithrix jacchus] | 0 | 115.705 | 58 | 18 | 65 | 18 | 1 | 418 | 47.4 | 5.21 | 1004 | 18 | 33165939.16 | 30235562.69 | 36531247.61 | 39740385.28 | 39871213.58 | 35465997.67 |
| 296219151 | PREDICTED: 26S proteasome non-ATPase regulatory subunit 13 isoform X2 [Callithrix jacchus] | 0 | 50.462 | 33 | 11 | 45 | 11 | 1 | 376 | 42.9 | 5.81 | 225 | 11 | 18642891.89 | 14936183.63 | 17352756.29 | 23844129.14 | 22897561.6 | 21587039.56 |
| 675750307 | PREDICTED: regulator of G-protein signaling 7 isoform X1 [Callithrix jacchus] | 0 | 141.68 | 40 | 20 | 114 | 17 | 1 | 494 | 57.6 | 7.72 | 1521 | 20 | 156193989.7 | 166694953.4 | 147153978.7 | 253807227.5 | 206269592.5 | 205742828.7 |
| 296195476 | PREDICTED: ATP-binding cassette sub-family E member 1 [Callithrix jacchus] | 0 | 15.625 | 8 | 4 | 11 | 4 | 1 | 599 | 67.3 | 8.34 | 28 | 4 | 3275442.197 | 5068329 | 4248298.333 | 3778056.276 | 4984266.707 | 4615590.587 |
| 1060931802 | PREDICTED: neuronal-specific septin-3 [Callithrix jacchus] | 0 | 199.103 | 59 | 22 | 338 | 22 | 1 | 358 | 40.7 | 7.18 | 4340 | 22 | 1064537740 | 1034747333 | 1135711795 | 1036806141 | 1044018848 | 1028125679 |
| 675689515 | PREDICTED: signal recognition particle 54 kDa protein isoform X1 [Callithrix jacchus] | 0 | 39.698 | 20 | 9 | 49 | 9 | 1 | 504 | 55.7 | 8.75 | 398 | 9 | 12569307.9 | 13448274.5 | 13689862.82 | 16088487.17 | 16596433.16 | 14778296.86 |
| 296193001 | PREDICTED: transcription initiation factor TFIID subunit 7 [Callithrix jacchus] | 0 | 9.499 | 7 | 2 | 7 | 2 | 1 | 349 | 40.2 | 5.34 | 95 | 2 | 1639534.194 | 1754707.313 | 1648533.175 | 666065.4269 | 584084.0389 | 1068896.785 |
| 532522194 | chloride transport protein 6 isoform 1 [Callithrix jacchus] | 0 | 50.799 | 16 | 12 | 58 | 12 | 1 | 870 | 97.3 | 6.9 | 707 | 12 | 30436470.23 | 24858632.63 | 27859675.27 | 26884900.75 | 23443515.66 | 18779375.78 |
| 296219202 | PREDICTED: peroxisomal 2,4-dienoyl-CoA reductase [Callithrix jacchus] | 0 | 27.844 | 26 | 7 | 26 | 7 | 1 | 292 | 30.8 | 8.81 | 102 | 7 | 21181279.61 | 15844619.72 | 24587733.08 | 40135347.96 | 22289603.41 | 19137929.42 |
| 532444827 | G protein-coupled receptor kinase 6 isoform B [Callithrix jacchus] | 0 | 20.097 | 7 | 3 | 7 | 3 | 1 | 589 | 67.4 | 8.37 | 150 | 3 | 5057364.095 | 6930188.938 | 6509326.083 | 3321993.937 | 6756017.087 | 1702920.297 |
| 296213521 | PREDICTED: dual specificity mitogen-activated protein kinase kinase 1 isoform X1 [Callithrix jacchus] | 0 | 79.744 | 37 | 14 | 84 | 9 | 1 | 393 | 43.4 | 6.62 | 716 | 14 | 39143789.67 | 37276006.13 | 35741550.39 | 31910891.38 | 34076056.11 | 42490543.26 |
| 675658057 | PREDICTED: dual specificity mitogen-activated protein kinase kinase 4 isoform X1 [Callithrix jacchus] | 0 | 17.044 | 12 | 3 | 7 | 3 | 1 | 410 | 45.6 | 8.03 | 66 | 3 | 2239133.491 | 4076203.75 | 6095442.581 | 3165821.924 | 4235060.118 | 2476906.911 |
| 532281698 | dual specificity mitogen-activated protein kinase kinase 7 [Callithrix jacchus] | 0 | 24.55 | 16 | 6 | 20 | 6 | 1 | 424 | 47.9 | 9.09 | 78 | 6 | 13151910.68 | 11148252.66 | 12847509.7 | 3348598.101 | 5613961.471 | 2893135.6 |
| 296199326 | PREDICTED: mitogen-activated protein kinase kinase kinase 5 [Callithrix jacchus] | 0 | 83.415 | 18 | 23 | 68 | 22 | 1 | 1373 | 154.3 | 5.85 | 314 | 23 | 67715152.3 | 64419524.13 | 61028139.41 | 47165533.82 | 38288735.86 | 55259846.38 |
| 1060936713 | PREDICTED: mitogen-activated protein kinase 1 [Callithrix jacchus] | 0 | 142.998 | 63 | 21 | 161 | 15 | 1 | 360 | 41.4 | 6.98 | 926 | 21 | 106205829.7 | 120050746.2 | 128706092.9 | 89907395.08 | 93243076.59 | 103983844.8 |
| 1060980669 | PREDICTED: mitogen-activated protein kinase 10 isoform X1 [Callithrix jacchus] | 0 | 43.738 | 27 | 11 | 44 | 6 | 1 | 464 | 52.5 | 6.74 | 352 | 11 | 30501565.6 | 29404113.31 | 29189873.35 | 28423226.63 | 28263774.5 | 25788180.16 |
| 296203704 | PREDICTED: neurobeachin isoform X4 [Callithrix jacchus] | 0 | 133.175 | 13 | 32 | 124 | 29 | 1 | 2946 | 327.5 | 6.19 | 1460 | 32 | 62807146.82 | 67917263.06 | 69535846.71 | 54995139.41 | 41092718.25 | 57948909.74 |
| 296217633 | PREDICTED: proteasome subunit alpha type-1 isoform X1 [Callithrix jacchus] | 0 | 64.533 | 49 | 13 | 49 | 13 | 1 | 263 | 29.5 | 6.61 | 573 | 13 | 39758996.45 | 37044469.31 | 38937270.34 | 53248649.66 | 55696361.16 | 61397177.22 |
| 1060959841 | PREDICTED: proteasome subunit alpha type-4 isoform X1 [Callithrix jacchus] | 0 | 22.496 | 25 | 6 | 29 | 6 | 1 | 261 | 29.5 | 7.72 | 205 | 6 | 26451210.24 | 18560260 | 27127788.99 | 47877142.48 | 43101469.7 | 38899873.39 |
| 296208738 | PREDICTED: proteasome subunit alpha type-5 isoform X1 [Callithrix jacchus] | 0 | 47.242 | 43 | 7 | 53 | 7 | 1 | 241 | 26.4 | 4.79 | 631 | 7 | 21487314.53 | 21333208.13 | 22717097.92 | 30355433.79 | 28402766.79 | 23542131.2 |
| 532448223 | proteasome subunit alpha type-6 [Callithrix jacchus] | 0 | 63.324 | 48 | 10 | 66 | 10 | 1 | 246 | 27.4 | 6.76 | 849 | 10 | 94099357.73 | 95474271.78 | 92428040.35 | 126107101.1 | 121033482.7 | 101105572.7 |
| 675657369 | PREDICTED: proteasome subunit alpha type-7 [Callithrix jacchus] | 0 | 58.524 | 44 | 9 | 52 | 9 | 1 | 248 | 27.9 | 8.46 | 899 | 9 | 26985685.79 | 30011627.02 | 26494713.4 | 42300346.66 | 40745578.37 | 37888701.06 |
| 532502369 | proteasome subunit beta type-3 [Callithrix jacchus] | 0 | 16.141 | 19 | 3 | 16 | 3 | 1 | 205 | 22.9 | 6.55 | 134 | 3 | 10560927.3 | 8300179 | 10041494.51 | 17746722.08 | 15497586.34 | 12040893.6 |
| 296217495 | PREDICTED: 60S ribosomal protein L27a [Callithrix jacchus] | 0 | 40.463 | 32 | 7 | 65 | 7 | 1 | 148 | 16.6 | 11.12 | 645 | 7 | 170670753.2 | 182988791.2 | 182911119.4 | 162042828.9 | 174464257.7 | 163663821.3 |
| 296205731 | PREDICTED: cullin-3 [Callithrix jacchus] | 0 | 37.437 | 14 | 10 | 19 | 9 | 1 | 768 | 88.9 | 8.48 | 100 | 10 | 7006301.09 | 4175035.375 | 5083567.952 | 8327695.789 | 7383492.788 | 8475553.621 |
| 296194253 | PREDICTED: homer protein homolog 1 [Callithrix jacchus] | 0 | 286.284 | 76 | 27 | 474 | 24 | 1 | 354 | 40.3 | 5.38 | 9385 | 27 | 1278312313 | 1368680376 | 1235796506 | 1376357516 | 1346915307 | 1305149681 |
| 675663254 | PREDICTED: homer protein homolog 2 isoform X1 [Callithrix jacchus] | 0 | 145.847 | 54 | 23 | 166 | 20 | 1 | 354 | 40.6 | 6.01 | 2156 | 23 | 87087885.18 | 93406754.21 | 89126573.08 | 110514588.9 | 107560230.6 | 105934289.9 |
| 1061010895 | PREDICTED: neurochondrin isoform X1 [Callithrix jacchus] | 0 | 147.287 | 40 | 22 | 143 | 22 | 1 | 712 | 77.2 | 5.54 | 2244 | 22 | 94957810.78 | 88773707.59 | 104524492.2 | 107632055.3 | 91488113.42 | 108651101.9 |
| 675639559 | PREDICTED: long-chain fatty acid transport protein 4 [Callithrix jacchus] | 0 | 24.284 | 11 | 5 | 12 | 4 | 1 | 643 | 72.1 | 8.47 | 94 | 5 | 2032069.093 | 4185276.5 | 1758352.491 | 621816.336 | 2525344.572 | 2019339.803 |
| 296237129 | PREDICTED: COP9 signalosome complex subunit 3 isoform X1 [Callithrix jacchus] | 0 | 35.155 | 18 | 6 | 40 | 6 | 1 | 423 | 47.8 | 6.65 | 719 | 6 | 9564282.691 | 8282948 | 9508217.878 | 7962101.33 | 10587440.83 | 12069291.22 |
| 390480433 | PREDICTED: reticulocalbin-2 isoform X1 [Callithrix jacchus] | 0 | 76.077 | 46 | 10 | 59 | 10 | 1 | 317 | 36.9 | 4.41 | 1623 | 10 | 17469529.88 | 19575480.41 | 13949491.63 | 31332715.13 | 24515545.92 | 19070165.56 |
| 1060956305 | PREDICTED: COP9 signalosome complex subunit 5 isoform X1 [Callithrix jacchus] | 0 | 31.95 | 17 | 6 | 26 | 6 | 1 | 390 | 43.4 | 7.66 | 242 | 6 | 17269133.85 | 14999613.81 | 16255921.97 | 23791062.82 | 22944645.86 | 18650844.48 |
| 1060948424 | PREDICTED: dihydropyrimidinase-related protein 4 [Callithrix jacchus] | 0 | 133.511 | 55 | 20 | 101 | 20 | 1 | 572 | 61.9 | 6.98 | 1254 | 20 | 60993997.69 | 57203231.92 | 76105711.48 | 91817283.05 | 59554039.58 | 55830024.08 |
| 390466812 | PREDICTED: LOW QUALITY PROTEIN: protein piccolo [Callithrix jacchus] | 0 | 1876.167 | 49 | 198 | 2682 | 193 | 1 | 4952 | 540.2 | 6.47 | 45849 | 198 | 2443941791 | 2448527794 | 2446465470 | 2201740406 | 2293496803 | 2569749451 |
| 675648765 | PREDICTED: COP9 signalosome complex subunit 4 [Callithrix jacchus] | 0 | 27.393 | 19 | 7 | 29 | 7 | 1 | 406 | 46.2 | 5.83 | 304 | 7 | 10619164.17 | 9916617.125 | 13037183.99 | 12389897.07 | 10584479.64 | 13213097.69 |
| 296192431 | PREDICTED: COP9 signalosome complex subunit 6 [Callithrix jacchus] | 0 | 23.515 | 20 | 5 | 24 | 5 | 1 | 324 | 35.9 | 5.73 | 32 | 5 | 6727848.644 | 6615498.516 | 7111120.088 | 15038055.61 | 8223063.853 | 9749877.859 |
| 296235140 | PREDICTED: eukaryotic translation initiation factor 2 subunit 3 isoform X1 [Callithrix jacchus] | 0 | 83.179 | 43 | 15 | 71 | 15 | 1 | 472 | 51.1 | 8.4 | 972 | 15 | 51459222.5 | 56192217 | 48188308.53 | 38666021.54 | 47485088.93 | 54506268.87 |
| 532515392 | G protein-regulated inducer of neurite outgrowth 1 [Callithrix jacchus] | 0 | 407.182 | 65 | 59 | 824 | 59 | 1 | 1012 | 103.6 | 6.27 | 10536 | 59 | 996477613.5 | 1036330474 | 1094188626 | 956783994.8 | 1038474644 | 1154301273 |
| 532512404 | apoptosis-inducing factor 1, mitochondrial isoform 1 precursor [Callithrix jacchus] | 0 | 100.917 | 38 | 18 | 126 | 18 | 1 | 612 | 66.8 | 9.06 | 2139 | 18 | 46033952.13 | 53226568.64 | 55383110.86 | 60432429 | 48708141.15 | 57324175.01 |
| 532329543 | erine/threonine-protein phosphatase 2A 56 kDa regulatory subunit gamma isoform isoform f [Callithrix jacchu | 0 | 100.721 | 29 | 14 | 104 | 8 | 1 | 540 | 62.8 | 8.7 | 1847 | 14 | 29952864.41 | 29770422.78 | 27830510.08 | 33650332.28 | 30926800.83 | 29647699.13 |
| 675690569 | EDICTED: serine/threonine-protein phosphatase 2A 56 kDa regulatory subunit epsilon isoform [Callithrix jacc | 0 | 107.795 | 42 | 16 | 130 | 13 | 1 | 467 | 54.7 | 6.95 | 1886 | 16 | 72389579.01 | 67495152.58 | 81013338.98 | 84926487.03 | 77505408.77 | 66753343.02 |
| 675772390 | PREDICTED: myosin phosphatase Rho-interacting protein isoform X1 [Callithrix jacchus] | 0 | 297.896 | 57 | 49 | 371 | 48 | 1 | 1034 | 117.7 | 6.33 | 5061 | 49 | 230738967.5 | 233382236 | 221495562 | 206648767.2 | 230796281.8 | 216442010 |
| 675661189 | PREDICTED: Na(+)/H(+) exchange regulatory cofactor NHE-RF1 isoform X1 [Callithrix jacchus] | 0 | 26.799 | 16 | 5 | 27 | 5 | 1 | 375 | 40.5 | 5.63 | 533 | 5 | 4597153.982 | 4708878.75 | 5805972.483 | 5618439.506 | 5191679.723 | 5073057.212 |
| 390462624 | PREDICTED: serine incorporator 3 [Callithrix jacchus] | 0 | 7.827 | 2 | 2 | 16 | 2 | 1 | 473 | 52.6 | 7.65 | 90 | 2 | 2743158.38 | 3604066.25 | 3918460.242 | 4781846.296 | 4333515.106 | 4169540.619 |
| 296201522 | PREDICTED: synaptic vesicle membrane protein VAT-1 homolog isoform X1 [Callithrix jacchus] | 0 | 16.343 | 14 | 5 | 16 | 5 | 1 | 401 | 42.7 | 6.55 | 161 | 5 | 4453856.751 | 3638260.063 | 5478447.565 | 4977615.033 | 5542300.066 | 5766278.425 |
| 675724464 | PREDICTED: visinin-like protein 1 [Callithrix jacchus] | 0 | 92.348 | 76 | 16 | 201 | 12 | 1 | 191 | 22.1 | 5.15 | 3683 | 16 | 275914332.6 | 279278561.6 | 267263736.5 | 328054057.1 | 311569699 | 340113945.3 |
| 296227008 | PREDICTED: 60S ribosomal protein L8 isoform X2 [Callithrix jacchus] | 0 | 93.909 | 48 | 12 | 157 | 12 | 1 | 257 | 28 | 11.03 | 2597 | 12 | 152958945.3 | 131230338.9 | 165989878.8 | 161090953.1 | 173154961.3 | 235832406.5 |
| 1060963638 | PREDICTED: ras GTPase-activating protein-binding protein 1 isoform X1 [Callithrix jacchus] | 0 | 20.933 | 10 | 4 | 38 | 3 | 1 | 466 | 52.1 | 5.52 | 535 | 4 | 1555523.342 | 5114724 | 6202512.845 | 5003235.151 | 4597919.964 | 4214558.304 |
| 390470016 | PREDICTED: 40S ribosomal protein S3 [Callithrix jacchus] | 0 | 108.567 | 80 | 20 | 274 | 20 | 1 | 243 | 26.7 | 9.66 | 2875 | 20 | 470131846.9 | 406327559.7 | 456816820.9 | 488263233 | 439931453.2 | 441887581.1 |
| 1061014026 | PREDICTED: asparagine synthetase [glutamine-hydrolyzing] [Callithrix jacchus] | 0 | 53.755 | 23 | 11 | 39 | 11 | 1 | 561 | 64.3 | 6.86 | 294 | 11 | 29470983.02 | 28243954.38 | 26158020.79 | 2629311.01 | 1971125.055 | 2824325.876 |
| 296230303 | PREDICTED: signal recognition particle 9 kDa protein [Callithrix jacchus] | 0 | 19.188 | 45 | 4 | 41 | 4 | 1 | 86 | 10.2 | 7.97 | 751 | 4 | 47948107.62 | 47872877 | 54095456.66 | 41701511.36 | 38169613.38 | 45476504.99 |
| 675770870 | PREDICTED: B-cell receptor-associated protein 31 [Callithrix jacchus] | 0 | 20.172 | 20 | 6 | 29 | 6 | 1 | 246 | 27.9 | 8.44 | 170 | 6 | 8553129.332 | 9414117.125 | 9834065.874 | 5275436.096 | 6618750.013 | 7341804.241 |
| 675641216 | PREDICTED: synapsin-3 isoform X1 [Callithrix jacchus] | 0 | 189.731 | 55 | 22 | 190 | 21 | 1 | 580 | 63.2 | 9.38 | 2717 | 22 | 147348838.8 | 140402328.7 | 159638101.8 | 118036263 | 118115712.9 | 115979728.1 |
| 532450671 | sodium/potassium-transporting ATPase subunit alpha-2 proprotein [Callithrix jacchus] | 0 | 536.42 | 58 | 59 | 890 | 37 | 1 | 1020 | 112.2 | 5.66 | 17891 | 59 | 528416213 | 546617902.4 | 534737583.4 | 519026258 | 521728601.8 | 557691109.4 |
| 532522060 | cysteine--tRNA ligase, cytoplasmic isoform c [Callithrix jacchus] | 0 | 21.117 | 8 | 7 | 21 | 7 | 1 | 831 | 94.7 | 6.86 | 28 | 7 | 10616140.85 | 10874263.44 | 10608946.43 | 13126500.39 | 13587793.01 | 15223731.68 |
| 1060942454 | DICTED: LOW QUALITY PROTEIN: pleckstrin homology domain-containing family B member 1 [Callithrix jac | 0 | 13.374 | 17 | 3 | 10 | 3 | 1 | 224 | 25.3 | 8.54 | 50 | 3 | 1289761.453 | 1675382.625 | 1772336.702 | 2143221.005 | 1705469.236 | 2194599.814 |
| 1060948611 | PREDICTED: paladin [Callithrix jacchus] | 0 | 9.714 | 4 | 3 | 6 | 3 | 1 | 900 | 101.4 | 7.68 | 0 | 3 | 6604382.347 | 6753396.25 | 5050576.966 | 2667438.003 | 2434800.48 | 3946592.345 |
| 296236921 | PREDICTED: 60S ribosomal protein L3 [Callithrix jacchus] | 0 | 208.624 | 54 | 22 | 299 | 22 | 1 | 403 | 46.1 | 10.18 | 3977 | 22 | 428275104.7 | 405256832.6 | 428219122.8 | 324263147.5 | 391982202.1 | 481582973.2 |
| 532438009 | transducin beta-like protein 2 precursor [Callithrix jacchus] | 0 | 16.41 | 12 | 5 | 9 | 5 | 1 | 446 | 49.6 | 9.2 | 32 | 5 | 6551838.628 | 6044746.625 | 5676931.224 | 5605522.268 | 6375170.087 | 8097890.861 |
| 296196056 | PREDICTED: 40S ribosomal protein S26 [Callithrix jacchus] | 0 | 23.48 | 50 | 5 | 52 | 5 | 1 | 115 | 13 | 11 | 442 | 5 | 44480079.29 | 56586966.13 | 69426067.58 | 71536762.54 | 83787679.12 | 57228519.95 |
| 532282756 | casein kinase I isoform epsilon [Callithrix jacchus] | 0 | 104.338 | 46 | 18 | 141 | 7 | 1 | 416 | 47.3 | 9.61 | 1670 | 18 | 52327640.86 | 52505820.53 | 54566673.77 | 26722482.11 | 20972192.41 | 22153408.75 |
| 675688692 | PREDICTED: protein arginine N-methyltransferase 5 isoform X1 [Callithrix jacchus] | 0 | 25.344 | 12 | 7 | 20 | 7 | 1 | 659 | 75.2 | 6.14 | 31 | 7 | 12134479.99 | 11321513.97 | 7558641.883 | 14289045.27 | 13899215.34 | 14850191.36 |
| 296206357 | PREDICTED: ATP-dependent zinc metalloprotease YME1L1 [Callithrix jacchus] | 0 | 12.835 | 6 | 4 | 11 | 4 | 1 | 716 | 79.7 | 8.66 | 74 | 4 | 62349610.73 | 70070113.63 | 74287940.86 | 34909964.02 | 36214634.95 | 60954814.38 |
| 296231948 | PREDICTED: 39S ribosomal protein L39, mitochondrial isoform X1 [Callithrix jacchus] | 0 | 38.541 | 27 | 8 | 40 | 8 | 1 | 338 | 38.7 | 7.71 | 553 | 8 | 11641381.27 | 11089538.63 | 12449010.65 | 17093379.37 | 16496037.22 | 12905267.56 |
| 296226481 | PREDICTED: 39S ribosomal protein L15, mitochondrial [Callithrix jacchus] | 0 | 22.681 | 19 | 5 | 19 | 5 | 1 | 296 | 33.6 | 10.04 | 69 | 5 | 15901122.85 | 18501038.13 | 16512038.89 | 17360641.8 | 12005240.01 | 14223698.06 |
| 296217427 | PREDICTED: 39S ribosomal protein L17, mitochondrial [Callithrix jacchus] | 0 | 12.319 | 26 | 5 | 13 | 5 | 1 | 175 | 19.9 | 9.79 | 42 | 5 | 13609619.64 | 11807917.33 | 4906327.052 | 15947260.17 | 15411658.14 | 13648830.65 |
| 675652680 | PREDICTED: 39S ribosomal protein L2, mitochondrial isoform X1 [Callithrix jacchus] | 0 | 32.613 | 22 | 6 | 20 | 6 | 1 | 313 | 34.4 | 11 | 120 | 6 | 7117763.067 | 4042780.5 | 3662503.976 | 3967447.314 | 3269094.606 | 1580232.337 |
| 532333613 | pyruvate dehydrogenase protein X component, mitochondrial isoform 1 precursor [Callithrix jacchus] | 0 | 83.883 | 34 | 14 | 141 | 14 | 1 | 502 | 54.1 | 8.43 | 2288 | 14 | 172603331 | 167449058.7 | 179869500.6 | 16723907 | 162718193.9 | 173782942.1 |
| 675674223 | PREDICTED: ATP-binding cassette sub-family F member 2 isoform X1 [Callithrix jacchus] | 0 | 51.15 | 19 | 10 | 32 | 10 | 1 | 624 | 71.4 | 7.24 | 348 | 10 | 19814498.82 | 16438027.38 | 12822697.44 | 10760557.03 | 8984971.793 | 9713833.703 |
| 1061019158 | PREDICTED: angiomotin [Callithrix jacchus] | 0 | 60.088 | 13 | 13 | 46 | 12 | 1 | 1094 | 119.1 | 8 | 393 | 13 | 24070638.92 | 18921697.34 | 21695644.65 | 9770870.228 | 10437834.69 | 10123353.06 |
| 532513496 | eukaryotic translation initiation factor 3 subunit B [Callithrix jacchus] | 0 | 156.851 | 41 | 24 | 146 | 24 | 1 | 810 | 92.2 | 5.08 | 2949 | 24 | 38174346.72 | 65040558.38 | 43206977.36 | 50487236.84 | 49974972.74 | 43921682.92 |
| 532519102 | EF-hand domain-containing protein D2 [Callithrix jacchus] | 0 | 73.74 | 50 | 13 | 195 | 12 | 1 | 241 | 26.7 | 5.2 | 4211 | 13 | 288342513.6 | 283596672.1 | 323219079.2 | 279941371.9 | 236633537.1 | 245653491.5 |
| 296218414 | PREDICTED: pre-mRNA-processing factor 19 isoform X1 [Callithrix jacchus] | 0 | 36.399 | 26 | 8 | 24 | 8 | 1 | 504 | 55.1 | 6.61 | 85 | 8 | 14202015.15 | 15387218.81 | 18436028.13 | 15670710.29 | 11927320.66 | 10194705.3 |
| 532338059 | WASH complex subunit FAM21A [Callithrix jacchus] | 0 | 68.87 | 13 | 13 | 40 | 13 | 1 | 1336 | 147.1 | 4.79 | 241 | 13 | 14078560.08 | 14382343.38 | 12873351.9 | 13348221.81 | 10441701.02 | 15366087.31 |
| 296190427 | PREDICTED: 39S ribosomal protein L50, mitochondrial [Callithrix jacchus] | 0 | 23.957 | 41 | 4 | 10 | 4 | 1 | 159 | 18.2 | 8.6 | 71 | 4 | 7359384.604 | 3429404.75 | 8771019.215 | 3645614.903 | 3904076.075 | 5180394.956 |
| 675754941 | PREDICTED: ATP synthase subunit O, mitochondrial isoform X1 [Callithrix jacchus] | 0 | 78.178 | 66 | 11 | 105 | 11 | 1 | 213 | 23.3 | 9.86 | 1622 | 11 | 156801831.8 | 162185614.5 | 161702797.3 | 167506681.1 | 167245596 | 159963202 |
| 532445345 | tRNA-splicing ligase RtcB homolog [Callithrix jacchus] | 0 | 205.029 | 61 | 27 | 172 | 27 | 1 | 505 | 55.2 | 7.23 | 2069 | 27 | 238601758.1 | 213050573.3 | 265052444.9 | 214333625.2 | 224476260.5 | 216681853 |
| 296226401 | PREDICTED: mitochondrial import receptor subunit TOM70 [Callithrix jacchus] | 0 | 247.345 | 62 | 26 | 205 | 26 | 1 | 608 | 67.3 | 6.74 | 3221 | 26 | 238679802.2 | 228342280.8 | 252441568.8 | 220970928 | 216006885.6 | 221610856.5 |
| 675755674 | PREDICTED: ES1 protein homolog, mitochondrial isoform X2 [Callithrix jacchus] | 0 | 71.646 | 56 | 10 | 59 | 10 | 1 | 268 | 28.1 | 8.35 | 951 | 10 | 47752750.93 | 49632180.5 | 43949204.1 | 30451977.54 | 22630589.31 | 22011731.63 |
| 1060939073 | PREDICTED: protein NDRG2 isoform X1 [Callithrix jacchus] | 0 | 89.169 | 45 | 10 | 70 | 10 | 1 | 371 | 40.7 | 5.3 | 1029 | 10 | 23764466.34 | 21133455.33 | 25236451.89 | 30099325.64 | 27524448.67 | 21646872.62 |
| 532346505 | huntingtin-interacting protein 1-related protein [Callithrix jacchus] | 0 | 175.767 | 32 | 31 | 160 | 29 | 1 | 1068 | 119.5 | 6.64 | 2410 | 31 | 70778206.49 | 70040547.75 | 78449370.06 | 43519947.18 | 43576497.53 | 56395184.65 |
| 532291968 | double-stranded RNA-binding protein Staufen homolog 2 isoform a [Callithrix jacchus] | 0 | 50.561 | 24 | 11 | 56 | 9 | 1 | 570 | 62.6 | 9.6 | 210 | 11 | 18119767.98 | 18546633.5 | 18400992.87 | 17898169.28 | 16470070.88 | 14243698.57 |
| 532508802 | ninjurin-2 [Callithrix jacchus] | 0 | 31.78 | 33 | 5 | 45 | 5 | 1 | 190 | 20.8 | 8.22 | 512 | 5 | 30942732.79 | 33477624.38 | 28055047.99 | 16831562.65 | 29088610.54 | 20213187.16 |
| 1061018356 | PREDICTED: calcium-binding protein 1 [Callithrix jacchus] | 0 | 27.789 | 24 | 7 | 46 | 7 | 1 | 369 | 39.6 | 7.85 | 254 | 7 | 9336774.456 | 9697726.375 | 10433009.59 | 10450588.42 | 11196631.69 | 11093470.28 |
| 1060954163 | PREDICTED: chondroitin sulfate proteoglycan 5 [Callithrix jacchus] | 0 | 28.378 | 13 | 5 | 34 | 5 | 1 | 566 | 60.1 | 4.53 | 407 | 5 | 31260606.83 | 40211533.5 | 27258916.41 | 19216229.11 | 22915065.8 | 28258486.65 |
| 532205813 | proSAAS precursor [Callithrix jacchus] | 0 | 41.037 | 28 | 6 | 26 | 6 | 1 | 260 | 27.4 | 7.24 | 449 | 6 | 3576559.044 | 3907895.672 | 7785005.785 | 4731308.493 | 3015387.781 | 3172553.297 |
| 675756476 | PREDICTED: mitochondrial import inner membrane translocase subunit Tim13 [Callithrix jacchus] | 0 | 46.865 | 71 | 6 | 65 | 6 | 1 | 95 | 10.5 | 8.18 | 1039 | 6 | 49522902.76 | 51814850.75 | 50754915 | 55276889.41 | 58375912.43 | 48265205.58 |
| 1060939791 | PREDICTED: mitochondrial import inner membrane translocase subunit Tim9 [Callithrix jacchus] | 0 | 27.487 | 62 | 5 | 26 | 5 | 1 | 89 | 10.4 | 7.21 | 219 | 5 | 30345753.68 | 31623592 | 33321918.87 | 34552030.38 | 39464201.79 | 32407637.87 |
| 675693784 | PREDICTED: mitochondrial import inner membrane translocase subunit Tim8 B [Callithrix jacchus] | 0 | 11.68 | 26 | 3 | 21 | 3 | 1 | 113 | 13 | 8.66 | 70 | 3 | 10267328.39 | 8801520.359 | 9389203.245 | 9315600.131 | 8649092.091 | 9201741.707 |
| 296235993 | PREDICTED: mitochondrial import inner membrane translocase subunit Tim8 A [Callithrix jacchus] | 0 | 20.791 | 46 | 4 | 24 | 4 | 1 | 97 | 11 | 5.16 | 237 | 4 | 21786178.96 | 22893498.7 | 21266139.94 | 26877673.67 | 40706200.79 | 28567347.7 |
| 296210092 | PREDICTED: cortactin-binding protein 2 [Callithrix jacchus] | 0 | 91.006 | 13 | 19 | 68 | 19 | 1 | 1662 | 181.2 | 7.93 | 476 | 19 | 47972123.63 | 46219417 | 43743466.75 | 37707438.58 | 32964397.34 | 34902943.81 |
| 296214900 | PREDICTED: peptidyl-prolyl cis-trans isomerase FKBP3 [Callithrix jacchus] | 0 | 34.132 | 26 | 6 | 50 | 6 | 1 | 224 | 25.2 | 9.28 | 650 | 6 | 32832494.5 | 36849466.5 | 33580646.28 | 27310384.37 | 26520412.64 | 25985831.15 |
| 675677581 | PREDICTED: myeloid leukemia factor 2 [Callithrix jacchus] | 0 | 45.411 | 38 | 7 | 73 | 7 | 1 | 248 | 28.2 | 6.9 | 985 | 7 | 41190848.23 | 46562246 | 34124769.49 | 46687586.61 | 27567476.77 | 36908068.6 |
| 296221476 | PREDICTED: glutaredoxin-3 [Callithrix jacchus] | 0 | 16.371 | 12 | 2 | 7 | 2 | 1 | 335 | 37.4 | 5.6 | 199 | 2 | 3017810.049 | 495636.4688 | 2422134.175 | 2997437.081 | 3094222.473 | 2414880.012 |
| 532519738 | myc box-dependent-interacting protein 1 isoform 1 [Callithrix jacchus] | 0 | 207.658 | 44 | 25 | 321 | 23 | 1 | 593 | 64.8 | 5.1 | 3735 | 25 | 147349308.4 | 144975167.3 | 147340680 | 179560984 | 179540025.5 | 176039706.2 |
| 532284832 | C-Jun-amino-terminal kinase-interacting protein 3 isoform 1 [Callithrix jacchus] | 0 | 141.256 | 24 | 26 | 138 | 21 | 1 | 1348 | 148.5 | 5.5 | 1414 | 26 | 37837214.74 | 48758130.81 | 47472592.62 | 45771957.04 | 48738018.36 | 44447823.41 |
| 532509233 | vesicle-associated membrane protein-associated protein A isoform 2 [Callithrix jacchus] | 0 | 142.001 | 71 | 15 | 186 | 14 | 1 | 249 | 27.9 | 8.62 | 1664 | 15 | 254044667.3 | 237446954.8 | 229037741 | 194026969.4 | 224830727.5 | 245203728.7 |
| 532522500 | 28S ribosomal protein S7, mitochondrial [Callithrix jacchus] | 0 | 43.738 | 45 | 11 | 37 | 11 | 1 | 242 | 28.4 | 10.08 | 131 | 11 | 14020308.57 | 12132666.31 | 8488211.172 | 15489883.36 | 16618211.45 | 8815939.888 |
| 1061000445 | PREDICTED: sacsin [Callithrix jacchus] | 0 | 320.044 | 18 | 72 | 262 | 72 | 1 | 4588 | 521.7 | 7.12 | 2709 | 72 | 63818057.92 | 67927218.7 | 83709187.68 | 71411970.31 | 81786340.91 | 72909074.36 |
| 675691235 | PREDICTED: regulator of G-protein signaling 6 isoform X3 [Callithrix jacchus] | 0 | 112.529 | 37 | 14 | 59 | 11 | 1 | 553 | 63.1 | 8.21 | 1255 | 14 | 22423050.88 | 17146727.03 | 20503334.05 | 22071354.88 | 27364828.41 | 26426742.14 |
| 532284120 | membrane-associated guanylate kinase, WW and PDZ domain-containing protein 2 [Callithrix jacchus] | 0 | 141.978 | 25 | 28 | 171 | 27 | 1 | 1453 | 158.5 | 6.42 | 1620 | 28 | 122411213.5 | 140058233.6 | 112208835.4 | 116064383.6 | 122674669.3 | 114555946.4 |
| 296224141 | PREDICTED: spastin isoform X1 [Callithrix jacchus] | 0 | 67.662 | 29 | 15 | 70 | 15 | 1 | 616 | 67.2 | 9.6 | 882 | 15 | 35113626.06 | 36757336.56 | 35799647.78 | 29784525.86 | 31304361.15 | 30306964.15 |
| 1060938685 | PREDICTED: tropomodulin-2 [Callithrix jacchus] | 0 | 309.683 | 74 | 23 | 395 | 22 | 1 | 351 | 39.5 | 5.27 | 8332 | 23 | 527076219.4 | 568044191.7 | 519716495.1 | 576092838.6 | 565366714.6 | 556653434.5 |
| 1060975705 | PREDICTED: myeloid-associated differentiation marker [Callithrix jacchus] | 0 | 44.646 | 28 | 5 | 26 | 5 | 1 | 322 | 35.3 | 8.15 | 931 | 5 | 45858382.46 | 43207101.88 | 48490117.96 | 69124279.32 | 59490262.26 | 49155663.94 |
| 296196170 | PREDICTED: heterogeneous nuclear ribonucleoprotein D-like isoform X1 [Callithrix jacchus] | 0 | 43.917 | 21 | 8 | 31 | 6 | 1 | 420 | 46.3 | 9.57 | 563 | 8 | 25881146.15 | 25199395.13 | 13484439.21 | 8494286.481 | 6679677.101 | 5627826.359 |
| 675658609 | PREDICTED: MAGUK p55 subfamily member 2 isoform X2 [Callithrix jacchus] | 0 | 211.075 | 61 | 27 | 233 | 26 | 1 | 552 | 61.5 | 6.44 | 2380 | 27 | 274216352.6 | 261652975.3 | 267940039.3 | 197557340.7 | 198372409.4 | 205507100.8 |
| 390459547 | PREDICTED: regulator of G-protein signaling 14 [Callithrix jacchus] | 0 | 39.572 | 19 | 9 | 30 | 9 | 1 | 567 | 61.6 | 8.34 | 214 | 9 | 9800093.419 | 8960173.453 | 8489144.111 | 14036418.33 | 9272270.056 | 12764146.84 |
| 296234514 | EDICTED: serine/threonine-protein phosphatase 2A 65 kDa regulatory subunit A alpha isoform [Callithrix jacc | 0 | 296.864 | 61 | 30 | 343 | 30 | 1 | 589 | 65.3 | 5.11 | 6738 | 30 | 666702053.7 | 642825909.4 | 655804565.2 | 726369715.7 | 720374479.9 | 683008694.8 |
| 1061011077 | PREDICTED: CTP synthase 1 [Callithrix jacchus] | 0 | 157.241 | 46 | 25 | 145 | 21 | 1 | 591 | 66.7 | 6.54 | 1829 | 25 | 89290225.48 | 85483206.13 | 94286933.19 | 91227032.39 | 93315254.59 | 89053580.02 |
| 532287088 | heterogeneous nuclear ribonucleoprotein U isoform a [Callithrix jacchus] | 0 | 141.04 | 33 | 25 | 202 | 25 | 1 | 825 | 90.6 | 6.01 | 1998 | 25 | 122539223.3 | 111898150.7 | 120283374.6 | 88694415.68 | 101512985.1 | 114027884.5 |
| 675700160 | PREDICTED: autophagy-related protein 13 isoform X1 [Callithrix jacchus] | 0 | 21.367 | 14 | 6 | 15 | 6 | 1 | 550 | 60 | 5.22 | 51 | 6 | 17143028.67 | 15600599.31 | 14961708.58 | 18844676 | 18063412.34 | 18731946.94 |
| 675720865 | PREDICTED: rab11 family-interacting protein 5 isoform X1 [Callithrix jacchus] | 0 | 207.66 | 29 | 27 | 187 | 27 | 1 | 1309 | 137.5 | 5.14 | 3421 | 27 | 49915981.1 | 44796772.61 | 53347027.7 | 59904881.01 | 64245853.51 | 61353717.95 |
| 675713156 | PREDICTED: 1-acyl-sn-glycerol-3-phosphate acyltransferase epsilon isoform X1 [Callithrix jacchus] | 0 | 24.402 | 19 | 7 | 37 | 7 | 1 | 364 | 42.2 | 9.28 | 444 | 7 | 39964571.96 | 38599446.94 | 34847032.2 | 16523955.49 | 12806462.21 | 17374838.59 |
| 1060955003 | REDICTED: LOW QUALITY PROTEIN: calcium/calmodulin-dependent protein kinase type 1 [Callithrix jacchu | 0 | 47.143 | 16 | 8 | 39 | 5 | 1 | 537 | 58.7 | 6.98 | 471 | 8 | 15133848.35 | 14873066.38 | 13334360.98 | 9529926.492 | 10729086.48 | 14572010.27 |
| 1060948446 | PREDICTED: adhesion G protein-coupled receptor A1 [Callithrix jacchus] | 0 | 45.814 | 16 | 7 | 39 | 7 | 1 | 554 | 60.2 | 8.09 | 533 | 7 | 17972807.13 | 20396161.5 | 15897484.11 | 20853710.95 | 19567376.53 | 18767959.66 |
| 675648939 | PREDICTED: septin-11 isoform X1 [Callithrix jacchus] | 0 | 338.274 | 63 | 32 | 556 | 17 | 1 | 437 | 50.3 | 6.39 | 11044 | 32 | 1022352916 | 1094317542 | 1064338994 | 1068335402 | 1070161791 | 1196175023 |
| 296217097 | PREDICTED: 39S ribosomal protein L48, mitochondrial isoform X1 [Callithrix jacchus] | 0 | 22.489 | 23 | 4 | 20 | 4 | 1 | 261 | 29.2 | 9.35 | 293 | 4 | 11067875 | 11043545.84 | 10168602.32 | 9178283.141 | 9823262.163 | 8171285.898 |
| 296222651 | PREDICTED: LOW QUALITY PROTEIN: 3-ketoacyl-CoA thiolase, mitochondrial [Callithrix jacchus] | 0 | 41.712 | 27 | 8 | 31 | 8 | 1 | 397 | 41.9 | 8.4 | 145 | 8 | 16868395.47 | 16674100.27 | 18841345.22 | 16737102.09 | 14899516.58 | 11533039.24 |
| 1060976120 | REDICTED: small glutamine-rich tetratricopeptide repeat-containing protein alpha isoform X1 [Callithrix jacchu | 0 | 12.725 | 11 | 3 | 8 | 3 | 1 | 353 | 37.3 | 6.76 | 33 | 3 | 28911105.16 | 32394531.06 | 26498470.35 | 34023693.62 | 29653501.43 | 26744864.37 |
| 1060957426 | PREDICTED: neurocalcin-delta [Callithrix jacchus] | 0 | 66.672 | 57 | 11 | 79 | 6 | 1 | 193 | 22.2 | 5.35 | 819 | 11 | 16151827.05 | 13697557.5 | 10602441.02 | 13763299.44 | 13241086.86 | 11532422.31 |
| 675674052 | PREDICTED: LOW QUALITY PROTEIN: extended synaptotagmin-2 [Callithrix jacchus] | 0 | 44.756 | 16 | 10 | 32 | 10 | 1 | 894 | 99.1 | 8.29 | 435 | 10 | 69722124.73 | 73896239.88 | 43940462.83 | 64541343.23 | 50139183.63 | 84086142.36 |
| 296220643 | PREDICTED: CDGSH iron-sulfur domain-containing protein 1 [Callithrix jacchus] | 0 | 63.958 | 57 | 7 | 99 | 7 | 1 | 108 | 12.2 | 9.09 | 1963 | 7 | 271215085.7 | 278086743.3 | 240114768.3 | 221517096 | 230497293.4 | 230883360.1 |
| 1060966945 | PREDICTED: probable D-lactate dehydrogenase, mitochondrial isoform X1 [Callithrix jacchus] | 0 | 18.713 | 11 | 5 | 20 | 5 | 1 | 484 | 51.7 | 6.32 | 253 | 5 | 6702909.283 | 7032693.75 | 5349380.26 | 7040521.48 | 5823544.556 | 5502515.235 |
| 532285020 | protein RUFY3 isoform 1 [Callithrix jacchus] | 0 | 127.918 | 42 | 25 | 149 | 24 | 1 | 620 | 70.6 | 5.66 | 1499 | 25 | 96277078.93 | 219274338 | 190618033.4 | 98541955.57 | 86981005.34 | 88815091.03 |
| 296191527 | PREDICTED: small G protein signaling modulator 1 isoform X1 [Callithrix jacchus] | 0 | 6.986 | 3 | 2 | 2 | 2 | 1 | 1147 | 129.4 | 5.83 | 16 | 2 | 1243058.015 | 1151289.875 | 1491123.241 | 1755459.77 | 1108756.61 | 921612.1972 |
| 296194482 | PREDICTED: regulator of G-protein signaling 7-binding protein [Callithrix jacchus] | 0 | 79.19 | 54 | 8 | 62 | 8 | 1 | 257 | 28.9 | 8.46 | 1280 | 8 | 57508879.43 | 59486564.69 | 65962204.39 | 108943988.4 | 82834309.01 | 57591695.49 |
| 1060994791 | PREDICTED: protein SCO1 homolog, mitochondrial [Callithrix jacchus] | 0 | 28.787 | 19 | 6 | 17 | 6 | 1 | 344 | 38.3 | 9.2 | 32 | 6 | 9576747.027 | 9710736.25 | 10772129.44 | 6710570.889 | 8064825.886 | 7984256.015 |
| 532333879 | disks large homolog 3 isoform a [Callithrix jacchus] | 0 | 387.07 | 59 | 44 | 454 | 40 | 1 | 831 | 91.6 | 6.95 | 6141 | 44 | 611885315.7 | 600984031.7 | 617829662 | 656058761.9 | 643467919.6 | 624567083.9 |
| 296218542 | PREDICTED: nuclear RNA export factor 1 [Callithrix jacchus] | 0.005 | 3.74 | 3 | 2 | 2 | 2 | 1 | 619 | 70.2 | 8.51 | 0 | 2 | 1407020.805 | 1595585.938 | 2278193.054 | 1021478.411 | 1873806.252 | 1115939.194 |
| 532524094 | contactin-associated protein 1 precursor [Callithrix jacchus] | 0 | 430.58 | 46 | 51 | 611 | 51 | 1 | 1377 | 155.5 | 7.08 | 12508 | 51 | 945290361.9 | 932804787.6 | 987438932.9 | 1243754600 | 1168182486 | 1127608893 |
| 296192510 | PREDICTED: neuronal pentraxin-2 [Callithrix jacchus] | 0 | 14.071 | 12 | 5 | 16 | 4 | 1 | 431 | 47.1 | 5.78 | 14 | 5 | 2422469.254 | 3009292.125 | 5335849.707 | 6601604.223 | 7492494.71 | 6396048.18 |
| 532511504 | membrane-associated progesterone receptor component 1 [Callithrix jacchus] | 0.001 | 6.519 | 11 | 2 | 5 | 2 | 1 | 195 | 21.7 | 4.7 | 46 | 2 | 7361212.811 | 10577722 | 7725418.933 | 7179418.262 | 6916056.751 | 6841448.361 |
| 1060988498 | PREDICTED: syntaxin-7 isoform X1 [Callithrix jacchus] | 0 | 55.675 | 50 | 9 | 84 | 9 | 1 | 261 | 29.8 | 5.55 | 1525 | 9 | 43085588.38 | 37272102 | 36082580.73 | 33157169.14 | 32885335.19 | 31109862.15 |
| 675770674 | PREDICTED: myotubularin-related protein 1 isoform X1 [Callithrix jacchus] | 0 | 28.822 | 15 | 9 | 14 | 8 | 1 | 689 | 77.3 | 6.54 | 26 | 9 | 12084520.72 | 11441528.38 | 15293402.02 | 10968718.14 | 10355903.25 | 12044976 |
| 296234064 | EDICTED: LOW QUALITY PROTEIN: mitochondrial import receptor subunit TOM40 homolog [Callithrix jacch | 0 | 126.314 | 43 | 11 | 153 | 11 | 1 | 361 | 37.9 | 7.25 | 2952 | 11 | 179347684.2 | 159764890.2 | 148713970.3 | 146631542.1 | 154116054.1 | 152814599.1 |
| 675757724 | PREDICTED: eukaryotic translation initiation factor 3 subunit G [Callithrix jacchus] | 0 | 44.577 | 27 | 9 | 53 | 9 | 1 | 356 | 39.2 | 6.6 | 905 | 9 | 25679418.8 | 28090262.98 | 20682445.49 | 25366748.27 | 16221257.95 | 21012383.65 |
| 1060956168 | PREDICTED: syntenin-1 isoform X1 [Callithrix jacchus] | 0 | 61.198 | 41 | 9 | 86 | 9 | 1 | 298 | 32.5 | 7.49 | 855 | 9 | 48208209.06 | 45945085.25 | 41066113.56 | 39199856.66 | 43035747.45 | 46881897.59 |
| 675672750 | PREDICTED: heterogeneous nuclear ribonucleoproteins A2/B1 isoform X1 [Callithrix jacchus] | 0 | 210.164 | 54 | 24 | 374 | 19 | 1 | 396 | 41.1 | 8.9 | 5783 | 24 | 474191350.3 | 464373586.4 | 451149675.6 | 220516681.8 | 218158457.6 | 217953198.6 |
| 1060949073 | PREDICTED: thioredoxin-like protein 1 isoform X1 [Callithrix jacchus] | 0 | 29.329 | 19 | 5 | 24 | 5 | 1 | 315 | 35.1 | 4.97 | 331 | 5 | 2463221.174 | 2992023.594 | 4414807.564 | 7224024.126 | 7485579.609 | 7158795.345 |
| 1060999355 | PREDICTED: exocyst complex component 7 isoform X1 [Callithrix jacchus] | 0 | 178.991 | 51 | 32 | 200 | 32 | 1 | 721 | 82.2 | 6.71 | 2565 | 32 | 130694476.7 | 132609998.2 | 125670489 | 143819586.8 | 143474228.8 | 133584094.2 |
| 675764106 | PREDICTED: synaptotagmin-5 isoform X1 [Callithrix jacchus] | 0 | 42.053 | 19 | 8 | 66 | 5 | 1 | 386 | 42.9 | 9.16 | 820 | 8 | 15288078.58 | 16215827.38 | 12648118.43 | 2586364.13 | 6974915.13 | 13068087.18 |
| 296190175 | PREDICTED: dynactin subunit 3 isoform X1 [Callithrix jacchus] | 0 | 31.622 | 41 | 8 | 64 | 8 | 1 | 186 | 21.2 | 5.64 | 750 | 8 | 75677276 | 78789726 | 70185359.96 | 60581552.66 | 68340235.78 | 64123786.12 |
| 675767727 | PREDICTED: non-POU domain-containing octamer-binding protein [Callithrix jacchus] | 0 | 151.214 | 51 | 21 | 231 | 19 | 1 | 471 | 54.2 | 8.95 | 2360 | 21 | 167235448.9 | 171877047.8 | 185759937.1 | 98964633.71 | 104233340.4 | 145208315.7 |
| 675690899 | PREDICTED: vesicle transport through interaction with t-SNAREs homolog 1B isoform X1 [Callithrix jacchus] | 0 | 18.576 | 19 | 4 | 28 | 4 | 1 | 232 | 26.7 | 9.04 | 181 | 4 | 12710823.91 | 12180272.81 | 12819397.07 | 7469842.059 | 6942842.411 | 6961036.225 |
| 532523134 | glutamate receptor 3 isoform 1 precursor [Callithrix jacchus] | 0 | 252.196 | 36 | 30 | 341 | 5 | 1 | 894 | 101 | 8.46 | 6631 | 30 | 42066631.37 | 48701841.75 | 41612483.07 | 70353613.43 | 63745298.22 | 53521725.62 |
| 532348827 | protein PRRC2A [Callithrix jacchus] | 0 | 78.674 | 11 | 17 | 70 | 16 | 1 | 2163 | 229 | 9.45 | 343 | 17 | 13212193.24 | 11625842.94 | 17673244.09 | 18181149.3 | 12925159.12 | 13458647.8 |
| 675651861 | PREDICTED: LOW QUALITY PROTEIN: spliceosome RNA helicase DDX39B [Callithrix jacchus] | 0 | 18.294 | 12 | 4 | 17 | 4 | 1 | 422 | 48.2 | 5.67 | 87 | 4 | 5199296.347 | 5761570.75 | 5550011.493 | 5797259.533 | 4925090.753 | 3600941.551 |
| 675640399 | PREDICTED: tubulin alpha-8 chain [Callithrix jacchus] | 0 | 356.76 | 65 | 25 | 931 | 10 | 1 | 449 | 50.1 | 5.06 | 24664 | 25 | 36235955.45 | 38084525.13 | 47555355.23 | 35510974.41 | 41290521.41 | 42985019.25 |
| 532273108 | septin-9 isoform b [Callithrix jacchus] | 0 | 261.566 | 69 | 34 | 416 | 34 | 1 | 579 | 64.8 | 7.59 | 5956 | 34 | 408423446.4 | 395458060.1 | 409506808.5 | 426546647.3 | 468551693.9 | 438492810.7 |
| 296208256 | PREDICTED: zinc finger Ran-binding domain-containing protein 2 isoform X1 [Callithrix jacchus] | 0 | 50.401 | 36 | 10 | 90 | 10 | 1 | 330 | 37.4 | 10.01 | 727 | 10 | 32321650.42 | 33607753.69 | 33484126.72 | 33020401.16 | 28115756.25 | 35264732.53 |
| 1060954883 | PREDICTED: contactin-6 [Callithrix jacchus] | 0 | 32.238 | 7 | 6 | 33 | 5 | 1 | 1053 | 116.5 | 6.62 | 585 | 6 | 5553988.133 | 7617945.75 | 9651505.459 | 10513857.42 | 10692550.53 | 9073605.836 |
| 1060943239 | PREDICTED: caprin-1 [Callithrix jacchus] | 0 | 113.793 | 29 | 14 | 98 | 14 | 1 | 709 | 78.3 | 5.25 | 1767 | 14 | 77055632.05 | 80907163.39 | 66987894.66 | 68882007.89 | 70381143.81 | 75130894.86 |
| 532347385 | cadherin EGF LAG seven-pass G-type receptor 2 precursor [Callithrix jacchus] | 0 | 67.205 | 6 | 12 | 32 | 12 | 1 | 2929 | 317.6 | 5.35 | 304 | 12 | 8886439.052 | 10008614.88 | 9523780.17 | 9727214.002 | 8894940.23 | 11097634.77 |
| 532519170 | rho guanine nucleotide exchange factor 7 isoform b [Callithrix jacchus] | 0 | 183.908 | 44 | 26 | 164 | 26 | 1 | 782 | 87.7 | 7.09 | 2539 | 26 | 115682820.6 | 136172637.3 | 103823877.5 | 86827173.06 | 85611238.43 | 103544502.5 |
| 675757339 | PREDICTED: 40S ribosomal protein S28 [Callithrix jacchus] | 0 | 19.226 | 46 | 3 | 59 | 3 | 1 | 69 | 7.8 | 10.7 | 1641 | 3 | 51874306.31 | 53670059.75 | 54517794.71 | 55220539.89 | 60013946.68 | 65533511.49 |
| 296221101 | PREDICTED: alpha-centractin [Callithrix jacchus] | 0 | 158.721 | 55 | 18 | 201 | 10 | 1 | 376 | 42.6 | 6.64 | 3015 | 18 | 51372725.23 | 46153223.71 | 64436505.4 | 59899562.72 | 58229027.3 | 55599027.12 |
| 675730786 | PREDICTED: coatomer subunit gamma-1 [Callithrix jacchus] | 0 | 25.612 | 10 | 7 | 13 | 7 | 1 | 874 | 97.7 | 5.43 | 64 | 7 | 3482259.571 | 3668455.75 | 4611485.969 | 1324558.658 | 735878.6389 | 2471583.109 |
| 296207548 | PREDICTED: ras-related GTP-binding protein C [Callithrix jacchus] | 0 | 24.9 | 17 | 6 | 24 | 6 | 1 | 399 | 44.2 | 5.1 | 228 | 6 | 8717283.277 | 8180782 | 9414718.582 | 14623079.16 | 13390660.13 | 10734186.6 |
| 1060997488 | PREDICTED: rab GTPase-binding effector protein 1 [Callithrix jacchus] | 0 | 40.218 | 13 | 10 | 30 | 10 | 1 | 862 | 99.3 | 5.02 | 271 | 10 | 50510298.37 | 43282212.19 | 42972925.59 | 39818266.2 | 43207109.08 | 42965163.1 |
| 296195333 | PREDICTED: guanylate cyclase soluble subunit beta-1 isoform X1 [Callithrix jacchus] | 0 | 68.66 | 28 | 15 | 88 | 15 | 1 | 619 | 70.4 | 5.35 | 836 | 15 | 41200868.46 | 43911738.41 | 38005290.89 | 52790455.41 | 50403564.68 | 42852197.77 |
| 166183799 | sorting nexin 3 (predicted) [Callithrix jacchus] | 0 | 21.47 | 36 | 6 | 25 | 6 | 1 | 162 | 18.8 | 8.66 | 352 | 6 | 41512033.97 | 38154612.75 | 30950985.39 | 6622603.112 | 10745774.43 | 6593347.656 |
| 1060982468 | PREDICTED: protocadherin-7 isoform X1 [Callithrix jacchus] | 0 | 58.941 | 12 | 11 | 40 | 11 | 1 | 1275 | 138.8 | 5.16 | 609 | 11 | 16977450.85 | 18072210.19 | 17099872.91 | 9892367.478 | 9990712.197 | 13069597.01 |
| 390478415 | PREDICTED: 60S ribosomal protein L36 [Callithrix jacchus] | 0 | 21.35 | 30 | 5 | 37 | 5 | 1 | 105 | 12.2 | 11.59 | 289 | 5 | 138246579.9 | 141123366.5 | 121385947.2 | 129641831.9 | 130317962.7 | 115960648 |
| 1060935347 | PREDICTED: band 4.1-like protein 4B [Callithrix jacchus] | 0 | 53.049 | 18 | 13 | 59 | 12 | 1 | 835 | 94 | 8.98 | 357 | 13 | 19479943.64 | 13586772.31 | 19699848.72 | 28327070.06 | 24368392.71 | 23666159.78 |
| 296219788 | PREDICTED: voltage-dependent calcium channel gamma-3 subunit [Callithrix jacchus] | 0 | 79.154 | 33 | 10 | 112 | 8 | 1 | 315 | 35.5 | 9.45 | 987 | 10 | 95741800.59 | 97669000.42 | 97232000.79 | 151169226 | 146916461.1 | 150464074 |
| 532505314 | voltage-dependent calcium channel gamma-4 subunit [Callithrix jacchus] | 0 | 7.058 | 6 | 2 | 12 | 2 | 1 | 327 | 36.5 | 9.1 | 24 | 2 | 4653067.275 | 3527967.25 | 4101797.609 | 3384050.124 | 2874649.898 | 2795763.208 |
| 675651692 | PREDICTED: gamma-aminobutyric acid type B receptor subunit 1 [Callithrix jacchus] | 0 | 190.862 | 33 | 26 | 219 | 5 | 1 | 964 | 108.6 | 8.13 | 3047 | 26 | 37382918.05 | 38363242 | 38874086.69 | 47710322.74 | 38695760.89 | 42743977.46 |
| 675656793 | PREDICTED: 14-3-3 protein beta/alpha [Callithrix jacchus] | 0 | 136.979 | 59 | 16 | 389 | 8 | 1 | 246 | 28.1 | 4.83 | 7045 | 16 | 126438764.7 | 129159294.1 | 118613830.4 | 106728417.8 | 89801130.04 | 115999217.1 |
| 532519730 | electrogenic sodium bicarbonate cotransporter 1 isoform 1 [Callithrix jacchus] | 0 | 73.005 | 16 | 12 | 50 | 12 | 1 | 1079 | 121.2 | 7.01 | 765 | 12 | 44029034.58 | 42886064.56 | 26063425.05 | 47166910.29 | 44086372.78 | 42939239.01 |
| 296236284 | PREDICTED: NADH dehydrogenase [ubiquinone] 1 alpha subcomplex subunit 1 [Callithrix jacchus] | 0 | 27.933 | 71 | 3 | 16 | 3 | 1 | 70 | 8 | 8.78 | 199 | 3 | 17924922.81 | 15557614.38 | 17244781.59 | 22272095.46 | 22581960.8 | 19859166.01 |
| 296230882 | PREDICTED: LOW QUALITY PROTEIN: formin-2 [Callithrix jacchus] | 0 | 301.792 | 32 | 39 | 297 | 39 | 1 | 1621 | 170.7 | 6.34 | 5688 | 39 | 131243127.5 | 137305828.2 | 156091578.7 | 157533502.8 | 194027850.8 | 150989633.9 |
| 390470941 | PREDICTED: toll-interacting protein isoform X2 [Callithrix jacchus] | 0 | 35.097 | 32 | 8 | 50 | 8 | 1 | 274 | 30.3 | 5.97 | 505 | 8 | 31090828.22 | 30754403.91 | 28356593.88 | 26448891.42 | 28534303.11 | 27835703.87 |
| 1061012827 | PREDICTED: LOW QUALITY PROTEIN: synaptotagmin-6 [Callithrix jacchus] | 0 | 36.192 | 28 | 11 | 41 | 10 | 1 | 443 | 50.3 | 8.09 | 87 | 11 | 10500480.96 | 10207881.94 | 12469528.56 | 11729931.02 | 15387970.74 | 16087402.01 |
| 1060944837 | PREDICTED: synaptotagmin-7 [Callithrix jacchus] | 0 | 230.932 | 43 | 27 | 294 | 27 | 1 | 760 | 83.3 | 9.32 | 4153 | 27 | 239358058 | 214918600 | 226364224.8 | 286018089.7 | 278218892.1 | 302982554.4 |
| 390479839 | PREDICTED: ubiquilin-2 [Callithrix jacchus] | 0 | 36.763 | 14 | 6 | 32 | 3 | 1 | 624 | 65.6 | 5.22 | 691 | 6 | 3251100.878 | 3388442.313 | 2782445.377 | 6691446.887 | 4420447.333 | 5173887.011 |
| 675766809 | PREDICTED: WD repeat domain phosphoinositide-interacting protein 4 isoform X1 [Callithrix jacchus] | 0 | 17.751 | 18 | 5 | 11 | 5 | 1 | 361 | 40 | 7.14 | 16 | 5 | 7064578.133 | 6410740.813 | 5867265.814 | 3731507.345 | 3452240.713 | 3148587.69 |
| 532523306 | GRIP1-associated protein 1 isoform 1 [Callithrix jacchus] | 0 | 190.837 | 40 | 32 | 171 | 32 | 1 | 841 | 95.9 | 5.15 | 1845 | 32 | 116292288.6 | 110855157.6 | 124944394.3 | 50860513.93 | 51627481.88 | 58858628.41 |
| 390470712 | PREDICTED: peroxiredoxin-5, mitochondrial isoform X1 [Callithrix jacchus] | 0 | 57.807 | 53 | 8 | 38 | 8 | 1 | 222 | 22.9 | 9.38 | 728 | 8 | 169618860.1 | 186711193.3 | 164991185.6 | 111181772.5 | 103409084.3 | 82734495.75 |
| 675669098 | PREDICTED: eukaryotic translation initiation factor 3 subunit I isoform X1 [Callithrix jacchus] | 0 | 87.014 | 57 | 13 | 92 | 13 | 1 | 325 | 36.5 | 5.64 | 992 | 13 | 37017846.36 | 34980241.06 | 25001650.76 | 31276351.37 | 31118547.43 | 46383437.85 |
| 390469686 | PREDICTED: cell adhesion molecule 1 isoform X6 [Callithrix jacchus] | 0.002 | 5.172 | 4 | 2 | 10 | 2 | 1 | 473 | 51.8 | 4.98 | 0 | 2 | 11604523.85 | 7773000.5 | 7408831.798 | 7494181.903 | 8185184.329 | 2414903.856 |
| 1061019643 | PREDICTED: CTP synthase 2 isoform X1 [Callithrix jacchus] | 0 | 68.8 | 21 | 18 | 96 | 12 | 1 | 904 | 103.3 | 4.89 | 780 | 18 | 22286347.23 | 19770365.94 | 29682555.33 | 32489914.2 | 31751006.28 | 28390353.17 |
| 296201276 | PREDICTED: syntaxin-8 [Callithrix jacchus] | 0 | 12.069 | 17 | 4 | 9 | 4 | 1 | 236 | 27 | 5.07 | 28 | 4 | 4205461.328 | 4086566.375 | 3758140.446 | 2499390.423 | 2735171.487 | 2253517.484 |
| 1060937236 | PREDICTED: eukaryotic translation initiation factor 3 subunit D [Callithrix jacchus] | 0 | 49.54 | 13 | 8 | 43 | 8 | 1 | 781 | 90.3 | 8.54 | 548 | 8 | 26078918.99 | 25296060.13 | 16243324.98 | 25849120.73 | 20973549.2 | 25246204.12 |
| 675708573 | PREDICTED: AP-3 complex subunit mu-1 isoform X1 [Callithrix jacchus] | 0 | 29.678 | 16 | 7 | 48 | 4 | 1 | 490 | 54.6 | 8.07 | 147 | 7 | 3495010.747 | 2340700.25 | 3621457.774 | 4254079.765 | 4112160.329 | 3396179.874 |
| 675665260 | PREDICTED: elongation factor 1-beta [Callithrix jacchus] | 0 | 47.128 | 33 | 7 | 55 | 7 | 1 | 225 | 24.7 | 4.67 | 516 | 7 | 70149948.33 | 61616114.25 | 72115454.82 | 58950460 | 54033773.96 | 47278496.45 |
| 532442639 | E3 ubiquitin-protein ligase PDZRN3 [Callithrix jacchus] | 0 | 35.229 | 11 | 9 | 21 | 8 | 1 | 1066 | 119.6 | 6 | 105 | 9 | 49725460.06 | 46319215.63 | 48345625.26 | 47657793.96 | 38852237.57 | 41126695 |
| 296217393 | PREDICTED: tripartite motif-containing protein 3 isoform X1 [Callithrix jacchus] | 0 | 110.661 | 32 | 20 | 100 | 19 | 1 | 744 | 80.7 | 7.72 | 1059 | 20 | 49544762.68 | 46293851.59 | 40783098.39 | 40811270.89 | 42934444.96 | 38566350.31 |
| 296196279 | PREDICTED: general vesicular transport factor p115 isoform X1 [Callithrix jacchus] | 0 | 60.683 | 14 | 10 | 51 | 10 | 1 | 973 | 109.1 | 4.91 | 1348 | 10 | 16957139.74 | 19183296 | 15818539.85 | 16810983.24 | 15287876.16 | 13918902.71 |
| 675672275 | PREDICTED: ras-related protein Ral-A [Callithrix jacchus] | 0 | 140.144 | 57 | 13 | 163 | 7 | 1 | 206 | 23.5 | 7.11 | 3679 | 13 | 130271242 | 148704694 | 151144076.3 | 193804417.7 | 192631388.4 | 186853567.9 |
| 675750399 | PREDICTED: galectin-8 isoform X2 [Callithrix jacchus] | 0 | 45.91 | 34 | 10 | 37 | 10 | 1 | 317 | 35.8 | 7.27 | 156 | 10 | 15211357.86 | 13054952.84 | 14853100.11 | 15590414.42 | 15938058.23 | 21365735.95 |
| 675643546 | PREDICTED: transcription elongation regulator 1 isoform X1 [Callithrix jacchus] | 0 | 44.436 | 15 | 14 | 50 | 14 | 1 | 1125 | 126.9 | 8.92 | 214 | 14 | 13206335.94 | 12425962.55 | 16687983.93 | 13362973.74 | 6950803.752 | 11986930.53 |
| 675660152 | PREDICTED: diacylglycerol kinase epsilon [Callithrix jacchus] | 0 | 35.009 | 16 | 9 | 42 | 9 | 1 | 567 | 64 | 7.44 | 417 | 9 | 12041200.31 | 13134447.69 | 18721534.97 | 20283317.58 | 19274755.98 | 18945538.51 |
| 1060942250 | PREDICTED: protein SET [Callithrix jacchus] | 0 | 47.629 | 32 | 7 | 59 | 5 | 1 | 277 | 32.1 | 4.22 | 1072 | 7 | 45482374.26 | 46030561.63 | 46001487.2 | 102279787.5 | 102984665.6 | 43498707.49 |
| 390474348 | PREDICTED: N-acetyl-D-glucosamine kinase isoform X1 [Callithrix jacchus] | 0 | 42.724 | 31 | 10 | 38 | 10 | 1 | 344 | 37.4 | 6.13 | 465 | 10 | 14394246.28 | 8747325.594 | 11963154.25 | 8304487.117 | 11120032.96 | 11728396.77 |
| 390479813 | PREDICTED: melanoma-associated antigen D2 [Callithrix jacchus] | 0 | 31.809 | 17 | 9 | 30 | 8 | 1 | 604 | 64.6 | 9.23 | 266 | 9 | 4996004.147 | 4564299.563 | 2643299.908 | 10619922.86 | 8184815.721 | 6328222.236 |
| 532348961 | polypyrimidine tract-binding protein 2 [Callithrix jacchus] | 0 | 20.258 | 11 | 6 | 28 | 5 | 1 | 537 | 58 | 8.81 | 172 | 6 | 15617178.77 | 11779339.63 | 16212648.66 | 2833328.053 | 4800684.837 | 7193705.9 |
| 532272836 | rho-related GTP-binding protein RhoG precursor [Callithrix jacchus] | 0 | 102.628 | 71 | 13 | 211 | 12 | 1 | 191 | 21.2 | 8.12 | 4073 | 13 | 91648451.28 | 92542555.06 | 76773521.71 | 106179096.1 | 93994226.27 | 96324231.88 |
| 296221389 | PREDICTED: LOW QUALITY PROTEIN: serine protease HTRA1 [Callithrix jacchus] | 0 | 58.189 | 26 | 11 | 83 | 11 | 1 | 484 | 51.7 | 7.49 | 1005 | 11 | 30683397.56 | 35846816.25 | 32598415.95 | 30197060.97 | 27932138.84 | 29075930.52 |
| 296220012 | PREDICTED: syntaxin-1B [Callithrix jacchus] | 0 | 174.789 | 55 | 17 | 262 | 16 | 1 | 288 | 33.2 | 5.38 | 4774 | 17 | 327464928.6 | 303616868.5 | 270556191.8 | 368407217 | 328373783.1 | 291034076.4 |
| 675768460 | PREDICTED: heterogeneous nuclear ribonucleoprotein H2 [Callithrix jacchus] | 0 | 84.173 | 34 | 11 | 112 | 6 | 1 | 449 | 49.2 | 6.3 | 2566 | 11 | 22860225.53 | 21607316 | 17868859.58 | 17086205.39 | 26625689.53 | 17503439.86 |
| 532503191 | peroxisomal membrane protein PEX14 [Callithrix jacchus] | 0 | 71.176 | 36 | 8 | 72 | 8 | 1 | 377 | 41.2 | 4.94 | 1439 | 8 | 42411944.05 | 33922689.44 | 39684173.57 | 32925012.45 | 34662049.13 | 28592895.14 |
| 532504872 | RNA-binding protein 14 isoform 1 [Callithrix jacchus] | 0 | 91.701 | 29 | 17 | 125 | 17 | 1 | 669 | 69.5 | 9.67 | 1188 | 17 | 74349675.05 | 74213311.5 | 73791957.78 | 32597870.19 | 38849810.62 | 45554127.58 |
| 390465990 | PREDICTED: 39S ribosomal protein L37, mitochondrial isoform X1 [Callithrix jacchus] | 0 | 41.553 | 20 | 7 | 28 | 7 | 1 | 423 | 48.5 | 9.06 | 443 | 7 | 13374969.54 | 13316342 | 13562859.78 | 13748347.74 | 14978289.28 | 14032702.84 |
| 675661670 | PREDICTED: 39S ribosomal protein L12, mitochondrial [Callithrix jacchus] | 0 | 66.341 | 57 | 10 | 68 | 10 | 1 | 198 | 21.4 | 8.56 | 659 | 10 | 110044860.7 | 91979851.56 | 105938914.1 | 64957073.47 | 69258059.81 | 73611754.93 |
| 296223464 | PREDICTED: 39S ribosomal protein L19, mitochondrial [Callithrix jacchus] | 0 | 36.536 | 34 | 8 | 30 | 8 | 1 | 292 | 33.7 | 9.33 | 205 | 8 | 16565165.7 | 17675510.13 | 10231974.43 | 36237772.82 | 14579870.45 | 12845706.02 |
| 532519572 | drebrin isoform a [Callithrix jacchus] | 0 | 358.193 | 60 | 34 | 582 | 34 | 1 | 650 | 71.6 | 4.46 | 10642 | 34 | 1288542319 | 1328696144 | 1369960796 | 1079571597 | 1161631623 | 1351857459 |
| 675707122 | PREDICTED: eukaryotic translation initiation factor 3 subunit C isoform X1 [Callithrix jacchus] | 0 | 114.282 | 28 | 23 | 126 | 23 | 1 | 914 | 105.5 | 5.68 | 1362 | 23 | 71814000.55 | 59793930.63 | 56589442 | 69057824.44 | 69948219.42 | 57928812.25 |
| 675680331 | PREDICTED: prostaglandin E synthase 3 isoform X1 [Callithrix jacchus] | 0 | 20.288 | 29 | 4 | 13 | 4 | 1 | 165 | 19.4 | 4.89 | 107 | 4 | 6462458.122 | 6220313.375 | 8610977.047 | 8830791.758 | 7412450.361 | 5079398.64 |
| 296202908 | PREDICTED: dnaJ homolog subfamily C member 7 isoform X1 [Callithrix jacchus] | 0 | 14.247 | 10 | 4 | 20 | 4 | 1 | 494 | 56.4 | 6.79 | 233 | 4 | 2484058.575 | 4725717.875 | 2991782.083 | 1723274.842 | 2525834.428 | 4260724.028 |
| 675765947 | PREDICTED: acyl-coenzyme A thioesterase 9, mitochondrial isoform X2 [Callithrix jacchus] | 0 | 35.093 | 22 | 11 | 35 | 11 | 1 | 445 | 50.6 | 8.4 | 34 | 11 | 25750525.76 | 30111173.88 | 26160775 | 15059213.4 | 14494078.81 | 18323718.26 |
| 296205121 | PREDICTED: tomoregulin-2 [Callithrix jacchus] | 0 | 21.143 | 17 | 6 | 25 | 6 | 1 | 374 | 41.4 | 5.15 | 59 | 6 | 9396168.373 | 9505347.75 | 10574846.96 | 4587150.469 | 6579244.643 | 8154999.721 |
| 1060955676 | PREDICTED: transgelin-3 [Callithrix jacchus] | 0 | 182.322 | 84 | 18 | 344 | 18 | 1 | 199 | 22.5 | 7.33 | 6239 | 18 | 889069805.2 | 900012369.1 | 915035442.5 | 904686115.7 | 897369276.5 | 933527121.3 |
| 296212908 | PREDICTED: actin-related protein 2/3 complex subunit 3 [Callithrix jacchus] | 0 | 44.232 | 35 | 6 | 70 | 6 | 1 | 178 | 20.5 | 8.59 | 905 | 6 | 95470430.14 | 102824097.9 | 114530384.8 | 90659693.67 | 108140734.6 | 106529671 |
| 296197022 | PREDICTED: LETM1 and EF-hand domain-containing protein 1, mitochondrial [Callithrix jacchus] | 0 | 109.944 | 31 | 16 | 104 | 16 | 1 | 742 | 83.3 | 6.35 | 1357 | 16 | 51151216.45 | 47700195.44 | 57237465.18 | 52097111 | 50944460.84 | 42021897.31 |
| 532515202 | calcineurin B homologous protein 1 [Callithrix jacchus] | 0 | 24.399 | 29 | 5 | 26 | 5 | 1 | 195 | 22.4 | 5.1 | 297 | 5 | 8820334.894 | 9098729.047 | 7376479.375 | 5238014.159 | 5452374.178 | 6042492.895 |
| 532449197 | heterogeneous nuclear ribonucleoprotein Q isoform 1 [Callithrix jacchus] | 0 | 104.4 | 33 | 19 | 119 | 12 | 1 | 624 | 69.7 | 8.59 | 1198 | 19 | 105955632.6 | 91894249.42 | 93188656.77 | 73968982.11 | 76687436.3 | 64632853.8 |
| 1060960222 | PREDICTED: double-stranded RNA-specific adenosine deaminase [Callithrix jacchus] | 0 | 27.337 | 6 | 7 | 21 | 7 | 1 | 1373 | 152 | 8.75 | 78 | 7 | 7727224.375 | 6712780.588 | 8178413.244 | 5837060.863 | 5152760 | 6224269.754 |
| 296209186 | PREDICTED: synaptobrevin homolog YKT6 [Callithrix jacchus] | 0 | 14.785 | 20 | 4 | 17 | 4 | 1 | 198 | 22.4 | 6.92 | 82 | 4 | 14577874.59 | 15653494.5 | 13530595.44 | 13323127.14 | 13842757.6 | 13856135.97 |
| 532442713 | 6-phosphofructokinase type C isoform 1 [Callithrix jacchus] | 0 | 220.6 | 45 | 31 | 253 | 28 | 1 | 784 | 85.4 | 7.28 | 3700 | 31 | 217608511.5 | 201014828.7 | 201647444.3 | 186049614 | 165891650.7 | 153034728.7 |
| 675704469 | PREDICTED: E3 ubiquitin-protein ligase CHIP isoform X1 [Callithrix jacchus] | 0 | 73.955 | 41 | 15 | 84 | 15 | 1 | 423 | 46.8 | 8.29 | 829 | 15 | 50679846.44 | 48445771.75 | 57140258.87 | 42764338.19 | 43300785.62 | 45774555.03 |
| 675737893 | PREDICTED: programmed cell death protein 10 [Callithrix jacchus] | 0 | 34.737 | 38 | 6 | 30 | 6 | 1 | 212 | 24.7 | 8.19 | 385 | 6 | 23784838.73 | 22900313 | 25797999.02 | 29453494.49 | 25834627.53 | 18136159.02 |
| 1061018449 | PREDICTED: LOW QUALITY PROTEIN: CAP-Gly domain-containing linker protein 1 [Callithrix jacchus] | 0 | 86.9 | 15 | 20 | 90 | 13 | 1 | 1440 | 162.4 | 5.38 | 397 | 20 | 22171024.38 | 21069278.5 | 11059897.22 | 10753887.55 | 13375251.94 | 32967795.55 |
| 675656277 | PREDICTED: destrin isoform X1 [Callithrix jacchus] | 0 | 50.367 | 60 | 10 | 111 | 9 | 1 | 165 | 18.5 | 7.85 | 1064 | 10 | 426805033 | 421943389.6 | 371303437.5 | 551375079.4 | 502366083.1 | 429412107.2 |
| 296212914 | PREDICTED: vacuolar protein sorting-associated protein 29 isoform X3 [Callithrix jacchus] | 0 | 50.808 | 48 | 8 | 59 | 8 | 1 | 182 | 20.5 | 6.79 | 864 | 8 | 49627269.97 | 50734238.88 | 50558408.34 | 50030296.07 | 47101901.15 | 49167444.21 |
| 675657404 | PREDICTED: proteasomal ubiquitin receptor ADRM1 isoform X1 [Callithrix jacchus] | 0 | 17.042 | 12 | 4 | 29 | 4 | 1 | 432 | 44.5 | 5.15 | 387 | 4 | 5797326.092 | 5386911.906 | 6942355.847 | 7889496.009 | 7311042.742 | 6506215.024 |
| 296191935 | PREDICTED: E3 ubiquitin-protein ligase RBX1 [Callithrix jacchus] | 0 | 24.648 | 25 | 3 | 35 | 3 | 1 | 108 | 12.3 | 6.96 | 843 | 3 | 2782724.498 | 3247389.375 | 3153232.993 | 3406369.478 | 3451678.367 | 3171655.18 |
| 390462005 | PREDICTED: serine incorporator 1 [Callithrix jacchus] | 0 | 20.083 | 9 | 3 | 14 | 3 | 1 | 453 | 50.5 | 5.85 | 167 | 3 | 7178601.252 | 8076178.75 | 7911719.633 | 4391100.158 | 4822947.231 | 6770770.503 |
| 390459074 | PREDICTED: actin-related protein 2/3 complex subunit 1A isoform X2 [Callithrix jacchus] | 0 | 176.876 | 72 | 20 | 189 | 19 | 1 | 370 | 41.6 | 8.18 | 3162 | 20 | 359434212.1 | 363965052.9 | 360913913.8 | 344246370.8 | 362749386.4 | 382576680.9 |
| 675690326 | PREDICTED: actin-related protein 10 [Callithrix jacchus] | 0 | 119.756 | 53 | 18 | 105 | 17 | 1 | 417 | 46.3 | 7.62 | 1249 | 18 | 71590124.31 | 70842905.74 | 62455654.2 | 53654359.78 | 58557579 | 52288974.49 |
| 296231010 | PREDICTED: dnaJ homolog subfamily A member 2 [Callithrix jacchus] | 0 | 181.672 | 59 | 22 | 206 | 22 | 1 | 412 | 45.7 | 6.48 | 2594 | 22 | 263586513.9 | 271986276 | 256731384.1 | 222923251.8 | 216770141.8 | 267373867.1 |
| 675720323 | REDICTED: succinyl-CoA ligase [ADP/GDP-forming] subunit alpha, mitochondrial isoform X1 [Callithrix jacchu | 0 | 32.213 | 20 | 5 | 24 | 5 | 1 | 370 | 38.6 | 8.76 | 461 | 5 | 15601458.55 | 14239538 | 12582751.45 | 13355247.53 | 15264773.13 | 21878352.29 |
| 675684070 | PREDICTED: dynein light chain 1, cytoplasmic [Callithrix jacchus] | 0 | 81.796 | 60 | 6 | 79 | 3 | 1 | 89 | 10.4 | 7.4 | 1693 | 6 | 434486509.5 | 418483038.9 | 473327492.1 | 274261010.4 | 348964350.8 | 479597712.3 |
| 296198039 | PREDICTED: mitochondrial carrier homolog 1 isoform X2 [Callithrix jacchus] | 0 | 133.893 | 56 | 18 | 209 | 18 | 1 | 389 | 41.5 | 9.32 | 2821 | 18 | 225301699.8 | 236519614.5 | 226828297.3 | 209083345.1 | 199689386.2 | 173830145 |
| 390467193 | PREDICTED: staphylococcal nuclease domain-containing protein 1 [Callithrix jacchus] | 0 | 338.867 | 59 | 41 | 403 | 41 | 1 | 910 | 102 | 7.17 | 6727 | 41 | 434291424.6 | 448964827.6 | 438099889.8 | 417909792 | 445135865 | 448153831.1 |
| 675680920 | PREDICTED: serine/threonine-protein kinase TBK1 isoform X1 [Callithrix jacchus] | 0 | 31.779 | 14 | 9 | 21 | 9 | 1 | 740 | 84.8 | 7.09 | 54 | 9 | 17161592.16 | 17691735.5 | 16365467.3 | 7563821.152 | 10917352.19 | 10602088.24 |
| 1060993710 | PREDICTED: vesicle-associated membrane protein-associated protein B/C [Callithrix jacchus] | 0 | 101.806 | 57 | 13 | 139 | 12 | 1 | 243 | 27.2 | 7.3 | 1660 | 13 | 179158582.3 | 156352737.3 | 174220152.5 | 158707039.6 | 165397749.6 | 137108255.9 |
| 675658892 | PREDICTED: Golgi SNAP receptor complex member 2 isoform X1 [Callithrix jacchus] | 0 | 13.084 | 12 | 2 | 10 | 2 | 1 | 264 | 30.3 | 9.83 | 43 | 2 | 2292529.205 | 2411818.5 | 1924218.876 | 1925083.768 | 2199359.197 | 1692297.911 |
| 296233029 | PREDICTED: ATPase ASNA1 [Callithrix jacchus] | 0 | 32.95 | 26 | 7 | 44 | 7 | 1 | 348 | 38.8 | 4.91 | 104 | 7 | 19310737.01 | 17030020.94 | 16393436.51 | 21549439.37 | 13675844.06 | 15990004.73 |
| 296225998 | PREDICTED: ruvB-like 1 [Callithrix jacchus] | 0 | 64.605 | 36 | 13 | 92 | 13 | 1 | 456 | 50.2 | 6.42 | 916 | 13 | 44468585.75 | 43011004.94 | 46001208.3 | 34366694.04 | 39442527.99 | 32914637.86 |
| 532515164 | rap guanine nucleotide exchange factor 4 isoform a [Callithrix jacchus] | 0 | 241.793 | 50 | 43 | 390 | 41 | 1 | 1011 | 115.4 | 6.84 | 5603 | 43 | 375121858.1 | 382748196.9 | 436507201.3 | 388307593.1 | 421162176.3 | 448886277.3 |
| 675672807 | PREDICTED: MAGUK p55 subfamily member 6 isoform X2 [Callithrix jacchus] | 0 | 77.941 | 45 | 19 | 105 | 18 | 1 | 540 | 61.1 | 6.39 | 860 | 19 | 53304313.87 | 62065217.5 | 59356396.83 | 59878410.8 | 61601826.43 | 54669688.81 |
| 532290566 | septin-6 isoform B [Callithrix jacchus] | 0 | 367.069 | 65 | 32 | 575 | 18 | 1 | 434 | 49.6 | 6.67 | 11152 | 32 | 836356635.3 | 870860810.5 | 852601559.8 | 934485282.9 | 962546360.9 | 859011570.7 |
| 675637173 | PREDICTED: intracellular hyaluronan-binding protein 4 isoform X1 [Callithrix jacchus] | 0 | 18.876 | 17 | 5 | 9 | 5 | 1 | 401 | 44.3 | 7.96 | 59 | 5 | 2103779.77 | 1956445.313 | 1573848.385 | 572696.568 | 861429.0957 | 801446.4632 |
| 296208841 | PREDICTED: potassium voltage-gated channel subfamily D member 3 isoform X1 [Callithrix jacchus] | 0 | 43.873 | 18 | 9 | 22 | 8 | 1 | 655 | 73.4 | 8.25 | 206 | 9 | 13234985.89 | 12676103.69 | 13765769.2 | 17412320.8 | 15598971.44 | 8740430.11 |
| 1060955511 | PREDICTED: glycogen synthase kinase-3 beta isoform X1 [Callithrix jacchus] | 0 | 108.647 | 53 | 16 | 147 | 12 | 1 | 433 | 48 | 8.78 | 1906 | 16 | 84880618.78 | 82779911.44 | 79075002.71 | 82197487.78 | 90216383.73 | 83062647.76 |
| 532288912 | cdc42 effector protein 4 [Callithrix jacchus] | 0 | 91.833 | 41 | 11 | 85 | 11 | 1 | 355 | 38.1 | 5.19 | 1910 | 11 | 35124695.26 | 33990549.38 | 38107864.01 | 54570931.36 | 37571321.76 | 40651014.61 |
| 675708892 | PREDICTED: dnaJ homolog subfamily B member 12 isoform X2 [Callithrix jacchus] | 0 | 23.021 | 17 | 5 | 13 | 5 | 1 | 410 | 45.5 | 9.28 | 43 | 5 | 11330227.15 | 9123398.25 | 9822610.528 | 7377256.264 | 4481198.138 | 5950670.682 |
| 296190683 | PREDICTED: BMP/retinoic acid-inducible neural-specific protein 1 [Callithrix jacchus] | 0 | 86.139 | 23 | 19 | 116 | 18 | 1 | 761 | 88.7 | 8.87 | 795 | 19 | 96407054.36 | 98503133.92 | 84227161.22 | 72795854.73 | 69982918.8 | 71136317.72 |
| 1060945476 | PREDICTED: target of rapamycin complex subunit LST8 [Callithrix jacchus] | 0 | 29.06 | 20 | 5 | 28 | 5 | 1 | 326 | 35.9 | 5.86 | 53 | 5 | 11533496.75 | 10656426 | 4669173.35 | 5585388.412 | 9703103.498 | 9417684.597 |
| 532506415 | serine/threonine-protein kinase mTOR [Callithrix jacchus] | 0 | 150.828 | 15 | 30 | 129 | 30 | 1 | 2549 | 288.8 | 7.17 | 849 | 30 | 31624531.41 | 33371881 | 29075082.06 | 38954557.38 | 39160184.91 | 31486429.03 |
| 532341723 | 4-trimethylaminobutyraldehyde dehydrogenase [Callithrix jacchus] | 0.001 | 6.732 | 6 | 3 | 7 | 3 | 1 | 518 | 56.4 | 7.18 | 16 | 3 | 1968743.944 | 2152279.984 | 2051345.476 | 3287996.581 | 3469683.291 | 3947292.016 |
| 1060957854 | PREDICTED: muscleblind-like protein 1 isoform X1 [Callithrix jacchus] | 0 | 8.027 | 7 | 3 | 12 | 3 | 1 | 399 | 42.9 | 8.82 | 42 | 3 | 12272531.88 | 11330821.25 | 9725514.599 | 9262746.024 | 10955997.17 | 9095213.02 |
| 532519156 | ral GTPase-activating protein subunit alpha-1 isoform 2 [Callithrix jacchus] | 0 | 110.168 | 14 | 27 | 120 | 26 | 1 | 2084 | 234.7 | 6.23 | 1111 | 27 | 63229670.19 | 54237875.45 | 53369105.63 | 49493107.44 | 61669646.79 | 52207552 |
| 1060940872 | PREDICTED: secretory carrier-associated membrane protein 5 [Callithrix jacchus] | 0 | 22.88 | 23 | 4 | 20 | 4 | 1 | 359 | 39 | 9.96 | 142 | 4 | 18659004.33 | 18204389.25 | 14593751.14 | 17865302.58 | 23049635.14 | 19448737.67 |
| 166831537 | DnaJ homolog, subfamily B, member 2 isoform b (predicted) [Callithrix jacchus] | 0 | 72.717 | 39 | 10 | 98 | 9 | 1 | 324 | 35.4 | 5.95 | 1310 | 10 | 45668077.05 | 49509136.78 | 51218097.34 | 29438788.35 | 28499255.25 | 41123469.6 |

| | | | | | | | | | | | |
|---|---|---|---|---|---|---|---|---|---|---|---|
| 391852756 | SH3 and multiple ankyrin repeat domains protein 3 [Callithrix jacchus] | 0 | 593.221 | 52 | 66 | 675 | 64 | 1 | 1731 | 184.8 | 8.88 |
| 296229617 | PREDICTED: syntaxin-6 [Callithrix jacchus] | 0 | 25.633 | 29 | 6 | 17 | 6 | 1 | 255 | 29.1 | 4.93 |
| 1060944170 | PREDICTED: acidic fibroblast growth factor intracellular-binding protein isoform X1 [Callithrix jacchus] | 0 | 49.914 | 34 | 11 | 45 | 11 | 1 | 364 | 41.9 | 6.61 |
| 675725372 | PREDICTED: PEX5-related protein isoform X6 [Callithrix jacchus] | 0 | 61.146 | 30 | 14 | 46 | 14 | 1 | 602 | 66.9 | 5.4 |
| 296231418 | PREDICTED: sphingomyelin phosphodiesterase 3 [Callithrix jacchus] | 0 | 77.788 | 27 | 14 | 102 | 14 | 1 | 655 | 71.2 | 5.97 |
| 532522830 | rho GTPase-activating protein 23 [Callithrix jacchus] | 0 | 92.399 | 17 | 21 | 107 | 20 | 1 | 1488 | 162 | 8.95 |
| 296199262 | PREDICTED: DBH-like monooxygenase protein 1 [Callithrix jacchus] | 0 | 110.107 | 35 | 18 | 135 | 18 | 1 | 612 | 69.6 | 6.58 |
| 1060963791 | PREDICTED: heterogeneous nuclear ribonucleoprotein H isoform X1 [Callithrix jacchus] | 0 | 111.176 | 35 | 12 | 130 | 6 | 1 | 472 | 51.2 | 6.8 |
| 296226560 | PREDICTED: ras-related protein Rab-2A isoform X1 [Callithrix jacchus] | 0 | 61.637 | 67 | 12 | 55 | 12 | 1 | 212 | 23.5 | 6.54 |
| 1060950629 | PREDICTED: ubiquitin carboxyl-terminal hydrolase 14 isoform X1 [Callithrix jacchus] | 0 | 62.686 | 27 | 12 | 54 | 12 | 1 | 492 | 55.7 | 5.3 |
| 1061019501 | PREDICTED: E3 ubiquitin-protein ligase HUWE1 [Callithrix jacchus] | 0 | 126.099 | 9 | 29 | 79 | 29 | 1 | 4374 | 481.5 | 5.21 |
| 675664073 | PREDICTED: 26S proteasome non-ATPase regulatory subunit 14 [Callithrix jacchus] | 0 | 16.35 | 14 | 4 | 19 | 4 | 1 | 310 | 34.6 | 6.52 |
| 675659602 | PREDICTED: mitochondrial Rho GTPase 1 isoform X1 [Callithrix jacchus] | 0 | 85.32 | 19 | 12 | 69 | 11 | 1 | 752 | 86.3 | 6.67 |
| 1061007545 | PREDICTED: serine/threonine-protein kinase 25 [Callithrix jacchus] | 0 | 27.972 | 19 | 7 | 23 | 5 | 1 | 426 | 48.1 | 6.74 |
| 296194653 | PREDICTED: 28S ribosomal protein S30, mitochondrial [Callithrix jacchus] | 0 | 32.839 | 20 | 6 | 12 | 6 | 1 | 432 | 49.5 | 8.84 |
| 675743370 | PREDICTED: tropomyosin alpha-3 chain isoform X3 [Callithrix jacchus] | 0 | 124.772 | 52 | 23 | 307 | 7 | 1 | 285 | 33.2 | 4.77 |
| 1060965314 | PREDICTED: LOW QUALITY PROTEIN: protein LAP2 [Callithrix jacchus] | 0 | 58.526 | 10 | 12 | 44 | 9 | 1 | 1459 | 163.4 | 5.55 |
| 1060941520 | PREDICTED: FXYD domain-containing ion transport regulator 6 isoform X1 [Callithrix jacchus] | 0 | 58.222 | 51 | 5 | 58 | 5 | 1 | 95 | 10.5 | 5.1 |
| 296219778 | PREDICTED: dynactin subunit 5 isoform X1 [Callithrix jacchus] | 0 | 20.298 | 26 | 5 | 38 | 5 | 1 | 182 | 20.1 | 8.02 |
| 296228547 | PREDICTED: RNA-binding protein 8A isoform X1 [Callithrix jacchus] | 0 | 11.035 | 16 | 2 | 7 | 2 | 1 | 174 | 19.9 | 5.72 |
| 675661343 | PREDICTED: 39S ribosomal protein L38, mitochondrial [Callithrix jacchus] | 0 | 37.29 | 20 | 6 | 31 | 6 | 1 | 380 | 44.9 | 7.59 |
| 532512896 | alpha-actinin-4 [Callithrix jacchus] | 0 | 648.738 | 74 | 66 | 1069 | 5 | 1 | 911 | 104.8 | 5.44 |
| 296203470 | PREDICTED: fructosamine-3-kinase [Callithrix jacchus] | 0 | 25.351 | 21 | 5 | 29 | 5 | 1 | 309 | 35.2 | 8.4 |
| 1060939723 | PREDICTED: glia maturation factor beta [Callithrix jacchus] | 0 | 32.279 | 42 | 4 | 25 | 4 | 1 | 142 | 16.7 | 5.29 |
| 675654870 | PREDICTED: nesprin-1 isoform X7 [Callithrix jacchus] | 0 | 1105.597 | 29 | 216 | 1011 | 216 | 1 | 8809 | 1011.5 | 5.59 |
| 532501762 | neurogranin [Callithrix jacchus] | 0 | 19.95 | 60 | 3 | 15 | 3 | 1 | 78 | 7.6 | 7.87 |
| 532523094 | synaptic vesicle glycoprotein 2A [Callithrix jacchus] | 0 | 81.881 | 21 | 12 | 113 | 12 | 1 | 745 | 83.1 | 5.69 |
| 675664642 | PREDICTED: ras-related protein Ral-B [Callithrix jacchus] | 0 | 112.9 | 53 | 10 | 103 | 4 | 1 | 206 | 23.4 | 6.62 |
| 1061002775 | PREDICTED: LOW QUALITY PROTEIN: synaptic vesicle glycoprotein 2B [Callithrix jacchus] | 0 | 33.64 | 12 | 7 | 35 | 7 | 1 | 683 | 77.3 | 5.44 |
| 1061007111 | PREDICTED: integral membrane protein 2C isoform X1 [Callithrix jacchus] | 0 | 34.473 | 33 | 6 | 22 | 6 | 1 | 267 | 30.2 | 8.41 |
| 675706581 | PREDICTED: G-protein coupled receptor family C group 5 member B [Callithrix jacchus] | 0 | 35.878 | 13 | 4 | 30 | 4 | 1 | 403 | 44.7 | 8.07 |
| 532517424 | NAD-dependent protein deacetylase sirtuin-2 isoform 1 [Callithrix jacchus] | 0 | 181.76 | 65 | 21 | 158 | 21 | 1 | 389 | 43.6 | 5.54 |
| 1060969400 | PREDICTED: LOW QUALITY PROTEIN: disco-interacting protein 2 homolog A [Callithrix jacchus] | 0 | 40.194 | 9 | 12 | 28 | 9 | 1 | 1581 | 171.5 | 7.99 |
| 675739344 | PREDICTED: 28S ribosomal protein S22, mitochondrial isoform X1 [Callithrix jacchus] | 0 | 65.167 | 41 | 15 | 62 | 15 | 1 | 360 | 41.4 | 6.64 |
| 296202391 | PREDICTED: 28S ribosomal protein S23, mitochondrial isoform X1 [Callithrix jacchus] | 0 | 36.268 | 50 | 8 | 48 | 8 | 1 | 189 | 21.7 | 8.9 |
| 296229727 | PREDICTED: 28S ribosomal protein S14, mitochondrial [Callithrix jacchus] | 0 | 18.05 | 29 | 3 | 13 | 3 | 1 | 128 | 15.1 | 11.3 |
| 1060951757 | PREDICTED: serine protease HTRA2, mitochondrial [Callithrix jacchus] | 0 | 45.854 | 23 | 8 | 36 | 8 | 1 | 509 | 54.9 | 9.85 |
| 532513274 | hamartin isoform 3 [Callithrix jacchus] | 0 | 154.694 | 31 | 27 | 107 | 27 | 1 | 1168 | 130.2 | 6.52 |

[illegible]

| | | | | | | | | | | | | | | | | | | | | | | | | | | | | | | | | | | | | | | | | | | |
|---|---|---|---|---|---|---|---|---|---|---|---|---|---|---|---|---|---|---|---|---|---|---|---|---|---|---|---|---|---|---|---|---|---|---|---|---|---|---|---|---|---|---|
| 532524114 | reticulon-4 isoform A [Callithrix jacchus] | 0 | 226.486 | 35 | 31 | 262 | 26 | 1 | 1194 | 129.6 | 4.56 | 3651 | 31 | 153118043 | 149924776.4 | 180066041 | 146883704.9 | 140012998.3 | 147185198.7 | 135174144.8 | 119983212.8 | 141452232.7 | 151345992 | 112062984.7 | 146357630.2 | 10.28179 | 2.8036 | 8.349685 | 15.65719 | 100402774 | RTN4 | 68585 | Rtn4 | 1.12 | 1.01 | 0.92 | 0.95 | 0.3056 | 0.352971736 | 1.218097 | 2 | No |
| 1060945306 | PREDICTED: LOW QUALITY PROTEIN: 39S ribosomal protein L28, mitochondrial [Callithrix jacchus] | 0 | 33.772 | 30 | 9 | 33 | 9 | 1 | 276 | 32.6 | 9.28 | 71 | 9 | 14867130.08 | 16410378.16 | 11427386.93 | 16187298.7 | 16031980.54 | 14373326.13 | 9389619.698 | 9795394.003 | 7171546.857 | 25333407.38 | 14426420.99 | 14824624.71 | 17.92021 | 6.473976 | 16.07634 | 33.99542 | 100393090 | MRPL28 | 68611 | Mrpl28 | 1.00 | 1.09 | 0.62 | 1.28 | 0.0173 | 0.046646804 | 2.071001 | 1 | Yes |
| 296211668 | PREDICTED: LETM1 domain-containing protein 1 isoform X1 [Callithrix jacchus] | 0 | 31.15 | 18 | 6 | 24 | 6 | 1 | 360 | 41.9 | 10.2 | 130 | 6 | 16761753.87 | 15961035.75 | 19198648.11 | 14706515.18 | 16003885.41 | 12154710.76 | 11242588.25 | 13016198.89 | 15364594.56 | 6967176.655 | 4726848.364 | 8346127.663 | 9.7434 | 13.70596 | 15.65493 | 27.34475 | 100385608 | LOC100385608 | 68614 | Letmd1 | 1.34 | 1.11 | 1.03 | 0.52 | 0.009548 | 0.032637302 | 2.59087 | 2 | Yes |
| 296218531 | PREDICTED: heterogeneous nuclear ribonucleoprotein U-like protein 2 isoform X1 [Callithrix jacchus] | 0 | 42.643 | 15 | 8 | 20 | 8 | 1 | 749 | 85 | 4.88 | 132 | 8 | 13565795.62 | 13940855.75 | 13954886.82 | 12897880.84 | 8335242.034 | 7590473.14 | 14093398.31 | 13767191.84 | 12696569.83 | 17886049.42 | 15760297.78 | 16472616.12 | 1.596924 | 29.90746 | 5.405164 | 6.476437 | 100397549 | HNRNPUL2 | 68693 | Hnrnpul2 | 1.03 | 0.72 | 1.01 | 1.25 | 0.0562 | 0.100033459 | 1.738817 | 1 | No |
| 532289436 | leucine-rich repeat-containing protein 16A isoform 1 [Callithrix jacchus] | 0 | 35.466 | 8 | 9 | 24 | 9 | 1 | 1375 | 151.6 | 7.85 | 120 | 9 | 11703918.53 | 11072079.63 | 19587614.18 | 14278267.27 | 13487986.5 | 13370926.22 | 13283621.31 | 13703133.89 | 14328593.97 | 12525272.52 | 7286115.079 | 18923054.31 | 33.59897 | 3.599256 | 3.818372 | 45.1387 | 100407163 | LRRC16A | 68732 | Carmil1 | 1.04 | 1.01 | 1.01 | 0.95 | 0.9938 | 0.994220923 | 1.093694 | 2 | No |
| [illegible] | | | | | | | | | | | | | | | | | | | | | | | | | | | | | | | | | | | | | | | | | | |

| | | | | | | | | | | | |
|---|---|---|---|---|---|---|---|---|---|---|---|
| 296196805 | PREDICTED: Kv channel-interacting protein 4 isoform X1 [Callithrix jacchus] | 0 | 8.321 | 9 | 2 | 5 | 2 | 1 | 250 | 28.7 | [illegible] |
| 675700229 | PREDICTED: protein kinase C and casein kinase substrate in neurons protein 3 [Callithrix jacchus] | 0 | 86.879 | 34 | 12 | 78 | 12 | 1 | 424 | 48.6 | [illegible] |
| 1060991921 | PREDICTED: vacuolar protein sorting-associated protein 16 homolog [Callithrix jacchus] | 0 | 97.481 | 30 | 20 | 99 | 20 | 1 | 839 | 94.6 | [illegible] |
| 1060977233 | EDICTED: leucine-rich repeat and fibronectin type III domain-containing protein 1 isoform X1 [Callithrix jacch | 0 | 20.147 | 5 | 3 | 7 | 3 | 1 | 837 | 89.7 | [illegible] |
| 1061012571 | PREDICTED: netrin-G1 isoform X4 [Callithrix jacchus] | 0 | 51.323 | 17 | 8 | 56 | 8 | 1 | 581 | 65.1 | [illegible] |
| 675647508 | PREDICTED: tripartite motif-containing protein 2 isoform X6 [Callithrix jacchus] | 0 | 169.731 | 39 | 21 | 122 | 20 | 1 | 748 | 81.9 | [illegible] |
| 675685168 | PREDICTED: serine beta-lactamase-like protein LACTB, mitochondrial [Callithrix jacchus] | 0 | 99.178 | 34 | 15 | 79 | 15 | 1 | 549 | 60.6 | [illegible] |
| 1060977693 | PREDICTED: LOW QUALITY PROTEIN: peroxisomal acyl-coenzyme A oxidase 3 [Callithrix jacchus] | 0 | 15.659 | 7 | 4 | 8 | 4 | 1 | 737 | 81.1 | [illegible] |
| 675727385 | PREDICTED: NCK-interacting protein with SH3 domain isoform X1 [Callithrix jacchus] | 0 | 128.403 | 33 | 21 | 158 | 21 | 1 | 714 | 78.1 | [illegible] |
| 532516526 | dnaJ homolog subfamily B member 1 [Callithrix jacchus] | 0 | 50.201 | 30 | 12 | 73 | 12 | 1 | 340 | 38.1 | [illegible] |
| 675763886 | PREDICTED: voltage-dependent calcium channel gamma-7 subunit [Callithrix jacchus] | 0 | 21.814 | 17 | 3 | 20 | 3 | 1 | 306 | 34.1 | [illegible] |
| 1060971370 | PREDICTED: voltage-dependent calcium channel gamma-8 subunit, partial [Callithrix jacchus] | 0 | 25.839 | 13 | 5 | 32 | 5 | 1 | 365 | 38.3 | [illegible] |
| 675656283 | PREDICTED: LOW QUALITY PROTEIN: ribosome-binding protein 1 [Callithrix jacchus] | 0 | 229.457 | 37 | 42 | 262 | 41 | 1 | 1382 | 149.9 | [illegible] |
| 532523870 | A-kinase anchor protein 12 isoform 1 [Callithrix jacchus] | 0 | 147.222 | 21 | 23 | 96 | 23 | 1 | 1771 | 189.7 | [illegible] |
| 296229144 | PREDICTED: ragulator complex protein LAMTOR2 isoform X1 [Callithrix jacchus] | 0 | 15.176 | 36 | 4 | 12 | 4 | 1 | 125 | 13.5 | [illegible] |
| 1060994952 | PREDICTED: nuclear distribution protein nudE-like 1 isoform X2 [Callithrix jacchus] | 0 | 44.546 | 27 | 9 | 34 | 6 | 1 | 379 | 42.3 | [illegible] |
| 675727413 | PREDICTED: WD repeat-containing protein 6 isoform X1 [Callithrix jacchus] | 0 | 7.054 | 3 | 3 | 10 | 3 | 1 | 1151 | 124.8 | [illegible] |
| 390473012 | PREDICTED: metal transporter CNNM1 isoform X1 [Callithrix jacchus] | 0 | 12.466 | 4 | 4 | 10 | 4 | 1 | 972 | 106.3 | [illegible] |
| 1060962615 | PREDICTED: actin-like protein 6B isoform X1 [Callithrix jacchus] | 0 | 19.608 | 8 | 3 | 19 | 3 | 1 | 439 | 47.8 | [illegible] |
| 296198979 | PREDICTED: wiskott-Aldrich syndrome protein family member 1 [Callithrix jacchus] | 0 | 140.166 | 31 | 18 | 228 | 15 | 1 | 559 | 61.6 | [illegible] |
| 296222725 | PREDICTED: E3 ubiquitin-protein ligase NEDD4-like isoform X10 [Callithrix jacchus] | 0 | 107.153 | 30 | 25 | 117 | 21 | 1 | 975 | 112 | [illegible] |
| 675705692 | PREDICTED: dnaJ homolog subfamily A member 3, mitochondrial isoform X1 [Callithrix jacchus] | 0 | 112.1 | 33 | 15 | 93 | 15 | 1 | 480 | 52.5 | [illegible] |
| 1060941909 | PREDICTED: junctional adhesion molecule C [Callithrix jacchus] | 0 | 16.595 | 15 | 5 | 18 | 5 | 1 | 333 | 37.3 | [illegible] |
| 675728894 | PREDICTED: sarcolemmal membrane-associated protein isoform X1 [Callithrix jacchus] | 0 | 40.305 | 13 | 9 | 20 | 9 | 1 | 847 | 97.2 | [illegible] |
| 675752861 | PREDICTED: lysine--tRNA ligase isoform X2 [Callithrix jacchus] | 0 | 82.027 | 31 | 18 | 85 | 18 | 1 | 597 | 68 | [illegible] |
| 296193209 | PREDICTED: casein kinase I isoform X3 [Callithrix jacchus] | 0 | 74.485 | 42 | 13 | 70 | 13 | 1 | 337 | 38.9 | [illegible] |
| 675645303 | PREDICTED: glutaredoxin-1 [Callithrix jacchus] | 0 | 11.816 | 31 | 3 | 12 | 3 | 1 | 106 | 11.7 | [illegible] |
| 532216943 | protocadherin gamma-C3 isoform 1 precursor [Callithrix jacchus] | 0 | 41.402 | 14 | 12 | 48 | 10 | 1 | 936 | 101.4 | [illegible] |
| 532292102 | protocadherin gamma-C5 isoform 1 precursor [Callithrix jacchus] | 0 | 49.897 | 15 | 10 | 39 | 8 | 1 | 944 | 101.9 | [illegible] |
| 675752849 | PREDICTED: gamma-aminobutyric acid receptor-associated protein-like 2 [Callithrix jacchus] | 0 | 36.392 | 47 | 7 | 74 | 6 | 1 | 117 | 13.7 | [illegible] |
| 296221519 | PREDICTED: enoyl-CoA hydratase, mitochondrial [Callithrix jacchus] | 0 | 25.662 | 26 | 5 | 16 | 5 | 1 | 290 | 31.5 | [illegible] |
| 532280108 | calmin [Callithrix jacchus] | 0 | 39.11 | 12 | 11 | 38 | 11 | 1 | 1005 | 111.9 | [illegible] |
| 675640342 | PREDICTED: LOW QUALITY PROTEIN: cat eye syndrome critical region protein 6 [Callithrix jacchus] | 0 | 10.786 | 6 | 3 | 20 | 3 | 1 | 575 | 58.8 | [illegible] |
| 675648899 | PREDICTED: 39S ribosomal protein L1, mitochondrial isoform X1 [Callithrix jacchus] | 0 | 34.648 | 24 | 7 | 19 | 7 | 1 | 325 | 36.7 | [illegible] |
| 296218373 | PREDICTED: 39S ribosomal protein L16, mitochondrial [Callithrix jacchus] | 0 | 39.873 | 33 | 7 | 37 | 7 | 1 | 251 | 28.4 | [illegible] |
| 296202488 | PREDICTED: 39S ribosomal protein L27, mitochondrial isoform X1 [Callithrix jacchus] | 0 | 34.84 | 50 | 6 | 26 | 6 | 1 | 148 | 16 | [illegible] |
| 675710921 | PREDICTED: 39S ribosomal protein L43, mitochondrial [Callithrix jacchus] | 0 | 20.085 | 30 | 5 | 21 | 5 | 1 | 159 | 17.9 | [illegible] |
| 1060959825 | PREDICTED: long-chain-fatty-acid--CoA ligase ACSBG1 isoform X2 [Callithrix jacchus] | 0 | 15.925 | 8 | 5 | 17 | 5 | 1 | 787 | 87.1 | [illegible] |
| 296226791 | PREDICTED: phosphoprotein associated with glycosphingolipid-enriched microdomains 1 [Callithrix jacchus] | 0 | 80.226 | 33 | 10 | 81 | 10 | 1 | 432 | 47 | [illegible] |
| 296199103 | PREDICTED: Golgi-associated PDZ and coiled-coil motif-containing protein isoform X1 [Callithrix jacchus] | 0 | 45.19 | 25 | 10 | 33 | 10 | 1 | 463 | 50.7 | [illegible] |
| [illegible] | [illegible] | [illegible] | [illegible] | [illegible] | [illegible] | [illegible] | [illegible] | [illegible] | [illegible] | [illegible] | [illegible] |

| | | | | | | | | | | | | | | | | |
|---|---|---|---|---|---|---|---|---|---|---|---|---|---|---|---|---|
| 296230768 | PREDICTED: putative transferase CAF17, mitochondrial isoform X1 [Callithrix jacchus] | 0 | 20.576 | 20 | 7 | 26 | 7 | 1 | 357 | 38.6 | 9.85 | 175 | 7 | 8256396.991 | 8813603.563 | [illegible] |
| 532331655 | transmembrane protein 11, mitochondrial [Callithrix jacchus] | 0 | 22.44 | 28 | 5 | 35 | 5 | 1 | 192 | 21.5 | 7.65 | 230 | 5 | 65285073.71 | 64366978.5 | [illegible] |
| 296201221 | PREDICTED: rho GTPase-activating protein 44 isoform X1 [Callithrix jacchus] | 0 | 208.587 | 40 | 28 | 218 | 28 | 1 | 814 | 88.8 | 6.55 | 3144 | 28 | 185746210 | 204905452 | [illegible] |
| 1060995176 | PREDICTED: LOW QUALITY PROTEIN: neuroligin-2 [Callithrix jacchus] | 0 | 88.598 | 27 | 15 | 92 | 11 | 1 | 760 | 83.1 | 6.32 | 1108 | 15 | 30079681.41 | 28993428.47 | [illegible] |
| 675659914 | PREDICTED: beta-arrestin-2 isoform X1 [Callithrix jacchus] | 0 | 11.453 | 9 | 3 | 10 | 2 | 1 | 440 | 49.3 | 8.85 | 19 | 3 | 2294668.175 | 2895092.875 | [illegible] |
| 532450019 | ARF GTPase-activating protein GIT1 isoform 1 [Callithrix jacchus] | 0 | 240.703 | 58 | 33 | 287 | 30 | 1 | 770 | 85.4 | 7.01 | 4101 | 33 | 242423957.3 | 232898399.9 | [illegible] |
| 532510456 | serine/threonine-protein kinase TAO1 isoform 1 [Callithrix jacchus] | 0 | 281.925 | 39 | 41 | 359 | 34 | 1 | 1001 | 116.1 | 7.65 | 4527 | 41 | 159404267.7 | 147288857.9 | [illegible] |
| 675660313 | PREDICTED: neurabin-2 isoform X1 [Callithrix jacchus] | 0 | 362.591 | 50 | 36 | 453 | 35 | 1 | 825 | 90.3 | 5 | 9260 | 36 | 659877403.5 | 678382673 | [illegible] |
| 675660311 | PREDICTED: sterile alpha motif domain-containing protein 14 [Callithrix jacchus] | 0 | 15.864 | 11 | 3 | 5 | 3 | 1 | 417 | 44.9 | 9.41 | 89 | 3 | 1913577.914 | 2067357.5 | [illegible] |
| 390463594 | CTED: LOW QUALITY PROTEIN: CDGSH iron-sulfur domain-containing protein 3, mitochondrial [Callithrix ja | 0 | 44.895 | 41 | 7 | 65 | 7 | 1 | 127 | 14.3 | 10.54 | 416 | 7 | 137493925 | 136319331 | [illegible] |
| 532515002 | protein FAM171A2 precursor [Callithrix jacchus] | 0 | 107.19 | 26 | 17 | 108 | 16 | 1 | 826 | 87.4 | 8.05 | 939 | 17 | 51947320.17 | 56611925 | [illegible] |
| 296203233 | PREDICTED: signal recognition particle subunit SRP68 isoform X1 [Callithrix jacchus] | 0 | 93.52 | 39 | 21 | 68 | 21 | 1 | 622 | 70.6 | 8.47 | 990 | 21 | 28856288.41 | 31719604.53 | [illegible] |
| 532214985 | ubiquitin-conjugating enzyme E2 O [Callithrix jacchus] | 0 | 75.627 | 14 | 14 | 69 | 14 | 1 | 1301 | 142.2 | 5.08 | 391 | 14 | 24380279.74 | 23635979.19 | [illegible] |
| 532328669 | syntaxin-binding protein 6 [Callithrix jacchus] | 0 | 33.405 | 38 | 7 | 33 | 7 | 1 | 210 | 23.5 | 9.04 | 224 | 7 | 10249049.52 | 8081005.188 | [illegible] |
| 1060940691 | PREDICTED: LOW QUALITY PROTEIN: serine/threonine-protein kinase MRCK beta [Callithrix jacchus] | 0 | 377.645 | 36 | 65 | 395 | 60 | 1 | 1715 | 194.4 | 6.38 | 3882 | 65 | 239306256.4 | 262151446.6 | [illegible] |
| 532215931 | protein KIAA0284 isoform 1 [Callithrix jacchus] | 0 | 253.864 | 40 | 48 | 253 | 7 | 1 | 1549 | 166.9 | 6.79 | 2562 | 48 | 30602717.53 | 46560660.75 | [illegible] |
| 532330999 | phosphofurin acidic cluster sorting protein 2 isoform 1 [Callithrix jacchus] | 0 | 28.447 | 8 | 6 | 25 | 6 | 1 | 900 | 99 | 6.95 | 245 | 6 | 8171967.87 | 8101437.5 | [illegible] |
| 296209103 | PREDICTED: vacuolar protein sorting-associated protein 41 homolog [Callithrix jacchus] | 0 | 53.774 | 17 | 14 | 73 | 14 | 1 | 854 | 98.4 | 5.67 | 168 | 14 | 35175142.74 | 36314611.95 | [illegible] |
| 532346601 | phosphatase and actin regulator 1 [Callithrix jacchus] | 0 | 132.085 | 38 | 21 | 166 | 20 | 1 | 580 | 66.3 | 6.96 | 1938 | 21 | 311343173.7 | 305107785.4 | [illegible] |
| 675637083 | PREDICTED: constitutive coactivator of PPAR-gamma-like protein 1 [Callithrix jacchus] | 0 | 197.17 | 33 | 27 | 204 | 26 | 1 | 1118 | 121.9 | 8.88 | 3711 | 27 | 112942935.7 | 103932565.9 | [illegible] |
| 675645740 | PREDICTED: F-BAR domain only protein 2 isoform X1 [Callithrix jacchus] | 0 | 26.779 | 8 | 5 | 19 | 5 | 1 | 840 | 92.1 | 6.24 | 179 | 5 | 3800286.145 | 3184751.375 | [illegible] |
| 1060946833 | PREDICTED: protein transport protein Sec24C isoform X1 [Callithrix jacchus] | 0 | 70.426 | 17 | 15 | 55 | 15 | 1 | 1118 | 120.9 | 6.87 | 828 | 15 | 23343916.75 | 22406996.25 | [illegible] |
| 296214371 | PREDICTED: type 1 phosphatidylinositol 4,5-bisphosphate 4-phosphatase isoform X1 [Callithrix jacchus] | 0 | 29.804 | 25 | 6 | 34 | 5 | 1 | 284 | 30.2 | 8.98 | 203 | 6 | 22902952.63 | 24644505.34 | [illegible] |
| 296203574 | PREDICTED: myotubularin-related protein 6 isoform X1 [Callithrix jacchus] | 0.008 | 3.361 | 3 | 2 | 9 | 2 | 1 | 621 | 72.1 | 7.65 | 40 | 2 | 967515.6298 | 1035896.063 | [illegible] |
| 1060948827 | PREDICTED: cell cycle and apoptosis regulator protein 2 isoform X1 [Callithrix jacchus] | 0 | 55.223 | 14 | 12 | 52 | 12 | 1 | 1037 | 115 | 5.47 | 308 | 12 | 59652104.49 | 45784640.25 | [illegible] |
| 675635888 | PREDICTED: FERM, RhoGEF and pleckstrin domain-containing protein 1 isoform X2 [Callithrix jacchus] | 0 | 237.26 | 38 | 32 | 262 | 32 | 1 | 1046 | 118.5 | 8.06 | 4590 | 32 | 228547879.5 | 223996025.6 | [illegible] |
| 390457484 | PREDICTED: LOW QUALITY PROTEIN: serine/threonine-protein kinase 24 [Callithrix jacchus] | 0 | 48.973 | 26 | 9 | 38 | 7 | 1 | 431 | 47.8 | 5.43 | 351 | 9 | 22558055.88 | 23406805.19 | [illegible] |
| 1060962330 | PREDICTED: LOW QUALITY PROTEIN: triple functional domain protein [Callithrix jacchus] | 0 | 437.195 | 29 | 77 | 424 | 59 | 1 | 3026 | 340.1 | 6.37 | 4511 | 77 | 224180047.6 | 225684788.4 | [illegible] |
| 532523988 | nuclear receptor-binding protein 2 [Callithrix jacchus] | 0 | 43.619 | 23 | 10 | 30 | 10 | 1 | 499 | 57.3 | 6.57 | 276 | 10 | 16643652.84 | 16771016.56 | [illegible] |
| 1060956787 | PREDICTED: rho GTPase-activating protein 39 isoform X2 [Callithrix jacchus] | 0 | 194.161 | 37 | 32 | 216 | 32 | 1 | 1083 | 121.2 | 7.5 | 2465 | 32 | 168031852.8 | 162770040.1 | [illegible] |
| 390458829 | PREDICTED: eukaryotic translation initiation factor 3 subunit L [Callithrix jacchus] | 0 | 88.212 | 34 | 16 | 81 | 16 | 1 | 564 | 66.6 | 6.34 | 1339 | 16 | 59266882.14 | 44142274.83 | [illegible] |
| 296191904 | PREDICTED: mitochondrial import receptor subunit TOM22 homolog [Callithrix jacchus] | 0 | 95.692 | 66 | 6 | 78 | 6 | 1 | 142 | 15.5 | 4.34 | 2704 | 6 | 43372148.61 | 42905173.88 | [illegible] |
| 296191996 | PREDICTED: malonyl-CoA-acyl carrier protein transacylase, mitochondrial isoform X1 [Callithrix jacchus] | 0 | 22.328 | 15 | 5 | 28 | 5 | 1 | 390 | 42.9 | 7.93 | 215 | 5 | 10015958.97 | 11165966.3 | [illegible] |
| 296211538 | PREDICTED: rho-related GTP-binding protein Rho6 [Callithrix jacchus] | 0 | 14.499 | 19 | 4 | 11 | 4 | 1 | 232 | 25.9 | 7.75 | 62 | 4 | 4248491.045 | 4503164.563 | [illegible] |
| 296211788 | PREDICTED: SPRY domain-containing protein 3 [Callithrix jacchus] | 0 | 11.783 | 7 | 3 | 8 | 3 | 1 | 442 | 49.7 | 6.16 | 65 | 3 | 3810741.089 | 3784581.125 | [illegible] |
| 1061016549 | REDICTED: LOW QUALITY PROTEIN: FYVE, RhoGEF and PH domain-containing protein 4 [Callithrix jacchu | 0 | 15.661 | 7 | 6 | 11 | 6 | 1 | 879 | 98.8 | 6.8 | 17 | 6 | 2707791.257 | 2523880.438 | [illegible] |
| 675640589 | PREDICTED: phosphatidylinositol 4-kinase alpha [Callithrix jacchus] | 0 | 424.561 | 39 | 76 | 530 | 76 | 1 | 2095 | 236.2 | 7.03 | 7358 | 76 | 312455660 | 295997007.8 | [illegible] |
| 1060930181 | PREDICTED: LOW QUALITY PROTEIN: kelch-like protein 22 [Callithrix jacchus] | 0 | 36.872 | 15 | 8 | 18 | 8 | 1 | 634 | 71.8 | 5.59 | 97 | 8 | 17080809.86 | 20335910.88 | [illegible] |
| 675704849 | PREDICTED: TBC1 domain family member 24 isoform X1 [Callithrix jacchus] | 0 | 135.602 | 40 | 20 | 160 | 20 | 1 | 559 | 62.8 | 7.52 | 3503 | 20 | 187659733.2 | 180142132.3 | [illegible] |
| 532329119 | LEM domain-containing protein 2 isoform 1 [Callithrix jacchus] | 0 | 15.486 | 10 | 5 | 14 | 5 | 1 | 503 | 56.9 | 9.1 | 83 | 5 | 2016224.069 | 1879336.25 | [illegible] |
| 296197960 | PREDICTED: UHRF1-binding protein 1 [Callithrix jacchus] | 0 | 17.744 | 5 | 6 | 13 | 6 | 1 | 1440 | 159.5 | 6.34 | 17 | 6 | 2408395.518 | 2120121.406 | [illegible] |
| 1060985495 | PREDICTED: ankyrin repeat and SAM domain-containing protein 1A [Callithrix jacchus] | 0 | 45.814 | 8 | 7 | 91 | 6 | 1 | 1153 | 125.5 | 6.46 | 2087 | 7 | 7413694.987 | 3389694.5 | [illegible] |
| 532510954 | vacuolar protein sorting-associated protein 52 homolog [Callithrix jacchus] | 0 | 46.335 | 19 | 11 | 38 | 11 | 1 | 723 | 82.2 | 5.91 | 285 | 11 | 13321905.25 | 9622133.188 | [illegible] |
| 1060984925 | PREDICTED: large proline-rich protein BAG6 isoform X1 [Callithrix jacchus] | 0 | 99.438 | 20 | 16 | 89 | 16 | 1 | 1177 | 124.1 | 5.69 | 1281 | 16 | 33842408.95 | 28845418 | [illegible] |
| 532444657 | ATP-binding cassette sub-family F member 1 isoform a [Callithrix jacchus] | 0 | 17.289 | 7 | 5 | 15 | 5 | 1 | 841 | 95.6 | 6.6 | 37 | 5 | 5021827.859 | 4458749.75 | [illegible] |
| 1060986182 | PREDICTED: CSC1-like protein 2 [Callithrix jacchus] | 0 | 56.155 | 17 | 10 | 47 | 10 | 1 | 819 | 93.1 | 7.3 | 417 | 10 | 33948460.75 | 32001854.38 | [illegible] |
| 296198257 | PREDICTED: leucine-rich repeat-containing protein 73 [Callithrix jacchus] | 0 | 64.886 | 34 | 6 | 51 | 6 | 1 | 316 | 33.4 | 4.89 | 1672 | 6 | 69967393.1 | 61992394.09 | [illegible] |
| 1060958455 | PREDICTED: inactive phospholipase C-like protein 2 isoform X1 [Callithrix jacchus] | 0 | 106.88 | 23 | 20 | 52 | 19 | 1 | 1157 | 129 | 6.98 | 489 | 20 | 9104450.748 | 12333597.58 | [illegible] |
| 532286380 | scaffold attachment factor B2 [Callithrix jacchus] | 0 | 21.94 | 8 | 5 | 8 | 5 | 1 | 953 | 108 | 5.67 | 65 | 5 | 14060288.23 | 14514454.13 | [illegible] |
| 1060971967 | PREDICTED: MICOS complex subunit MIC13 isoform X1 [Callithrix jacchus] | 0 | 47.884 | 43 | 3 | 16 | 3 | 1 | 171 | 18.3 | 9.66 | 619 | 3 | 1633878.044 | 1470536.625 | [illegible] |
| 532287860 | disks large-associated protein 1 isoform 3 [Callithrix jacchus] | 0 | 249.744 | 41 | 33 | 299 | 10 | 1 | 929 | 103.6 | 6.77 | 5941 | 33 | 60992912.34 | 58710395.63 | [illegible] |
| 296224047 | PREDICTED: serine/arginine-rich splicing factor 7 isoform X1 [Callithrix jacchus] | 0 | 36.055 | 22 | 6 | 42 | 5 | 1 | 238 | 27.4 | 11.82 | 324 | 6 | 19686479.66 | 17123934 | [illegible] |
| 296224030 | PREDICTED: mitogen-activated protein kinase kinase kinase kinase 3 isoform X1 [Callithrix jacchus] | 0 | 39.024 | 12 | 9 | 33 | 9 | 1 | 893 | 100.8 | 7.56 | 662 | 9 | 12405156.24 | 12503692.83 | [illegible] |
| 296222558 | PREDICTED: phosphatidylinositol 3-kinase catalytic subunit type 3 [Callithrix jacchus] | 0 | 25.743 | 7 | 5 | 31 | 5 | 1 | 887 | 101.5 | 6.81 | 141 | 5 | 2381057.936 | 19695255.25 | [illegible] |
| 296192858 | PREDICTED: eukaryotic peptide chain release factor subunit 1 isoform X1 [Callithrix jacchus] | 0 | 7.34 | 5 | 2 | 12 | 2 | 1 | 437 | 49 | 5.71 | 70 | 2 | 1689805.791 | 1464542 | [illegible] |
| 296218846 | PREDICTED: ras and Rab interactor 1 [Callithrix jacchus] | 0 | 16.102 | 9 | 6 | 19 | 6 | 1 | 783 | 84.1 | 7.66 | 44 | 6 | 10736459.11 | 11409831.13 | [illegible] |
| 1060944344 | PREDICTED: NADH dehydrogenase [ubiquinone] iron-sulfur protein 8, mitochondrial [Callithrix jacchus] | 0 | 28.247 | 30 | 6 | 30 | 6 | 1 | 210 | 23.7 | 6.87 | 307 | 6 | 50988492.92 | 41419502.5 | [illegible] |
| 675710443 | PREDICTED: opalin isoform X1 [Callithrix jacchus] | 0 | 46.202 | 32 | 4 | 34 | 4 | 1 | 141 | 15.6 | 5.55 | 504 | 4 | 19834513.79 | 17838006.13 | [illegible] |
| 296221128 | PREDICTED: alpha-internexin [Callithrix jacchus] | 0 | 328.377 | 81 | 42 | 702 | 36 | 1 | 499 | 55.3 | 5.4 | 13503 | 42 | 1720397407 | 1403187363 | [illegible] |
| 296221267 | PREDICTED: actin-binding LIM protein 1 isoform X10 [Callithrix jacchus] | 0 | 269.057 | 62 | 37 | 324 | 37 | 1 | 748 | 84.4 | 8.32 | 4494 | 37 | 275533684.3 | 306490849.9 | [illegible] |
| 532290326 | protein FAM160B1 isoform a [Callithrix jacchus] | 0 | 9.49 | 4 | 3 | 5 | 3 | 1 | 764 | 86.3 | 5.22 | 20 | 3 | 2612979.929 | 2503827.125 | [illegible] |
| 296204889 | PREDICTED: aspartate--tRNA ligase, cytoplasmic [Callithrix jacchus] | 0 | 123.308 | 56 | 25 | 121 | 25 | 1 | 501 | 57.1 | 6.44 | 1395 | 25 | 137042780.7 | 151682268.4 | [illegible] |
| 675746008 | PREDICTED: ras GTPase-activating protein nGAP isoform X2 [Callithrix jacchus] | 0 | 302.167 | 48 | 50 | 340 | 43 | 1 | 1287 | 144.6 | 7.36 | 5529 | 50 | 160515039.6 | 118581079.6 | [illegible] |
| 675746367 | PREDICTED: protein PRRC2C isoform X1 [Callithrix jacchus] | 0 | 84.724 | 9 | 22 | 100 | 21 | 1 | 2902 | 317.6 | 9.06 | 410 | 22 | 27652144.27 | 26674929.75 | [illegible] |
| 296229450 | PREDICTED: NADH dehydrogenase [ubiquinone] iron-sulfur protein 2, mitochondrial [Callithrix jacchus] | 0 | 242.047 | 60 | 21 | 165 | 21 | 1 | 463 | 52.6 | 7.18 | 2342 | 21 | 263115881.2 | 260309780.7 | [illegible] |
| 675747956 | PREDICTED: serine/threonine-protein kinase MRCK alpha isoform X1 [Callithrix jacchus] | 0 | 212.525 | 26 | 39 | 215 | 35 | 1 | 1845 | 209.7 | 6.47 | 2968 | 39 | 92672364.4 | 89120291.39 | [illegible] |
| 390477223 | PREDICTED: WD repeat-containing protein 26 [Callithrix jacchus] | 0 | 30.972 | 13 | 8 | 16 | 8 | 1 | 658 | 71.9 | 6.16 | 76 | 8 | 11326928.78 | 10214013.03 | [illegible] |
| 532509886 | serine/threonine-protein kinase MARK1 [Callithrix jacchus] | 0 | 279.327 | 51 | 37 | 296 | 27 | 1 | 795 | 88.8 | 9.35 | 3421 | 37 | 156554986.5 | 155412494.7 | [illegible] |
| 675645703 | PREDICTED: beta-centractin [Callithrix jacchus] | 0 | 118.601 | 51 | 15 | 145 | 7 | 1 | 376 | 42.3 | 6.4 | 1830 | 15 | 45868060.18 | 53556074.13 | [illegible] |
| 532511542 | zinc transporter ZIP10 precursor [Callithrix jacchus] | 0 | 19.692 | 6 | 6 | 37 | 6 | 1 | 831 | 94.1 | 6.8 | 247 | 6 | 74550753.23 | 79669436.5 | [illegible] |
| 675664936 | PREDICTED: inactive phospholipase C-like protein 1 isoform X1 [Callithrix jacchus] | 0 | 8.385 | 4 | 3 | 5 | 2 | 1 | 1095 | 122.6 | 5.67 | 21 | 3 | 1855289.381 | 1992433.75 | [illegible] |
| 675665262 | PREDICTED: NADH-ubiquinone oxidoreductase 75 kDa subunit, mitochondrial isoform X1 [Callithrix jacchus] | 0 | 353.304 | 63 | 38 | 368 | 38 | 1 | 729 | 79.5 | 6.35 | 6074 | 38 | 410911043 | 435752834.5 | [illegible] |
| 675717594 | REDICTED: lisH domain and HEAT repeat-containing protein KIAA1468 homolog isoform X5 [Callithrix jacchu | 0 | 34.017 | 8 | 9 | 34 | 9 | 1 | 1203 | 132.9 | 5.38 | 285 | 9 | 12988604.15 | 13495269.04 | [illegible] |
| 532516950 | calcium/calmodulin-dependent protein kinase type 1D isoform 2 [Callithrix jacchus] | 0 | 47.173 | 28 | 9 | 37 | 6 | 1 | 385 | 42.9 | 7.21 | 546 | 9 | 22047630.47 | 18899204.25 | [illegible] |
| 532502685 | complement C1q-like protein 3 precursor [Callithrix jacchus] | 0 | 28.239 | 24 | 5 | 25 | 5 | 1 | 255 | 26.7 | 6.79 | 222 | 5 | 10671997.61 | 10605433.75 | [illegible] |
| 532218343 | tubulin beta-4B chain [Callithrix jacchus] | 0 | 583.079 | 80 | 31 | 1605 | 3 | 1 | 445 | 49.8 | 4.89 | 38351 | 31 | 714185004 | 717926653.6 | [illegible] |
| 532280676 | rab-like protein 6 isoform 1 [Callithrix jacchus] | 0 | 30.783 | 13 | 9 | 41 | 9 | 1 | 732 | 79.8 | 5.35 | 293 | 9 | 29128088.73 | 43133132.75 | [illegible] |
| 532439993 | calmodulin-regulated spectrin-associated protein 1 [Callithrix jacchus] | 0 | 56.343 | 11 | 14 | 42 | 13 | 1 | 1602 | 178 | 6.79 | 539 | 14 | 18565003.96 | 20870803.66 | [illegible] |
| 1060936357 | PREDICTED: protein transport protein Sec16A isoform X2 [Callithrix jacchus] | 0 | 179.247 | 19 | 30 | 163 | 30 | 1 | 2376 | 253.3 | 5.99 | 2594 | 30 | 54936394.02 | 51546769.69 | [illegible] |
| 675639520 | PREDICTED: calcium-binding mitochondrial carrier protein SCaMC-2 isoform X3 [Callithrix jacchus] | 0 | 51.215 | 28 | 12 | 45 | 11 | 1 | 501 | 56 | 8.95 | 456 | 12 | 15336982.63 | 15142363.98 | [illegible] |
| 1060935840 | :DICTED: LOW QUALITY PROTEIN: phosphatidylinositol 4-phosphate 5-kinase-like protein 1 [Callithrix jaccl | 0 | 47.558 | 27 | 10 | 48 | 10 | 1 | 407 | 45.7 | 9.48 | 315 | 10 | 12569758.92 | 64299110.5 | [illegible] |
| 532276392 | gelsolin isoform c [Callithrix jacchus] | 0 | 234.392 | 41 | 27 | 245 | 27 | 1 | 742 | 82 | 5.77 | 3753 | 27 | 344104683.8 | 342790641.2 | [illegible] |
| 1060935650 | PREDICTED: rab GTPase-activating protein 1 isoform X1 [Callithrix jacchus] | 0 | 53.583 | 12 | 11 | 42 | 10 | 1 | 1069 | 121.6 | 5.25 | 653 | 11 | 8516248.245 | 10346301.94 | [illegible] |
| 1060935666 | PREDICTED: DENN domain-containing protein 1A isoform X1 [Callithrix jacchus] | 0 | 117.707 | 23 | 17 | 131 | 17 | 1 | 1074 | 118.3 | 6.89 | 1668 | 17 | 52533383.55 | 58428621.88 | [illegible] |
| 532448201 | plakophilin-4 isoform a [Callithrix jacchus] | 0 | 292.2 | 48 | 45 | 325 | 40 | 1 | 1192 | 131.9 | 8.97 | 4690 | 45 | 213255198.7 | 200617049.8 | [illegible] |
| 1061003489 | PREDICTED: SEC14 domain and spectrin repeat-containing protein 1 isoform X1 [Callithrix jacchus] | 0 | 110.729 | 36 | 20 | 122 | 20 | 1 | 696 | 79.3 | 5.12 | 1238 | 20 | 72596568.55 | 78142115.03 | [illegible] |
| 1060943644 | PREDICTED: palmitoyltransferase ZDHHC5 isoform X1 [Callithrix jacchus] | 0 | 56.68 | 21 | 11 | 42 | 11 | 1 | 715 | 77.4 | 9.01 | 447 | 11 | 16180696.07 | 19333032.31 | [illegible] |
| 532288512 | 182 kDa tankyrase-1-binding protein [Callithrix jacchus] | 0 | 188.188 | 23 | 29 | 128 | 29 | 1 | 1731 | 182 | 4.86 | 2198 | 29 | 140739750.7 | 134056911.6 | [illegible] |
| 1060943466 | PREDICTED: MAP kinase-activating death domain protein isoform X1 [Callithrix jacchus] | 0 | 142.884 | 21 | 26 | 124 | 26 | 1 | 1660 | 184.3 | 6.3 | 1908 | 26 | 45365567.41 | 55610596.67 | [illegible] |
| 532332949 | vacuolar protein sorting-associated protein 18 homolog [Callithrix jacchus] | 0 | 78.298 | 17 | 16 | 83 | 16 | 1 | 973 | 110.2 | 6.14 | 940 | 16 | 41725052.74 | 45021390.13 | [illegible] |
| 532510940 | inositol-trisphosphate 3-kinase A [Callithrix jacchus] | 0 | 195.86 | 57 | 23 | 195 | 23 | 1 | 461 | 50.9 | 7.85 | 3964 | 23 | 258818580.2 | 258027881.1 | [illegible] |
| 390468605 | PREDICTED: FERM domain-containing protein 5 isoform X1 [Callithrix jacchus] | 0 | 29.092 | 15 | 7 | 24 | 7 | 1 | 570 | 65 | 8.35 | 133 | 7 | 4533009.178 | 4230831.063 | [illegible] |
| 532517336 | disks large-associated protein 4 isoform a [Callithrix jacchus] | 0 | 261.772 | 49 | 35 | 273 | 32 | 1 | 985 | 107.1 | 7.43 | 5105 | 35 | 154538618 | 150960372.3 | [illegible] |
| 675656633 | PREDICTED: ral GTPase-activating protein subunit beta isoform X4 [Callithrix jacchus] | 0 | 164.923 | 27 | 30 | 169 | 30 | 1 | 1503 | 167.8 | 6.83 | 1847 | 30 | 121533961.3 | 112154712 | [illegible] |
| 532504126 | protein phosphatase 1 regulatory inhibitor subunit 16B isoform 1 [Callithrix jacchus] | 0 | 30.996 | 17 | 8 | 40 | 8 | 1 | 567 | 63.5 | 6.42 | 170 | 8 | 14934464.07 | 14041996.25 | [illegible] |
| 675656744 | PREDICTED: ganglioside-induced differentiation-associated protein 1-like 1 [Callithrix jacchus] | 0 | 125.085 | 50 | 15 | 87 | 15 | 1 | 367 | 41.9 | 6.6 | 1585 | 15 | 86792914.39 | 88918345.78 | [illegible] |
| 675657271 | PREDICTED: syntaxin-16 isoform X1 [Callithrix jacchus] | 0 | 19.543 | 20 | 5 | 20 | 5 | 1 | 339 | 38.3 | 5.81 | 456 | 5 | 45968549.95 | 41998310.38 | [illegible] |
| 1060993969 | REDICTED: LOW QUALITY PROTEIN: ADP-ribosylation factor GTPase-activating protein 1 [Callithrix jacchu | 0 | 118.424 | 46 | 16 | 127 | 16 | 1 | 416 | 45.8 | 5.66 | 2145 | 16 | 114501055.7 | 101773031.5 | [illegible] |
| 296226575 | PREDICTED: YTH domain-containing family protein 3 isoform X2 [Callithrix jacchus] | 0 | 32.807 | 14 | 7 | 37 | 5 | 1 | 585 | 63.8 | 9.04 | 191 | 7 | 2048668.81 | 3825085.875 | [illegible] |
| 390475397 | DICTED: LOW QUALITY PROTEIN: acyl-CoA dehydrogenase family member 9, mitochondrial [Callithrix jacc | 0 | 31.274 | 14 | 8 | 21 | 8 | 1 | 621 | 68.8 | 7.44 | 114 | 8 | 44762350.07 | 52321360.5 | [illegible] |
| 532515864 | uncharacterized protein KIAA1109 [Callithrix jacchus] | 0 | 34.241 | 2 | 10 | 15 | 10 | 1 | 5004 | 555.5 | 6.6 | 102 | 10 | 6200494.234 | 2654515.75 | [illegible] |
| 532215783 | heterogeneous nuclear ribonucleoprotein A3 [Callithrix jacchus] | 0 | 173.568 | 45 | 19 | 233 | 4 | 1 | 378 | 39.6 | 9.01 | 3557 | 19 | 40875197.02 | 46958706.92 | [illegible] |
| 390476220 | PREDICTED: eukaryotic translation initiation factor 2A isoform X1 [Callithrix jacchus] | 0 | 30.475 | 17 | 8 | 24 | 8 | 1 | 585 | 65 | 8.95 | 46 | 8 | 19798207.11 | 20106795.25 | [illegible] |
| 532513954 | synaptotagmin-11 [Callithrix jacchus] | 0 | 69.952 | 25 | 10 | 74 | 10 | 1 | 435 | 48.6 | 9.11 | 838 | 10 | 49189597.58 | 66697299.88 | [illegible] |
| 675743530 | PREDICTED: pre-B-cell leukemia transcription factor-interacting protein 1 isoform X1 [Callithrix jacchus] | 0 | 56.513 | 19 | 13 | 67 | 13 | 1 | 721 | 79.5 | 5.08 | 816 | 13 | 23102858.52 | 30112727.75 | [illegible] |
| 675743296 | PREDICTED: DENN domain-containing protein 4B [Callithrix jacchus] | 0 | 15.204 | 4 | 4 | 6 | 4 | 1 | 1544 | 168.9 | 7.39 | 35 | 4 | 5804974.64 | 6086677.5 | [illegible] |
| 1061012835 | PREDICTED: cold shock domain-containing protein E1 [Callithrix jacchus] | 0 | 89.054 | 26 | 19 | 93 | 19 | 1 | 844 | 93.7 | 6.6 | 684 | 19 | 30597141.3 | 32230461.28 | [illegible] |
| 675671664 | PREDICTED: striatin-interacting protein 1 isoform X1 [Callithrix jacchus] | 0 | 37.358 | 12 | 9 | 27 | 5 | 1 | 838 | 95.6 | 6.29 | 184 | 9 | 2867780.662 | 4291772.438 | [illegible] |
| 296208769 | PREDICTED: adenosylhomocysteinase 2 isoform X1 [Callithrix jacchus] | 0 | 206.893 | 59 | 28 | 275 | 12 | 1 | 530 | 58.9 | 6.89 | 3884 | 28 | 53371301.67 | 46021807.88 | [illegible] |
| 1061012548 | PREDICTED: LOW QUALITY PROTEIN: noelin-3 [Callithrix jacchus] | 0 | 50.945 | 32 | 13 | 59 | 13 | 1 | 458 | 52.7 | 8.15 | 338 | 13 | 24931862.49 | 27455699.41 | [illegible] |
| 390466235 | PREDICTED: phospholipid phosphatase-related protein type 4 [Callithrix jacchus] | 0 | 140.52 | 36 | 21 | 197 | 21 | 1 | 715 | 78.3 | 8.69 | 3558 | 21 | 159204596.9 | 158002569.1 | [illegible] |
| 675648482 | PREDICTED: rap1 GTPase-GDP dissociation stimulator 1 isoform X2 [Callithrix jacchus] | 0 | 121.119 | 41 | 20 | 101 | 20 | 1 | 607 | 66.4 | 5.31 | 1058 | 20 | 50468942.41 | 45609025.97 | [illegible] |
| 296208305 | PREDICTED: adenylate kinase isoenzyme 5 isoform X1 [Callithrix jacchus] | 0 | 123.787 | 43 | 22 | 175 | 21 | 1 | 562 | 63.2 | 5.17 | 2785 | 22 | 341667105.6 | 338498312.8 | [illegible] |
| 296190087 | PREDICTED: NADH dehydrogenase [ubiquinone] 1 beta subcomplex subunit 6 isoform X2 [Callithrix jacchus | 0 | 35.805 | 52 | 5 | 31 | 5 | 1 | 128 | 15.4 | 9.5 | 275 | 5 | 19918990.19 | 20149457.38 | [illegible] |
| 532512040 | protein KIAA1045 [Callithrix jacchus] | 0 | 108.876 | 45 | 15 | 148 | 15 | 1 | 400 | 45.2 | 5.64 | 2289 | 15 | 104530183.7 | 101658477.5 | [illegible] |
| 1060935345 | PREDICTED: DOMON domain-containing protein FRRS1L [Callithrix jacchus] | 0 | 29.804 | 28 | 5 | 59 | 5 | 1 | 167 | 19 | 5.06 | 884 | 5 | 34774744.85 | 32327117.25 | [illegible] |
| 1061011007 | PREDICTED: polyadenylate-binding protein 4 isoform X1 [Callithrix jacchus] | 0 | 83.004 | 30 | 18 | 81 | 10 | 1 | 660 | 72.4 | 9.35 | 999 | 18 | 25731749.33 | 19680121.13 | [illegible] |
| 296207491 | PREDICTED: thyroid hormone receptor-associated protein 3 [Callithrix jacchus] | 0 | 74.529 | 17 | 14 | 59 | 13 | 1 | 955 | 108.6 | 10.17 | 436 | 14 | 20042225.36 | 17646731.08 | [illegible] |
| 675669074 | PREDICTED: adhesion G protein-coupled receptor B2 [Callithrix jacchus] | 0 | 228.538 | 26 | 31 | 201 | 31 | 1 | 1583 | 172.7 | 6.8 | 3771 | 31 | 124433766 | 126836731.1 | [illegible] |
| 1061010274 | PREDICTED: eukaryotic translation initiation factor 4 gamma 3 isoform X1 [Callithrix jacchus] | 0 | 101.023 | 17 | 28 | 140 | 23 | 1 | 1678 | 186.9 | 5.68 | 872 | 28 | 66267996.78 | 76110712.63 | [illegible] |
| 1061010271 | PREDICTED: SH2 domain-containing protein 5 isoform X1 [Callithrix jacchus] | 0 | 37.598 | 25 | 10 | 31 | 10 | 1 | 465 | 51.1 | 8.75 | 115 | 10 | 9875432.545 | 8183655.75 | [illegible] |
| 532288136 | immunoglobulin superfamily member 21 precursor [Callithrix jacchus] | 0 | 129.343 | 34 | 15 | 268 | 15 | 1 | 468 | 51.7 | 6.47 | 5918 | 15 | 704078265.2 | 736754151.9 | [illegible] |
| 296206720 | PREDICTED: F-box only protein 2 [Callithrix jacchus] | 0 | 100.337 | 49 | 11 | 75 | 11 | 1 | 297 | 33.6 | 4.36 | 1407 | 11 | 53341752.64 | 47063880.5 | [illegible] |
| 296206694 | PREDICTED: TAR DNA-binding protein 43 [Callithrix jacchus] | 0 | 57.721 | 24 | 9 | 58 | 9 | 1 | 414 | 44.6 | 6.43 | 1209 | 9 | 32342165.05 | 30655240.38 | [illegible] |
| 296206602 | PREDICTED: dnaJ homolog subfamily C member 11 [Callithrix jacchus] | 0 | 166.684 | 54 | 26 | 279 | 26 | 1 | 559 | 63.2 | 8.4 | 4482 | 26 | 377630698.6 | 423818966.7 | [illegible] |
| 532510558 | adherens junction-associated protein 1 precursor [Callithrix jacchus] | 0 | 39.184 | 18 | 4 | 17 | 4 | 1 | 412 | 44.5 | 9.5 | 192 | 4 | 14560093.52 | 14864792.75 | [illegible] |
| 675723886 | PREDICTED: trifunctional enzyme subunit beta, mitochondrial [Callithrix jacchus] | 0 | 143.728 | 47 | 21 | 263 | 21 | 1 | 475 | 51.4 | 9.47 | 3707 | 21 | 446255373.2 | 456239976.2 | [illegible] |
| 532452349 | actin-binding LIM protein 2 isoform 1 [Callithrix jacchus] | 0 | 267.779 | 62 | 33 | 348 | 2 | 1 | 645 | 71.7 | 8.12 | 6303 | 33 | 9129769.847 | 10578801 | [illegible] |
| 296196376 | PREDICTED: G-rich sequence factor 1 [Callithrix jacchus] | 0 | 28.499 | 13 | 5 | 34 | 5 | 1 | 479 | 53 | 6.27 | 313 | 5 | 31408454.89 | 27411237.41 | [illegible] |
| 532446747 | cyclin-G-associated kinase [Callithrix jacchus] | 0 | 208.116 | 32 | 37 | 246 | 37 | 1 | 1311 | 143.1 | 5.72 | 2690 | 37 | 174291383.2 | 177436617.2 | [illegible] |
| 532334589 | translational activator GCN1 [Callithrix jacchus] | 0 | 38.352 | 4 | 10 | 22 | 10 | 1 | 2671 | 292.3 | 7.58 | 102 | 10 | 6430970.767 | 4731607.688 | [illegible] |
| 532517638 | TRAF-type zinc finger domain-containing protein 1 [Callithrix jacchus] | 0 | 7.381 | 4 | 2 | 2 | 2 | 1 | 581 | 64.9 | 5.34 | 16 | 2 | 1692942.653 | 1274486.125 | [illegible] |
| 1061018590 | PREDICTED: RIMS-binding protein 2 [Callithrix jacchus] | 0 | 310.184 | 34 | 35 | 292 | 35 | 1 | 1528 | 168.2 | 6.8 | 5070 | 35 | 205087908.9 | 209501527.8 | [illegible] |
| 675642443 | PREDICTED: uncharacterized protein C7orf43 homolog [Callithrix jacchus] | 0 | 20.561 | 11 | 5 | 18 | 5 | 1 | 580 | 62.6 | 8.59 | 141 | 5 | 2483171.982 | 3056817.625 | [illegible] |
| 1060963747 | PREDICTED: arf-GAP with dual PH domain-containing protein 1 [Callithrix jacchus] | 0 | 38.267 | 30 | 9 | 43 | 9 | 1 | 374 | 43.5 | 8.9 | 381 | 9 | 4409718.202 | 4520488 | [illegible] |
| 675642653 | EDICTED: aminoacyl tRNA synthase complex-interacting multifunctional protein 2 isoform X1 [Callithrix jacch | 0 | 18.491 | 16 | 5 | 22 | 5 | 1 | 320 | 35.2 | 8.02 | 198 | 5 | 6547055.291 | 9392381.25 | [illegible] |
| 1060962819 | PREDICTED: serine/threonine-protein kinase LMTK2 isoform X1 [Callithrix jacchus] | 0 | 115.242 | 16 | 18 | 100 | 18 | 1 | 1502 | 165.3 | 4.51 | 1002 | 18 | 77838507.48 | 83731410.09 | [illegible] |
| 296223392 | PREDICTED: S-adenosylmethionine synthase [Callithrix jacchus] | 0 | 18.35 | 14 | 5 | 12 | 5 | 1 | 395 | 43.6 | 6.48 | 98 | 5 | 7305237.66 | 6547474.5 | [illegible] |
| 675730487 | PREDICTED: IQ motif and SEC7 domain-containing protein 1 [Callithrix jacchus] | 0 | 421.525 | 46 | 48 | 654 | 42 | 1 | 1129 | 126.1 | 7.97 | 10783 | 48 | 832382586.2 | 811105678.7 | [illegible] |
| 390475353 | PREDICTED: histone deacetylase 11 isoform X5 [Callithrix jacchus] | 0 | 10.712 | 10 | 3 | 13 | 3 | 1 | 347 | 39.1 | 7.08 | 41 | 3 | 4226471.918 | 5177193.375 | [illegible] |
| 1060955077 | PREDICTED: sodium- and chloride-dependent GABA transporter 1 [Callithrix jacchus] | 0 | 48.114 | 16 | 6 | 69 | 6 | 1 | 599 | 67 | 8.09 | 1282 | 6 | 56263804.47 | 58680370.5 | [illegible] |
| 532346705 | serine/threonine-protein kinase WNK1 isoform 4 [Callithrix jacchus] | 0 | 48.88 | 6 | 14 | 47 | 9 | 1 | 2641 | 279.4 | 6.21 | 684 | 14 | 5551778.009 | 5455326.625 | [illegible] |
| 532441423 | 28S ribosomal protein S35, mitochondrial isoform 1 precursor [Callithrix jacchus] | 0 | 34.438 | 28 | 7 | 43 | 7 | 1 | 322 | 36.7 | 9.01 | 485 | 7 | 12327924.38 | 12703058.22 | [illegible] |
| 675675826 | PREDICTED: LOW QUALITY PROTEIN: coiled-coil domain-containing protein 136 [Callithrix jacchus] | 0 | 120.459 | 15 | 17 | 108 | 17 | 1 | 1158 | 134.3 | 4.63 | 1801 | 17 | 43453190.82 | 42221149.63 | [illegible] |
| 675774079 | PREDICTED: protein phosphatase 1 regulatory subunit 12C isoform X1 [Callithrix jacchus] | 0 | 60.063 | 20 | 12 | 58 | 12 | 1 | 781 | 84.7 | 5.72 | 688 | 12 | 14805375.8 | 12964808.25 | [illegible] |
| 1060971400 | PREDICTED: LOW QUALITY PROTEIN: protein shisa-7, partial [Callithrix jacchus] | 0 | 115.823 | 40 | 20 | 191 | 20 | 1 | 494 | 51.7 | 9.7 | 1481 | 20 | 236165980.1 | 232534010.1 | [illegible] |
| 532202281 | rho GTPase-activating protein 35 [Callithrix jacchus] | 0 | 36.905 | 7 | 10 | 53 | 10 | 1 | 1499 | 170.4 | 6.64 | 221 | 10 | 13976023.33 | 14235245.28 | [illegible] |
| 675762489 | PREDICTED: AP-2 complex subunit sigma [Callithrix jacchus] | 0 | 58.277 | 44 | 7 | 343 | 7 | 1 | 142 | 17 | 6.18 | 4029 | 7 | 669991277.7 | 722095800.9 | [illegible] |
| 532446439 | MAP/microtubule affinity-regulating kinase 4 isoform 2 [Callithrix jacchus] | 0 | 81.284 | 27 | 14 | 124 | 12 | 1 | 688 | 75.2 | 9.79 | 1082 | 14 | 41686926.63 | 43774331.5 | [illegible] |
| 675761661 | PREDICTED: sodium/potassium-transporting ATPase subunit alpha-3 isoform X1 [Callithrix jacchus] | 0 | 680.772 | 52 | 56 | 1100 | 33 | 1 | 1114 | 122.8 | 6 | 22377 | 56 | 1381894291 | 1440460790 | [illegible] |
| 296233871 | PREDICTED: heterogeneous nuclear ribonucleoprotein U-like protein 1 [Callithrix jacchus] | 0 | 11.083 | 5 | 3 | 3 | 3 | 1 | 854 | 95.5 | 6.92 | 49 | 3 | 5692690.5 | 4975534.25 | [illegible] |
| 1061002432 | PREDICTED: myotubularin-related protein 10 [Callithrix jacchus] | 0 | 27.132 | 9 | 6 | 35 | 6 | 1 | 777 | 88.4 | 8.41 | 181 | 6 | 7266464.251 | 9749377.656 | [illegible] |
| 296204031 | PREDICTED: vacuolar protein sorting-associated protein 33B isoform X1 [Callithrix jacchus] | 0 | 11.98 | 7 | 5 | 10 | 5 | 1 | 617 | 70.6 | 6.9 | 40 | 5 | 14305880.45 | 17324487.75 | [illegible] |
| 675706706 | PREDICTED: DCN1-like protein 3 [Callithrix jacchus] | 0 | 30.977 | 24 | 6 | 37 | 6 | 1 | 304 | 34.3 | 5.12 | 409 | 6 | 12184935.18 | 14475589.94 | [illegible] |
| 296219856 | PREDICTED: elongation factor Tu, mitochondrial isoform X1 [Callithrix jacchus] | 0 | 292.588 | 62 | 25 | 388 | 25 | 1 | 455 | 49.7 | 7.55 | 7042 | 25 | 1235724626 | 1357339641 | [illegible] |
| 532291308 | ataxin-2-like protein isoform C [Callithrix jacchus] | 0 | 97.328 | 23 | 21 | 91 | 20 | 1 | 1098 | 115.6 | 8.66 | 552 | 21 | 60476110.79 | 55460550.69 | [illegible] |
| 675707382 | PREDICTED: LOW QUALITY PROTEIN: seizure 6-like protein 2 [Callithrix jacchus] | 0 | 17.597 | 6 | 6 | 15 | 5 | 1 | 931 | 99.6 | 4.86 | 35 | 6 | 5973788.489 | 4532467.5 | [illegible] |
| 1236589234 | FUS RNA binding protein [Callithrix jacchus] | 0 | 60.812 | 13 | 7 | 56 | 7 | 1 | 527 | 53.4 | 9.36 | 641 | 7 | 62055919.08 | 62779257.44 | [illegible] |
| 1060944424 | PREDICTED: liprin-alpha-1 isoform X1 [Callithrix jacchus] | 0 | 130.127 | 21 | 24 | 183 | 9 | 1 | 1268 | 143.2 | 6.32 | 1683 | 24 | 22583908.24 | 21544902.19 | [illegible] |
| 390460228 | PREDICTED: neuronal membrane glycoprotein M6-a isoform X2 [Callithrix jacchus] | 0 | 92.404 | 26 | 9 | 171 | 8 | 1 | 278 | 31.2 | 5.27 | 3487 | 9 | 378854476.5 | 297196129.7 | [illegible] |
| 1060949961 | PREDICTED: PH and SEC7 domain-containing protein 3 isoform X1 [Callithrix jacchus] | 0 | 333.484 | 31 | 41 | 616 | 41 | 1 | 1291 | 142.1 | 6.89 | 9002 | 41 | 1467527868 | 1432724583 | [illegible] |
| 675759030 | PREDICTED: coiled-coil domain-containing protein 124 [Callithrix jacchus] | 0 | 38.214 | 35 | 8 | 65 | 8 | 1 | 223 | 25.8 | 9.55 | 590 | 8 | 26814057.13 | 27972082.66 | [illegible] |
| 532524196 | CCR4-NOT transcription complex subunit 1 isoform a [Callithrix jacchus] | 0 | 64.297 | 8 | 17 | 50 | 17 | 1 | 2383 | 267.7 | 7.11 | 271 | 17 | 16473463.82 | 13359245.81 | [illegible] |
| 296231279 | PREDICTED: cytoplasmic dynein 1 light intermediate chain 2 [Callithrix jacchus] | 0 | 105.151 | 41 | 16 | 109 | 15 | 1 | 490 | 53.9 | 6.38 | 1335 | 16 | 72314198.92 | 47630187.81 | [illegible] |
| 1060967540 | PREDICTED: UPF0183 protein C16orf70 homolog [Callithrix jacchus] | 0 | 38.552 | 23 | 7 | 21 | 7 | 1 | 422 | 47.6 | 7.75 | 242 | 7 | 10374556.43 | 9567192.125 | [illegible] |
| 675752028 | REDICTED: LOW QUALITY PROTEIN: capping protein, Arp2/3 and myosin-I linker protein 2 [Callithrix jacchu | 0 | 25.817 | 5 | 6 | 14 | 6 | 1 | 1374 | 148.8 | 6.54 | 206 | 6 | 3192130.153 | 3728249.25 | [illegible] |
| 390477803 | PREDICTED: enhancer of mRNA-decapping protein 4 isoform X2 [Callithrix jacchus] | 0 | 215.499 | 31 | 31 | 176 | 31 | 1 | 1403 | 151.8 | 5.82 | 2557 | 31 | 111413146.1 | 109171679.3 | [illegible] |
| 1060966997 | PREDICTED: LOW QUALITY PROTEIN: protein VAC14 homolog [Callithrix jacchus] | 0 | 37.729 | 11 | 7 | 43 | 7 | 1 | 782 | 88 | 6.54 | 300 | 7 | 15426950.86 | 14106314.69 | [illegible] |
| 1060967746 | PREDICTED: L-fucose kinase isoform X1 [Callithrix jacchus] | 0 | 297.758 | 51 | 32 | 193 | 32 | 1 | 1084 | 117.7 | 6.27 | 3586 | 32 | 152634068.2 | 154302291.6 | [illegible] |
| 675752684 | PREDICTED: alanine--tRNA ligase, cytoplasmic [Callithrix jacchus] | 0 | 24.882 | 8 | 6 | 29 | 6 | 1 | 968 | 106.7 | 5.45 | 196 | 6 | 10935368.34 | 11297563.19 | [illegible] |
| 1060968123 | PREDICTED: uncharacterized protein KIAA0513 homolog [Callithrix jacchus] | 0 | 41.027 | 19 | 7 | 37 | 7 | 1 | 411 | 46.7 | 5.1 | 456 | 7 | 11511521.67 | 11713987.38 | [illegible] |
| 296231815 | PREDICTED: charged multivesicular body protein 1a [Callithrix jacchus] | 0 | 22.268 | 30 | 8 | 82 | 8 | 1 | 196 | 21.7 | 8.06 | 1117 | 8 | 116974783.8 | 117789945.5 | [illegible] |
| 1060941217 | PREDICTED: guanylate cyclase soluble subunit alpha-2 [Callithrix jacchus] | 0 | 96.631 | 33 | 18 | 91 | 18 | 1 | 637 | 72.3 | 6.89 | 916 | 18 | 57961224.16 | 53972791.63 | [illegible] |
| 1060974516 | PREDICTED: phospholipid phosphatase-related protein type 2 isoform X1 [Callithrix jacchus] | 0 | 43.552 | 20 | 6 | 34 | 6 | 1 | 471 | 50.4 | 9.42 | 993 | 6 | 23352958.81 | 22998773.63 | [illegible] |
| 1061013204 | PREDICTED: septin-7 isoform X5 [Callithrix jacchus] | 0 | 373.019 | 78 | 32 | 836 | 2 | 1 | 384 | 44.8 | 8.22 | 12687 | 32 | 47293238.09 | 44250044 | [illegible] |
| 675695521 | PREDICTED: LOW QUALITY PROTEIN: protein turtle homolog B [Callithrix jacchus] | 0 | 142.239 | 25 | 23 | 81 | 23 | 1 | 1437 | 156.8 | 7.18 | 1468 | 23 | 13233076.17 | 15859245.17 | [illegible] |
| 1060941902 | PREDICTED: neurotrimin [Callithrix jacchus] | 0 | 117.494 | 40 | 14 | 378 | 10 | 1 | 367 | 40.5 | 7.64 | 8691 | 14 | 951740399 | 968597134.6 | [illegible] |
| 296216564 | PREDICTED: fasciculation and elongation protein zeta-1 [Callithrix jacchus] | 0 | 26.23 | 17 | 6 | 20 | 6 | 1 | 392 | 45.1 | 4.32 | 166 | 6 | 11646363.69 | 11982566.25 | [illegible] |
| 1060941710 | PREDICTED: GRAM domain-containing protein 1B [Callithrix jacchus] | 0 | 30.077 | 10 | 7 | 28 | 6 | 1 | 888 | 102.3 | 6.73 | 134 | 7 | 7837526.055 | 7032397.719 | [illegible] |
| 296216278 | PREDICTED: RING finger protein 214 isoform X1 [Callithrix jacchus] | 0 | 47.041 | 17 | 10 | 35 | 10 | 1 | 704 | 77.5 | 6.8 | 640 | 10 | 7844837.348 | 8168976.313 | [illegible] |
| 1060941356 | ıydrolipoyllysine-residue acetyltransferase component of pyruvate dehydrogenase complex, mitochondrial [C | 0 | 242.462 | 51 | 23 | 206 | 23 | 1 | 647 | 68.5 | 8.13 | 3091 | 23 | 532356751.2 | 492436574.9 | [illegible] |
| 296214014 | PREDICTED: dmX-like protein 2 isoform X2 [Callithrix jacchus] | 0 | 848.537 | 48 | 112 | 766 | 109 | 1 | 3037 | 339.7 | 6.38 | 10265 | 112 | 483400594.3 | 463674567.9 | [illegible] |
| 1061021310 | :D: leucine-rich repeat and immunoglobulin-like domain-containing nogo receptor-interacting protein 1 [Callithr | 0 | 42.395 | 18 | 9 | 40 | 9 | 1 | 620 | 69.8 | 8.4 | 193 | 9 | 20101118.18 | 23796093.88 | [illegible] |
| 532518664 | coronin-2B isoform 1 [Callithrix jacchus] | 0 | 166.714 | 49 | 20 | 200 | 20 | 1 | 480 | 54.9 | 8.27 | 2894 | 20 | 343103847.9 | 369545200.8 | [illegible] |
| 296213405 | PREDICTED: ras-related protein Rab-8B isoform X1 [Callithrix jacchus] | 0 | 30.123 | 30 | 6 | 54 | 2 | 1 | 207 | 23.6 | 9.07 | 509 | 6 | 3124743.629 | 3738227.75 | [illegible] |
| 532283588 | dnaJ homolog subfamily C member 13 [Callithrix jacchus] | 0 | 117.896 | 15 | 28 | 91 | 28 | 1 | 2248 | 254.7 | 6.7 | 721 | 28 | 35626907.9 | 35760968.94 | [illegible] |
| 532518406 | calcium-transporting ATPase type 2C member 1 isoform 2a [Callithrix jacchus] | 0 | 9.397 | 2 | 2 | 11 | 2 | 1 | 983 | 108.1 | 6.83 | 47 | 2 | 2196804.526 | 2928080.375 | [illegible] |
| 296225243 | PREDICTED: caM kinase-like vesicle-associated protein isoform X2 [Callithrix jacchus] | 0 | 257.356 | 71 | 25 | 272 | 2 | 1 | 467 | 51.5 | 5.63 | 7747 | 25 | 9949214.847 | 10496889.5 | [illegible] |
| 532339861 | plexin-B1 precursor [Callithrix jacchus] | 0 | 22.185 | 4 | 6 | 14 | 6 | 1 | 2142 | 233.4 | 5.68 | 117 | 6 | 1225061.465 | 9457556.625 | [illegible] |
| 296228295 | PREDICTED: cytoplasmic dynein 1 light intermediate chain 1 [Callithrix jacchus] | 0 | 194.692 | 59 | 25 | 275 | 24 | 1 | 523 | 56.6 | 6.42 | 3387 | 25 | 175989335.4 | 164451516.8 | [illegible] |
| 675669288 | PREDICTED: protein argonaute-1 isoform X1 [Callithrix jacchus] | 0 | 54.139 | 15 | 11 | 63 | 6 | 1 | 860 | 97.5 | 9.19 | 896 | 11 | 5153849.629 | 5902179 | [illegible] |
| 296228521 | PREDICTED: D-3-phosphoglycerate dehydrogenase [Callithrix jacchus] | 0 | 130.995 | 42 | 18 | 181 | 18 | 1 | 533 | 56.5 | 6.54 | 2878 | 18 | 275172162.3 | 253562920.3 | [illegible] |
| 1061018942 | PREDICTED: LOW QUALITY PROTEIN: ubiquitin carboxyl-terminal hydrolase 11 [Callithrix jacchus] | 0 | 12.147 | 4 | 4 | 10 | 4 | 1 | 970 | 110.7 | 5.63 | 14 | 4 | 6828608.387 | 7350514.938 | [illegible] |
| 1061020464 | PREDICTED: sodium/hydrogen exchanger 6 isoform X1 [Callithrix jacchus] | 0 | 10.512 | 5 | 3 | 10 | 3 | 1 | 707 | 78.5 | 5.9 | 45 | 3 | 5025479.571 | 4649618.75 | [illegible] |
| 296235150 | PREDICTED: pyruvate dehydrogenase kinase, isozyme 3 isoform X1 [Callithrix jacchus] | 0 | 97.537 | 43 | 15 | 101 | 14 | 1 | 415 | 48 | 8.91 | 2059 | 15 | 61026836.89 | 60238392.63 | [illegible] |
| 1061020030 | PREDICTED: rho guanine nucleotide exchange factor 9 isoform X1 [Callithrix jacchus] | 0 | 88.446 | 40 | 18 | 113 | 16 | 1 | 516 | 61 | 5.72 | 712 | 18 | 57325370.81 | 60161942.67 | [illegible] |
| 675756023 | PREDICTED: C2 calcium-dependent domain-containing protein 4C [Callithrix jacchus] | 0 | 44.547 | 32 | 11 | 54 | 11 | 1 | 418 | 44 | 9.95 | 341 | 11 | 23468888.92 | 23805665.52 | [illegible] |
| 532509707 | cyclin-dependent kinase 17 isoform 2 [Callithrix jacchus] | 0 | 53.711 | 24 | 12 | 72 | 9 | 1 | 523 | 59.5 | 8.81 | 682 | 12 | 26073113.48 | 27131603.19 | [illegible] |
| 1061017481 | PREDICTED: oxysterol-binding protein-related protein 8 isoform X1 [Callithrix jacchus] | 0 | 72.119 | 20 | 14 | 51 | 14 | 1 | 889 | 101.2 | 6.96 | 620 | 14 | 23058573.87 | 23915030.31 | [illegible] |
| 532520740 | echinoderm microtubule-associated protein-like 6 [Callithrix jacchus] | 0 | 48.314 | 8 | 13 | 23 | 11 | 1 | 1958 | 217.8 | 7.42 | 141 | 13 | 2325211.046 | 1696085.5 | [illegible] |
| 532523860 | sterile alpha and TIR motif-containing protein 1 precursor [Callithrix jacchus] | 0 | 72.4 | 25 | 14 | 53 | 14 | 1 | 724 | 79.5 | 6.61 | 456 | 14 | 26218301.24 | 21518510.56 | [illegible] |
| 675690612 | PREDICTED: A-kinase anchor protein 5 [Callithrix jacchus] | 0 | 95.089 | 46 | 16 | 82 | 16 | 1 | 426 | 47 | 4.96 | 1895 | 16 | 40407064.95 | 43660827.97 | [illegible] |
| 1060962307 | PREDICTED: LOW QUALITY PROTEIN: transportin-1 [Callithrix jacchus] | 0 | 38.305 | 11 | 9 | 33 | 6 | 1 | 940 | 107.1 | 5.15 | 270 | 9 | 8689360.6 | 8961951.438 | [illegible] |
| 1060966467 | PREDICTED: cAMP-specific 3',5'-cyclic phosphodiesterase 4D isoform X1 [Callithrix jacchus] | 0 | 99.42 | 25 | 15 | 157 | 11 | 1 | 748 | 84.6 | 4.96 | 1859 | 15 | 26638876.5 | 28518448.91 | [illegible] |
| 532520494 | ERC protein 2 [Callithrix jacchus] | 0 | 412.11 | 58 | 66 | 688 | 2 | 1 | 957 | 110.5 | 7.05 | 9811 | 66 | 27160543.58 | 16765302.5 | [illegible] |
| 532214265 | 2-oxoglutarate dehydrogenase-like, mitochondrial isoform a [Callithrix jacchus] | 0 | 421.045 | 54 | 51 | 548 | 41 | 1 | 1010 | 114.6 | 6.87 | 8278 | 51 | 285004274.3 | 270587525.2 | [illegible] |
| 1060949768 | PREDICTED: paraneoplastic antigen Ma2 [Callithrix jacchus] | 0 | 44.645 | 29 | 8 | 45 | 8 | 1 | 364 | 41.5 | 4.84 | 1162 | 8 | 33079531.75 | 36709137.53 | [illegible] |
| 1060933510 | PREDICTED: BTB/POZ domain-containing protein KCTD12 isoform X1 [Callithrix jacchus] | 0 | 72.484 | 43 | 12 | 78 | 9 | 1 | 331 | 36 | 5.49 | 1412 | 12 | 52447363.81 | 56865636.38 | [illegible] |
| 532343035 | voltage-dependent T-type calcium channel subunit alpha-1I isoform b [Callithrix jacchus] | 0 | 22.909 | 3 | 5 | 12 | 5 | 1 | 2196 | 242.4 | 6.57 | 129 | 5 | 3414974.989 | 3118338.563 | [illegible] |
| 1061016735 | PREDICTED: disco-interacting protein 2 homolog B isoform X1 [Callithrix jacchus] | 0 | 212.135 | 32 | 37 | 212 | 35 | 1 | 1621 | 176.4 | 8.31 | 2202 | 37 | 128197956.5 | 125069909 | [illegible] |
| 1060970457 | PREDICTED: cell adhesion molecule 2 isoform X1 [Callithrix jacchus] | 0 | 66.688 | 25 | 9 | 88 | 9 | 1 | 437 | 47.5 | 5.33 | 955 | 9 | 21355671.71 | 20537234.31 | [illegible] |
| 675652173 | PREDICTED: ras/Rap GTPase-activating protein SynGAP [Callithrix jacchus] | 0 | 816.793 | 60 | 63 | 1161 | 61 | 1 | 1339 | 148 | 8.94 | 22721 | 63 | 2634041569 | 2572687177 | [illegible] |
| 1060985640 | PREDICTED: copine-5 [Callithrix jacchus] | 0 | 21.792 | 11 | 6 | 21 | 5 | 1 | 610 | 67.5 | 5.88 | 440 | 6 | 6156660.02 | 6428212.063 | [illegible] |
| 1060974801 | PREDICTED: fibronectin type III and SPRY domain-containing protein 1 [Callithrix jacchus] | 0 | 33.443 | 18 | 8 | 31 | 8 | 1 | 496 | 55.7 | 6.86 | 177 | 8 | 37605329.49 | 36881285.88 | [illegible] |
| 1061009461 | PREDICTED: junctional protein associated with coronary artery disease isoform X2 [Callithrix jacchus] | 0 | 195.996 | 35 | 34 | 159 | 34 | 1 | 1352 | 146.9 | 6.87 | 1537 | 34 | 79680509.97 | 86196648.73 | [illegible] |
| 1060961374 | PREDICTED: pleckstrin homology domain-containing family A member 6 isoform X2 [Callithrix jacchus] | 0 | 137.52 | 28 | 27 | 117 | 27 | 1 | 1296 | 145.3 | 8.32 | 1110 | 27 | 70074629.92 | 81062354.99 | [illegible] |
| 296229708 | PREDICTED: BMP/retinoic acid-inducible neural-specific protein 2 [Callithrix jacchus] | 0 | 19.878 | 8 | 6 | 14 | 5 | 1 | 783 | 88.9 | 8.03 | 188 | 6 | 16012830.67 | 14661927.88 | [illegible] |
| 296206317 | PREDICTED: probable G-protein coupled receptor 158 [Callithrix jacchus] | 0 | 386.424 | 42 | 46 | 459 | 46 | 1 | 1215 | 135.5 | 8.15 | 8820 | 46 | 399293013.1 | 359140755.8 | [illegible] |
| 296190963 | PREDICTED: LOW QUALITY PROTEIN: volume-regulated anion channel subunit LRRC8A [Callithrix jacchus | 0 | 26.744 | 10 | 7 | 23 | 7 | 1 | 810 | 94.2 | 7.94 | 118 | 7 | 5507544.306 | 5193222.219 | [illegible] |
| 532515064 | protein FAM171B precursor [Callithrix jacchus] | 0 | 67.322 | 18 | 10 | 61 | 10 | 1 | 827 | 92.1 | 8.56 | 677 | 10 | 26706346.03 | 34668059.69 | [illegible] |
| 1060943290 | PREDICTED: leucine-rich repeat-containing protein 4C [Callithrix jacchus] | 0 | 28.422 | 11 | 7 | 30 | 5 | 1 | 640 | 71.9 | 7.15 | 175 | 7 | 8483536.525 | 6464568.875 | [illegible] |
| 1060943218 | PREDICTED: UPF0606 protein KIAA1549L homolog isoform X1 [Callithrix jacchus] | 0 | 273.106 | 22 | 29 | 289 | 29 | 1 | 1896 | 204.4 | 9.14 | 6153 | 29 | 191325708.6 | 220509587.7 | [illegible] |
| 390462402 | PREDICTED: leucine zipper putative tumor suppressor 3 isoform X1 [Callithrix jacchus] | 0 | 92.726 | 23 | 14 | 106 | 14 | 1 | 703 | 74.8 | 7.4 | 1828 | 14 | 139417928.1 | 110208254.2 | [illegible] |
| 296200170 | PREDICTED: serine/threonine-protein kinase PAK 7 [Callithrix jacchus] | 0 | 169.709 | 41 | 24 | 165 | 20 | 1 | 719 | 80.9 | 7.96 | 2779 | 24 | 182329765.8 | 192718531.5 | [illegible] |
| 1060991802 | PREDICTED: syntaphilin [Callithrix jacchus] | 0 | 248.579 | 69 | 26 | 292 | 2 | 1 | 571 | 61.1 | 6.06 | 5981 | 26 | 798449.2712 | 2025023 | [illegible] |
| 532508458 | regulating synaptic membrane exocytosis protein 4 isoform 2 [Callithrix jacchus] | 0 | 38.017 | 29 | 5 | 24 | 5 | 1 | 269 | 29.3 | 5.8 | 525 | 5 | 48543230.39 | 53617280.5 | [illegible] |
| 1061011971 | PREDICTED: LOW QUALITY PROTEIN: leucine-rich repeat-containing protein 7 [Callithrix jacchus] | 0 | 616.006 | 57 | 69 | 586 | 66 | 1 | 1594 | 179.2 | 7.11 | 9636 | 69 | 706097182.6 | 680511474.1 | [illegible] |
| 1060932767 | REDICTED: LOW QUALITY PROTEIN: gamma-aminobutyric acid type B receptor subunit 2 [Callithrix jacchu | 0 | 153.888 | 28 | 22 | 258 | 22 | 1 | 1007 | 112.8 | 8.48 | 4463 | 22 | 337165864.9 | 358303470.6 | [illegible] |
| 1061010856 | PREDICTED: disks large-associated protein 3 [Callithrix jacchus] | 0 | 259.361 | 42 | 33 | 401 | 32 | 1 | 979 | 106 | 8.76 | 4901 | 33 | 298486702.7 | 289503784.7 | [illegible] |
| 296210164 | PREDICTED: actin-related protein 3B isoform X1 [Callithrix jacchus] | 0 | 123.516 | 54 | 19 | 189 | 15 | 1 | 418 | 47.6 | 5.88 | 2797 | 19 | 111667524.2 | 128772637.6 | [illegible] |
| 296196643 | PREDICTED: BTB/POZ domain-containing protein KCTD8 [Callithrix jacchus] | 0 | 53.716 | 24 | 11 | 93 | 9 | 1 | 472 | 52.3 | 8.37 | 331 | 11 | 26033158.25 | 25751854.19 | [illegible] |
| 1060962634 | EDICTED: neuronal tyrosine-phosphorylated phosphoinositide-3-kinase adapter 1 isoform X1 [Callithrix jacch | 0 | 52.544 | 15 | 11 | 38 | 11 | 1 | 845 | 88.5 | 9.6 | 669 | 11 | 20032228.19 | 20220184.25 | [illegible] |
| 532286692 | protein phosphatase 1 regulatory subunit 28 precursor [Callithrix jacchus] | 0 | 74.72 | 20 | 12 | 50 | 9 | 1 | 831 | 90.6 | 8.4 | 258 | 12 | 12324089.26 | 12058528.64 | [illegible] |
| 675648577 | PREDICTED: G protein-regulated inducer of neurite outgrowth 3 [Callithrix jacchus] | 0 | 86.37 | 26 | 15 | 57 | 15 | 1 | 776 | 82.9 | 7.64 | 1371 | 15 | 33128981.82 | 27876064.31 | [illegible] |
| 1060951723 | PREDICTED: leucine-rich repeat transmembrane neuronal protein 4 isoform X1 [Callithrix jacchus] | 0 | 35.283 | 16 | 11 | 129 | 8 | 1 | 591 | 67.2 | 8.88 | 996 | 11 | 29892331.19 | 29088380 | [illegible] |
| 296225958 | PREDICTED: histone H1x [Callithrix jacchus] | 0 | 10.583 | 10 | 2 | 7 | 2 | 1 | 214 | 22.7 | 10.65 | 48 | 2 | 7591692.605 | 7492712 | [illegible] |
| 1060954789 | PREDICTED: prickle-like protein 2 [Callithrix jacchus] | 0 | 289.884 | 52 | 35 | 281 | 34 | 1 | 844 | 95.6 | 7.36 | 5769 | 35 | 366486778.3 | 336008591.6 | [illegible] |
| 1060955071 | PREDICTED: sodium- and chloride-dependent GABA transporter 3 [Callithrix jacchus] | 0 | 16.679 | 8 | 4 | 21 | 4 | 1 | 632 | 70.5 | 6.61 | 232 | 4 | 9286787.694 | 7292192.75 | [illegible] |
| 532275048 | IQ motif and SEC7 domain-containing protein 3 isoform 1 [Callithrix jacchus] | 0 | 233.056 | 38 | 38 | 263 | 36 | 1 | 1186 | 128.4 | 6.27 | 3924 | 38 | 101789022.1 | 91126349.42 | [illegible] |
| 532345869 | plexin-A4 isoform 1 precursor [Callithrix jacchus] | 0 | 97.939 | 14 | 24 | 106 | 16 | 1 | 1894 | 212.6 | 7.05 | 687 | 24 | 46825674.17 | 43833363.75 | [illegible] |
| 296233537 | PREDICTED: leucine-rich repeat LGI family member 4 [Callithrix jacchus] | 0 | 14.787 | 8 | 4 | 16 | 4 | 1 | 538 | 59.3 | 7.66 | 146 | 4 | 4951020.689 | 4546206.125 | [illegible] |
| 532510317 | repulsive guidance molecule A isoform 3 [Callithrix jacchus] | 0 | 23.502 | 12 | 5 | 25 | 5 | 1 | 451 | 49.3 | 7.27 | 278 | 5 | 23036286.74 | 26468292 | [illegible] |
| 296216963 | PREDICTED: probable asparagine--tRNA ligase, mitochondrial isoform X1 [Callithrix jacchus] | 0 | 13.494 | 7 | 3 | 18 | 3 | 1 | 477 | 54.1 | 7.69 | 70 | 3 | 4155900.39 | 4223653.625 | [illegible] |
| 1060949591 | PREDICTED: LOW QUALITY PROTEIN: disks large-associated protein 2 [Callithrix jacchus] | 0 | 232.727 | 31 | 29 | 285 | 25 | 1 | 1117 | 124.3 | 7.01 | 5055 | 29 | 204383208.9 | 207513622.8 | [illegible] |
| 675714664 | PREDICTED: erlin-2 [Callithrix jacchus] | 0 | 46.761 | 35 | 10 | 55 | 7 | 1 | 339 | 37.8 | 5.5 | 651 | 10 | 19137275.64 | 16592671.58 | [illegible] |
| 296221904 | PREDICTED: tyrosine-protein kinase SgK223 [Callithrix jacchus] | 0 | 86.061 | 18 | 19 | 87 | 19 | 1 | 1398 | 149.3 | 7.53 | 818 | 19 | 39498882.76 | 35276501.88 | [illegible] |
| 1060966993 | PREDICTED: LOW QUALITY PROTEIN: MTSS1-like protein [Callithrix jacchus] | 0 | 72.687 | 24 | 12 | 74 | 12 | 1 | 742 | 79.4 | 7.66 | 621 | 12 | 27369499.77 | 27269866.64 | [illegible] |
| 296230957 | PREDICTED: signal-induced proliferation-associated 1-like protein 2 isoform X1 [Callithrix jacchus] | 0 | 87.57 | 14 | 21 | 97 | 19 | 1 | 1722 | 190.4 | 6.74 | 765 | 21 | 27458429.02 | 31526429.63 | [illegible] |
| 296216730 | PREDICTED: contactin-5 [Callithrix jacchus] | 0 | 91.708 | 18 | 17 | 76 | 15 | 1 | 1100 | 120.9 | 6.24 | 1384 | 17 | 44714692.59 | 19942049.11 | [illegible] |
| 296232843 | PREDICTED: noelin-2 isoform X1 [Callithrix jacchus] | 0 | 36.463 | 20 | 8 | 46 | 8 | 1 | 454 | 51.4 | 7.94 | 390 | 8 | 24259137.28 | 24652032.25 | [illegible] |
| 1061021272 | PREDICTED: pseudopodium-enriched atypical kinase 1 [Callithrix jacchus] | 0 | 41.27 | 7 | 9 | 21 | 9 | 1 | 1743 | 192.4 | 6.86 | 236 | 9 | 4322387.72 | 3628510.344 | [illegible] |
| 1061020753 | PREDICTED: neuroligin-3 [Callithrix jacchus] | 0 | 69.998 | 20 | 11 | 78 | 7 | 1 | 848 | 93.9 | 6.39 | 1284 | 11 | 31051299.31 | 30259262.34 | [illegible] |
| 532518556 | IQ motif and SEC7 domain-containing protein 2 isoform 1 [Callithrix jacchus] | 0 | 597.766 | 52 | 61 | 803 | 55 | 1 | 1488 | 162.8 | 8.56 | 14114 | 61 | 645672716.5 | 694653370.9 | [illegible] |
| 1061019675 | PREDICTED: connector enhancer of kinase suppressor of ras 2 [Callithrix jacchus] | 0 | 261.328 | 41 | 36 | 304 | 36 | 1 | 1032 | 117.3 | 6.89 | 4486 | 36 | 381830511.9 | 388655774.5 | [illegible] |
| 675665844 | PREDICTED: autophagy-related protein 9A [Callithrix jacchus] | 0 | 26.557 | 8 | 6 | 30 | 6 | 1 | 839 | 94.4 | 6.67 | 274 | 6 | 3280879.759 | 3679694.156 | [illegible] |
| 532523698 | MAP7 domain-containing protein 1 [Callithrix jacchus] | 0 | 126.008 | 32 | 23 | 177 | 23 | 1 | 840 | 92.8 | 10.08 | 2243 | 23 | 61729533.36 | 54345414.79 | [illegible] |
| 1061000538 | PREDICTED: wiskott-Aldrich syndrome protein family member 3 isoform X1 [Callithrix jacchus] | 0 | 85.273 | 34 | 13 | 96 | 9 | 1 | 502 | 55.2 | 6.43 | 1315 | 13 | 41066544.55 | 43813038.47 | [illegible] |
| 296196787 | PREDICTED: leucine-rich repeat LGI family member 2 [Callithrix jacchus] | 0 | 20.331 | 12 | 6 | 13 | 6 | 1 | 545 | 62.2 | 6.98 | 84 | 6 | 12358711.58 | 7973335.063 | [illegible] |
| 296199889 | PREDICTED: dual specificity protein phosphatase 15 isoform X1 [Callithrix jacchus] | 0 | 43.87 | 55 | 9 | 30 | 9 | 1 | 235 | 26.2 | 9.55 | 116 | 9 | 16573096.23 | 19703069 | [illegible] |
| 532523636 | ubiquitin carboxyl-terminal hydrolase 7 [Callithrix jacchus] | 0 | 85.668 | 20 | 18 | 80 | 18 | 1 | 1101 | 128.2 | 5.55 | 459 | 18 | 46532606.05 | 46466780.81 | [illegible] |
| 532504241 | WD repeat-containing protein mio [Callithrix jacchus] | 0 | 24.709 | 9 | 7 | 27 | 7 | 1 | 875 | 98.5 | 6.71 | 196 | 7 | 3047636.053 | 3654354.25 | [illegible] |
| 675636219 | PREDICTED: gamma-tubulin complex component 3 [Callithrix jacchus] | 0 | 30.325 | 9 | 8 | 39 | 8 | 1 | 907 | 103.5 | 8.13 | 128 | 8 | 10477040.2 | 12409575.13 | [illegible] |
| 532437503 | EH domain-containing protein 2 [Callithrix jacchus] | 0 | 46.775 | 22 | 12 | 51 | 10 | 1 | 543 | 61.1 | 6.46 | 319 | 12 | 35732447 | 37583793.88 | [illegible] |
| 532513710 | SLIT-ROBO Rho GTPase-activating protein 3 isoform b [Callithrix jacchus] | 0 | 367.303 | 56 | 52 | 450 | 47 | 1 | 1075 | 121.8 | 6.58 | 8968 | 52 | 287883210.1 | 292283541.8 | [illegible] |
| 1060985178 | PREDICTED: LOW QUALITY PROTEIN: proline-rich transmembrane protein 1 [Callithrix jacchus] | 0 | 24.969 | 23 | 4 | 15 | 4 | 1 | 306 | 31.5 | 7.62 | 104 | 4 | 11826391.08 | 13088452.13 | [illegible] |
| 532443759 | cell adhesion molecule 4 precursor [Callithrix jacchus] | 0 | 69.05 | 26 | 7 | 39 | 7 | 1 | 388 | 42.8 | 6.3 | 880 | 7 | 10864448.08 | 9882834.125 | [illegible] |
| 1061017498 | PREDICTED: neuron navigator 3 isoform X1 [Callithrix jacchus] | 0 | 88.638 | 11 | 23 | 65 | 22 | 1 | 2363 | 252.9 | 8.57 | 431 | 23 | 38185392.77 | 39872716.69 | [illegible] |
| 532524314 | spermatogenesis-associated protein 2 [Callithrix jacchus] | 0 | 27.469 | 16 | 6 | 12 | 6 | 1 | 520 | 58.4 | 8.66 | 34 | 6 | 7740089.736 | 6873353.438 | [illegible] |
| 1061013371 | PREDICTED: peptidyl-prolyl cis-trans isomerase A isoform X1 [Callithrix jacchus] | 0 | 106.945 | 65 | 10 | 96 | 10 | 1 | 164 | 17.9 | 8.16 | 1872 | 10 | 225776842.1 | 215008163.4 | [illegible] |
| 296202178 | PREDICTED: 60S ribosomal protein L23a [Callithrix jacchus] | 0 | 60.238 | 51 | 10 | 123 | 10 | 1 | 156 | 17.7 | 10.45 | 1202 | 10 | 136477679.3 | 140431011.6 | [illegible] |
| 532519604 | 4-aminobutyrate aminotransferase, mitochondrial precursor [Callithrix jacchus] | 0 | 49.875 | 25 | 10 | 33 | 10 | 1 | 500 | 56.4 | 7.5 | 470 | 10 | 26459140.42 | 26139979.06 | [illegible] |

| | | | | | | | | | | | |
|---|---|---|---|---|---|---|---|---|---|---|---|
| 390475482 | PREDICTED: limbic system-associated membrane protein isoform X1 [Callithrix jacchus] | 0 | 136.333 | 45 | 16 | 585 | 16 | 1 | 361 | 39.8 | [illegible] |
| 675705006 | PREDICTED: caskin-1, partial [Callithrix jacchus] | 0 | 364.031 | 46 | 46 | 451 | 44 | 1 | 1339 | 140.3 | [illegible] |
| 1060952534 | PREDICTED: striatin isoform X1 [Callithrix jacchus] | 0 | 79.487 | 24 | 15 | 61 | 14 | 1 | 809 | 89.4 | [illegible] |
| 390470649 | PREDICTED: sn1-specific diacylglycerol lipase alpha [Callithrix jacchus] | 0 | 88.392 | 24 | 20 | 99 | 20 | 1 | 1042 | 114.8 | [illegible] |
| 1061008060 | PREDICTED: inactive dipeptidyl peptidase 10 [Callithrix jacchus] | 0 | 33.911 | 12 | 9 | 36 | 8 | 1 | 800 | 91.3 | [illegible] |
| 1060951215 | PREDICTED: type I inositol 3,4-bisphosphate 4-phosphatase isoform X1 [Callithrix jacchus] | 0 | 91.853 | 22 | 20 | 86 | 20 | 1 | 1005 | 113.1 | [illegible] |
| 675666958 | PREDICTED: protein FAM171A1 [Callithrix jacchus] | 0 | 131.113 | 34 | 25 | 104 | 24 | 1 | 890 | 97.9 | [illegible] |
| 1060935806 | PREDICTED: 60S ribosomal protein L12 [Callithrix jacchus] | 0 | 87.69 | 45 | 9 | 121 | 9 | 1 | 233 | 25.5 | [illegible] |
| 296218252 | PREDICTED: reticulon-4 receptor-like 2 [Callithrix jacchus] | 0 | 46.382 | 28 | 11 | 96 | 11 | 1 | 420 | 46.1 | [illegible] |
| 296238032 | PREDICTED: vam6/Vps39-like protein isoform X1 [Callithrix jacchus] | 0 | 74.1 | 21 | 16 | 86 | 16 | 1 | 886 | 101.8 | [illegible] |
| 1060934952 | PREDICTED: transitional endoplasmic reticulum ATPase [Callithrix jacchus] | 0 | 289.233 | 60 | 39 | 301 | 39 | 1 | 806 | 89.3 | [illegible] |
| 1061010620 | PREDICTED: protein 4.1 isoform X1 [Callithrix jacchus] | 0 | 67.908 | 17 | 13 | 110 | 9 | 1 | 1016 | 113 | [illegible] |
| 1061010331 | PREDICTED: leucine zipper protein 1 [Callithrix jacchus] | 0 | 110.401 | 29 | 27 | 95 | 27 | 1 | 1074 | 120 | [illegible] |
| 1061009758 | PREDICTED: pantothenate kinase 4 [Callithrix jacchus] | 0 | 54.022 | 17 | 11 | 45 | 11 | 1 | 781 | 86.9 | [illegible] |
| 296196931 | PREDICTED: protein phosphatase 2, regulatory subunit B, gamma isoform X4 [Callithrix jacchus] | 0 | 28.278 | 13 | 6 | 33 | 3 | 1 | 447 | 51.3 | [illegible] |
| 532506300 | Golgin subfamily A member 3 isoform 1 [Callithrix jacchus] | 0 | 168.415 | 27 | 35 | 153 | 4 | 1 | 1503 | 167.5 | [illegible] |
| 675683637 | PREDICTED: LOW QUALITY PROTEIN: probable E3 ubiquitin-protein ligase HECTD4 [Callithrix jacchus] | 0 | 292.043 | 18 | 64 | 247 | 64 | 1 | 4418 | 484.3 | [illegible] |
| 532516114 | AP2-associated protein kinase 1 [Callithrix jacchus] | 0 | 409.778 | 55 | 41 | 431 | 41 | 1 | 964 | 104.2 | [illegible] |
| 675729820 | PREDICTED: contactin-4 isoform X1 [Callithrix jacchus] | 0 | 66.153 | 14 | 13 | 63 | 10 | 1 | 1026 | 113.4 | [illegible] |
| 532274852 | mitogen-activated protein kinase kinase kinase 10 [Callithrix jacchus] | 0 | 13.81 | 5 | 4 | 11 | 4 | 1 | 955 | 103.9 | [illegible] |
| 532200161 | isocitrate dehydrogenase [NADP], mitochondrial precursor [Callithrix jacchus] | 0 | 153.77 | 52 | 23 | 197 | 23 | 1 | 452 | 50.7 | [illegible] |
| 675707025 | PREDICTED: germ cell-specific gene 1-like protein [Callithrix jacchus] | 0 | 28.938 | 17 | 7 | 32 | 7 | 1 | 332 | 36.9 | [illegible] |
| 532509638 | microtubule-associated protein 1S [Callithrix jacchus] | 0 | 117.502 | 25 | 18 | 90 | 18 | 1 | 1063 | 112.6 | [illegible] |
| 675758416 | PREDICTED: glutaryl-CoA dehydrogenase, mitochondrial isoform X1 [Callithrix jacchus] | 0 | 26.567 | 14 | 5 | 25 | 5 | 1 | 438 | 48.3 | [illegible] |
| 390477946 | PREDICTED: synaptic vesicle membrane protein VAT-1 homolog-like [Callithrix jacchus] | 0.002 | 5.569 | 5 | 2 | 3 | 2 | 1 | 419 | 45.8 | [illegible] |
| 532523784 | 60S ribosomal protein L13 isoform 1 [Callithrix jacchus] | 0 | 62.466 | 36 | 10 | 271 | 10 | 1 | 211 | 24.3 | [illegible] |
| 675694591 | PREDICTED: NLR family member X1 [Callithrix jacchus] | 0 | 65.74 | 23 | 15 | 60 | 15 | 1 | 794 | 87.9 | [illegible] |
| 675693481 | PREDICTED: ELMO domain-containing protein 1 isoform X1 [Callithrix jacchus] | 0 | 47.566 | 28 | 8 | 36 | 8 | 1 | 326 | 38 | [illegible] |
| 532333787 | unconventional myosin-IXa [Callithrix jacchus] | 0 | 41.306 | 6 | 14 | 48 | 14 | 1 | 2548 | 292.6 | [illegible] |
| 296228059 | PREDICTED: ras-related protein Rab-6B [Callithrix jacchus] | 0 | 41.45 | 41 | 9 | 72 | 2 | 1 | 208 | 23.4 | [illegible] |
| 532338109 | vacuolar protein sorting-associated protein 13A isoform A [Callithrix jacchus] | 0 | 36.138 | 4 | 11 | 33 | 11 | 1 | 3175 | 359.9 | [illegible] |
| 1060980077 | PREDICTED: TBC domain-containing protein kinase-like protein isoform X1 [Callithrix jacchus] | 0 | 13.488 | 7 | 6 | 10 | 6 | 1 | 909 | 102.4 | [illegible] |
| 296234437 | PREDICTED: leucine-rich repeat-containing protein 4B [Callithrix jacchus] | 0 | 58.699 | 18 | 10 | 58 | 8 | 1 | 711 | 76.4 | [illegible] |
| 1061002467 | PREDICTED: probable threonine--tRNA ligase 2, cytoplasmic [Callithrix jacchus] | 0 | 32.118 | 12 | 9 | 27 | 9 | 1 | 808 | 93.2 | [illegible] |
| 532508648 | eukaryotic translation initiation factor 5A-1 isoform A [Callithrix jacchus] | 0 | 28.268 | 27 | 4 | 34 | 4 | 1 | 184 | 20.2 | [illegible] |
| 1060990609 | LOW QUALITY PROTEIN: phosphatidylinositol 3,4,5-trisphosphate-dependent Rac exchanger 1 protein [Cal | 0 | 60.317 | 9 | 13 | 66 | 13 | 1 | 1706 | 191.2 | [illegible] |
| 1060948927 | PREDICTED: 60S ribosomal protein L17 [Callithrix jacchus] | 0 | 75.48 | 51 | 10 | 140 | 10 | 1 | 184 | 21.4 | [illegible] |
| 532345129 | UPF0515 protein C19orf66 [Callithrix jacchus] | 0 | 38.362 | 33 | 8 | 27 | 8 | 1 | 292 | 33.3 | [illegible] |
| 1060981622 | PREDICTED: adhesion G protein-coupled receptor L3 isoform X1 [Callithrix jacchus] | 0 | 212.968 | 27 | 35 | 183 | 32 | 1 | 1586 | 176.7 | [illegible] |
| 296212220 | PREDICTED: protein phosphatase 1H [Callithrix jacchus] | 0 | 40.982 | 24 | 9 | 44 | 9 | 1 | 514 | 56.4 | [illegible] |
| 675673847 | PREDICTED: neuronal cell adhesion molecule isoform X1 [Callithrix jacchus] | 0 | 129.155 | 22 | 23 | 149 | 23 | 1 | 1311 | 144.5 | [illegible] |
| 1060957316 | PREDICTED: syntabulin isoform X2 [Callithrix jacchus] | 0 | 49.196 | 20 | 12 | 63 | 10 | 1 | 663 | 72.5 | [illegible] |
| 296196740 | PREDICTED: NACHT and WD repeat domain-containing protein 2 isoform X1 [Callithrix jacchus] | 0 | 305.386 | 32 | 46 | 269 | 46 | 1 | 1742 | 197.4 | [illegible] |
| 296189785 | PREDICTED: amyloid beta A4 precursor protein-binding family A member 1 isoform X1 [Callithrix jacchus] | 0 | 36.653 | 11 | 7 | 41 | 6 | 1 | 837 | 92.9 | [illegible] |
| 1060942790 | PREDICTED: myotubularin-related protein 13 isoform X1 [Callithrix jacchus] | 0 | 10.744 | 2 | 3 | 9 | 2 | 1 | 1881 | 212.1 | [illegible] |
| 296214588 | PREDICTED: junctophilin-4 [Callithrix jacchus] | 0 | 12.934 | 5 | 3 | 8 | 3 | 1 | 628 | 65.9 | [illegible] |
| 1061015071 | PREDICTED: diacylglycerol kinase iota isoform X4 [Callithrix jacchus] | 0 | 154.067 | 29 | 22 | 141 | 21 | 1 | 1005 | 111.7 | [illegible] |
| 390473699 | REDICTED: LOW QUALITY PROTEIN: transforming acidic coiled-coil-containing protein 1 [Callithrix jacchus | 0 | 21.578 | 7 | 6 | 27 | 6 | 1 | 807 | 87.8 | [illegible] |
| 532511346 | protein Hook homolog 3 [Callithrix jacchus] | 0 | 33.655 | 14 | 9 | 23 | 9 | 1 | 720 | 83.3 | [illegible] |
| 675639363 | PREDICTED: protein SCAI isoform X1 [Callithrix jacchus] | 0 | 265.294 | 56 | 28 | 328 | 28 | 1 | 606 | 70.4 | [illegible] |
| 532275556 | protein furry homolog [Callithrix jacchus] | 0 | 146.013 | 12 | 31 | 139 | 31 | 1 | 3022 | 339.5 | [illegible] |
| 532516884 | calcium-dependent secretion activator 2 isoform c [Callithrix jacchus] | 0 | 98.182 | 20 | 21 | 71 | 16 | 1 | 1306 | 148.6 | [illegible] |
| 532273056 | protein phosphatase 1E [Callithrix jacchus] | 0 | 32.013 | 8 | 6 | 30 | 6 | 1 | 761 | 84.8 | [illegible] |
| 1060952521 | PREDICTED: HEAT repeat-containing protein 5B isoform X1 [Callithrix jacchus] | 0 | 25.267 | 3 | 5 | 10 | 5 | 1 | 2091 | 226.4 | [illegible] |
| 675699452 | PREDICTED: T-complex protein 11-like protein 1 isoform X1 [Callithrix jacchus] | 0 | 69.422 | 35 | 13 | 49 | 13 | 1 | 509 | 57 | [illegible] |
| 675769733 | PREDICTED: inositol polyphosphate 5-phosphatase OCRL-1 isoform X6 [Callithrix jacchus] | 0 | 11.754 | 4 | 4 | 10 | 4 | 1 | 1015 | 115.9 | [illegible] |
| 1060939556 | EDICTED: MAM domain-containing glycosylphosphatidylinositol anchor protein 2 isoform X1 [Callithrix jacch | 0 | 59.565 | 16 | 14 | 80 | 14 | 1 | 1025 | 115.2 | [illegible] |
| 296208258 | PREDICTED: neuronal growth regulator 1 isoform X1 [Callithrix jacchus] | 0 | 169.423 | 49 | 16 | 496 | 16 | 1 | 354 | 38.7 | [illegible] |
| 532513182 | cadherin-18 isoform 1 preproprotein [Callithrix jacchus] | 0 | 18.773 | 7 | 4 | 10 | 4 | 1 | 790 | 87.9 | [illegible] |
| 532509605 | cadherin-10 preproprotein [Callithrix jacchus] | 0 | 52.473 | 17 | 12 | 80 | 11 | 1 | 788 | 88.4 | [illegible] |
| 296195082 | PREDICTED: ectonucleotide pyrophosphatase/phosphodiesterase family member 6 [Callithrix jacchus] | 0 | 150.251 | 38 | 13 | 170 | 13 | 1 | 429 | 48.7 | [illegible] |
| 532515318 | tropomyosin alpha-4 chain isoform 2 [Callithrix jacchus] | 0 | 163.019 | 71 | 25 | 346 | 12 | 1 | 248 | 28.5 | [illegible] |
| 675681683 | PREDICTED: liprin-alpha-2 isoform X2 [Callithrix jacchus] | 0 | 384.028 | 51 | 61 | 476 | 41 | 1 | 1257 | 143.2 | [illegible] |
| 1061017371 | PREDICTED: fibroblast growth factor receptor substrate 2 [Callithrix jacchus] | 0 | 11.878 | 8 | 4 | 12 | 3 | 1 | 508 | 57.1 | [illegible] |
| 532513610 | glucose-fructose oxidoreductase domain-containing protein 1 isoform 1 precursor [Callithrix jacchus] | 0 | 28.432 | 19 | 7 | 70 | 7 | 1 | 390 | 43.1 | [illegible] |
| 296205138 | PREDICTED: E3 ubiquitin-protein ligase HECW2 [Callithrix jacchus] | 0 | 131.496 | 19 | 25 | 143 | 23 | 1 | 1567 | 175.3 | [illegible] |
| 675665179 | PREDICTED: abl interactor 2 isoform X1 [Callithrix jacchus] | 0 | 103.078 | 37 | 15 | 127 | 3 | 1 | 542 | 58.7 | [illegible] |
| 675748618 | PREDICTED: protein phosphatase 1 regulatory subunit 12B isoform X1 [Callithrix jacchus] | 0 | 114.988 | 23 | 24 | 147 | 24 | 1 | 1062 | 118.5 | [illegible] |
| 675738455 | PREDICTED: protein C3orf33 homolog [Callithrix jacchus] | 0 | 16.803 | 15 | 4 | 12 | 4 | 1 | 301 | 34.8 | [illegible] |
| 296213054 | PREDICTED: serine/threonine-protein kinase TAO3 isoform X1 [Callithrix jacchus] | 0 | 68.627 | 19 | 16 | 89 | 11 | 1 | 898 | 105.4 | [illegible] |
| 1061015040 | PREDICTED: UPF0606 protein KIAA1549 homolog isoform X1 [Callithrix jacchus] | 0 | 165.659 | 15 | 24 | 171 | 24 | 1 | 1910 | 206.4 | [illegible] |
| 532513434 | F-box only protein 41 [Callithrix jacchus] | 0 | 285.583 | 51 | 39 | 304 | 39 | 1 | 876 | 94.5 | [illegible] |
| 675712635 | PREDICTED: dedicator of cytokinesis protein 1 isoform X1 [Callithrix jacchus] | 0 | 13.572 | 2 | 5 | 8 | 5 | 1 | 1886 | 217.7 | [illegible] |
| 532276932 | hyaluronan and proteoglycan link protein 4 precursor [Callithrix jacchus] | 0 | 166.627 | 63 | 21 | 101 | 20 | 1 | 402 | 42.8 | [illegible] |
| 532518610 | latrophilin-1 isoform 1 precursor [Callithrix jacchus] | 0 | 234.965 | 32 | 36 | 171 | 34 | 1 | 1481 | 163.3 | [illegible] |
| 532524006 | opioid-binding protein/cell adhesion molecule isoform a preproprotein [Callithrix jacchus] | 0 | 131.858 | 39 | 14 | 401 | 2 | 1 | 354 | 38.8 | [illegible] |
| 1060944749 | PREDICTED: rho GTPase-activating protein 32 [Callithrix jacchus] | 0 | 347.288 | 40 | 58 | 417 | 56 | 1 | 2087 | 230.2 | [illegible] |
| 1061019343 | PREDICTED: interleukin-1 receptor accessory protein-like 1 [Callithrix jacchus] | 0 | 42.544 | 16 | 9 | 37 | 9 | 1 | 696 | 79.9 | [illegible] |
| 1061018779 | PREDICTED: kinase suppressor of Ras 2 [Callithrix jacchus] | 0 | 23.791 | 8 | 7 | 24 | 6 | 1 | 1003 | 113.4 | [illegible] |
| 675774283 | PREDICTED: FERM and PDZ domain-containing protein 4 isoform X1 [Callithrix jacchus] | 0 | 47.391 | 8 | 10 | 30 | 10 | 1 | 1761 | 192.8 | [illegible] |
| 532510221 | FK506-binding protein 15 [Callithrix jacchus] | 0 | 60.175 | 12 | 11 | 58 | 11 | 1 | 1219 | 133.6 | [illegible] |
| 1060960998 | PREDICTED: LOW QUALITY PROTEIN: melanoma inhibitory activity protein 3 [Callithrix jacchus] | 0 | 77.802 | 11 | 15 | 51 | 13 | 1 | 1837 | 205.2 | [illegible] |
| 675659568 | PREDICTED: unconventional myosin-Id isoform X1 [Callithrix jacchus] | 0 | 352.858 | 58 | 57 | 416 | 57 | 1 | 1006 | 116.1 | [illegible] |
| 1060932657 | PREDICTED: LOW QUALITY PROTEIN: sodium leak channel non-selective protein [Callithrix jacchus] | 0 | 11.795 | 2 | 3 | 5 | 3 | 1 | 1938 | 222.5 | [illegible] |
| 1060938280 | PREDICTED: enhancer of mRNA-decapping protein 3 [Callithrix jacchus] | 0 | 39.085 | 23 | 8 | 23 | 8 | 1 | 508 | 56 | [illegible] |
| 1060944409 | PREDICTED: 39S ribosomal protein L21, mitochondrial isoform X1 [Callithrix jacchus] | 0 | 27.087 | 25 | 4 | 24 | 4 | 1 | 201 | 22.8 | [illegible] |
| 675743700 | PREDICTED: tripartite motif-containing protein 46 [Callithrix jacchus] | 0 | 83.618 | 28 | 18 | 95 | 18 | 1 | 759 | 83.4 | [illegible] |
| 296199361 | PREDICTED: LOW QUALITY PROTEIN: headcase protein homolog [Callithrix jacchus] | 0 | 8.974 | 5 | 2 | 3 | 2 | 1 | 543 | 58.8 | [illegible] |
| 1060932865 | PREDICTED: neurofilament heavy polypeptide [Callithrix jacchus] | 0 | 190.244 | 38 | 31 | 245 | 27 | 1 | 893 | 97.4 | [illegible] |
| 675721349 | PREDICTED: CB1 cannabinoid receptor-interacting protein 1 isoform X1 [Callithrix jacchus] | 0 | 111.271 | 73 | 9 | 127 | 9 | 1 | 164 | 18.6 | [illegible] |
| 675725023 | PREDICTED: protein TSSC1 isoform X2 [Callithrix jacchus] | 0 | 34.342 | 22 | 6 | 26 | 6 | 1 | 387 | 43.6 | [illegible] |
| 675691063 | PREDICTED: LOW QUALITY PROTEIN: coiled-coil domain-containing protein 177 [Callithrix jacchus] | 0 | 87.374 | 24 | 19 | 130 | 19 | 1 | 704 | 79.7 | [illegible] |
| 296215621 | PREDICTED: SRA stem-loop-interacting RNA-binding protein, mitochondrial [Callithrix jacchus] | 0 | 29.361 | 62 | 6 | 25 | 6 | 1 | 108 | 12.3 | [illegible] |
| 532512068 | brain-enriched guanylate kinase-associated protein [Callithrix jacchus] | 0 | 149.831 | 49 | 23 | 184 | 23 | 1 | 593 | 64.8 | [illegible] |
| 532516510 | LIM domain only protein 7 isoform 2 [Callithrix jacchus] | 0 | 362.725 | 50 | 61 | 396 | 2 | 1 | 1384 | 158.2 | [illegible] |
| 532510107 | MAP6 domain-containing protein 1 [Callithrix jacchus] | 0 | 57.129 | 63 | 8 | 42 | 5 | 1 | 195 | 20.8 | [illegible] |
| 296222484 | PREDICTED: GRB2-associated and regulator of MAPK protein 1 isoform X1 [Callithrix jacchus] | 0 | 21.491 | 7 | 5 | 10 | 5 | 1 | 876 | 97 | [illegible] |
| 296230484 | PREDICTED: plasma membrane calcium-transporting ATPase 4 isoform X3 [Callithrix jacchus] | 0 | 293.735 | 30 | 31 | 291 | 14 | 1 | 1169 | 129.2 | [illegible] |
| 296230264 | PREDICTED: protein stum homolog [Callithrix jacchus] | 0 | 8.867 | 21 | 2 | 19 | 2 | 1 | 141 | 15 | [illegible] |
| 675640140 | PREDICTED: uncharacterized protein C9orf172 homolog [Callithrix jacchus] | 0 | 236.822 | 51 | 38 | 285 | 38 | 1 | 976 | 106.6 | [illegible] |
| 1060956639 | PREDICTED: pyruvate dehyrogenase phosphatase catalytic subunit 1 isoform X1 [Callithrix jacchus] | 0 | 50.326 | 17 | 8 | 35 | 8 | 1 | 596 | 67.4 | [illegible] |
| 296206491 | PREDICTED: transmembrane protein 240 [Callithrix jacchus] | 0 | 34.812 | 44 | 6 | 38 | 6 | 1 | 173 | 19.9 | [illegible] |
| 1060962573 | PREDICTED: neurosecretory protein VGF [Callithrix jacchus] | 0 | 97.798 | 34 | 17 | 85 | 17 | 1 | 616 | 67.5 | [illegible] |
| 675674886 | PREDICTED: single-stranded DNA-binding protein, mitochondrial isoform X2 [Callithrix jacchus] | 0 | 78.132 | 59 | 9 | 80 | 9 | 1 | 148 | 17.1 | [illegible] |
| 296211116 | PREDICTED: protein arginine N-methyltransferase 8 isoform X1 [Callithrix jacchus] | 0 | 82.704 | 37 | 15 | 105 | 12 | 1 | 394 | 45.3 | [illegible] |
| 532448845 | serine/threonine-protein kinase TAO2 isoform 2 [Callithrix jacchus] | 0 | 184.178 | 29 | 29 | 194 | 26 | 1 | 1049 | 119.2 | [illegible] |
| 532514220 | serine/threonine-protein kinase BRSK1 [Callithrix jacchus] | 0 | 181.432 | 45 | 24 | 172 | 18 | 1 | 778 | 85.1 | [illegible] |
| 675762730 | PREDICTED: serine/threonine-protein kinase LMTK3 [Callithrix jacchus] | 0 | 149.04 | 23 | 24 | 164 | 23 | 1 | 1454 | 153.5 | [illegible] |
| 532519246 | CREB-regulated transcription coactivator 1 isoform 3 [Callithrix jacchus] | 0 | 14.562 | 6 | 2 | 6 | 2 | 1 | 650 | 68.7 | [illegible] |
| 1061019725 | PREDICTED: cyclin-dependent kinase-like 5 [Callithrix jacchus] | 0 | 198.483 | 44 | 35 | 258 | 34 | 1 | 960 | 107.6 | [illegible] |
| 296194247 | PREDICTED: metaxin-3 isoform X1 [Callithrix jacchus] | 0 | 58.081 | 40 | 12 | 135 | 12 | 1 | 312 | 35.1 | [illegible] |
| 1060957420 | PREDICTED: ribonucleoside-diphosphate reductase subunit M2 B isoform X1 [Callithrix jacchus] | 0 | 9.66 | 7 | 3 | 9 | 3 | 1 | 423 | 48.2 | [illegible] |
| 675643520 | PREDICTED: BTB/POZ domain-containing protein KCTD16 [Callithrix jacchus] | 0 | 138.031 | 49 | 24 | 272 | 22 | 1 | 428 | 49.1 | [illegible] |
| 532450625 | E3 ubiquitin-protein ligase ZNRF2 [Callithrix jacchus] | 0 | 11.748 | 14 | 3 | 14 | 2 | 1 | 242 | 24.1 | [illegible] |
| 296215663 | PREDICTED: leucine-rich repeat transmembrane protein FLRT2 [Callithrix jacchus] | 0 | 61.321 | 24 | 13 | 66 | 13 | 1 | 660 | 74 | [illegible] |
| 1060966333 | PREDICTED: NADH dehydrogenase [ubiquinone] iron-sulfur protein 6, mitochondrial [Callithrix jacchus] | 0 | 68.081 | 60 | 7 | 87 | 7 | 1 | 124 | 13.6 | [illegible] |
| 1061015198 | PREDICTED: adenylate cyclase type 1 [Callithrix jacchus] | 0 | 48.869 | 15 | 12 | 64 | 12 | 1 | 914 | 101.6 | [illegible] |
| 1060994185 | PREDICTED: cytospin-B isoform X1 [Callithrix jacchus] | 0 | 93.652 | 22 | 21 | 106 | 21 | 1 | 1106 | 122.5 | [illegible] |
| 296193482 | PREDICTED: proline-rich protein 7 [Callithrix jacchus] | 0 | 28.515 | 25 | 5 | 29 | 5 | 1 | 272 | 30.7 | [illegible] |
| 296206890 | PREDICTED: MICOS complex subunit MIC10 [Callithrix jacchus] | 0.003 | 4.691 | 19 | 2 | 9 | 2 | 1 | 78 | 8.8 | [illegible] |
| 296196581 | PREDICTED: OCIA domain-containing protein 2 isoform X1 [Callithrix jacchus] | 0 | 29.851 | 54 | 8 | 28 | 8 | 1 | 154 | 16.9 | [illegible] |
| 532506465 | transmembrane protein 178B precursor [Callithrix jacchus] | 0 | 23.162 | 15 | 4 | 23 | 4 | 1 | 294 | 33.4 | [illegible] |
| 296234179 | PREDICTED: PNMA-like protein 2 [Callithrix jacchus] | 0 | 10.082 | 5 | 3 | 17 | 3 | 1 | 635 | 68.2 | [illegible] |
| 675733153 | PREDICTED: XK-related protein 4 [Callithrix jacchus] | 0 | 38.566 | 11 | 7 | 50 | 7 | 1 | 651 | 71.5 | [illegible] |
| 532289440 | kalirin isoform 1 [Callithrix jacchus] | 0 | 714.421 | 42 | 106 | 632 | 32 | 1 | 2985 | 340 | [illegible] |
| 675750189 | PREDICTED: centrosomal protein of 170 kDa isoform X4 [Callithrix jacchus] | 0 | 309.784 | 39 | 48 | 319 | 46 | 1 | 1558 | 172.1 | [illegible] |
| 1060959745 | PREDICTED: ankyrin repeat domain-containing protein 34A [Callithrix jacchus] | 0 | 42.813 | 25 | 9 | 43 | 9 | 1 | 496 | 52.6 | [illegible] |
| 1060973483 | PREDICTED: microtubule-associated serine/threonine-protein kinase 3, partial [Callithrix jacchus] | 0 | 152.489 | 24 | 21 | 134 | 18 | 1 | 1314 | 143.8 | [illegible] |
| 296207566 | PREDICTED: NADH dehydrogenase [ubiquinone] iron-sulfur protein 5 [Callithrix jacchus] | 0 | 80.678 | 74 | 10 | 74 | 10 | 1 | 106 | 12.8 | [illegible] |
| 675761697 | PREDICTED: glycogen synthase kinase-3 alpha [Callithrix jacchus] | 0 | 89.101 | 36 | 13 | 134 | 9 | 1 | 483 | 51 | [illegible] |
| 1060972621 | PREDICTED: phospholipid hydroperoxide glutathione peroxidase, mitochondrial isoform X1 [Callithrix jacchus | 0 | 75.436 | 40 | 11 | 115 | 11 | 1 | 234 | 26.9 | [illegible] |
| 1060974690 | PREDICTED: dapper homolog 3 [Callithrix jacchus] | 0 | 60.787 | 24 | 12 | 69 | 12 | 1 | 631 | 65.1 | [illegible] |
| 675745084 | PREDICTED: mitochondrial import receptor subunit TOM40B [Callithrix jacchus] | 0 | 44.607 | 34 | 8 | 31 | 8 | 1 | 308 | 33.9 | [illegible] |
| 675737555 | PREDICTED: TRAF2 and NCK-interacting protein kinase [Callithrix jacchus] | 0 | 249.948 | 33 | 39 | 307 | 29 | 1 | 1360 | 154.8 | [illegible] |
| 1060952867 | PREDICTED: protein EFR3 homolog B [Callithrix jacchus] | 0 | 163.75 | 33 | 26 | 156 | 25 | 1 | 888 | 100.3 | [illegible] |
| 675746776 | PREDICTED: LOW QUALITY PROTEIN: immunoglobulin-like domain-containing receptor 2 [Callithrix jacchus | 0 | 141.475 | 39 | 23 | 165 | 23 | 1 | 636 | 70.7 | [illegible] |
| 1061019732 | PREDICTED: 40S ribosomal protein S4, X isoform [Callithrix jacchus] | 0 | 122.95 | 54 | 20 | 217 | 20 | 1 | 335 | 37.7 | [illegible] |
| 675671249 | PREDICTED: uncharacterized protein KIAA1107 homolog isoform X1 [Callithrix jacchus] | 0 | 204.648 | 22 | 33 | 219 | 33 | 1 | 1776 | 197.4 | [illegible] |
| 296208534 | PREDICTED: 60S ribosomal protein L5 isoform X1 [Callithrix jacchus] | 0 | 88.122 | 35 | 13 | 133 | 13 | 1 | 297 | 34.4 | [illegible] |
| 296205168 | PREDICTED: 10 kDa heat shock protein, mitochondrial isoform X4 [Callithrix jacchus] | 0 | 23.3 | 53 | 5 | 41 | 5 | 1 | 102 | 10.9 | [illegible] |
| 675706185 | PREDICTED: 40S ribosomal protein S20, partial [Callithrix jacchus] | 0 | 38.187 | 28 | 6 | 55 | 6 | 1 | 138 | 15.2 | [illegible] |
| 296208015 | PREDICTED: 40S ribosomal protein SA [Callithrix jacchus] | 0 | 113.817 | 55 | 12 | 90 | 12 | 1 | 295 | 32.8 | [illegible] |
| 296210617 | PREDICTED: 60S ribosomal protein L32 [Callithrix jacchus] | 0 | 58.745 | 54 | 11 | 75 | 11 | 1 | 135 | 15.9 | [illegible] |
| 1060938502 | PREDICTED: ferritin heavy chain [Callithrix jacchus] | 0 | 69.124 | 54 | 10 | 74 | 10 | 1 | 183 | 21.3 | [illegible] |
| 296212580 | PREDICTED: ubiquitin-conjugating enzyme E2 N isoform X1 [Callithrix jacchus] | 0 | 32.38 | 57 | 7 | 30 | 7 | 1 | 152 | 17.1 | [illegible] |
| 296196229 | PREDICTED: 60S ribosomal protein L7a isoform X1 [Callithrix jacchus] | 0 | 143.739 | 42 | 17 | 208 | 17 | 1 | 266 | 30 | [illegible] |
| 675753928 | PREDICTED: LOW QUALITY PROTEIN: tubulin beta-3 chain-like [Callithrix jacchus] | 0 | 431.426 | 40 | 25 | 1126 | 2 | 1 | 799 | 88.7 | [illegible] |
| 296228641 | PREDICTED: histone H2A type 2-A [Callithrix jacchus] | 0 | 28.094 | 49 | 4 | 28 | 2 | 1 | 130 | 14.1 | [illegible] |
| 532285854 | kazrin isoform E [Callithrix jacchus] | 0 | 23.995 | 7 | 5 | 13 | 5 | 1 | 775 | 86.3 | [illegible] |
| 167427222 | ribosomal protein S25 (predicted) [Callithrix jacchus] | 0 | 22.317 | 30 | 5 | 70 | 5 | 1 | 125 | 13.8 | [illegible] |
| 532342253 | tyrosine-protein phosphatase non-receptor type substrate 1 precursor [Callithrix jacchus] | 0 | 122.063 | 39 | 16 | 177 | 16 | 1 | 506 | 54.9 | [illegible] |
| 296217690 | PREDICTED: ATP synthase subunit beta, mitochondrial [Callithrix jacchus] | 0 | 218.691 | 40 | 17 | 199 | 3 | 1 | 529 | 56.4 | [illegible] |
| 1060931222 | PREDICTED: LOW QUALITY PROTEIN: myelin-associated glycoprotein [Callithrix jacchus] | 0 | 135.763 | 31 | 15 | 164 | 15 | 1 | 626 | 69.1 | [illegible] |
| 1060931379 | PREDICTED: ADP/ATP translocase 3 [Callithrix jacchus] | 0 | 224.193 | 74 | 25 | 561 | 10 | 1 | 298 | 32.8 | [illegible] |
| 296228277 | PREDICTED: glyceraldehyde-3-phosphate dehydrogenase [Callithrix jacchus] | 0 | 280.684 | 71 | 19 | 311 | 5 | 1 | 335 | 36.1 | [illegible] |
| 1060943807 | PREDICTED: ATP synthase subunit g, mitochondrial [Callithrix jacchus] | 0 | 10.774 | 21 | 2 | 23 | 2 | 1 | 120 | 13.4 | [illegible] |
| 675646566 | PREDICTED: LOW QUALITY PROTEIN: eukaryotic initiation factor 4A-II-like [Callithrix jacchus] | 0 | 86.955 | 38 | 14 | 108 | 3 | 1 | 408 | 46.5 | [illegible] |
| 1060983585 | PREDICTED: uncharacterized protein LOC100406435 [Callithrix jacchus] | 0 | 45.014 | 34 | 8 | 98 | 7 | 1 | 340 | 37.7 | [illegible] |
| 1061020833 | PREDICTED: creatine kinase B-type [Callithrix jacchus] | 0 | 181.016 | 57 | 15 | 227 | 15 | 1 | 419 | 46.9 | [illegible] |
| 296206714 | PREDICTED: 40S ribosomal protein S11-like [Callithrix jacchus] | 0 | 66.139 | 53 | 10 | 117 | 10 | 1 | 158 | 18.4 | [illegible] |
| 1060930458 | PREDICTED: heat shock protein HSP 90-beta-like [Callithrix jacchus] | 0 | 133.511 | 52 | 20 | 222 | 5 | 1 | 337 | 39.8 | [illegible] |
| 1060946011 | PREDICTED: ADP-ribosylation factor-like protein 6-interacting protein 1 isoform X1 [Callithrix jacchus] | 0 | 63.788 | 32 | 9 | 109 | 9 | 1 | 296 | 33.9 | [illegible] |
| 675687221 | PREDICTED: LOW QUALITY PROTEIN: heat shock protein HSP 90-beta-like [Callithrix jacchus] | 0 | 216.919 | 33 | 27 | 317 | 9 | 1 | 723 | 82.7 | [illegible] |
| 296222312 | PREDICTED: myosin regulatory light polypeptide 9 isoform X1 [Callithrix jacchus] | 0 | 136.401 | 52 | 10 | 134 | 10 | 1 | 205 | 23.5 | [illegible] |
| 296217282 | PREDICTED: hemoglobin subunit beta [Callithrix jacchus] | 0 | 128.668 | 95 | 13 | 108 | 12 | 1 | 147 | 16.1 | [illegible] |
| 675737987 | PREDICTED: LOW QUALITY PROTEIN: heat shock protein HSP 90-alpha-like [Callithrix jacchus] | 0 | 257.679 | 44 | 33 | 360 | 28 | 1 | 736 | 85.2 | [illegible] |
| 296189334 | PREDICTED: 60S ribosomal protein L30-like [Callithrix jacchus] | 0 | 60.057 | 67 | 9 | 100 | 9 | 1 | 115 | 12.8 | [illegible] |
| 326368312 | paralemmin-2 [Callithrix jacchus] | 0 | 100.296 | 40 | 17 | 167 | 8 | 1 | 413 | 45.8 | [illegible] |
| 675671819 | PREDICTED: elongation factor 1-alpha 1 [Callithrix jacchus] | 0 | 166.307 | 56 | 17 | 304 | 9 | 1 | 462 | 50.1 | [illegible] |
| 532516200 | hemoglobin subunit alpha [Callithrix jacchus] | 0 | 46.986 | 69 | 8 | 72 | 8 | 1 | 142 | 15.3 | [illegible] |
| 532452937 | tubulin beta chain [Callithrix jacchus] | 0 | 632.13 | 82 | 32 | 1678 | 6 | 1 | 444 | 49.6 | [illegible] |
| 675670214 | PREDICTED: mapk-regulated corepressor-interacting protein 1-like, partial [Callithrix jacchus] | 0 | 26.723 | 48 | 5 | 28 | 5 | 1 | 116 | 12.8 | [illegible] |
| 1060932811 | PREDICTED: noelin-like [Callithrix jacchus] | 0 | 126.716 | 25 | 10 | 149 | 10 | 1 | 468 | 52.9 | [illegible] |
| 1060932648 | PREDICTED: 40S ribosomal protein S27-like [Callithrix jacchus] | 0 | 37.271 | 30 | 5 | 59 | 3 | 1 | 115 | 13 | [illegible] |
| 1060987816 | PREDICTED: 60S ribosomal protein L31 [Callithrix jacchus] | 0 | 27.619 | 38 | 6 | 89 | 6 | 1 | 125 | 14.5 | [illegible] |
| 1060943855 | EDICTED: LOW QUALITY PROTEIN: neuroblast differentiation-associated protein AHNAK-like [Callithrix jacc | 0 | 162.272 | 29 | 38 | 193 | 38 | 1 | 3118 | 331.5 | [illegible] |
| 1060936750 | PREDICTED: breakpoint cluster region protein-like [Callithrix jacchus] | 0 | 62.68 | 27 | 9 | 53 | 6 | 1 | 504 | 54.8 | [illegible] |
| 532444627 | beta-adrenergic receptor kinase 1 [Callithrix jacchus] | 0 | 99.749 | 31 | 16 | 53 | 9 | 1 | 689 | 79.5 | [illegible] |
| 532513956 | FAD-linked sulfhydryl oxidase ALR [Callithrix jacchus] | 0 | 14.919 | 19 | 3 | 9 | 3 | 1 | 206 | 23.8 | [illegible] |
| 532514088 | inositol 1,4,5-trisphosphate receptor type 2 [Callithrix jacchus] | 0 | 72.768 | 7 | 17 | 59 | 9 | 1 | 2704 | 308.2 | [illegible] |
| 532276140 | brefeldin A-inhibited guanine nucleotide-exchange protein 3 [Callithrix jacchus] | 0 | 134.826 | 15 | 24 | 107 | 24 | 1 | 2177 | 240.6 | [illegible] |
| 296232919 | PREDICTED: 60S acidic ribosomal protein P1 [Callithrix jacchus] | 0 | 63.842 | 80 | 3 | 21 | 3 | 1 | 114 | 11.5 | [illegible] |
| 296237736 | PREDICTED: factor VIII intron 22 protein [Callithrix jacchus] | 0 | 16.606 | 14 | 3 | 3 | 3 | 1 | 377 | 39.6 | [illegible] |
| 675768684 | PREDICTED: G-protein coupled receptor-associated sorting protein 2 [Callithrix jacchus] | 0 | 32.695 | 10 | 8 | 33 | 3 | 1 | 874 | 97.4 | [illegible] |
| 675776241 | PREDICTED: LOW QUALITY PROTEIN: cat eye syndrome critical region protein 6-like [Callithrix jacchus] | 0 | 35.948 | 12 | 5 | 27 | 5 | 1 | 570 | 58.3 | [illegible] |
| 390461946 | PREDICTED: 60S ribosomal protein L7 [Callithrix jacchus] | 0 | 102.879 | 51 | 21 | 237 | 21 | 1 | 248 | 29.2 | [illegible] |
| 1061016309 | PREDICTED: solute carrier family 2, facilitated glucose transporter member 3 [Callithrix jacchus] | 0 | 45.536 | 11 | 5 | 41 | 5 | 1 | 496 | 54 | [illegible] |
| 532219359 | septin-4 isoform 1 [Callithrix jacchus] | 0 | 176.749 | 45 | 16 | 284 | 2 | 1 | 470 | 53.9 | [illegible] |
| 675652084 | PREDICTED: LOW QUALITY PROTEIN: breakpoint cluster region protein-like [Callithrix jacchus] | 0 | 15.736 | 12 | 4 | 31 | 2 | 1 | 383 | 42.2 | [illegible] |
| 296208761 | PREDICTED: glutathione S-transferase Mu 5 [Callithrix jacchus] | 0 | 8.064 | 14 | 3 | 8 | 3 | 1 | 218 | 25.6 | [illegible] |
| 532522768 | neuroligin-4, X-linked [Callithrix jacchus] | 0 | 86.135 | 21 | 13 | 94 | 8 | 1 | 816 | 91.8 | [illegible] |
| 1060956953 | PREDICTED: uncharacterized protein LOC100414591 [Callithrix jacchus] | 0 | 20.177 | 14 | 2 | 65 | 2 | 1 | 149 | 15.6 | [illegible] |
| 1060946688 | PREDICTED: growth hormone-inducible transmembrane protein [Callithrix jacchus] | 0 | 15.871 | 9 | 4 | 28 | 4 | 1 | 350 | 37.7 | [illegible] |
| 1060983956 | PREDICTED: histone H2B type 1-M [Callithrix jacchus] | 0 | 39.395 | 32 | 5 | 79 | 2 | 1 | 142 | 15.5 | [illegible] |
| 296207834 | PREDICTED: guanine nucleotide-binding protein G(I)/G(S)/G(O) subunit gamma-5 [Callithrix jacchus] | 0 | 17.519 | 49 | 5 | 33 | 5 | 1 | 68 | 7.3 | [illegible] |
| 532524190 | piggyBac transposable element-derived protein 5 [Callithrix jacchus] | 0 | 30.625 | 13 | 6 | 35 | 6 | 1 | 524 | 58.2 | [illegible] |
| 1060931111 | PREDICTED: janus kinase and microtubule-interacting protein 3 [Callithrix jacchus] | 0 | 39.754 | 12 | 11 | 39 | 10 | 1 | 935 | 107.5 | [illegible] |
| 1060934128 | PREDICTED: cytochrome c oxidase subunit 7C, mitochondrial-like [Callithrix jacchus] | 0 | 13.537 | 62 | 3 | 56 | 3 | 1 | 63 | 7.3 | [illegible] |

**Supplementary Table 6 LC-MS/MS quantitation and comparison of 1,740 PSD proteins in marmoset cerebellum**

| Description of protein | | Detection by LC-MS/MS | | | | | | | | | | | | LFQ intensity normalized with total peptide abundance | | | | | | | | | | | | Coefficient of Variation (CV) | | | | Marmoset Gene ID | | Mouse Gene ID | | Mean relative abundance | | | | | | | | |
|---|---|---|---|---|---|---|---|---|---|---|---|---|---|---|---|---|---|---|---|---|---|---|---|---|---|---|---|---|---|---|---|---|---|---|---|---|---|---|---|---|---|---|
| GI Number | Description | q-value: Combined | Sum PEP Score | Coverage [%] | # Peptides | # PSMs | # Unique Peptides | # Protein Groups | # AAs | MW [kDa] | calc. pI | Score Mascot: Mascot | # Peptides (by Search Engine) | (Normalized): F48: Sample | (Normalized): F49: Sample | (Normalized): F50: Sample | (Normalized): F51: Sample | (Normalized): F52: Sample | (Normalized): F53: Sample | (Normalized): F54: Sample | (Normalized): F55: Sample | (Normalized): F56: Sample | (Normalized): F57: Sample | (Normalized): F58: Sample | (Normalized): F59: Sample | CV 2M | CV 3M | CV 6M | CV 12M | Gene ID | Gene Symbol | Gene ID | Gene Symbol | 2M | 3M | 6M | 24M | P-value | q-value (BH) | Fold | Cluster | DE |
| 532218031 | ATP-binding cassette sub-family A member 2 isoform a [Callithrix jacchus] | 0 | 13.881 | 2 | 3 | 15 | 3 | 1 | 2434 | 270 | 6.51 | 66 | 3 | 2898137.097 | 7407066.217 | 8020218.839 | 4391454 | 6270521.836 | 3941650.116 | 4192277.057 | 2326833 | 1937772.059 | 1217641.126 | 2443506.455 | 232117.4104 | 45.790247 | 25.37805 | 42.7509 | 85.36813 | 100411477 | ABCA2 | 11305 | Abca2 | 1.62 | 1.29 | 0.75 | 0.34 | 0.0816 | 0.18126 | 4.70695 | 2 | No |
| 296206333 | PREDICTED: abl interactor 1 isoform X1 [Callithrix jacchus] | 0 | 51.327 | 17 | 7 | 37 | 6 | 1 | 508 | 55.1 | 7.06 | 483 | 7 | 15962779.33 | 27708081.21 | 22063995.13 | 18694641.13 | 34433212.27 | 32211695.82 | 44389530 | 29167150.98 | 20158640.87 | 16405880.18 | 9896060.72 | 7292575.537 | 26.808304 | 29.94428 | 39.2066 | 41.91857 | 100408906 | ABI1 | 11308 | Abi1 | 0.94 | 1.23 | 1.35 | 0.48 | 0.1018 | 0.20815 | 2.7896 | 3 | No |
| 532503413 | very long-chain specific acyl-CoA dehydrogenase, mitochondrial isoform 1 precursor [Callithrix jacchus] | 0 | 35.402 | 13 | 7 | 39 | 7 | 1 | 655 | 70.2 | 8.76 | 475 | 7 | 10085329.41 | 32264277.56 | 23413907.46 | 34136152.58 | 57786124.9 | 53982795.68 | 34360557.38 | 45181717.34 | 58201270.38 | 21739408.3 | 22823217.71 | 25610707.75 | 50.930537 | 26.112 | 25.9988 | 8.538082 | 100399197 | ACADVL | 11370 | Acadvl | 0.63 | 1.39 | 1.31 | 0.67 | 0.08581 | 0.18781 | 2.21863 | 3 | No |
| 532349499 | microtubule-actin cross-linking factor 1 isoform a [Callithrix jacchus] | 0 | 462.139 | 19 | 86 | 478 | 84 | 1 | 5436 | 620.8 | 5.39 | 6286 | 86 | 336951734.7 | 346408973 | 328542924.8 | 362549702.6 | 327055233 | 361656948.7 | 233934289.3 | 260756967.8 | 251053612.5 | 222361664 | 255088484.5 | 231587124.3 | 2.6499014 | 5.775901 | 5.46345 | 7.13925 | 100398371 | MACF1 | 11426 | Macf1 | 1.15 | 1.20 | 0.85 | 0.81 | 0.00151 | 0.02479 | 1.48267 | 2 | No |
| 1060937427 | PREDICTED: aconitate hydratase, mitochondrial [Callithrix jacchus] | 0 | 92.747 | 19 | 15 | 107 | 15 | 1 | 857 | 94.2 | 7.91 | 1142 | 15 | 108593042.9 | 130981234.3 | 117623834.9 | 164587680.8 | 173833767.1 | 182451241.8 | 177728459.6 | 197577612.2 | 181543681.3 | 98464187.25 | 125783727 | 125326146.1 | 9.4599252 | 5.145378 | 5.67446 | 14.49801 | 100389367 | ACO2 | 11429 | Aco2 | 0.80 | 1.17 | 1.25 | 0.78 | 0.004538 | 0.03673 | 1.6021 | 3 | Yes |
| 675658800 | PREDICTED: disintegrin and metalloproteinase domain-containing protein 11 [Callithrix jacchus] | 0 | 8.74 | 3 | 2 | 3 | 2 | 1 | 777 | 84.2 | 7.08 | 22 | 2 | 4142559.722 | 2863457.764 | 2601590.511 | 2792145.406 | 2830946.646 | 2503588.162 | 2702430.672 | 1173910.733 | 1812197.648 | 5237188.389 | 5541127.019 | 4749963.917 | 25.746676 | 6.602507 | 40.4873 | 7.710532 | 100398230 | ADAM11 | 11488 | Adam11 | 0.99 | 0.83 | 0.58 | 1.59 | 0.005423 | 0.04085 | 2.72975 | 1 | Yes |
| 675673469 | PREDICTED: disintegrin and metalloproteinase domain-containing protein 22 isoform X1 [Callithrix jacchus] | 0 | 46.706 | 9 | 9 | 30 | 9 | 1 | 1024 | 113.5 | 7.14 | 223 | 9 | 7733832.551 | 2054465.031 | 3122314.799 | 23572191.97 | 6860086.853 | 5296811.507 | 5524703.353 | 2355623.899 | 3473978.327 | 13372065 | 12153061.48 | 921377.0758 | 70.135842 | 85.05857 | 42.4659 | 77.85862 | 100395237 | ADAM22 | 11496 | Adam22 | 0.60 | 1.65 | 0.53 | 1.22 | 0.5497 | 0.65776 | 3.14674 | 3 | No |
| 675650424 | PREDICTED: alpha-adducin isoform X5 [Callithrix jacchus] | 0 | 112.037 | 30 | 16 | 364 | 15 | 1 | 693 | 76.7 | 6.11 | 6415 | 16 | 727347535.3 | 837488038.2 | 967480363.6 | 846547059.1 | 854461665.1 | 900423955.7 | 608732121.7 | 690107990.5 | 756925654.2 | 822824017.4 | 744862042.4 | 756322452.6 | 14.240296 | 3.354863 | 10.8304 | 5.433913 | 100401343 | ADD1 | 11518 | Add1 | 1.06 | 1.09 | 0.86 | 0.98 | 0.06712 | 0.1589 | 1.26543 | 3 | No |
| 296223606 | PREDICTED: beta-adducin [Callithrix jacchus] | 0 | 96.214 | 25 | 15 | 171 | 14 | 1 | 726 | 80.8 | 5.92 | 2663 | 15 | 178413685.5 | 194312934.6 | 169943653.4 | 224821444.8 | 173234805.3 | 160702510.1 | 170390498.9 | 107130530.7 | 55505567.74 | 266177351.9 | 144533815.8 | 212605774.2 | 6.8394792 | 18.24616 | 51.8342 | 29.34252 | 100402289 | ADD2 | 11519 | Add2 | 1.05 | 1.09 | 0.65 | 1.21 | 0.4119 | 0.53766 | 1.87167 | 1 | No |
| 296230268 | PREDICTED: poly [ADP-ribose] polymerase 1 [Callithrix jacchus] | 0 | 41.057 | 10 | 8 | 35 | 8 | 1 | 1013 | 113 | 8.9 | 267 | 8 | 45334431.02 | 43568909.4 | 44024759.14 | 35087270.7 | 40925672.02 | 46364869.19 | 28310203.08 | 28781693.55 | 29152719.76 | 28303871 | 36086354.73 | 25401037.82 | 2.0684673 | 13.82598 | 1.46881 | 18.46027 | 100403519 | PARP1 | 11545 | Parp1 | 1.23 | 1.13 | 0.80 | 0.83 | 0.0001815 | 0.01425 | 1.54129 | 2 | Yes |
| 1061002891 | PREDICTED: LOW QUALITY PROTEIN: aggrecan core protein [Callithrix jacchus] | 0 | 20.942 | 2 | 4 | 31 | 4 | 1 | 2264 | 234.8 | 4.31 | 239 | 4 | 101211274.7 | 15906938.95 | 68950394.56 | 53801000.75 | 43798257.37 | 16340786.71 | 27834736.39 | 14929392.57 | 15972346.13 | 38174445.66 | 20074304.54 | 22981119.21 | 69.445412 | 51.06903 | 36.6152 | 35.89912 | 100385452 | ACAN | 11595 | Acan | 1.69 | 1.04 | 0.53 | 0.74 | 0.3777 | 0.50476 | 3.16785 | 2 | No |
| 532524482 | adenylate kinase isoenzyme 1 [Callithrix jacchus] | 0 | 14.522 | 21 | 4 | 45 | 4 | 1 | 194 | 21.6 | 8.62 | 500 | 4 | 42554709.72 | 58009057.01 | 74538453.08 | 68080072.44 | 105492198.7 | 120584416.2 | 71679325.44 | 79769231.17 | 95629811.9 | 68038441.04 | 95795663.37 | 100895316.3 | 27.403792 | 27.56164 | 14.7931 | 20.03849 | 100400464 | AK1 | 11636 | Ak1 | 0.71 | 1.20 | 1.01 | 1.08 | 0.2598 | 0.38604 | 1.6798 | 3 | No |
| 1061008114 | PREDICTED: alkaline phosphatase, tissue-nonspecific isozyme isoform X1 [Callithrix jacchus] | 0 | 7.287 | 4 | 2 | 2 | 2 | 1 | 524 | 57.6 | 6.76 | 55 | 2 | 8831059.807 | 7074884.384 | 7390158.291 | 5082278 | 5032254.367 | 6789705.138 | 2891075.779 | 3024858.613 | 4252381.966 | 4156255.033 | 4484864.738 | 3808932.454 | 12.057153 | 17.75656 | 22.137 | 8.144768 | 100401643 | ALPL | 11647 | Alpl | 1.48 | 1.08 | 0.65 | 0.79 | 0.017 | 0.06876 | 2.29105 | 2 | No |
| 296198349 | PREDICTED: serum albumin [Callithrix jacchus] | 0 | 83.645 | 26 | 15 | 102 | 15 | 1 | 608 | 69.4 | 5.99 | 1085 | 15 | 52461644.9 | 38057694.15 | 30913242.95 | 79772536.61 | 66588291.57 | 50070750.33 | 107163241.3 | 80435315.83 | 59411730.98 | 88866095.31 | 53277523.8 | 23541513.92 | 27.116565 | 22.72858 | 29.0666 | 59.21948 | 100391910 | ALB | 11657 | Alb | 0.66 | 1.08 | 1.35 | 0.91 | 0.1785 | 0.29893 | 2.03414 | 3 | No |
| 296189731 | PREDICTED: retinal dehydrogenase 1 [Callithrix jacchus] | 0 | 16.34 | 7 | 3 | 11 | 2 | 1 | 501 | 54.9 | 6.55 | 106 | 3 | 9569588.245 | 10062747.27 | 11271611.93 | 14185118.56 | 15336190.19 | 14811544.85 | 1012798.169 | 879499.0163 | 1089938.563 | 651919.659 | 1089925.936 | 682243.1893 | 8.501171 | 3.899717 | 10.7096 | 30.27114 | 100407242 | ALDH1A1 | 11668 | Aldh1a1 | 1.53 | 2.20 | 0.15 | 0.12 | 2.60E-05 | 0.00566 | 18.2885 | 2 | Yes |
| 1061018166 | PREDICTED: aldehyde dehydrogenase, mitochondrial [Callithrix jacchus] | 0 | 38.521 | 20 | 8 | 38 | 7 | 1 | 517 | 56.2 | 6.65 | 634 | 8 | 11039152.29 | 10139233.22 | 10967273.7 | 18455195 | 17669640.01 | 20980996.79 | 24396972.01 | 18453461.15 | 13547030.05 | 16348240.4 | 16221676.53 | 6136756.996 | 4.6673049 | 9.089516 | 28.9014 | 45.41394 | 100388478 | ALDH2 | 11669 | Aldh2 | 0.70 | 1.24 | 1.22 | 0.84 | 0.01416 | 0.06441 | 1.77647 | 3 | No |
| 675658716 | PREDICTED: fatty aldehyde dehydrogenase isoform X1 [Callithrix jacchus] | 0 | 16.232 | 6 | 2 | 17 | 2 | 1 | 527 | 59.7 | 8.88 | 447 | 2 | 3069362.308 | 3541605.428 | 3665277.86 | 1806162.125 | 2694722.499 | 3714379.83 | 2699469.762 | 2267032.672 | 2625367.684 | 1615210.986 | 1567616.846 | 2020963.517 | 9.181077 | 34.86896 | 9.1386 | 14.36297 | 100411274 | ALDH3A2 | 11671 | Aldh3a2 | 1.31 | 1.05 | 0.97 | 0.67 | 0.01175 | 0.06027 | 1.97476 | 2 | No |
| 1060946290 | PREDICTED: fructose-bisphosphate aldolase A isoform X2 [Callithrix jacchus] | 0 | 43.665 | 24 | 9 | 124 | 8 | 1 | 364 | 39.4 | 8.24 | 2579 | 9 | 99399036.5 | 137054739 | 140551423 | 175167213.5 | 170471388.8 | 205161015.9 | 185349081.3 | 218143226 | 204569716.4 | 122270394.8 | 143281464.5 | 130618708.3 | 18.156527 | 10.2503 | 8.12972 | 8.011023 | 103796602 | ALDOA | 11674 | Aldoa | 0.78 | 1.14 | 1.26 | 0.82 | 0.01336 | 0.06386 | 1.61287 | 3 | No |
| 296202196 | PREDICTED: fructose-bisphosphate aldolase C [Callithrix jacchus] | 0 | 44.559 | 16 | 7 | 148 | 6 | 1 | 364 | 39.4 | 6.44 | 3532 | 7 | 185852430.6 | 172406321.1 | 136988316.6 | 263249471 | 170417326.9 | 169188634 | 233129116.3 | 177899381.3 | 166313724.4 | 170665719.4 | 145849928.6 | 213462303.2 | 15.290515 | 26.84965 | 18.5528 | 19.36083 | 100408520 | ALDOC | 11676 | Aldoc | 0.90 | 1.09 | 1.05 | 0.96 | 0.7218 | 0.79742 | 1.21728 | 3 | No |
| 1060950263 | PREDICTED: ankyrin-1 isoform X3 [Callithrix jacchus] | 0 | 171.394 | 22 | 35 | 283 | 33 | 1 | 1908 | 209.1 | 6.73 | 3374 | 35 | 345285392.9 | 279833802.7 | 274816211.3 | 356778151.8 | 249114466.6 | 270308492.1 | 249584167.4 | 221997428 | 247333887.3 | 403918223.2 | 384509811.2 | 355926905.9 | 13.106631 | 19.52791 | 6.39254 | 6.328816 | 100388861 | ANK1 | 11733 | Ank1 | 0.99 | 0.96 | 0.79 | 1.26 | 0.007577 | 0.04829 | 1.59178 | 1 | Yes |
| 532519378 | ankyrin-3 isoform 1 [Callithrix jacchus] | 0 | 182.556 | 11 | 39 | 251 | 24 | 1 | 4382 | 480.6 | 6.52 | 3157 | 39 | 86774119.24 | 89812699.93 | 79722527.49 | 100460721.6 | 103966906.4 | 98094332 | 59916970.57 | 69150775.89 | 53230359.78 | 97261360.7 | 94259699.35 | 115101317.4 | 6.0587497 | 2.930033 | 13.1555 | 11.02348 | 100408683 | ANK3 | 11735 | Ank3 | 0.98 | 1.15 | 0.70 | 1.17 | 0.007357 | 0.04759 | 1.68198 | 3 | Yes |
| 675657852 | PREDICTED: rabankyrin-5 [Callithrix jacchus] | 0 | 7.336 | 2 | 3 | 8 | 3 | 1 | 1169 | 128.7 | 6.06 | 47 | 3 | 3074531.162 | 5892594.263 | 5764234.413 | 3958321.375 | 4238461.389 | 8145828.162 | 6590778.883 | 1981764.291 | 1852464.436 | 2221740.893 | 963249.1824 | 3162546.658 | 32.405359 | 42.97324 | 77.6723 | 52.15249 | 100415707 | ANKFY1 | 11736 | Ankfy1 | 1.23 | 1.37 | 0.87 | 0.53 | 0.214 | 0.33606 | 2.57464 | 2 | No |
| 296213563 | PREDICTED: acidic leucine-rich nuclear phosphoprotein 32 family member A [Callithrix jacchus] | 0 | 7.461 | 7 | 3 | 31 | 3 | 1 | 249 | 28.6 | 4.09 | 303 | 3 | 156379616.5 | 156794654 | 163168523.8 | 97541960.63 | 90309429.7 | 92466248.71 | 110284403.4 | 111452805.3 | 130081923.8 | 94330004.68 | 130518960.4 | 125615535.9 | 2.3966536 | 3.973851 | 9.47207 | 16.80504 | 100398495 | ANP32A | 11737 | Anp32a | 1.31 | 0.77 | 0.96 | 0.96 | 0.000208 | 0.01455 | 1.6993 | 2 | Yes |
| 296236267 | PREDICTED: ADP/ATP translocase 2 [Callithrix jacchus] | 0 | 51.163 | 33 | 11 | 166 | 4 | 1 | 298 | 32.9 | 9.74 | 1865 | 11 | 25064173.64 | 36204120.77 | 33387104.83 | 68972415.32 | 55990559.78 | 37971808.86 | 11658751.35 | 34027465.21 | 31814189.48 | 8563394.641 | 4960724.713 | 13153043.55 | 18.358099 | 28.66488 | 47.7112 | 46.17496 | 100387000 | SLC25A5 | 11740 | Slc25a5 | 1.05 | 1.80 | 0.86 | 0.29 | 0.01666 | 0.06821 | 6.10765 | 2 | No |
| 532503130 | annexin A6 isoform 1 [Callithrix jacchus] | 0 | 48.608 | 17 | 10 | 33 | 10 | 1 | 673 | 75.8 | 5.55 | 226 | 10 | 27879866.61 | 34050763.53 | 56347480.59 | 18228840.78 | 27033944.28 | 26788888.41 | 25581585.59 | 35006031.2 | 53640267.44 | 26599404.29 | 23078093.1 | 37093569.22 | 37.984276 | 20.87828 | 37.5014 | 25.20805 | 100411129 | ANXA6 | 11749 | Anxa6 | 1.21 | 0.74 | 1.17 | 0.89 | 0.3908 | 0.51724 | 1.64157 | 3 | No |
| [illegible] | [illegible] | [illegible] | [illegible] | [illegible] | [illegible] | [illegible] | [illegible] | [illegible] | [illegible] | [illegible] | [illegible] | [illegible] | [illegible] | [illegible] | [illegible] | [illegible] | [illegible] | [illegible] | [illegible] | [illegible] | [illegible] | [illegible] | [illegible] | [illegible] | [illegible] | [illegible] | [illegible] | [illegible] | [illegible] | [illegible] | [illegible] | [illegible] | [illegible] | [illegible] | [illegible] | [illegible] | [illegible] | [illegible] | [illegible] | [illegible] | [illegible] | [illegible] |

| | | | | | | | | |
|---|---|---|---|---|---|---|---|---|
| 296217670 | LOW QUALITY PROTEIN: phosphatidylinositol 4-phosphate 3-kinase C2 domain-containing subunit alpha [Call | 0 | 14.118 | 2 | 4 | 17 | 4 | 1 |
| 675758843 | PREDICTED: phosphatidylinositol 4-phosphate 5-kinase type-1 gamma [Callithrix jacchus] | 0 | 16.567 | 7 | 5 | 22 | 5 | 1 |
| 296206295 | PREDICTED: phosphatidylinositol 5-phosphate 4-kinase type-2 alpha isoform X1 [Callithrix jacchus] | 0 | 39.754 | 20 | 7 | 62 | 4 | 1 |
| 675703149 | PREDICTED: membrane-associated phosphatidylinositol transfer protein 1 isoform X1 [Callithrix jacchus] | 0 | 10.754 | 3 | 4 | 10 | 4 | 1 |
| 1060938161 | PREDICTED: pyruvate kinase PKM isoform X1 [Callithrix jacchus] | 0 | 91.049 | 31 | 15 | 172 | 15 | 1 |
| 296208352 | PREDICTED: cAMP-dependent protein kinase catalytic subunit beta isoform X1 [Callithrix jacchus] | 0 | 24.878 | 11 | 3 | 16 | 2 | 1 |
| 532344533 | protein kinase C beta type isoform 2 [Callithrix jacchus] | 0 | 40.821 | 14 | 7 | 32 | 3 | 1 |
| 532524322 | protein kinase C gamma type [Callithrix jacchus] | 0 | 155.258 | 34 | 18 | 470 | 16 | 1 |
| 532286372 | protein kinase C epsilon type [Callithrix jacchus] | 0 | 37.016 | 18 | 12 | 46 | 11 | 1 |
| 675737598 | PREDICTED: protein kinase C iota type [Callithrix jacchus] | 0 | 12.615 | 7 | 3 | 17 | 3 | 1 |
| 1060992206 | PREDICTED: 1-phosphatidylinositol 4,5-bisphosphate phosphodiesterase beta-1 [Callithrix jacchus] | 0 | 93.452 | 18 | 20 | 153 | 19 | 1 |
| 1060943976 | PREDICTED: 1-phosphatidylinositol 4,5-bisphosphate phosphodiesterase beta-3 [Callithrix jacchus] | 0 | 53.974 | 11 | 14 | 81 | 14 | 1 |
| 532276962 | 1-phosphatidylinositol 4,5-bisphosphate phosphodiesterase beta-4 isoform a [Callithrix jacchus] | 0 | 216.284 | 33 | 37 | 327 | 37 | 1 |
| 1060955933 | PREDICTED: LOW QUALITY PROTEIN: plectin [Callithrix jacchus] | 0 | 395.585 | 21 | 91 | 689 | 91 | 1 |
| 532513324 | proliferation-associated protein 2G4 [Callithrix jacchus] | 0 | 19.295 | 13 | 4 | 32 | 4 | 1 |
| 296236083 | PREDICTED: myelin proteolipid protein isoform X1 [Callithrix jacchus] | 0 | 50.009 | 33 | 10 | 440 | 10 | 1 |
| 1060936574 | PREDICTED: septin-5 [Callithrix jacchus] | 0 | 41.748 | 23 | 8 | 88 | 2 | 1 |
| 532449211 | serine/threonine-protein phosphatase PP1-alpha catalytic subunit isoform 1 [Callithrix jacchus] | 0 | 29.659 | 22 | 7 | 33 | 3 | 1 |
| 675723479 | PREDICTED: serine/threonine-protein phosphatase PP1-beta catalytic subunit isoform X1 [Callithrix jacchus] | 0 | 40.63 | 24 | 7 | 34 | 3 | 1 |
| 390468177 | PREDICTED: serine/threonine-protein phosphatase PP1-gamma catalytic subunit [Callithrix jacchus] | 0 | 24.397 | 15 | 5 | 46 | 2 | 1 |
| 296195900 | PREDICTED: serine/threonine-protein phosphatase 2B catalytic subunit alpha isoform [Callithrix jacchus] | 0 | 49.485 | 26 | 11 | 87 | 8 | 1 |
| 675708690 | REDICTED: serine/threonine-protein phosphatase 2B catalytic subunit beta isoform isoform X1 [Callithrix jacchu | 0 | 33.287 | 23 | 9 | 47 | 6 | 1 |
| 296234171 | PREDICTED: serine/threonine-protein phosphatase 5 isoform X1 [Callithrix jacchus] | 0 | 14.597 | 7 | 3 | 11 | 3 | 1 |
| 296211554 | PREDICTED: 5'-AMP-activated protein kinase subunit gamma-1 [Callithrix jacchus] | 0 | 14.531 | 13 | 4 | 13 | 4 | 1 |
| 1060998742 | PREDICTED: cAMP-dependent protein kinase type I-alpha regulatory subunit [Callithrix jacchus] | 0 | 64.537 | 27 | 11 | 81 | 9 | 1 |
| 1060963768 | ICTED: LOW QUALITY PROTEIN: cAMP-dependent protein kinase type I-beta regulatory subunit [Callithrix jac | 0 | 38.638 | 23 | 8 | 72 | 6 | 1 |
| 390475024 | PREDICTED: cAMP-dependent protein kinase type II-alpha regulatory subunit isoform X1 [Callithrix jacchus] | 0 | 16.277 | 12 | 5 | 38 | 5 | 1 |
| 474367 | prion protein [Callithrix jacchus] | 0 | 7.899 | 8 | 2 | 43 | 2 | 1 |
| 1061020560 | PREDICTED: ribose-phosphate pyrophosphokinase 1 [Callithrix jacchus] | 0 | 19.321 | 15 | 4 | 33 | 4 | 1 |
| 296233836 | PREDICTED: periaxin isoform X1 [Callithrix jacchus] | 0 | 204.898 | 29 | 28 | 154 | 28 | 1 |
| 532519674 | puromycin-sensitive aminopeptidase [Callithrix jacchus] | 0 | 20.036 | 5 | 4 | 19 | 4 | 1 |
| 1060999653 | PREDICTED: cytohesin-1 isoform X1 [Callithrix jacchus] | 0 | 10.679 | 8 | 3 | 17 | 3 | 1 |
| 1060962843 | PREDICTED: cytohesin-3 isoform X1 [Callithrix jacchus] | 0 | 20.368 | 11 | 3 | 16 | 3 | 1 |
| 296215138 | PREDICTED: proteasome subunit alpha type-3 isoform X1 [Callithrix jacchus] | 0 | 8.289 | 9 | 2 | 12 | 2 | 1 |
| 296202288 | PREDICTED: proteasome subunit beta type-6 isoform X1 [Callithrix jacchus] | 0 | 11.905 | 14 | 3 | 11 | 3 | 1 |
| 296215716 | PREDICTED: 26S protease regulatory subunit 4 [Callithrix jacchus] | 0 | 18.809 | 13 | 5 | 24 | 4 | 1 |
| 390466836 | PREDICTED: 26S protease regulatory subunit 7 [Callithrix jacchus] | 0 | 63.92 | 32 | 12 | 58 | 12 | 1 |
| 1060996156 | PREDICTED: LOW QUALITY PROTEIN: 26S protease regulatory subunit 8 [Callithrix jacchus] | 0 | 25.123 | 18 | 6 | 29 | 5 | 1 |
| 532447313 | twinfilin-1 isoform 2 [Callithrix jacchus] | 0 | 26.985 | 15 | 5 | 48 | 4 | 1 |
| 675683661 | PREDICTED: tyrosine-protein phosphatase non-receptor type 11 isoform X1 [Callithrix jacchus] | 0 | 10.788 | 5 | 3 | 10 | 3 | 1 |
| 532205061 | tyrosine-protein phosphatase non-receptor type 13 isoform 4 [Callithrix jacchus] | 0 | 5.193 | 1 | 2 | 4 | 2 | 1 |
| 1060932637 | PREDICTED: receptor-type tyrosine-protein phosphatase delta [Callithrix jacchus] | 0 | 75.333 | 9 | 14 | 62 | 7 | 1 |

[illegible]

| Protein | | | | | | | | | | | | | | | | | | | |
|---|---|---|---|---|---|---|---|---|---|---|---|---|---|---|---|---|---|---|---|
| [illegible]x jacchus] | 0 | 50.241 | 9 | 12 | 55 | 11 | 1 | 1731 | 184.8 | 8.88 | 368 | 12 | 140848875.2 | 35567153.89 | 29076704.18 | 18756290.5 | 21130773.65 | 23198525.42 | 45023173.97 |
| [illegible]form X1 [Callithrix jacchus] | 0 | 13.429 | 8 | 3 | 11 | 3 | 1 | 364 | 41.9 | 6.61 | 95 | 3 | 546920.8624 | 5335554.198 | 2980743.337 | 6730950.688 | 4568481.58 | 10850351.02 | 2628592.06 |
| [illegible]Callithrix jacchus] | 0 | 12.008 | 12 | 3 | 5 | 3 | 1 | 365 | 41.3 | 10.7 | 27 | 3 | 62678868.77 | 50298257.13 | 56194924.13 | 38098439.84 | 39508686.39 | 44182431.42 | 49495605.08 |
| [illegible]rix jacchus] | 0 | 14.409 | 18 | 3 | 16 | 3 | 1 | 212 | 23.5 | 6.54 | 80 | 3 | 14368083.51 | 15316822.54 | 14484026.19 | 15252211 | 19027996.81 | 19774430.64 | 9918651.952 |
| [illegible][Callithrix jacchus] | 0 | 20.414 | 13 | 6 | 26 | 6 | 1 | 492 | 55.7 | 5.3 | 178 | 6 | 15669533.61 | 18661272.54 | 15344455.58 | 14171106.59 | 10881798.33 | 16028576.83 | 11596113.93 |
| [illegible] jacchus] | 0 | 75.32 | 4 | 13 | 69 | 13 | 1 | 4374 | 481.5 | 5.21 | 1471 | 13 | 39633950.82 | 33926757.69 | 26363618.67 | 40375021.75 | 26771271.95 | 26708487.31 | 33756440.8 |
| [illegible]

| | | | | | | | | | | | | | | | | | | | | | | | | | | | | | | | | | | | | | | | | | | |
|---|---|---|---|---|---|---|---|---|---|---|---|---|---|---|---|---|---|---|---|---|---|---|---|---|---|---|---|---|---|---|---|---|---|---|---|---|---|---|---|---|---|---|
| 296215361 | PREDICTED: LOW QUALITY PROTEIN: exonuclease 3'-5' domain-containing protein 2 [Callithrix jacchus] | 0 | 16.282 | 8 | 4 | 20 | 4 | 1 | 621 | 70.2 | 8.09 | 454 | 4 | 8476179.538 | 8176573.032 | 6250076.772 | 3465669.75 | 6346676.742 | 10535292.5 | 4476323.643 | 859900.0493 | 6005282.373 | 8632364.272 | 6373215.145 | 5506526.302 | 15.824368 | 52.4126 | 69.8934 | 23.60239 | 100408563 | EXD2 | 97827 | Exd2 | 1.22 | 1.08 | 0.60 | 1.09 | 0.3756 | 0.5035 | 2.01938 | 2 | No |
| 296202480 | PREDICTED: leucine-rich repeat-containing protein 59 [Callithrix jacchus] | 0 | 31.165 | 25 | 6 | 60 | 6 | 1 | 307 | 35 | 9.58 | 640 | 6 | 91407976.14 | 119268987 | 85684774.69 | 130508950.7 | 123186828.4 | 86509851.65 | 119559810.8 | 115095062.2 | 64866898.34 | 80636483.75 | 106135642.6 | 67272103.03 | 18.187626 | 20.73831 | 30.4188 | 23.3168 | 100411408 | LRRC59 | 98238 | Lrrc59 | 1.00 | 1.14 | 1.01 | 0.85 | 0.6041 | 0.7031 | 1.33951 | 3 | No |
| 675666288 | PREDICTED: EF-hand domain-containing protein D1 [Callithrix jacchus] | 0 | 23.792 | 19 | 4 | 15 | 3 | 1 | 233 | 26.2 | 5.29 | 197 | 4 | 25536540.7 | 7215913.522 | 6148377.848 | 8957978.25 | 8592291.912 | 10776028.82 | 26764240.67 | 11208342.94 | 12766820.62 | 10976308.19 | 8368469.687 | 11070183.66 | 84.049631 | 12.38704 | 50.6519 | 15.12533 | 100398958 | EFHD1 | 98363 | Efhd1 | 1.05 | 0.76 | 1.37 | 0.82 | 0.6061 | 0.70496 | 1.79125 | 3 | No |
| 1060949182 | CTED: LOW QUALITY PROTEIN: PH domain leucine-rich repeat-containing protein phosphatase 1 [Callithrix ja | 0 | 5.211 | 2 | 2 | 3 | 2 | 1 | 1714 | 184.4 | 6.46 | 14 | 2 | 13277531.23 | 15383305.73 | 11830642.62 | 14465870 | 9323687.746 | 10894601.93 | 12036317.9 | 10255702.06 | 10221773.87 | 14743311.31 | 14717749.6 | 18795136.72 | 13.236012 | 22.79259 | 9.57721 | 14.58923 | 100395153 | PHLPP1 | 98432 | Phlpp1 | 1.04 | 0.89 | 0.83 | 1.24 | 0.1114 | 0.21927 | 1.48418 | 1 | No |
| 1060960932 | PREDICTED: rab3 GTPase-activating protein non-catalytic subunit isoform X2 [Callithrix jacchus] | 0 | 11.642 | 2 | 3 | 4 | 3 | 1 | 1392 | 155.8 | 5.64 | 18 | 3 | 4671332.215 | 4537310.661 | 4903036.845 | 9868888.125 | 3987988.551 | 7915263.875 | 8774454.755 | 8492354.834 | 9050162.778 | 1545336.488 | 3208489.812 | 3139720.264 | 3.9334338 | 41.27024 | 3.17943 | 35.76333 | 100386852 | RAB3GAP2 | 98732 | Rab3gap2 | 0.81 | 1.24 | 1.50 | 0.45 | 0.0002551 | 0.01455 | 3.33399 | 3 | Yes |
| 675687504 | PREDICTED: EH domain-containing protein 4 [Callithrix jacchus] | 0 | 47.605 | 24 | 11 | 45 | 10 | 1 | 541 | 61.2 | 6.76 | 457 | 11 | 17513988.58 | 24685071.69 | 15378224.07 | 15514691.06 | 21953552.67 | 20550400.42 | 26308876.94 | 30333094.07 | 30796662.25 | 11726172.61 | 18310130.32 | 22300899.99 | 25.401496 | 17.50768 | 8.46803 | 30.60974 | 100408190 | EHD4 | 98878 | Ehd4 | 0.90 | 0.91 | 1.37 | 0.82 | 0.04456 | 0.12115 | 1.67068 | 3 | No |
| 296206142 | PREDICTED: LOW QUALITY PROTEIN: USP6 N-terminal-like protein [Callithrix jacchus] | 0 | 23.626 | 8 | 5 | 19 | 5 | 1 | 845 | 95.8 | 9.04 | 77 | 5 | 18543030.07 | 16440240.5 | 20428111.73 | 7292590.375 | 15391829.75 | 14719218.96 | 8820225.284 | 23099736.54 | 26460678.75 | 10089214.99 | 6930954.052 | 14720302.18 | 10.800628 | 36.04889 | 48.1317 | 37.02981 | 100392325 | USP6NL | 98910 | Usp6nl | 1.21 | 0.82 | 1.28 | 0.69 | 0.1499 | 0.26588 | 1.83932 | 3 | No |
| 1060990612 | :DICTED: LOW QUALITY PROTEIN: brefeldin A-inhibited guanine nucleotide-exchange protein 2 [Callithrix jacc | 0 | 20.144 | 3 | 4 | 15 | 3 | 1 | 1814 | 204.7 | 6.43 | 310 | 4 | 3467093.346 | 2381865.262 | 2442689.183 | 4349543.5 | 2027077.205 | 3801471.754 | 2067190.28 | 3509532.592 | 3654783.784 | 2999744.705 | 3316307.816 | 2044832.934 | 22.059181 | 35.7826 | 28.5222 | 23.75009 | 100410913 | ARFGEF2 | 99371 | Arfgef2 | 0.92 | 1.13 | 1.02 | 0.93 | 0.8802 | 0.9149 | 1.22748 | 3 | No |
| 675639549 | PREDICTED: LOW QUALITY PROTEIN: golgin subfamily A member 2 [Callithrix jacchus] | 0 | 26.113 | 6 | 6 | 15 | 6 | 1 | 1001 | 113 | 5.05 | 267 | 6 | 8754801.664 | 10833844.65 | 7367143.845 | 5460108.188 | 4273937.142 | 3579237.262 | 4973892.56 | 5283392.974 | 2961831.183 | 2826795.613 | 2964250.272 | 1446233.387 | 19.418498 | 21.43146 | 28.6073 | 34.8017 | 100404592 | GOLGA2 | 99412 | Golga2 | 1.78 | 0.88 | 0.87 | 0.48 | 0.02421 | 0.08362 | 3.72458 | 2 | No |
| 675671569 | PREDICTED: WD repeat-containing protein 47 isoform X1 [Callithrix jacchus] | 0 | 13.677 | 4 | 3 | 12 | 3 | 1 | 929 | 103.1 | 6.11 | 64 | 3 | 4113579.598 | 3865923.749 | 4001835.185 | 5222580 | 4800101.551 | 5805012.427 | 7621088.339 | 4019382.806 | 3669282.071 | 2410548.462 | 2908272.028 | 2210755.72 | 3.1054367 | 9.56373 | 42.8653 | 14.31181 | 100399455 | WDR47 | 99512 | Wdr47 | 0.95 | 1.25 | 1.21 | 0.59 | 0.01247 | 0.06164 | 2.10207 | 3 | No |
| 1060979916 | PREDICTED: protein transport protein Sec24B isoform X1 [Callithrix jacchus] | 0 | 12.968 | 4 | 4 | 11 | 4 | 1 | 1282 | 140.1 | 6.8 | 20 | 4 | 6523481.639 | 8106210.385 | 7787571.815 | 3867713.125 | 5769991.152 | 6860696.556 | 5881045.064 | 7677310.397 | 1081858.338 | 11483189.93 | 9255245.905 | 5625824.693 | 11.202615 | 27.54302 | 69.8711 | 33.64207 | 100392878 | SEC24B | 99683 | Sec24b | 1.12 | 0.83 | 0.73 | 1.32 | 0.3368 | 0.46621 | 1.80081 | 1 | No |
| 1060952789 | PREDICTED: microtubule-associated protein RP/EB family member 3 isoform X1 [Callithrix jacchus] | 0 | 35.114 | 26 | 8 | 54 | 2 | 1 | 281 | 32 | 5.54 | 412 | 8 | 29637393.54 | 38924538.31 | 47523991 | 34599404 | 39734938.44 | 35734676.46 | 33416571.7 | 36972900.04 | 38867144.05 | 47903965.78 | 47069081.84 | 57562076.16 | 23.117795 | 7.352678 | 7.59826 | 11.47031 | 100386237 | MAPRE3 | 100732 | Mapre3 | 0.95 | 0.90 | 0.90 | 1.25 | 0.09779 | 0.20247 | 1.39612 | 1 | No |
| 675673405 | PREDICTED: A-kinase anchor protein 9 isoform X1 [Callithrix jacchus] | 0 | 32.094 | 3 | 10 | 21 | 10 | 1 | 3964 | 459.5 | 5.05 | 106 | 10 | 17205112.57 | 13164108.94 | 7868547.009 | 12394747 | 13787357.85 | 14091687.59 | 14389828.7 | 10391441.03 | 12059762.03 | 9179613.383 | 10689949.73 | 8931066.195 | 36.735745 | 6.7396 | 16.3538 | 11.03229 | 100389678 | AKAP9 | 100986 | Akap9 | 1.06 | 1.12 | 1.02 | 0.80 | 0.06273 | 0.15118 | 1.38872 | 3 | No |
| 532452821 | WD repeat-containing protein 91 [Callithrix jacchus] | 0 | 6.392 | 3 | 2 | 11 | 2 | 1 | 747 | 83.4 | 6.49 | 32 | 2 | 2767199.246 | 2397028.6 | 2091608.385 | 1924968.875 | 2156790.555 | 811213.9625 | 1870643.506 | 1656984.658 | 2809268.928 | 1908962.913 | 1095130.704 | 2704423.206 | 13.987865 | 44.10495 | 29.0193 | 42.28754 | 100415131 | WDR91 | 101240 | Wdr91 | 1.20 | 0.81 | 1.05 | 0.94 | 0.5098 | 0.61945 | 1.48291 | 3 | No |
| 532508674 | protein BRICK1 [Callithrix jacchus] | 0 | 9.564 | 39 | 3 | 14 | 3 | 1 | 75 | 8.7 | 5.45 | 89 | 3 | 28717210.75 | 38759522.32 | 42252107.56 | 28476900 | 34063132.94 | 33184570.17 | 36316450.02 | 41689317.45 | 43413012.88 | 26857205.65 | 10212858.22 | 36244690.23 | 19.210842 | 9.414154 | 9.14525 | 53.94584 | 100403256 | BRK1 | 101314 | Brk1 | 1.10 | 0.96 | 1.21 | 0.73 | 0.1554 | 0.27275 | 1.65613 | 3 | No |
| 296230968 | PREDICTED: exocyst complex component 8 [Callithrix jacchus] | 0 | 48.33 | 15 | 9 | 56 | 9 | 1 | 725 | 81.8 | 5.44 | 476 | 9 | 248550098.2 | 209174423 | 200669810.7 | 224642366.8 | 205633610 | 200946709.4 | 177794356.2 | 170182195.2 | 166725860.4 | 205333958.7 | 202777963.2 | 162251474.3 | 11.639699 | 5.963886 | 3.30062 | 12.71276 | 100413887 | EXOC8 | 102058 | Exoc8 | 1.11 | 1.06 | 0.87 | 0.96 | 0.04031 | 0.11545 | 1.27917 | 3 | No |
| 296233064 | PREDICTED: growth arrest and DNA damage-inducible proteins-interacting protein 1 [Callithrix jacchus] | 0 | 23.671 | 18 | 3 | 22 | 3 | 1 | 222 | 25.8 | 10.1 | 340 | 3 | 67378291.47 | 53468237.33 | 60155909.26 | 45269419.53 | 48702349.31 | 49314021.57 | 65074080.53 | 66333201.82 | 68469295.53 | 99134256.1 | 109184214.3 | 69653544.95 | 11.530353 | 4.564589 | 2.57616 | 22.03733 | 100402687 | GADD45GIP1 | 102060 | Gadd45gip1 | 0.90 | 0.71 | 1.00 | 1.39 | 0.002226 | 0.03038 | 1.94138 | 1 | Yes |
| 296231019 | PREDICTED: phosphorylase b kinase regulatory subunit beta isoform X2 [Callithrix jacchus] | 0 | 24.51 | 6 | 7 | 23 | 7 | 1 | 1093 | 124.9 | 6.99 | 145 | 7 | 19423414.46 | 31513445.48 | 29225497.94 | 22983439.84 | 33748796.15 | 62698900.4 | 26557342.95 | 44605569.42 | 19026273.02 | 64839323.16 | 26088554.45 | 12307820.92 | 24.035273 | 51.59428 | 43.725 | 79.14963 | 100395554 | PHKB | 102093 | Phkb | 0.82 | 1.22 | 0.92 | 1.05 | 0.806 | 0.86272 | 1.48987 | 3 | No |
| 1060941577 | PREDICTED: pleckstrin homology-like domain family B member 1 isoform X4 [Callithrix jacchus] | 0 | 45.045 | 8 | 11 | 57 | 11 | 1 | 1459 | 159.6 | 8.59 | 478 | 11 | 40644457.53 | 32747440.48 | 33863139.73 | 32032457.19 | 33221735.12 | 25848987.32 | 28429773.91 | 21411099.23 | 19931681.35 | 22821815.11 | 22981971.72 | 25219885.4 | 11.954218 | 13.03444 | 19.5204 | 5.662986 | 100407349 | PHLDB1 | 102693 | Phldb1 | 1.26 | 1.07 | 0.82 | 0.84 | 0.05007 | 0.1318 | 1.53721 | 2 | No |
| 1061019656 | PREDICTED: plastin-3 isoform X1 [Callithrix jacchus] | 0 | 18.985 | 8 | 4 | 16 | 4 | 1 | 639 | 71.8 | 5.62 | 143 | 4 | 6504774.728 | 5589306.659 | 5736141.003 | 6687648.531 | 7764346.607 | 8758938.83 | 9006096.781 | 6946624.782 | 7117412.366 | 5904972.85 | 4737216.871 | 4877968.79 | 8.2725374 | 13.38916 | 14.8625 | 12.32209 | 100403677 | PLS3 | 102866 | Pls3 | 0.90 | 1.17 | 1.16 | 0.78 | 0.07027 | 0.16435 | 1.49553 | 3 | No |
| 296223770 | PREDICTED: exportin-1 isoform X1 [Callithrix jacchus] | 0 | 16.061 | 3 | 3 | 14 | 3 | 1 | 1071 | 123.3 | 6.06 | 181 | 3 | 4234274.83 | 3668191.173 | 6957746.426 | 5711420.313 | 2254292.305 | 9458581.463 | 3034586.481 | 8449538.558 | 4818762.044 | 13110646.1 | 13040997.31 | 53655302.39 | 35.505611 | 62.0361 | 50.7785 | 88.06978 | 100410392 | XPO1 | 103573 | Xpo1 | 0.46 | 0.54 | 0.51 | 2.49 | 0.6133 | 0.70859 | 5.37051 | 1 | No |
| 532447989 | TBC1 domain family member 10A isoform 2 [Callithrix jacchus] | 0 | 35.271 | 15 | 5 | 22 | 5 | 1 | 509 | 57.3 | 7.88 | 214 | 5 | 7524537.045 | 7117904.001 | 7530555.702 | 10248052.25 | 8236302.677 | 6194367.967 | 7156028.451 | 6573043.25 | 5197032.035 | 10642342.93 | 6799075.7 | 5644233.364 | 3.2001937 | 24.63697 | 15.9446 | 34.00538 | 100388268 | TBC1D10A | 103724 | Tbc1d10a | 1.00 | 1.11 | 0.85 | 1.04 | 0.5073 | 0.61831 | 1.30395 | 3 | No |
| 296201464 | PREDICTED: tubulin gamma-1 chain [Callithrix jacchus] | 0 | 13.696 | 6 | 3 | 14 | 3 | 1 | 451 | 51.2 | 6.14 | 73 | 3 | 11812837.12 | 8333155.783 | 7369031.769 | 3164867.297 | 6527816.074 | 4915428.238 | 2775848.427 | 4618756.776 | 5435189.956 | 12480527.27 | 10984710.64 | 10170450.81 | 25.486731 | 34.54137 | 31.8545 | 10.45029 | 100410082 | TUBG1 | 103768 | Tubg2 | 1.24 | 0.66 | 0.58 | 1.52 | 0.01065 | 0.05825 | 2.62169 | 1 | No |
| 296225987 | PREDICTED: dolichyl-diphosphooligosaccharide--protein glycosyltransferase subunit 1 [Callithrix jacchus] | 0 | 24.345 | 12 | 7 | 32 | 7 | 1 | 607 | 68.7 | 6.33 | 275 | 7 | 26198921.45 | 21760137.8 | 21491797.07 | 19488188.19 | 25152716.78 | 31110034.22 | 26170667.16 | 20017347.96 | 21827740.58 | 15532394.42 | 19633263.26 | 21064087.42 | 11.419311 | 23.01571 | 13.9482 | 15.31878 | 100399378 | RPN1 | 103963 | Rpn1 | 1.03 | 1.12 | 1.01 | 0.83 | 0.3586 | 0.48595 | 1.34717 | 3 | No |
| 296229794 | PREDICTED: dynamin-3 isoform X3 [Callithrix jacchus] | 0 | 74.755 | 18 | 15 | 170 | 7 | 1 | 863 | 97.1 | 8.35 | 3275 | 15 | 27772067.54 | 28418848.05 | 37091117.01 | 39663907.42 | 27202656.39 | 39147001.43 | 29425686.54 | 22212388.51 | 22234118.34 | 22325081.55 | 64637129.44 | 30584341.34 | 16.735392 | 19.95036 | 16.8873 | 57.24055 | 100406310 | DNM3 | 103967 | Dnm3 | 0.95 | 1.09 | 0.76 | 1.20 | 0.2932 | 0.41851 | 1.59121 | 1 | No |
| 1060940928 | PREDICTED: LOW QUALITY PROTEIN: reticulon-1 [Callithrix jacchus] | 0 | 51.902 | 19 | 9 | 58 | 8 | 1 | 839 | 91 | 4.84 | 491 | 9 | 50160819.21 | 57470216.79 | 64896860.62 | 51751758.28 | 65160185.29 | 30580848.2 | 51566918.46 | 33563649.17 | 38664975.98 | 34226064.59 | 32086392.82 | 47400334.98 | 12.812012 | 35.46128 | 22.4863 | 21.87916 | 100390775 | RTN1 | 104001 | Rtn1 | 1.24 | 1.06 | 0.89 | 0.82 | 0.1648 | 0.28363 | 1.51723 | 2 | No |
| 675754752 | PREDICTED: synaptojanin-1 isoform X1 [Callithrix jacchus] | 0 | 75.969 | 11 | 14 | 81 | 14 | 1 | 1613 | 177.5 | 7.99 | 1131 | 14 | 44243307.1 | 40999891.39 | 36176812.13 | 59694783.88 | 49241392.14 | 52059924.3 | 70837340.81 | 73215410.23 | 57133842.26 | 51802110.85 | 34019700.69 | 50250423.53 | 10.028687 | 10.07812 | 12.9433 | 21.71497 | 100395303 | SYNJ1 | 104015 | Synj1 | 0.78 | 1.04 | 1.30 | 0.88 | 0.03971 | 0.11459 | 1.65695 | 3 | No |
| 532446003 | WD repeat-containing protein 7 isoform 1 [Callithrix jacchus] | 0 | 80.233 | 11 | 15 | 122 | 15 | 1 | 1490 | 163.7 | 6.84 | 1182 | 15 | 64025122.28 | 53266215.48 | 75497885.67 | 79833731.53 | 71840070.93 | 83993375.02 | 91717146.85 | 85676433.15 | 94098811.2 | 77158447.21 | 50489384.64 | 96434777.33 | 17.300363 | 7.862728 | 4.79864 | 30.88609 | 103787702 | WDR7 | 104082 | Wdr7 | 0.83 | 1.02 | 1.18 | 0.97 | 0.08066 | 0.1797 | 1.40822 | 3 | No |
| 532210711 | ATP-citrate synthase isoform 1 [Callithrix jacchus] | 0 | 57.573 | 12 | 12 | 72 | 12 | 1 | 1101 | 120.7 | 7.23 | 609 | 12 | 84789669 | 69525847.09 | 78120985.42 | 100638010.5 | 70008348.86 | 75580819.85 | 67143092.95 | 40340568.17 | 40093758.52 | 50418424.66 | 48220634.79 | 44901412.28 | 9.8764338 | 19.87608 | 31.6028 | 5.804852 | 100405725 | ACLY | 104112 | Acly | 1.21 | 1.28 | 0.77 | 0.75 | 0.02131 | 0.07757 | 1.71539 | 2 | No |
| 532452699 | diacylglycerol kinase zeta isoform 4 [Callithrix jacchus] | 0 | 73.167 | 14 | 10 | 61 | 8 | 1 | 1121 | 124.9 | 9.06 | 691 | 10 | 40313405.94 | 28418185.38 | 20724057.31 | 33298044 | 19901689.09 | 12663867.98 | 23075208.71 | 15526295.68 | 18061324.99 | 22925776.74 | 16248212.56 | 13426687.99 | 33.098419 | 47.68207 | 20.3397 | 27.82232 | 100415148 | DGKZ | 104418 | Dgkz | 1.35 | 1.00 | 0.86 | 0.80 | 0.4716 | 0.59077 | 1.70066 | 2 | No |
| 675641404 | PREDICTED: cdc42 effector protein 1 [Callithrix jacchus] | 0 | 31.221 | 22 | 6 | 52 | 6 | 1 | 368 | 38.3 | 8.46 | 442 | 6 | 21235472.25 | 25488323.5 | 19722711.69 | 14751898.5 | 11815110.37 | 6388643.068 | 12424539.21 | 5903348.173 | 8907081.326 | 9059811.574 | 6757860.553 | 11017642.18 | 13.49667 | 38.6241 | 35.9534 | 23.83657 | 100398181 | CDC42EP1 | 104445 | Cdc42ep1 | 1.73 | 0.86 | 0.71 | 0.70 | 0.01754 | 0.07016 | 2.47608 | 2 | No |
| 296192722 | PREDICTED: arginine--tRNA ligase, cytoplasmic [Callithrix jacchus] | 0 | 43.103 | 16 | 9 | 45 | 9 | 1 | 660 | 75.5 | 7.05 | 538 | 9 | 19346271.12 | 20855848.82 | 25664010.17 | 10707634.78 | 16630284.98 | 16287925.01 | 18000466.59 | 17559306.55 | 15258063.35 | 11200685.53 | 5311760.312 | 8438543.594 | 15.027159 | 22.86504 | 8.69334 | 35.42557 | 100413748 | RARS | 104458 | Rars | 1.42 | 0.94 | 1.10 | 0.54 | 0.03386 | 0.10373 | 2.63982 | 2 | No |
| 532513702 | tetratricopeptide repeat protein 7B [Callithrix jacchus] | 0 | 21.82 | 7 | 5 | 20 | 5 | 1 | 851 | 95 | 6.61 | 296 | 5 | 13025580.95 | 3492730.191 | 3812735.727 | 27277126.5 | 28381618.76 | 19524594.63 | 29821158.3 | 20395287.9 | 17471285.32 | 19736350.38 | 27906293.44 | 22294327.26 | 79.884444 | 19.25878 | 28.1895 | 17.92624 | 100390538 | TTC7B | 104718 | Ttc7b | 0.35 | 1.29 | 1.16 | 1.20 | 0.04998 | 0.13177 | 3.69796 | 3 | No |
| 296224457 | PREDICTED: ATP-dependent RNA helicase DDX1 [Callithrix jacchus] | 0 | 42.029 | 15 | 9 | 113 | 9 | 1 | 740 | 82.4 | 7.23 | 940 | 9 | 129883638.5 | 131435164.4 | 141429194.4 | 120686922.1 | 146801963.9 | 99769313.46 | 115204069.9 | 119692666.8 | 117359417 | 108968623.1 | 127983414.8 | 128847820.5 | 4.6675456 | 19.25035 | 1.91186 | 9.21491 | 100397694 | DDX1 | 104721 | Ddx1 | 1.08 | 0.99 | 0.95 | 0.98 | 0.09304 | 0.19854 | 1.14334 | 3 | No |
| [illegible] | [illegible] | [illegible] | [illegible] | [illegible] | [illegible] | [illegible] | [illegible] | [illegible] | [illegible] | [illegible] | [illegible] | [illegible] | [illegible] | [illegible] | [illegible] | [illegible] | [illegible] | [illegible] | [illegible] | [illegible] | [illegible] | [illegible] | [illegible] | [illegible] | [illegible] | [illegible] | [illegible] | [illegible] | [illegible] | [illegible] | [illegible] | [illegible] | [illegible] | [illegible] | [illegible] | [illegible] | [illegible] | [illegible] | [illegible] | [illegible] | [illegible] | [illegible] |

| | | | | | | | | | | | | | | | | | | | | | | | | | | | | | | | | | | | | | | | | | | |
|---|---|---|---|---|---|---|---|---|---|---|---|---|---|---|---|---|---|---|---|---|---|---|---|---|---|---|---|---|---|---|---|---|---|---|---|---|---|---|---|---|---|---|
| 532340489 | vacuolar protein sorting-associated protein 13C isoform 2A [Callithrix jacchus] | 0 | 36 | 2 | 6 | 23 | 6 | 1 | 3754 | 421.3 | 6.86 | 356 | 6 | 9801438.627 | 6882064.411 | 8511590.906 | 6399744.953 | 7063920.636 | 7226923.429 | 7235495.131 | 8379706.601 | 7695022.454 | 6675529.279 | 6070384.934 | 8398004.156 | 17.419784 | 6.353084 | 7.4103 | 17.13512 | 100391369 | VPS13C | 320528 | Vps13c | 1.12 | 0.92 | 1.03 | 0.94 | 0.3204 | 0.44959 | 1.21771 | 3 | No |
| 675699452 | PREDICTED: T-complex protein 11-like protein 1 isoform X1 [Callithrix jacchus] | 0 | 24.217 | 8 | 3 | 13 | 3 | 1 | 509 | 57 | 5.68 | 429 | 3 | 9112351.341 | 5297284.716 | 5246678.095 | 4631153.531 | 3393031.363 | 1781255.092 | 5854271.578 | 6155801.683 | 1343607.319 | 6175588.388 | 5950801.231 | 1221052.477 | 33.842301 | 43.72142 | 60.5563 | 62.88555 | 100408079 | TCP11L1 | 320554 | Tcp11l1 | 1.40 | 0.70 | 0.95 | 0.95 | 0.415 | 0.54049 | 2.00463 | 2 | No |
| 296222971 | PREDICTED: U5 small nuclear ribonucleoprotein 200 kDa helicase [Callithrix jacchus] | 0 | 17.923 | 3 | 5 | 17 | 5 | 1 | 2136 | 244.4 | 6.06 | 138 | 5 | 12663993.28 | 13291513.5 | 12740884.87 | 11150719.13 | 13054832.61 | 8033996.19 | 7003885.266 | 10043713.69 | 9310447.102 | 9643530.842 | 18366172.55 | 12098044.25 | 2.6534906 | 23.58631 | 18.0551 | 33.64528 | 100413728 | SNRNP200 | 320632 | Snrnp200 | 1.13 | 0.94 | 0.77 | 1.17 | 0.08952 | 0.19326 | 1.52165 | 1 | No |
| 1060991114 | PREDICTED: LOW QUALITY PROTEIN: protein SOGA1 [Callithrix jacchus] | 0 | 28.65 | 7 | 9 | 26 | 8 | 1 | 1458 | 163.5 | 6.19 | 133 | 9 | 10350899.08 | 15538486.71 | 11475348.56 | 4645823.375 | 6281980.991 | 7406527.629 | 1733393.457 | 2067465.184 | 6098595.621 | 4114613.01 | 1690989.252 | 4611036.075 | 21.91101 | 22.71526 | 73.6271 | 44.99801 | 100411046 | SOGA1 | 320706 | Soga1 | 1.97 | 0.96 | 0.52 | 0.55 | 0.03965 | 0.11459 | 3.77442 | 2 | No |
| 1061020766 | PREDICTED: LOW QUALITY PROTEIN: plasma membrane calcium-transporting ATPase 3 [Callithrix jacchus] | 0 | 74.526 | 12 | 10 | 101 | 5 | 1 | 1261 | 138.7 | 5.9 | 1537 | 10 | 14144303.75 | 16612144.77 | 15375039.34 | 12927460.56 | 15778370.15 | 25349459.96 | 14809742.57 | 16740931.91 | 10441387.53 | 12974355.99 | 12843565.21 | 25374652.99 | 8.0243795 | 36.11254 | 23.0571 | 42.1782 | 100403194 | ATP2B3 | 320707 | Atp2b3 | 0.95 | 1.12 | 0.87 | 1.06 | 0.8284 | 0.87891 | 1.28727 | 3 | No |
| 1060981743 | PREDICTED: uncharacterized protein KIAA1211 homolog isoform X1 [Callithrix jacchus] | 0 | 5.211 | 2 | 2 | 6 | 2 | 1 | 1235 | 136.8 | 5.6 | 49 | 2 | 1677767.534 | 759869.5429 | 935521.021 | 845225.1875 | 1515619.269 | 994949.5043 | 851147.7383 | 789198.705 | 766978.7926 | 915930.8208 | 1482573.536 | 1402233.224 | 43.332398 | 31.45777 | 5.43581 | 24.20072 | 100403030 | CRACD | 320827 | Cracd | 1.04 | 1.04 | 0.74 | 1.18 | 0.2232 | 0.34707 | 1.57882 | 1 | No |
| 296208258 | PREDICTED: neuronal growth regulator 1 isoform X1 [Callithrix jacchus] | 0 | 40.932 | 21 | 6 | 132 | 6 | 1 | 354 | 38.7 | 6.21 | 2013 | 6 | 442460528.2 | 409179158.8 | 424263680 | 571878050.8 | 539668142.3 | 524573485 | 392776391.3 | 362409342.2 | 397716175.3 | 392613541.9 | 425823254.9 | 394189312.6 | 3.9183817 | 4.430509 | 4.97491 | 4.635062 | 100402487 | NEGR1 | 320840 | Negr1 | 0.97 | 1.24 | 0.87 | 0.92 | 0.003711 | 0.03561 | 1.41913 | 3 | No |
| 296195082 | PREDICTED: ectonucleotide pyrophosphatase/phosphodiesterase family member 6 [Callithrix jacchus] | 0 | 55.866 | 20 | 8 | 72 | 8 | 1 | 429 | 48.7 | 6.54 | 897 | 8 | 255837014.1 | 365513518.6 | 406016389.8 | 290569858.3 | 425503033.6 | 454882403 | 327587649.5 | 398765594.6 | 322290601.2 | 274251737.1 | 292273571 | 260010688.3 | 22.688921 | 22.44959 | 12.2175 | 5.868463 | 100412799 | ENPP6 | 320981 | Enpp6 | 1.01 | 1.15 | 1.03 | 0.81 | 0.1467 | 0.26261 | 1.4167 | 3 | No |
| 532515316 | tropomyosin alpha-4 chain isoform 2 [Callithrix jacchus] | 0 | 41.127 | 41 | 9 | 128 | 3 | 1 | 248 | 28.6 | 4.7 | 1864 | 9 | 8048697.392 | 6696760.558 | 7737673.076 | 41702778.5 | 41583416.69 | 48784824.04 | 35884031.61 | 37604046.38 | 39495703.37 | 30484602.66 | 31509218.83 | 28907976.22 | 9.4476862 | 9.367028 | 4.79675 | 4.324491 | 100412931 | TPM4 | 326618 | Tpm4 | 0.25 | 1.47 | 1.26 | 1.01 | 2.97E-05 | 0.00574 | 5.87423 | 3 | Yes |
| 675681683 | PREDICTED: liprin-alpha-2 isoform X2 [Callithrix jacchus] | 0 | 67.919 | 13 | 15 | 73 | 7 | 1 | 1257 | 143.2 | 6.11 | 883 | 15 | 12329688.71 | 7285574.193 | 12640378.38 | 13453201.66 | 11404075.96 | 12333002.29 | 12793572.75 | 9637675.511 | 8962519.151 | 10527989.87 | 4296429.025 | 7544975.774 | 27.957212 | 8.276756 | 19.5422 | 41.79894 | 100398322 | PPFIA2 | 327814 | Ppfia2 | 1.05 | 1.21 | 1.02 | 0.73 | 0.2471 | 0.3742 | 1.66255 | 3 | No |
| 532348731 | membrane-associated phosphatidylinositol transfer protein 3 isoform 1 [Callithrix jacchus] | 0 | 19.353 | 6 | 4 | 19 | 4 | 1 | 973 | 106.6 | 6.93 | 260 | 4 | 4166100.848 | 3660038.604 | 3770617.042 | 4213873.563 | 4636311.438 | 4134422.748 | 5056632.424 | 7339867.419 | 6918206.065 | 5657017.997 | 7104840.493 | 4441222.931 | 6.8828391 | 6.232865 | 18.8706 | 23.25442 | 100385086 | PITPNM3 | 327958 | Pitpnm3 | 0.76 | 0.85 | 1.26 | 1.13 | 0.0777 | 0.17592 | 1.66553 | 1 | No |
| 1060944069 | PREDICTED: LOW QUALITY PROTEIN: autophagy-related protein 2 homolog A [Callithrix jacchus] | 0 | 4.515 | 1 | 2 | 6 | 2 | 1 | 1930 | 211.7 | 5.78 | 14 | 2 | 1434074.531 | 2231023.149 | 2443972.258 | 1468011.375 | 2137467.811 | 6088589.395 | 5193034.754 | 3543542.356 | 3084929.39 | 2484832.712 | 3504829.575 | 4584925.679 | 26.14219 | 77.27325 | 28.1359 | 29.79378 | 100407000 | ATG2A | 329015 | Atg2a | 0.64 | 1.02 | 1.24 | 1.11 | 0.1907 | 0.30924 | 1.93507 | 3 | No |
| 675665179 | PREDICTED: abl interactor 2 isoform X1 [Callithrix jacchus] | 0 | 32.391 | 13 | 5 | 37 | 4 | 1 | 542 | 58.7 | 6.02 | 363 | 5 | 21728313.37 | 14584396.27 | 15714845.51 | 20394413.75 | 17471277.27 | 12688537.2 | 19226409.8 | 15997056.84 | 17356340.87 | 16969066.68 | 16229772.72 | 10341538.44 | 22.142321 | 23.08505 | 9.25105 | 25.02401 | 100405981 | ABI2 | 329165 | Abi2 | 1.05 | 1.02 | 1.06 | 0.88 | 0.7375 | 0.80809 | 1.20761 | 3 | No |
| 675748618 | PREDICTED: protein phosphatase 1 regulatory subunit 12B isoform X1 [Callithrix jacchus] | 0 | 11.505 | 3 | 3 | 9 | 3 | 1 | 1062 | 118.5 | 5.83 | 122 | 3 | 4514932.049 | 3924617.565 | 1594169.476 | 2284517 | 3742701.805 | 1536153.852 | 2528925.426 | 2244812.739 | 1687432.061 | 3139970.244 | 474946.2837 | 1583556.313 | 46.175145 | 44.50955 | 19.8762 | 77.25929 | 100404247 | PPP1R12B | 329251 | Ppp1r12b | 1.37 | 1.03 | 0.88 | 0.71 | 0.6684 | 0.75472 | 1.93013 | 2 | No |
| 675738455 | PREDICTED: protein C3orf33 homolog [Callithrix jacchus] | 0 | 15.277 | 14 | 4 | 12 | 4 | 1 | 301 | 34.8 | 9.96 | 33 | 4 | 19288238.28 | 26219494.64 | 25691524.58 | 14956891.16 | 22307711.15 | 23211583.21 | 22962599.56 | 20173496.87 | 28367377.35 | 34879130.57 | 22537667.29 | 27475828.12 | 16.25744 | 22.45943 | 17.4786 | 21.95113 | 100388364 | C17H3orf33 | 329659 | E130311K13Rik | 0.99 | 0.84 | 0.99 | 1.18 | 0.5162 | 0.62591 | 1.40374 | 1 | No |
| 532513434 | F-box only protein 41 [Callithrix jacchus] | 0 | 20.377 | 7 | 6 | 16 | 6 | 1 | 876 | 94.5 | 8.24 | 145 | 6 | 21082098.1 | 14361437.36 | 20470349.4 | 21337607.25 | 22144575.04 | 22146236.81 | 24237902.52 | 14962193.63 | 13355607.14 | 15533969.92 | 10266775.46 | 10727083.22 | 19.938812 | 2.131928 | 33.5318 | 23.95896 | 100395524 | FBXO41 | 330369 | Fbxo41 | 1.06 | 1.25 | 1.00 | 0.69 | 0.04346 | 0.11946 | 1.79667 | 3 | No |
| 675762505 | PREDICTED: zinc finger CCCH domain-containing protein 4 [Callithrix jacchus] | 0 | 16.66 | 4 | 3 | 14 | 3 | 1 | 1306 | 140.6 | 6.25 | 358 | 3 | 3391114.659 | 2488910.01 | 2691732.631 | 2498950.75 | 2083263.85 | 3220051.69 | 3556636.78 | 3808706.722 | 3393510.1 | 3173121.081 | 2698804.44 | 2717837.112 | 16.565901 | 22.11631 | 5.83278 | 9.378184 | 100408999 | ZC3H4 | 330474 | Zc3h4 | 0.96 | 0.87 | 1.20 | 0.96 | 0.07252 | 0.16812 | 1.37894 | 3 | No |
| 675712635 | PREDICTED: dedicator of cytokinesis protein 1 isoform X1 [Callithrix jacchus] | 0 | 53.115 | 9 | 12 | 38 | 11 | 1 | 1886 | 217.7 | 7.85 | 214 | 12 | 21654503.98 | 20353298.48 | 20357008.73 | 17297863.02 | 20049042.68 | 26522294.18 | 18886965.7 | 19482825.76 | 23305242.99 | 19317806.18 | 23639714.48 | 23661991.28 | 3.608682 | 22.24414 | 11.6618 | 11.26567 | 100411936 | DOCK1 | 330662 | Dock1 | 0.98 | 1.00 | 0.97 | 1.05 | 0.8668 | 0.90367 | 1.08017 | 3 | No |
| 532276932 | hyaluronan and proteoglycan link protein 4 precursor [Callithrix jacchus] | 0 | 44.461 | 26 | 9 | 79 | 9 | 1 | 402 | 42.8 | 8.95 | 932 | 9 | 114774564.4 | 120520842.2 | 122425107.1 | 124496494 | 140581726.2 | 141627746.9 | 151905095.4 | 168434331.9 | 168137652.4 | 219316798.3 | 225415621.6 | 156657341.1 | 3.3401594 | 7.083515 | 5.80908 | 18.98572 | 100392931 | HAPLN4 | 330790 | Hapln4 | 0.77 | 0.88 | 1.05 | 1.30 | 0.01073 | 0.05825 | 1.68117 | 1 | No |
| 532524006 | opioid-binding protein/cell adhesion molecule isoform a preproprotein [Callithrix jacchus] | 0 | 26.044 | 23 | 8 | 88 | 4 | 1 | 354 | 38.8 | 6.87 | 768 | 8 | 97808485.02 | 122345016.9 | 168166091.8 | 242898671.5 | 299454967 | 266078485.9 | 145889598.3 | 122851051.5 | 124022700.4 | 182958429.2 | 96515838.25 | 127370322.5 | 27.589147 | 10.5504 | 9.91155 | 32.30256 | 100413595 | OPCML | 330908 | Opcml | 0.78 | 1.62 | 0.79 | 0.82 | 0.01226 | 0.0613 | 2.08187 | 3 | No |
| 1060944749 | PREDICTED: rho GTPase-activating protein 32 [Callithrix jacchus] | 0 | 30.205 | 6 | 11 | 32 | 11 | 1 | 2087 | 230.2 | 6.79 | 76 | 11 | 11524461.44 | 22801684.3 | 30660934.49 | 5267096.172 | 25783630.39 | 24806558.35 | 15699420.13 | 16133591.01 | 13929046.57 | 19182293.05 | 36268059.37 | 29085367.2 | 44.404099 | 62.15923 | 7.65578 | 30.44477 | 100409288 | ARHGAP32 | 330914 | Arhgap32 | 1.04 | 0.89 | 0.73 | 1.35 | 0.2824 | 0.40914 | 1.84729 | 1 | No |
| 1061019343 | PREDICTED: interleukin-1 receptor accessory protein-like 1 [Callithrix jacchus] | 0 | 23.801 | 9 | 5 | 20 | 5 | 1 | 696 | 79.9 | 6.23 | 287 | 5 | 10654059.07 | 11075976.99 | 10889211.81 | 5034378.625 | 4414983.574 | 7746776.833 | 4796595.797 | 5834173.309 | 5405801.904 | 6391614.768 | 6845741.359 | 3385368.347 | 1.944443 | 30.91531 | 9.75412 | 33.93868 | 100399196 | IL1RAPL1 | 331461 | Il1rapl1 | 1.58 | 0.83 | 0.78 | 0.81 | 0.000831 | 0.01981 | 2.03405 | 2 | Yes |
| 1061018779 | PREDICTED: kinase suppressor of Ras 2 [Callithrix jacchus] | 0 | 10.095 | 3 | 2 | 6 | 2 | 1 | 1003 | 113.4 | 8.19 | 125 | 2 | 801997.6922 | 860215.489 | 620862.2348 | 665136.25 | 585161.2916 | 395585.8403 | 633927.8101 | 376226.8893 | 425028.6915 | 546336.4003 | 249865.8229 | 288218.6756 | 16.402364 | 25.2337 | 28.6141 | 44.60636 | 100399468 | KSR2 | 333050 | Ksr2 | 1.42 | 1.02 | 0.89 | 0.67 | 0.1208 | 0.232 | 2.10534 | 2 | No |
| 675689735 | PREDICTED: cTAGE family member 5 isoform X1 [Callithrix jacchus] | 0 | 25.277 | 6 | 7 | 16 | 7 | 1 | 1407 | 159.6 | 4.81 | 217 | 7 | 5976821.069 | 7504472.633 | 7081546.606 | 7541796.938 | 6178390.976 | 6958351.449 | 4335163.58 | 5358330.853 | 4957445.473 | 2866378.477 | 782774.0321 | 2553238.788 | 11.507786 | 9.924192 | 10.5569 | 54.34372 | 100385732 | CTAGE5 | 338320 | Mia2 | 1.32 | 1.33 | 0.94 | 0.40 | 0.01173 | 0.06027 | 3.33396 | 2 | No |
| 532510221 | FK506-binding protein 15 [Callithrix jacchus] | 0 | 60.071 | 15 | 12 | 51 | 12 | 1 | 1219 | 133.6 | 5.16 | 977 | 12 | 31926619.81 | 29870732.75 | 27952486.29 | 27882920.56 | 25851948.68 | 25775077.19 | 20613857.52 | 21254712.61 | 20953068.34 | 17152436.16 | 18126952.04 | 20214486.02 | 6.6433455 | 4.510346 | 1.53105 | 8.457018 | 100390679 | FKBP15 | 338355 | Fkbp15 | 1.25 | 1.11 | 0.87 | 0.77 | 0.004845 | 0.03801 | 1.61729 | 2 | Yes |
| 1060960998 | PREDICTED: LOW QUALITY PROTEIN: melanoma inhibitory activity protein 3 [Callithrix jacchus] | 0 | 47.369 | 7 | 10 | 57 | 9 | 1 | 1837 | 205.2 | 4.88 | 738 | 10 | 48352657.12 | 40035461.55 | 31620914.24 | 46644281.88 | 43613569.95 | 28723011.42 | 37017959.81 | 36379328.87 | 24043083.83 | 21206834.89 | 18513570.76 | 14021301.27 | 20.913222 | 24.18643 | 22.5174 | 20.26419 | 100410418 | MIA3 | 338366 | Mia3 | 1.23 | 1.22 | 1.00 | 0.55 | 0.03619 | 0.10857 | 2.23307 | 2 | No |
| 675659568 | PREDICTED: unconventional myosin-Id isoform X1 [Callithrix jacchus] | 0 | 67.526 | 17 | 18 | 153 | 18 | 1 | 1006 | 116.1 | 9.38 | 1261 | 18 | 238347995 | 266107444.9 | 280042955.2 | 275951884.8 | 250400499.1 | 244250924.6 | 252958481.3 | 271477929.2 | 251227809.3 | 198241180.8 | 198208467.8 | 189532957.9 | 8.1170289 | 6.54458 | 4.34152 | 2.569161 | 100388686 | MYO1D | 338367 | Myo1d | 1.08 | 1.06 | 1.06 | 0.80 | 0.003725 | 0.03561 | 1.33877 | 3 | No |
| 532450891 | arf-GAP with GTPase, ANK repeat and PH domain-containing protein 1 isoform 1 [Callithrix jacchus] | 0 | 27.389 | 10 | 7 | 30 | 5 | 1 | 857 | 94.4 | 7.94 | 451 | 7 | 26648437.06 | 51364740.11 | 59288341.82 | 36026483.88 | 49084121.98 | 50580824.32 | 12182148.02 | 22312169.25 | 21160340.86 | 19452687.75 | 22682663.21 | 26996476.65 | 37.198496 | 17.70033 | 29.8953 | 16.42448 | 100404056 | AGAP1 | 347722 | Agap1 | 1.38 | 1.36 | 0.56 | 0.70 | 0.04125 | 0.11617 | 2.46703 | 2 | No |
| 532205955 | glycine--tRNA ligase precursor [Callithrix jacchus] | 0 | 25.977 | 11 | 6 | 14 | 6 | 1 | 739 | 83 | 7.03 | 109 | 6 | 15866357.69 | 30283371.9 | 27833131.53 | 23733233 | 24515576.54 | 22983906.67 | 9966627.41 | 12742815.91 | 14075304.34 | 16132819.62 | 15799639.31 | 12667298.39 | 31.28122 | 3.2256 | 17.0956 | 12.86045 | 100388470 | GARS | 353172 | Gars | 1.31 | 1.26 | 0.65 | 0.79 | 0.004044 | 0.03587 | 2.01124 | 2 | Yes |
| 1060938280 | PREDICTED: enhancer of mRNA-decapping protein 3 [Callithrix jacchus] | 0 | 12.234 | 8 | 3 | 6 | 3 | 1 | 508 | 56 | 7.11 | 41 | 3 | 12840650.27 | 9313064.222 | 10263432.51 | 5471462 | 4108648.568 | 3238785.488 | 3038906.682 | 7414222.63 | 3807271.723 | 2880786.138 | 10004972.53 | 11063431.51 | 16.891477 | 26.33702 | 49.1449 | 55.74654 | 100393915 | EDC3 | 353190 | Edc3 | 1.55 | 0.61 | 0.68 | 1.15 | 0.0395 | 0.11436 | 2.52886 | 2 | No |
| 1060944409 | PREDICTED: 39S ribosomal protein L21, mitochondrial isoform X1 [Callithrix jacchus] | 0 | 17.525 | 15 | 2 | 11 | 2 | 1 | 201 | 22.8 | 9.82 | 545 | 2 | 6551661.819 | 1104184.682 | 4868970.213 | 5566681.75 | 7503951.194 | 6848100.535 | 4886005.243 | 3140345.056 | 940423.6682 | 4521233.174 | 2245130.024 | 1260634.426 | 66.809772 | 14.84017 | 66.149 | 62.50379 | 100407361 | MRPL21 | 353242 | Mrpl21 | 1.01 | 1.61 | 0.73 | 0.65 | 0.08964 | 0.19326 | 2.48147 | 2 | No |
| 532516730 | unconventional myosin-XVIIIa isoform a [Callithrix jacchus] | 0 | 194.357 | 20 | 35 | 239 | 35 | 1 | 2055 | 233.2 | 6.29 | 3124 | 35 | 232906584.6 | 246091625.7 | 303332069.4 | 227000724.4 | 237536996.7 | 258397281.5 | 252837497 | 269766781.6 | 214565546.1 | 227666220.5 | 240579896.1 | 220866024.6 | 14.356716 | 6.630752 | 11.5088 | 4.359382 | 100401613 | MYO18A | 360013 | Myo18a | 1.07 | 0.99 | 1.01 | 0.94 | 0.5627 | 0.6656 | 1.13527 | 3 | No |
| 675743700 | PREDICTED: tripartite motif-containing protein 46 [Callithrix jacchus] | 0 | 31.049 | 10 | 7 | 48 | 7 | 1 | 759 | 83.4 | 7.8 | 760 | 7 | 12136866.67 | 27543799.69 | 22698747.25 | 20348858.63 | 16854358.29 | 20723786.84 | 21401613.89 | 24856174.72 | 23965771 | 16320958.94 | 28262625.83 | 22156663.19 | 37.888727 | 11.05201 | 7.66233 | 26.84142 | 100407511 | TRIM46 | 360213 | Trim46 | 0.97 | 0.90 | 1.09 | 1.04 | 0.311 | 0.43924 | 1.21228 | 3 | No |
| 1060932865 | PREDICTED: neurofilament heavy polypeptide [Callithrix jacchus] | 0 | 55.326 | 16 | 12 | 106 | 10 | 1 | 893 | 97.4 | 5.63 | 1127 | 12 | 75678981.33 | 204231998.6 | 194160873 | 45875242.63 | 157975051.1 | 193736546.9 | 130703977.3 | 215494777.2 | 212683656.9 | 163609899.2 | 221399746.2 | 195826300.8 | 45.240231 | 58.21001 | 25.8532 | 14.95697 | 100411710 | NEFH | 380684 | Nefh | 0.94 | 0.79 | 1.11 | 1.16 | 0.6901 | 0.77072 | 1.4609 | 1 | No |
| 532279464 | CB1 cannabinoid receptor-interacting protein 1 isoform CRIP1a [Callithrix jacchus] | 0 | 17.152 | 21 | 3 | 31 | 3 | 1 | 164 | 18.6 | 8.38 | 700 | 3 | 30075069.89 | 41572281.48 | 34963725.48 | 36052832.81 | 35832236.51 | 28029662.76 | 28369598.32 | 65926355.47 | 30195193.38 | 20259958.46 | 7996990.249 | 27991247.3 | 16.236605 | 13.72119 | 51.0303 | 53.7743 | 100387425 | CNRIP1 | 380686 | Cnrip1 | 1.10 | 1.03 | 1.29 | 0.58 | 0.2865 | 0.41199 | 2.21325 | 3 | No |
| 1060990712 | PREDICTED: LOW QUALITY PROTEIN: protein shisa-6 homolog [Callithrix jacchus] | 0 | 23.691 | 13 | 6 | 31 | 6 | 1 | 526 | 59.2 | 9.06 | 213 | 6 | 26299849.02 | 27984391.79 | 23522036.33 | 25737113 | 29800638.92 | 23532325.38 | 23084807.76 | 20430076.36 | 19234628.61 | 30271582.74 | 21630646.23 | 39014856.45 | 8.6884557 | 12.06432 | 9.42147 | 28.68159 | 100386882 | SHISA6 | 380702 | Shisa6 | 1.00 | 1.02 | 0.81 | 1.17 | 0.1369 | 0.25154 | 1.44889 | 1 | No |
| 675636820 | REDICTED: leucine-rich repeat and calponin homology domain-containing protein 1 isoform X1 [Callithrix jacchu | 0 | 20.322 | 7 | 4 | 20 | 3 | 1 | 764 | 84.7 | 5.82 | 173 | 4 | 8888392.809 | 10131810.71 | 12570895.99 | 7590630.219 | 11840290.13 | 11302336.64 | 10780654.08 | 11178054 | 10195355.9 | 8777793.656 | 12648635.52 | 7294730.639 | 17.789731 | 22.58727 | 4.61218 | 28.87354 | 100402285 | LRCH1 | 380916 | Lrch1 | 1.03 | 1.00 | 1.04 | 0.93 | 0.9241 | 0.94473 | 1.11953 | 3 | No |
| 1060953611 | PREDICTED: presenilins-associated rhomboid-like protein, mitochondrial isoform X1 [Callithrix jacchus] | 0 | 14.275 | 11 | 4 | 16 | 3 | 1 | 408 | 44.7 | 10.14 | 324 | 4 | 13832406.29 | 14208708.72 | 14058494.51 | 14233120.5 | 17154256.58 | 16281669.44 | 10001362.29 | 13065339.84 | 13167170.71 | 8204143.235 | 10498077.39 | 11899806.69 | 1.3498116 | 9.43694 | 14.8958 | 18.28992 | 100412777 | .OC10041277 | 381038 | Parl | 1.08 | 1.22 | 0.93 | 0.78 | 0.09468 | 0.19945 | 1.55771 | 3 | No |
| 675722722 | PREDICTED: rho guanine nucleotide exchange factor 33 isoform X1 [Callithrix jacchus] | 0 | 167.674 | 26 | 18 | 273 | 18 | 1 | 875 | 98 | 6.81 | 5565 | 18 | 373714680.5 | 371789690.1 | 312167734.7 | 282281401.9 | 272424844.2 | 272103879 | 215537060 | 251313619.8 | 220818836.9 | 308557684.4 | 306939948.6 | 318750974.2 | 9.9251049 | 2.099237 | 8.42511 | 2.056217 | 100415291 | ARHGEF33 | 381112 | Arhgef33 | 1.21 | 0.94 | 0.78 | 1.07 | 0.005135 | 0.03936 | 1.53805 | 2 | Yes |
| 675748744 | PREDICTED: plasma membrane calcium-transporting ATPase 4 isoform X4 [Callithrix jacchus] | 0 | 72.461 | 11 | 10 | 100 | 3 | 1 | 1241 | 137.7 | 6.47 | 1534 | 10 | 14077944.09 | 10522418.33 | 7807286.635 | 15499677.88 | 5496287.752 | 2987855.434 | 2342759.892 | 7213966.745 | 6404829.706 | 7121906.089 | 6877124.835 | 11316963.69 | 29.110725 | 82.79942 | 49.0621 | 29.57438 | 100410901 | ATP2B4 | 381290 | Atp2b4 | 1.33 | 0.98 | 0.65 | 1.04 | 0.3502 | 0.47755 | 2.03036 | 2 | No |
| 675640140 | PREDICTED: uncharacterized protein C9orf172 homolog [Callithrix jacchus] | 0 | 34.743 | 9 | 8 | 63 | 8 | 1 | 976 | 106.6 | 9.09 | 783 | 8 | 145288141.4 | 173942433 | 138045086.4 | 180540778.3 | 172191777 | 189218097.9 | 136482641.8 | 174117025.2 | 145666814.3 | 175694545.7 | 178056474.9 | 161836057.9 | 12.454045 | 4.712803 | 12.9016 | 5.098833 | 100393358 | AJM1 | 381353 | Ajm1 | 0.93 | 1.10 | 0.93 | 1.05 | 0.2045 | 0.32348 | 1.18779 | 3 | No |
| 1060962573 | PREDICTED: neurosecretory protein VGF [Callithrix jacchus] | 0 | 27.532 | 7 | 3 | 16 | 3 | 1 | 616 | 67.5 | 4.79 | 795 | 3 | 5262267.173 | 4452209.391 | 3404383.439 | 2398772.875 | 4979650.695 | 1690412.529 | 6644378.769 | 7125693.491 | 8674254.729 | 1506305.405 | 4036151.084 | 3027804.656 | 21.300803 | 57.26781 | 14.1773 | 44.58093 | 100415421 | VGF | 381677 | Vgf | 0.99 | 0.68 | 1.69 | 0.64 | 0.03344 | 0.10283 | 2.61886 | 3 | No |
| 1061014953 | PREDICTED: single-stranded DNA-binding protein, mitochondrial isoform X1 [Callithrix jacchus] | 0 | 11.431 | 17 | 3 | 32 | 3 | 1 | 151 | 17.3 | 10.17 | 227 | 3 | 24282539.11 | 121536282.1 | 90294106.73 | 120577288 | 102291516.2 | 67631502.29 | 80124364.91 | 81696049.46 | 95956242.77 | 81533029.62 | 115346697 | 111750155.5 | 63.086686 | 27.77096 | 10.151 | 18.05208 | 100400548 | SSBP1 | 381760 | Ssbp1 | 0.86 | 1.06 | 0.94 | 1.13 | 0.6484 | 0.73981 | 1.30713 | 3 | No |
| 296211116 | PREDICTED: protein arginine N-methyltransferase 8 isoform X1 [Callithrix jacchus] | 0 | 24.873 | 14 | 6 | 27 | 4 | 1 | 394 | 45.3 | 6.93 | 262 | 6 | 20129625.16 | 16488919.85 | 17847711.33 | 22952742.38 | 12907166.61 | 14021654.07 | 15741036.62 | 10178717.86 | 9757616.667 | 12860591.78 | 9523799.561 | 9886313.406 | 10.133363 | 33.11661 | 28.0818 | 17.02018 | 100414176 | PRMT8 | 381813 | Prmt8 | 1.26 | 1.16 | 0.83 | 0.75 | 0.04414 | 0.12057 | 1.68779 | 2 | No |
| 532448845 | serine/threonine-protein kinase TAO2 isoform 2 [Callithrix jacchus] | 0 | 77.484 | 18 | 16 | 144 | 15 | 1 | 1049 | 119.2 | 7.47 | 1874 | 16 | 112295688.8 | 97793198.59 | 77458887.7 | 94298225.34 | 108923049 | 100321641.1 | 58592058.03 | 102555701.5 | 128920839.9 | 62952222.79 | 125933644.5 | 69003220.25 | 18.257384 | 7.264393 | 36.7459 | 40.42149 | 100396700 | TAOK2 | 381921 | Taok2 | 1.01 | 1.07 | 1.02 | 0.91 | 0.9002 | 0.92574 | 1.17703 | 3 | No |
| 532514220 | serine/threonine-protein kinase BRSK1 [Callithrix jacchus] | 0 | 21.571 | 8 | 4 | 27 | 4 | 1 | 778 | 85.1 | 9.32 | 513 | 4 | 9650902.706 | 8522540.833 | 7552211.284 | 11824261.88 | 13852933.28 | 13692375.12 | 11937158.57 | 13200745.63 | 7111343.98 | 11736649.67 | 9647990.617 | 8721907.093 | 12.248515 | 8.593686 | 29.8953 | 15.38815 | 100392299 | BRSK1 | 381979 | Brsk1 | 0.81 | 1.24 | 1.01 | 0.94 | 0.04285 | 0.11855 | 1.53036 | 3 | No |
| 675762730 | PREDICTED: serine/threonine-protein kinase LMTK3 [Callithrix jacchus] | 0 | 60.639 | 10 | 9 | 58 | 9 | 1 | 1454 | 153.5 | 4.86 | 532 | 9 | 28010702.4 | 37563431.69 | 32415903.71 | 22984795.44 | 24640273.04 | 28653643.34 | 19009445.36 | 21339462.11 | 24063584.62 | 54944783.44 | 26504656.01 | 38543116.15 | 14.637721 | 11.46456 | 11.7817 | 35.69156 | 100402217 | LMTK3 | 381983 | Lmtk3 | 1.09 | 0.85 | 0.72 | 1.34 | 0.09286 | 0.19854 | 1.86288 | 1 | No |
| 1060973926 | PREDICTED: protein unc-13 homolog A [Callithrix jacchus] | 0 | 87.102 | 12 | 18 | 151 | 15 | 1 | 1721 | 194.8 | 5.34 | 1775 | 18 | 94763688.36 | 79824367.61 | 96413745.06 | 89700344.5 | 98550210.59 | 119178439.8 | 66456612.59 | 77860044.5 | 80814474.09 | 81098157.91 | 79494721.25 | 100527406.9 | 10.116761 | 14.76063 | 10.1034 | 13.45108 | 100411765 | UNC13A | 382018 | Unc13a | 1.02 | 1.16 | 0.85 | 0.98 | 0.1785 | 0.29893 | 1.36556 | 3 | No |
| 296194247 | PREDICTED: metaxin-3 isoform X1 [Callithrix jacchus] | 0 | 26.527 | 18 | 4 | 33 | 4 | 1 | 312 | 35.1 | 8.38 | 615 | 4 | 14752075.17 | 7467642.396 | 13848156.98 | 12932762.75 | 14134580.37 | 16257971.63 | 19187290.1 | 22585854.49 | 20058747 | 9834950.388 | 14551782.33 | 668752.0127 | 33.025544 | 11.65892 | 8.5646 | 84.52456 | 100391899 | MTX3 | 382793 | Mtx3 | 0.87 | 1.04 | 1.49 | 0.60 | 0.04058 | 0.11555 | 2.4678 | 3 | No |
| 296199982 | PREDICTED: NSFL1 cofactor p47 isoform X1 [Callithrix jacchus] | 0 | 28.078 | 15 | 4 | 23 | 4 | 1 | 372 | 40.8 | 5.15 | 278 | 4 | 8982707.661 | 9302019.21 | 9703862.342 | 9417859.125 | 12164089.45 | 11869904.36 | 9459706.319 | 10178003.54 | 10451781.68 | 14491554.28 | 11941874.5 | 11623207.54 | 3.8733312 | 13.52216 | 5.10843 | 12.3932 | 100398828 | NSFL1C | 386649 | Nsfl1c | 0.86 | 1.03 | 0.93 | 1.17 | 0.08951 | 0.19326 | 1.35972 | 1 | No |
| 532450625 | E3 ubiquitin-protein ligase ZNRF2 [Callithrix jacchus] | 0 | 10.437 | 10 | 2 | 8 | 2 | 1 | 242 | 24.1 | 7.09 | 43 | 2 | 6055486.608 | 9888590.823 | 11969209.7 | 518870.75 | 6053969.316 | 11758768.57 | 9362983.55 | 6148775.571 | 17073773.44 | 10921012.04 | 10552619.38 | 7619078.457 | 32.240853 | 91.97495 | 51.6911 | 18.65855 | 100405996 | ZNRF2 | 387524 | Znrf2 | 1.03 | 0.68 | 1.21 | 1.08 | 0.8059 | 0.86272 | 1.77756 | 1 | No |
| 1060966333 | PREDICTED: NADH dehydrogenase [ubiquinone] iron-sulfur protein 6, mitochondrial [Callithrix jacchus] | 0 | 13.656 | 18 | 2 | 17 | 2 | 1 | 124 | 13.6 | 8.5 | 158 | 2 | 27301736.14 | 64252492.58 | 53280361.2 | 62376049 | 48929216.23 | 49813418.05 | 62747332.75 | 53139417.32 | 52406577.67 | 29319147.63 | 43770960.66 | 25441783.44 | 39.30647 | 14.0045 | 10.2862 | 29.41059 | 100894557 | NDUFS6 | 407785 | Ndufs6 | 1.01 | 1.13 | 1.18 | 0.69 | 0.1276 | 0.24055 | 1.70801 | 3 | No |
| 1061015198 | PREDICTED: adenylate cyclase type 1 [Callithrix jacchus] | 0 | 4.466 | 2 | 2 | 7 | 2 | 1 | 914 | 101.6 | 8.27 | 33 | 2 | 5651879.505 | 3526243.138 | 7972933.42 | 6993652.25 | 9819165.08 | 7117053.073 | 7968026.788 | 4980957.987 | 8723976.397 | 6171949.259 | 7583752.351 | 9123716.062 | 38.902456 | 20.0195 | 27.3968 | 19.35819 | 100394265 | ADCY1 | 432530 | Adcy1 | 0.80 | 1.12 | 1.01 | 1.07 | 0.666 | 0.75249 | 1.39524 | 3 | No |
| 1060994185 | PREDICTED: cytospin-B isoform X1 [Callithrix jacchus] | 0 | 9.416 | 3 | 3 | 9 | 3 | 1 | 1106 | 122.5 | 7.05 | 28 | 3 | 2472334.196 | 3082053.458 | 1730725.948 | 1581068.68 | 1265337.831 | 2819253.031 | 707822.1893 | 84514.56944 | 1441758.551 | 522986.8134 | 1346289.285 | 1045485.64 | 27.867881 | 43.48959 | 91.2279 | 42.87791 | 100388084 | SPECC1 | 432572 | Specc1 | 1.61 | 1.25 | 0.49 | 0.64 | 0.1216 | 0.23277 | 3.26088 | 2 | No |
| 675662426 | PREDICTED: kelch repeat and BTB domain-containing protein 6 [Callithrix jacchus] | 0 | 5.491 | 3 | 2 | 8 | 2 | 1 | 679 | 76.5 | 6.21 | 110 | 2 | 2944287.295 | 5734321.381 | 5465042.475 | 3050682 | 3348338.018 | 3841048.858 | 3165985.375 | 2355369.695 | 1839101.959 | 3271727.226 | 2527477.143 | 3477434.327 | 32.643486 | 11.69449 | 27.2617 | 16.16233 | 100415716 | KBTBD6 | 432879 | Kbtbd6 | 1.38 | 1.00 | 0.72 | 0.90 | 0.2593 | 0.38604 | 1.92157 | 2 | No |
| 296194304 | PREDICTED: ras GTPase-activating-like protein IQGAP2 isoform X1 [Callithrix jacchus] | 0 | 18.5 | 4 | 5 | 6 | 5 | 1 | 1575 | 180.4 | 5.67 | 58 | 5 | 11956161 | 11608246.36 | 10343028.78 | 7171819.438 | 7530780.118 | 7965976.073 | 5307106.139 | 5348427.312 | 6430788.077 | 6092814.336 | 13364835.37 | 9274913.858 | 7.5109403 | 5.26307 | 11.1873 | 38.06248 | 100400958 | IQGAP2 | 544963 | Iqgap2 | 1.32 | 0.89 | 0.67 | 1.12 | 0.006013 | 0.04243 | 1.98448 | 2 | Yes |
| 532289440 | kalirin isoform 1 [Callithrix jacchus] | 0 | 15.851 | 2 | 6 | 19 | 2 | 1 | 2985 | 340 | 6.07 | 126 | 6 | 1952209.449 | 2399677.085 | 2553315.648 | 2223090 | 2085454.971 | 2361822.901 | 1790275.068 | 1814156.146 | 1906907.596 | 1455697.431 | 1743498.366 | 1639653.397 | 13.567719 | 6.214844 | 3.35374 | 9.036047 | 100400950 | KALRN | 545156 | Kalrn | 1.15 | 1.12 | 0.92 | 0.81 | 0.03159 | 0.10067 | 1.42703 | 2 | No |
| 1060961839 | PREDICTED: centrosomal protein of 170 kDa isoform X1 [Callithrix jacchus] | 0 | 47.1 | 9 | 11 | 57 | 11 | 1 | 1594 | 176.4 | 7.06 | 601 | 11 | 32889782.9 | 41472522.66 | 41744973.43 | 27446931.88 | 31206746.86 | 42398572.93 | 25639316.06 | 35804061.76 | 36471985.13 | 32428215.57 | 36196872.29 | 44042875.05 | 13.011432 | 23.08979 | 18.5996 | 15.77754 | 100399400 | CEP170 | 545389 | Cep170 | 1.09 | 0.94 | 0.92 | 1.05 | 0.6592 | 0.74772 | 1.18579 | 3 | No |
| 296207566 | PREDICTED: NADH dehydrogenase [ubiquinone] iron-sulfur protein 5 [Callithrix jacchus] | 0 | 14.638 | 35 | 4 | 14 | 4 | 1 | 106 | 12.8 | 8.72 | 141 | 4 | 60989309.05 | 34807461.72 | 28657662.02 | 47124205 | 34612927.98 | 35779302.79 | 28077734.07 | 1040204.657 | 27095689.32 | 53734585.16 | 31887890.75 | 35629163.03 | 41.386199 | 17.64349 | 81.8369 | 28.90823 | 100398013 | NDUFS5 | 595136 | Ndufs5 | 1.19 | 1.12 | 0.54 | 1.16 | 0.3958 | 0.52121 | 2.21395 | 2 | No |
| 296207300 | PREDICTED: nucleolar protein of 40 kDa [Callithrix jacchus] | 0 | 11.267 | 15 | 3 | 9 | 3 | 1 | 241 | 27.6 | 9.74 | 17 | 3 | 7555148.375 | 7271357.754 | 8787972.079 | 8025539 | 7771868.614 | 9475092.68 | 7259567.702 | 10167846.23 | 5936250.247 | 9566468.504 | 9778332.052 | 2036034.869 | 10.243019 | 10.90819 | 27.7958 | 61.87967 | 100409512 | ZCCHC17 | 619605 | Zcchc17 | 1.01 | 1.08 | 1.00 | 0.91 | 0.8879 | 0.91906 | 1.18202 | 3 | No |
| 1060972621 | PREDICTED: phospholipid hydroperoxide glutathione peroxidase, mitochondrial isoform X1 [Callithrix jacchus] | 0 | 15.511 | 17 | 4 | 27 | 4 | 1 | 234 | 26.9 | 9.83 | 265 | 4 | 55829942.52 | 29948757.73 | 10157591.74 | 52498078 | 35348032.93 | 6597911.555 | 59120150.54 | 40419428.12 | 17343206.2 | 90530077.16 | 51093119.75 | 6108519.535 | 71.621738 | 73.67252 | 53.7118 | 85.77944 | 100391793 | GPX4 | 625249 | Gpx4 | 0.84 | 0.83 | 1.03 | 1.30 | 0.9294 | 0.94889 | 1.56423 | 1 | No |
| 532286750 | mitochondrial import receptor subunit TOM40B [Callithrix jacchus] | 0 | 11.879 | 5 | 2 | 7 | 2 | 1 | 308 | 33.8 | 7.46 | 40 | 2 | 2258448.986 | 5430383.867 | 5321086.245 | 6913734.438 | 5156795.432 | 6664534.337 | 1789332.456 | 2767354.651 | 3176660.792 | 1795360.072 | 2174276.444 | 1723028.746 | 41.520522 | 15.22224 | 27.6527 | 12.77229 | 100385187 | TOMM40L | 641376 | Tomm40l | 1.15 | 1.66 | 0.68 | 0.50 | 0.01412 | 0.06441 | 3.29109 | 2 | No |
| 296214716 | PREDICTED: RNA-binding protein Nova-1 isoform X1 [Callithrix jacchus] | 0 | 20.714 | 11 | 4 | 27 | 4 | 1 | 507 | 51.7 | 8.72 | 179 | 4 | 8981172.458 | 10325175.2 | 12406220.37 | 12581794.69 | 13071140.92 | 25381567.4 | 15929422.38 | 15442760.49 | 27365149.02 | 11348430.49 | 12171133.11 | 16244093.66 | 16.324983 | 42.63479 | 34.4617 | 19.77806 | 100389932 | NOVA1 | 664883 | Nova1 | 0.70 | 1.13 | 1.30 | 0.88 | 0.2575 | 0.38492 | 1.85218 | 3 | No |
| 675737555 | PREDICTED: TRAF2 and NCK-interacting protein kinase [Callithrix jacchus] | 0 | 28.505 | 7 | 8 | 28 | 5 | 1 | 1360 | 154.8 | 7.17 | 106 | 8 | 15986276.38 | 5697453.277 | 7477630.38 | 12746119.25 | 7030438.947 | 5211640.99 | 8324968.285 | 5647444.331 | 5459609.99 | 7458930.32 | 5043048.62 | 4578730.203 | 56.570182 | 47.20161 | 24.7454 | 27.16 | 100393247 | TNIK | 665113 | Tnik | 1.29 | 1.10 | 0.86 | 0.75 | 0.6382 | 0.7325 | 1.70727 | 2 | No |
| 1060952867 | PREDICTED: protein EFR3 homolog B [Callithrix jacchus] | 0 | 36.538 | 9 | 6 | 47 | 5 | 1 | 888 | 100.3 | 6.62 | 404 | 6 | 10962776.6 | 19019450.82 | 32642195.95 | 37600937 | 33761316.55 | 37918131.37 | 31678226.55 | 26819652.49 | 45781946.47 | 36448611.98 | 23768306.17 | 20347529.04 | 52.494619 | 6.351956 | 28.6239 | 31.58724 | 100395986 | EFR3B | 668212 | Efr3b | 0.70 | 1.23 | 1.17 | 0.90 | 0.245 | 0.37264 | 1.74501 | 3 | No |
| 1060963260 | PREDICTED: LOW QUALITY PROTEIN: pro-neuregulin-2, membrane-bound isoform [Callithrix jacchus] | 0 | 10.927 | 5 | 3 | 5 | 3 | 1 | 839 | 90.9 | 9.39 | 42 | 3 | 360686.3271 | 1204346.386 | 2163373.425 | 814670.875 | 310178.2395 | 1306313.259 | 3183123.839 | 1177317.567 | 1567975.45 | 2012743.258 | 737949.4157 | 1458885.02 | 72.574599 | 61.46212 | 53.8107 | 45.55463 | 100391893 | NRG2 | 100042150 | Nrg2 | 0.92 | 0.60 | 1.46 | 1.03 | 0.4952 | 0.60851 | 2.43851 | 1 | No |
| 1061020819 | PREDICTED: 60S ribosomal protein L10 isoform X1 [Callithrix jacchus] | 0 | 21.514 | 15 | 3 | 51 | 3 | 1 | 231 | 26.4 | 10.04 | 950 | 3 | 50082075.15 | 3369995.149 | 24250028.95 | 85689163.25 | 65567362.03 | 54171307.64 | 98685687.65 | 33795425.56 | 59785083.26 | 60059414.74 | 19948684.7 | 56273619.5 | 90.344071 | 23.30595 | 50.9583 | 48.75075 | 100415355 | RPL10 | 100043346 | Rpl10-ps3 | 0.51 | 1.34 | 1.26 | 0.89 | 0.2596 | 0.38604 | 2.64379 | 3 | No |
| 1061019732 | PREDICTED: 40S ribosomal protein S4, X isoform [Callithrix jacchus] | 0 | 24.606 | 16 | 6 | 39 | 6 | 1 | 335 | 37.7 | 10.33 | 352 | 6 | 108246884 | 226387048.4 | 298064703.2 | 152532390.1 | 200126597.3 | 197565388.8 | 244749621.5 | 262944102.4 | 274408940.4 | 198078424.5 | 230524573 | 255707972.1 | 45.449107 | 14.59577 | 5.73698 | 12.66571 | 100395458 | RPS4X | 100417274 | Gm15013 | 0.96 | 0.83 | 1.18 | 1.03 | 0.0751 | 0.17192 | 1.42142 | 3 | No |
| 675671249 | PREDICTED: uncharacterized protein KIAA1107 homolog isoform X1 [Callithrix jacchus] | 0 | 19.537 | 3 | 4 | 12 | 4 | 1 | 1776 | 197.4 | 6.14 | 152 | 4 | 1692115.455 | 5933439.165 | 8016611.565 | 7018074 | 4955754.041 | 5452892.164 | 3994718.55 | 4578386.987 | 4541224.833 | 4246085.301 | 3656081.878 | 3951435.278 | 61.814339 | 18.52788 | 7.47538 | 7.46613 | 100386425 | KIAA1107 | 100503185 | Btbd8 | 1.08 | 1.20 | 0.90 | 0.82 | 0.197 | 0.31506 | 1.47016 | 3 | No |
| 296208534 | PREDICTED: 60S ribosomal protein L5 isoform X1 [Callithrix jacchus] | 0 | 63.881 | 19 | 7 | 158 | 7 | 1 | 297 | 34.4 | 9.69 | 2479 | 7 | 244453499.1 | 206981615.5 | 287453060.3 | 241346767 | 203345894.4 | 253822314.1 | 332699682.8 | 192185957.2 | 196524718.5 | 202169035.6 | 138882807 | 180799772.4 | 16.349166 | 11.2919 | 33.2277 | 18.50771 | 100401520 | RPL5 | 100503670 | Rpl5 | 1.10 | 1.04 | 1.08 | 0.78 | 0.2345 | 0.3595 | 1.4159 | 3 | No |
| 296205170 | PREDICTED: MOB-like protein phocein isoform X1 [Callithrix jacchus] | 0 | 6.52 | 10 | 2 | 17 | 2 | 1 | 225 | 26 | 5.78 | 73 | 2 | 25794604.42 | 18730059.63 | 16672027.86 | 17601891.25 | 16889335.75 | 15121341.68 | 13522027.29 | 13779255.11 | 9651764.629 | 19641695.97 | 15925097.57 | 10341827.46 | 23.45607 | 7.722737 | 18.7725 | 30.58937 | 100385103 | .OC10038510 | - | - | 1.26 | 1.02 | 0.76 | 0.95 | 0.1865 | 0.30585 | 1.65607 | 2 | No |
| 1061003411 | PREDICTED: actin-3-like [Callithrix jacchus] | 0 | 75.683 | 37 | 9 | 456 | 2 | 1 | 375 | 41.6 | 6.15 | 7011 | 9 | 11820600.97 | 11560947.68 | 12195676.98 | 31023558 | 27973969.38 | 14530376.17 | 19288853.73 | 22738592.73 | 19379305.79 | 6349813.7 | 9272486.367 | 3862392.913 | 2.6908418 | 35.80473 | 9.60538 | 41.6937 | 100385454 | .OC10038545 | - | - | 0.75 | 1.55 | 1.29 | 0.41 | 0.01407 | 0.06441 | 3.77362 | 3 | No |
| 296237487 | PREDICTED: actin, partial [Callithrix jacchus] | 0 | 57.18 | 43 | 6 | 463 | 2 | 1 | 185 | 20.4 | 5.52 | 6395 | 6 | 8248388.889 | 6739125.116 | 6591292.167 | 19763097 | 9350754.606 | 9981537.714 | 8515561.226 | 6562930.508 | 4752061.991 | 24989093.63 | 23150275.37 | 12163539.66 | 12.749084 | 44.79814 | 28.4742 | 34.50197 | 100388094 | .OC10038809 | - | - | 0.61 | 1.11 | 0.56 | 1.71 | 0.1442 | 0.25947 | 3.04091 | 1 | No |
| 296211574 | PREDICTED: tubulin alpha-1A chain [Callithrix jacchus] | 0 | 145.47 | 52 | 17 | 426 | 6 | 1 | 451 | 50.1 | 5.02 | 7772 | 17 | 77339480.31 | 57744521.1 | 57337305.73 | 137383290.3 | 129011748.6 | 145530020.7 | 129851766.9 | 125304180.2 | 134816142.7 | 43405189.7 | 44896103.24 | 41820666.98 | 17.824202 | 6.015214 | 3.65988 | 3.545802 | 100392817 | .OC10039281 | - | - | 0.68 | 1.47 | 1.39 | 0.46 | 8.31E-05 | 0.00964 | 3.16568 | 3 | Yes |
| 296193086 | PREDICTED: 40S ribosomal protein S3a [Callithrix jacchus] | 0 | 21.397 | 27 | 8 | 52 | 8 | 1 | 264 | 29.9 | 9.57 | 292 | 8 | 173420723.4 | 213433957.7 | 149418448.5 | 193458056.3 | 173259435.1 | 148219255.5 | 219165848.1 | 291798441.7 | 248461847.2 | 175656558.8 | 266794370.8 | 207690227.6 | 18.091383 | 13.2031 | 14.4353 | 21.33419 | 100394558 | .OC10039455 | - | - | 0.87 | 0.84 | 1.23 | 1.06 | 0.1434 | 0.25893 | 1.47479 | 3 | No |
| 1060935277 | PREDICTED: eukaryotic initiation factor 4A-III [Callithrix jacchus] | 0 | 19.382 | 10 | 5 | 18 | 5 | 1 | 411 | 46.7 | 6.24 | 50 | 5 | 3308470.605 | 3905644.895 | 4014084.779 | 6218235.75 | 7813905.406 | 5552306.015 | 10584015.7 | 9182556.181 | 7258810.021 | 16376591.16 | 4481186.358 | 5137149.71 | 10.152232 | 17.80313 | 18.5317 | 77.16702 | 100397693 | .OC10039769 | - | - | 0.54 | 0.93 | 1.29 | 1.24 | 0.02888 | 0.0956 | 2.40692 | 1 | No |
| 1060997095 | PREDICTED: protein SET [Callithrix jacchus] | 0 | 53.272 | 23 | 5 | 89 | 4 | 1 | 291 | 33.7 | 4.37 | 1687 | 5 | 418536650.2 | 428680012.1 | 425806107.1 | 229686284.8 | 283333595.1 | 347254141.9 | 357368023.5 | 337694176.2 | 378316350 | 425129283.2 | 481475703.3 | 410499737 | 1.2320291 | 20.52556 | 5.6777 | 8.535908 | 100400147 | .OC10040014 | - | - | 1.13 | 0.76 | 0.95 | 1.16 | 0.02747 | 0.09157 | 1.53103 | 1 | No |
| 675725269 | EDICTED: NADH dehydrogenase [ubiquinone] 1 beta subcomplex subunit 5, mitochondrial-like [Callithrix jacch | 0 | 14.545 | 16 | 3 | 31 | 3 | 1 | 204 | 23.5 | 9.1 | 148 | 3 | 12637275.94 | 18526034.84 | 20845244.86 | 15521017.5 | 18537540.97 | 23873565.17 | 15681483.55 | 16271756.51 | 13885875.02 | 11871041.58 | 12245108.16 | 8833513.818 | 24.407736 | 21.90293 | 8.13268 | 17.03574 | 100401813 | .OC10040181 | - | - | 1.10 | 1.23 | 0.97 | 0.70 | 0.09604 | 0.20134 | 1.7582 | 3 | No |
| 1060943807 | PREDICTED: ATP synthase subunit g, mitochondrial [Callithrix jacchus] | 0 | 10.038 | 21 | 2 | 39 | 2 | 1 | 120 | 13.4 | 9.85 | 539 | 2 | 35980715.44 | 38586865.1 | 38875200.35 | 40073670 | 46728726.57 | 40968980.59 | 32046157.09 | 29039432.5 | 34498679.17 | 24985157.56 | 24605302.67 | 26835525.17 | 4.2164717 | 8.480056 | 8.58188 | 4.683661 | 100405644 | .OC10040564 | - | - | 1.10 | 1.24 | 0.93 | 0.74 | 0.001752 | 0.02698 | 1.67183 | 3 | Yes |
| 675646136 | PREDICTED: NADH dehydrogenase [ubiquinone] 1 alpha subcomplex subunit 5-like [Callithrix jacchus] | 0 | 15.881 | 22 | 3 | 18 | 3 | 1 | 153 | 17.5 | 6.33 | 214 | 3 | 70756878.07 | 59076444.55 | 56487999.15 | 61883301.75 | 61197090.05 | 63846372.7 | 58528032.36 | 54073959.44 | 63869877.36 | 49935003.55 | 60188519.9 | 57064441.59 | 12.239987 | 2.206696 | 8.33786 | 9.430451 | 100405683 | .OC10040568 | - | - | 1.04 | 1.04 | 0.98 | 0.93 | 0.3866 | 0.51325 | 1.11806 | 3 | No |
| 1061020833 | PREDICTED: creatine kinase B-type [Callithrix jacchus] | 0 | 57.103 | 16 | 5 | 110 | 5 | 1 | 419 | 46.9 | 5.52 | 1690 | 5 | 312884485.7 | 171008508.8 | 157406297.7 | 322537843.1 | 202941434.3 | 216442716.2 | 318653088.5 | 203389890.3 | 222405928.7 | 234750514.1 | 174586549.6 | 171228153.7 | 40.281237 | 26.4854 | 24.9018 | 18.4706 | 100406585 | .OC10040658 | - | - | 0.95 | 1.10 | 1.10 | 0.86 | 0.6082 | 0.70657 | 1.28228 | 3 | No |
| 296222312 | PREDICTED: myosin regulatory light polypeptide 9 isoform X1 [Callithrix jacchus] | 0 | 39.325 | 26 | 5 | 119 | 2 | 1 | 205 | 23.5 | 4.94 | 1950 | 5 | 292610010.2 | 191637616.6 | 207294279.6 | 6124739.5 | 189363886 | 180210373.4 | 3076146.197 | 152805040.4 | 166908387.3 | 178455264.5 | 177465968.9 | 176479135.8 | 23.574949 | 82.44813 | 84.3815 | 0.55676 | 100410505 | .OC10041050 | - | - | 1.44 | 0.78 | 0.67 | 1.11 | 0.4099 | 0.53586 | 2.14239 | 2 | No |
| 326368312 | paralemmin-2 [Callithrix jacchus] | 0 | 15.78 | 13 | 4 | 20 | 3 | 1 | 413 | 45.8 | 5.12 | 183 | 4 | 6838837.172 | 8481102.311 | 5892243.107 | 4450306.063 | 7028020.06 | 8644239.86 | 6087286.719 | 7093343.229 | 5240772.495 | 18771507.68 | 23773607.18 | 7845740.876 | 18.52589 | 31.53559 | 15.1036 | 48.49363 | 100412199 | PALM2AKAP2 | - | - | 0.77 | 0.73 | 0.67 | 1.83 | 0.3484 | 0.47584 | 2.73545 | 1 | No |
| 1061015155 | PREDICTED: MICOS complex subunit MIC19 isoform X1 [Callithrix jacchus] | 0 | 34.409 | 24 | 5 | 84 | 5 | 1 | 238 | 27.1 | 8.12 | 955 | 5 | 73132580.89 | 89900099.07 | 89746753.34 | 81445525.25 | 97290234.44 | 103121383.9 | 96626842.74 | 112564634.6 | 116247451.8 | 53797745.79 | 67539788.73 | 76302843.97 | 11.43697 | 11.93882 | 9.61239 | 17.21911 | 100412273 | .OC10041227 | - | - | 0.96 | 1.07 | 1.23 | 0.75 | 0.04683 | 0.12536 | 1.64663 | 3 | No |
| 1061019786 | PREDICTED: neuroligin-4, X-linked [Callithrix jacchus] | 0 | 20.275 | 6 | 4 | 20 | 3 | 1 | 816 | 91.8 | 6.15 | 296 | 4 | 328679.6929 | 5369385.529 | 4443158.583 | 4777237.5 | 9682610.261 | 8659874.349 | 2069534.795 | 2946181.024 | 5775908.673 | 1087568.218 | 4095287.906 | 2238393.713 | 79.373359 | 33.5805 | 53.8488 | 61.34854 | 100412584 | NLGN4X | - | - | 0.79 | 1.80 | 0.84 | 0.58 | 0.205 | 0.32398 | 3.11534 | 3 | No |
| 675718851 | PREDICTED: T-complex protein 1 subunit gamma-like [Callithrix jacchus] | 0 | 72.603 | 25 | 13 | 123 | 13 | 1 | 569 | 62.8 | 6.86 | 1692 | 13 | 202767400.7 | 126825326.2 | 150964834.1 | 220213516.2 | 152372379.8 | 145841685 | 232006338 | 157116462.8 | 167873475 | 187658462.5 | 140556505.5 | 144178790.9 | 24.222911 | 23.83149 | 21.8087 | 16.64585 | 100412770 | .OC10041277 | - | - | 0.95 | 1.02 | 1.10 | 0.93 | 0.8249 | 0.87788 | 1.17909 | 3 | No |
| 296191756 | PREDICTED: histone-binding protein RBBP4-like [Callithrix jacchus] | 0 | 12.826 | 6 | 3 | 15 | 3 | 1 | 425 | 47.5 | 4.97 | 273 | 3 | 31532785.88 | 19751313.11 | 17509855.18 | 10709098 | 33972429.62 | 33776077.72 | 18447237.47 | 14354711.2 | 9298435.593 | 17504784.37 | 13093231.29 | 14167606.7 | 32.849947 | 51.14139 | 32.6566 | 15.41673 | 100413976 | .OC10041397 | - | - | 1.18 | 1.34 | 0.72 | 0.76 | 0.3869 | 0.51325 | 1.86358 | 2 | No |
| 675650659 | PREDICTED: histone H4-like [Callithrix jacchus] | 0 | 12.832 | 16 | 3 | 31 | 3 | 1 | 193 | 21.6 | 11.52 | 429 | 3 | 170969450.8 | 212664819.4 | 187150464.8 | 142790498 | 199158272.9 | 216880332.7 | 127003115 | 156342973.9 | 130089303.4 | 586663851.6 | 411680714.6 | 499616191.9 | 11.048508 | 20.76929 | 11.6989 | 17.52221 | 103792192 | .OC10379219 | - | - | 0.75 | 0.74 | 0.54 | 1.97 | 0.0119 | 0.06027 | 3.6232 | 1 | No |
| 675660734 | PREDICTED: keratin, type I cytoskeletal 10-like [Callithrix jacchus] | 0 | 11.205 | 27 | 3 | 14 | 3 | 1 | 171 | 20 | 6.35 | 159 | 3 | 10315871.07 | 6172645.67 | 6706065.519 | 12594461.75 | 17683005.89 | 15674541.55 | 5458391.538 | 9409063.52 | 3325106.975 | 6091290.185 | 11871340.88 | 6943383.401 | 29.15262 | 16.73274 | 50.9036 | 37.5856 | 103793316 | .OC10379331 | - | - | 0.83 | 1.64 | 0.65 | 0.89 | 0.05834 | 0.14493 | 2.52587 | 3 | No |
| 1060987816 | PREDICTED: 60S ribosomal protein L31 [Callithrix jacchus] | 0 | 12.011 | 19 | 3 | 35 | 3 | 1 | 125 | 14.5 | 10.54 | 419 | 3 | 115443360.6 | 106825025.2 | 126917492 | 94113368.25 | 73840293.42 | 104556022 | 131403316.2 | 234119298.6 | 153073422.4 | 213133747.4 | 195289329.8 | 198037204.7 | 8.6601456 | 17.19335 | 31.3207 | 4.752816 | 108591274 | .OC10859127 | - | - | 0.80 | 0.62 | 1.19 | 1.39 | 0.00136 | 0.02375 | 2.22546 | 1 | Yes |
| 532276140 | brefeldin A-inhibited guanine nucleotide-exchange protein 3 [Callithrix jacchus] | 0 | 43.845 | 6 | 11 | 44 | 11 | 1 | 2177 | 240.6 | 5.71 | 513 | 11 | 18290583.17 | 10229655.39 | 11476221.66 | 10557634.06 | 14451618.01 | 10862676.92 | 18607110.38 | 11434022.48 | 10275257.86 | 14012144.15 | 11303845.54 | 5786312.913 | 32.546314 | 18.11036 | 33.5836 | 40.5668 | - | KIAA1244 | - | - | 1.09 | 0.97 | 1.10 | 0.84 | 0.8507 | 0.89439 | 1.29709 | 3 | No |
| 532444627 | beta-adrenergic receptor kinase 1 [Callithrix jacchus] | 0 | 9.777 | 4 | 3 | 9 | 3 | 1 | 689 | 79.5 | 7.28 | 153 | 3 | 12296218.28 | 14825321.79 | 10992815.79 | 11103949 | 8966256.285 | 11941737.14 | 6282241.469 | 8711547.72 | 8477778.017 | 4803227.475 | 7789192.264 | 8973988.315 | 15.337889 | 14.37903 | 17.1295 | 29.89682 | - | ADRBK1 | - | - | 1.32 | 1.11 | 0.82 | 0.75 | 0.07841 | 0.17673 | 1.7673 | 2 | No |
| 532513956 | FAD-linked sulfhydryl oxidase ALR [Callithrix jacchus] | 0 | 10.512 | 12 | 2 | 14 | 2 | 1 | 206 | 23.8 | 7.28 | 195 | 2 | 5420052.496 | 3900009.394 | 3772852.401 | 5795707.25 | 3697278.642 | 3451636.843 | 5920049.274 | 4438015.059 | 3335046.942 | 7678550.027 | 4617906.639 | 3182303.989 | 21.000189 | 29.85735 | 28.4185 | 44.51026 | - | GFER | - | - | 0.95 | 0.94 | 0.99 | 1.12 | 0.9598 | 0.9704 | 1.19577 | 1 | No |
| 532514088 | inositol 1,4,5-trisphosphate receptor type 2 [Callithrix jacchus] | 0 | 62.443 | 7 | 16 | 75 | 9 | 1 | 2704 | 308.2 | 6.47 | 443 | 16 | 8008276.176 | 9030723.61 | 5362465.192 | 7367065.625 | 4636101.872 | 6903059.328 | 7286017.388 | 5037208.648 | 6904551.4 | 6937341.271 | 6734466.492 | 5823682.494 | 25.351693 | 23.18776 | 18.7766 | 9.127409 | - | ITPR2 | - | - | 1.12 | 0.94 | 0.96 | 0.97 | 0.8834 | 0.91755 | 1.18487 | 3 | No |
| 296208015 | PREDICTED: 40S ribosomal protein SA [Callithrix jacchus] | 0 | 27.375 | 17 | 4 | 42 | 4 | 1 | 295 | 32.8 | 4.87 | 706 | 4 | 43768904.87 | 29245186.24 | 25295818.28 | 36980987.75 | 28133250.5 | 22334779.28 | 45701125.3 | 36355888.17 | 40738332.67 | 20640855.1 | 10549586.97 | 9066145.386 | 29.685374 | 25.30326 | 11.423 | 46.93584 | 100386315 | .OC10038631 | - | - | 1.13 | 1.00 | 1.41 | 0.46 | 0.02324 | 0.08193 | 3.05032 | 3 | No |
| 1060938502 | PREDICTED: ferritin heavy chain [Callithrix jacchus] | 0 | 18.5 | 22 | 4 | 23 | 4 | 1 | 183 | 21.3 | 5.85 | 156 | 4 | 42814743.84 | 36856169.47 | 40564382.89 | 42835750.44 | 60237510.14 | 56215647.19 | 83104064.66 | 103468719.2 | 51710131.35 | 154618951.7 | 92909719.68 | 153585418.3 | 7.5074397 | 17.15863 | 32.8279 | 26.42635 | 100387518 | .OC10038751 | - | - | 0.52 | 0.69 | 1.04 | 1.75 | 0.04663 | 0.12529 | 3.33608 | 1 | No |
| 1060932431 | PREDICTED: septin-4 isoform X1 [Callithrix jacchus] | 0 | 85.26 | 30 | 12 | 247 | 11 | 1 | 494 | 57 | 5.59 | 4183 | 12 | 370674283 | 365798472.6 | 313571433.9 | 388955812.5 | 289773969.8 | 308306304.9 | 372808848.8 | 362958952.4 | 345023966 | 464967377.2 | 418917422.5 | 435124889.8 | 9.0438587 | 16.02778 | 3.91023 | 5.312847 | 100388347 | .OC10038834 | - | - | 0.95 | 0.89 | 0.97 | 1.19 | 0.03752 | 0.11099 | 1.33633 | 1 | No |
| 296212580 | PREDICTED: ubiquitin-conjugating enzyme E2 N isoform X1 [Callithrix jacchus] | 0 | 16.224 | 29 | 4 | 14 | 4 | 1 | 152 | 17.1 | 6.57 | 118 | 4 | 9814328.078 | 12813496.84 | 10998105.94 | 15799729.5 | 15268057.51 | 15743049.55 | 16107793.02 | 11280884.32 | 14861003.97 | 14823866.82 | 14982550.41 | 14222842.11 | 13.477352 | 1.871217 | 17.792 | 2.73052 | 100390645 | .OC10039064 | - | - | 0.81 | 1.12 | 1.01 | 1.06 | 0.03615 | 0.10857 | 1.39211 | 3 | No |
| 296196229 | PREDICTED: 60S ribosomal protein L7a isoform X1 [Callithrix jacchus] | 0 | 24.258 | 21 | 5 | 73 | 5 | 1 | 266 | 30 | 10.61 | 966 | 5 | 113907322.5 | 139155169.1 | 126054454.5 | 116910808.4 | 99246914.65 | 93586722.78 | 81849098.7 | 137408603 | 138370001.2 | 109427891.6 | 184846968.9 | 62832459.24 | 9.9918437 | 11.78323 | 27.1442 | 51.72568 | 100392033 | .OC10039203 | - | - | 1.08 | 0.88 | 1.02 | 1.02 | 0.3784 | 0.50492 | 1.22397 | 3 | No |
| 1060931563 | PREDICTED: translin-like [Callithrix jacchus] | 0 | 16.969 | 19 | 3 | 11 | 3 | 1 | 228 | 26.2 | 6.44 | 104 | 3 | 3975195.893 | 4409965.856 | 5256305.385 | 5193900.469 | 6441249.589 | 6553279.008 | 7001209.309 | 7590167.33 | 8285471.487 | 3858946.81 | 3947307.014 | 3217118.741 | 14.327198 | 12.44603 | 8.43033 | 10.84578 | 100399437 | .OC10039943 | - | - | 0.83 | 1.11 | 1.39 | 0.67 | 0.004277 | 0.03648 | 2.0753 | 3 | Yes |
| 167427222 | ribosomal protein S25 (predicted) [Callithrix jacchus] | 0 | 13.245 | 16 | 3 | 27 | 3 | 1 | 125 | 13.8 | 10.11 | 275 | 3 | 40417424.86 | 37248394.94 | 40059356.48 | 48532933.5 | 41885167.95 | 60914050.43 | 77780062.79 | 112400269.3 | 141520402 | 40115744.05 | 41010644.1 | 37937037.46 | 4.4226749 | 19.14462 | 28.8601 | 3.983233 | 100400186 | .OC10040018 | - | - | 0.65 | 0.84 | 1.84 | 0.66 | 0.08449 | 0.18539 | 2.81759 | 3 | No |
| 1060931222 | PREDICTED: LOW QUALITY PROTEIN: myelin-associated glycoprotein [Callithrix jacchus] | 0 | 28.179 | 9 | 5 | 65 | 5 | 1 | 626 | 69.1 | 5.05 | 394 | 5 | 197337740.4 | 247249251 | 285723182.9 | 199325769.5 | 246871765.1 | 273898588.8 | 261701139.8 | 332543122.4 | 385990224.4 | 278734751.6 | 361102481.6 | 185518528.7 | 18.204275 | 15.72872 | 19.0813 | 31.93089 | 100402349 | .OC10040234 | - | - | 0.90 | 0.88 | 1.20 | 1.01 | 0.4074 | 0.53339 | 1.36126 | 3 | No |
| 675646566 | PREDICTED: LOW QUALITY PROTEIN: eukaryotic initiation factor 4A-II-like [Callithrix jacchus] | 0 | 28.435 | 12 | 5 | 35 | 4 | 1 | 408 | 46.5 | 5.68 | 358 | 5 | 41670522.52 | 40392848.76 | 25645492.74 | 26933993.75 | 23230054.07 | 25291860.48 | 28153212.63 | 26967802.16 | 40536270.09 | 15606244.56 | 25171843.66 | 25968298.57 | 24.806215 | 7.378858 | 23.5684 | 25.91774 | 100406173 | .OC10040617 | - | - | 1.25 | 0.87 | 1.11 | 0.77 | 0.2967 | 0.42247 | 1.6137 | 3 | No |
| 296206714 | PREDICTED: 40S ribosomal protein S11-like [Callithrix jacchus] | 0 | 24.361 | 30 | 6 | 96 | 6 | 1 | 158 | 18.4 | 10.24 | 1209 | 6 | 217399675.2 | 258083188.8 | 269107496.3 | 193221399.8 | 207140403.7 | 198112132.9 | 335707782.1 | 317272553.5 | 327764560.1 | 226171336.6 | 238056738.3 | 248229868 | 10.973038 | 3.539628 | 2.82854 | 4.648837 | 100407326 | .OC10040732 | - | - | 0.98 | 0.79 | 1.29 | 0.94 | 0.0002642 | 0.01455 | 1.63874 | 3 | Yes |
| 1060930458 | PREDICTED: heat shock protein HSP 90-beta-like [Callithrix jacchus] | 0 | 46.573 | 27 | 8 | 135 | 2 | 1 | 337 | 39.8 | 5.34 | 2196 | 8 | 39799453.93 | 13359408.79 | 17378337.76 | 21291912 | 19806195.68 | 25222453.28 | 21940012.62 | 24202791.2 | 25887149.2 | 14816113.25 | 17605233.43 | 18030314.35 | 60.595354 | 12.65934 | 8.24915 | 10.38218 | 100409130 | .OC10040913 | - | - | 1.09 | 1.02 | 1.11 | 0.78 | 0.05294 | 0.13687 | 1.4277 | 3 | No |
| 296194351 | PREDICTED: 60S ribosomal protein L26-like [Callithrix jacchus] | 0 | 16.397 | 24 | 5 | 48 | 2 | 1 | 145 | 17.1 | 10.46 | 518 | 5 | 35130560.84 | 41109986.09 | 25176089.76 | 47661156 | 62895824.27 | 24857062.49 | 78006663.86 | 97493086.65 | 141192699.3 | 4652111.773 | 146577509.5 | 112904584.6 | 23.810183 | 42.4132 | 30.6515 | 84.22614 | 100409208 | .OC10040920 | - | - | 0.50 | 0.66 | 1.55 | 1.29 | 0.121 | 0.23213 | 3.12269 | 1 | No |
| 1060946011 | PREDICTED: ADP-ribosylation factor-like protein 6-interacting protein 1 isoform X1 [Callithrix jacchus] | 0 | 25.998 | 14 | 5 | 75 | 5 | 1 | 296 | 33.9 | 9.58 | 1125 | 5 | 188949726.7 | 186537760.5 | 226574395.7 | 219640841.2 | 179162569.6 | 220453154.3 | 278398536.9 | 263387818 | 227538879.9 | 258510139.6 | 201479273.1 | 231111985.7 | 11.187206 | 11.437 | 10.1901 | 12.38142 | 100409293 | .OC10040929 | - | - | 0.90 | 0.92 | 1.15 | 1.03 | 0.195 | 0.31346 | 1.27782 | 3 | No |
| [illegible] | [illegible] | [illegible] | [illegible] | [illegible] | [illegible] | [illegible] | [illegible] | [illegible] | [illegible] | [illegible] | [illegible] | [illegible] | [illegible] | [illegible] | [illegible] | [illegible] | [illegible] | [illegible] | [illegible] | [illegible] | [illegible] | [illegible] | [illegible] | [illegible] | [illegible] | [illegible] | [illegible] | [illegible] | [illegible] | [illegible] | [illegible] | [illegible] | [illegible] | [illegible] | [illegible] | [illegible] | [illegible] | [illegible] | [illegible] | [illegible] | [illegible] | [illegible] |
